# Distinct cohorts of *Aspergillus fumigatus* transcription factors are required for epithelial damage occurring via contact- or soluble effector-mediated mechanisms

**DOI:** 10.1101/2022.04.07.487466

**Authors:** S Rahman, N van Rhijn, P Papastamoulis, D.D Thomson, Z Carter, L Gregson, R Fortune-Grant, M Rattray, J M Bromley, M E Bignell

## Abstract

Damage to the lung epithelium is a unifying feature of disease caused by the saprophytic fungus *Aspergillus fumigatus*. However, the mechanistic basis and the regulatory control of such damage is poorly characterized. Previous studies have identified *A. fumigatus* mediated pathogenesis as occurring at early (≤ 16 hours) or late (>16 hours) phases of the fungal interaction with epithelial cells, and respectively involve direct contact with the host cell or the action of soluble factors produced by mature fungal hyphae. Both early and late phases of epithelial damage have been shown to be subject to genetic regulation by the pH-responsive transcription factor PacC. This study sought to determine whether other transcriptional regulators play a role in modulating epithelial damage. In particular, whether the early and late phases of epithelial damage are governed by same or distinct regulators. Furthermore, whether processes such as spore uptake and hyphal adhesion, that have previously been documented to promote epithelial damage, are governed by the same cohorts of epithelial regulators. Using 479 strains from the recently constructed library of *A. fumigatus* transcription factor null mutants, two high-throughput screens assessing epithelial cell detachment and epithelial cell lysis were conducted. A total of 17 transcription factor mutants were found to exhibit reproducible deficits in epithelial damage causation. Of these, 10 mutants were defective in causing early phase damage via epithelial detachment and 8 mutants were defective in causing late phase damage via epithelial lysis. Remarkably only one transcription factor, PacC, was required for causation of both phases of epithelial damage. The 17 mutants exhibited varied and often unique phenotypic profiles with respect to fitness, epithelial adhesion, cell wall defects, and rates of spore uptake by epithelial cells. Strikingly, 9 out of 10 mutants deficient in causing early phase damage also exhibited reduced rates of hyphal extension, and culture supernatants of 7 out of 8 mutants deficient in late phase damage were significantly less cytotoxic. Our study delivers the first high-level overview of *A. fumigatus* regulatory genes governing lung epithelial damage, suggesting highly coordinated genetic orchestration of host-damaging activities that govern epithelial damage in both space and time.

**Contribution to the Field Statement:** *Aspergillus fumigatus* is a soil dwelling fungus that can cause lethal lung infections in individuals with a compromised immune system. Disease initiates with inhalation of the fungal spores, followed by growth of the fungus leading to destruction of the lung. Our understanding of the *A. fumigatus* genes and mechanisms driving lung damage leading to establishment of disease is limited. This study has identified the genes moderating lung damage by assessing 479 regulatory mutants of *A. fumigatus* for their ability to cause epithelial damage using a lung cell line model. We observed that distinct cohorts of transcriptional regulators are required for driving early, and late phases of epithelial damage and that early- and late-occurring damage are associated respectively with hyphal growth rates and secreted fungal products. This study is the first to reveal that mechanistically distinct programs of host damage elicited during early and late stage of fungal interaction with epithelial cells are genetically regulated via distinct cohorts of *A. fumigatus* transcription factors.

## Introduction

The lung is the major portal of human exposure to airborne particles, including the spores of various fungal species, some of which can cause severe infections. Spores of the saprophytic pathogen, *Aspergillus fumigatus* are small (2-3 μm) and ubiquitous in indoor and outdoor environments (Kwon-Chung and Sugui, 2013; van Rhijn *et al*., 2021). The spores upon inhalation, easily reach the alveolar regions of the human lung (Escobar *et al*., 2016). Interaction of *A. fumigatus* spores with the respiratory epithelium can sometimes cause disease, most often in hosts having immune dysfunction. The resultant spectrum of pulmonary syndromes, called aspergilloses, can be broadly categorised into allergic, chronic and invasive diseases. The most severe manifestation is invasive pulmonary aspergillosis (IPA) occurring in individuals suffering extreme immunocompromise suffering prolonged neutropenia in haematological malignancy, post-stem cell or solid organ transplantation, or on immunosuppressive treatments (Kosmidis and Denning, 2015). IPA can also complicate the clinical course of COVID-19 and influenza and is associated with a significant increase in mortality, especially in critically ill patients admitted to intensive care units (Schauwvlieghe *et al*., 2018; Casalini *et al*., 2021). The annual burden of fatal disease is estimated at over 200,000 per annum for IPA (Brown *et al*., 2012; Bongomin *et al*., 2017). Understanding the pathogenic mechanisms leading to lung damage, as well as identifying *A. fumigatus* genes and attributes involved in the process could lead to a more targeted therapeutic strategy that subverts adverse pathology.

The pathogenesis of *A. fumigatus*-mediated disease occurs in a stepwise fashion that involves the morphological transition of the inhaled fungal spore into a hyphal form. Broadly speaking, epithelial damage can be considered as occurring at early (spore) or late (hyphal) phases of the fungal interaction with epithelial cells. Contact-mediated damage leads to detachment of the epithelial cells within 16 hours of *A. fumigatus* infection *in vitro*, wherein even a mere contact with spores leads to loss of actin polymerisation, rounding and epithelial detachment (Kogan *et al*., 2004), or via activation of host immune signaling (Okaa *et al*., 2021). *In vitro* studies have shown that alveolar epithelial cells internalise 30-50% of encountered spores (Wasylnka and Moore, 2003) leading to fungal killing or intracellular occupancy that might aid immune evasion and dissemination (reviewed in Bertuzzi *et al*., 2018) and actin-mediated internalization of fungal spores might also disrupt epithelial integrity by provoking the detachment of epithelial cells during early infection (Bertuzzi *et al*., 2014). If the spore form is not rapidly neutralised by the combined activities of epithelial cells and professional phagocytes, spore germination initiates with the breaking of dormancy and isotropic swelling of the spores (Baltussen *et al*., 2019). Spore germination leads to outgrowth of an apically extending cell called a primary hypha. Hyphal growth within, or close to, epithelial monolayers is associated with fungal invasion of epithelial cells, and transcriptomic analysis of this process both *in vitro* and *in vivo* suggests the involvement of fungal proteases and toxic secondary metabolites (Bertuzzi *et al*., 2014; Escobar *et al*., 2018). Any alterations in cell wall composition has been implied in deficit host-fungal interaction and altered virulence in invasive fungi (Bulawa *et al*., 1995; Valiante *et al*., 2015). *In vitro* infection studies have revealed specific cell wall adhesins mediating adhesion to the lung epithelium at the distinct morphological growth stages of the fungus (Levdansky *et al*., 2010; Liu *et al*., 2016; Voltersen *et al*., 2018). The secreted and hyphal exopolysaccharide galactosaminogalactan (GAG), that mediates adherence of *A. fumigatus* hyphae to host cells is also critical for host damage and for virulence in murine IPA models (Gravelat *et al*., 2013). *A. fumigatus* secreted proteases cause destruction of the mammalian F-actin cytoskeleton and loss of focal adhesion (Kogan *et al*., 2004; Namvar *et al*., 2015), and the secreted secondary metabolite gliotoxin has been shown to exert a directly cytotoxic and immune-modulatory effect on several host cells, including airway epithelial cells (Scharf *et al*., 2012; Zhang *et al*., 2019). However, strains lacking certain protease activities or deficit in gliotoxin biosynthesis still retain virulence *in vivo*, (Bok *et al*., 2006; Sharon, Hagag and Osherov, 2009; Cacho Teixeira *et al*., 2017) suggesting that the coordinated activities of multiple fungal attributes combine to elicit fatal tissue damage in the host. Supporting this theory, Bertuzzi et al (2014) revealed that *A. fumigatus* mutants lacking the pH-responsive transcription factor PacC fail to activate the expression of a multitude of secreted proteases and secondary metabolites during host colonisation. *In vitro* dissection of this phenotype revealed a 50% reduction in detachment of A549 cells from cultured monolayers at an early stage of infection with a *ΔpacC* strain, as well as a reduction in epithelial cell lysis occurring at a later timepoint. Moreover, the mutants were unable to penetrate the lung epithelium and demonstrated reduced virulence in a murine model of aspergillosis, in comparison to the parental strain (Bertuzzi *et al*., 2014).

By negating the effects of redundancy of gene products involved in host-pathogen interactions of invasive fungi in the establishment of infection, transcription factor (TF) mutants have proven to be powerful tools for resolving the complexity of host-pathogen interactions that drive fungal diseases of humans (Nobile *et al*., 2012; Jung *et al*., 2015). Although multiple *A. fumigatus* TFs regulating fungal physiological and growth processes have been implicated in *in vivo* virulence (Bultman, Kowalski and Cramer, 2017), no high throughput or genome wide studies on epithelial damage or host-pathogen interaction employing transcription factors have been previously carried out in the *Aspergillus* genus. The recently generated library of *A. fumigatus* transcription factor mutants (TFKOs) by (Furukawa *et al*., 2020) has provided a methodological toolkit that allowed, for the first time, a genome-scale census of *A. fumigatus* TFs driving epithelial damage. This study sought to identify the transcriptional regulators that are required for causation of epithelial damage and via phenotypic analyses to identify the underlying causal events during the host-pathogen interaction.

## Material and Methods

### *A. fumigatus* strains and growth conditions

A collection of 479 *A. fumigatus* transcription factor knock-out mutants (Furukawa et al, 2020) constructed in the A1160+ genetic background (Fraczek *et al*., 2013; Bertuzzi *et al*., 2021) was used in this study. The TFKO strains were stored at −80 °C and cultured on Aspergillus Complete Media agar (ACM)(Pontecorvo *et al*., 1953), pH 6.5 containing Hygromycin B (Alfa Aesar, UK) at a concentration of 100 μg/ml for selective growth. The fungus was cultured on ACM at 37 °C for 3-5 days and the spores were harvested in sterile distilled water and the spore suspensions were filtered through sterile miracloth (Calbiochem, UK) to remove mycelial fragments. Following two washes of the spore suspension with sterile water, the spores/ml were enumerated using OD_600_ measurement (Furukawa et al, 2020). Accuracy of spore counts in infecting inocula was assessed via serial dilution and enumeration of viable fungal colonies following culture on ACM solid agar.

### Culture and maintenance of epithelial cells

Commercially sourced human carcinomic alveolar basal epithelial A549 cells were used in this study (American type culture collection, CCL-185). A549 cells were routinely cultivated and maintained in RPMI-1640 medium with L-glutamine, (Sigma-Aldrich) supplemented with 10% fetal bovine serum (Gibco, UK) and 1% penicillin-streptomycin (Sigma-Aldrich) at 37ºC with 5% CO_2_. Cells were seeded at 5×10^4^/ml for a 24 well plate and at 7.5×10^4^/ml for a 96 well plate and incubated for 2 days at 37ºC and 5% CO_2_ for a 90-100% confluent A549 monolayer. On infection day, media was replaced, and all infections were performed in supplemented RPMI-1640 medium (as above, with additional trace elements (Na_2_B4O_7_.10H_2_O (0.04 mg/l), CuSO_4_. 5H_2_O (0.4 mg/l), FeCl_3_.6H_2_O (1.16 mg/l), MnSO_4_. 2H_2_O (0.8 mg/l), Na_2_MoO_4_. 2H_2_O (0.8 mg/l), ZnSO_4_. 7H_2_O (8.0 mg/l)) at 37°C and 5% CO_2_.

### High-throughput screens for epithelial damage

The parental strain (A1160p+) and the PBS challenge were included as controls for each infection in both the detachment and cell lysis assays. To ensure the measurement of damage from the high-throughput screens was not an artefact of inoculum size, A549 cells were challenged with a 10 times serial dilution of spores of the *A. fumigatus* parental strain (A1160p+) to calculate dose-dependency of epithelial damage phenotypes. According to standard curves generated by plotting viable spore density versus epithelial detachment or lysis, the phenotypic data were retrospectively adjusted for small deviances from desired inoculum sizes.

### Detachment assay

As detailed in Rahman et al, 2021, A549 monolayers were cultured to confluence in 96 well glass bottom plates (Greiner Bio-one) and challenged with 20 μl of a 10^7^ spores/ml suspension (200,000 spores per well), with five technical replicates (one technical replicate per plate). After 16 hours, the A549 monolayer was washed once with pre-warmed phosphate buffered saline (PBS) (Thermo-Fisher Scientific) to remove detached A549 cells. This was followed by fixing the remaining adherent A549 cells in the wells with 4% formaldehyde in PBS for 10 min (Alfa Aesar, UK), permeabilizing the cells with 0.2% Triton-X100 (VWR) for 2 min and staining nuclei of the adherent A549 cells with DAPI (Alfa Aesar) at 300 nM for 5 min, protected from direct light (Rahman *et al*, 2021). DAPI fluorescence was excited with a 405nm LED and its emission was captured on a HyD detector (405-600 nm). Image capture was performed in high throughput via automated confocal microscopy (Leica SP8x; Leica Microsystems, Germany) from 9 fields of view per well at 40x magnification using a HC PL APOCS 40x 0.85 DRY objective. The number of adherent A549 cells in each image was quantified using a cell segmentation macro written for FIJI (Rahman *et al*, 2021). Viable counts of inocula used for infection were determined by plating 10^3^ spores on ACM agar for 48 hr. The difference between the observed and predicted counts was calculated for each mutant.

### Cell lysis assay

Fully confluent A549 cells grown in 24 well plates (Greiner Bio-one) were challenged with 50 μl of 10^7^ spores/ml suspension (500,000 spores per well) with at least 3 technical replicates in one plate. For fungal culture filtrates a 10^6^ spores/ml culture of RPMI was incubated at 37 ºC and 5% CO_2_ for 48 hours and then filtered through five layers of Miracloth (Calbiochem, UK) and then a 0.45 μm syringe filter to remove any spores and hyphal fragments. Elimination of all live spores from the culture filtrate was confirmed by plating of the filtrates on solid ACM agar for up to 48 hours. Fully confluent A549 monolayers in 24 well plates were challenged with a 1:5 dilution (in supplemented RPMI) of the filtered culture filtrates for 24 hours. Following incubation with *A. fumigatus* spores or culture filtrates for 24 hours, cell culture supernatants were collected to measure LDH released by epithelial cells on lysis, using the Cytox 96 non-radioactive cytotoxicity assay kit (Promega), as per manufacturer’s instructions. The LDH assay was carried out in 96 or 384 well plate format and a recombinant porcine LDH enzyme (Sigma-Aldrich) was used in each LDH assay plate in a serial dilution to extrapolate a standard curve which was used to estimate LDH released on infection with *A. fumigatus*. Viable counts of inocula used for infection were determined by plating 100 spores on solid ACM agar for up to 48 hours.

### Internalization of spores by epithelial cells

*A. fumigatus* spores were stained with Fluorescein isothiocyanate (FITC, Sigma) for 30 min at 37 °C while shaking. After three washes with PBS, 3×10^5^ spores were added onto fully confluent A549 monolayers in 24 well glass bottom plates. Following 4 hours of infection, the wells were washed two times with PBS to remove non-adherent spores, and the non-internalised spores were stained with 0.1 mg/ml of Calcofluor White in PBS (CFW; Sigma) for 5 min at 37 °C. The wells were washed again twice with PBS, fixed for 15 min at 37 °C using 5% formaldehyde and stored in PBS protected from light. Fluorescence images from 9 fields of view were captured in each well using a 1.5 pin hole and a 40 x objective of a Leica SP8x confocal microscope (Leica Microsystems, Germany). External CFW stained spore fluorescence was excited with a 405nm LED and collected on a HyD detector (410-450 nm) and all spore FITC fluorescence was excited with a 488nm Argon laser and collected on a HyD detector (495-570nm). The number of external and internal spores within the total population in each field of view of pixel size (388.26 μm x 388.26 μm) were counted manually in FIJI (Schindelin *et al*., 2012) for all strains and expressed as % spores internalised. The % uptake relative to the parental strain was calculated of each strain.

### Germination efficiency and hyphal extension

Fungal growth was assessed by time-lapse transmitted light imaging (Leica SP8; Leica Microsystems, Germany) of strains cultured in supplemented RPMI (as described above) at 37 °C. Approximately 5×10^4^ spores were inoculated per well of a 24 well glass bottom plate (Greiner Bio-one) and allowed to settle for 45-60 min at 37 °C. Transmitted light images were captured using a 10x/0.4NA lens and a 514nm argon laser, in each well of the plate every hour up to 48 hours. Approximately 100 spores were captured in each image. The total number of germinated spores were counted at every time point using FIJI and the germination rate was calculated. The lengths of individual hyphae were measured at each time-point starting from germination, by using the segmented line measurement tool in FIJI (Schindelin *et al*., 2012). Hyphae were measured up until the moment they grew out of the optical plane, of approximately 16-18 hours for the parental strain.

### Adhesion to epithelial cells

Germlings were generated by growth of 5×10^4^ spores/ml in supplemented RPMI (as described above) for 7 hours at 37 °C and added to fully confluent A549 cells in a 24-well glass bottom plate (Greiner Bio-one). Germlings for the TF mutants with a delay in germination were grown for a longer length of time until they achieve approximately similar visual hyphal growths as the parental strain. Following 30 min of incubation, non-adherent germlings were removed by a PBS rinse and the remaining adherent germlings were stained with 10 μg/ml of CFW for 5 min, prior to fixation with 4% formaldehyde. A 1441 μm^2^ field of view was captured in each well using a 10x lens objective for CFW germling fluorescence using a confocal microscope (Leica SP8x; Leica Microsystems, Germany). Fluorescence was excited with a 405nm LED and collected on a HyD detector (410-450nm). The number of adherent germlings remaining in each well were counted manually in FIJI (Schindelin *et al*., 2012).

### Cell wall sensitivity assay

Sensitivity to the cell wall destabilising agent, CFW, was tested by inoculating serial spore concentrations (10^6^, 10^5^, 10^4^, 10^3^) onto solid aspergillus minimal media agar (Bertuzzi *et al*., 2014) with CFW (200 ug/ml).

### Statistics

Log_2_ transformed LDH data were filtered for assay plate-dependent batch effects and analyzed using a Bayesian modelling approach (Kruschke, 2013) that describes data using a Student’s t distribution and performs posterior inference by Markov chain Monte Carlo sampling. We compared the posterior distributions for each TFKO mutant to the corresponding parental strain of each assay plate. Significant difference between the two means, based on a 95% Bayesian credible interval, was deemed “significant” and is denoted by TRUE in Table S3. Data are presented as the mean value ± standard error of mean (SEM). Other statistical tests were performed using Prism 9.00 software (GraphPad Software, La Jolla, CA). Quantitative differences between groups were tested using ANOVA one way or two-way analysis. To correct for multiple testing, Fishers LSD t-test or Dunnett’s test (comparisons of selected groups to the parental strain) was applied; p < 0.01 was considered statistically significant for these tests.

## Results

### Distinct cohorts of transcriptional regulators drive early and late phases of *A. fumigatus*-mediated epithelial damage

In order to identify transcriptional regulators required for epithelial damage during *A. fumigatus* infection 479 *A. fumigatus* knock-out mutants (TFKOs), each lacking an individual transcription factor-encoding gene (Furukawa *et al*., 2020), were incubated with A549 alveolar epithelial monolayers. Damage was measured at 16 hours and 24 hours of infection, respectively using high throughput quantitation of epithelial cell detachment and epithelial cell lysis (Bertuzzi *et al*., 2014; Okaa *et al*., 2021; Rahman, Thomson and Bertuzzi, 2021) as indicators of host cell damage. Assay formats were optimised for high throughput use by assessing damage caused by *A. fumigatus* A1160p+ versus that of a Δ*pacC* null mutant which is deficient in causing both early and late stages of the host damage (**Fig S1)**.

Detachment of epithelial cells was quantified following a 16-hour infection of A549 monolayers with *A. fumigatus* spores (**Fig 1A**). Of the 479 TFKO mutants initially screened (**Table S1**), 18 and 28 TFKOs exhibited an increased and decreased capacity, respectively, to cause detachment of epithelial cells during a 16-hour infection (**Fig 1B**). For the purposes of this study, our interest focused upon those TFKOs demonstrating reduced epithelial detachment. In a follow-up low throughput assay, 10 of the 18 originally identified TFKOs that were identified as being defective in eliciting epithelial cell detachment exhibited reproducible phenotypes (**Fig 1C and Table 1**). Among these, 6 TFs have been previously characterised, including the pH-responsive transcription factor PacC (AFUB_037210) previously reported to regulate *A. fumigatus* epithelial invasion (Bertuzzi *et al*., 2014), CreA (AFUB_027530) the broad domain regulator of carbon catabolism (Beattie *et al*., 2017), AreA (AFUB_096370) the broad domain regulator of nitrogen metabolism (Hensel *et al*., 1998; Krappmann and Braus, 2005), a C6 zinc cluster family TF (AFUB_033930) located in the uncharacterised small peptide secondary metabolite gene cluster 12 (Bignell *et al*., 2016) and two TFs NsdD and NsdC (AFUB_035330 and AFUB_089440) that repress the asexual sporulation programme in *Aspergillus* species (Wu *et al*., 2018; de Castro *et al*., 2021).

**Figure 1.**
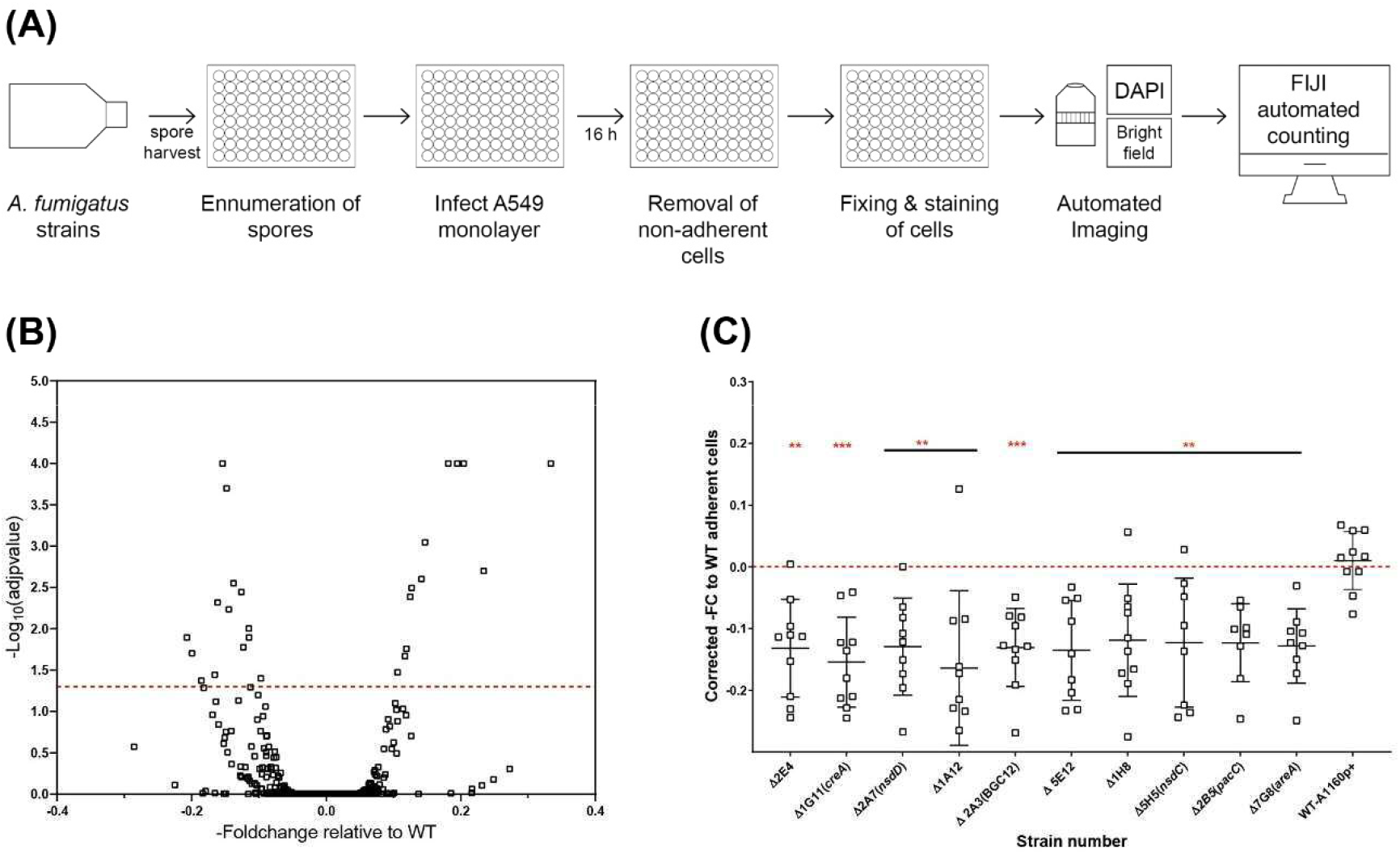
High throughput screening identifies *A. fumigatus* transcription factors required for pneumocyte detachment. A) Experimental design to analyse detachment of epithelial cells in high throughput. Harvested spores of *A. fumigatus* TFKOs were counted using OD600 measurement and incubated with A549 cell monolayers for 16 hours, followed by automated imaging and quantification of remaining number of A549 cells in the monolayer to determine epithelial cell damage via detachment of cells. **B) Volcano plot showing the output from the detachment screen**. Data is transformed as -fold change number of adherent cells for 479 TF mutants. The cut off (dotted line) signifies p value<0.1 from the Dunnett’s multiple comparison of one-way ANOVA relative to the parental strain **C) Fold change number of adherent cells, relative to the parental strain for the 10 cell detachment mutants**. The 10 cell detachment mutants showing reproducibility of phenotypes when assay was conducted in low throughput. Data shown is technical replicates of two biological replicates. Error bars show ±SEM. Data was analysed by one-way ANOVA with Dunnett’s multiple comparisons test relative to the parental strain (WT-A1160p+). *P<0.1, **P<0.01 ***P<0.001, ****P<0.0001.

**TABLE 1:** List of cell detachment and cell lysis *A. fumigatus* regulators. Annotation of the identified epithelial damage TFKOs with the gene name, TF number, generic name, main function and references.

| TABLE 1: List of cell detachment and cell lysis *A. fumigatus* regulators |  |  |  |  |
| --- | --- | --- | --- | --- |
| CELL DETACHMENT TFKOs |  |  |  |  |
| AFUB number | AFUA number | TF number /<br>Generic name | Main Function | References |
| AFUB_046160 | AFUA_3G02210 | 2E4 | Unknown | ~ |
| AFUB_027530 | AFUA_2G11780 | 1G11 (*creA*) | Hypoxic adaptation, carbon<br>catabolite repression | Beattie et al. 2017 |
| AFUB_019830 | AFUA_2G02730 | 5E12 | Unknown | ~ |
| AFUB_035330 | AFUA_3G13870 | 2A7 (*nsdD*) | Early stage of mating | Castro et al. 2021 |
| AFUB_037210 | AFUA_3G11970 | 2B5 (*pacC*) | pH response, epithelial<br>invasion | Bertuzzi et al. 2014 |
| AFUB_031000 | AFUA_2G15340 | 1H8 | Unknown | ~ |
| AFUB_096370 | AFUA_6G01970 | 7G8 (*areA*) | Nitrogen utilization | Krappmann & Braus, 2005 |
| AFUB_033930 | AFUA_3G15290 | 2A3 (BGC12) | Secondary metabolite<br>production | Elaine et al. 2016 |
| AFUB_089440 | AFUA_7G03910 | 5H5 (*nsdC*) | Early stage of mating | Castro et al. 2021 |
| AFUB_005510 | AFUA_1G05150 | 1A12 | Unknown | ~ |
| CELL LYSIS TFKOs |  |  |  |  |
| AFUB number | AFUA number | TF number /<br>Generic name | Main Function | References |
| AFUB_052420 | AFUA_5G03920 | 2G3 (*hapX*) | Iron homeostasis | Gsaller et al. 2014 |
| AFUB_033470 | AFUA_2G17800 | 1H12 | Unknown | ~ |
| AFUB_031980 | AFUA_2G16310 | 1H10 | Unknown | ~ |
| AFUB_041100 | AFUA_3G08010 | 2C7 (*sltA*) | Conidial formation, cell wall<br>architecture | Liu et al. 2021 |
| AFUB_078150 | AFUA_6G12150 | 3E3 (*atfD*) | Stress response to conidia | Silva et al. 2021 |
| AFUB_078520 | AFUA_6G12522 | 3E5 | Unknown | ~ |
| AFUB_037210 | AFUA_3G11970 | 2B5 (*pacC*) | pH response, epithelial<br>invasion | Bertuzzi et al. 2014 |
| AFUB_015750 | AFUA_1G16410 | 1D11 | Unknown | ~ |

Epithelial cell lysis was measured by infecting fully confluent A549 alveolar monolayers with spores from the TFKO null mutants for 24 hours, after which LDH released by the epithelial cells lysed because of hyphal mediated damage was quantified (**Fig 2A**). *A. fumigatus* was not found to secrete endogenous LDH enzyme at this time-point, therefore confirming the sole source of LDH as being epithelial cells (**Fig S1**). The quantification of LDH was performed via a standard curve generated from a serial dilution of the standard LDH enzyme for each assay plate. A Bayesian approach for calculation of the posterior probability that mutants deviate from the wild type was applied to overcome batch-related variations in LDH quantitation (**Table S2**). After filtering for plate and batch effects, 34 TFKOs caused a decreased LDH release by the A549 epithelial monolayers after a 24-hour infection (**Fig 2B** and **Table S1**). Of these 34, only 8 TFs demonstrated reproducible reductions in LDH activity (**Fig 2C and Table 1**). Among these 8 TFKOs, only 4 TF-encoding genes have been previously characterised. This includes HapX (AFUB_052420), a component of the CCAAT-binding complex (Hortschansky *et al*., 2017) which is involved in iron acquisition and metabolism in *Aspergillus* (Gsaller *et al*., 2014), PacC (AFUB_037210) previously reported to regulate *A. fumigatus* epithelial invasion (Bertuzzi *et al*., 2014), Ace1/SltA (AFUB_041100) which regulates conidial formation including cell wall architecture and secretion of mycotoxins and secondary metabolites (H. Liu *et al*., 2021) and AtfD (AFUB_078150) which is involved in conidial stress responses (Silva, Horta and Goldman, 2021). PacC (Bertuzzi *et al*., 2014) was found to be the only transcription factor that is required for causation of both epithelial cell detachment and cell lysis (**Table 1)**. Conclusively, our analysis identified distinct cohorts of transcriptional regulators involved in the early and late causation of epithelial damage.

**Figure 2.**
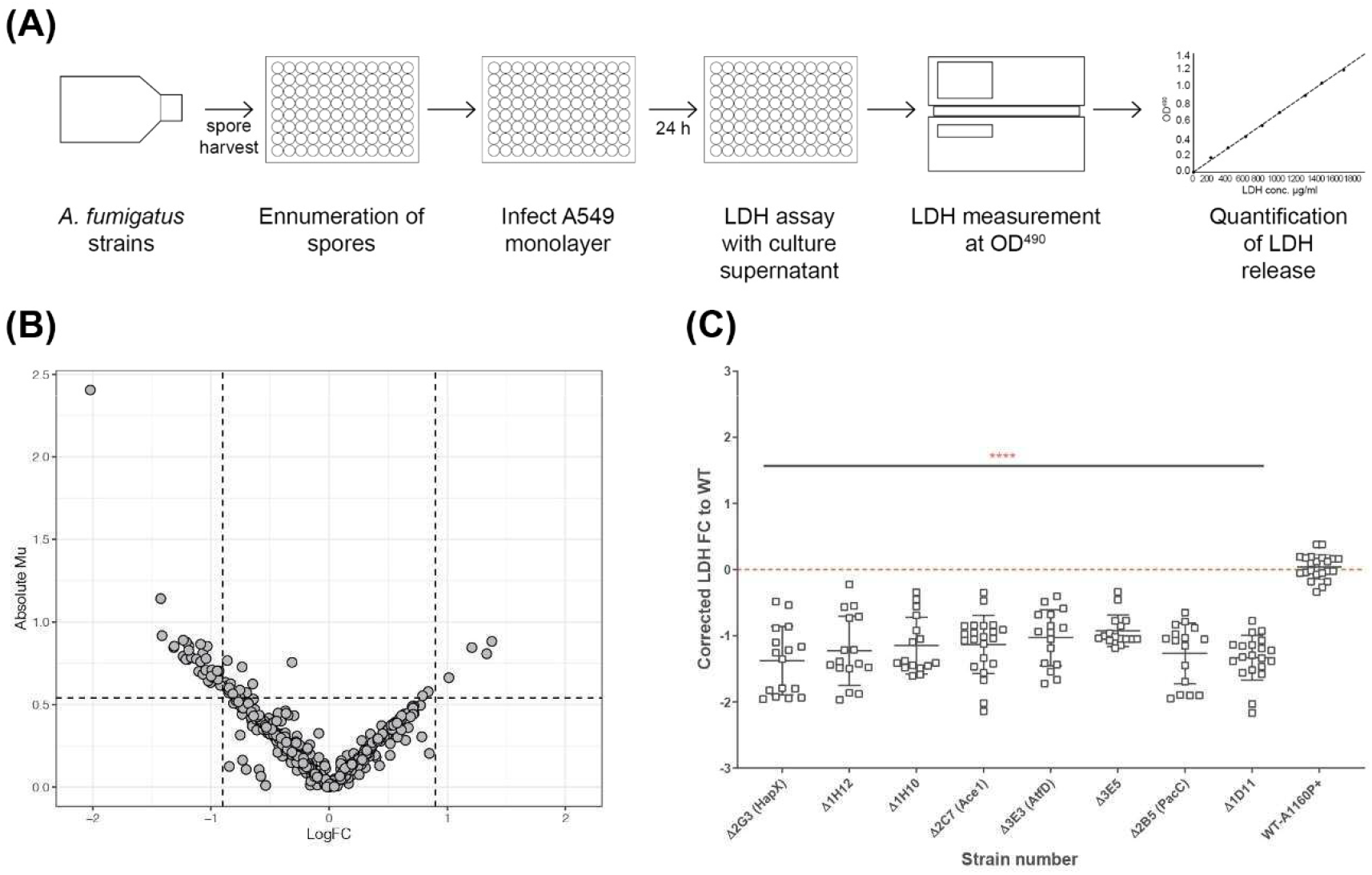
High throughput screening identifies *A. fumigatus* transcription factors required for cytolytic death of pneumocytes. **A**) Experimental design to analyse cell lysis of epithelial monolayers in high throughput using LDH release assay. Harvested spores of *A. fumigatus* TF mutants were counted using OD600 measurement and incubated with A549 cell monolayers for 24 hours, followed by measurement of LDH in the supernatant to determine epithelial cell damage via cell lysis. **B) Volcano plot showing the output from the cell lysis screen**. Data is filtered to remove plate/batch errors and shows the Absolute Mu (derived from Bayesian Modelling) versus log Fold change for each mutant relative to the parental strain. The cut-offs for the plot were based upon the average fold change and the average Mu difference between the controls (parental strain and the *ΔpacC*) throughout the screen. **C) LDH fold change relative to the parental strain for the 8 cell lysis mutants**. The 8 cell lysis mutants showing reproducibility of phenotypes when assay conducted in low throughput. Data shown is technical replicates of at least two biological replicates. Error bars show ±SEM. Data was analysed by one-way ANOVA with Dunnett’s multiple comparisons test relative to the parental strain (WT-A1160p+). ****P<0.0001.

### Hyphal growth rates distinguish between *A. fumigatus* TFKO mutants deficient in epithelial detachment and cell lysis

An obvious explanation for the observed deficits in epithelial detachment and cytolysis might be reduced fitness of the respective TFKOs. At early time points of host interaction this might derive from germination defects, at later time points from reductions in hyphal growth rates, or both. To explore this possibility, the germination efficiencies and hyphal growth rates of the TFKOs were measured at hourly intervals using time-lapse confocal imaging. Of the 17 mutants identified as having significant deficits in epithelial damage, four (*ΔpacC, ΔcreA, ΔhapX*, and Δ*sltA*) exhibited at least 50% reductions in germination and/or hyphal growth rates relative to the parental strain (**Fig 3 and Fig S2**). Enumeration of spore germination in the parental isolate A1160p+ revealed that spore germination commenced at 4 hours reaching 100% by 10-11 hours. Amongst the TFKOs causing reduced epithelial cell detachment (**Table 1)**, most exhibited similar germination efficiencies to that of the parental strain (**Fig S2 A, C**). Notable exceptions are the *ΔpacC* and *ΔcreA* strains that exhibited 70% germination at 11 hours (**Fig S2**), achieving 100% germination by 16 hours and the *ΔhapX*, and Δ*sltA* mutants where only 20-30% of spores had germinated at 11 hours (**Fig 3**). Even after 20 hours, germination efficiencies fell well short of 100% for both the *ΔhapX*, and Δ*sltA* mutants.

**Figure 3.**
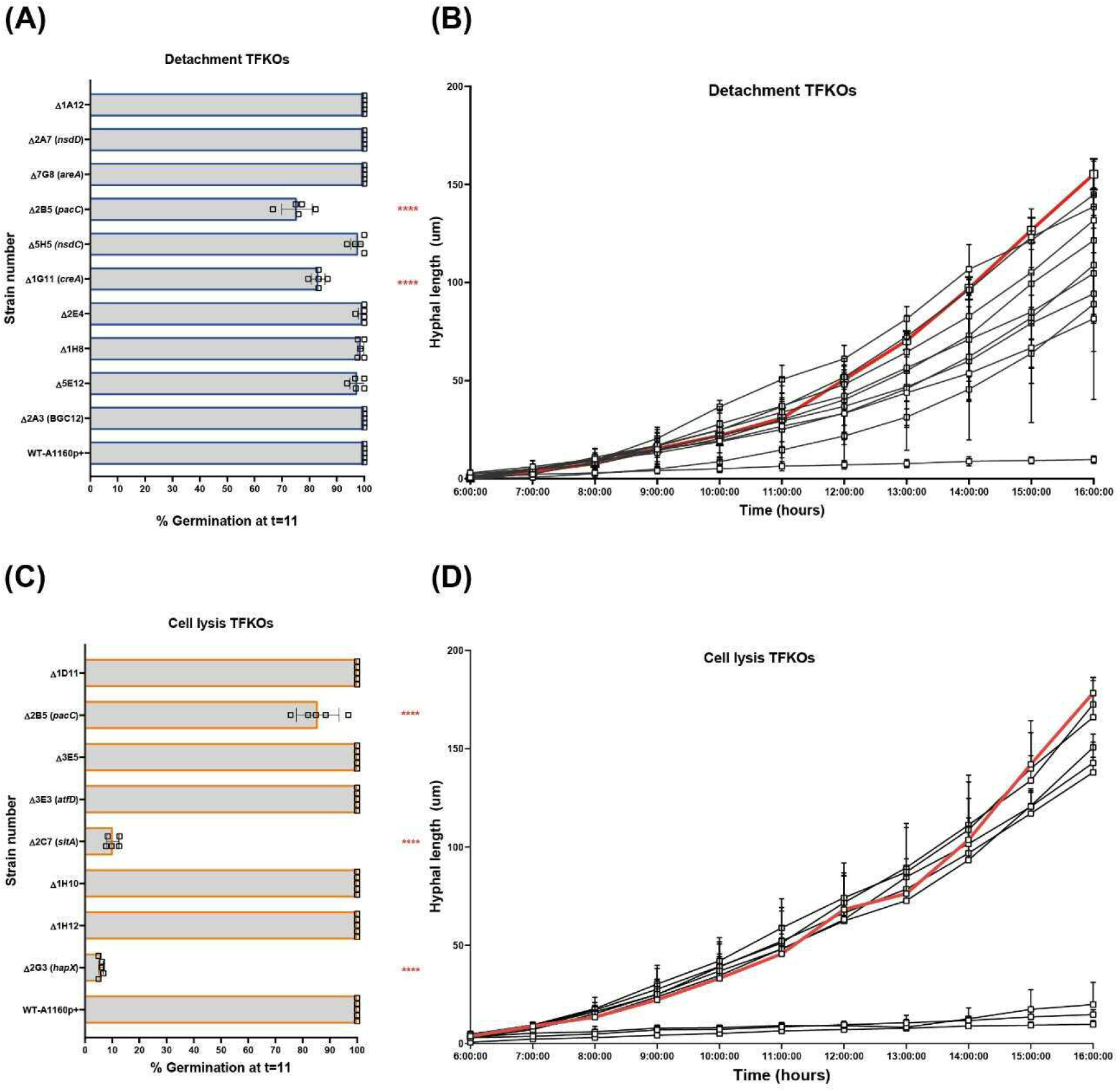
Percentage germination and hyphal lengths of *A. fumigatus* TFKOs measured by assessing growth using time-lapse confocal microscopy. Mutants of the cell damage TFKOs were grown in supplemented RPMI-1640 for up to 24 hours. Imaged were captured every hour at 40x magnification. **A) Percentage germination at 11 hours of growth for the cell detachment TFKOs**. Percentage germination was calculated by enumerating total germinated spores in at least three fields of view **B)** Hyphal lengths over time for cell detachment TFKOs. Hyphae were measured from the images at each hour, using FIJI. **B) Percentage germination at 11 hours of growth for the cell lysis TFKOs**. Percentage germination was calculated by enumerating total germinated spores in at least three fields of view **D) Hyphal lengths over time for cell lysis TFKOs**. Hyphae were measured from the images at each hour, using FIJI. The WT parental strain is highlighted in red. Error bars show ±SEM. Data was analysed by one-way ANOVA with uncorrected Fisher’s LSD multiple comparisons test relative to the parental strain (WT-A1160p+). *P<0.1, **P<0.01 ***P<0.001, ****P<0.0001.

The parental strain achieved hyphal lengths of ~ 150 μm by 16 hours of growth in supplemented RPMI, representing a mean rate of hyphal extension of 20 μm hour of growth (**Fig S2**). Hyphal extension rates of TFKOs defective in epithelial cell detachment were reduced with 8 of the 10 TFKOs exhibiting hyphal extension rates of 15-20 μm/hour or less (**Figs 3 and S2)**. Consistent with previous reports of hyper-branching growth (Bertuzzi *et al*., 2014), the *ΔpacC* strain achieved less than 20 μm in hyphal length by 16 hours. In summary, analyses of germination and hyphal growth rates revealed that the spores of most mutants germinated as efficiently as those of the parental strain, but hyphal extension rates frequently differed from those of the parental isolate. Categorically, we find that the phenotype that associates most robustly with reductions in epithelial detachment is that of reduced hyphal extension rates.

### Deficits in epithelial uptake and adhesion are common amongst *A. fumigatus* TFKOs defective in epithelial detachment and cytolysis

The Dectin-1 ligand, β-1,3-glucan, that becomes exposed upon the surface of swollen *A. fumigatus* conidia has been demonstrated to activate host-derived phospholipase D, a critical driver of actin-mediated endocytosis (Han *et al*., 2011). In cultured monolayers of immortalised human pneumocytes, epithelial cell detachment is mitigated by antibody-mediated blockade of the Dectin-1 receptor suggesting that actin-mediated uptake of fungal spores is an important driver of epithelial cell detachment (Bertuzzi *et al*., 2014). To assess the contribution of spore internalisation on epithelial cell detachment, the uptake of TFKO spores was quantified via by differential fluorescence imaging following incubation of FITC-stained spores with A549 cells from 30 minutes to 6 hours, followed by CFW staining of externally adherent spores (**Fig S3**). The proportion of infecting A1160p+ spores internalised by A549 cells after a 4-hour infection was 20% (**Fig 4B, C**). Of the 10 TFKOs causing reduced epithelial detachment, five (*ΔpacC, ΔnsdD, ΔnsdC*, ΔAFUB_031000-Δ1H8 and ΔAFUB_019830-Δ5 E12) showed half the spore uptake relative to the parental isolate. Of the 8 TFKOs defective for epithelial lysis, 6 mutants (*ΔpacC, ΔhapX*, Δ*sltA, ΔatfD*, ΔAFUB_015750-Δ1D11, ΔAFUB_078520-Δ3 E5) demonstrated significantly reduced rates of spore uptake relative to the parental strain. Of these, spores of the *ΔpacC, ΔhapX*, and Δ*sltA* mutants exhibited ≥ 50% reduction in epithelial uptake relative to the parental isolate.

**Figure 4.**
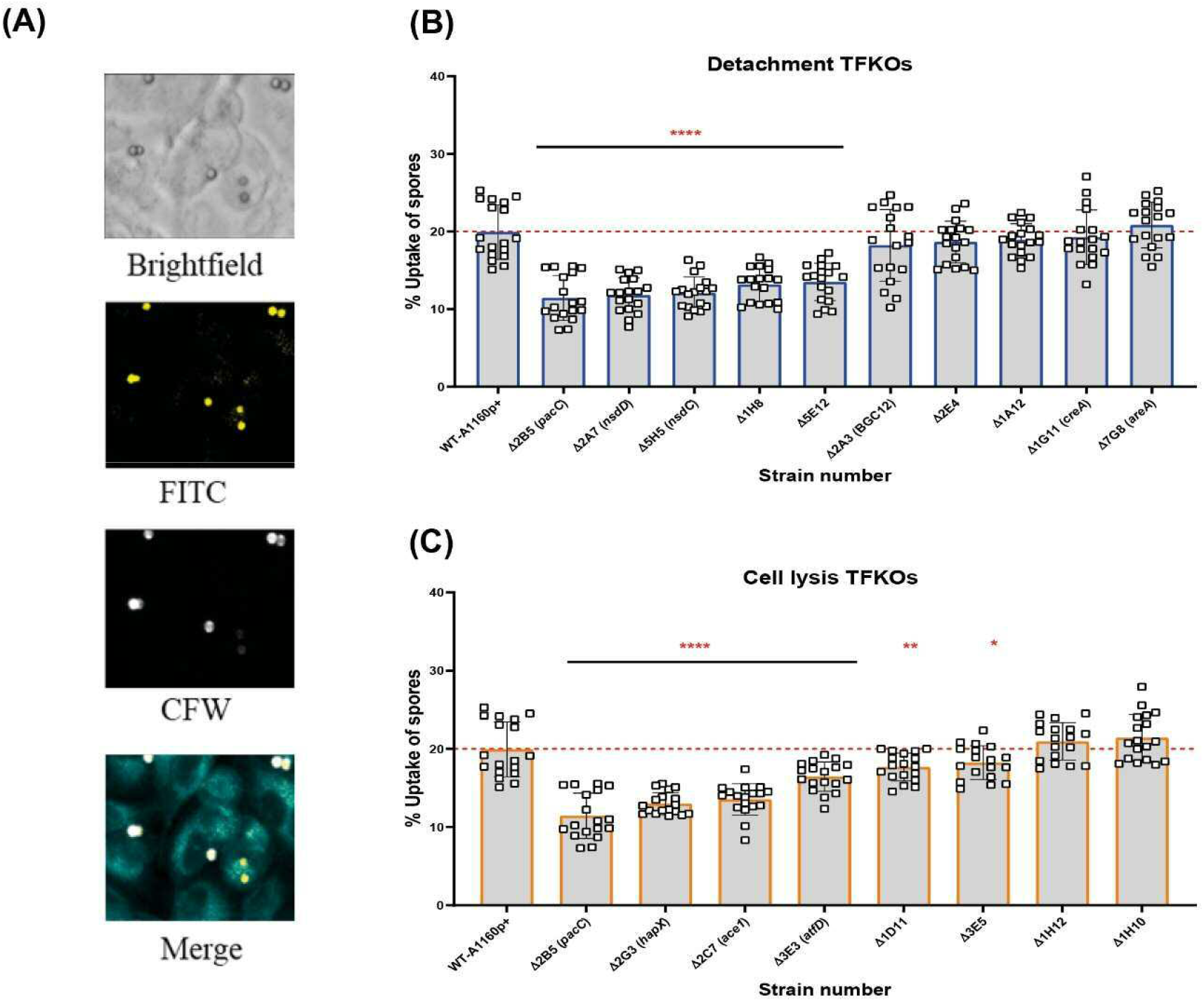
Uptake capacity of A549 cells to cell detachment and cell lysis *A. fumigatus* TFKO spores. FITC stained spores incubated with A549 cells for 4 hours to enable internalisation were washed with PBS to stain with calcofluor white to visualise the internal and external spores **A) Representative images of brightfield, FITC (all spores), calcofluor white (external adherent spores)**. Images were captured using the TCS-SP8 confocal microscope at 40x magnification **B) Percentage spore uptake for the cell detachment TFKOs. C) Percentage spore uptake for the cell lysis TFKOs**. WT=parental strain A1160p+. Error bars show ±SEM. Data was analysed by one-way ANOVA with Fisher’s LSD multiple comparisons test. UI=Un-infected. *P<0.1, **P<0.01 ***P<0.001, ****P<0.0001. Data shown is technical replicates of two biological replicates.

Adhesion of *A. fumigatus* to host pneumocytes is thought to prevent easy removal of *A. fumigatus* from the mucosal surfaces and provide a close contact for manipulation and invasion of host cells (Sheppard, 2011; Brunke *et al*., 2016). *In vitro* observations report *A. fumigatus* morphotypes adhere to epithelial cells, from as early as 30 minutes post-infection (DeHart *et al*., 1997). The expression of a hyphal exopolysaccharide, galactosaminogalactan (GAG) has been demonstrated as critical for hyphal adherence to epithelial cells. Moreover, a null mutant of a transcriptional regulator for GAG synthesis, MedA, is hypovirulent in a murine models of IA (Gravelat *et al*., 2010) implying an important role for adherence to host cells during infection of the mammalian lung. To address the relevance of reduced hyphal adherence in TFKOs causing reduced epithelial damage, germlings of the 17 TFKOs of interest were incubated with A549 cells for 30 min, after which the non-adhered germlings were removed by rinsing and those remaining were CFW stained for visualisation and enumeration of adherence by differential fluorescence microscopy (**Fig 5A**). To obviate the confounding effects of fitness deficits, TFKOs defective in germination and/or hyphal elongation were cultured for longer durations to achieve similar morphogenic states to the parental strain. Amongst the 10 TFKOs defective in causing epithelial detachment, 6 mutants (*ΔpacC, ΔcreA, ΔareA*, ΔAFUB_005510-Δ1A12, *ΔnsdC*, and ΔAFUB_019830- Δ5E12) also exhibited significantly reduced adherence of germlings to epithelial cells, compared to the parental strain, with *ΔcreA* and *ΔpacC* strains exhibiting a a 50% reduction in adherence relative to the parental strain (**Fig 5B**). Similarly, 5 out of the 8 mutants (*ΔpacC*, ΔAFUB_031980-Δ1H10, Δ*sltA*, ΔAFUB_015750-Δ1D11, ΔAFUB_078520-Δ3 E5) mutants defective in epithelial lysis displayed significantly reduced adherence compared to the parental strain (**Fig 5C**).

**Figure 5.**
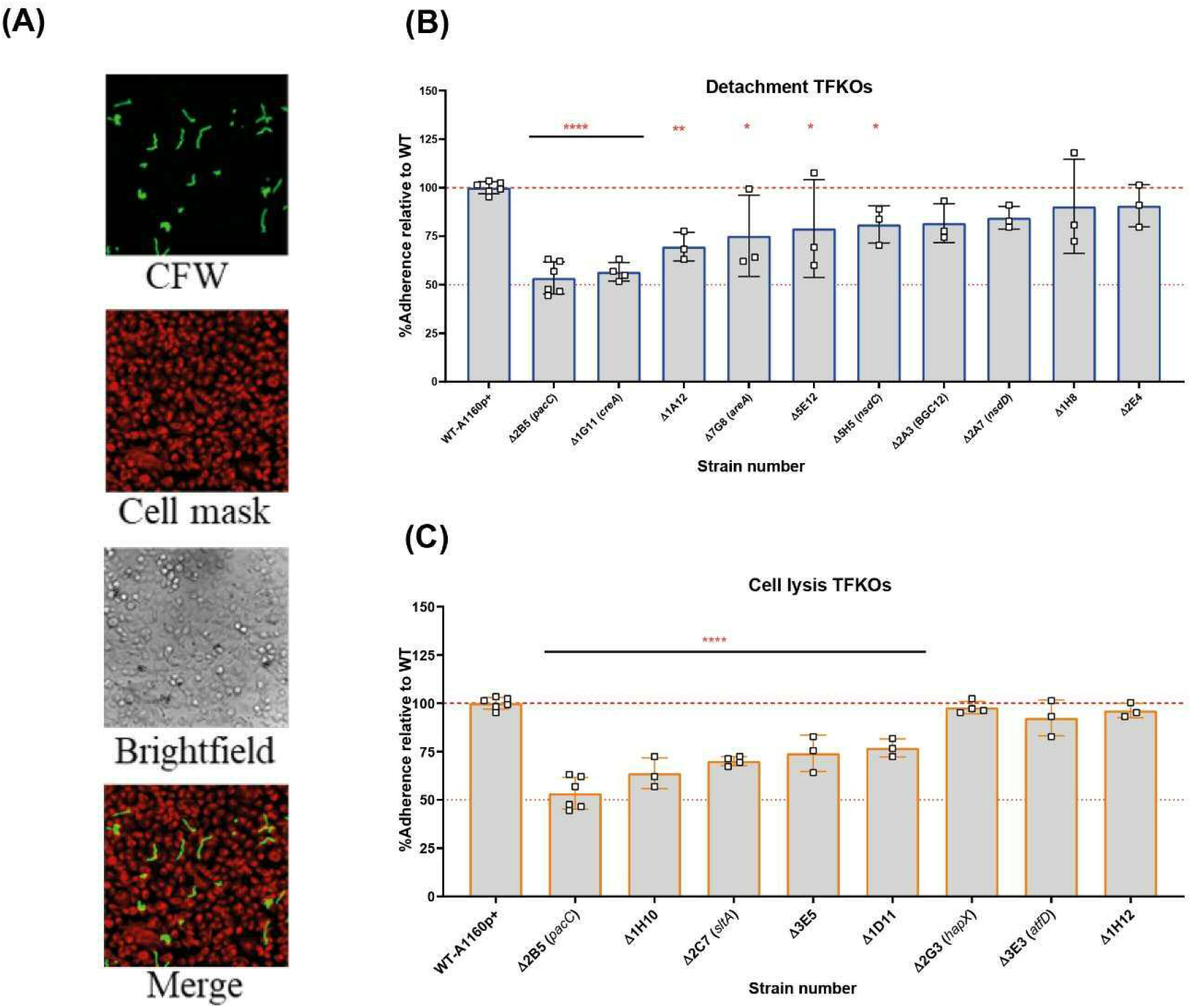
Adhesion capacity of the hyphal of *A. fumigatus* TFKOs to epithelial cells. Germlings of TFKOs incubated with A549 cells for 30 min to enable adhesion were rinsed with PBS and stained with calcofluor white to visualise only the adherent germlings **A) Representative images of calcofluor white stained germlings, cell mask stained A549 cells, brightfield and a merged image**. Images captured with TCS-SP8 confocal microscope at 40x magnification **B) Percentage adherence of the cell detachment TFKOs C) Percentage adherence of the cell lysis TFKOs**. Error bars show ±SEM. Data was analysed by one-way ANOVA with Fisher’s LSD multiple comparisons test. Data shown is technical reps of 3 biological reps. WT=A1160p+. *P<0.1, **P<0.01 ***P<0.001, ****P<0.000.

### Calcofluor white sensitivity associates predominantly with *A. fumigatus* TFKOs defective in causation of epithelial damage

The cell wall of *A. fumigatus* is the outermost cellular structure that mediates host interactions including pathogen recognition and stimulation of the host immune response and pathogen adherence to host cells (Lee and Sheppard, 2016). Therefore, changes in cell wall structure and/or composition often affect pathogenicity. Previous studies have demonstrated that *A. fumigatus* spores and germlings are internalised by epithelial cells in a contact-, actin-, cell wall- and Dectin-1 dependent manner and *ΔpacC* mutants, which aberrantly remodel the cell wall during germinative growth, are less able than wild type counterparts to gain entry into epithelial cells (Bertuzzi *et al*., 2018). In fungal cells suffering cell wall aberrancies that involve chitin redistribution, sensitivity to the anionic dye Calcofluor white (CFW) that binds to chitin and interferes with the construction and stress tolerance of the cell wall, is a common phenotype (Roncero *et al*., 1988). To test the hypothesis that alterations in fungal cell wall contribute to differential damage in epithelial cells, the 17 TFKOs of interest were tested for sensitivity/resistance to CFW by growing serial dilutions (10^7^,10^6^,10^5^,10^4^) spores/ml of the mutants on solid AMM containing 200 μg/ml of CFW. Amongst the 10 mutants inducing reduced epithelial detachment (**Fig 6**), 9 TFKOs exhibited sensitivity to CFW relative to the parental strain. Furthermore, 6 of the 8 TFKOs isolates effecting reduced cytotoxicity to epithelial cells (**Fig 6**) exhibited sensitivity to CFW. The Δ*sltA* strain was resistant to CFW relative to the parental strain (**Fig 6**), as reported previously (Liu *et al*., 2021). Our findings demonstrate a robust association between cell wall aberrances and inability to cause epithelial damage.

**Figure 6.**
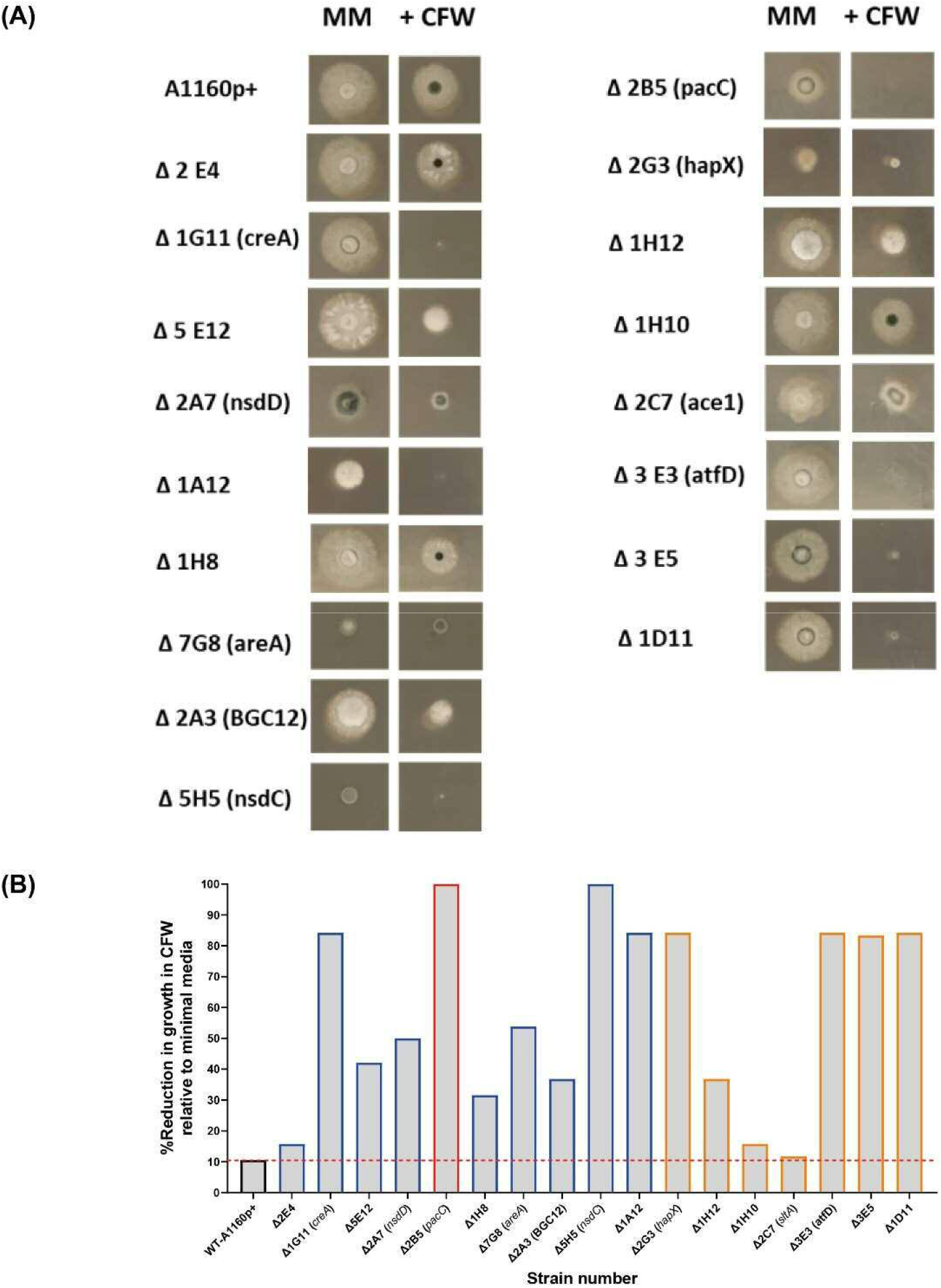
Sensitivity to calcofluor white in *A*.*fumigatus* TFKOs. Cell wall sensitivity was assessed in the *A. fumigatus* TFKOs by growing 10^6^ TFKO spores on minimal media agar or minimal media agar containing CFW (200 mg/L) for 48 hours. **A) Images of the TFKO colonies grown in Aspergillus minimal media with and without CFW**. 10^6^ TFKO spores were inoculated on agar and images taken after 48 hours. **B) Colony measurements taken in mm**. Percentage reduction in growth of TFKO in CFW calculated relative to growth in minimal media. WT=parental strain A1160p+.

### Reduced cytotoxicity of the *A. fumigatus* culture filtrates associates exclusively *with A. fumigatus* TFKOs defective in late causation of epithelial damage

*A. fumigatus* secretes a wide range of proteases (serine proteases, metalloproteinases and aspartic proteases), and other enzymes, proteins and toxins during growth in the host environment (Latgé and Chamilos, 2020). Gliotoxin, one of the most widely studied secreted toxins of *A. fumigatus* is associated with induction of apoptotic cell death in several mammalian cell types (Kwon-Chung and Sugui, 2009; Raffa and Keller, 2019). Additionally, culture filtrates from mutants lacking the PrtT regulator that governs expression of six secreted proteases in *A. fumigatus* is associated with a reduced epithelial damage capacity *in vitro* (Sugui *et al*., 2007; Cacho Teixeira *et al*., 2017). Further, the host-adapting transcriptome of the non-invasive *ΔpacC* isolate revealed dysregulation of several genes encoding putatively secreted proteins suggesting profound importance of *A. fumigatus* secretions during epithelial invasion (Bertuzzi *et al*., 2014). To measure cytotoxicity of TFKO culture filtrates, epithelial cell lysis was measured after challenging A549 cells with the culture filtrates of the TFKO mutants. To align with previous studies that had identified the expression of cytotoxic factors at >16 hours of fungal culture (Bertuzzi *et al*., 2014), hyphae were grown to maturity (48 hours) before culture filtrates were harvested and co-incubated with A549 cells for 24 hours. Interestingly, with the exception of *ΔpacC*, none of the TFKOs defective in epithelial detachment exhibited significantly reduced epithelial damage relative to the parental strain (**Fig 7A**). However, in stark contrast, the culture filtrates of 7 out of 8 of the TFKOs defective in late causation of epithelial damage exhibited reduced capacity to cause damage relative to the parental strain (**Fig 7B**). These results corroborated soluble factors to be major contributors to epithelial damage during the late phase of epithelial damage infection.

**Figure 7.**
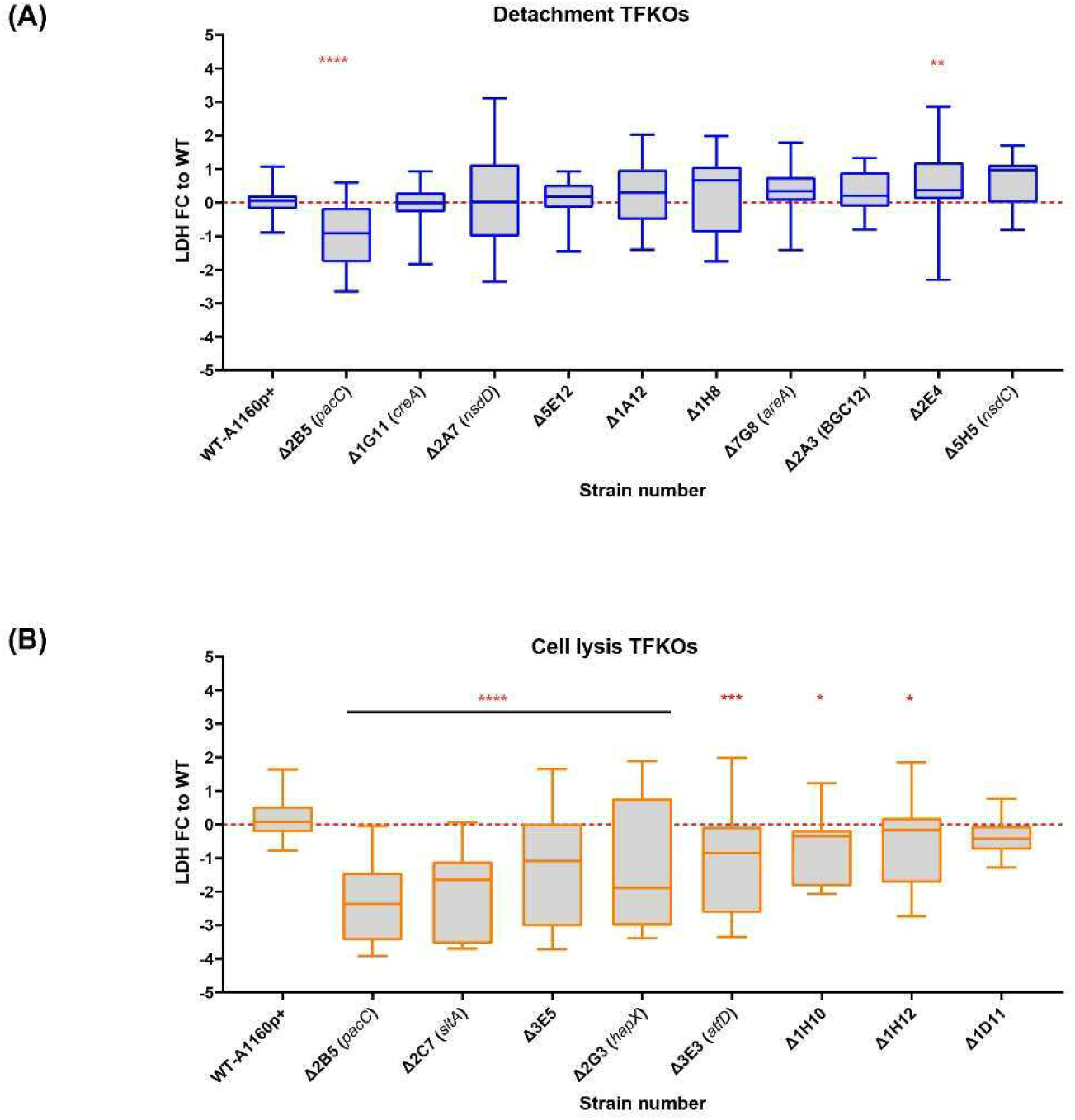
Culture filtrate damage caused by the *A*.*fumigatus* TFKOs. Damage caused by soluble factors of TFKOs to epithelial cells was assessed by measuring LDH release after 24-hour challenge of A549 cells with 48-hour culture filtrate of the TFKOs. **A) Fold change LDH of the cell detachment TFKOs B) Fold change LDH of cell lysis TFKOs**. WT=parental strain A1160+. Error bars show ±SEM. Data was analysed by one-way ANOVA with Fisher’s LSD multiple comparisons test. Data shown is technical reps of 3 biological reps. WT=A1160p+. *P<0.1, **P<0.01 ***P<0.001, ****P<0.0001.

## Discussion

*Aspergillus fumigatus* is a ubiquitous filamentous fungus and a potential cause of life-threatening lung disease in the immunocompromised. Interactions between the inhaled *A. fumigatus* spores and host lung pneumocytes are dynamic, complex, and poorly understood. Several recent reports identify important roles for lung epithelial cells during *A. fumigatus* infection, the nature of which can result in either expelling of the fungus or destruction of the lung (Croft *et al*., 2016; Bertuzzi *et al*., 2018). Previous *invitro* studies have identified a biphasic basis of fungus-induced lung epithelial damage, commencing with contact-dependent perturbation causing epithelial detachment, followed by hyphal mediated damage causing epithelial cell lysis, that in large part derives from the fungal secretome (Bertuzzi *et al*., 2014; Okaa *et al*., 2021). Despite the central importance of transcriptional regulation in driving expression of secreted fungal gene products, and the translational potential of inhibiting this process, very few transcriptional regulators have been identified as governing epithelial invasion.

To address this knowledge gap, a genome-scale census of *A. fumigatus* transcription factors was conducted (**Fig 1 and Fig 2**) by challenging a laboratory cultured pneumocyte cell line with 479 null mutants of *A. fumigatus* transcription factors (Furukawa *et al*., 2020). This identified 17 regulators of epithelial damage, 8 of which have not been previously characterised (**Table 1**). A critical discovery is that distinct cohorts of *A. fumigatus* genetic regulators drive early-acting epithelial detachment and late-acting cytolytic modes of epithelial damage. Only one *A. fumigatus* transcription factor, the pH-responsive transcription factor PacC, was found to govern causation of both early- and late-occurring damage. This finding is consistent with the idea that distinct genes could be employed at each phase of infection as the fungus establishes its niche in the host environment. A similar scenario has been reported for host kidney colonisation by *C. albicans* at 12, 24, and 48 hr post infection (Xu *et al*., 2015).

Several fungal attributes have been acknowledged to contribute towards the pathogenesis of *A. fumigatus* during establishment of infection (Dagenais and Keller, 2009; Latgé and Chamilos, 2020). This study has probed the regulatory landscape during the temporal pathogenic profile of *A. fumigatus* mediated damage of the alveolar epithelium (**Schematic in Fig 8a**). Interestingly, almost every mutant deficient in causing epithelial damage exhibited a unique pattern of phenotypes (**Summarised in Fig 8b and 8c**). Nonetheless, phenotypic traits were identified that correlate with early- and/or late-causation of damage. For example, amongst 10 mutants exhibiting reduced epithelial detachment, all but one isolate exhibited reduced rates of hyphal extension (**Fig 3**). The hyphal forms of the fungus have been implied as critical for invasion in other invasive fungi such as *C*.*albicans* (Desai, 2018). An obvious explanation, negated by our data, is that reduced rates of hyphal extension reduce the capacity of extending hyphae to secrete soluble mediators of epithelial damage. If this were true, we might expect to have seen significant numbers of TFKOs defective in both early- and late-causation of epithelial damage rather than exclusivity of transcriptional regulators to governance of particular traits (**Fig 7**). An alternative explanation is that slower hyphal growth rates place a limitation upon the number of host cells that the nascent hyphae can come into direct contact with. The latter argument aligns well with previous observations that early-acting damage occurs in a contact-dependent manner (Bertuzzi *et al*., 2014), as well as with the phenotypic analyses conducted in this study that revealed deficits in rate of uptake into and/or adhesion to epithelial cells (**Fig 4 and 5**). Aberrancies of cell wall organisation, as indicated by CFW sensitivity (**Fig 6**) might also contribute to reduced causation of epithelial detachment.

**Figure 8.**
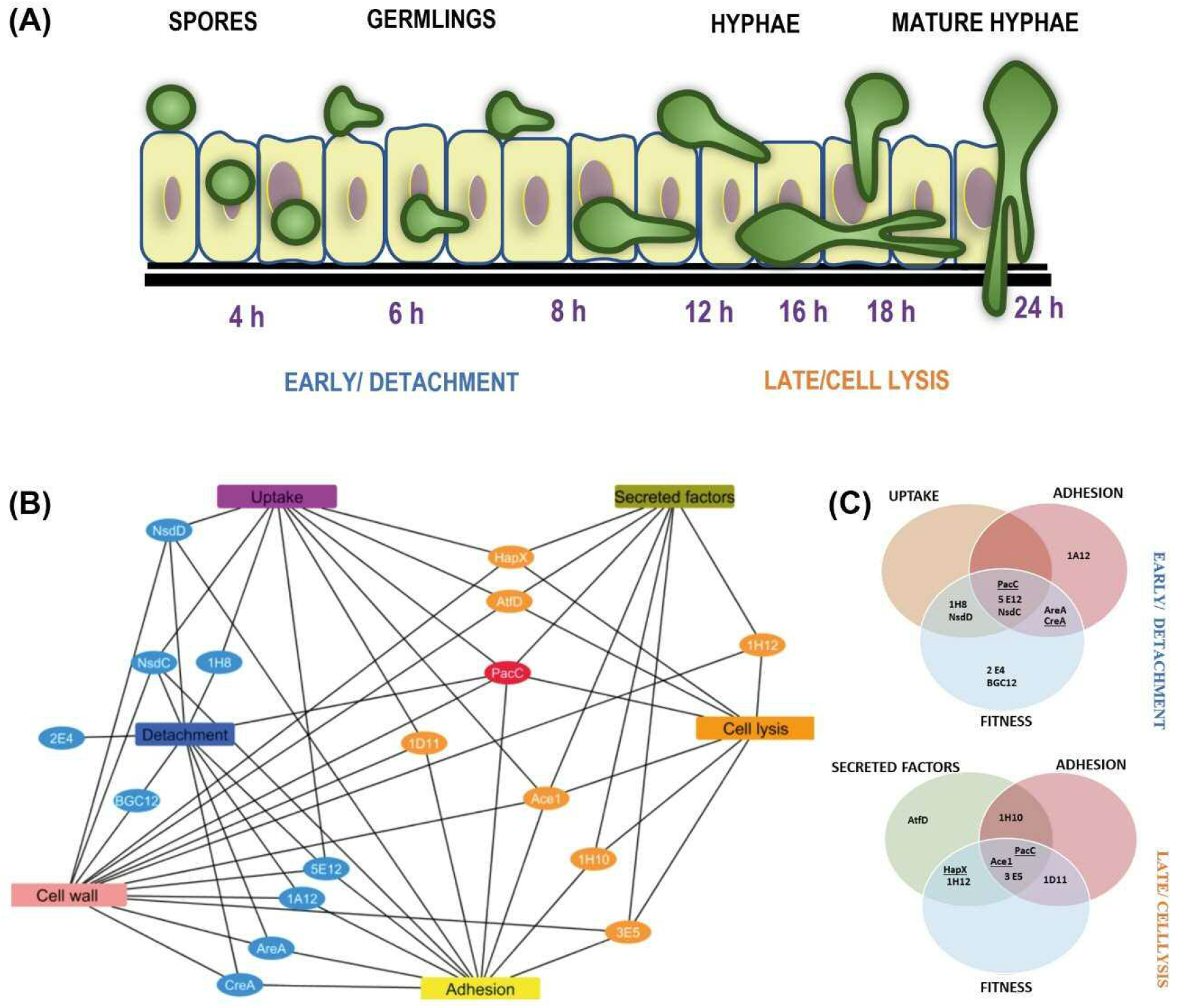
Temporal mechanistic profile of *A. fumigatus* mediated damage of alveolar epithelium regulated by the *A. fumigatus* transcription factors. **A) Schematic showing the temporal mechanistic profile of *A. fumigatus* mediated damage of alveolar epithelium**. *A. fumigatus* early interaction with the lung epithelium begins with spore uptake by the epithelial cells between 2-6 hours post infection. The spores germinate extracellular or intracellularly progressing infection between 8 and 12 hours. Adhesion of the germlings to the epithelium most likely enables close contact for invasion. Cell wall mediated damage occurs at the early stage directed by the detachment regulators. Following hyphal extension post 16 hours, the cell lysis regulators facilitate invasion via secretion of proteases and toxins, The cumulative effect of these epithelial interactions moderated by the *A. fumigatus* regulators leads to epithelial invasion. **B) and C) Unique phenotypic and mechanistic profile of the *A. fumigatus* regulators**, B) shows a cytoscape network of the unique mechanistic attributes of the cell detachment and cell lysis regulators. C) shows the attributes of the cell detachment and cell lysis mutants for uptake, adhesion, fitness and secreted factors (culture filtrates). The TFs underlined exhibited altered germination rates.

The characterised functions of previously studied transcription factors yield further clues to the mechanisms underlying the epithelial detachment phenomenon. Among the 8 TFs required for efficient pneumocyte detachment 4 TFs (PacC (AFUB_037210), AreA (AFUB_096370) NsdD and NsdC (AFUB_035330 and AFUB_089440) have been implicated in co-ordination of sporulation and secondary metabolism (Calvo *et al*., 2002; Bayram and Braus, 2012; de Castro *et al*., 2021). Consistent with such functionality, the colonial phenotypes of all four mutants exhibit compact morphology (**Fig 3 and Fig 6**). It is therefore feasible that the secondary metabolite composition of the spores of these mutants differs sufficiently from that of the parental isolate to cause deficits in epithelial detachment, either via aberrancies in one or several of contact-mediated toxicity, epithelial uptake of and phagolysomal fusion, the latter of which has recently been shown to determine efficiency of intracellular fungal killing (Ben-Ghazzi *et al*., 2021).

Remarkably, TFKOs that are deficient in causing epithelial cell lysis universally exhibit culture filtrates that are less cytolytic than the parental progenitor (**Fig7**). In 3 out of 8 instances PacC, HapX and SltA (Bertuzzi *et al*., 2014; Gsaller *et al*., 2014; H. Liu *et al*., 2021), it might be argued that this occurs as a direct consequence of radically reduced hyphal growth. However, in the case of PacC mature hyphae (generated via extended incubation times) produce a cytolytically inert secretome, presumably due to loss of gliotoxin biosynthesis and multiple secreted fungal proteases (Bertuzzi *et al*., 2014). Of the remaining 5 TFKOs deficient in causing cytolytic damage to epithelial cells, the functionality of the TFs remains uncharacterised. It will be interesting to identify commonalities/distinctions between the secretomes of these mutants to pinpoint the causal agents of cytolytic host cell death. In summary, our finding supports the idea that virulence is multifactorial, hence there could be a combination of the fungal attributes that contributed to reduced epithelial invasion. For example, i*nvivo* transcriptomic studies have shown PacC to regulate secreted and cell wall genes and deficit in causing uptake but not spore epithelial adhesion (Bertuzzi *et al*., 2014) and SltA regulates the expression of multiple secondary metabolite gene clusters and mycotoxin as well being sensitive to cell wall perturbing agents (H. Liu *et al*., 2021)

Surprisingly *A. fumigatus* transcription factor PrtT reported to regulate protease production (Sharon, Hagag and Osherov, 2009) did not show significant differences in epithelial cytotoxicity compared to the parental strain. (Okaa *et al*., 2021). Further, the GliZ TF that regulates gliotoxin production (Bok *et al*., 2006; Schoberle *et al*., 2014) did not contribute to epithelial damage during our high throughput screening. Our study also did not identify TFKO mutants previously reported to have deficient adhesion and damage to epithelial cells namely DvrA, MedA and SomA (Ejzykowicz *et al*., 2010; Gravelat *et al*., 2010; Lin *et al*., 2015) to cause highly significant differences in epithelial damage. One plausible explanation is the disparate genetic backgrounds of the constructed mutants that could have an effect on their capacity to cause damage.

It remains to be seen, in the context of mammalian disease, whether the effects of early- and late-acting damage processes have mutually exclusive, additive or synergistic effects upon pathogenicity. Of note, the only TF that orchestrates both modes of damage, PacC, has been demonstrated to be a key regulator of epithelial invasion that is critical for mammalian pathogenicity (Bertuzzi *et al*., 2014). Given the broad regulatory activity of TFs, it is likely that the phenotypes associated with null mutants will be more pleiotropic in nature than we have discovered in this study, and it will therefore be necessary to identify the individual effectors of epithelial damage that function under their regulatory control. Nonetheless, the distinctive spatial and temporal contributions of TFs to epithelial damage indicate that early- and late-acting damage might be differentially significant to diseases caused predominantly by spore or hyphal forms of *A. fumigatus*. It is important to note that in this study, we are inherently studying the interaction between the fungus and the epithelial cells, we are not considering whether host inflammatory responses dampen or worsen the damage elicited by *A. fumigatus* infection. Acknowledging that pathogenicity is a dual phenomenon moderated by both host and pathogen factors, it could be likely that the fungus must elicit host damage which serves as a signal to amplify residual host inflammatory response, in turn driving pathogenicity.

In conclusion, the discovery of the regulators of epithelial damage and essentially their phenotypic and mechanistically profile moderating damage has allowed a birds-eye scan in understanding the highly dynamic host-pathogen turmoil. This is necessary to fully understand the pathogenesis of IPA and also facilitates new gene targets for therapeutics, both of which would help us overcome the morbidity and mortality caused by the human killer fungus *A. fumigatus*.

## ACKNOWLEDGEMENTS

This research work from this paper was funded from project grants MR/M02010X/1 to EMB, MJB and MR; and MR/S001824/1 and MR/L000822/1 to EMB.

## AUTHOR CONTRIBUTIONS

Conceived and designed the experiments: SR EMB MR. Performed the experiments: SR LG ZC. Analyzed the data: SR EMB PP NVR MR. Prepared cell lines: RFG LG. Assisted with microscopy and technical issues: DDT. Wrote the paper: SR EMB. Critically revised the manuscript: SR EMB NVR MJB DDT PP

## COMPETING INTERESTS

MJB is the director and shareholder of PIQ Laboratories Ltd. The remaining authors declare no competing interests.

## MATERIALS AND CORRESPONDENCE

Correspondence to Elaine M Bignell

## SUPPLEMENTARY INFORMATION

### SUPPLEMENTARY FIGURES

**Figure S1.**
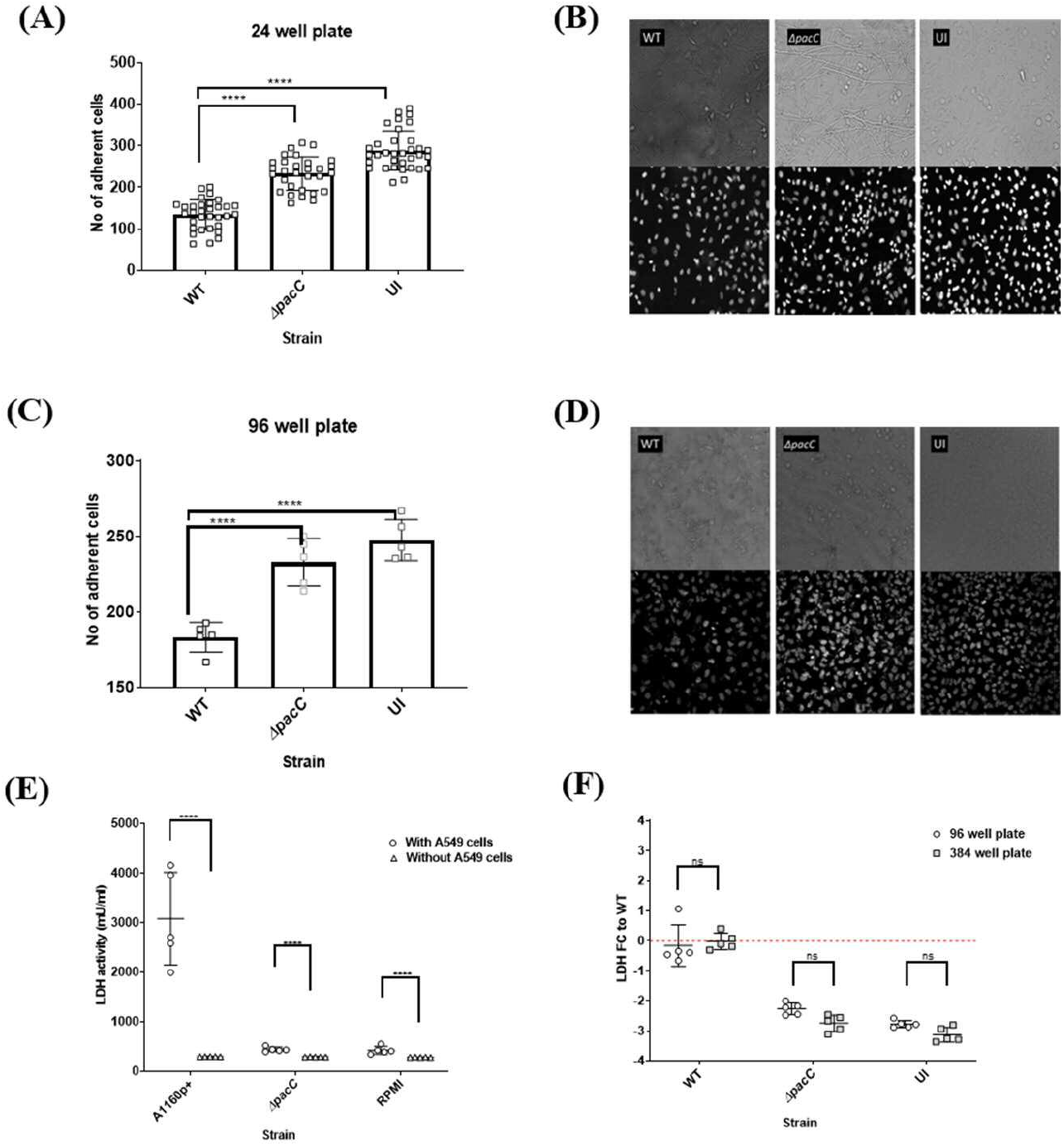
Optimisation of epithelial damage assays for high throughput screening. Performance of detachment assay in a low throughput (A and B) and high throughput plate format (C and D) A) Number of adherent cells per image enumerated in a 24 well plate. The total number of adherent cells per field of view were enumerated using a FIJI macro counting DAPI stained cells after *A. fumigatus* A1160+ and *ΔpacC* strains infection. Data was analysed by multiple t-tests. ****P<0.0001. **B) Representative images A1160p+ or *ΔpacC* after 16-hour infection of A549 monolayer in a 24 well plate**. Images captured manually at 20 x magnification using NIKON TE-2000E **C) Number of adherent cells per image enumerated in a 96 well plate**. The total number of adherent cells per image were enumerated using a FIJI macro counting DAPI stained cells after *A. fumigatus* A1160+ and *ΔpacC* strains infection. Data was analysed by multiple t-tests. ****P<0.0001. **D) Representative images A1160p+ and *ΔpacC* strains after 16-hour infection of A549 monolayer in a 96 well plate**. Images captured automatedly at 40x magnification using confocal microscope. **Suitability of LDH assay for high-throughput cell cytotoxicity screening (E and F) E) *A. fumigatus* does not produce measurable LDH enzyme at 24 hours**. Absolute LDH released by 500,000 spores of A1160p+, *ΔpacC* and un-infected in a 24 well plate after 24 hours challenge with and without A549 cells. **F) Performance of LDH assay in 96 versus 384 well format**. LDH assay performed in a 96 well and a 384 well plate showing LDH fold change. Error bars show ±SEM. Data was analysed by one-way ANOVA with Fisher’s LSD multiple comparisons test. UI=Un-infected. ***P<0.001, ****P<0.0001.

**Figure S2.**
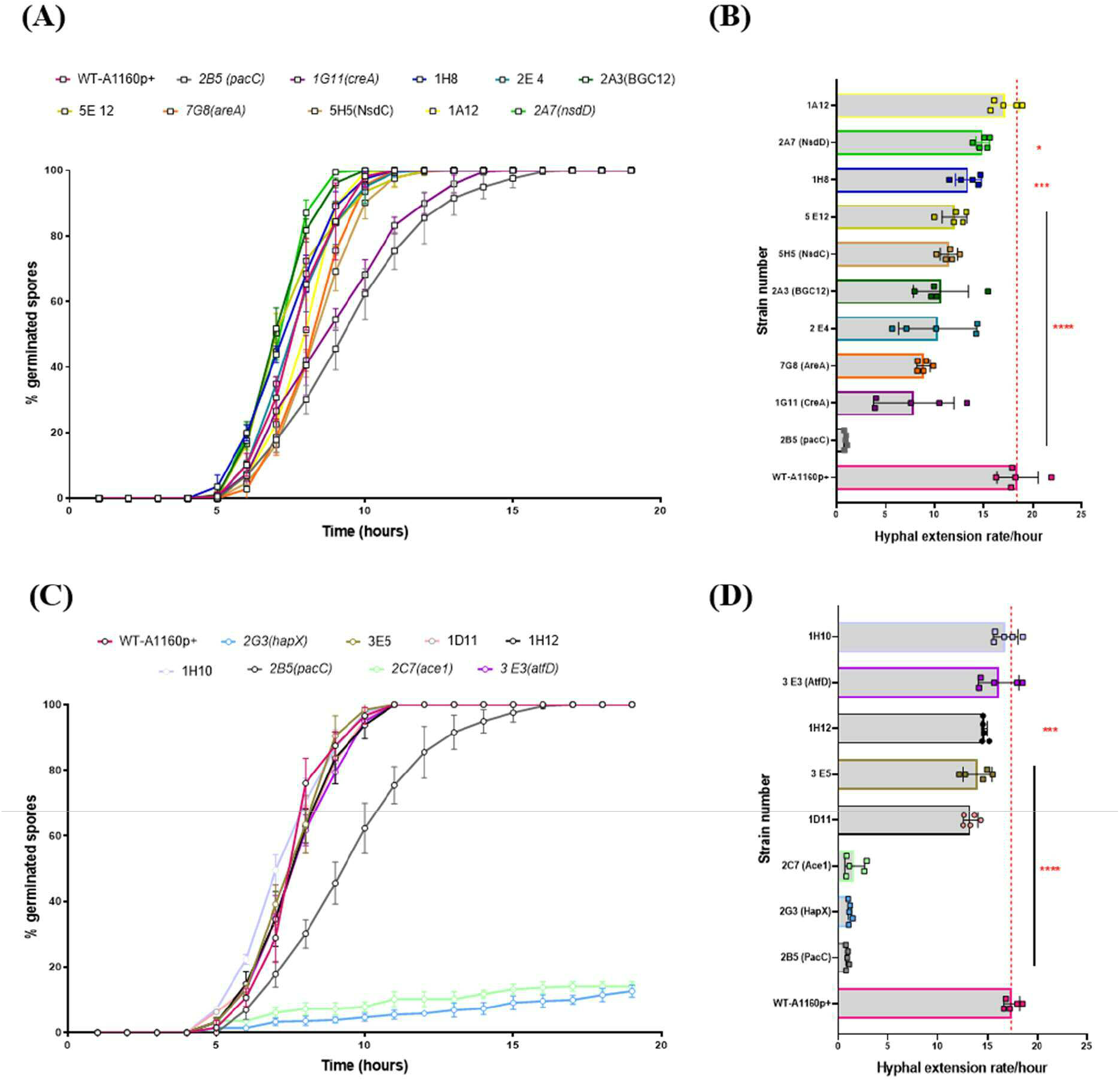
Hyphal extension rate and germination potential of the *A. fumigatus* TFKOS measured by assessing growth using time-lapse confocal microscopy. **A)** Hyphal extension rate of the eRegs (cell detachment) mutants **B)** Percentage germination of the eRegs (cell detachment) mutants. **C)** Percentage germination of the eRegs (cell lysis) mutants. **D)** Hyphal extension rate of the eRegs (cell lysis) mutants. Data was analysed by one-way ANOVA with Fisher’s LSD multiple comparisons test. WT=A1160p+. *P<0.1, **P<0.01 ***P<0.001, ****P<0.0001.

**Figure S3.**
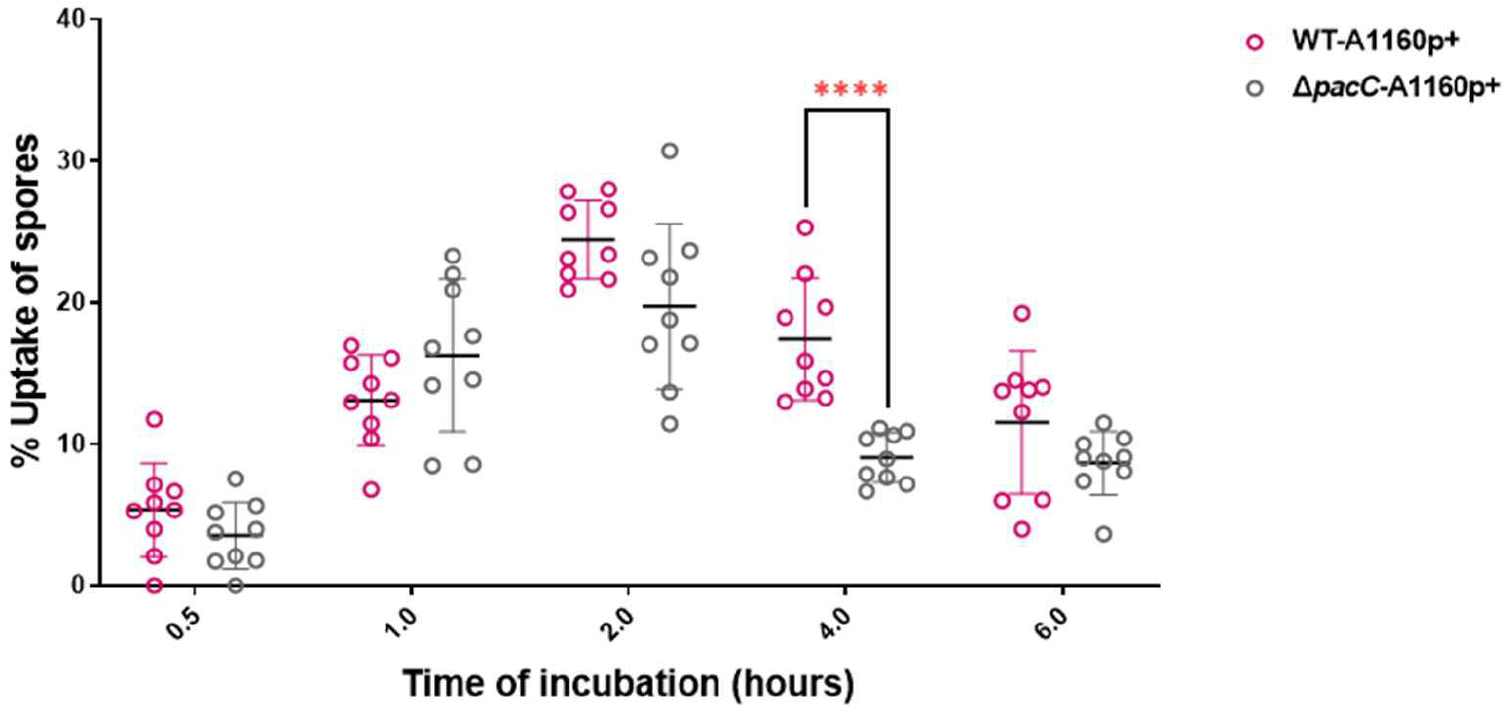
Uptake capacity of A549 cells to the *A. fumigatus* parental strain (A1160p+) and *ΔpacC* strain over time. FITC stained spores incubated with A549 cells for 30min, 1 hour, 2 hours and 4 hours to enable A549 internalisation, after which the infected cells, were washed with PBS to stain the externally adherent spores with calcofluor white. Images were captured using the TCS-SP8 confocal microscope at 40x magnification Error bars show ±SEM. Data was analysed by two-way ANOVA Sidak’s comparisons test of means. ***P<0.001, ****P<0.0001. Data shown is 3 technical reps with three biological replicates.

### SUPPLEMENTARY TABLE

**TABLE S1:** Output of the detachment (sheet 1) and cell lysis (sheet 2) screening for the TFKO mutants used for the generation of the volcano plot and the analysis

### SUPPLEMENTARY DATA

**TABLE S2:** Output of the Boolean analysis performed for the LDH epithelial cell lysis screening for the TFKO mutants

