## Supplementary material for "Distinct cohorts of *Aspergillus fumigatus* transcription factors are required for epithelial damage occurring via contact- or soluble effector-mediated mechanisms": TABLE S2

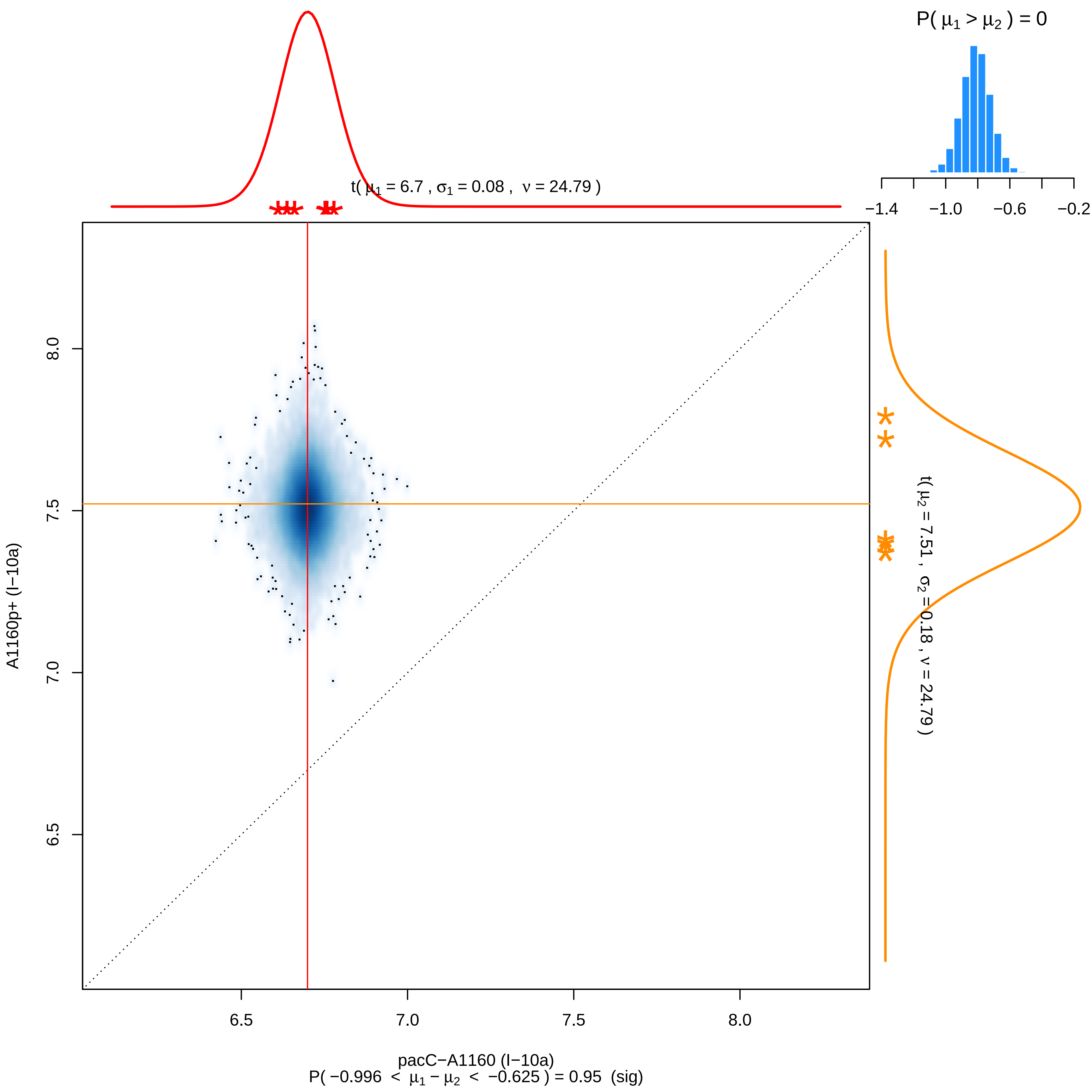
P( μ1 > μ2 ) = 0
−1.4
−1.0
−0.6
−0.2
t( μ1 = 6.7 , σ1 = 0.08 , ν = 24.79 )
t( μ2 = 7.51 , σ2 = 0.18 , ν = 24.79 )
A1160p+ (I−10a)
8.0
7.5
7.0
6.5
6.5
7.0
7.5
8.0
pacC−A1160 (I−10a)
P( −0.996 < μ1 − μ2 < −0.625 ) = 0.95 (sig)

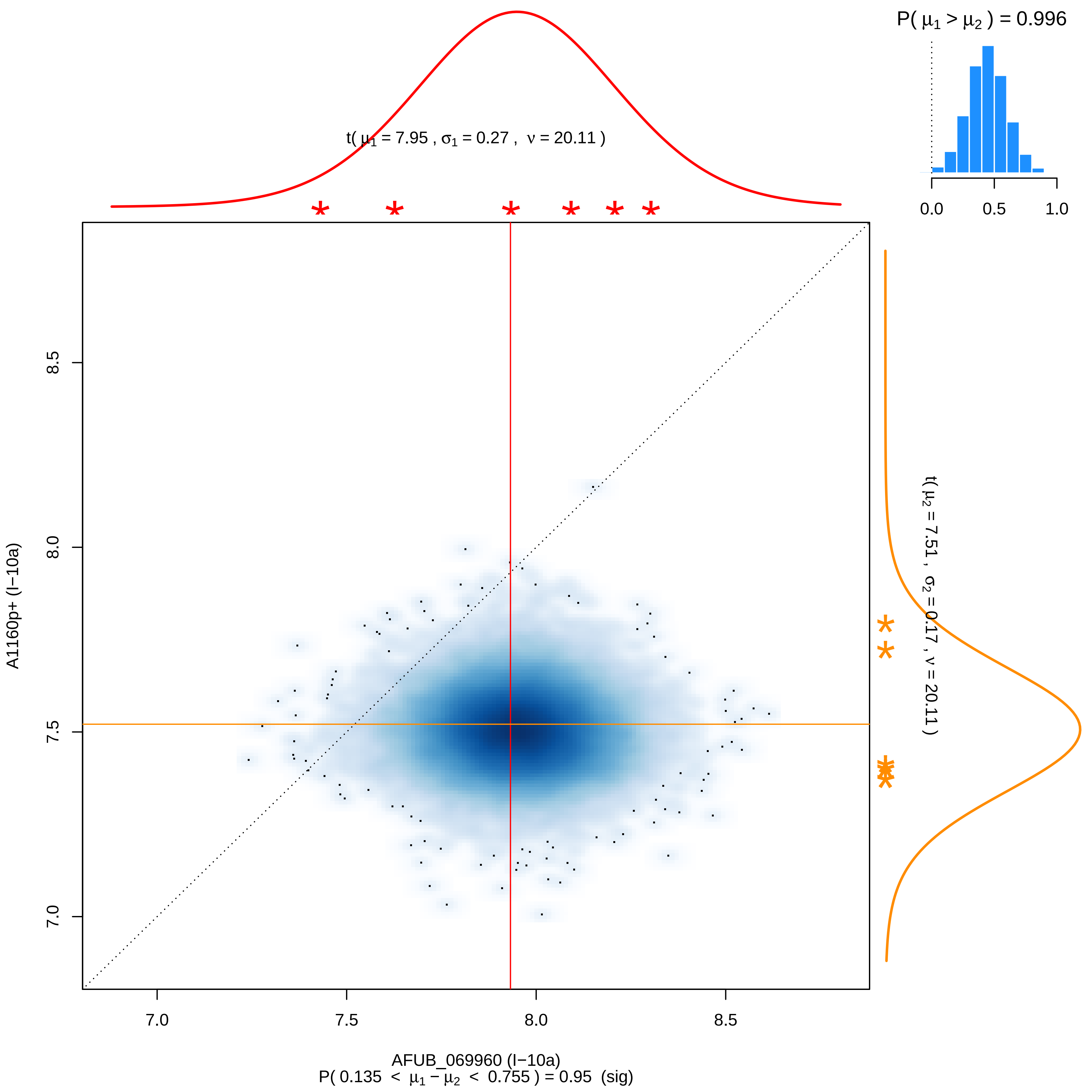

P( $\mu_1 > \mu_2$ ) = 0.996
0.0
0.5
1.0
t( $\mu_1$ = 7.95 , $\sigma_1$ = 0.27 , $\nu$ = 20.11 )
t( $\mu_2$ = 7.51 , $\sigma_2$ = 0.17 , $\nu$ = 20.11 )
A1160p+ (I−10a)
8.5
8.0
7.5
7.0
7.0
7.5
8.0
8.5
AFUB_069960 (I−10a)
P( 0.135 < $\mu_1 - \mu_2$ < 0.755 ) = 0.95 (sig)

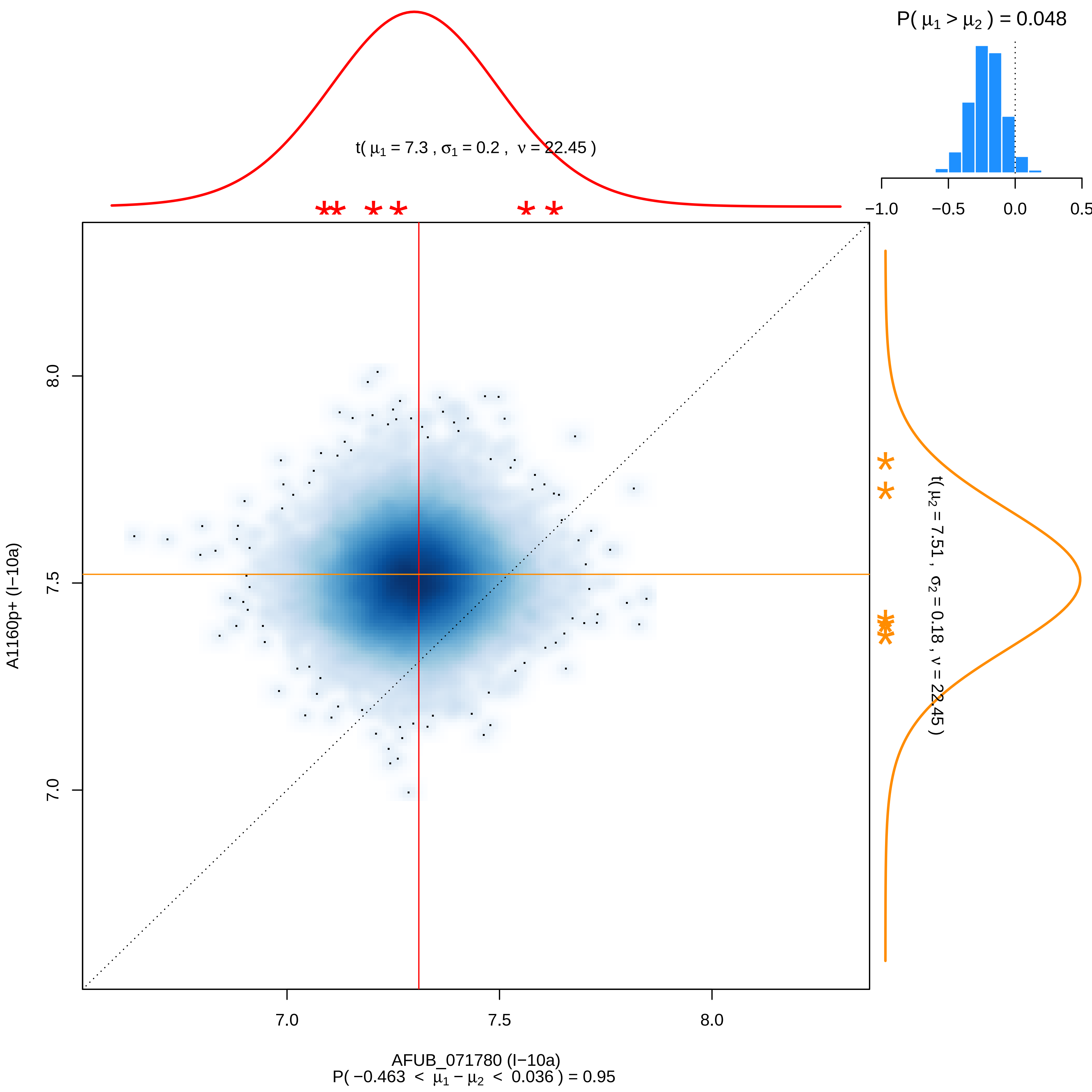
P( μ1 > μ2 ) = 0.048
−1.0
−0.5
0.0
0.5
t( μ1 = 7.3 , σ1 = 0.2 , ν = 22.45 )
t( μ2 = 7.51 , σ2 = 0.18 , ν = 22.45 )
8.0
7.5
7.0
A1160p+ (I−10a)
7.0
7.5
8.0
AFUB_071780 (I−10a)
P( −0.463 < μ1 − μ2 < 0.036 ) = 0.95

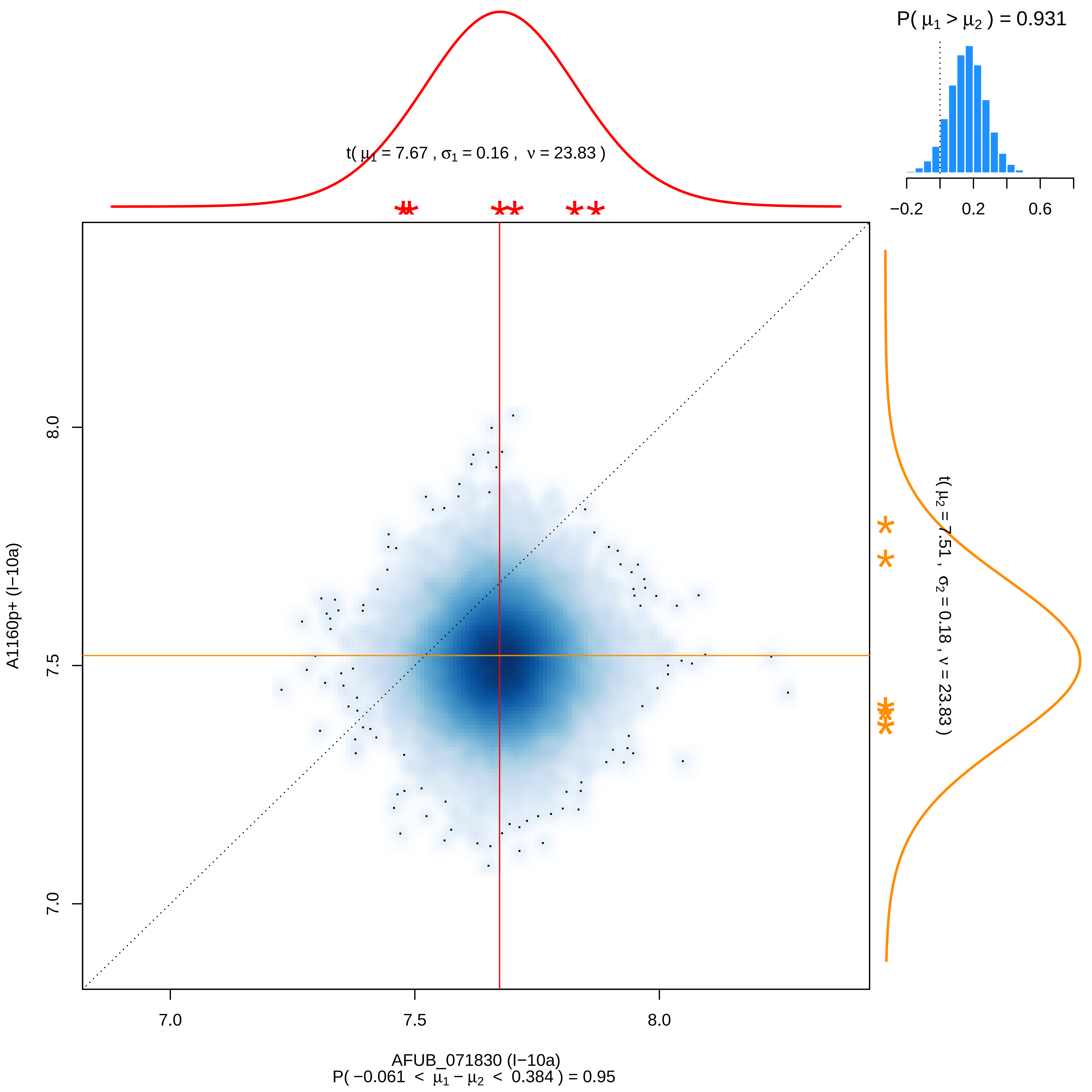
P( μ1 > μ2 ) = 0.931
−0.2
0.2
0.6
t( μ1 = 7.67 , σ1 = 0.16 , ν = 23.83 )
t( μ2 = 7.51 , σ2 = 0.18 , ν = 23.83 )
A1160p+ (I−10a)
7.0
7.5
8.0
AFUB_071830 (I−10a)
P( −0.061 < μ1 − μ2 < 0.384 ) = 0.95

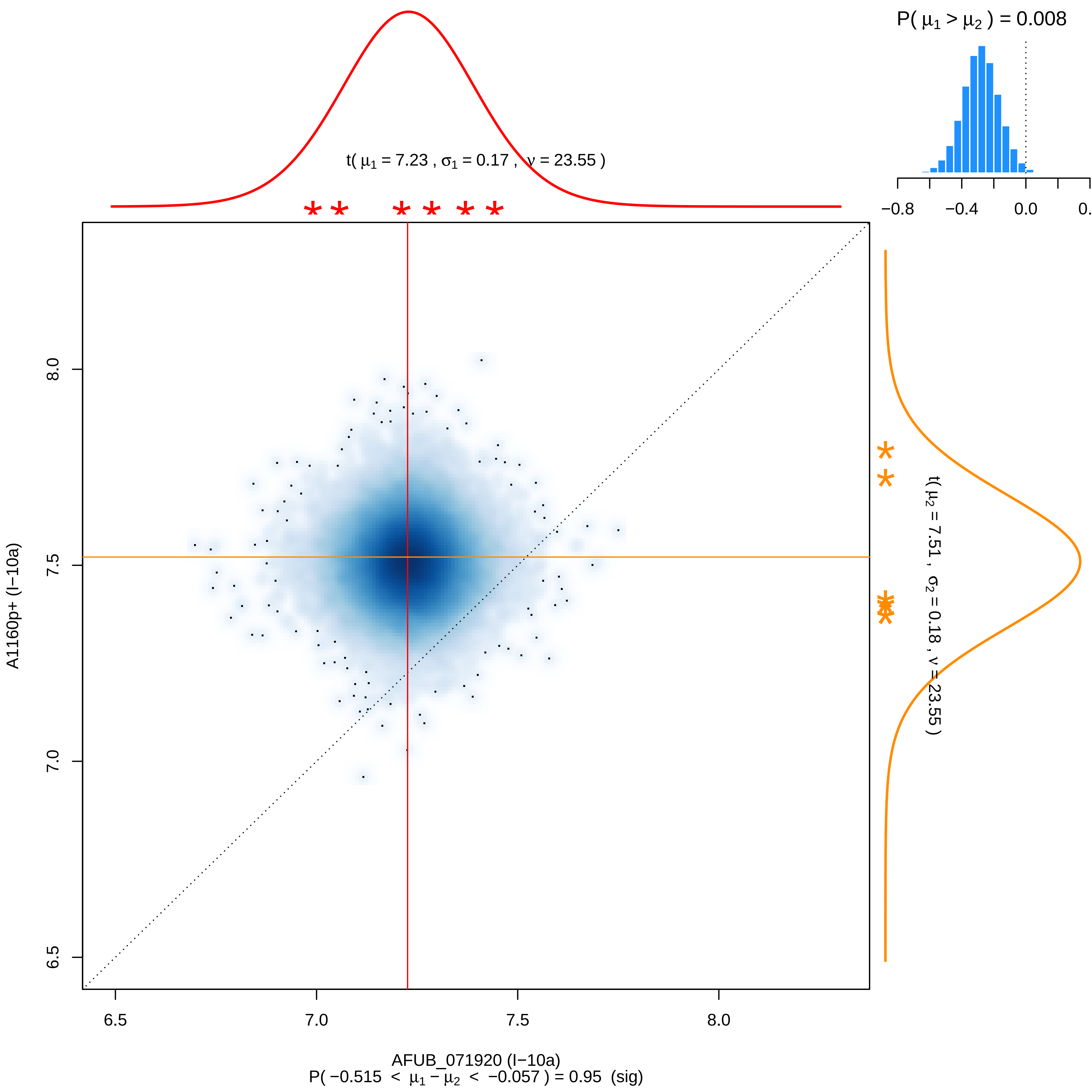
P( μ1 > μ2 ) = 0.008
−0.8
−0.4
0.0
t( μ1 = 7.23 , σ1 = 0.17 , ν = 23.55 )
t( μ2 = 7.51 , σ2 = 0.18 , ν = 23.55 )
8.0
7.5
7.0
6.5
A1160p+ (I−10a)
6.5
7.0
7.5
8.0
AFUB_071920 (I−10a)
P( −0.515 < μ1 − μ2 < −0.057 ) = 0.95 (sig)

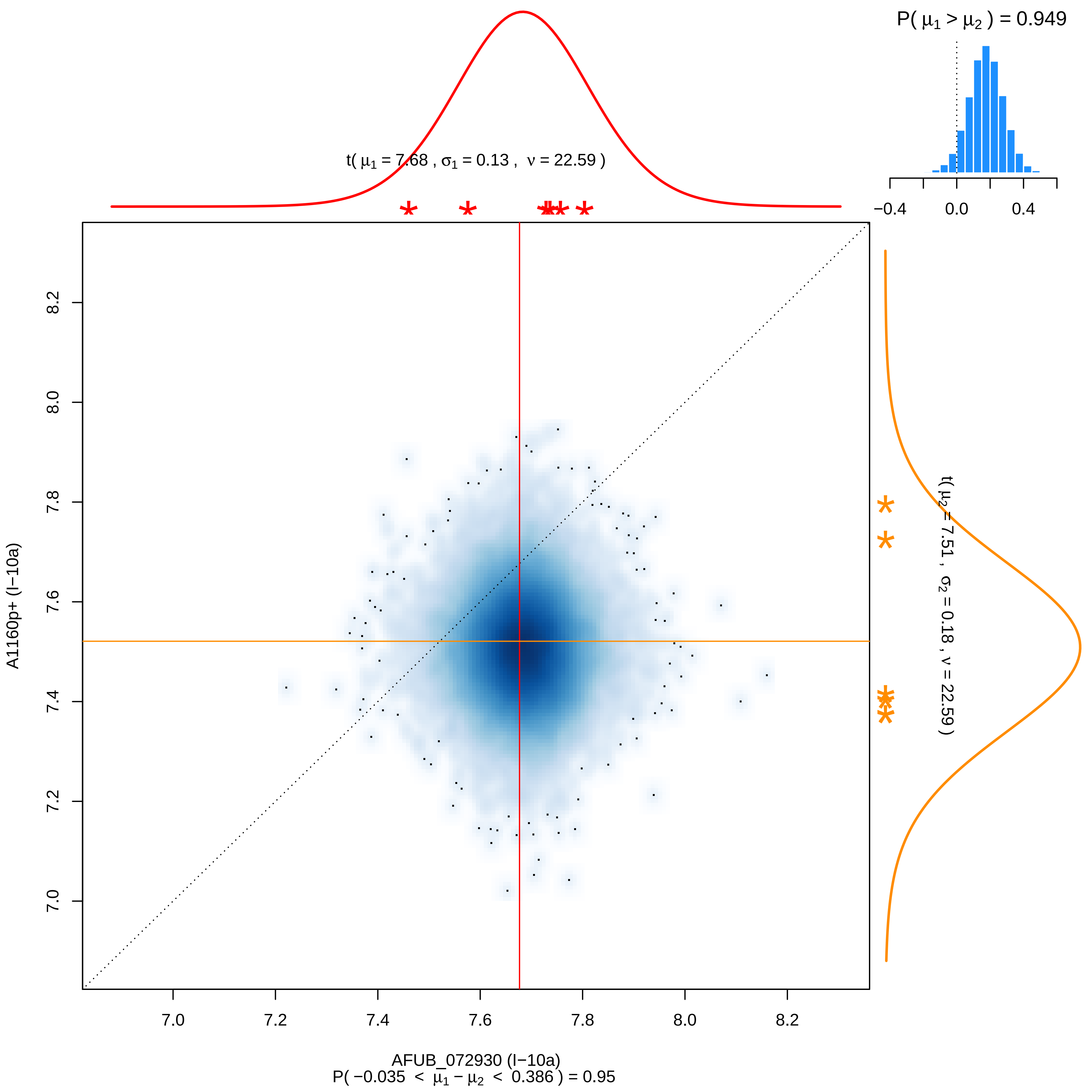
P( μ1 > μ2 ) = 0.949
−0.4
0.0
0.4
t( μ1 = 7.68 , σ1 = 0.13 , ν = 22.59 )
t( μ2 = 7.51 , σ2 = 0.18 , ν = 22.59 )
A1160p+ (I−10a)
8.2
8.0
7.8
7.6
7.4
7.2
7.0
7.0
7.2
7.4
7.6
7.8
8.0
8.2
AFUB_072930 (I−10a)
P( −0.035 < μ1 − μ2 < 0.386 ) = 0.95

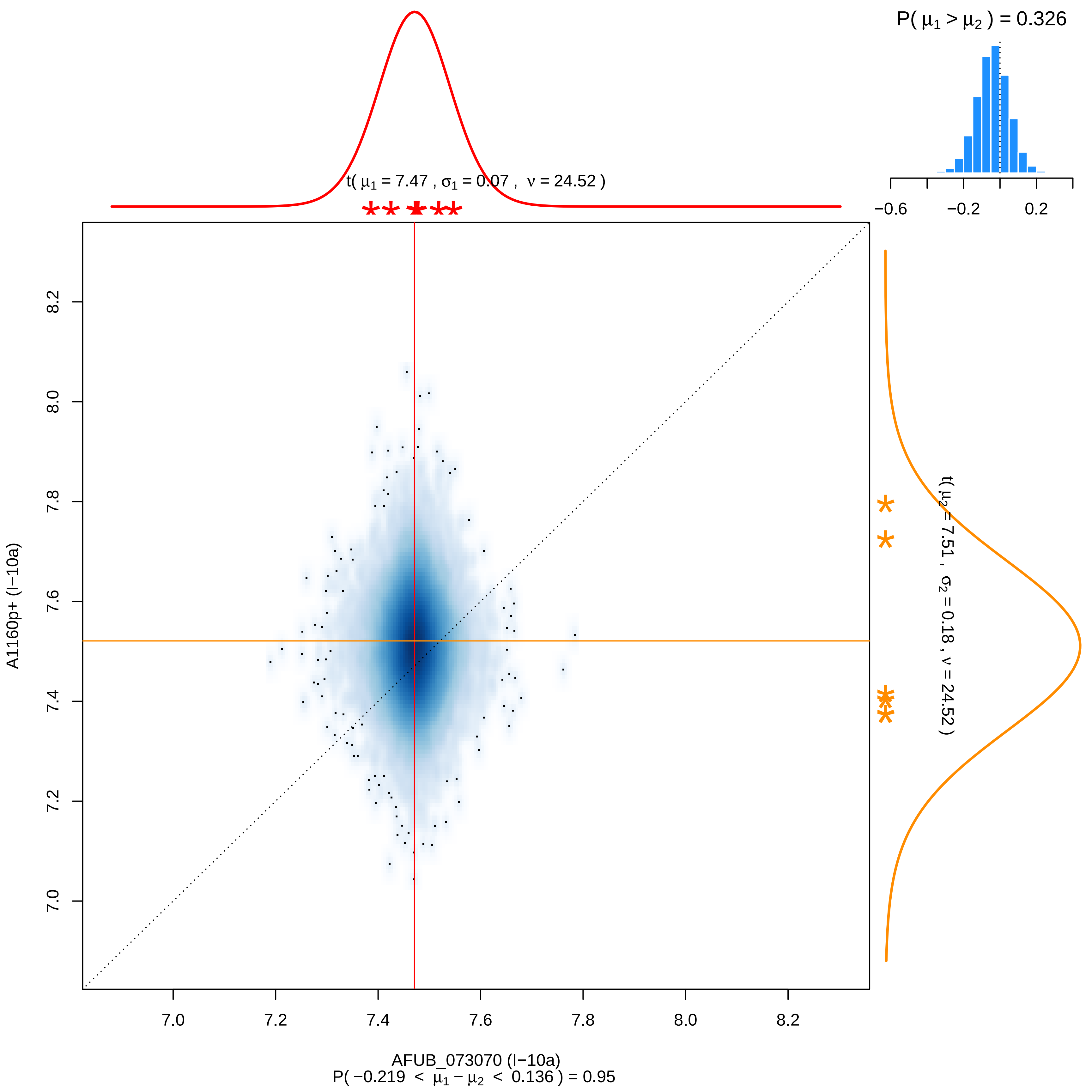

P( μ1 > μ2 ) = 0.326
−0.6
−0.2
0.2
t( μ1 = 7.47 , σ1 = 0.07 , ν = 24.52 )
t( μ2 = 7.51 , σ2 = 0.18 , ν = 24.52 )
8.2
8.0
7.8
7.6
7.4
7.2
7.0
A1160p+ (I−10a)
7.0
7.2
7.4
7.6
7.8
8.0
8.2
AFUB_073070 (I−10a)
P( −0.219 < μ1 − μ2 < 0.136 ) = 0.95

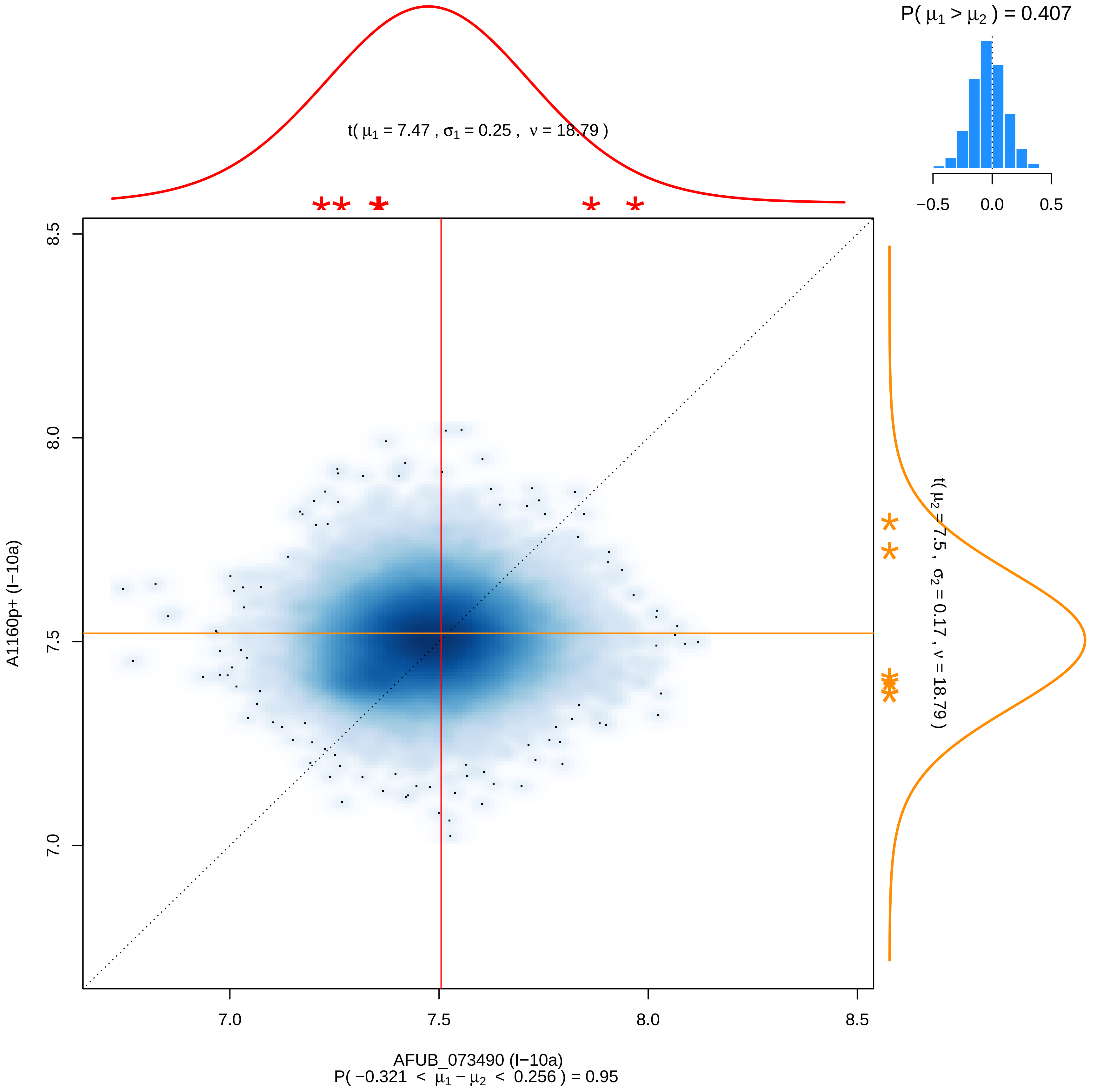

P( $\mu_1 > \mu_2$ ) = 0.407
−0.5
0.0
0.5
t( $\mu_1$ = 7.47 , $\sigma_1$ = 0.25 , $\nu$ = 18.79 )
t( $\mu_2$ = 7.5 , $\sigma_2$ = 0.17 , $\nu$ = 18.79 )
A1160p+ (I−10a)
7.0
7.5
8.0
8.5
AFUB_073490 (I−10a)
P( −0.321 < $\mu_1 - \mu_2$ < 0.256 ) = 0.95

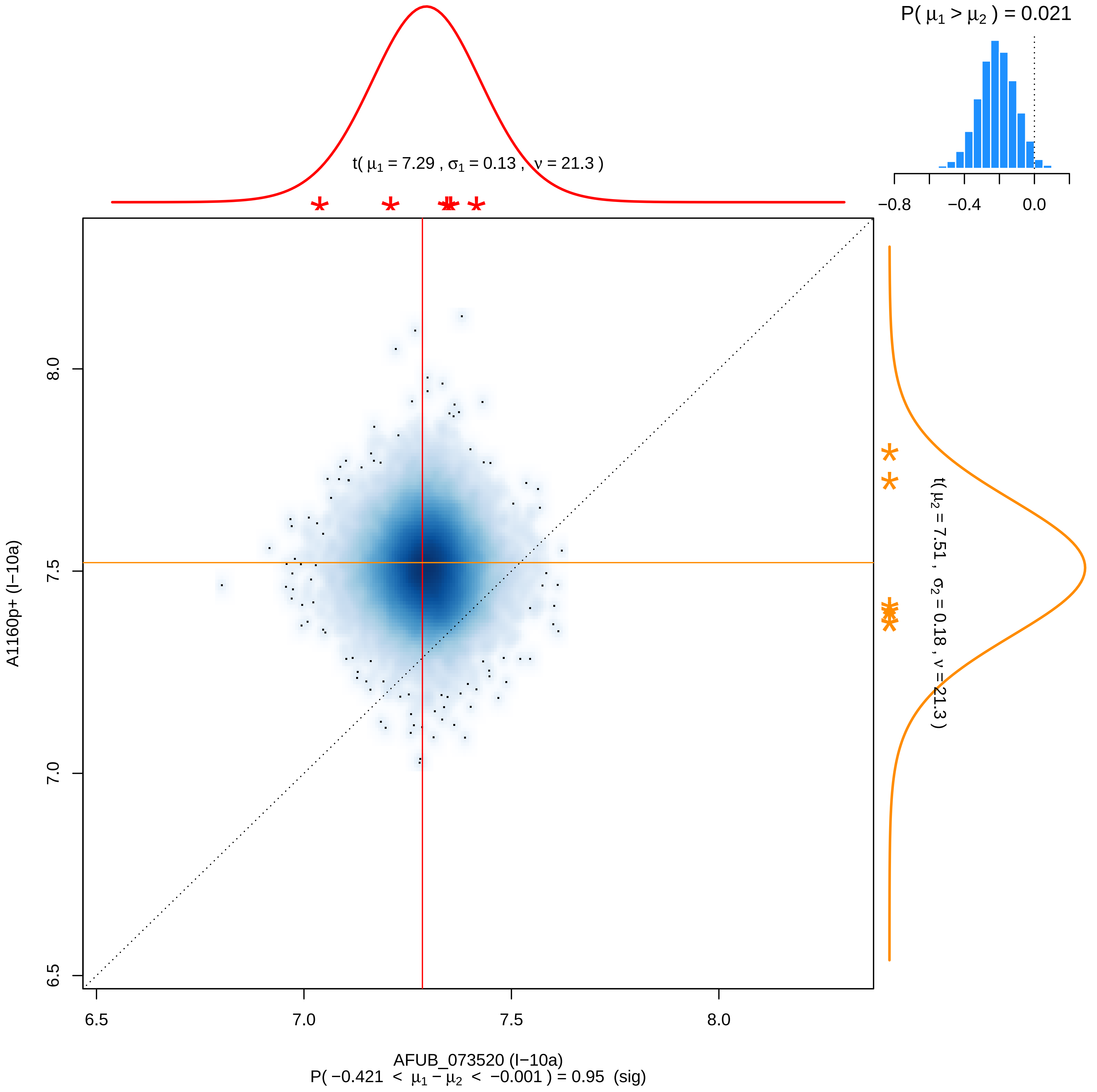

P( μ1 > μ2 ) = 0.021
−0.8
−0.4
0.0
t( μ1 = 7.29 , σ1 = 0.13 , ν = 21.3 )
t( μ2 = 7.51 , σ2 = 0.18 , ν = 21.3 )
A1160p+ (I−10a)
8.0
7.5
7.0
6.5
6.5
7.0
7.5
8.0
AFUB_073520 (I−10a)
P( −0.421 < μ1 − μ2 < −0.001 ) = 0.95 (sig)

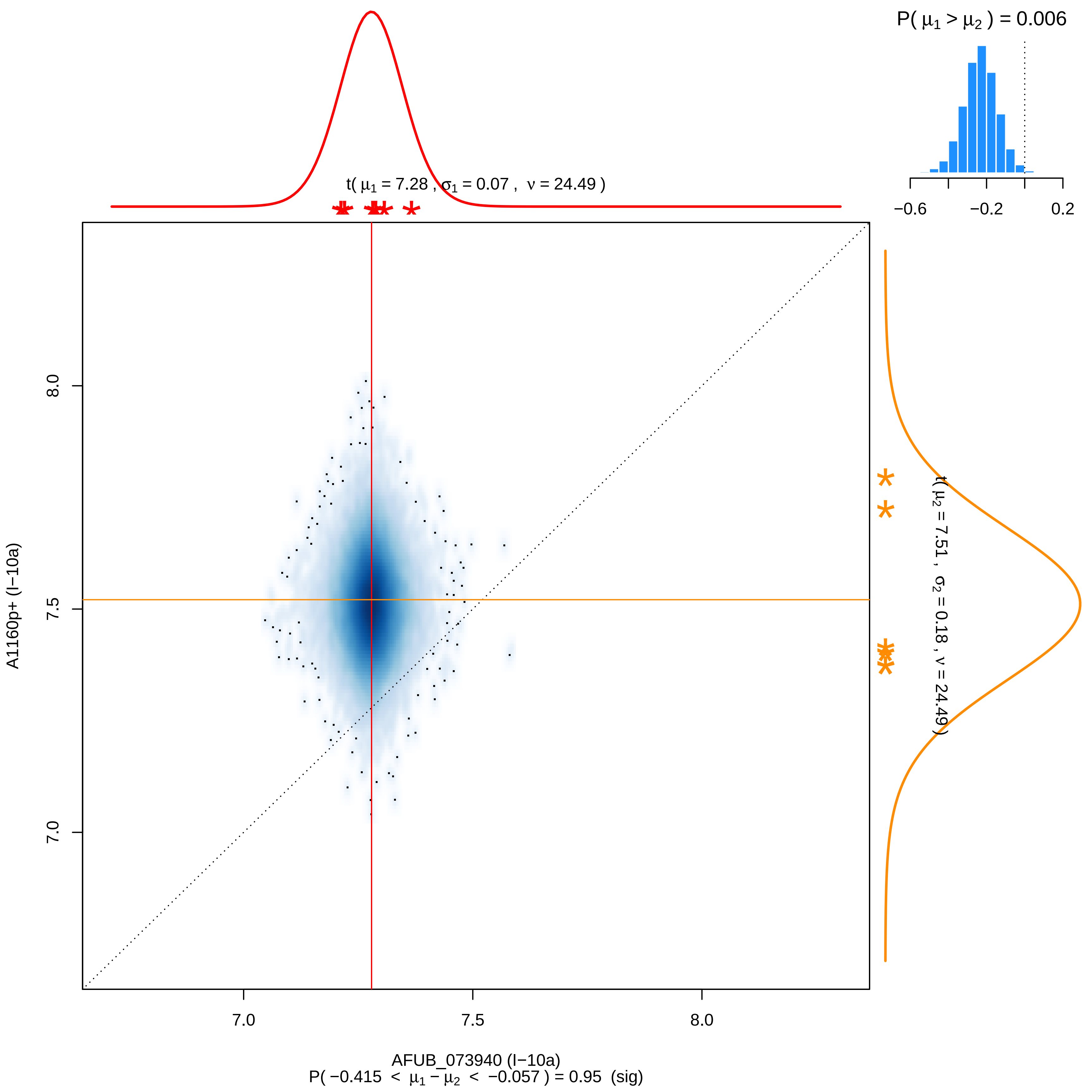

P( μ1 > μ2 ) = 0.006
−0.6
−0.2
0.2
t( μ1 = 7.28 , σ1 = 0.07 , ν = 24.49 )
t( μ2 = 7.51 , σ2 = 0.18 , ν = 24.49 )
8.0
7.5
7.0
A1160p+ (I−10a)
7.0
7.5
8.0
AFUB_073940 (I−10a)
P( −0.415 < μ1 − μ2 < −0.057 ) = 0.95 (sig)

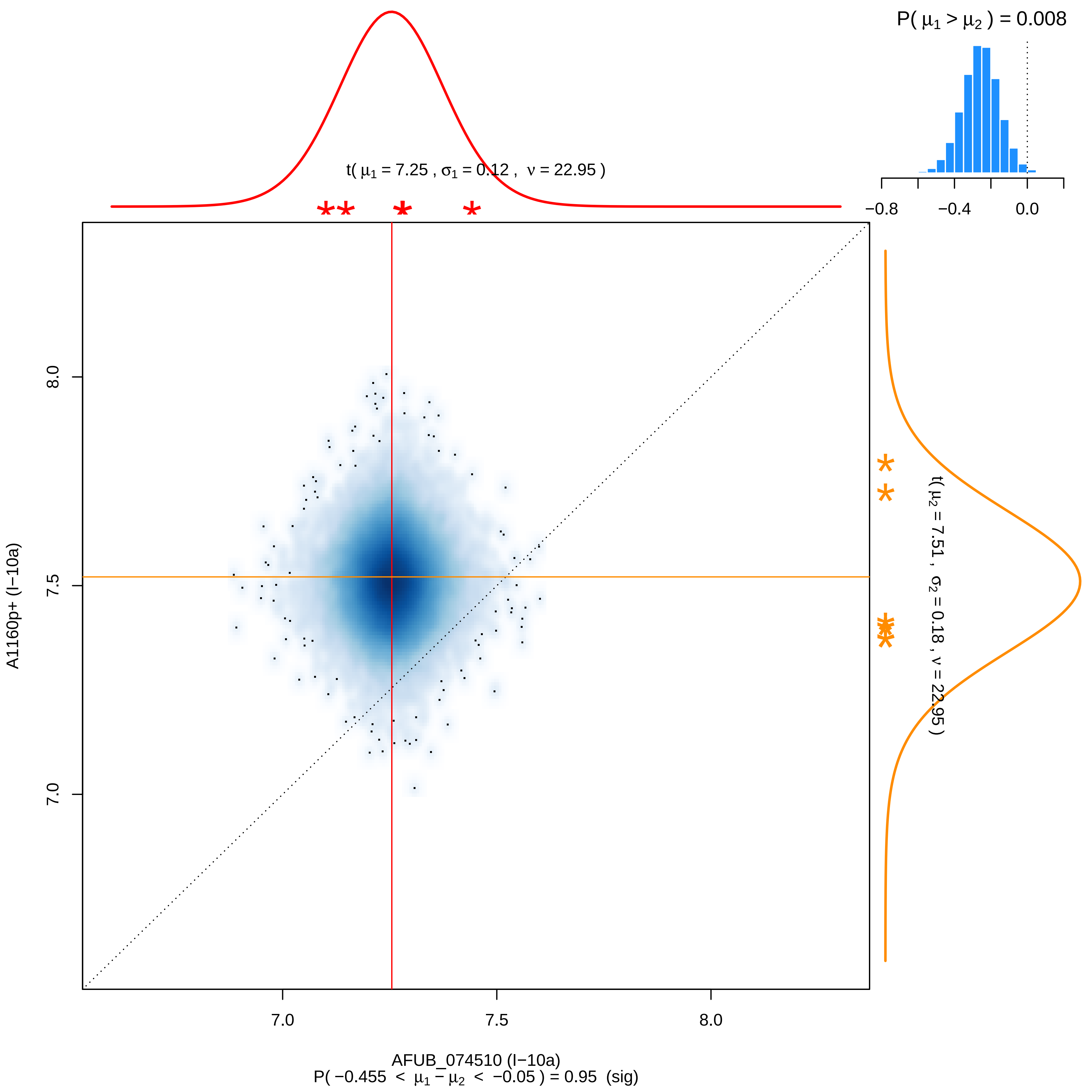

P( μ1 > μ2 ) = 0.008
−0.8
−0.4
0.0
t( μ1 = 7.25 , σ1 = 0.12 , ν = 22.95 )
t( μ2 = 7.51 , σ2 = 0.18 , ν = 22.95 )
A1160p+ (I−10a)
8.0
7.5
7.0
AFUB_074510 (I−10a)
P( −0.455 < μ1 − μ2 < −0.05 ) = 0.95 (sig)

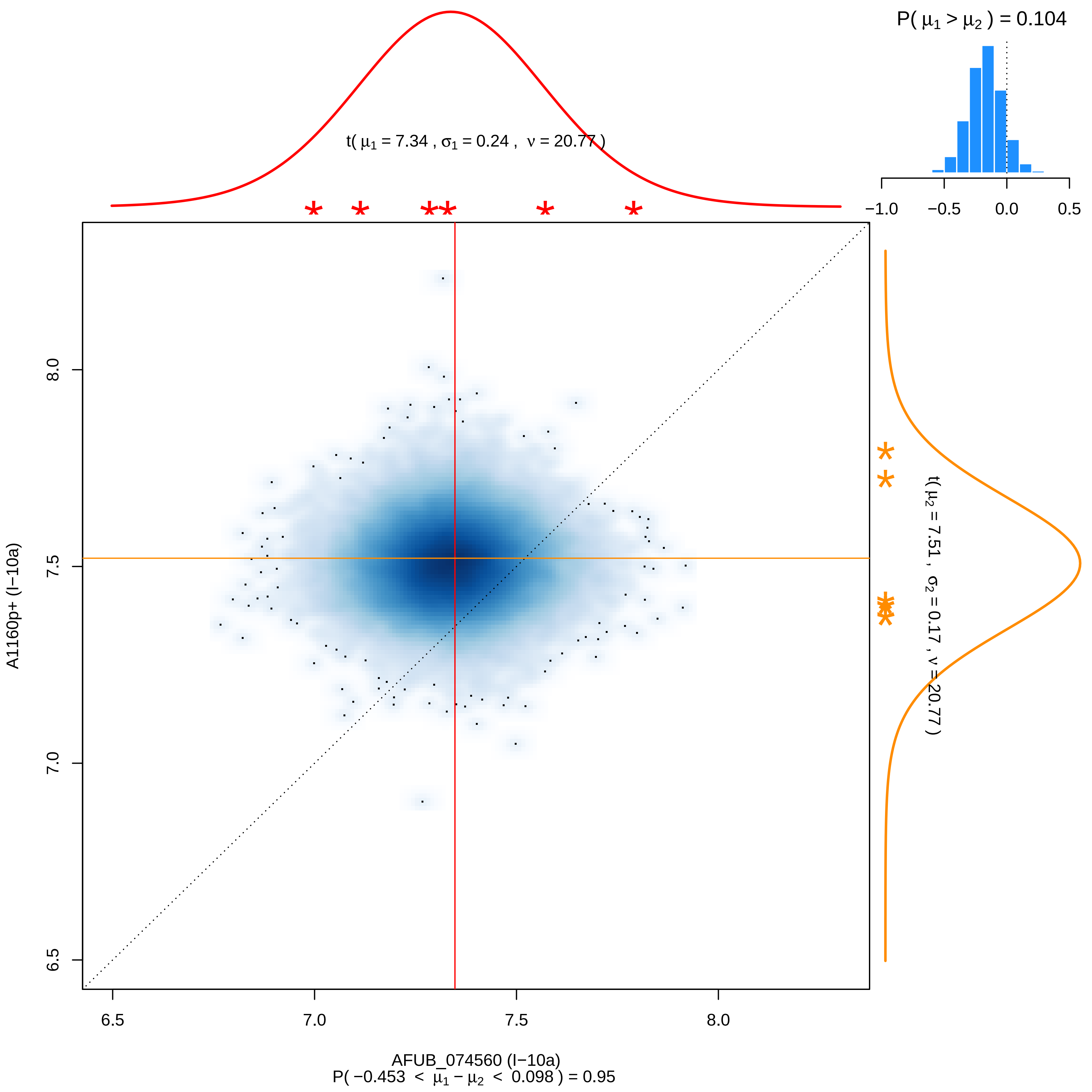

P( μ1 > μ2 ) = 0.104
−1.0
−0.5
0.0
0.5
t( μ1 = 7.34 , σ1 = 0.24 , ν = 20.77 )
t( μ2 = 7.51 , σ2 = 0.17 , ν = 20.77 )
8.0
7.5
7.0
6.5
A1160p+ (I−10a)
6.5
7.0
7.5
8.0
AFUB_074560 (I−10a)
P( −0.453 < μ1 − μ2 < 0.098 ) = 0.95

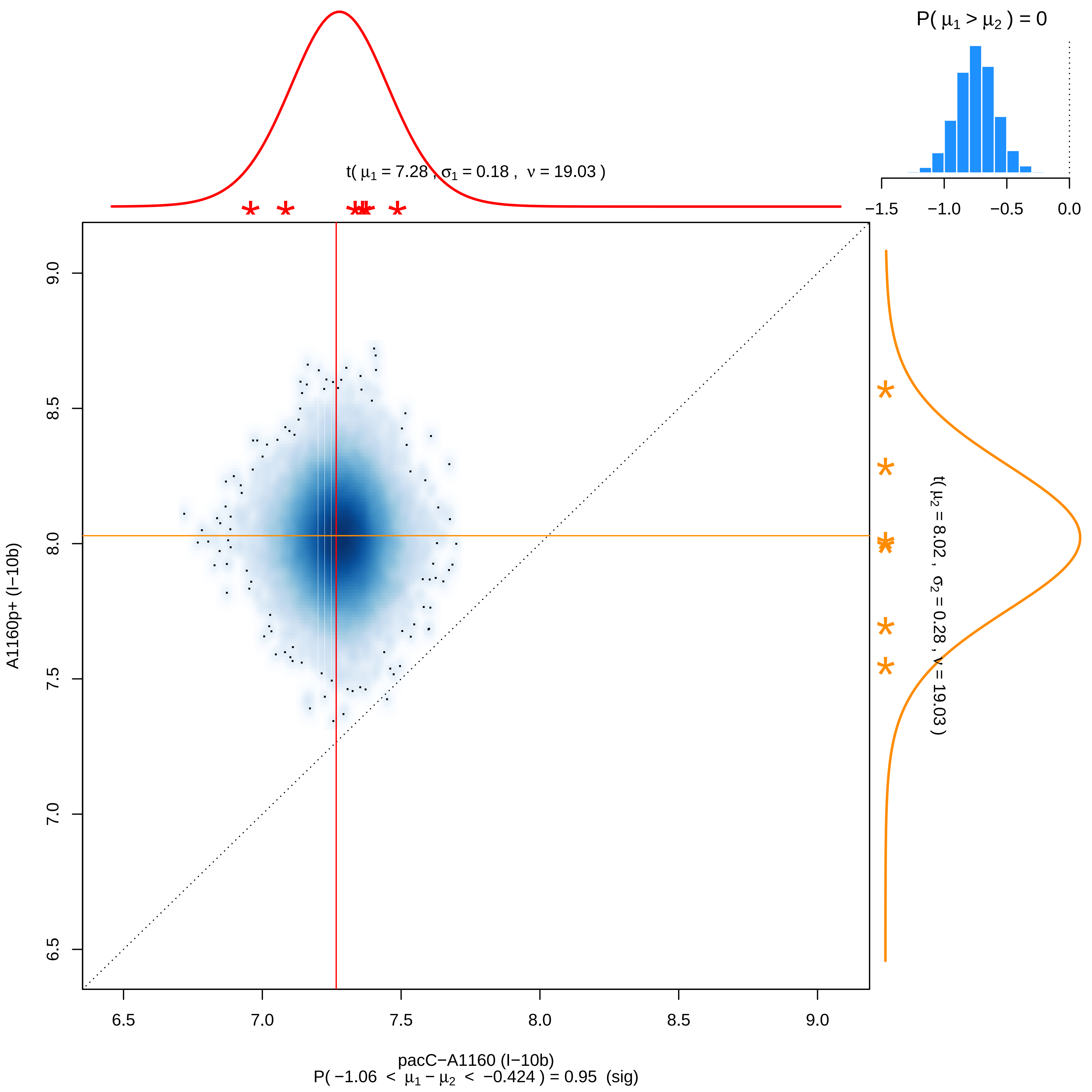

P( μ1 > μ2 ) = 0
−1.5
−1.0
−0.5
0.0
t( μ1 = 7.28 , σ1 = 0.18 , ν = 19.03 )
t( μ2 = 8.02 , σ2 = 0.28 , ν = 19.03 )
A1160p+ (I−10b)
9.0
8.5
8.0
7.5
7.0
6.5
pacC−A1160 (I−10b)
P( −1.06 < μ1 − μ2 < −0.424 ) = 0.95 (sig)

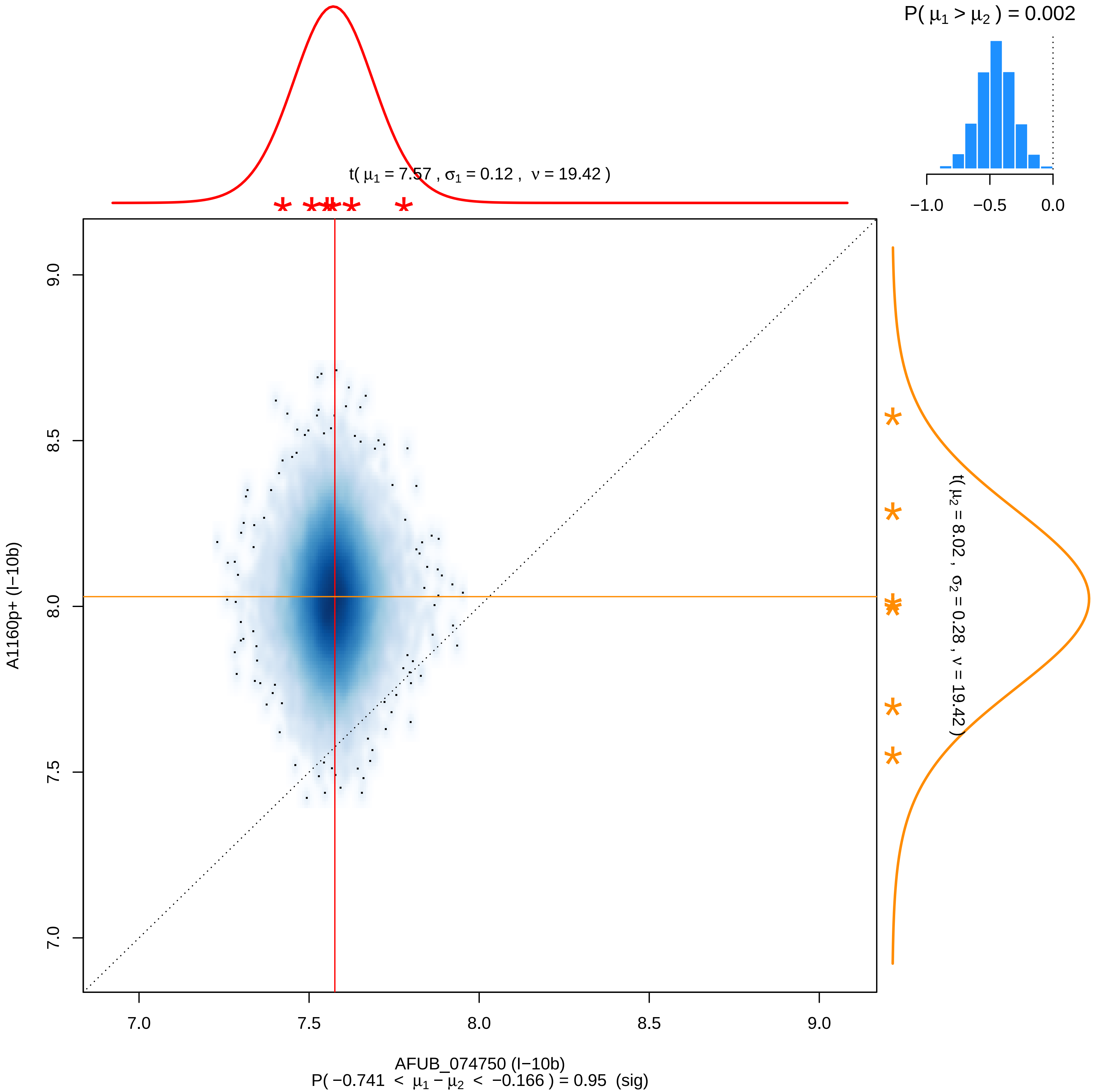

P( μ1 > μ2 ) = 0.002
−1.0
−0.5
0.0
t( μ1 = 7.57 , σ1 = 0.12 , ν = 19.42 )
9.0
8.5
8.0
7.5
7.0
A1160p+ (I−10b)
t( μ2 = 8.02 , σ2 = 0.28 , ν = 19.42 )
7.0
7.5
8.0
8.5
9.0
AFUB_074750 (I−10b)
P( −0.741 < μ1 − μ2 < −0.166 ) = 0.95 (sig)

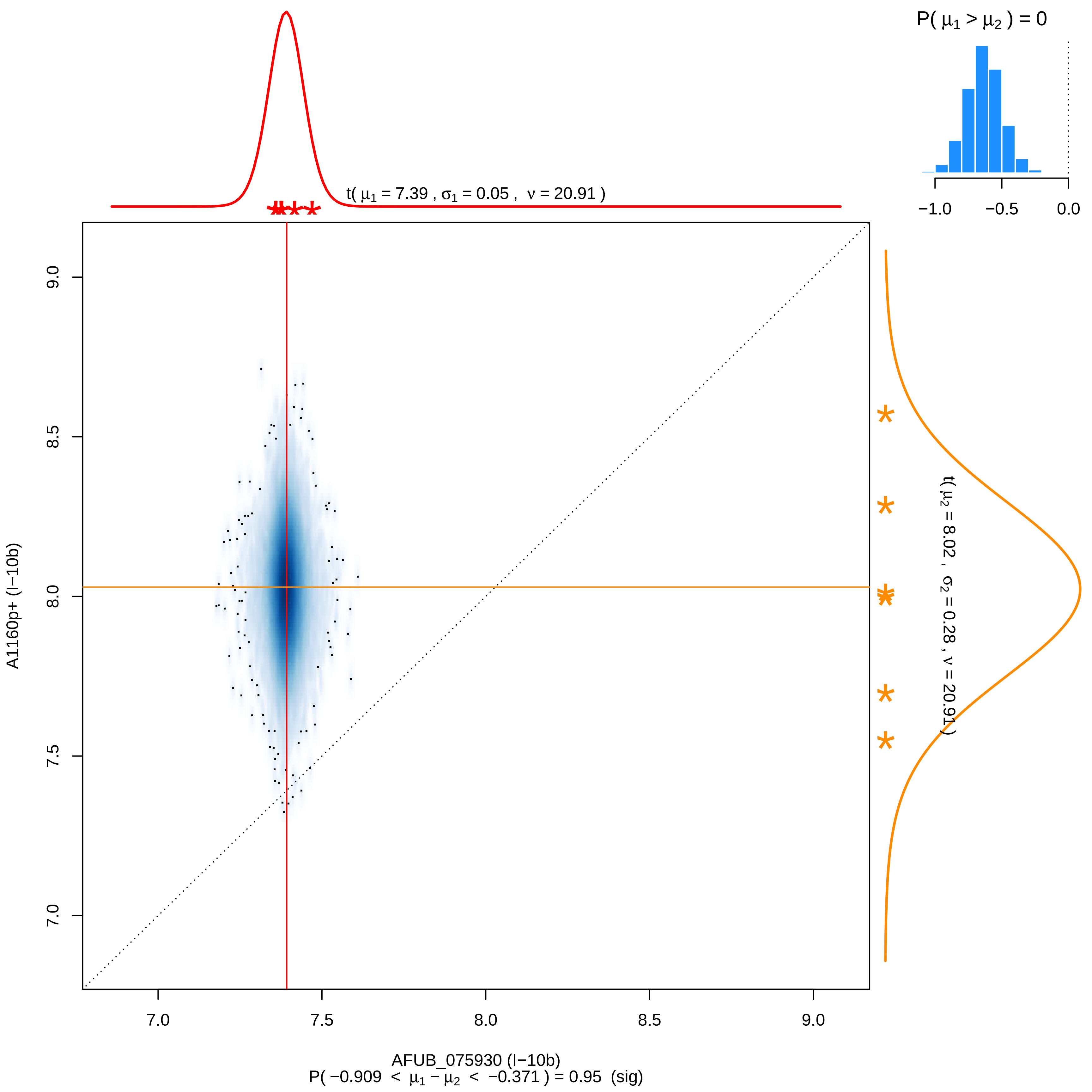

P( μ1 > μ2 ) = 0
−1.0
−0.5
0.0
t( μ1 = 7.39 , σ1 = 0.05 , ν = 20.91 )
9.0
8.5
8.0
7.5
7.0
A1160p+ (I−10b)
t( μ2 = 8.02 , σ2 = 0.28 , ν = 20.91 )
AFUB_075930 (I−10b)
P( −0.909 < μ1 − μ2 < −0.371 ) = 0.95 (sig)

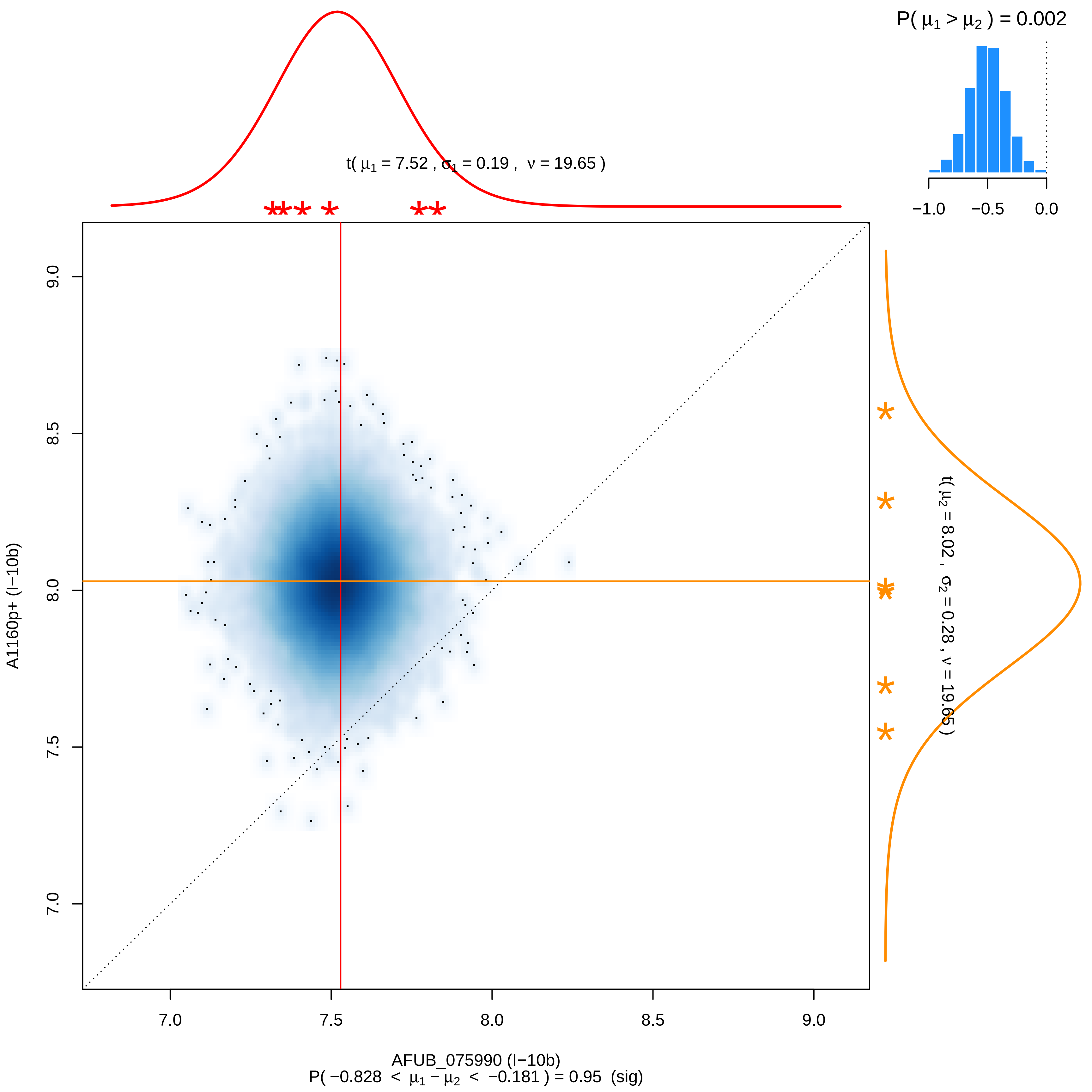

P( μ1 > μ2 ) = 0.002
−1.0
−0.5
0.0
t( μ1 = 7.52 , σ1 = 0.19 , ν = 19.65 )
t( μ2 = 8.02 , σ2 = 0.28 , ν = 19.65 )
A1160p+ (I−10b)
9.0
8.5
8.0
7.5
7.0
7.0
7.5
8.0
8.5
9.0
AFUB_075990 (I−10b)
P( −0.828 < μ1 − μ2 < −0.181 ) = 0.95 (sig)

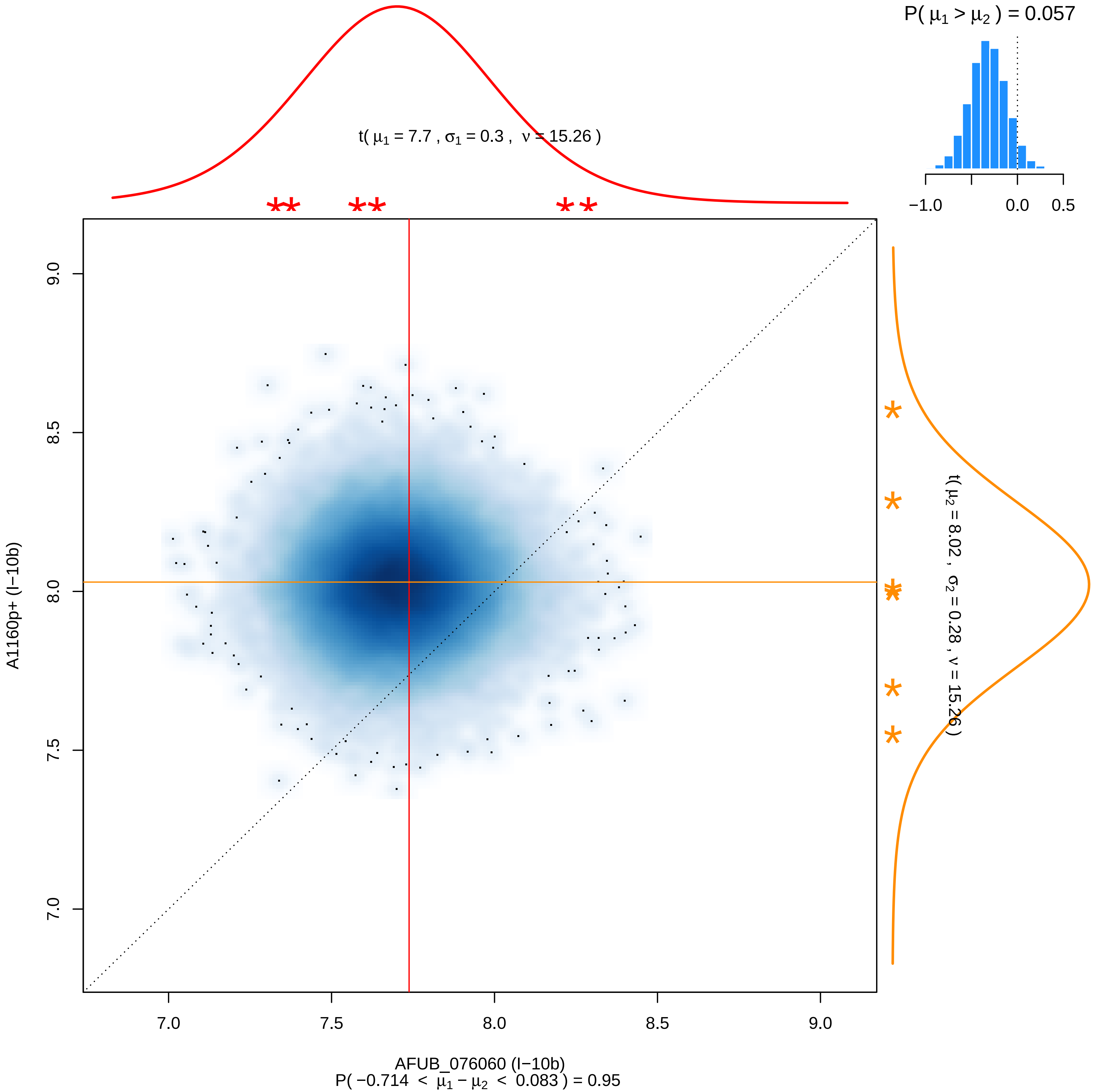

P( μ1 > μ2 ) = 0.057
−1.0
0.0
0.5
t( μ1 = 7.7 , σ1 = 0.3 , ν = 15.26 )
9.0
8.5
8.0
7.5
7.0
A1160p+ (I−10b)
t( μ2 = 8.02 , σ2 = 0.28 , ν = 15.26 )
7.0
7.5
8.0
8.5
9.0
AFUB_076060 (I−10b)
P( −0.714 < μ1 − μ2 < 0.083 ) = 0.95

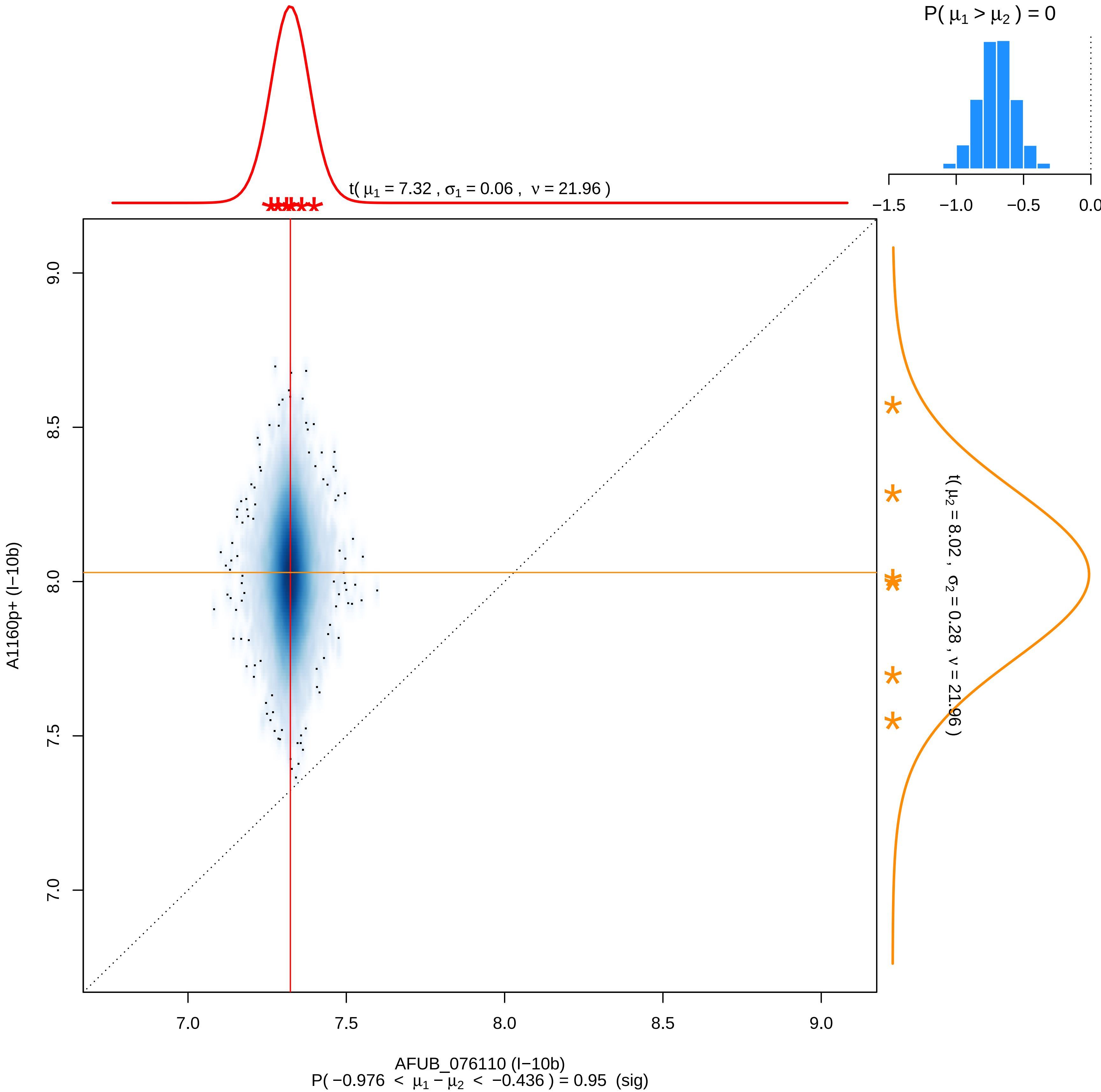

P( μ1 > μ2 ) = 0
−1.5
−1.0
−0.5
0.0
t( μ1 = 7.32 , σ1 = 0.06 , ν = 21.96 )
9.0
8.5
8.0
7.5
7.0
A1160p+ (I−10b)
t( μ2 = 8.02 , σ2 = 0.28 , ν = 21.96 )
7.0
7.5
8.0
8.5
9.0
AFUB_076110 (I−10b)
P( −0.976 < μ1 − μ2 < −0.436 ) = 0.95 (sig)

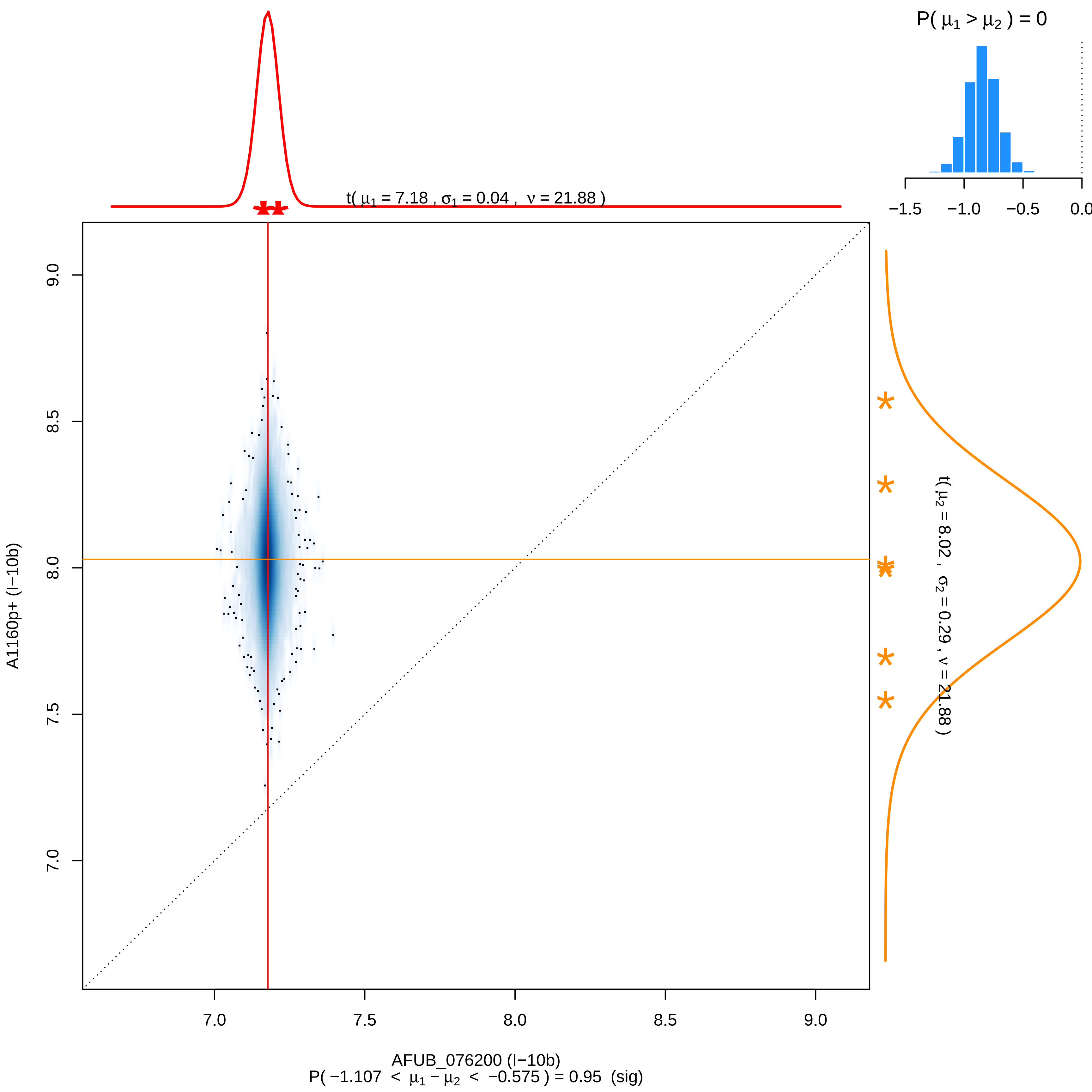

P( μ1 > μ2 ) = 0
−1.5
−1.0
−0.5
0.0
t( μ1 = 7.18 , σ1 = 0.04 , ν = 21.88 )
t( μ2 = 8.02 , σ2 = 0.29 , ν = 21.88 )
A1160p+ (I−10b)
9.0
8.5
8.0
7.5
7.0
AFUB_076200 (I−10b)
P( −1.107 < μ1 − μ2 < −0.575 ) = 0.95 (sig)

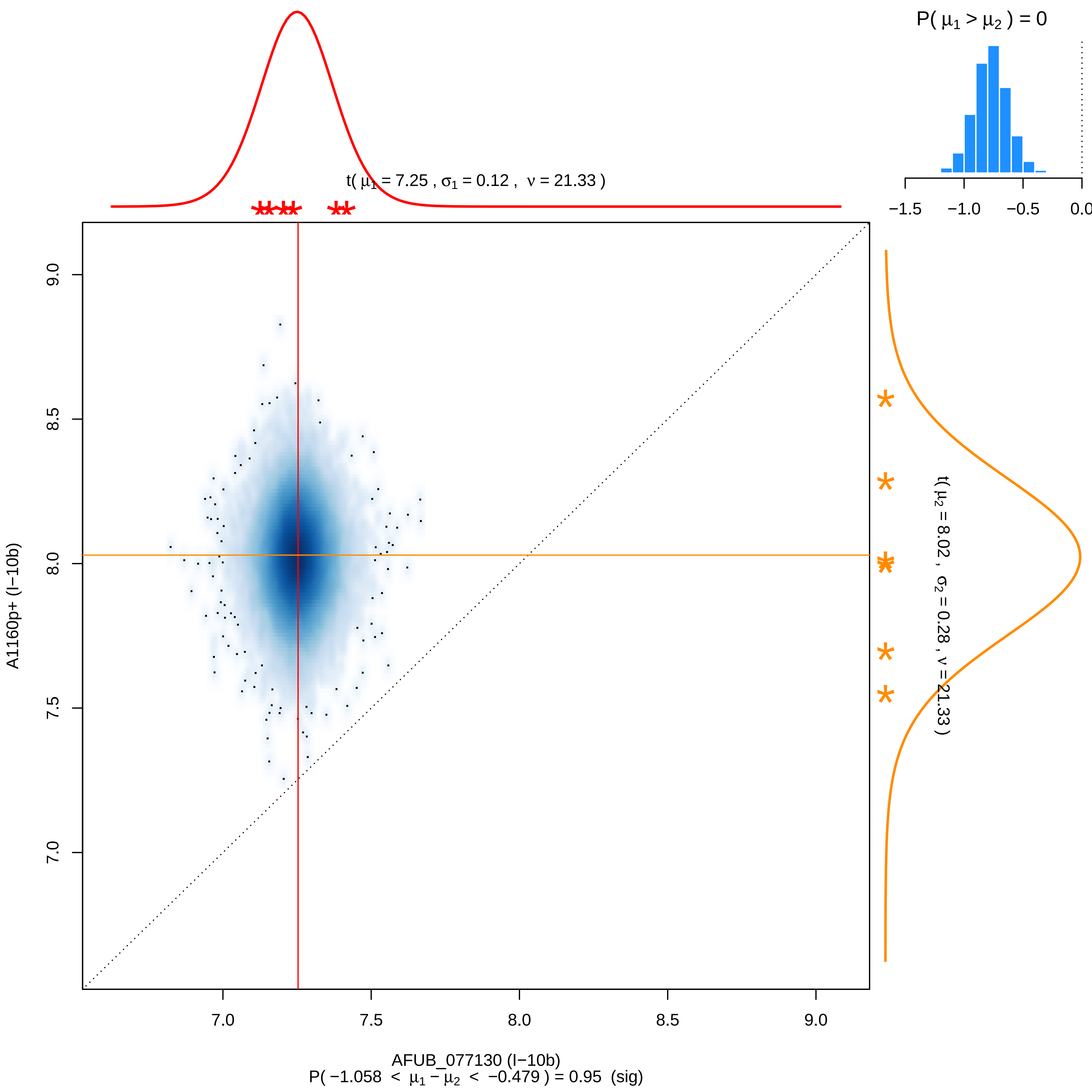

P( μ1 > μ2 ) = 0
−1.5
−1.0
−0.5
0.0
t( μ1 = 7.25 , σ1 = 0.12 , ν = 21.33 )
9.0
8.5
8.0
7.5
7.0
A1160p+ (I−10b)
t( μ2 = 8.02 , σ2 = 0.28 , ν = 21.33 )
7.0
7.5
8.0
8.5
9.0
AFUB_077130 (I−10b)
P( −1.058 < μ1 − μ2 < −0.479 ) = 0.95 (sig)

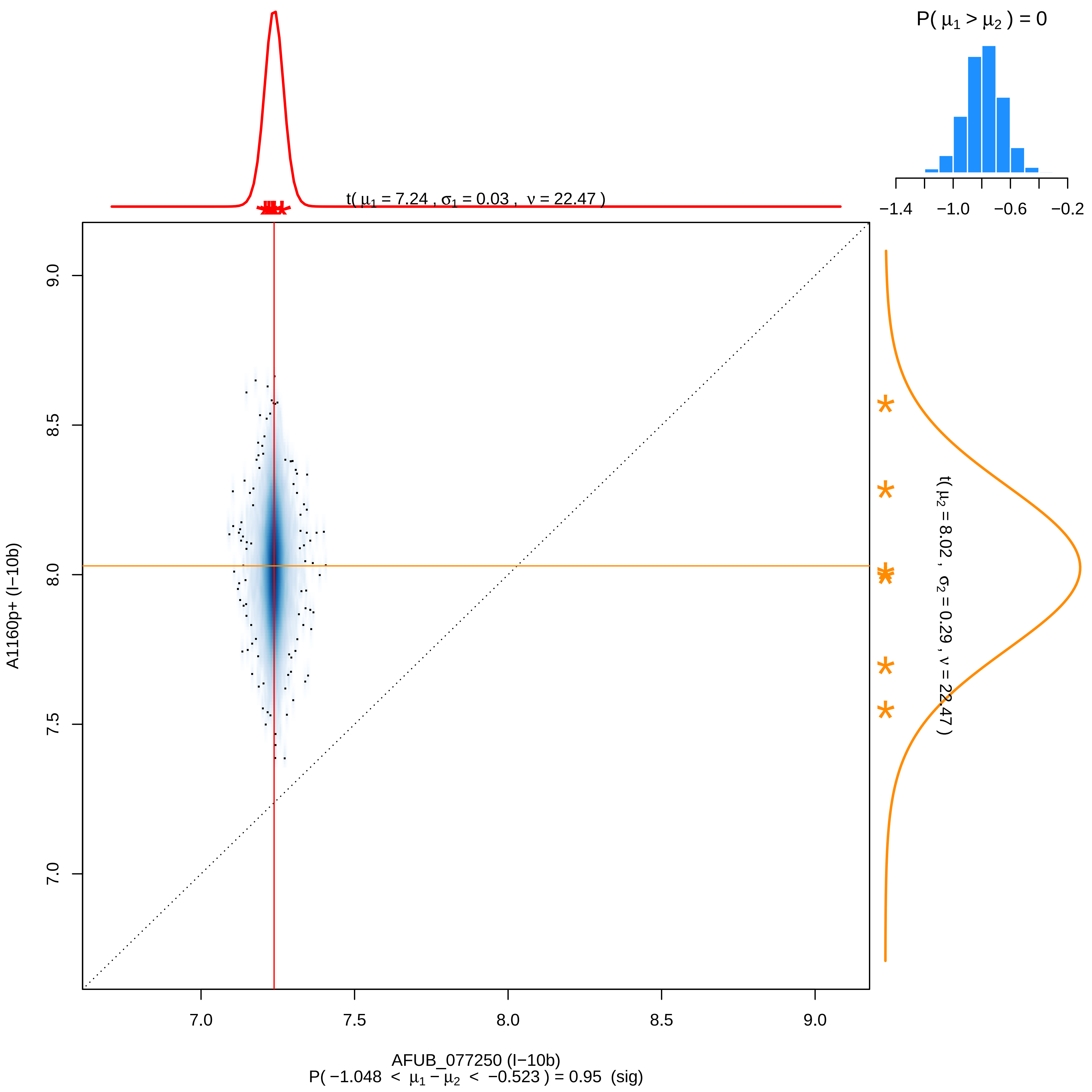

P( μ1 > μ2 ) = 0
−1.4
−1.0
−0.6
−0.2
t( μ1 = 7.24 , σ1 = 0.03 , ν = 22.47 )
9.0
8.5
8.0
7.5
7.0
A1160p+ (I−10b)
t( μ2 = 8.02 , σ2 = 0.29 , ν = 22.47 )
7.0
7.5
8.0
8.5
9.0
AFUB_077250 (I−10b)
P( −1.048 < μ1 − μ2 < −0.523 ) = 0.95 (sig)

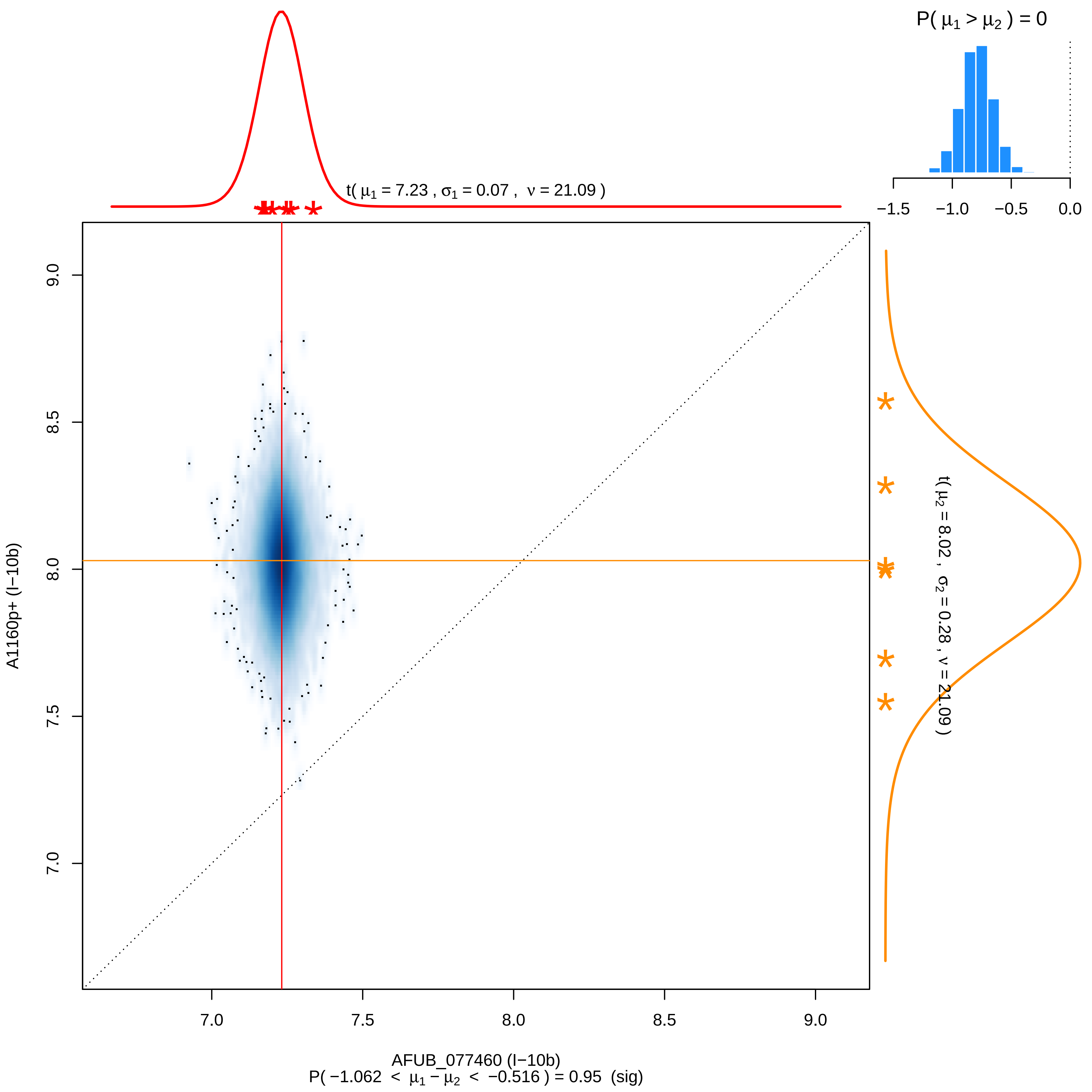

P( μ1 > μ2 ) = 0
−1.5 −1.0 −0.5 0.0
t( μ1 = 7.23 , σ1 = 0.07 , ν = 21.09 )
t( μ2 = 8.02 , σ2 = 0.28 , ν = 21.09 )
9.0 8.5 8.0 7.5 7.0
A1160p+ (I−10b)
7.0 7.5 8.0 8.5 9.0
AFUB_077460 (I−10b)
P( −1.062 < μ1 − μ2 < −0.516 ) = 0.95 (sig)

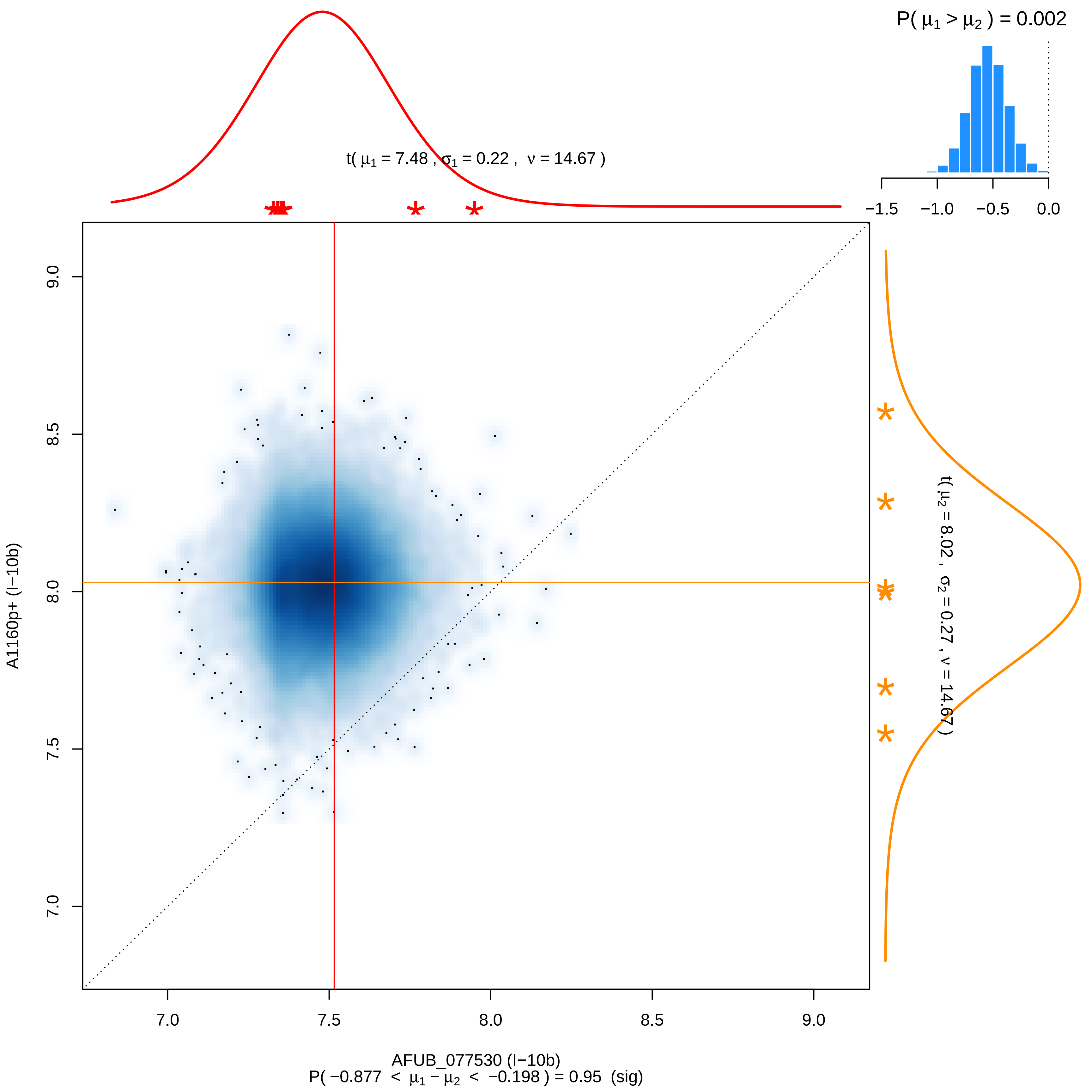

P( μ1 > μ2 ) = 0.002
−1.5
−1.0
−0.5
0.0
t( μ1 = 7.48 , σ1 = 0.22 , ν = 14.67 )
t( μ2 = 8.02 , σ2 = 0.27 , ν = 14.67 )
9.0
8.5
8.0
7.5
7.0
A1160p+ (I−10b)
7.0
7.5
8.0
8.5
9.0
AFUB_077530 (I−10b)
P( −0.877 < μ1 − μ2 < −0.198 ) = 0.95 (sig)

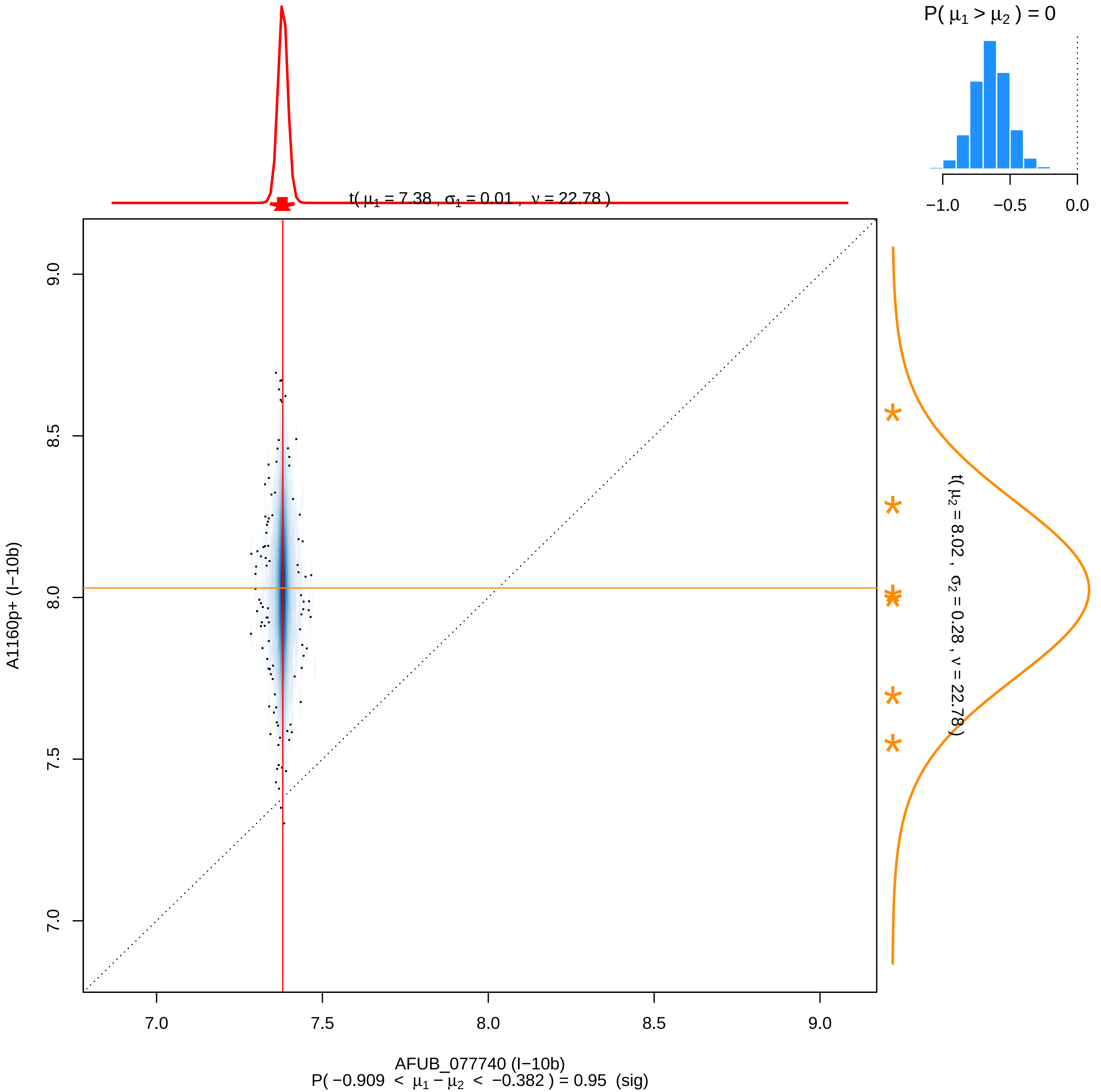

P( $\mu_1 > \mu_2$ ) = 0
−1.0
−0.5
0.0
t( $\mu_1$ = 7.38 , $\sigma_1$ = 0.01 , $\nu$ = 22.78 )
t( $\mu_2$ = 8.02 , $\sigma_2$ = 0.28 , $\nu$ = 22.78 )
A1160p+ (I−10b)
9.0
8.5
8.0
7.5
7.0
7.0
7.5
8.0
8.5
9.0
AFUB_077740 (I−10b)
P( −0.909 < $\mu_1 - \mu_2$ < −0.382 ) = 0.95 (sig)

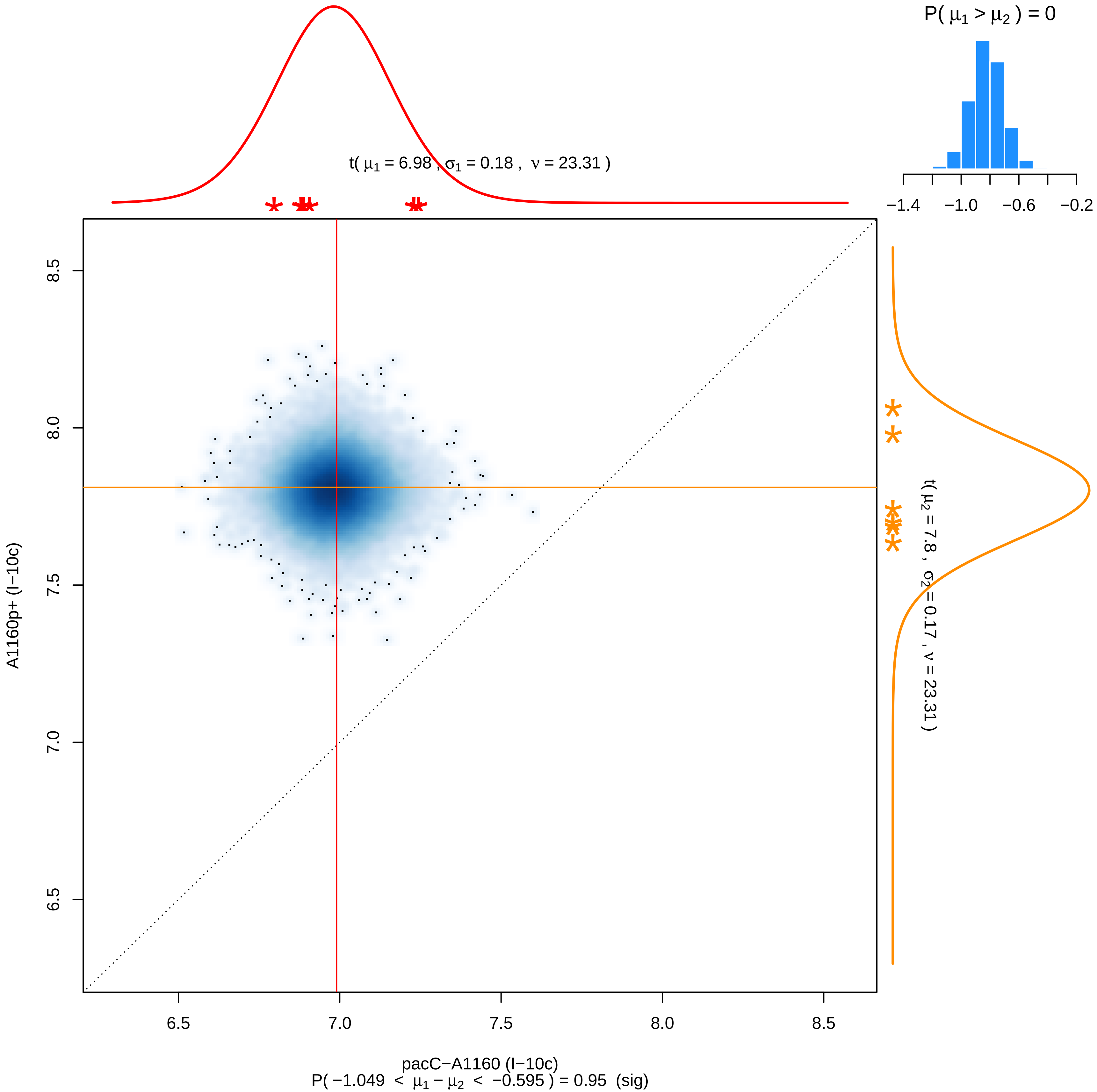

P( μ1 > μ2 ) = 0
−1.4 −1.0 −0.6 −0.2
t( μ1 = 6.98 , σ1 = 0.18 , ν = 23.31 )
t( μ2 = 7.8 , σ2 = 0.17 , ν = 23.31 )
8.5
8.0
7.5
7.0
6.5
A1160p+ (I−10c)
6.5 7.0 7.5 8.0 8.5
pacC−A1160 (I−10c)
P( −1.049 < μ1 − μ2 < −0.595 ) = 0.95 (sig)

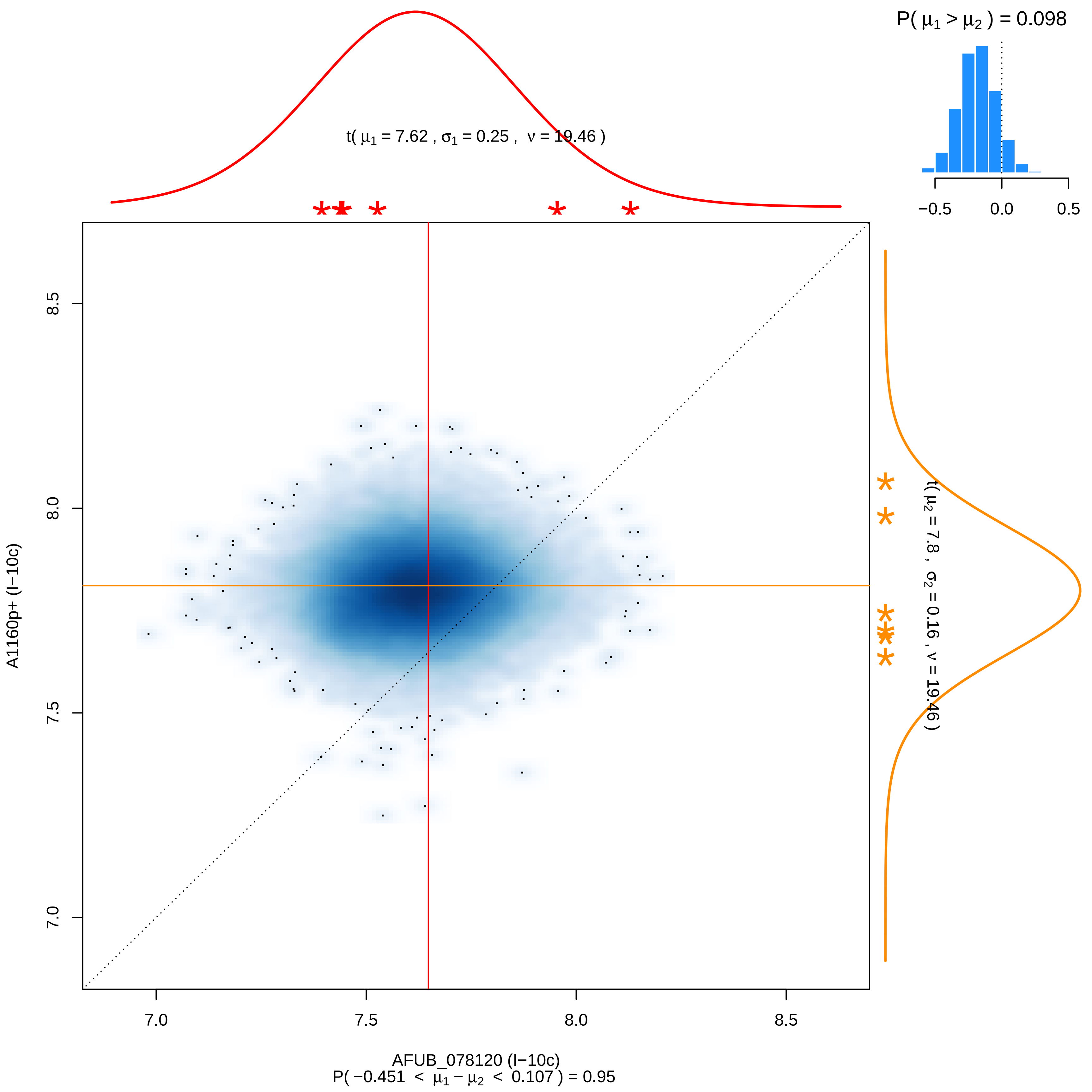

P( μ1 > μ2 ) = 0.098
−0.5
0.0
0.5
t( μ1 = 7.62 , σ1 = 0.25 , ν = 19.46 )
t( μ2 = 7.8 , σ2 = 0.16 , ν = 19.46 )
A1160p+ (I−10c)
8.5
8.0
7.5
7.0
7.0
7.5
8.0
8.5
AFUB_078120 (I−10c)
P( −0.451 < μ1 − μ2 < 0.107 ) = 0.95

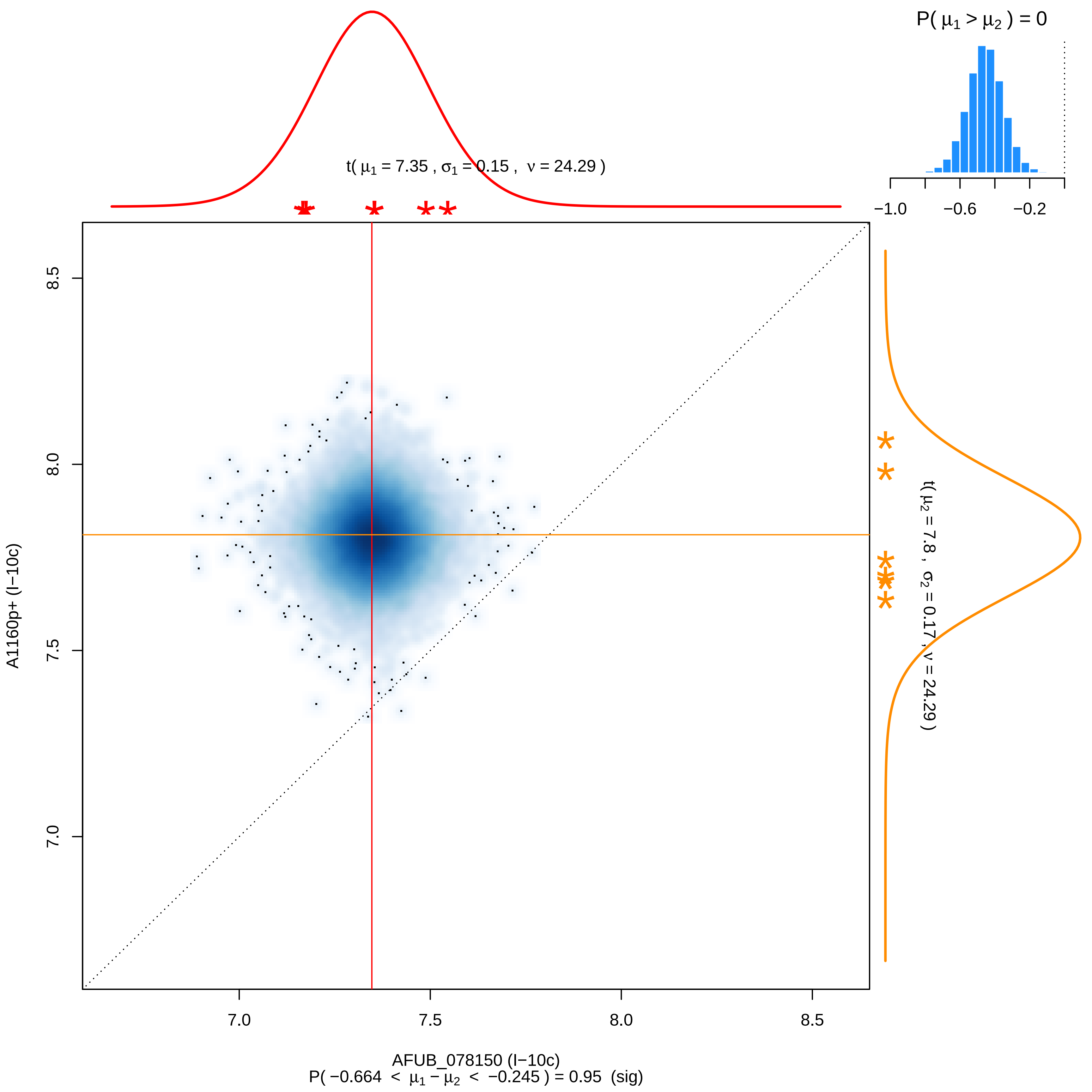
P( μ1 > μ2 ) = 0
−1.0
−0.6
−0.2
t( μ1 = 7.35 , σ1 = 0.15 , ν = 24.29 )
8.5
8.0
7.5
7.0
A1160p+ (I−10c)
t( μ2 = 7.8 , σ2 = 0.17 , ν = 24.29 )
AFUB_078150 (I−10c)
P( −0.664 < μ1 − μ2 < −0.245 ) = 0.95 (sig)

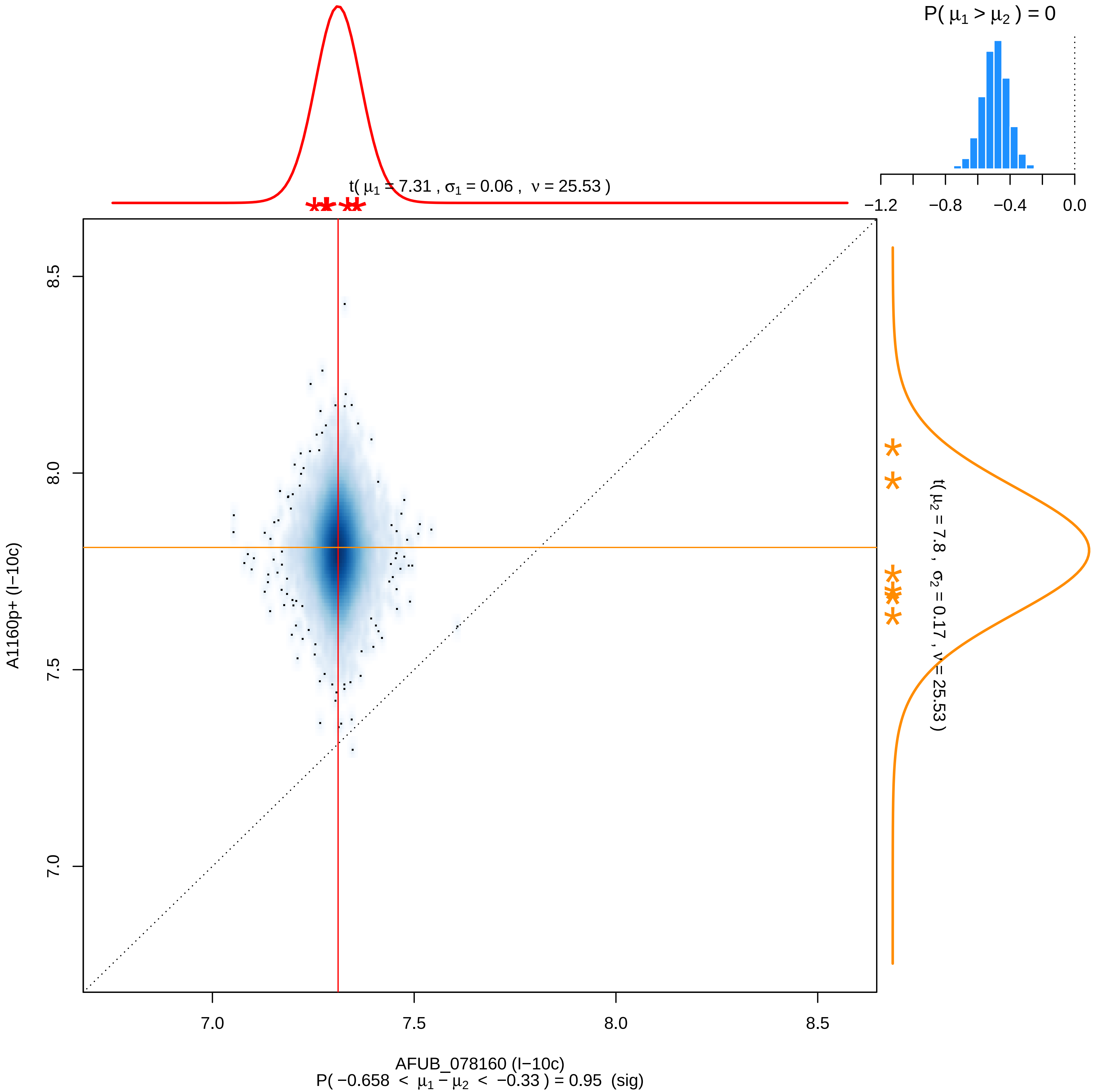

P( μ1 > μ2 ) = 0
−1.2
−0.8
−0.4
0.0
t( μ1 = 7.31 , σ1 = 0.06 , ν = 25.53 )
t( μ2 = 7.8 , σ2 = 0.17 , ν = 25.53 )
A1160p+ (I−10c)
8.5
8.0
7.5
7.0
AFUB_078160 (I−10c)
P( −0.658 < μ1 − μ2 < −0.33 ) = 0.95 (sig)

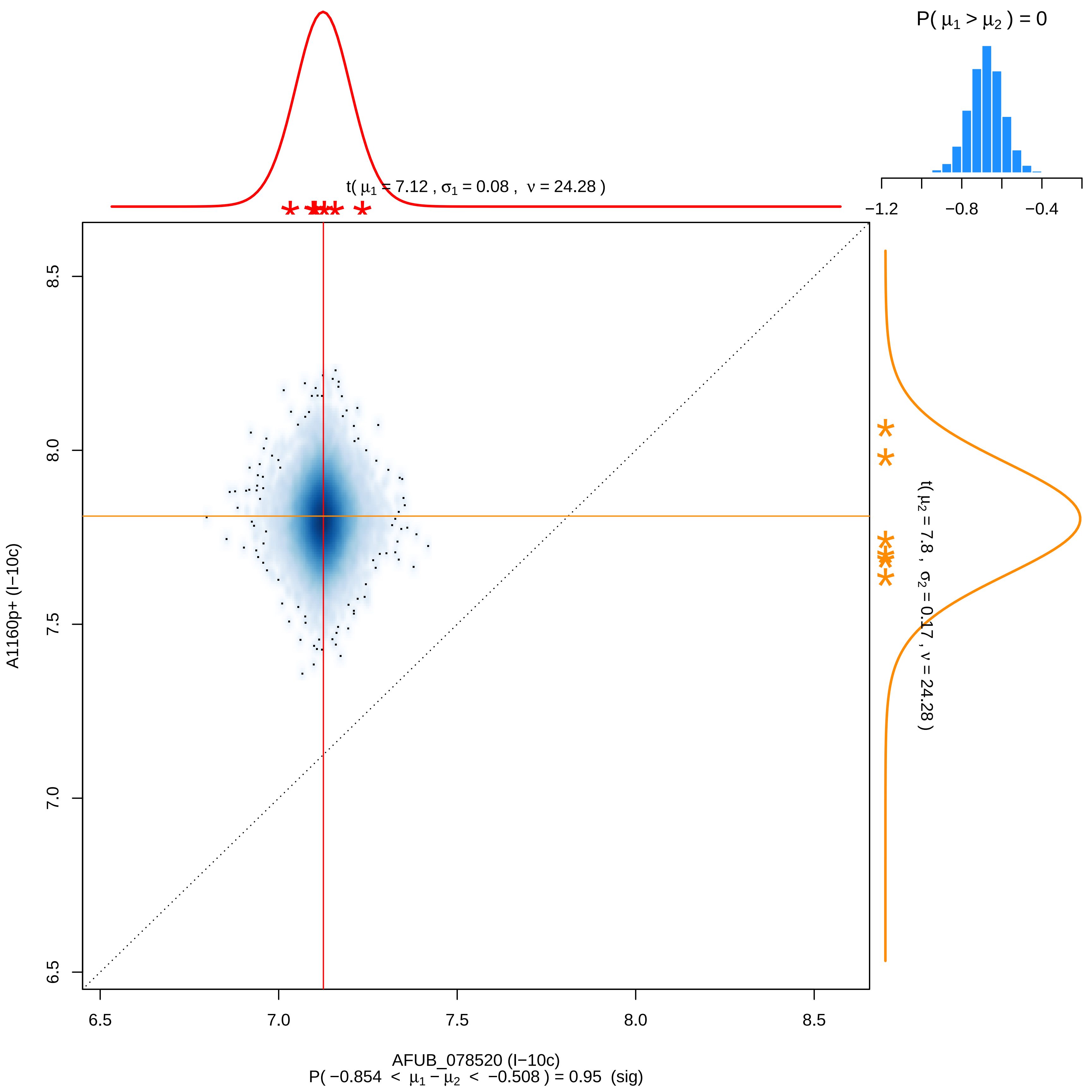

P( μ1 > μ2 ) = 0
−1.2
−0.8
−0.4
t( μ1 = 7.12 , σ1 = 0.08 , ν = 24.28 )
t( μ2 = 7.8 , σ2 = 0.17 , ν = 24.28 )
A1160p+ (I−10c)
6.5
7.0
7.5
8.0
8.5
AFUB_078520 (I−10c)
P( −0.854 < μ1 − μ2 < −0.508 ) = 0.95 (sig)

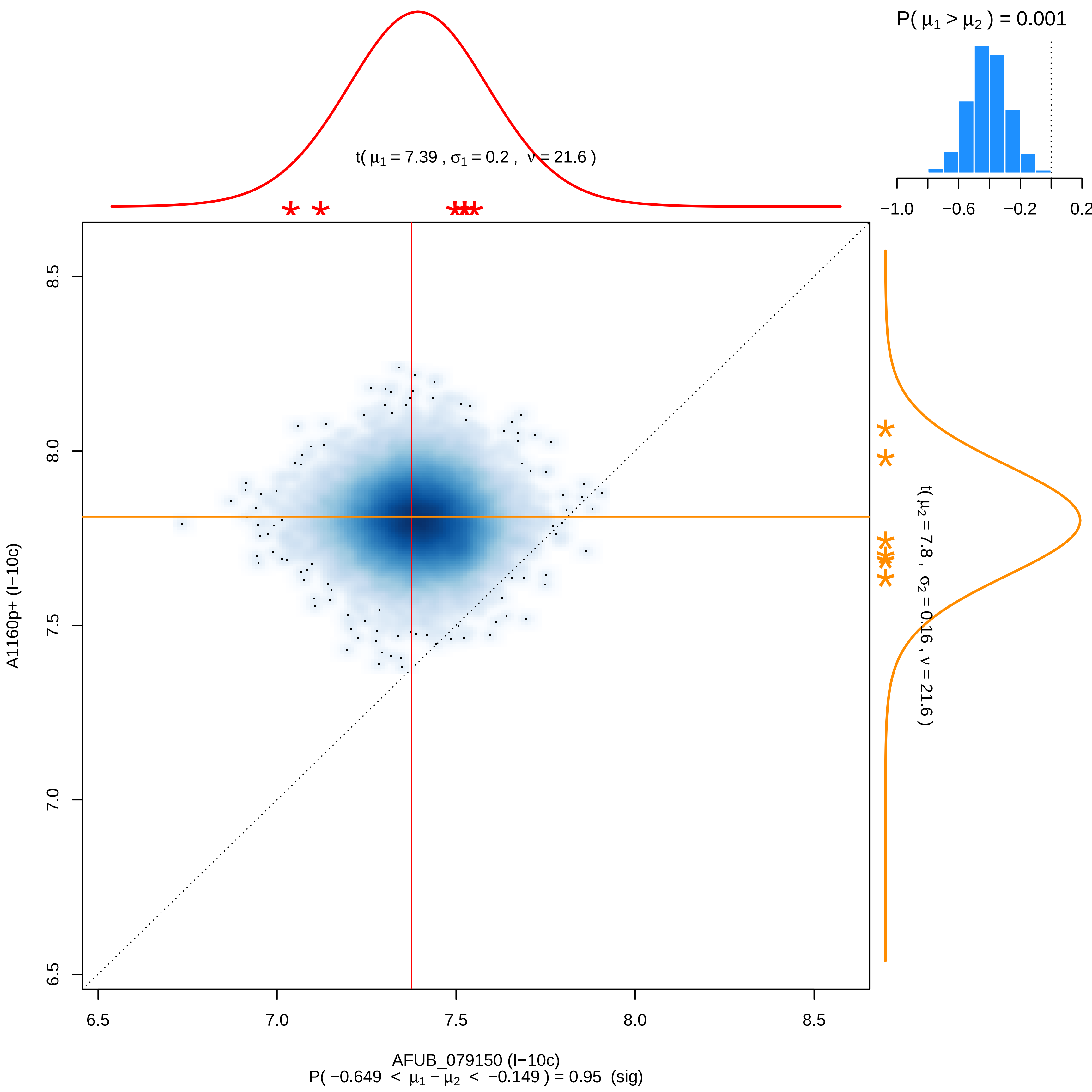

P( μ1 > μ2 ) = 0.001
−1.0
−0.6
−0.2
0.2
t( μ1 = 7.39 , σ1 = 0.2 , ν = 21.6 )
t( μ2 = 7.8 , σ2 = 0.16 , ν = 21.6 )
8.5
8.0
7.5
7.0
6.5
A1160p+ (I−10c)
6.5
7.0
7.5
8.0
8.5
AFUB_079150 (I−10c)
P( −0.649 < μ1 − μ2 < −0.149 ) = 0.95 (sig)

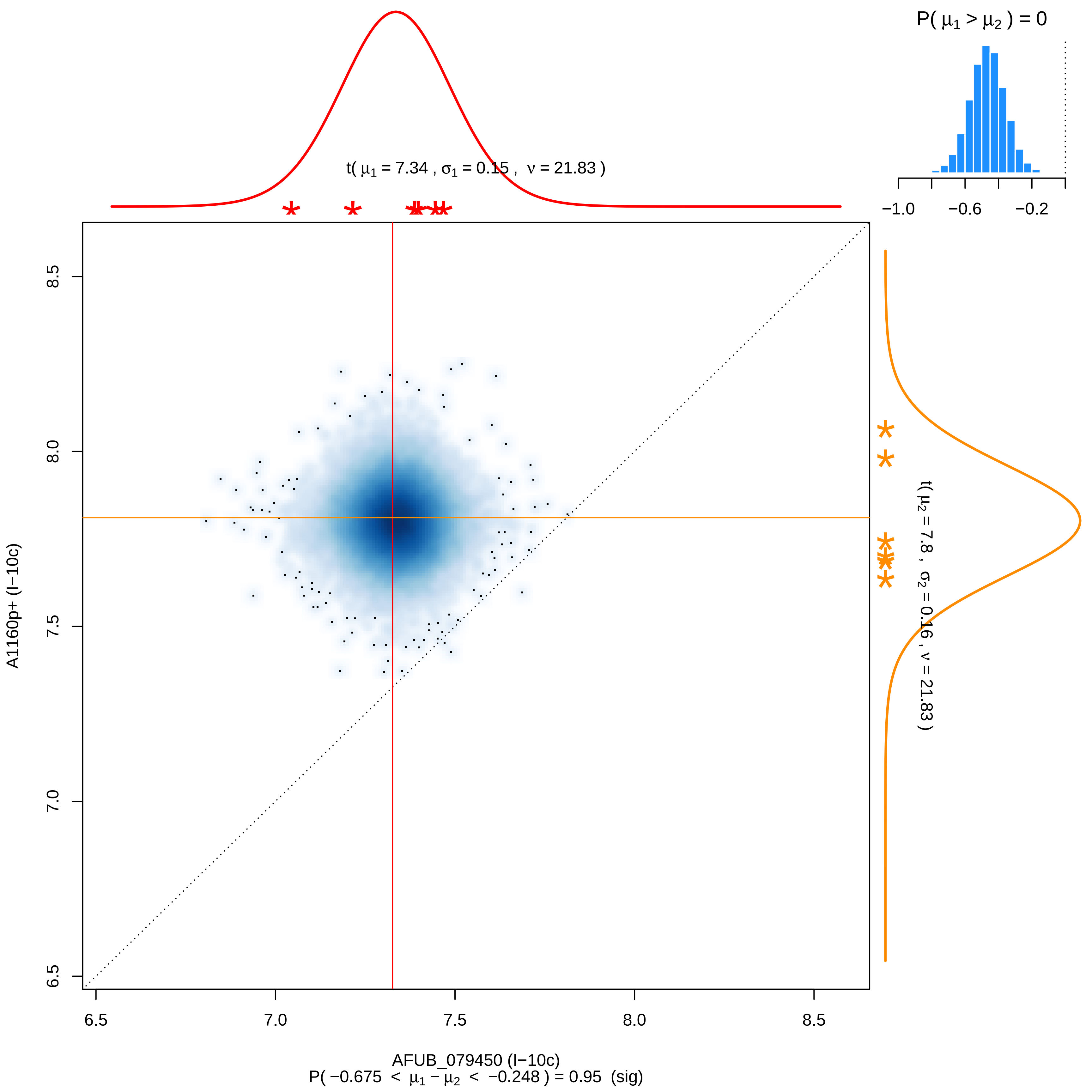

P( μ1 > μ2 ) = 0
−1.0
−0.6
−0.2
t( μ1 = 7.34 , σ1 = 0.15 , ν = 21.83 )
8.5
8.0
7.5
7.0
6.5
A1160p+ (I−10c)
t( μ2 = 7.8 , σ2 = 0.16 , ν = 21.83 )
6.5
7.0
7.5
8.0
8.5
AFUB_079450 (I−10c)
P( −0.675 < μ1 − μ2 < −0.248 ) = 0.95 (sig)

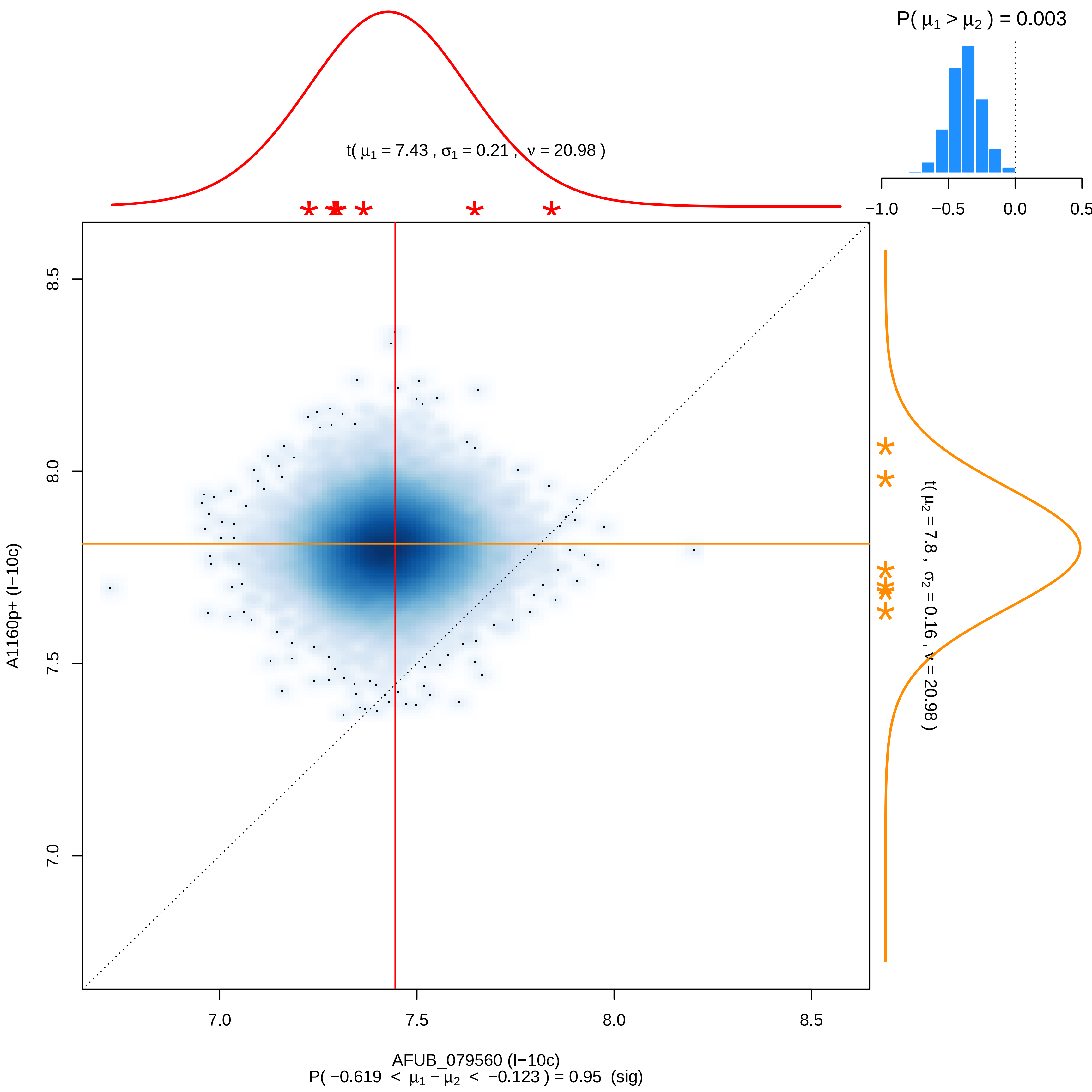

P( μ1 > μ2 ) = 0.003
−1.0
−0.5
0.0
0.5
t( μ1 = 7.43 , σ1 = 0.21 , ν = 20.98 )
t( μ2 = 7.8 , σ2 = 0.16 , ν = 20.98 )
A1160p+ (I−10c)
8.5
8.0
7.5
7.0
AFUB_079560 (I−10c)
P( −0.619 < μ1 − μ2 < −0.123 ) = 0.95 (sig)

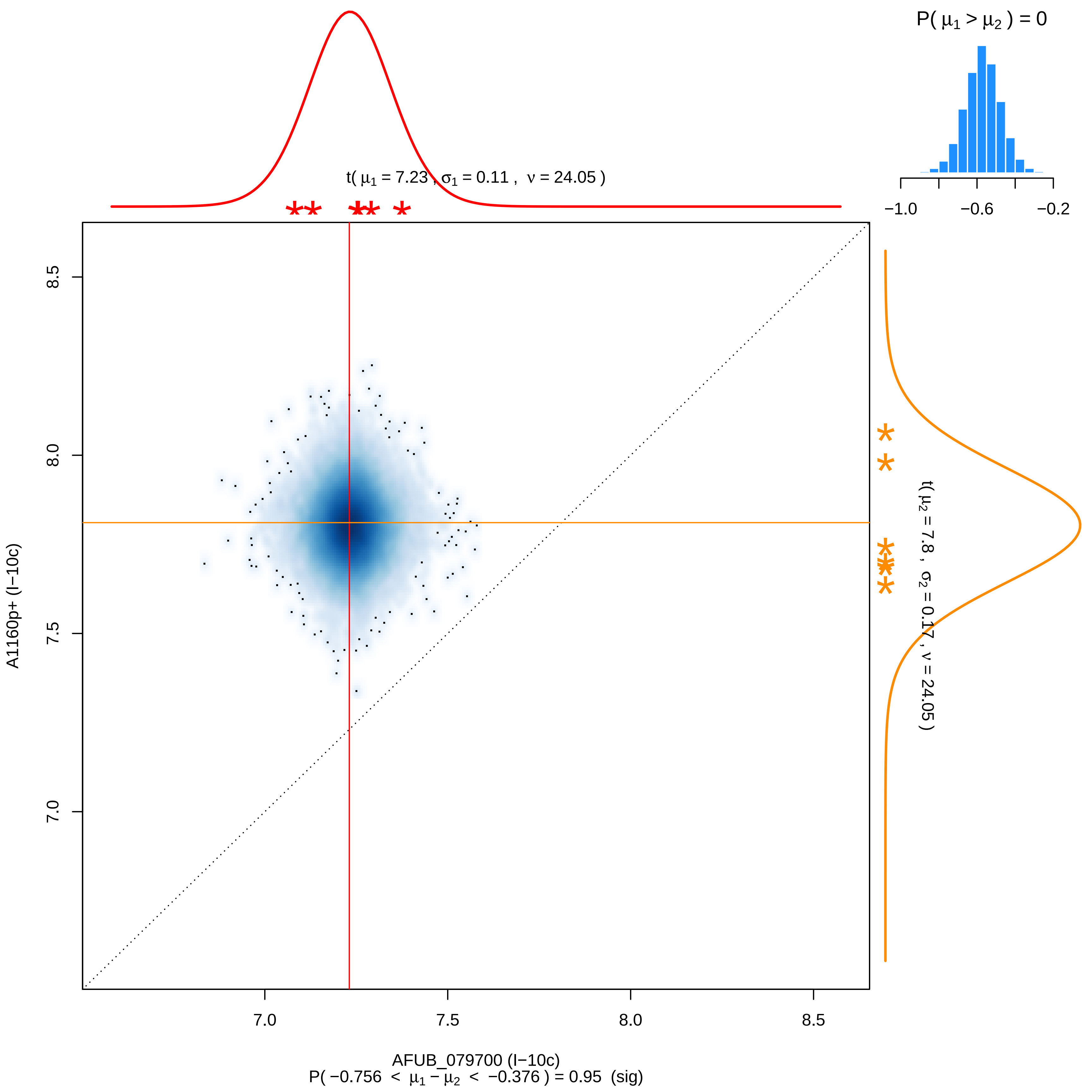

P( μ1 > μ2 ) = 0
−1.0
−0.6
−0.2
t( μ1 = 7.23 , σ1 = 0.11 , ν = 24.05 )
t( μ2 = 7.8 , σ2 = 0.17 , ν = 24.05 )
8.5
8.0
7.5
7.0
A1160p+ (I−10c)
7.0
7.5
8.0
8.5
AFUB_079700 (I−10c)
P( −0.756 < μ1 − μ2 < −0.376 ) = 0.95 (sig)

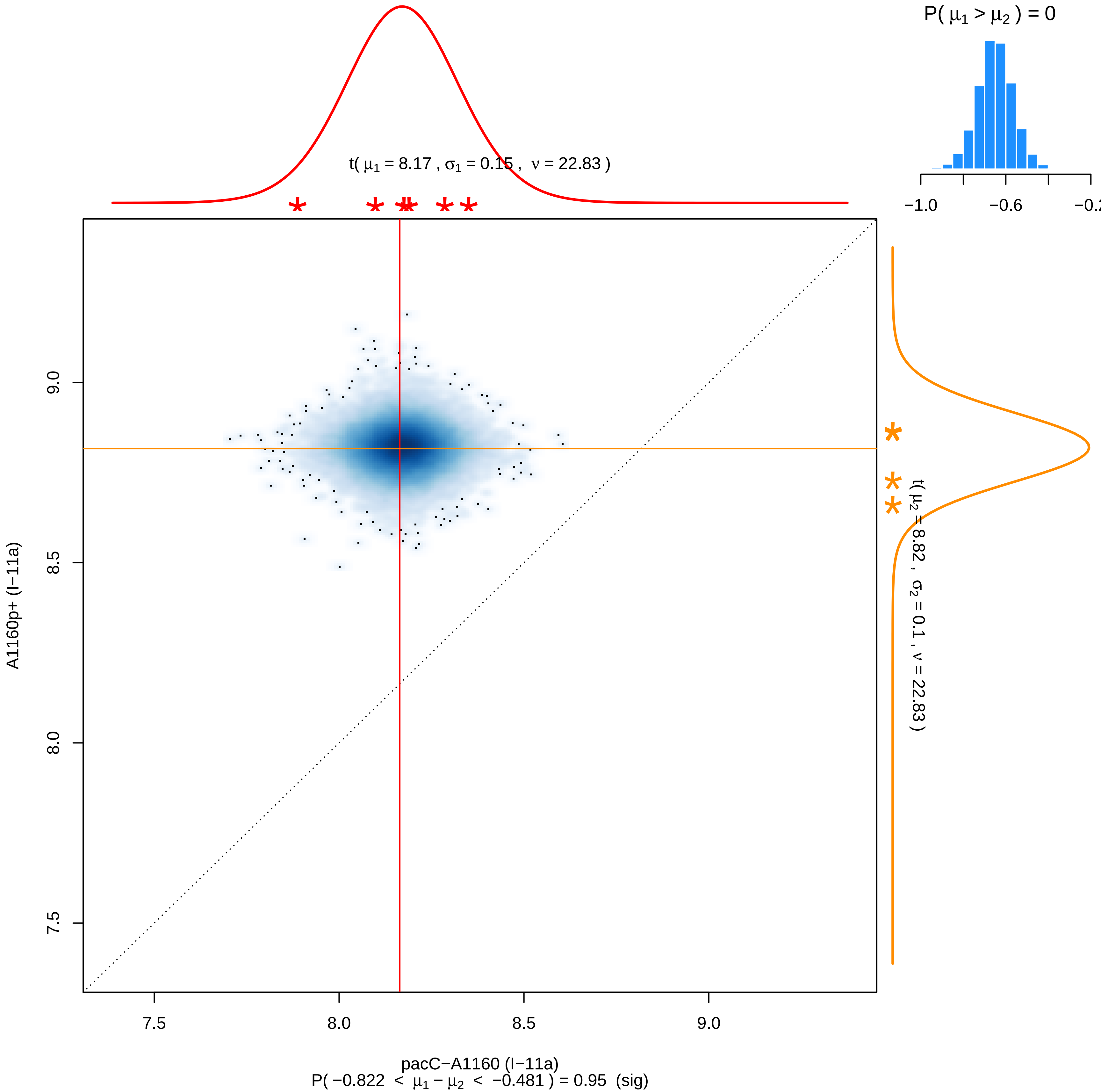

P( μ1 > μ2 ) = 0
−1.0
−0.6
−0.2
t( μ1 = 8.17 , σ1 = 0.15 , ν = 22.83 )
t( μ2 = 8.82 , σ2 = 0.1 , ν = 22.83 )
9.0
8.5
8.0
7.5
A1160p+ (I−11a)
7.5
8.0
8.5
9.0
pacC−A1160 (I−11a)
P( −0.822 < μ1 − μ2 < −0.481 ) = 0.95 (sig)

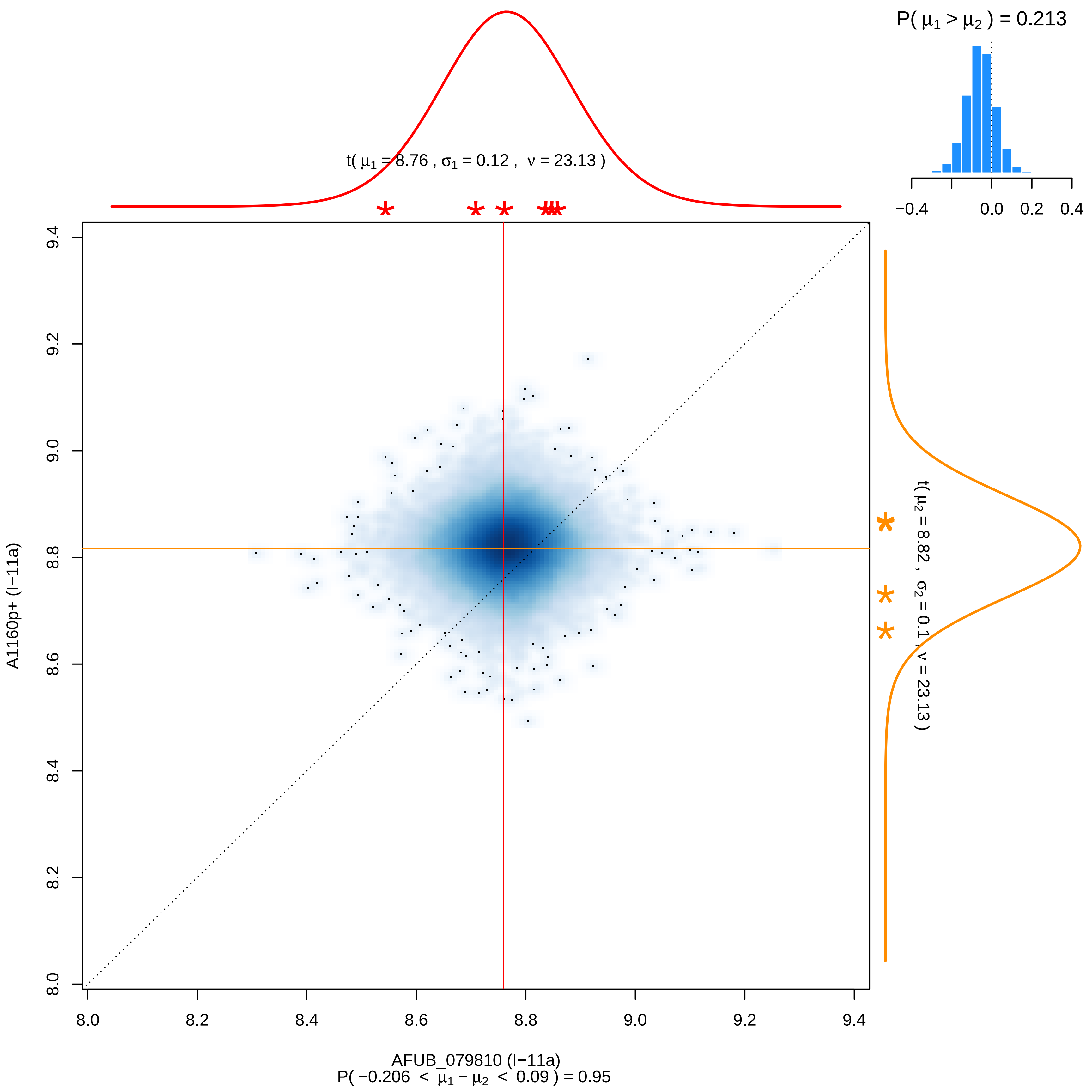

P( μ1 > μ2 ) = 0.213
−0.4
0.0
0.2
0.4
t( μ1 = 8.76 , σ1 = 0.12 , ν = 23.13 )
t( μ2 = 8.82 , σ2 = 0.1 , ν = 23.13 )
A1160p+ (I−11a)
9.4
9.2
9.0
8.8
8.6
8.4
8.2
8.0
8.0
8.2
8.4
8.6
8.8
9.0
9.2
9.4
AFUB_079810 (I−11a)
P( −0.206 < μ1 − μ2 < 0.09 ) = 0.95

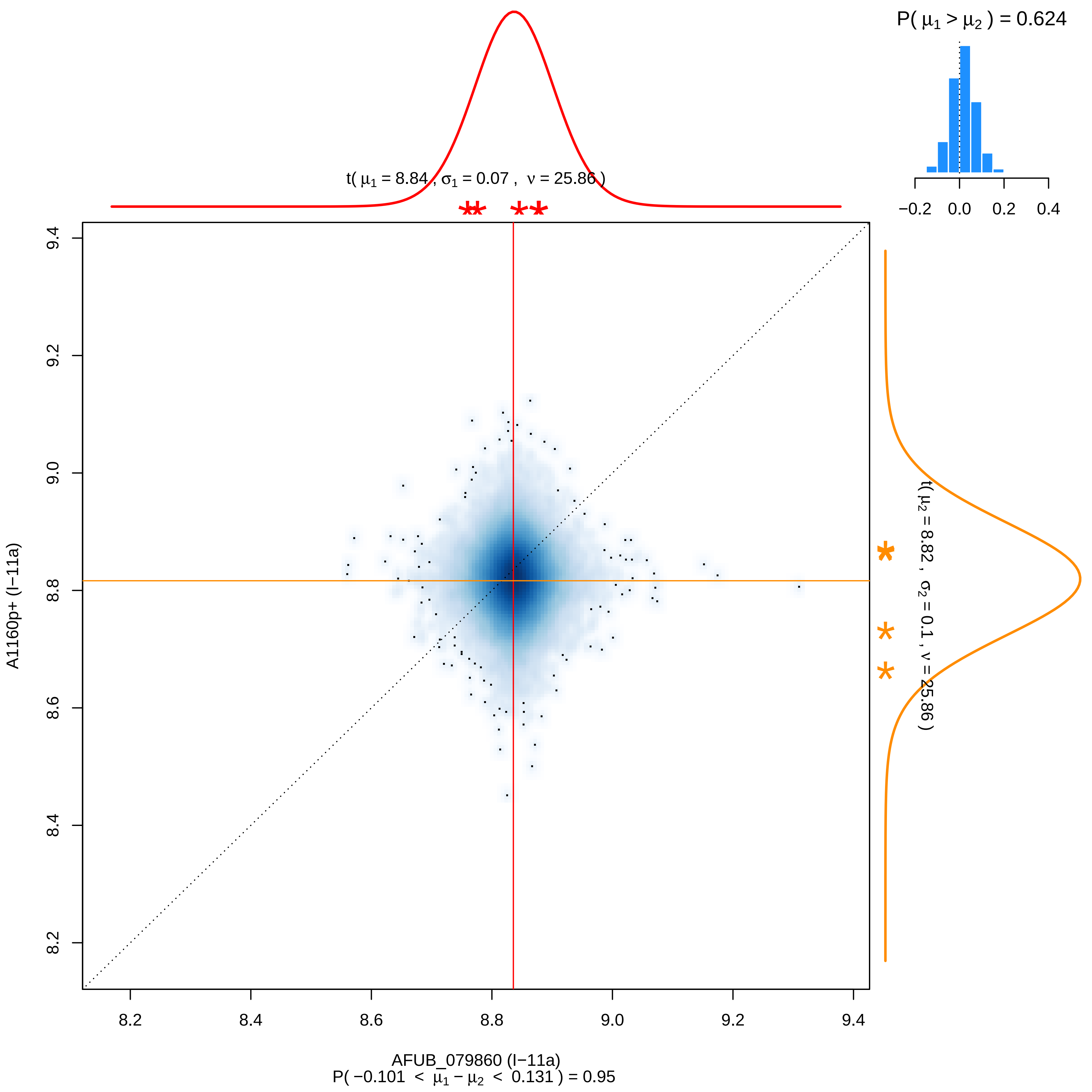

P( μ1 > μ2 ) = 0.624
−0.2 0.0 0.2 0.4
t( μ1 = 8.84 , σ1 = 0.07 , ν = 25.86 )
t( μ2 = 8.82 , σ2 = 0.1 , ν = 25.86 )
A1160p+ (I−11a)
AFUB_079860 (I−11a)
P( −0.101 < μ1 − μ2 < 0.131 ) = 0.95
8.2 8.4 8.6 8.8 9.0 9.2 9.4

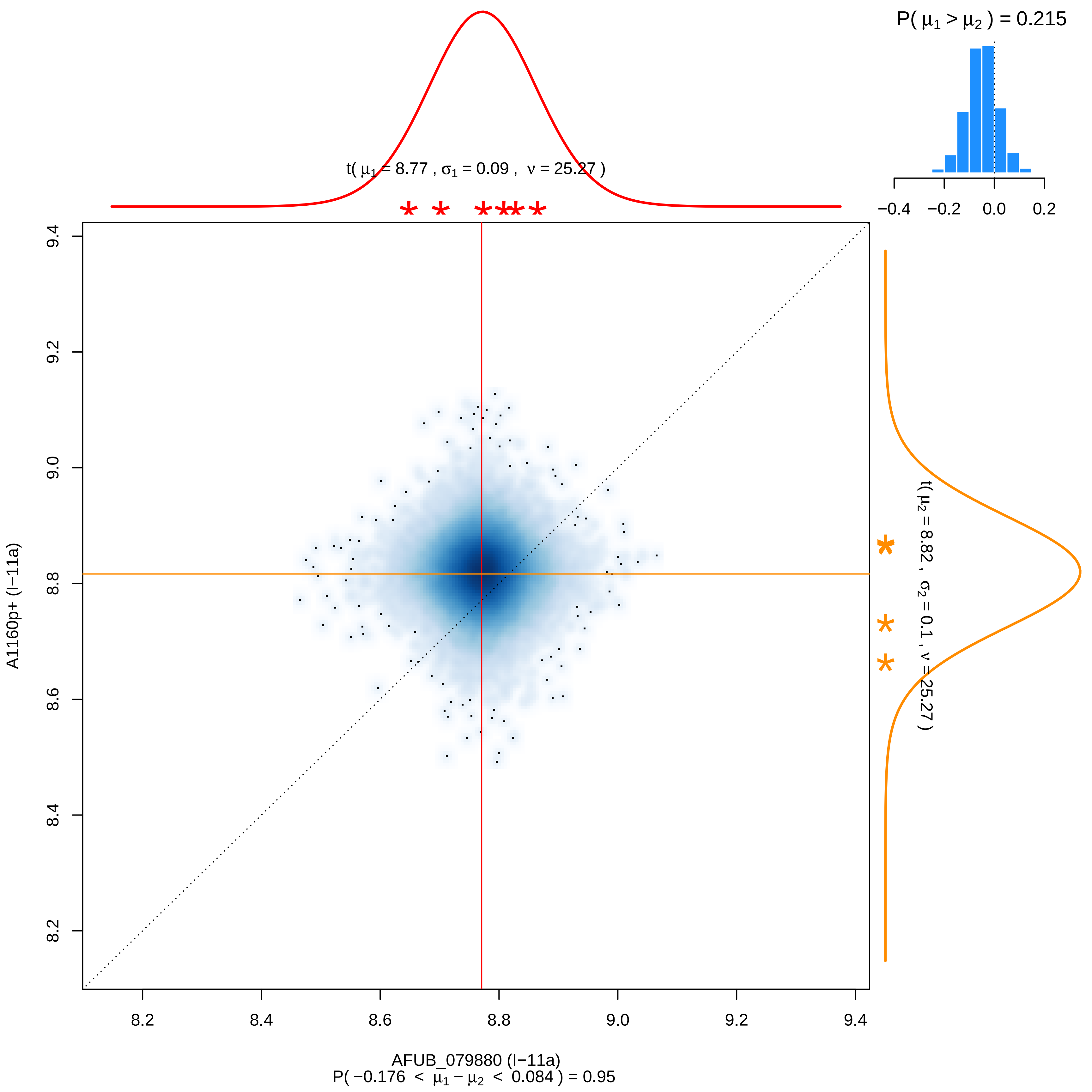

P( μ1 > μ2 ) = 0.215
t( μ1 = 8.77 , σ1 = 0.09 , ν = 25.27 )
t( μ2 = 8.82 , σ2 = 0.1 , ν = 25.27 )
−0.4 −0.2 0.0 0.2
A1160p+ (I−11a)
AFUB_079880 (I−11a)
P( −0.176 < μ1 − μ2 < 0.084 ) = 0.95
8.2 8.4 8.6 8.8 9.0 9.2 9.4

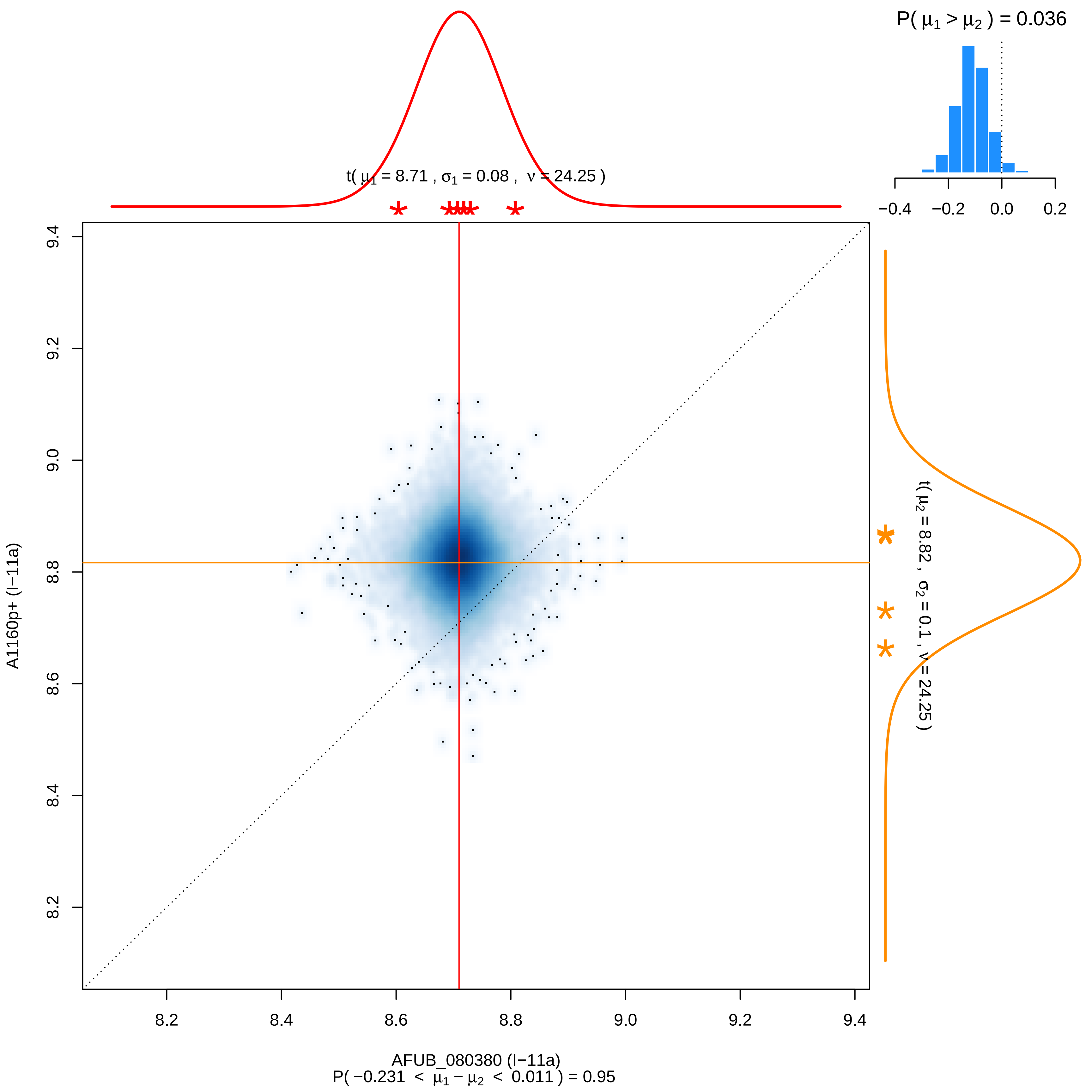

P( μ1 > μ2 ) = 0.036
−0.4
−0.2
0.0
0.2
t( μ1 = 8.71 , σ1 = 0.08 , ν = 24.25 )
t( μ2 = 8.82 , σ2 = 0.1 , ν = 24.25 )
A1160p+ (I−11a)
9.4
9.2
9.0
8.8
8.6
8.4
8.2
AFUB_080380 (I−11a)
P( −0.231 < μ1 − μ2 < 0.011 ) = 0.95

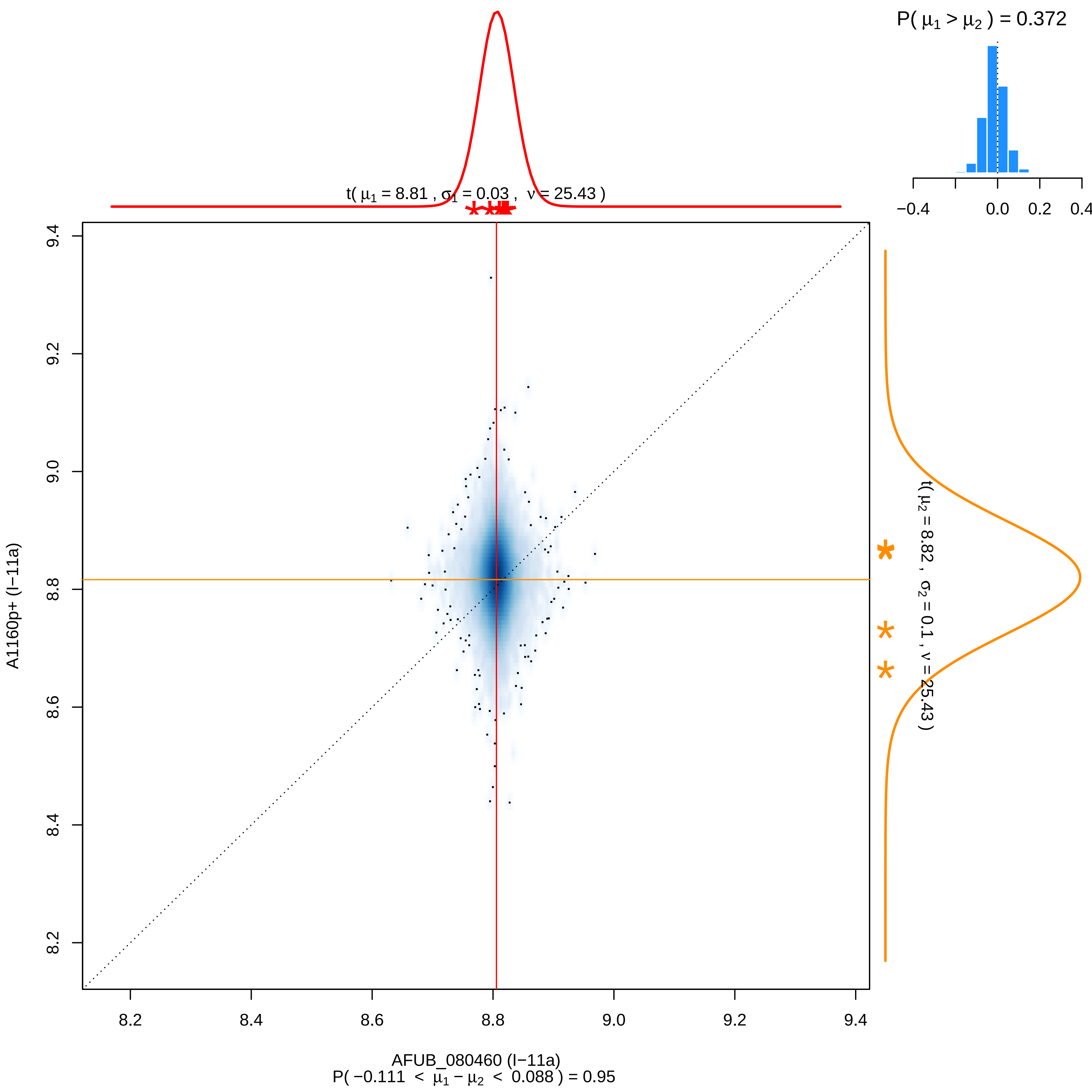

P( μ1 > μ2 ) = 0.372
−0.4
0.0
0.2
0.4
t( μ1 = 8.81 , σ1 = 0.03 , ν = 25.43 )
t( μ2 = 8.82 , σ2 = 0.1 , ν = 25.43 )
A1160p+ (I−11a)
9.4
9.2
9.0
8.8
8.6
8.4
8.2
8.2
8.4
8.6
8.8
9.0
9.2
9.4
AFUB_080460 (I−11a)
P( −0.111 < μ1 − μ2 < 0.088 ) = 0.95

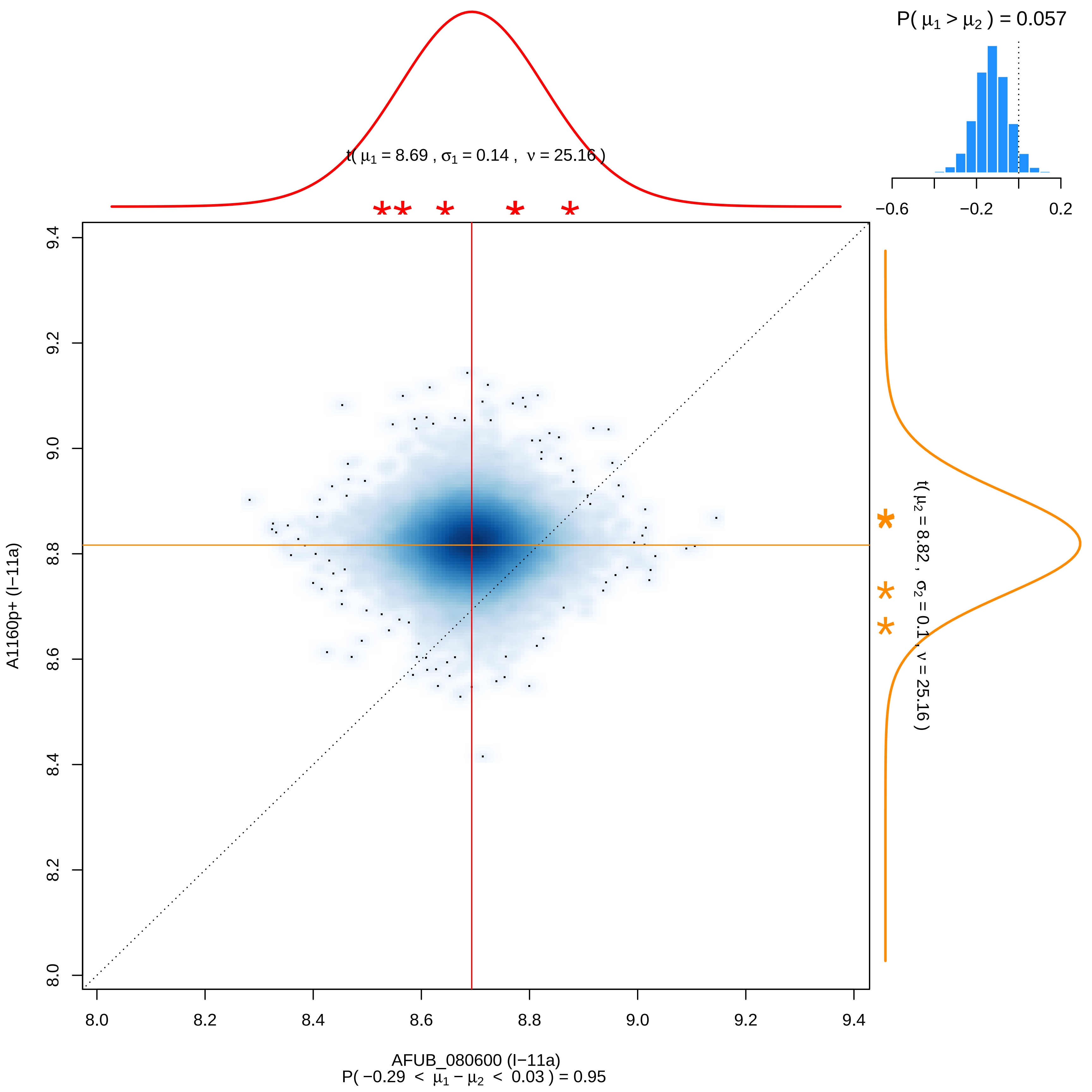

P( μ1 > μ2 ) = 0.057
−0.6
−0.2
0.2
t( μ1 = 8.69 , σ1 = 0.14 , ν = 25.16 )
t( μ2 = 8.82 , σ2 = 0.1 , ν = 25.16 )
9.4
9.2
9.0
8.8
8.6
8.4
8.2
8.0
A1160p+ (I−11a)
AFUB_080600 (I−11a)
P( −0.29 < μ1 − μ2 < 0.03 ) = 0.95

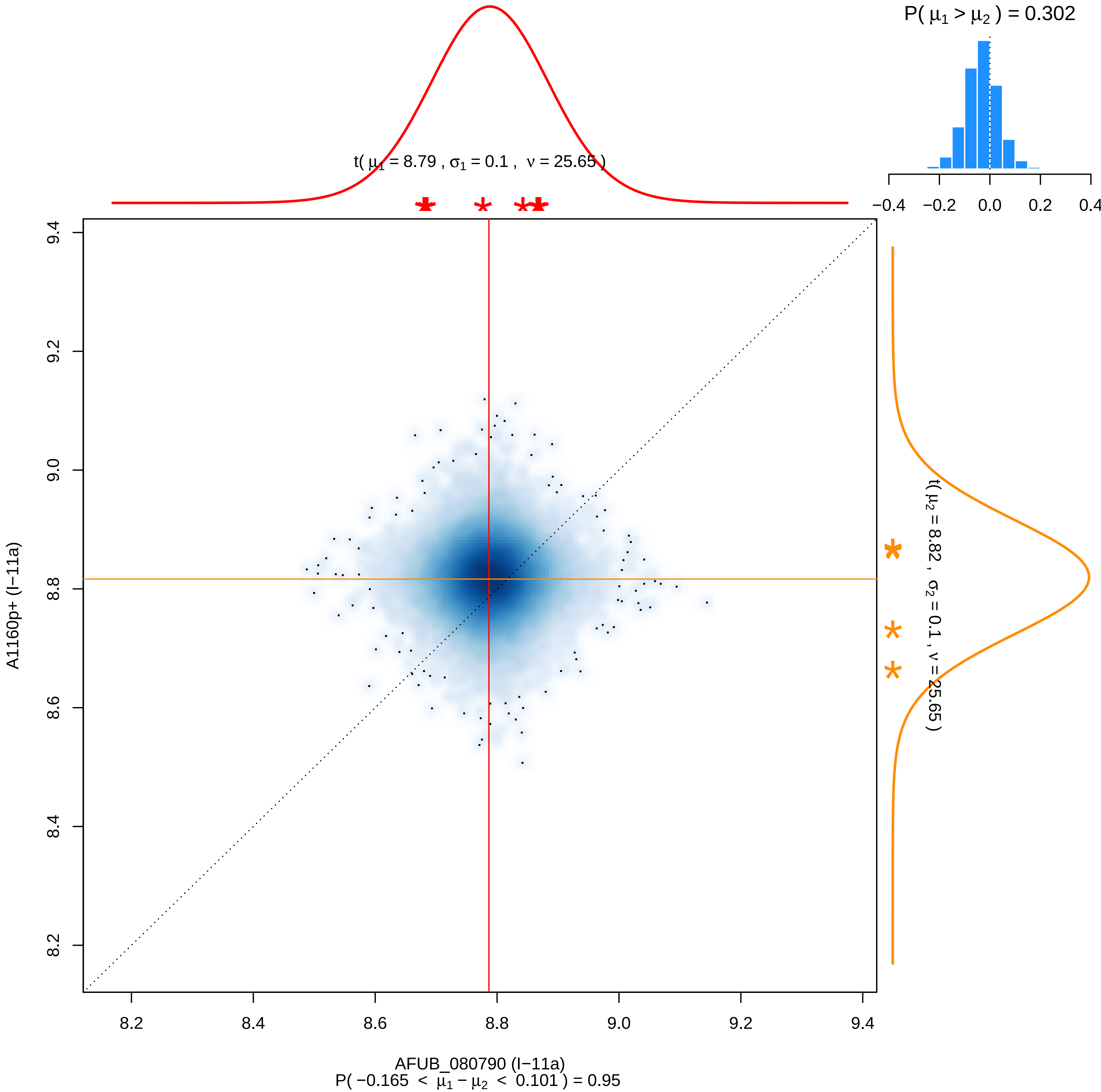

P( μ1 > μ2 ) = 0.302
−0.4
−0.2
0.0
0.2
0.4
t( μ1 = 8.79 , σ1 = 0.1 , ν = 25.65 )
t( μ2 = 8.82 , σ2 = 0.1 , ν = 25.65 )
A1160p+ (I−11a)
8.2
8.4
8.6
8.8
9.0
9.2
9.4
AFUB_080790 (I−11a)
P( −0.165 < μ1 − μ2 < 0.101 ) = 0.95

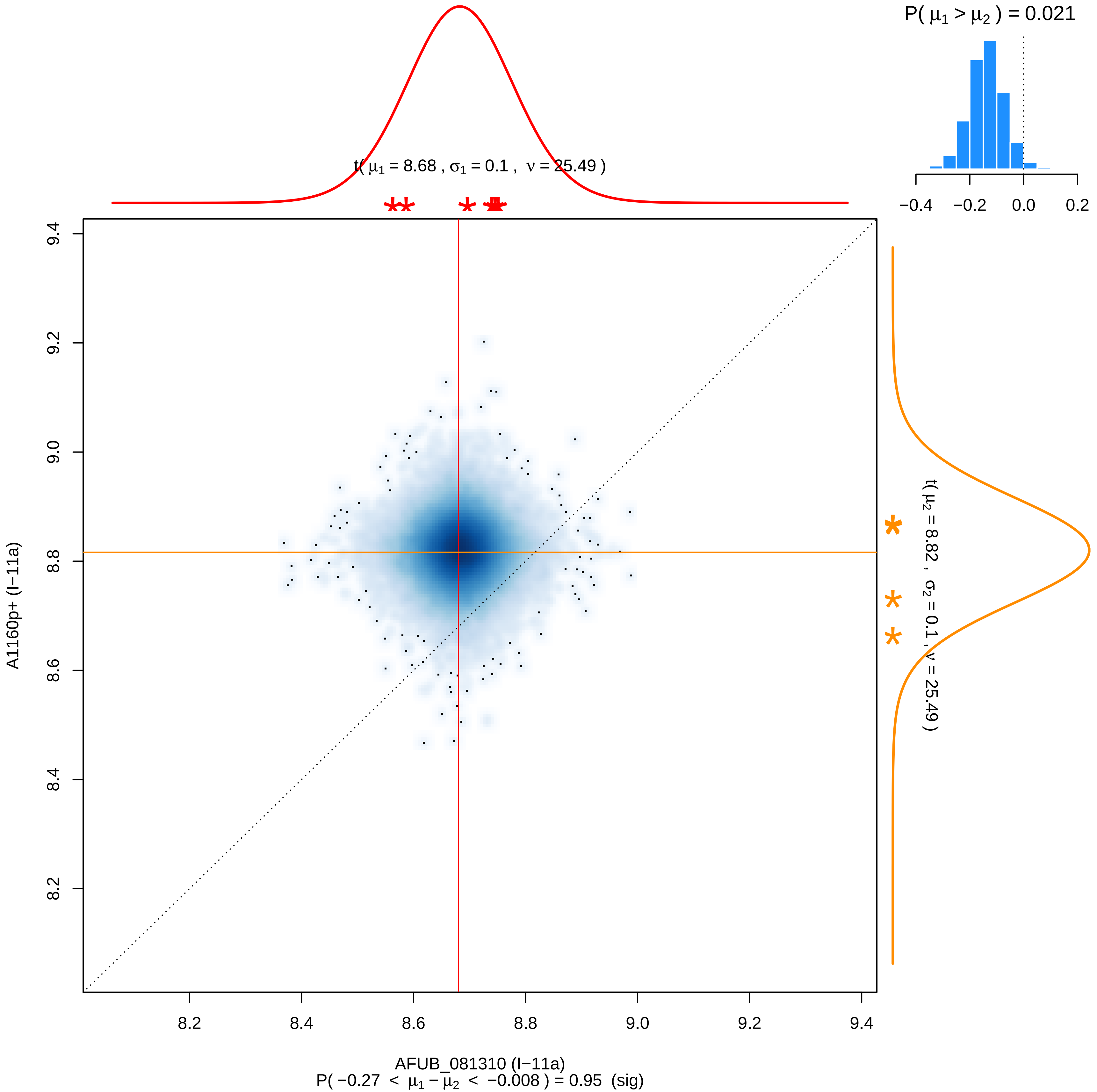

P( μ1 > μ2 ) = 0.021
−0.4 −0.2 0.0 0.2
t( μ1 = 8.68 , σ1 = 0.1 , ν = 25.49 )
t( μ2 = 8.82 , σ2 = 0.1 , ν = 25.49 )
9.4 9.2 9.0 8.8 8.6 8.4 8.2
A1160p+ (I−11a)
8.2 8.4 8.6 8.8 9.0 9.2 9.4
AFUB_081310 (I−11a)
P( −0.27 < μ1 − μ2 < −0.008 ) = 0.95 (sig)

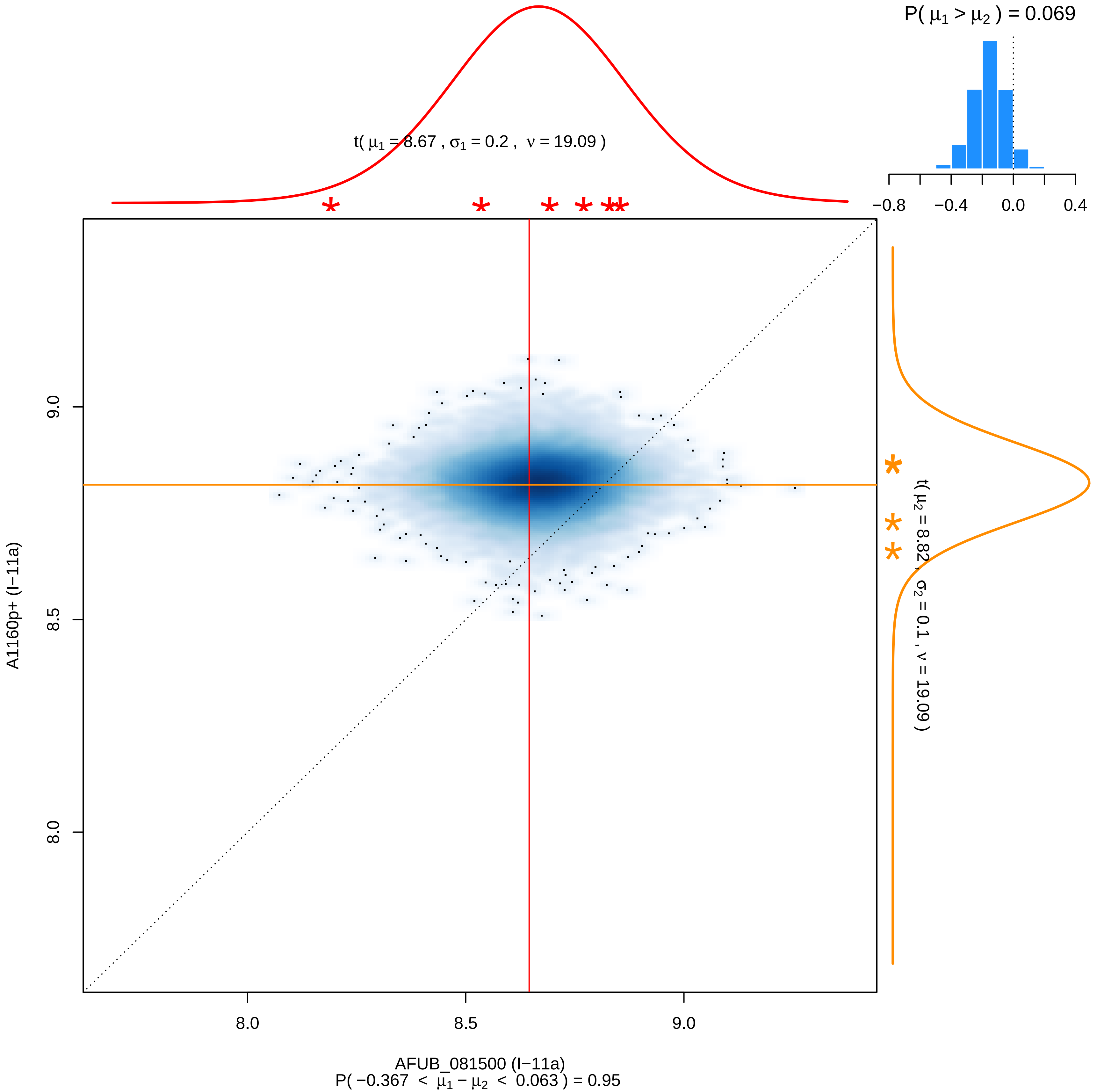

P( μ1 > μ2 ) = 0.069
−0.8
−0.4
0.0
0.4
t( μ1 = 8.67 , σ1 = 0.2 , ν = 19.09 )
t( μ2 = 8.82 , σ2 = 0.1 , ν = 19.09 )
9.0
8.5
8.0
A1160p+ (I−11a)
8.0
8.5
9.0
AFUB_081500 (I−11a)
P( −0.367 < μ1 − μ2 < 0.063 ) = 0.95

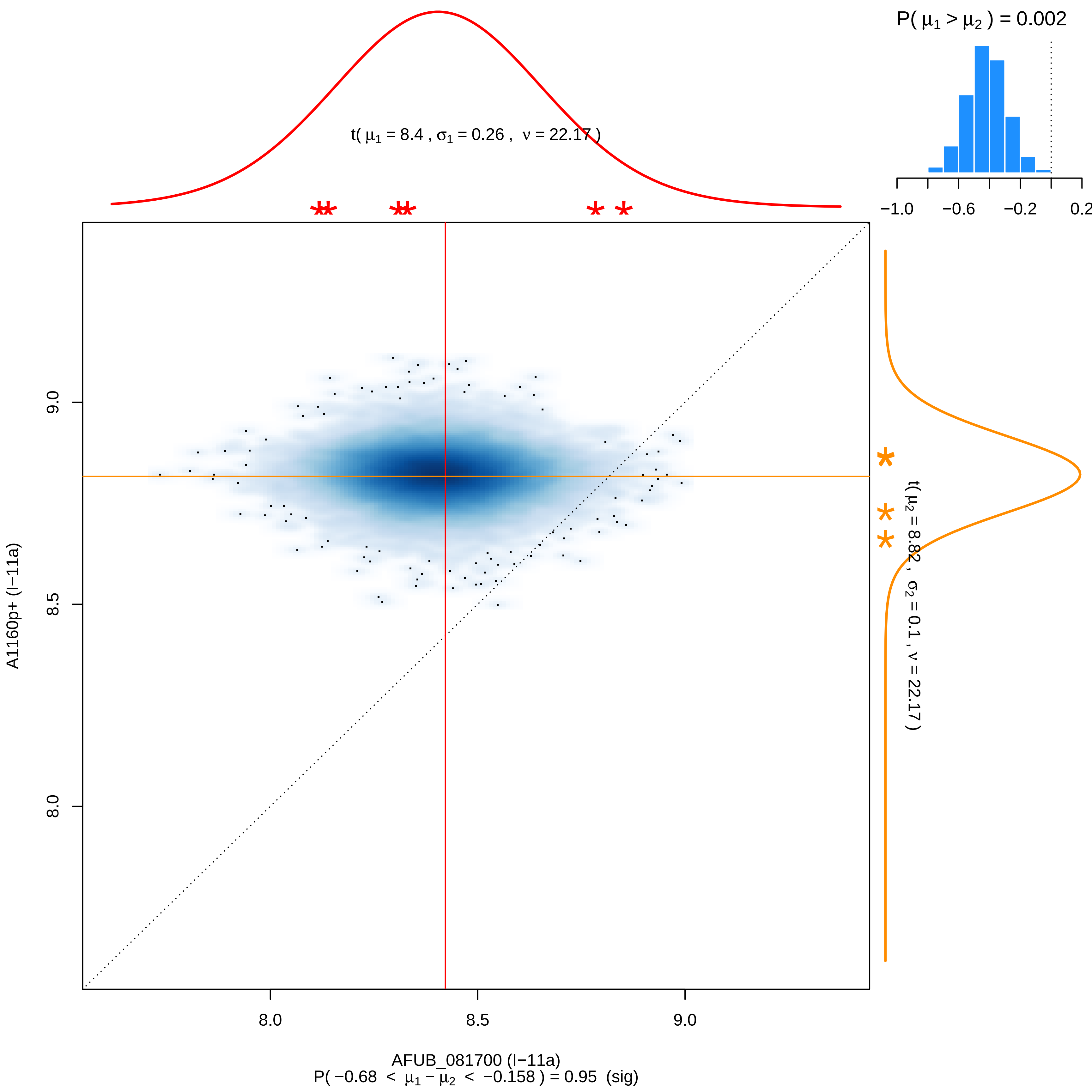

P( μ1 > μ2 ) = 0.002
−1.0
−0.6
−0.2
0.2
t( μ1 = 8.4 , σ1 = 0.26 , ν = 22.17 )
t( μ2 = 8.82 , σ2 = 0.1 , ν = 22.17 )
9.0
8.5
8.0
A1160p+ (I−11a)
AFUB_081700 (I−11a)
P( −0.68 < μ1 − μ2 < −0.158 ) = 0.95 (sig)

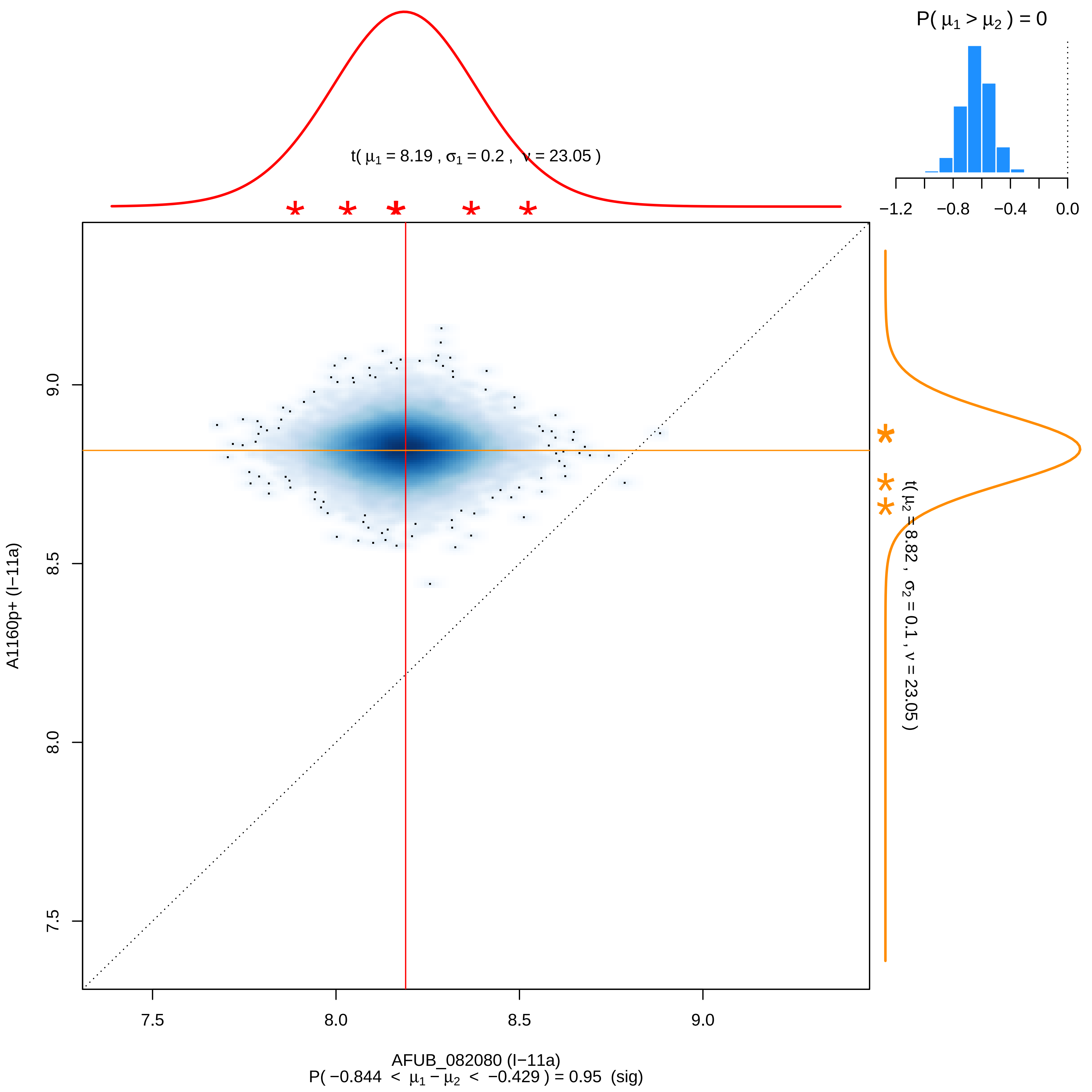

P( $\mu_1 > \mu_2$ ) = 0
−1.2
−0.8
−0.4
0.0
t( $\mu_1$ = 8.19 , $\sigma_1$ = 0.2 , $\nu$ = 23.05 )
t( $\mu_2$ = 8.82 , $\sigma_2$ = 0.1 , $\nu$ = 23.05 )
A1160p+ (I−11a)
9.0
8.5
8.0
7.5
7.5
8.0
8.5
9.0
AFUB_082080 (I−11a)
P( −0.844 < $\mu_1 - \mu_2$ < −0.429 ) = 0.95 (sig)

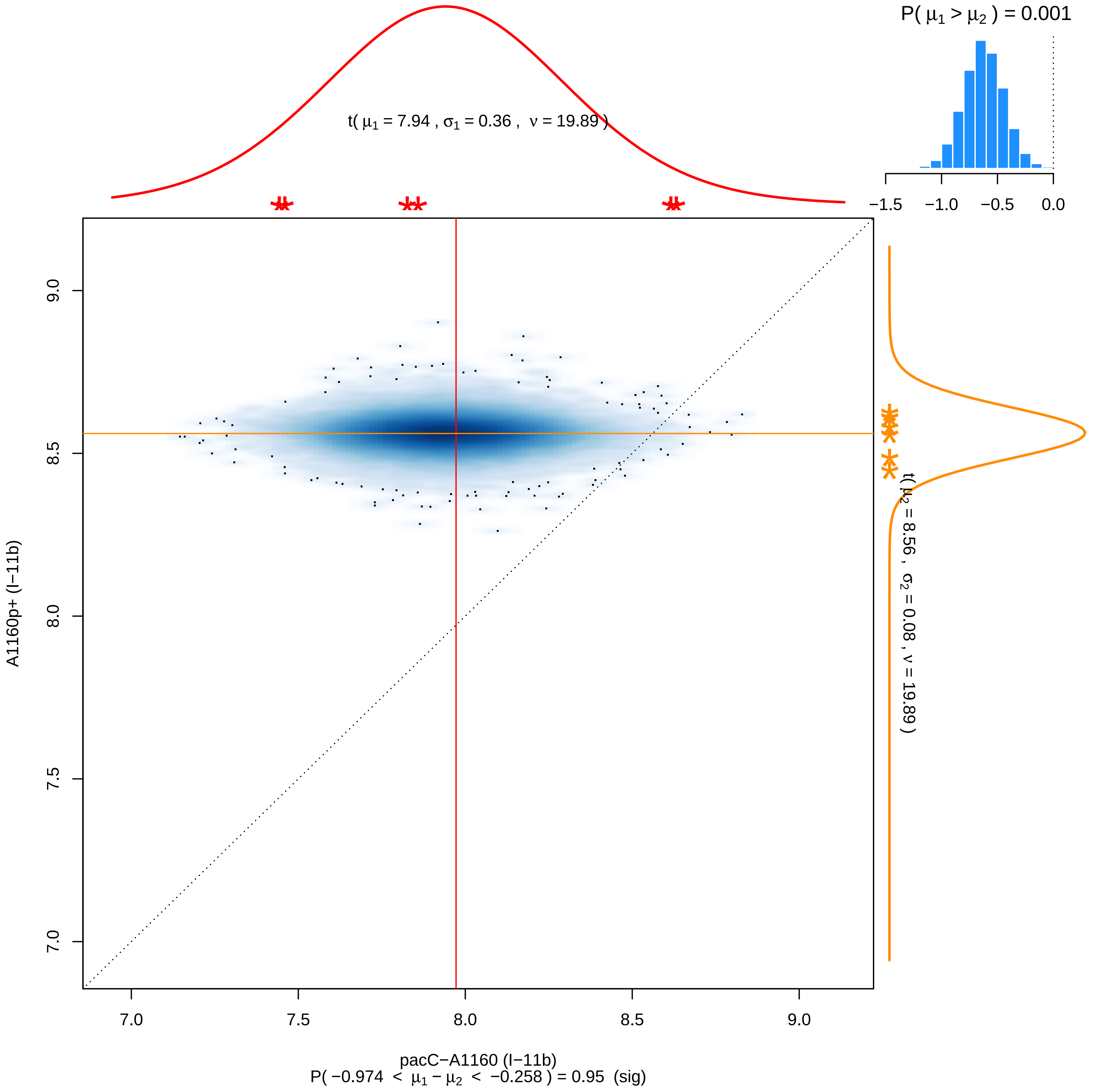

P( $\mu_1 > \mu_2$ ) = 0.001
−1.5
−1.0
−0.5
0.0
t( $\mu_1$ = 7.94 , $\sigma_1$ = 0.36 , $\nu$ = 19.89 )
t( $\mu_2$ = 8.56 , $\sigma_2$ = 0.08 , $\nu$ = 19.89 )
9.0
8.5
8.0
7.5
7.0
A1160p+ (I−11b)
7.0
7.5
8.0
8.5
9.0
pacC−A1160 (I−11b)
P( −0.974 < $\mu_1 - \mu_2$ < −0.258 ) = 0.95 (sig)

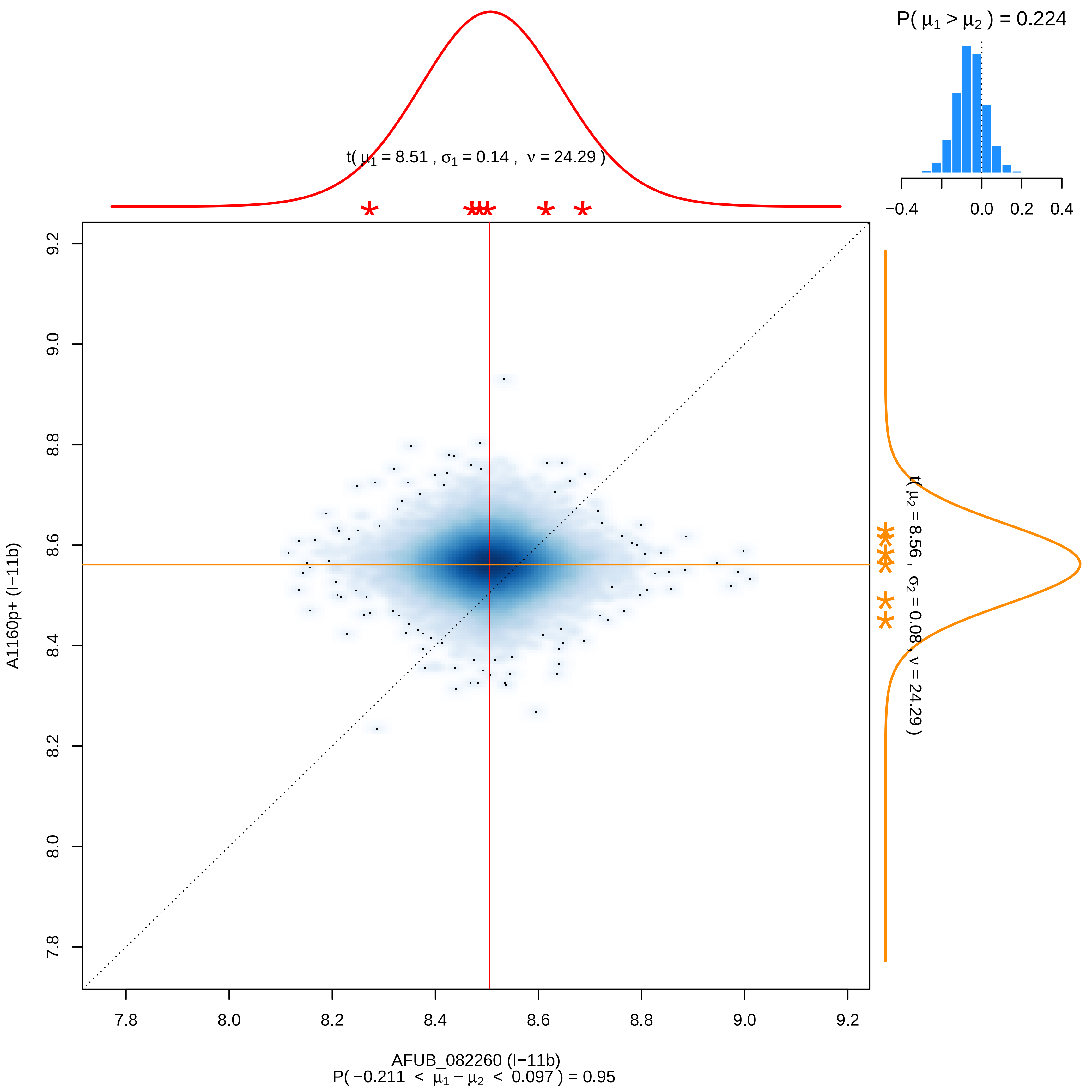

P( μ1 > μ2 ) = 0.224
−0.4
0.0
0.2
0.4
t( μ1 = 8.51 , σ1 = 0.14 , ν = 24.29 )
t( μ2 = 8.56 , σ2 = 0.08 , ν = 24.29 )
A1160p+ (I−11b)
7.8
8.0
8.2
8.4
8.6
8.8
9.0
9.2
AFUB_082260 (I−11b)
P( −0.211 < μ1 − μ2 < 0.097 ) = 0.95

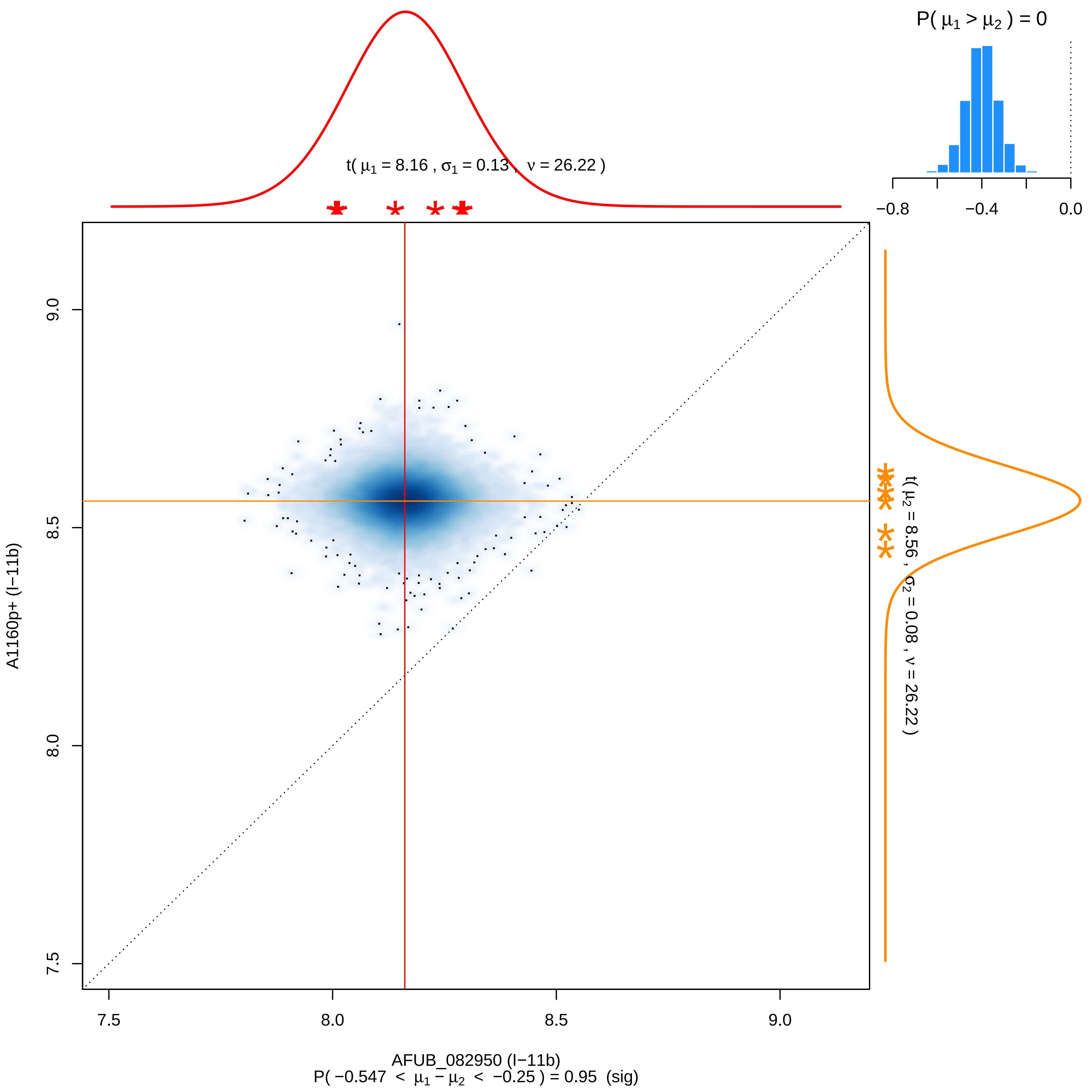
P( μ1 > μ2 ) = 0
−0.8
−0.4
0.0
t( μ1 = 8.16 , σ1 = 0.13 , ν = 26.22 )
t( μ2 = 8.56 , σ2 = 0.08 , ν = 26.22 )
9.0
8.5
8.0
7.5
A1160p+ (I−11b)
7.5
8.0
8.5
9.0
AFUB_082950 (I−11b)
P( −0.547 < μ1 − μ2 < −0.25 ) = 0.95 (sig)

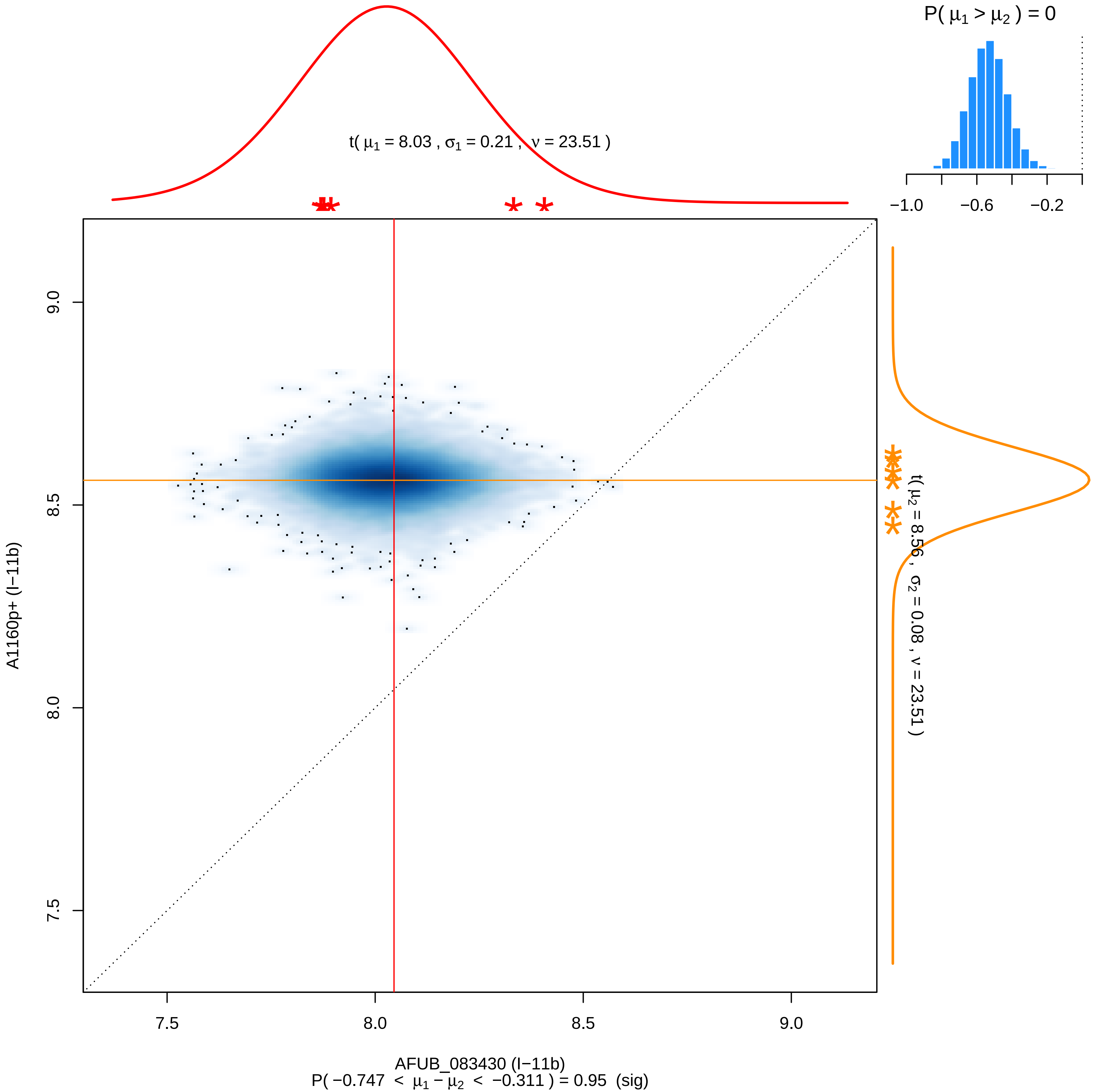

P( μ1 > μ2 ) = 0
−1.0
−0.6
−0.2
t( μ1 = 8.03 , σ1 = 0.21 , ν = 23.51 )
t( μ2 = 8.56 , σ2 = 0.08 , ν = 23.51 )
9.0
8.5
8.0
7.5
A1160p+ (I−11b)
7.5
8.0
8.5
9.0
AFUB_083430 (I−11b)
P( −0.747 < μ1 − μ2 < −0.311 ) = 0.95 (sig)

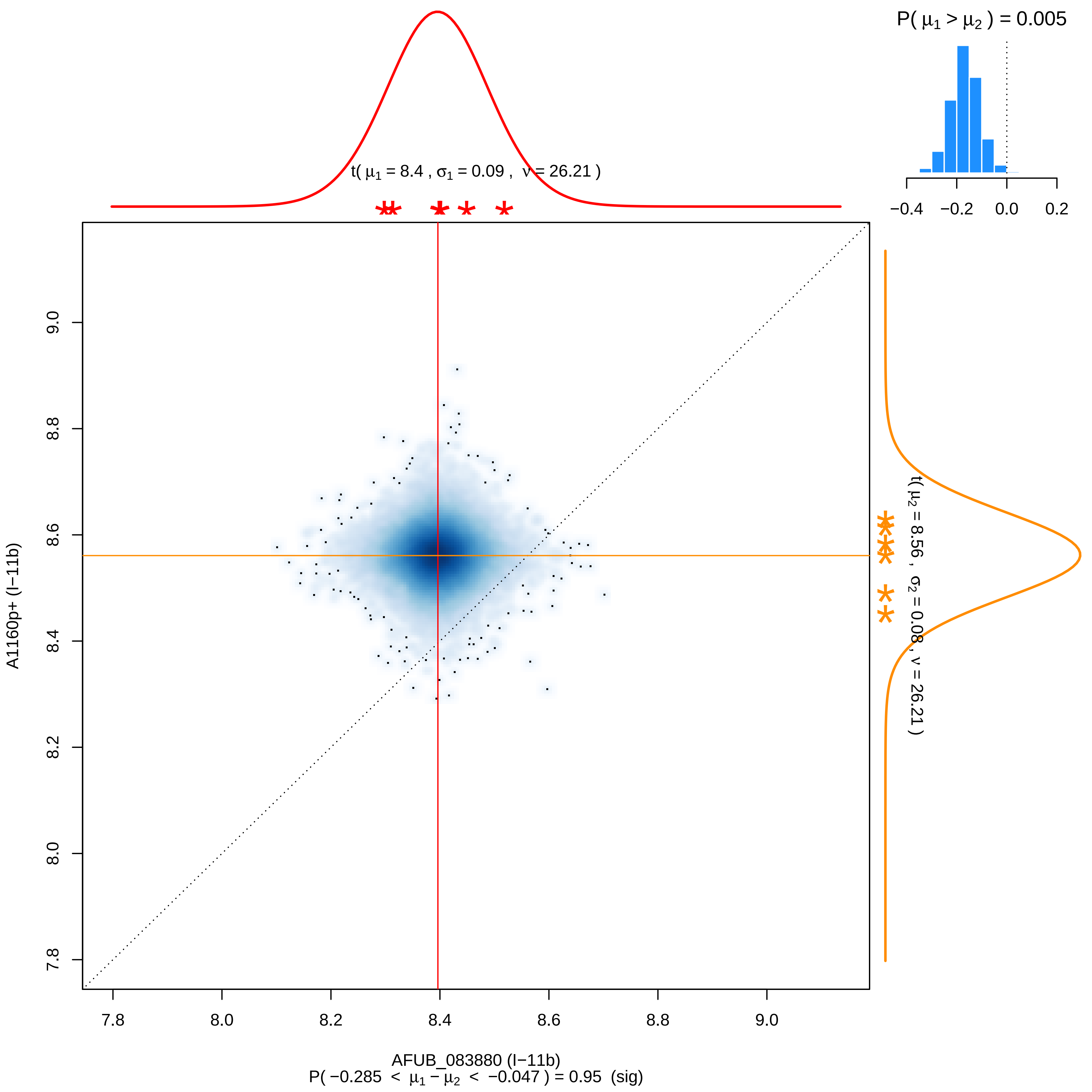

P( μ1 > μ2 ) = 0.005
−0.4
−0.2
0.0
0.2
t( μ1 = 8.4 , σ1 = 0.09 , ν = 26.21 )
t( μ2 = 8.56 , σ2 = 0.08 , ν = 26.21 )
A1160p+ (I−11b)
9.0
8.8
8.6
8.4
8.2
8.0
7.8
AFUB_083880 (I−11b)
P( −0.285 < μ1 − μ2 < −0.047 ) = 0.95 (sig)

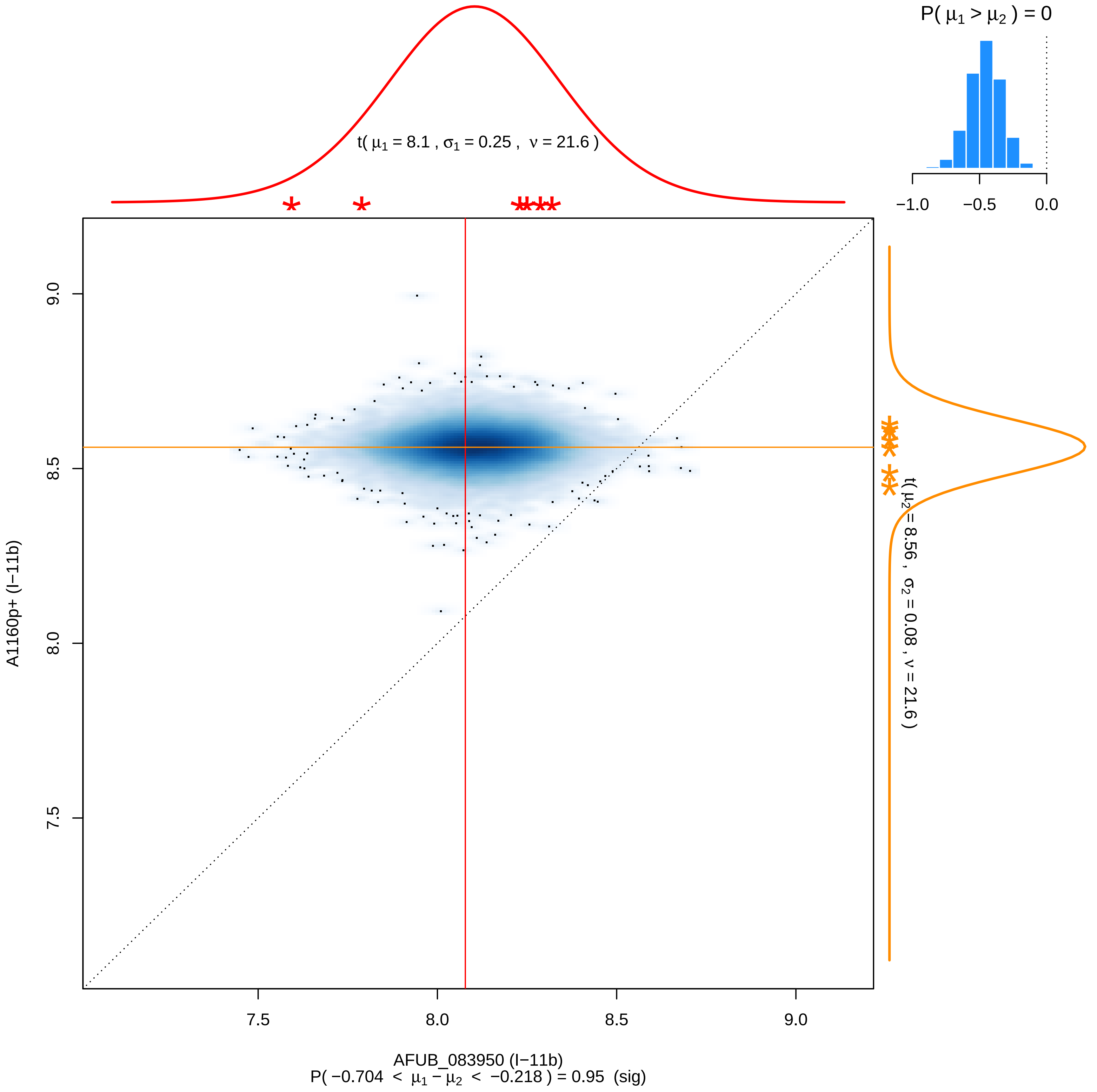
P( μ1 > μ2 ) = 0
−1.0
−0.5
0.0
t( μ1 = 8.1 , σ1 = 0.25 , ν = 21.6 )
9.0
8.5
8.0
7.5
A1160p+ (I−11b)
t( μ2 = 8.56 , σ2 = 0.08 , ν = 21.6 )
7.5
8.0
8.5
9.0
AFUB_083950 (I−11b)
P( −0.704 < μ1 − μ2 < −0.218 ) = 0.95 (sig)

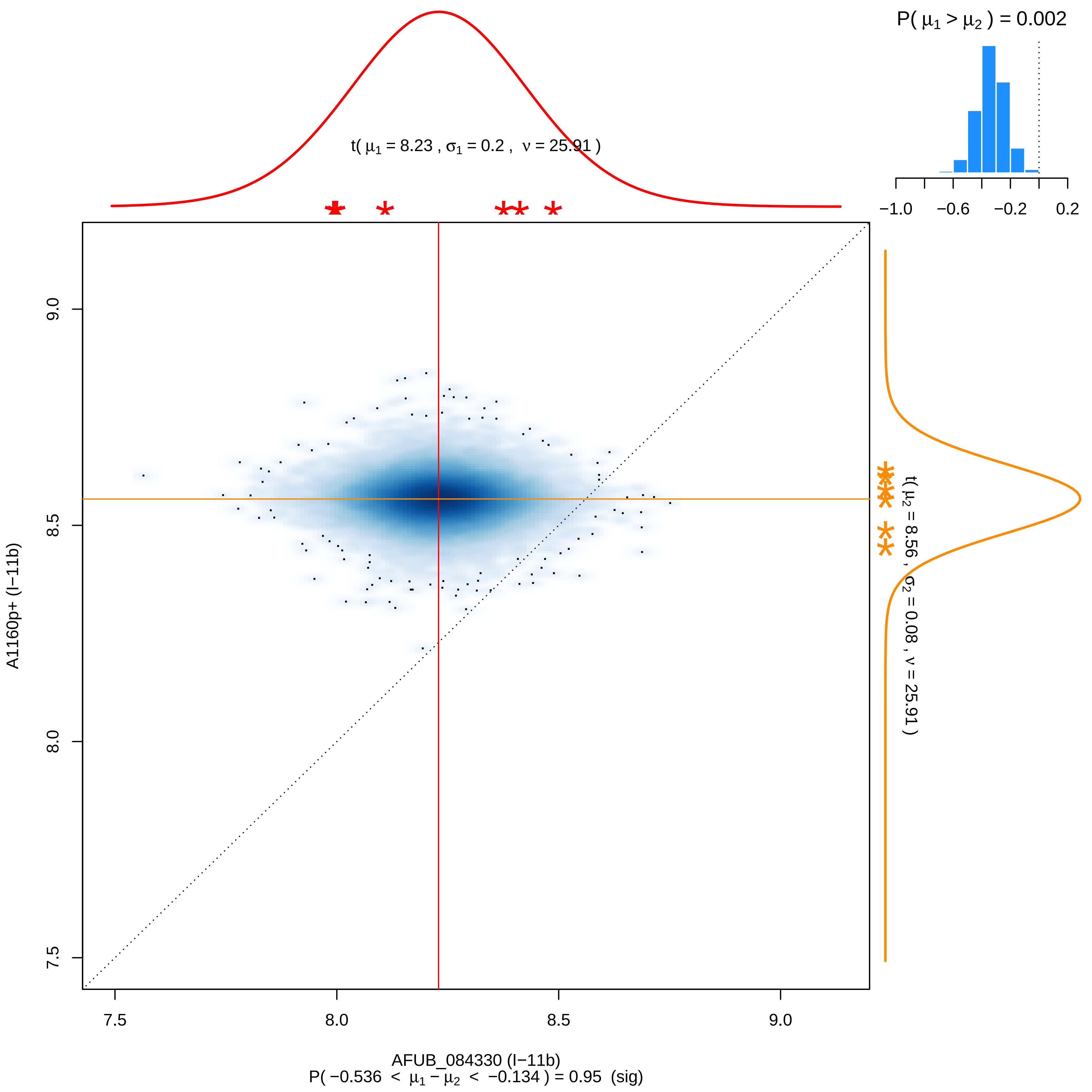

$P(\mu_1 > \mu_2) = 0.002$
−1.0 −0.6 −0.2 0.2
$t(\mu_1 = 8.23, \sigma_1 = 0.2, \nu = 25.91)$
$t(\mu_2 = 8.56, \sigma_2 = 0.08, \nu = 25.91)$
9.0
8.5
8.0
7.5
A1160p+ (I−11b)
7.5 8.0 8.5 9.0
AFUB_084330 (I−11b)
$P(-0.536 < \mu_1 - \mu_2 < -0.134) = 0.95$ (sig)

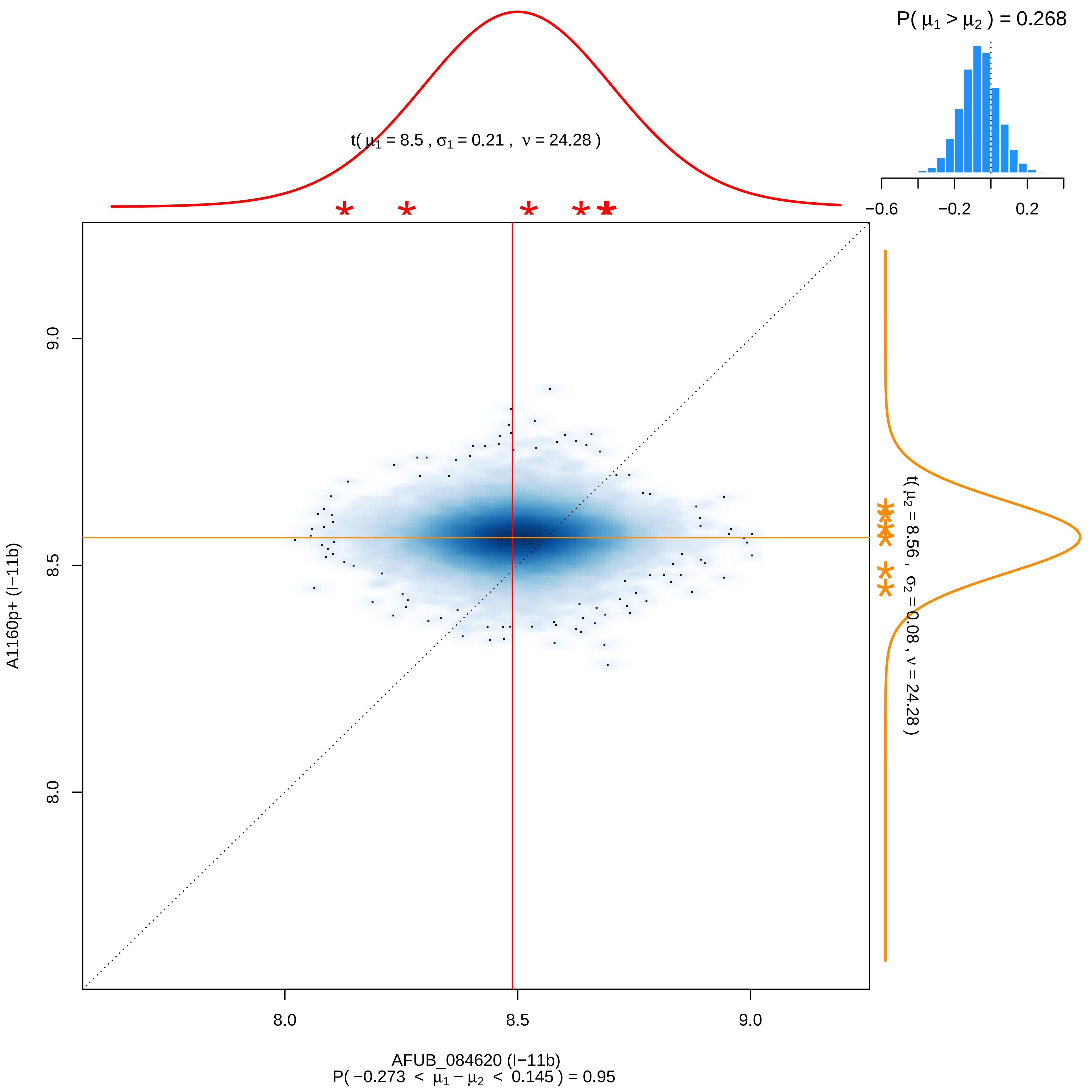
P( μ1 > μ2 ) = 0.268
−0.6
−0.2
0.2
t( μ1 = 8.5 , σ1 = 0.21 , ν = 24.28 )
t( μ2 = 8.56 , σ2 = 0.08 , ν = 24.28 )
9.0
8.5
8.0
A1160p+ (I−11b)
AFUB_084620 (I−11b)
P( −0.273 < μ1 − μ2 < 0.145 ) = 0.95

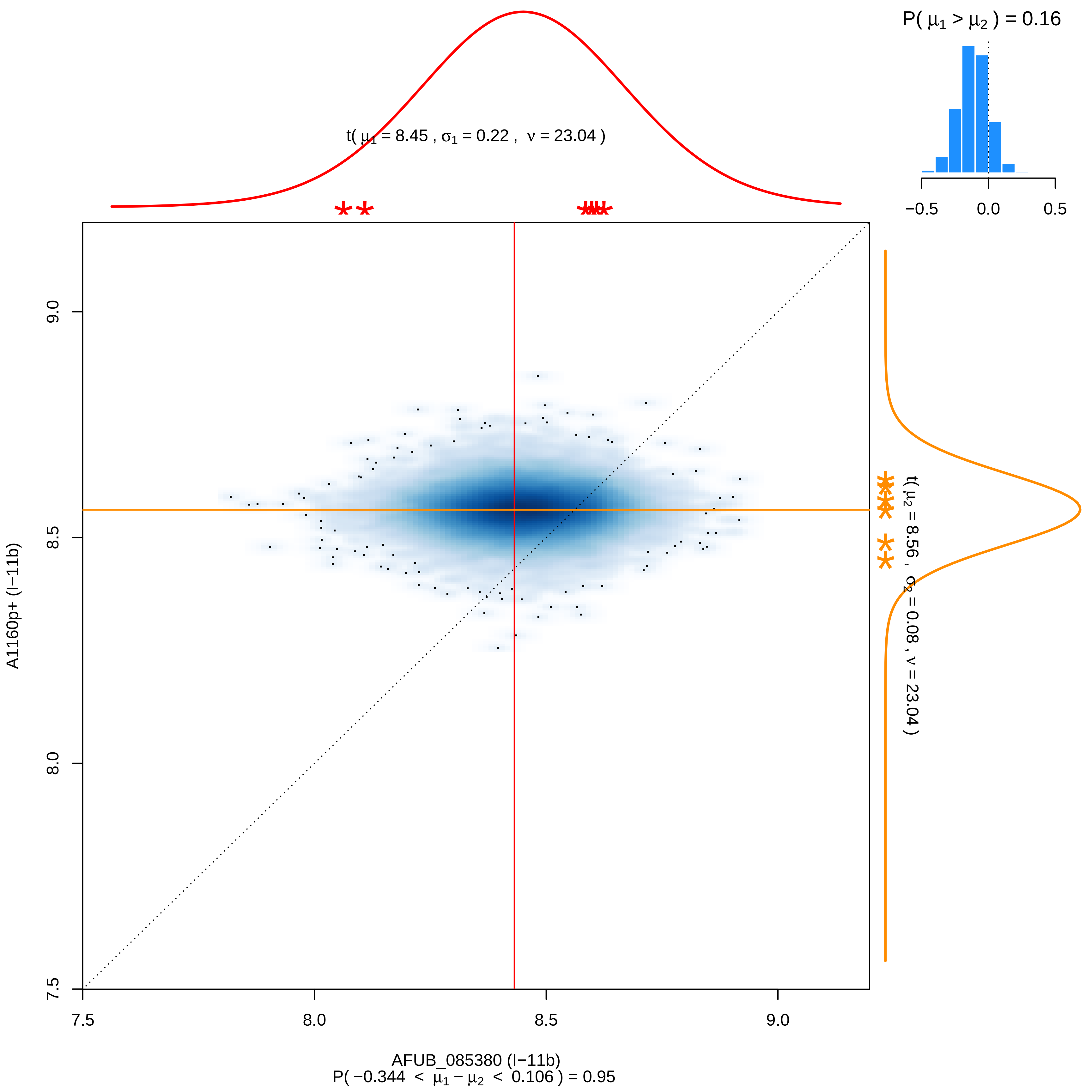

P( μ1 > μ2 ) = 0.16
−0.5
0.0
0.5
t( μ1 = 8.45 , σ1 = 0.22 , ν = 23.04 )
t( μ2 = 8.56 , σ2 = 0.08 , ν = 23.04 )
9.0
8.5
8.0
7.5
A1160p+ (I−11b)
7.5
8.0
8.5
9.0
AFUB_085380 (I−11b)
P( −0.344 < μ1 − μ2 < 0.106 ) = 0.95

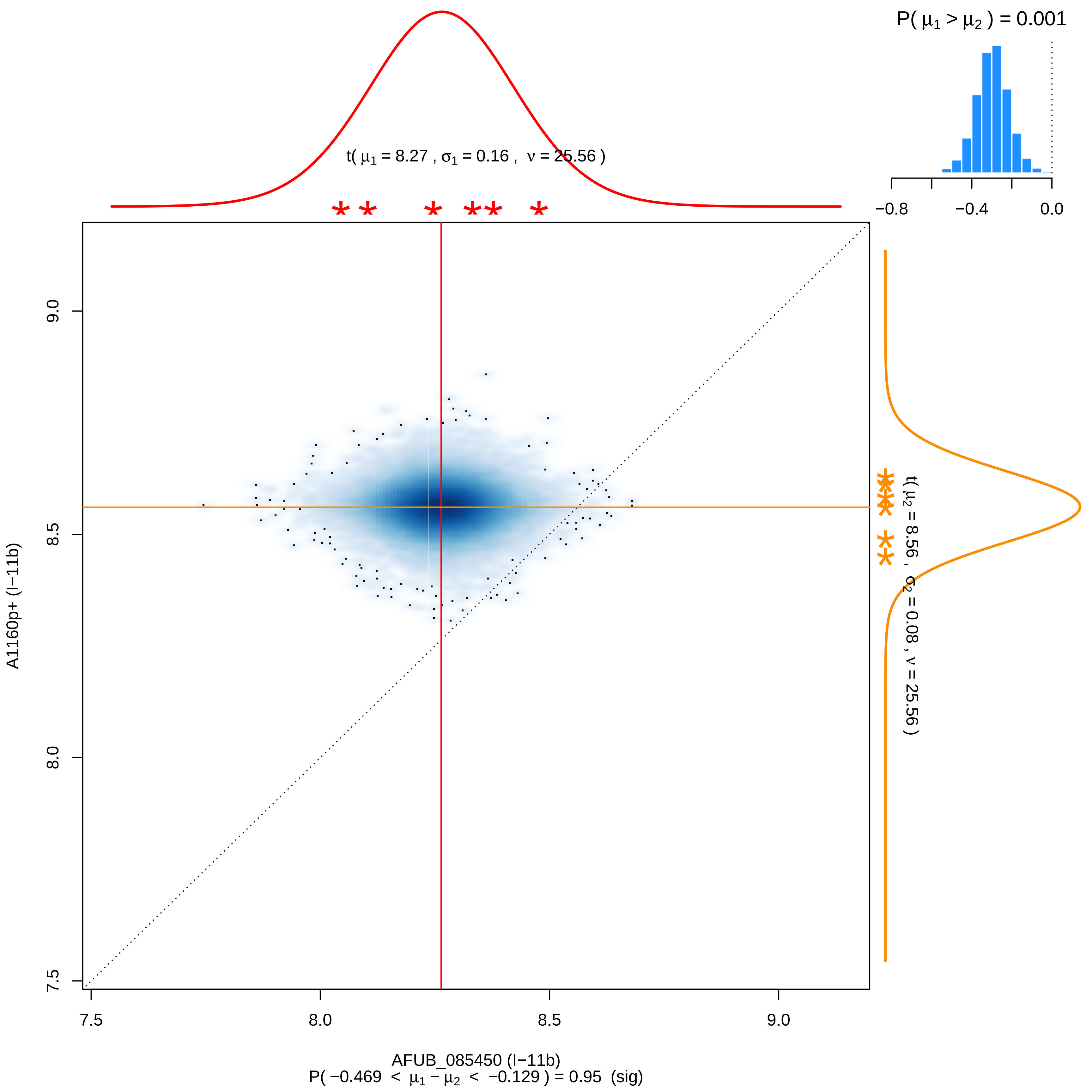

P( μ1 > μ2 ) = 0.001
−0.8
−0.4
0.0
t( μ1 = 8.27 , σ1 = 0.16 , ν = 25.56 )
t( μ2 = 8.56 , σ2 = 0.08 , ν = 25.56 )
A1160p+ (I−11b)
7.5
8.0
8.5
9.0
AFUB_085450 (I−11b)
P( −0.469 < μ1 − μ2 < −0.129 ) = 0.95 (sig)

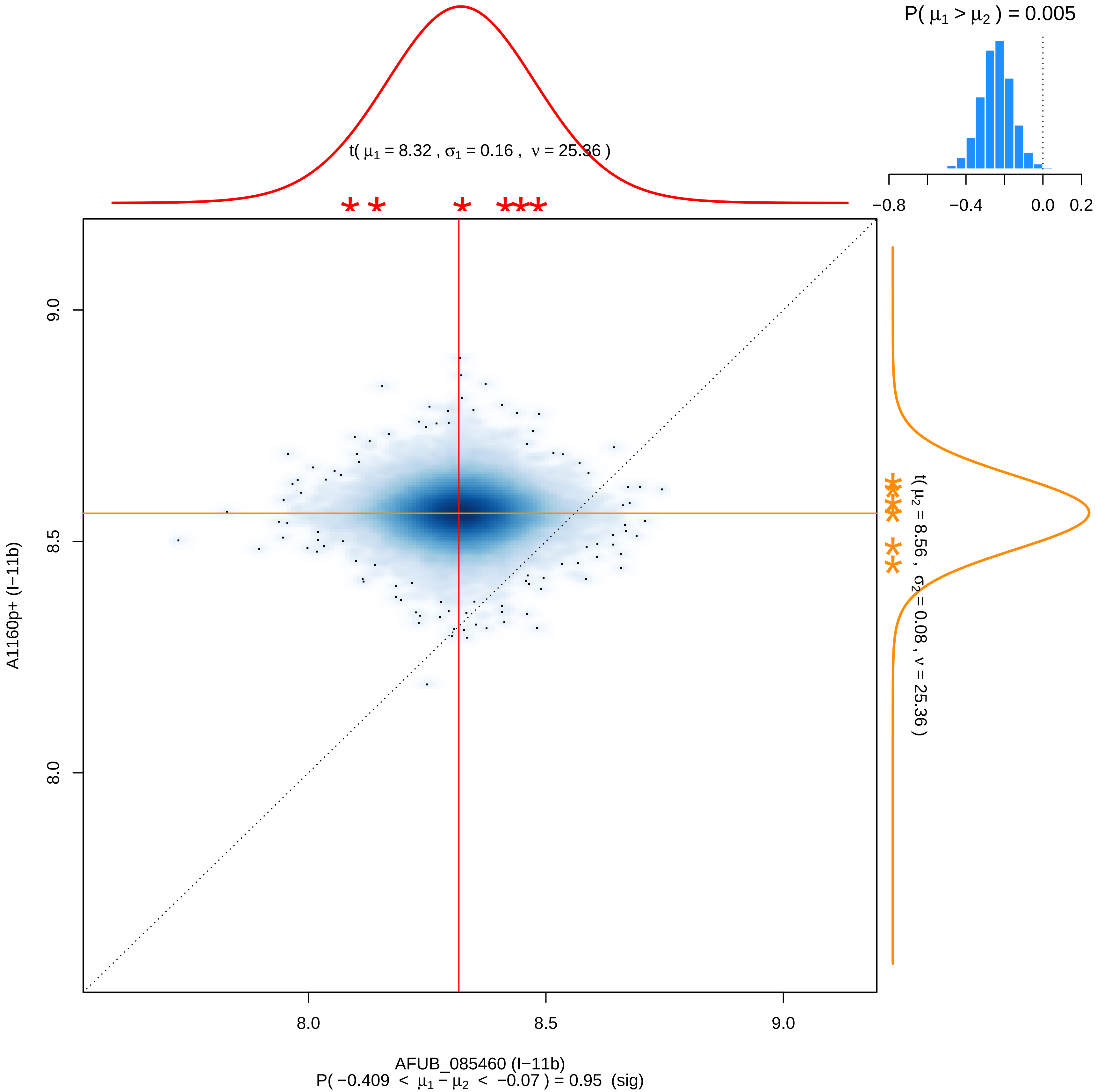

P( μ1 > μ2 ) = 0.005
−0.8 −0.4 0.0 0.2
t( μ1 = 8.32 , σ1 = 0.16 , ν = 25.36 )
t( μ2 = 8.56 , σ2 = 0.08 , ν = 25.36 )
9.0
8.5
8.0
A1160p+ (I−11b)
8.0 8.5 9.0
AFUB_085460 (I−11b)
P( −0.409 < μ1 − μ2 < −0.07 ) = 0.95 (sig)

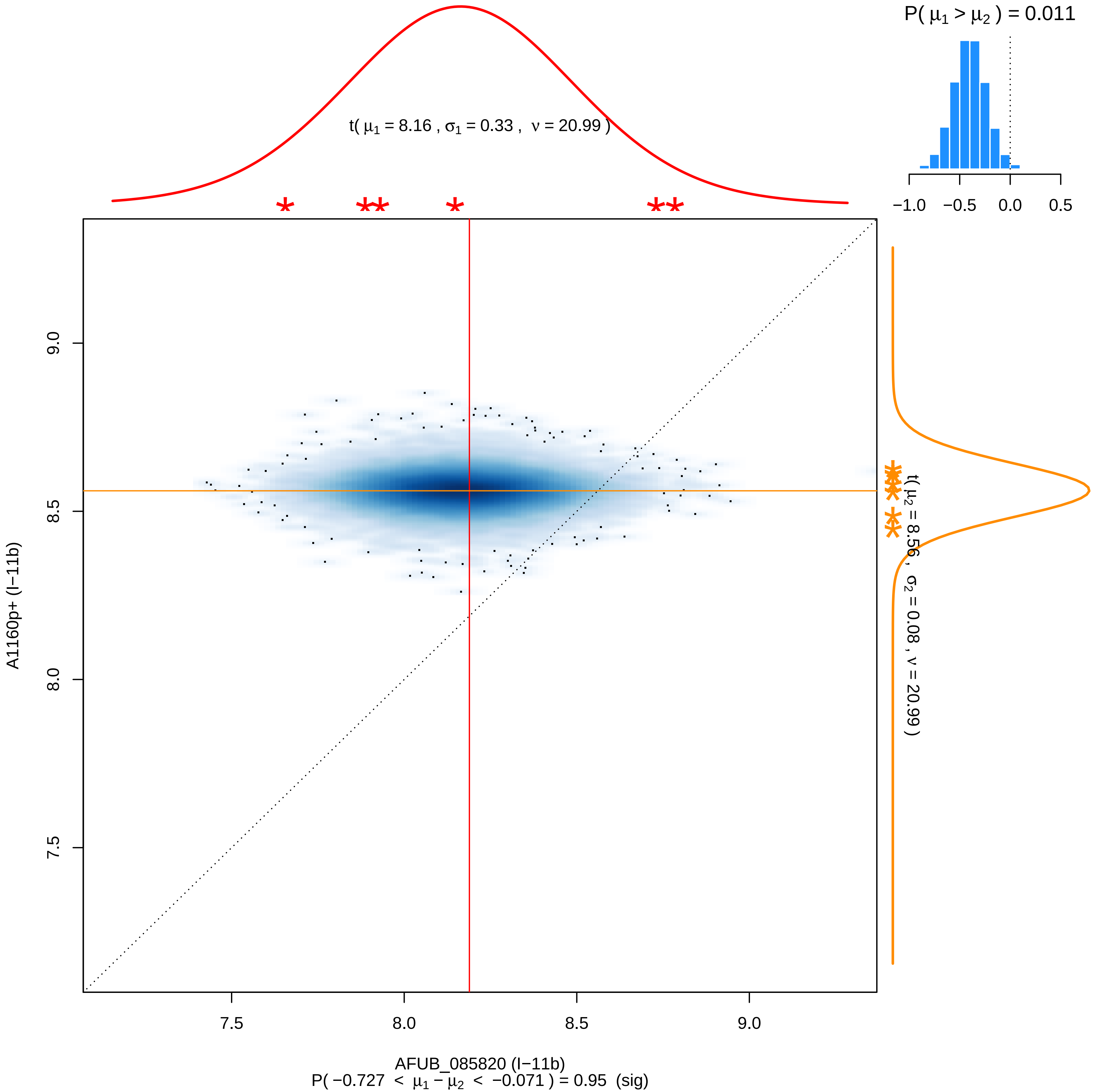
P( $\mu_1 > \mu_2$ ) = 0.011
−1.0 −0.5 0.0 0.5
t( $\mu_1$ = 8.16 , $\sigma_1$ = 0.33 , $\nu$ = 20.99 )
t( $\mu_2$ = 8.56 , $\sigma_2$ = 0.08 , $\nu$ = 20.99 )
A1160p+ (I−11b)
7.5 8.0 8.5 9.0
AFUB_085820 (I−11b)
P( −0.727 < $\mu_1 - \mu_2$ < −0.071 ) = 0.95 (sig)

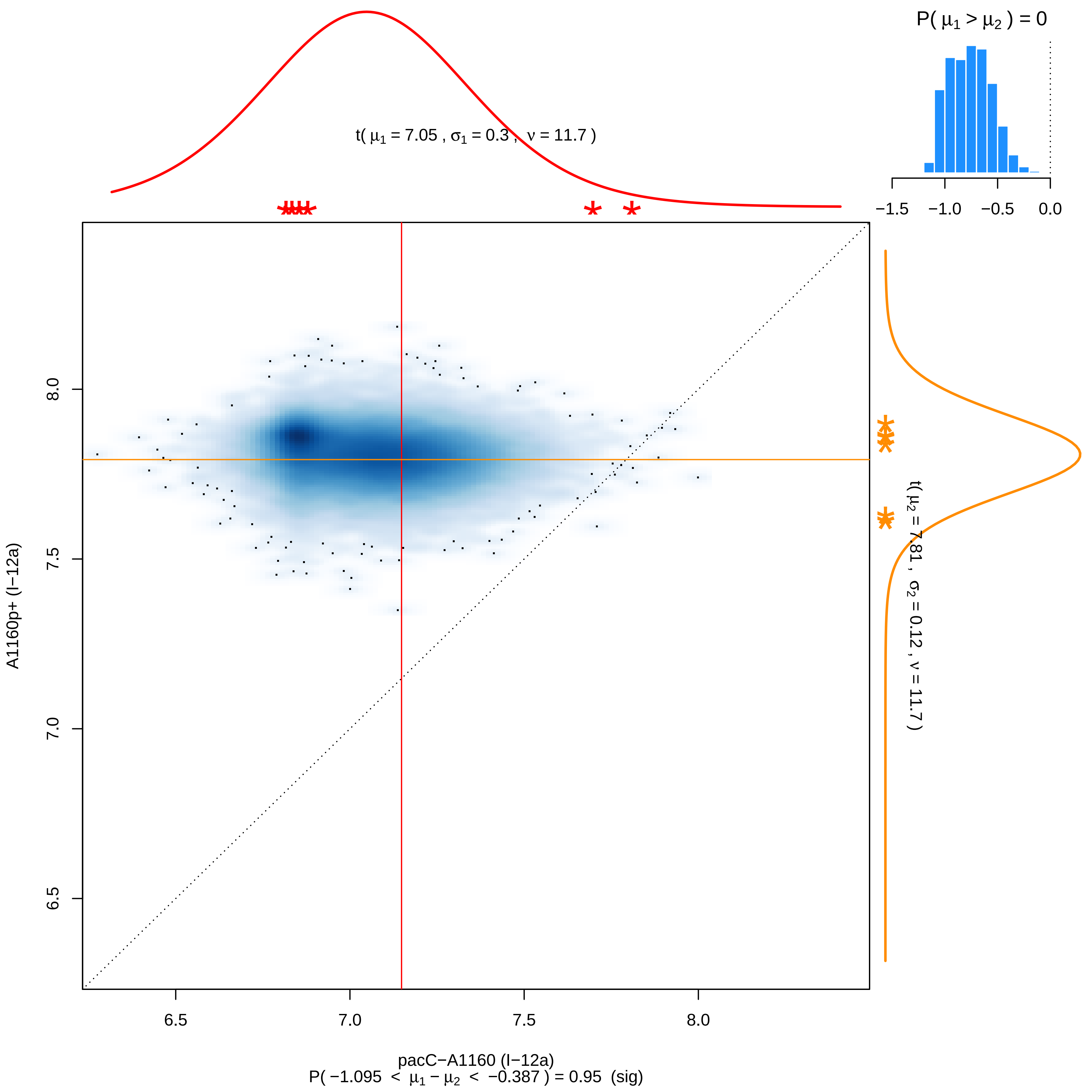

P( μ1 > μ2 ) = 0
−1.5 −1.0 −0.5 0.0
t( μ1 = 7.05 , σ1 = 0.3 , ν = 11.7 )
8.0
7.5
7.0
6.5
A1160p+ (I−12a)
t( μ2 = 7.81 , σ2 = 0.12 , ν = 11.7 )
6.5 7.0 7.5 8.0
pacC−A1160 (I−12a)
P( −1.095 < μ1 − μ2 < −0.387 ) = 0.95 (sig)

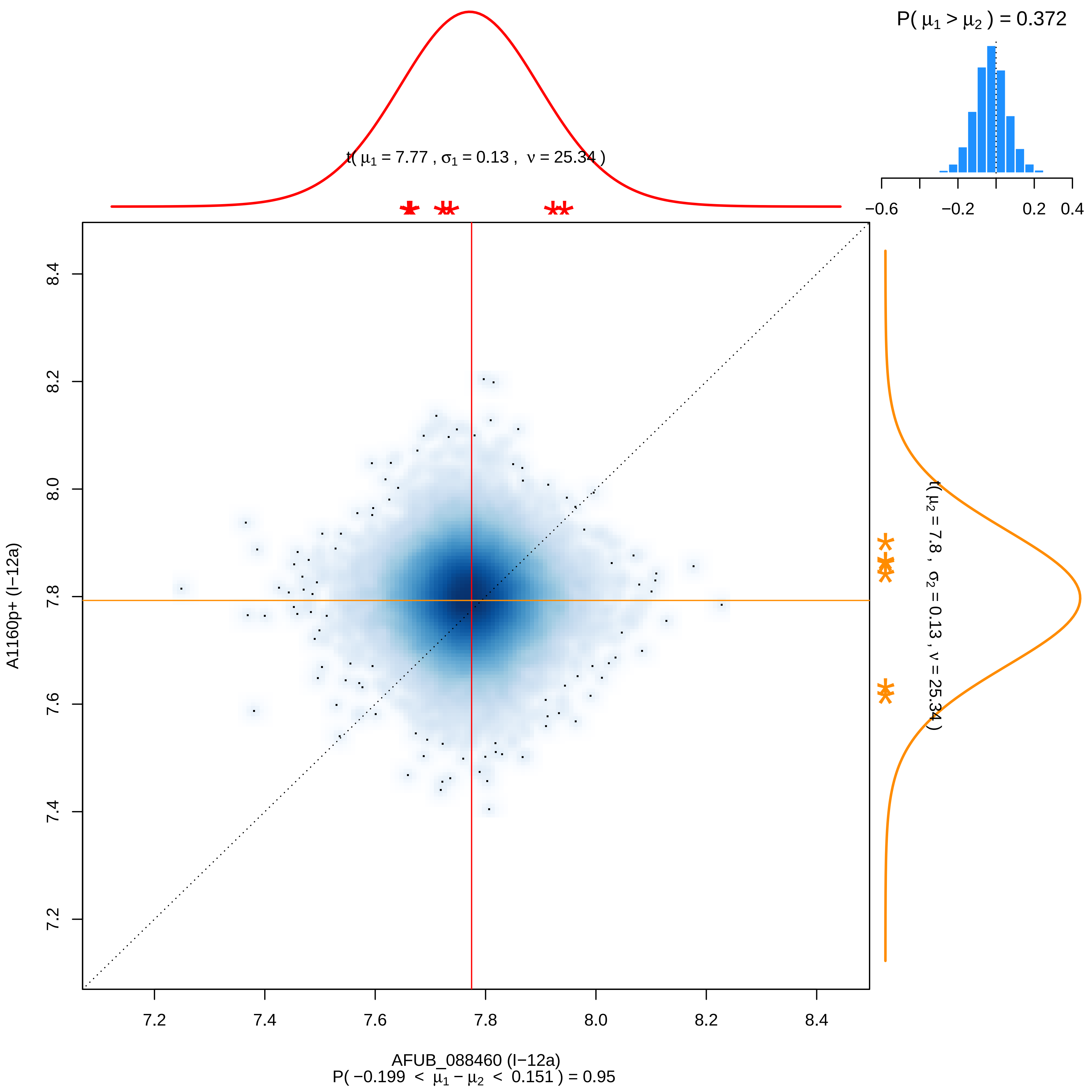

P( μ1 > μ2 ) = 0.372
−0.6
−0.2
0.2
0.4
t( μ1 = 7.77 , σ1 = 0.13 , ν = 25.34 )
t( μ2 = 7.8 , σ2 = 0.13 , ν = 25.34 )
A1160p+ (I−12a)
8.4
8.2
8.0
7.8
7.6
7.4
7.2
AFUB_088460 (I−12a)
P( −0.199 < μ1 − μ2 < 0.151 ) = 0.95

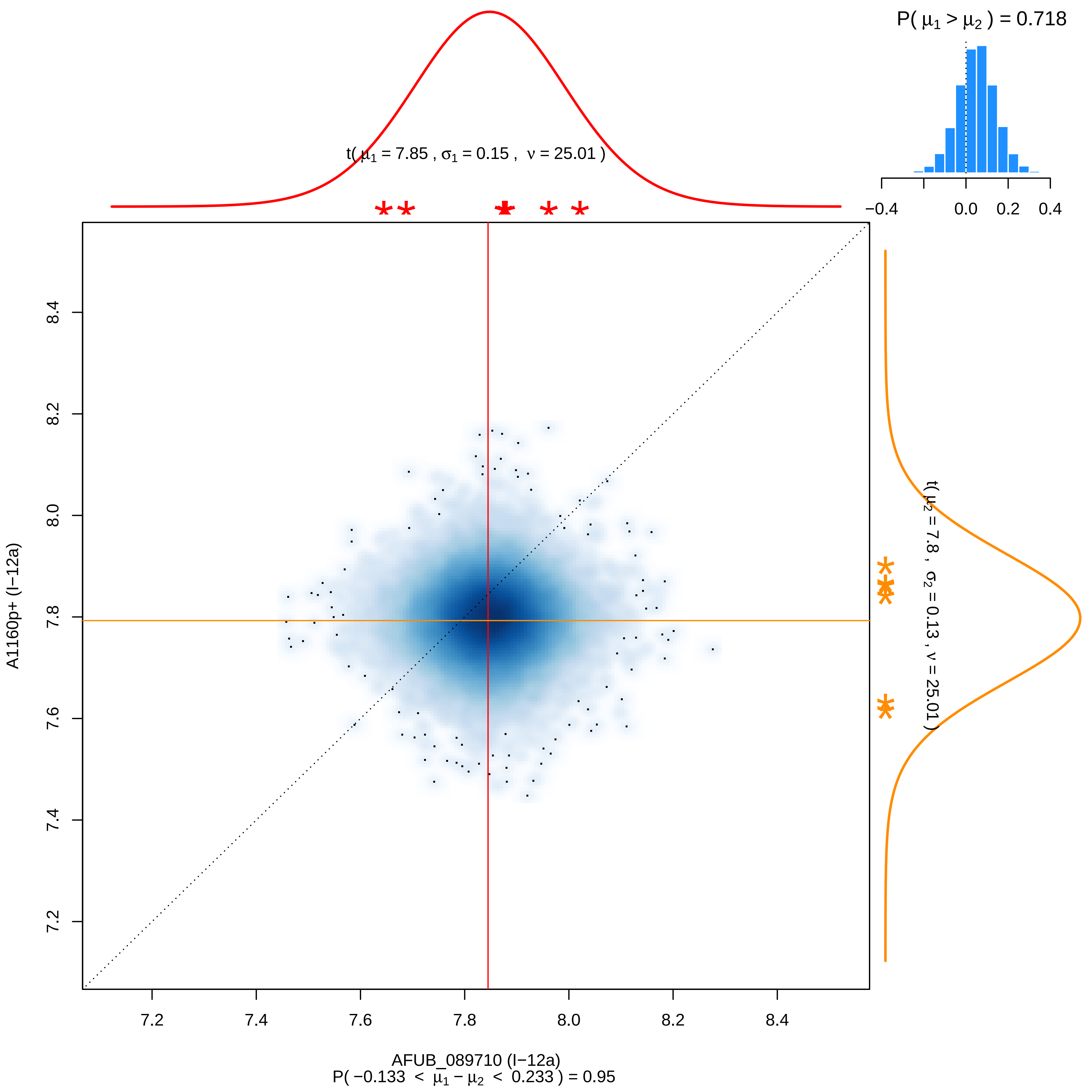

P( μ1 > μ2 ) = 0.718
−0.4
0.0
0.2
0.4
t( μ1 = 7.85 , σ1 = 0.15 , ν = 25.01 )
t( μ2 = 7.8 , σ2 = 0.13 , ν = 25.01 )
A1160p+ (I−12a)
7.2
7.4
7.6
7.8
8.0
8.2
8.4
AFUB_089710 (I−12a)
P( −0.133 < μ1 − μ2 < 0.233 ) = 0.95

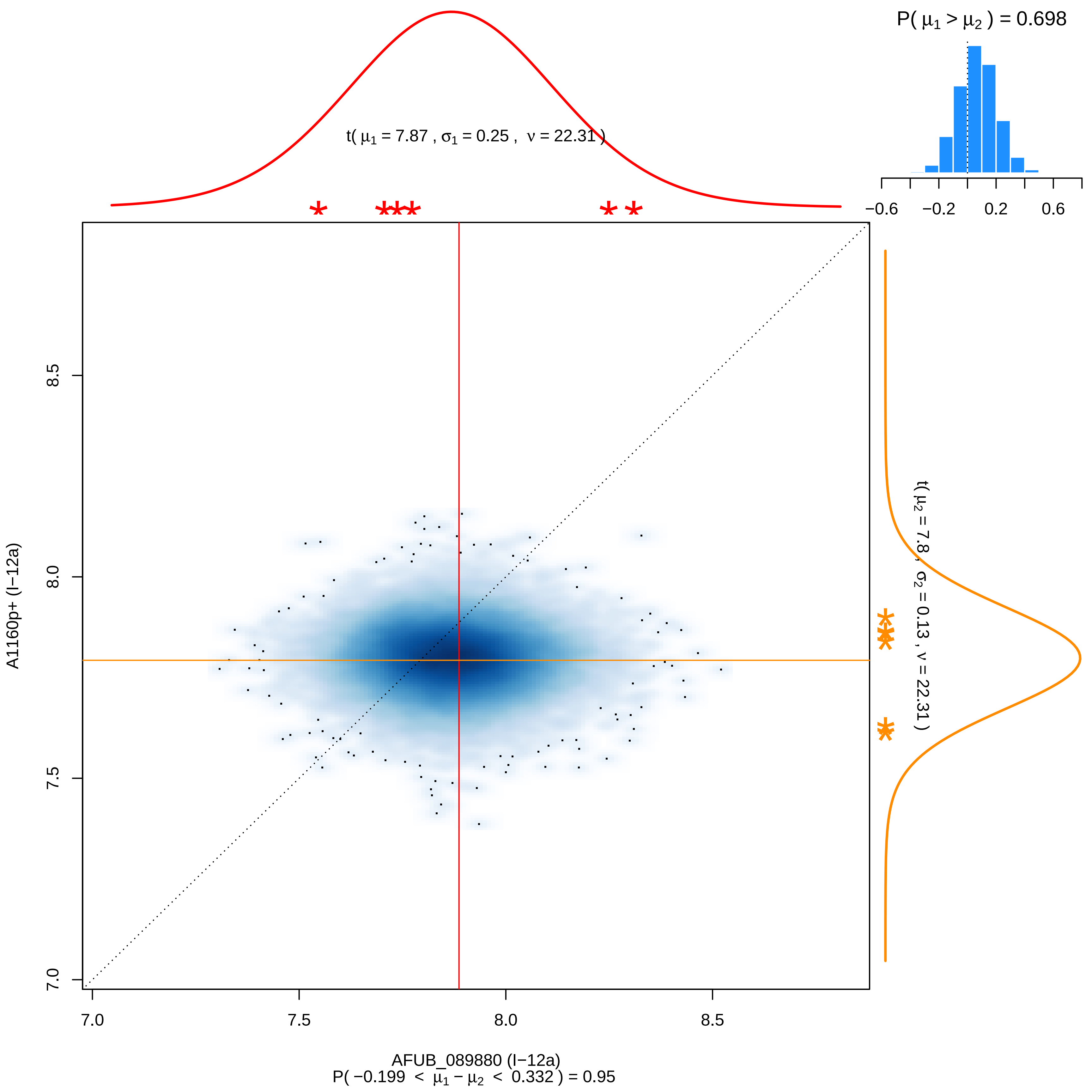

P( μ1 > μ2 ) = 0.698
−0.6
−0.2
0.2
0.6
t( μ1 = 7.87 , σ1 = 0.25 , ν = 22.31 )
t( μ2 = 7.8 , σ2 = 0.13 , ν = 22.31 )
A1160p+ (I−12a)
8.5
8.0
7.5
7.0
7.0
7.5
8.0
8.5
AFUB_089880 (I−12a)
P( −0.199 < μ1 − μ2 < 0.332 ) = 0.95

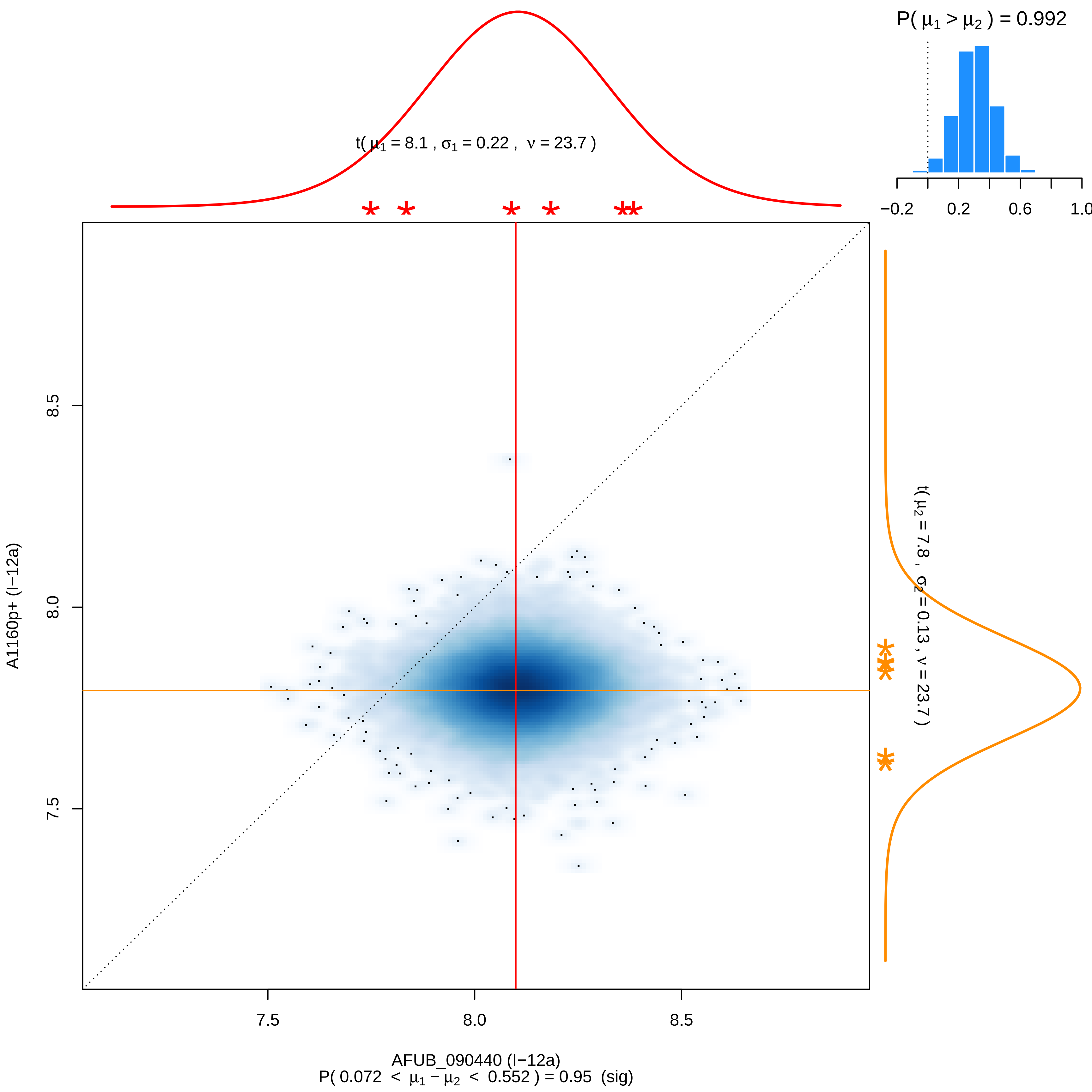

P( μ1 > μ2 ) = 0.992
−0.2
0.2
0.6
1.0
t( μ1 = 8.1 , σ1 = 0.22 , ν = 23.7 )
t( μ2 = 7.8 , σ2 = 0.13 , ν = 23.7 )
8.5
8.0
7.5
A1160p+ (I−12a)
7.5
8.0
8.5
AFUB_090440 (I−12a)
P( 0.072 < μ1 − μ2 < 0.552 ) = 0.95 (sig)

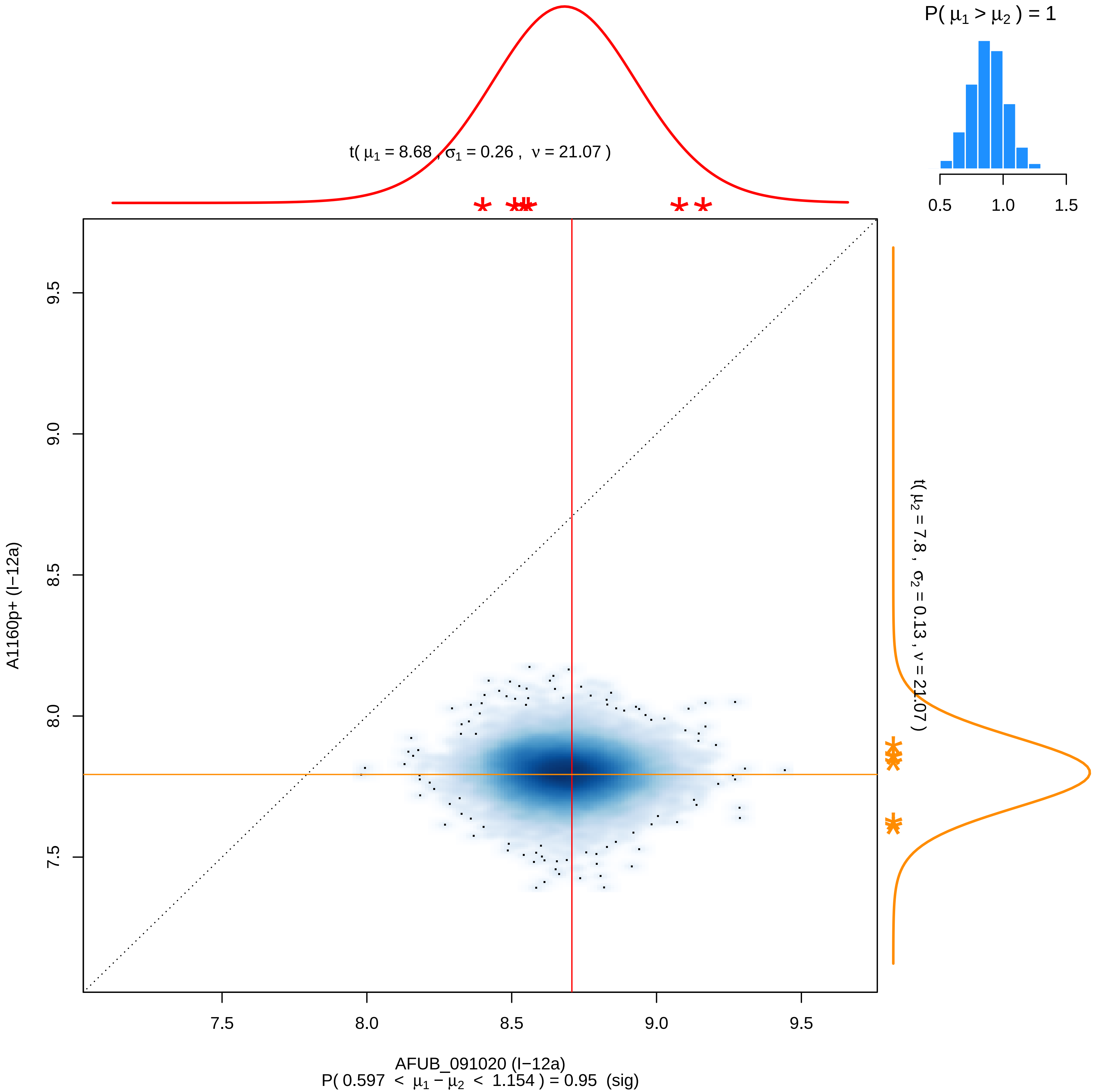

t( $\mu_1$ = 8.68 , $\sigma_1$ = 0.26 , $\nu$ = 21.07 )
P( $\mu_1$ > $\mu_2$ ) = 1
0.5
1.0
1.5
9.5
9.0
8.5
8.0
7.5
A1160p+ (I−12a)
t( $\mu_2$ = 7.8 , $\sigma_2$ = 0.13 , $\nu$ = 21.07 )
7.5
8.0
8.5
9.0
9.5
AFUB_091020 (I−12a)
P( 0.597 < $\mu_1 - \mu_2$ < 1.154 ) = 0.95 (sig)

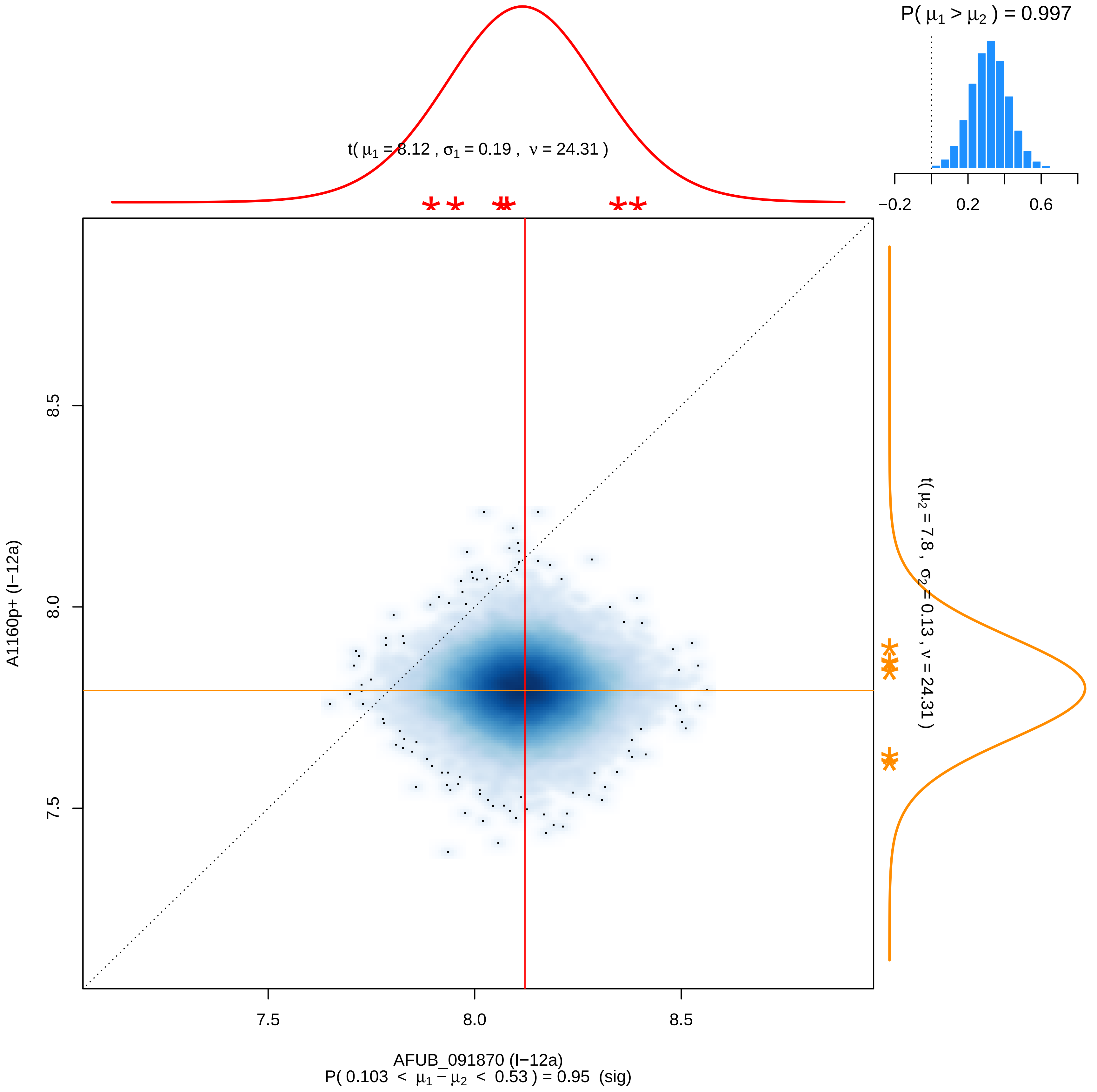

P( μ1 > μ2 ) = 0.997
−0.2
0.2
0.6
t( μ1 = 8.12 , σ1 = 0.19 , ν = 24.31 )
t( μ2 = 7.8 , σ2 = 0.13 , ν = 24.31 )
8.5
8.0
7.5
A1160p+ (I−12a)
7.5
8.0
8.5
AFUB_091870 (I−12a)
P( 0.103 < μ1 − μ2 < 0.53 ) = 0.95 (sig)

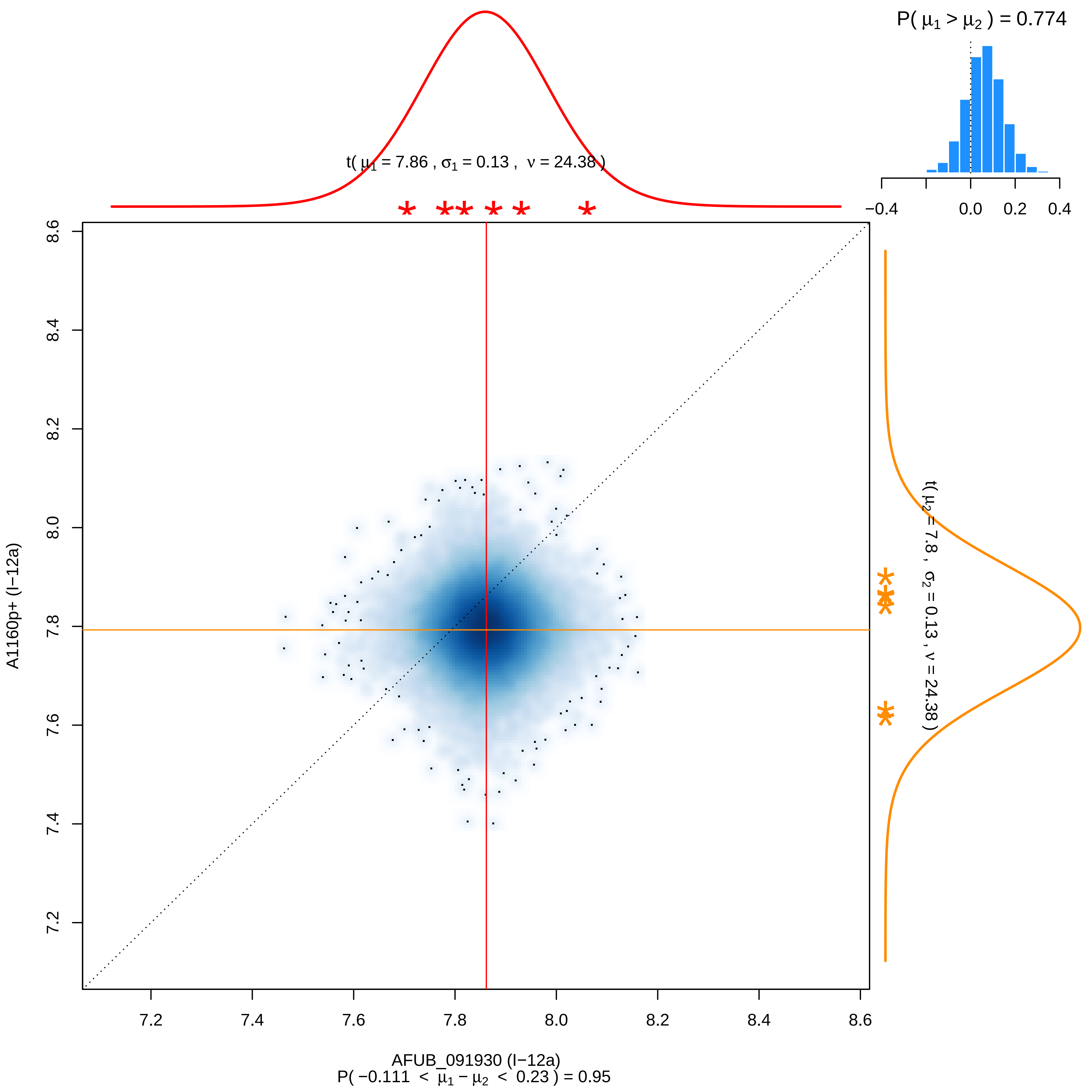

P( μ1 > μ2 ) = 0.774
−0.4
0.0
0.2
0.4
t( μ1 = 7.86 , σ1 = 0.13 , ν = 24.38 )
t( μ2 = 7.8 , σ2 = 0.13 , ν = 24.38 )
A1160p+ (I−12a)
AFUB_091930 (I−12a)
P( −0.111 < μ1 − μ2 < 0.23 ) = 0.95
7.2
7.4
7.6
7.8
8.0
8.2
8.4
8.6

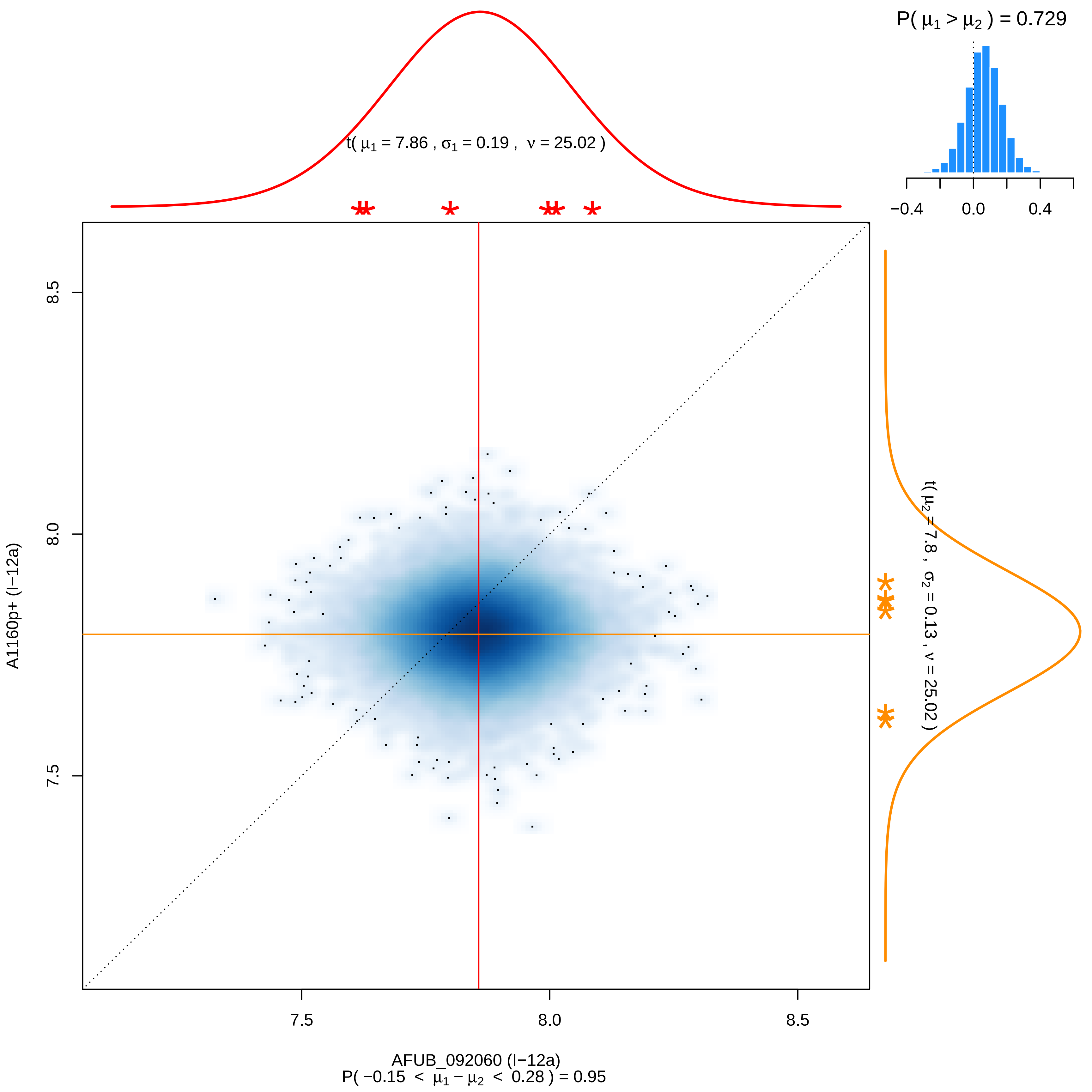
P( μ1 > μ2 ) = 0.729
−0.4
0.0
0.4
t( μ1 = 7.86 , σ1 = 0.19 , ν = 25.02 )
t( μ2 = 7.8 , σ2 = 0.13 , ν = 25.02 )
8.5
8.0
7.5
A1160p+ (I−12a)
7.5
8.0
8.5
AFUB_092060 (I−12a)
P( −0.15 < μ1 − μ2 < 0.28 ) = 0.95

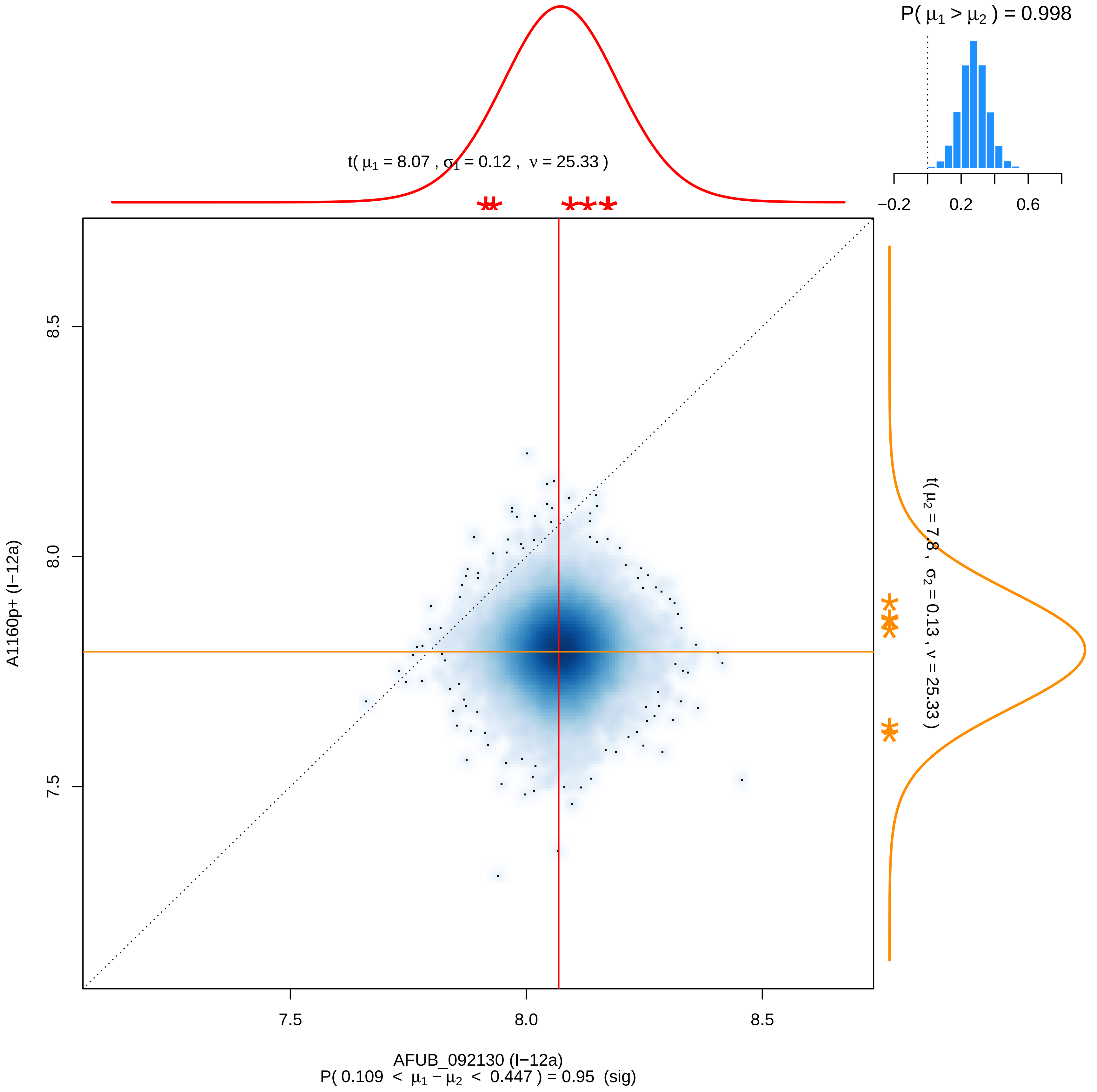

P( μ1 > μ2 ) = 0.998
−0.2
0.2
0.6
t( μ1 = 8.07 , σ1 = 0.12 , ν = 25.33 )
t( μ2 = 7.8 , σ2 = 0.13 , ν = 25.33 )
8.5
8.0
7.5
A1160p+ (I−12a)
7.5
8.0
8.5
AFUB_092130 (I−12a)
P( 0.109 < μ1 − μ2 < 0.447 ) = 0.95 (sig)

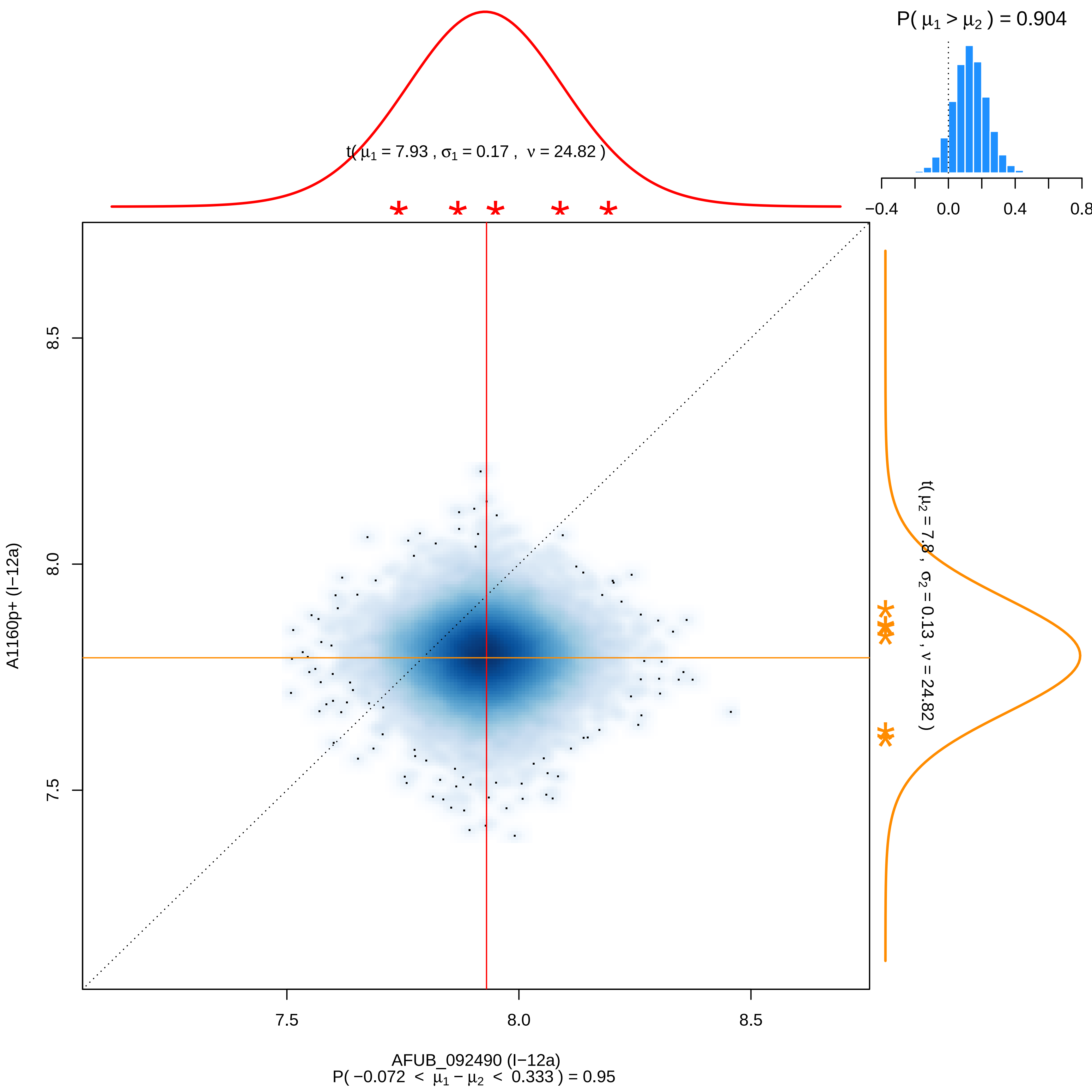

P( μ1 > μ2 ) = 0.904
−0.4 0.0 0.4 0.8
t( μ1 = 7.93 , σ1 = 0.17 , ν = 24.82 )
t( μ2 = 7.8 , σ2 = 0.13 , ν = 24.82 )
8.5
8.0
7.5
A1160p+ (I−12a)
7.5 8.0 8.5
AFUB_092490 (I−12a)
P( −0.072 < μ1 − μ2 < 0.333 ) = 0.95

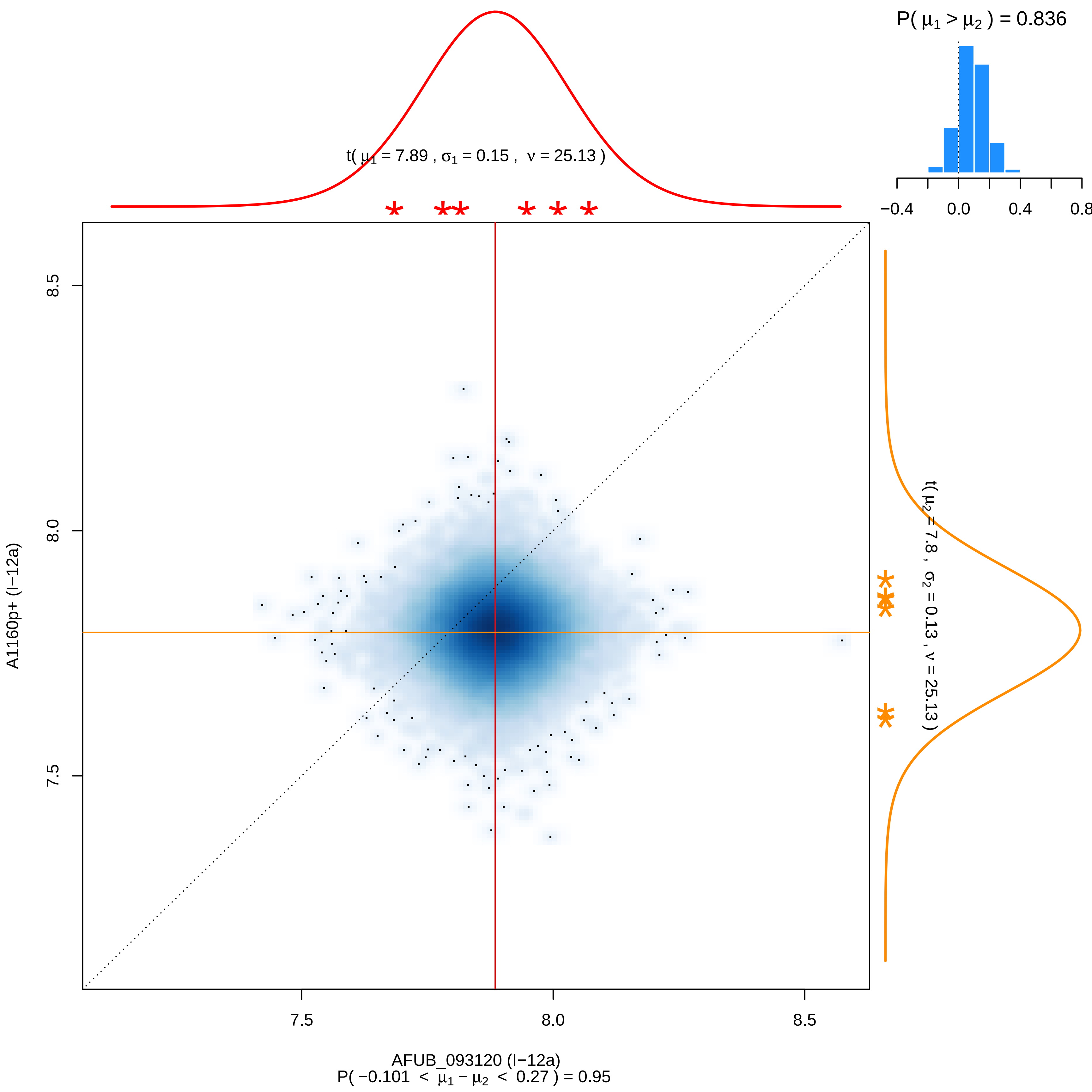
P( μ1 > μ2 ) = 0.836
−0.4
0.0
0.4
0.8
t( μ1 = 7.89 , σ1 = 0.15 , ν = 25.13 )
t( μ2 = 7.8 , σ2 = 0.13 , ν = 25.13 )
8.5
8.0
7.5
A1160p+ (I−12a)
7.5
8.0
8.5
AFUB_093120 (I−12a)
P( −0.101 < μ1 − μ2 < 0.27 ) = 0.95

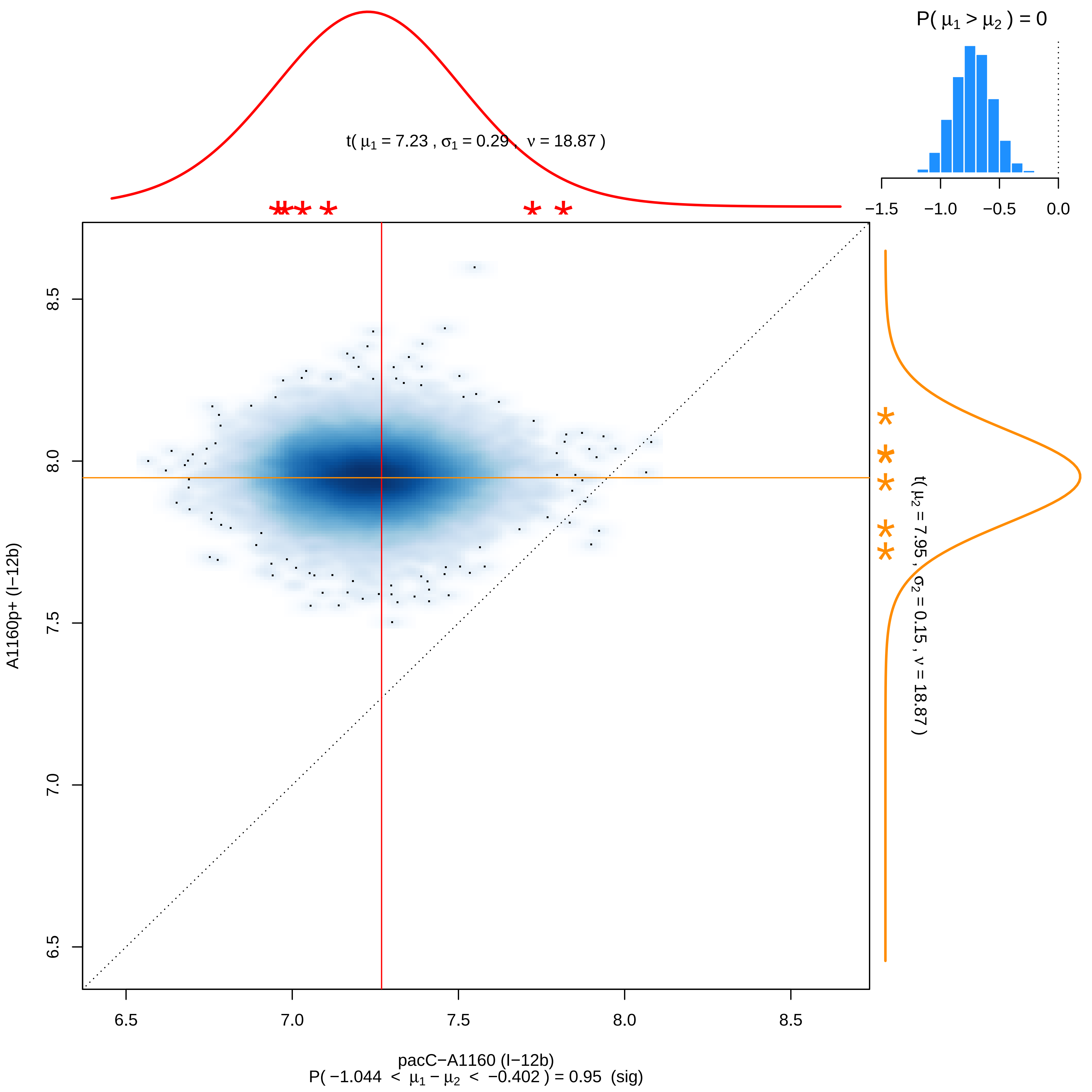

P( μ1 > μ2 ) = 0
−1.5
−1.0
−0.5
0.0
t( μ1 = 7.23 , σ1 = 0.29 , ν = 18.87 )
t( μ2 = 7.95 , σ2 = 0.15 , ν = 18.87 )
A1160p+ (I−12b)
8.5
8.0
7.5
7.0
6.5
6.5
7.0
7.5
8.0
8.5
pacC−A1160 (I−12b)
P( −1.044 < μ1 − μ2 < −0.402 ) = 0.95 (sig)

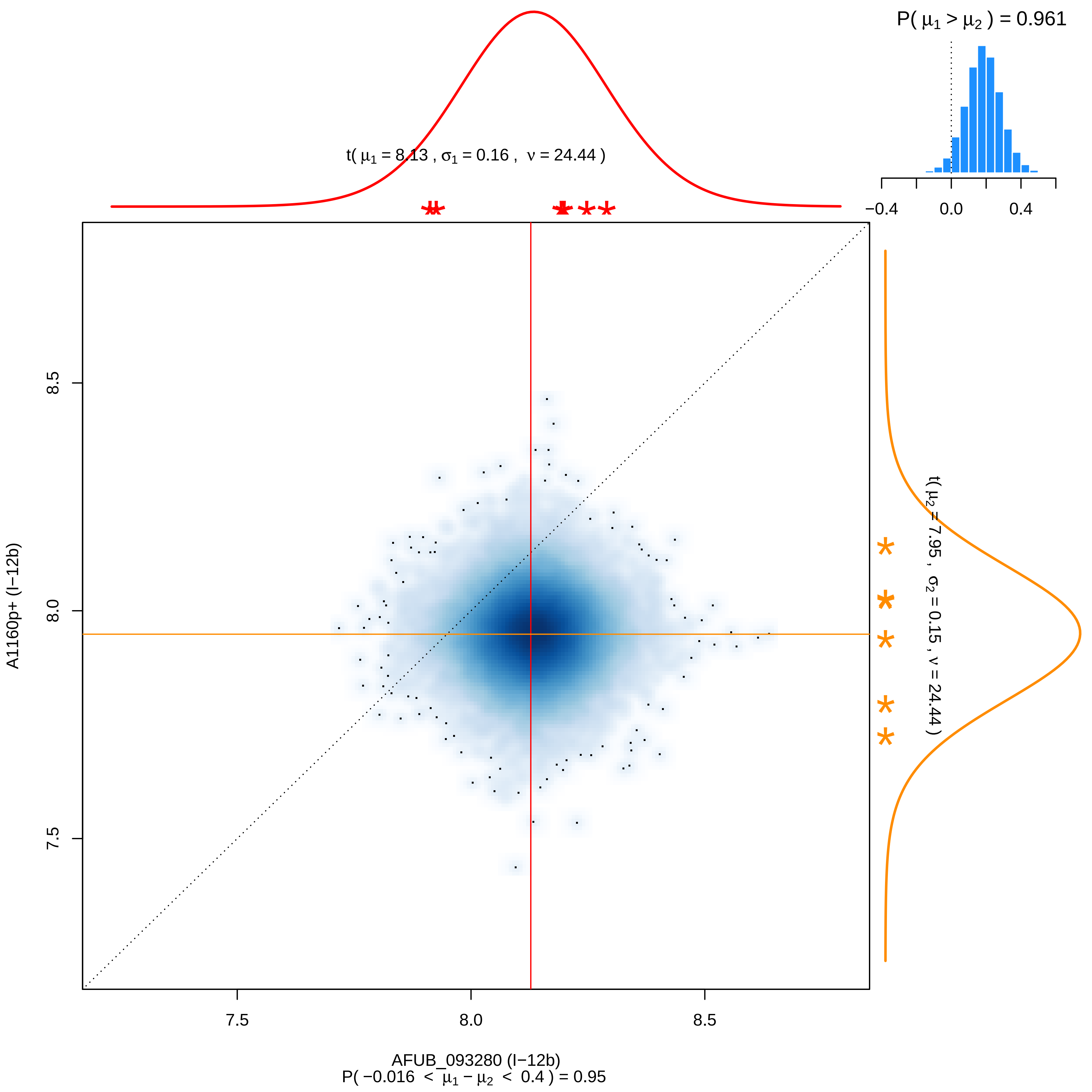
P( μ1 > μ2 ) = 0.961
t( μ1 = 8.13 , σ1 = 0.16 , ν = 24.44 )
−0.4
0.0
0.4
A1160p+ (I−12b)
8.5
8.0
7.5
7.5
8.0
8.5
t( μ2 = 7.95 , σ2 = 0.15 , ν = 24.44 )
AFUB_093280 (I−12b)
P( −0.016 < μ1 − μ2 < 0.4 ) = 0.95

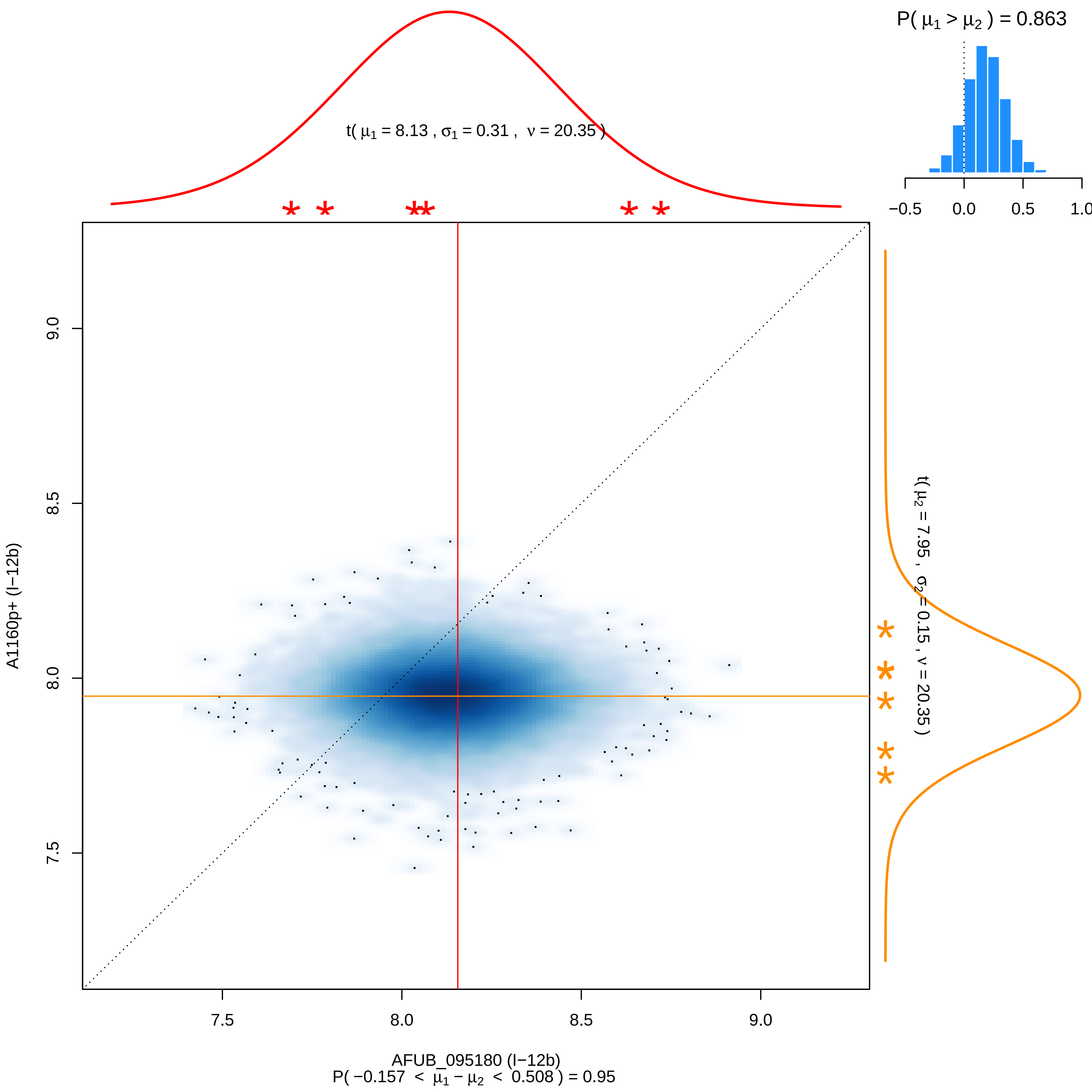

P( μ1 > μ2 ) = 0.863
−0.5
0.0
0.5
1.0
t( μ1 = 8.13 , σ1 = 0.31 , ν = 20.35 )
t( μ2 = 7.95 , σ2 = 0.15 , ν = 20.35 )
9.0
8.5
8.0
7.5
A1160p+ (I−12b)
7.5
8.0
8.5
9.0
AFUB_095180 (I−12b)
P( −0.157 < μ1 − μ2 < 0.508 ) = 0.95

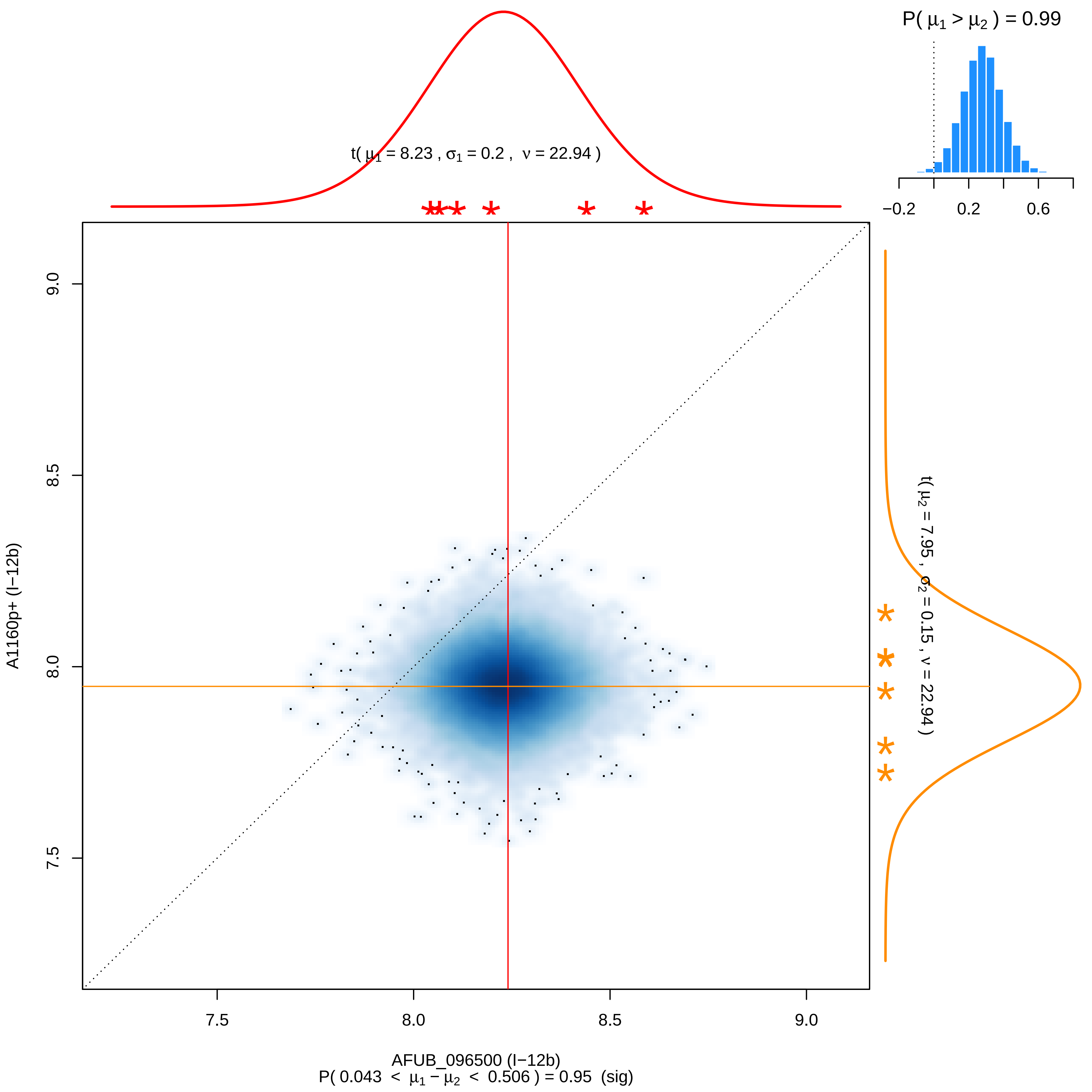

P( μ1 > μ2 ) = 0.99
−0.2
0.2
0.6
t( μ1 = 8.23 , σ1 = 0.2 , ν = 22.94 )
t( μ2 = 7.95 , σ2 = 0.15 , ν = 22.94 )
9.0
8.5
8.0
7.5
A1160p+ (I−12b)
7.5
8.0
8.5
9.0
AFUB_096500 (I−12b)
P( 0.043 < μ1 − μ2 < 0.506 ) = 0.95 (sig)

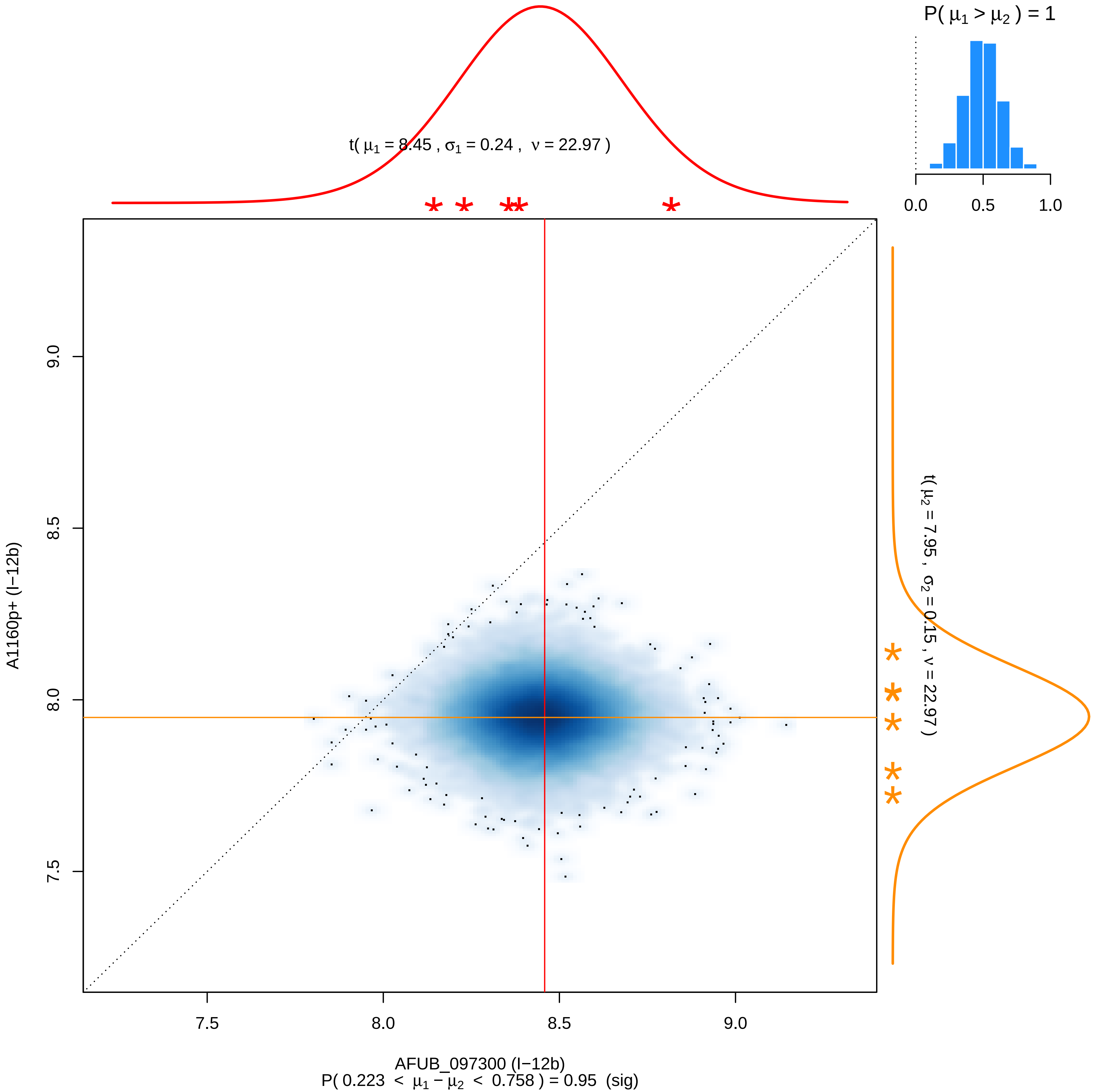

P( μ1 > μ2 ) = 1
0.0
0.5
1.0
t( μ1 = 8.45 , σ1 = 0.24 , ν = 22.97 )
t( μ2 = 7.95 , σ2 = 0.15 , ν = 22.97 )
A1160p+ (I−12b)
9.0
8.5
8.0
7.5
7.5
8.0
8.5
9.0
AFUB_097300 (I−12b)
P( 0.223 < μ1 − μ2 < 0.758 ) = 0.95 (sig)

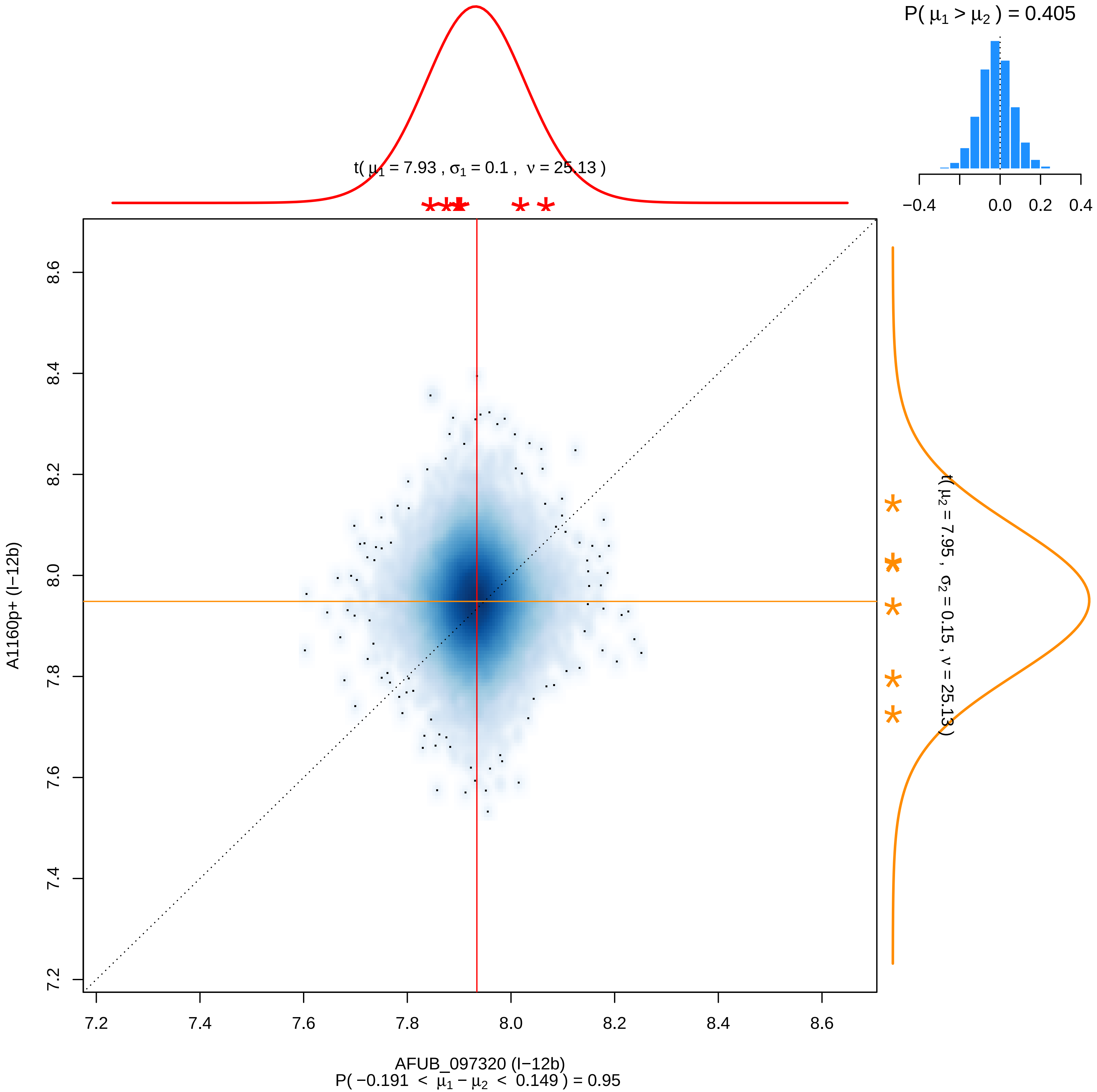

P( μ1 > μ2 ) = 0.405
−0.4
0.0
0.2
0.4
t( μ1 = 7.93 , σ1 = 0.1 , ν = 25.13 )
t( μ2 = 7.95 , σ2 = 0.15 , ν = 25.13 )
8.6
8.4
8.2
8.0
7.8
7.6
7.4
7.2
A1160p+ (I−12b)
7.2
7.4
7.6
7.8
8.0
8.2
8.4
8.6
AFUB_097320 (I−12b)
P( −0.191 < μ1 − μ2 < 0.149 ) = 0.95

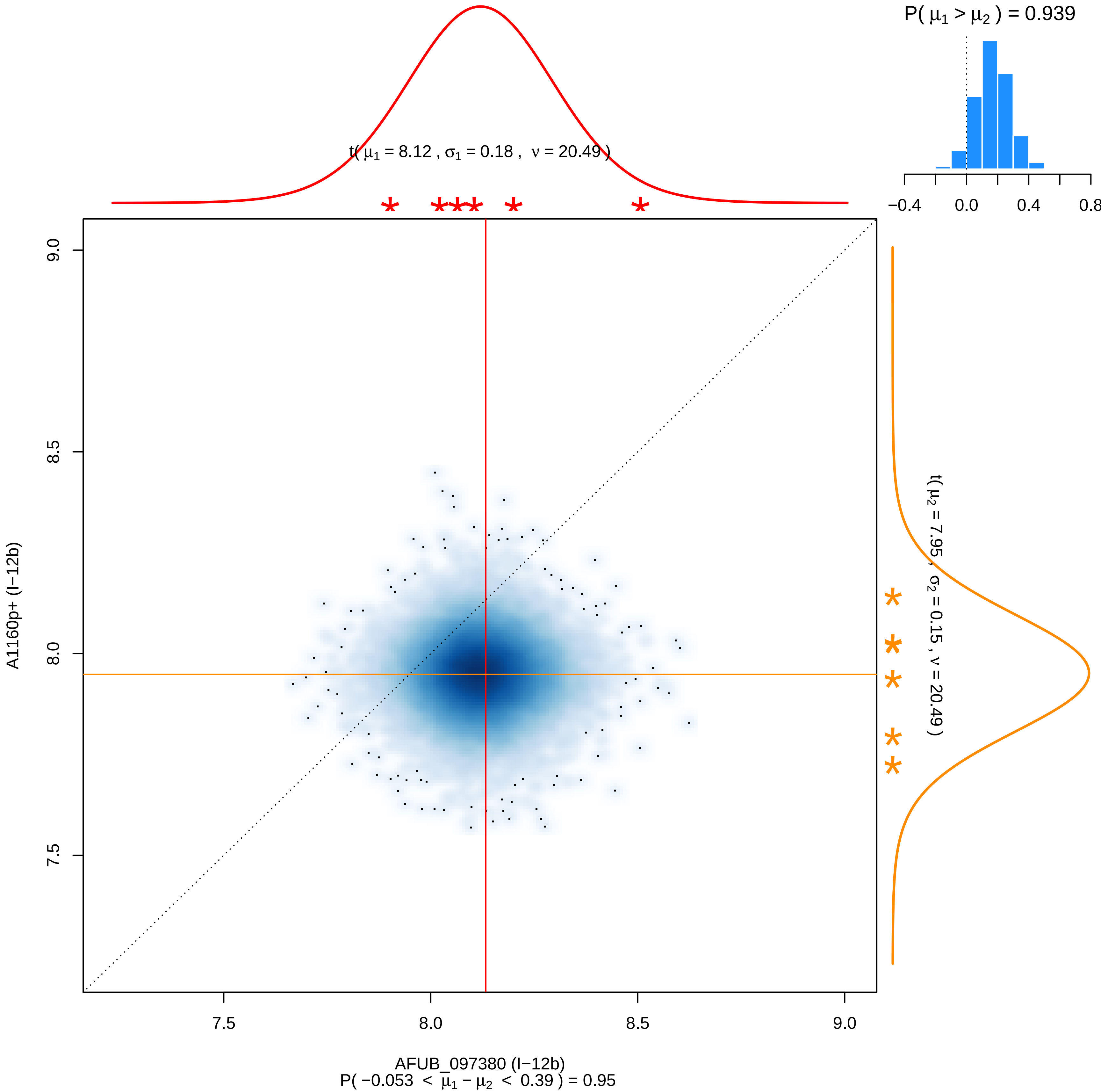

P( μ1 > μ2 ) = 0.939
−0.4
0.0
0.4
0.8
t( μ1 = 8.12 , σ1 = 0.18 , ν = 20.49 )
t( μ2 = 7.95 , σ2 = 0.15 , ν = 20.49 )
9.0
8.5
8.0
7.5
A1160p+ (I−12b)
7.5
8.0
8.5
9.0
AFUB_097380 (I−12b)
P( −0.053 < μ1 − μ2 < 0.39 ) = 0.95

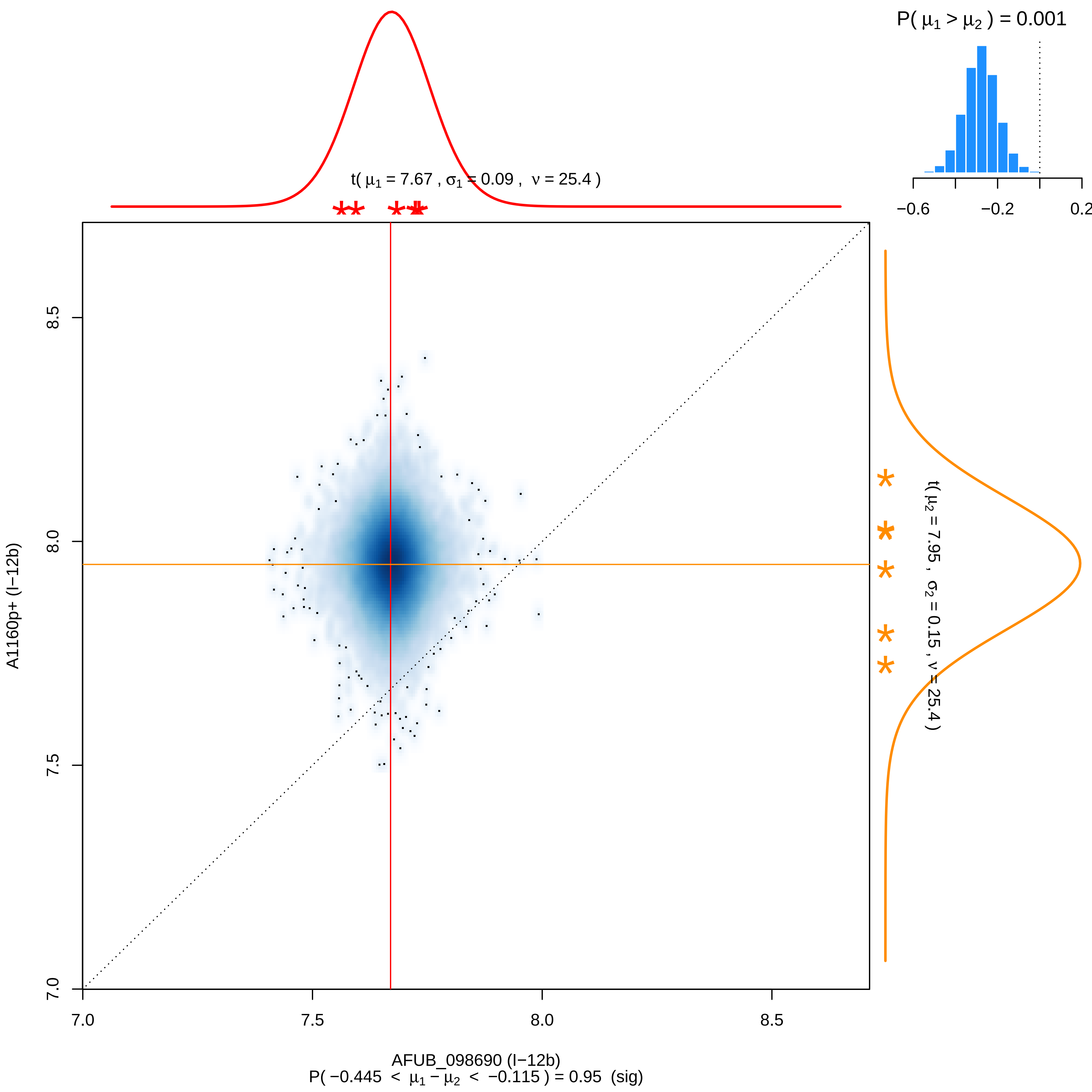

P( μ1 > μ2 ) = 0.001
−0.6
−0.2
0.2
t( μ1 = 7.67 , σ1 = 0.09 , ν = 25.4 )
t( μ2 = 7.95 , σ2 = 0.15 , ν = 25.4 )
8.5
8.0
7.5
7.0
A1160p+ (I−12b)
7.0
7.5
8.0
8.5
AFUB_098690 (I−12b)
P( −0.445 < μ1 − μ2 < −0.115 ) = 0.95 (sig)

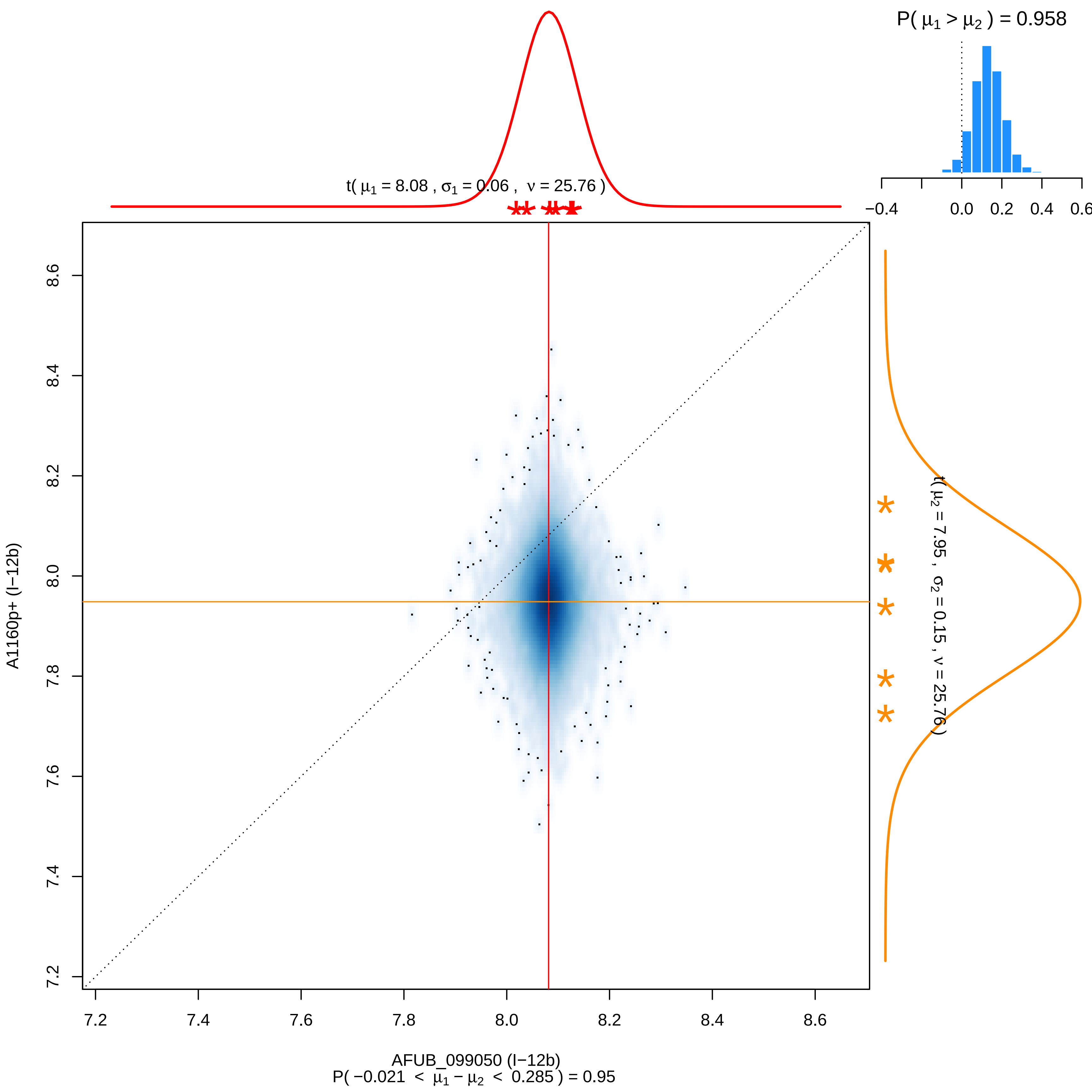

P( μ1 > μ2 ) = 0.958
−0.4
0.0
0.2
0.4
0.6
t( μ1 = 8.08 , σ1 = 0.06 , ν = 25.76 )
t( μ2 = 7.95 , σ2 = 0.15 , ν = 25.76 )
8.6
8.4
8.2
8.0
7.8
7.6
7.4
7.2
A1160p+ (I−12b)
7.2
7.4
7.6
7.8
8.0
8.2
8.4
8.6
AFUB_099050 (I−12b)
P( −0.021 < μ1 − μ2 < 0.285 ) = 0.95

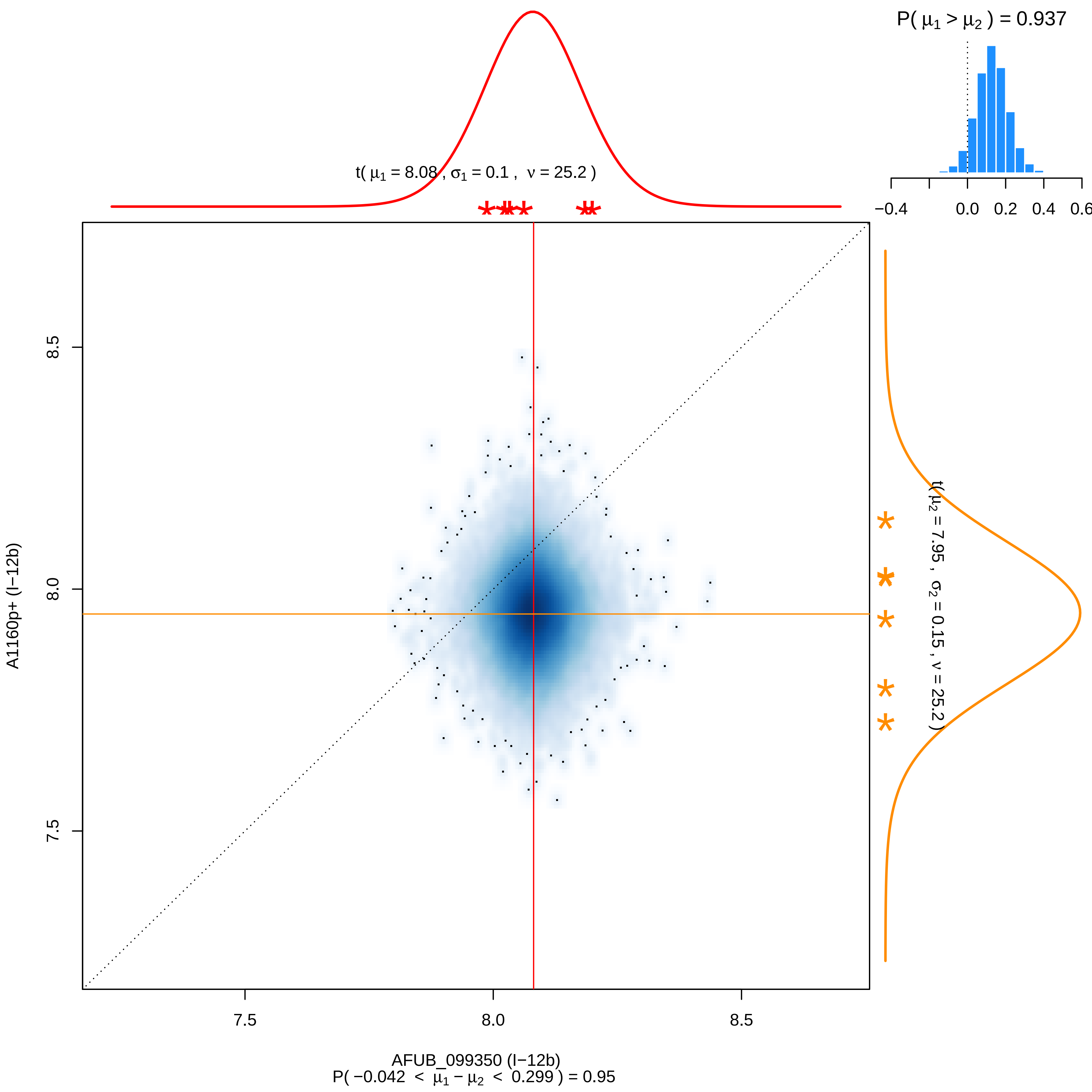

P( μ1 > μ2 ) = 0.937
−0.4
0.0
0.2
0.4
0.6
t( μ1 = 8.08 , σ1 = 0.1 , ν = 25.2 )
t( μ2 = 7.95 , σ2 = 0.15 , ν = 25.2 )
8.5
8.0
7.5
A1160p+ (I−12b)
7.5
8.0
8.5
AFUB_099350 (I−12b)
P( −0.042 < μ1 − μ2 < 0.299 ) = 0.95

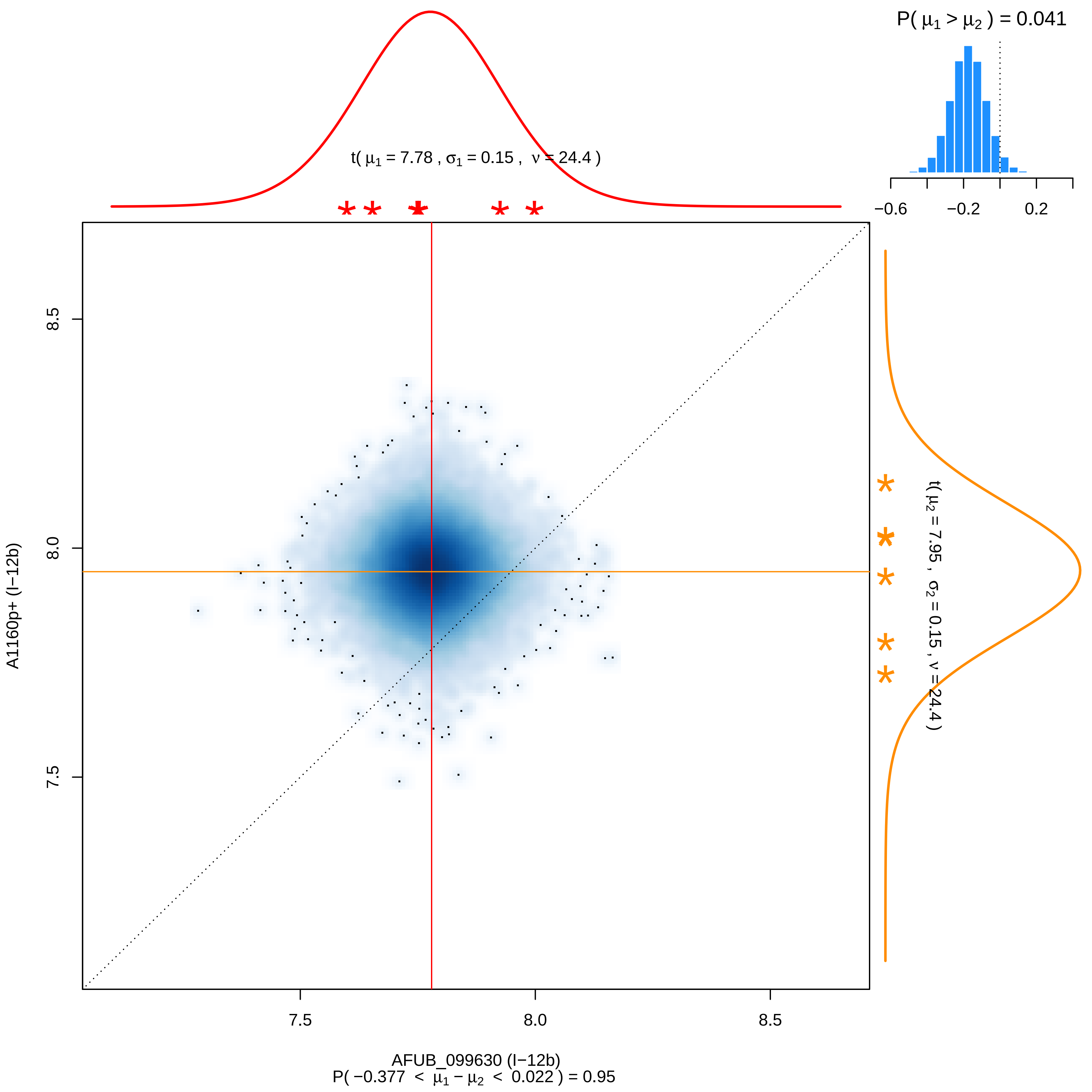

P( $\mu_1 > \mu_2$ ) = 0.041
−0.6
−0.2
0.2
t( $\mu_1$ = 7.78 , $\sigma_1$ = 0.15 , $\nu$ = 24.4 )
t( $\mu_2$ = 7.95 , $\sigma_2$ = 0.15 , $\nu$ = 24.4 )
8.5
8.0
7.5
A1160p+ (I−12b)
7.5
8.0
8.5
AFUB_099630 (I−12b)
P( −0.377 < $\mu_1 - \mu_2$ < 0.022 ) = 0.95

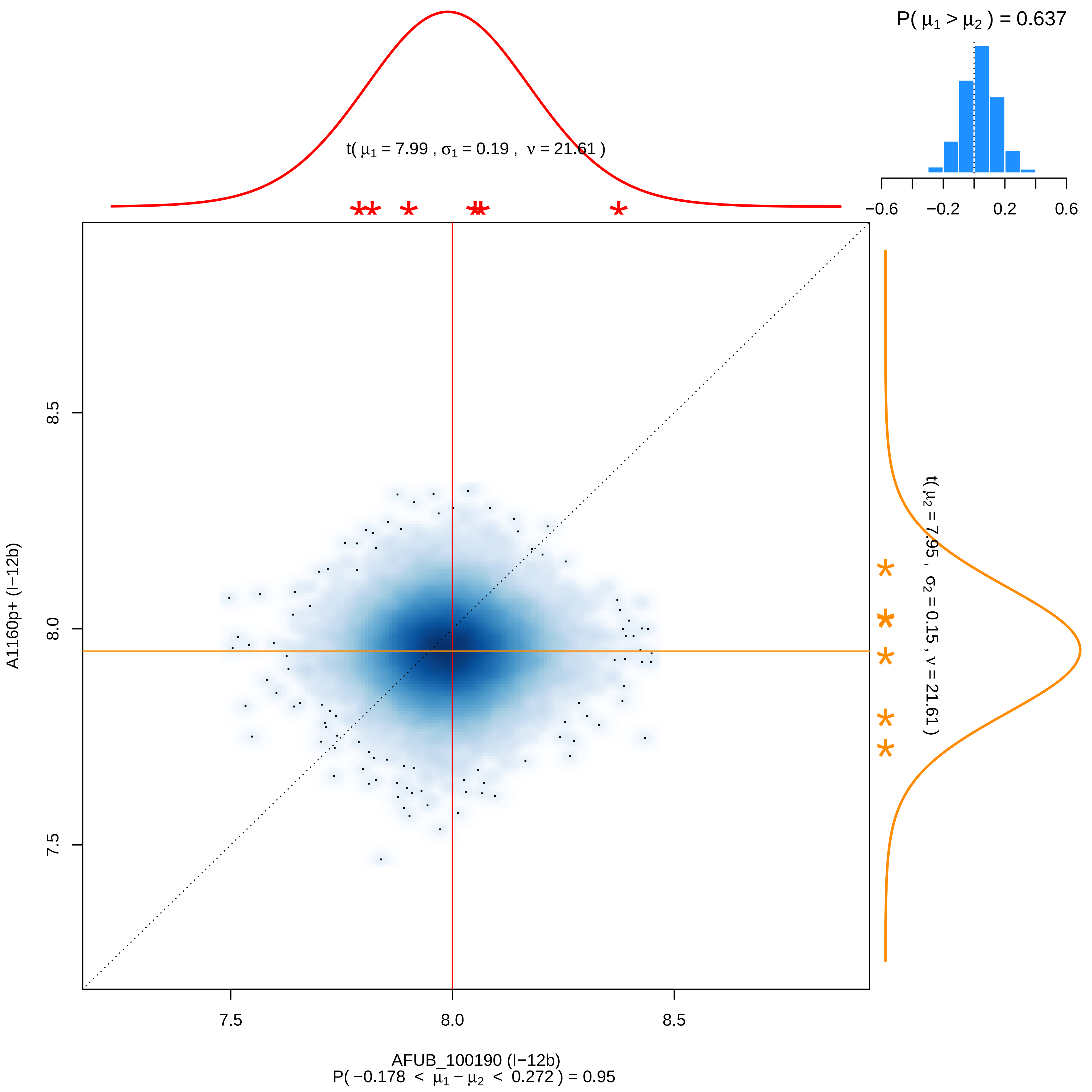

P( $\mu_1 > \mu_2$ ) = 0.637
−0.6
−0.2
0.2
0.6
t( $\mu_1$ = 7.99 , $\sigma_1$ = 0.19 , $\nu$ = 21.61 )
t( $\mu_2$ = 7.95 , $\sigma_2$ = 0.15 , $\nu$ = 21.61 )
A1160p+ (I−12b)
7.5
8.0
8.5
AFUB_100190 (I−12b)
P( −0.178 < $\mu_1 - \mu_2$ < 0.272 ) = 0.95

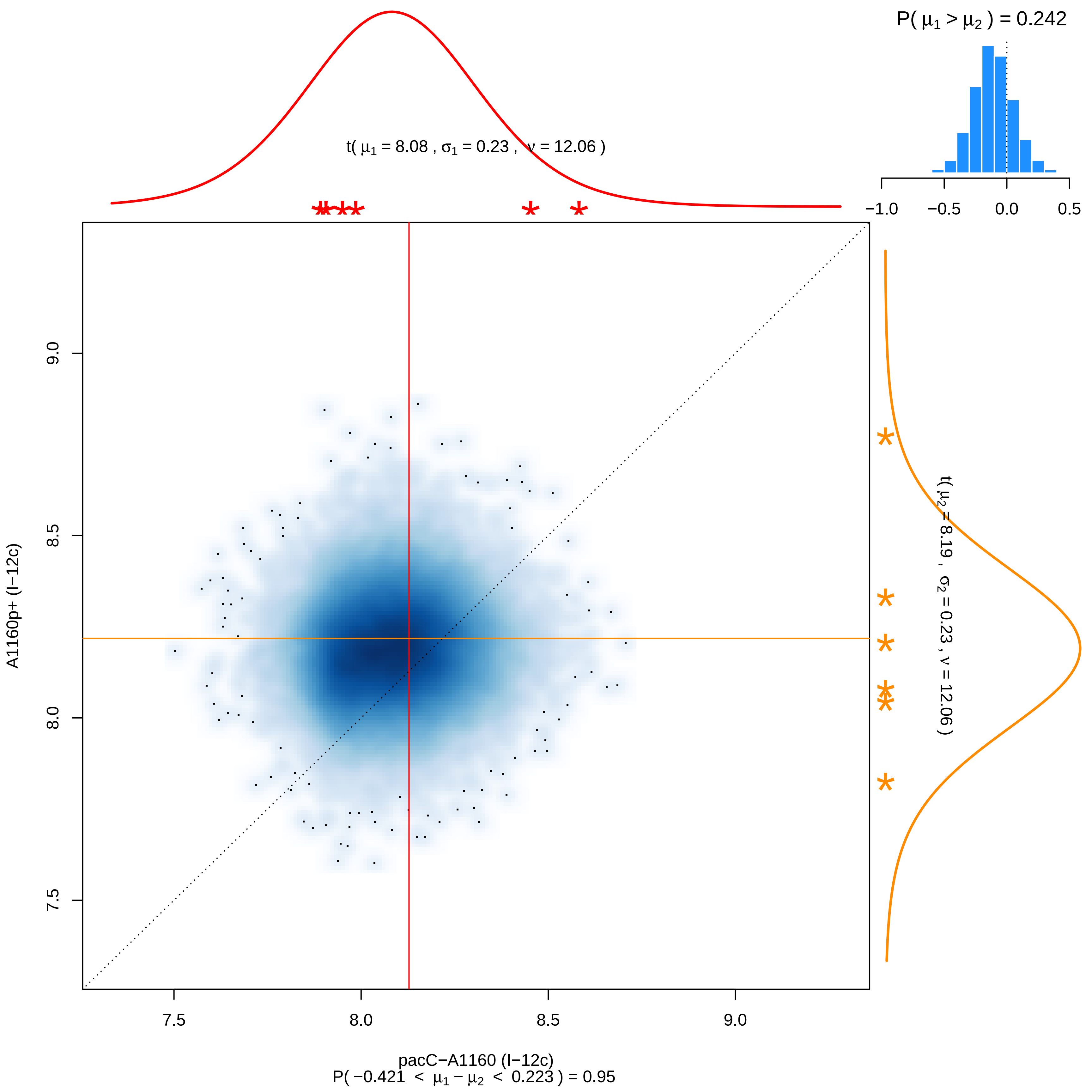

P( μ1 > μ2 ) = 0.242
−1.0
−0.5
0.0
0.5
t( μ1 = 8.08 , σ1 = 0.23 , ν = 12.06 )
t( μ2 = 8.19 , σ2 = 0.23 , ν = 12.06 )
9.0
8.5
8.0
7.5
A1160p+ (I−12c)
7.5
8.0
8.5
9.0
pacC−A1160 (I−12c)
P( −0.421 < μ1 − μ2 < 0.223 ) = 0.95

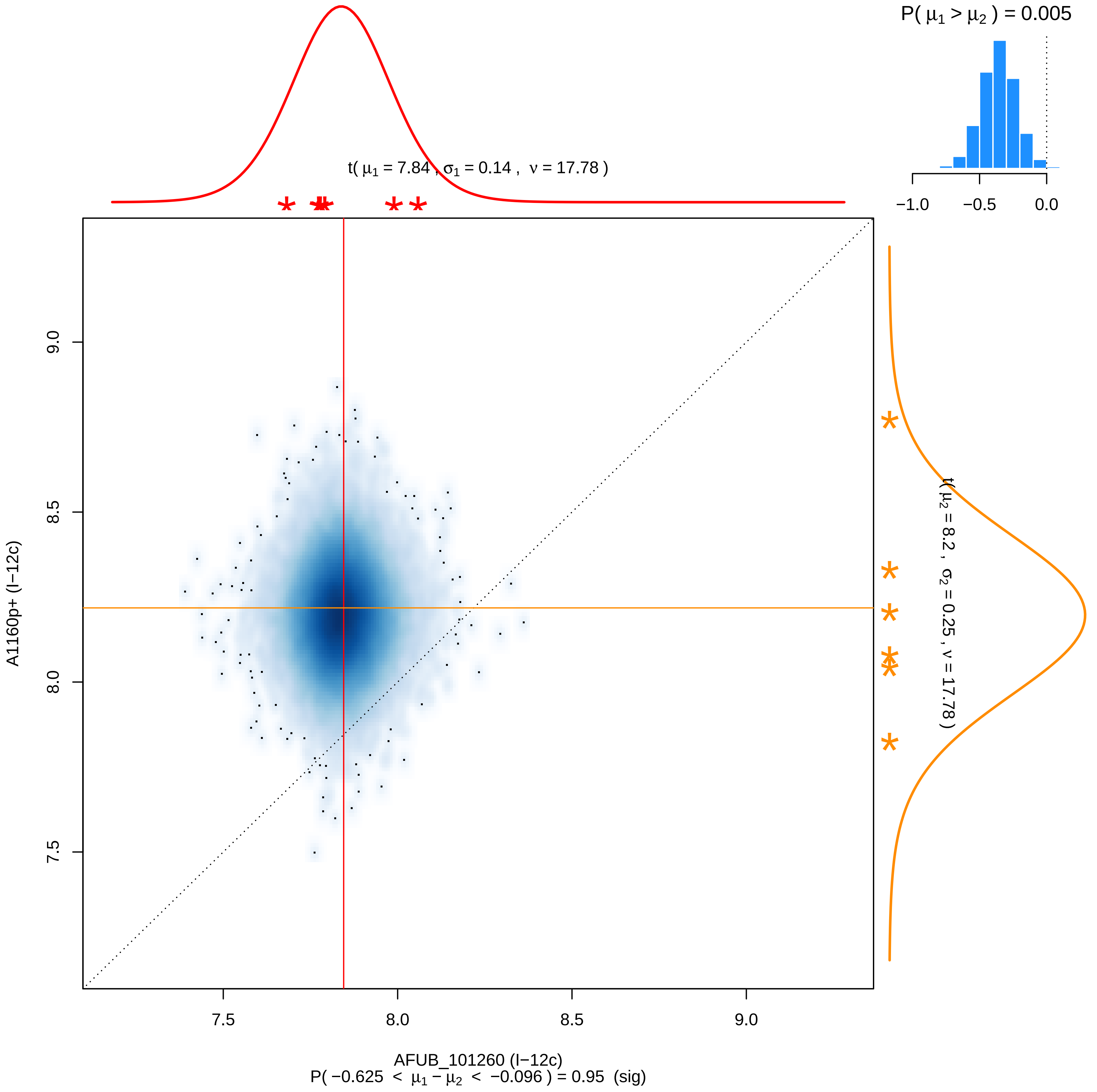

P( μ1 > μ2 ) = 0.005
−1.0
−0.5
0.0
t( μ1 = 7.84 , σ1 = 0.14 , ν = 17.78 )
t( μ2 = 8.2 , σ2 = 0.25 , ν = 17.78 )
A1160p+ (I−12c)
9.0
8.5
8.0
7.5
7.5
8.0
8.5
9.0
AFUB_101260 (I−12c)
P( −0.625 < μ1 − μ2 < −0.096 ) = 0.95 (sig)

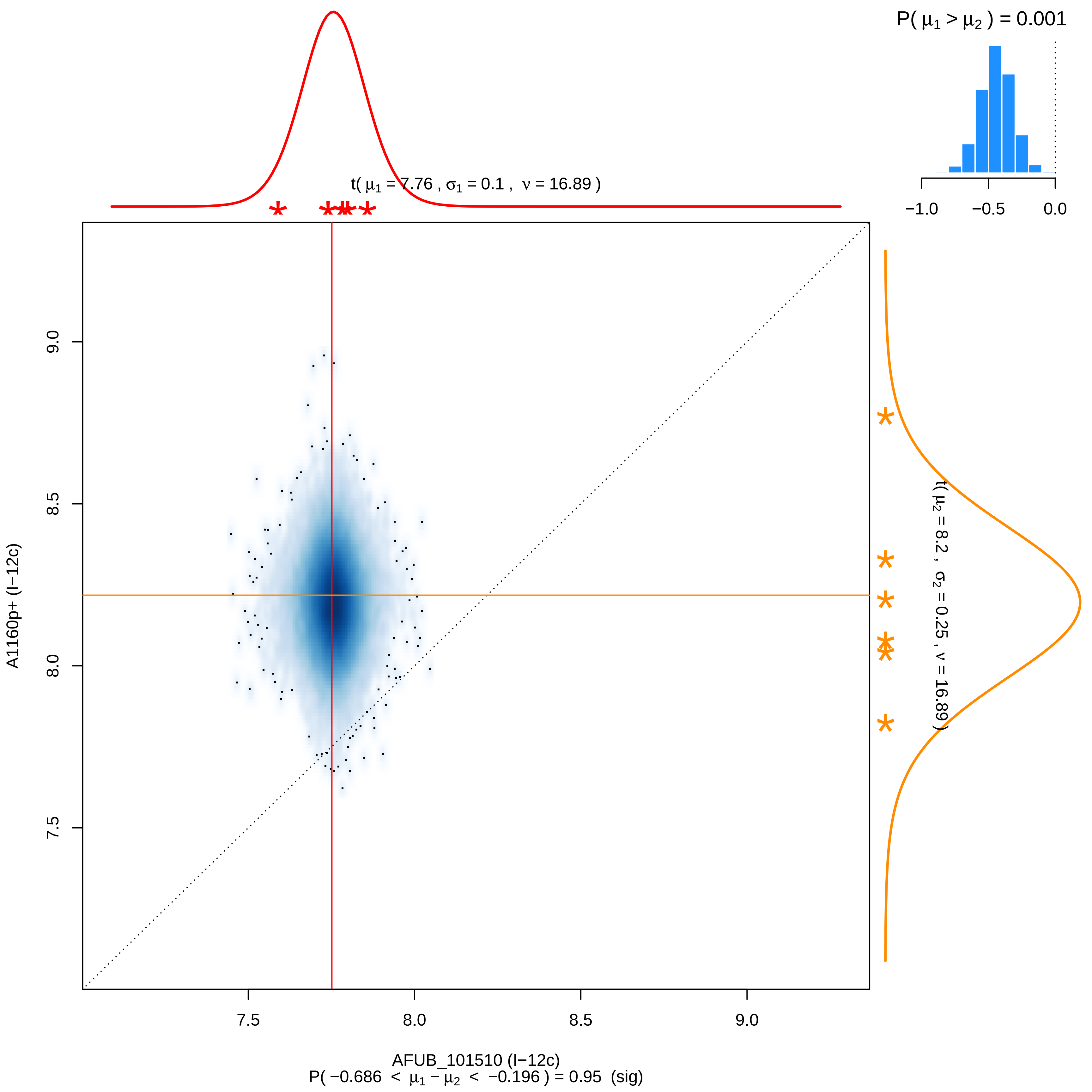

P( μ1 > μ2 ) = 0.001
−1.0
−0.5
0.0
t( μ1 = 7.76 , σ1 = 0.1 , ν = 16.89 )
t( μ2 = 8.2 , σ2 = 0.25 , ν = 16.89 )
9.0
8.5
8.0
7.5
A1160p+ (I−12c)
7.5
8.0
8.5
9.0
AFUB_101510 (I−12c)
P( −0.686 < μ1 − μ2 < −0.196 ) = 0.95 (sig)

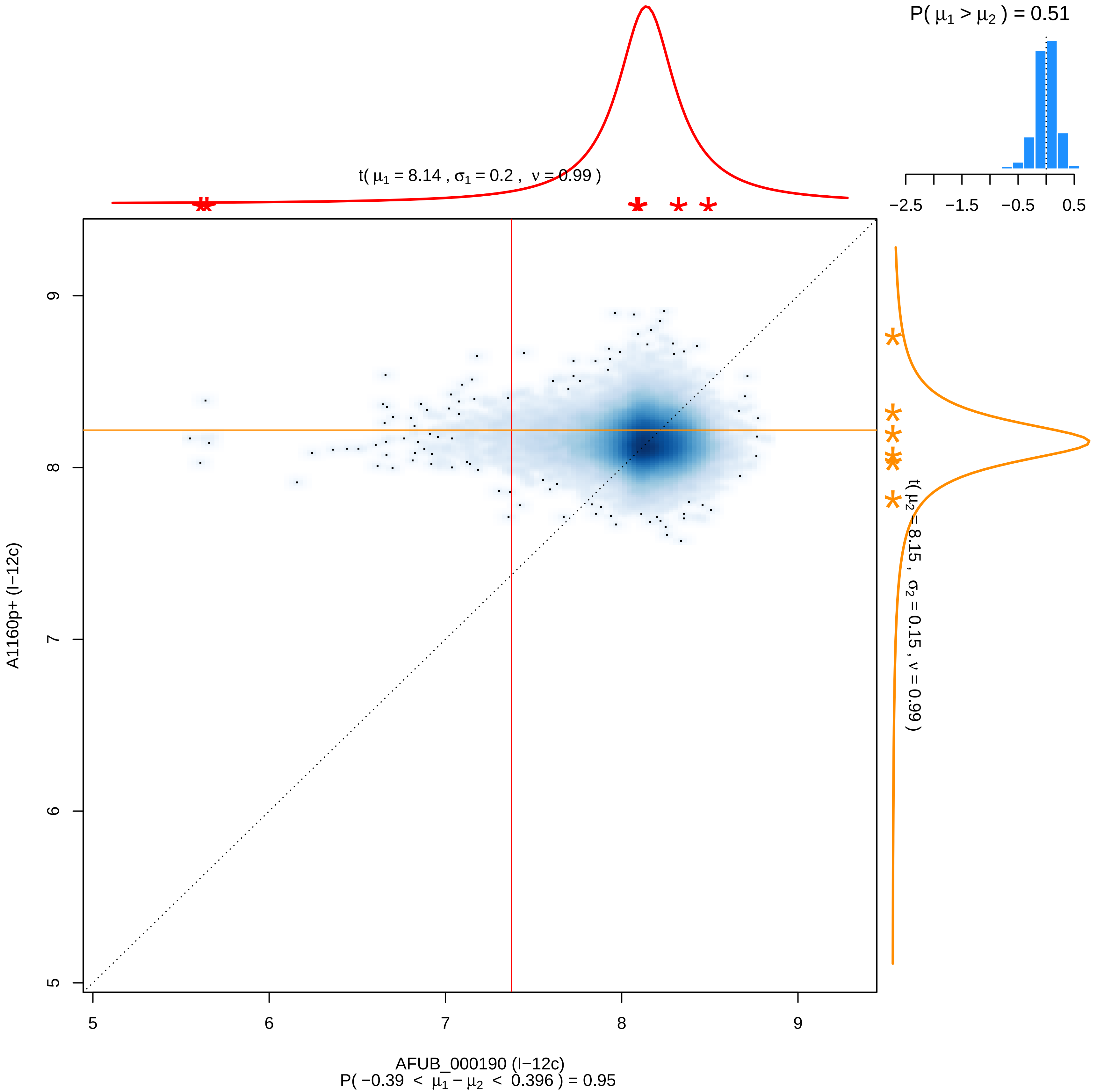

P( μ1 > μ2 ) = 0.51
−2.5
−1.5
−0.5
0.5
t( μ1 = 8.14 , σ1 = 0.2 , ν = 0.99 )
t( μ2 = 8.15 , σ2 = 0.15 , ν = 0.99 )
A1160p+ (I−12c)
5
6
7
8
9
AFUB_000190 (I−12c)
P( −0.39 < μ1 − μ2 < 0.396 ) = 0.95

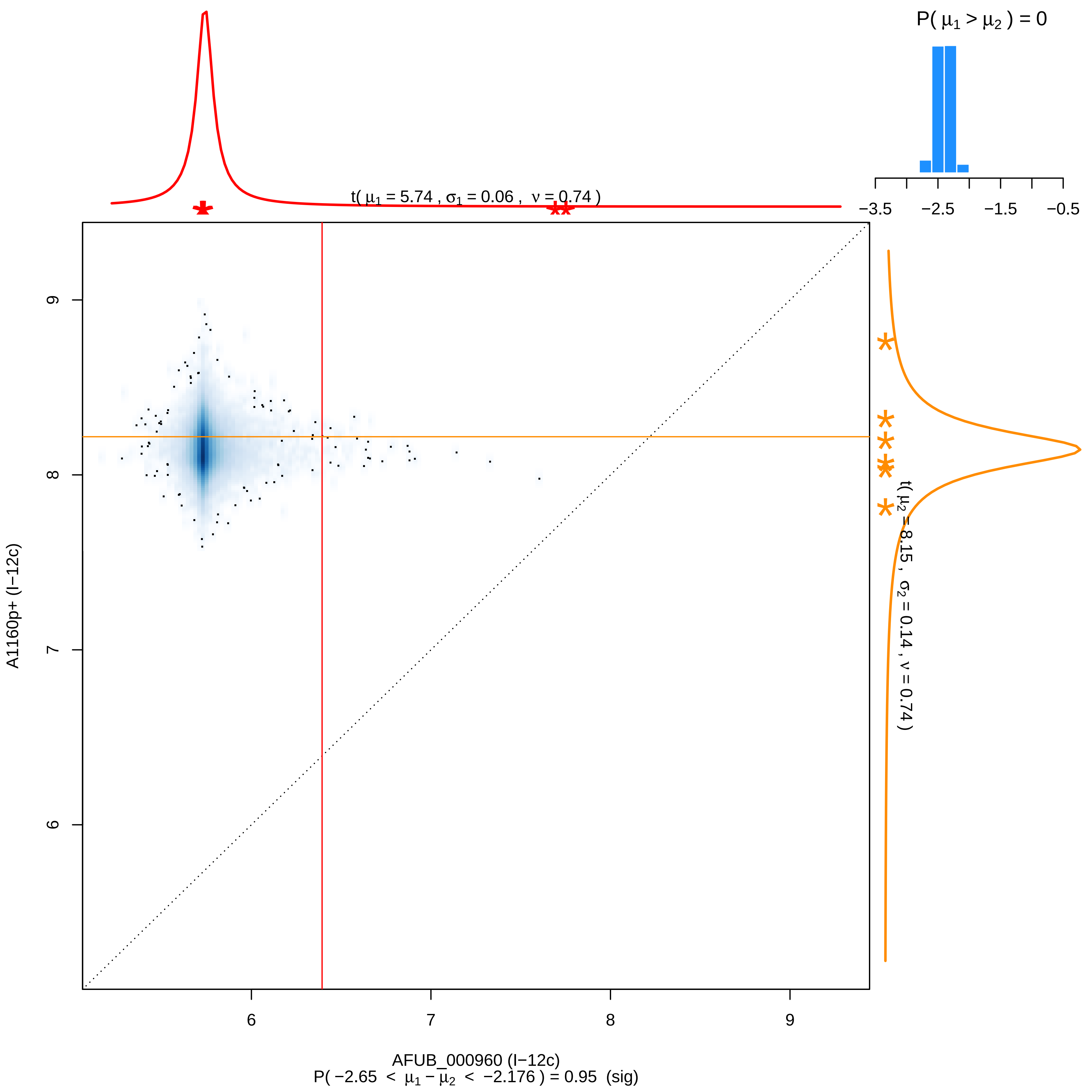

P( μ1 > μ2 ) = 0
−3.5
−2.5
−1.5
−0.5
t( μ1 = 5.74 , σ1 = 0.06 , ν = 0.74 )
9
8
7
6
A1160p+ (I−12c)
t( μ2 = 8.15 , σ2 = 0.14 , ν = 0.74 )
6
7
8
9
AFUB_000960 (I−12c)
P( −2.65 < μ1 − μ2 < −2.176 ) = 0.95 (sig)

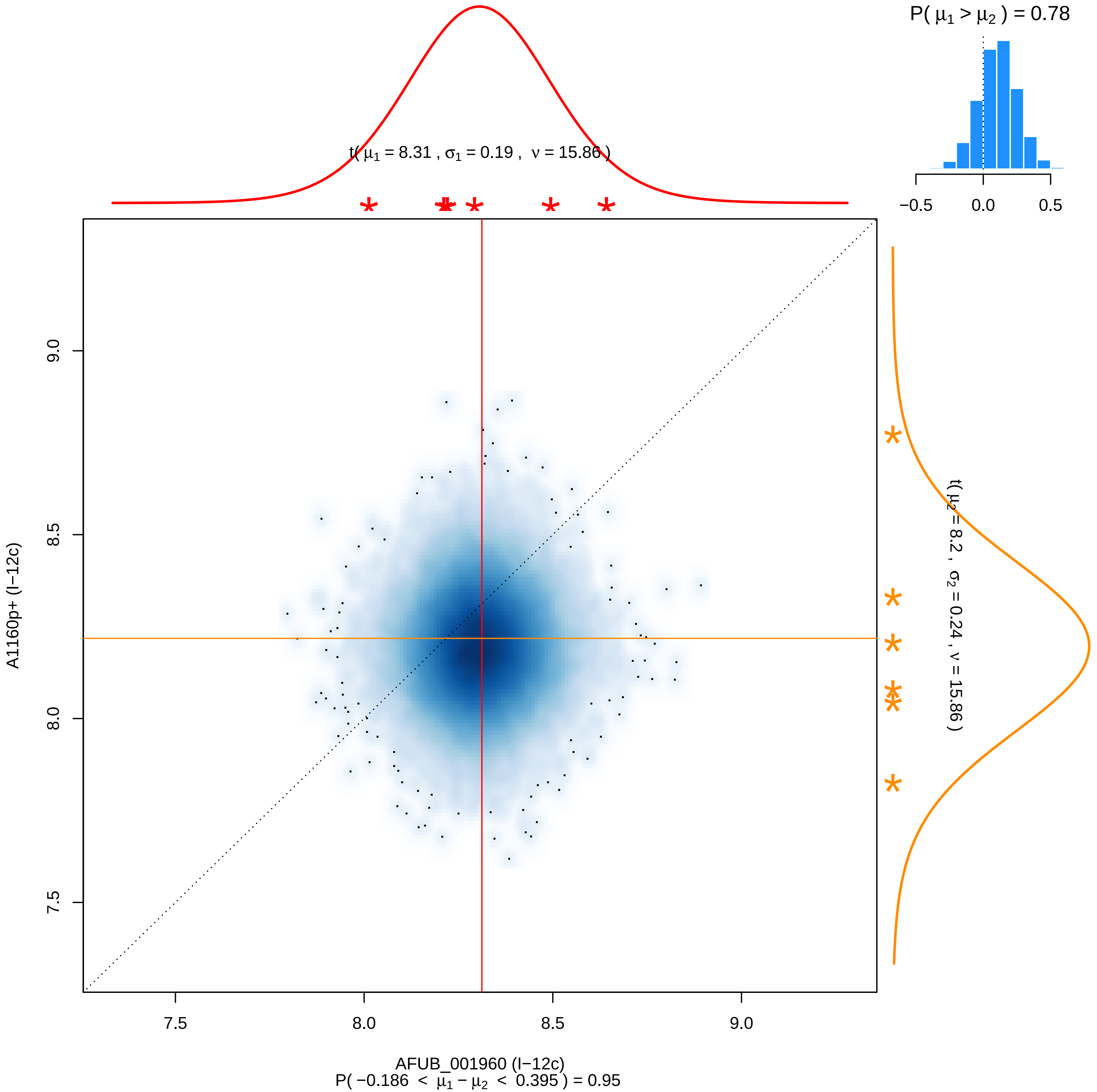

P( μ1 > μ2 ) = 0.78
−0.5
0.0
0.5
t( μ1 = 8.31 , σ1 = 0.19 , ν = 15.86 )
t( μ2 = 8.2 , σ2 = 0.24 , ν = 15.86 )
9.0
8.5
8.0
7.5
A1160p+ (I−12c)
7.5
8.0
8.5
9.0
AFUB_001960 (I−12c)
P( −0.186 < μ1 − μ2 < 0.395 ) = 0.95

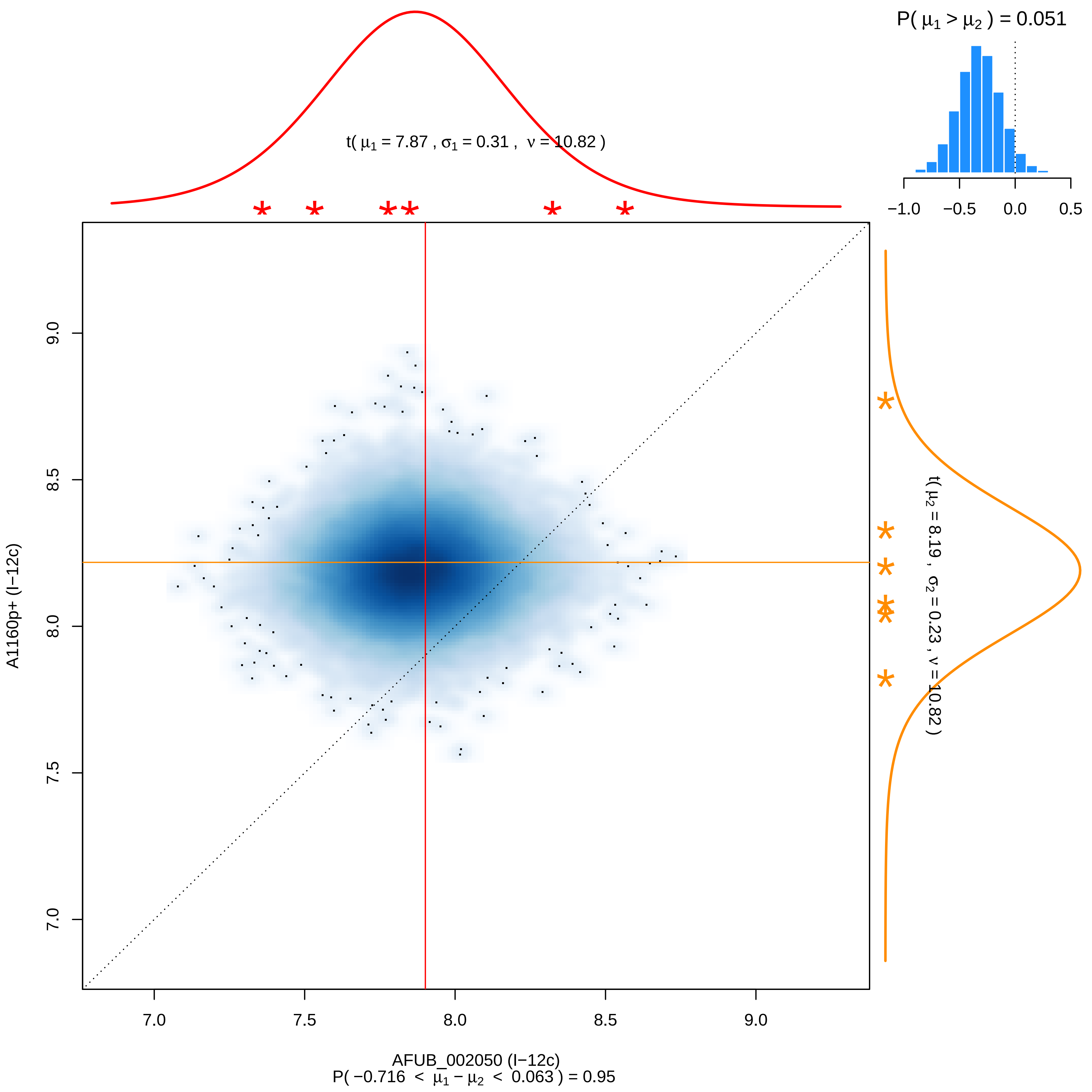

P( μ1 > μ2 ) = 0.051
−1.0 −0.5 0.0 0.5
t( μ1 = 7.87 , σ1 = 0.31 , ν = 10.82 )
t( μ2 = 8.19 , σ2 = 0.23 , ν = 10.82 )
9.0
8.5
8.0
7.5
7.0
A1160p+ (I−12c)
7.0 7.5 8.0 8.5 9.0
AFUB_002050 (I−12c)
P( −0.716 < μ1 − μ2 < 0.063 ) = 0.95

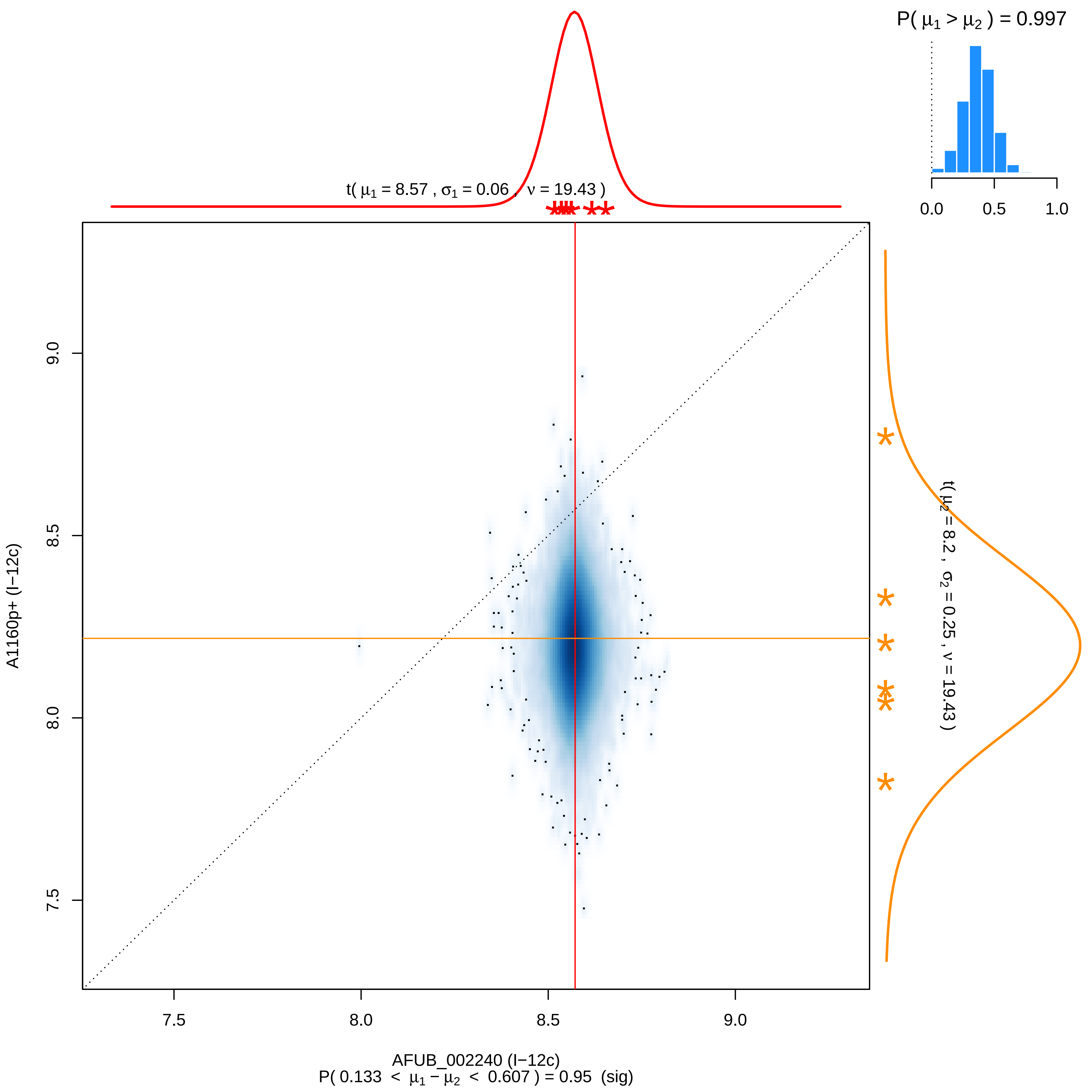

P( μ1 > μ2 ) = 0.997
0.0
0.5
1.0
t( μ1 = 8.57 , σ1 = 0.06 , ν = 19.43 )
t( μ2 = 8.2 , σ2 = 0.25 , ν = 19.43 )
A1160p+ (I−12c)
7.5
8.0
8.5
9.0
AFUB_002240 (I−12c)
P( 0.133 < μ1 − μ2 < 0.607 ) = 0.95 (sig)

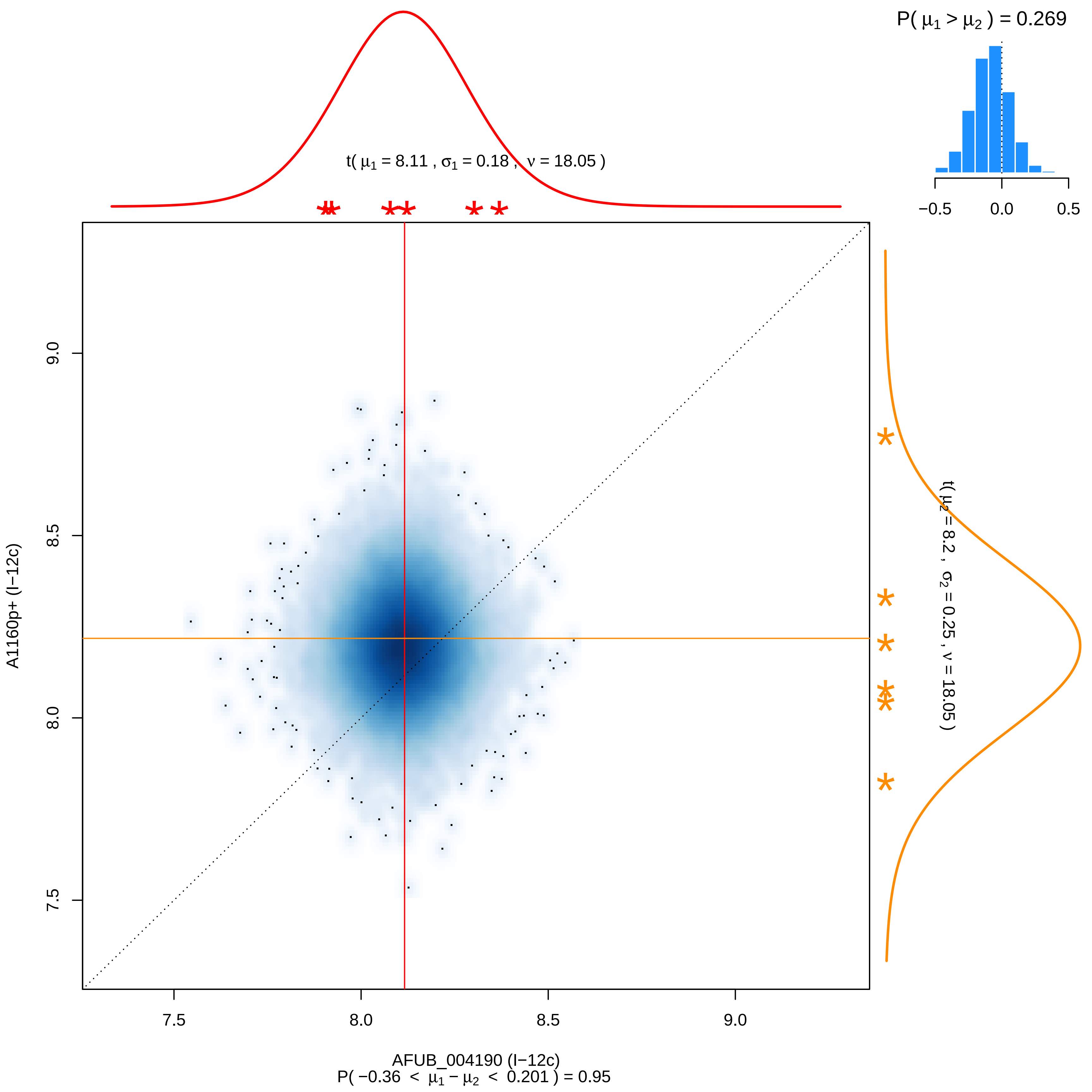

P( μ1 > μ2 ) = 0.269
−0.5
0.0
0.5
t( μ1 = 8.11 , σ1 = 0.18 , ν = 18.05 )
t( μ2 = 8.2 , σ2 = 0.25 , ν = 18.05 )
9.0
8.5
8.0
7.5
A1160p+ (I−12c)
7.5
8.0
8.5
9.0
AFUB_004190 (I−12c)
P( −0.36 < μ1 − μ2 < 0.201 ) = 0.95

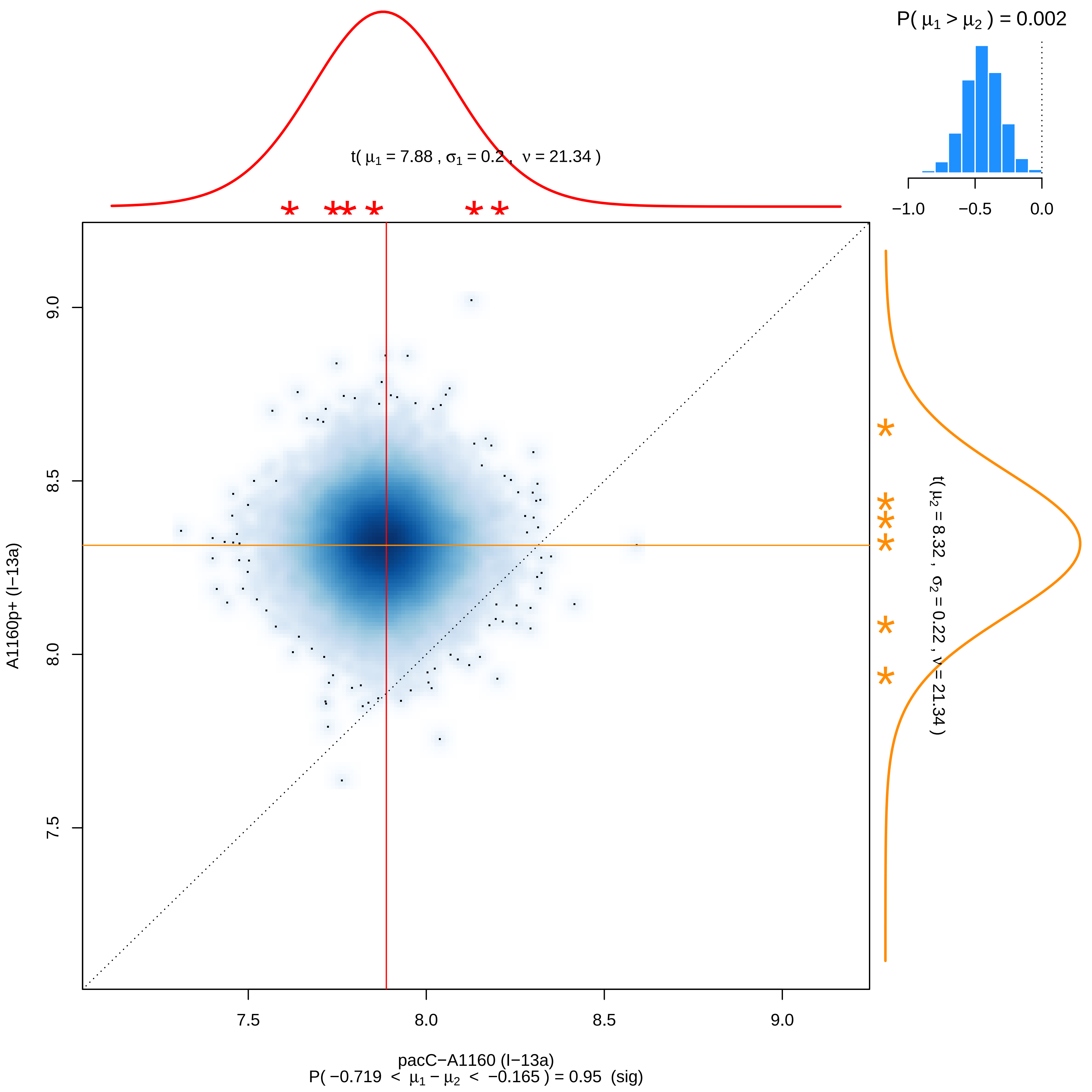

P( μ1 > μ2 ) = 0.002
−1.0
−0.5
0.0
t( μ1 = 7.88 , σ1 = 0.2 , ν = 21.34 )
t( μ2 = 8.32 , σ2 = 0.22 , ν = 21.34 )
A1160p+ (I−13a)
9.0
8.5
8.0
7.5
7.5
8.0
8.5
9.0
pacC−A1160 (I−13a)
P( −0.719 < μ1 − μ2 < −0.165 ) = 0.95 (sig)

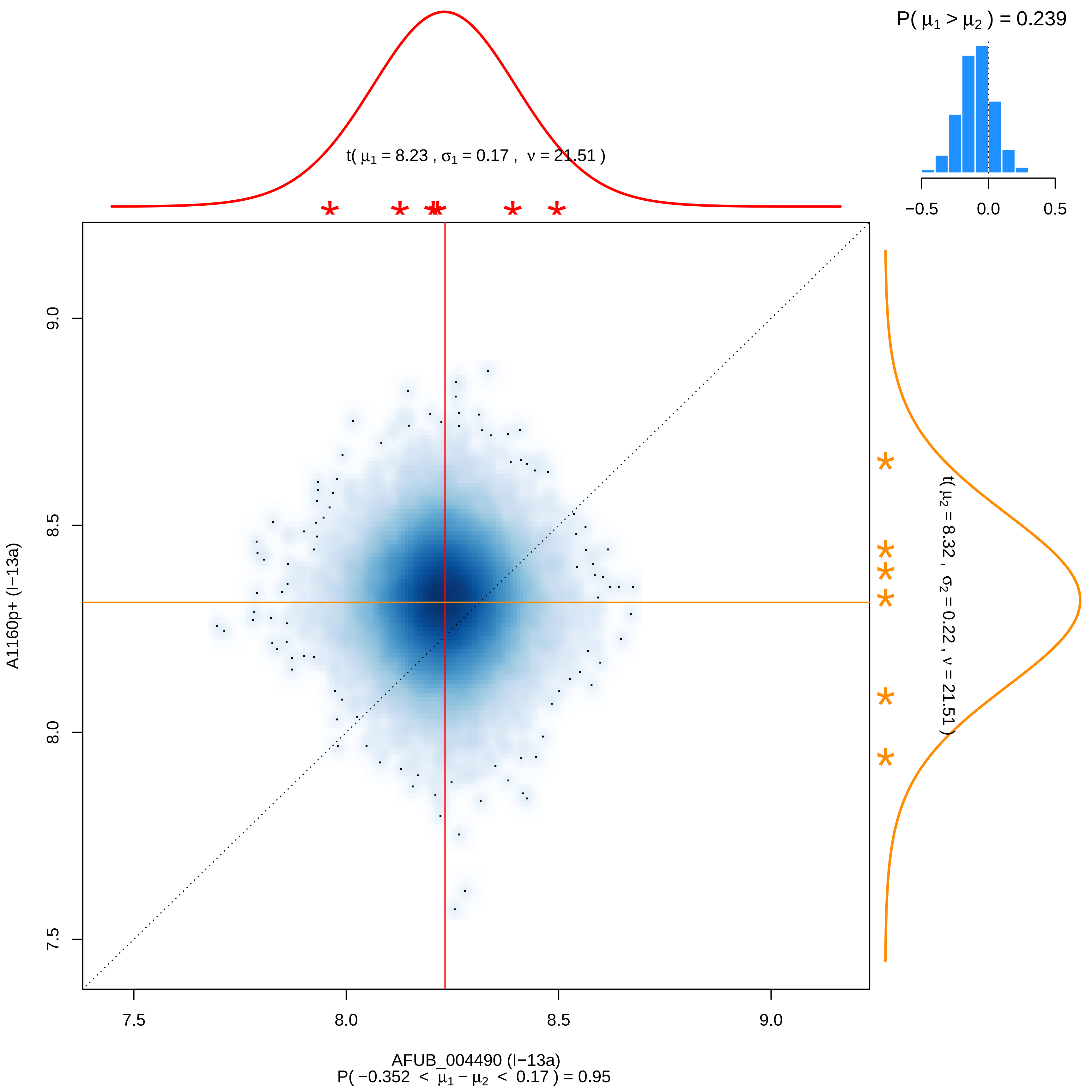

P( μ1 > μ2 ) = 0.239
−0.5
0.0
0.5
t( μ1 = 8.23 , σ1 = 0.17 , ν = 21.51 )
t( μ2 = 8.32 , σ2 = 0.22 , ν = 21.51 )
9.0
8.5
8.0
7.5
A1160p+ (I−13a)
7.5
8.0
8.5
9.0
AFUB_004490 (I−13a)
P( −0.352 < μ1 − μ2 < 0.17 ) = 0.95

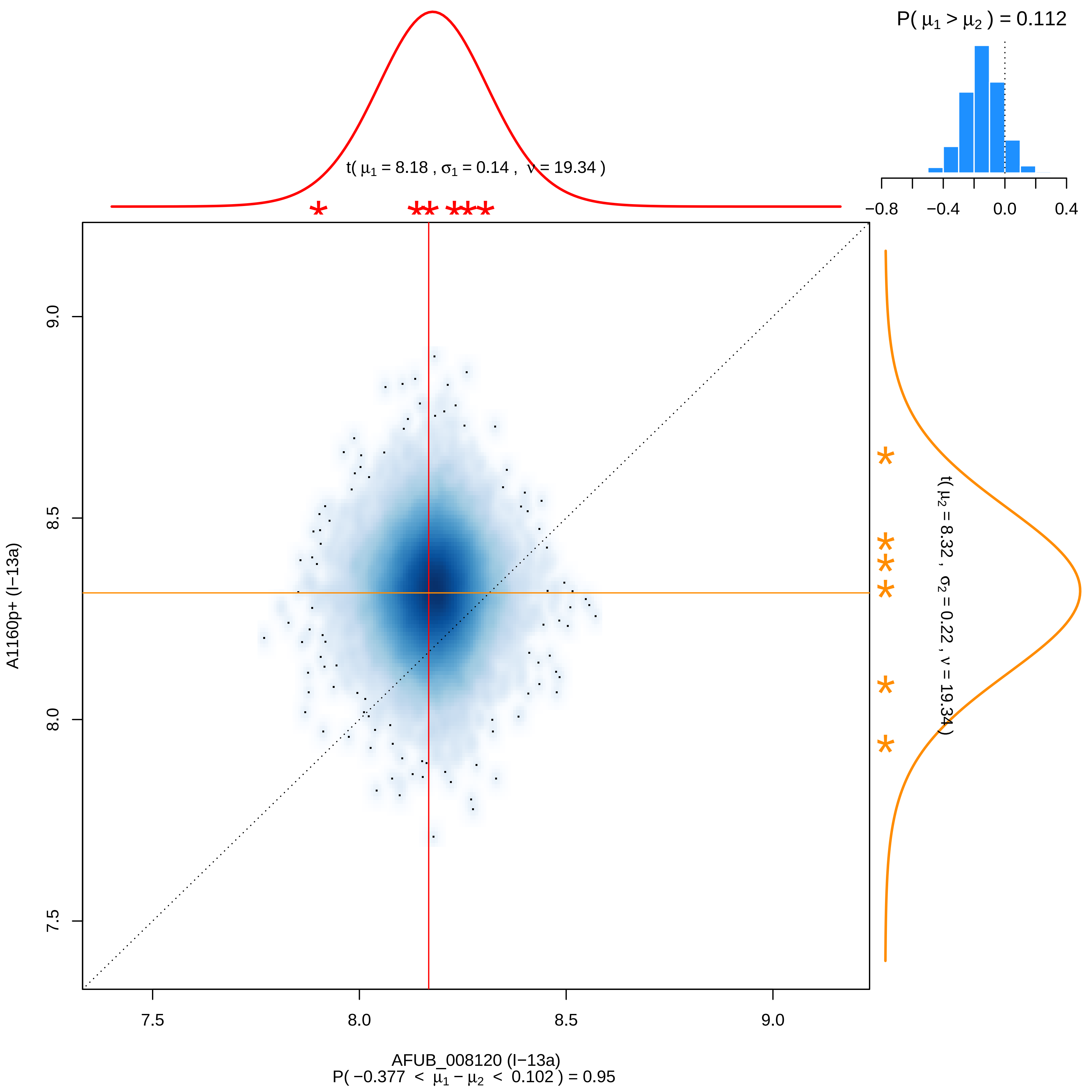
P( μ1 > μ2 ) = 0.112
−0.8
−0.4
0.0
0.4
t( μ1 = 8.18 , σ1 = 0.14 , ν = 19.34 )
t( μ2 = 8.32 , σ2 = 0.22 , ν = 19.34 )
9.0
8.5
8.0
7.5
A1160p+ (I−13a)
7.5
8.0
8.5
9.0
AFUB_008120 (I−13a)
P( −0.377 < μ1 − μ2 < 0.102 ) = 0.95

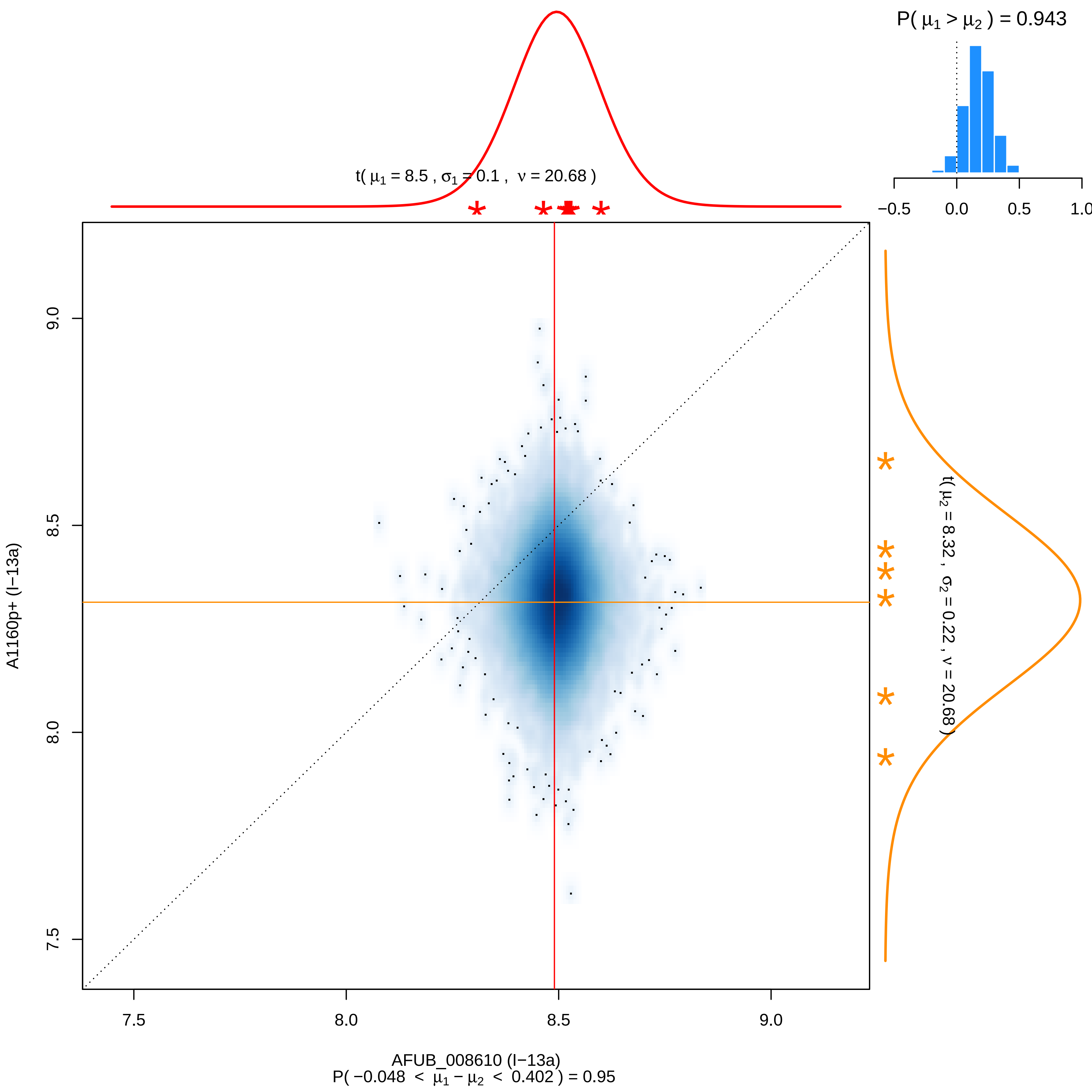

P( μ1 > μ2 ) = 0.943
−0.5
0.0
0.5
1.0
t( μ1 = 8.5 , σ1 = 0.1 , ν = 20.68 )
t( μ2 = 8.32 , σ2 = 0.22 , ν = 20.68 )
9.0
8.5
8.0
7.5
A1160p+ (I−13a)
7.5
8.0
8.5
9.0
AFUB_008610 (I−13a)
P( −0.048 < μ1 − μ2 < 0.402 ) = 0.95

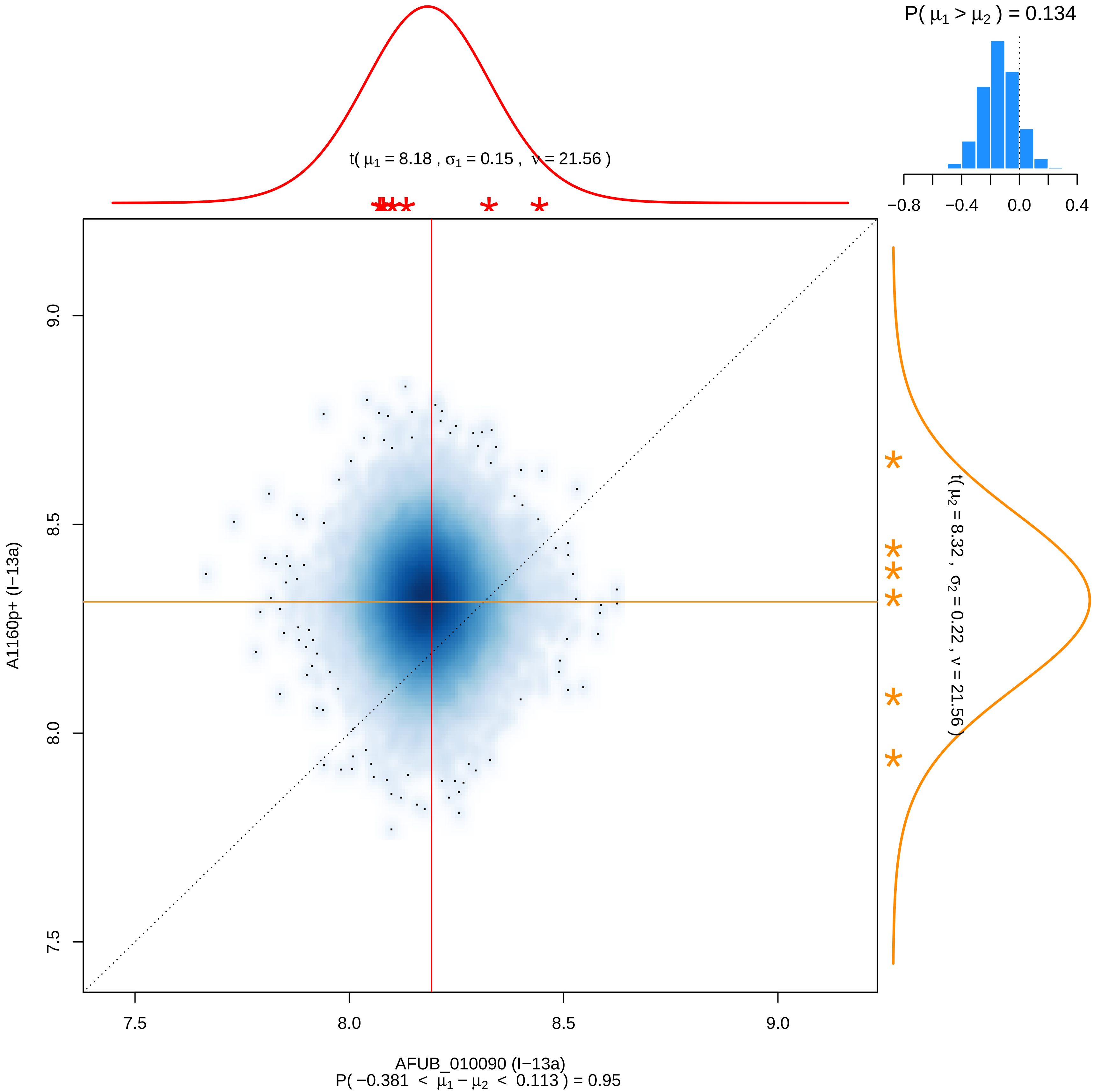

P( μ1 > μ2 ) = 0.134
−0.8
−0.4
0.0
0.4
t( μ1 = 8.18 , σ1 = 0.15 , ν = 21.56 )
t( μ2 = 8.32 , σ2 = 0.22 , ν = 21.56 )
A1160p+ (I−13a)
9.0
8.5
8.0
7.5
7.5
8.0
8.5
9.0
AFUB_010090 (I−13a)
P( −0.381 < μ1 − μ2 < 0.113 ) = 0.95

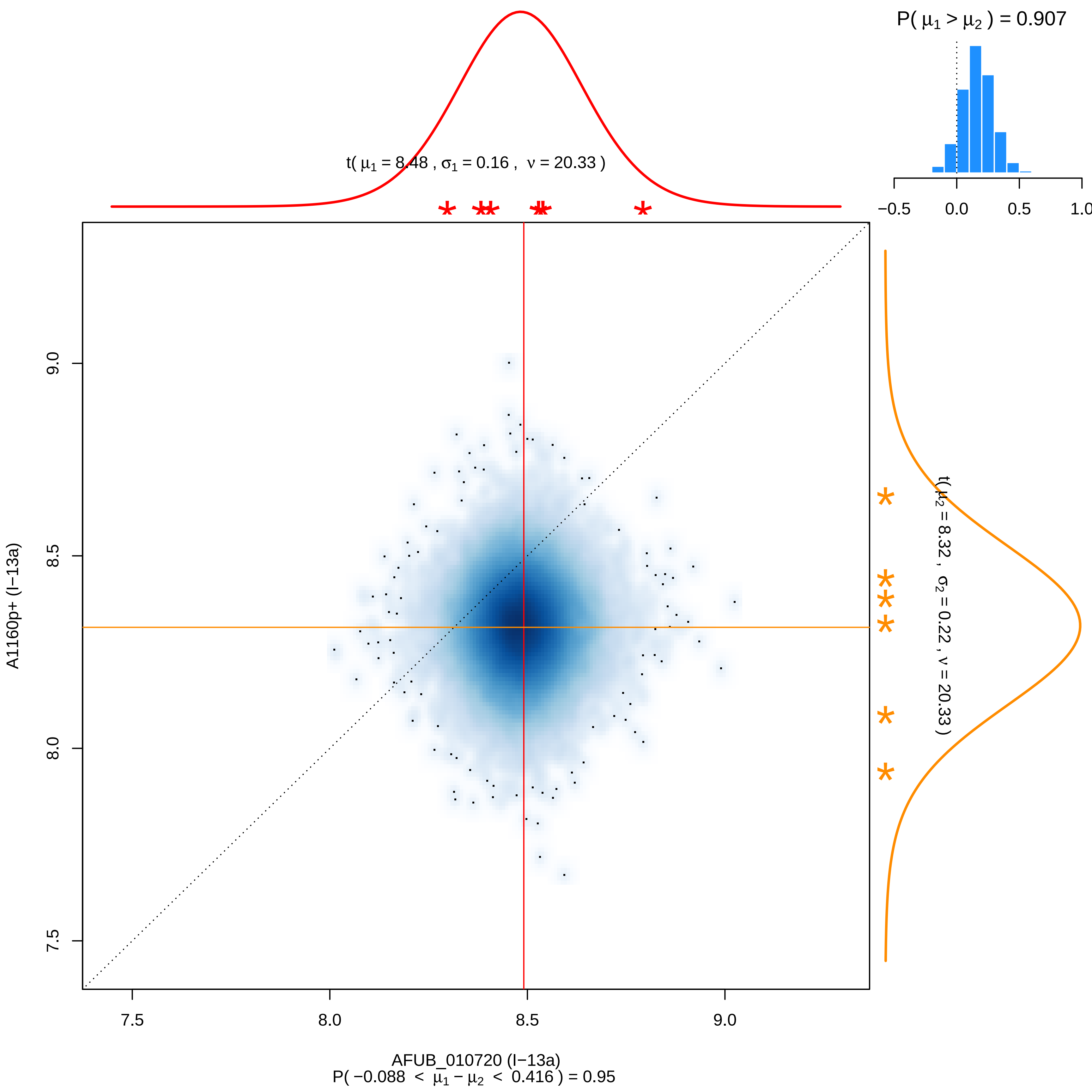

P( μ1 > μ2 ) = 0.907
−0.5
0.0
0.5
1.0
t( μ1 = 8.48 , σ1 = 0.16 , ν = 20.33 )
t( μ2 = 8.32 , σ2 = 0.22 , ν = 20.33 )
A1160p+ (I−13a)
9.0
8.5
8.0
7.5
7.5
8.0
8.5
9.0
AFUB_010720 (I−13a)
P( −0.088 < μ1 − μ2 < 0.416 ) = 0.95

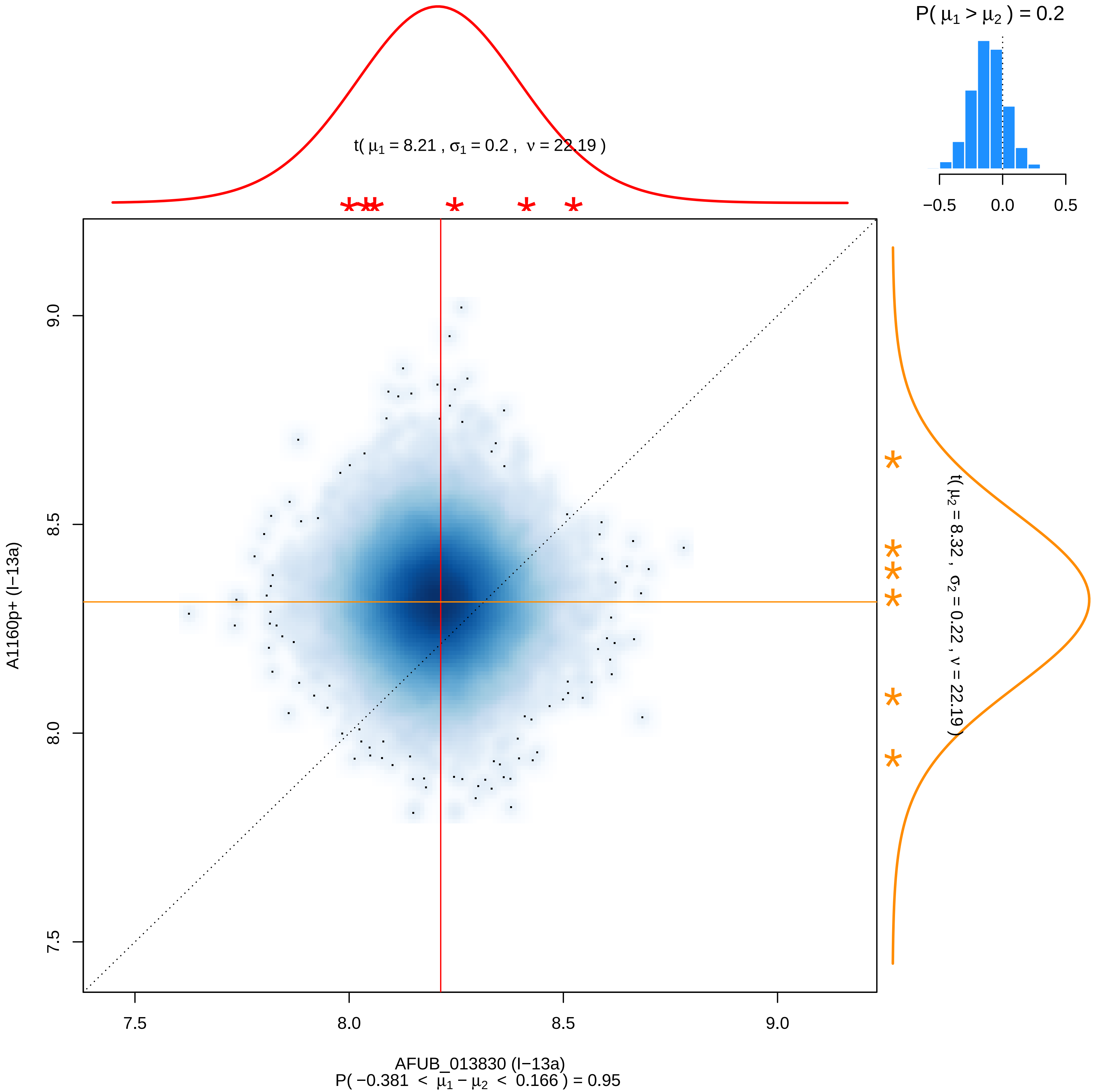

P( μ1 > μ2 ) = 0.2
−0.5
0.0
0.5
t( μ1 = 8.21 , σ1 = 0.2 , ν = 22.19 )
t( μ2 = 8.32 , σ2 = 0.22 , ν = 22.19 )
9.0
8.5
8.0
7.5
A1160p+ (I−13a)
7.5
8.0
8.5
9.0
AFUB_013830 (I−13a)
P( −0.381 < μ1 − μ2 < 0.166 ) = 0.95

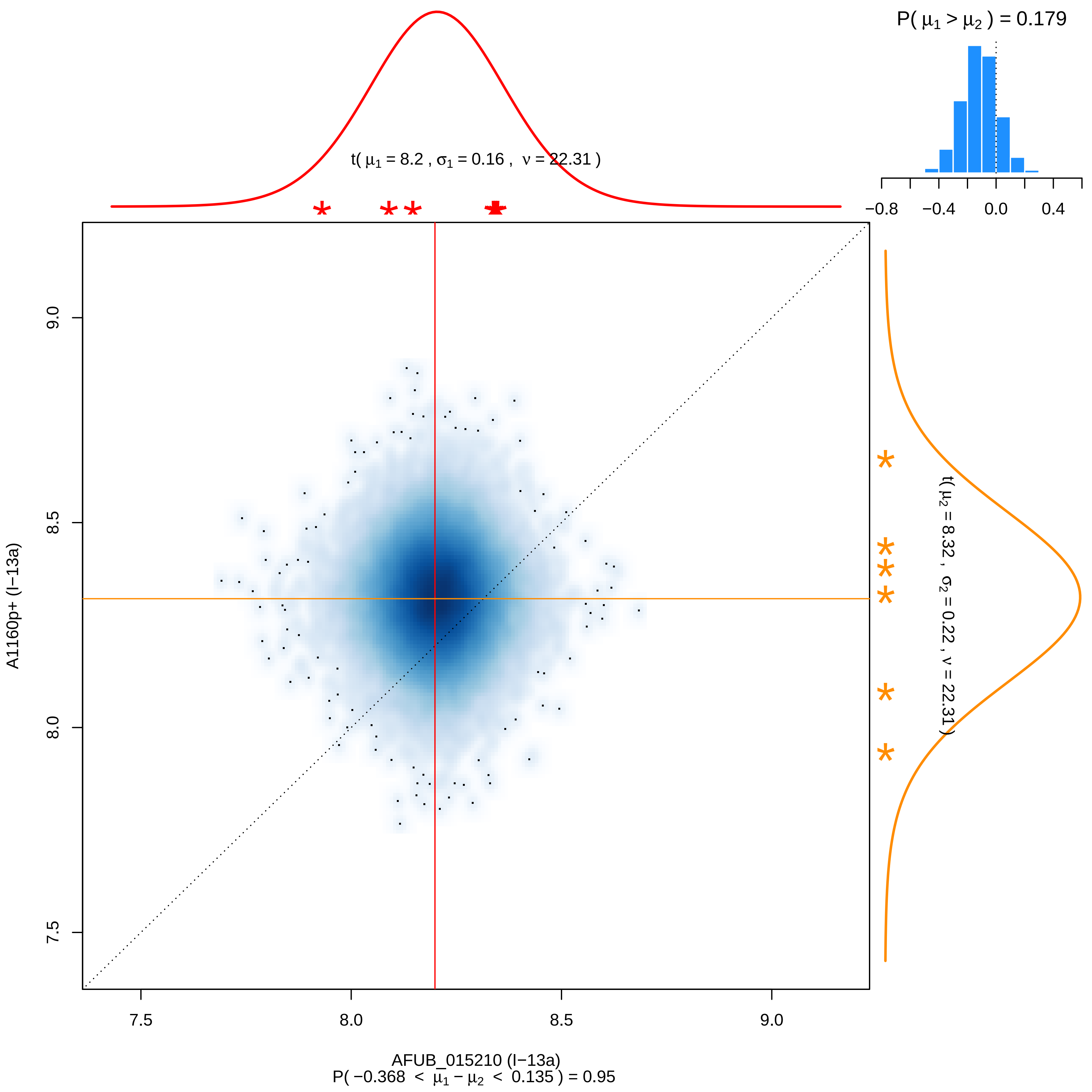

P( μ1 > μ2 ) = 0.179
−0.8
−0.4
0.0
0.4
t( μ1 = 8.2 , σ1 = 0.16 , ν = 22.31 )
t( μ2 = 8.32 , σ2 = 0.22 , ν = 22.31 )
A1160p+ (I−13a)
9.0
8.5
8.0
7.5
7.5
8.0
8.5
9.0
AFUB_015210 (I−13a)
P( −0.368 < μ1 − μ2 < 0.135 ) = 0.95

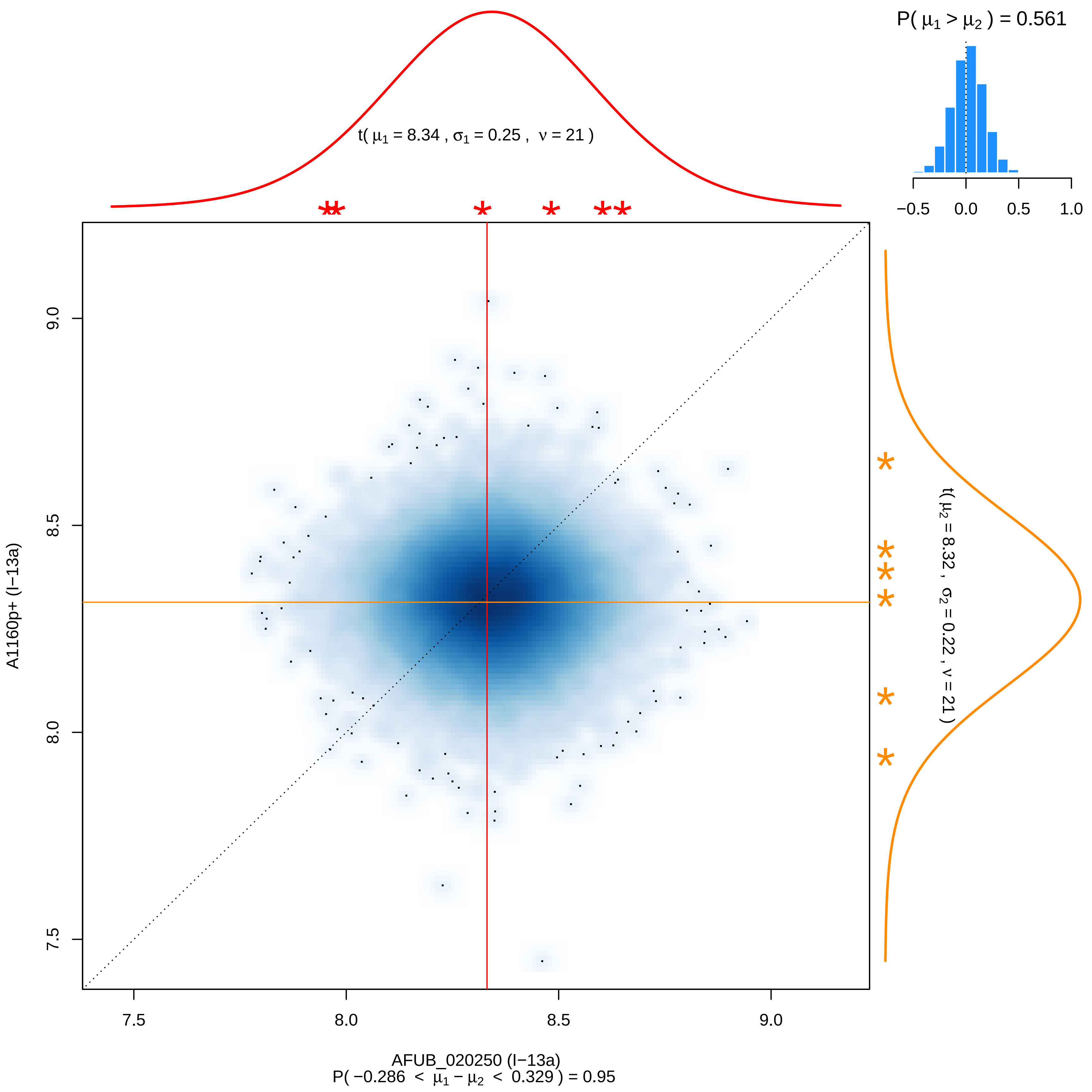

P( μ1 > μ2 ) = 0.561
−0.5 0.0 0.5 1.0
t( μ1 = 8.34 , σ1 = 0.25 , ν = 21 )
t( μ2 = 8.32 , σ2 = 0.22 , ν = 21 )
9.0
8.5
8.0
7.5
A1160p+ (I−13a)
7.5 8.0 8.5 9.0
AFUB_020250 (I−13a)
P( −0.286 < μ1 − μ2 < 0.329 ) = 0.95

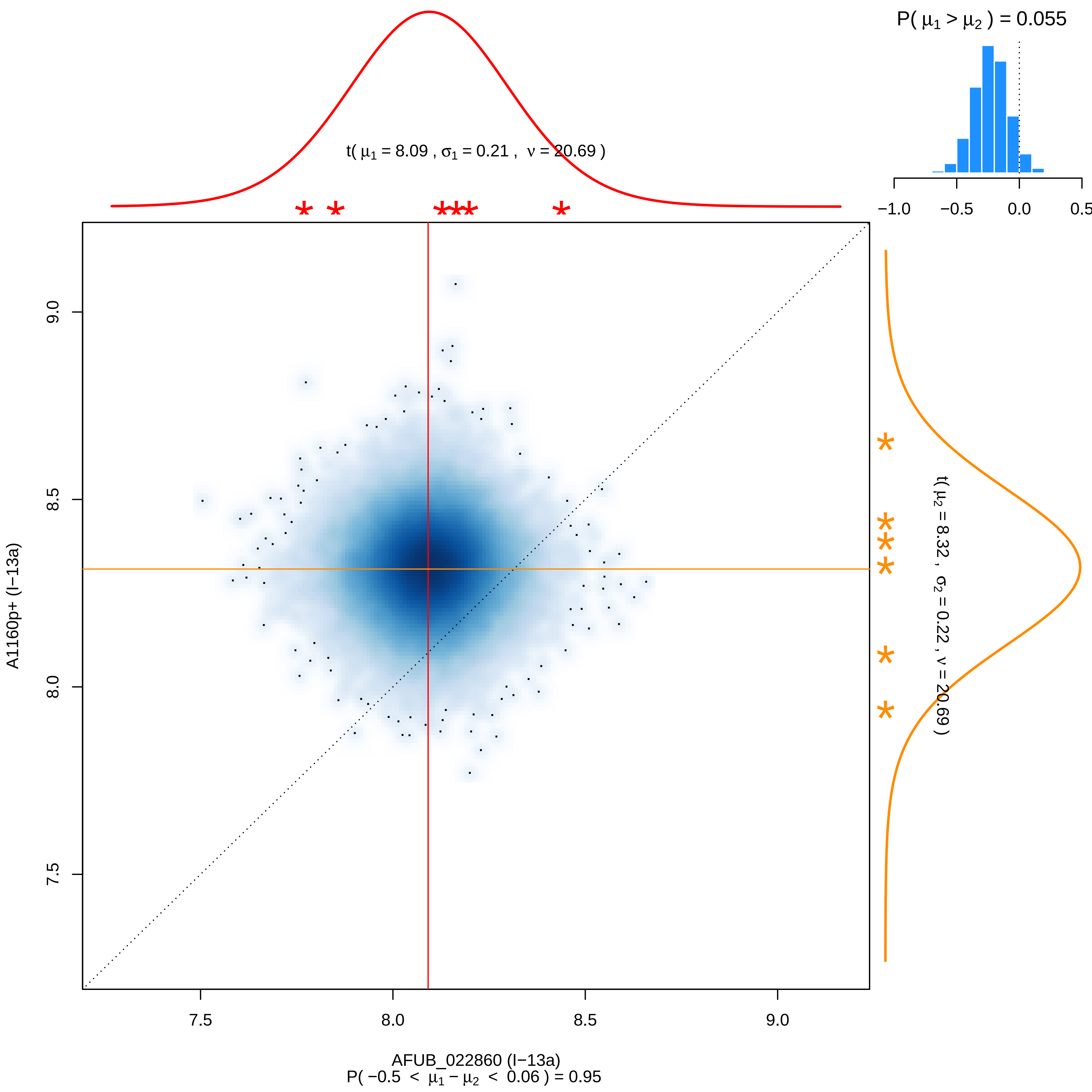

P( μ1 > μ2 ) = 0.055
−1.0 −0.5 0.0 0.5
t( μ1 = 8.09 , σ1 = 0.21 , ν = 20.69 )
t( μ2 = 8.32 , σ2 = 0.22 , ν = 20.69 )
A1160p+ (I−13a)
9.0
8.5
8.0
7.5
7.5 8.0 8.5 9.0
AFUB_022860 (I−13a)
P( −0.5 < μ1 − μ2 < 0.06 ) = 0.95

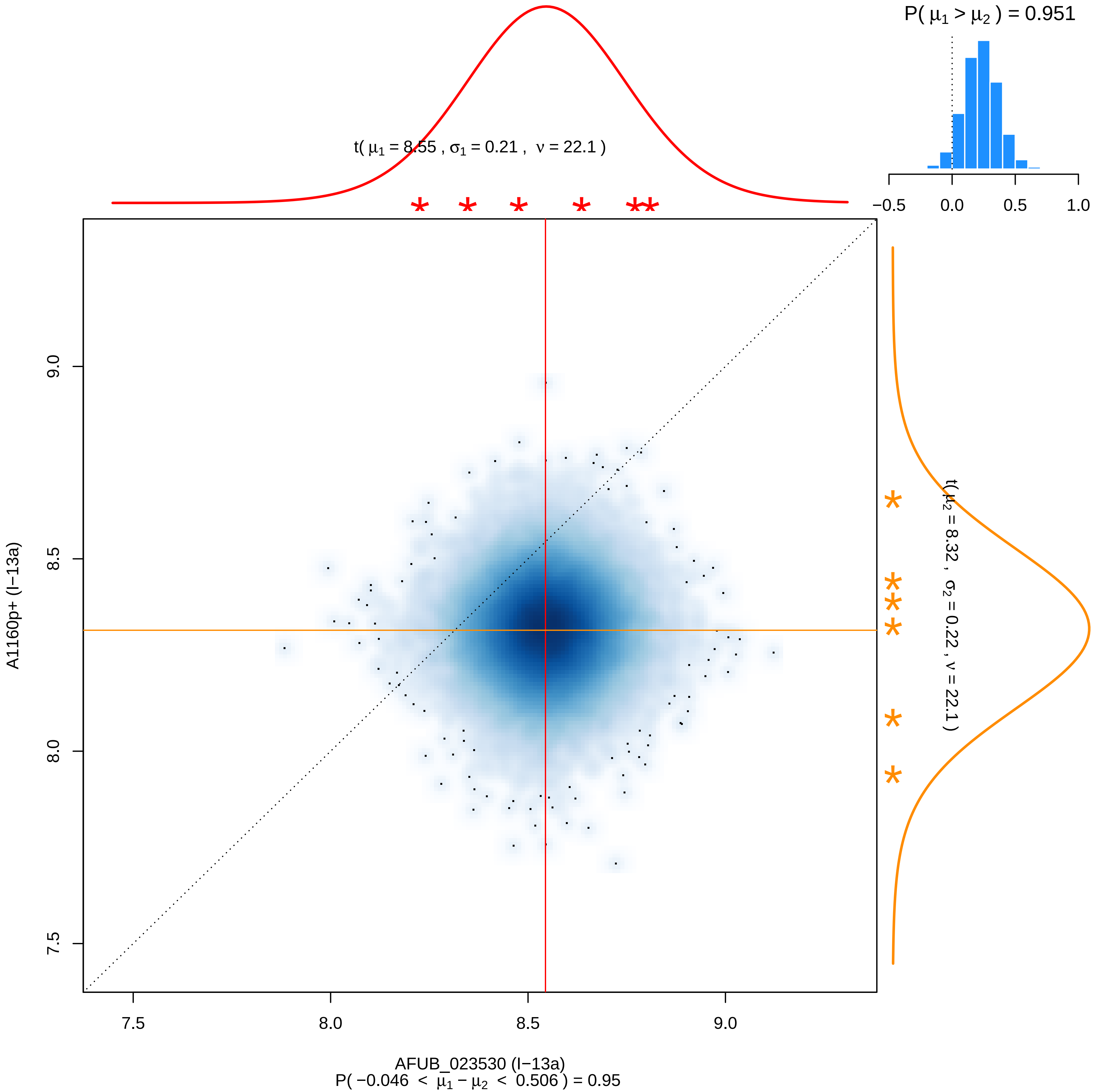

P( $\mu_1 > \mu_2$ ) = 0.951
−0.5
0.0
0.5
1.0
t( $\mu_1$ = 8.55 , $\sigma_1$ = 0.21 , $\nu$ = 22.1 )
t( $\mu_2$ = 8.32 , $\sigma_2$ = 0.22 , $\nu$ = 22.1 )
9.0
8.5
8.0
7.5
A1160p+ (I−13a)
7.5
8.0
8.5
9.0
AFUB_023530 (I−13a)
P( −0.046 < $\mu_1 - \mu_2$ < 0.506 ) = 0.95

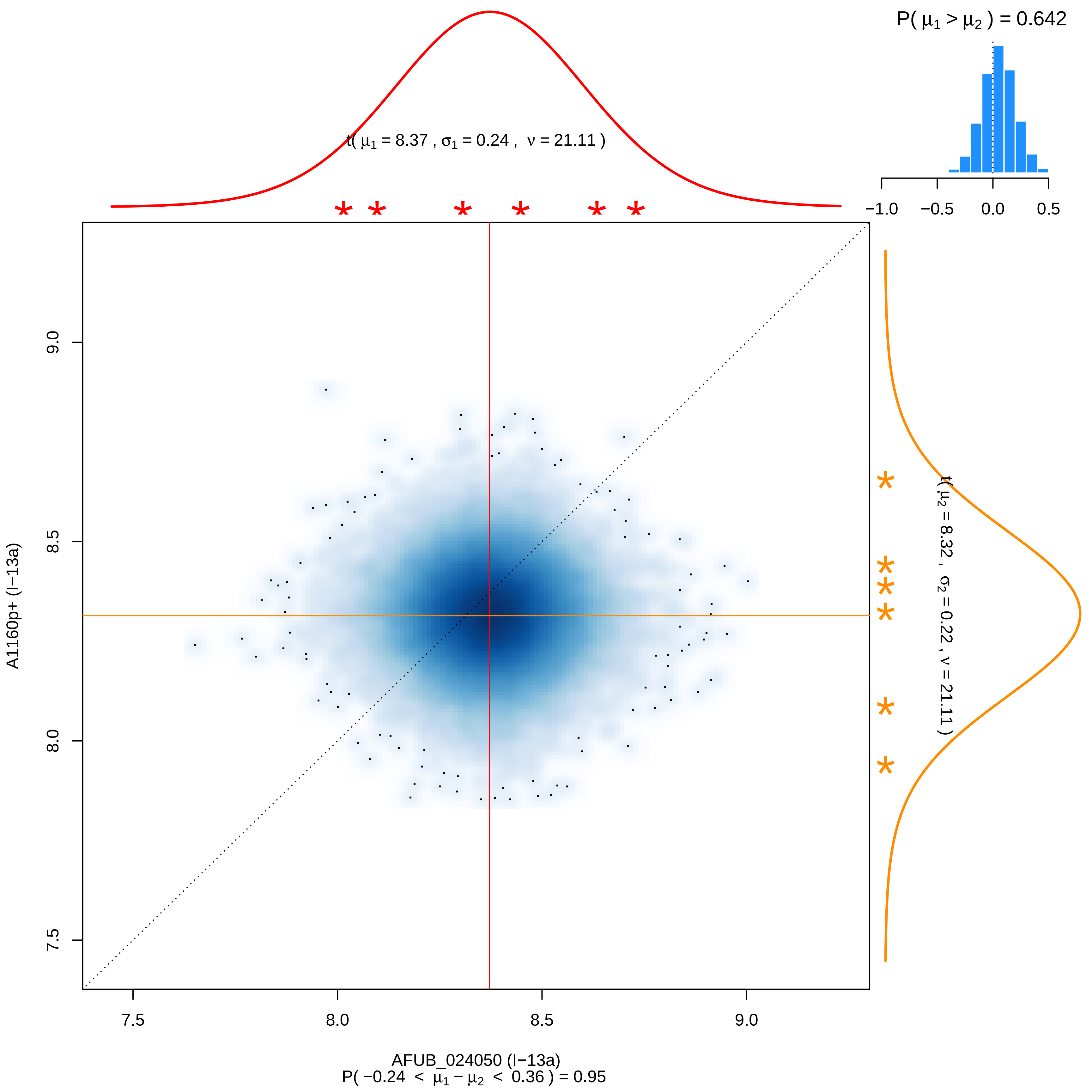

P( μ1 > μ2 ) = 0.642
−1.0
−0.5
0.0
0.5
t( μ1 = 8.37 , σ1 = 0.24 , ν = 21.11 )
t( μ2 = 8.32 , σ2 = 0.22 , ν = 21.11 )
9.0
8.5
8.0
7.5
A1160p+ (I−13a)
7.5
8.0
8.5
9.0
AFUB_024050 (I−13a)
P( −0.24 < μ1 − μ2 < 0.36 ) = 0.95

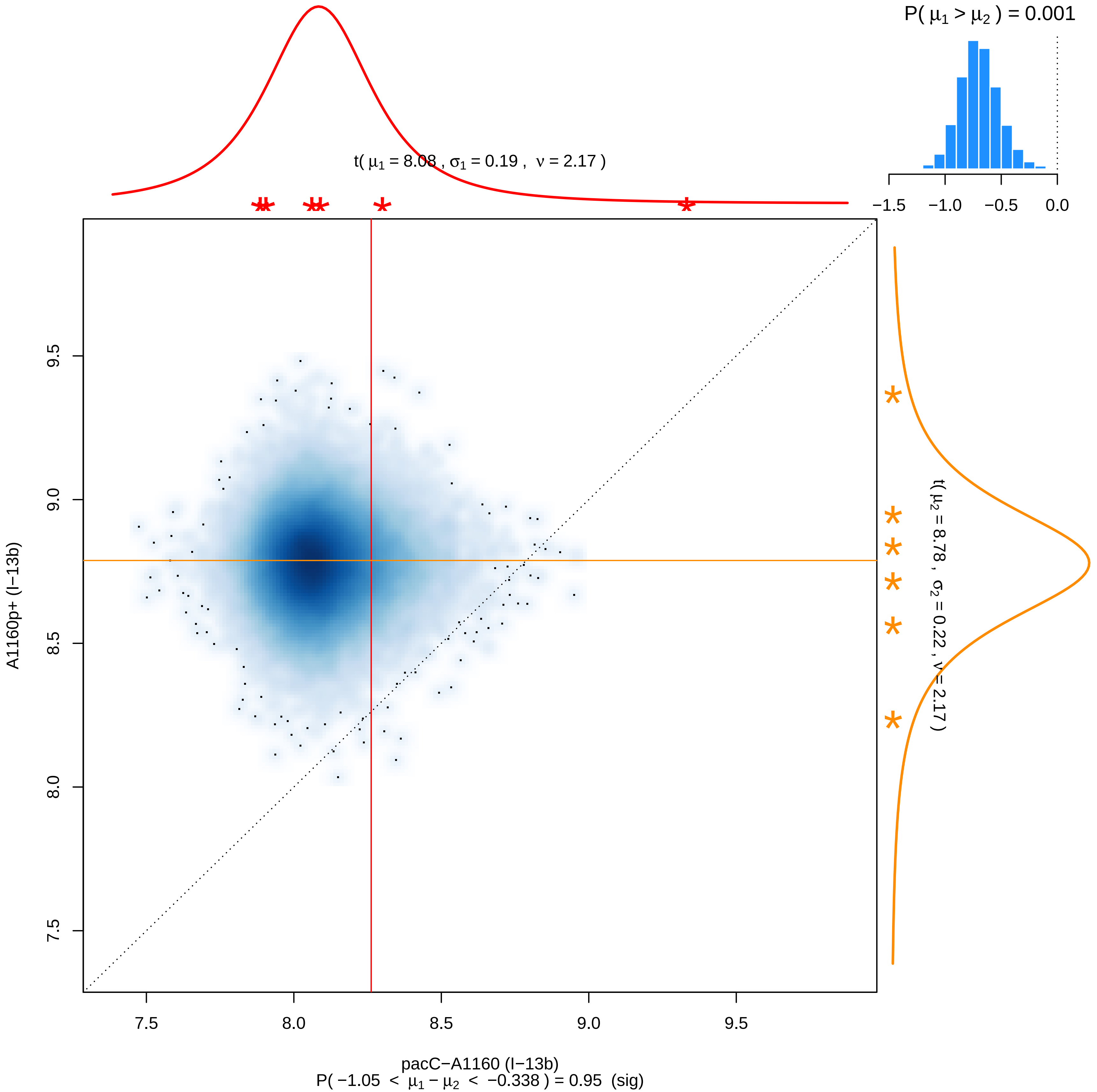

P( μ1 > μ2 ) = 0.001
−1.5 −1.0 −0.5 0.0
t( μ1 = 8.08 , σ1 = 0.19 , ν = 2.17 )
t( μ2 = 8.78 , σ2 = 0.22 , ν = 2.17 )
A1160p+ (I−13b)
9.5
9.0
8.5
8.0
7.5
7.5 8.0 8.5 9.0 9.5
pacC−A1160 (I−13b)
P( −1.05 < μ1 − μ2 < −0.338 ) = 0.95 (sig)

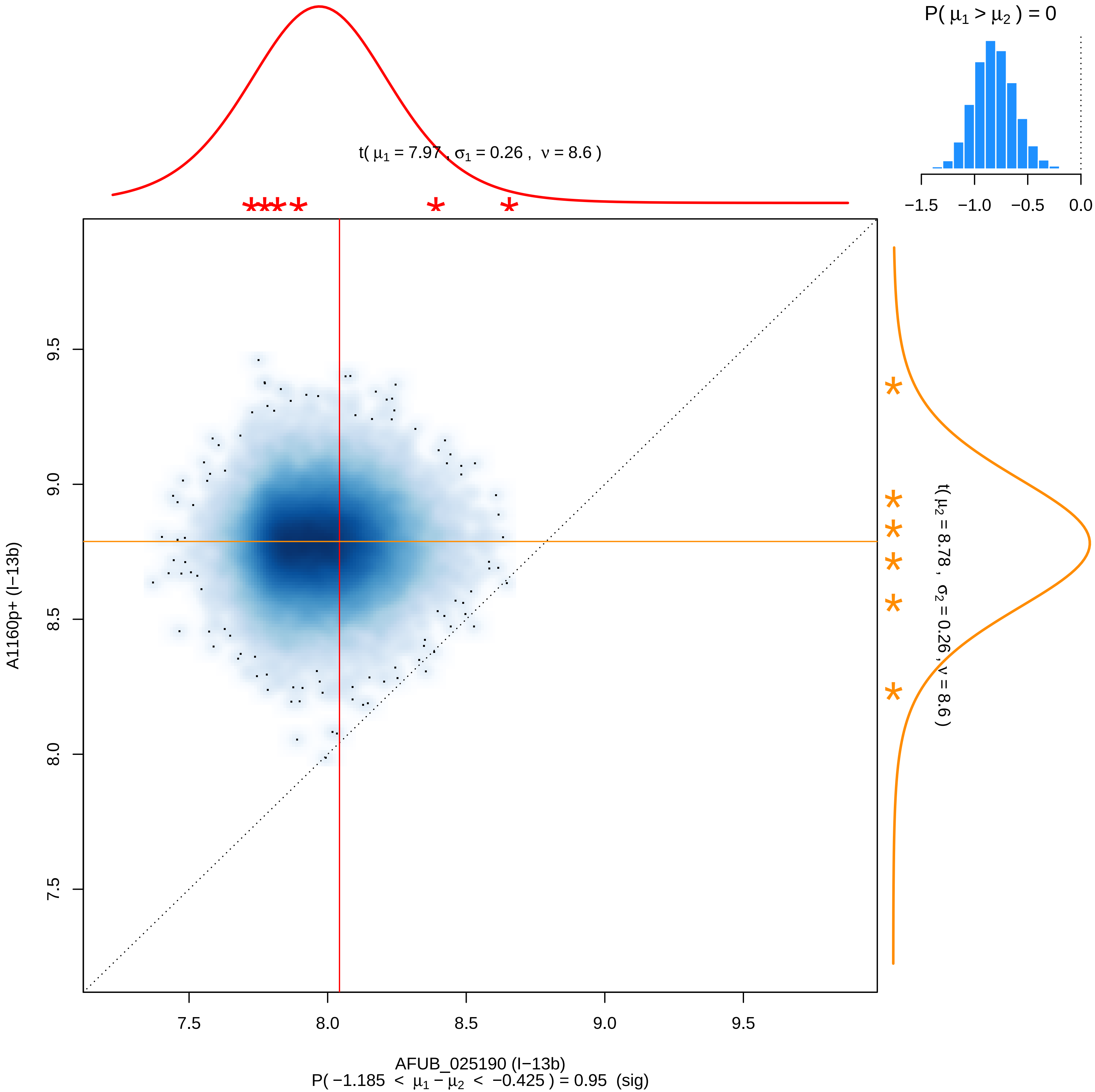

P( μ1 > μ2 ) = 0
−1.5 −1.0 −0.5 0.0
t( μ1 = 7.97 , σ1 = 0.26 , ν = 8.6 )
t( μ2 = 8.78 , σ2 = 0.26 , ν = 8.6 )
9.5
9.0
8.5
8.0
7.5
A1160p+ (I−13b)
7.5
8.0
8.5
9.0
9.5
AFUB_025190 (I−13b)
P( −1.185 < μ1 − μ2 < −0.425 ) = 0.95 (sig)

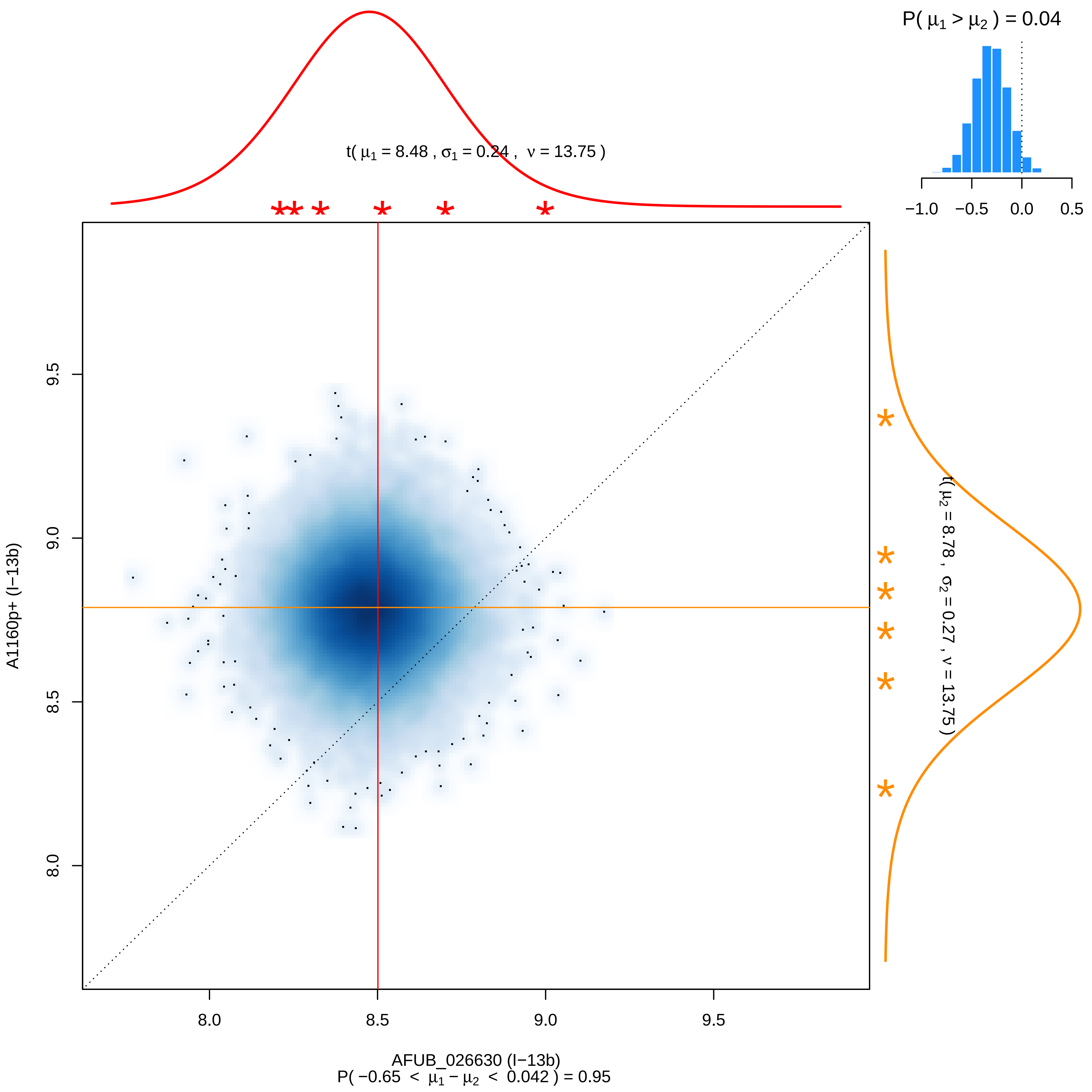

P( μ1 > μ2 ) = 0.04
−1.0 −0.5 0.0 0.5
t( μ1 = 8.48 , σ1 = 0.24 , ν = 13.75 )
t( μ2 = 8.78 , σ2 = 0.27 , ν = 13.75 )
9.5
9.0
8.5
8.0
A1160p+ (I−13b)
8.0 8.5 9.0 9.5
AFUB_026630 (I−13b)
P( −0.65 < μ1 − μ2 < 0.042 ) = 0.95

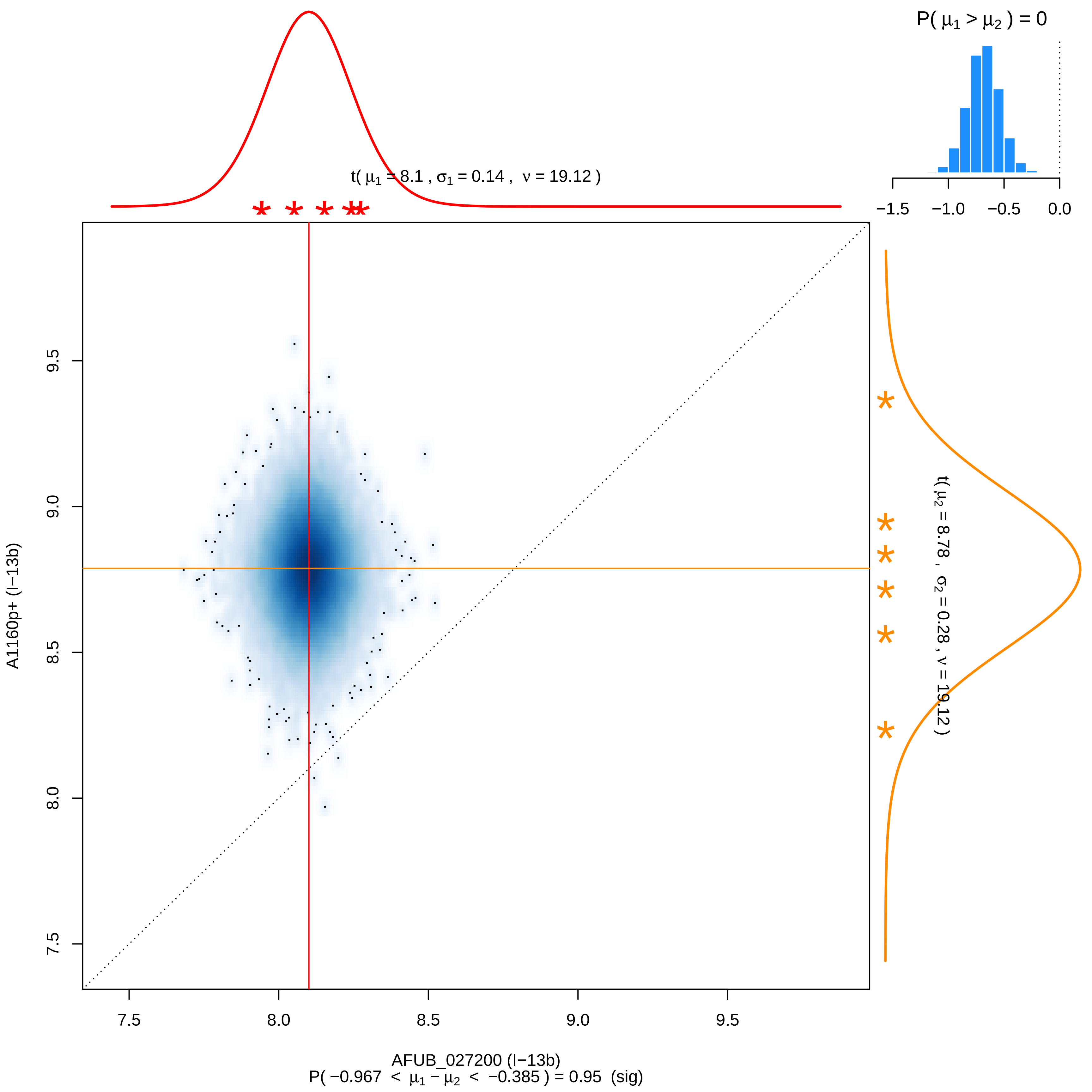

P( μ1 > μ2 ) = 0
−1.5 −1.0 −0.5 0.0
t( μ1 = 8.1 , σ1 = 0.14 , ν = 19.12 )
t( μ2 = 8.78 , σ2 = 0.28 , ν = 19.12 )
A1160p+ (I−13b)
7.5 8.0 8.5 9.0 9.5
AFUB_027200 (I−13b)
P( −0.967 < μ1 − μ2 < −0.385 ) = 0.95 (sig)

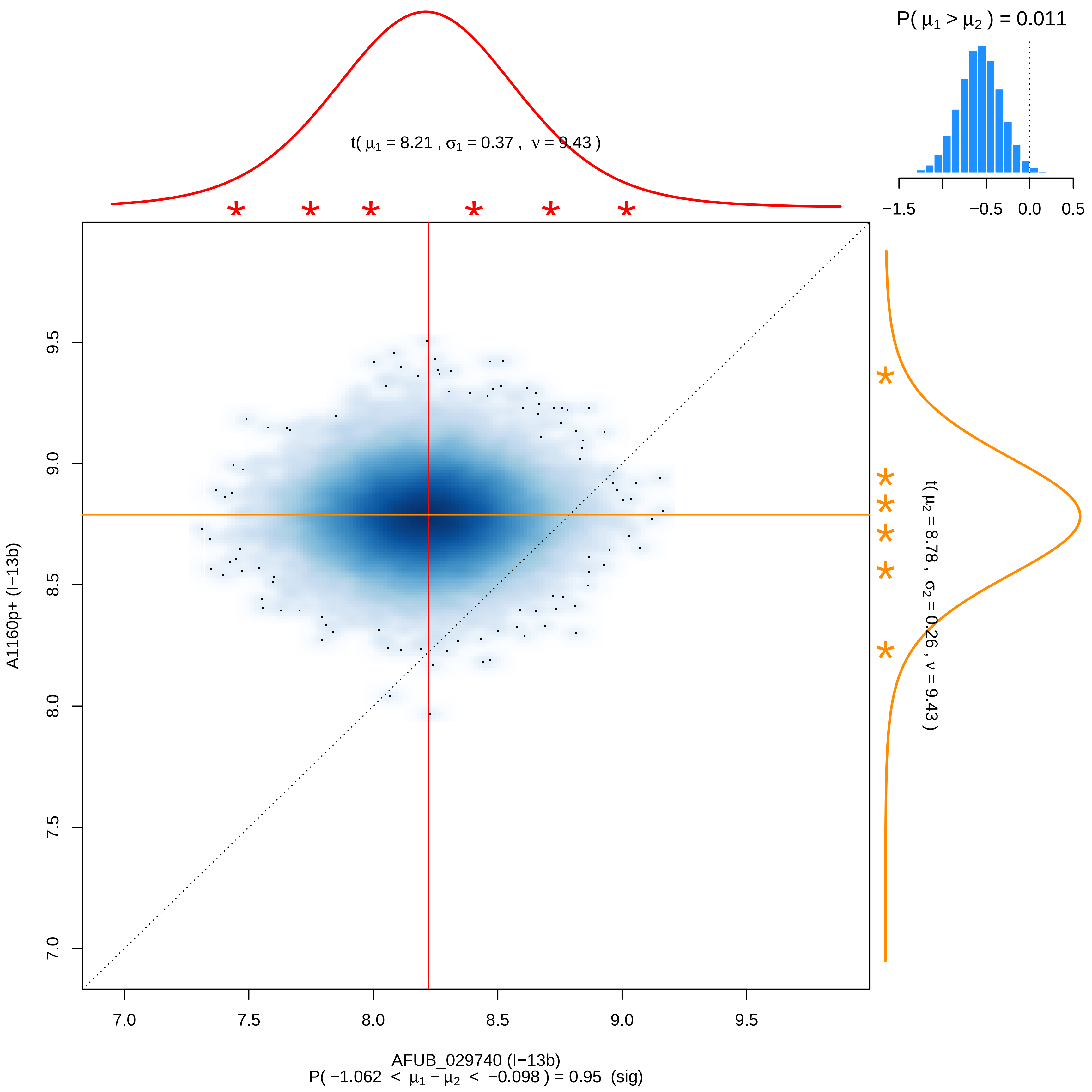

P( μ1 > μ2 ) = 0.011
−1.5
−0.5
0.0
0.5
t( μ1 = 8.21 , σ1 = 0.37 , ν = 9.43 )
t( μ2 = 8.78 , σ2 = 0.26 , ν = 9.43 )
A1160p+ (I−13b)
9.5
9.0
8.5
8.0
7.5
7.0
7.0
7.5
8.0
8.5
9.0
9.5
AFUB_029740 (I−13b)
P( −1.062 < μ1 − μ2 < −0.098 ) = 0.95 (sig)

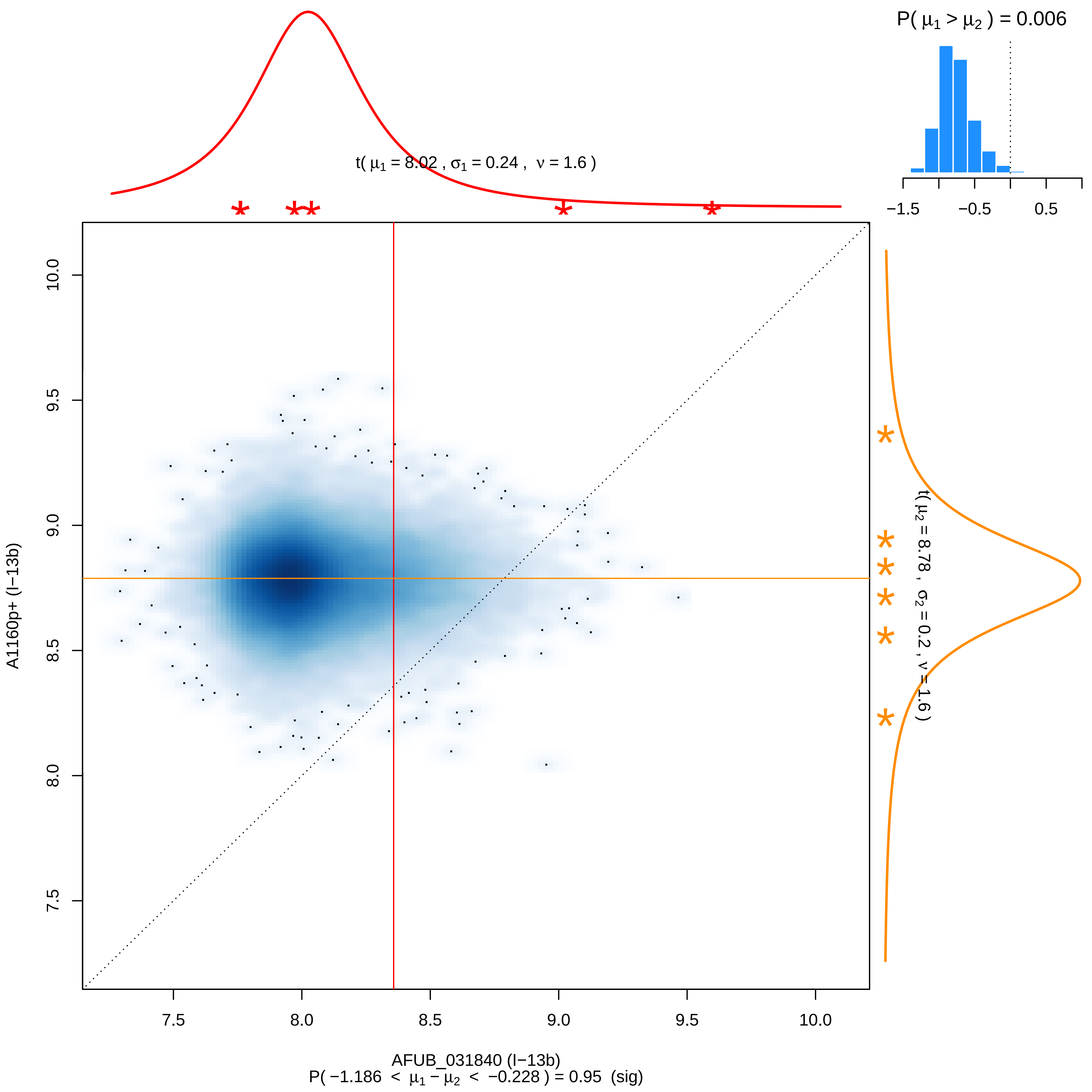

P( μ1 > μ2 ) = 0.006
−1.5
−0.5
0.5
t( μ1 = 8.02 , σ1 = 0.24 , ν = 1.6 )
t( μ2 = 8.78 , σ2 = 0.2 , ν = 1.6 )
A1160p+ (I−13b)
10.0
9.5
9.0
8.5
8.0
7.5
7.5
8.0
8.5
9.0
9.5
10.0
AFUB_031840 (I−13b)
P( −1.186 < μ1 − μ2 < −0.228 ) = 0.95 (sig)

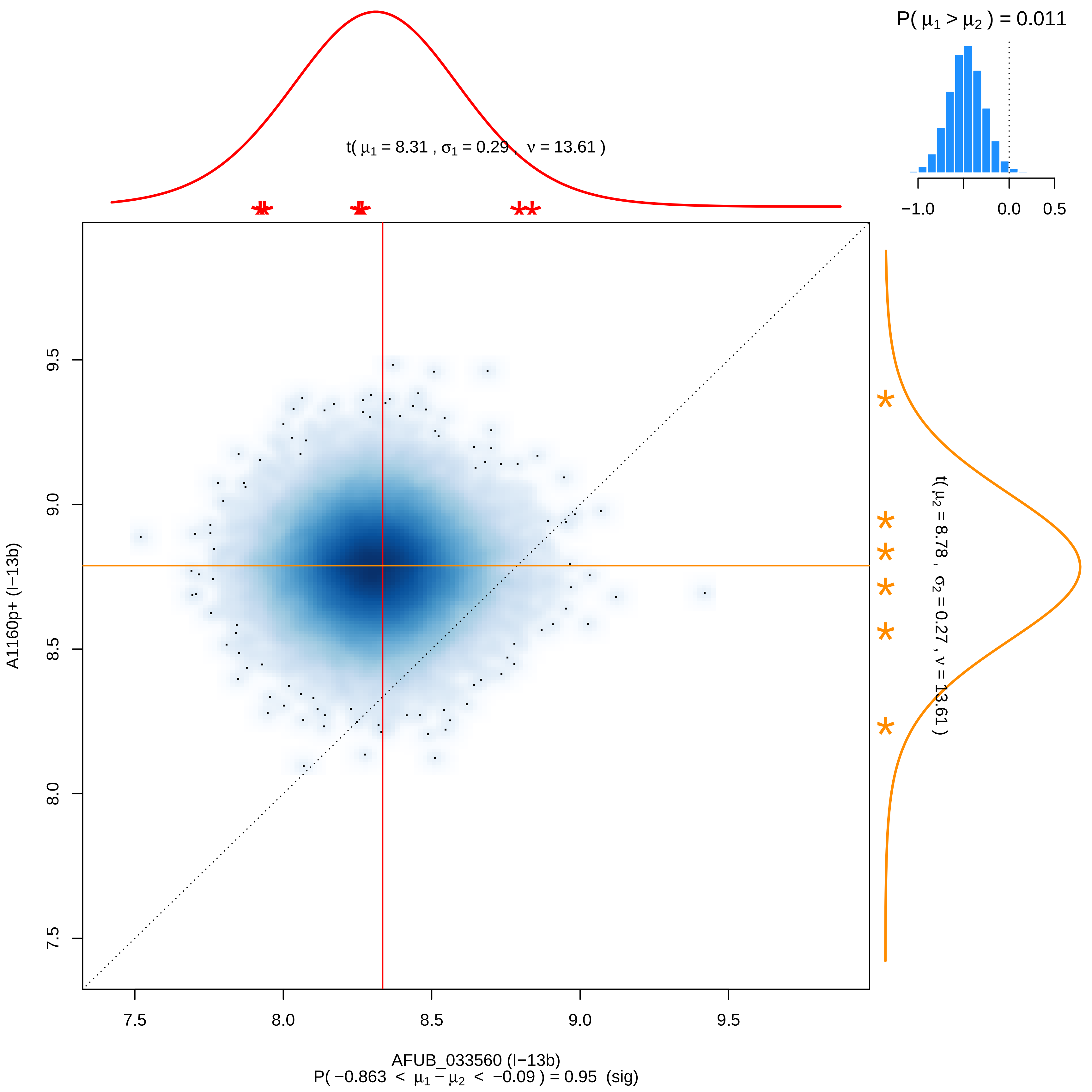

P( $\mu_1$ > $\mu_2$ ) = 0.011
−1.0
0.0
0.5
t( $\mu_1$ = 8.31 , $\sigma_1$ = 0.29 , $\nu$ = 13.61 )
t( $\mu_2$ = 8.78 , $\sigma_2$ = 0.27 , $\nu$ = 13.61 )
A1160p+ (I−13b)
9.5
9.0
8.5
8.0
7.5
7.5
8.0
8.5
9.0
9.5
AFUB_033560 (I−13b)
P( −0.863 < $\mu_1 - \mu_2$ < −0.09 ) = 0.95 (sig)

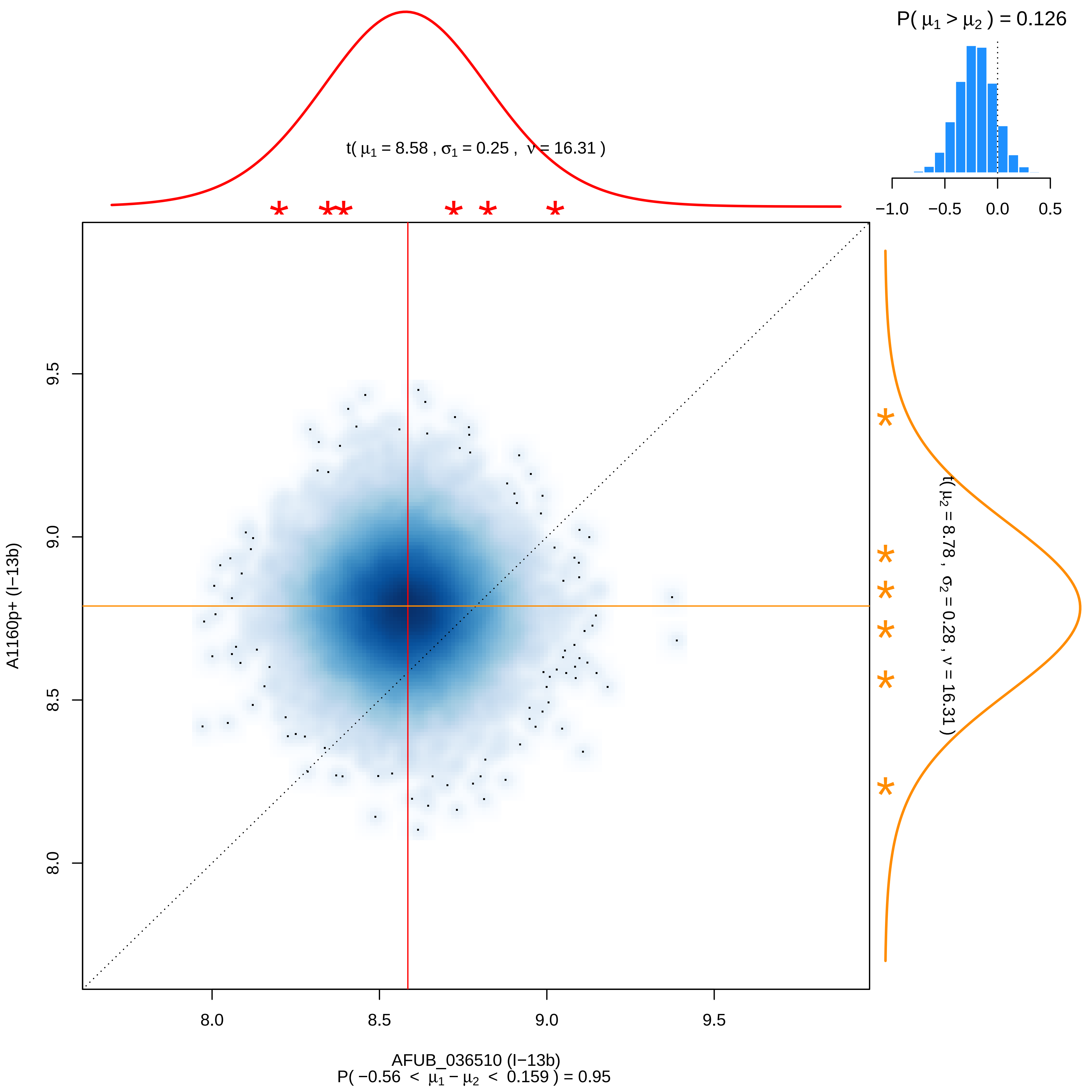

P( μ1 > μ2 ) = 0.126
−1.0
−0.5
0.0
0.5
t( μ1 = 8.58 , σ1 = 0.25 , ν = 16.31 )
t( μ2 = 8.78 , σ2 = 0.28 , ν = 16.31 )
A1160p+ (I−13b)
9.5
9.0
8.5
8.0
AFUB_036510 (I−13b)
P( −0.56 < μ1 − μ2 < 0.159 ) = 0.95

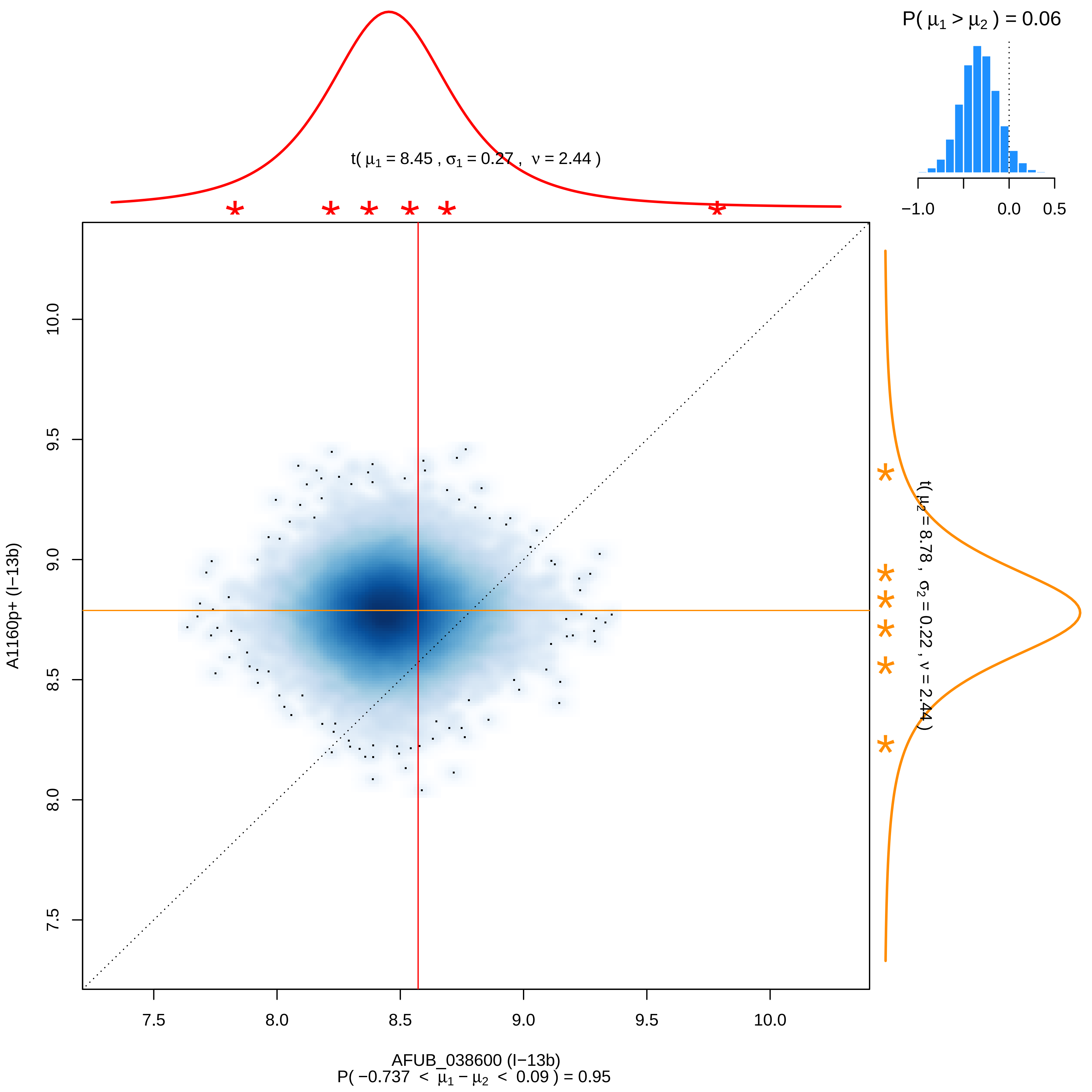

P( μ1 > μ2 ) = 0.06
−1.0
0.0
0.5
t( μ1 = 8.45 , σ1 = 0.27 , ν = 2.44 )
t( μ2 = 8.78 , σ2 = 0.22 , ν = 2.44 )
A1160p+ (I−13b)
7.5
8.0
8.5
9.0
9.5
10.0
AFUB_038600 (I−13b)
P( −0.737 < μ1 − μ2 < 0.09 ) = 0.95

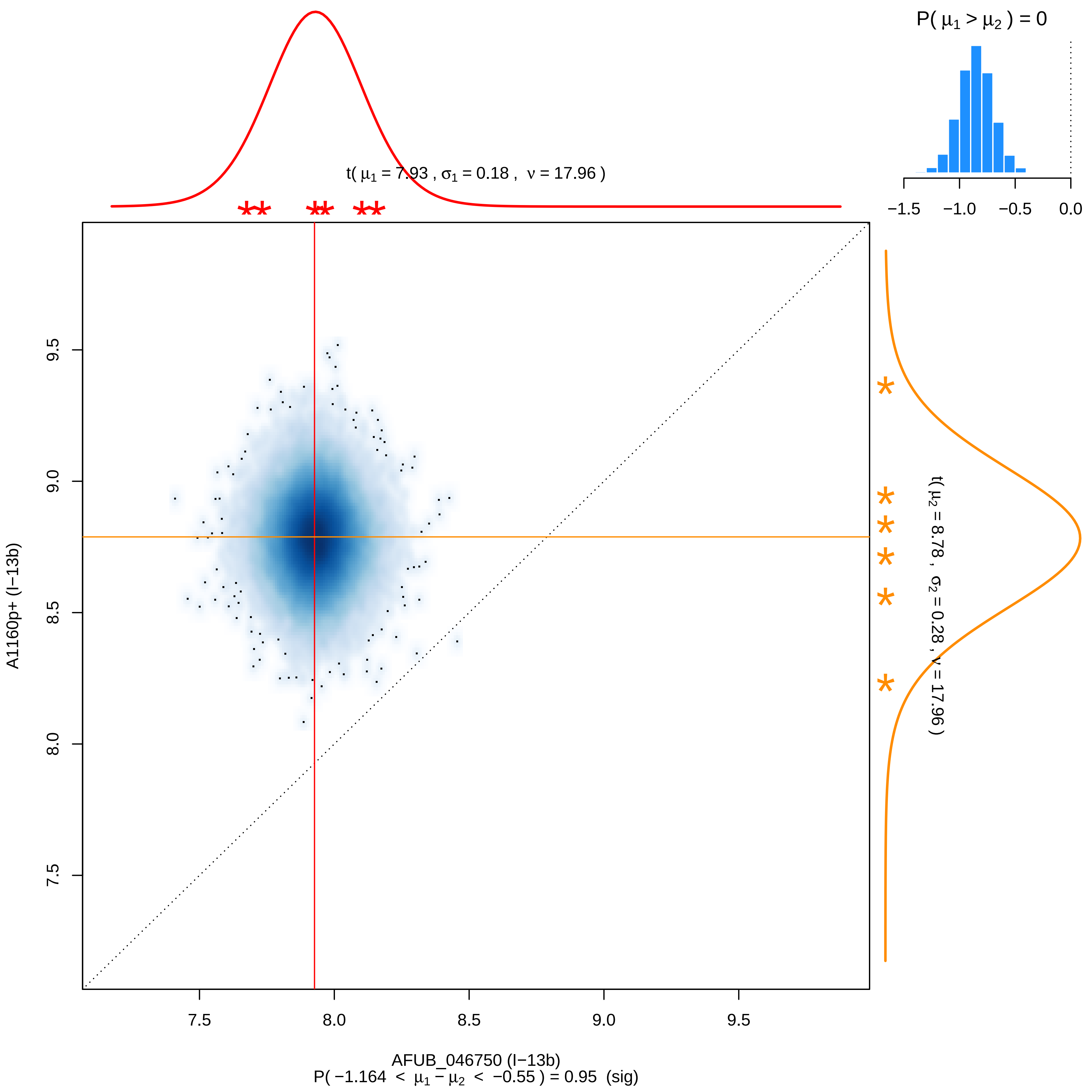

P( μ1 > μ2 ) = 0
−1.5
−1.0
−0.5
0.0
t( μ1 = 7.93 , σ1 = 0.18 , ν = 17.96 )
t( μ2 = 8.78 , σ2 = 0.28 , ν = 17.96 )
9.5
9.0
8.5
8.0
7.5
A1160p+ (I−13b)
7.5
8.0
8.5
9.0
9.5
AFUB_046750 (I−13b)
P( −1.164 < μ1 − μ2 < −0.55 ) = 0.95 (sig)

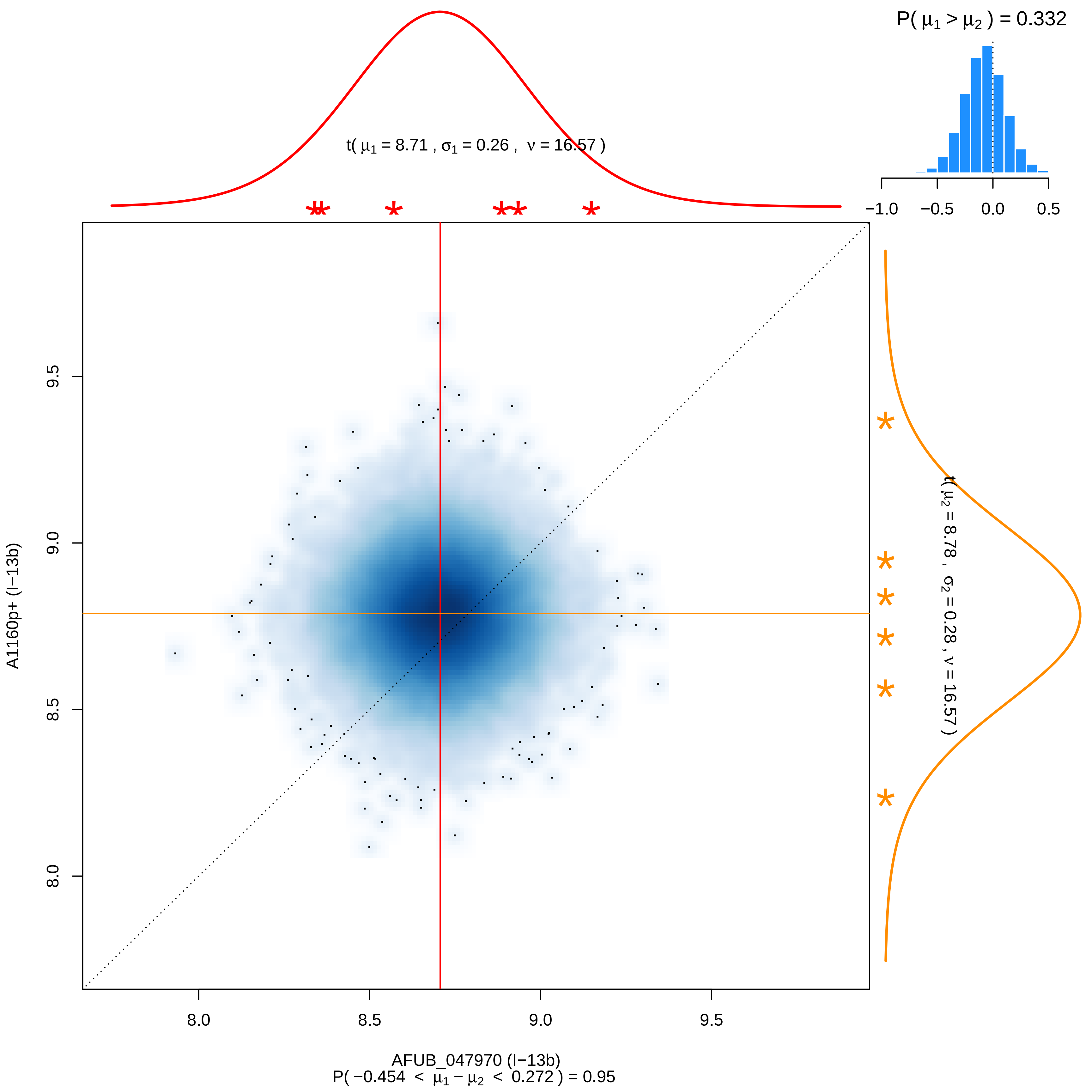

P( μ1 > μ2 ) = 0.332
t( μ1 = 8.71 , σ1 = 0.26 , ν = 16.57 )
−1.0 −0.5 0.0 0.5
t( μ2 = 8.78 , σ2 = 0.28 , ν = 16.57 )
A1160p+ (I−13b)
8.0 8.5 9.0 9.5
AFUB_047970 (I−13b)
P( −0.454 < μ1 − μ2 < 0.272 ) = 0.95

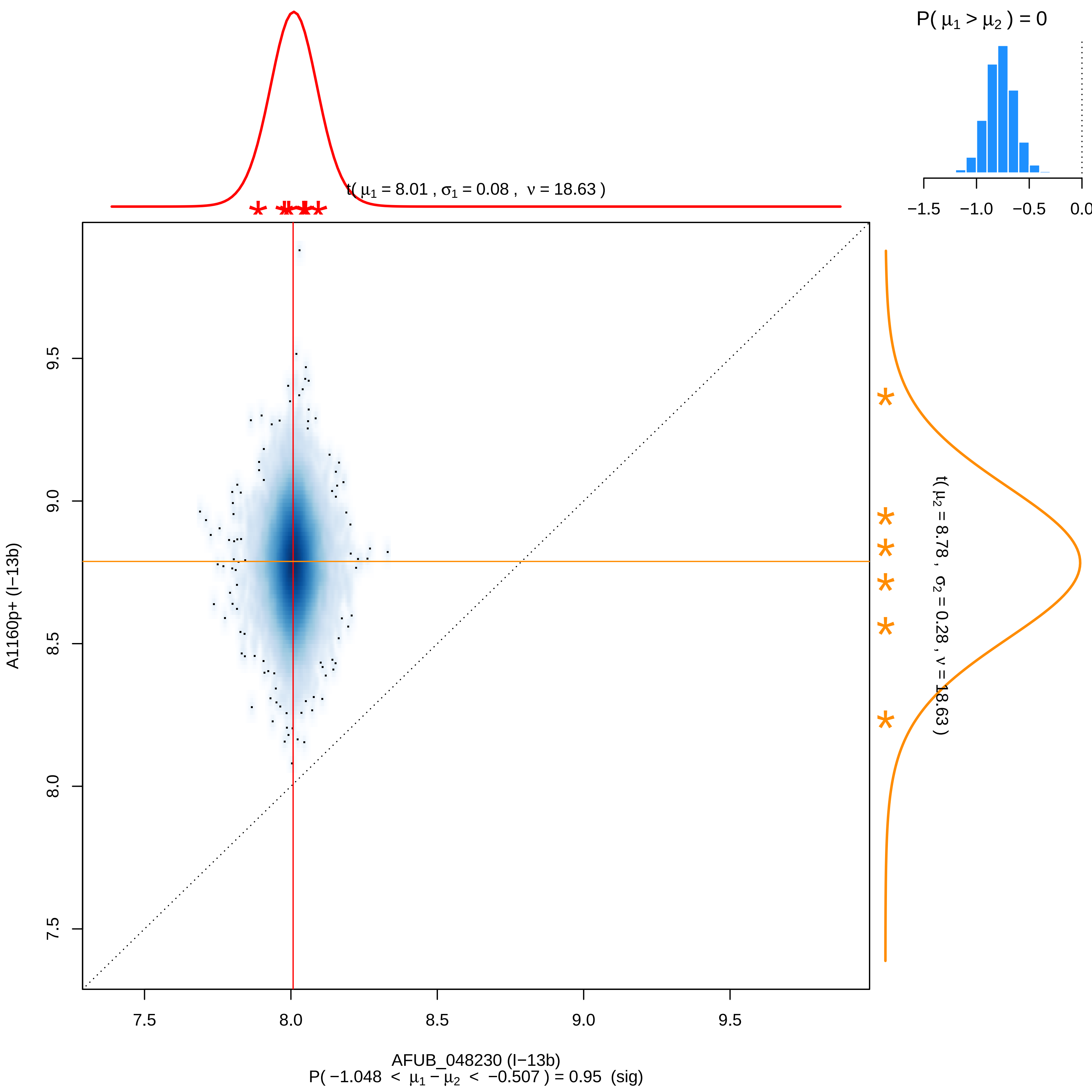

P( μ1 > μ2 ) = 0
−1.5
−1.0
−0.5
0.0
t( μ1 = 8.01 , σ1 = 0.08 , ν = 18.63 )
t( μ2 = 8.78 , σ2 = 0.28 , ν = 18.63 )
A1160p+ (I−13b)
9.5
9.0
8.5
8.0
7.5
7.5
8.0
8.5
9.0
9.5
AFUB_048230 (I−13b)
P( −1.048 < μ1 − μ2 < −0.507 ) = 0.95 (sig)

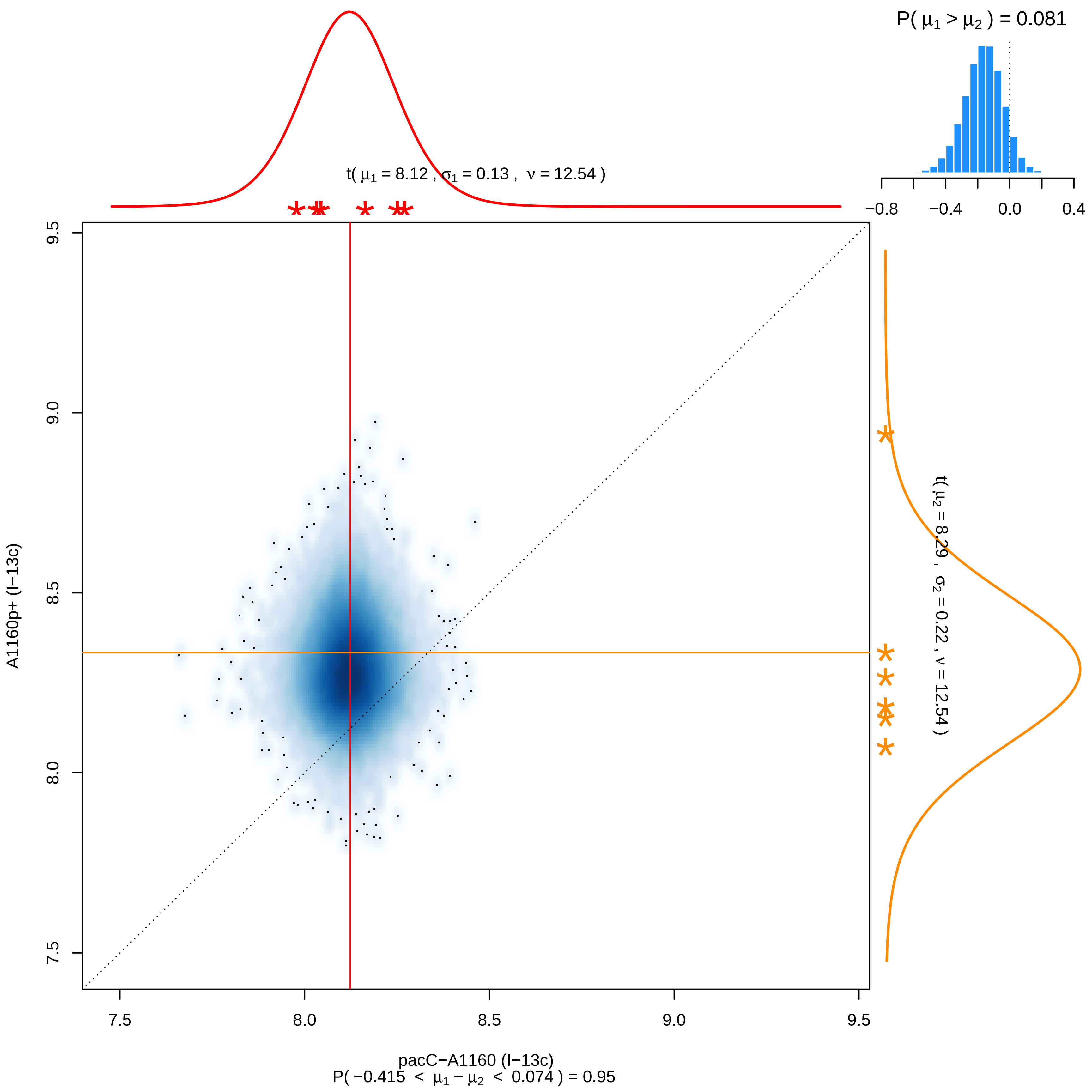

P( μ1 > μ2 ) = 0.081
−0.8
−0.4
0.0
0.4
t( μ1 = 8.12 , σ1 = 0.13 , ν = 12.54 )
t( μ2 = 8.29 , σ2 = 0.22 , ν = 12.54 )
A1160p+ (I−13c)
9.5
9.0
8.5
8.0
7.5
7.5
8.0
8.5
9.0
9.5
pacC−A1160 (I−13c)
P( −0.415 < μ1 − μ2 < 0.074 ) = 0.95

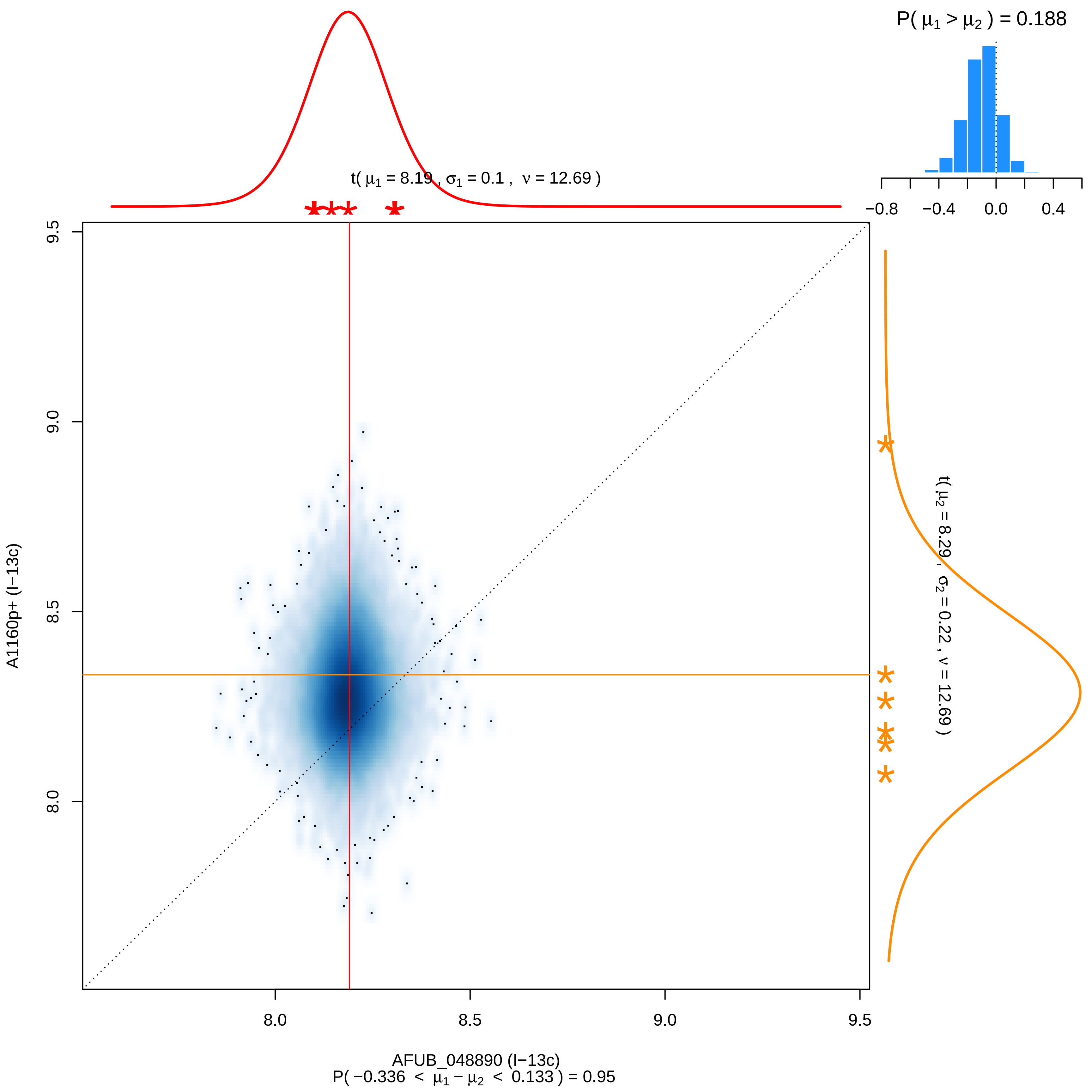

P( μ1 > μ2 ) = 0.188
−0.8
−0.4
0.0
0.4
t( μ1 = 8.19 , σ1 = 0.1 , ν = 12.69 )
t( μ2 = 8.29 , σ2 = 0.22 , ν = 12.69 )
A1160p+ (I−13c)
9.5
9.0
8.5
8.0
AFUB_048890 (I−13c)
P( −0.336 < μ1 − μ2 < 0.133 ) = 0.95

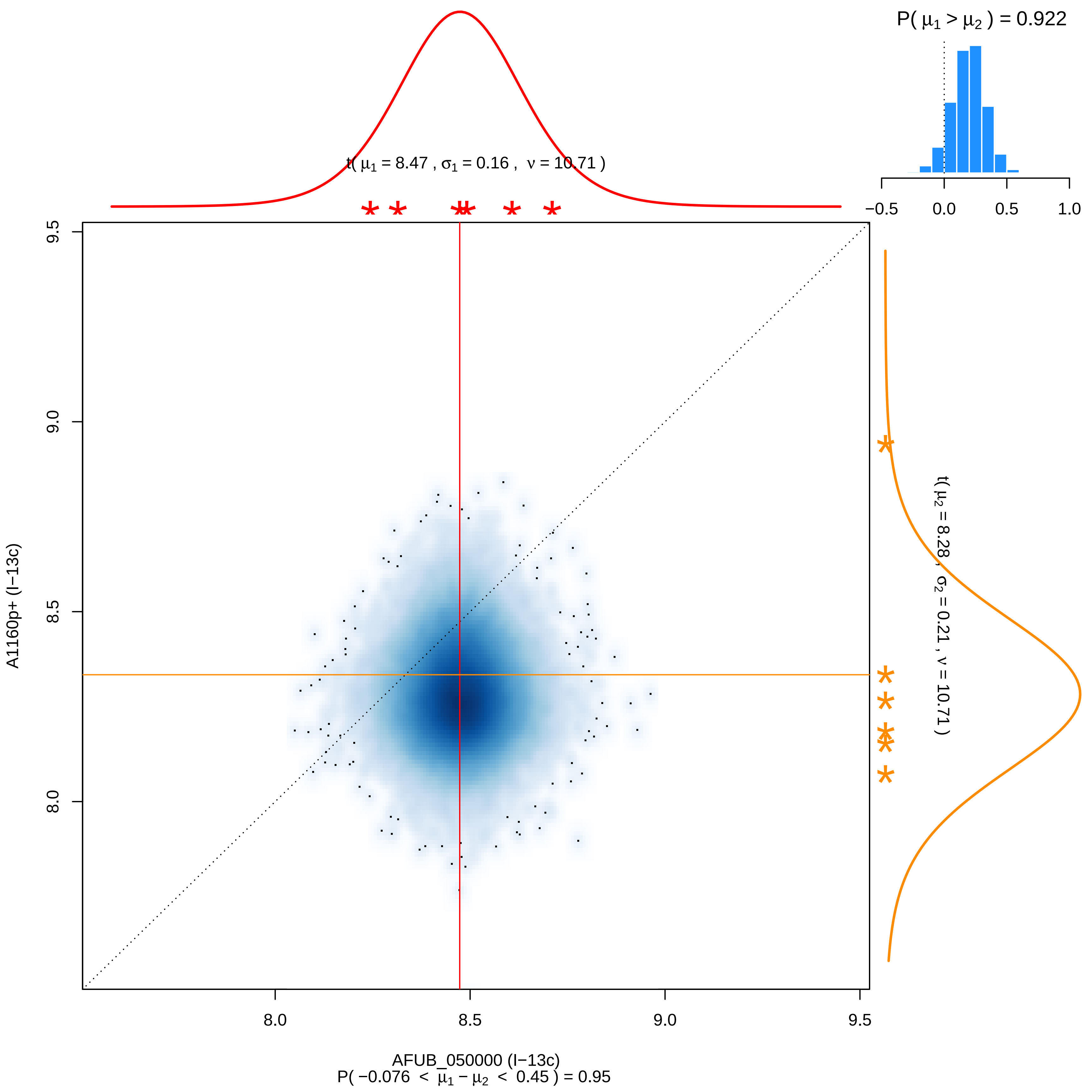

P( μ1 > μ2 ) = 0.922
−0.5
0.0
0.5
1.0
t( μ1 = 8.47 , σ1 = 0.16 , ν = 10.71 )
t( μ2 = 8.28 , σ2 = 0.21 , ν = 10.71 )
9.5
9.0
8.5
8.0
A1160p+ (I−13c)
8.0
8.5
9.0
9.5
AFUB_050000 (I−13c)
P( −0.076 < μ1 − μ2 < 0.45 ) = 0.95

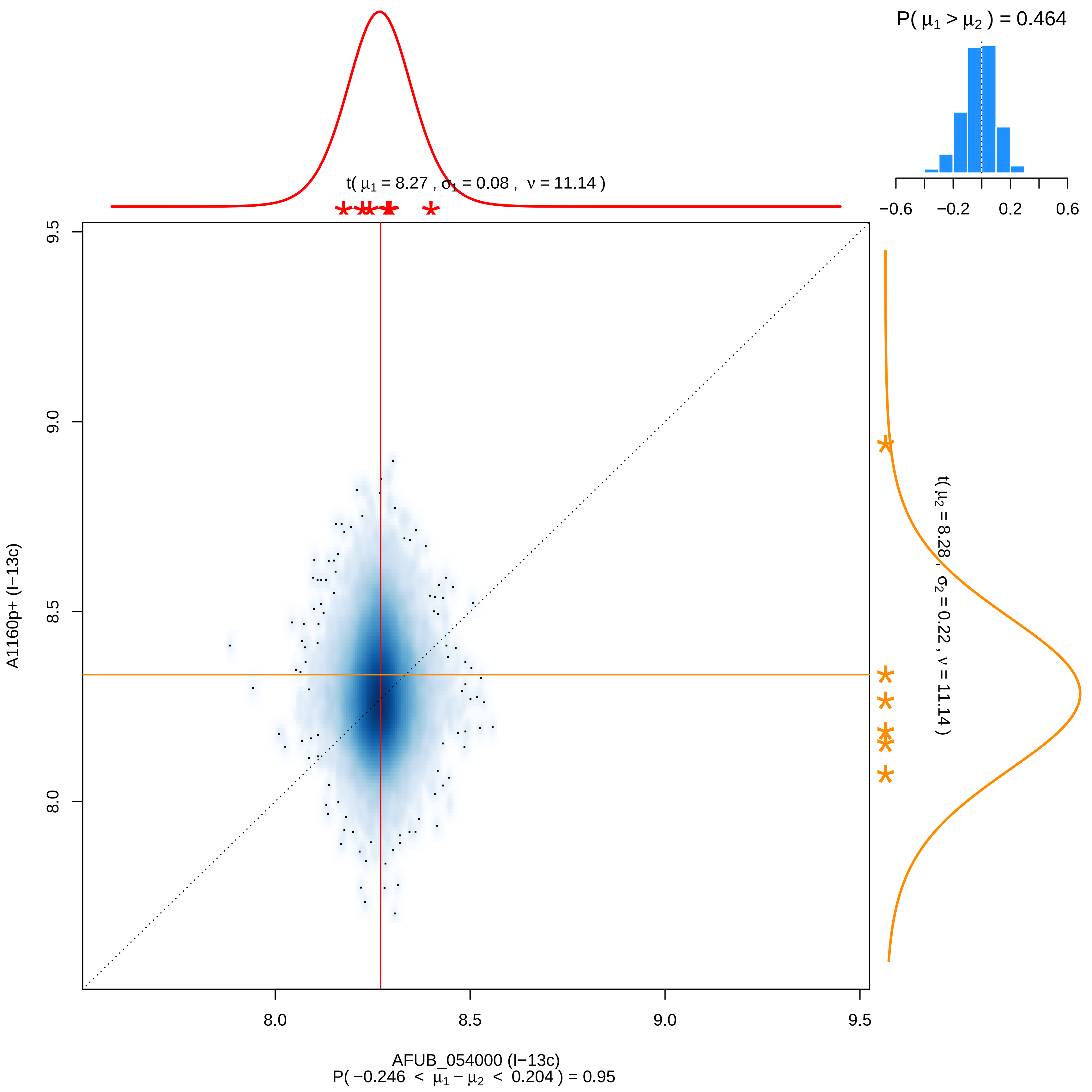

P( μ1 > μ2 ) = 0.464
−0.6
−0.2
0.2
0.6
t( μ1 = 8.27 , σ1 = 0.08 , ν = 11.14 )
t( μ2 = 8.28 , σ2 = 0.22 , ν = 11.14 )
A1160p+ (I−13c)
9.5
9.0
8.5
8.0
AFUB_054000 (I−13c)
P( −0.246 < μ1 − μ2 < 0.204 ) = 0.95

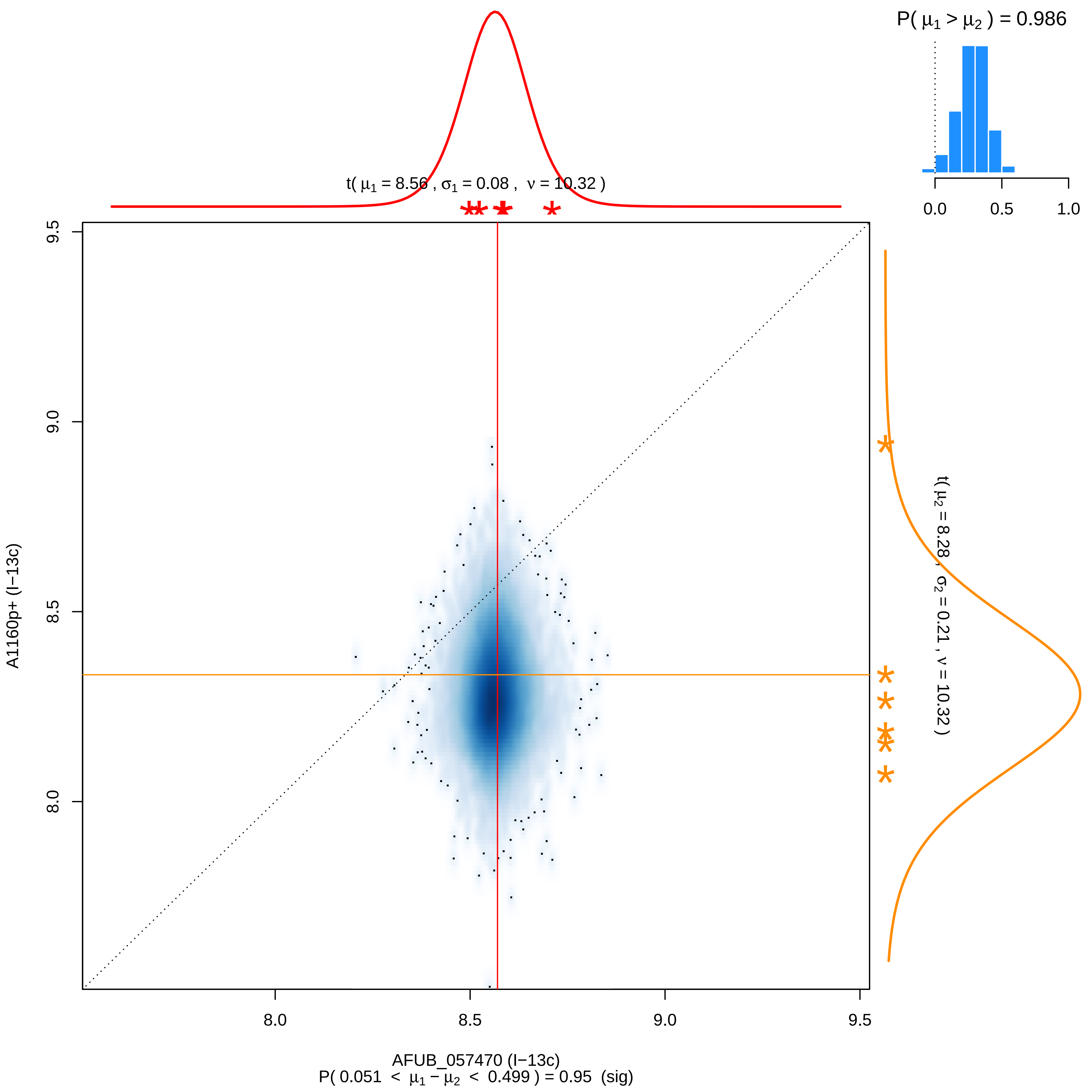

P( μ1 > μ2 ) = 0.986
0.0
0.5
1.0
t( μ1 = 8.56 , σ1 = 0.08 , ν = 10.32 )
9.5
9.0
8.5
8.0
A1160p+ (I−13c)
t( μ2 = 8.28 , σ2 = 0.21 , ν = 10.32 )
8.0
8.5
9.0
9.5
AFUB_057470 (I−13c)
P( 0.051 < μ1 − μ2 < 0.499 ) = 0.95 (sig)

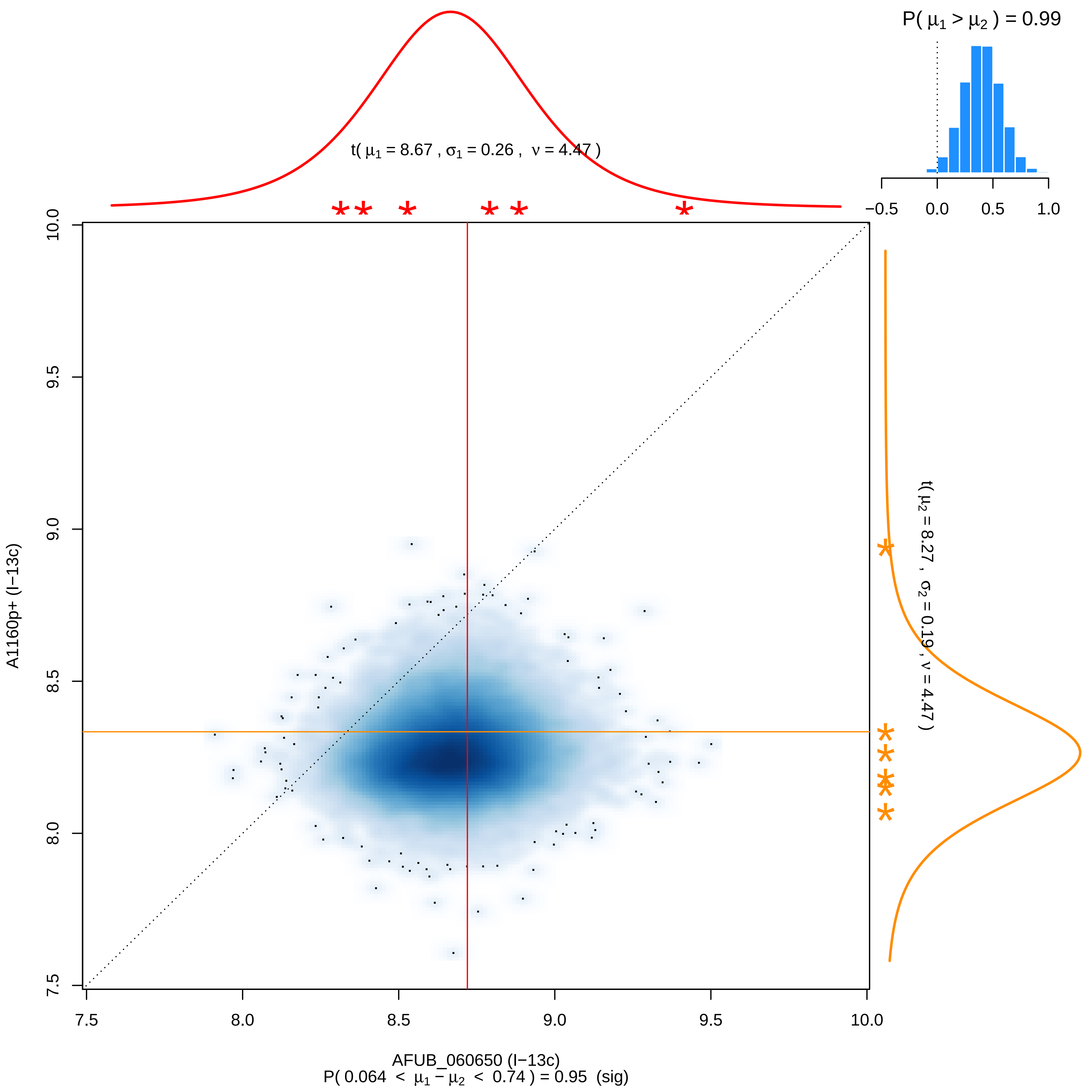

P( μ1 > μ2 ) = 0.99
−0.5
0.0
0.5
1.0
t( μ1 = 8.67 , σ1 = 0.26 , ν = 4.47 )
t( μ2 = 8.27 , σ2 = 0.19 , ν = 4.47 )
A1160p+ (I−13c)
AFUB_060650 (I−13c)
P( 0.064 < μ1 − μ2 < 0.74 ) = 0.95 (sig)
7.5
8.0
8.5
9.0
9.5
10.0

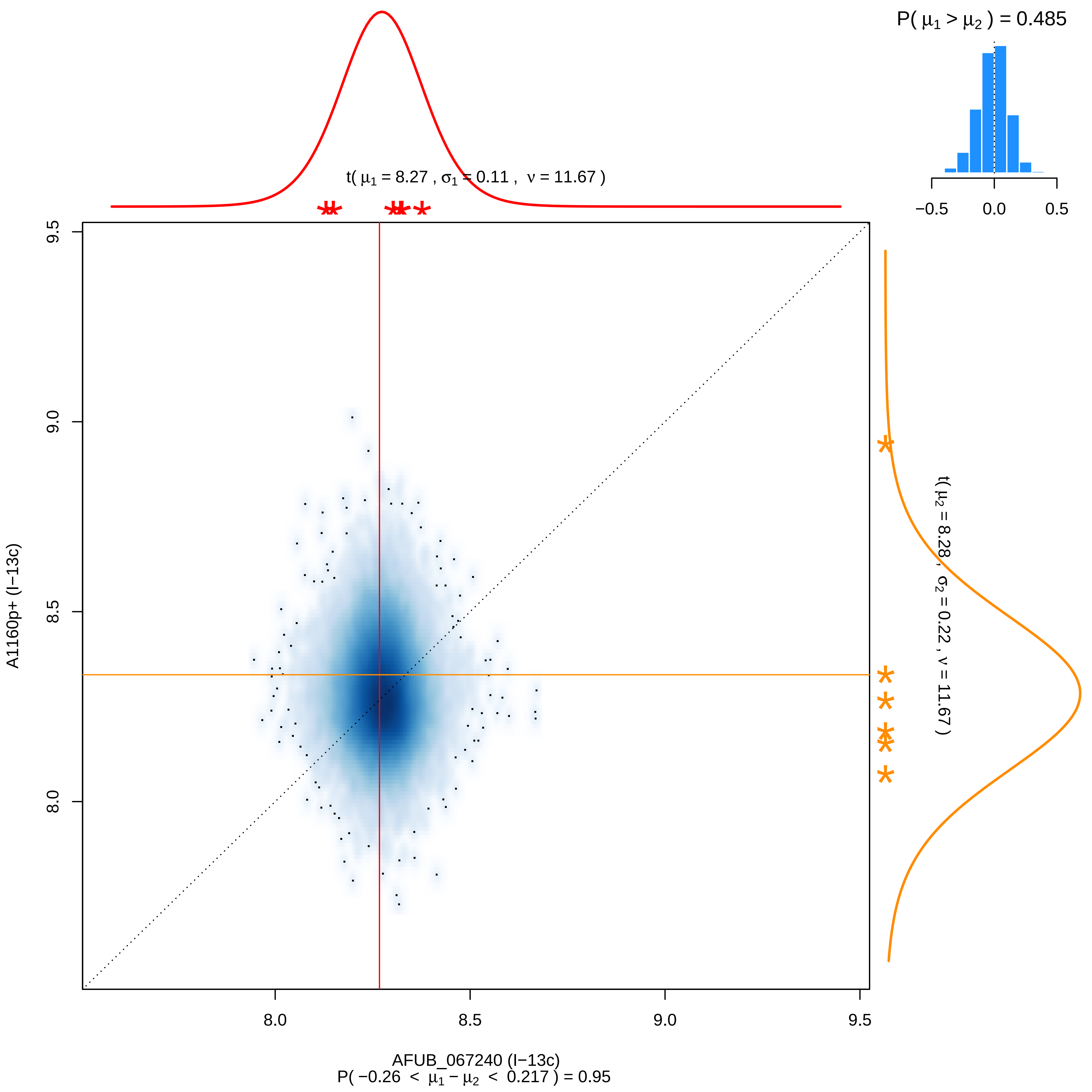

P( μ1 > μ2 ) = 0.485
−0.5
0.0
0.5
t( μ1 = 8.27 , σ1 = 0.11 , ν = 11.67 )
t( μ2 = 8.28 , σ2 = 0.22 , ν = 11.67 )
9.5
9.0
8.5
8.0
A1160p+ (I−13c)
AFUB_067240 (I−13c)
P( −0.26 < μ1 − μ2 < 0.217 ) = 0.95

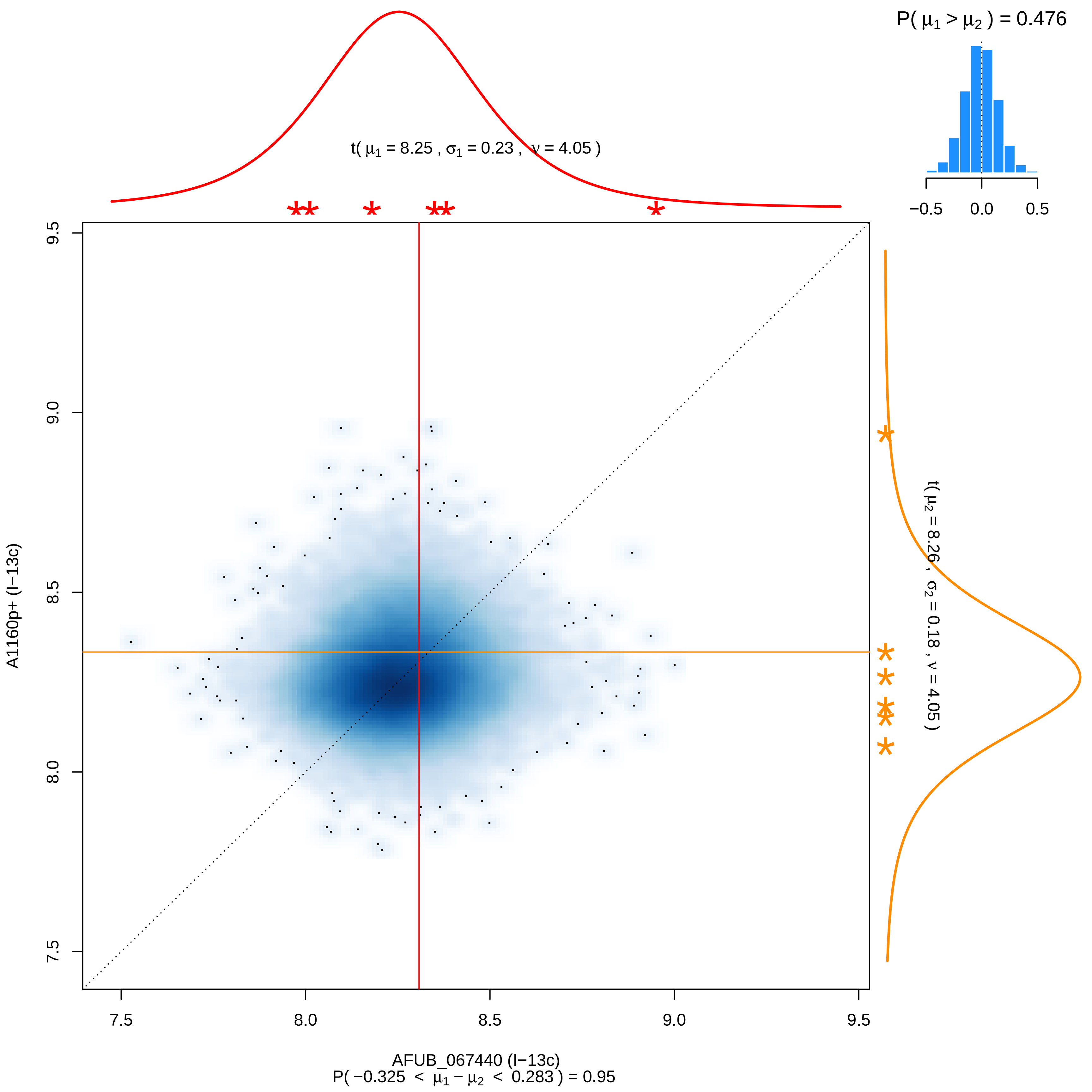

P( $\mu_1 > \mu_2$ ) = 0.476
−0.5
0.0
0.5
t( $\mu_1$ = 8.25 , $\sigma_1$ = 0.23 , $\nu$ = 4.05 )
t( $\mu_2$ = 8.26 , $\sigma_2$ = 0.18 , $\nu$ = 4.05 )
9.5
9.0
8.5
8.0
7.5
A1160p+ (I−13c)
7.5
8.0
8.5
9.0
9.5
AFUB_067440 (I−13c)
P( −0.325 < $\mu_1 - \mu_2$ < 0.283 ) = 0.95

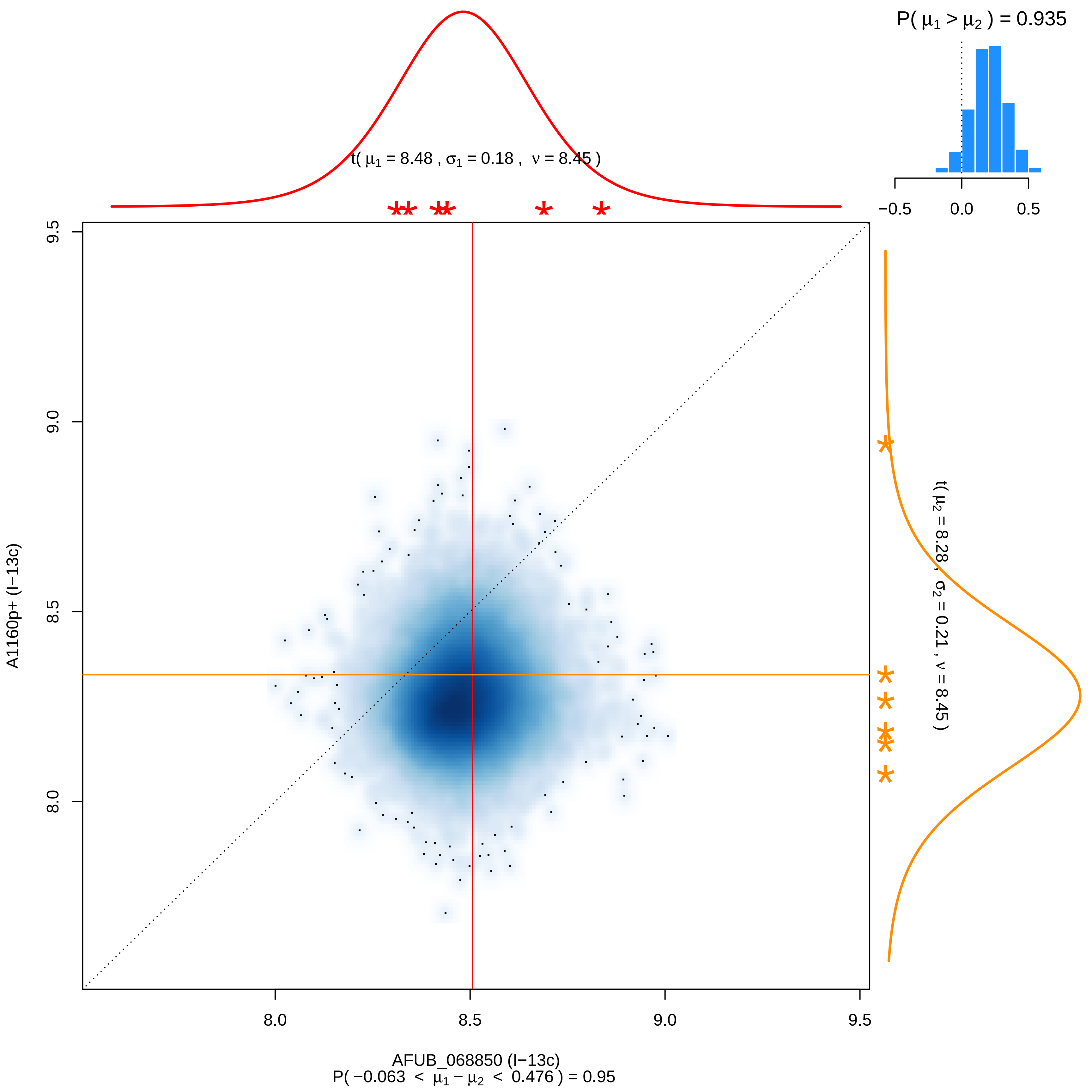

P( μ1 > μ2 ) = 0.935
−0.5
0.0
0.5
t( μ1 = 8.48 , σ1 = 0.18 , ν = 8.45 )
t( μ2 = 8.28 , σ2 = 0.21 , ν = 8.45 )
A1160p+ (I−13c)
AFUB_068850 (I−13c)
8.0
8.5
9.0
9.5
P( −0.063 < μ1 − μ2 < 0.476 ) = 0.95

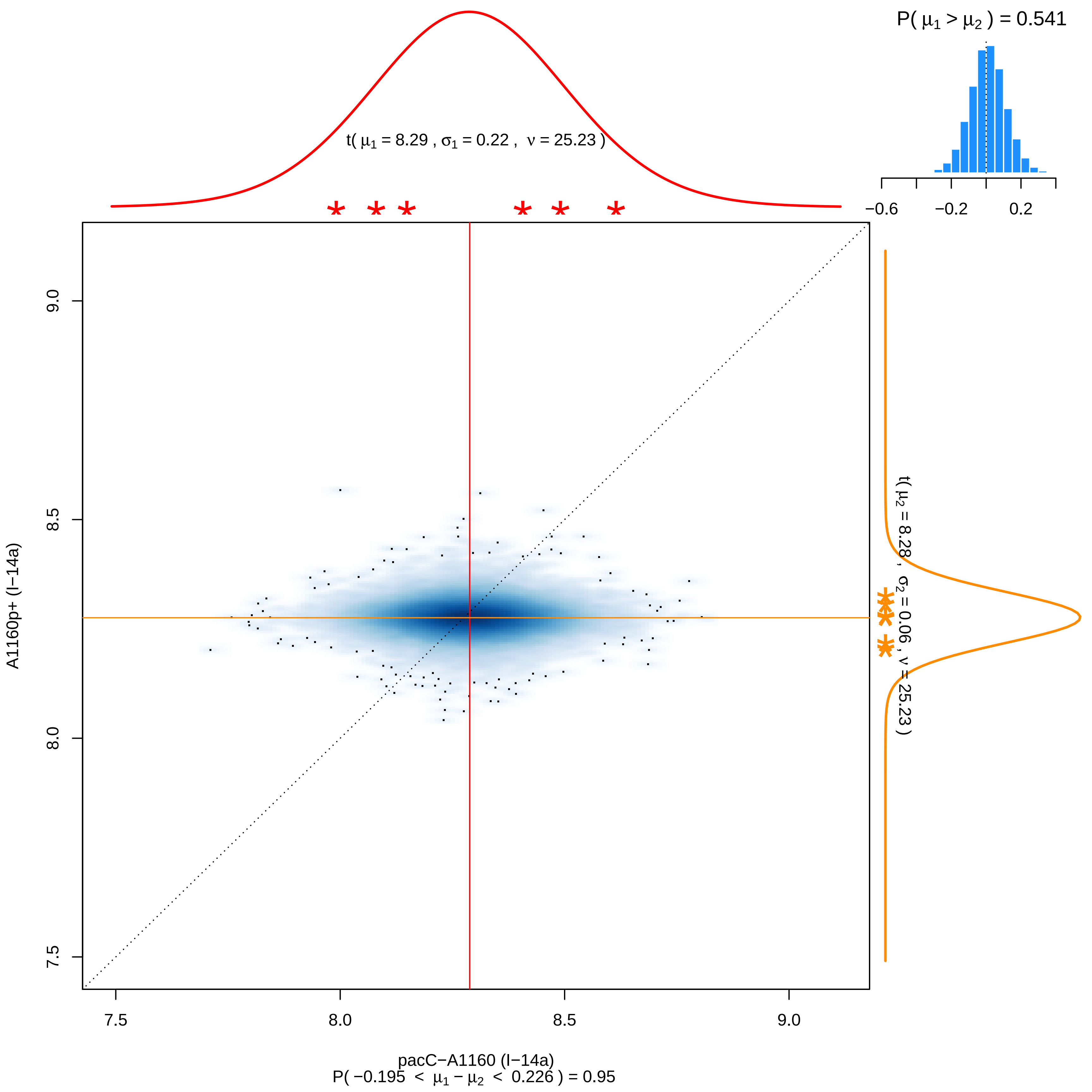

P( μ1 > μ2 ) = 0.541
−0.6
−0.2
0.2
t( μ1 = 8.29 , σ1 = 0.22 , ν = 25.23 )
t( μ2 = 8.28 , σ2 = 0.06 , ν = 25.23 )
9.0
8.5
8.0
7.5
A1160p+ (I−14a)
7.5
8.0
8.5
9.0
pacC−A1160 (I−14a)
P( −0.195 < μ1 − μ2 < 0.226 ) = 0.95

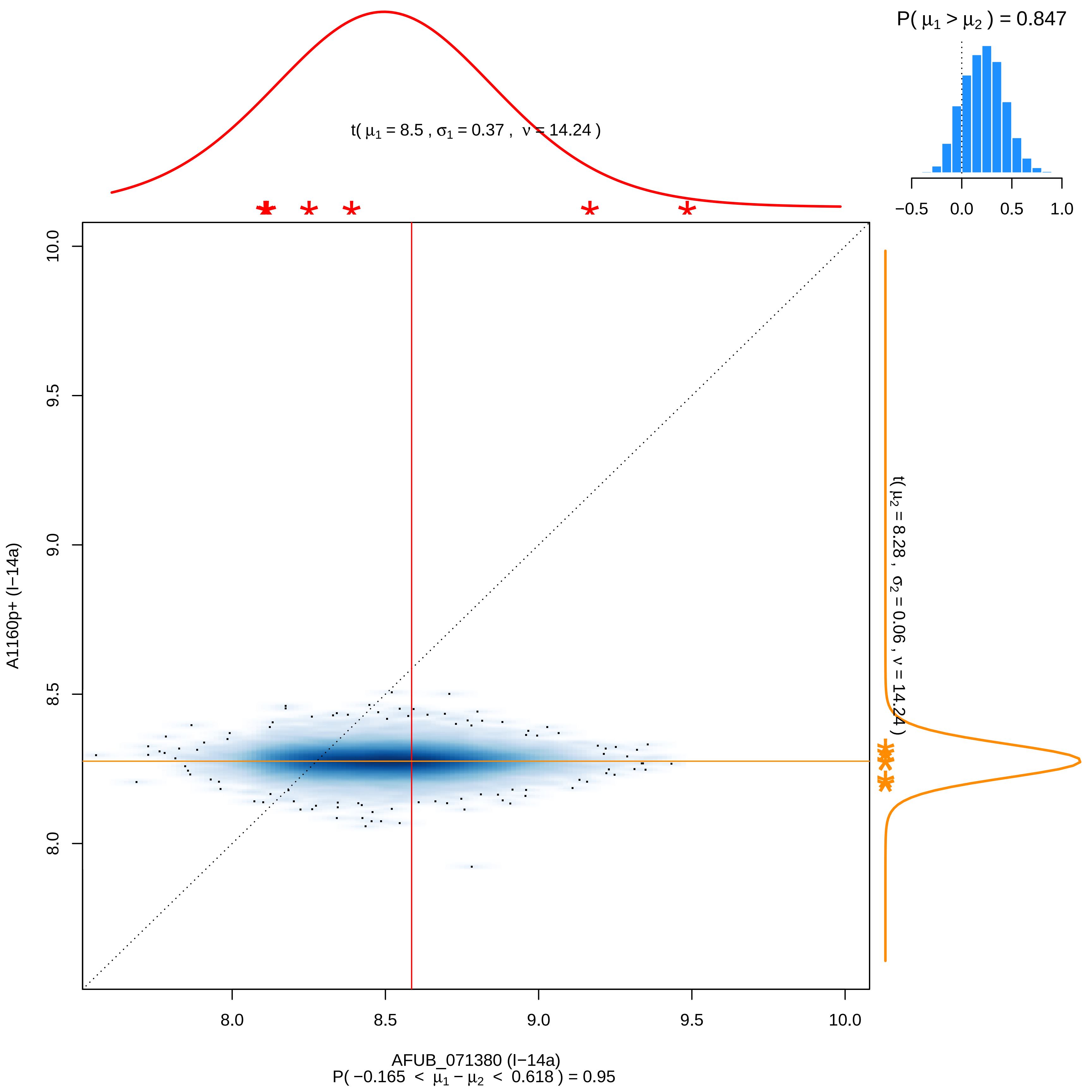
P( μ1 > μ2 ) = 0.847
−0.5 0.0 0.5 1.0
t( μ1 = 8.5 , σ1 = 0.37 , ν = 14.24 )
t( μ2 = 8.28 , σ2 = 0.06 , ν = 14.24 )
A1160p+ (I−14a)
8.0 8.5 9.0 9.5 10.0
AFUB_071380 (I−14a)
P( −0.165 < μ1 − μ2 < 0.618 ) = 0.95

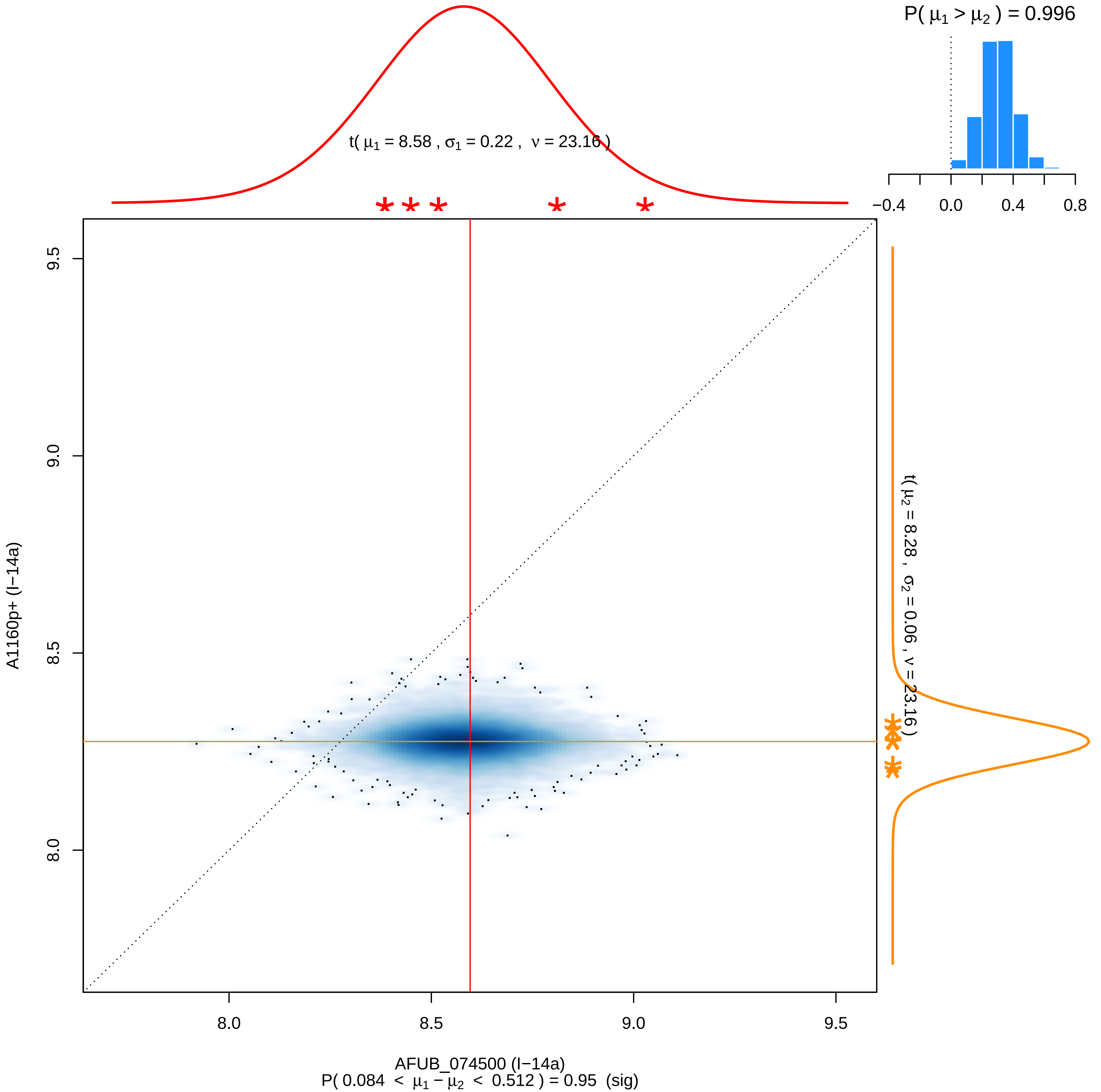

P( μ1 > μ2 ) = 0.996
−0.4
0.0
0.4
0.8
t( μ1 = 8.58 , σ1 = 0.22 , ν = 23.16 )
t( μ2 = 8.28 , σ2 = 0.06 , ν = 23.16 )
A1160p+ (I−14a)
9.5
9.0
8.5
8.0
AFUB_074500 (I−14a)
P( 0.084 < μ1 − μ2 < 0.512 ) = 0.95 (sig)

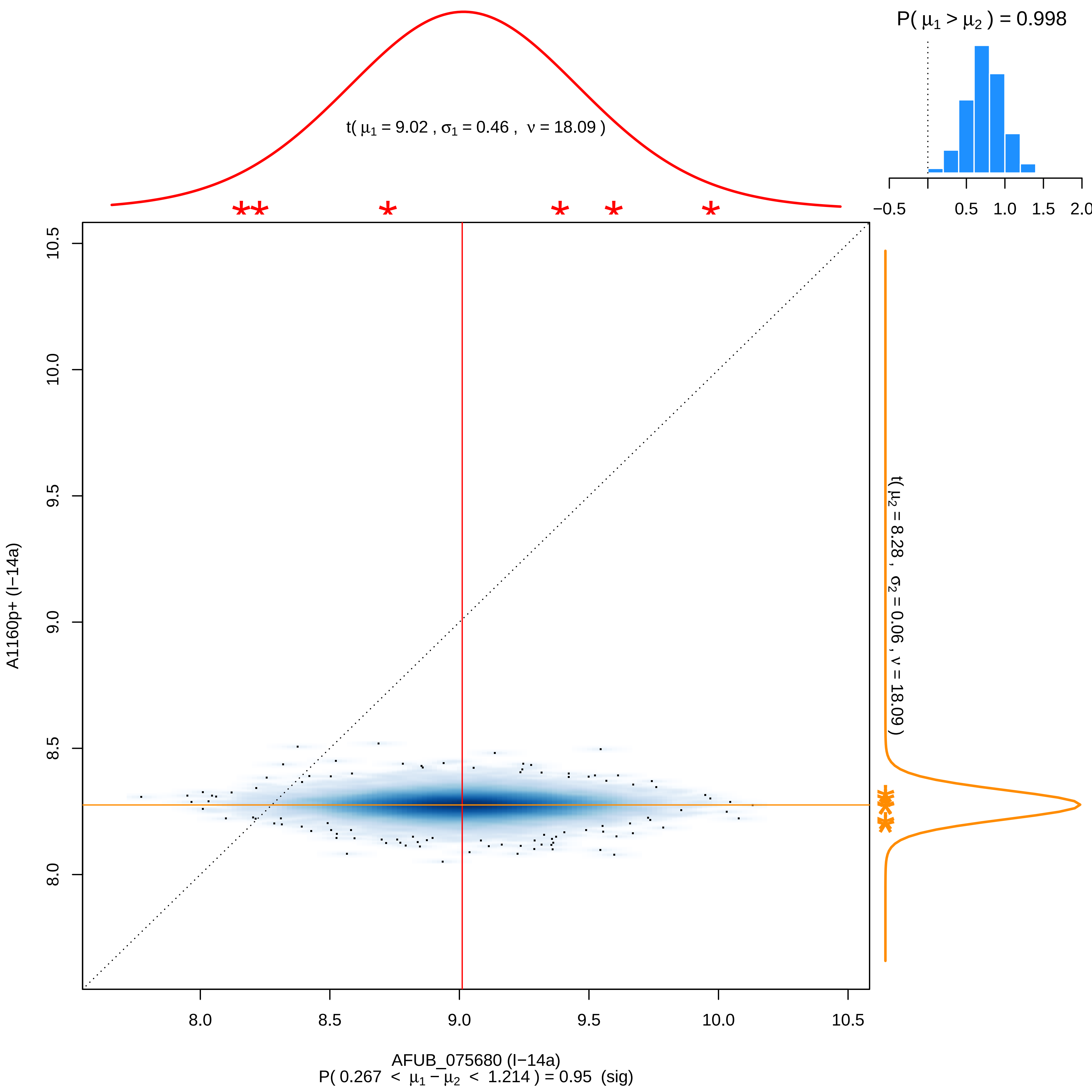
P( μ1 > μ2 ) = 0.998
t( μ1 = 9.02 , σ1 = 0.46 , ν = 18.09 )
−0.5
0.5
1.0
1.5
2.0
10.5
10.0
9.5
9.0
8.5
8.0
A1160p+ (I−14a)
t( μ2 = 8.28 , σ2 = 0.06 , ν = 18.09 )
8.0
8.5
9.0
9.5
10.0
10.5
AFUB_075680 (I−14a)
P( 0.267 < μ1 − μ2 < 1.214 ) = 0.95 (sig)

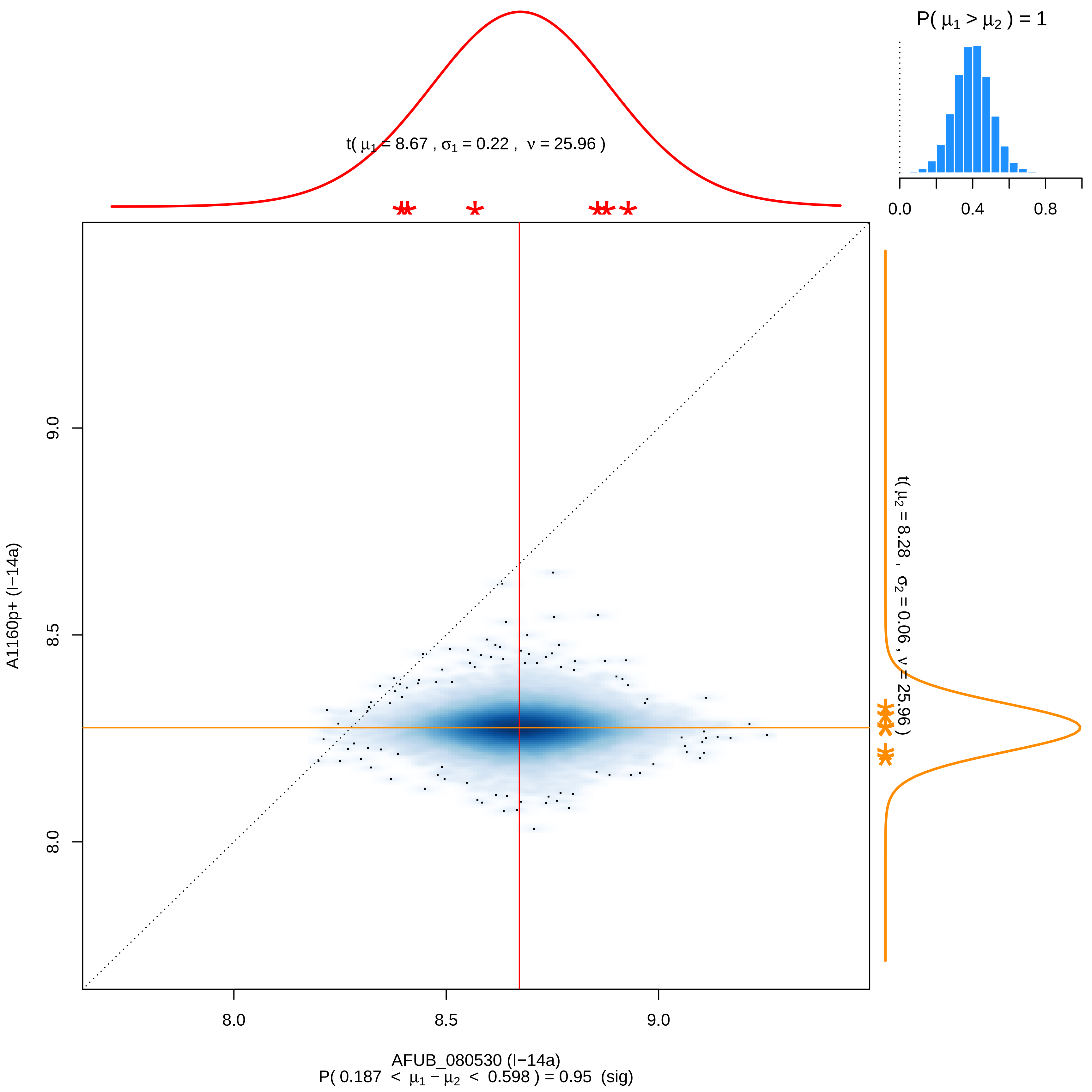

P( μ1 > μ2 ) = 1
0.0
0.4
0.8
t( μ1 = 8.67 , σ1 = 0.22 , ν = 25.96 )
t( μ2 = 8.28 , σ2 = 0.06 , ν = 25.96 )
A1160p+ (I−14a)
9.0
8.5
8.0
8.0
8.5
9.0
AFUB_080530 (I−14a)
P( 0.187 < μ1 − μ2 < 0.598 ) = 0.95 (sig)

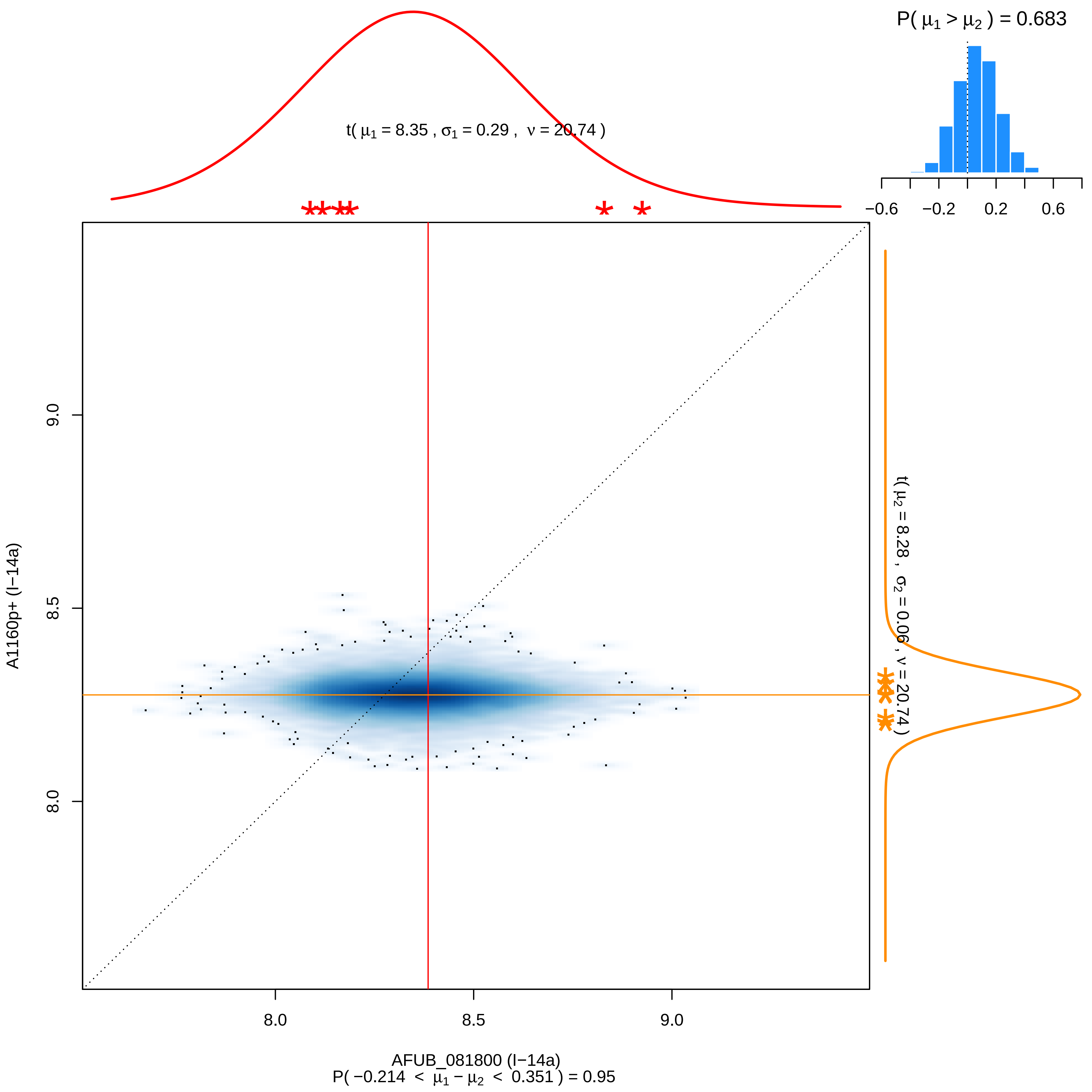

P( μ1 > μ2 ) = 0.683
−0.6
−0.2
0.2
0.6
t( μ1 = 8.35 , σ1 = 0.29 , ν = 20.74 )
t( μ2 = 8.28 , σ2 = 0.06 , ν = 20.74 )
A1160p+ (I−14a)
8.0
8.5
9.0
AFUB_081800 (I−14a)
P( −0.214 < μ1 − μ2 < 0.351 ) = 0.95

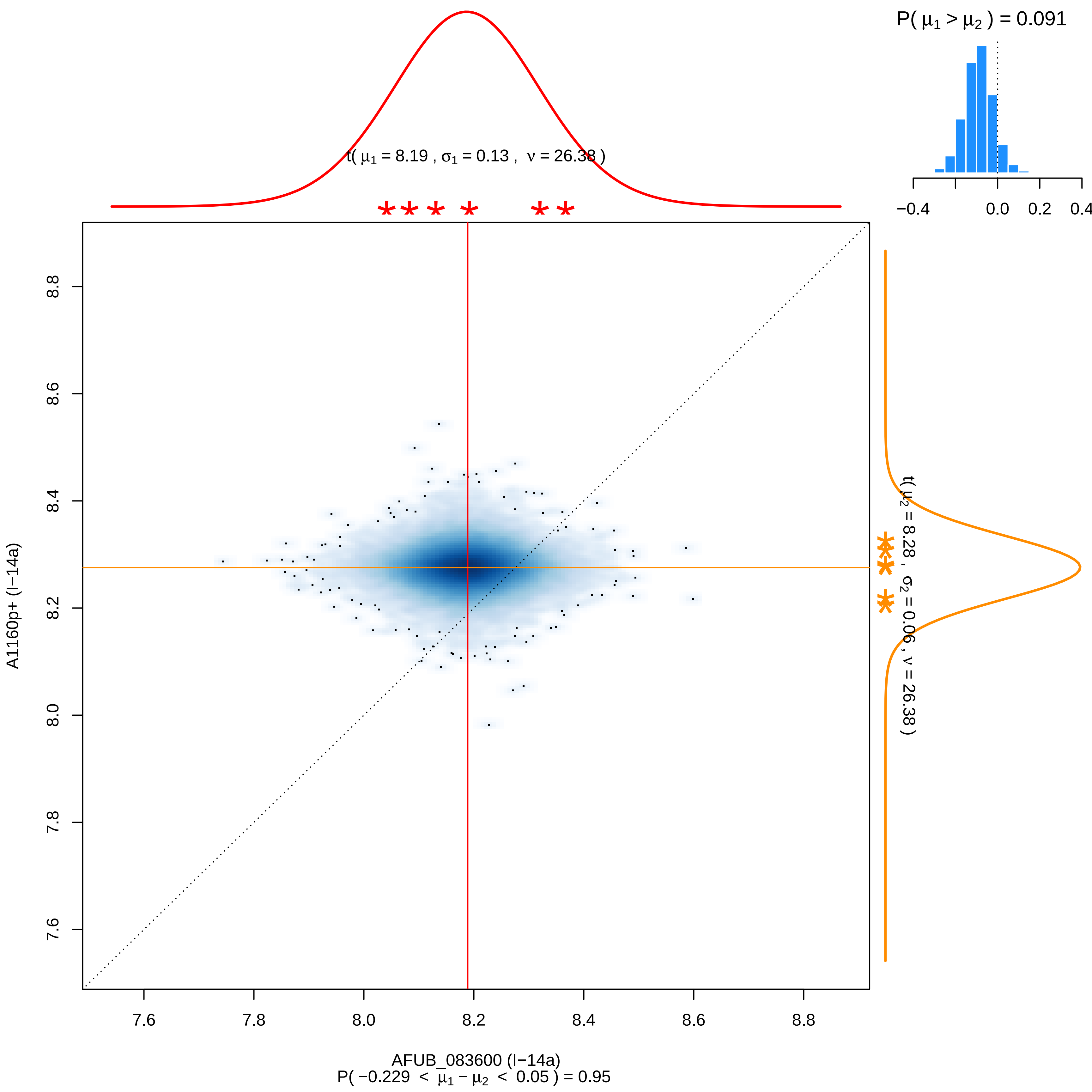

P( μ1 > μ2 ) = 0.091
−0.4
0.0
0.2
0.4
t( μ1 = 8.19 , σ1 = 0.13 , ν = 26.38 )
t( μ2 = 8.28 , σ2 = 0.06 , ν = 26.38 )
A1160p+ (I−14a)
8.8
8.6
8.4
8.2
8.0
7.8
7.6
7.6
7.8
8.0
8.2
8.4
8.6
8.8
AFUB_083600 (I−14a)
P( −0.229 < μ1 − μ2 < 0.05 ) = 0.95

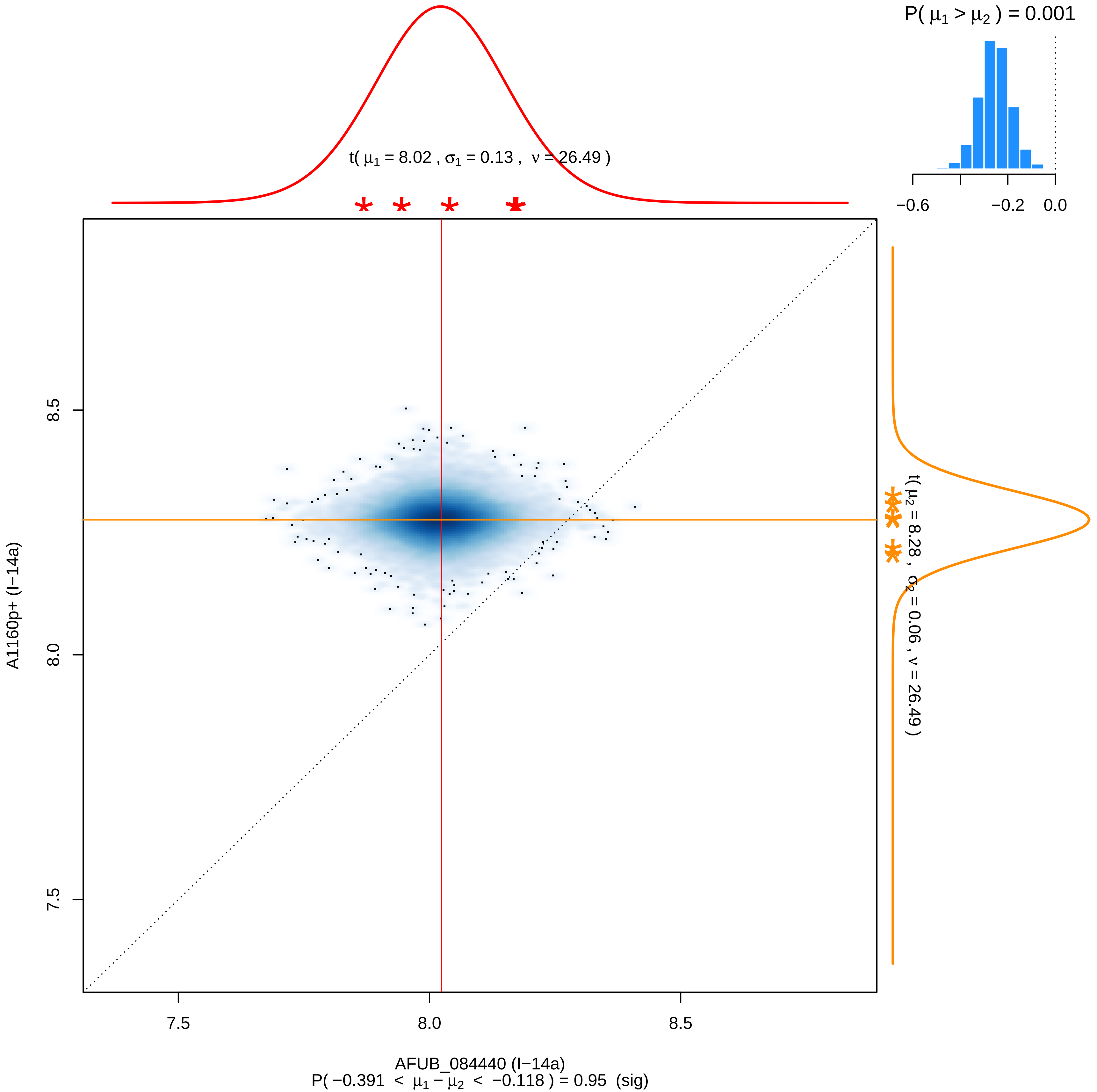

P( μ1 > μ2 ) = 0.001
−0.6
−0.2
0.0
t( μ1 = 8.02 , σ1 = 0.13 , ν = 26.49 )
t( μ2 = 8.28 , σ2 = 0.06 , ν = 26.49 )
A1160p+ (I−14a)
8.5
8.0
7.5
7.5
8.0
8.5
AFUB_084440 (I−14a)
P( −0.391 < μ1 − μ2 < −0.118 ) = 0.95 (sig)

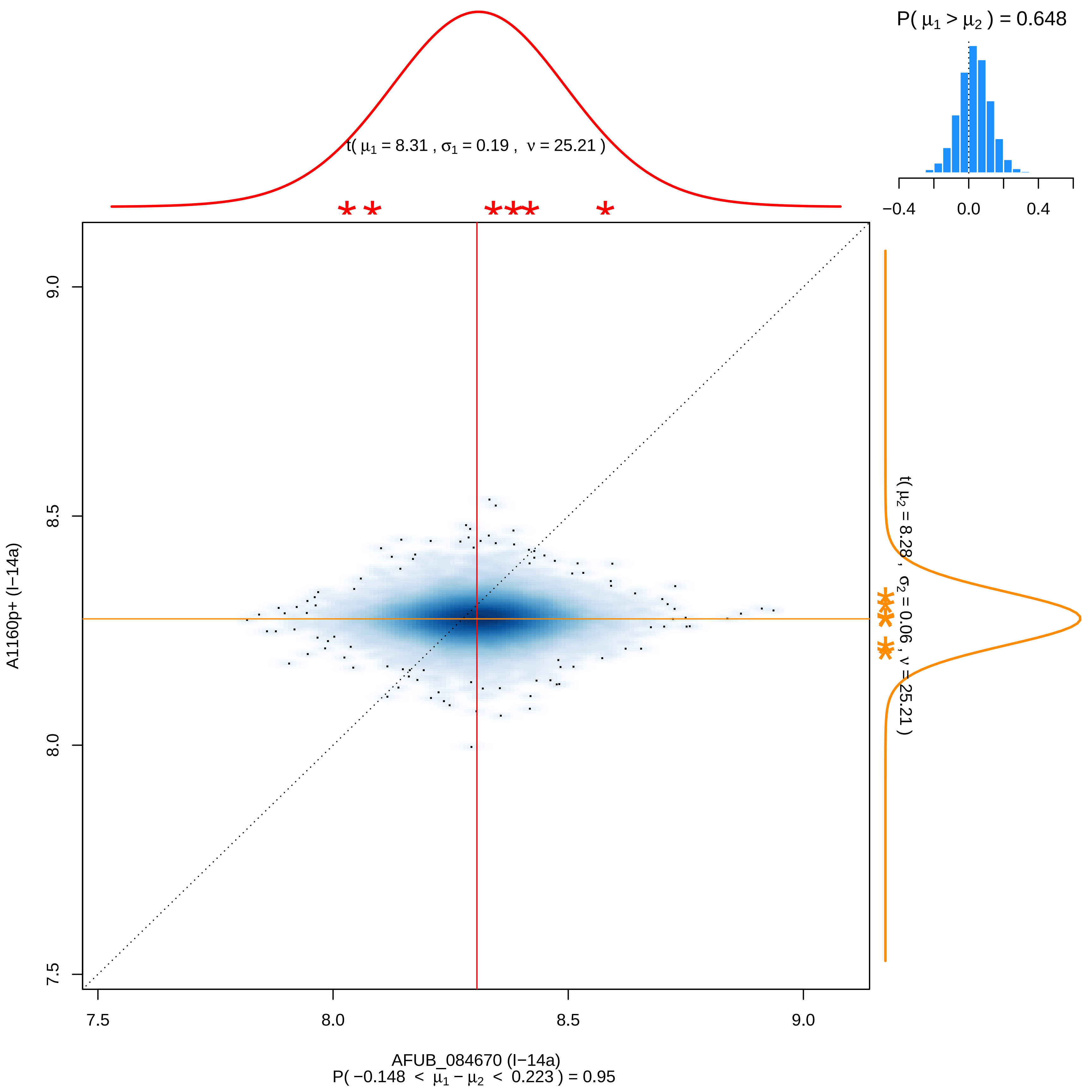

P( μ1 > μ2 ) = 0.648
−0.4
0.0
0.4
t( μ1 = 8.31 , σ1 = 0.19 , ν = 25.21 )
t( μ2 = 8.28 , σ2 = 0.06 , ν = 25.21 )
A1160p+ (I−14a)
9.0
8.5
8.0
7.5
7.5
8.0
8.5
9.0
AFUB_084670 (I−14a)
P( −0.148 < μ1 − μ2 < 0.223 ) = 0.95

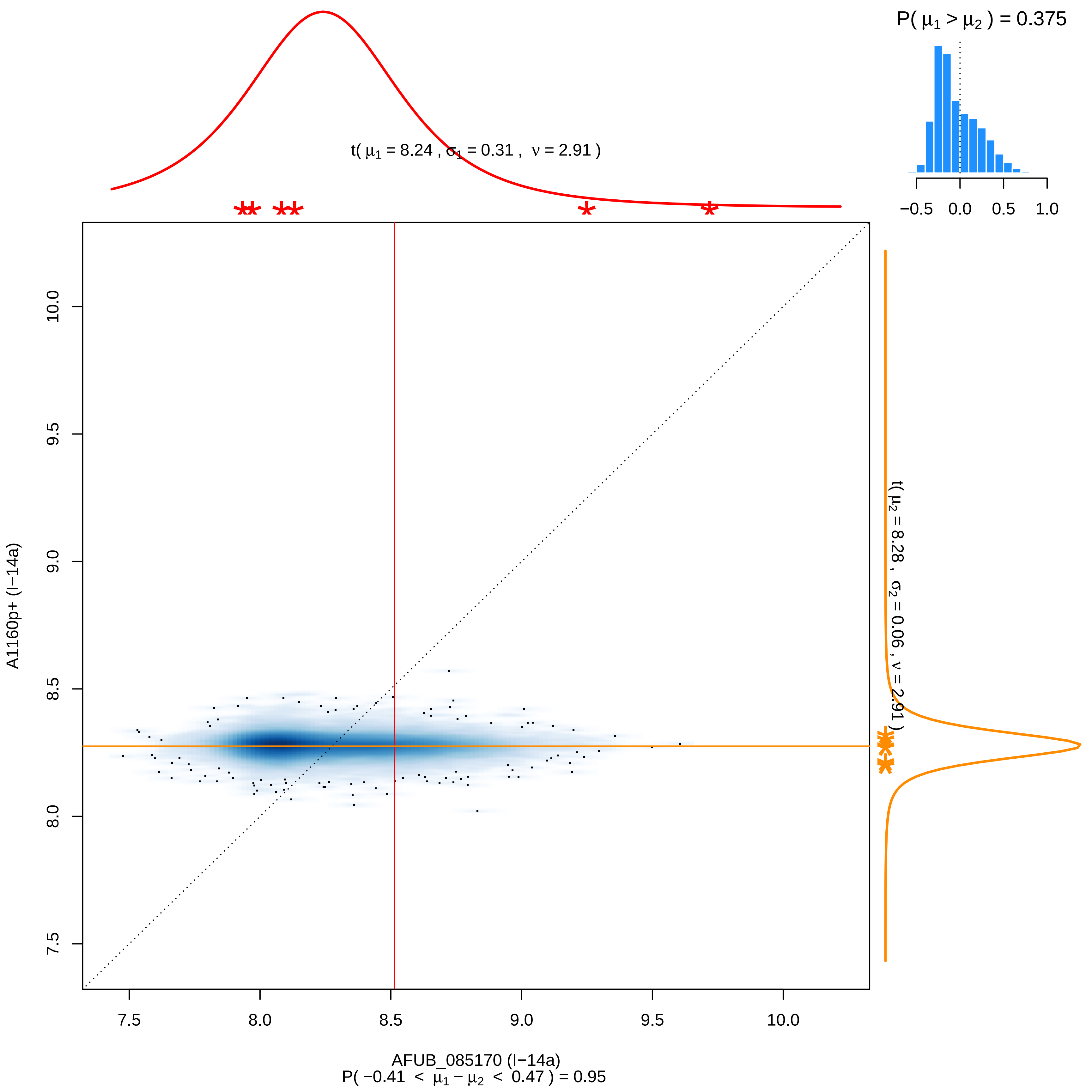

P( μ1 > μ2 ) = 0.375
t( μ1 = 8.24 , σ1 = 0.31 , ν = 2.91 )
t( μ2 = 8.28 , σ2 = 0.06 , ν = 2.91 )
A1160p+ (I−14a)
AFUB_085170 (I−14a)
P( −0.41 < μ1 − μ2 < 0.47 ) = 0.95

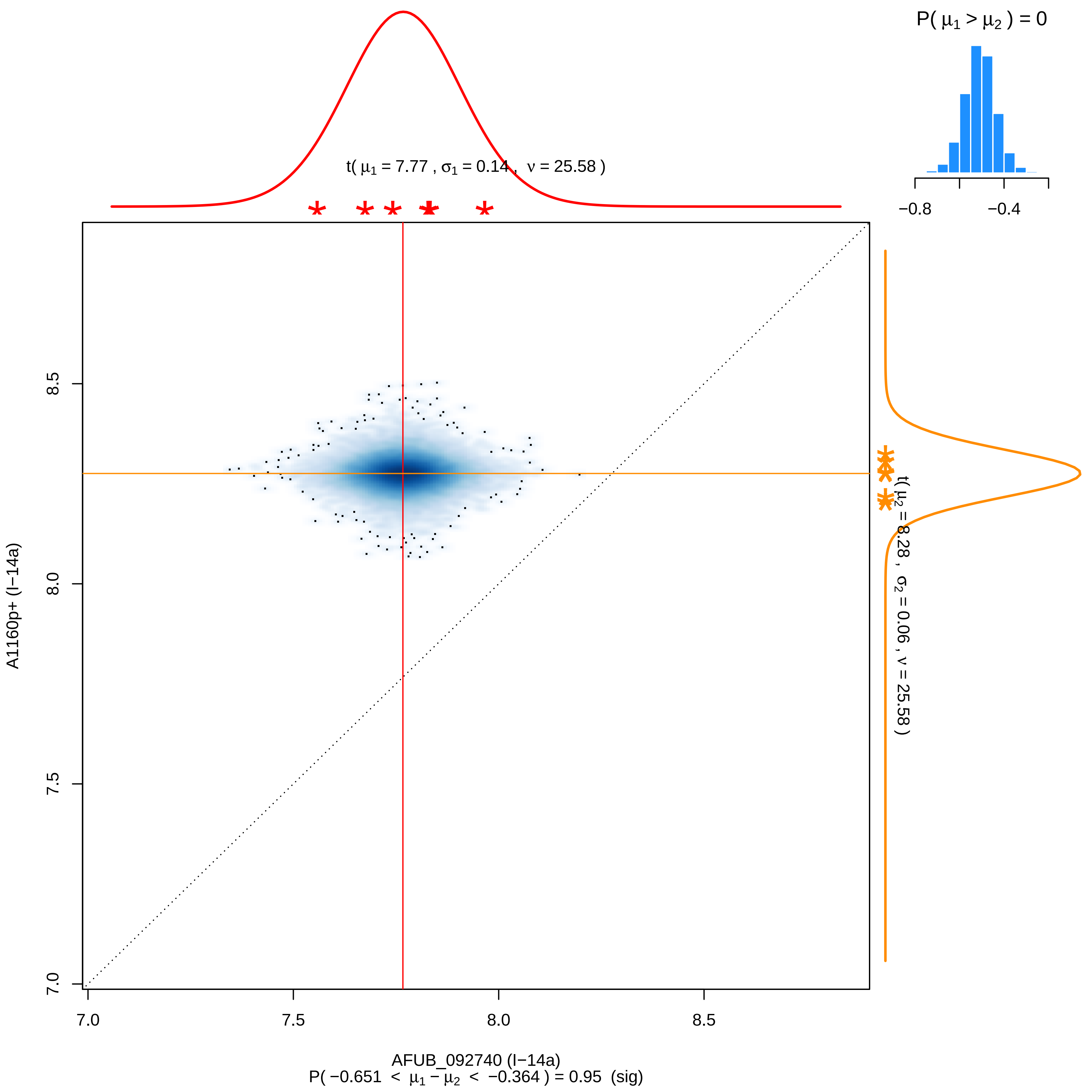
P( μ1 > μ2 ) = 0
−0.8
−0.4
t( μ1 = 7.77 , σ1 = 0.14 , ν = 25.58 )
8.5
8.0
7.5
7.0
A1160p+ (I−14a)
t( μ2 = 8.28 , σ2 = 0.06 , ν = 25.58 )
7.0
7.5
8.0
8.5
AFUB_092740 (I−14a)
P( −0.651 < μ1 − μ2 < −0.364 ) = 0.95 (sig)

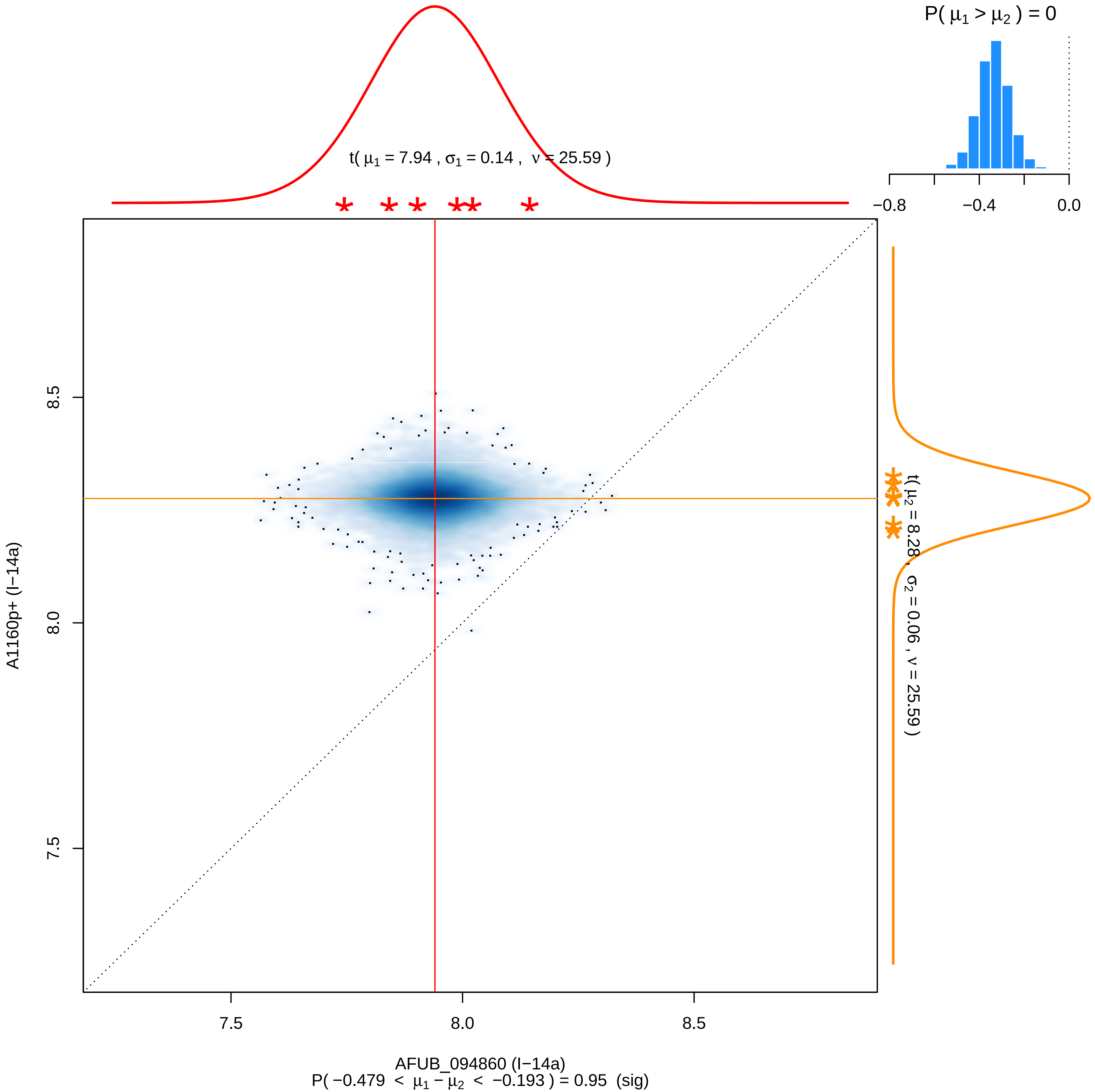

P( $\mu_1 > \mu_2$ ) = 0
−0.8
−0.4
0.0
t( $\mu_1$ = 7.94 , $\sigma_1$ = 0.14 , $\nu$ = 25.59 )
t( $\mu_2$ = 8.28 , $\sigma_2$ = 0.06 , $\nu$ = 25.59 )
A1160p+ (I−14a)
8.5
8.0
7.5
AFUB_094860 (I−14a)
P( −0.479 < $\mu_1 - \mu_2$ < −0.193 ) = 0.95 (sig)

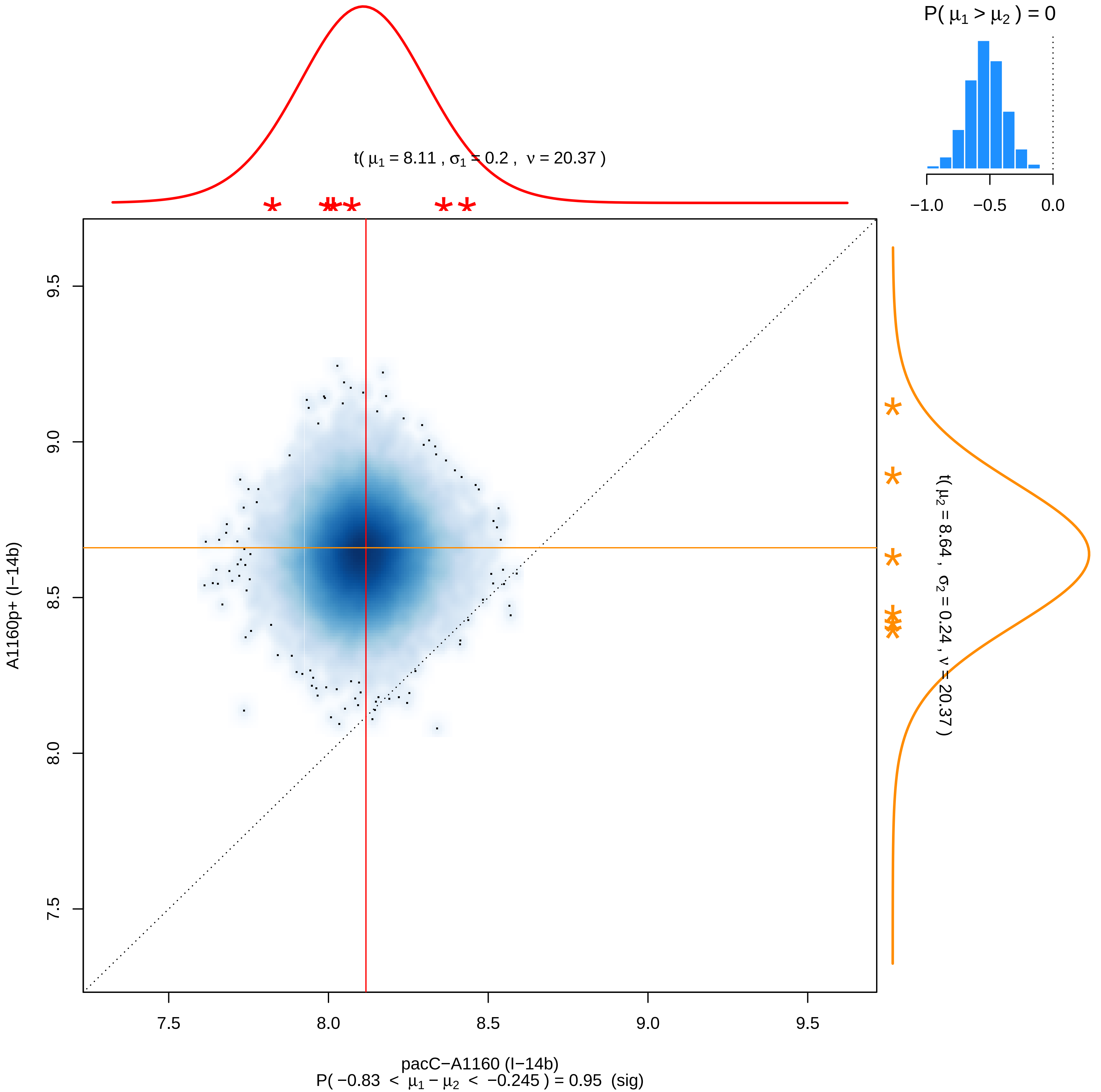

P( μ1 > μ2 ) = 0
−1.0
−0.5
0.0
t( μ1 = 8.11 , σ1 = 0.2 , ν = 20.37 )
t( μ2 = 8.64 , σ2 = 0.24 , ν = 20.37 )
A1160p+ (I−14b)
9.5
9.0
8.5
8.0
7.5
pacC−A1160 (I−14b)
P( −0.83 < μ1 − μ2 < −0.245 ) = 0.95 (sig)

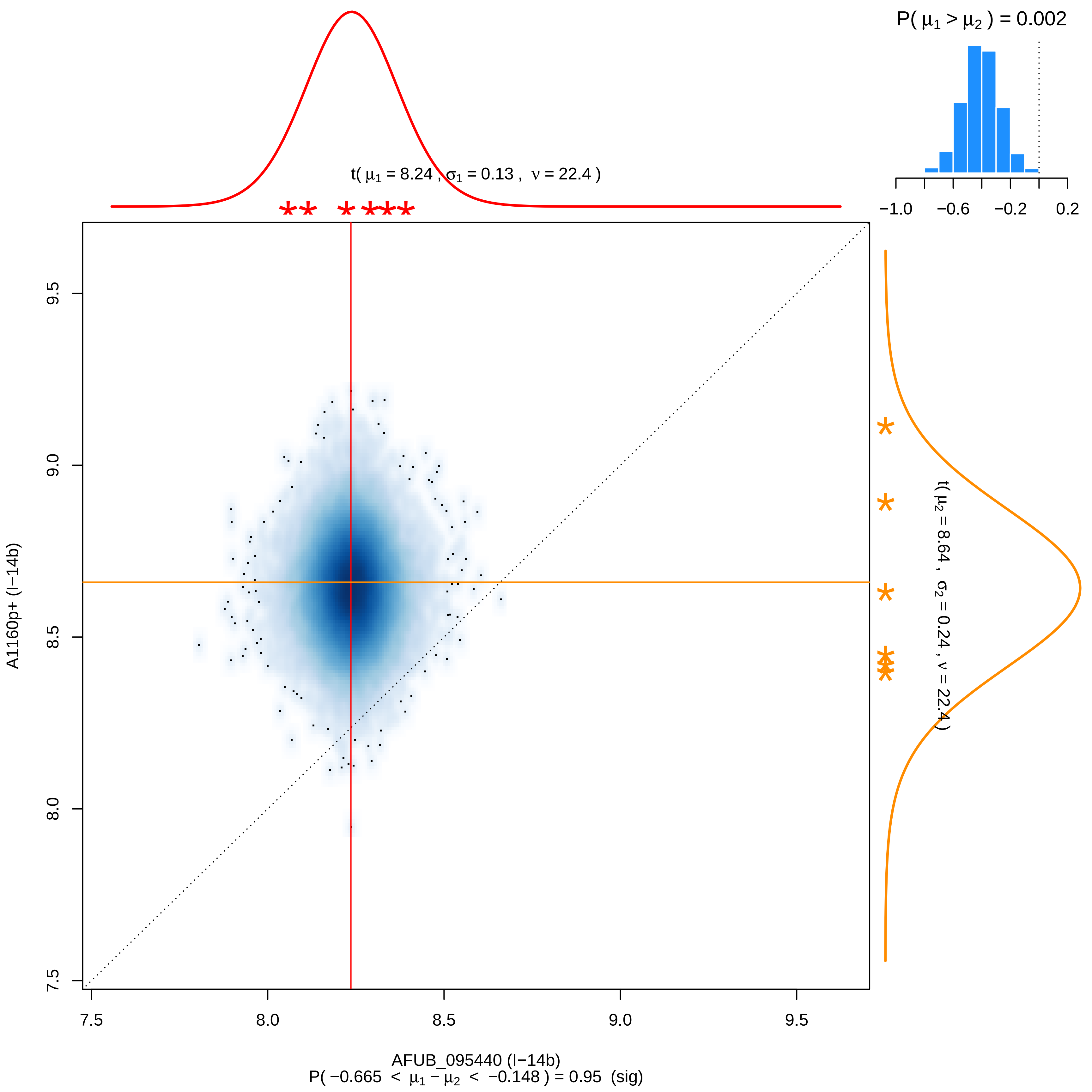

P( μ1 > μ2 ) = 0.002
−1.0 −0.6 −0.2 0.2
t( μ1 = 8.24 , σ1 = 0.13 , ν = 22.4 )
t( μ2 = 8.64 , σ2 = 0.24 , ν = 22.4 )
9.5
9.0
8.5
8.0
7.5
A1160p+ (I−14b)
7.5 8.0 8.5 9.0 9.5
AFUB_095440 (I−14b)
P( −0.665 < μ1 − μ2 < −0.148 ) = 0.95 (sig)

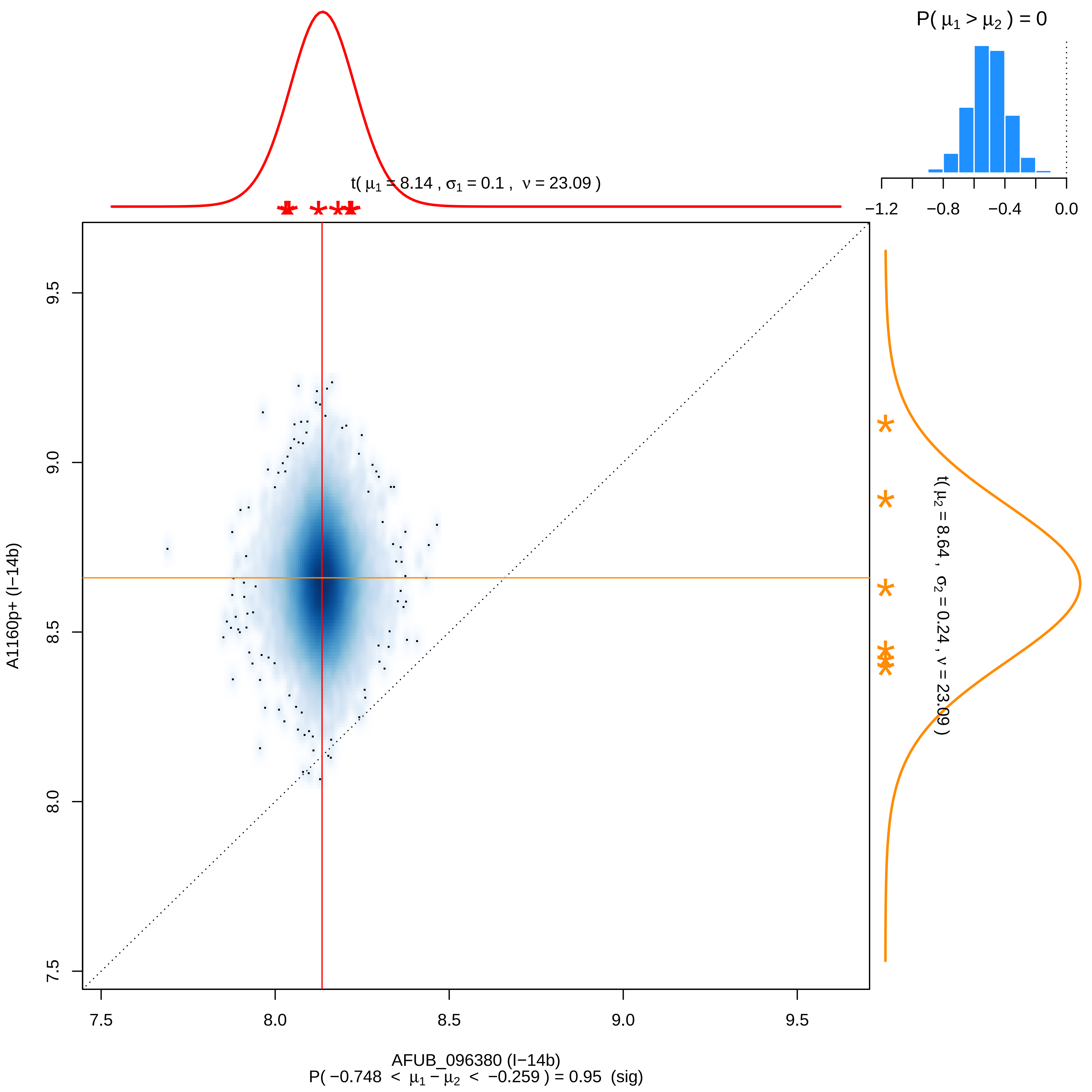

P( μ1 > μ2 ) = 0
−1.2
−0.8
−0.4
0.0
t( μ1 = 8.14 , σ1 = 0.1 , ν = 23.09 )
t( μ2 = 8.64 , σ2 = 0.24 , ν = 23.09 )
A1160p+ (I−14b)
9.5
9.0
8.5
8.0
7.5
7.5
8.0
8.5
9.0
9.5
AFUB_096380 (I−14b)
P( −0.748 < μ1 − μ2 < −0.259 ) = 0.95 (sig)

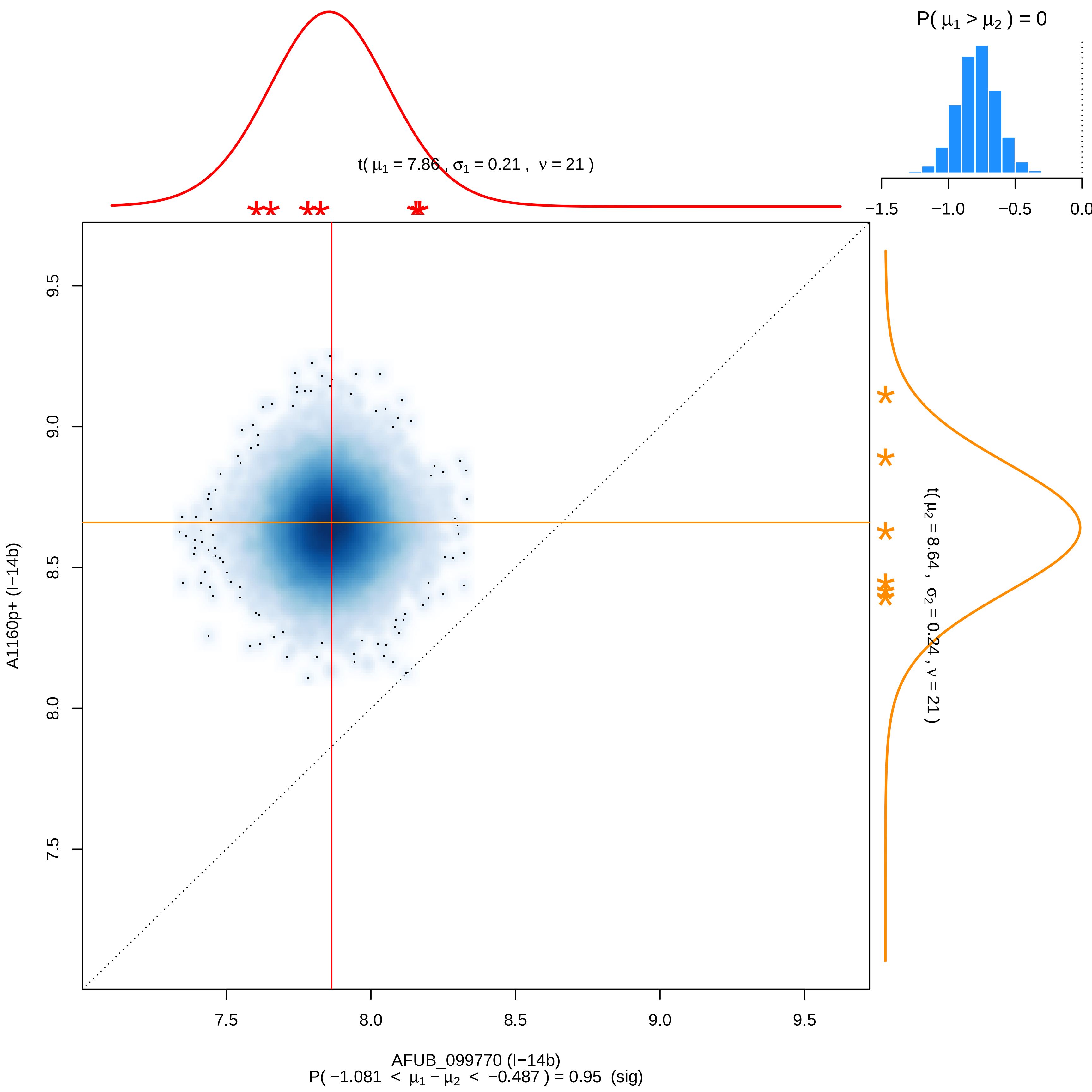

P( μ1 > μ2 ) = 0
−1.5
−1.0
−0.5
0.0
t( μ1 = 7.86 , σ1 = 0.21 , ν = 21 )
t( μ2 = 8.64 , σ2 = 0.24 , ν = 21 )
9.5
9.0
8.5
8.0
7.5
A1160p+ (I−14b)
7.5
8.0
8.5
9.0
9.5
AFUB_099770 (I−14b)
P( −1.081 < μ1 − μ2 < −0.487 ) = 0.95 (sig)

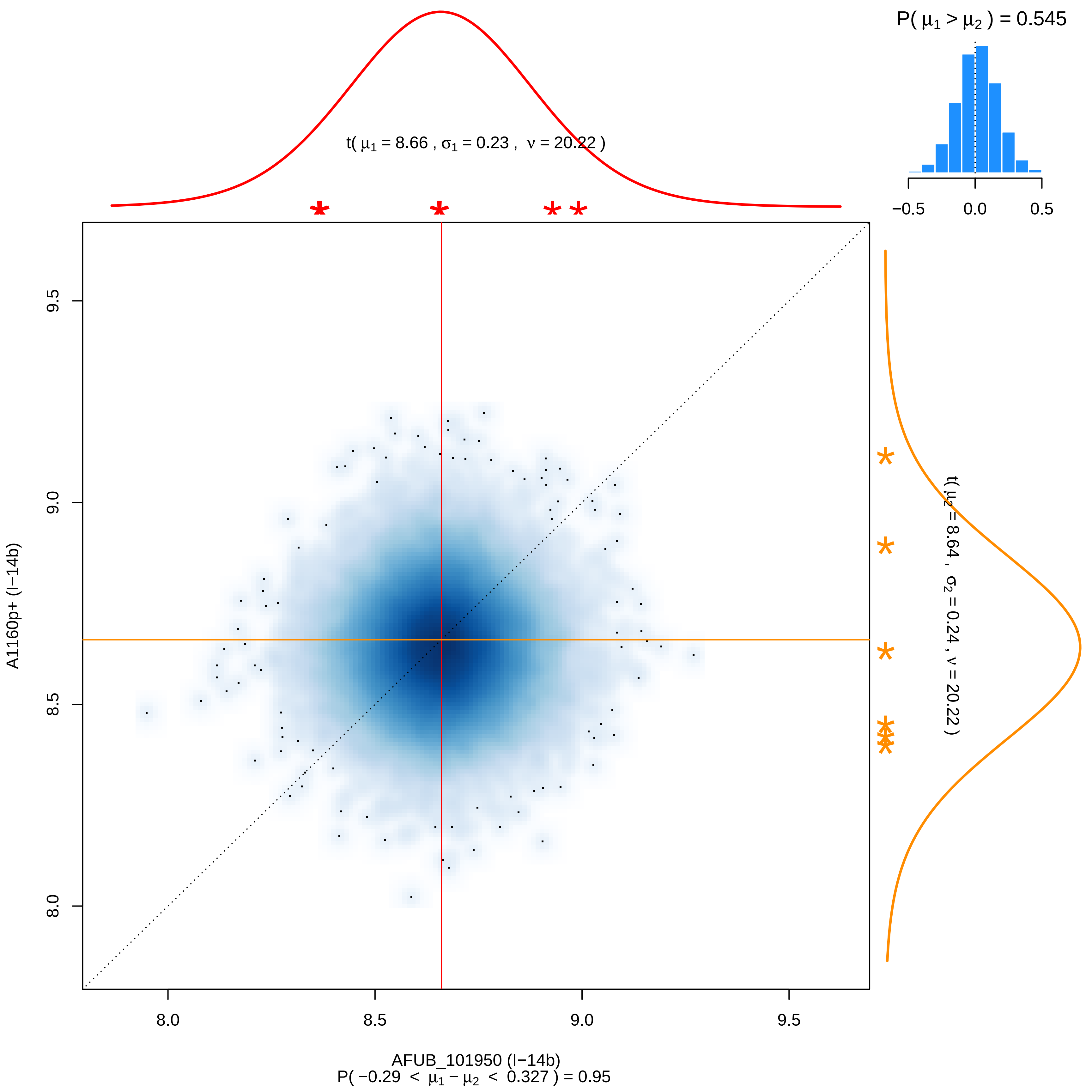

P( μ1 > μ2 ) = 0.545
−0.5
0.0
0.5
t( μ1 = 8.66 , σ1 = 0.23 , ν = 20.22 )
t( μ2 = 8.64 , σ2 = 0.24 , ν = 20.22 )
9.5
9.0
8.5
8.0
A1160p+ (I−14b)
8.0
8.5
9.0
9.5
AFUB_101950 (I−14b)
P( −0.29 < μ1 − μ2 < 0.327 ) = 0.95

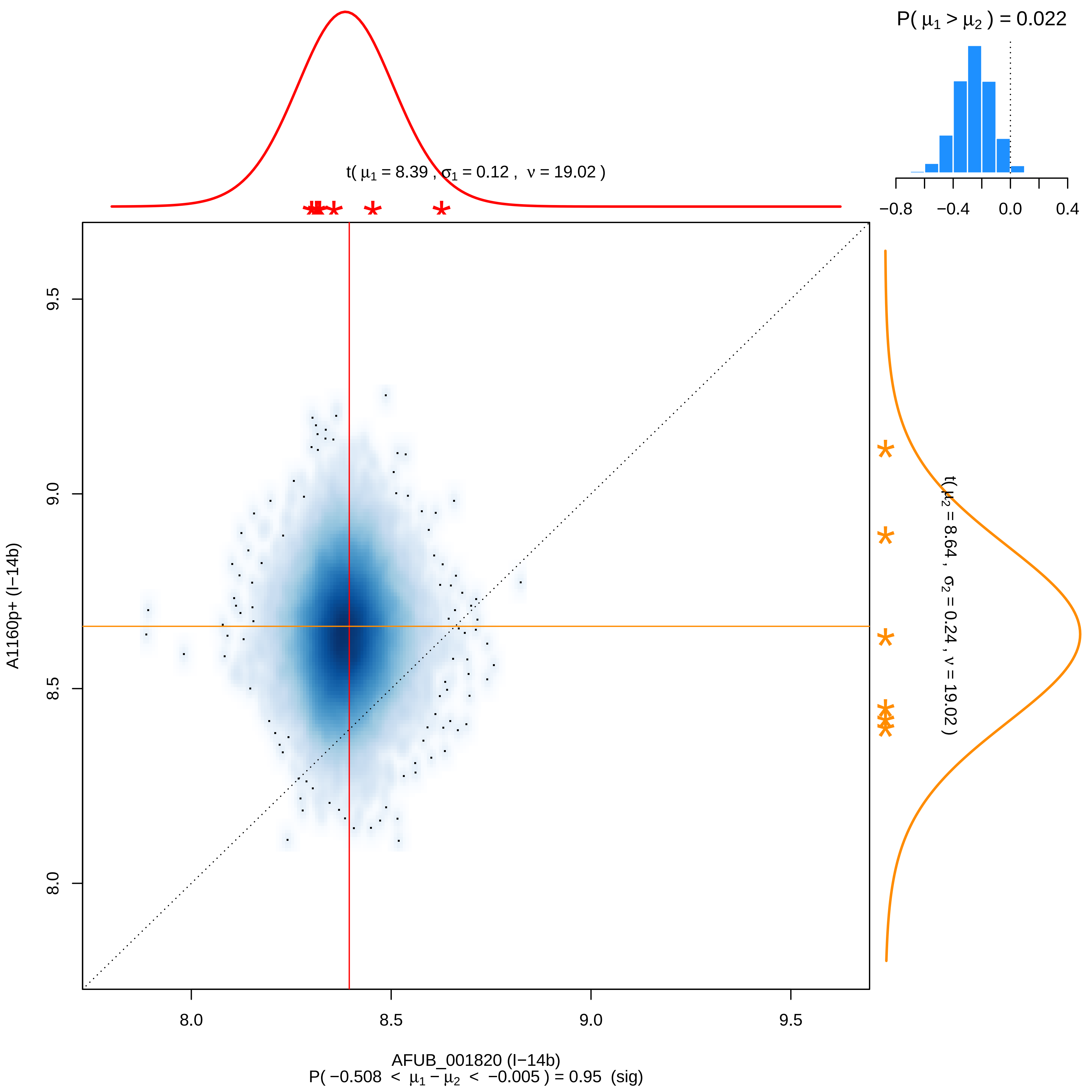

P( μ1 > μ2 ) = 0.022
−0.8
−0.4
0.0
0.4
t( μ1 = 8.39 , σ1 = 0.12 , ν = 19.02 )
t( μ2 = 8.64 , σ2 = 0.24 , ν = 19.02 )
9.5
9.0
8.5
8.0
A1160p+ (I−14b)
8.0
8.5
9.0
9.5
AFUB_001820 (I−14b)
P( −0.508 < μ1 − μ2 < −0.005 ) = 0.95 (sig)

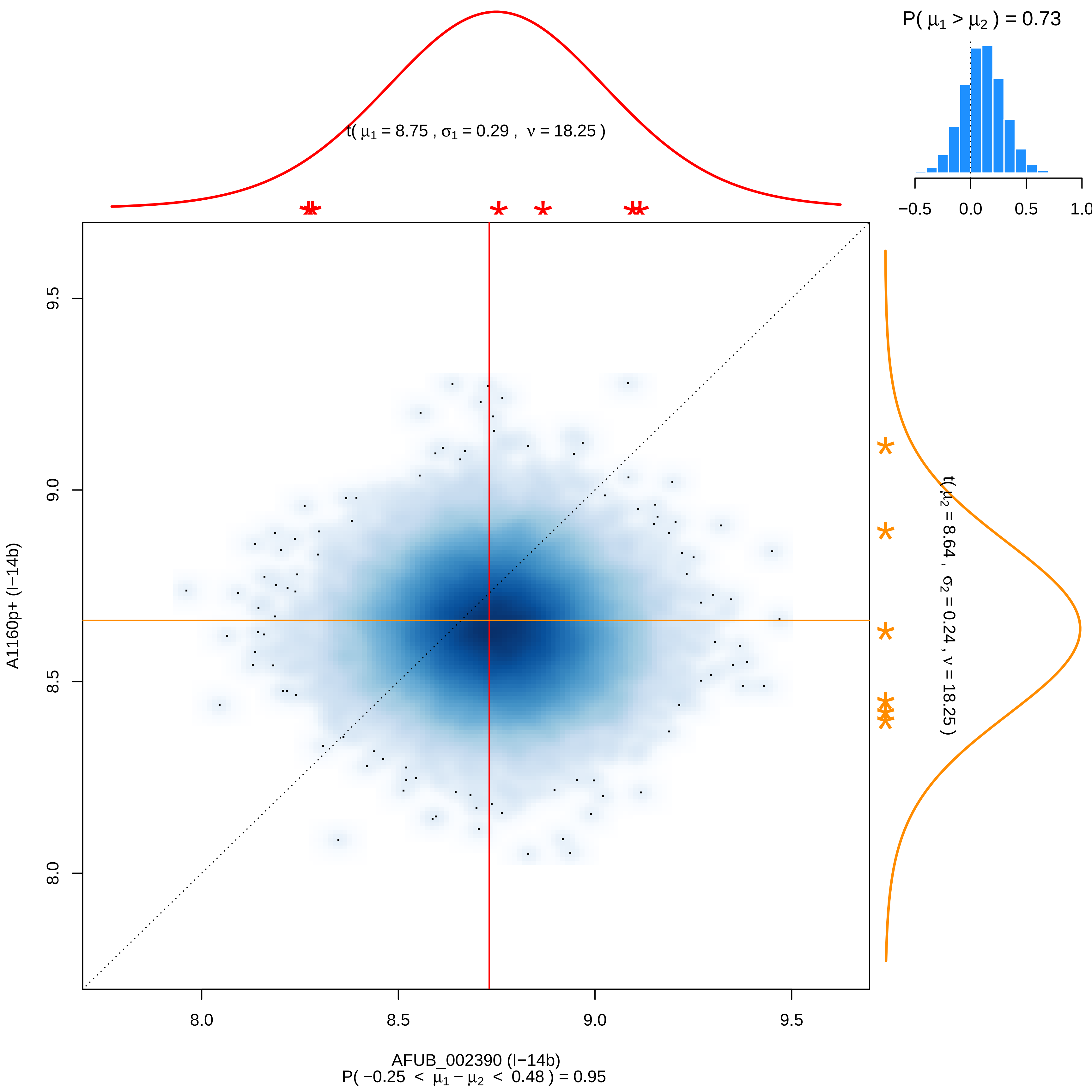
P( μ1 > μ2 ) = 0.73
−0.5 0.0 0.5 1.0
t( μ1 = 8.75 , σ1 = 0.29 , ν = 18.25 )
t( μ2 = 8.64 , σ2 = 0.24 , ν = 18.25 )
9.5
9.0
8.5
8.0
A1160p+ (I−14b)
8.0 8.5 9.0 9.5
AFUB_002390 (I−14b)
P( −0.25 < μ1 − μ2 < 0.48 ) = 0.95

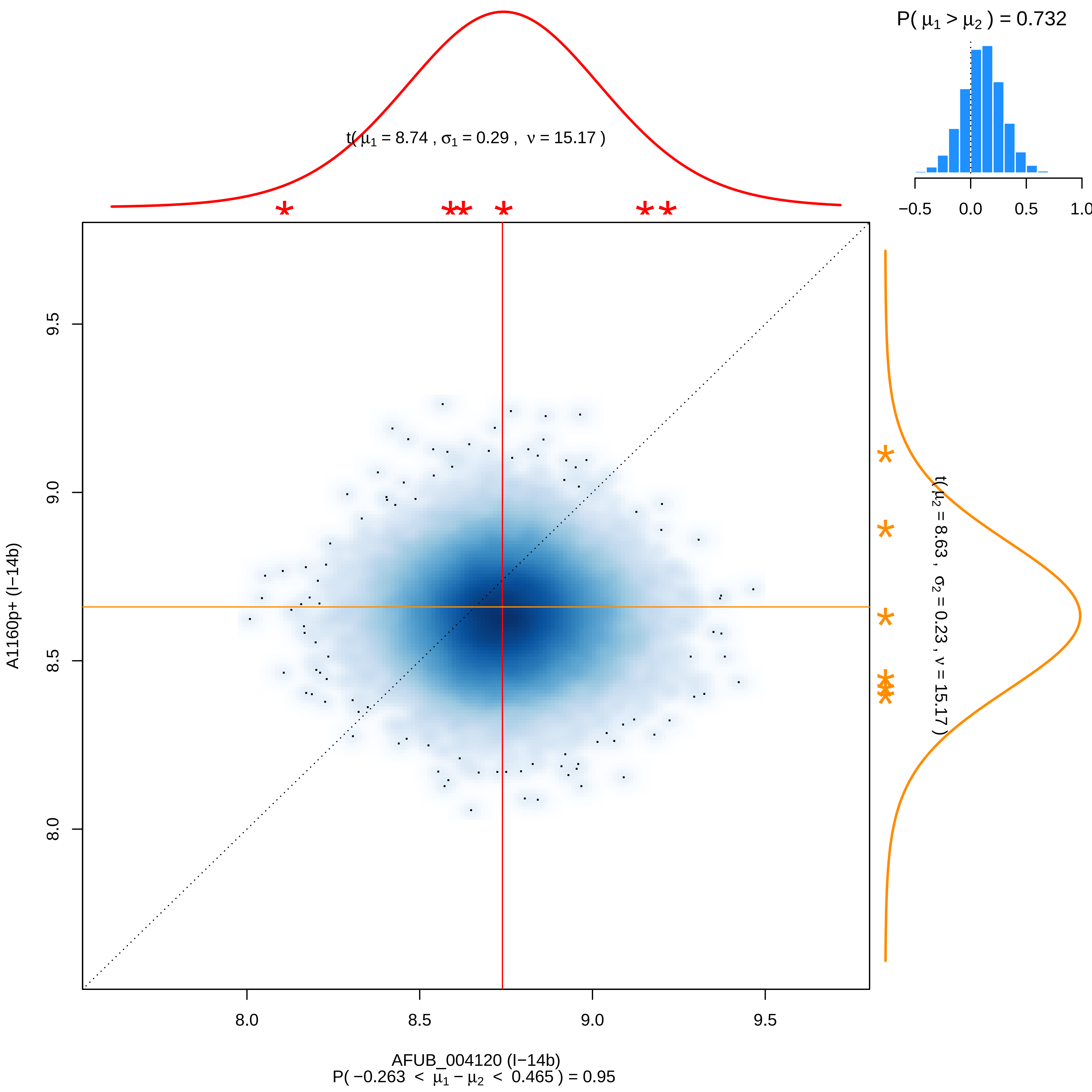

P( μ1 > μ2 ) = 0.732
−0.5
0.0
0.5
1.0
t( μ1 = 8.74 , σ1 = 0.29 , ν = 15.17 )
t( μ2 = 8.63 , σ2 = 0.23 , ν = 15.17 )
9.5
9.0
8.5
8.0
A1160p+ (I−14b)
AFUB_004120 (I−14b)
P( −0.263 < μ1 − μ2 < 0.465 ) = 0.95

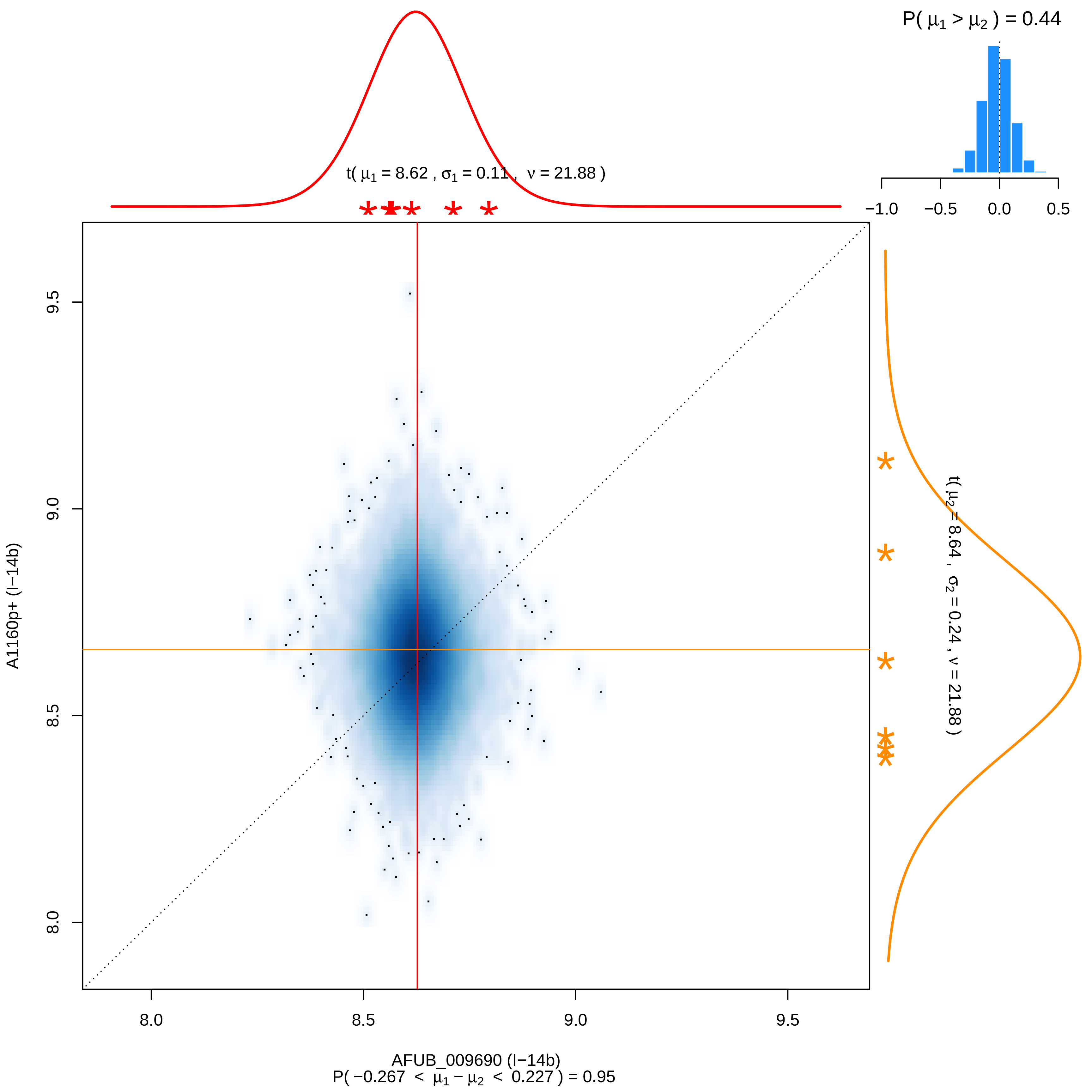

P( μ1 > μ2 ) = 0.44
−1.0
−0.5
0.0
0.5
t( μ1 = 8.62 , σ1 = 0.11 , ν = 21.88 )
t( μ2 = 8.64 , σ2 = 0.24 , ν = 21.88 )
A1160p+ (I−14b)
9.5
9.0
8.5
8.0
8.0
8.5
9.0
9.5
AFUB_009690 (I−14b)
P( −0.267 < μ1 − μ2 < 0.227 ) = 0.95

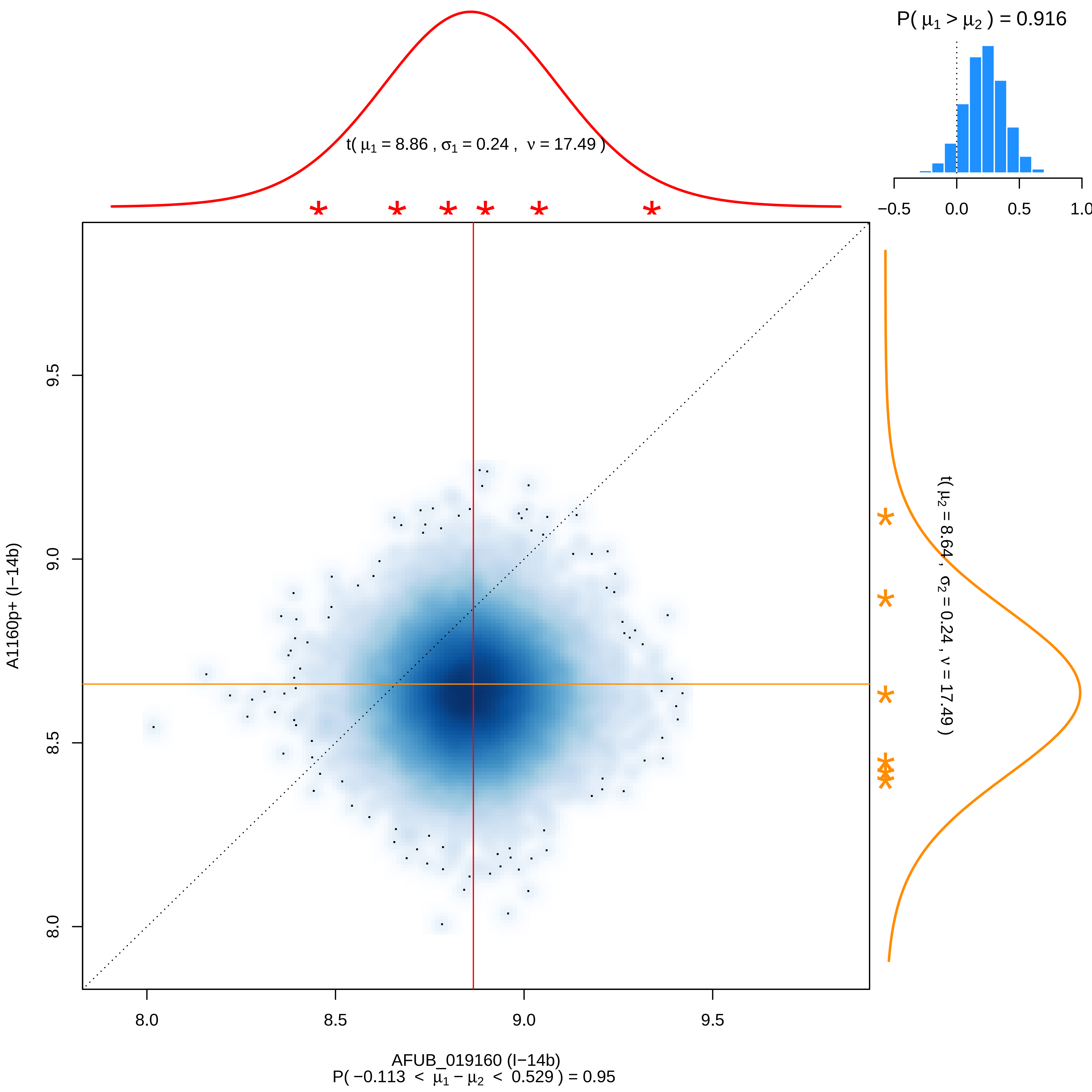
P( μ1 > μ2 ) = 0.916
−0.5
0.0
0.5
1.0
t( μ1 = 8.86 , σ1 = 0.24 , ν = 17.49 )
9.5
9.0
8.5
8.0
A1160p+ (I−14b)
t( μ2 = 8.64 , σ2 = 0.24 , ν = 17.49 )
8.0
8.5
9.0
9.5
AFUB_019160 (I−14b)
P( −0.113 < μ1 − μ2 < 0.529 ) = 0.95

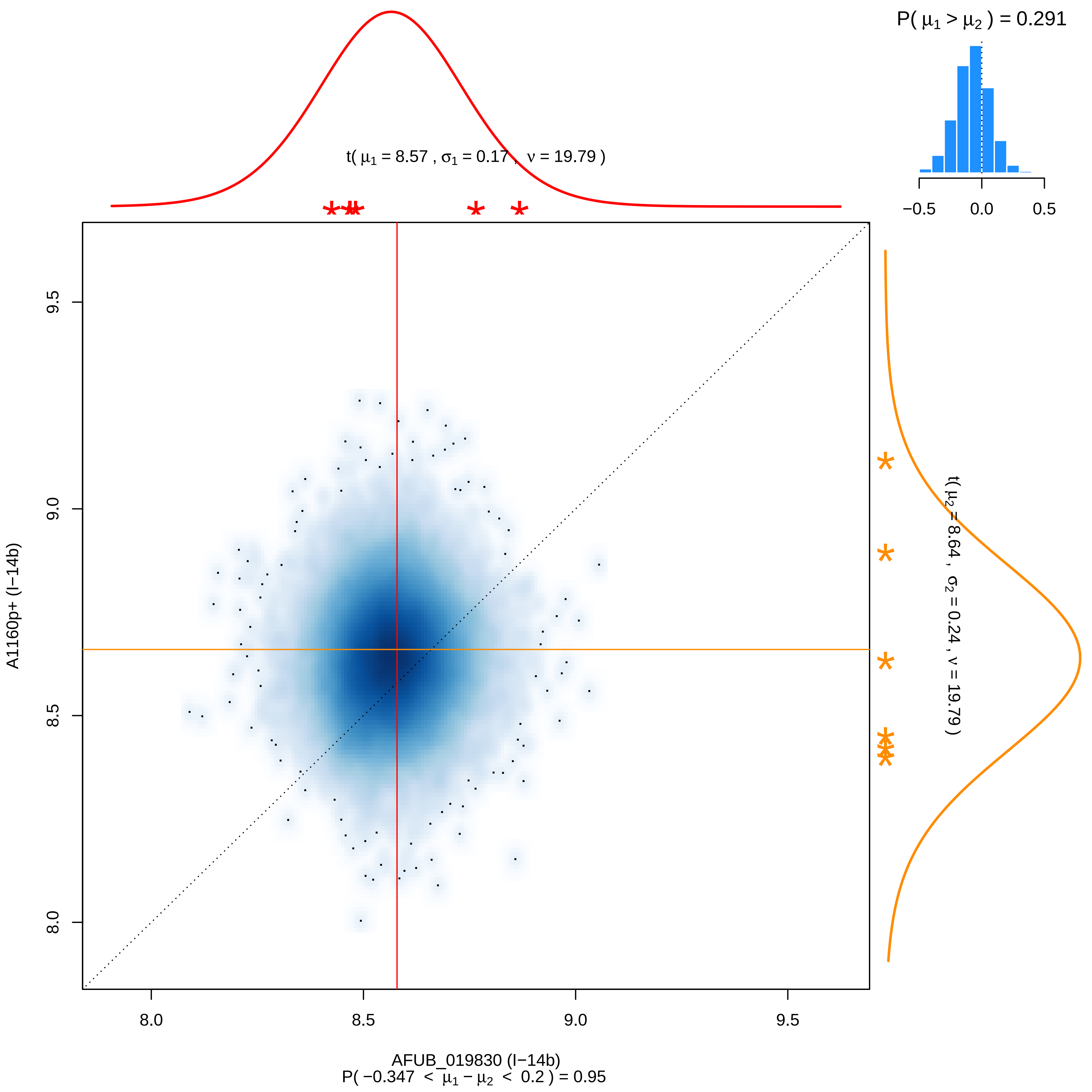

P( μ1 > μ2 ) = 0.291
−0.5
0.0
0.5
t( μ1 = 8.57 , σ1 = 0.17 , ν = 19.79 )
t( μ2 = 8.64 , σ2 = 0.24 , ν = 19.79 )
9.5
9.0
8.5
8.0
A1160p+ (I−14b)
8.0
8.5
9.0
9.5
AFUB_019830 (I−14b)
P( −0.347 < μ1 − μ2 < 0.2 ) = 0.95

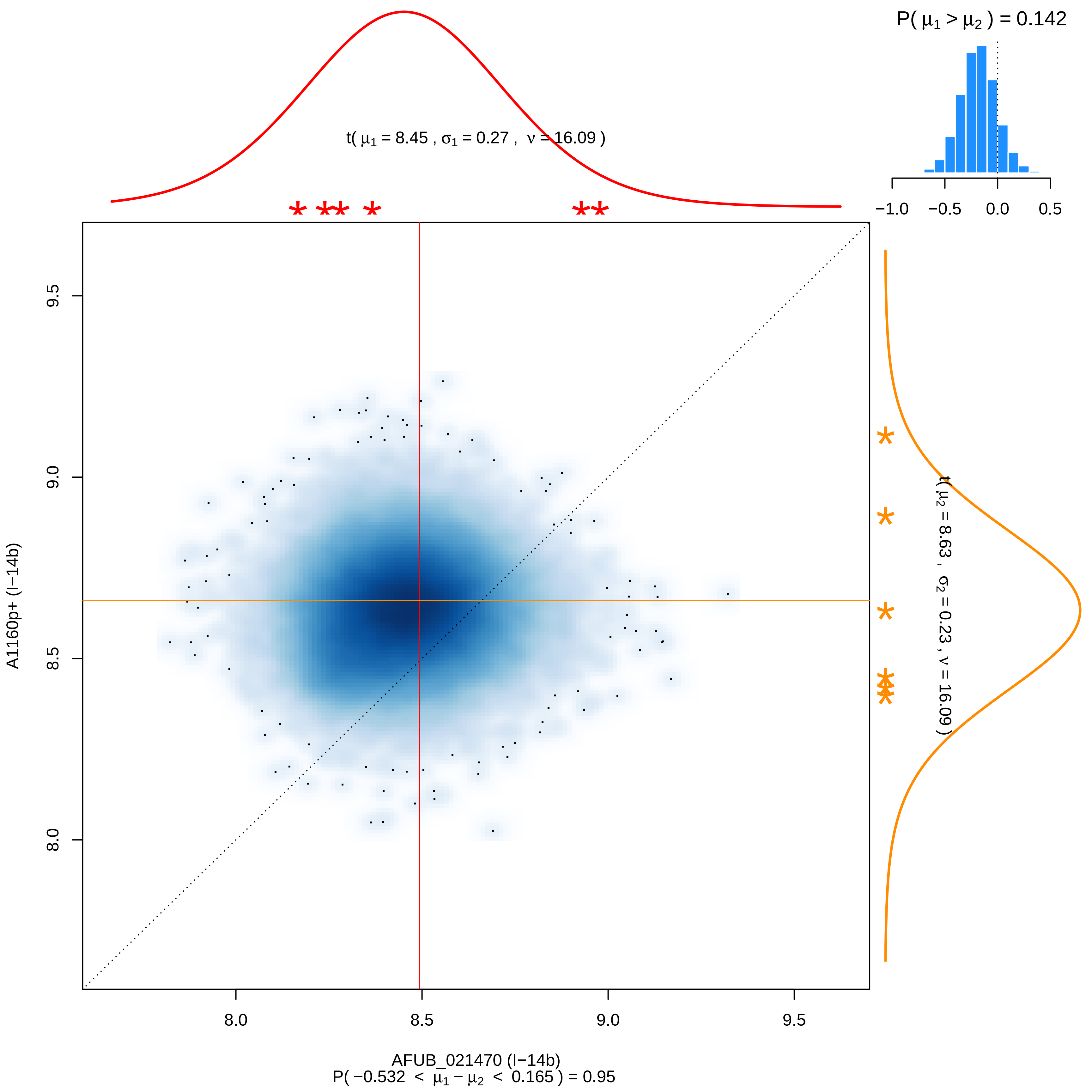

P( μ1 > μ2 ) = 0.142
−1.0 −0.5 0.0 0.5
t( μ1 = 8.45 , σ1 = 0.27 , ν = 16.09 )
t( μ2 = 8.63 , σ2 = 0.23 , ν = 16.09 )
9.5
9.0
8.5
8.0
A1160p+ (I−14b)
8.0 8.5 9.0 9.5
AFUB_021470 (I−14b)
P( −0.532 < μ1 − μ2 < 0.165 ) = 0.95

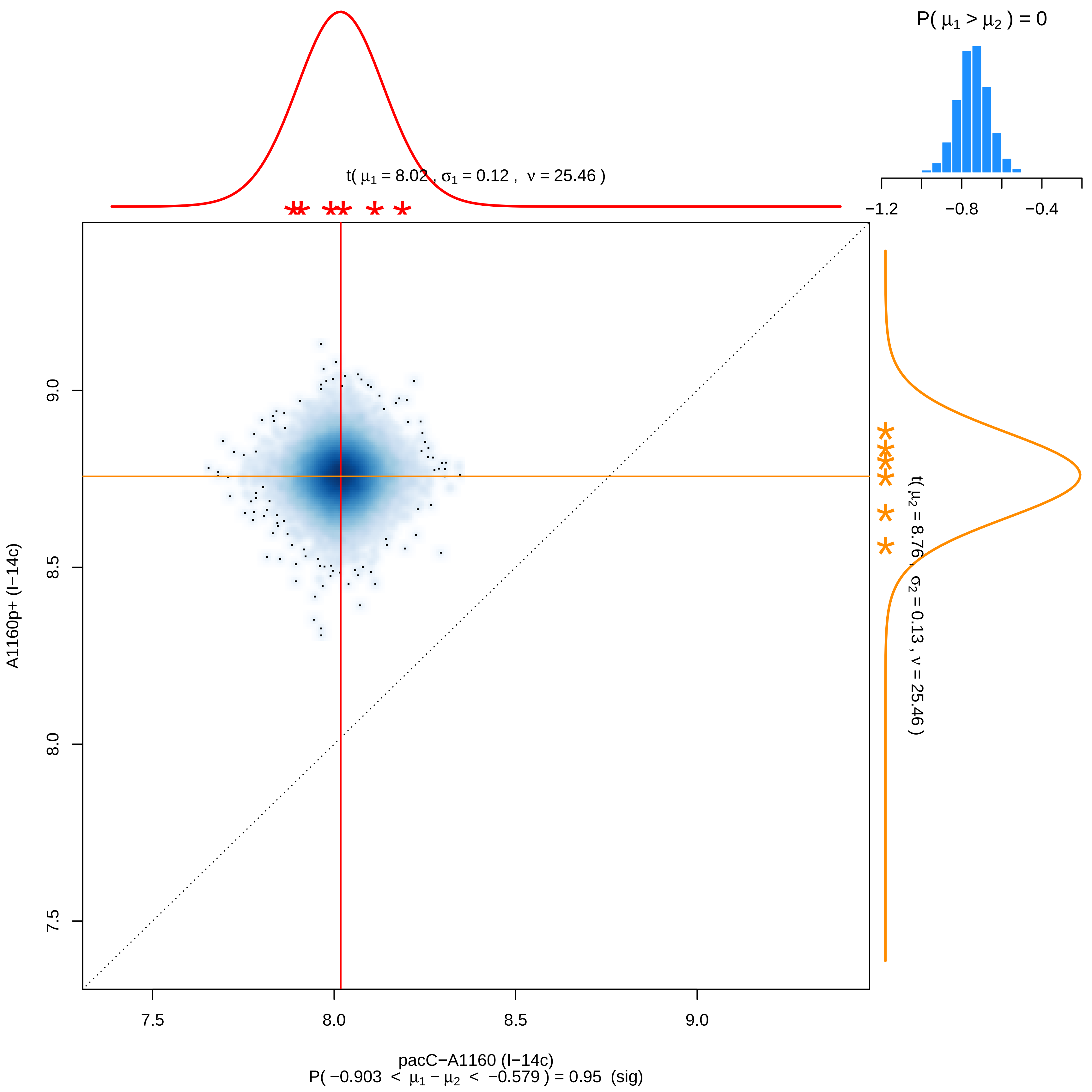

P( $\mu_1 > \mu_2$ ) = 0
−1.2
−0.8
−0.4
t( $\mu_1$ = 8.02 , $\sigma_1$ = 0.12 , $\nu$ = 25.46 )
t( $\mu_2$ = 8.76 , $\sigma_2$ = 0.13 , $\nu$ = 25.46 )
A1160p+ (I−14c)
9.0
8.5
8.0
7.5
7.5
8.0
8.5
9.0
pacC−A1160 (I−14c)
P( −0.903 < $\mu_1 - \mu_2$ < −0.579 ) = 0.95 (sig)

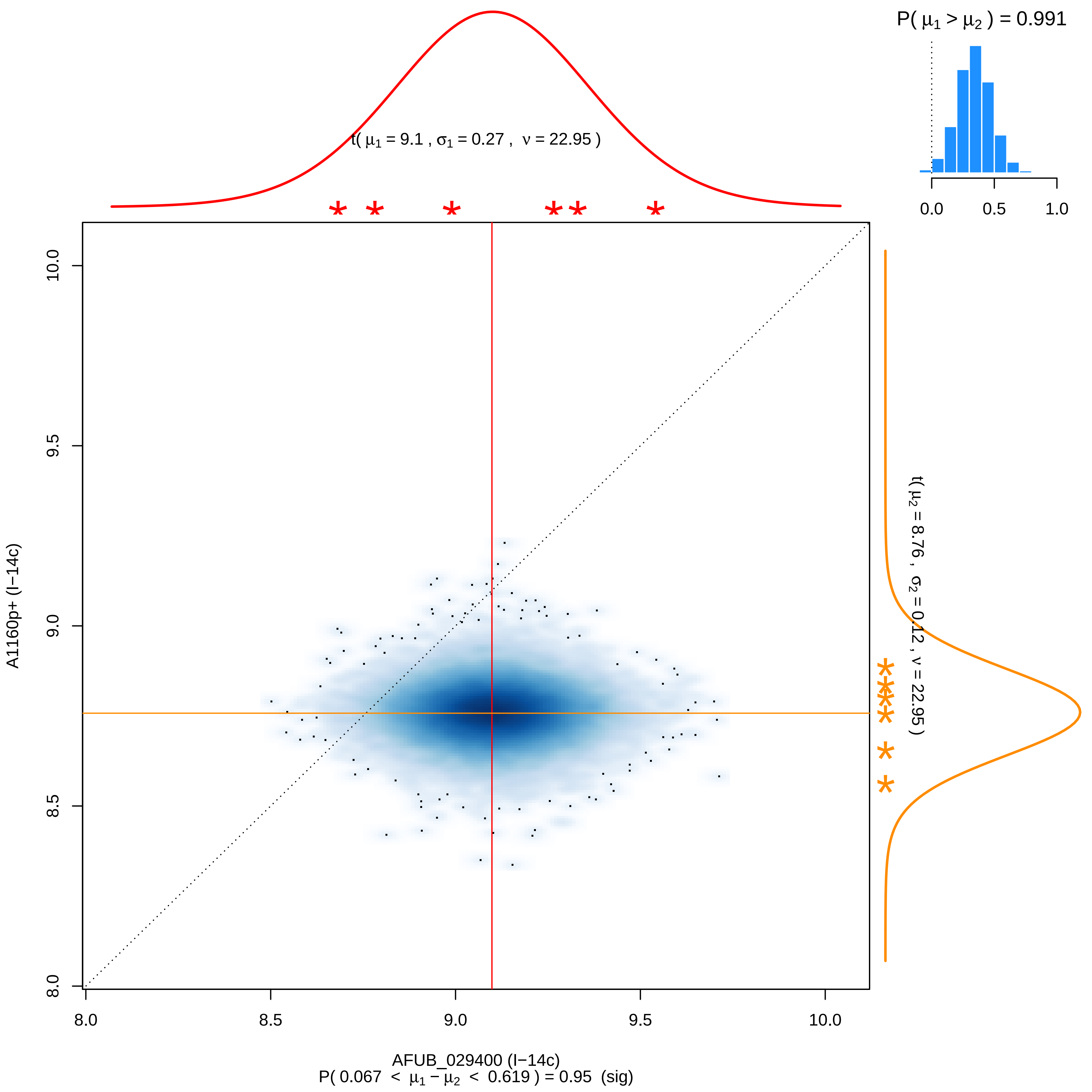

P( $\mu_1 > \mu_2$ ) = 0.991
0.0
0.5
1.0
t( $\mu_1$ = 9.1 , $\sigma_1$ = 0.27 , $\nu$ = 22.95 )
t( $\mu_2$ = 8.76 , $\sigma_2$ = 0.12 , $\nu$ = 22.95 )
A1160p+ (I−14c)
10.0
9.5
9.0
8.5
8.0
8.0
8.5
9.0
9.5
10.0
AFUB_029400 (I−14c)
P( 0.067 < $\mu_1 - \mu_2$ < 0.619 ) = 0.95 (sig)

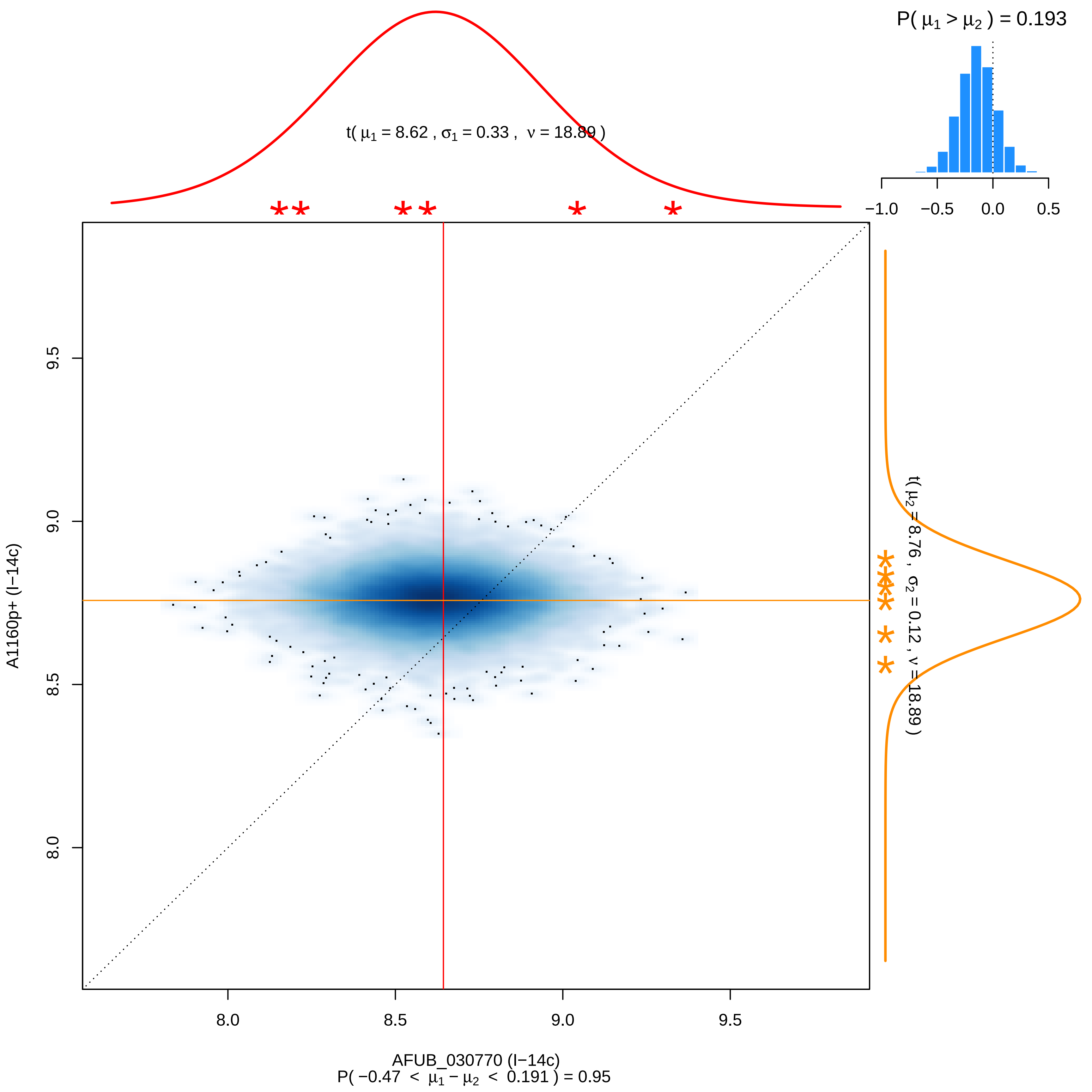

P( μ1 > μ2 ) = 0.193
−1.0
−0.5
0.0
0.5
t( μ1 = 8.62 , σ1 = 0.33 , ν = 18.89 )
t( μ2 = 8.76 , σ2 = 0.12 , ν = 18.89 )
9.5
9.0
8.5
8.0
A1160p+ (I−14c)
8.0
8.5
9.0
9.5
AFUB_030770 (I−14c)
P( −0.47 < μ1 − μ2 < 0.191 ) = 0.95

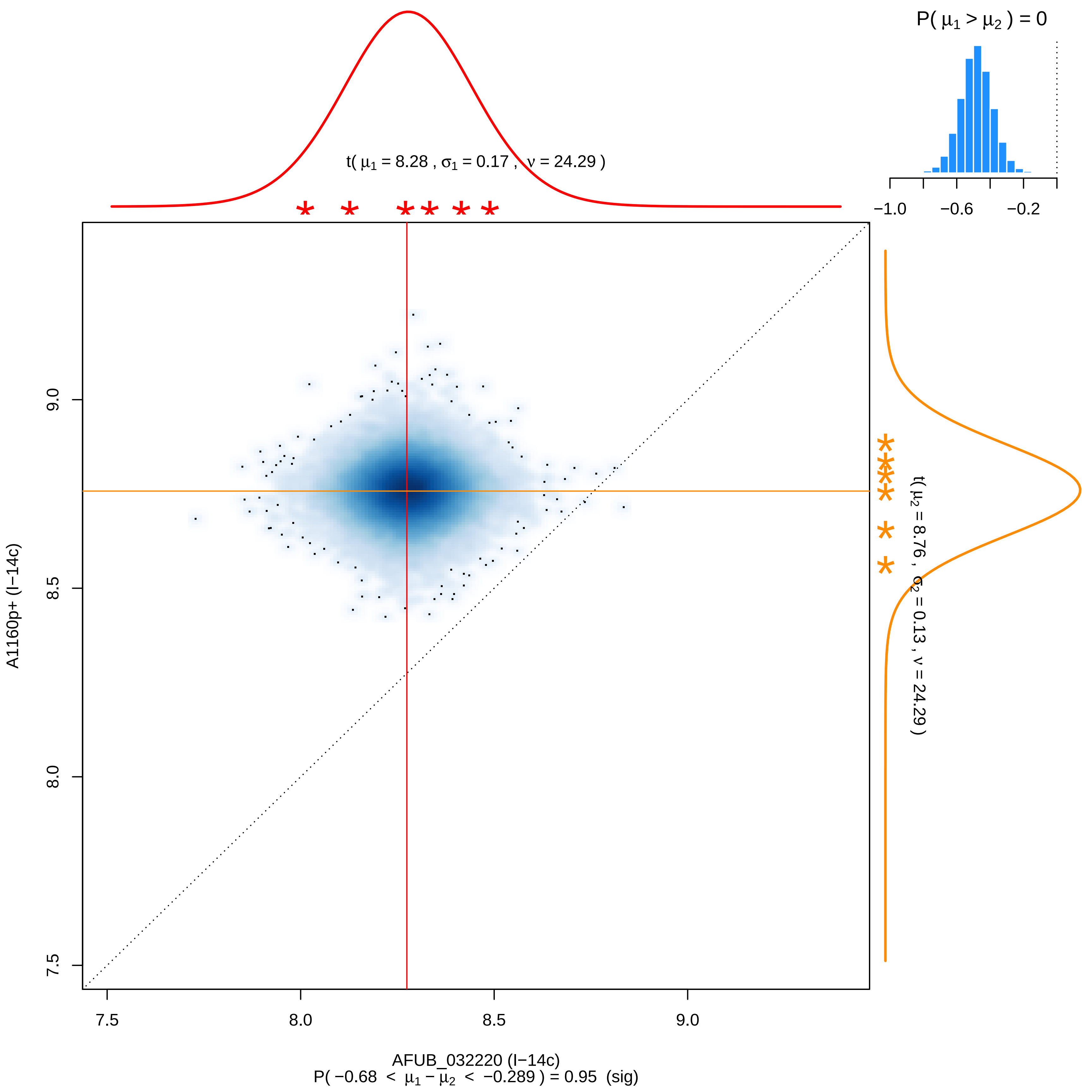

P( μ1 > μ2 ) = 0
−1.0
−0.6
−0.2
t( μ1 = 8.28 , σ1 = 0.17 , ν = 24.29 )
t( μ2 = 8.76 , σ2 = 0.13 , ν = 24.29 )
A1160p+ (I−14c)
9.0
8.5
8.0
7.5
7.5
8.0
8.5
9.0
AFUB_032220 (I−14c)
P( −0.68 < μ1 − μ2 < −0.289 ) = 0.95 (sig)

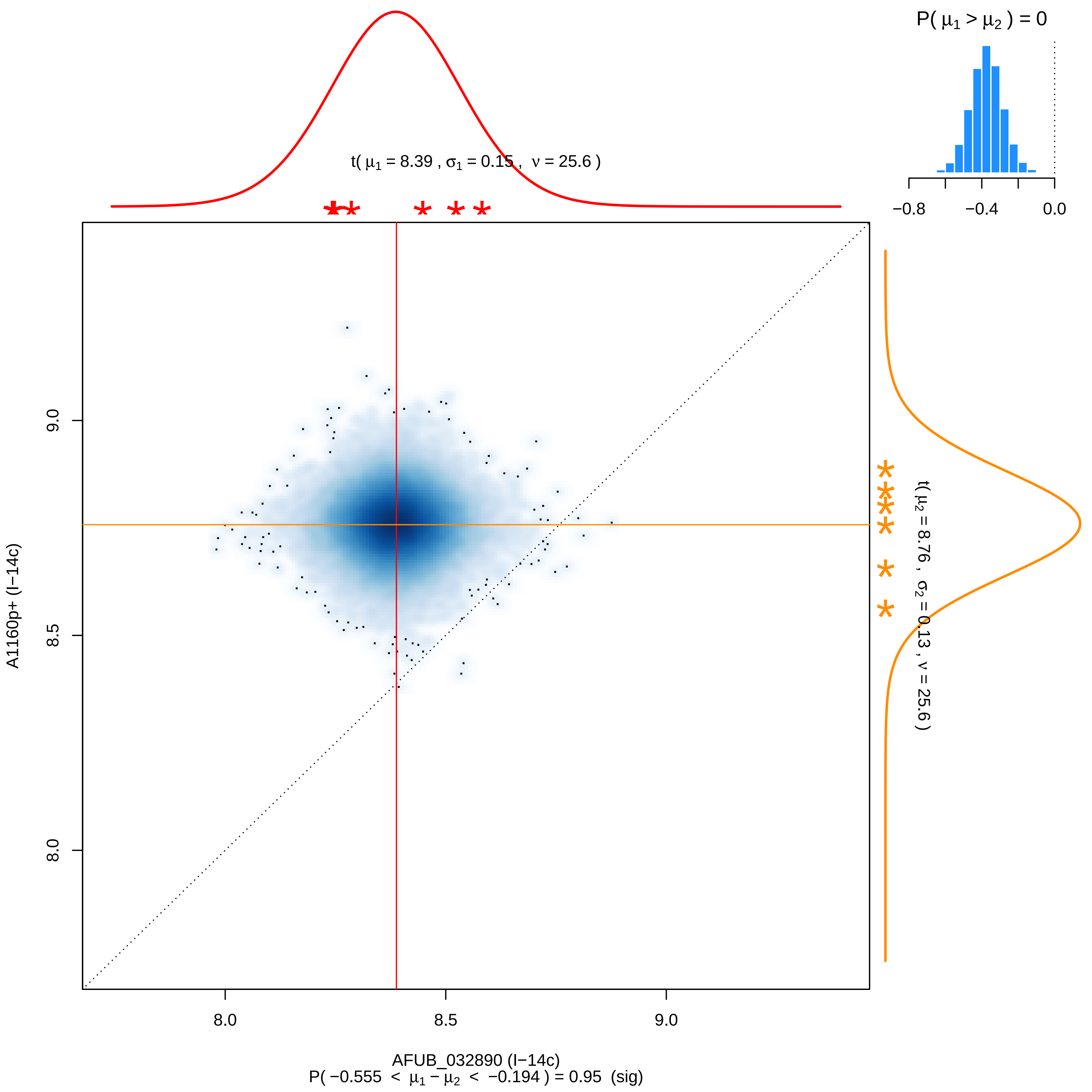

P( μ1 > μ2 ) = 0
−0.8
−0.4
0.0
t( μ1 = 8.39 , σ1 = 0.15 , ν = 25.6 )
t( μ2 = 8.76 , σ2 = 0.13 , ν = 25.6 )
A1160p+ (I−14c)
9.0
8.5
8.0
8.0
8.5
9.0
AFUB_032890 (I−14c)
P( −0.555 < μ1 − μ2 < −0.194 ) = 0.95 (sig)

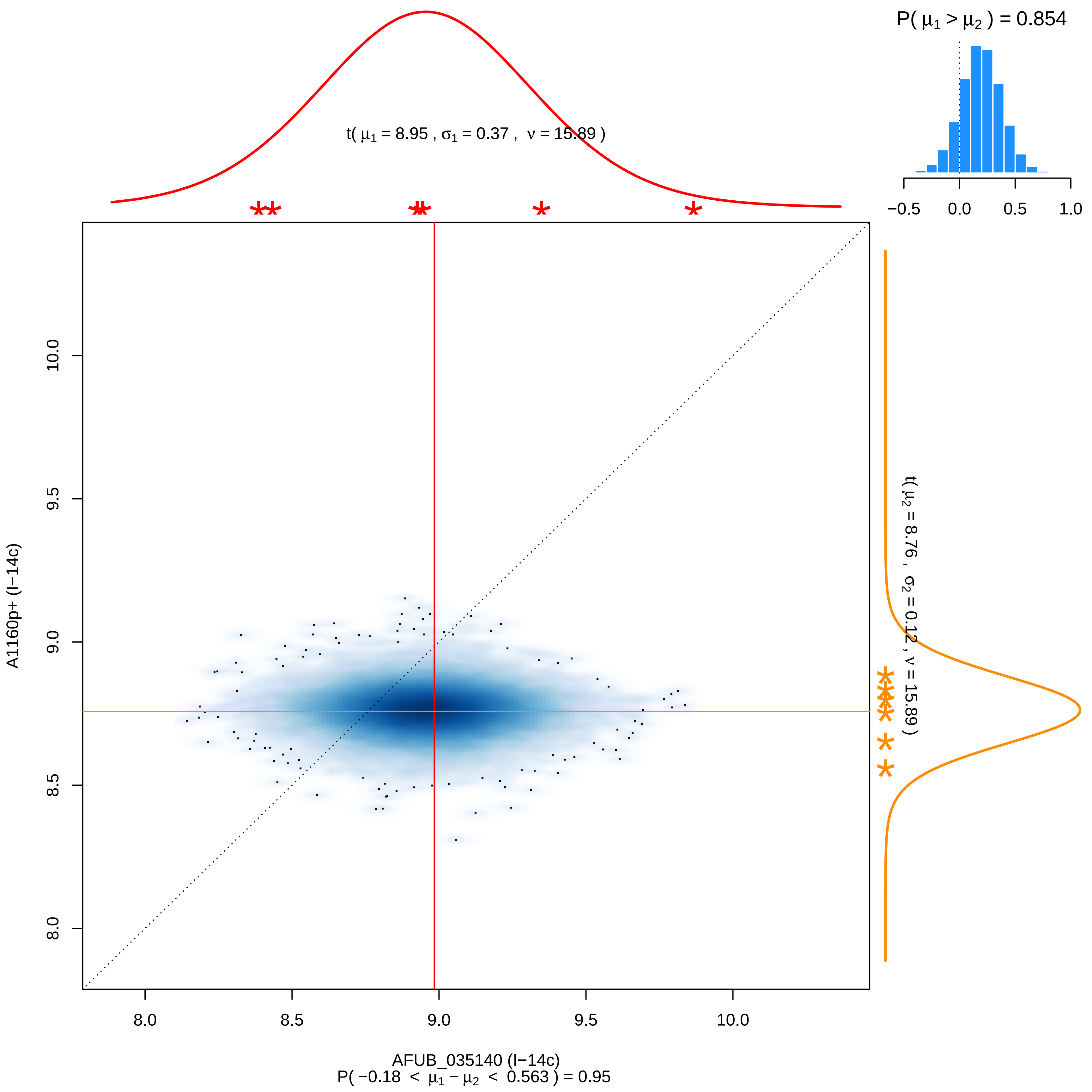

P( μ1 > μ2 ) = 0.854
−0.5
0.0
0.5
1.0
t( μ1 = 8.95 , σ1 = 0.37 , ν = 15.89 )
t( μ2 = 8.76 , σ2 = 0.12 , ν = 15.89 )
10.0
9.5
9.0
8.5
8.0
A1160p+ (I−14c)
8.0
8.5
9.0
9.5
10.0
AFUB_035140 (I−14c)
P( −0.18 < μ1 − μ2 < 0.563 ) = 0.95

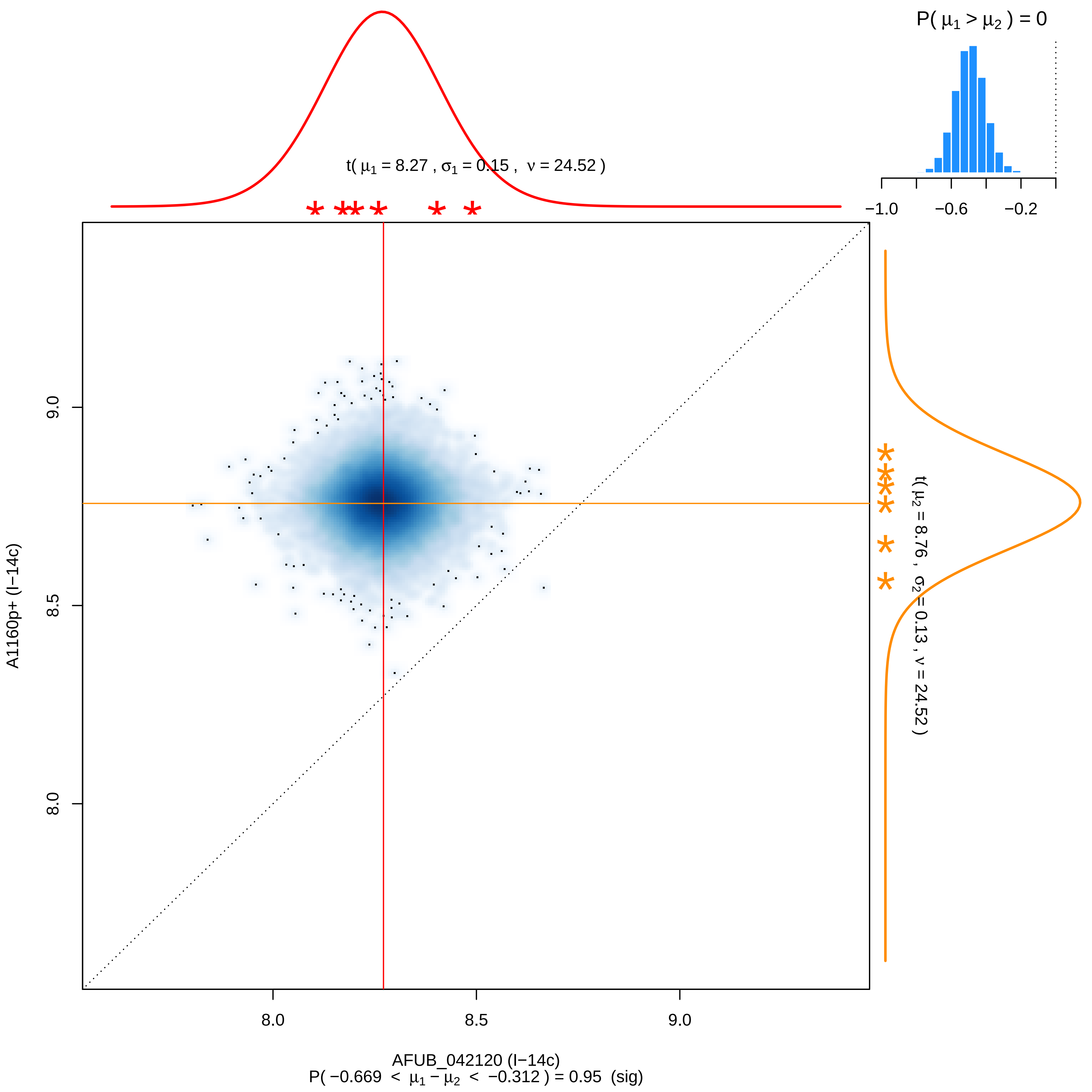

P( μ1 > μ2 ) = 0
−1.0
−0.6
−0.2
t( μ1 = 8.27 , σ1 = 0.15 , ν = 24.52 )
9.0
8.5
8.0
A1160p+ (I−14c)
t( μ2 = 8.76 , σ2 = 0.13 , ν = 24.52 )
8.0
8.5
9.0
AFUB_042120 (I−14c)
P( −0.669 < μ1 − μ2 < −0.312 ) = 0.95 (sig)

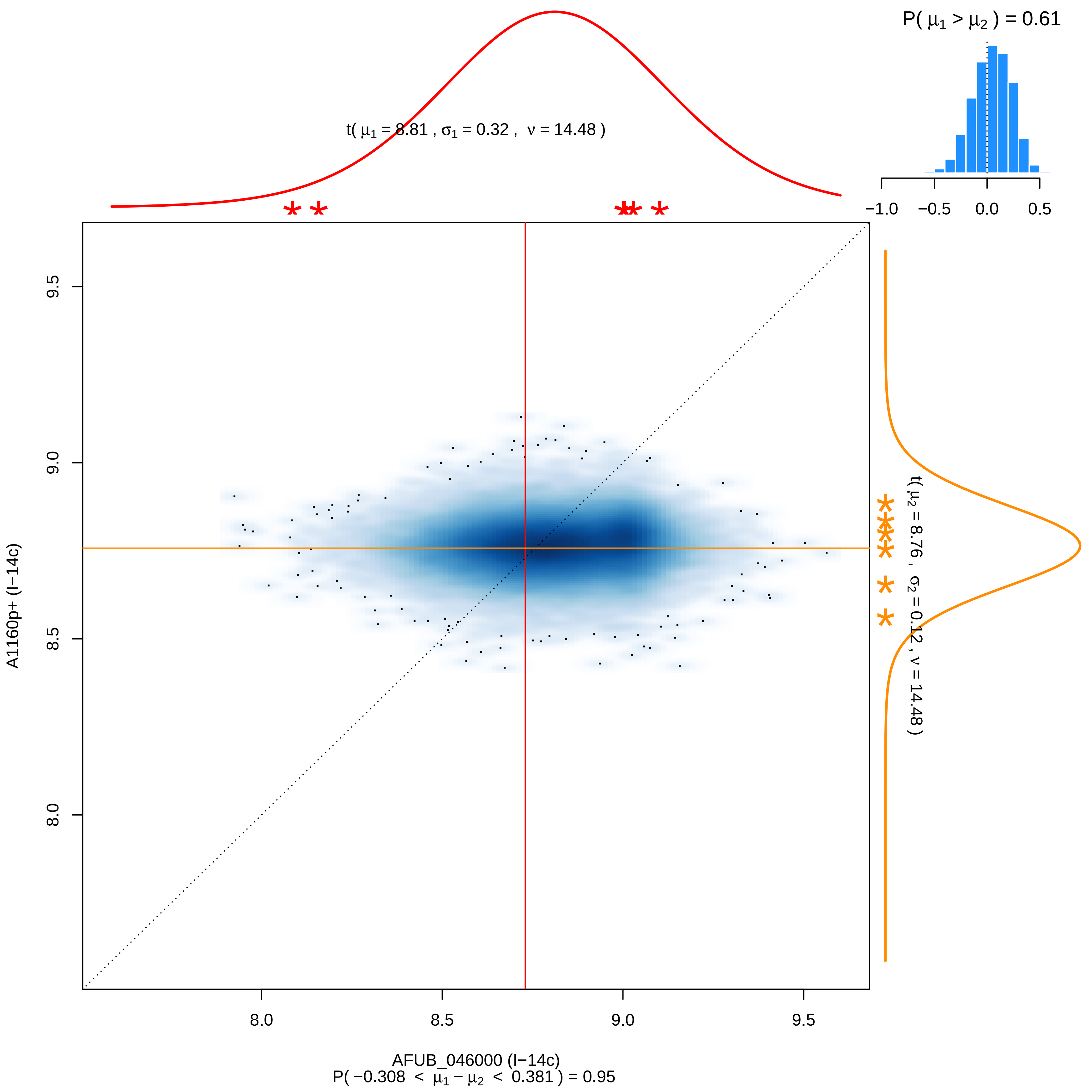

P( μ1 > μ2 ) = 0.61
t( μ1 = 8.81 , σ1 = 0.32 , ν = 14.48 )
−1.0 −0.5 0.0 0.5
9.5
9.0
8.5
8.0
A1160p+ (I−14c)
t( μ2 = 8.76 , σ2 = 0.12 , ν = 14.48 )
8.0 8.5 9.0 9.5
AFUB_046000 (I−14c)
P( −0.308 < μ1 − μ2 < 0.381 ) = 0.95

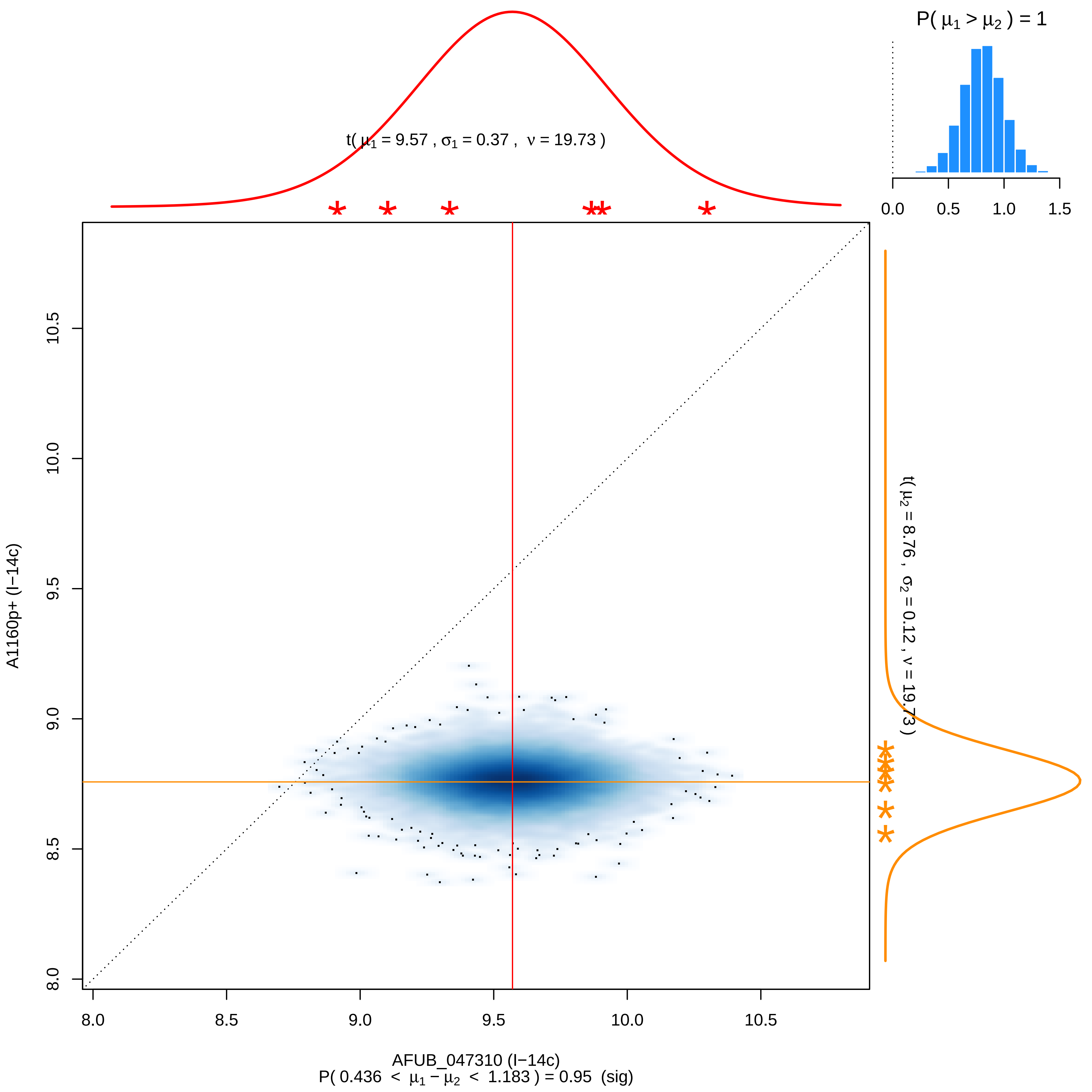
P( μ1 > μ2 ) = 1
0.0
0.5
1.0
1.5
t( μ1 = 9.57 , σ1 = 0.37 , ν = 19.73 )
t( μ2 = 8.76 , σ2 = 0.12 , ν = 19.73 )
A1160p+ (I−14c)
8.0
8.5
9.0
9.5
10.0
10.5
AFUB_047310 (I−14c)
P( 0.436 < μ1 − μ2 < 1.183 ) = 0.95 (sig)

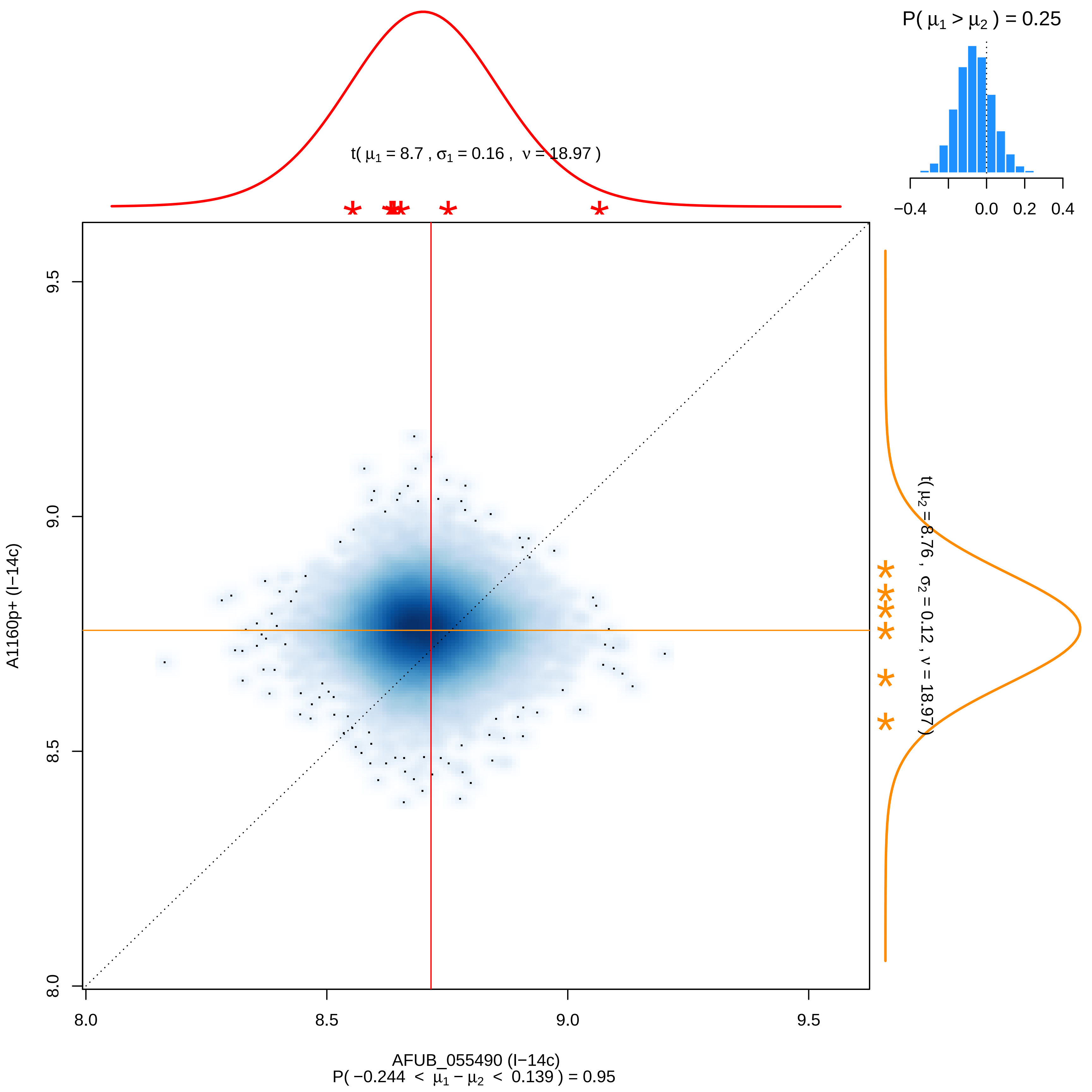

P( μ1 > μ2 ) = 0.25
−0.4
0.0
0.2
0.4
t( μ1 = 8.7 , σ1 = 0.16 , ν = 18.97 )
t( μ2 = 8.76 , σ2 = 0.12 , ν = 18.97 )
A1160p+ (I−14c)
9.5
9.0
8.5
8.0
8.0
8.5
9.0
9.5
AFUB_055490 (I−14c)
P( −0.244 < μ1 − μ2 < 0.139 ) = 0.95

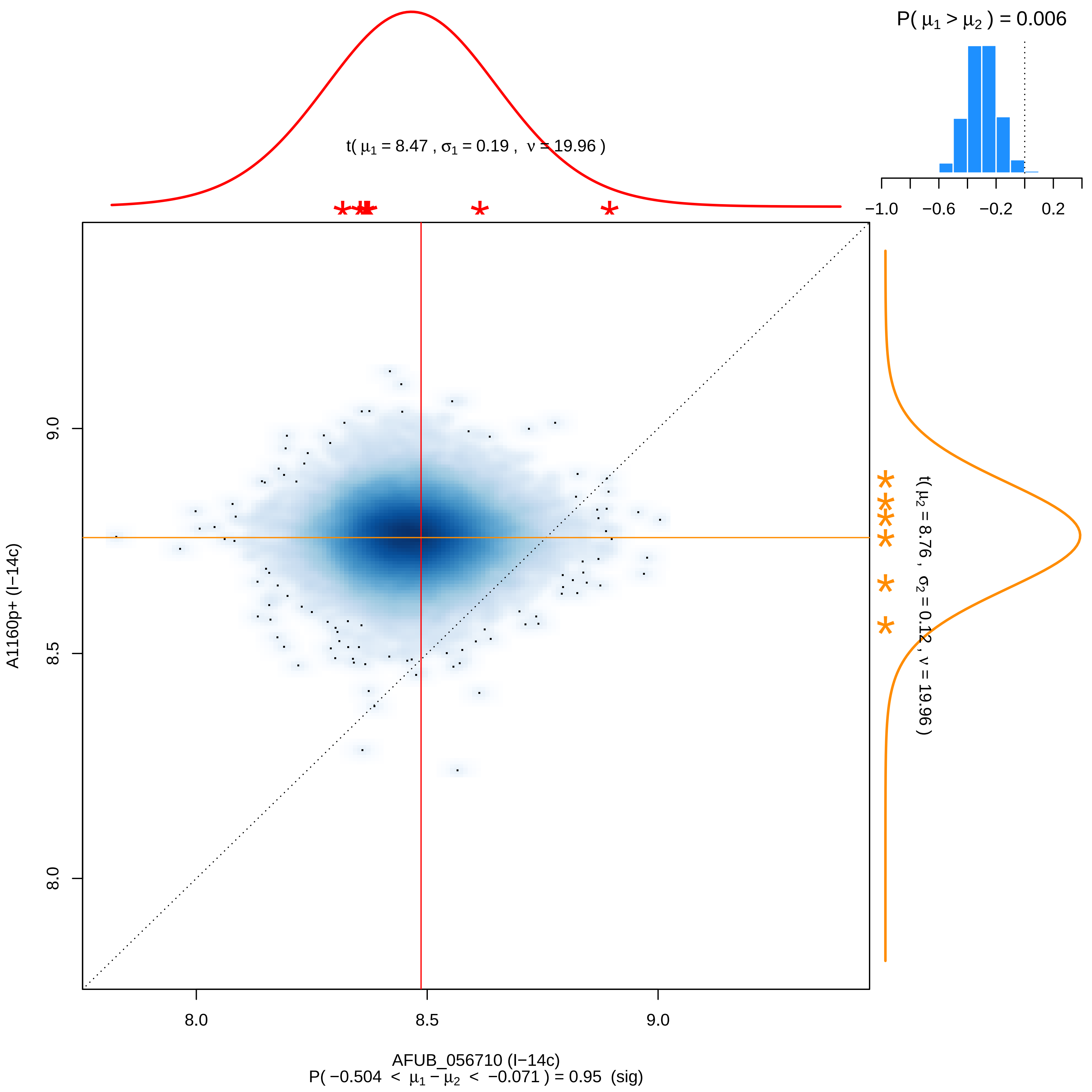

P( μ1 > μ2 ) = 0.006
−1.0
−0.6
−0.2
0.2
t( μ1 = 8.47 , σ1 = 0.19 , ν = 19.96 )
t( μ2 = 8.76 , σ2 = 0.12 , ν = 19.96 )
9.0
8.5
8.0
A1160p+ (I−14c)
8.0
8.5
9.0
AFUB_056710 (I−14c)
P( −0.504 < μ1 − μ2 < −0.071 ) = 0.95 (sig)

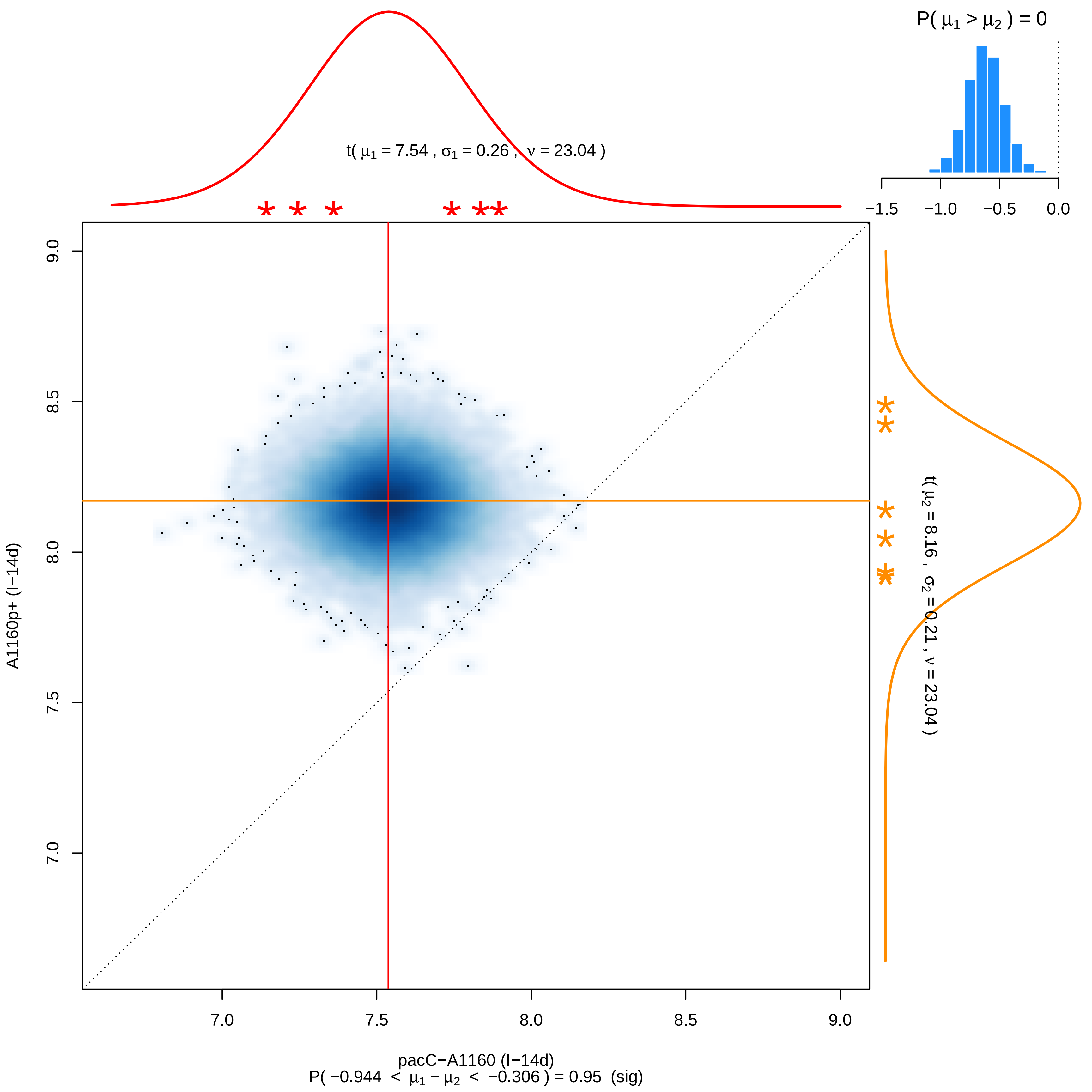
P( μ1 > μ2 ) = 0
−1.5
−1.0
−0.5
0.0
t( μ1 = 7.54 , σ1 = 0.26 , ν = 23.04 )
9.0
8.5
8.0
7.5
7.0
A1160p+ (I−14d)
t( μ2 = 8.16 , σ2 = 0.21 , ν = 23.04 )
7.0
7.5
8.0
8.5
9.0
pacC−A1160 (I−14d)
P( −0.944 < μ1 − μ2 < −0.306 ) = 0.95 (sig)

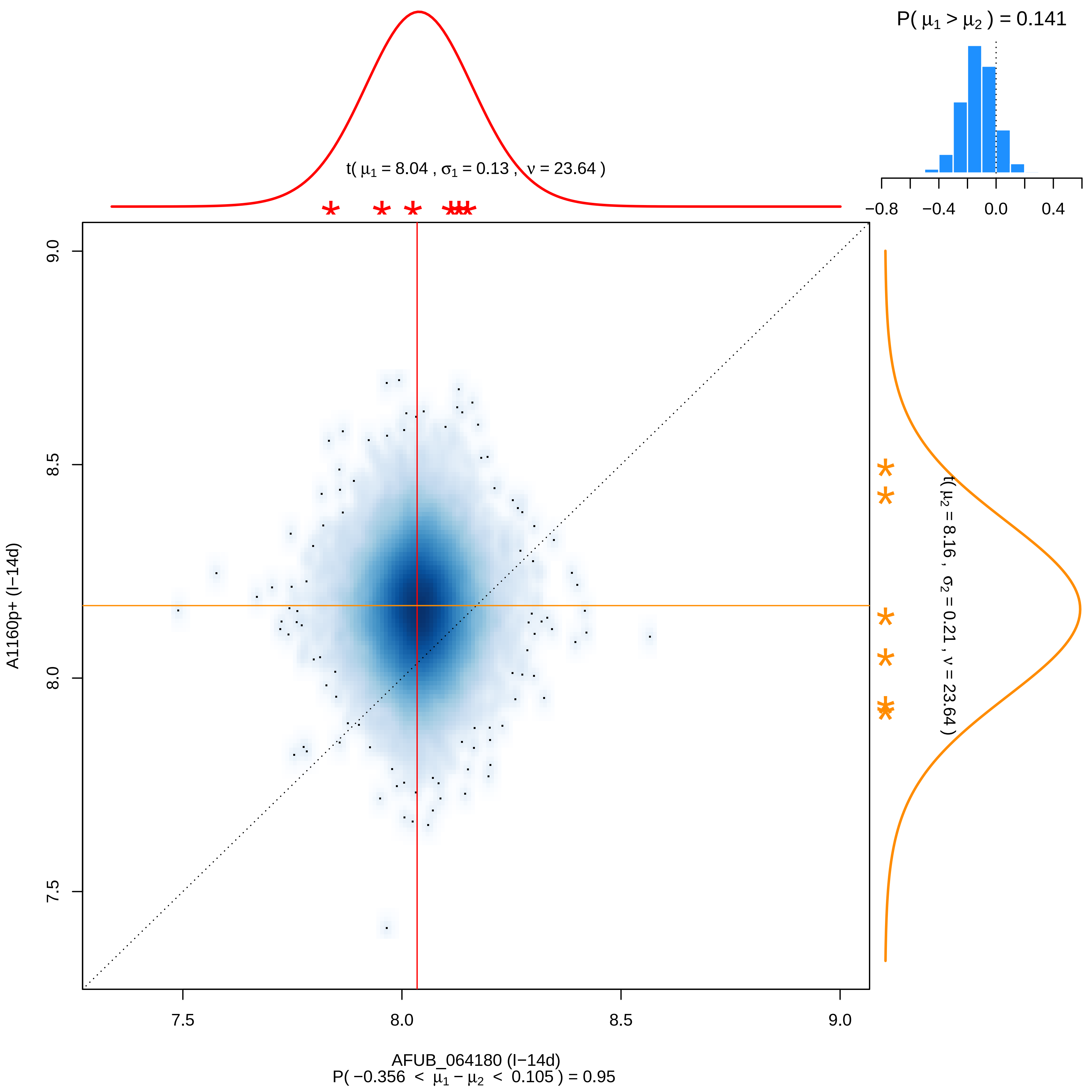

P( μ1 > μ2 ) = 0.141
−0.8
−0.4
0.0
0.4
t( μ1 = 8.04 , σ1 = 0.13 , ν = 23.64 )
t( μ2 = 8.16 , σ2 = 0.21 , ν = 23.64 )
9.0
8.5
8.0
7.5
A1160p+ (I−14d)
7.5
8.0
8.5
9.0
AFUB_064180 (I−14d)
P( −0.356 < μ1 − μ2 < 0.105 ) = 0.95

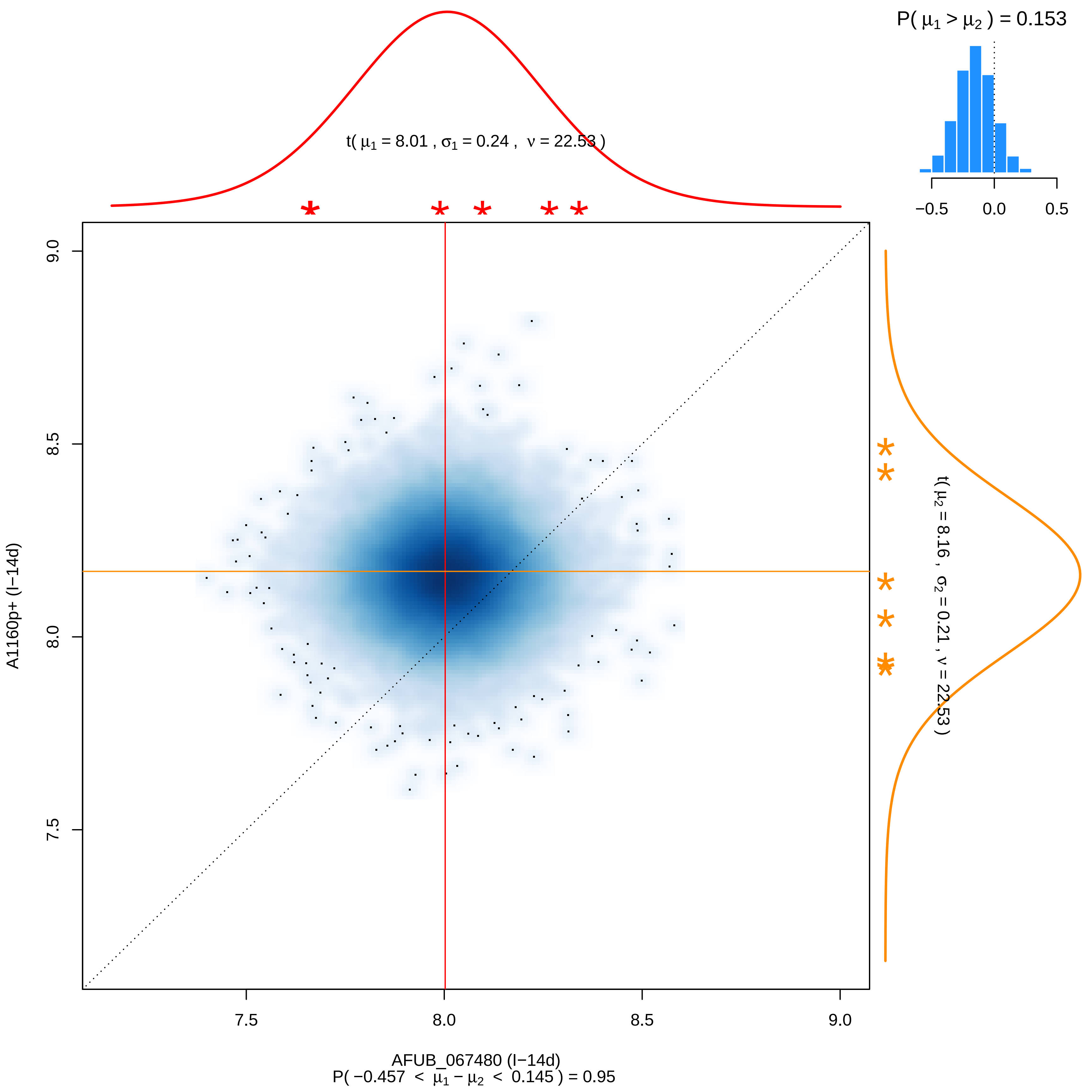
P( μ1 > μ2 ) = 0.153
−0.5
0.0
0.5
t( μ1 = 8.01 , σ1 = 0.24 , ν = 22.53 )
9.0
8.5
8.0
7.5
A1160p+ (I−14d)
t( μ2 = 8.16 , σ2 = 0.21 , ν = 22.53 )
7.5
8.0
8.5
9.0
AFUB_067480 (I−14d)
P( −0.457 < μ1 − μ2 < 0.145 ) = 0.95

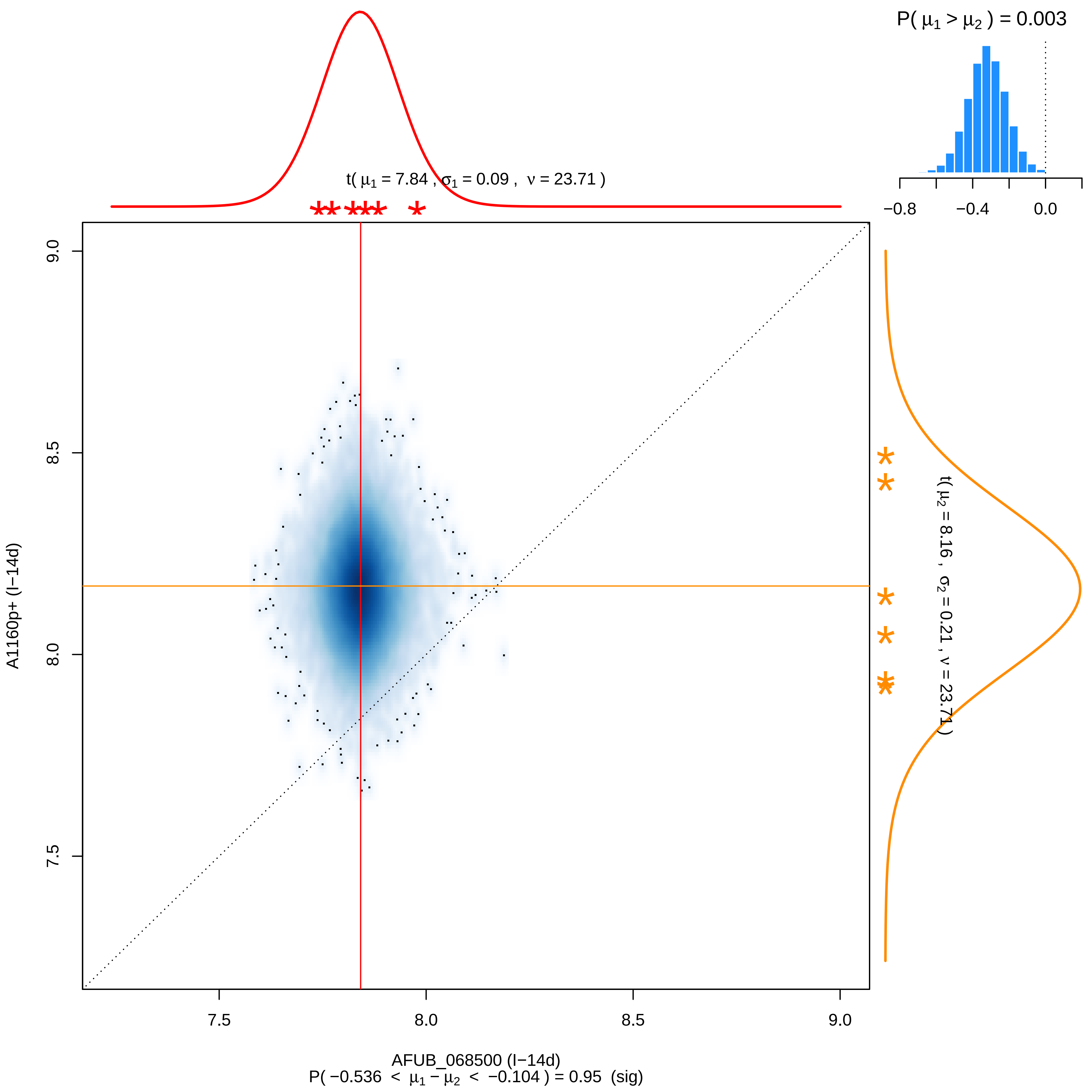

P( μ1 > μ2 ) = 0.003
−0.8
−0.4
0.0
t( μ1 = 7.84 , σ1 = 0.09 , ν = 23.71 )
9.0
8.5
8.0
7.5
A1160p+ (I−14d)
t( μ2 = 8.16 , σ2 = 0.21 , ν = 23.71 )
7.5
8.0
8.5
9.0
AFUB_068500 (I−14d)
P( −0.536 < μ1 − μ2 < −0.104 ) = 0.95 (sig)

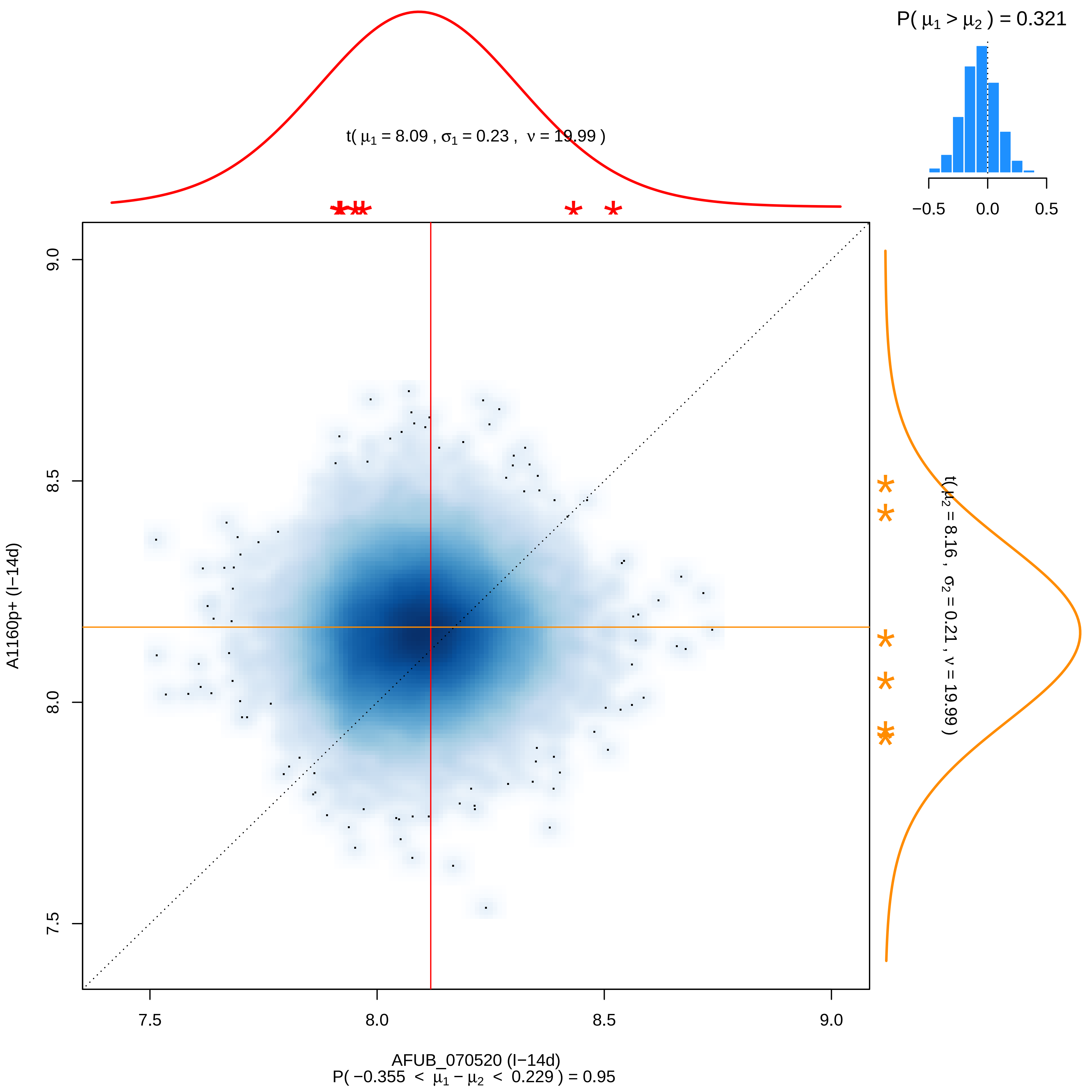

P( μ1 > μ2 ) = 0.321
−0.5
0.0
0.5
t( μ1 = 8.09 , σ1 = 0.23 , ν = 19.99 )
t( μ2 = 8.16 , σ2 = 0.21 , ν = 19.99 )
9.0
8.5
8.0
7.5
A1160p+ (I−14d)
7.5
8.0
8.5
9.0
AFUB_070520 (I−14d)
P( −0.355 < μ1 − μ2 < 0.229 ) = 0.95

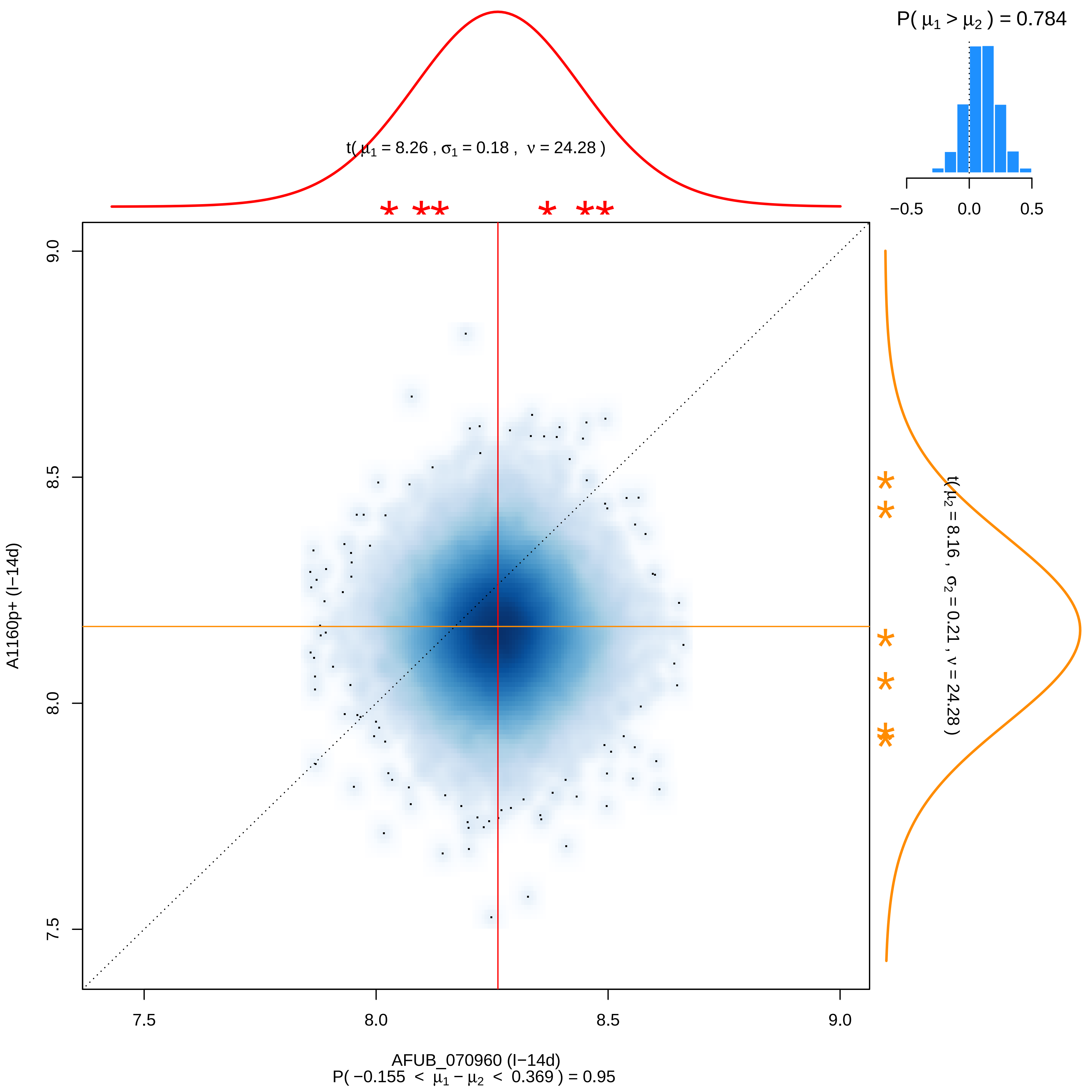
P( μ1 > μ2 ) = 0.784
−0.5
0.0
0.5
t( μ1 = 8.26 , σ1 = 0.18 , ν = 24.28 )
t( μ2 = 8.16 , σ2 = 0.21 , ν = 24.28 )
9.0
8.5
8.0
7.5
A1160p+ (I−14d)
7.5
8.0
8.5
9.0
AFUB_070960 (I−14d)
P( −0.155 < μ1 − μ2 < 0.369 ) = 0.95

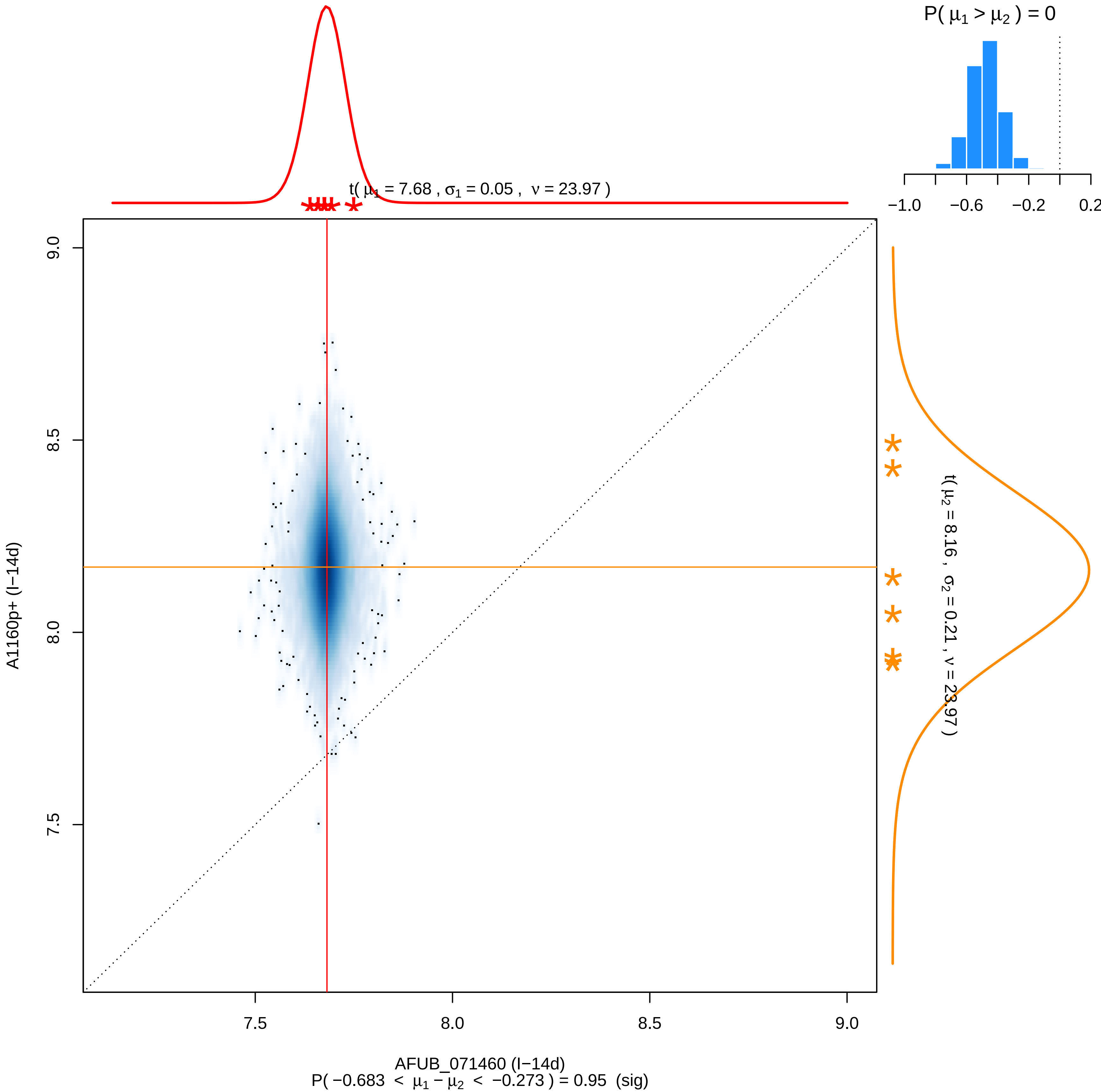

P( μ1 > μ2 ) = 0
−1.0
−0.6
−0.2
0.2
t( μ1 = 7.68 , σ1 = 0.05 , ν = 23.97 )
9.0
8.5
8.0
7.5
A1160p+ (I−14d)
7.5
8.0
8.5
9.0
t( μ2 = 8.16 , σ2 = 0.21 , ν = 23.97 )
AFUB_071460 (I−14d)
P( −0.683 < μ1 − μ2 < −0.273 ) = 0.95 (sig)

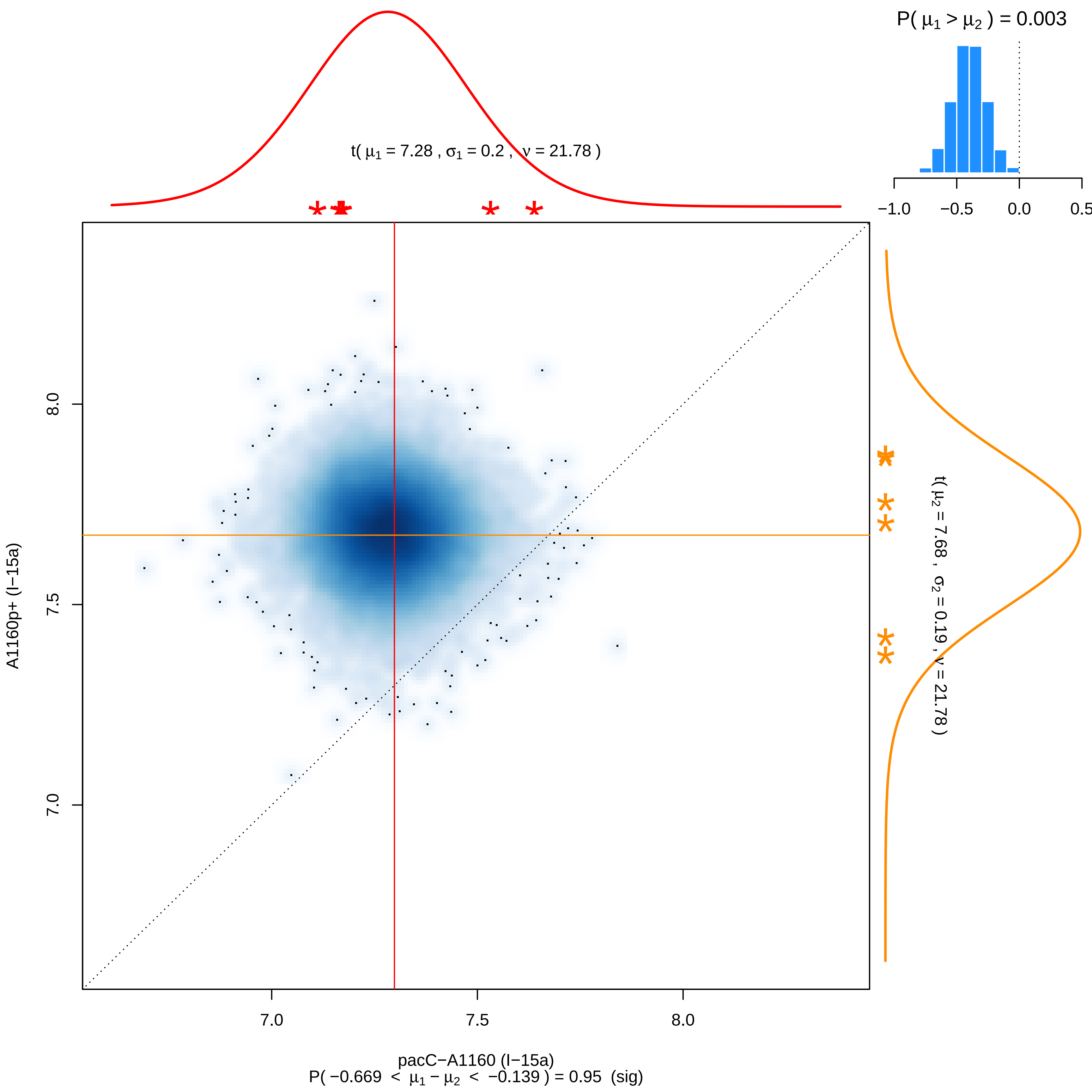

P( μ1 > μ2 ) = 0.003
−1.0
−0.5
0.0
0.5
t( μ1 = 7.28 , σ1 = 0.2 , ν = 21.78 )
t( μ2 = 7.68 , σ2 = 0.19 , ν = 21.78 )
A1160p+ (I−15a)
8.0
7.5
7.0
7.0
7.5
8.0
pacC−A1160 (I−15a)
P( −0.669 < μ1 − μ2 < −0.139 ) = 0.95 (sig)

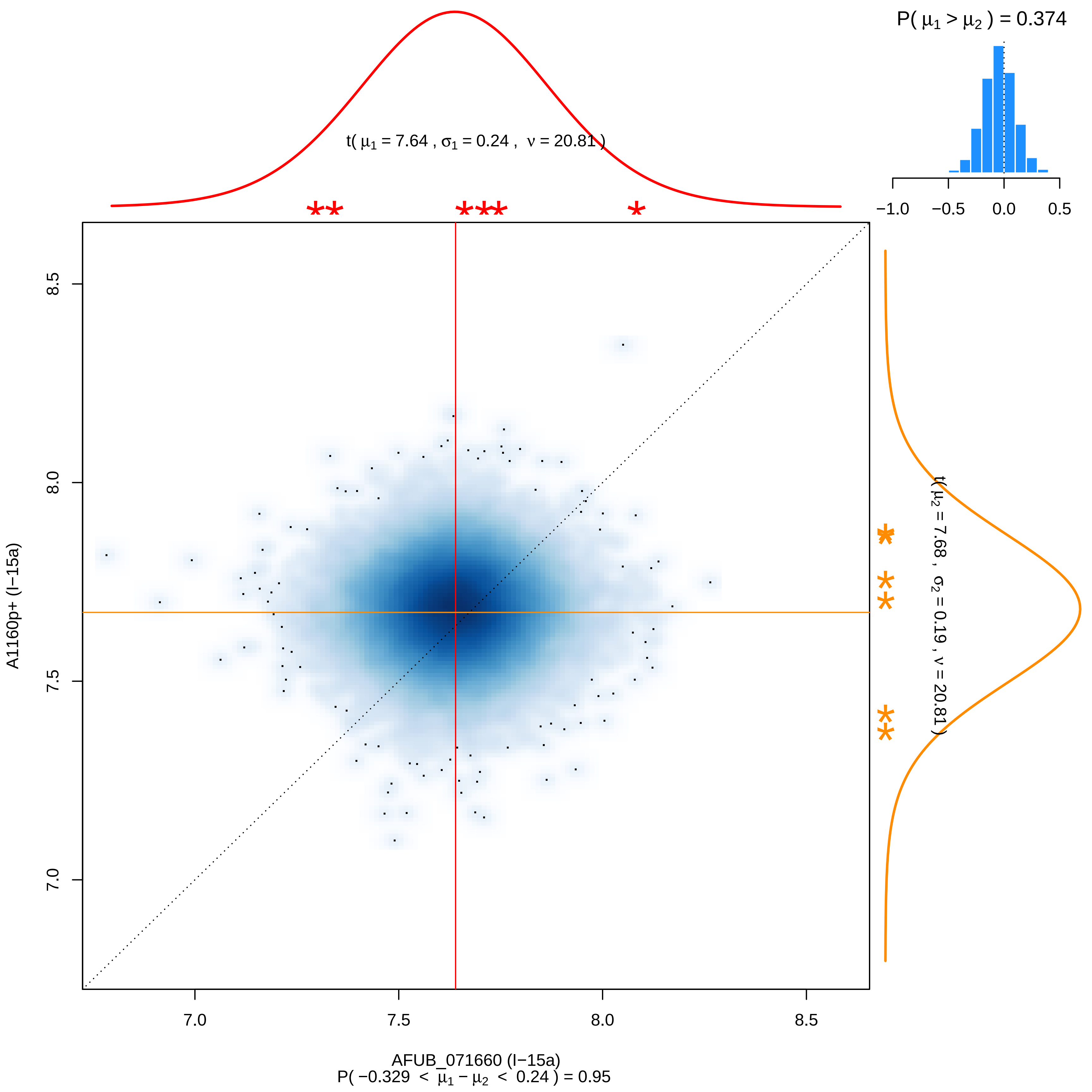

P( μ1 > μ2 ) = 0.374
−1.0
−0.5
0.0
0.5
t( μ1 = 7.64 , σ1 = 0.24 , ν = 20.81 )
t( μ2 = 7.68 , σ2 = 0.19 , ν = 20.81 )
A1160p+ (I−15a)
8.5
8.0
7.5
7.0
AFUB_071660 (I−15a)
P( −0.329 < μ1 − μ2 < 0.24 ) = 0.95

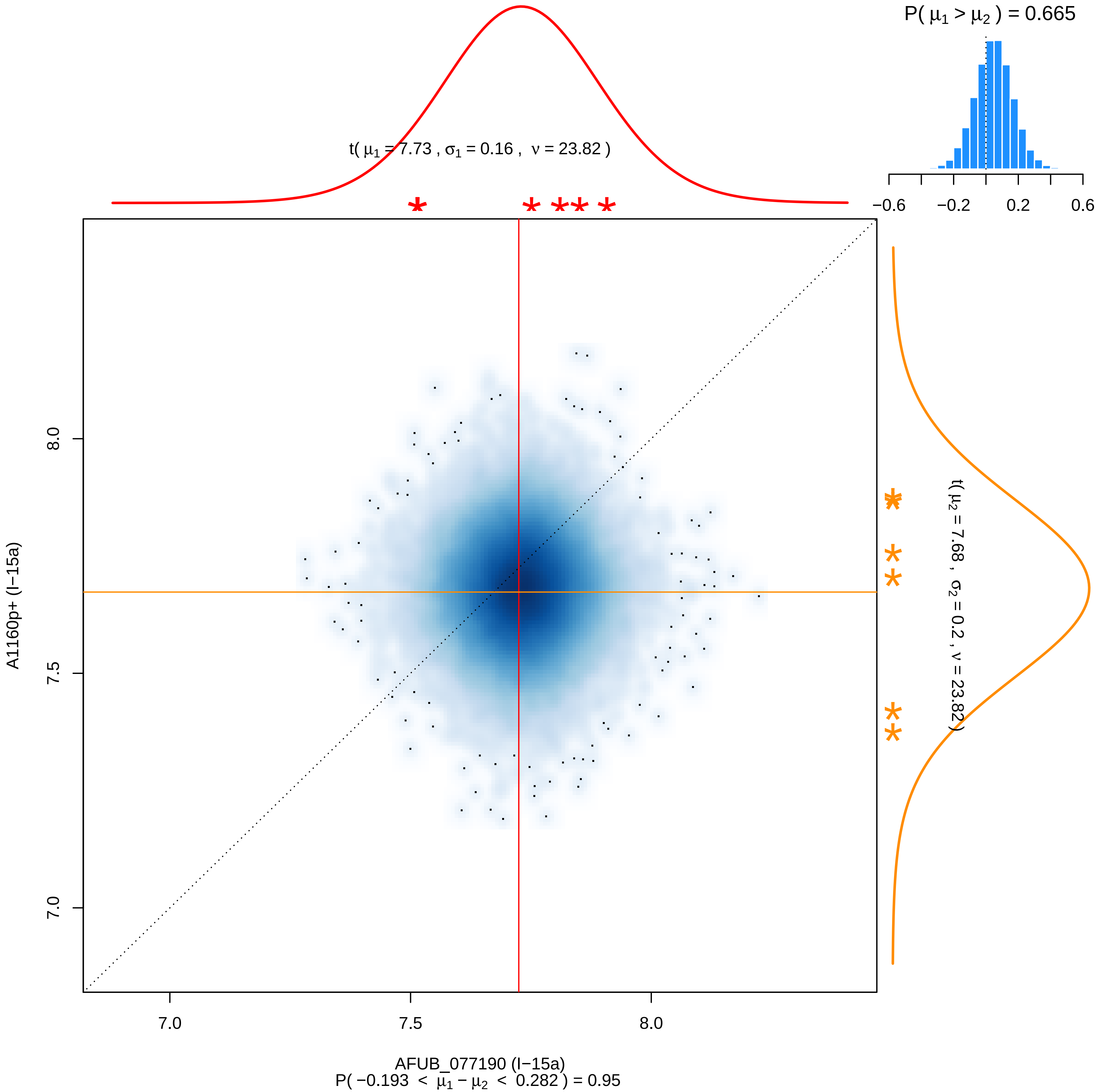

P( μ1 > μ2 ) = 0.665
−0.6
−0.2
0.2
0.6
t( μ1 = 7.73 , σ1 = 0.16 , ν = 23.82 )
t( μ2 = 7.68 , σ2 = 0.2 , ν = 23.82 )
A1160p+ (I−15a)
8.0
7.5
7.0
7.0
7.5
8.0
AFUB_077190 (I−15a)
P( −0.193 < μ1 − μ2 < 0.282 ) = 0.95

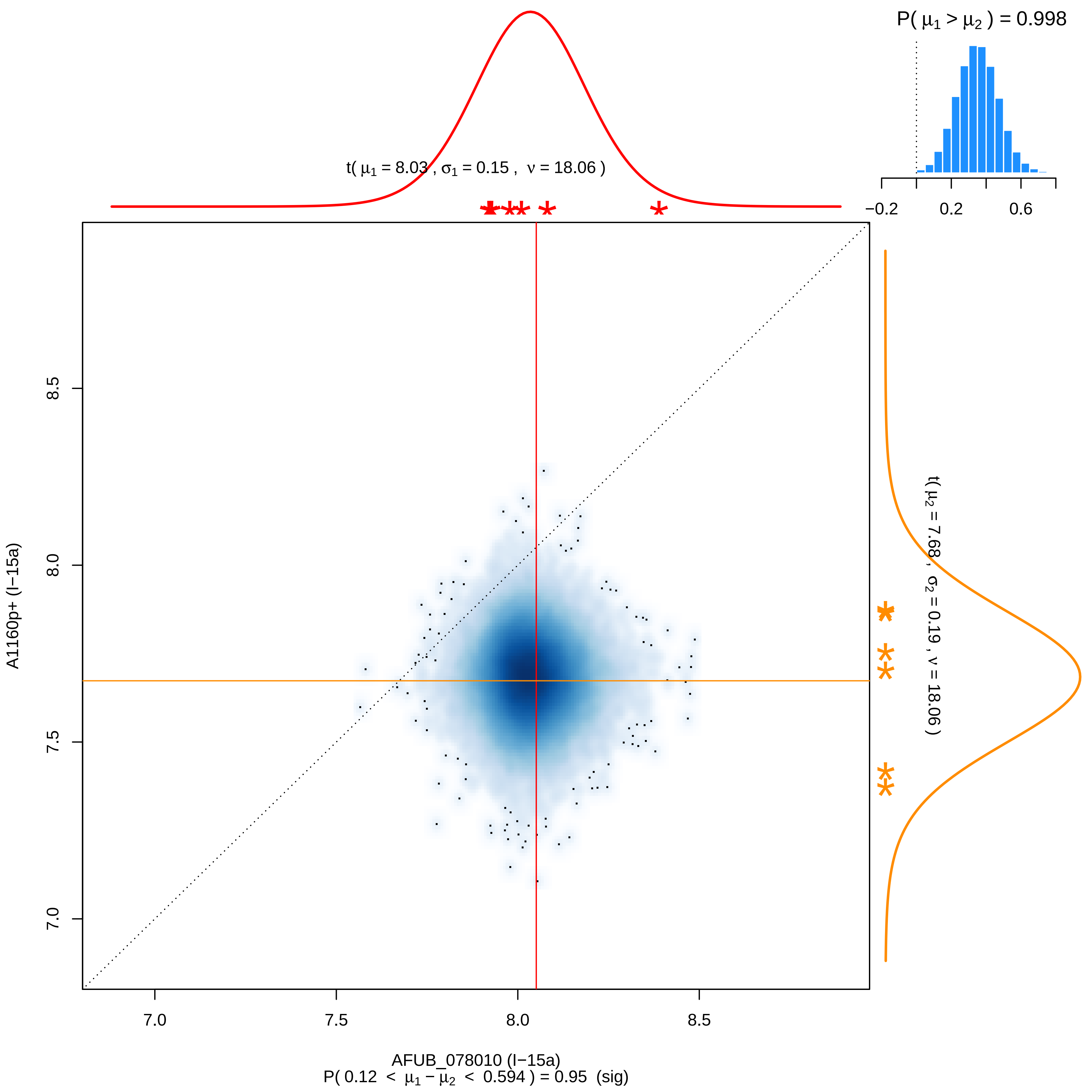
P( μ1 > μ2 ) = 0.998
−0.2
0.2
0.6
t( μ1 = 8.03 , σ1 = 0.15 , ν = 18.06 )
t( μ2 = 7.68 , σ2 = 0.19 , ν = 18.06 )
8.5
8.0
7.5
7.0
A1160p+ (I−15a)
7.0
7.5
8.0
8.5
AFUB_078010 (I−15a)
P( 0.12 < μ1 − μ2 < 0.594 ) = 0.95 (sig)

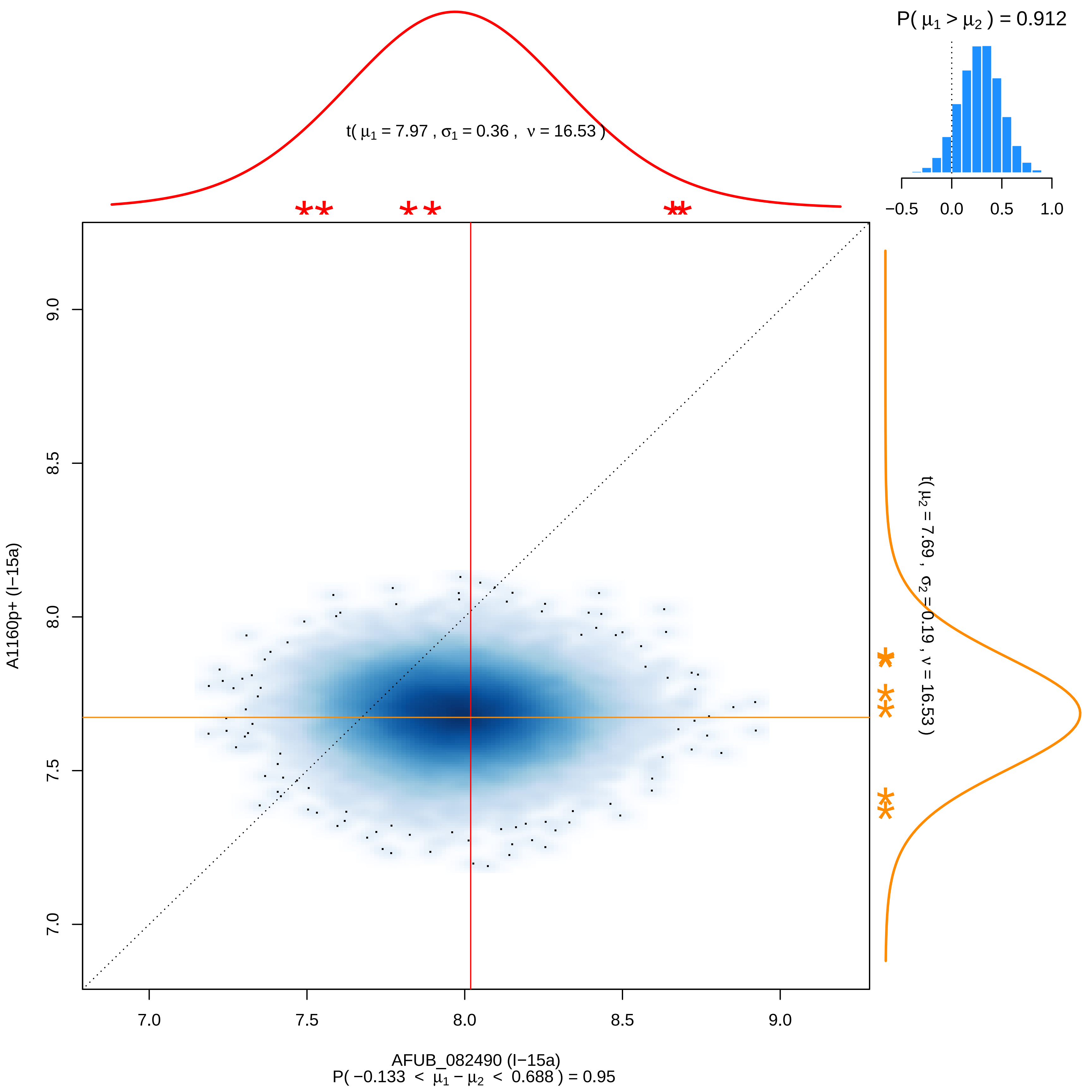

P( μ1 > μ2 ) = 0.912
−0.5 0.0 0.5 1.0
t( μ1 = 7.97 , σ1 = 0.36 , ν = 16.53 )
t( μ2 = 7.69 , σ2 = 0.19 , ν = 16.53 )
A1160p+ (I−15a)
9.0
8.5
8.0
7.5
7.0
7.0 7.5 8.0 8.5 9.0
AFUB_082490 (I−15a)
P( −0.133 < μ1 − μ2 < 0.688 ) = 0.95

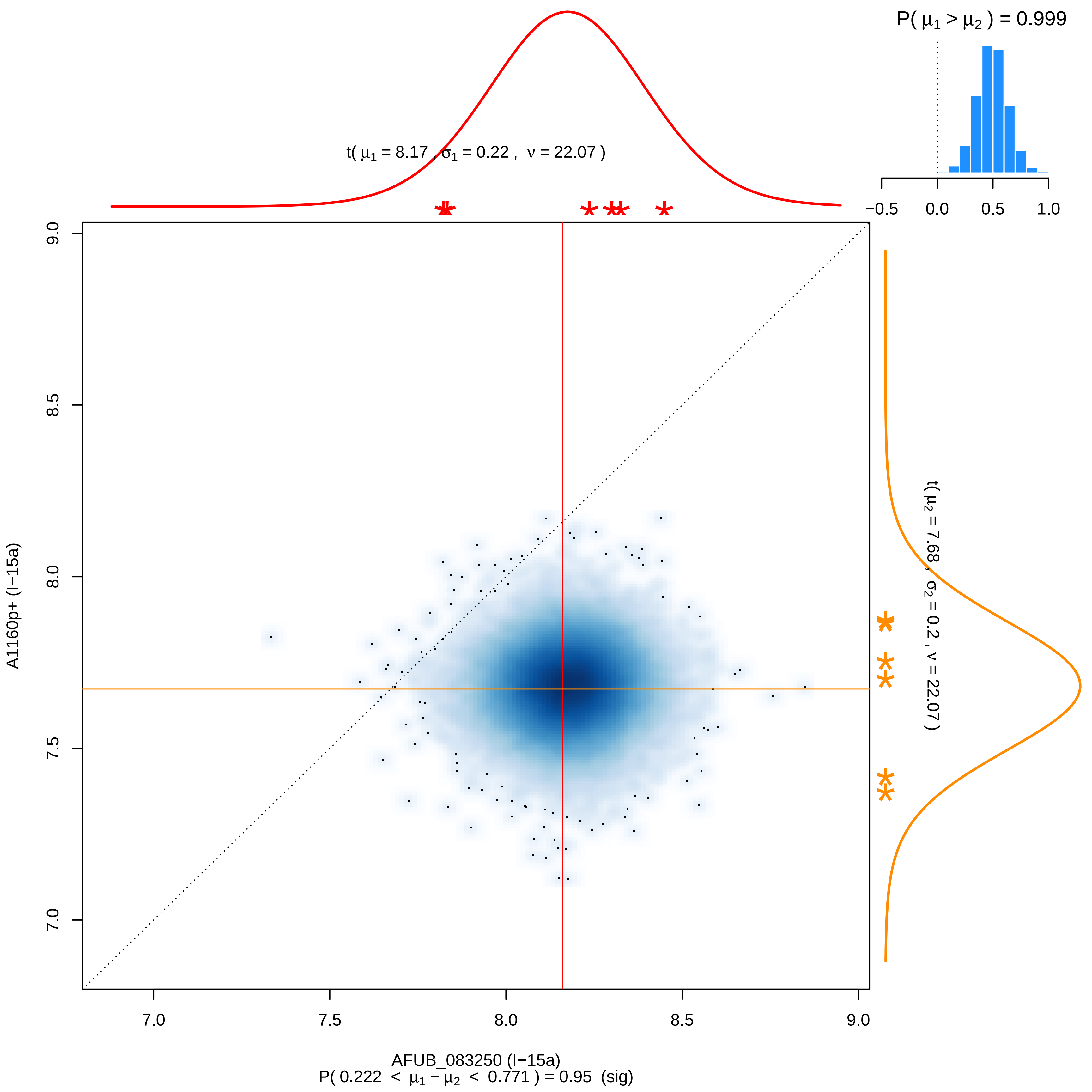

P( μ1 > μ2 ) = 0.999
t( μ1 = 8.17 , σ1 = 0.22 , ν = 22.07 )
−0.5
0.0
0.5
1.0
9.0
8.5
8.0
7.5
7.0
A1160p+ (I−15a)
t( μ2 = 7.68 , σ2 = 0.2 , ν = 22.07 )
7.0
7.5
8.0
8.5
9.0
AFUB_083250 (I−15a)
P( 0.222 < μ1 − μ2 < 0.771 ) = 0.95 (sig)

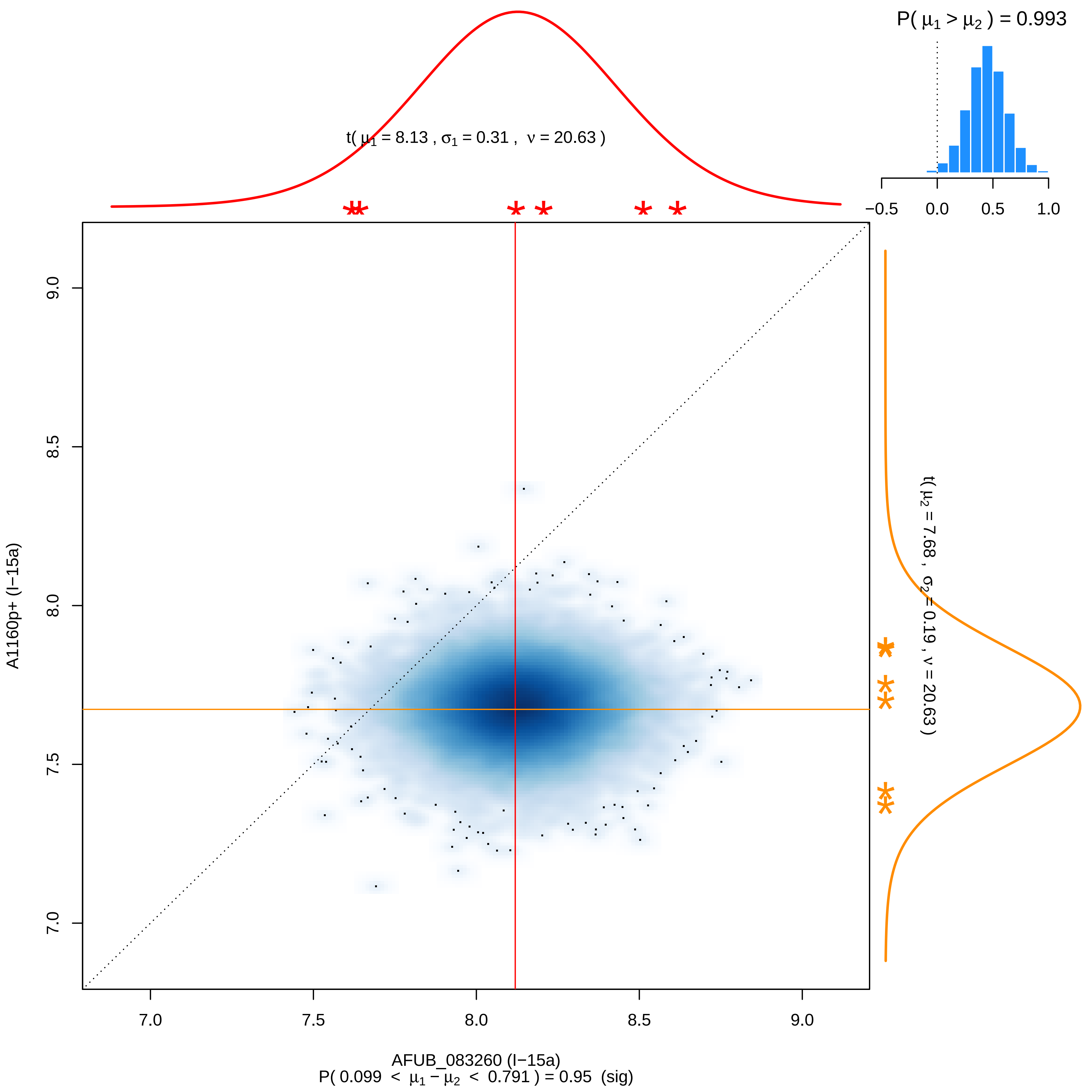

P( μ1 > μ2 ) = 0.993
−0.5
0.0
0.5
1.0
t( μ1 = 8.13 , σ1 = 0.31 , ν = 20.63 )
t( μ2 = 7.68 , σ2 = 0.19 , ν = 20.63 )
A1160p+ (I−15a)
7.0
7.5
8.0
8.5
9.0
AFUB_083260 (I−15a)
P( 0.099 < μ1 − μ2 < 0.791 ) = 0.95 (sig)

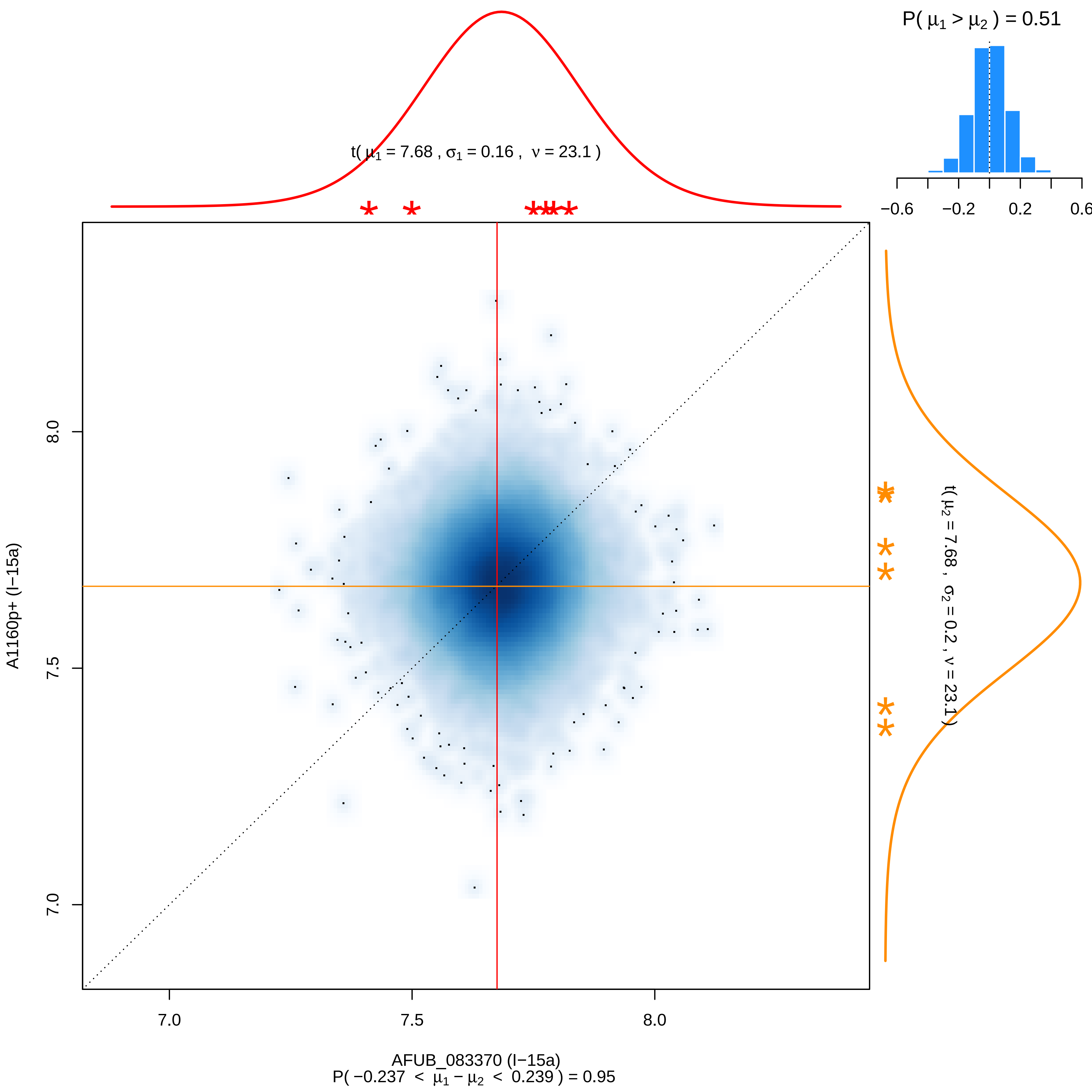

P( μ1 > μ2 ) = 0.51
−0.6
−0.2
0.2
0.6
t( μ1 = 7.68 , σ1 = 0.16 , ν = 23.1 )
t( μ2 = 7.68 , σ2 = 0.2 , ν = 23.1 )
8.0
7.5
7.0
A1160p+ (I−15a)
7.0
7.5
8.0
AFUB_083370 (I−15a)
P( −0.237 < μ1 − μ2 < 0.239 ) = 0.95

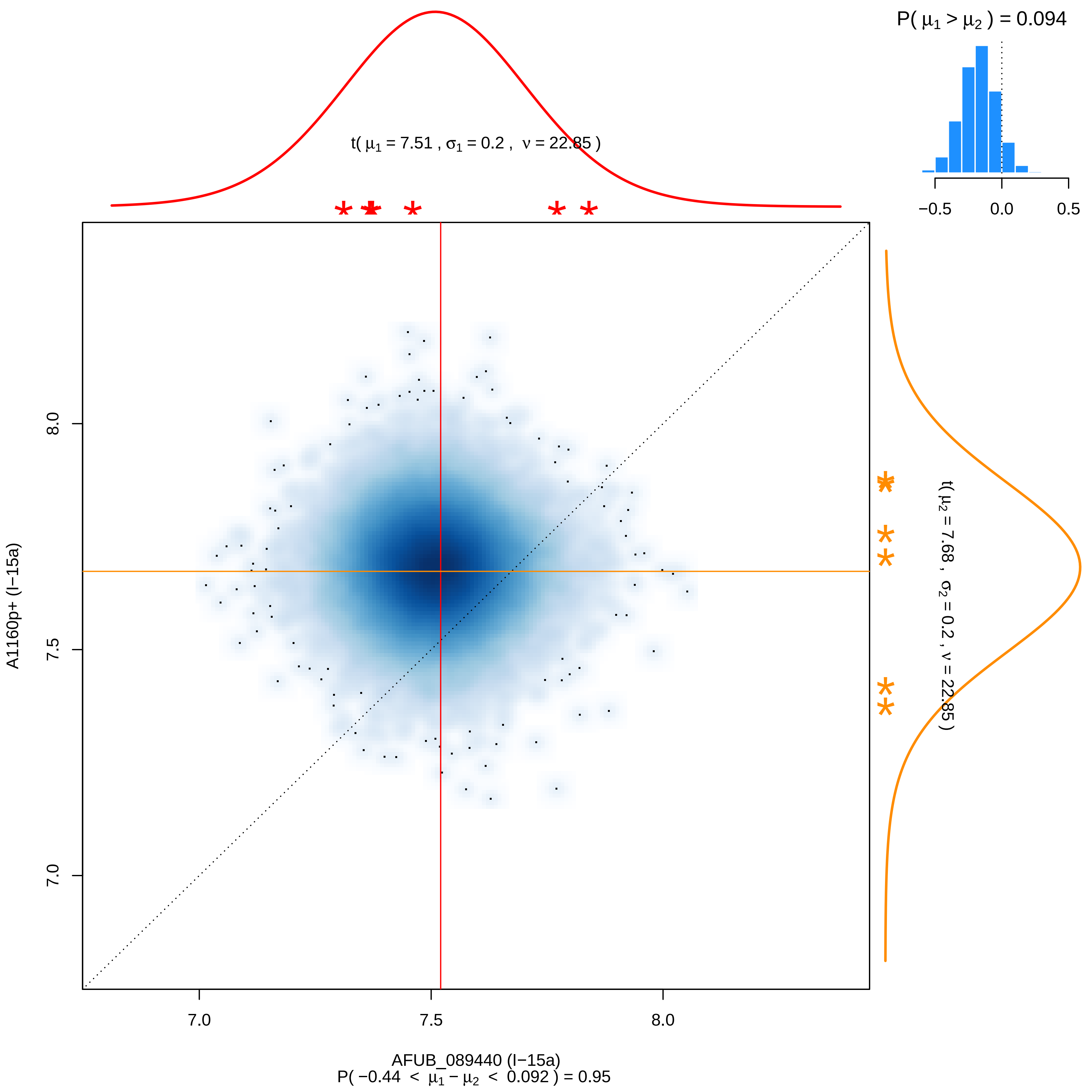

P( μ1 > μ2 ) = 0.094
−0.5
0.0
0.5
t( μ1 = 7.51 , σ1 = 0.2 , ν = 22.85 )
t( μ2 = 7.68 , σ2 = 0.2 , ν = 22.85 )
8.0
7.5
7.0
A1160p+ (I−15a)
7.0
7.5
8.0
AFUB_089440 (I−15a)
P( −0.44 < μ1 − μ2 < 0.092 ) = 0.95

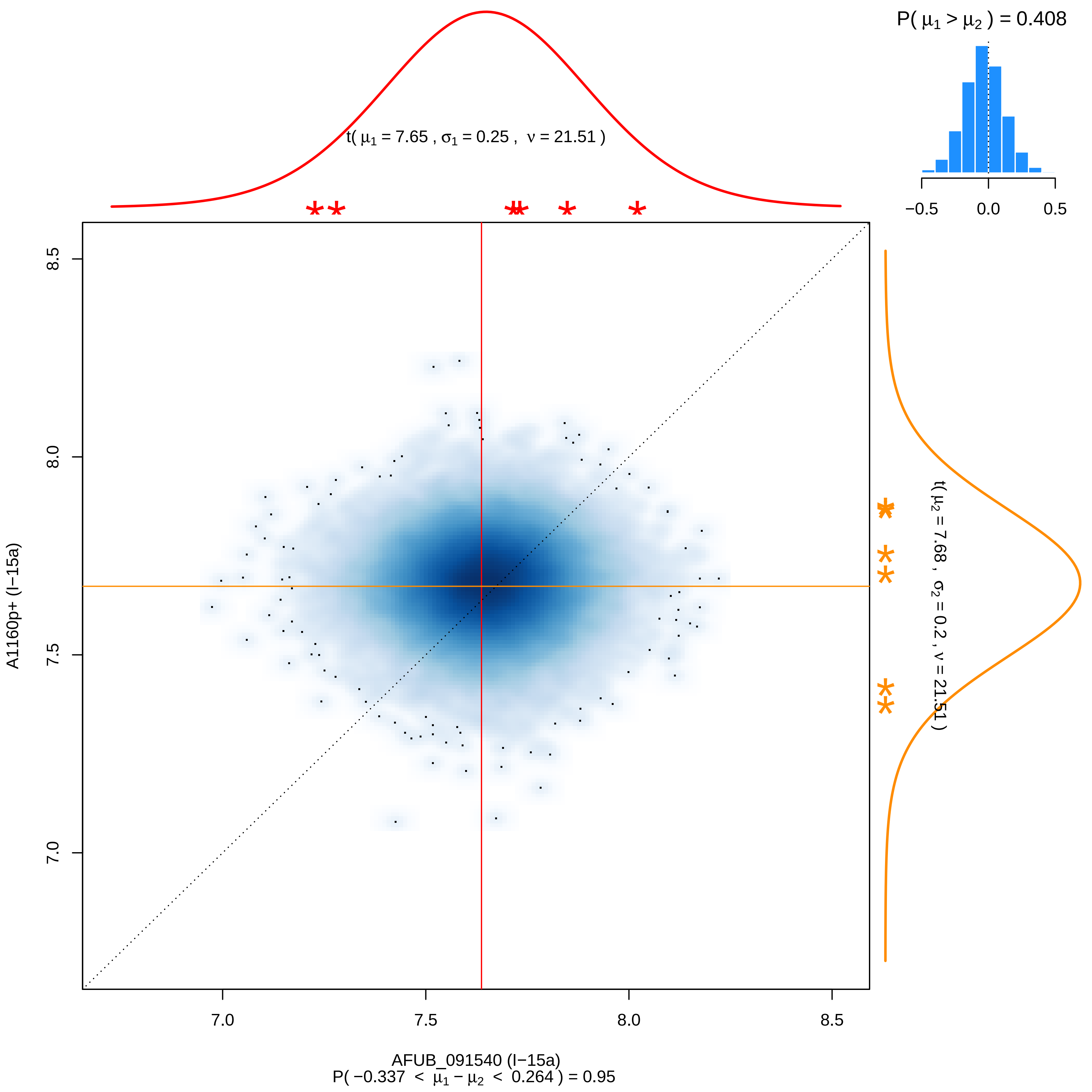

P( μ1 > μ2 ) = 0.408
−0.5
0.0
0.5
t( μ1 = 7.65 , σ1 = 0.25 , ν = 21.51 )
t( μ2 = 7.68 , σ2 = 0.2 , ν = 21.51 )
8.5
8.0
7.5
7.0
A1160p+ (I−15a)
AFUB_091540 (I−15a)
P( −0.337 < μ1 − μ2 < 0.264 ) = 0.95

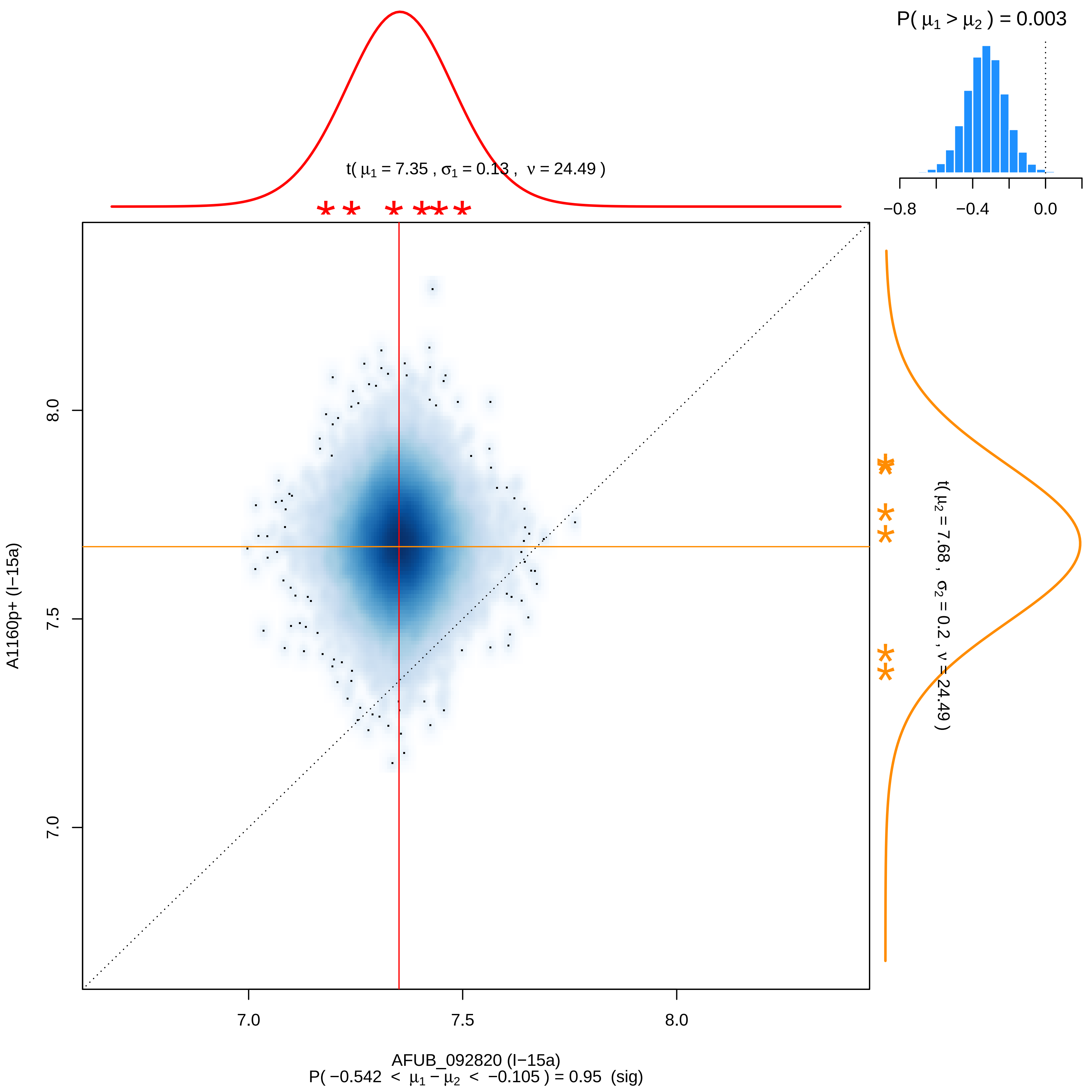

P( μ1 > μ2 ) = 0.003
−0.8
−0.4
0.0
t( μ1 = 7.35 , σ1 = 0.13 , ν = 24.49 )
8.0
7.5
7.0
A1160p+ (I−15a)
t( μ2 = 7.68 , σ2 = 0.2 , ν = 24.49 )
7.0
7.5
8.0
AFUB_092820 (I−15a)
P( −0.542 < μ1 − μ2 < −0.105 ) = 0.95 (sig)

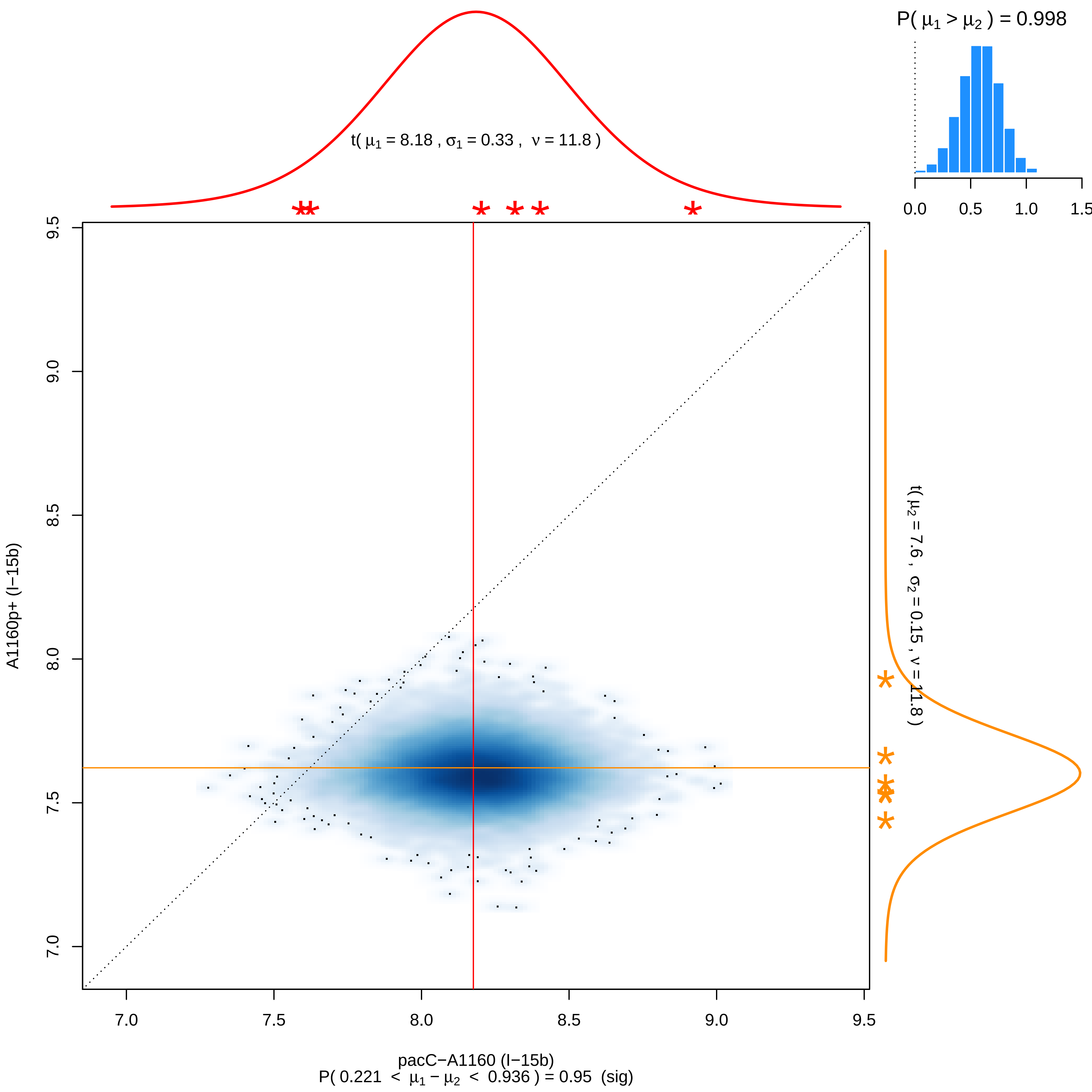

P( μ1 > μ2 ) = 0.998
0.0 0.5 1.0 1.5
t( μ1 = 8.18 , σ1 = 0.33 , ν = 11.8 )
t( μ2 = 7.6 , σ2 = 0.15 , ν = 11.8 )
A1160p+ (I−15b)
7.0 7.5 8.0 8.5 9.0 9.5
pacC−A1160 (I−15b)
P( 0.221 < μ1 − μ2 < 0.936 ) = 0.95 (sig)

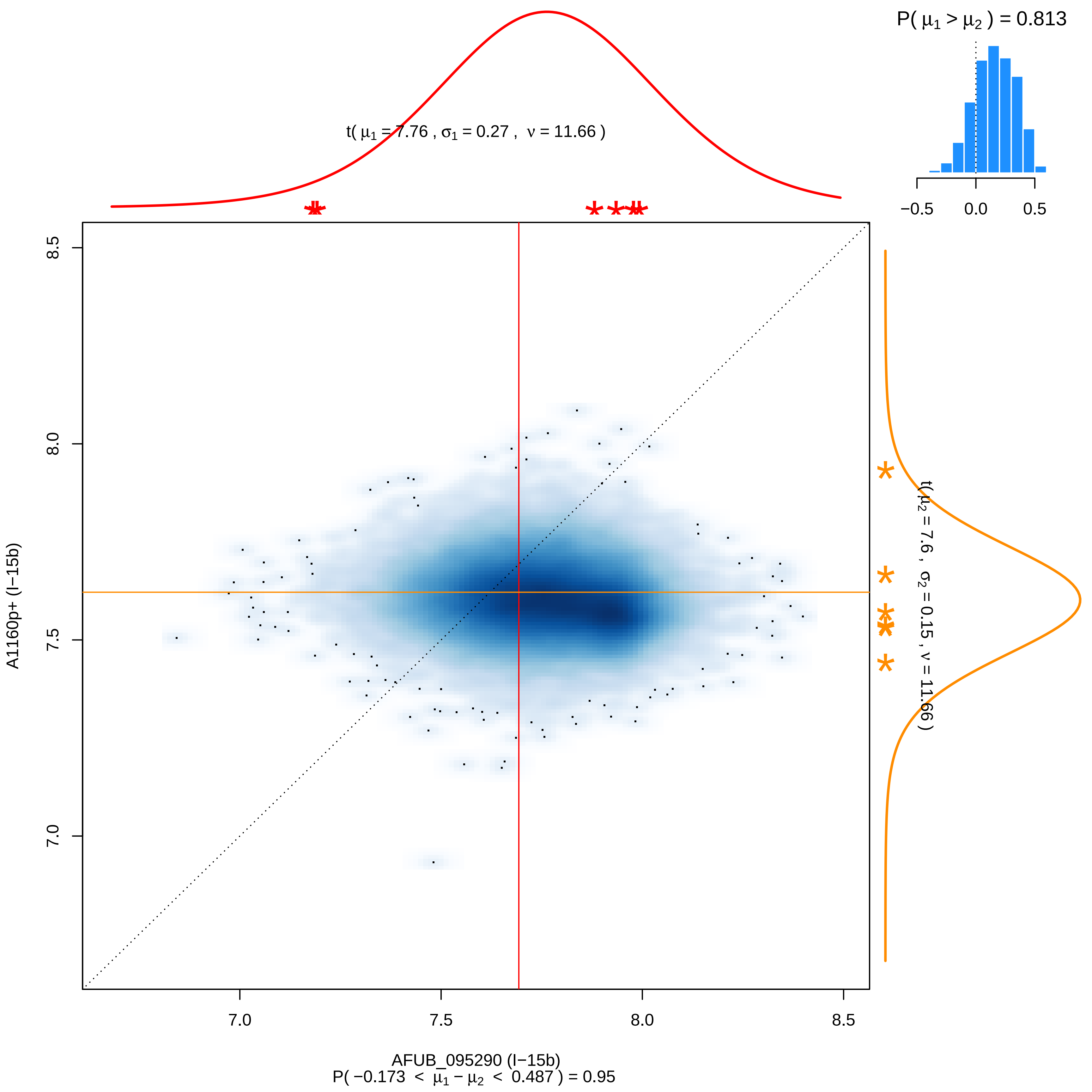

P( μ1 > μ2 ) = 0.813
−0.5
0.0
0.5
t( μ1 = 7.76 , σ1 = 0.27 , ν = 11.66 )
t( μ2 = 7.6 , σ2 = 0.15 , ν = 11.66 )
8.5
8.0
7.5
7.0
A1160p+ (I−15b)
AFUB_095290 (I−15b)
P( −0.173 < μ1 − μ2 < 0.487 ) = 0.95

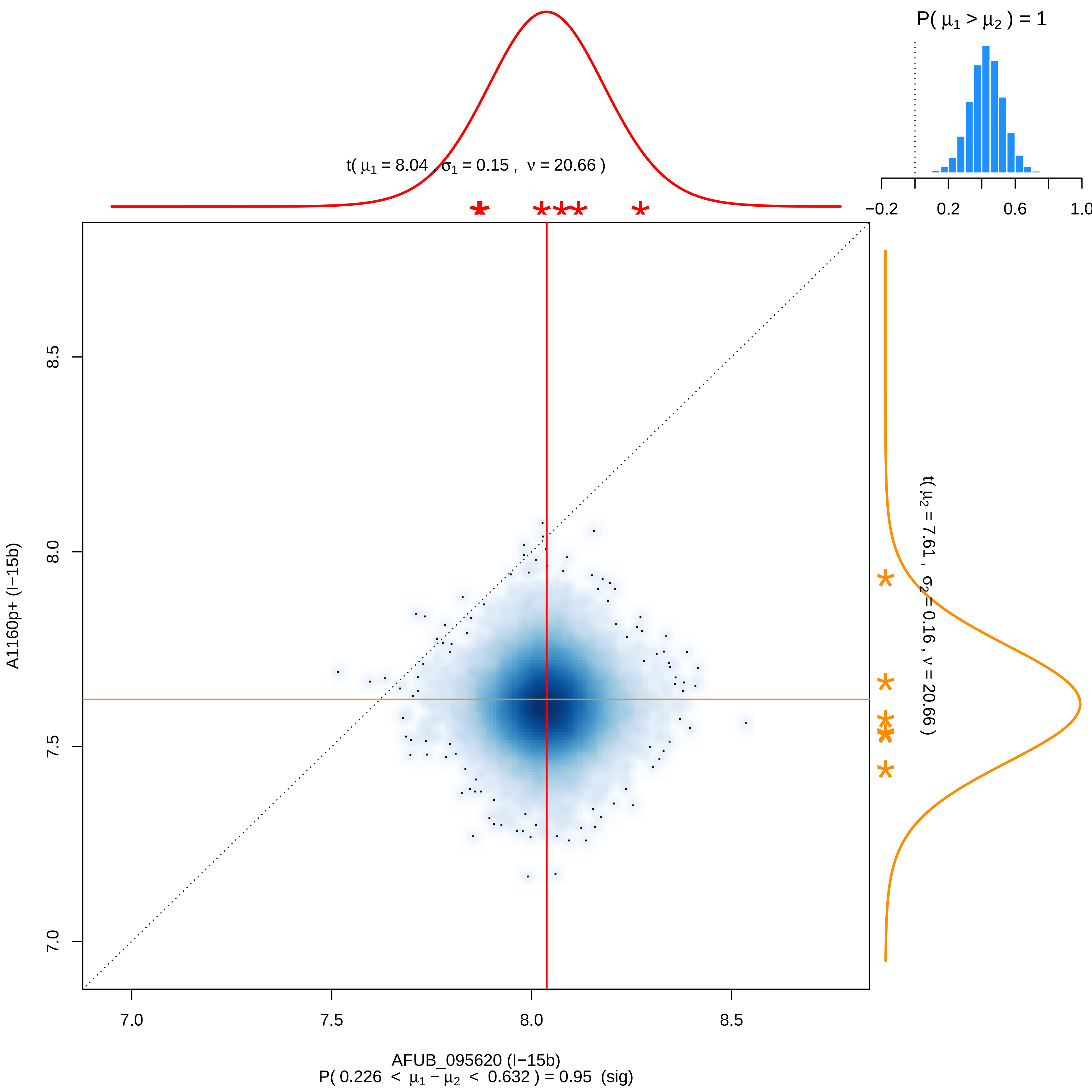

P( μ1 > μ2 ) = 1
−0.2
0.2
0.6
1.0
t( μ1 = 8.04 , σ1 = 0.15 , ν = 20.66 )
8.5
8.0
7.5
7.0
A1160p+ (I−15b)
t( μ2 = 7.61 , σ2 = 0.16 , ν = 20.66 )
7.0
7.5
8.0
8.5
AFUB_095620 (I−15b)
P( 0.226 < μ1 − μ2 < 0.632 ) = 0.95 (sig)

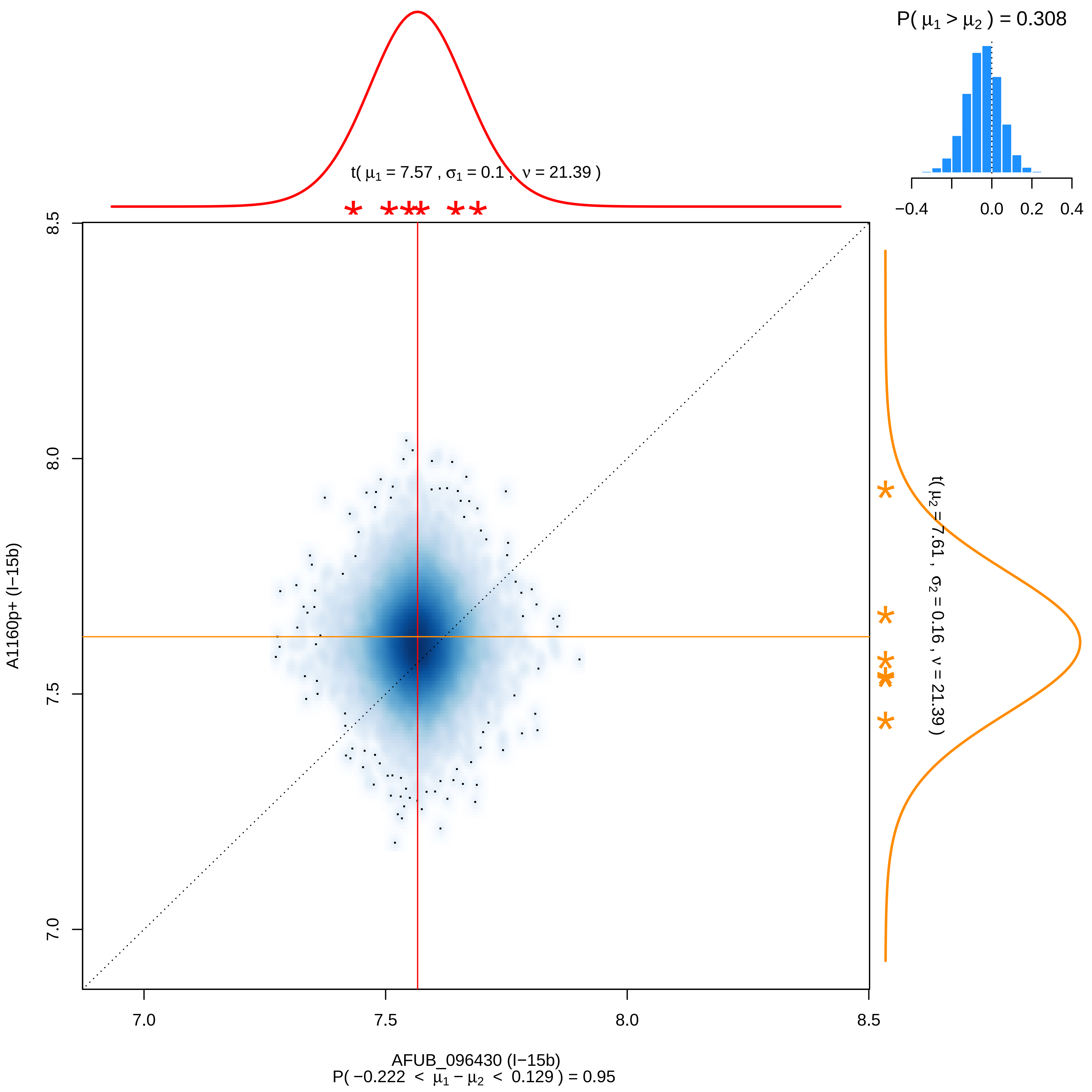

P( μ1 > μ2 ) = 0.308
−0.4
0.0
0.2
0.4
t( μ1 = 7.57 , σ1 = 0.1 , ν = 21.39 )
t( μ2 = 7.61 , σ2 = 0.16 , ν = 21.39 )
8.5
8.0
7.5
7.0
A1160p+ (I−15b)
7.0
7.5
8.0
8.5
AFUB_096430 (I−15b)
P( −0.222 < μ1 − μ2 < 0.129 ) = 0.95

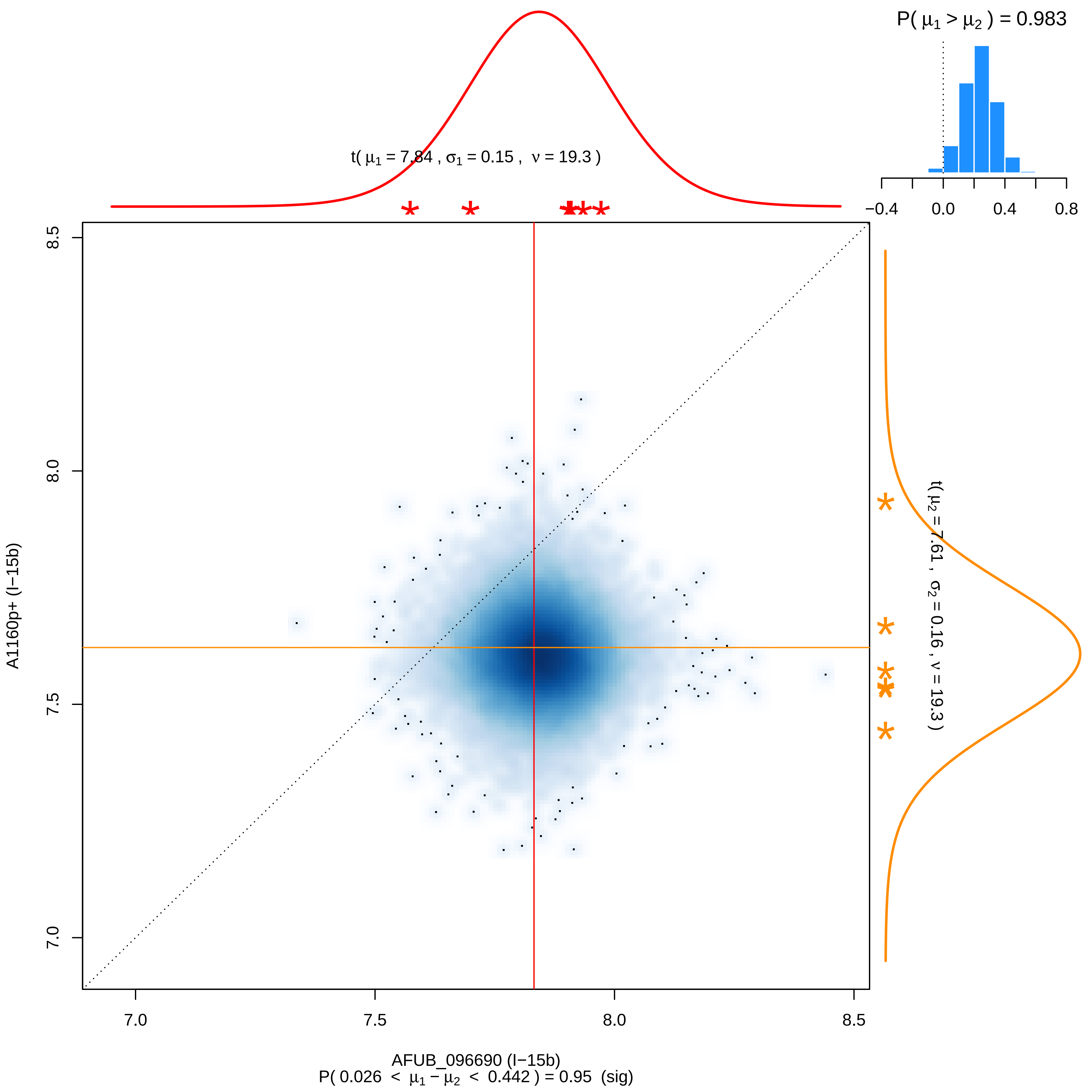

P( μ1 > μ2 ) = 0.983
−0.4
0.0
0.4
0.8
t( μ1 = 7.84 , σ1 = 0.15 , ν = 19.3 )
t( μ2 = 7.61 , σ2 = 0.16 , ν = 19.3 )
8.5
8.0
7.5
7.0
A1160p+ (I−15b)
7.0
7.5
8.0
8.5
AFUB_096690 (I−15b)
P( 0.026 < μ1 − μ2 < 0.442 ) = 0.95 (sig)

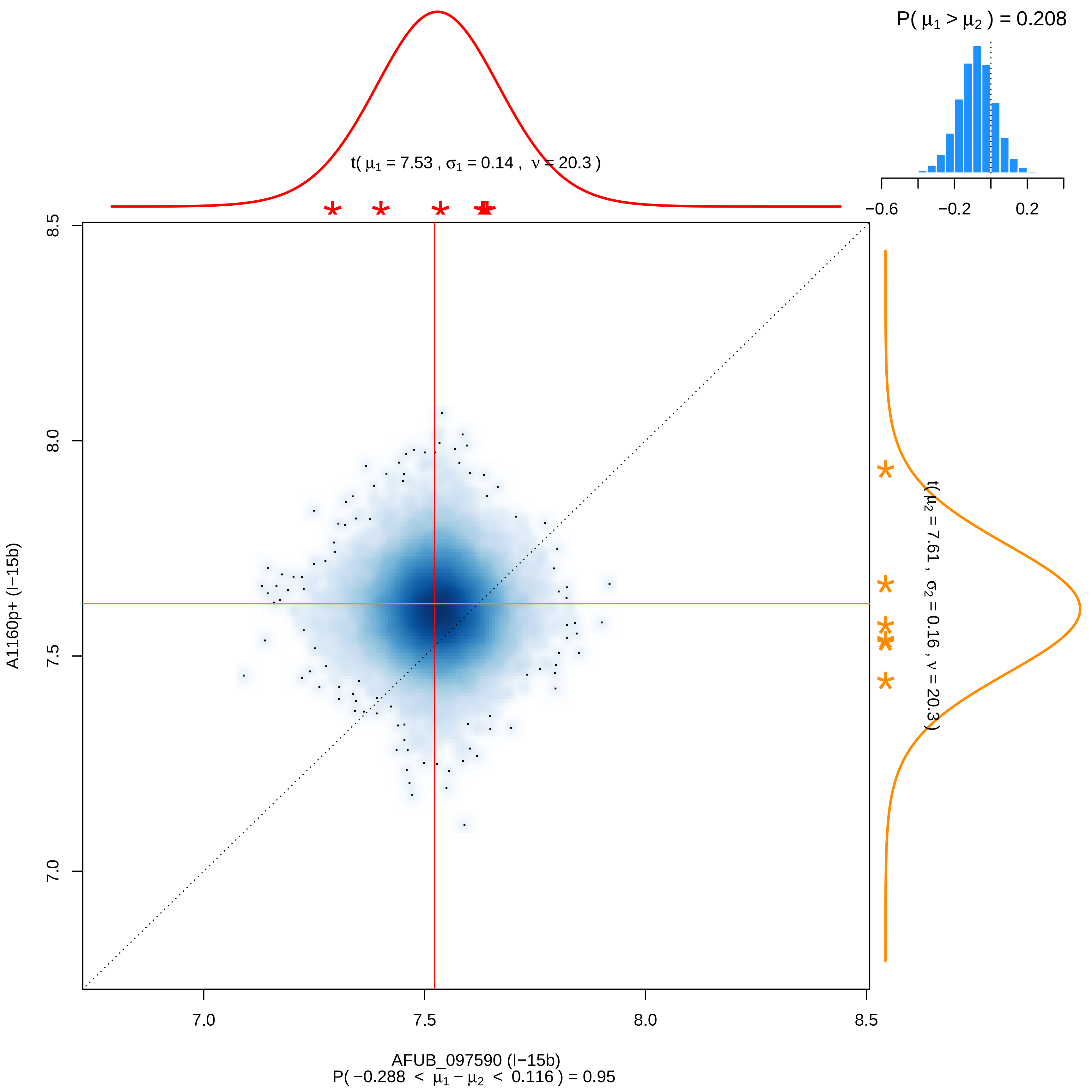

P( μ1 > μ2 ) = 0.208
−0.6
−0.2
0.2
t( μ1 = 7.53 , σ1 = 0.14 , ν = 20.3 )
t( μ2 = 7.61 , σ2 = 0.16 , ν = 20.3 )
8.5
8.0
7.5
7.0
A1160p+ (I−15b)
7.0
7.5
8.0
8.5
AFUB_097590 (I−15b)
P( −0.288 < μ1 − μ2 < 0.116 ) = 0.95

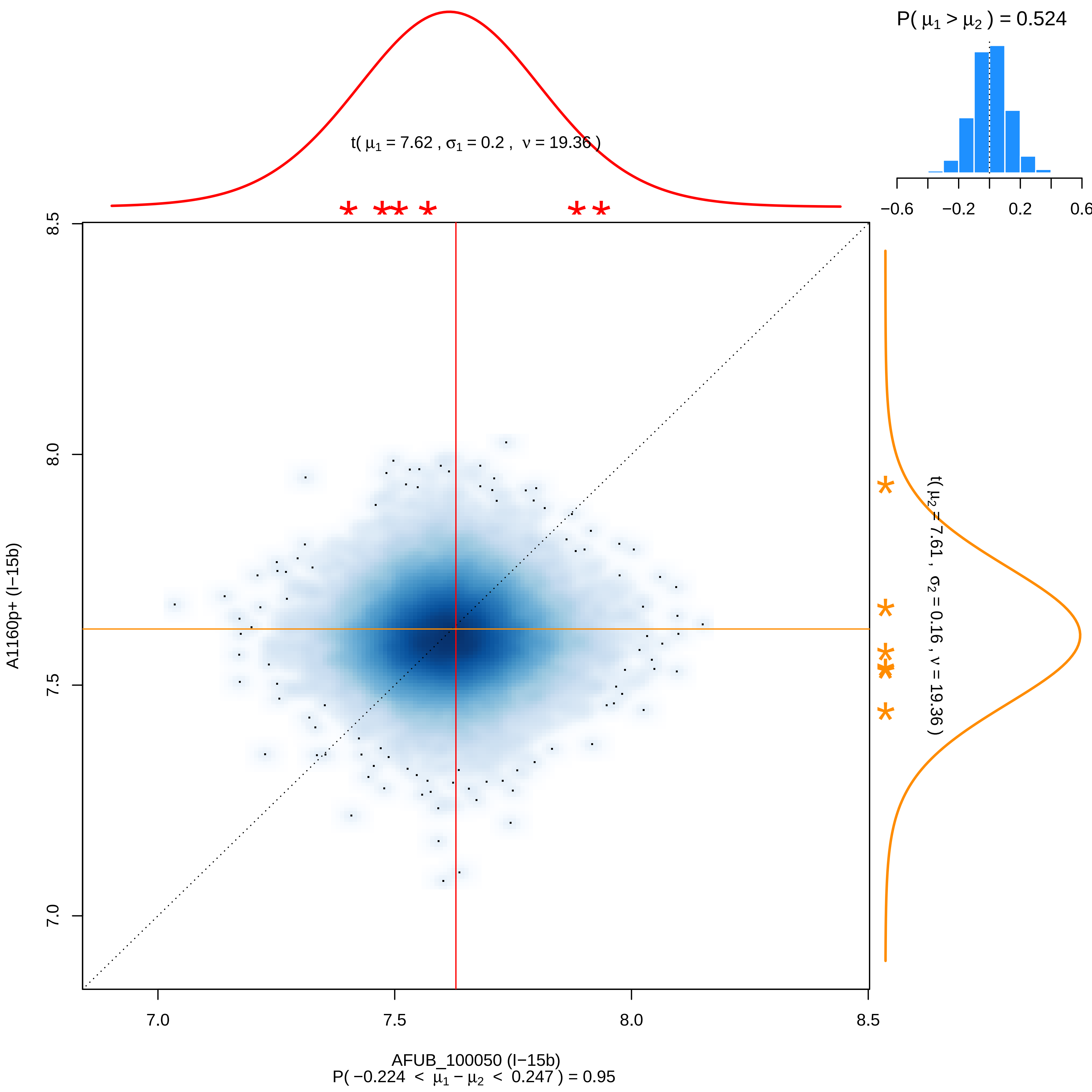

P( μ1 > μ2 ) = 0.524
−0.6
−0.2
0.2
0.6
t( μ1 = 7.62 , σ1 = 0.2 , ν = 19.36 )
t( μ2 = 7.61 , σ2 = 0.16 , ν = 19.36 )
A1160p+ (I−15b)
8.5
8.0
7.5
7.0
7.0
7.5
8.0
8.5
AFUB_100050 (I−15b)
P( −0.224 < μ1 − μ2 < 0.247 ) = 0.95

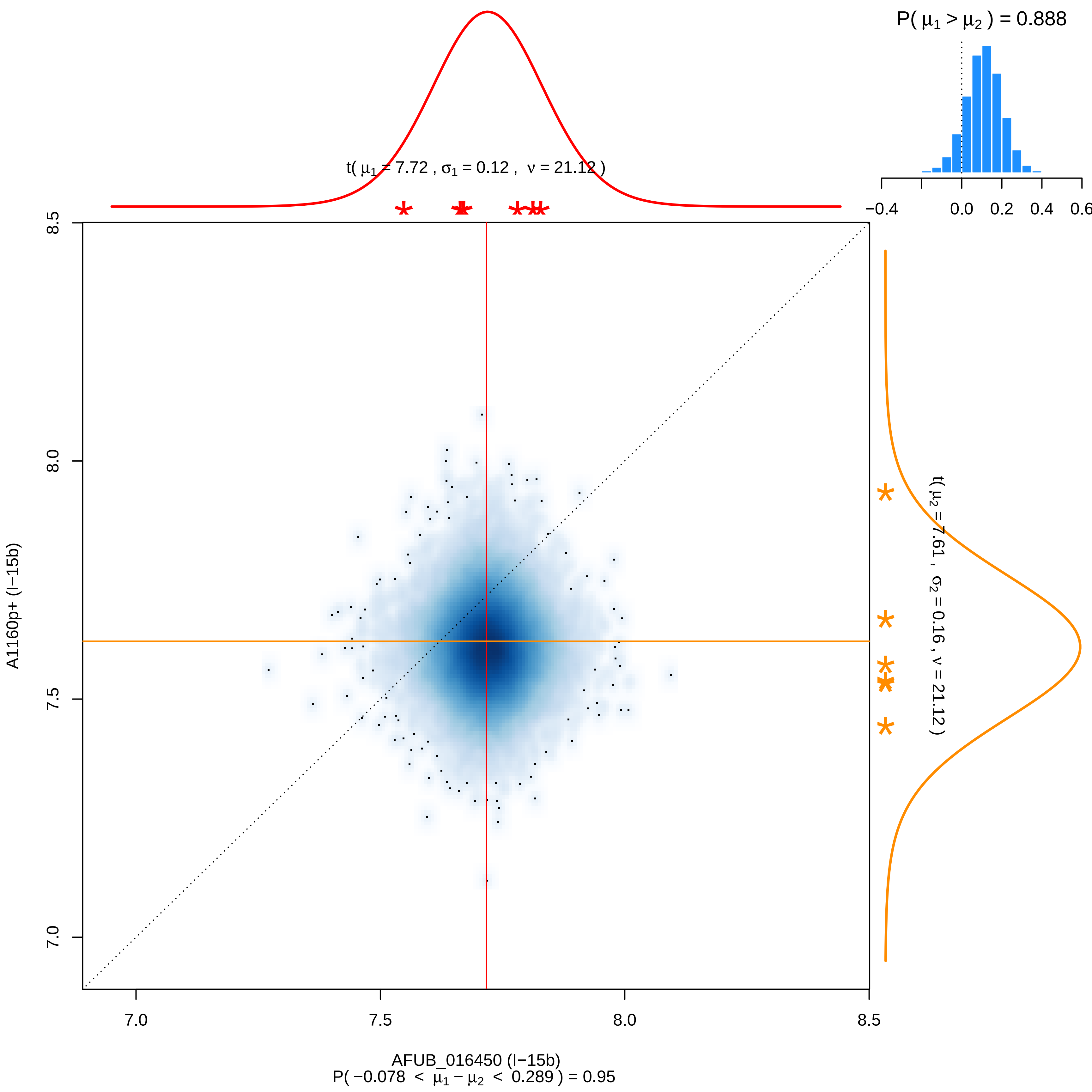

P( μ1 > μ2 ) = 0.888
−0.4
0.0
0.2
0.4
0.6
t( μ1 = 7.72 , σ1 = 0.12 , ν = 21.12 )
t( μ2 = 7.61 , σ2 = 0.16 , ν = 21.12 )
8.5
8.0
7.5
7.0
A1160p+ (I−15b)
7.0
7.5
8.0
8.5
AFUB_016450 (I−15b)
P( −0.078 < μ1 − μ2 < 0.289 ) = 0.95

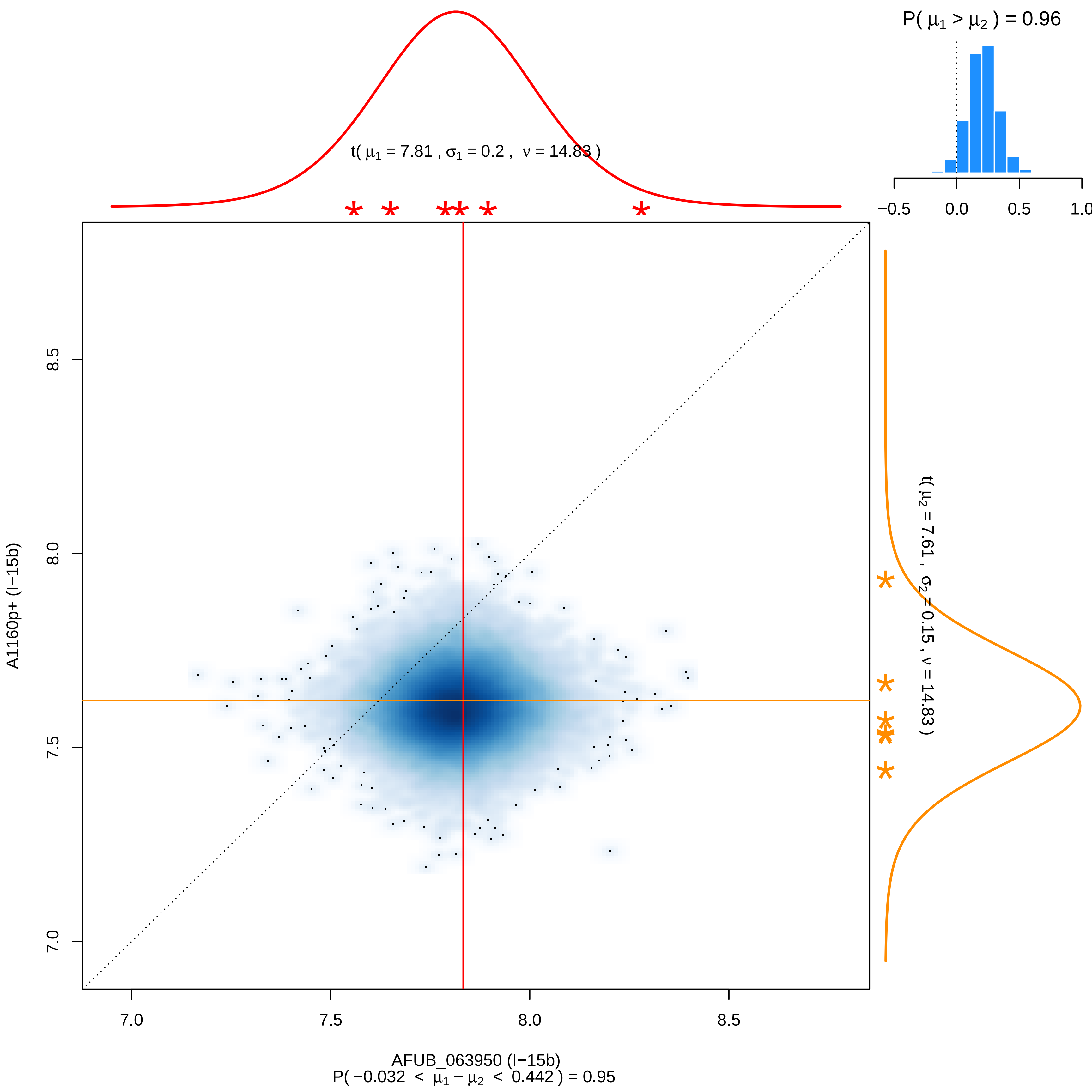

P( μ1 > μ2 ) = 0.96
−0.5
0.0
0.5
1.0
t( μ1 = 7.81 , σ1 = 0.2 , ν = 14.83 )
t( μ2 = 7.61 , σ2 = 0.15 , ν = 14.83 )
A1160p+ (I−15b)
8.5
8.0
7.5
7.0
AFUB_063950 (I−15b)
P( −0.032 < μ1 − μ2 < 0.442 ) = 0.95

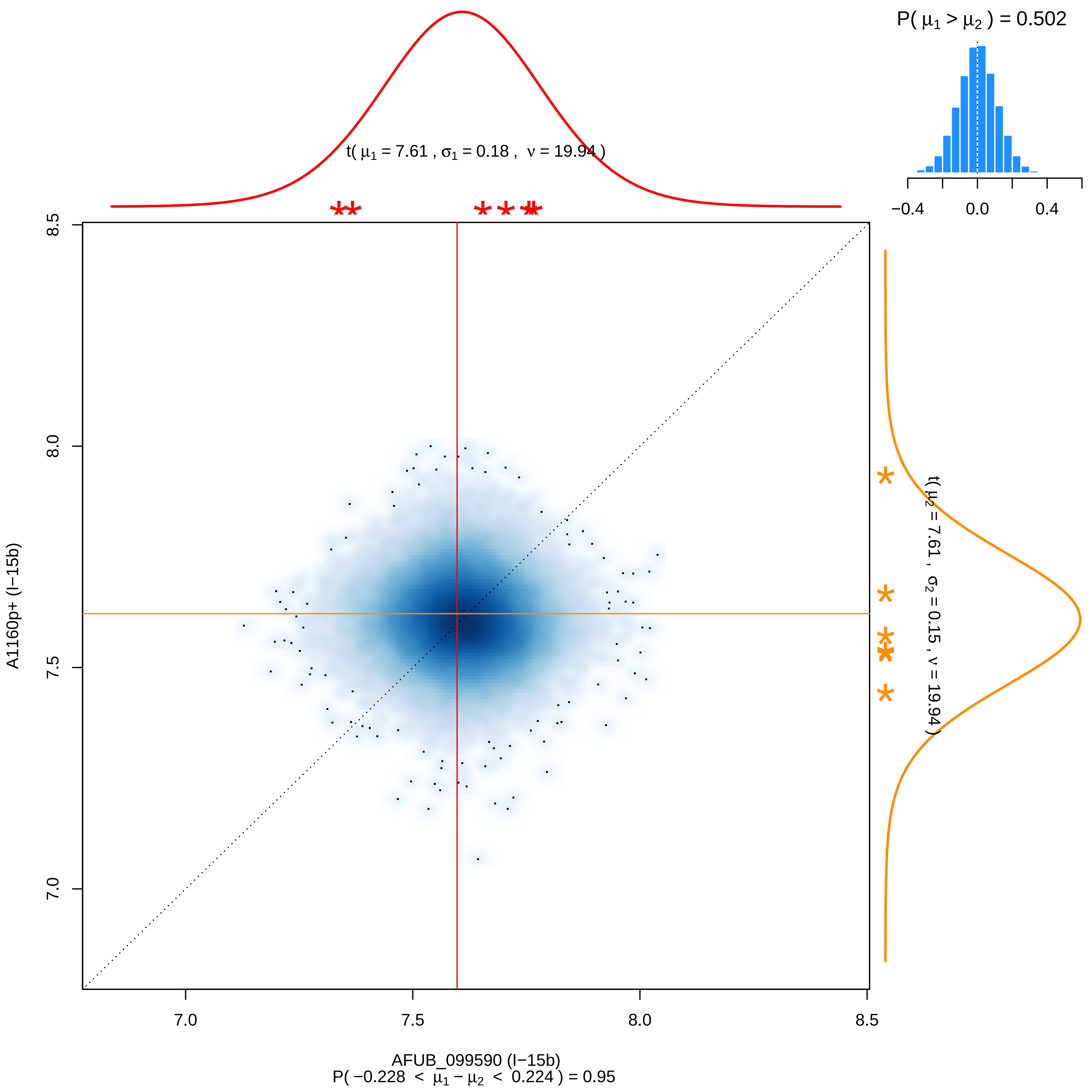

P( μ1 > μ2 ) = 0.502
−0.4
0.0
0.4
t( μ1 = 7.61 , σ1 = 0.18 , ν = 19.94 )
t( μ2 = 7.61 , σ2 = 0.15 , ν = 19.94 )
8.5
8.0
7.5
7.0
A1160p+ (I−15b)
7.0
7.5
8.0
8.5
AFUB_099590 (I−15b)
P( −0.228 < μ1 − μ2 < 0.224 ) = 0.95

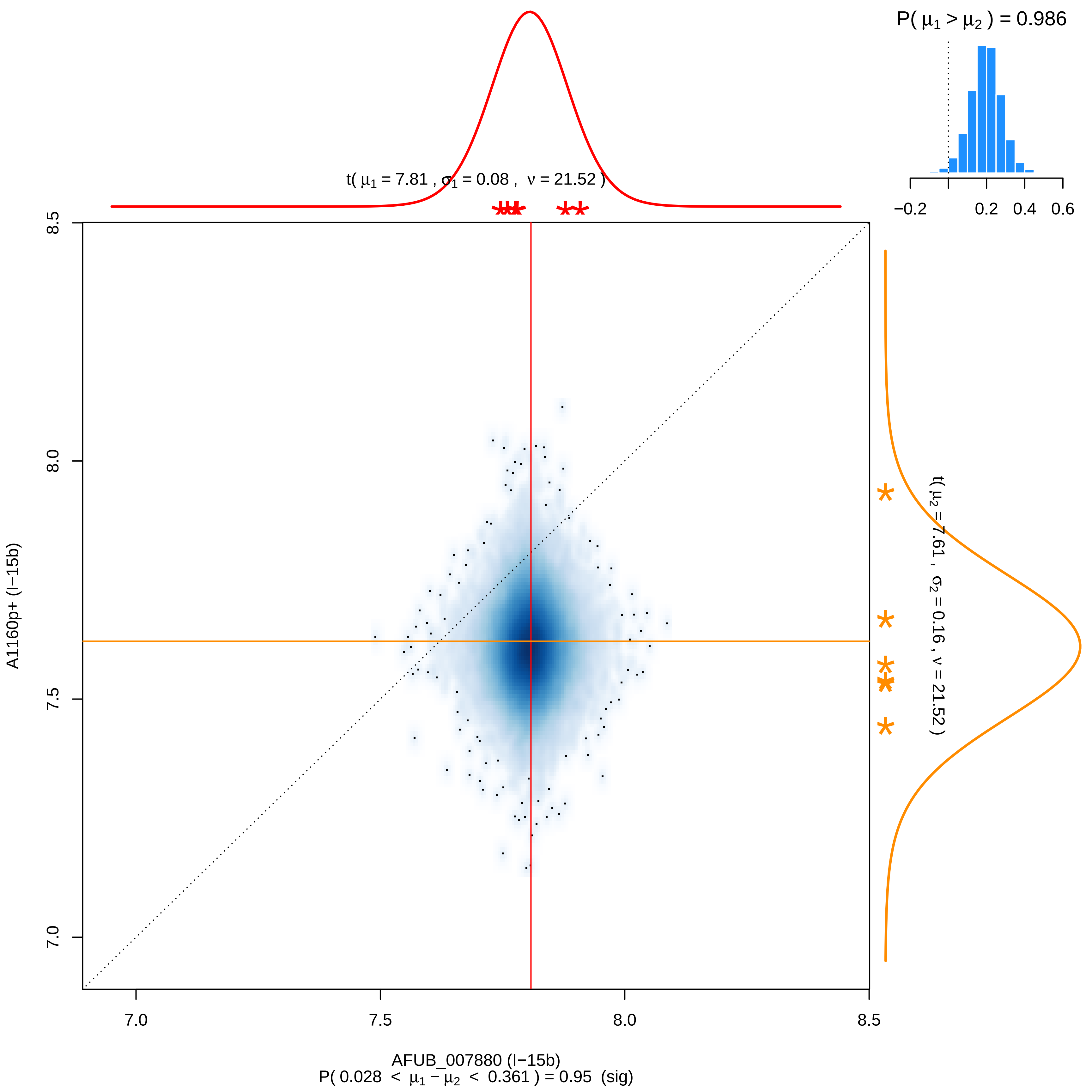

P( $\mu_1 > \mu_2$ ) = 0.986
−0.2
0.2
0.4
0.6
t( $\mu_1$ = 7.81 , $\sigma_1$ = 0.08 , $\nu$ = 21.52 )
t( $\mu_2$ = 7.61 , $\sigma_2$ = 0.16 , $\nu$ = 21.52 )
A1160p+ (I−15b)
8.5
8.0
7.5
7.0
AFUB_007880 (I−15b)
P( 0.028 < $\mu_1 - \mu_2$ < 0.361 ) = 0.95 (sig)

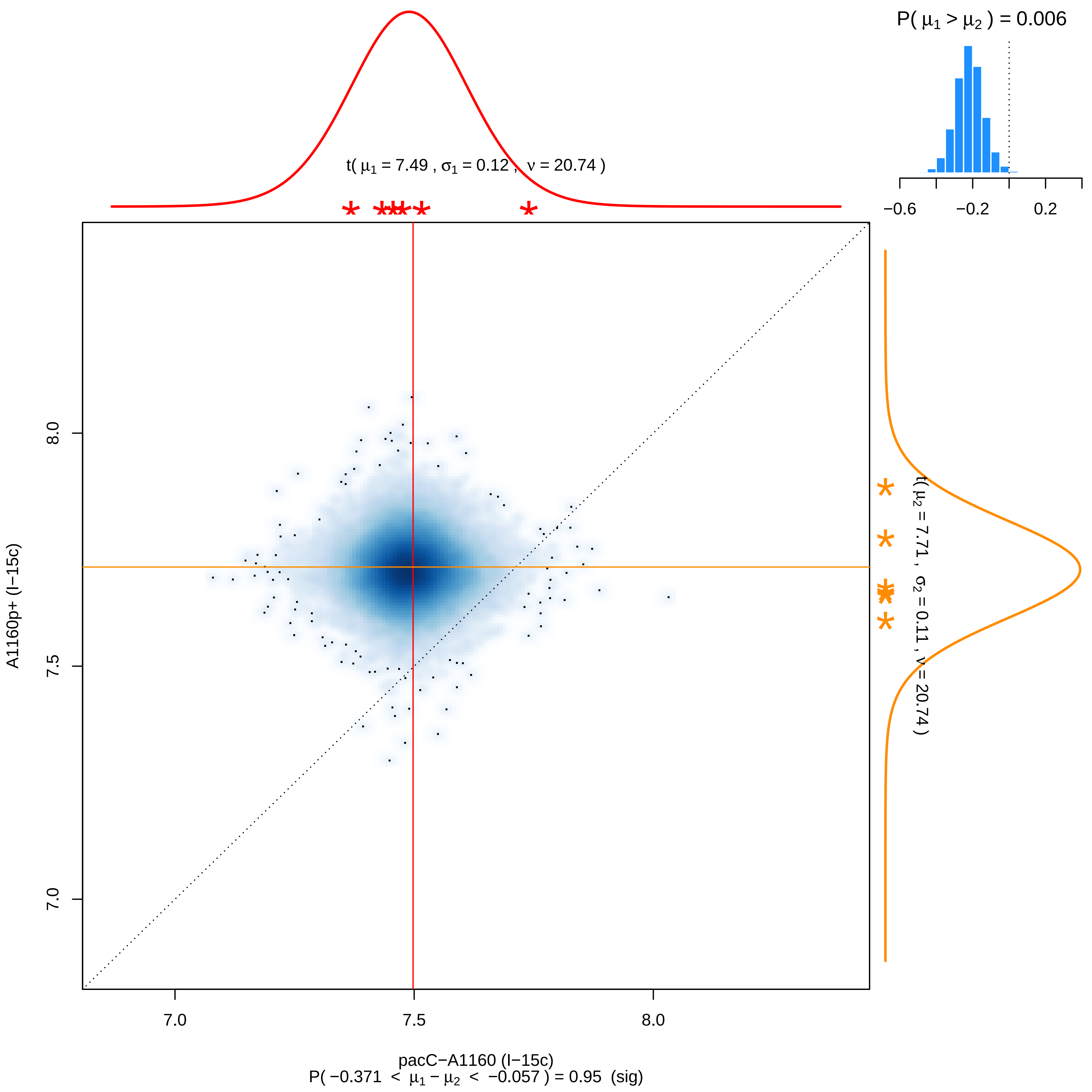

P( μ1 > μ2 ) = 0.006
−0.6
−0.2
0.2
t( μ1 = 7.49 , σ1 = 0.12 , ν = 20.74 )
t( μ2 = 7.71 , σ2 = 0.11 , ν = 20.74 )
A1160p+ (I−15c)
8.0
7.5
7.0
7.0
7.5
8.0
pacC−A1160 (I−15c)
P( −0.371 < μ1 − μ2 < −0.057 ) = 0.95 (sig)

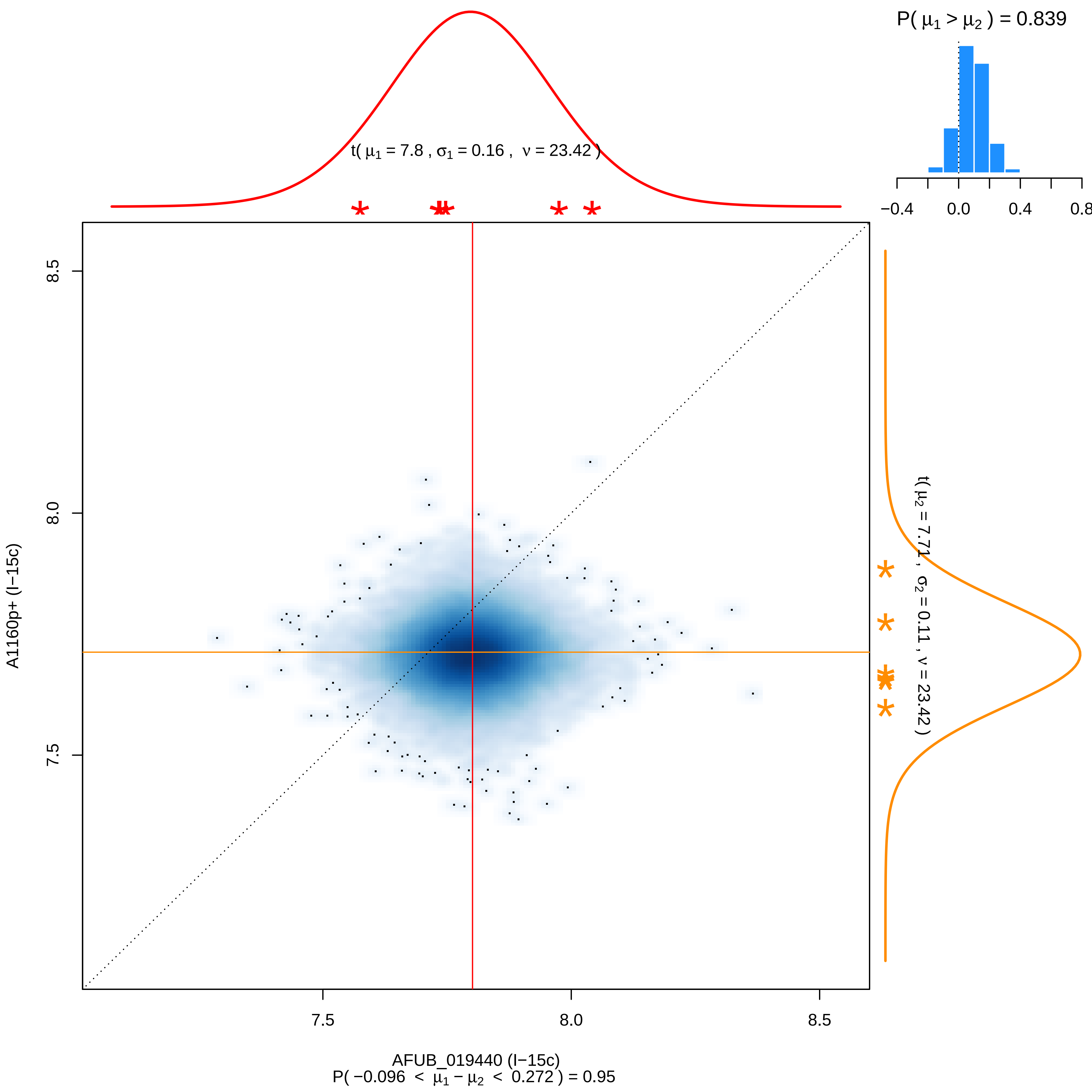
P( μ1 > μ2 ) = 0.839
−0.4
0.0
0.4
0.8
t( μ1 = 7.8 , σ1 = 0.16 , ν = 23.42 )
8.5
8.0
7.5
A1160p+ (I−15c)
t( μ2 = 7.71 , σ2 = 0.11 , ν = 23.42 )
7.5
8.0
8.5
AFUB_019440 (I−15c)
P( −0.096 < μ1 − μ2 < 0.272 ) = 0.95

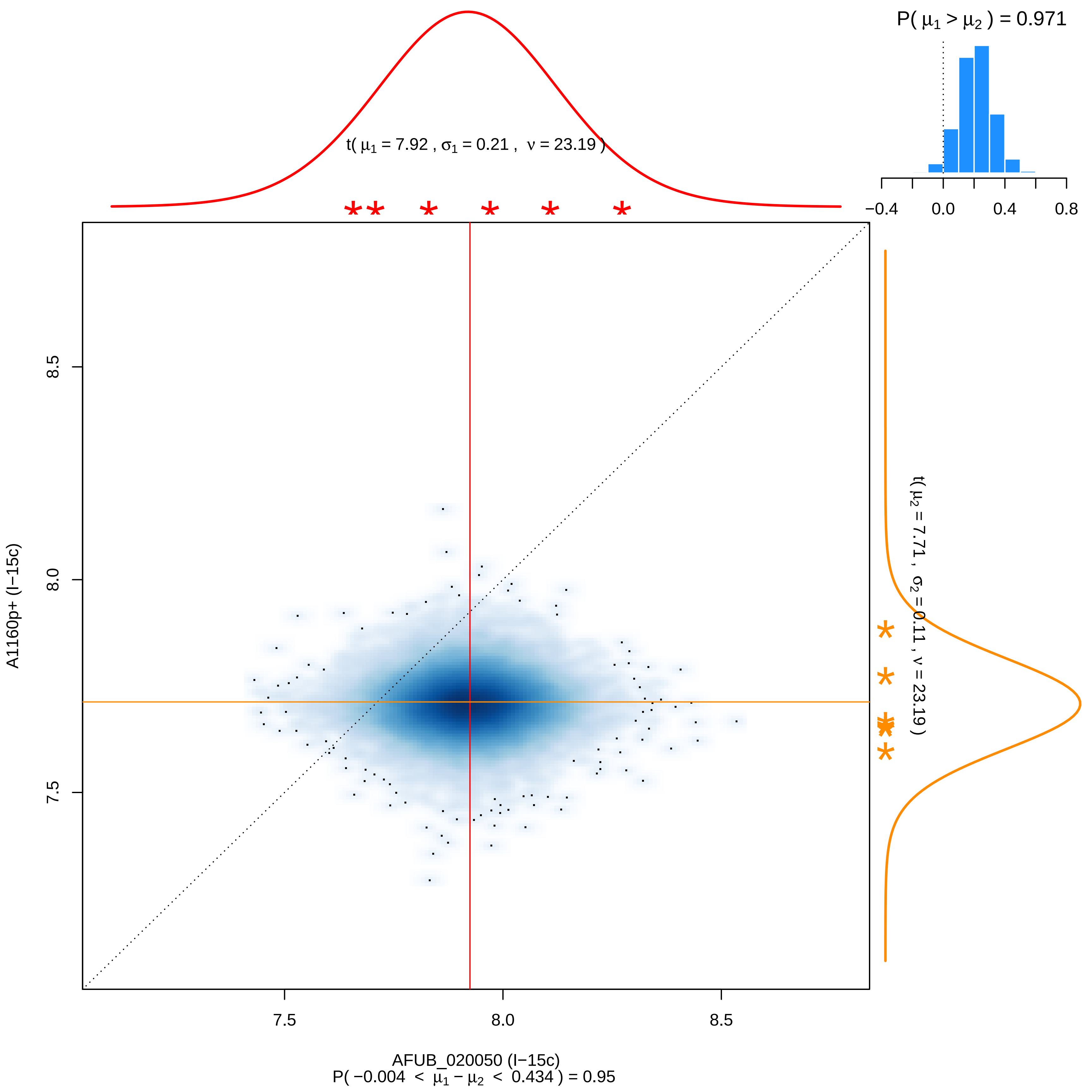

P( μ1 > μ2 ) = 0.971
−0.4
0.0
0.4
0.8
t( μ1 = 7.92 , σ1 = 0.21 , ν = 23.19 )
t( μ2 = 7.71 , σ2 = 0.11 , ν = 23.19 )
8.5
8.0
7.5
A1160p+ (I−15c)
AFUB_020050 (I−15c)
P( −0.004 < μ1 − μ2 < 0.434 ) = 0.95

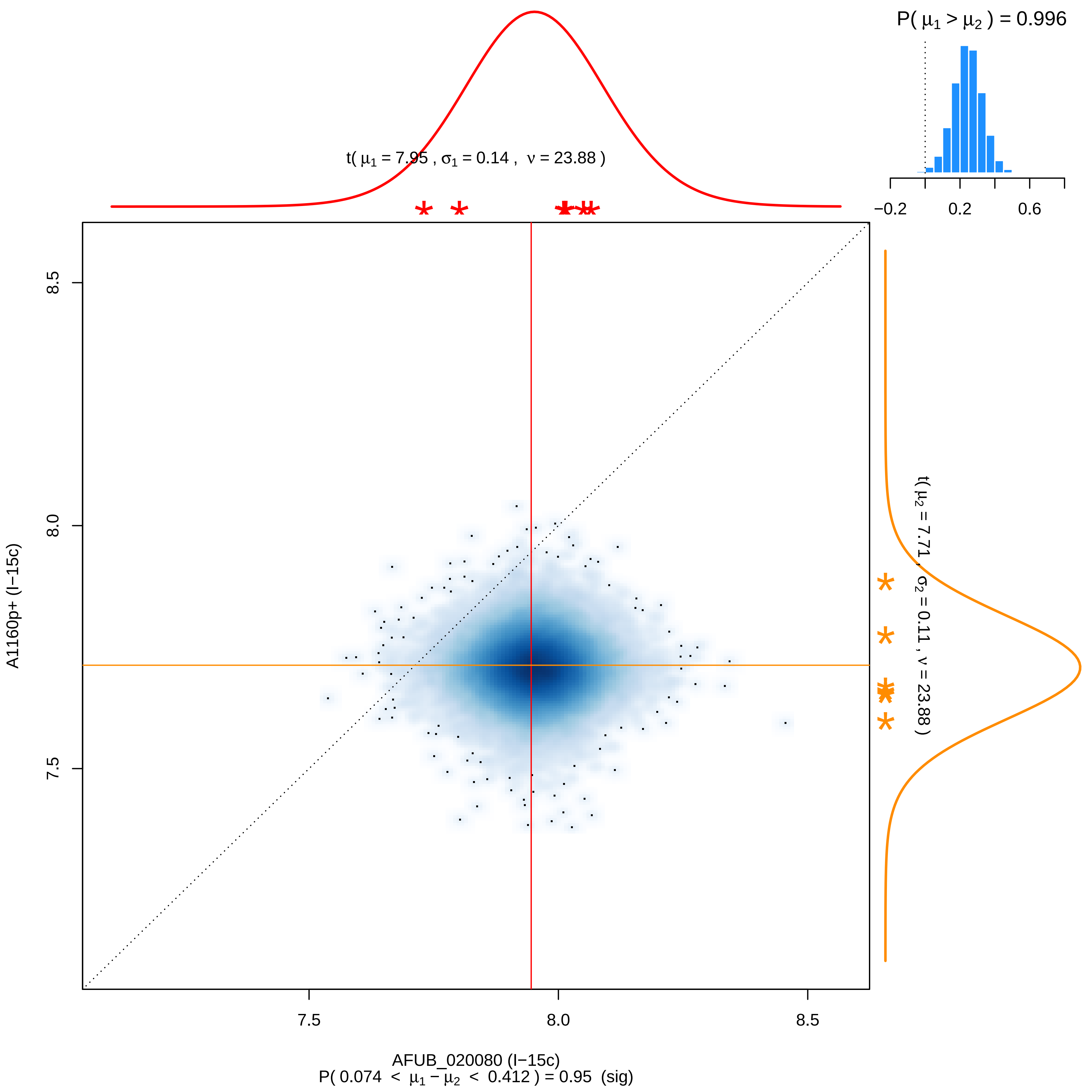

P( μ1 > μ2 ) = 0.996
−0.2
0.2
0.6
t( μ1 = 7.95 , σ1 = 0.14 , ν = 23.88 )
8.5
8.0
7.5
A1160p+ (I−15c)
t( μ2 = 7.71 , σ2 = 0.11 , ν = 23.88 )
7.5
8.0
8.5
AFUB_020080 (I−15c)
P( 0.074 < μ1 − μ2 < 0.412 ) = 0.95 (sig)

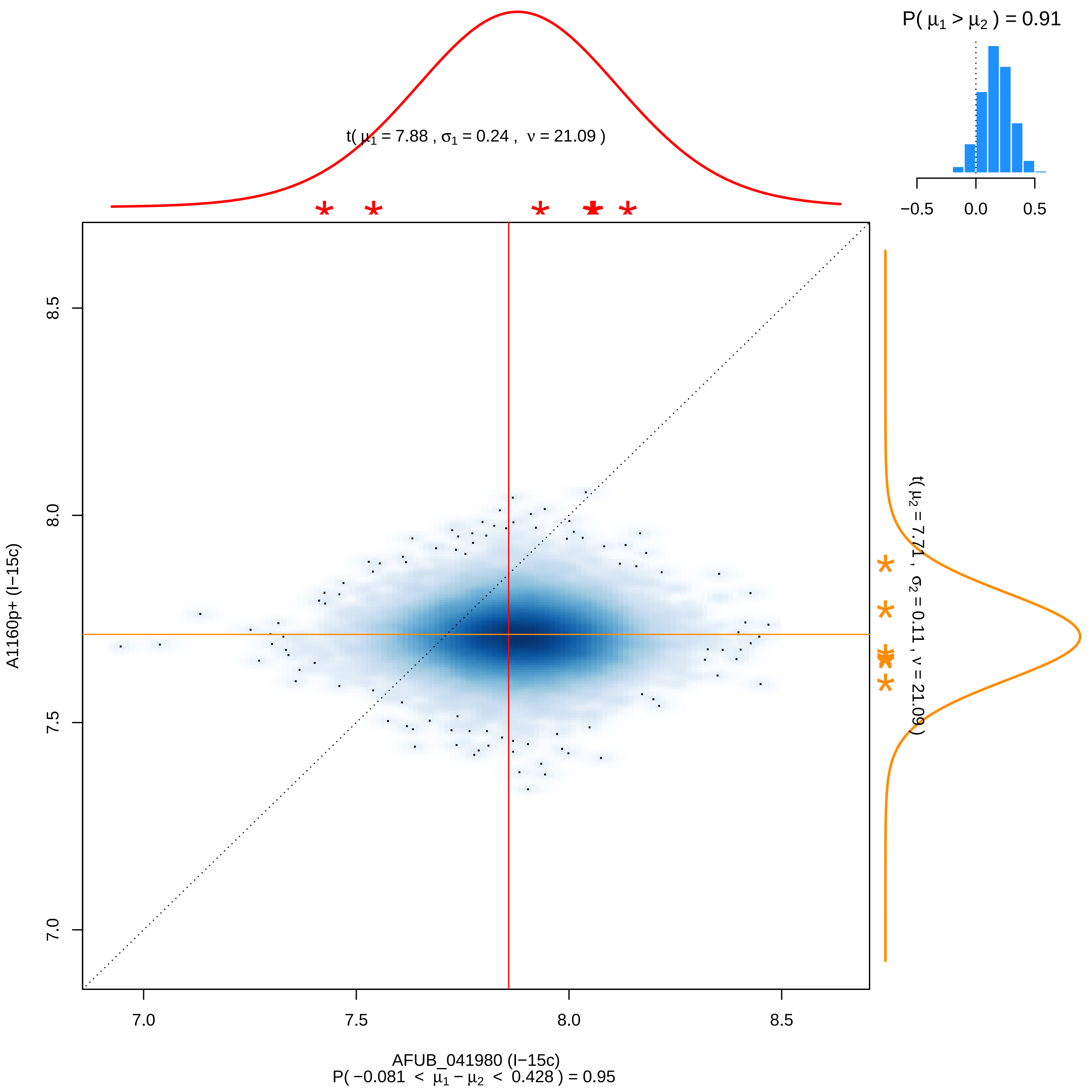
P( μ1 > μ2 ) = 0.91
−0.5
0.0
0.5
t( μ1 = 7.88 , σ1 = 0.24 , ν = 21.09 )
t( μ2 = 7.71 , σ2 = 0.11 , ν = 21.09 )
A1160p+ (I−15c)
8.5
8.0
7.5
7.0
7.0
7.5
8.0
8.5
AFUB_041980 (I−15c)
P( −0.081 < μ1 − μ2 < 0.428 ) = 0.95

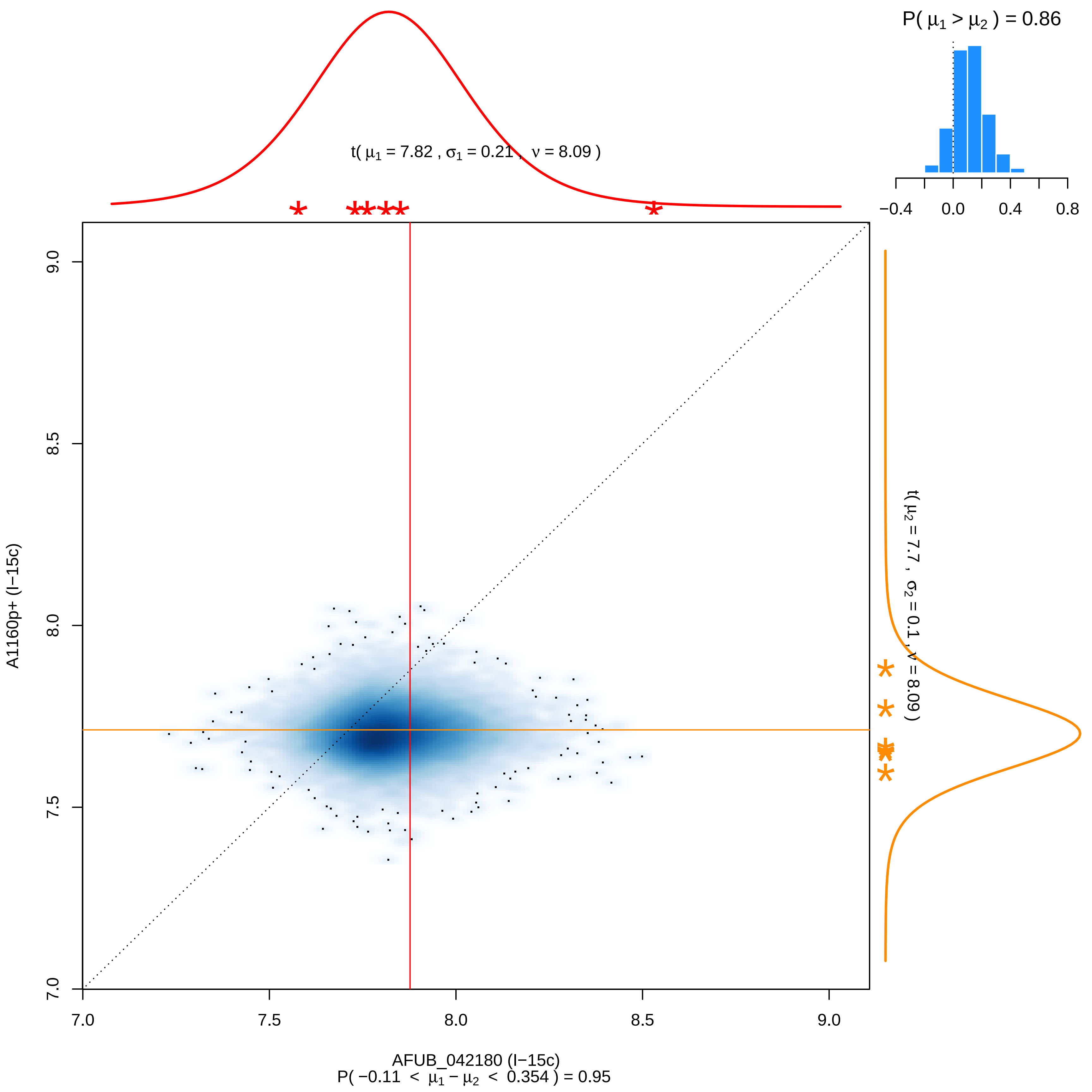

P( μ1 > μ2 ) = 0.86
−0.4
0.0
0.4
0.8
t( μ1 = 7.82 , σ1 = 0.21 , ν = 8.09 )
t( μ2 = 7.7 , σ2 = 0.1 , ν = 8.09 )
A1160p+ (I−15c)
7.0
7.5
8.0
8.5
9.0
AFUB_042180 (I−15c)
P( −0.11 < μ1 − μ2 < 0.354 ) = 0.95

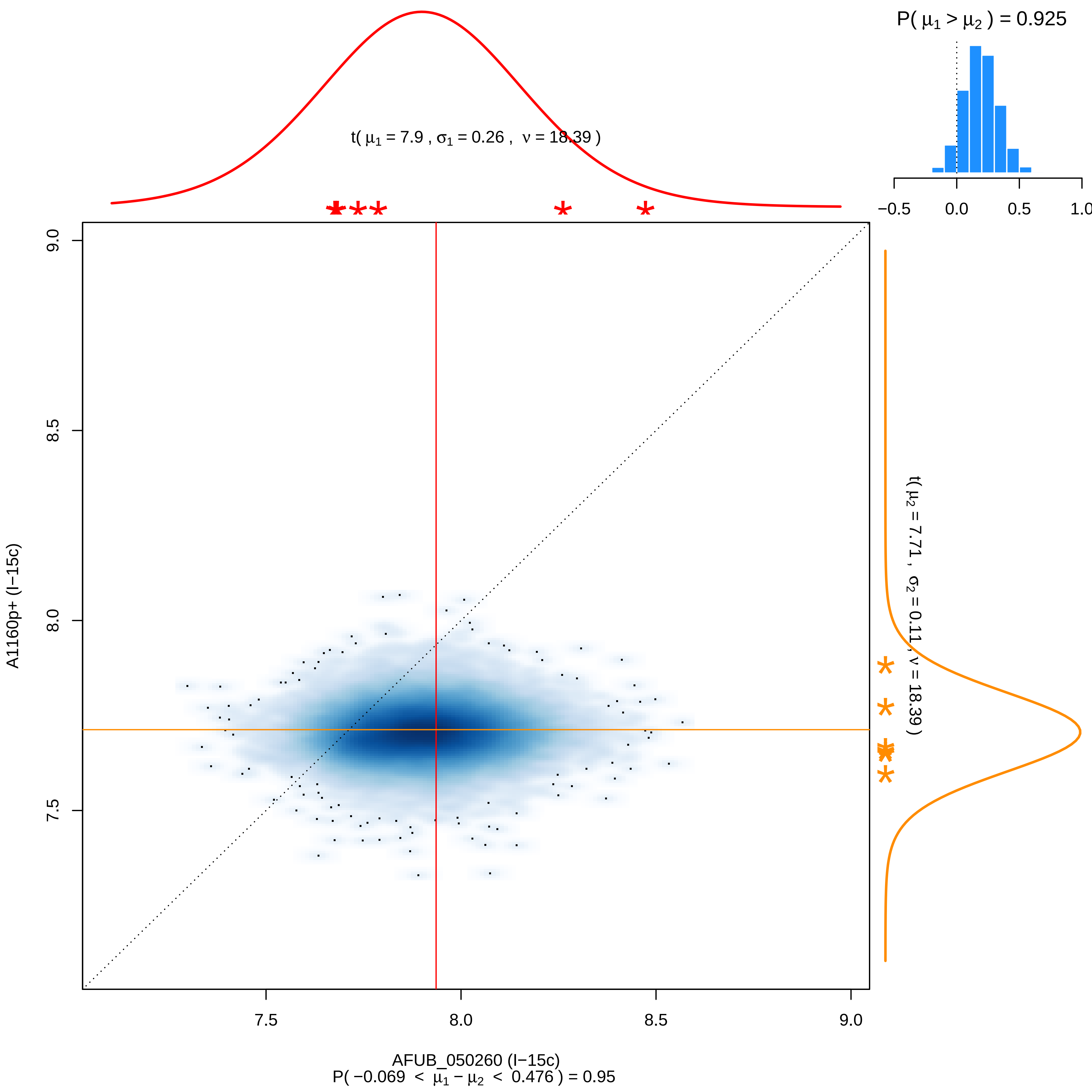

P( μ1 > μ2 ) = 0.925
−0.5
0.0
0.5
1.0
t( μ1 = 7.9 , σ1 = 0.26 , ν = 18.39 )
9.0
8.5
8.0
7.5
A1160p+ (I−15c)
t( μ2 = 7.71 , σ2 = 0.11 , ν = 18.39 )
7.5
8.0
8.5
9.0
AFUB_050260 (I−15c)
P( −0.069 < μ1 − μ2 < 0.476 ) = 0.95

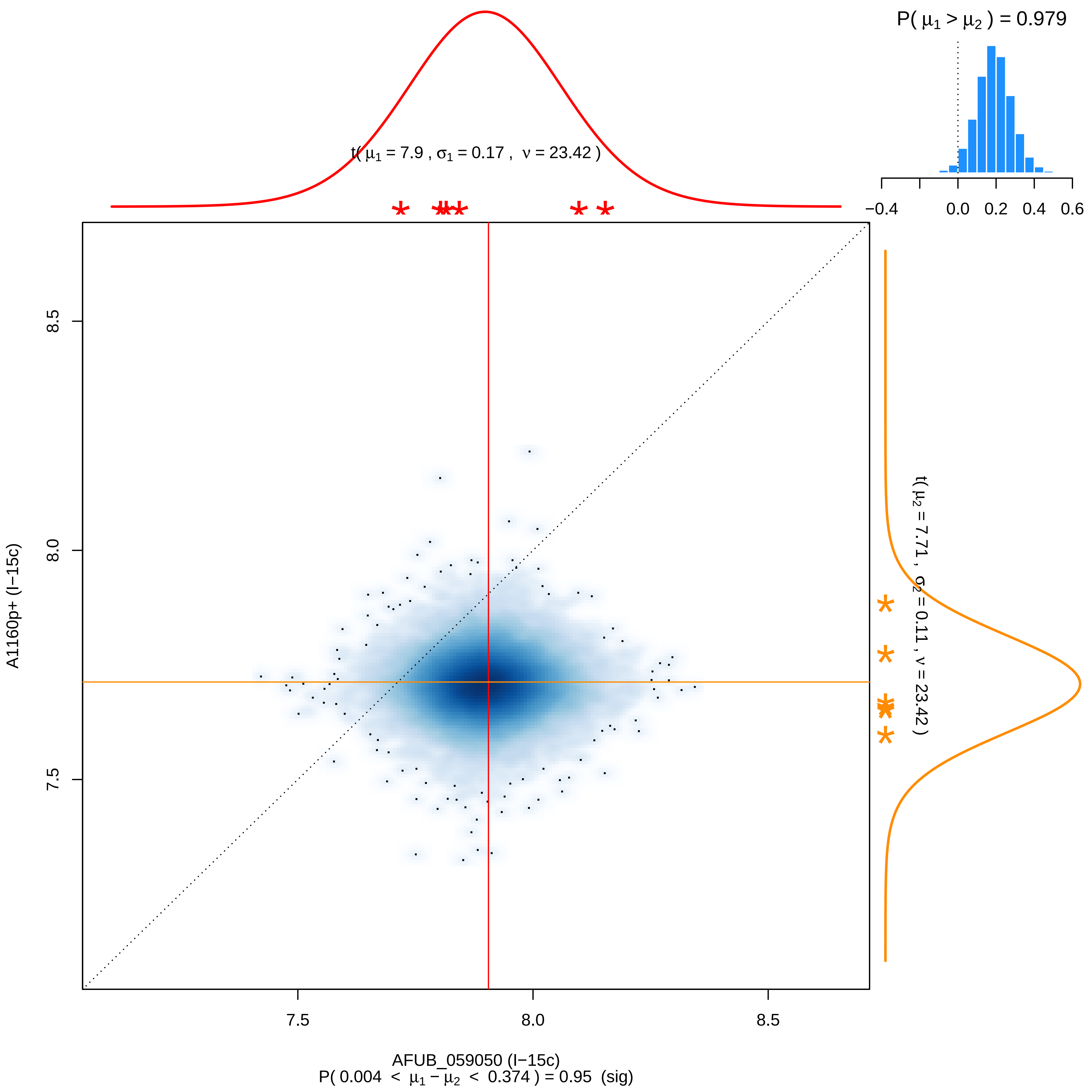

P( μ1 > μ2 ) = 0.979
−0.4
0.0
0.2
0.4
0.6
t( μ1 = 7.9 , σ1 = 0.17 , ν = 23.42 )
t( μ2 = 7.71 , σ2 = 0.11 , ν = 23.42 )
8.5
8.0
7.5
A1160p+ (I−15c)
7.5
8.0
8.5
AFUB_059050 (I−15c)
P( 0.004 < μ1 − μ2 < 0.374 ) = 0.95 (sig)

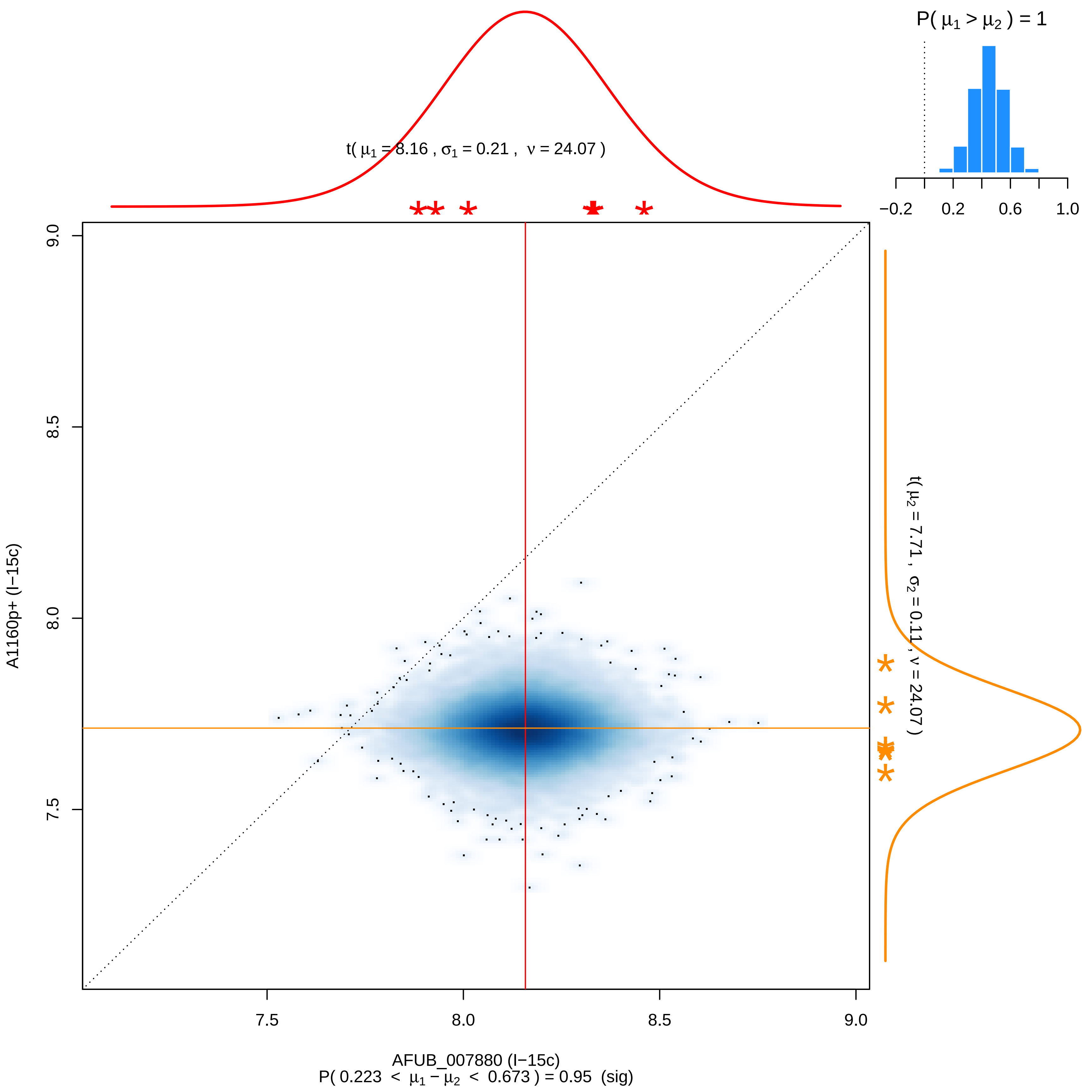

P( μ1 > μ2 ) = 1
−0.2
0.2
0.6
1.0
t( μ1 = 8.16 , σ1 = 0.21 , ν = 24.07 )
t( μ2 = 7.71 , σ2 = 0.11 , ν = 24.07 )
9.0
8.5
8.0
7.5
A1160p+ (I−15c)
7.5
8.0
8.5
9.0
AFUB_007880 (I−15c)
P( 0.223 < μ1 − μ2 < 0.673 ) = 0.95 (sig)

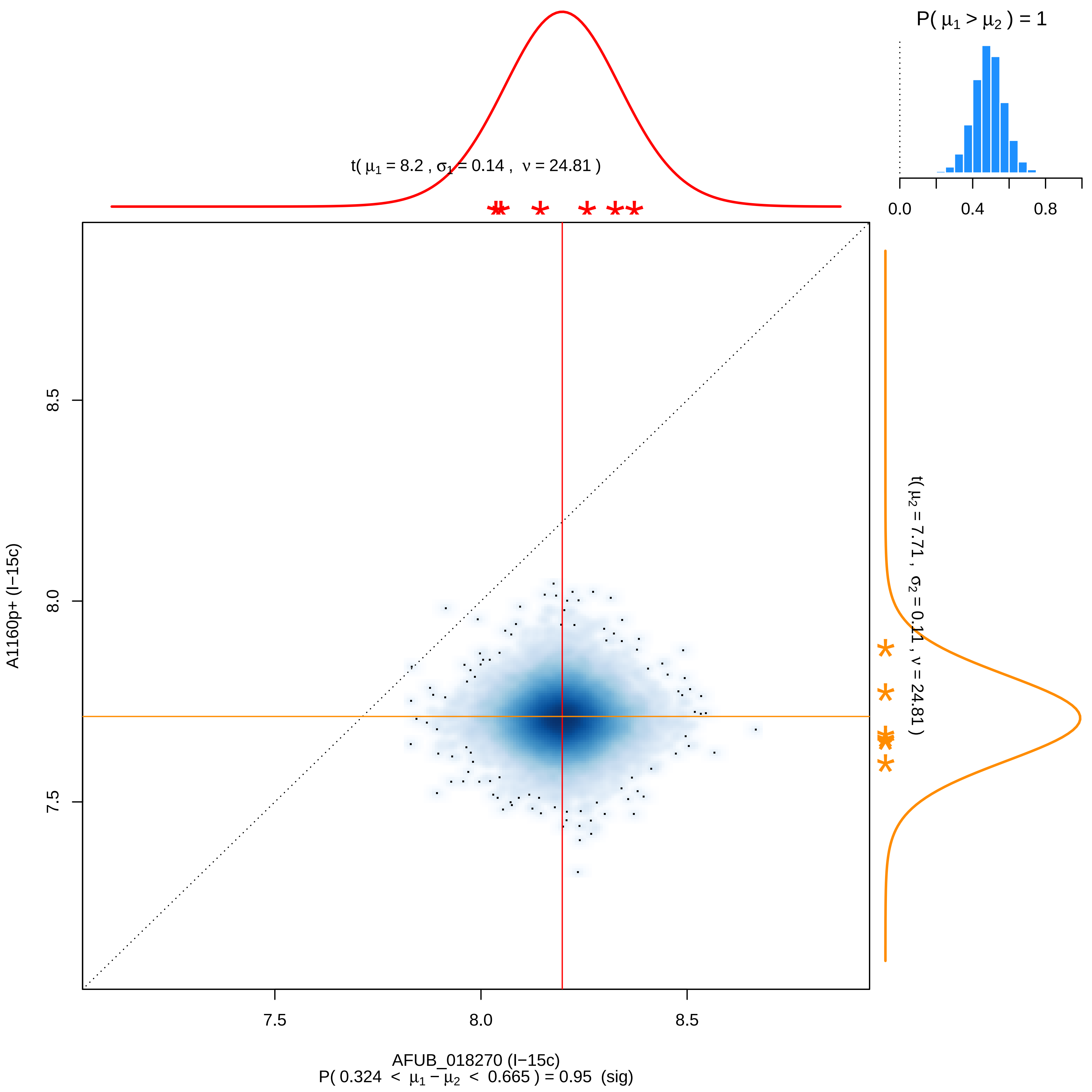
P( μ1 > μ2 ) = 1
0.0
0.4
0.8
t( μ1 = 8.2 , σ1 = 0.14 , ν = 24.81 )
8.5
8.0
7.5
A1160p+ (I−15c)
t( μ2 = 7.71 , σ2 = 0.11 , ν = 24.81 )
7.5
8.0
8.5
AFUB_018270 (I−15c)
P( 0.324 < μ1 − μ2 < 0.665 ) = 0.95 (sig)

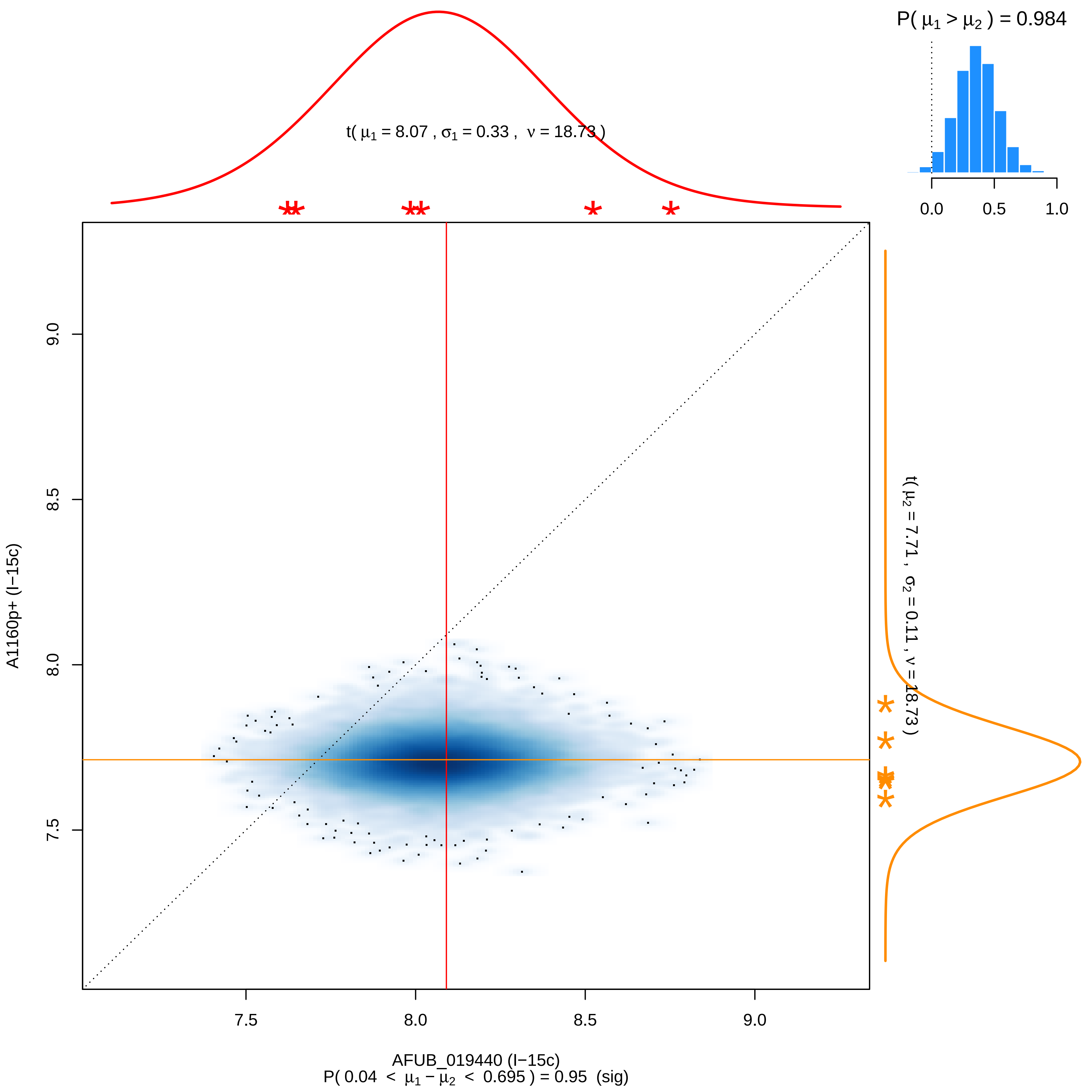

P( $\mu_1 > \mu_2$ ) = 0.984
0.0
0.5
1.0
t( $\mu_1$ = 8.07 , $\sigma_1$ = 0.33 , $\nu$ = 18.73 )
t( $\mu_2$ = 7.71 , $\sigma_2$ = 0.11 , $\nu$ = 18.73 )
9.0
8.5
8.0
7.5
A1160p+ (I−15c)
7.5
8.0
8.5
9.0
AFUB_019440 (I−15c)
P( 0.04 < $\mu_1 - \mu_2$ < 0.695 ) = 0.95 (sig)

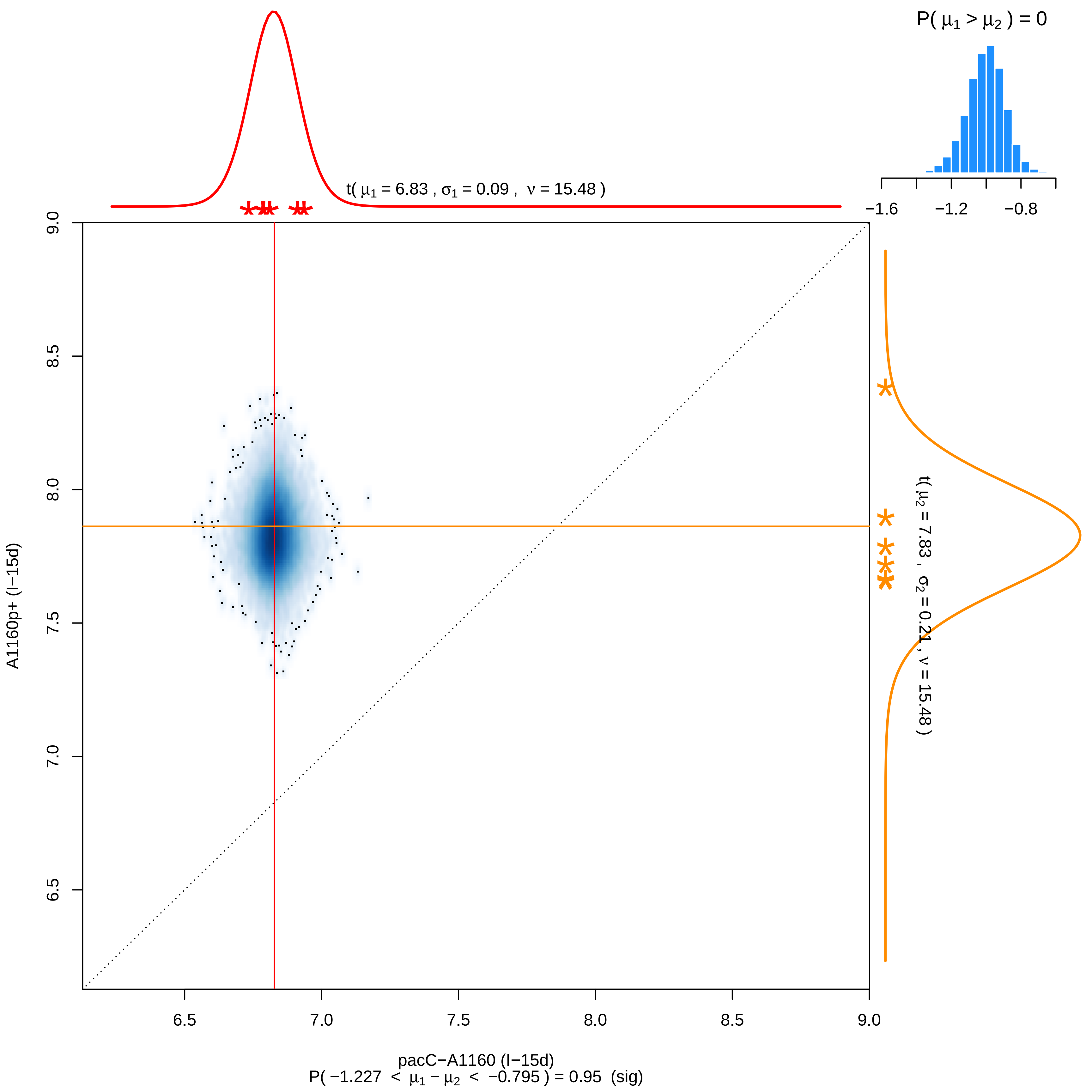

P( μ1 > μ2 ) = 0
−1.6
−1.2
−0.8
t( μ1 = 6.83 , σ1 = 0.09 , ν = 15.48 )
t( μ2 = 7.83 , σ2 = 0.21 , ν = 15.48 )
9.0
8.5
8.0
7.5
7.0
6.5
A1160p+ (I−15d)
6.5
7.0
7.5
8.0
8.5
9.0
pacC−A1160 (I−15d)
P( −1.227 < μ1 − μ2 < −0.795 ) = 0.95 (sig)

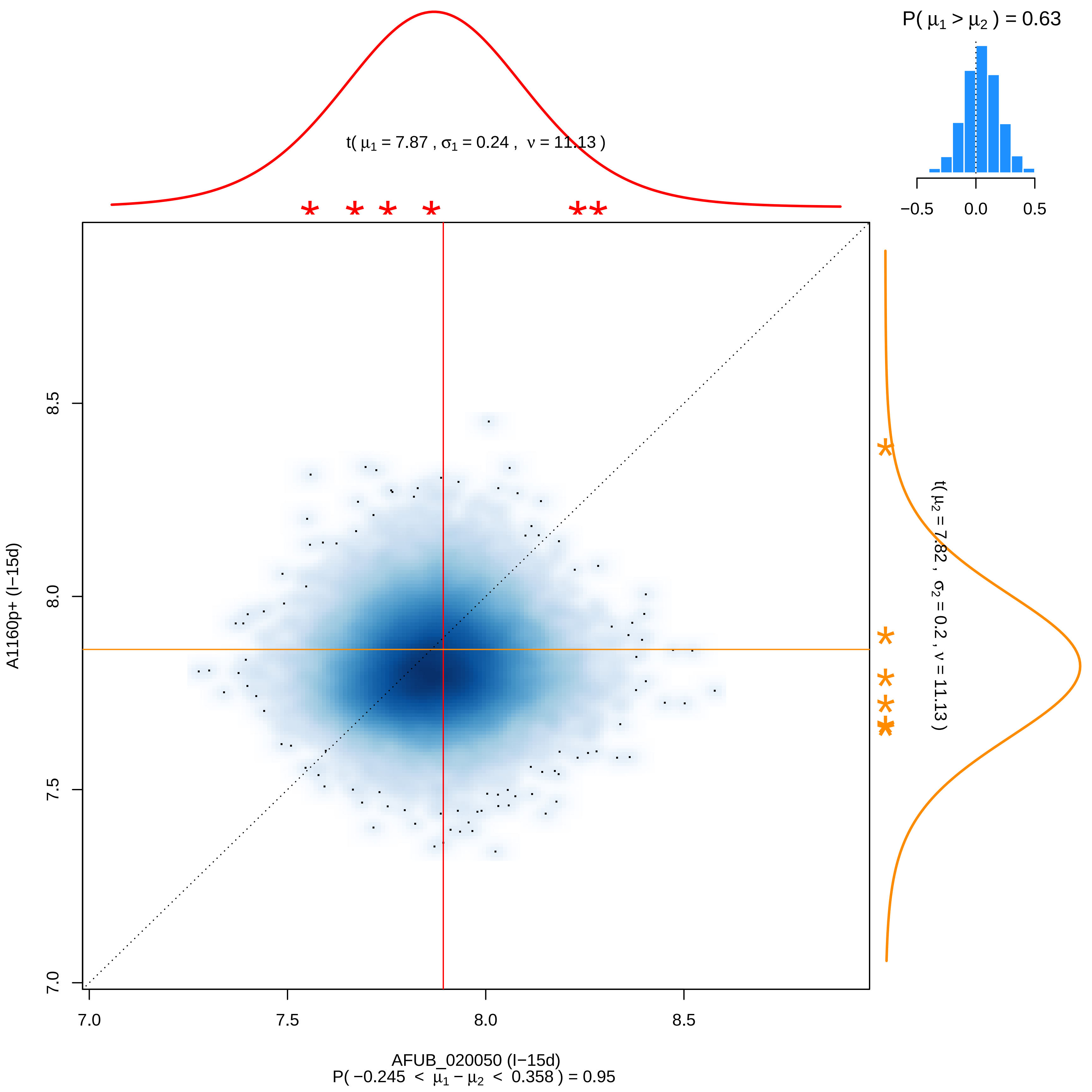

P( μ1 > μ2 ) = 0.63
−0.5 0.0 0.5
t( μ1 = 7.87 , σ1 = 0.24 , ν = 11.13 )
t( μ2 = 7.82 , σ2 = 0.2 , ν = 11.13 )
A1160p+ (I−15d)
8.5
8.0
7.5
7.0
7.0 7.5 8.0 8.5
AFUB_020050 (I−15d)
P( −0.245 < μ1 − μ2 < 0.358 ) = 0.95

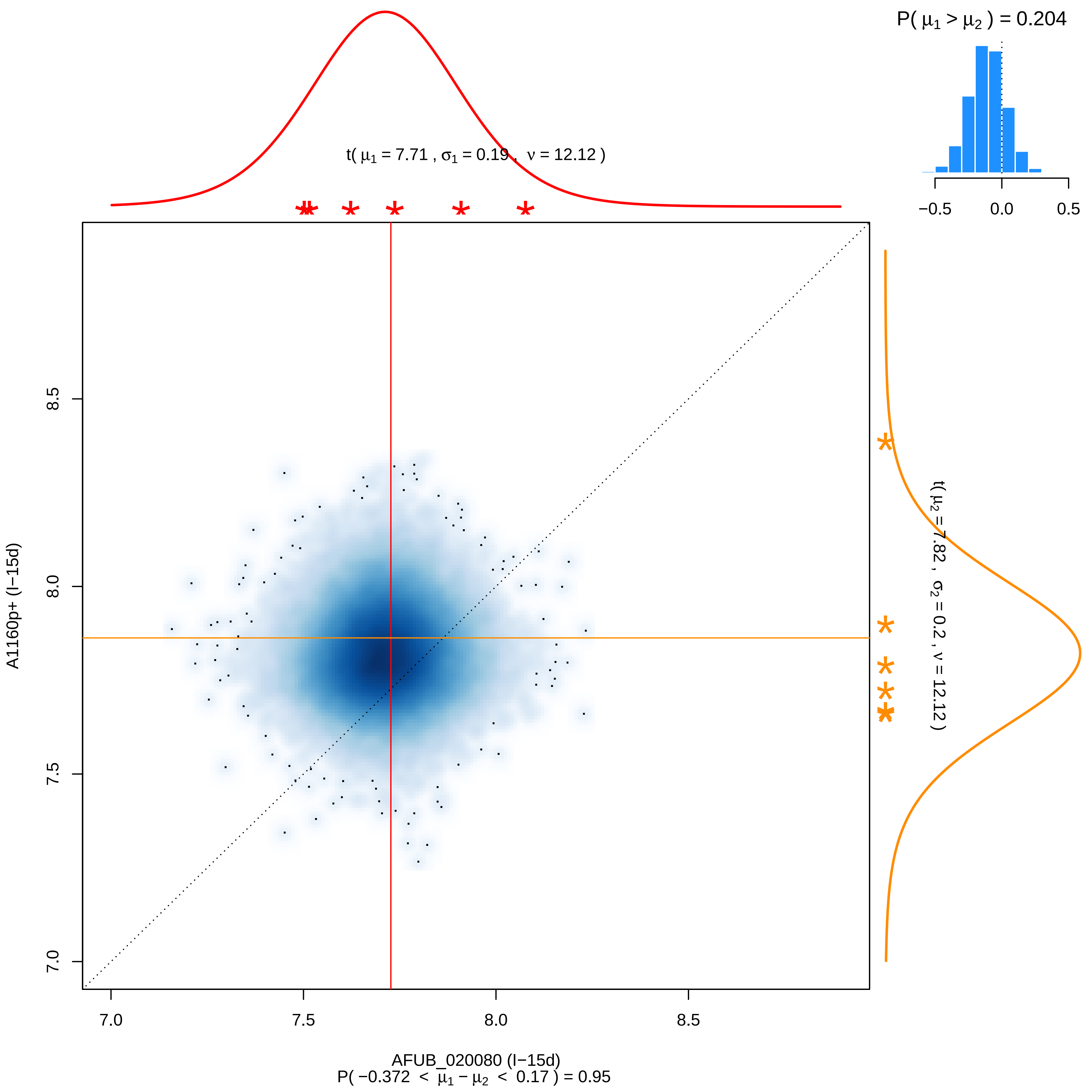
P( μ1 > μ2 ) = 0.204
−0.5
0.0
0.5
t( μ1 = 7.71 , σ1 = 0.19 , ν = 12.12 )
t( μ2 = 7.82 , σ2 = 0.2 , ν = 12.12 )
A1160p+ (I−15d)
8.5
8.0
7.5
7.0
AFUB_020080 (I−15d)
P( −0.372 < μ1 − μ2 < 0.17 ) = 0.95

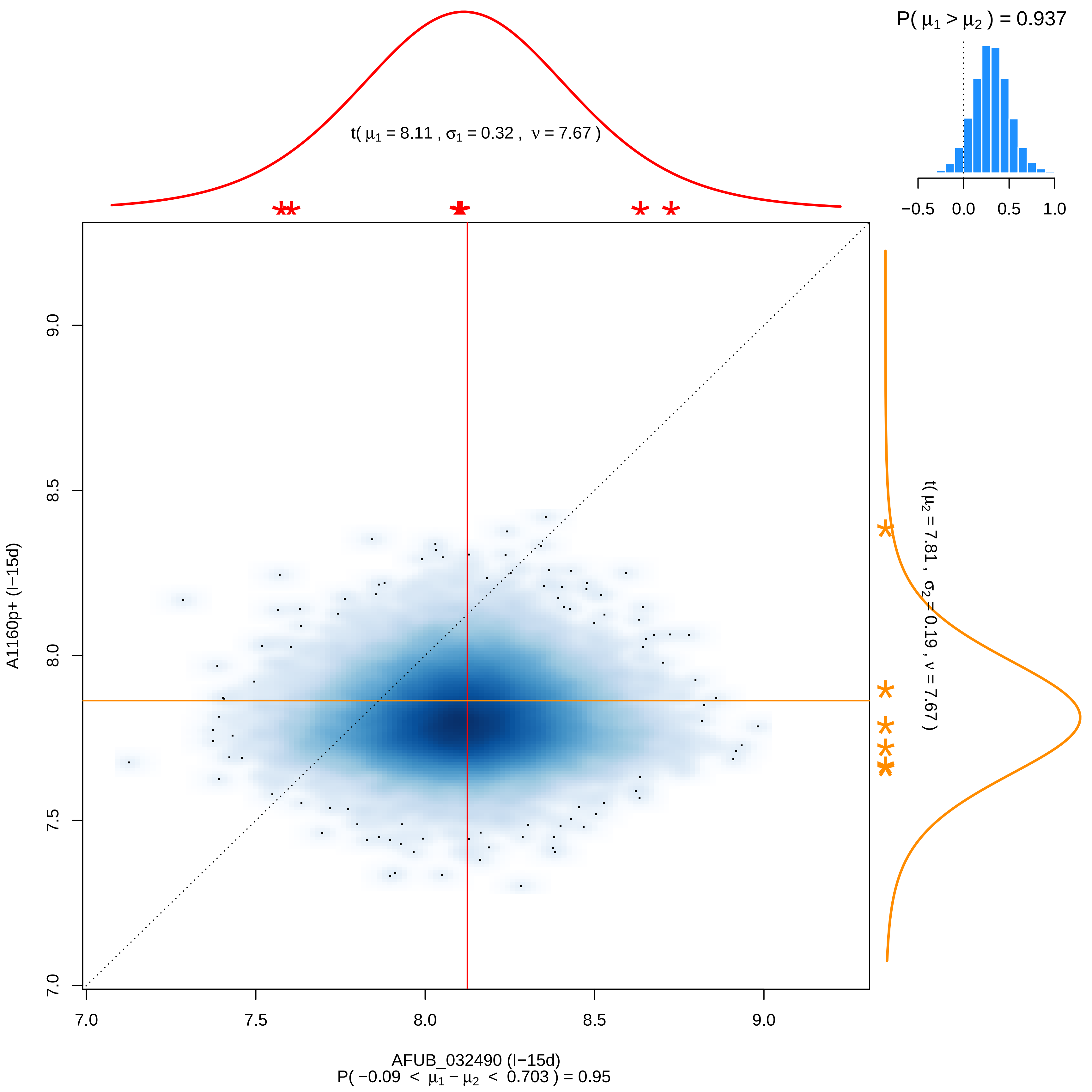

P( μ1 > μ2 ) = 0.937
−0.5
0.0
0.5
1.0
t( μ1 = 8.11 , σ1 = 0.32 , ν = 7.67 )
t( μ2 = 7.81 , σ2 = 0.19 , ν = 7.67 )
A1160p+ (I−15d)
7.0
7.5
8.0
8.5
9.0
AFUB_032490 (I−15d)
P( −0.09 < μ1 − μ2 < 0.703 ) = 0.95

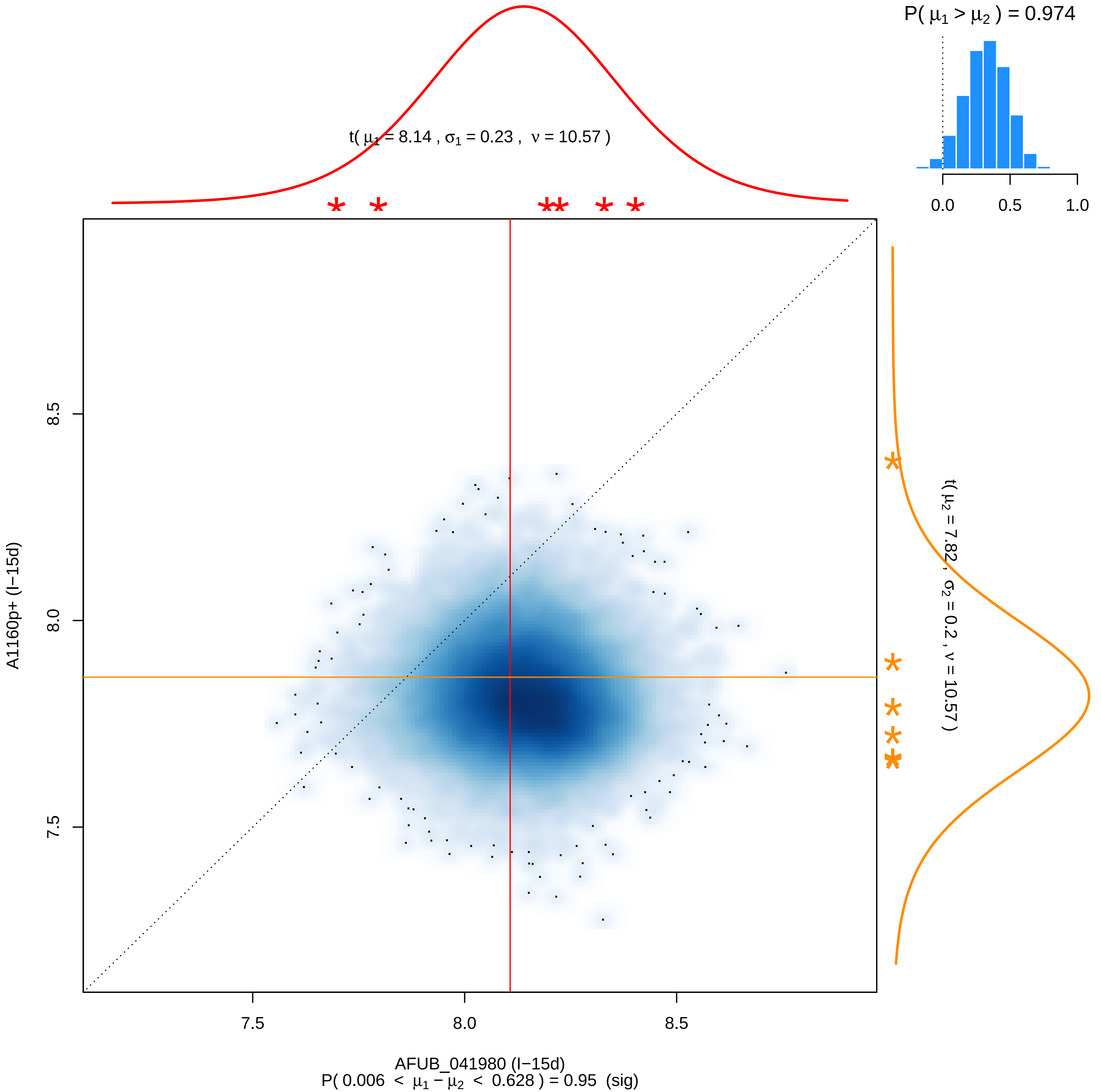

P( μ1 > μ2 ) = 0.974
0.0
0.5
1.0
t( μ1 = 8.14 , σ1 = 0.23 , ν = 10.57 )
t( μ2 = 7.82 , σ2 = 0.2 , ν = 10.57 )
8.5
8.0
7.5
A1160p+ (I−15d)
7.5
8.0
8.5
AFUB_041980 (I−15d)
P( 0.006 < μ1 − μ2 < 0.628 ) = 0.95 (sig)

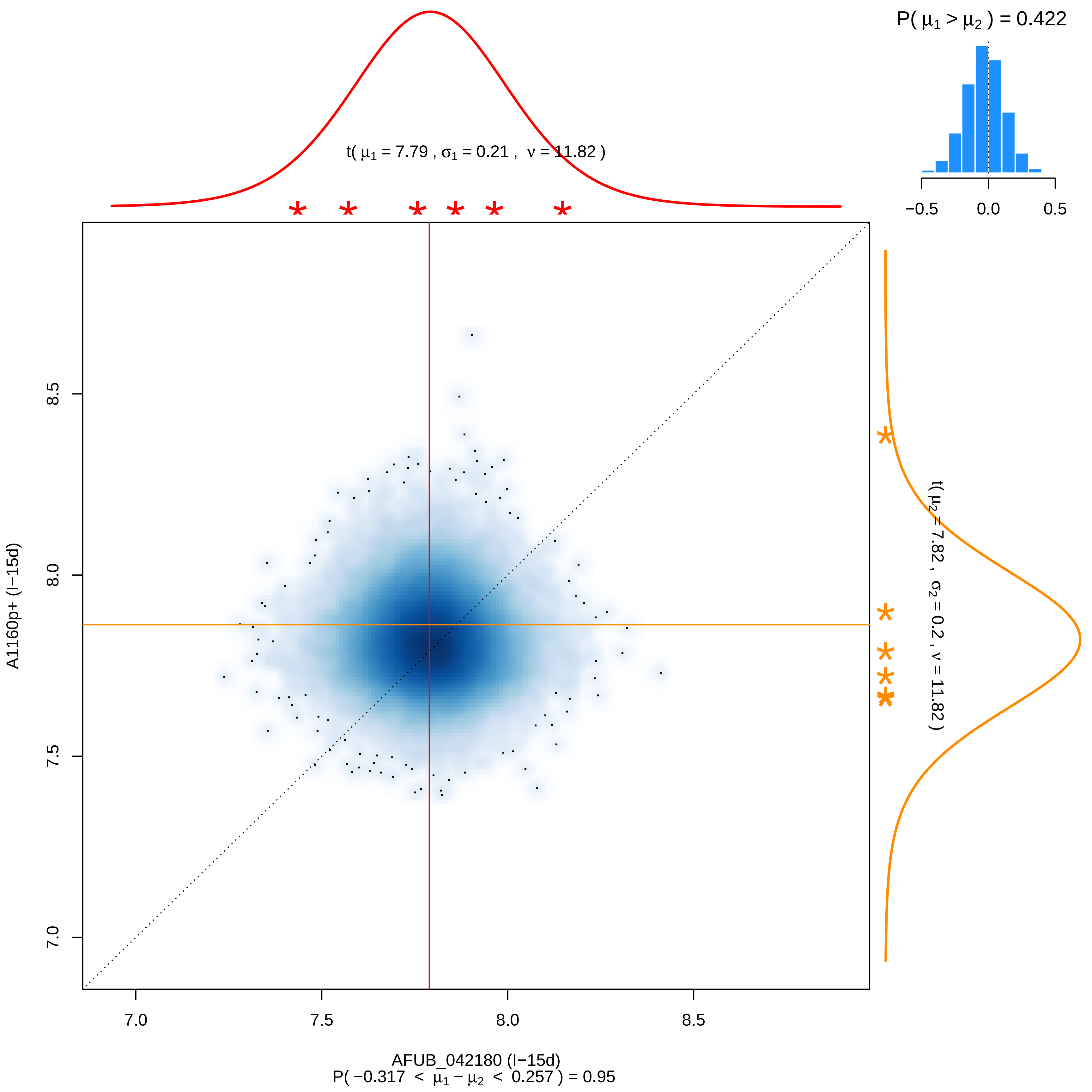

P( μ1 > μ2 ) = 0.422
−0.5
0.0
0.5
t( μ1 = 7.79 , σ1 = 0.21 , ν = 11.82 )
t( μ2 = 7.82 , σ2 = 0.2 , ν = 11.82 )
A1160p+ (I−15d)
8.5
8.0
7.5
7.0
7.0
7.5
8.0
8.5
AFUB_042180 (I−15d)
P( −0.317 < μ1 − μ2 < 0.257 ) = 0.95

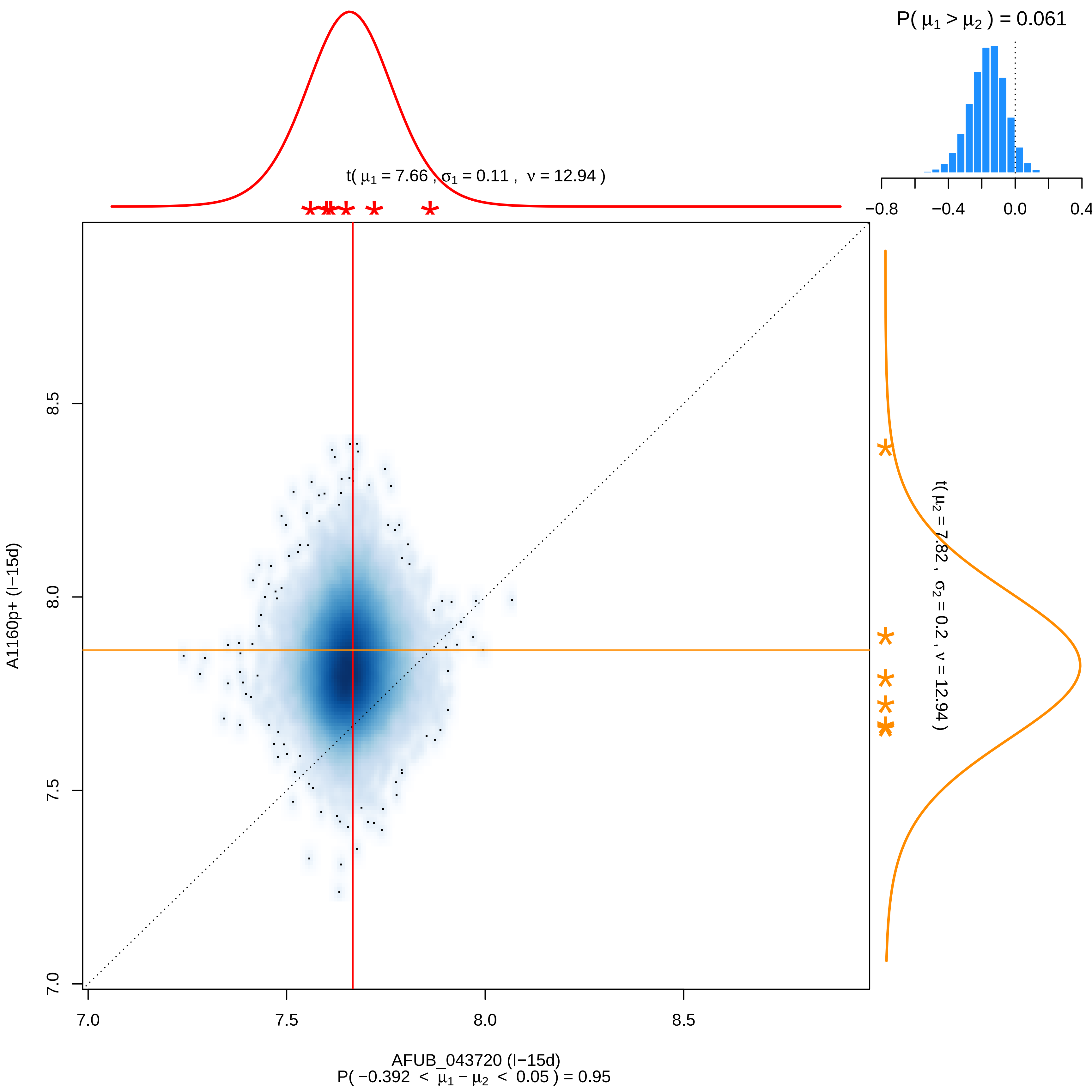
P( μ1 > μ2 ) = 0.061
−0.8
−0.4
0.0
0.4
t( μ1 = 7.66 , σ1 = 0.11 , ν = 12.94 )
t( μ2 = 7.82 , σ2 = 0.2 , ν = 12.94 )
A1160p+ (I−15d)
8.5
8.0
7.5
7.0
7.0
7.5
8.0
8.5
AFUB_043720 (I−15d)
P( −0.392 < μ1 − μ2 < 0.05 ) = 0.95

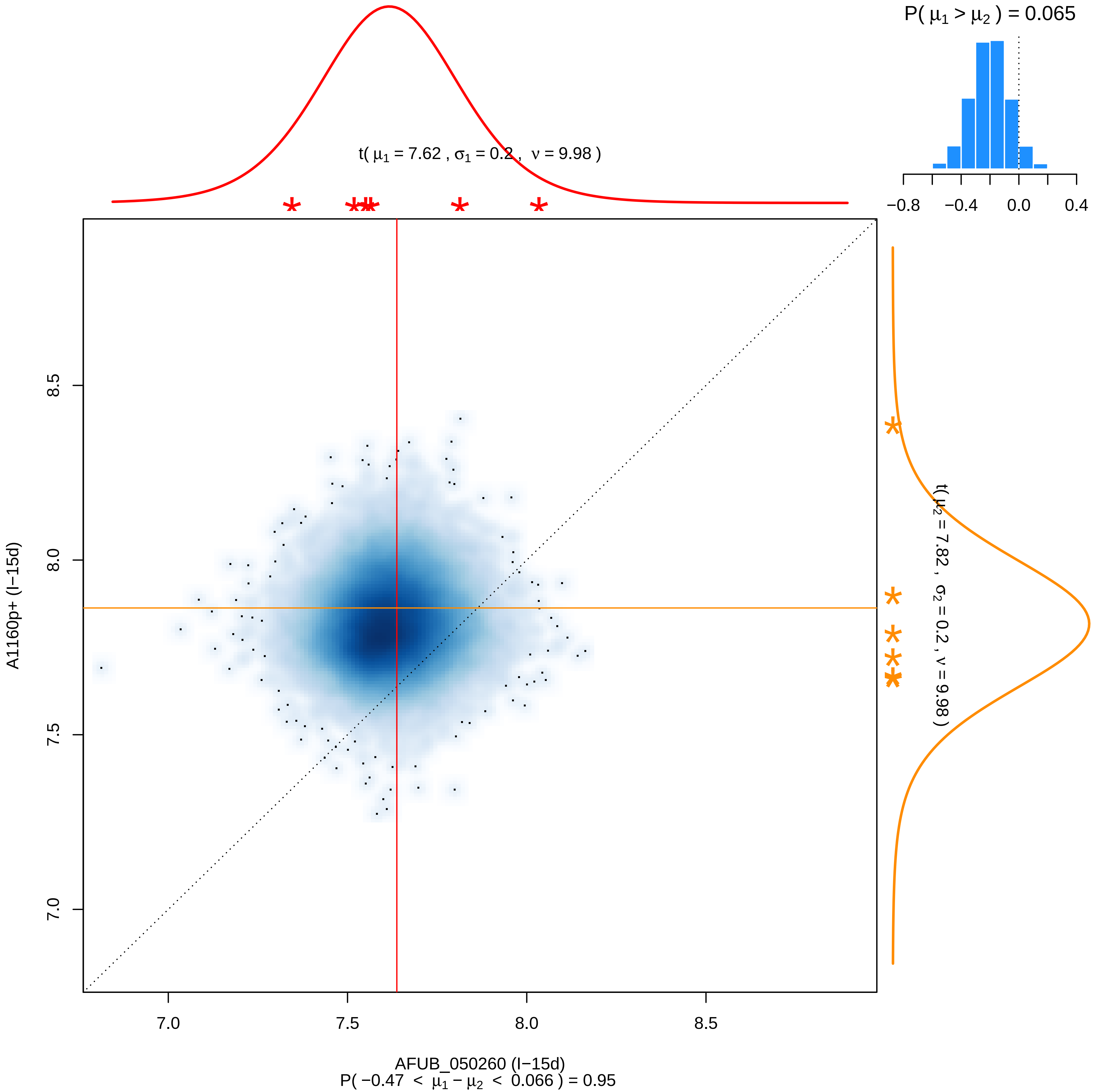

P( μ1 > μ2 ) = 0.065
−0.8
−0.4
0.0
0.4
t( μ1 = 7.62 , σ1 = 0.2 , ν = 9.98 )
t( μ2 = 7.82 , σ2 = 0.2 , ν = 9.98 )
A1160p+ (I−15d)
8.5
8.0
7.5
7.0
AFUB_050260 (I−15d)
P( −0.47 < μ1 − μ2 < 0.066 ) = 0.95

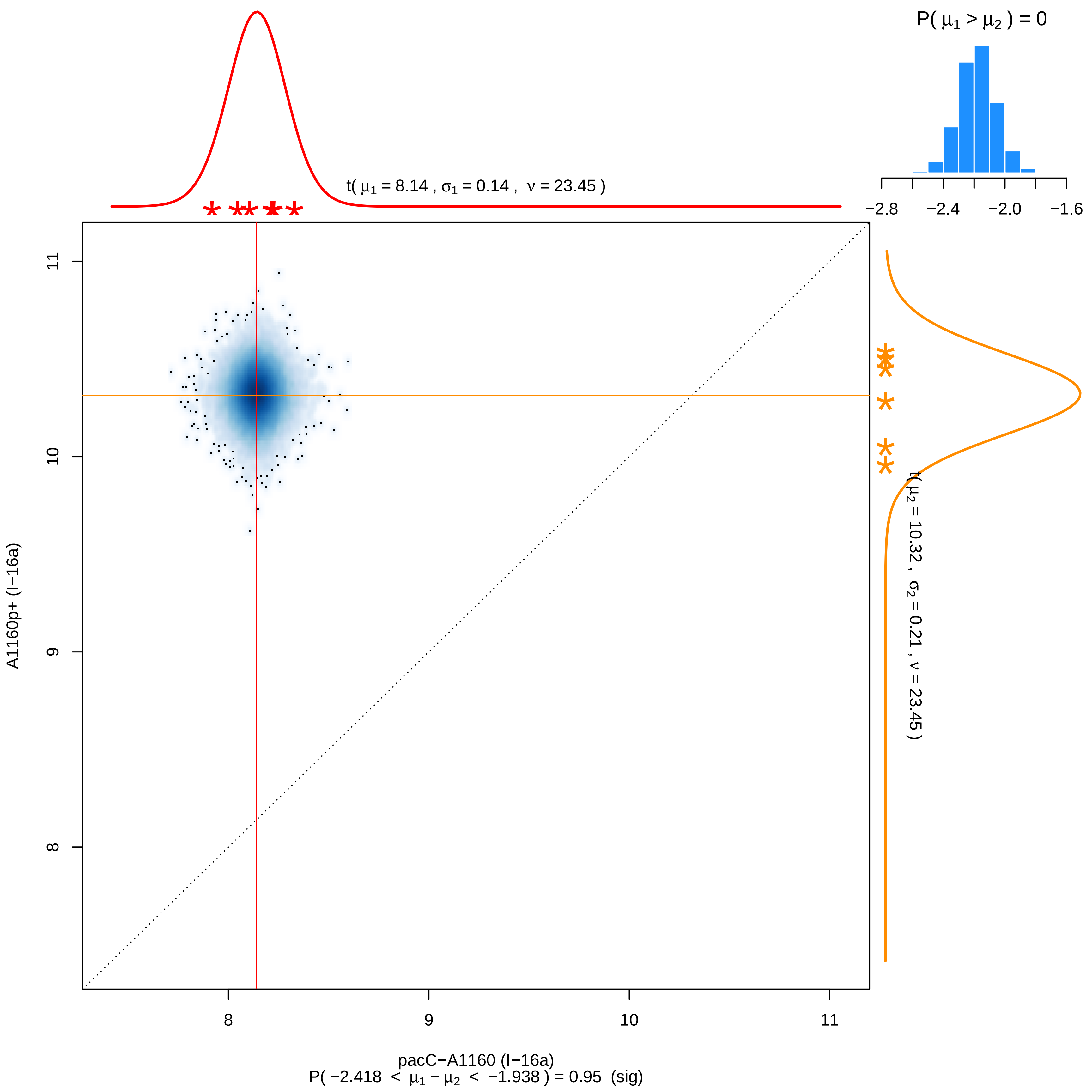

P( μ1 > μ2 ) = 0
−2.8
−2.4
−2.0
−1.6
t( μ1 = 8.14 , σ1 = 0.14 , ν = 23.45 )
t( μ2 = 10.32 , σ2 = 0.21 , ν = 23.45 )
A1160p+ (I−16a)
8
9
10
11
pacC−A1160 (I−16a)
P( −2.418 < μ1 − μ2 < −1.938 ) = 0.95 (sig)

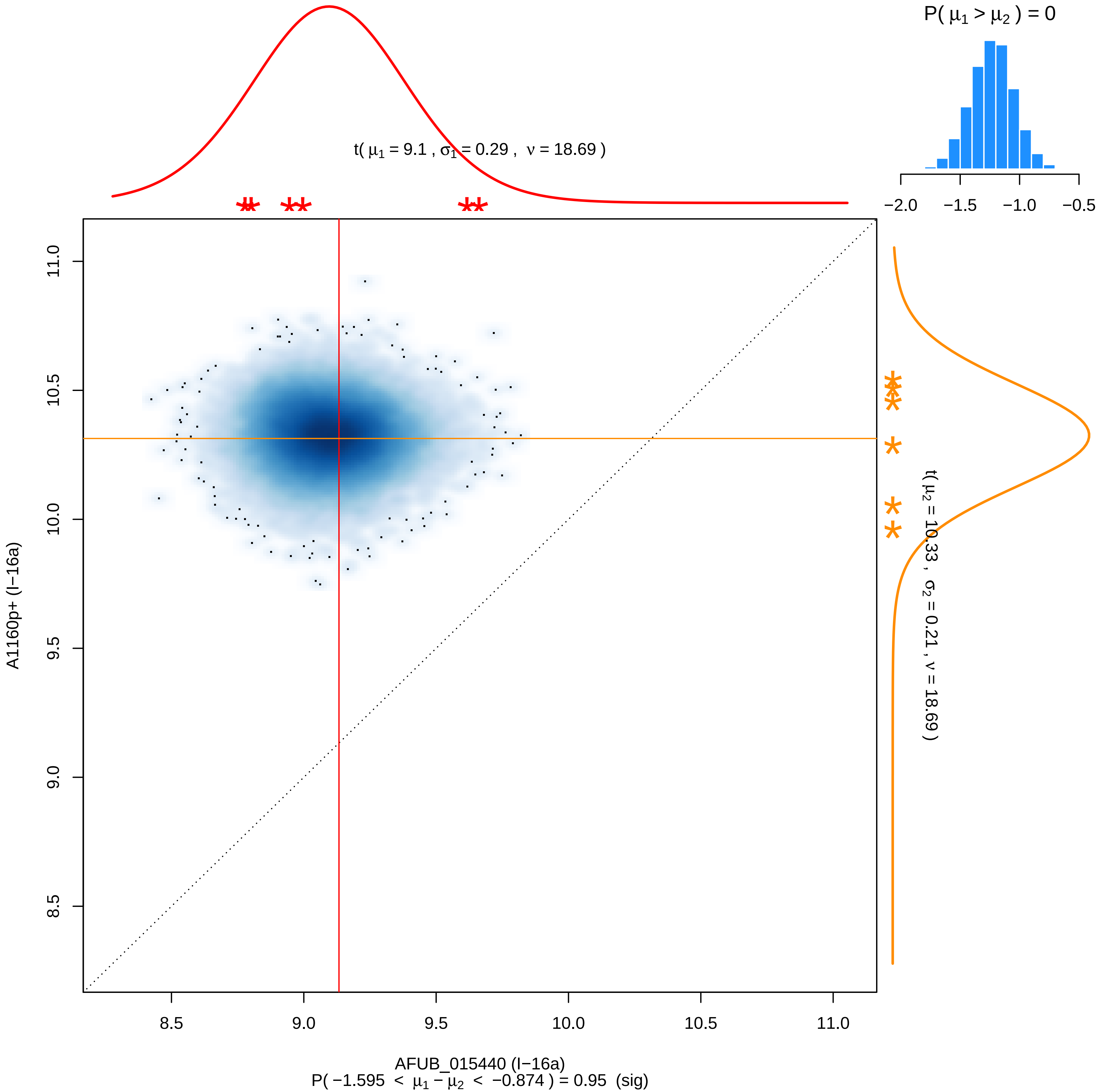

P( μ1 > μ2 ) = 0
−2.0 −1.5 −1.0 −0.5
t( μ1 = 9.1 , σ1 = 0.29 , ν = 18.69 )
t( μ2 = 10.33 , σ2 = 0.21 , ν = 18.69 )
A1160p+ (I−16a)
11.0 10.5 10.0 9.5 9.0 8.5
8.5 9.0 9.5 10.0 10.5 11.0
AFUB_015440 (I−16a)
P( −1.595 < μ1 − μ2 < −0.874 ) = 0.95 (sig)

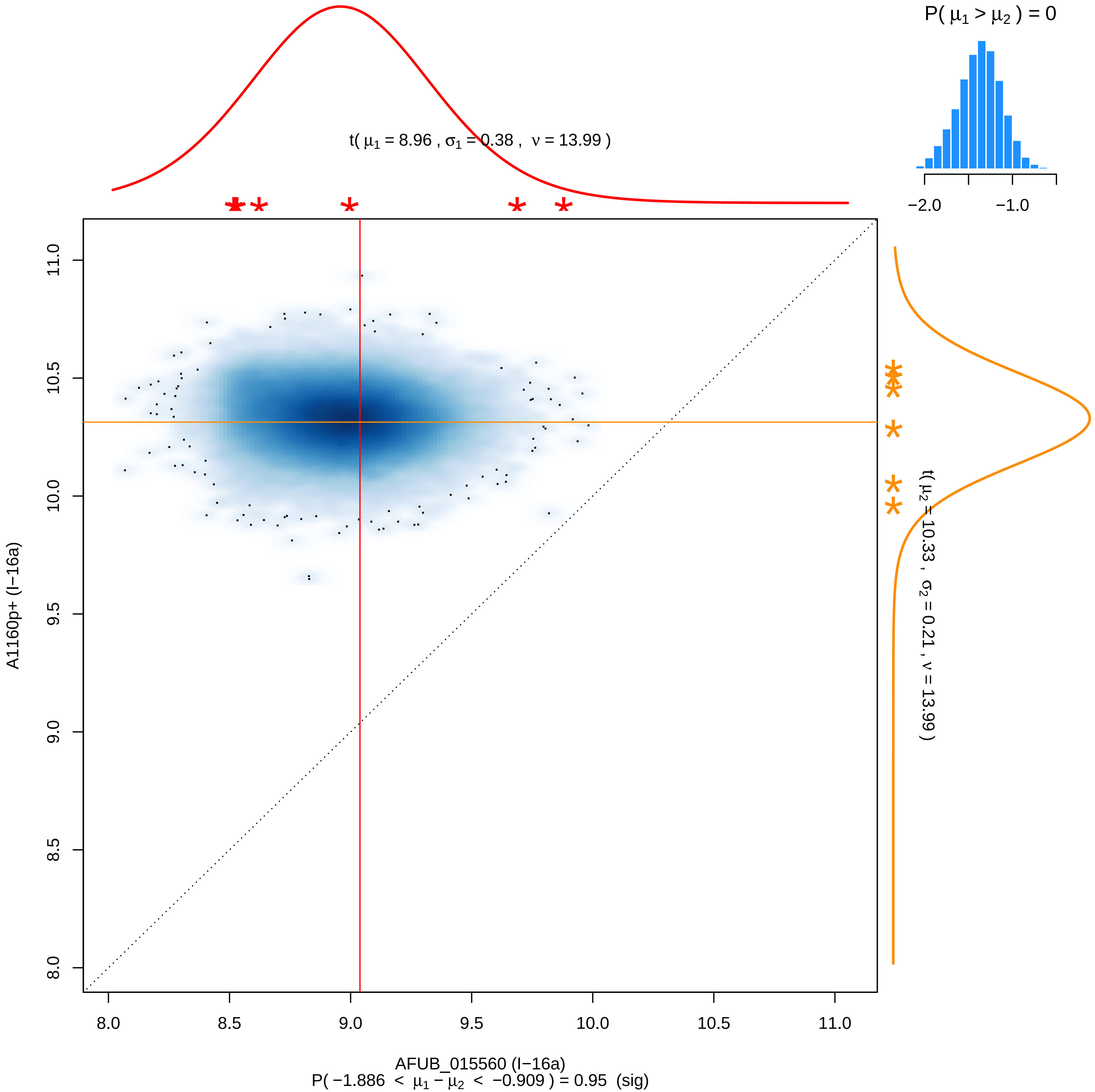

P( μ1 > μ2 ) = 0
−2.0
−1.0
t( μ1 = 8.96 , σ1 = 0.38 , ν = 13.99 )
t( μ2 = 10.33 , σ2 = 0.21 , ν = 13.99 )
A1160p+ (I−16a)
AFUB_015560 (I−16a)
P( −1.886 < μ1 − μ2 < −0.909 ) = 0.95 (sig)

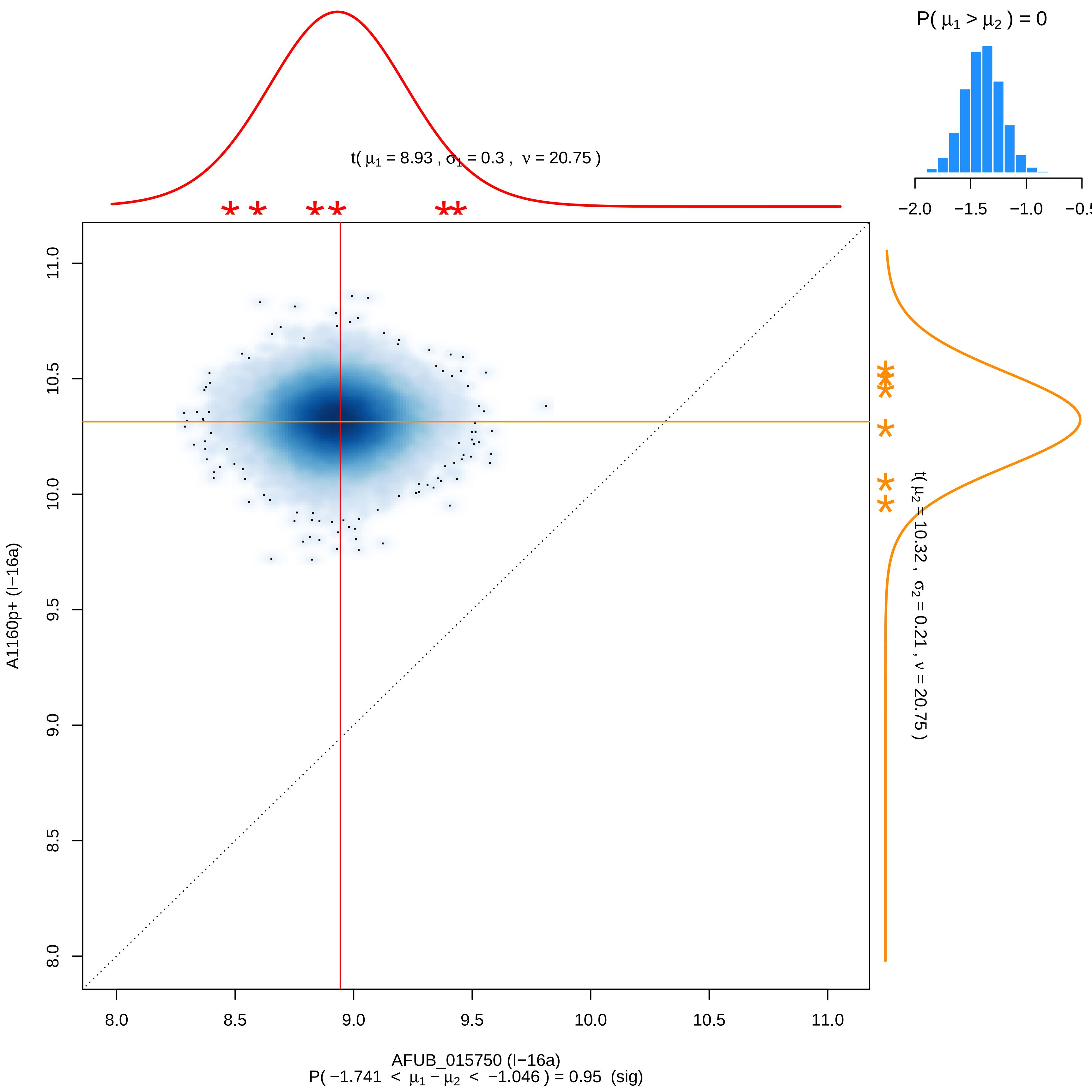

P( μ1 > μ2 ) = 0
−2.0 −1.5 −1.0 −0.5
t( μ1 = 8.93 , σ1 = 0.3 , ν = 20.75 )
11.0
10.5
10.0
9.5
9.0
8.5
8.0
A1160p+ (I−16a)
t( μ2 = 10.32 , σ2 = 0.21 , ν = 20.75 )
8.0 8.5 9.0 9.5 10.0 10.5 11.0
AFUB_015750 (I−16a)
P( −1.741 < μ1 − μ2 < −1.046 ) = 0.95 (sig)

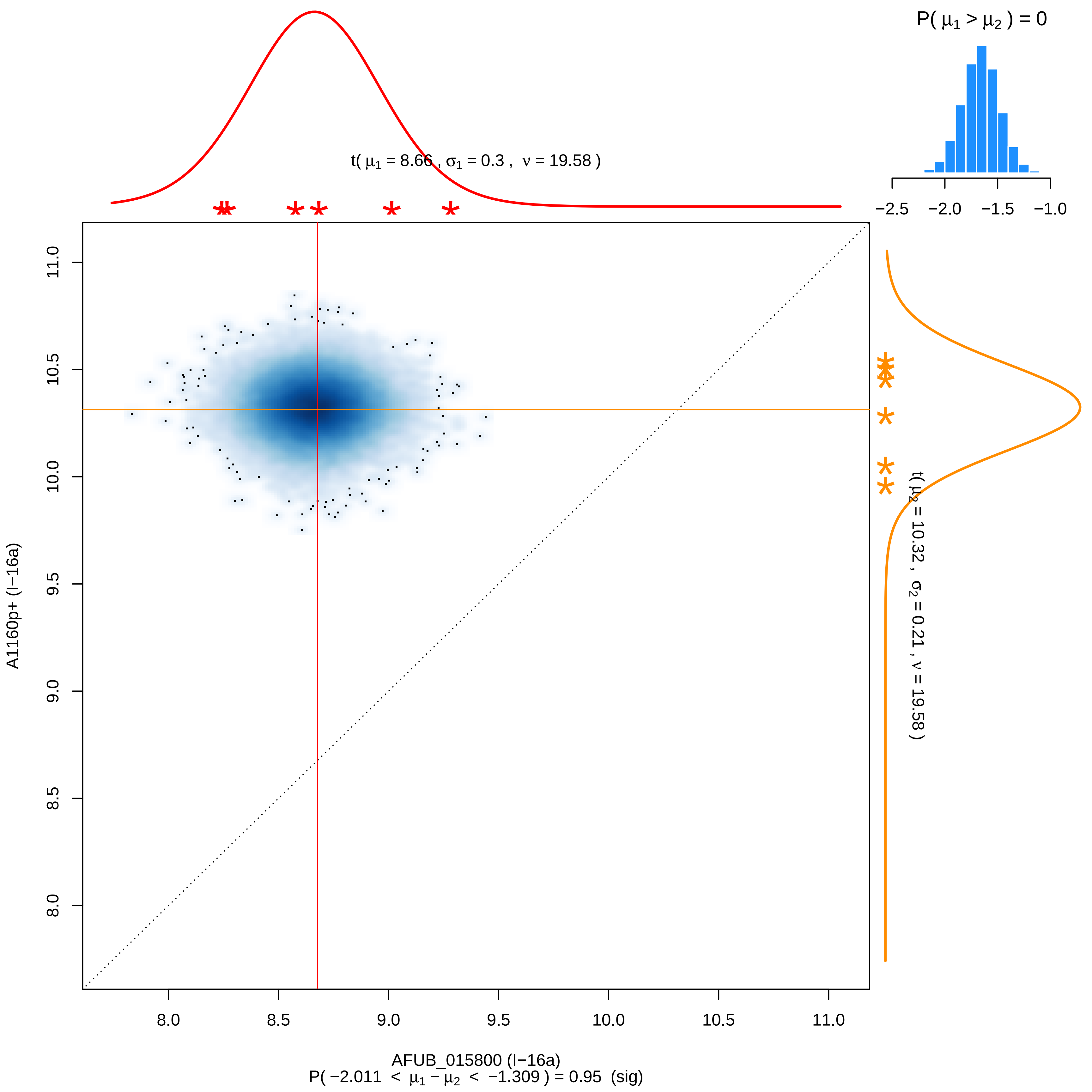

P( μ1 > μ2 ) = 0
−2.5 −2.0 −1.5 −1.0
t( μ1 = 8.66 , σ1 = 0.3 , ν = 19.58 )
11.0
10.5
10.0
9.5
9.0
8.5
8.0
A1160p+ (I−16a)
t( μ2 = 10.32 , σ2 = 0.21 , ν = 19.58 )
8.0 8.5 9.0 9.5 10.0 10.5 11.0
AFUB_015800 (I−16a)
P( −2.011 < μ1 − μ2 < −1.309 ) = 0.95 (sig)

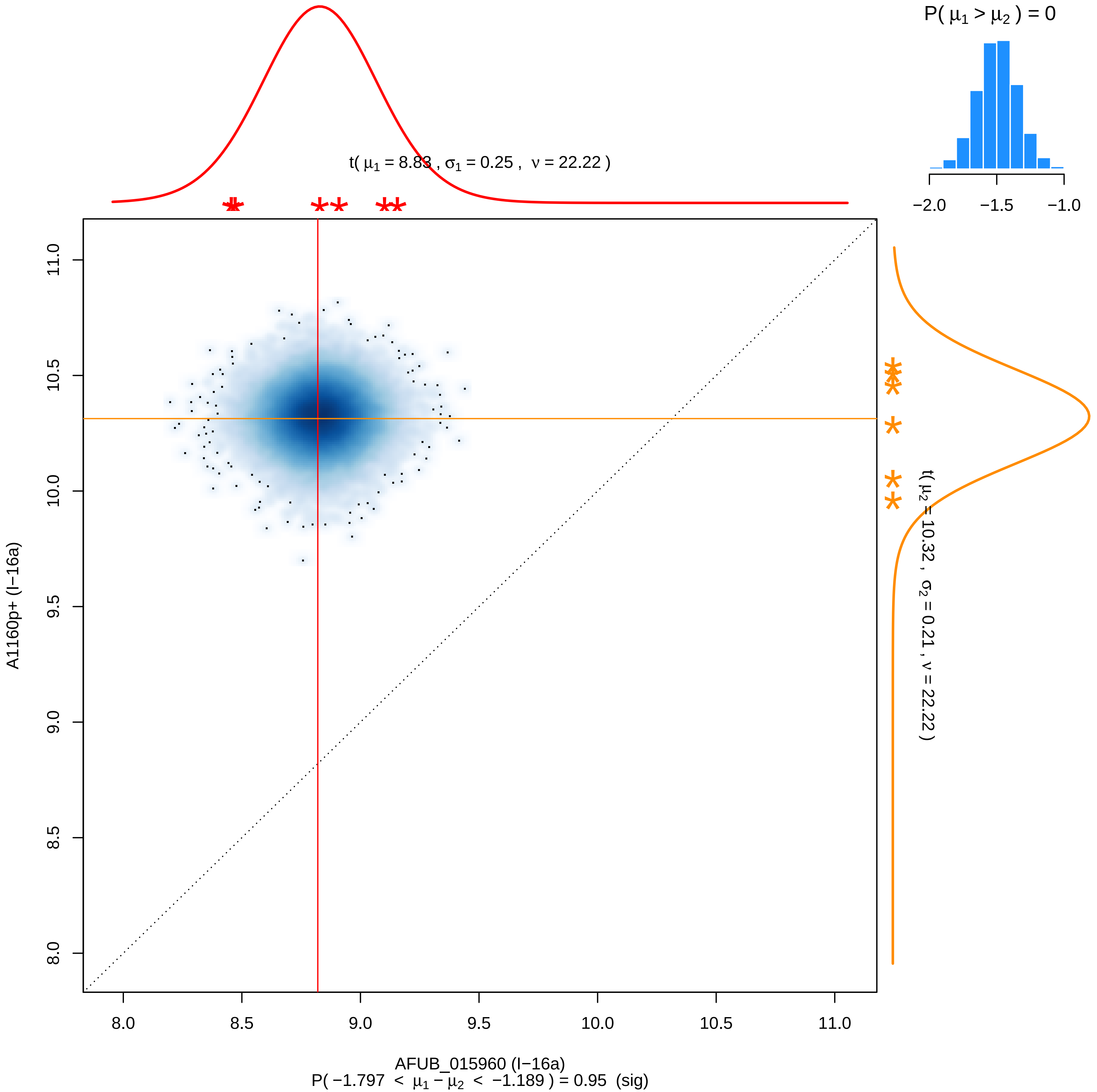

P( μ1 > μ2 ) = 0
−2.0
−1.5
−1.0
t( μ1 = 8.83 , σ1 = 0.25 , ν = 22.22 )
t( μ2 = 10.32 , σ2 = 0.21 , ν = 22.22 )
11.0
10.5
10.0
9.5
9.0
8.5
8.0
A1160p+ (I−16a)
8.0
8.5
9.0
9.5
10.0
10.5
11.0
AFUB_015960 (I−16a)
P( −1.797 < μ1 − μ2 < −1.189 ) = 0.95 (sig)

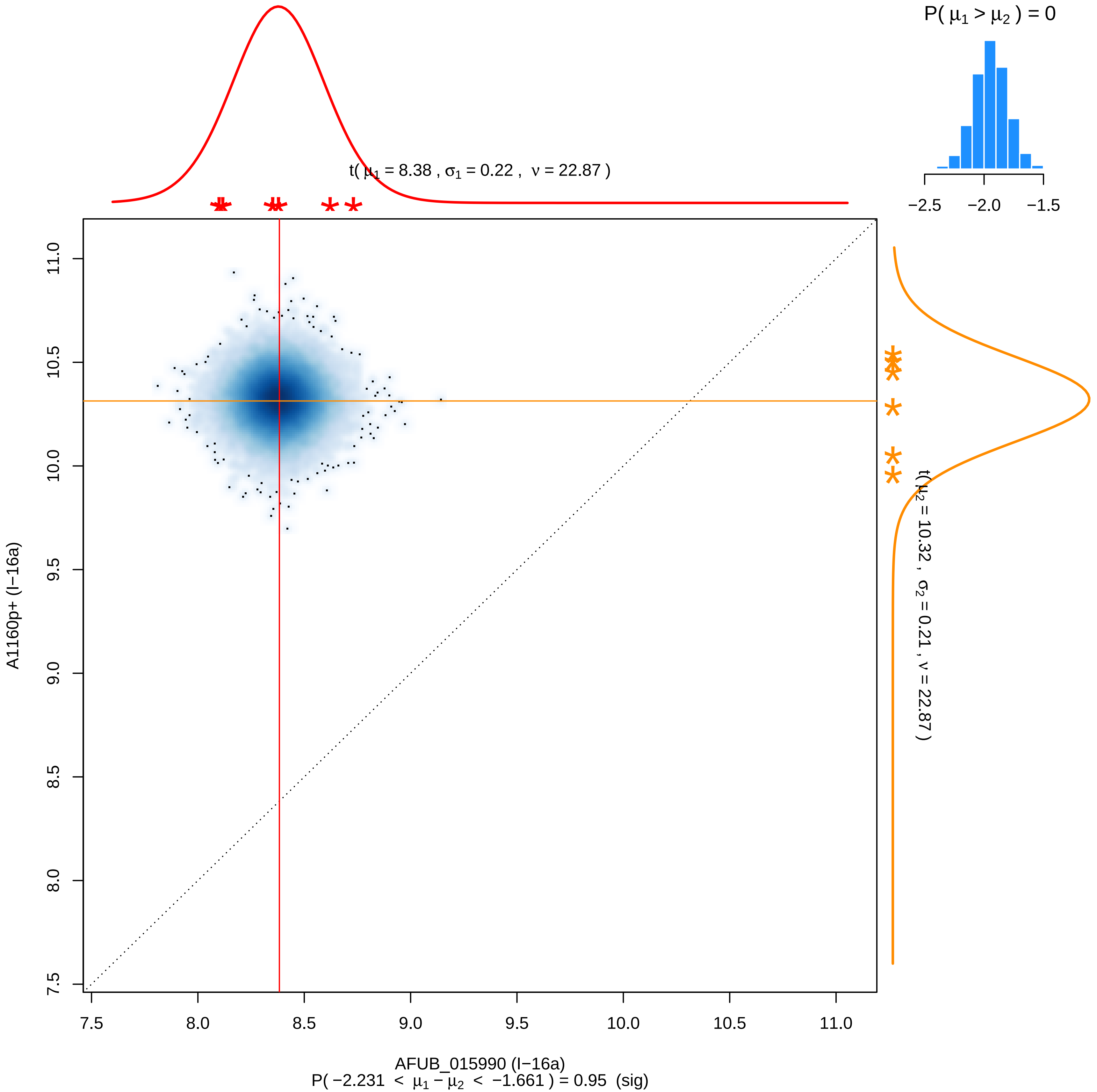

P( μ1 > μ2 ) = 0
−2.5
−2.0
−1.5
t( μ1 = 8.38 , σ1 = 0.22 , ν = 22.87 )
t( μ2 = 10.32 , σ2 = 0.21 , ν = 22.87 )
A1160p+ (I−16a)
AFUB_015990 (I−16a)
P( −2.231 < μ1 − μ2 < −1.661 ) = 0.95 (sig)
7.5
8.0
8.5
9.0
9.5
10.0
10.5
11.0

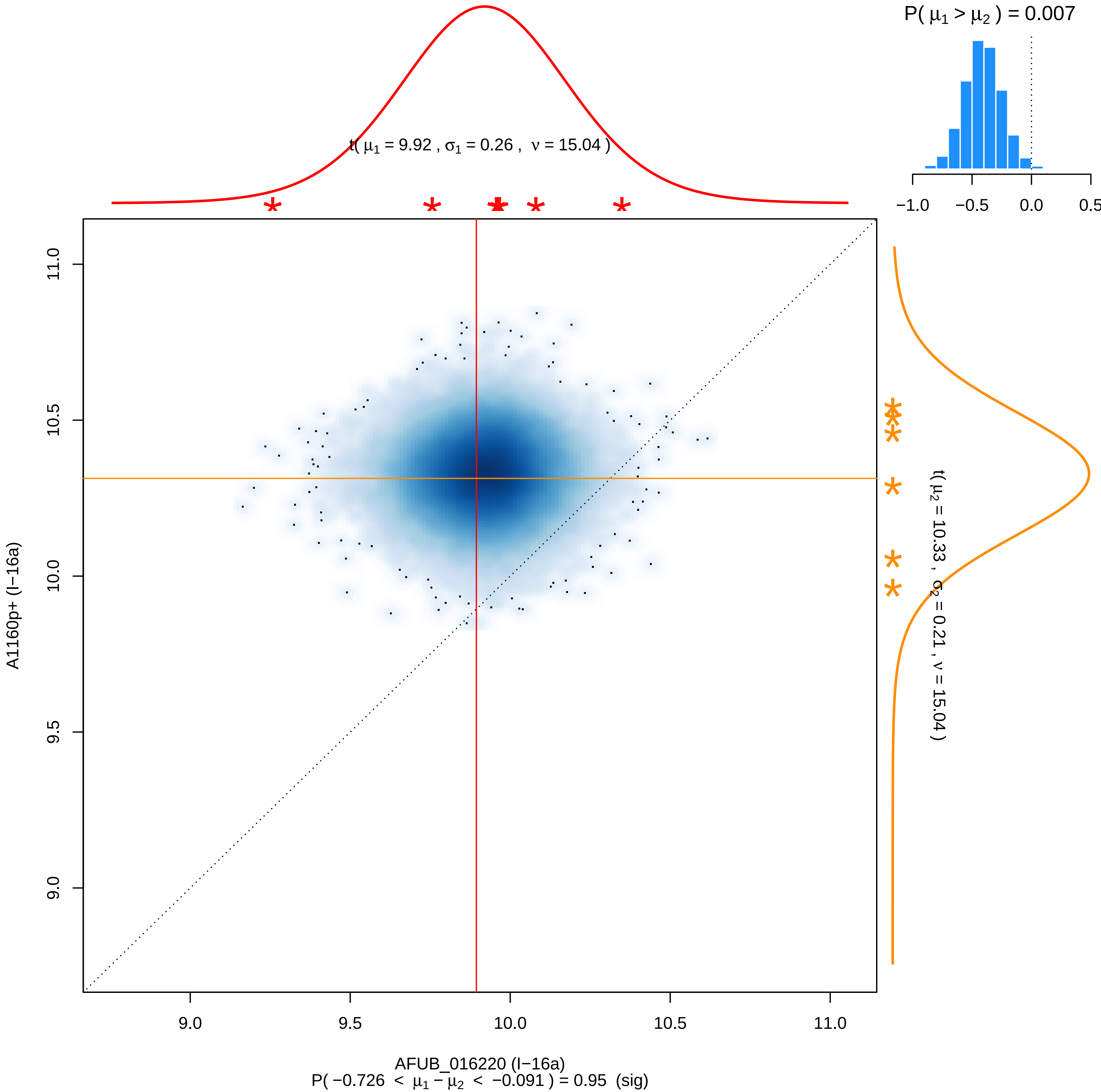

P( μ1 > μ2 ) = 0.007
−1.0
−0.5
0.0
0.5
t( μ1 = 9.92 , σ1 = 0.26 , ν = 15.04 )
11.0
10.5
10.0
9.5
9.0
A1160p+ (I−16a)
t( μ2 = 10.33 , σ2 = 0.21 , ν = 15.04 )
9.0
9.5
10.0
10.5
11.0
AFUB_016220 (I−16a)
P( −0.726 < μ1 − μ2 < −0.091 ) = 0.95 (sig)

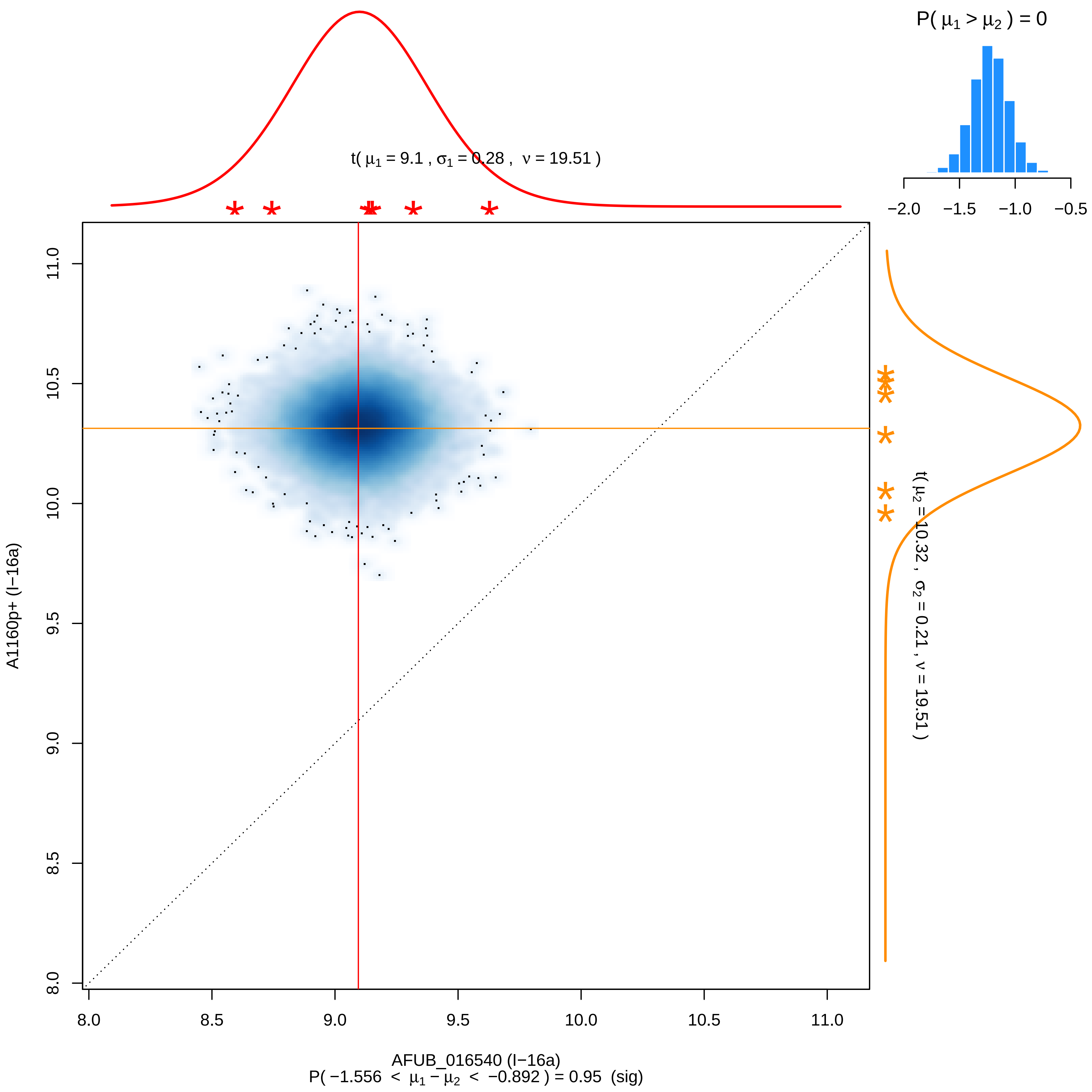

P( $\mu_1 > \mu_2$ ) = 0
−2.0
−1.5
−1.0
−0.5
t( $\mu_1$ = 9.1 , $\sigma_1$ = 0.28 , $\nu$ = 19.51 )
t( $\mu_2$ = 10.32 , $\sigma_2$ = 0.21 , $\nu$ = 19.51 )
8.0
8.5
9.0
9.5
10.0
10.5
11.0
A1160p+ (I−16a)
AFUB_016540 (I−16a)
P( −1.556 < $\mu_1 - \mu_2$ < −0.892 ) = 0.95 (sig)

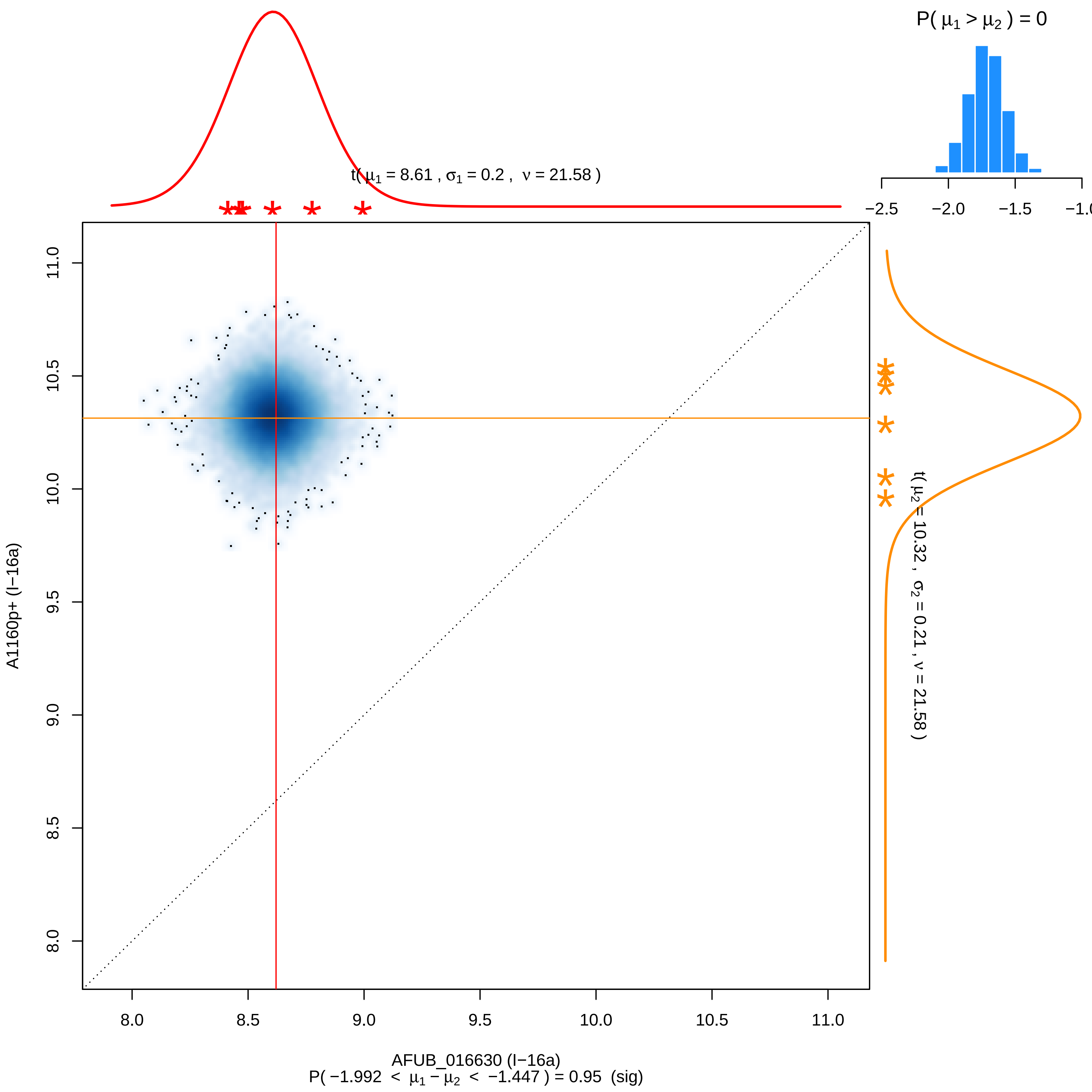
P( μ1 > μ2 ) = 0
−2.5
−2.0
−1.5
t( μ1 = 8.61 , σ1 = 0.2 , ν = 21.58 )
t( μ2 = 10.32 , σ2 = 0.21 , ν = 21.58 )
A1160p+ (I−16a)
11.0
10.5
10.0
9.5
9.0
8.5
8.0
8.0
8.5
9.0
9.5
10.0
10.5
11.0
AFUB_016630 (I−16a)
P( −1.992 < μ1 − μ2 < −1.447 ) = 0.95 (sig)

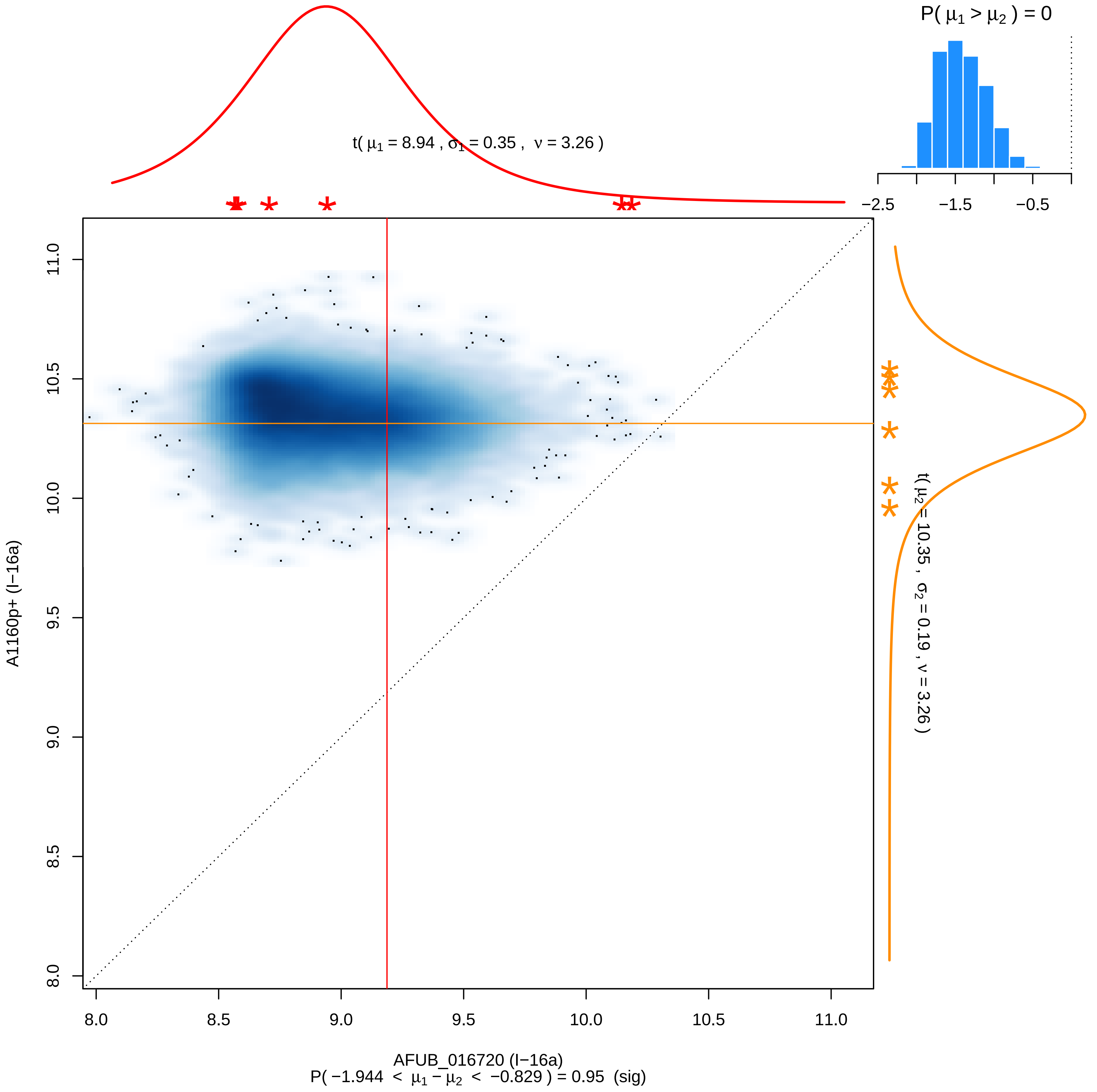

P( μ1 > μ2 ) = 0
−2.5
−1.5
−0.5
t( μ1 = 8.94 , σ1 = 0.35 , ν = 3.26 )
t( μ2 = 10.35 , σ2 = 0.19 , ν = 3.26 )
11.0
10.5
10.0
9.5
9.0
8.5
8.0
A1160p+ (I−16a)
8.0
8.5
9.0
9.5
10.0
10.5
11.0
AFUB_016720 (I−16a)
P( −1.944 < μ1 − μ2 < −0.829 ) = 0.95 (sig)

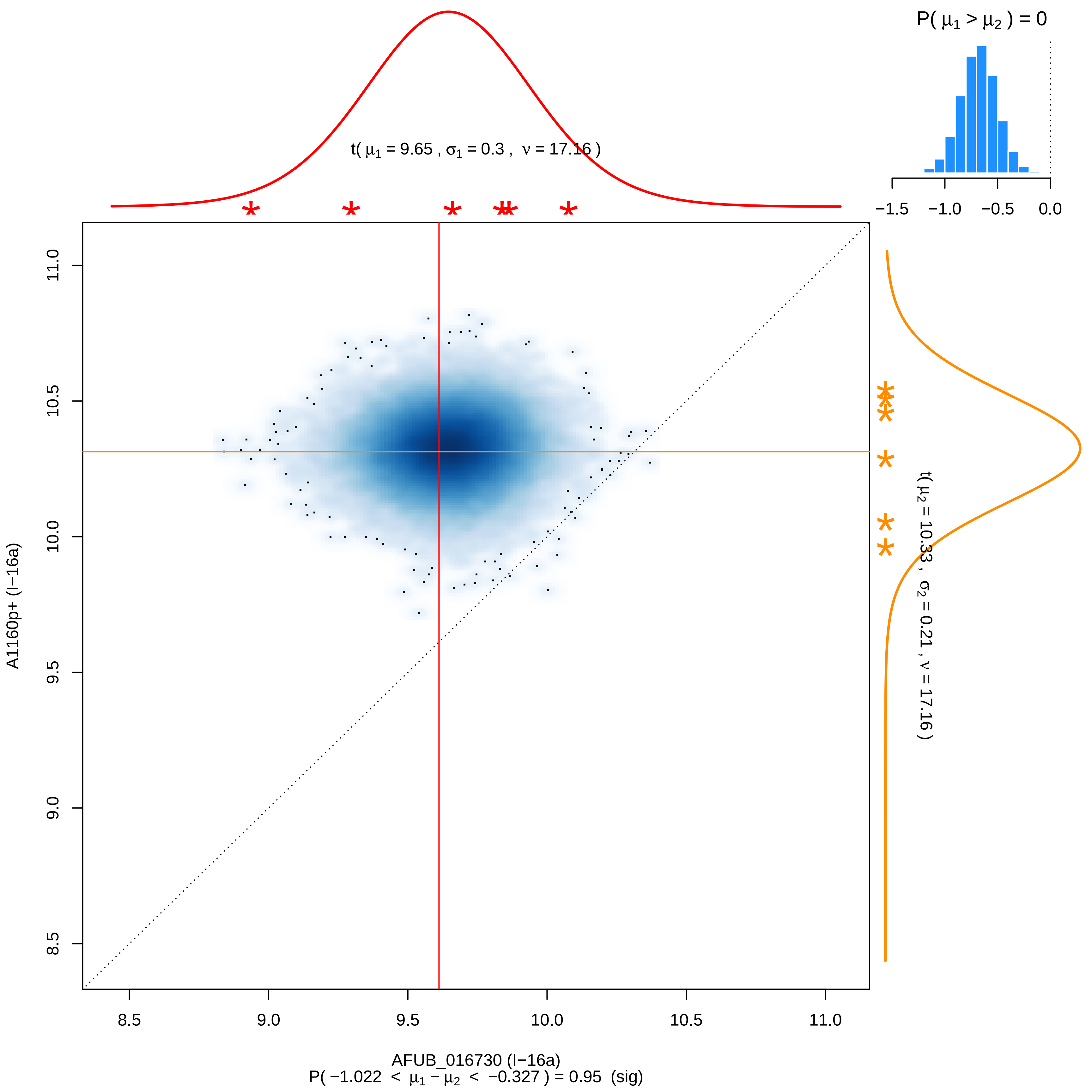

P( μ1 > μ2 ) = 0
−1.5 −1.0 −0.5 0.0
t( μ1 = 9.65 , σ1 = 0.3 , ν = 17.16 )
t( μ2 = 10.33 , σ2 = 0.21 , ν = 17.16 )
A1160p+ (I−16a)
8.5 9.0 9.5 10.0 10.5 11.0
AFUB_016730 (I−16a)
P( −1.022 < μ1 − μ2 < −0.327 ) = 0.95 (sig)

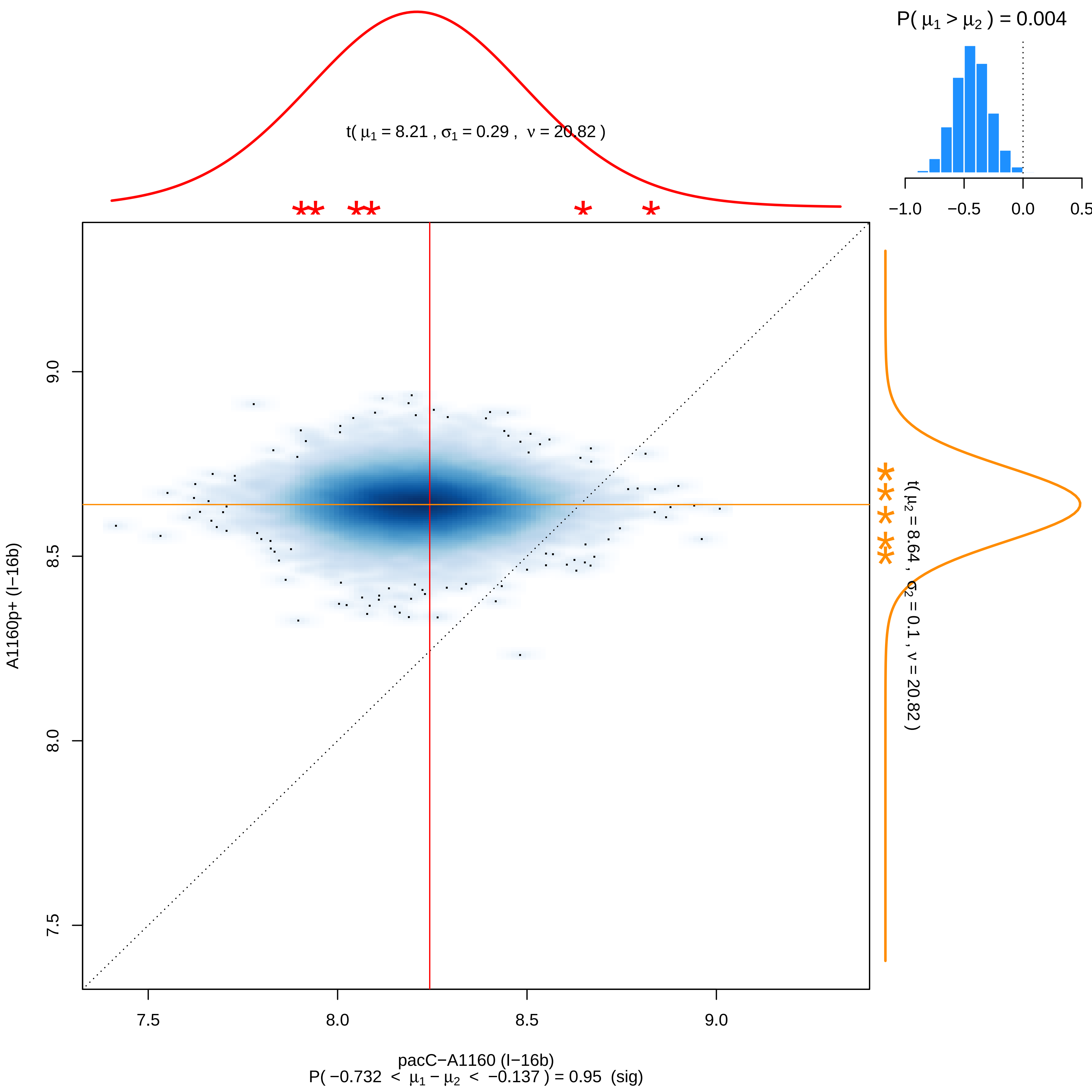

P( μ1 > μ2 ) = 0.004
−1.0
−0.5
0.0
0.5
t( μ1 = 8.21 , σ1 = 0.29 , ν = 20.82 )
9.0
8.5
8.0
7.5
A1160p+ (I−16b)
t( μ2 = 8.64 , σ2 = 0.1 , ν = 20.82 )
7.5
8.0
8.5
9.0
pacC−A1160 (I−16b)
P( −0.732 < μ1 − μ2 < −0.137 ) = 0.95 (sig)

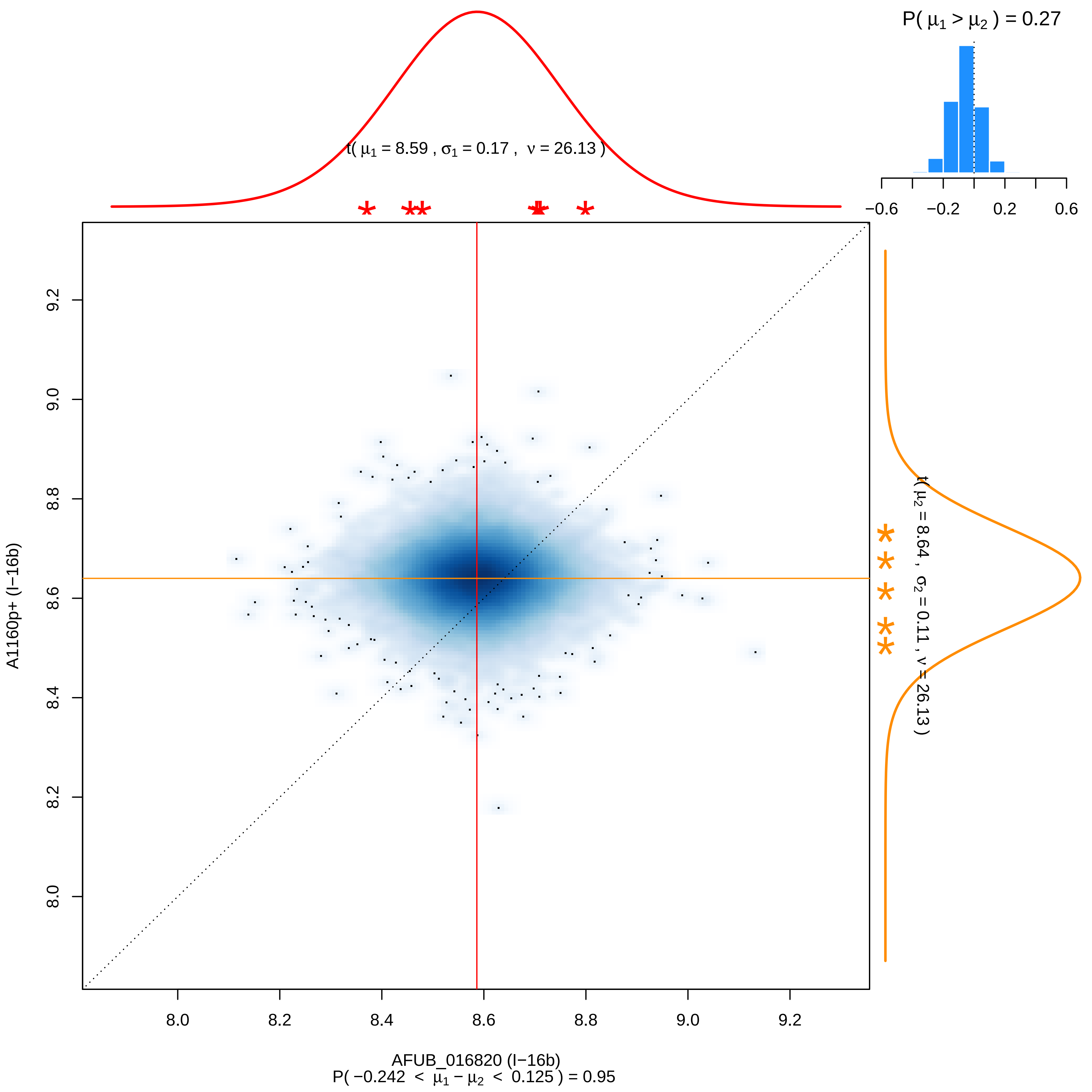

P( μ1 > μ2 ) = 0.27
−0.6
−0.2
0.2
0.6
t( μ1 = 8.59 , σ1 = 0.17 , ν = 26.13 )
t( μ2 = 8.64 , σ2 = 0.11 , ν = 26.13 )
A1160p+ (I−16b)
9.2
9.0
8.8
8.6
8.4
8.2
8.0
8.0
8.2
8.4
8.6
8.8
9.0
9.2
AFUB_016820 (I−16b)
P( −0.242 < μ1 − μ2 < 0.125 ) = 0.95

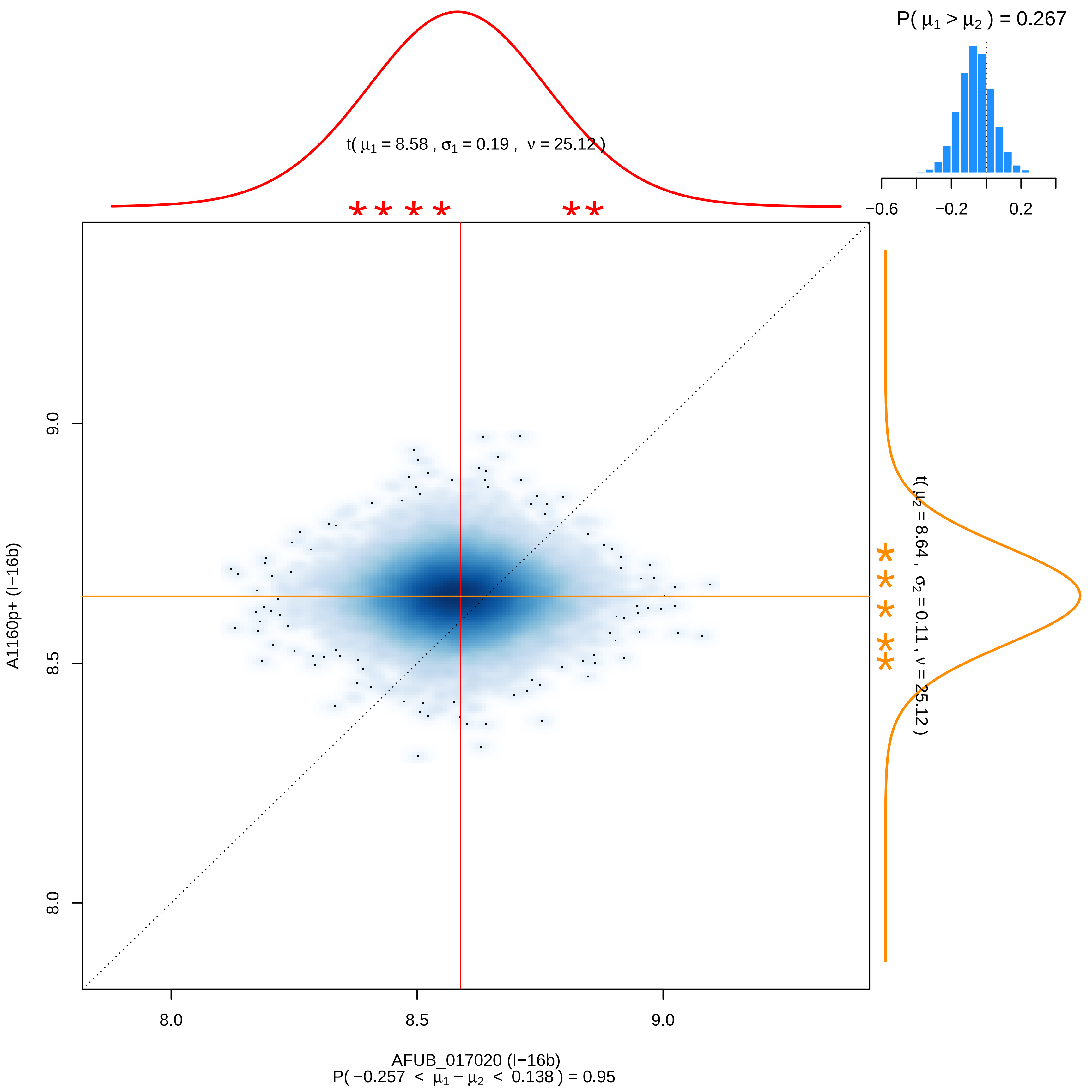

P( μ1 > μ2 ) = 0.267
−0.6
−0.2
0.2
t( μ1 = 8.58 , σ1 = 0.19 , ν = 25.12 )
t( μ2 = 8.64 , σ2 = 0.11 , ν = 25.12 )
9.0
8.5
8.0
A1160p+ (I−16b)
8.0
8.5
9.0
AFUB_017020 (I−16b)
P( −0.257 < μ1 − μ2 < 0.138 ) = 0.95

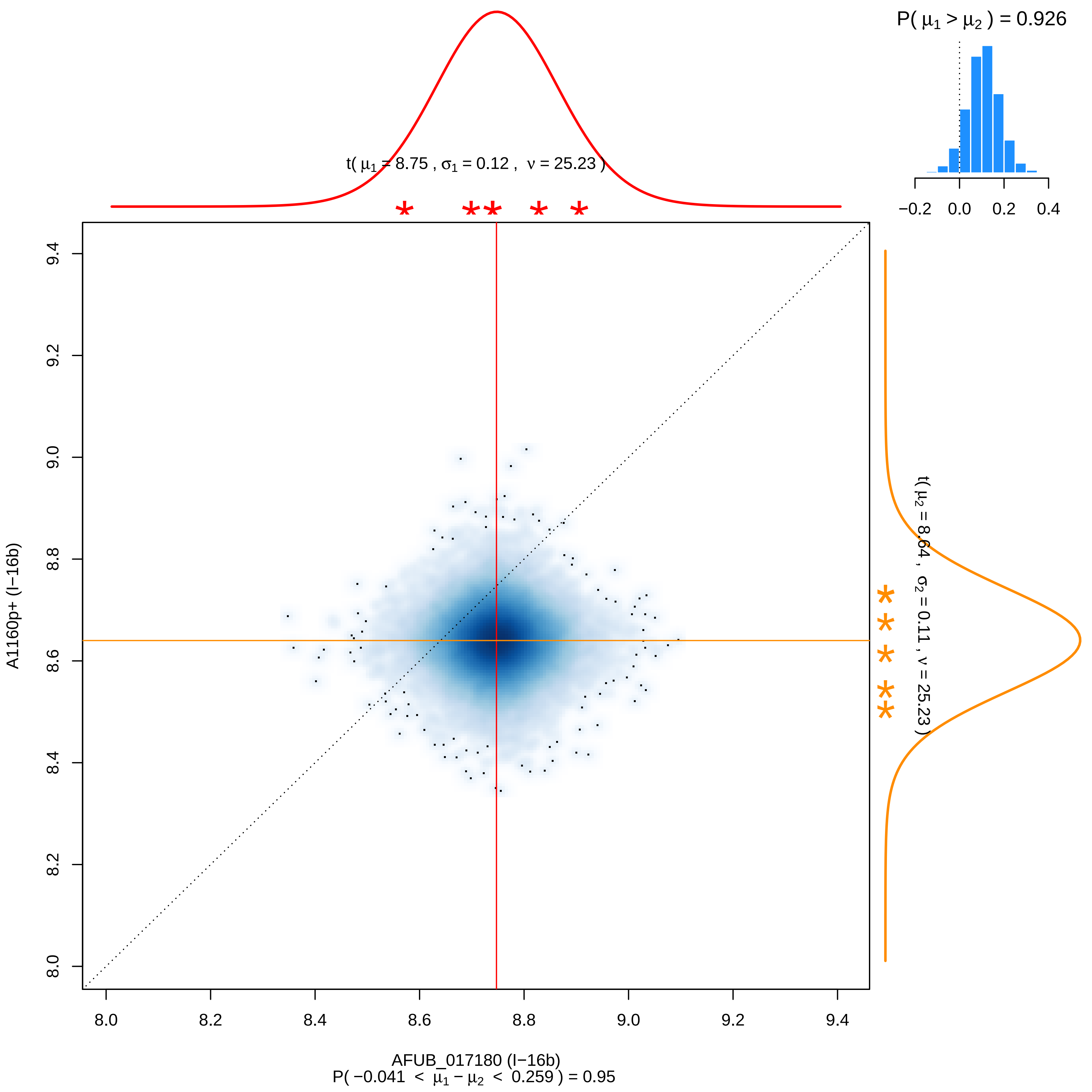
P( μ1 > μ2 ) = 0.926
−0.2 0.0 0.2 0.4
t( μ1 = 8.75 , σ1 = 0.12 , ν = 25.23 )
t( μ2 = 8.64 , σ2 = 0.11 , ν = 25.23 )
A1160p+ (I−16b)
AFUB_017180 (I−16b)
8.0 8.2 8.4 8.6 8.8 9.0 9.2 9.4
P( −0.041 < μ1 − μ2 < 0.259 ) = 0.95

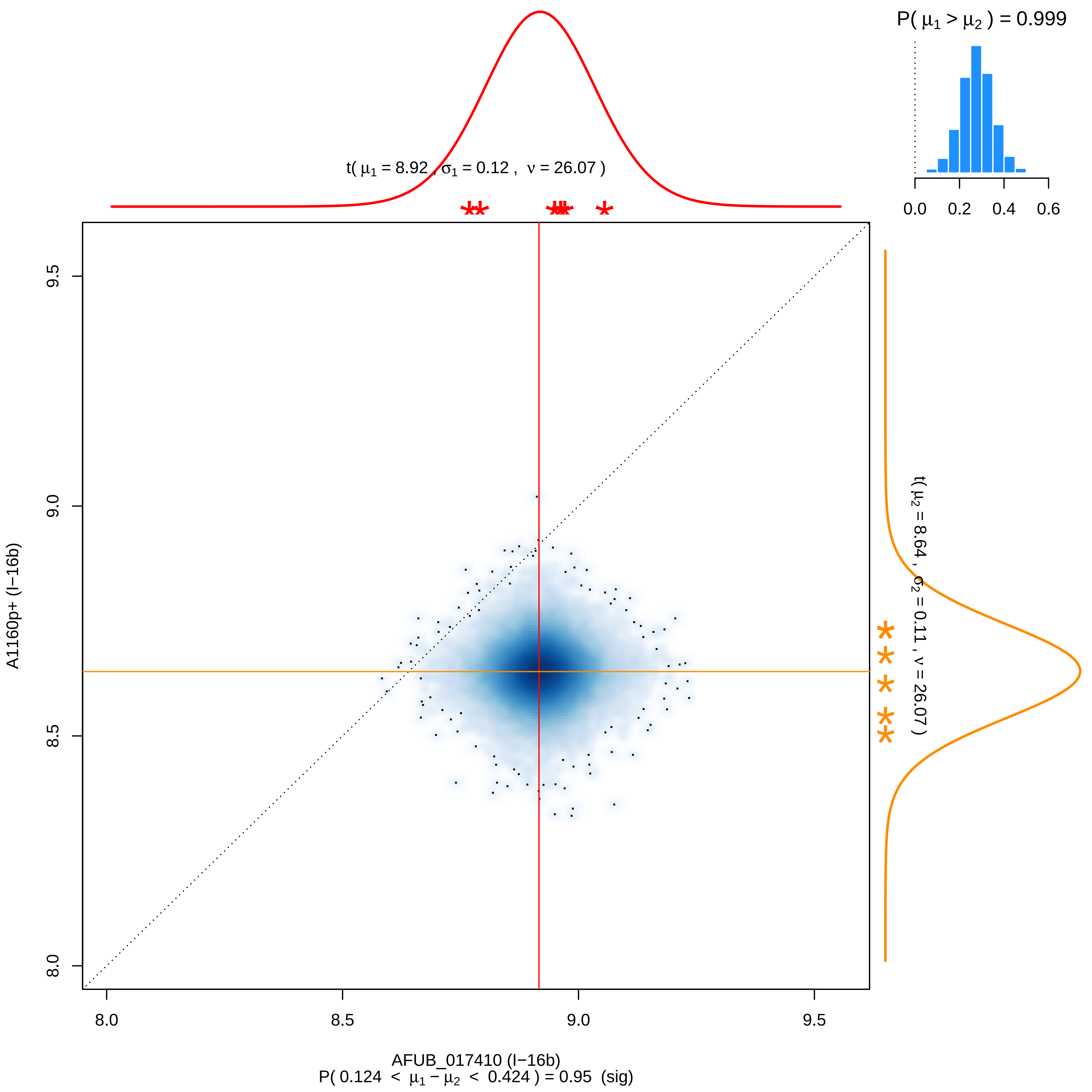

P( μ1 > μ2 ) = 0.999
0.0 0.2 0.4 0.6
t( μ1 = 8.92 , σ1 = 0.12 , ν = 26.07 )
t( μ2 = 8.64 , σ2 = 0.11 , ν = 26.07 )
9.5
9.0
8.5
8.0
A1160p+ (I−16b)
8.0 8.5 9.0 9.5
AFUB_017410 (I−16b)
P( 0.124 < μ1 − μ2 < 0.424 ) = 0.95 (sig)

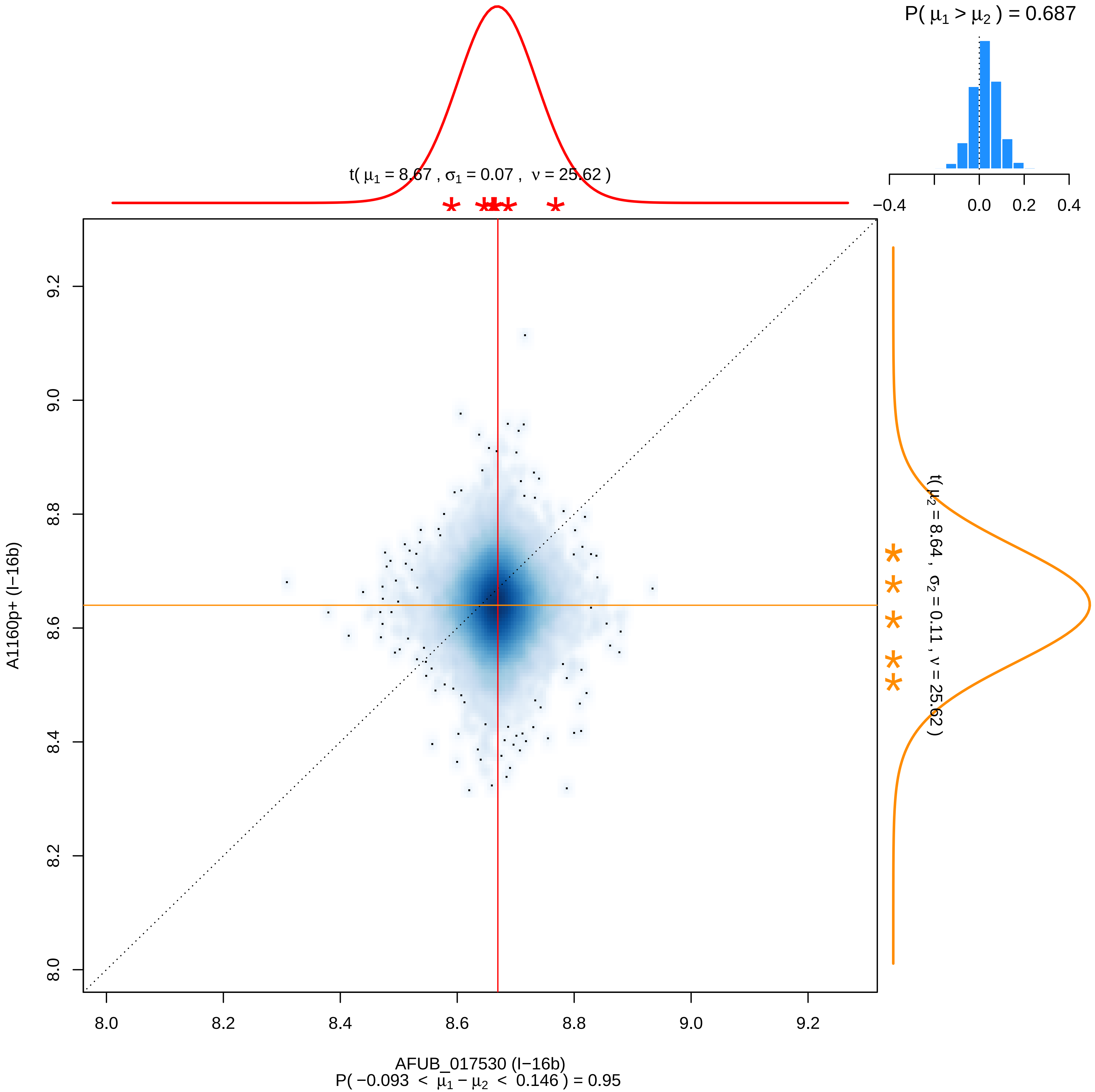

P( μ1 > μ2 ) = 0.687
t( μ1 = 8.67 , σ1 = 0.07 , ν = 25.62 )
−0.4
0.0
0.2
0.4
t( μ2 = 8.64 , σ2 = 0.11 , ν = 25.62 )
A1160p+ (I−16b)
9.2
9.0
8.8
8.6
8.4
8.2
8.0
8.0
8.2
8.4
8.6
8.8
9.0
9.2
AFUB_017530 (I−16b)
P( −0.093 < μ1 − μ2 < 0.146 ) = 0.95

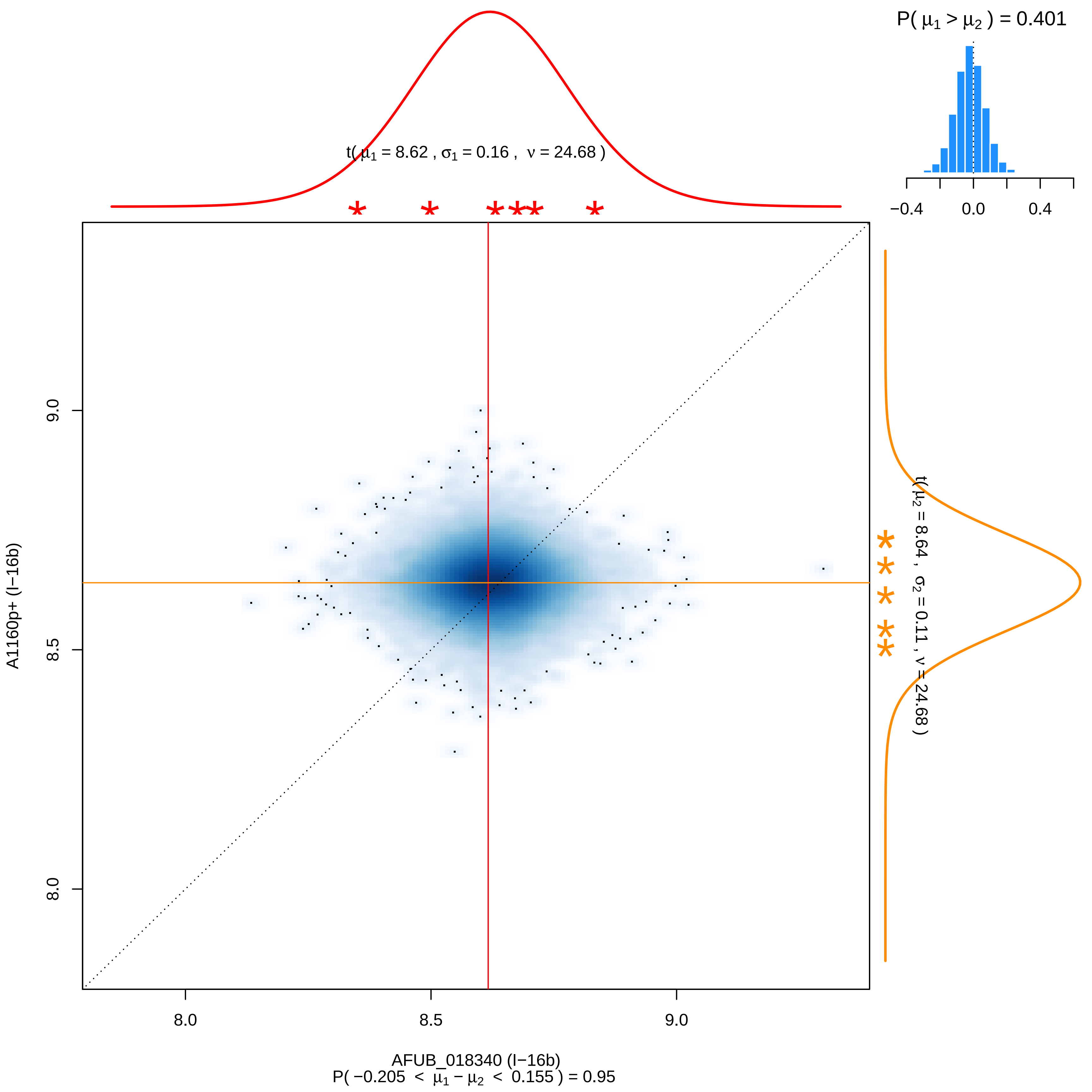

P( $\mu_1 > \mu_2$ ) = 0.401
−0.4
0.0
0.4
t( $\mu_1$ = 8.62 , $\sigma_1$ = 0.16 , $\nu$ = 24.68 )
t( $\mu_2$ = 8.64 , $\sigma_2$ = 0.11 , $\nu$ = 24.68 )
9.0
8.5
8.0
A1160p+ (I−16b)
8.0
8.5
9.0
AFUB_018340 (I−16b)
P( −0.205 < $\mu_1 - \mu_2$ < 0.155 ) = 0.95

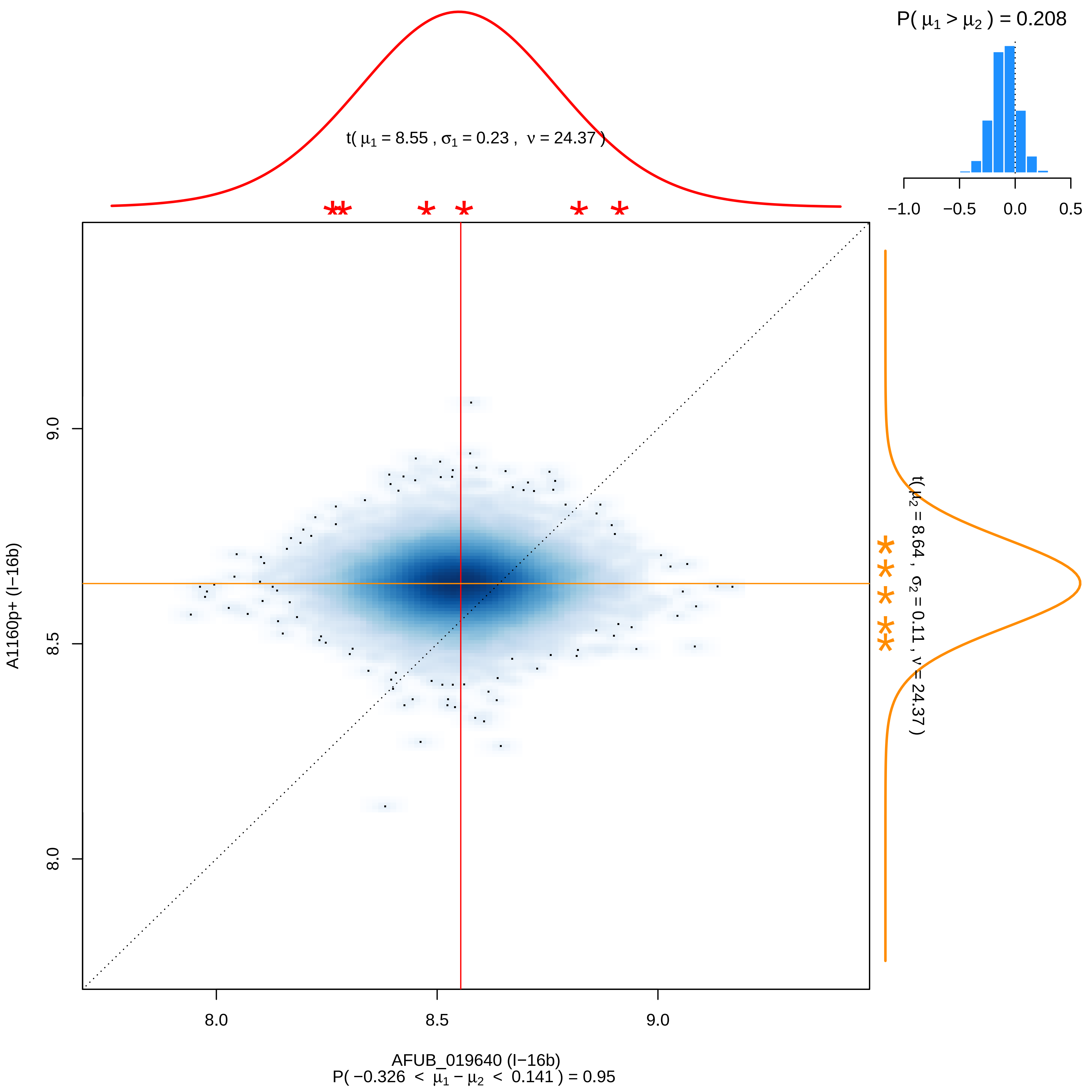

P( μ1 > μ2 ) = 0.208
−1.0 −0.5 0.0 0.5
t( μ1 = 8.55 , σ1 = 0.23 , ν = 24.37 )
t( μ2 = 8.64 , σ2 = 0.11 , ν = 24.37 )
9.0
8.5
8.0
A1160p+ (I−16b)
8.0 8.5 9.0
AFUB_019640 (I−16b)
P( −0.326 < μ1 − μ2 < 0.141 ) = 0.95

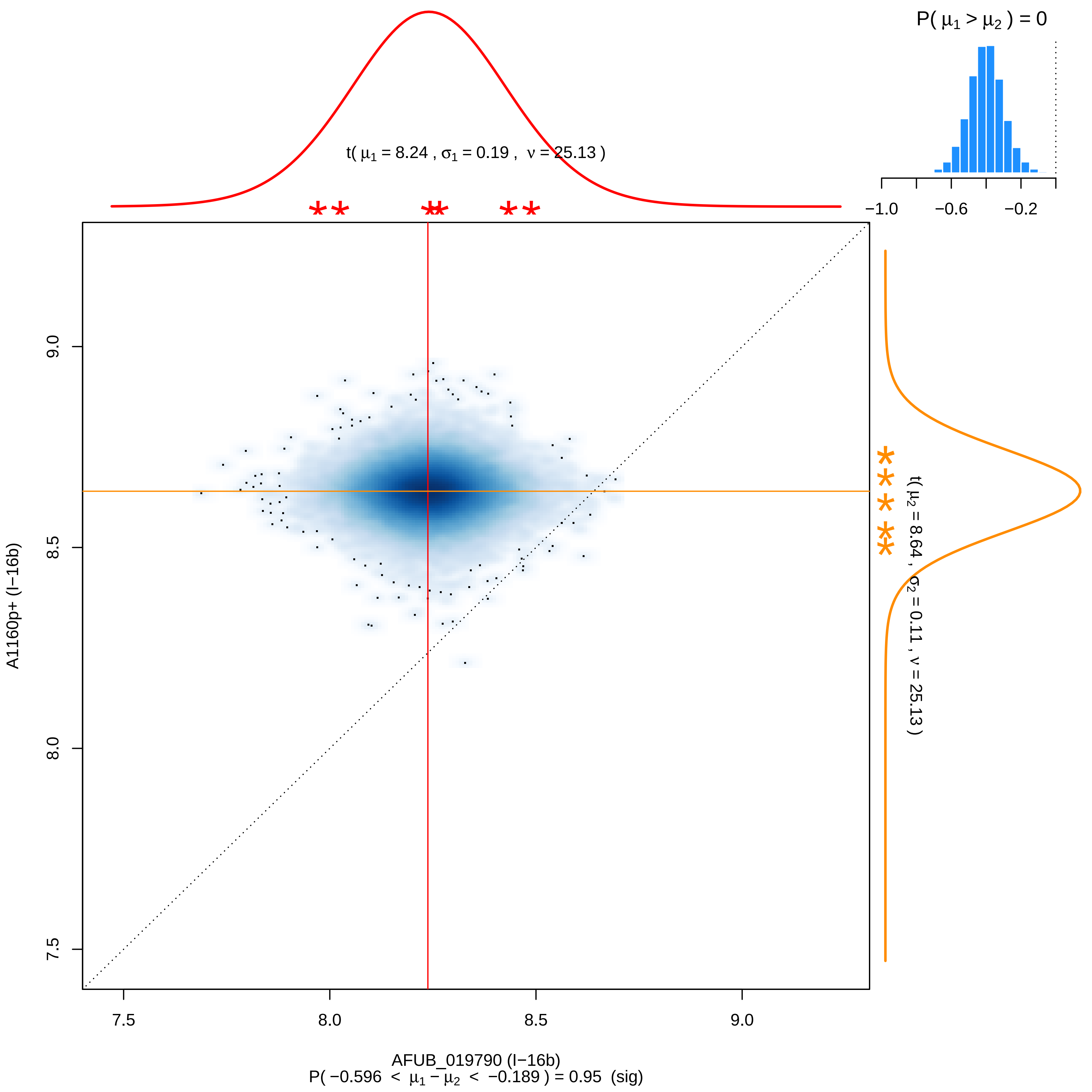

P( μ1 > μ2 ) = 0
−1.0
−0.6
−0.2
t( μ1 = 8.24 , σ1 = 0.19 , ν = 25.13 )
t( μ2 = 8.64 , σ2 = 0.11 , ν = 25.13 )
A1160p+ (I−16b)
9.0
8.5
8.0
7.5
7.5
8.0
8.5
9.0
AFUB_019790 (I−16b)
P( −0.596 < μ1 − μ2 < −0.189 ) = 0.95 (sig)

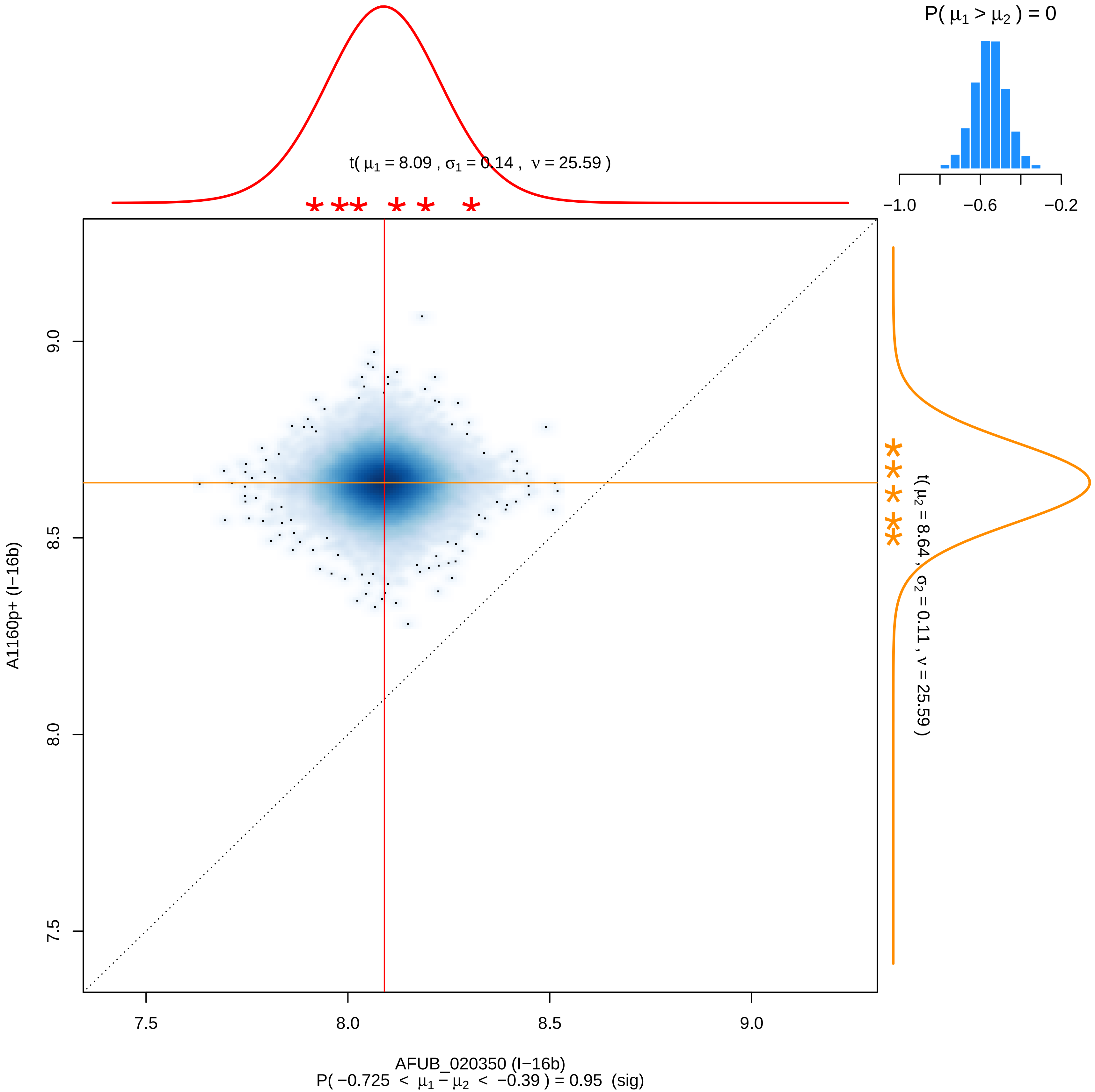

P( μ1 > μ2 ) = 0
−1.0
−0.6
−0.2
t( μ1 = 8.09 , σ1 = 0.14 , ν = 25.59 )
t( μ2 = 8.64 , σ2 = 0.11 , ν = 25.59 )
9.0
8.5
8.0
7.5
A1160p+ (I−16b)
7.5
8.0
8.5
9.0
AFUB_020350 (I−16b)
P( −0.725 < μ1 − μ2 < −0.39 ) = 0.95 (sig)

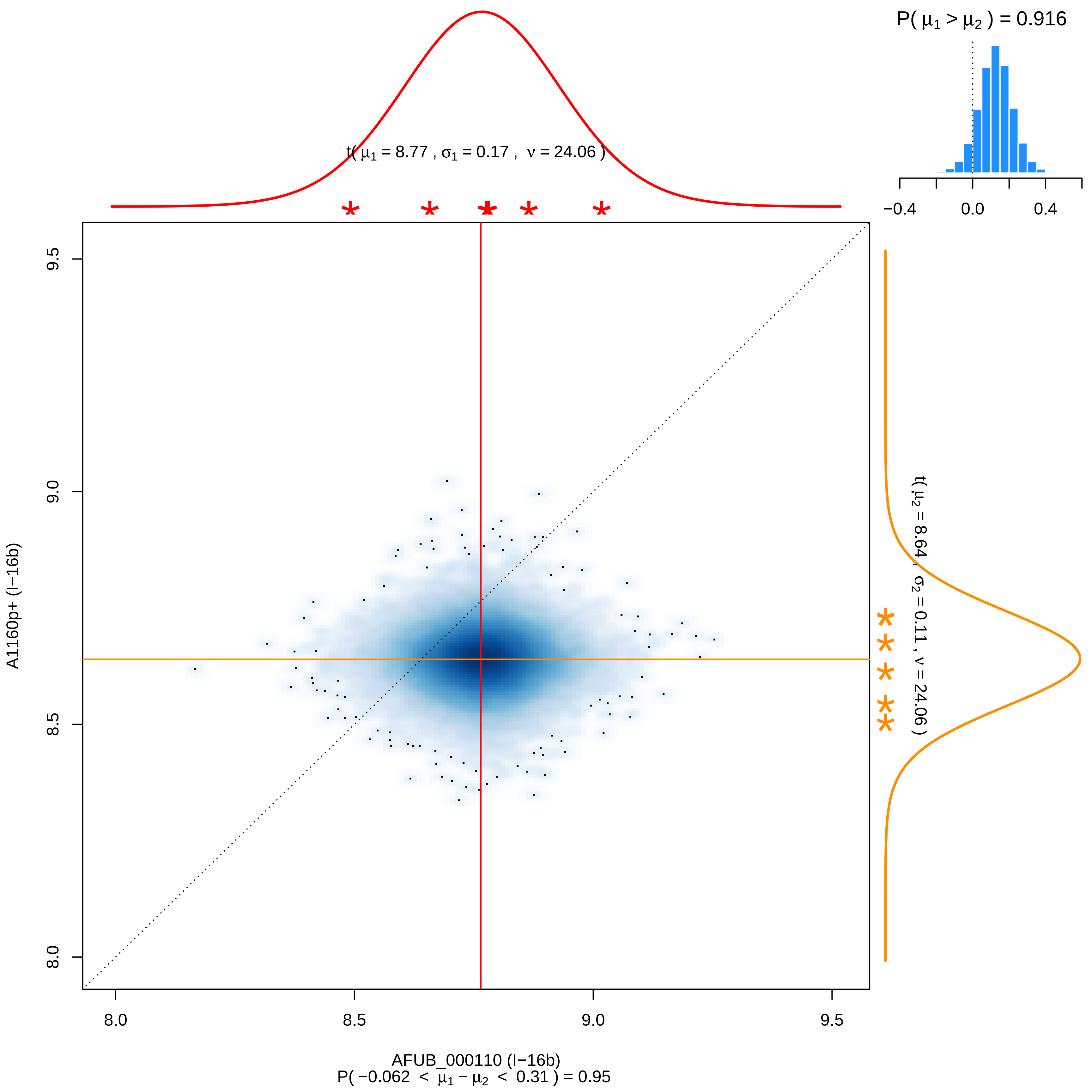

P( μ1 > μ2 ) = 0.916
−0.4
0.0
0.4
t( μ1 = 8.77 , σ1 = 0.17 , ν = 24.06 )
9.5
9.0
8.5
8.0
A1160p+ (I−16b)
8.0
8.5
9.0
9.5
t( μ2 = 8.64 , σ2 = 0.11 , ν = 24.06 )
AFUB_000110 (I−16b)
P( −0.062 < μ1 − μ2 < 0.31 ) = 0.95

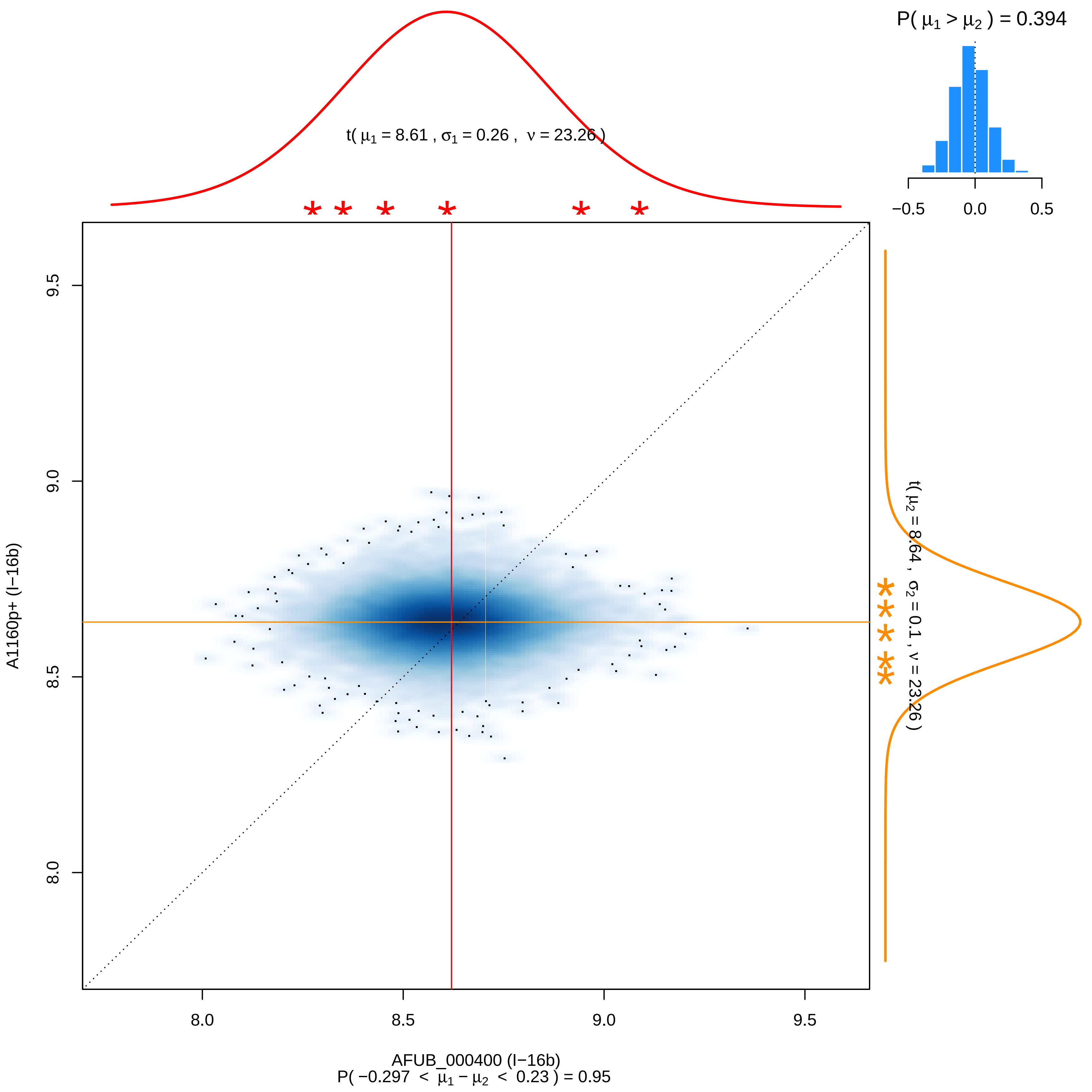

P( μ1 > μ2 ) = 0.394
−0.5
0.0
0.5
t( μ1 = 8.61 , σ1 = 0.26 , ν = 23.26 )
t( μ2 = 8.64 , σ2 = 0.1 , ν = 23.26 )
9.5
9.0
8.5
8.0
A1160p+ (I−16b)
8.0
8.5
9.0
9.5
AFUB_000400 (I−16b)
P( −0.297 < μ1 − μ2 < 0.23 ) = 0.95

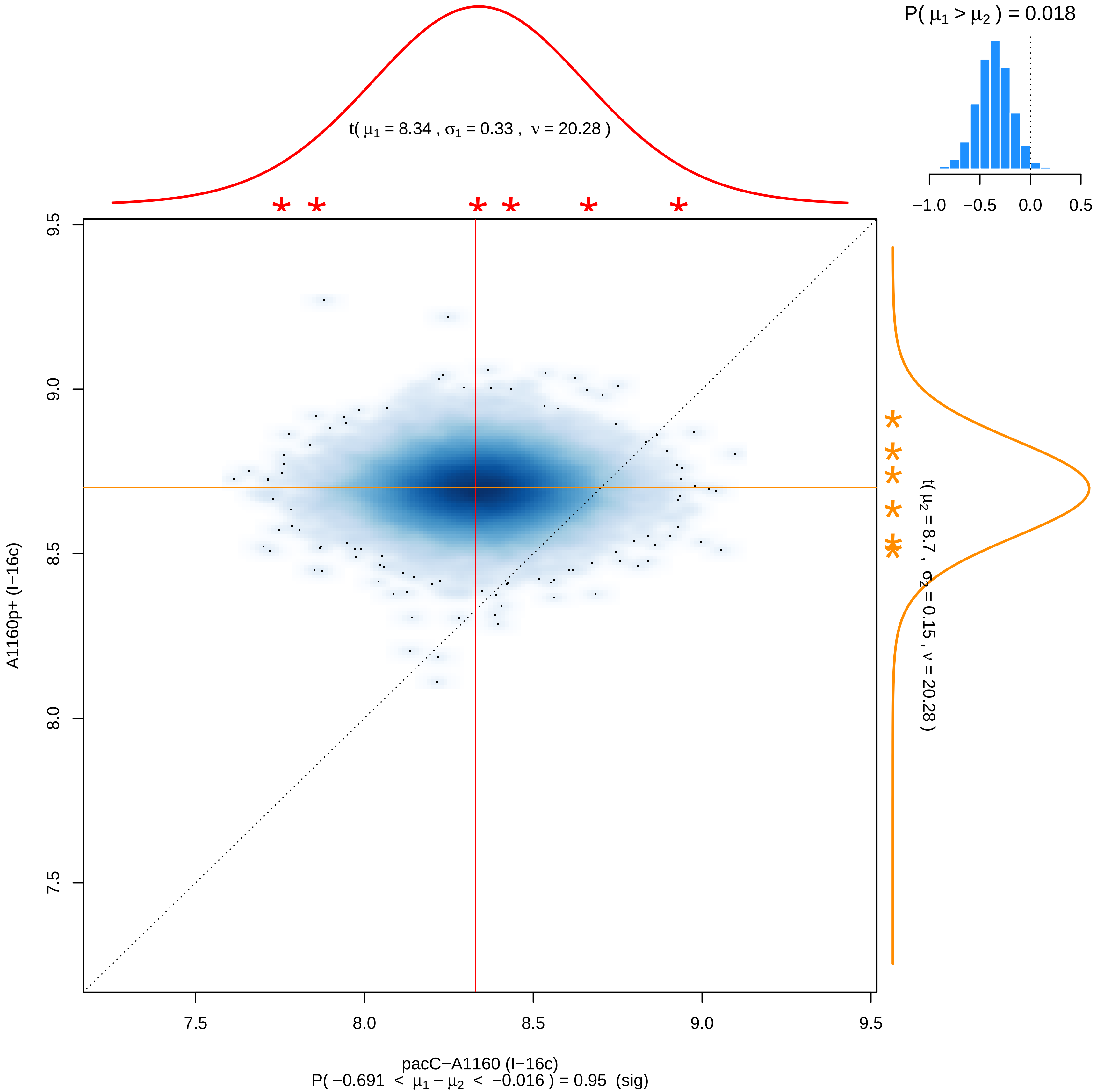

P( μ1 > μ2 ) = 0.018
t( μ1 = 8.34 , σ1 = 0.33 , ν = 20.28 )
t( μ2 = 8.7 , σ2 = 0.15 , ν = 20.28 )
A1160p+ (I−16c)
pacC−A1160 (I−16c)
P( −0.691 < μ1 − μ2 < −0.016 ) = 0.95 (sig)

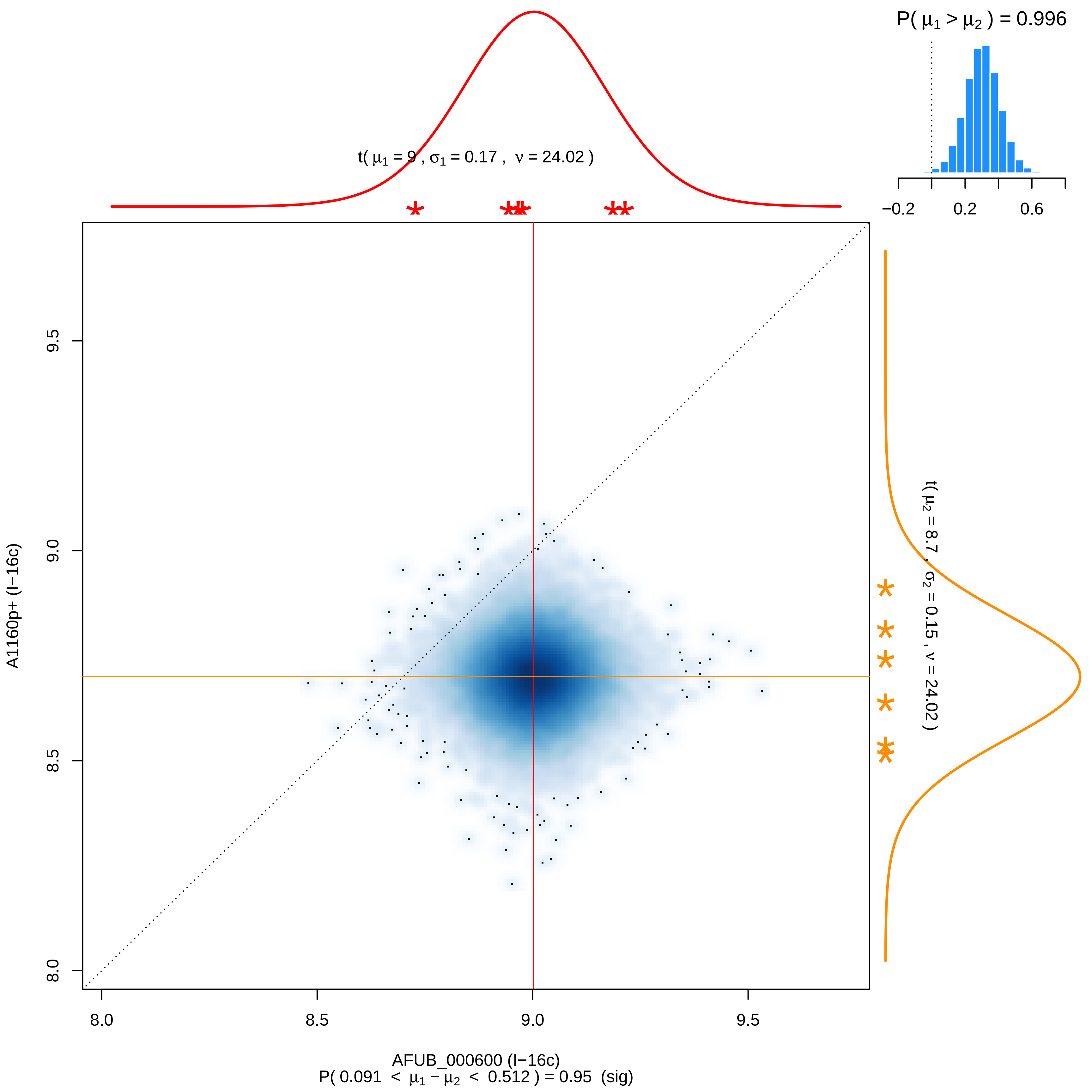

P( μ1 > μ2 ) = 0.996
−0.2
0.2
0.6
t( μ1 = 9 , σ1 = 0.17 , ν = 24.02 )
t( μ2 = 8.7 , σ2 = 0.15 , ν = 24.02 )
A1160p+ (I−16c)
8.0
8.5
9.0
9.5
AFUB_000600 (I−16c)
P( 0.091 < μ1 − μ2 < 0.512 ) = 0.95 (sig)

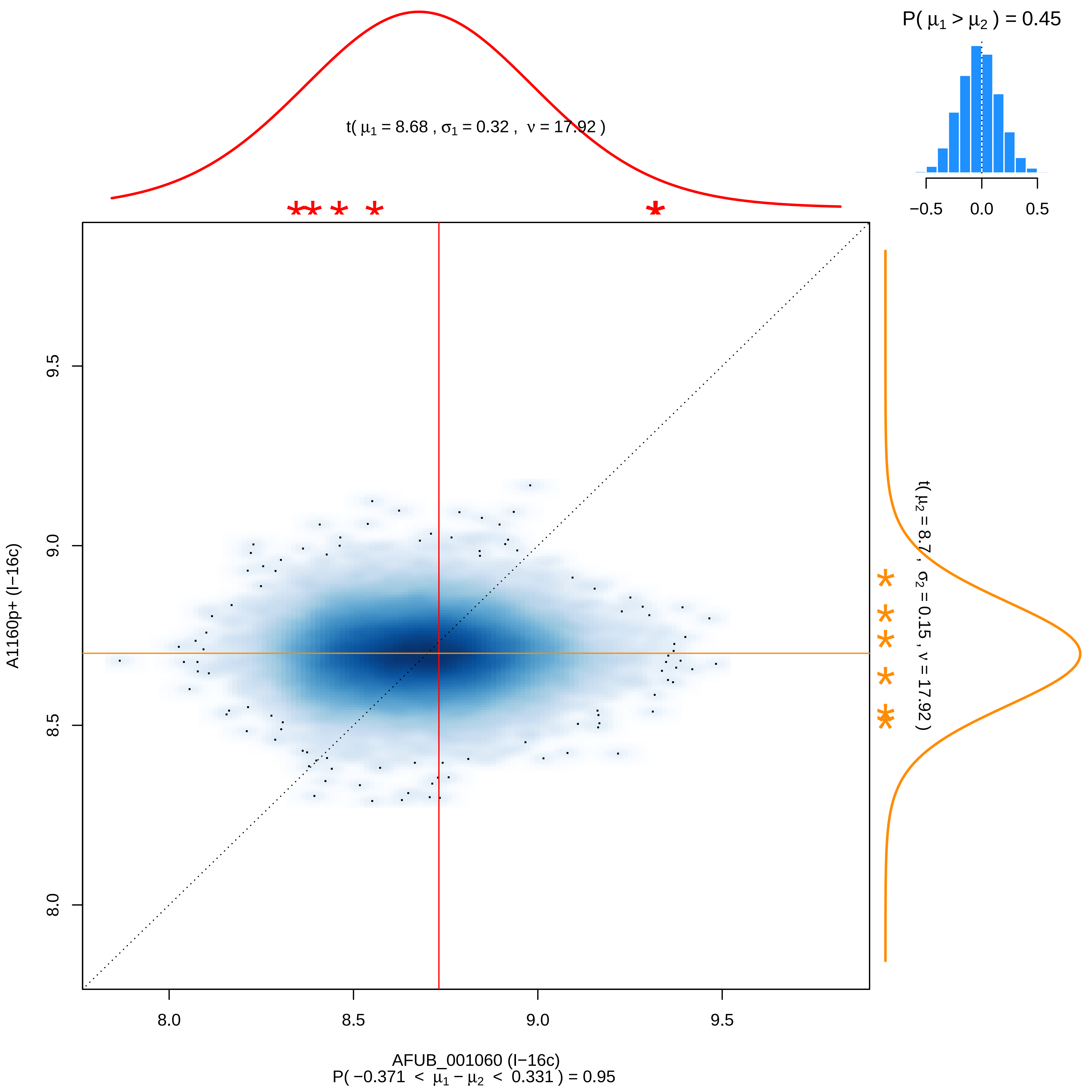

P( μ1 > μ2 ) = 0.45
−0.5
0.0
0.5
t( μ1 = 8.68 , σ1 = 0.32 , ν = 17.92 )
t( μ2 = 8.7 , σ2 = 0.15 , ν = 17.92 )
9.5
9.0
8.5
8.0
A1160p+ (I−16c)
8.0
8.5
9.0
9.5
AFUB_001060 (I−16c)
P( −0.371 < μ1 − μ2 < 0.331 ) = 0.95

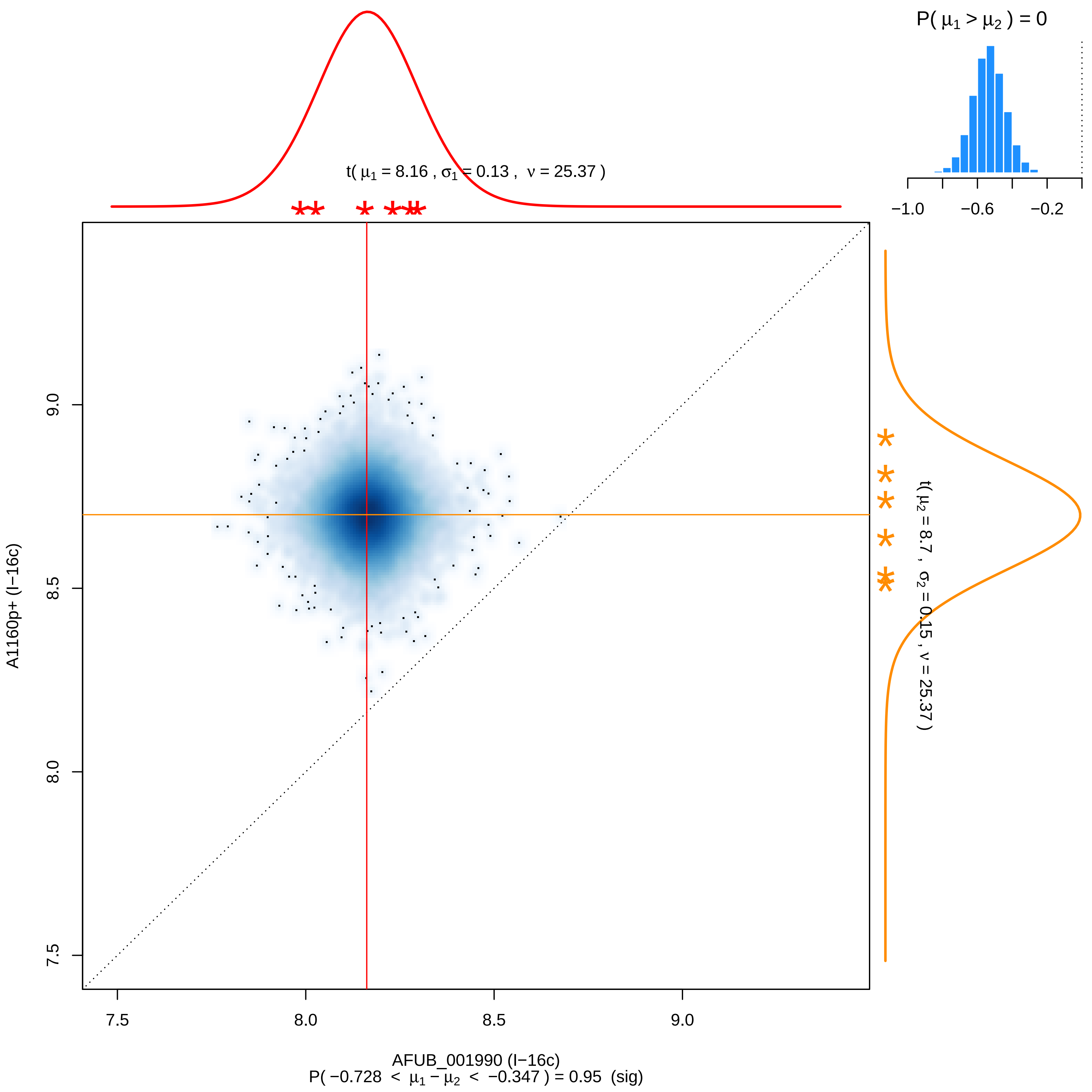

P( μ1 > μ2 ) = 0
−1.0
−0.6
−0.2
t( μ1 = 8.16 , σ1 = 0.13 , ν = 25.37 )
t( μ2 = 8.7 , σ2 = 0.15 , ν = 25.37 )
9.0
8.5
8.0
7.5
A1160p+ (I−16c)
7.5
8.0
8.5
9.0
AFUB_001990 (I−16c)
P( −0.728 < μ1 − μ2 < −0.347 ) = 0.95 (sig)

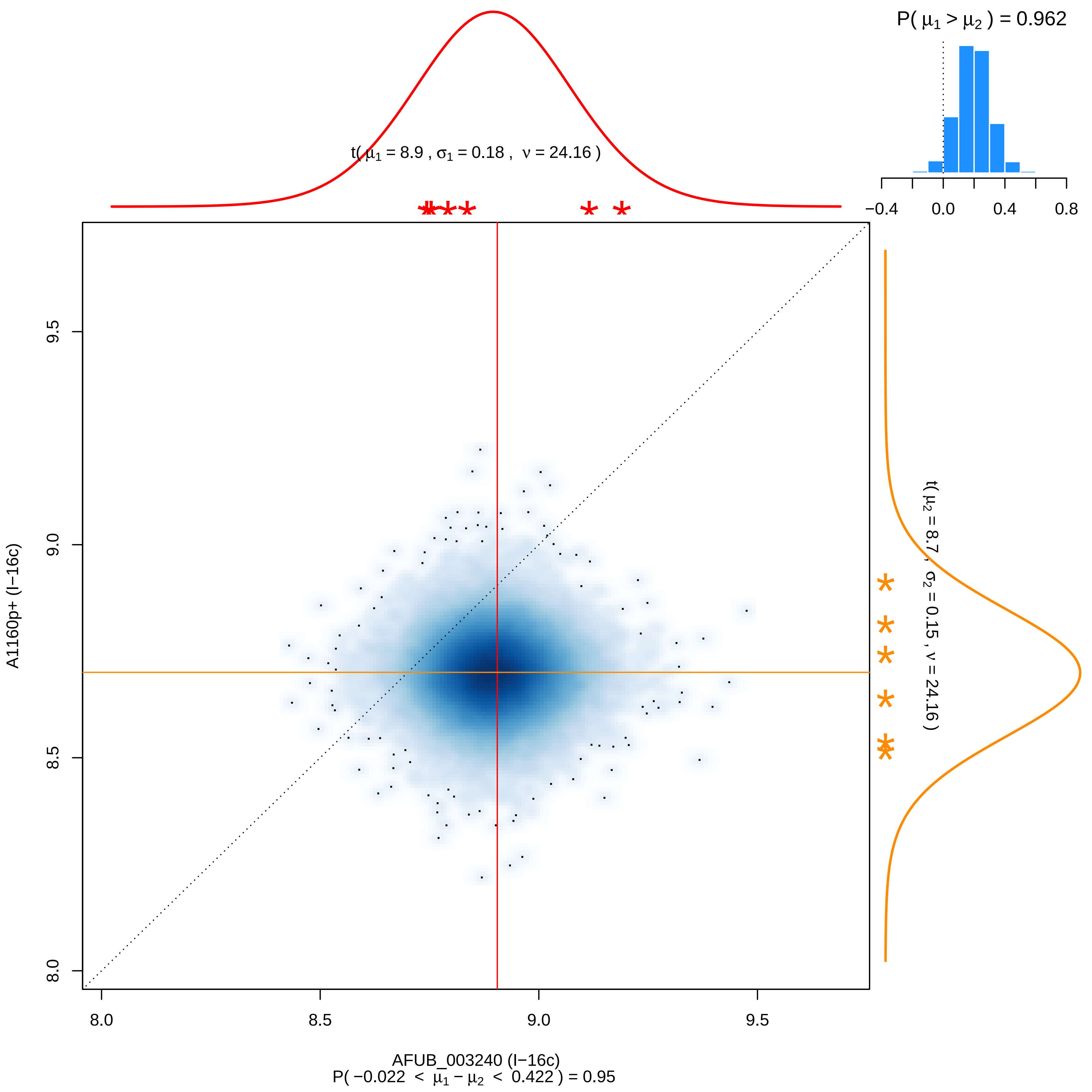

P( μ1 > μ2 ) = 0.962
−0.4
0.0
0.4
0.8
t( μ1 = 8.9 , σ1 = 0.18 , ν = 24.16 )
t( μ2 = 8.7 , σ2 = 0.15 , ν = 24.16 )
9.5
9.0
8.5
8.0
A1160p+ (I−16c)
8.0
8.5
9.0
9.5
AFUB_003240 (I−16c)
P( −0.022 < μ1 − μ2 < 0.422 ) = 0.95

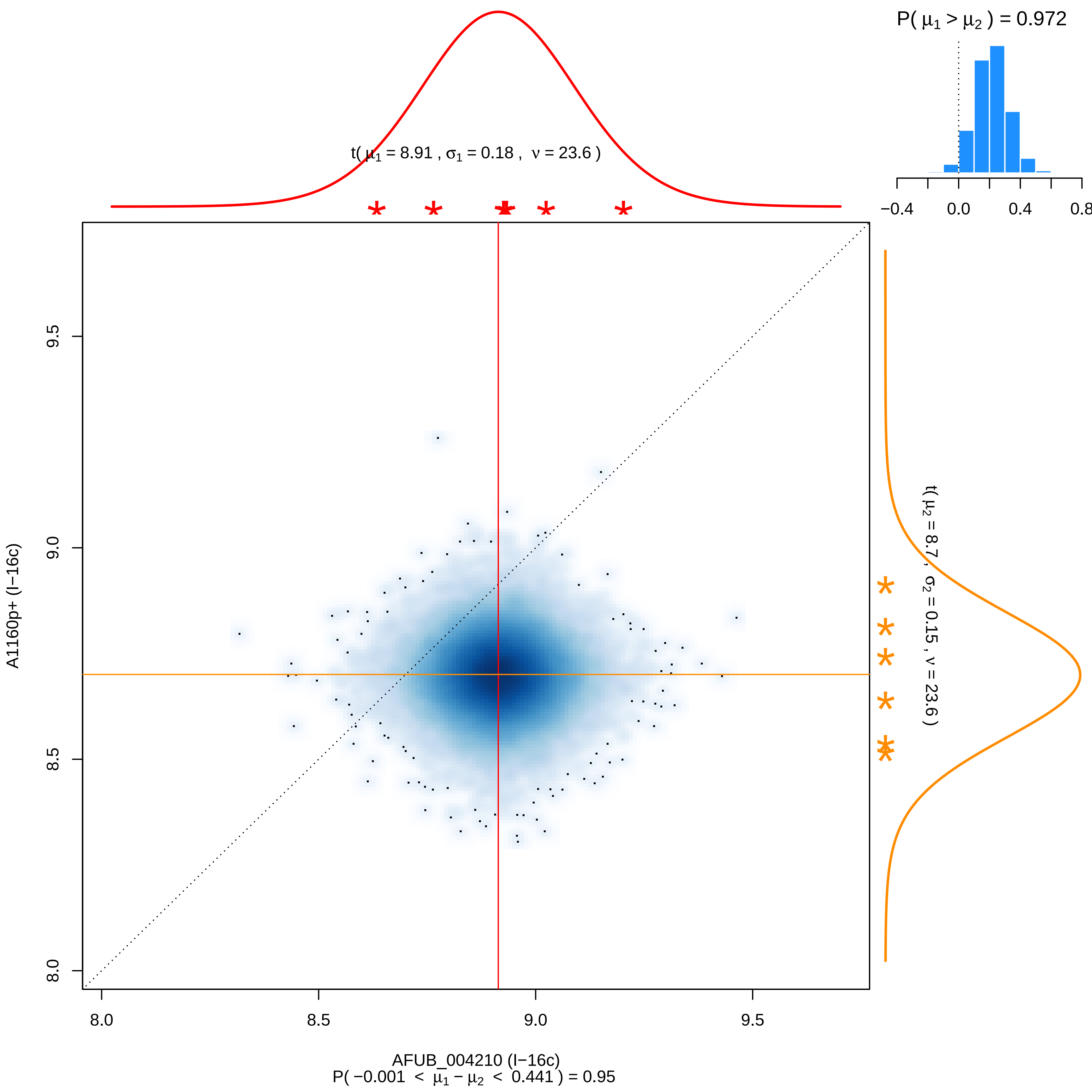

P( μ1 > μ2 ) = 0.972
t( μ1 = 8.91 , σ1 = 0.18 , ν = 23.6 )
−0.4
0.0
0.4
0.8
9.5
9.0
8.5
8.0
A1160p+ (I−16c)
t( μ2 = 8.7 , σ2 = 0.15 , ν = 23.6 )
8.0
8.5
9.0
9.5
AFUB_004210 (I−16c)
P( −0.001 < μ1 − μ2 < 0.441 ) = 0.95

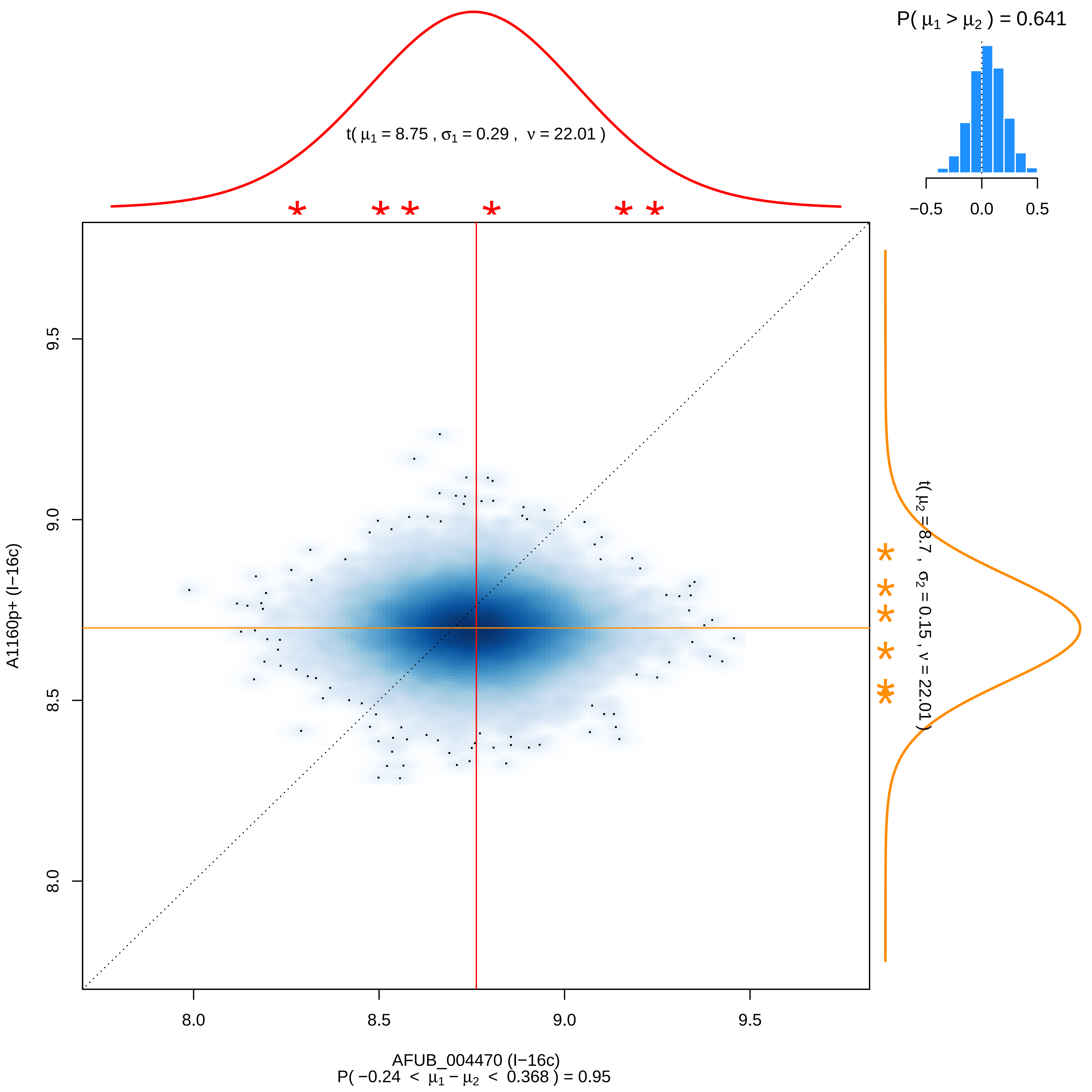

P( μ1 > μ2 ) = 0.641
−0.5
0.0
0.5
t( μ1 = 8.75 , σ1 = 0.29 , ν = 22.01 )
t( μ2 = 8.7 , σ2 = 0.15 , ν = 22.01 )
A1160p+ (I−16c)
8.0
8.5
9.0
9.5
AFUB_004470 (I−16c)
P( −0.24 < μ1 − μ2 < 0.368 ) = 0.95

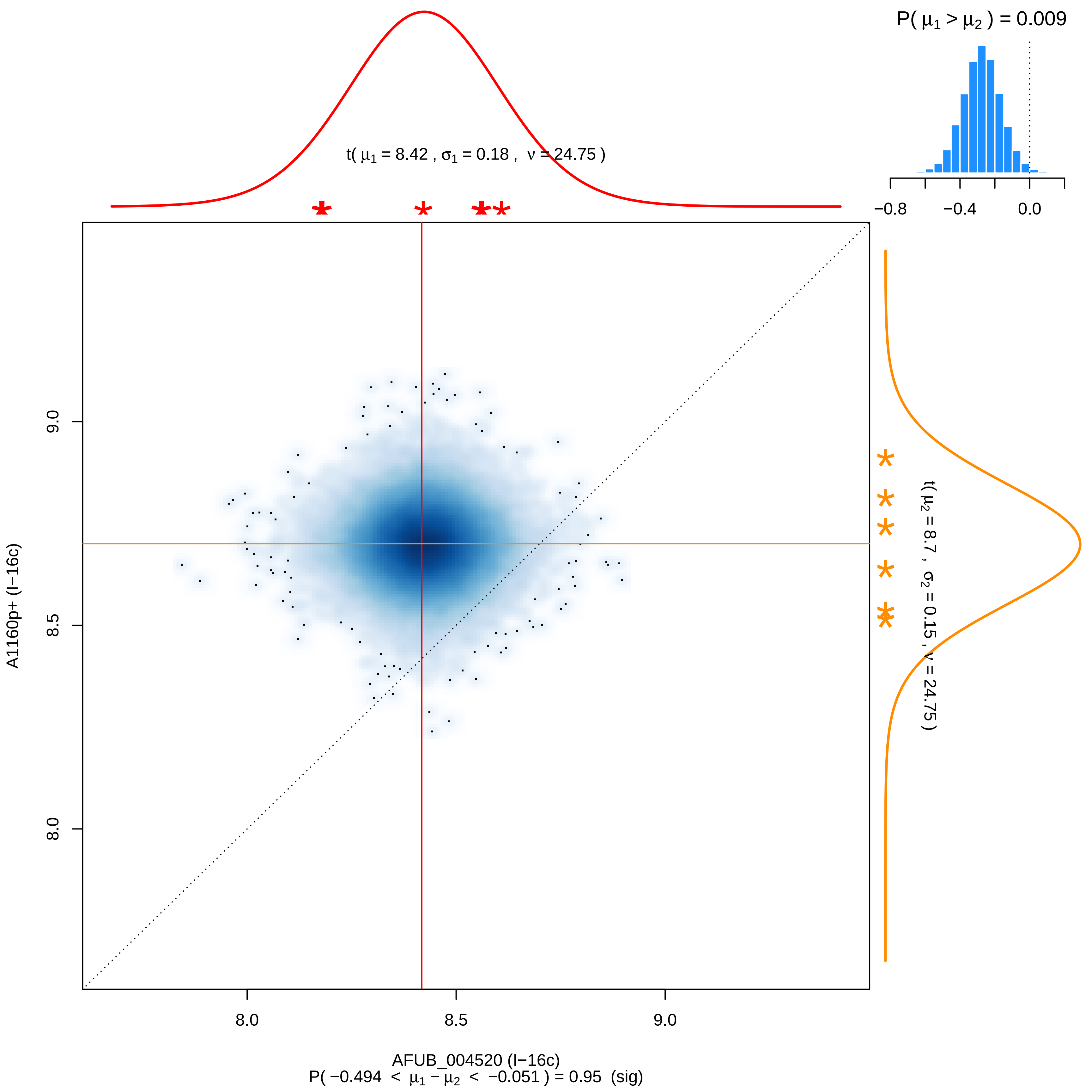

P( μ1 > μ2 ) = 0.009
−0.8
−0.4
0.0
t( μ1 = 8.42 , σ1 = 0.18 , ν = 24.75 )
t( μ2 = 8.7 , σ2 = 0.15 , ν = 24.75 )
A1160p+ (I−16c)
9.0
8.5
8.0
8.0
8.5
9.0
AFUB_004520 (I−16c)
P( −0.494 < μ1 − μ2 < −0.051 ) = 0.95 (sig)

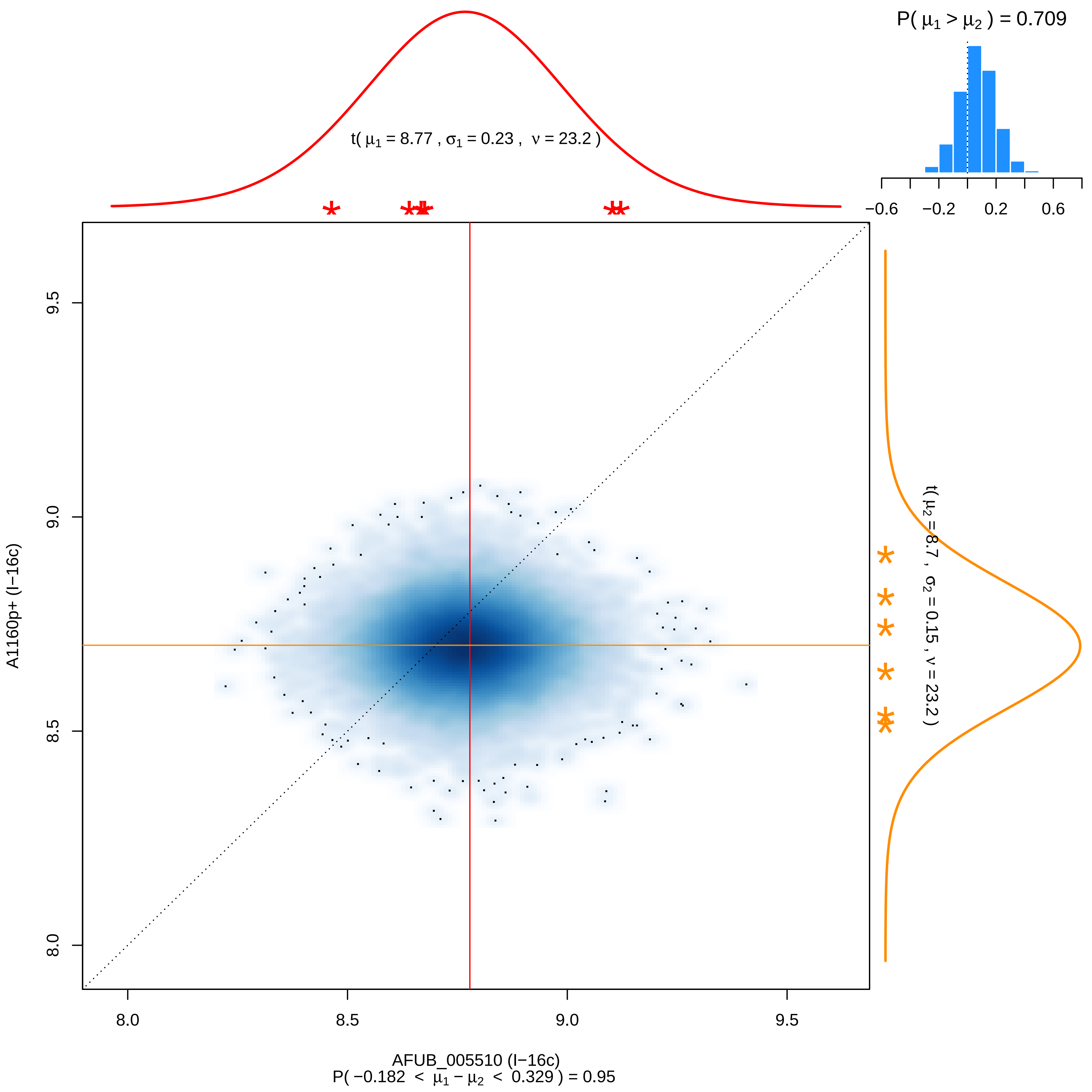

P( μ1 > μ2 ) = 0.709
t( μ1 = 8.77 , σ1 = 0.23 , ν = 23.2 )
−0.6
−0.2
0.2
0.6
t( μ2 = 8.7 , σ2 = 0.15 , ν = 23.2 )
9.5
9.0
8.5
8.0
A1160p+ (I−16c)
8.0
8.5
9.0
9.5
AFUB_005510 (I−16c)
P( −0.182 < μ1 − μ2 < 0.329 ) = 0.95

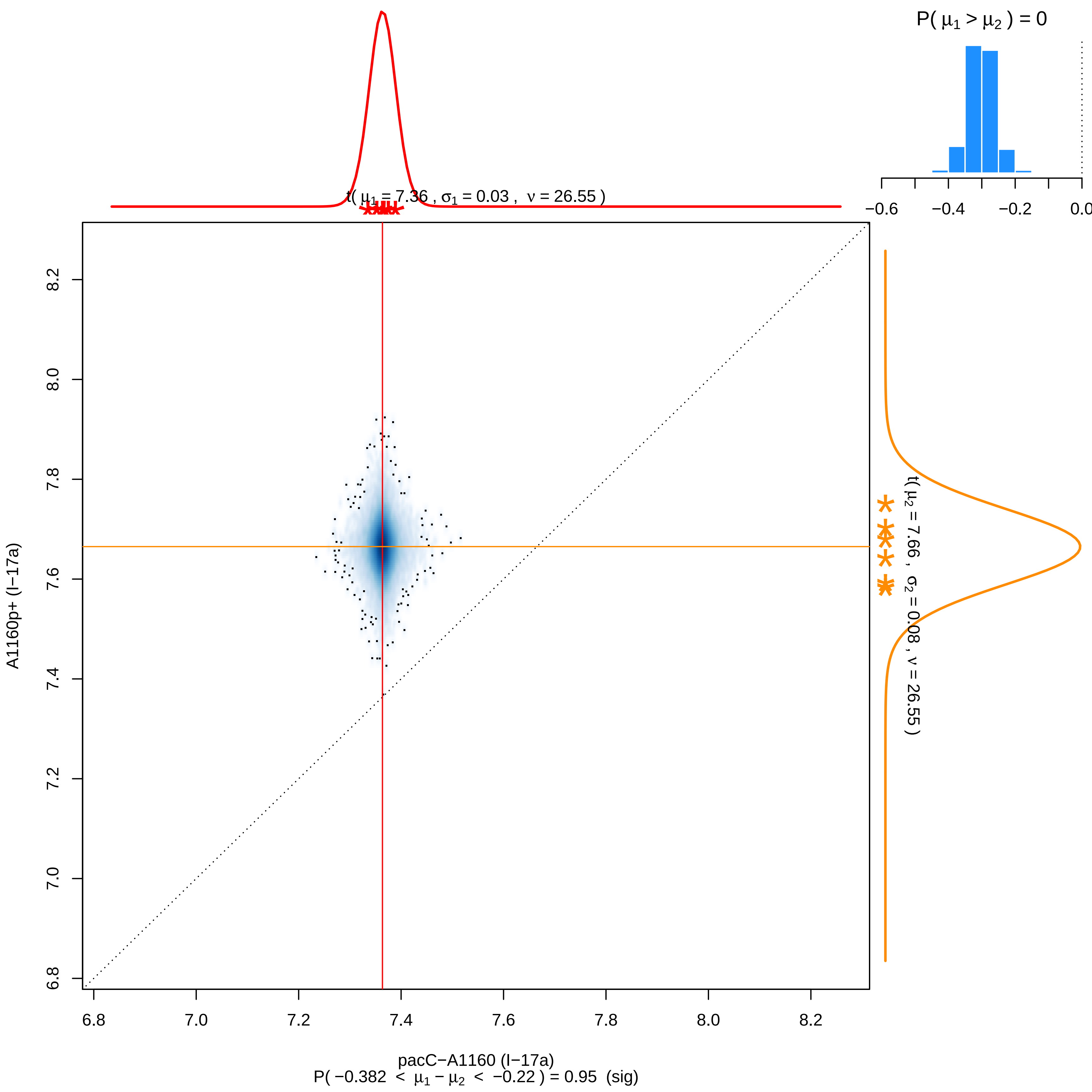

P( $\mu_1$ > $\mu_2$ ) = 0
−0.6
−0.4
−0.2
0.0
t( $\mu_1$ = 7.36 , $\sigma_1$ = 0.03 , $\nu$ = 26.55 )
t( $\mu_2$ = 7.66 , $\sigma_2$ = 0.08 , $\nu$ = 26.55 )
A1160p+ (I−17a)
6.8
7.0
7.2
7.4
7.6
7.8
8.0
8.2
pacC−A1160 (I−17a)
P( −0.382 < $\mu_1 - \mu_2$ < −0.22 ) = 0.95 (sig)

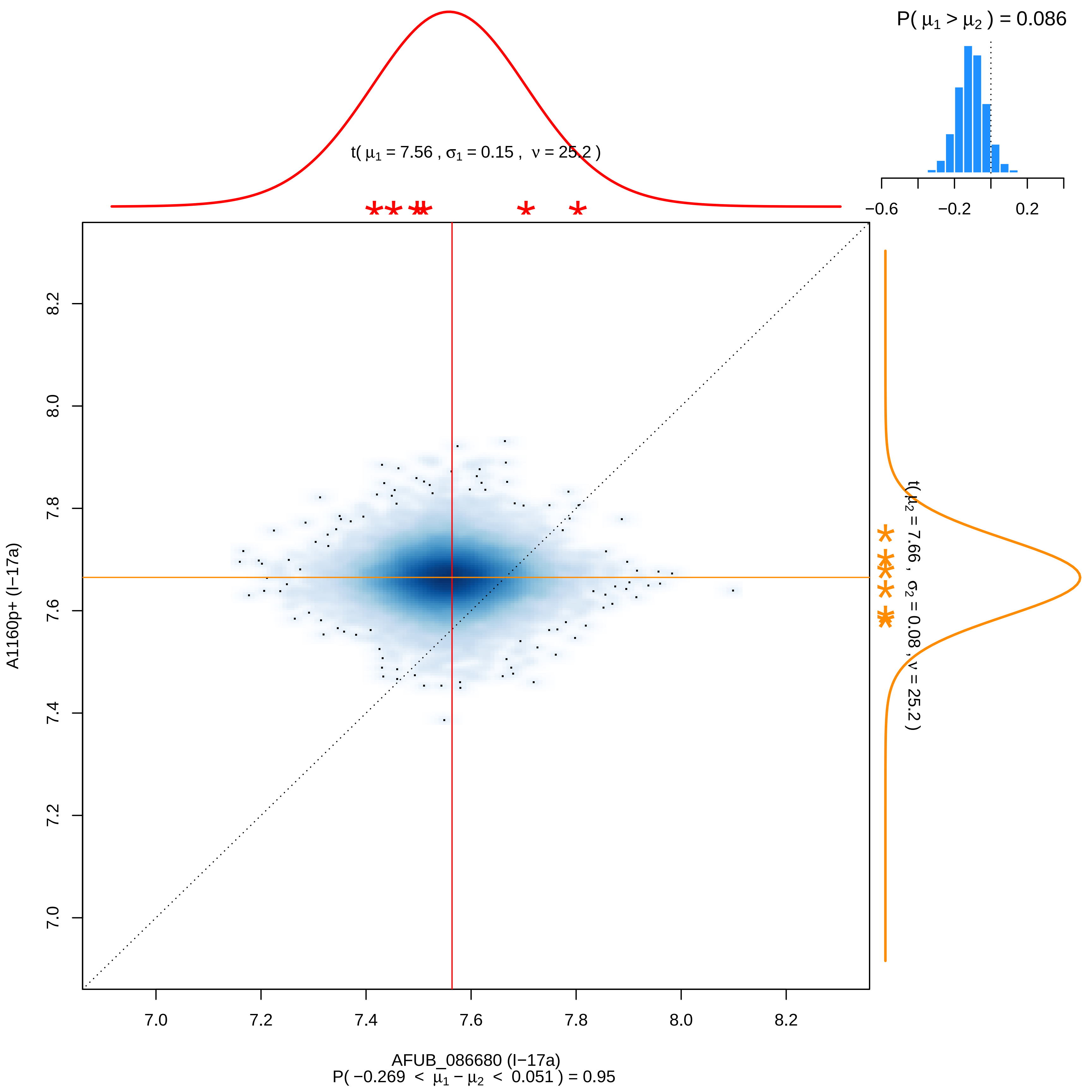

P( $\mu_1 > \mu_2$ ) = 0.086
−0.6
−0.2
0.2
t( $\mu_1$ = 7.56 , $\sigma_1$ = 0.15 , $\nu$ = 25.2 )
t( $\mu_2$ = 7.66 , $\sigma_2$ = 0.08 , $\nu$ = 25.2 )
A1160p+ (I−17a)
8.2
8.0
7.8
7.6
7.4
7.2
7.0
AFUB_086680 (I−17a)
P( −0.269 < $\mu_1 - \mu_2$ < 0.051 ) = 0.95

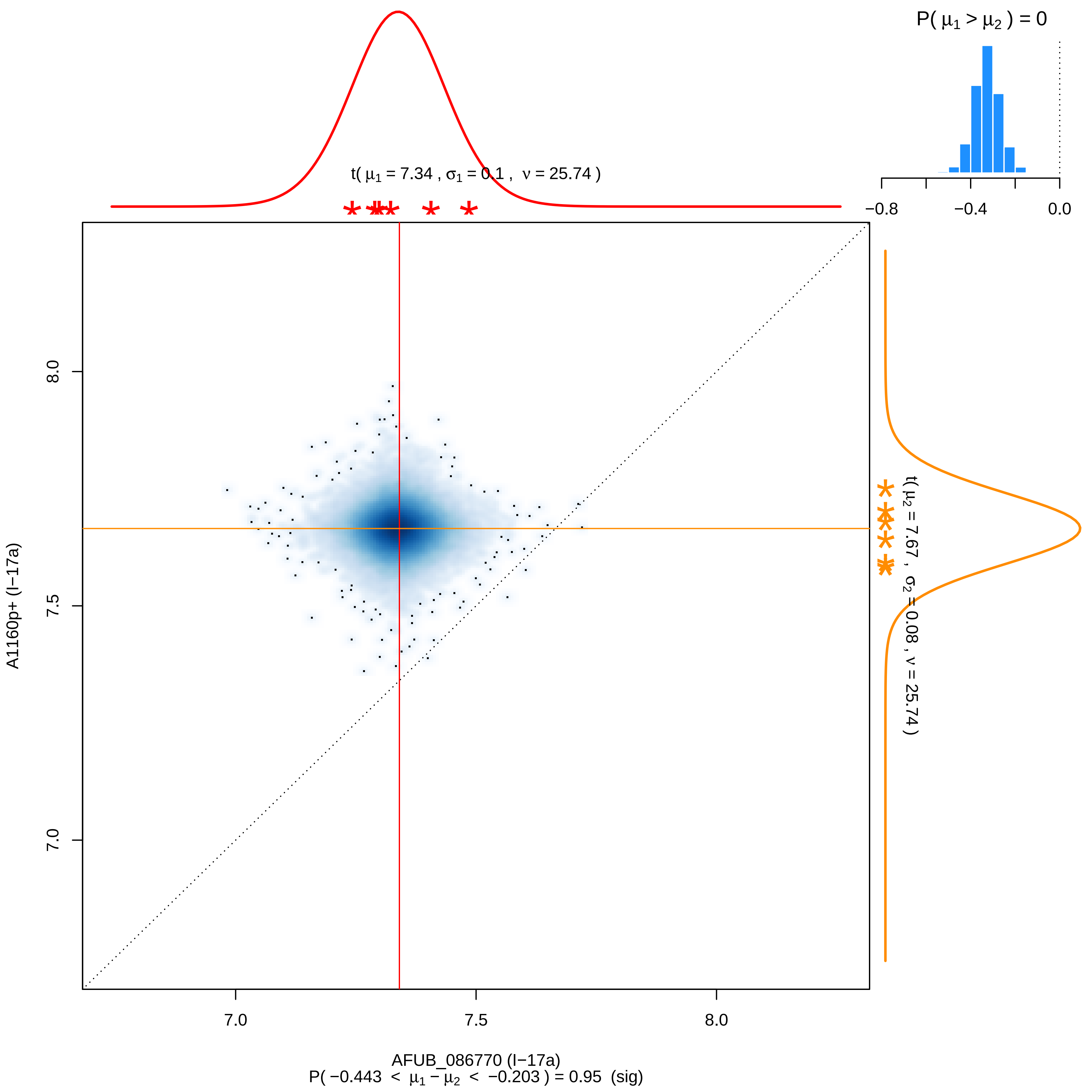

P( μ1 > μ2 ) = 0
−0.8
−0.4
0.0
t( μ1 = 7.34 , σ1 = 0.1 , ν = 25.74 )
t( μ2 = 7.67 , σ2 = 0.08 , ν = 25.74 )
8.0
7.5
7.0
A1160p+ (I−17a)
7.0
7.5
8.0
AFUB_086770 (I−17a)
P( −0.443 < μ1 − μ2 < −0.203 ) = 0.95 (sig)

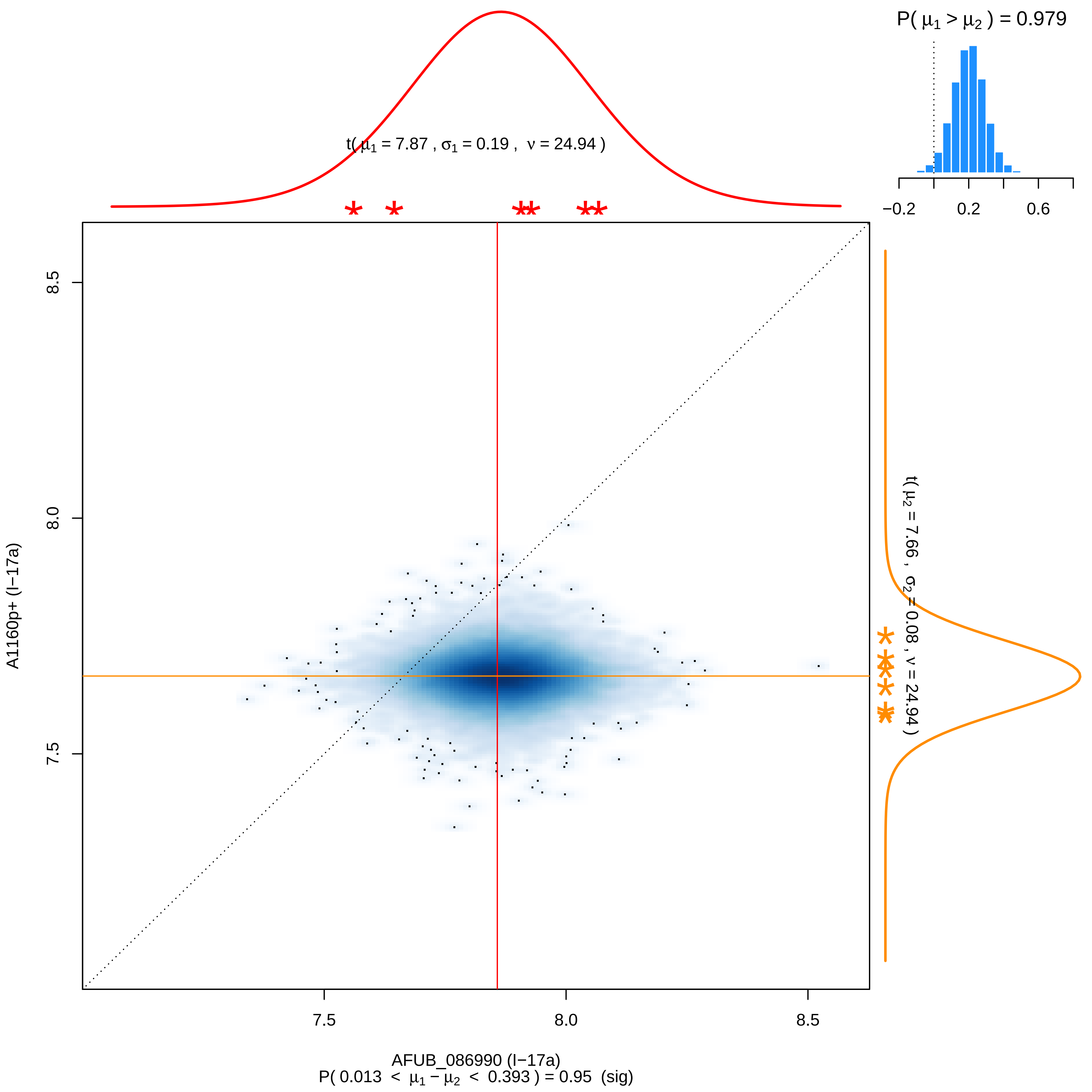

P( μ1 > μ2 ) = 0.979
−0.2
0.2
0.6
t( μ1 = 7.87 , σ1 = 0.19 , ν = 24.94 )
t( μ2 = 7.66 , σ2 = 0.08 , ν = 24.94 )
A1160p+ (I−17a)
8.5
8.0
7.5
7.5
8.0
8.5
AFUB_086990 (I−17a)
P( 0.013 < μ1 − μ2 < 0.393 ) = 0.95 (sig)

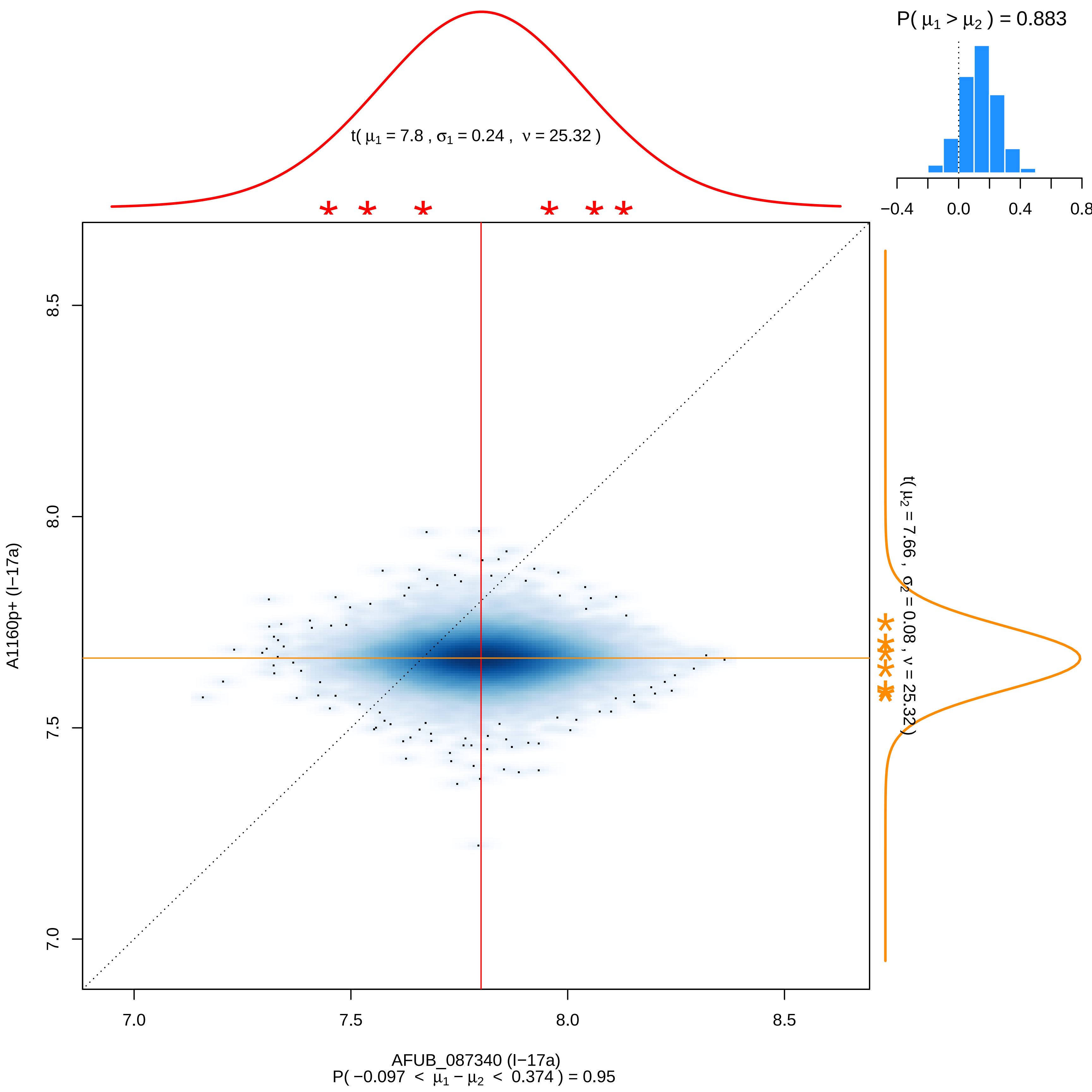

P( μ1 > μ2 ) = 0.883
−0.4
0.0
0.4
0.8
t( μ1 = 7.8 , σ1 = 0.24 , ν = 25.32 )
t( μ2 = 7.66 , σ2 = 0.08 , ν = 25.32 )
8.5
8.0
7.5
7.0
A1160p+ (I−17a)
7.0
7.5
8.0
8.5
AFUB_087340 (I−17a)
P( −0.097 < μ1 − μ2 < 0.374 ) = 0.95

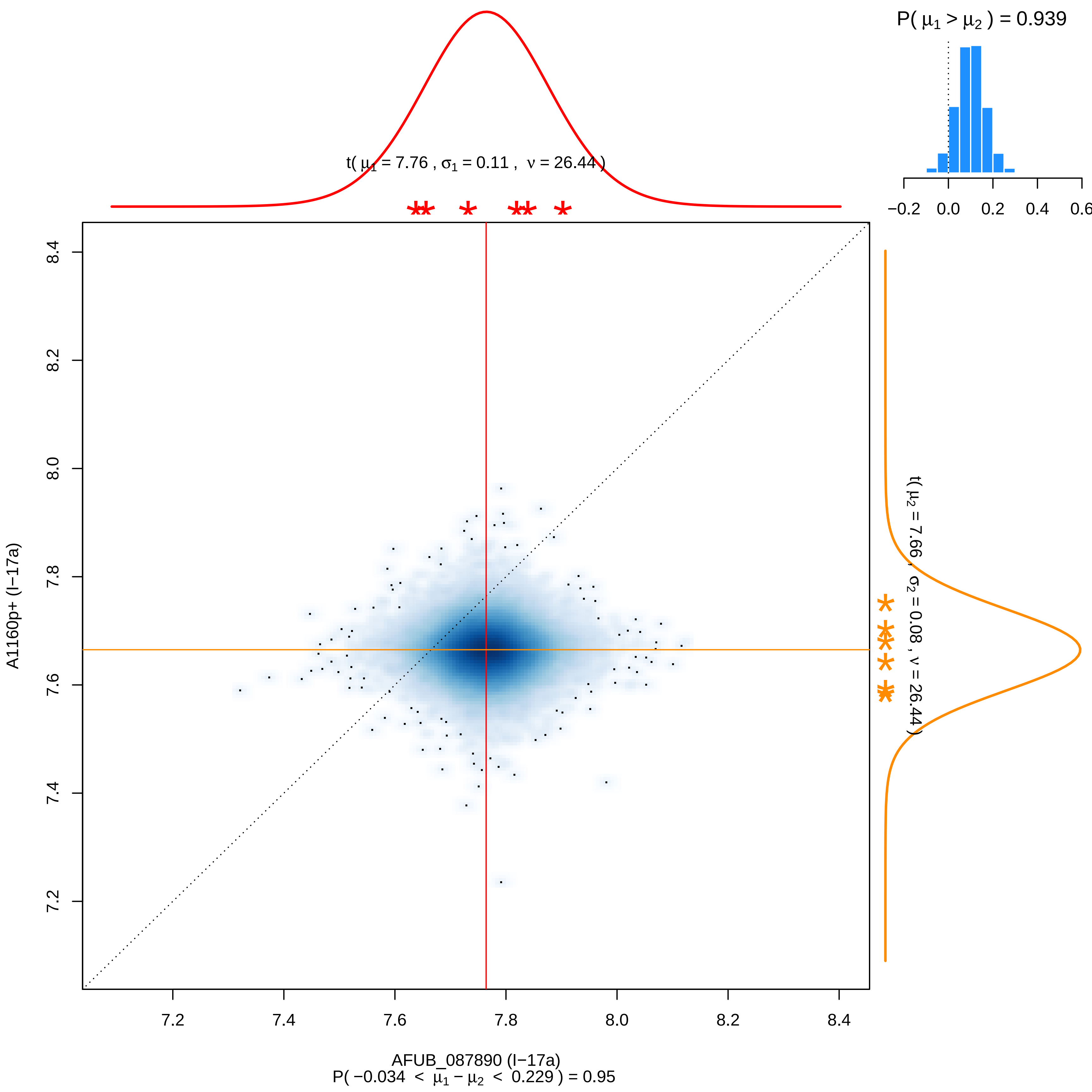

P( μ1 > μ2 ) = 0.939
−0.2 0.0 0.2 0.4 0.6
t( μ1 = 7.76 , σ1 = 0.11 , ν = 26.44 )
t( μ2 = 7.66 , σ2 = 0.08 , ν = 26.44 )
7.2 7.4 7.6 7.8 8.0 8.2 8.4
A1160p+ (I−17a)
AFUB_087890 (I−17a)
P( −0.034 < μ1 − μ2 < 0.229 ) = 0.95

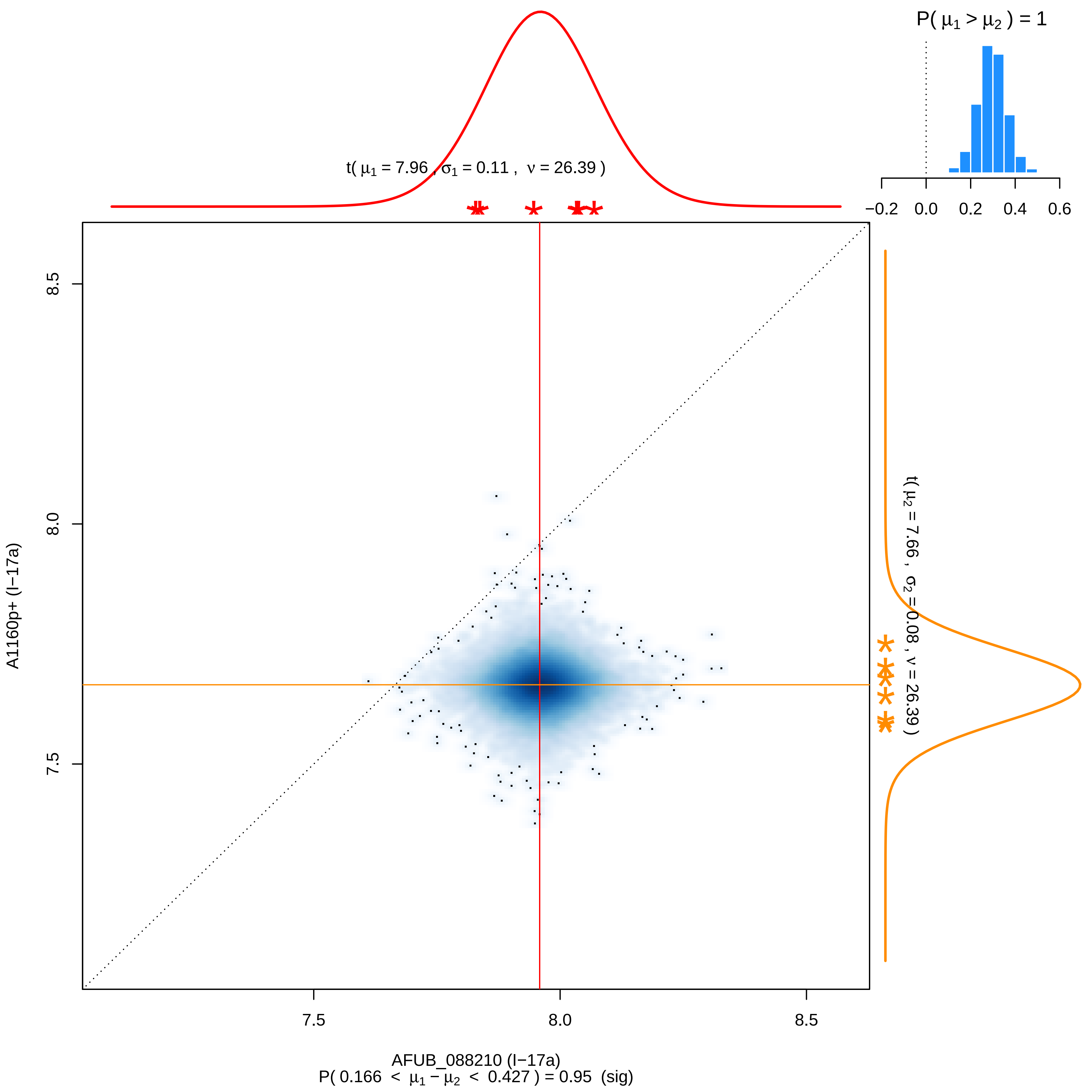

P( μ1 > μ2 ) = 1
−0.2 0.0 0.2 0.4 0.6
t( μ1 = 7.96 , σ1 = 0.11 , ν = 26.39 )
t( μ2 = 7.66 , σ2 = 0.08 , ν = 26.39 )
A1160p+ (I−17a)
8.5
8.0
7.5
7.5 8.0 8.5
AFUB_088210 (I−17a)
P( 0.166 < μ1 − μ2 < 0.427 ) = 0.95 (sig)

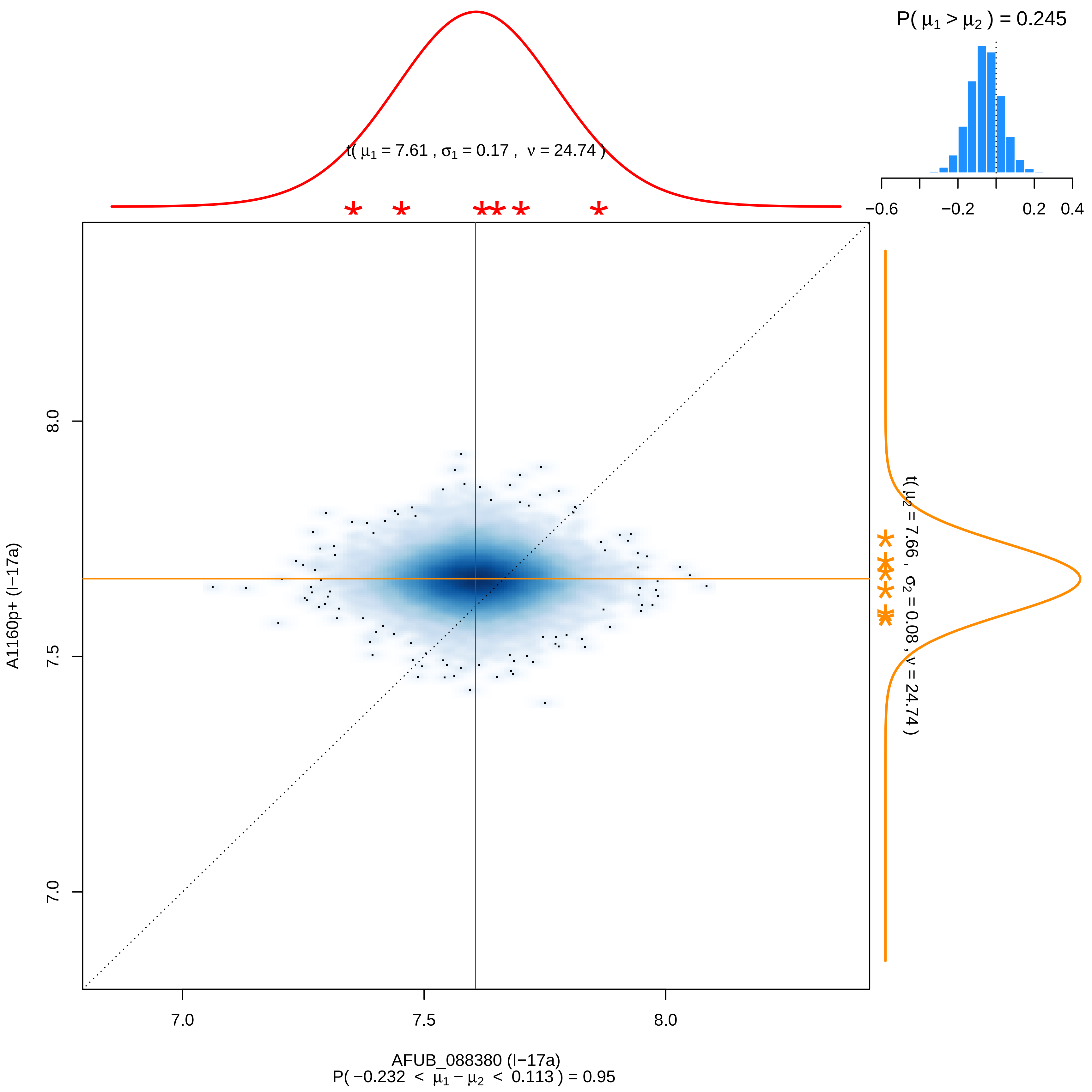
P( $\mu_1 > \mu_2$ ) = 0.245
−0.6
−0.2
0.2
0.4
t( $\mu_1$ = 7.61 , $\sigma_1$ = 0.17 , $\nu$ = 24.74 )
t( $\mu_2$ = 7.66 , $\sigma_2$ = 0.08 , $\nu$ = 24.74 )
8.0
7.5
7.0
A1160p+ (I−17a)
7.0
7.5
8.0
AFUB_088380 (I−17a)
P( −0.232 < $\mu_1 - \mu_2$ < 0.113 ) = 0.95

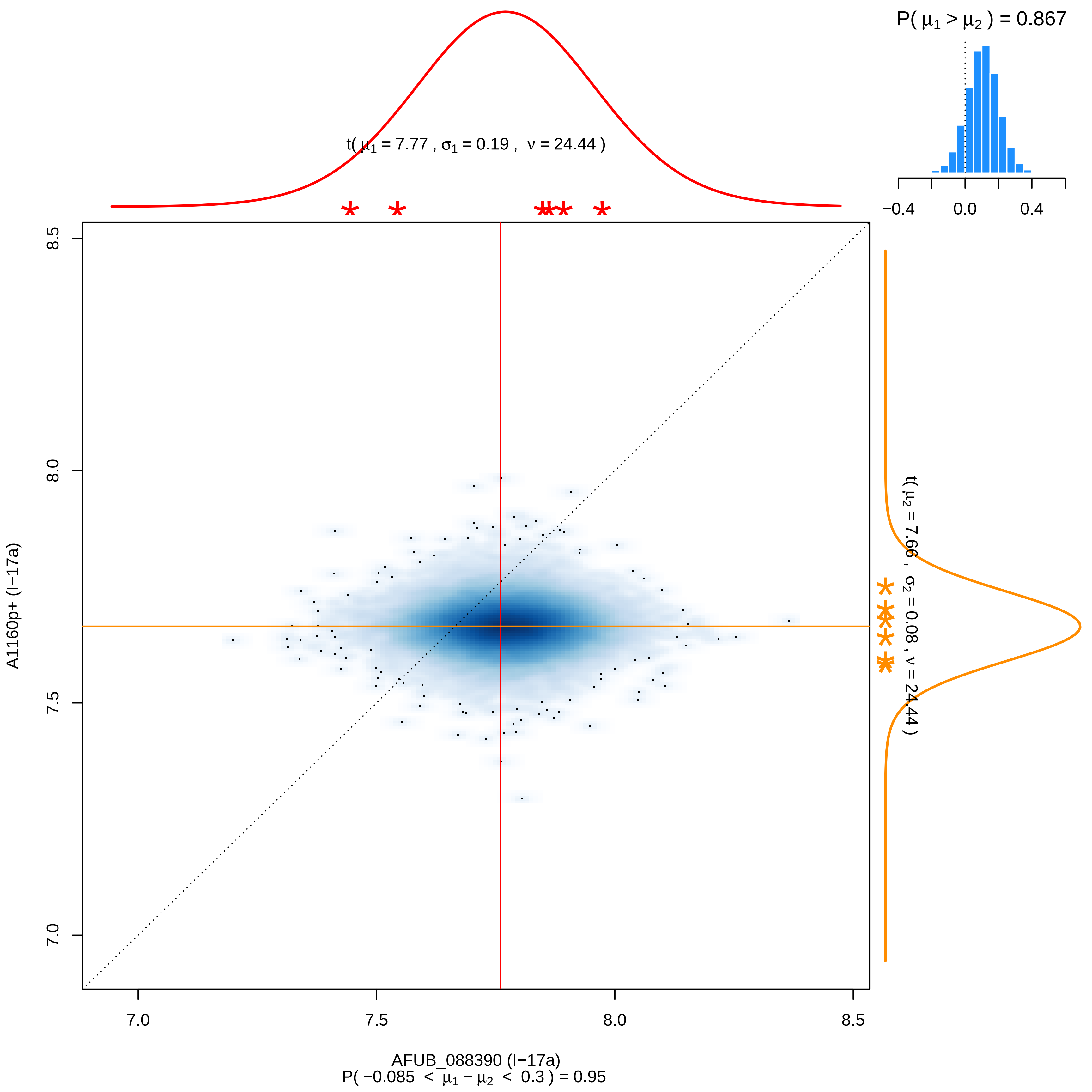
P( μ1 > μ2 ) = 0.867
−0.4
0.0
0.4
t( μ1 = 7.77 , σ1 = 0.19 , ν = 24.44 )
8.5
8.0
7.5
7.0
A1160p+ (I−17a)
t( μ2 = 7.66 , σ2 = 0.08 , ν = 24.44 )
7.0
7.5
8.0
8.5
AFUB_088390 (I−17a)
P( −0.085 < μ1 − μ2 < 0.3 ) = 0.95

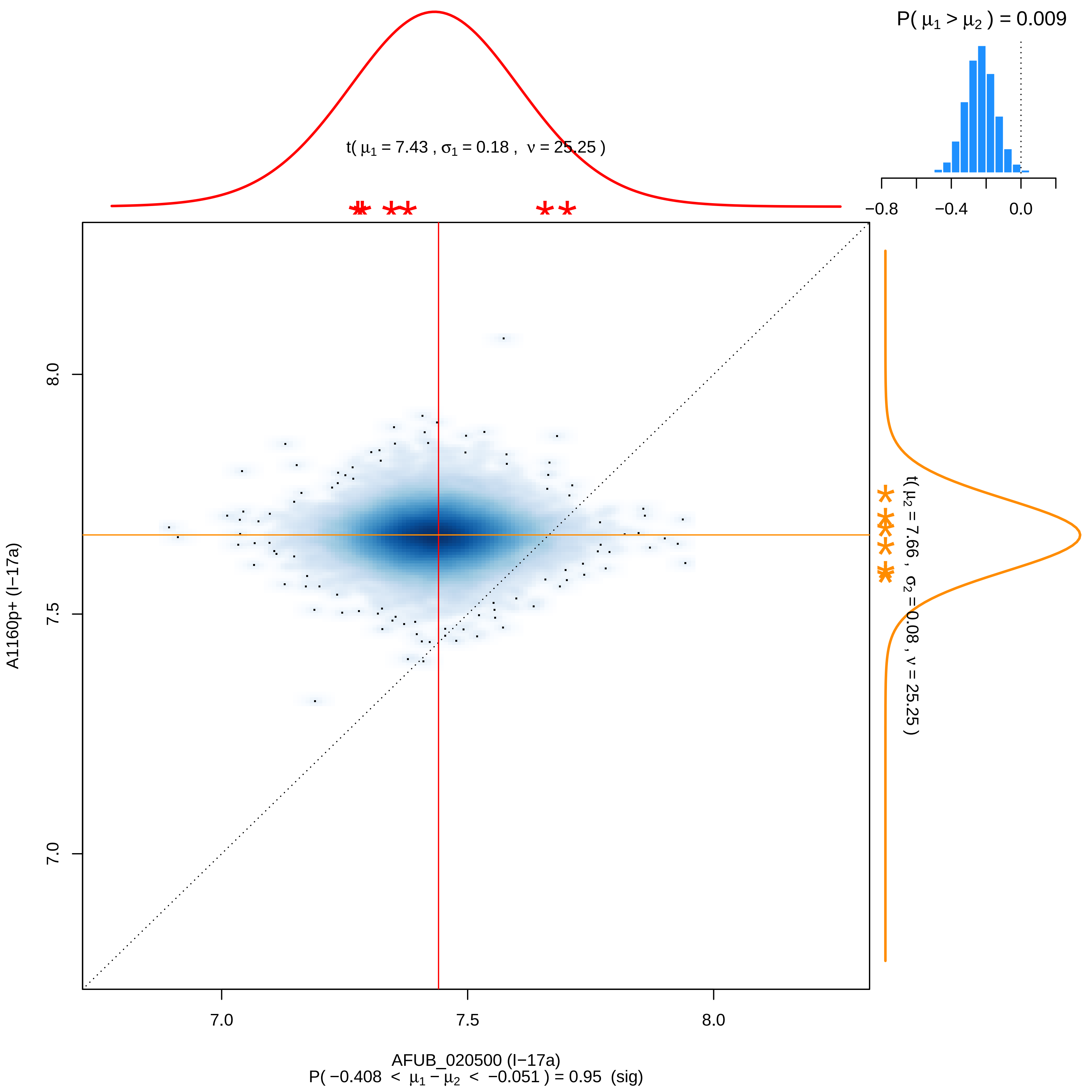

P( μ1 > μ2 ) = 0.009
−0.8 −0.4 0.0
t( μ1 = 7.43 , σ1 = 0.18 , ν = 25.25 )
t( μ2 = 7.66 , σ2 = 0.08 , ν = 25.25 )
8.0
7.5
7.0
A1160p+ (I−17a)
7.0 7.5 8.0
AFUB_020500 (I−17a)
P( −0.408 < μ1 − μ2 < −0.051 ) = 0.95 (sig)

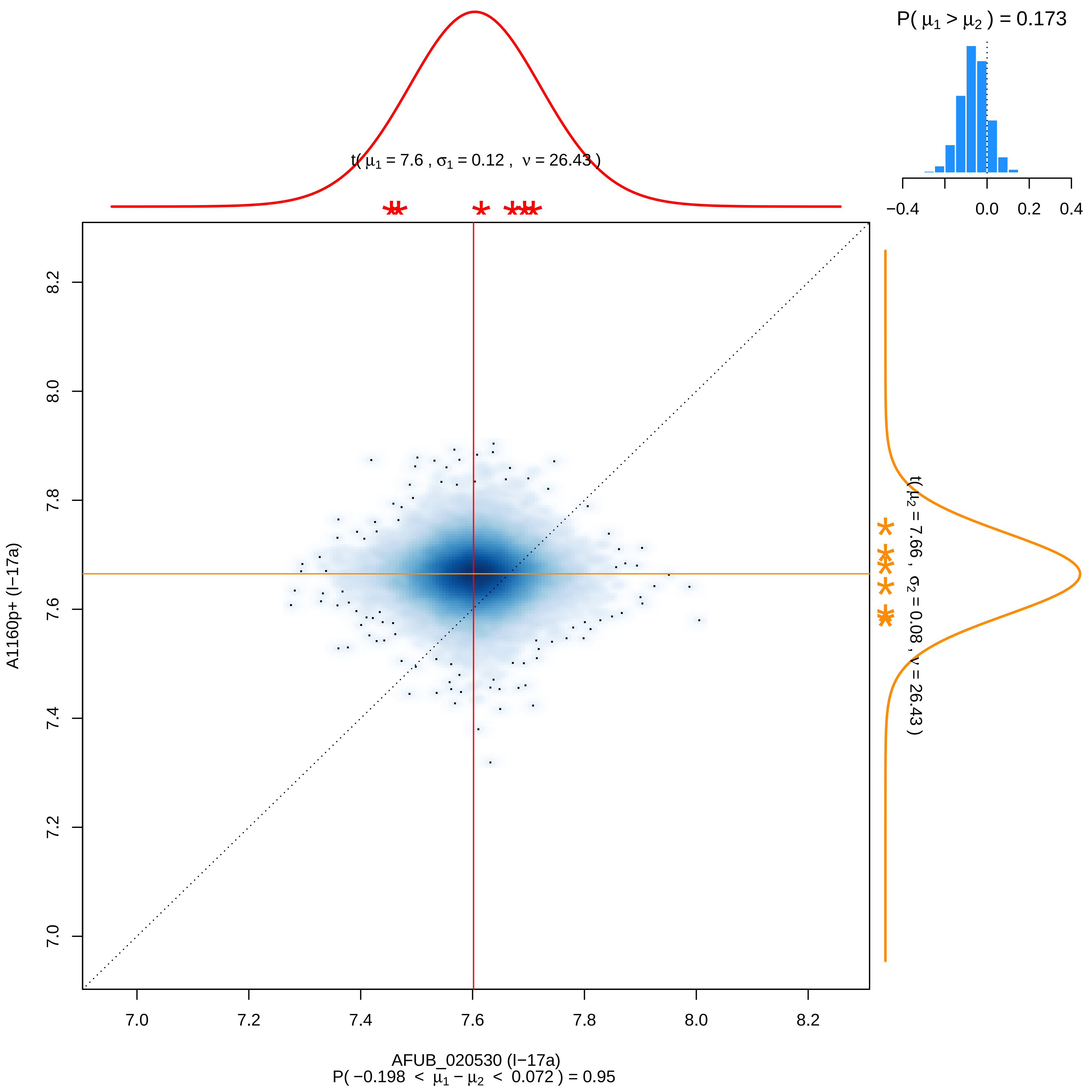

P( μ1 > μ2 ) = 0.173
−0.4
0.0
0.2
0.4
t( μ1 = 7.6 , σ1 = 0.12 , ν = 26.43 )
t( μ2 = 7.66 , σ2 = 0.08 , ν = 26.43 )
8.2
8.0
7.8
7.6
7.4
7.2
7.0
A1160p+ (I−17a)
7.0
7.2
7.4
7.6
7.8
8.0
8.2
AFUB_020530 (I−17a)
P( −0.198 < μ1 − μ2 < 0.072 ) = 0.95

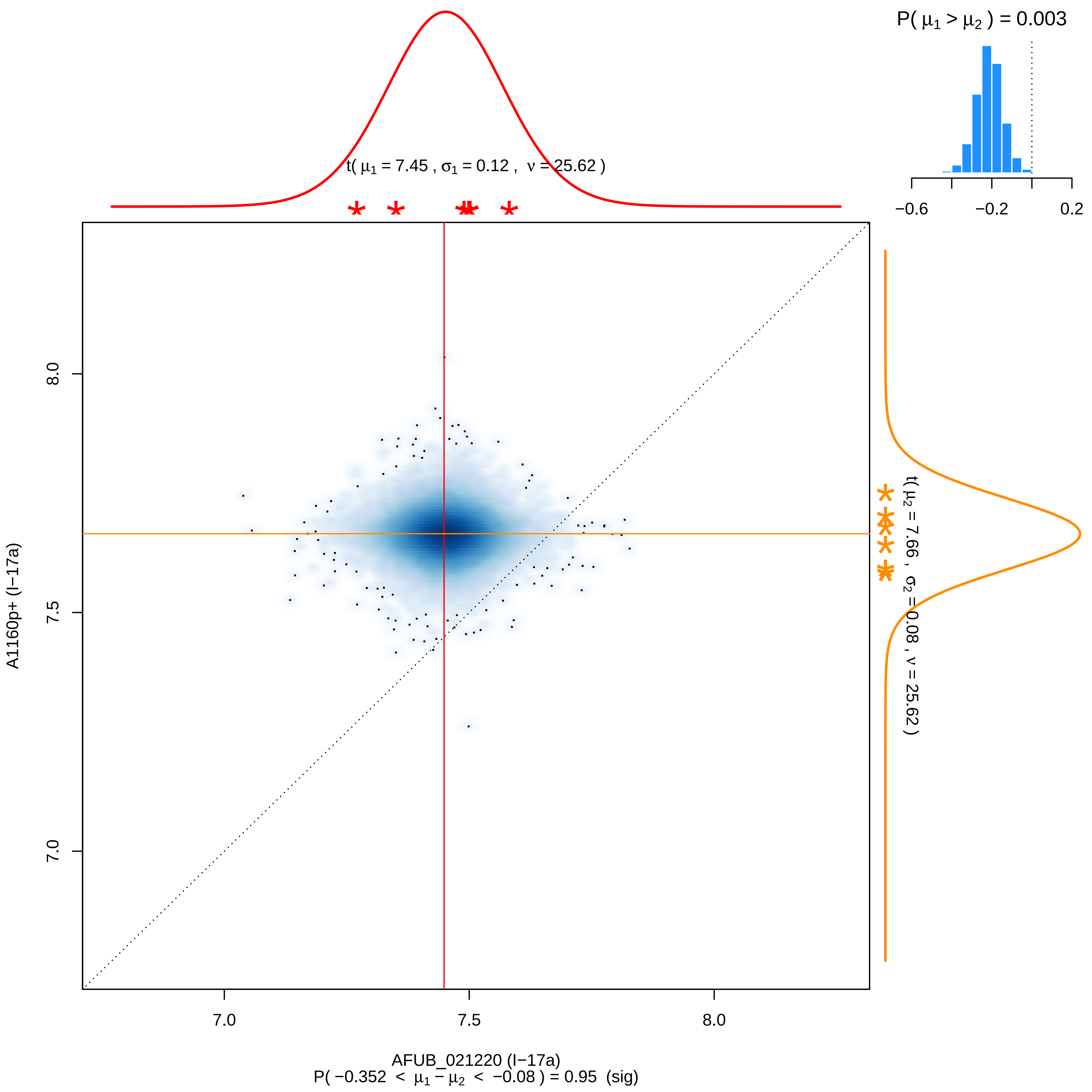

P( μ1 > μ2 ) = 0.003
−0.6
−0.2
0.2
t( μ1 = 7.45 , σ1 = 0.12 , ν = 25.62 )
t( μ2 = 7.66 , σ2 = 0.08 , ν = 25.62 )
A1160p+ (I−17a)
8.0
7.5
7.0
7.0
7.5
8.0
AFUB_021220 (I−17a)
P( −0.352 < μ1 − μ2 < −0.08 ) = 0.95 (sig)

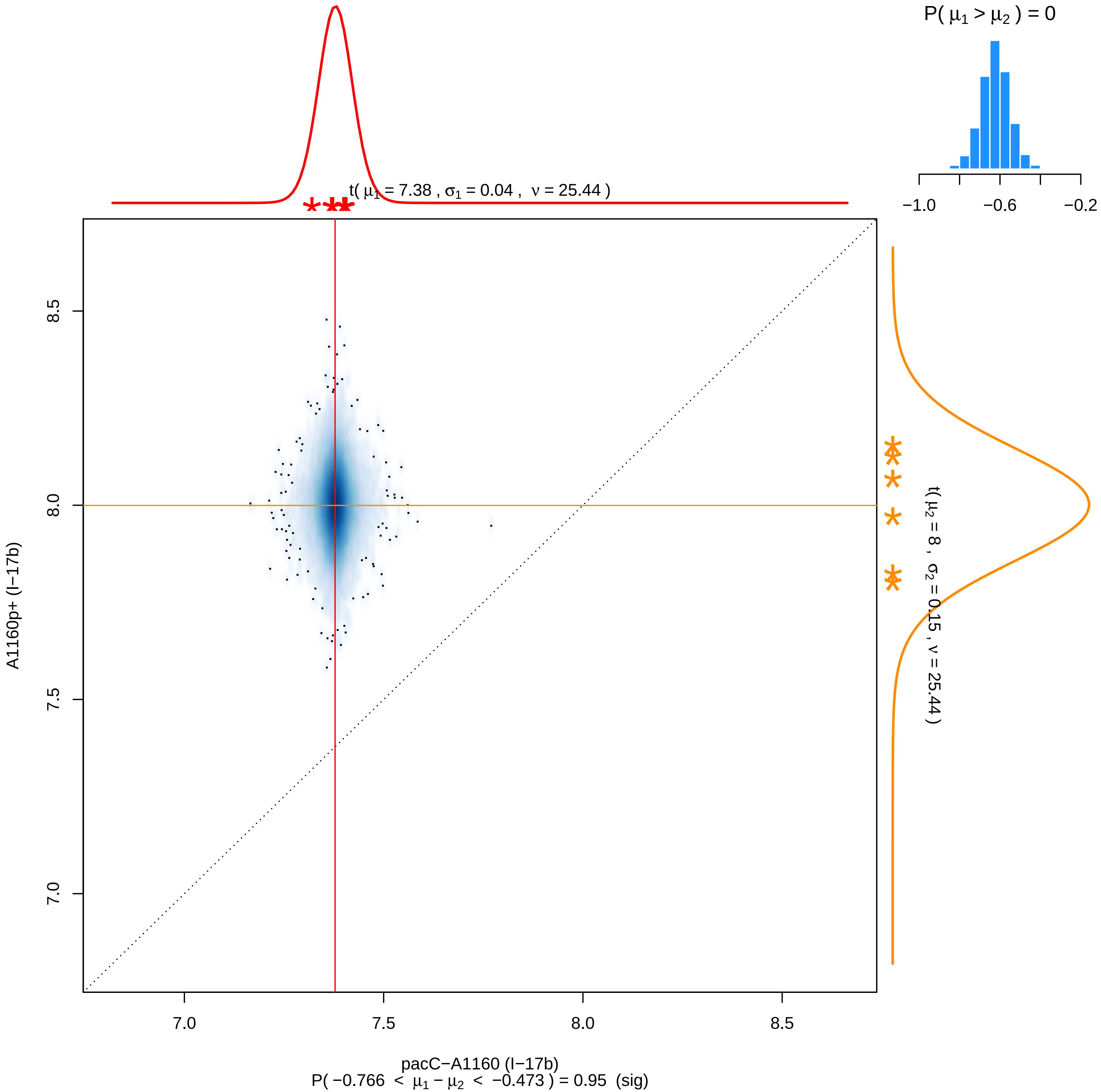

P( μ1 > μ2 ) = 0
−1.0
−0.6
−0.2
t( μ1 = 7.38 , σ1 = 0.04 , ν = 25.44 )
t( μ2 = 8 , σ2 = 0.15 , ν = 25.44 )
8.5
8.0
7.5
7.0
A1160p+ (I−17b)
7.0
7.5
8.0
8.5
pacC−A1160 (I−17b)
P( −0.766 < μ1 − μ2 < −0.473 ) = 0.95 (sig)

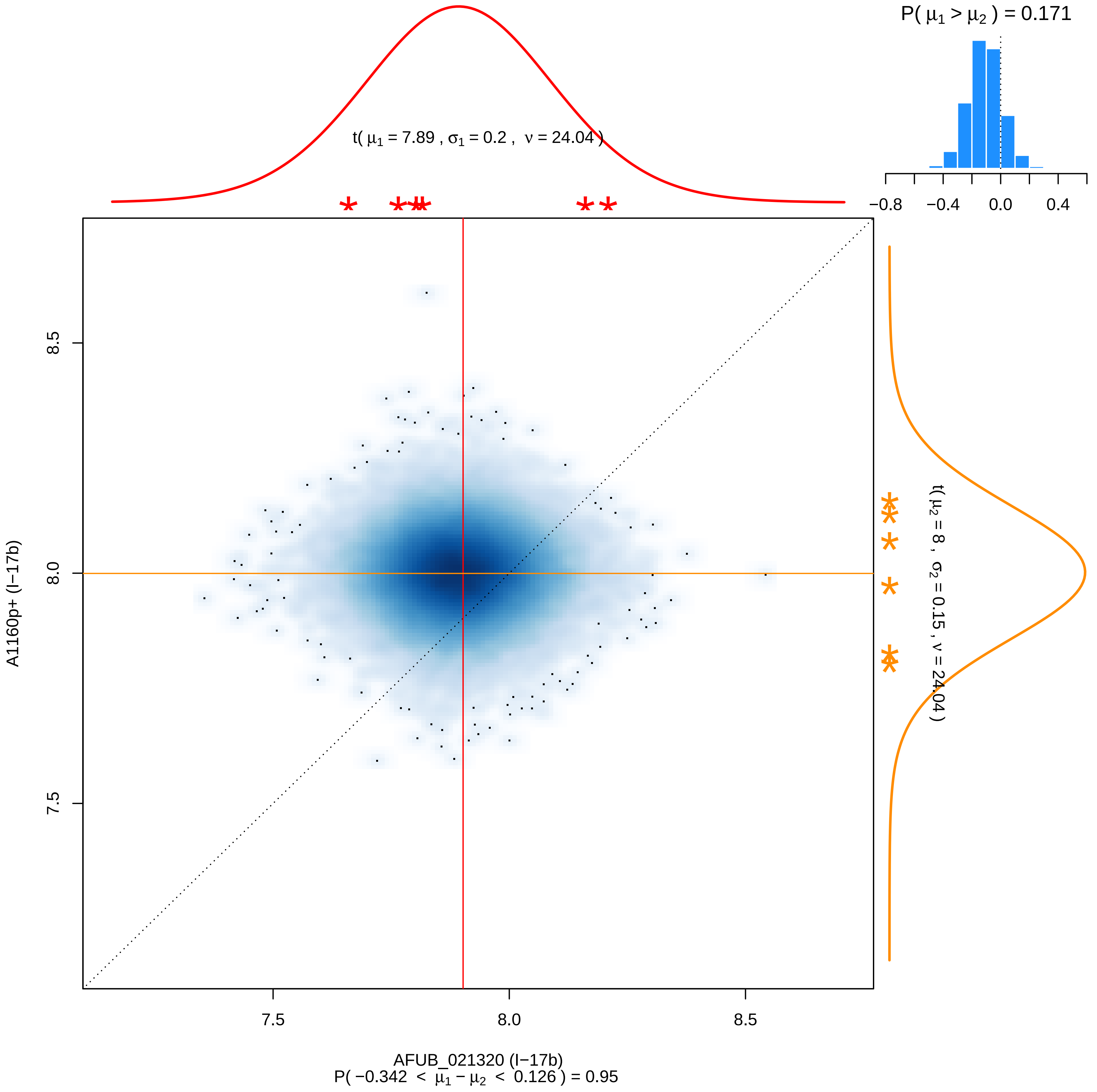

P( μ1 > μ2 ) = 0.171
−0.8
−0.4
0.0
0.4
t( μ1 = 7.89 , σ1 = 0.2 , ν = 24.04 )
t( μ2 = 8 , σ2 = 0.15 , ν = 24.04 )
8.5
8.0
7.5
A1160p+ (I−17b)
7.5
8.0
8.5
AFUB_021320 (I−17b)
P( −0.342 < μ1 − μ2 < 0.126 ) = 0.95

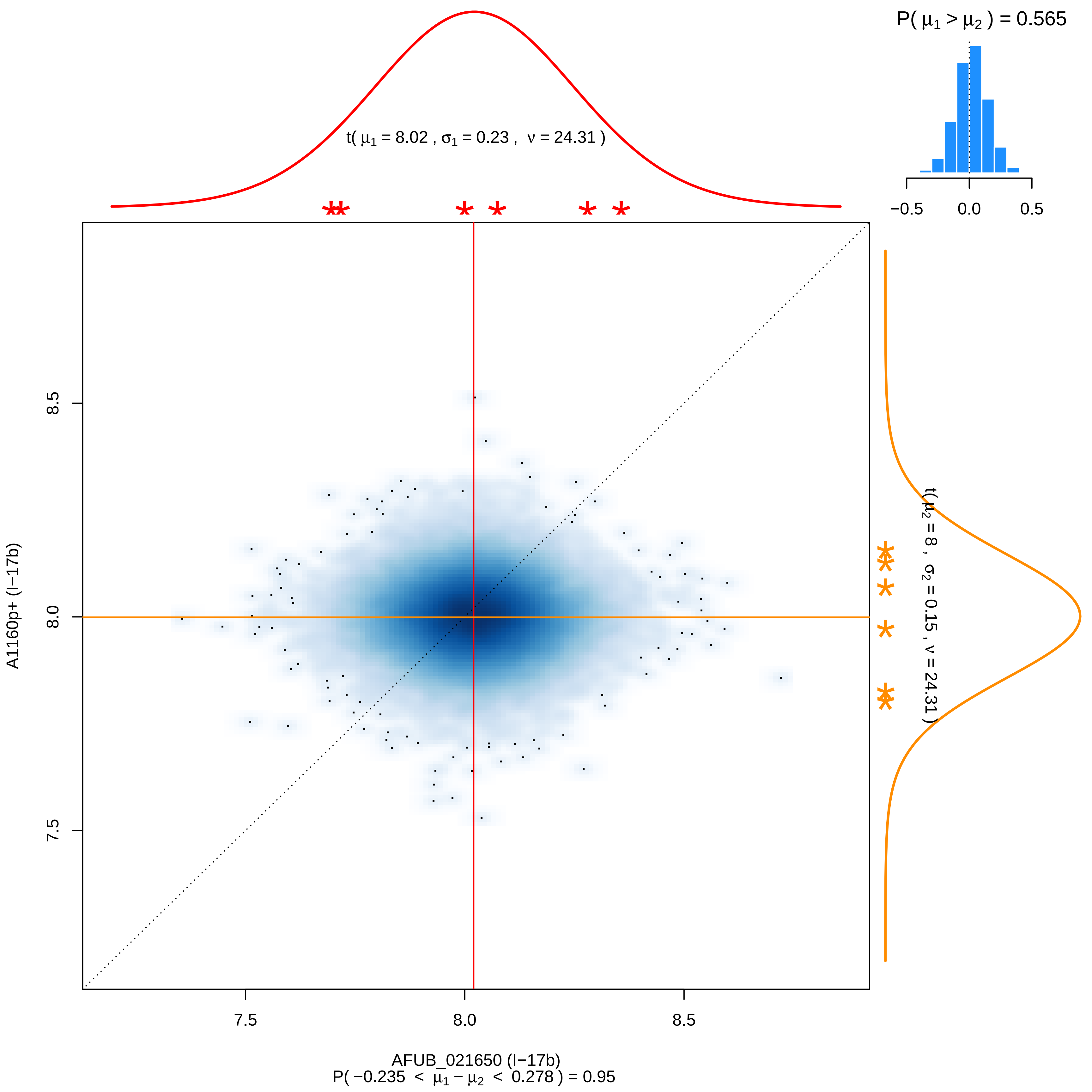

P( μ1 > μ2 ) = 0.565
−0.5
0.0
0.5
t( μ1 = 8.02 , σ1 = 0.23 , ν = 24.31 )
t( μ2 = 8 , σ2 = 0.15 , ν = 24.31 )
8.5
8.0
7.5
A1160p+ (I−17b)
7.5
8.0
8.5
AFUB_021650 (I−17b)
P( −0.235 < μ1 − μ2 < 0.278 ) = 0.95

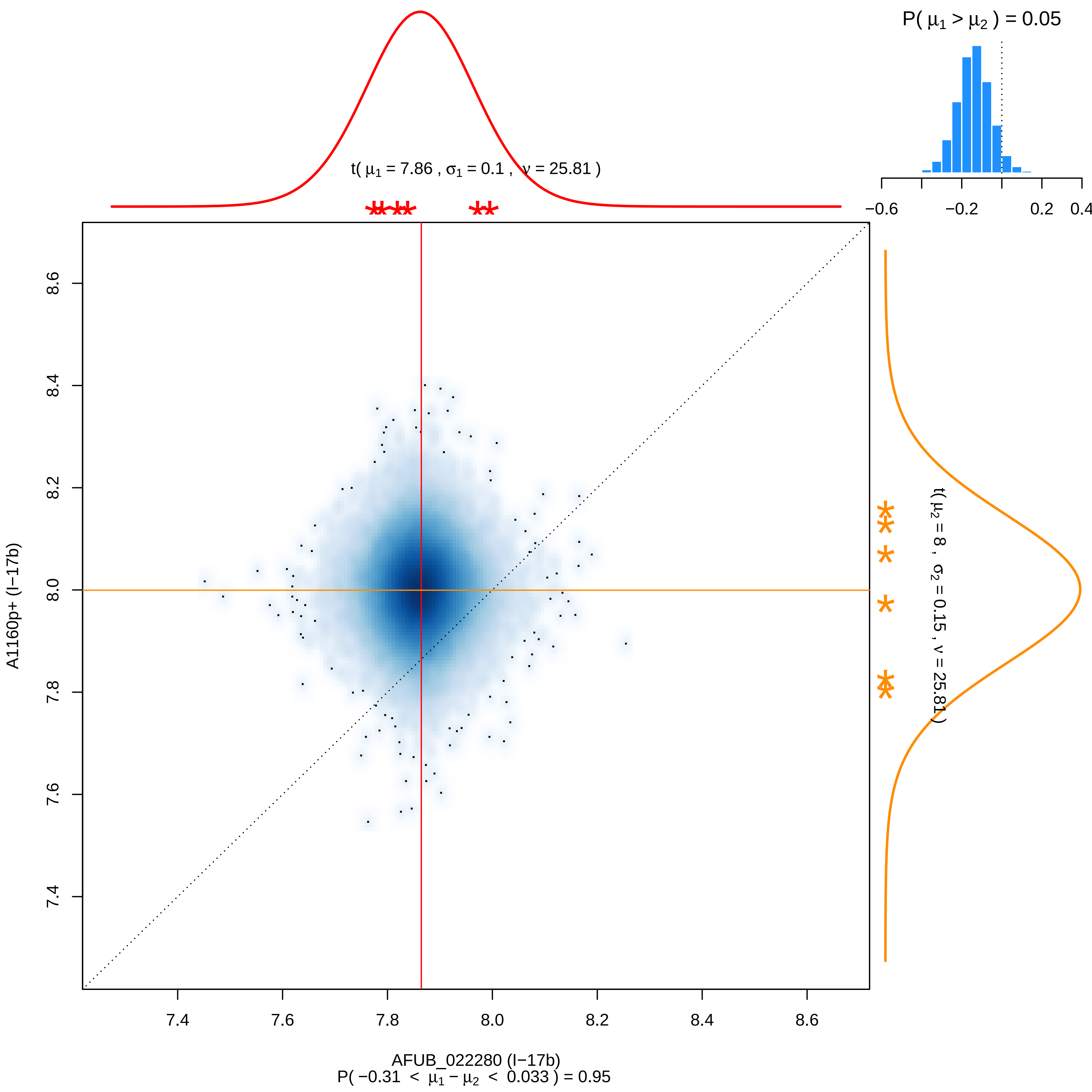

P( μ1 > μ2 ) = 0.05
−0.6
−0.2
0.2
0.4
t( μ1 = 7.86 , σ1 = 0.1 , ν = 25.81 )
t( μ2 = 8 , σ2 = 0.15 , ν = 25.81 )
A1160p+ (I−17b)
8.6
8.4
8.2
8.0
7.8
7.6
7.4
7.4
7.6
7.8
8.0
8.2
8.4
8.6
AFUB_022280 (I−17b)
P( −0.31 < μ1 − μ2 < 0.033 ) = 0.95

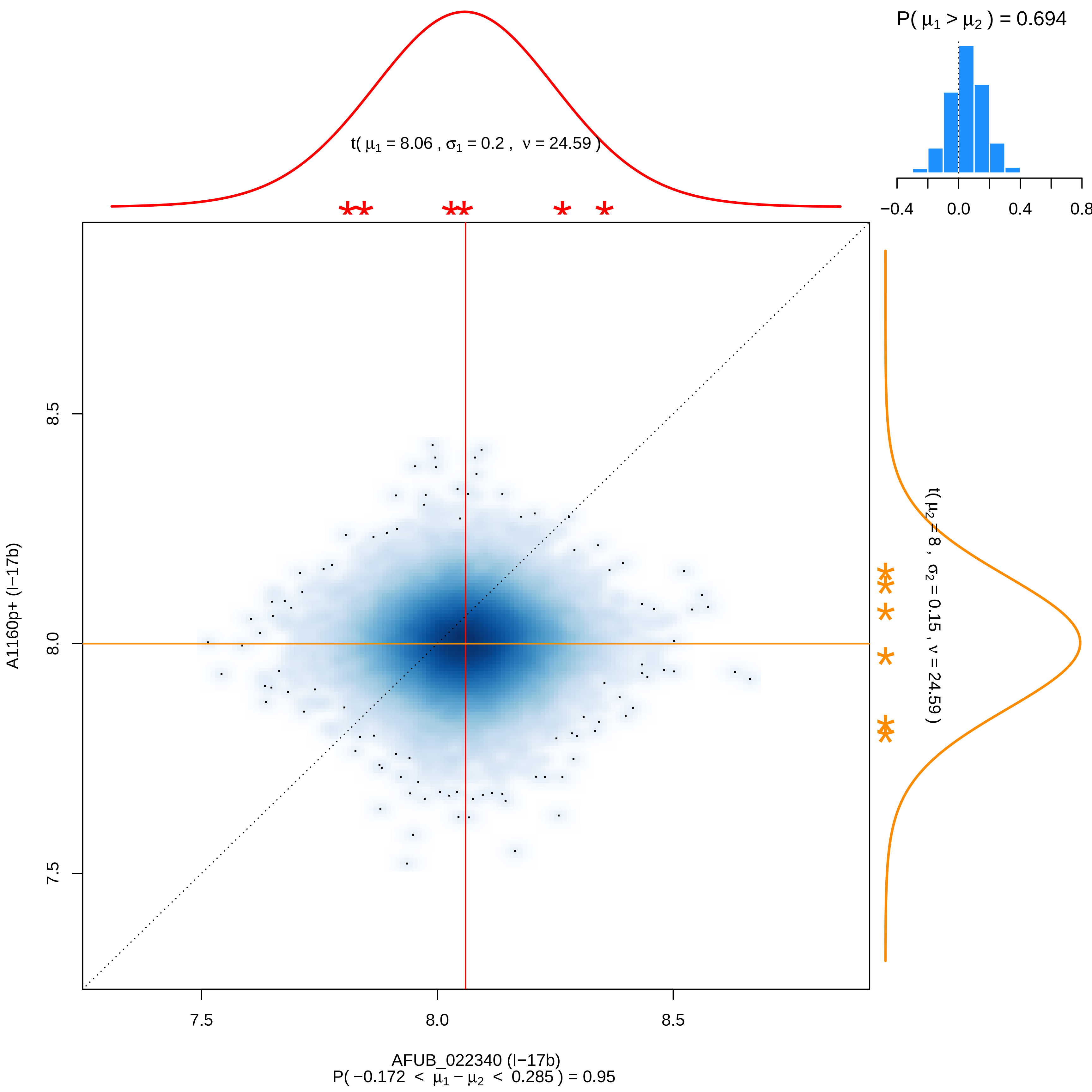

P( μ1 > μ2 ) = 0.694
−0.4
0.0
0.4
0.8
t( μ1 = 8.06 , σ1 = 0.2 , ν = 24.59 )
t( μ2 = 8 , σ2 = 0.15 , ν = 24.59 )
A1160p+ (I−17b)
8.5
8.0
7.5
7.5
8.0
8.5
AFUB_022340 (I−17b)
P( −0.172 < μ1 − μ2 < 0.285 ) = 0.95

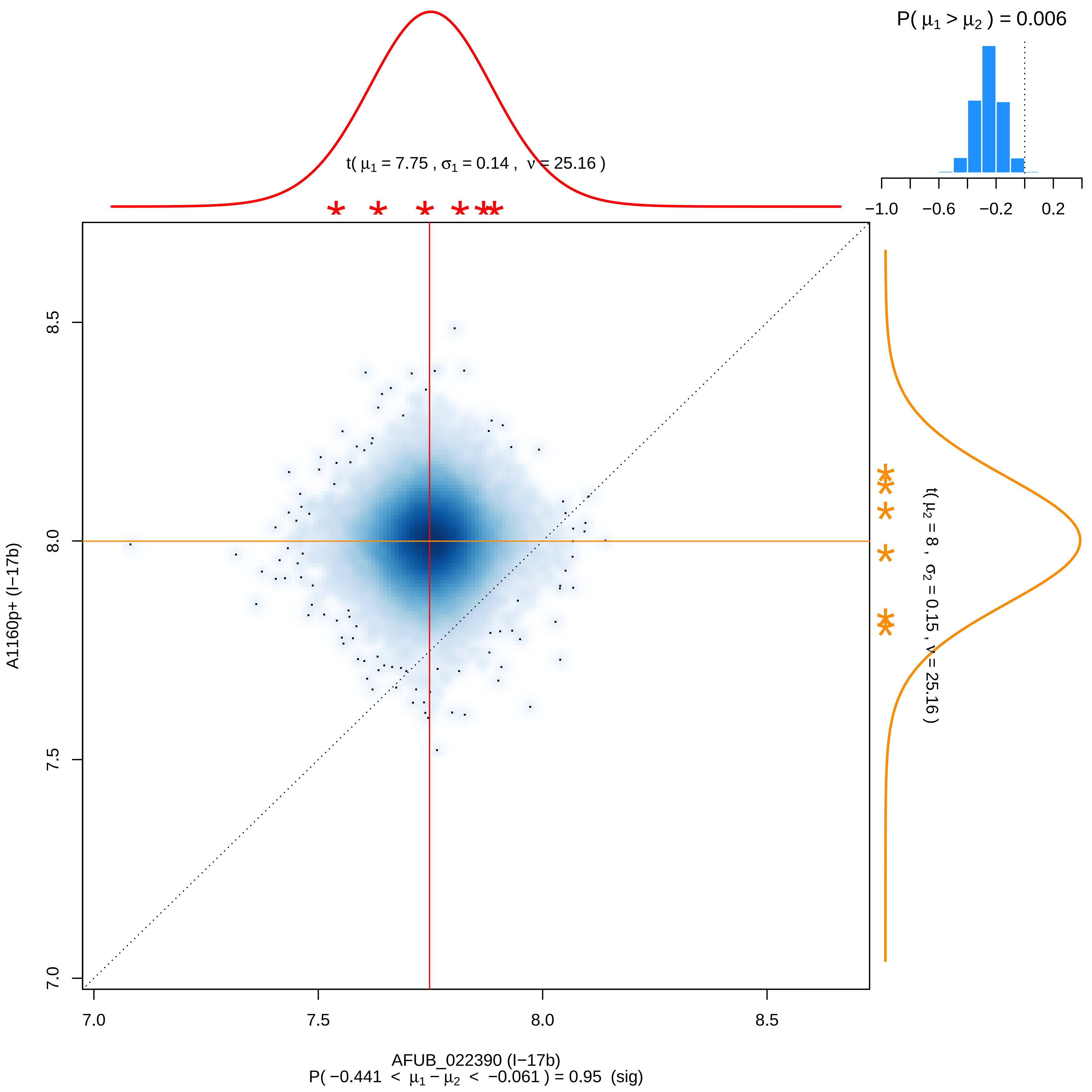

P( μ1 > μ2 ) = 0.006
−1.0
−0.6
−0.2
0.2
t( μ1 = 7.75 , σ1 = 0.14 , ν = 25.16 )
8.5
8.0
7.5
7.0
A1160p+ (I−17b)
t( μ2 = 8 , σ2 = 0.15 , ν = 25.16 )
7.0
7.5
8.0
8.5
AFUB_022390 (I−17b)
P( −0.441 < μ1 − μ2 < −0.061 ) = 0.95 (sig)

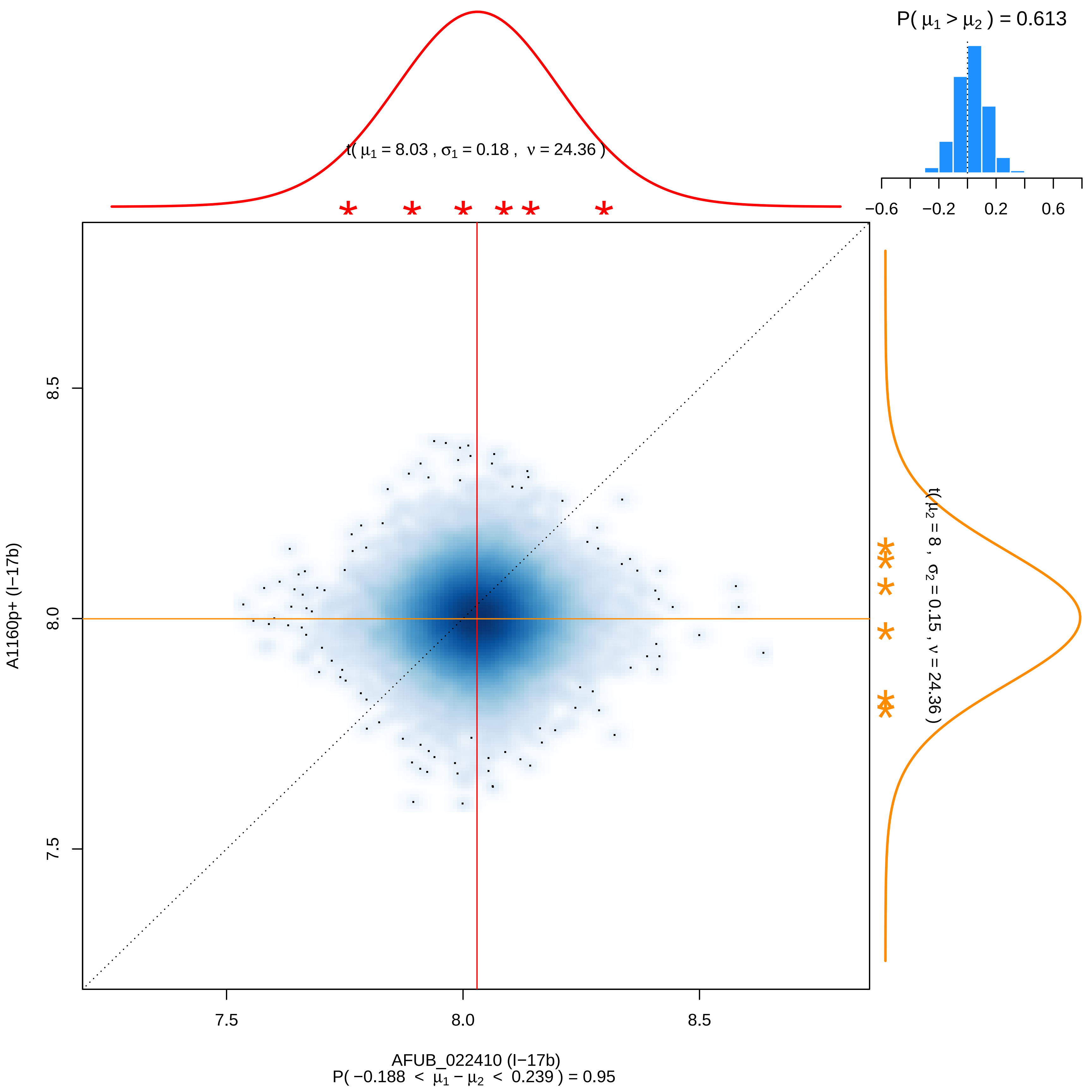
P( μ1 > μ2 ) = 0.613
−0.6
−0.2
0.2
0.6
t( μ1 = 8.03 , σ1 = 0.18 , ν = 24.36 )
t( μ2 = 8 , σ2 = 0.15 , ν = 24.36 )
8.5
8.0
7.5
A1160p+ (I−17b)
AFUB_022410 (I−17b)
P( −0.188 < μ1 − μ2 < 0.239 ) = 0.95

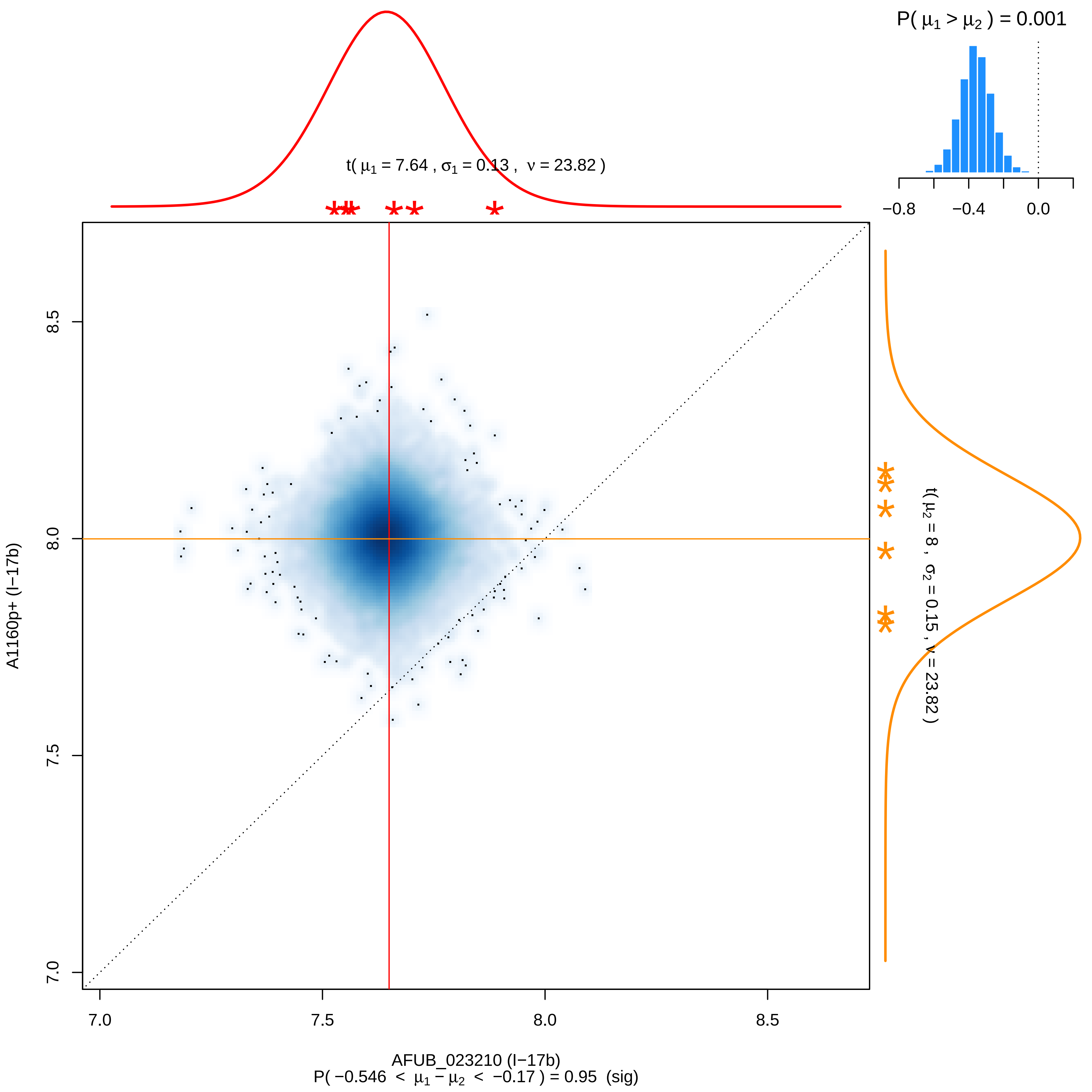

P( μ1 > μ2 ) = 0.001
−0.8
−0.4
0.0
t( μ1 = 7.64 , σ1 = 0.13 , ν = 23.82 )
t( μ2 = 8 , σ2 = 0.15 , ν = 23.82 )
8.5
8.0
7.5
7.0
A1160p+ (I−17b)
7.0
7.5
8.0
8.5
AFUB_023210 (I−17b)
P( −0.546 < μ1 − μ2 < −0.17 ) = 0.95 (sig)

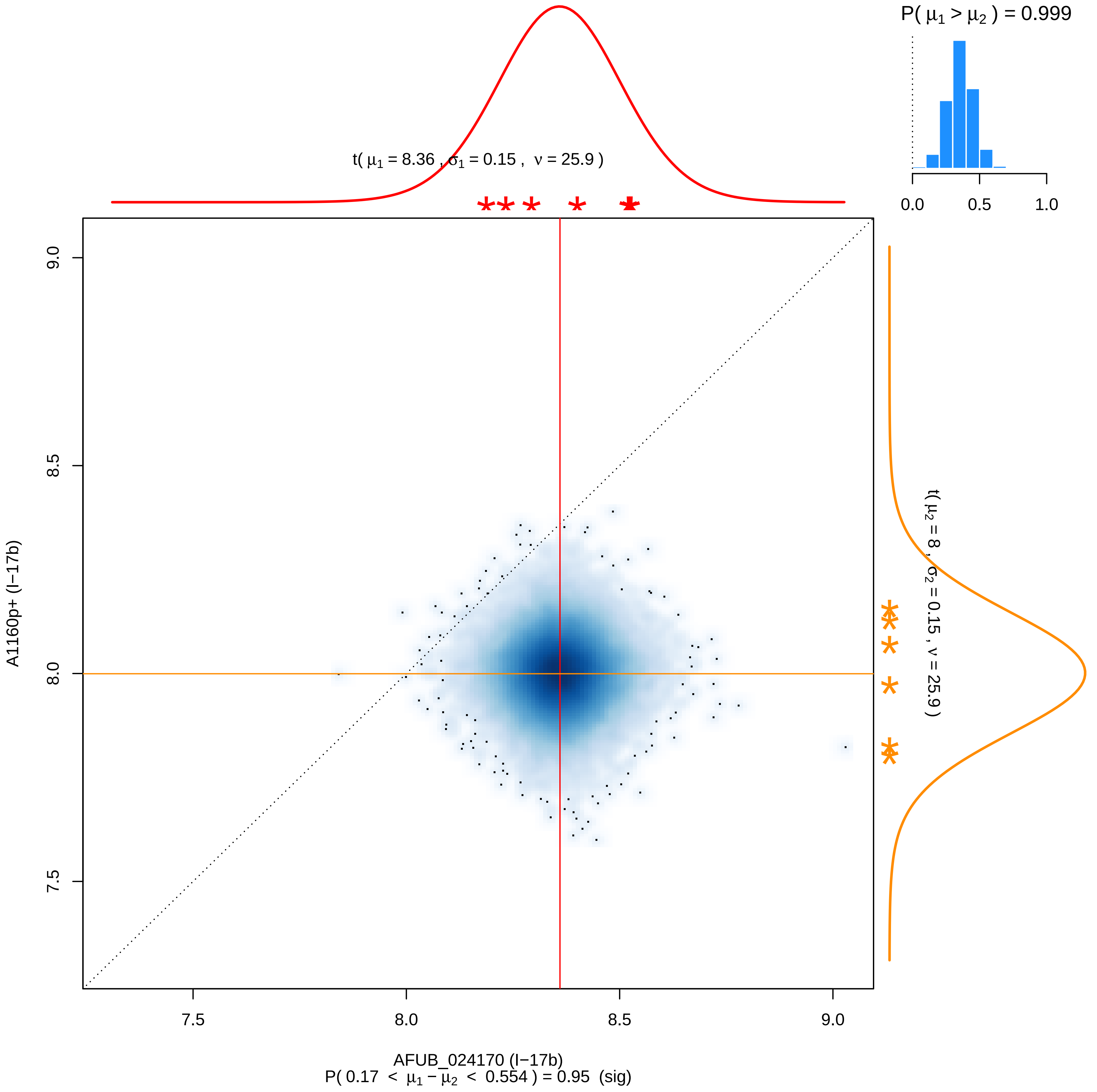
P( μ1 > μ2 ) = 0.999
0.0
0.5
1.0
t( μ1 = 8.36 , σ1 = 0.15 , ν = 25.9 )
t( μ2 = 8 , σ2 = 0.15 , ν = 25.9 )
9.0
8.5
8.0
7.5
A1160p+ (I−17b)
7.5
8.0
8.5
9.0
AFUB_024170 (I−17b)
P( 0.17 < μ1 − μ2 < 0.554 ) = 0.95 (sig)

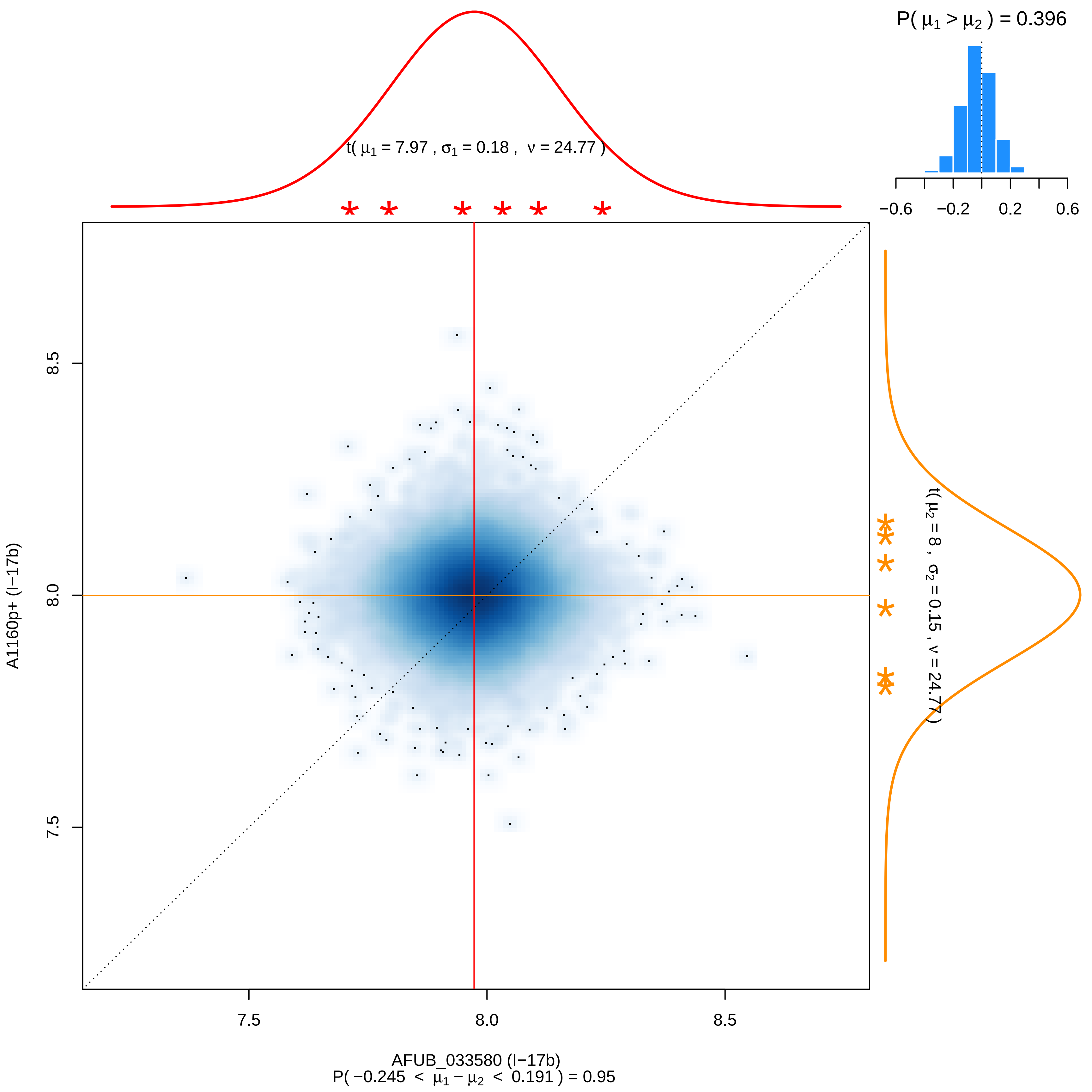

P( μ1 > μ2 ) = 0.396
−0.6 −0.2 0.2 0.6
t( μ1 = 7.97 , σ1 = 0.18 , ν = 24.77 )
t( μ2 = 8 , σ2 = 0.15 , ν = 24.77 )
8.5
8.0
7.5
A1160p+ (I−17b)
7.5 8.0 8.5
AFUB_033580 (I−17b)
P( −0.245 < μ1 − μ2 < 0.191 ) = 0.95

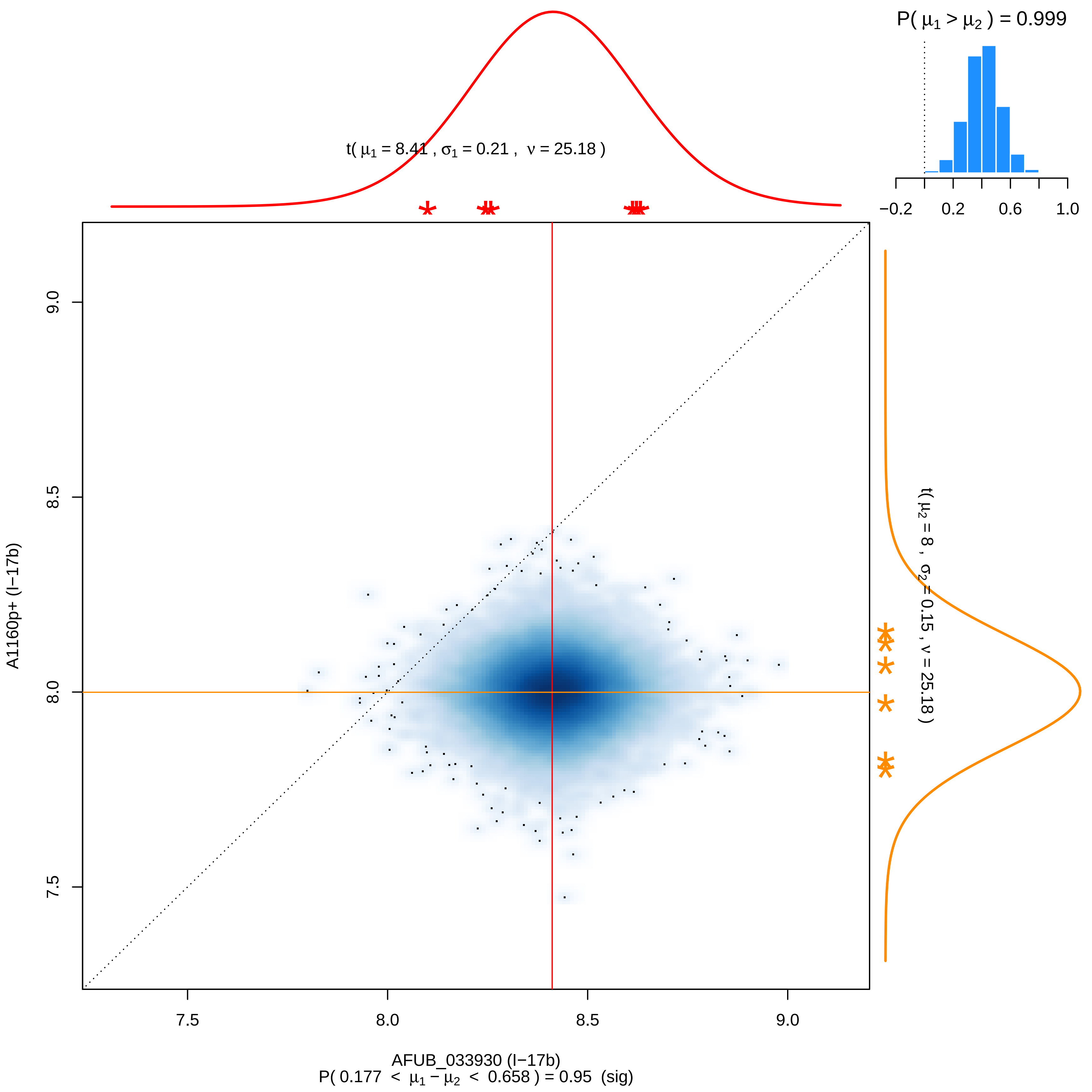

P( μ1 > μ2 ) = 0.999
t( μ1 = 8.41 , σ1 = 0.21 , ν = 25.18 )
−0.2
0.2
0.6
1.0
t( μ2 = 8 , σ2 = 0.15 , ν = 25.18 )
A1160p+ (I−17b)
9.0
8.5
8.0
7.5
7.5
8.0
8.5
9.0
AFUB_033930 (I−17b)
P( 0.177 < μ1 − μ2 < 0.658 ) = 0.95 (sig)

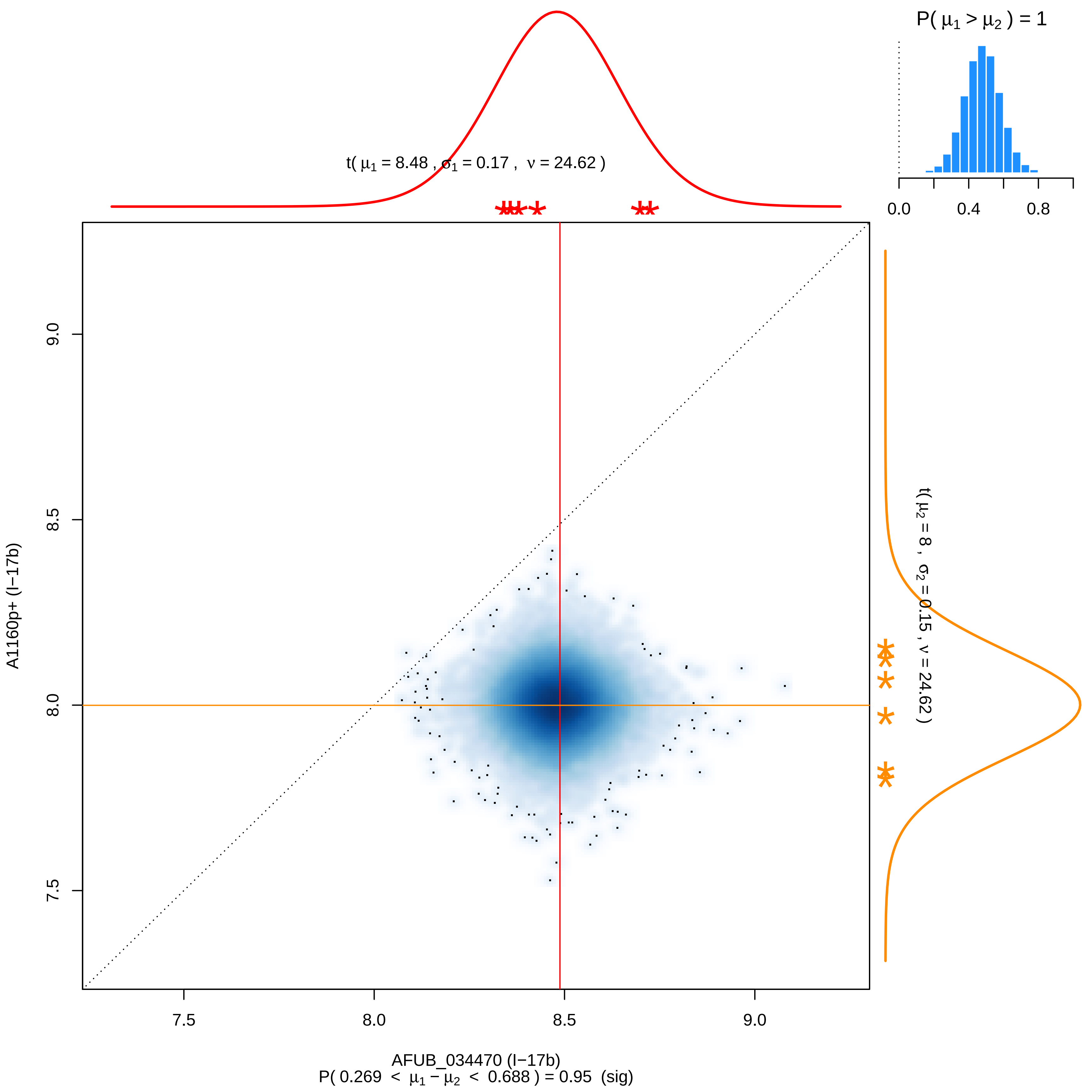

P( μ1 > μ2 ) = 1
0.0
0.4
0.8
t( μ1 = 8.48 , σ1 = 0.17 , ν = 24.62 )
t( μ2 = 8 , σ2 = 0.15 , ν = 24.62 )
A1160p+ (I−17b)
9.0
8.5
8.0
7.5
7.5
8.0
8.5
9.0
AFUB_034470 (I−17b)
P( 0.269 < μ1 − μ2 < 0.688 ) = 0.95 (sig)

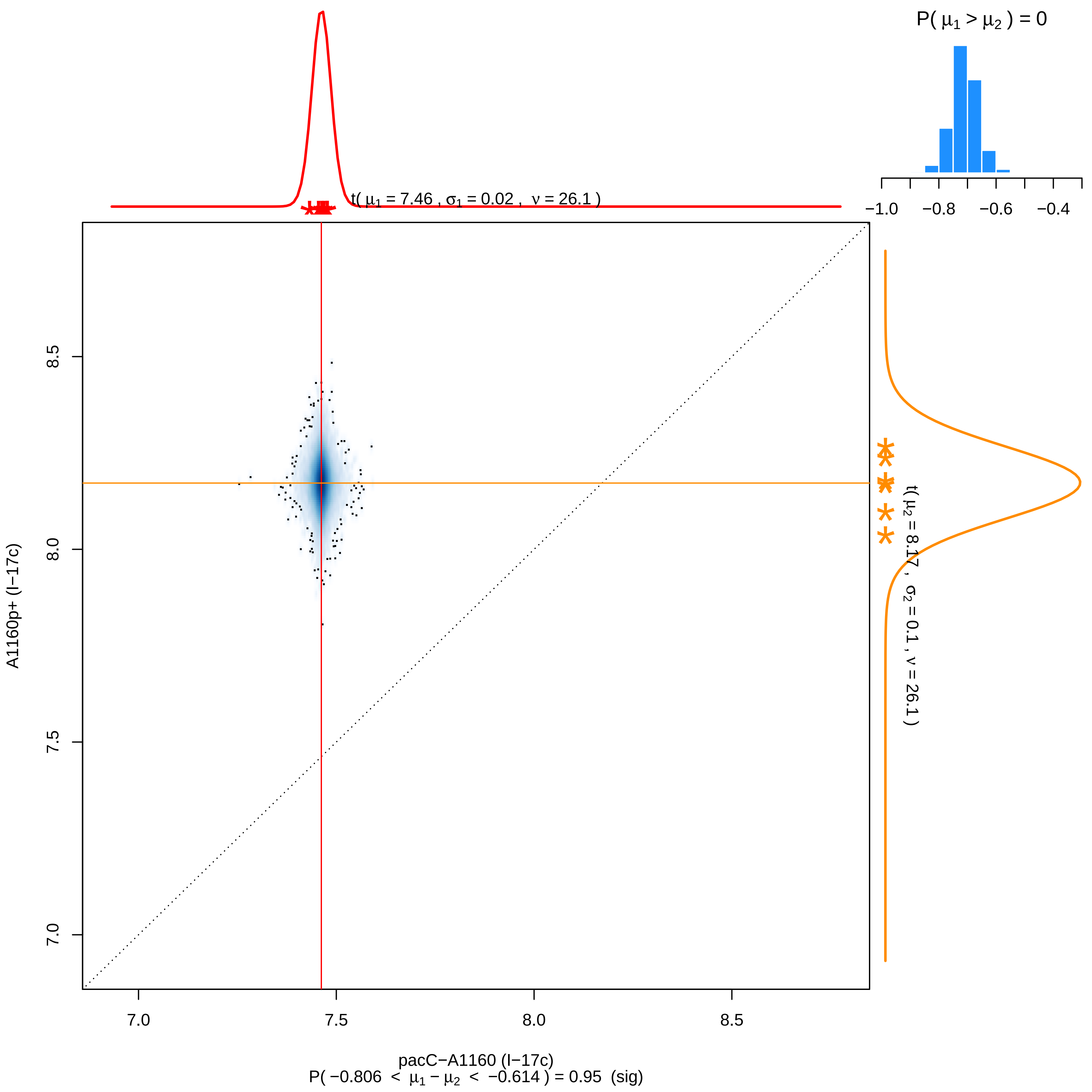

P( μ1 > μ2 ) = 0
−1.0
−0.8
−0.6
−0.4
t( μ1 = 7.46 , σ1 = 0.02 , ν = 26.1 )
t( μ2 = 8.17 , σ2 = 0.1 , ν = 26.1 )
A1160p+ (I−17c)
8.5
8.0
7.5
7.0
7.0
7.5
8.0
8.5
pacC−A1160 (I−17c)
P( −0.806 < μ1 − μ2 < −0.614 ) = 0.95 (sig)

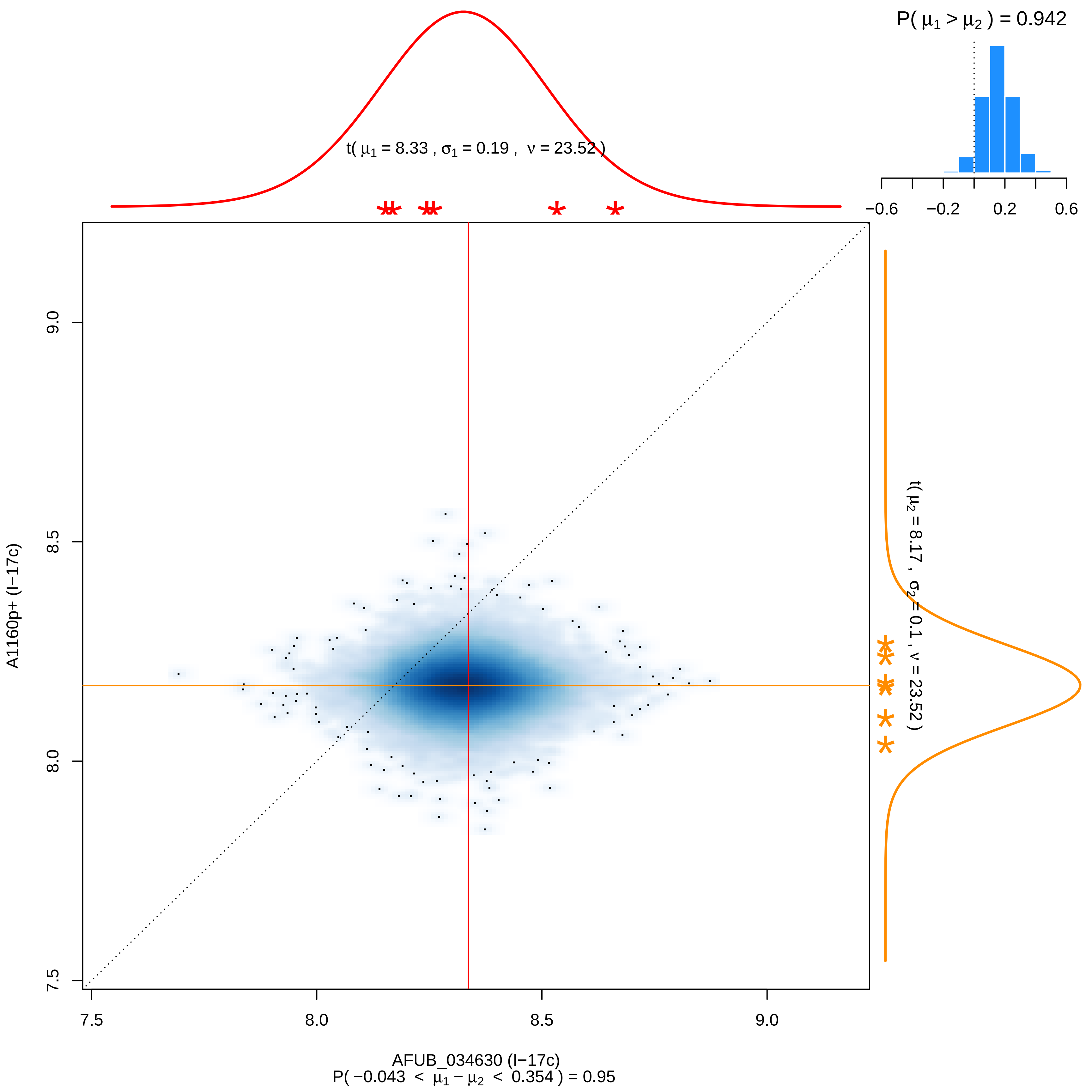

P( μ1 > μ2 ) = 0.942
−0.6
−0.2
0.2
0.6
t( μ1 = 8.33 , σ1 = 0.19 , ν = 23.52 )
t( μ2 = 8.17 , σ2 = 0.1 , ν = 23.52 )
A1160p+ (I−17c)
9.0
8.5
8.0
7.5
7.5
8.0
8.5
9.0
AFUB_034630 (I−17c)
P( −0.043 < μ1 − μ2 < 0.354 ) = 0.95

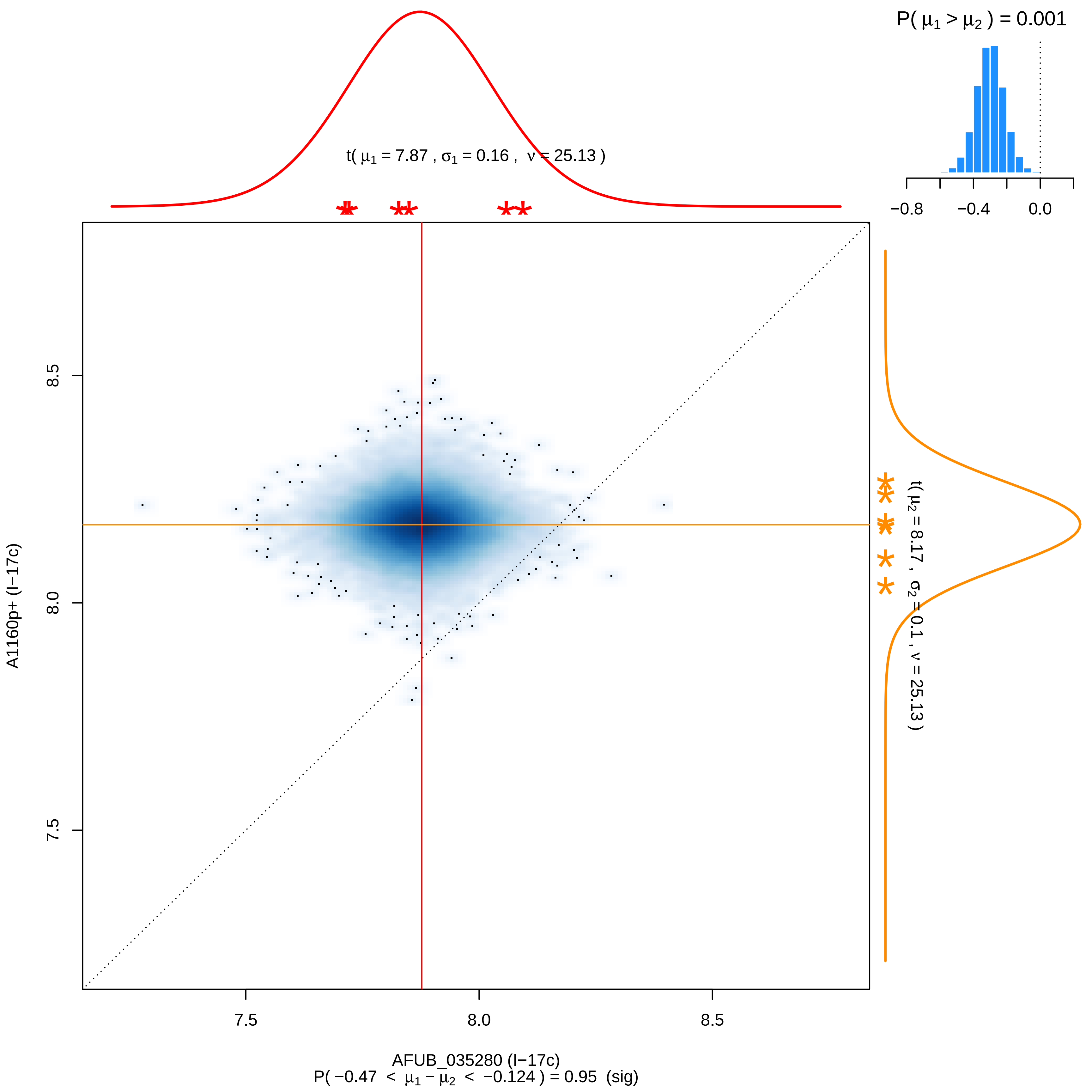

P( μ1 > μ2 ) = 0.001
−0.8
−0.4
0.0
t( μ1 = 7.87 , σ1 = 0.16 , ν = 25.13 )
t( μ2 = 8.17 , σ2 = 0.1 , ν = 25.13 )
8.5
8.0
7.5
A1160p+ (I−17c)
7.5
8.0
8.5
AFUB_035280 (I−17c)
P( −0.47 < μ1 − μ2 < −0.124 ) = 0.95 (sig)

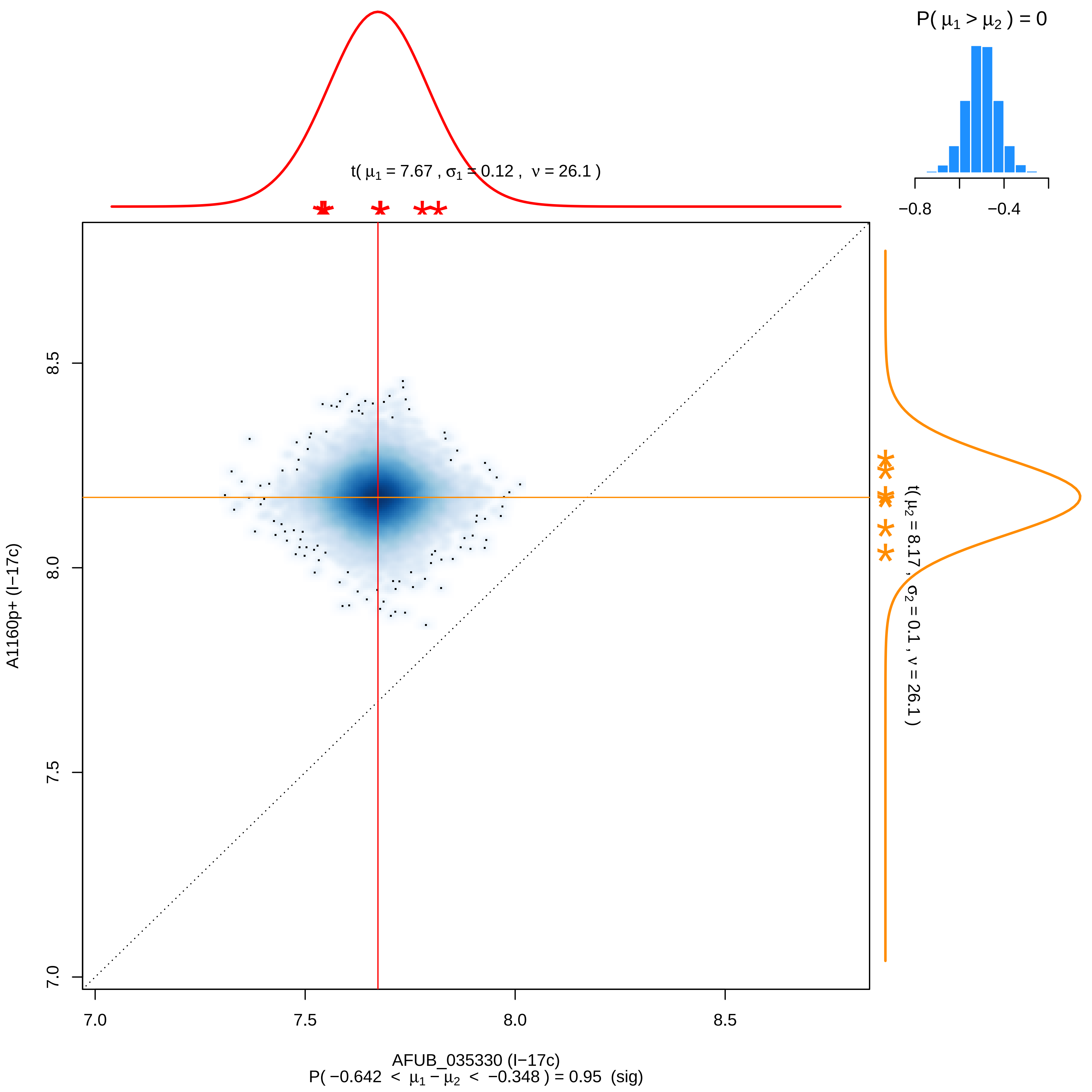

P( μ1 > μ2 ) = 0
−0.8
−0.4
t( μ1 = 7.67 , σ1 = 0.12 , ν = 26.1 )
t( μ2 = 8.17 , σ2 = 0.1 , ν = 26.1 )
8.5
8.0
7.5
7.0
A1160p+ (I−17c)
7.0
7.5
8.0
8.5
AFUB_035330 (I−17c)
P( −0.642 < μ1 − μ2 < −0.348 ) = 0.95 (sig)

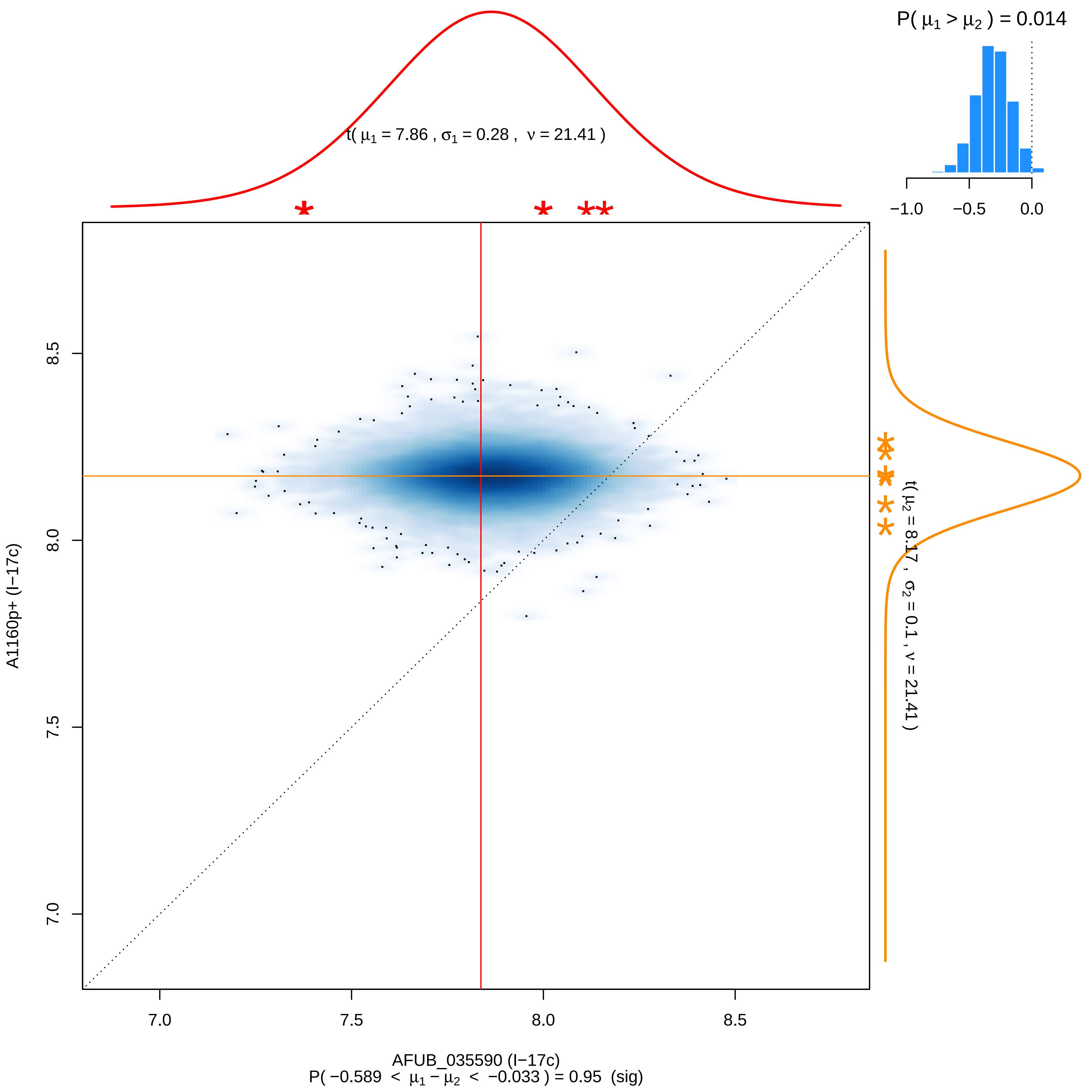

P( μ1 > μ2 ) = 0.014
−1.0
−0.5
0.0
t( μ1 = 7.86 , σ1 = 0.28 , ν = 21.41 )
t( μ2 = 8.17 , σ2 = 0.1 , ν = 21.41 )
A1160p+ (I−17c)
8.5
8.0
7.5
7.0
7.0
7.5
8.0
8.5
AFUB_035590 (I−17c)
P( −0.589 < μ1 − μ2 < −0.033 ) = 0.95 (sig)

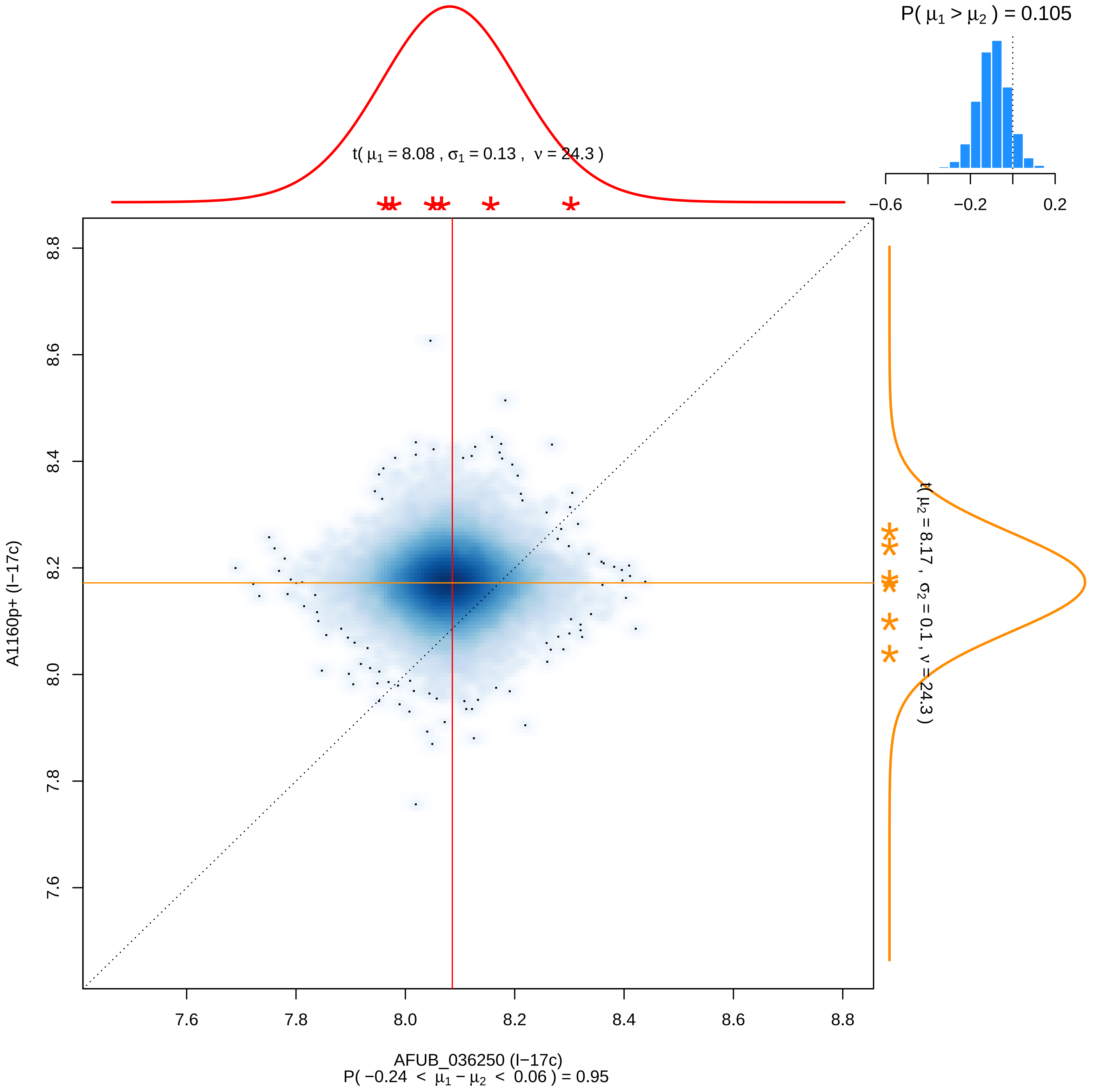

P( μ1 > μ2 ) = 0.105
−0.6
−0.2
0.2
t( μ1 = 8.08 , σ1 = 0.13 , ν = 24.3 )
t( μ2 = 8.17 , σ2 = 0.1 , ν = 24.3 )
8.8
8.6
8.4
8.2
8.0
7.8
7.6
A1160p+ (I−17c)
7.6
7.8
8.0
8.2
8.4
8.6
8.8
AFUB_036250 (I−17c)
P( −0.24 < μ1 − μ2 < 0.06 ) = 0.95

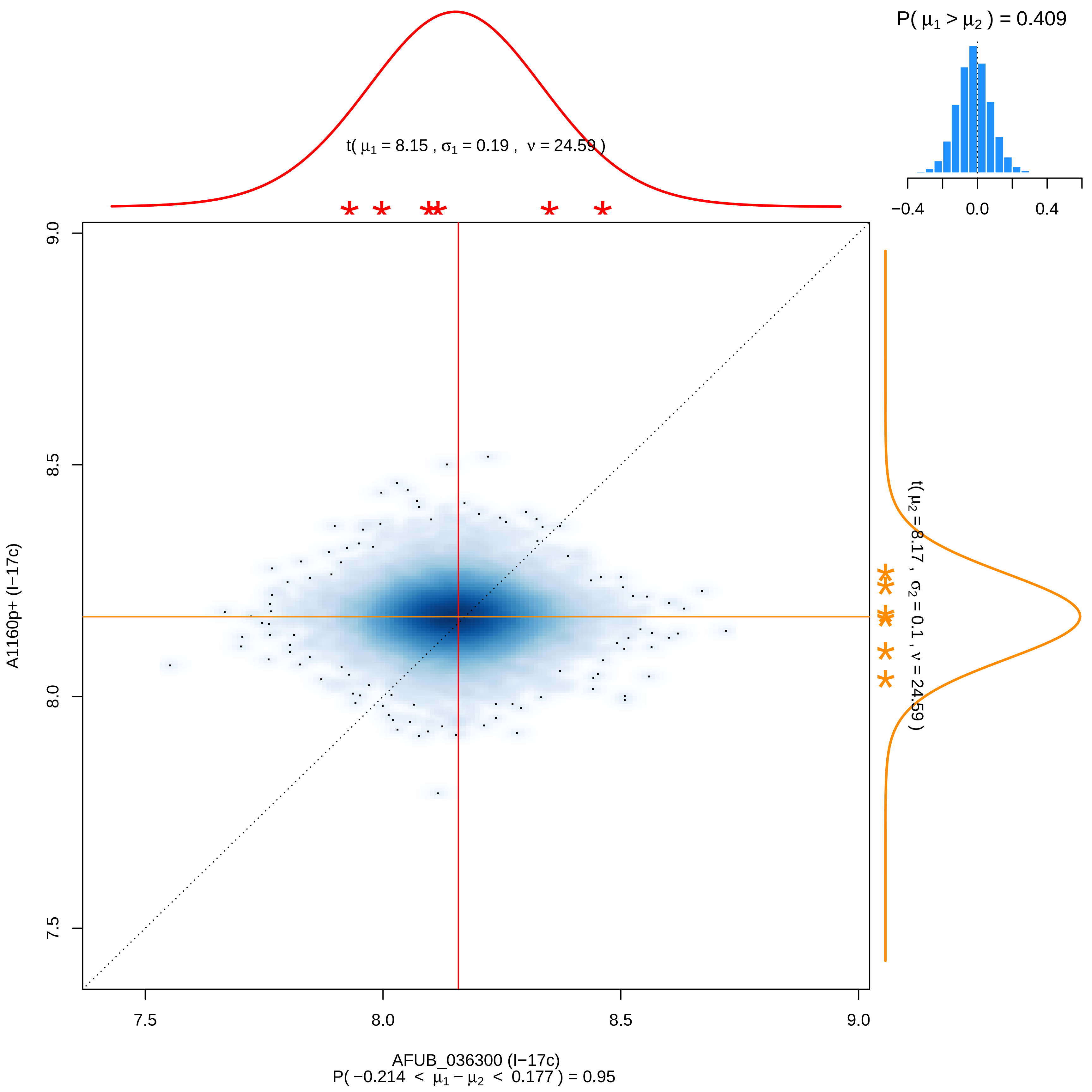

P( μ1 > μ2 ) = 0.409
−0.4
0.0
0.4
t( μ1 = 8.15 , σ1 = 0.19 , ν = 24.59 )
t( μ2 = 8.17 , σ2 = 0.1 , ν = 24.59 )
9.0
8.5
8.0
7.5
A1160p+ (I−17c)
7.5
8.0
8.5
9.0
AFUB_036300 (I−17c)
P( −0.214 < μ1 − μ2 < 0.177 ) = 0.95

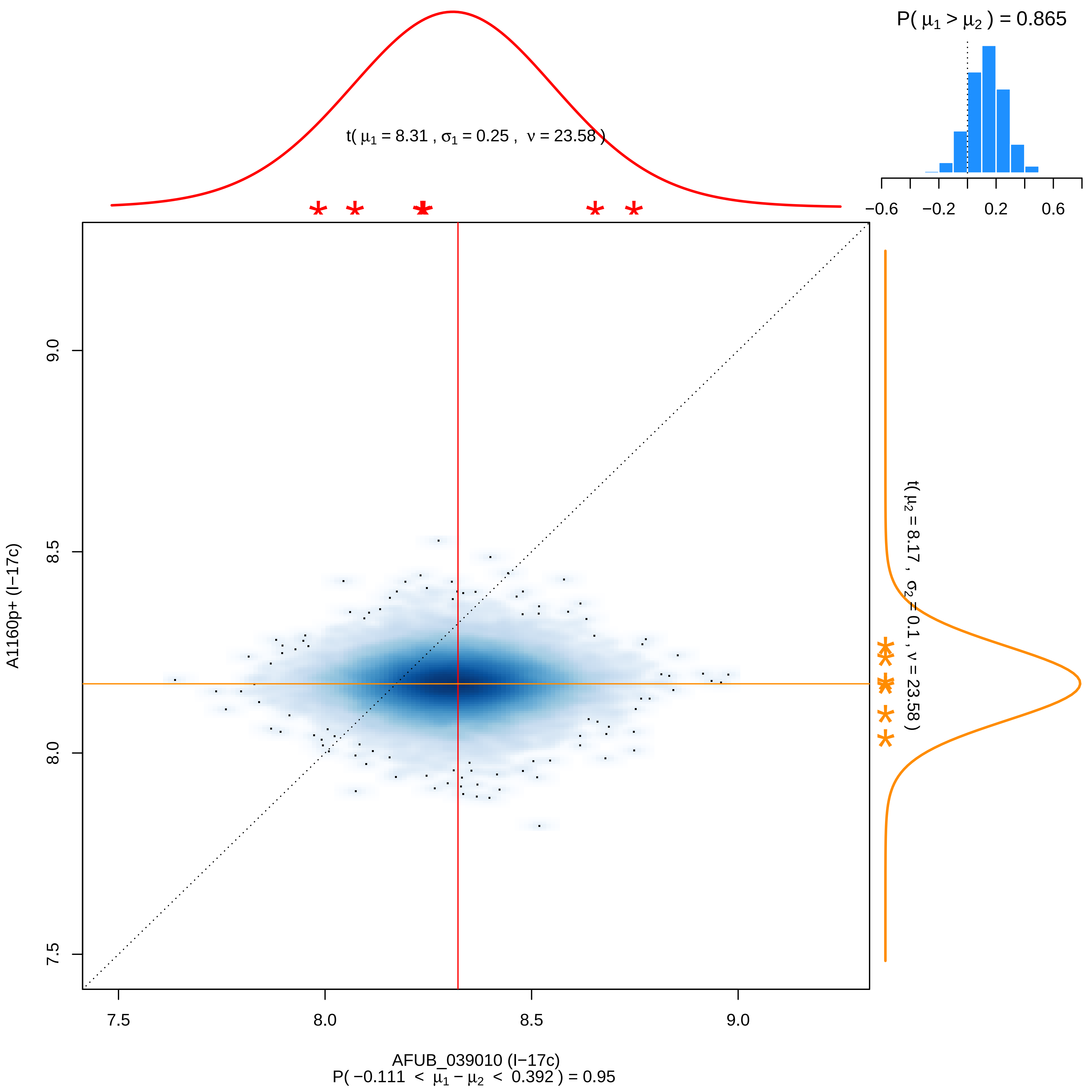

P( μ1 > μ2 ) = 0.865
−0.6
−0.2
0.2
0.6
t( μ1 = 8.31 , σ1 = 0.25 , ν = 23.58 )
t( μ2 = 8.17 , σ2 = 0.1 , ν = 23.58 )
A1160p+ (I−17c)
9.0
8.5
8.0
7.5
7.5
8.0
8.5
9.0
AFUB_039010 (I−17c)
P( −0.111 < μ1 − μ2 < 0.392 ) = 0.95

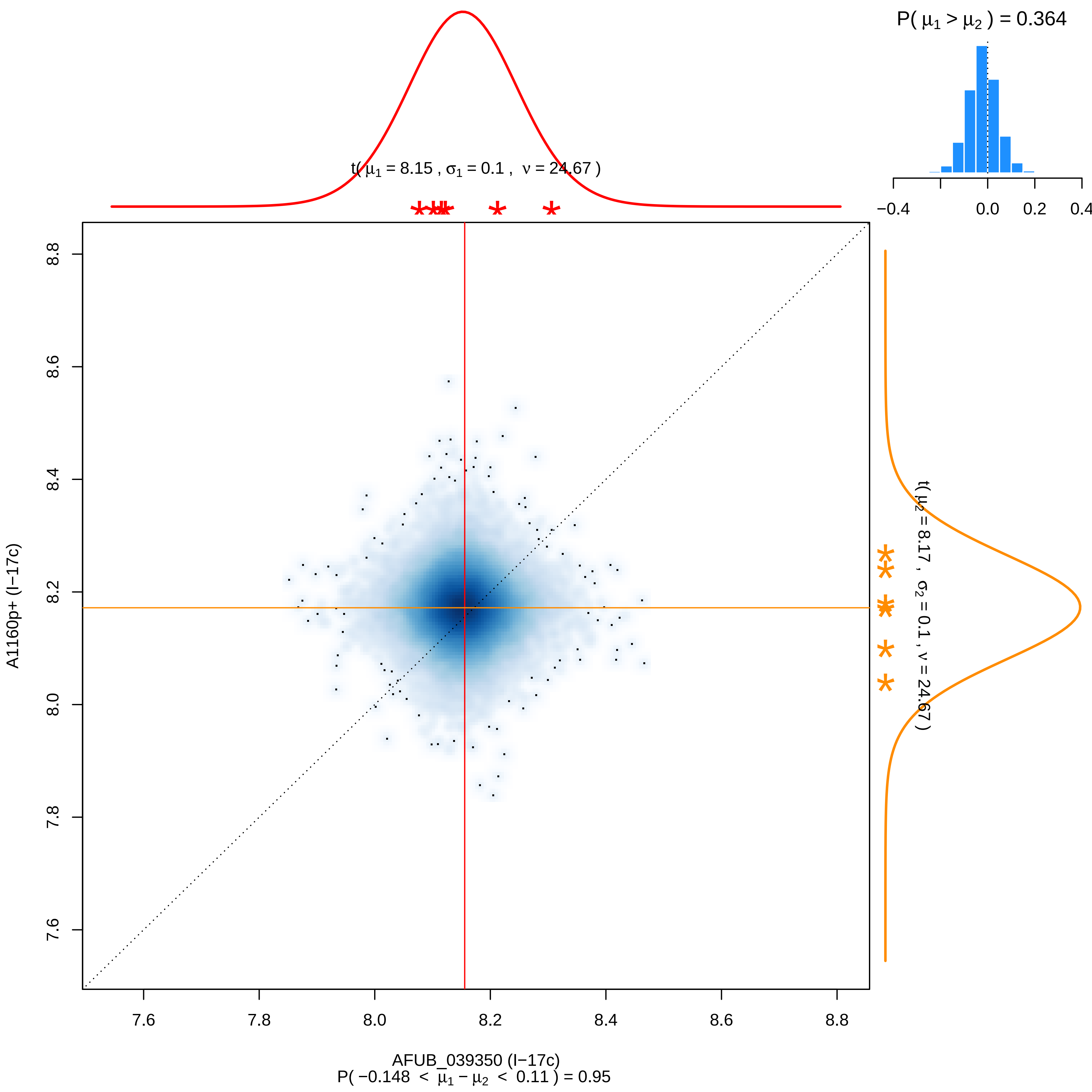

P( μ1 > μ2 ) = 0.364
−0.4
0.0
0.2
0.4
t( μ1 = 8.15 , σ1 = 0.1 , ν = 24.67 )
t( μ2 = 8.17 , σ2 = 0.1 , ν = 24.67 )
8.8
8.6
8.4
8.2
8.0
7.8
7.6
A1160p+ (I−17c)
7.6
7.8
8.0
8.2
8.4
8.6
8.8
AFUB_039350 (I−17c)
P( −0.148 < μ1 − μ2 < 0.11 ) = 0.95

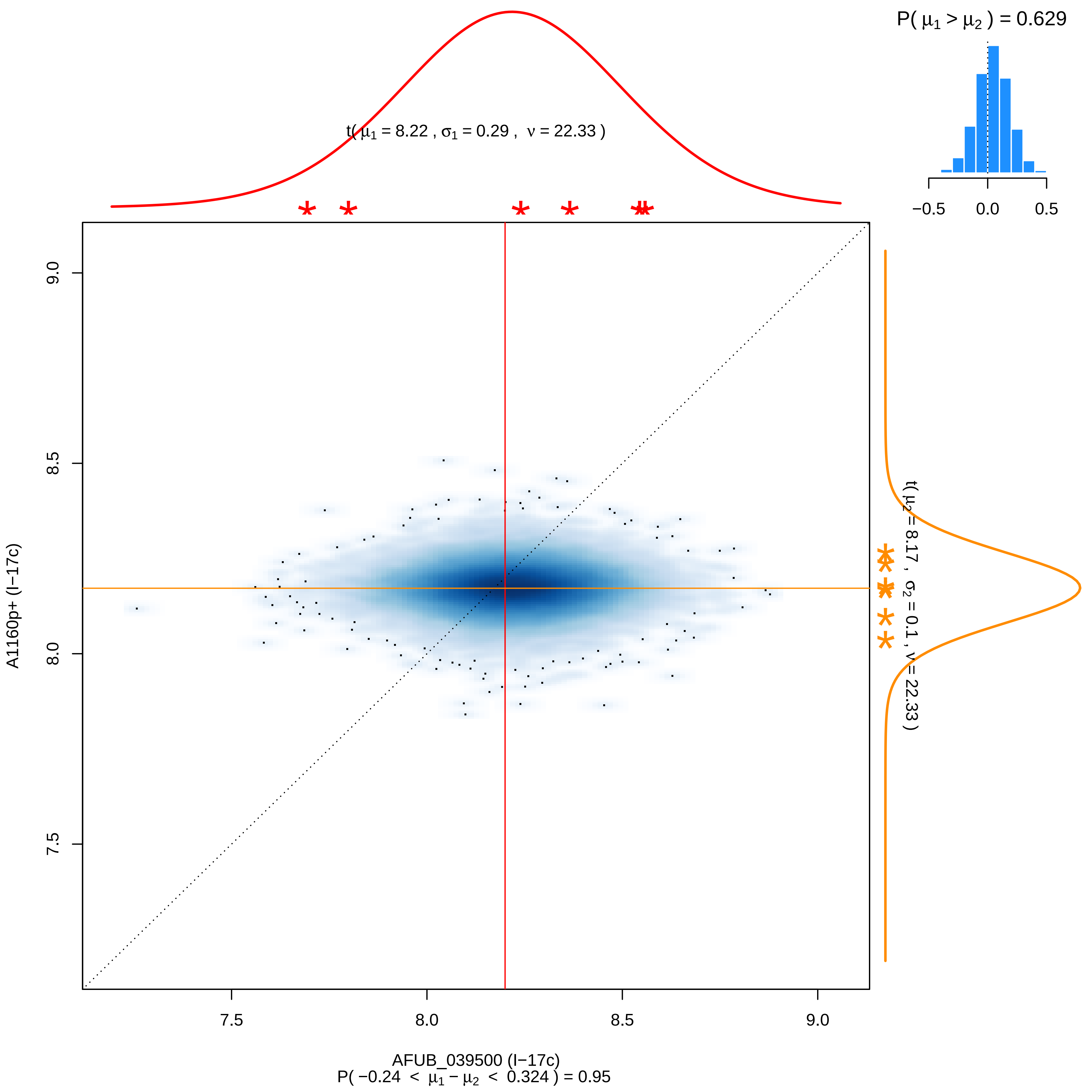
P( μ1 > μ2 ) = 0.629
−0.5
0.0
0.5
t( μ1 = 8.22 , σ1 = 0.29 , ν = 22.33 )
t( μ2 = 8.17 , σ2 = 0.1 , ν = 22.33 )
9.0
8.5
8.0
7.5
A1160p+ (I−17c)
7.5
8.0
8.5
9.0
AFUB_039500 (I−17c)
P( −0.24 < μ1 − μ2 < 0.324 ) = 0.95

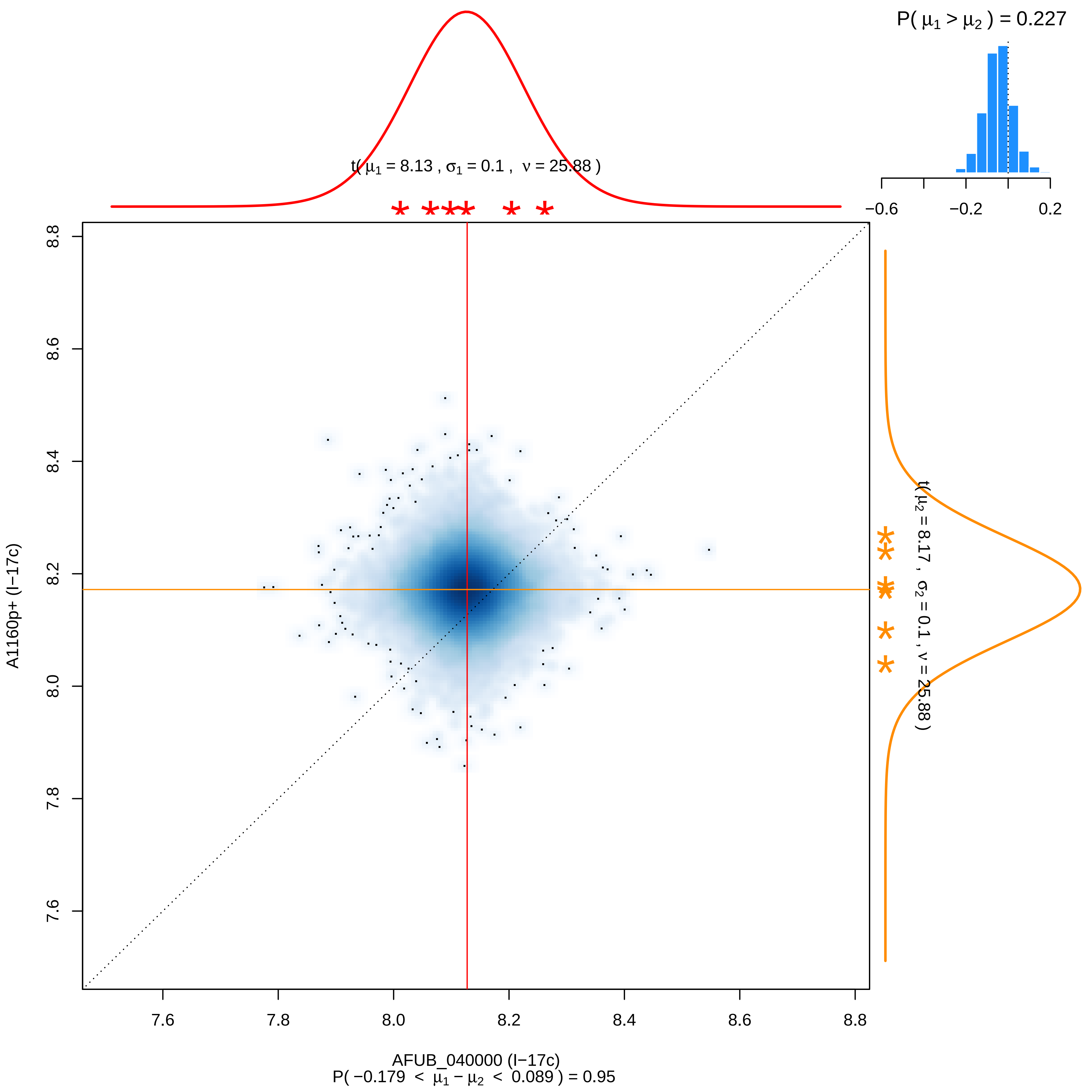

P( μ1 > μ2 ) = 0.227
−0.6
−0.2
0.2
t( μ1 = 8.13 , σ1 = 0.1 , ν = 25.88 )
t( μ2 = 8.17 , σ2 = 0.1 , ν = 25.88 )
8.8
8.6
8.4
8.2
8.0
7.8
7.6
A1160p+ (I−17c)
7.6
7.8
8.0
8.2
8.4
8.6
8.8
AFUB_040000 (I−17c)
P( −0.179 < μ1 − μ2 < 0.089 ) = 0.95

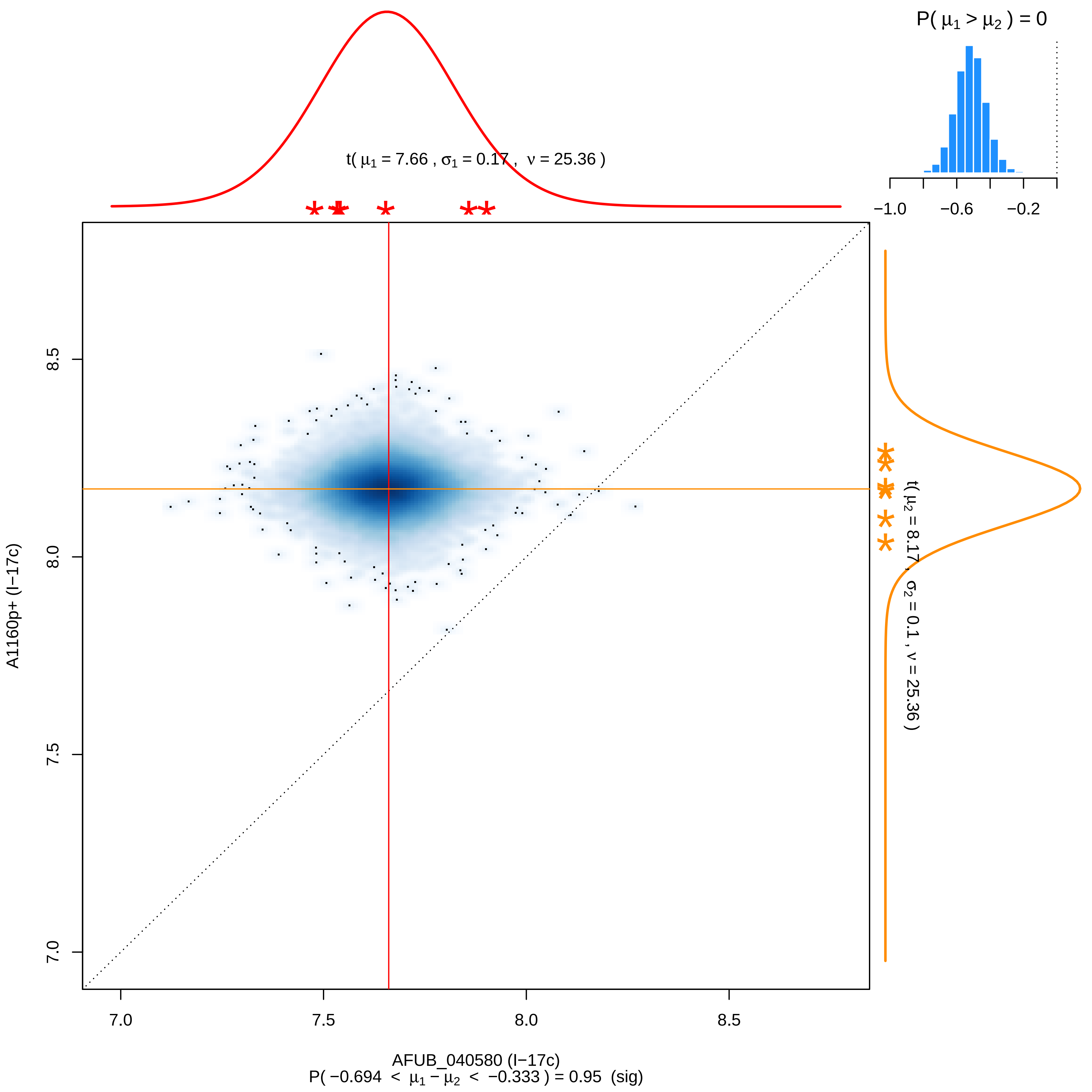

P( μ1 > μ2 ) = 0
−1.0
−0.6
−0.2
t( μ1 = 7.66 , σ1 = 0.17 , ν = 25.36 )
t( μ2 = 8.17 , σ2 = 0.1 , ν = 25.36 )
8.5
8.0
7.5
7.0
A1160p+ (I−17c)
7.0
7.5
8.0
8.5
AFUB_040580 (I−17c)
P( −0.694 < μ1 − μ2 < −0.333 ) = 0.95 (sig)

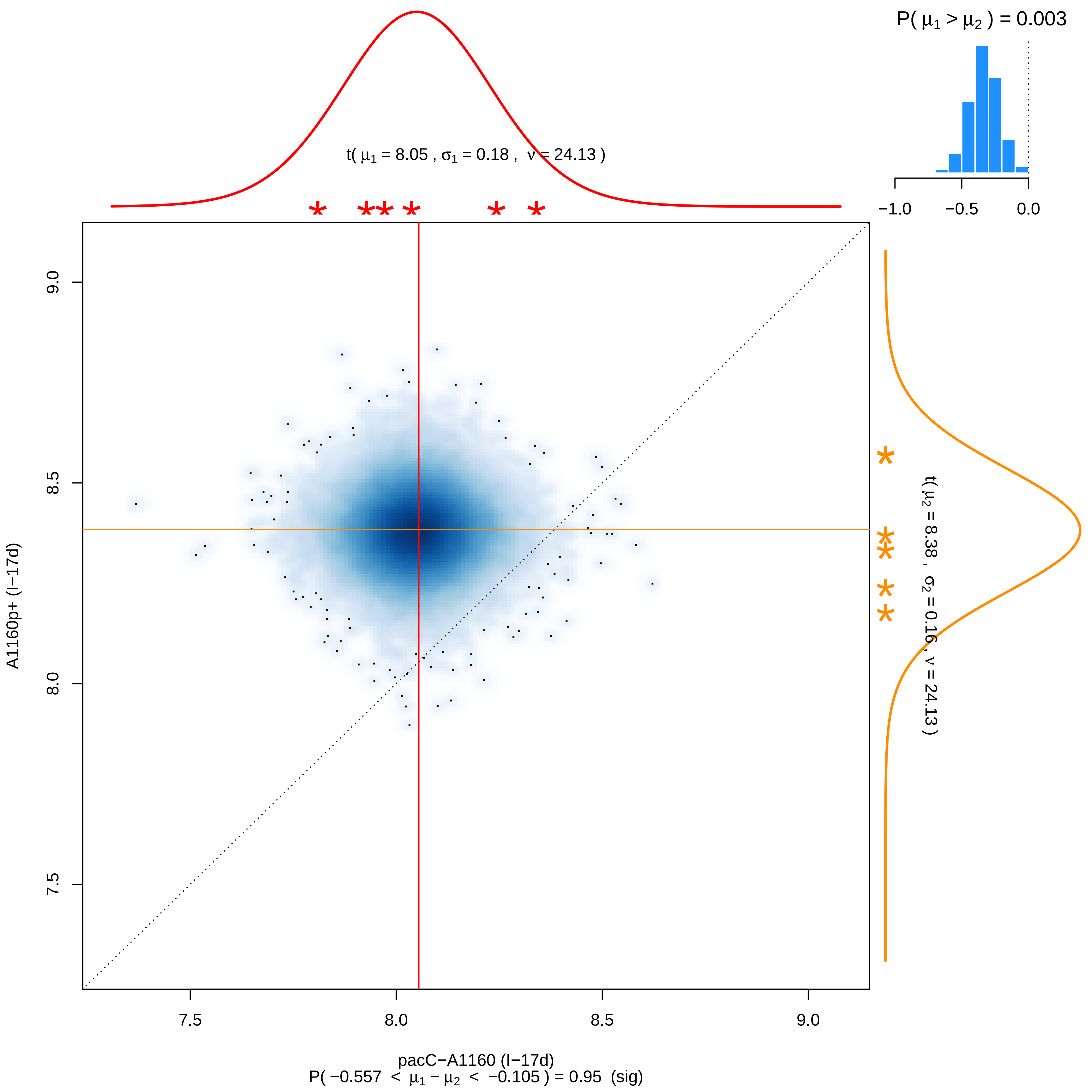

P( $\mu_1 > \mu_2$ ) = 0.003
−1.0
−0.5
0.0
t( $\mu_1$ = 8.05 , $\sigma_1$ = 0.18 , $\nu$ = 24.13 )
9.0
8.5
8.0
7.5
A1160p+ (I−17d)
t( $\mu_2$ = 8.38 , $\sigma_2$ = 0.16 , $\nu$ = 24.13 )
7.5
8.0
8.5
9.0
pacC−A1160 (I−17d)
P( −0.557 < $\mu_1 - \mu_2$ < −0.105 ) = 0.95 (sig)

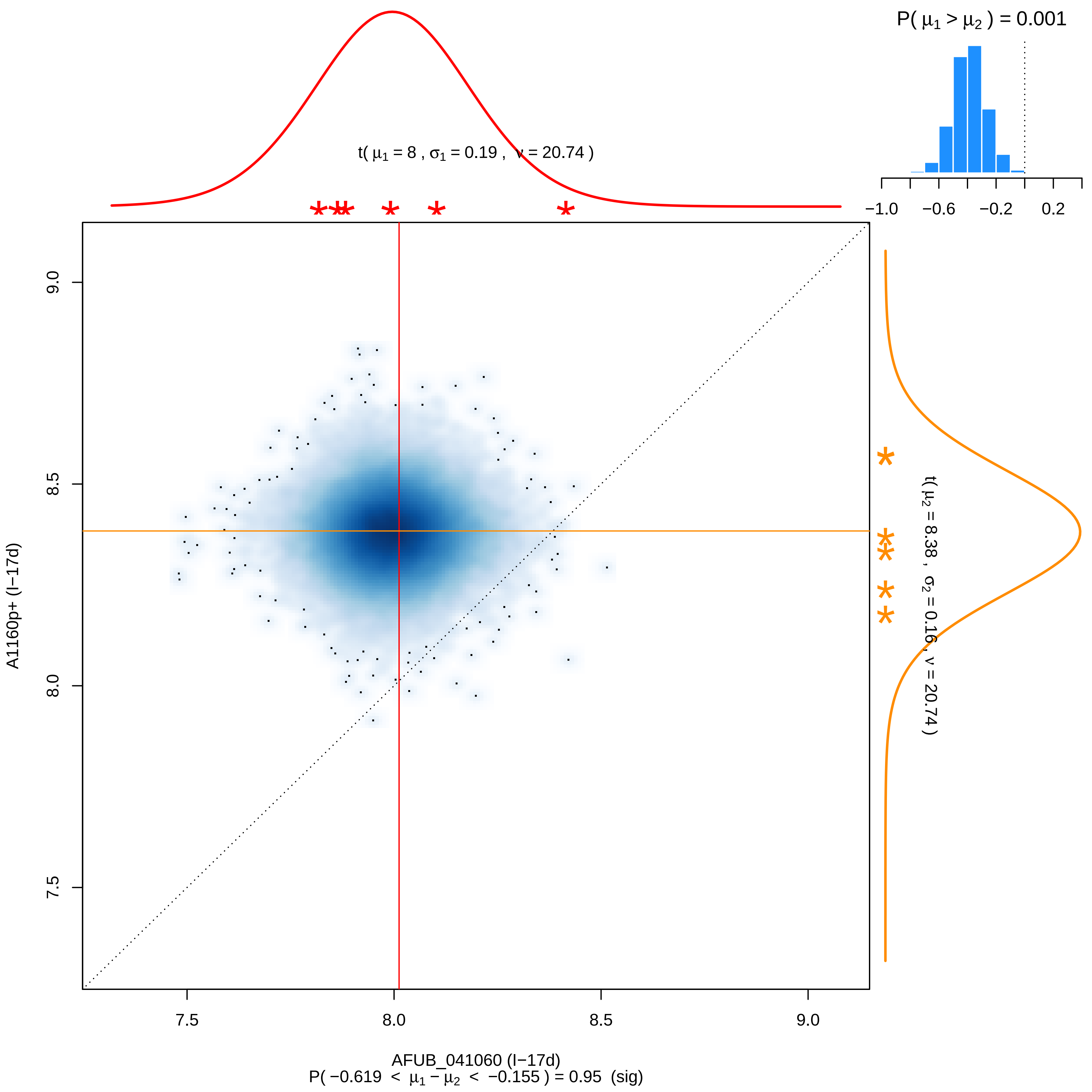

P( μ1 > μ2 ) = 0.001
t( μ1 = 8 , σ1 = 0.19 , ν = 20.74 )
−1.0
−0.6
−0.2
0.2
9.0
8.5
8.0
7.5
A1160p+ (I−17d)
t( μ2 = 8.38 , σ2 = 0.16 , ν = 20.74 )
7.5
8.0
8.5
9.0
AFUB_041060 (I−17d)
P( −0.619 < μ1 − μ2 < −0.155 ) = 0.95 (sig)

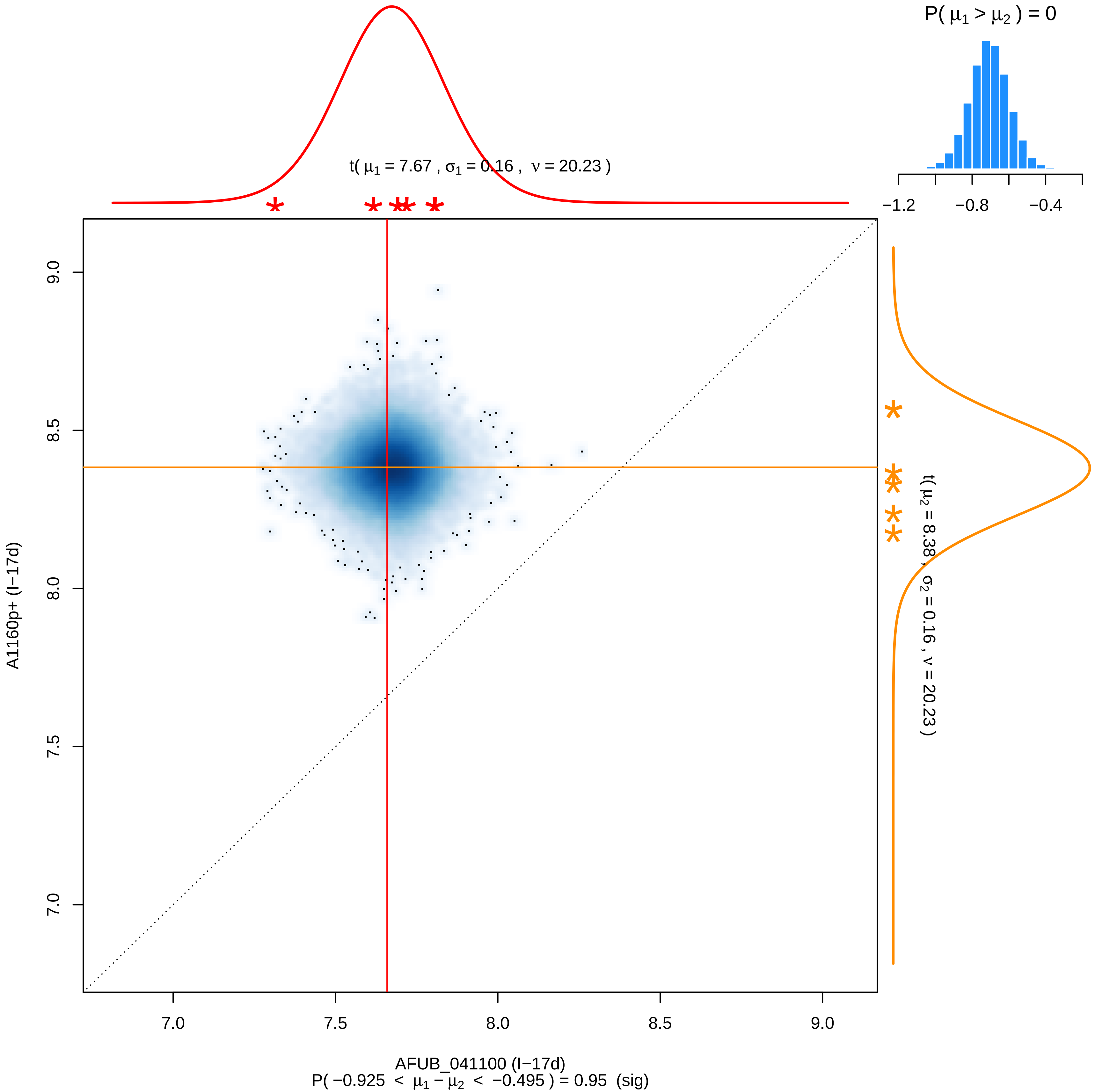

P( μ1 > μ2 ) = 0
−1.2
−0.8
−0.4
t( μ1 = 7.67 , σ1 = 0.16 , ν = 20.23 )
t( μ2 = 8.38 , σ2 = 0.16 , ν = 20.23 )
9.0
8.5
8.0
7.5
7.0
A1160p+ (I−17d)
7.0
7.5
8.0
8.5
9.0
AFUB_041100 (I−17d)
P( −0.925 < μ1 − μ2 < −0.495 ) = 0.95 (sig)

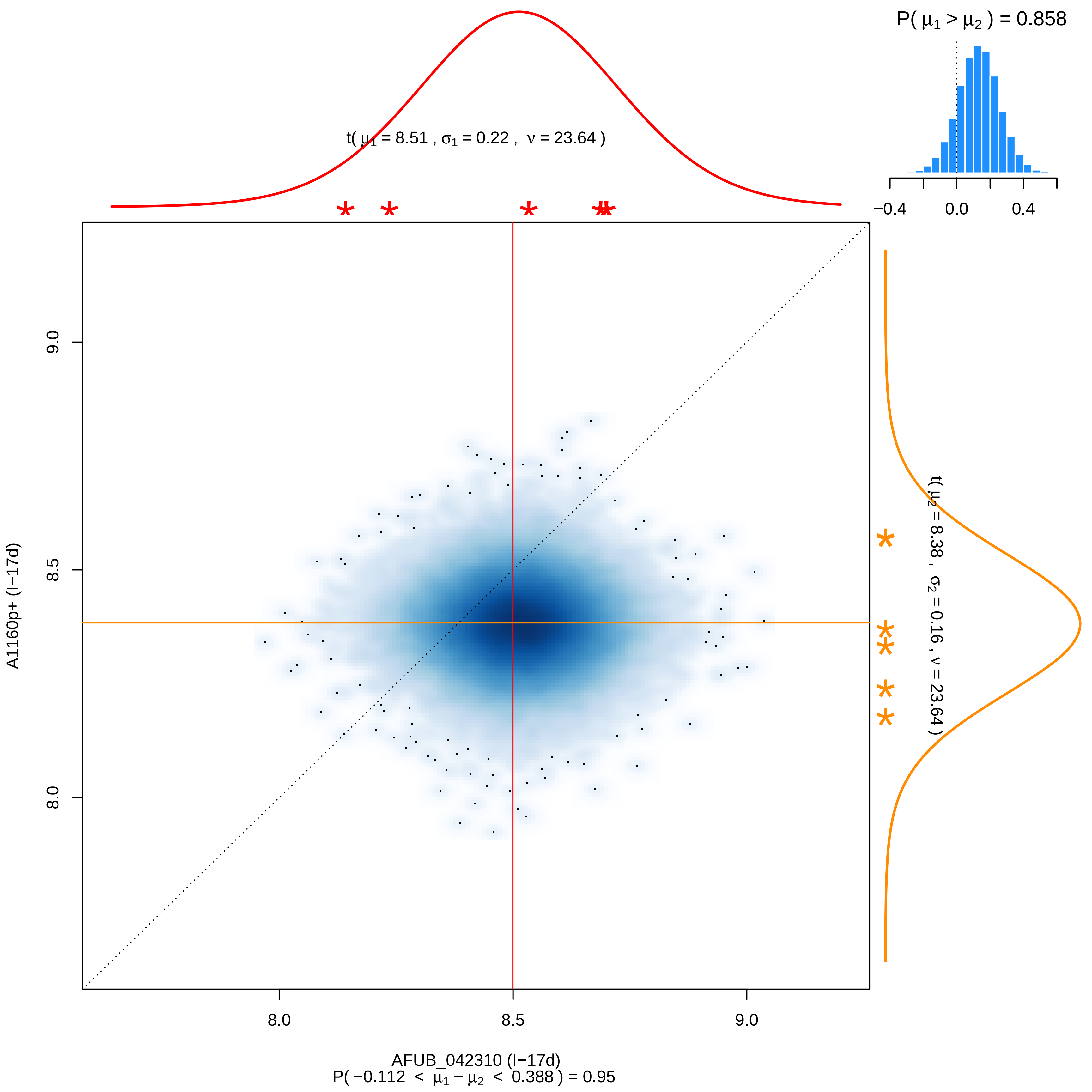

P( μ1 > μ2 ) = 0.858
−0.4
0.0
0.4
t( μ1 = 8.51 , σ1 = 0.22 , ν = 23.64 )
t( μ2 = 8.38 , σ2 = 0.16 , ν = 23.64 )
9.0
8.5
8.0
A1160p+ (I−17d)
8.0
8.5
9.0
AFUB_042310 (I−17d)
P( −0.112 < μ1 − μ2 < 0.388 ) = 0.95

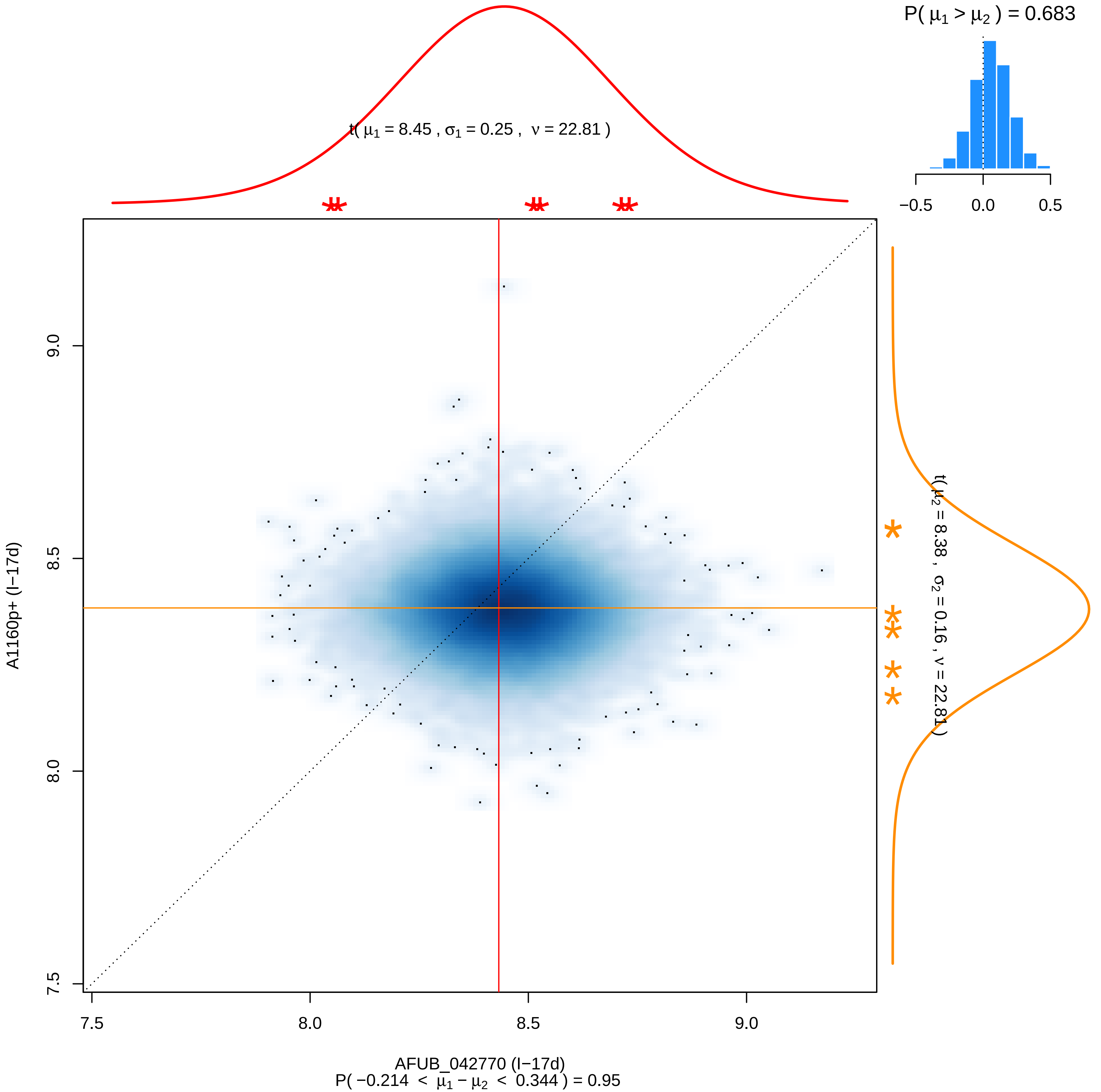
P( μ1 > μ2 ) = 0.683
−0.5
0.0
0.5
t( μ1 = 8.45 , σ1 = 0.25 , ν = 22.81 )
t( μ2 = 8.38 , σ2 = 0.16 , ν = 22.81 )
A1160p+ (I−17d)
7.5
8.0
8.5
9.0
AFUB_042770 (I−17d)
P( −0.214 < μ1 − μ2 < 0.344 ) = 0.95

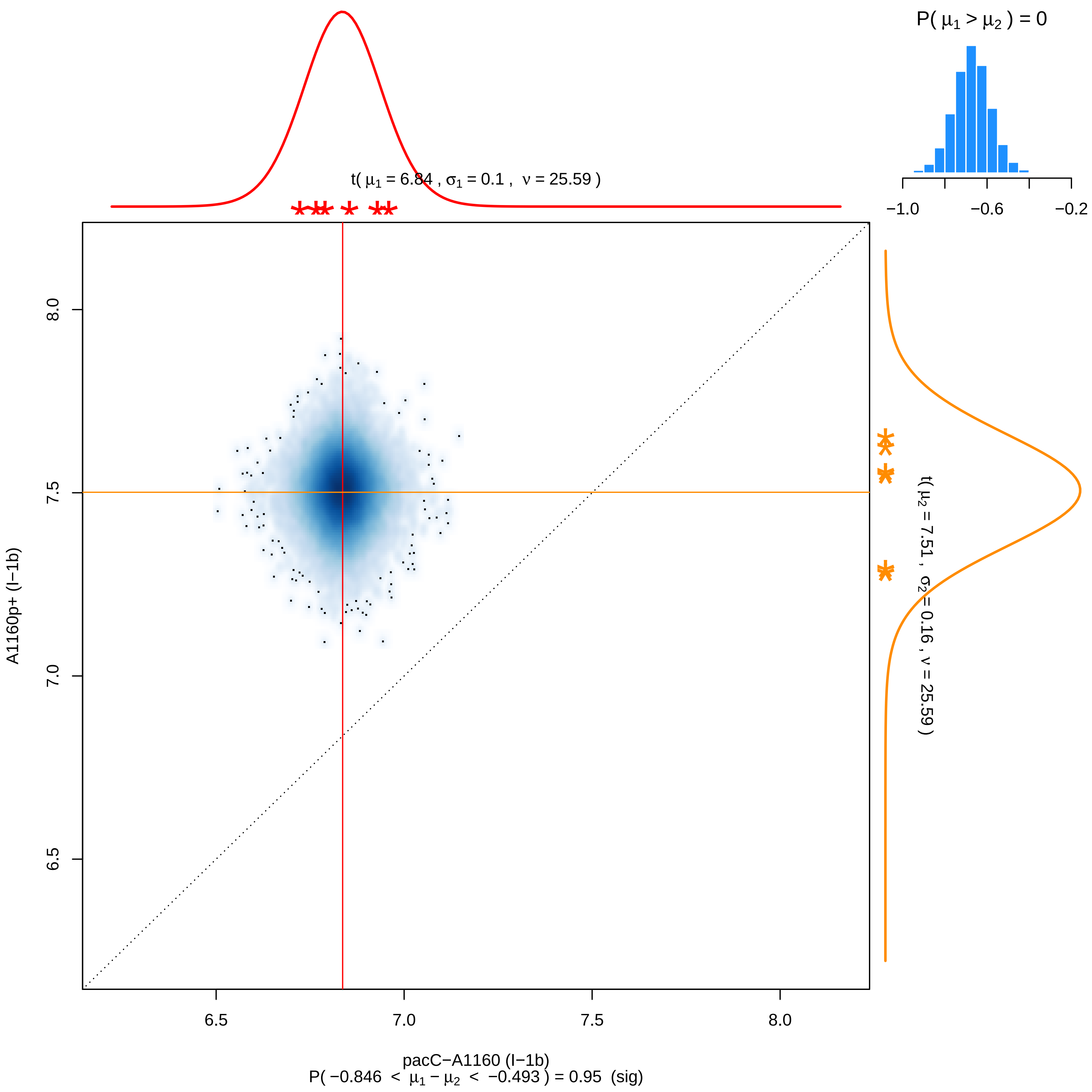

P( μ1 > μ2 ) = 0
−1.0
−0.6
−0.2
t( μ1 = 6.84 , σ1 = 0.1 , ν = 25.59 )
t( μ2 = 7.51 , σ2 = 0.16 , ν = 25.59 )
A1160p+ (I−1b)
8.0
7.5
7.0
6.5
pacC−A1160 (I−1b)
P( −0.846 < μ1 − μ2 < −0.493 ) = 0.95 (sig)

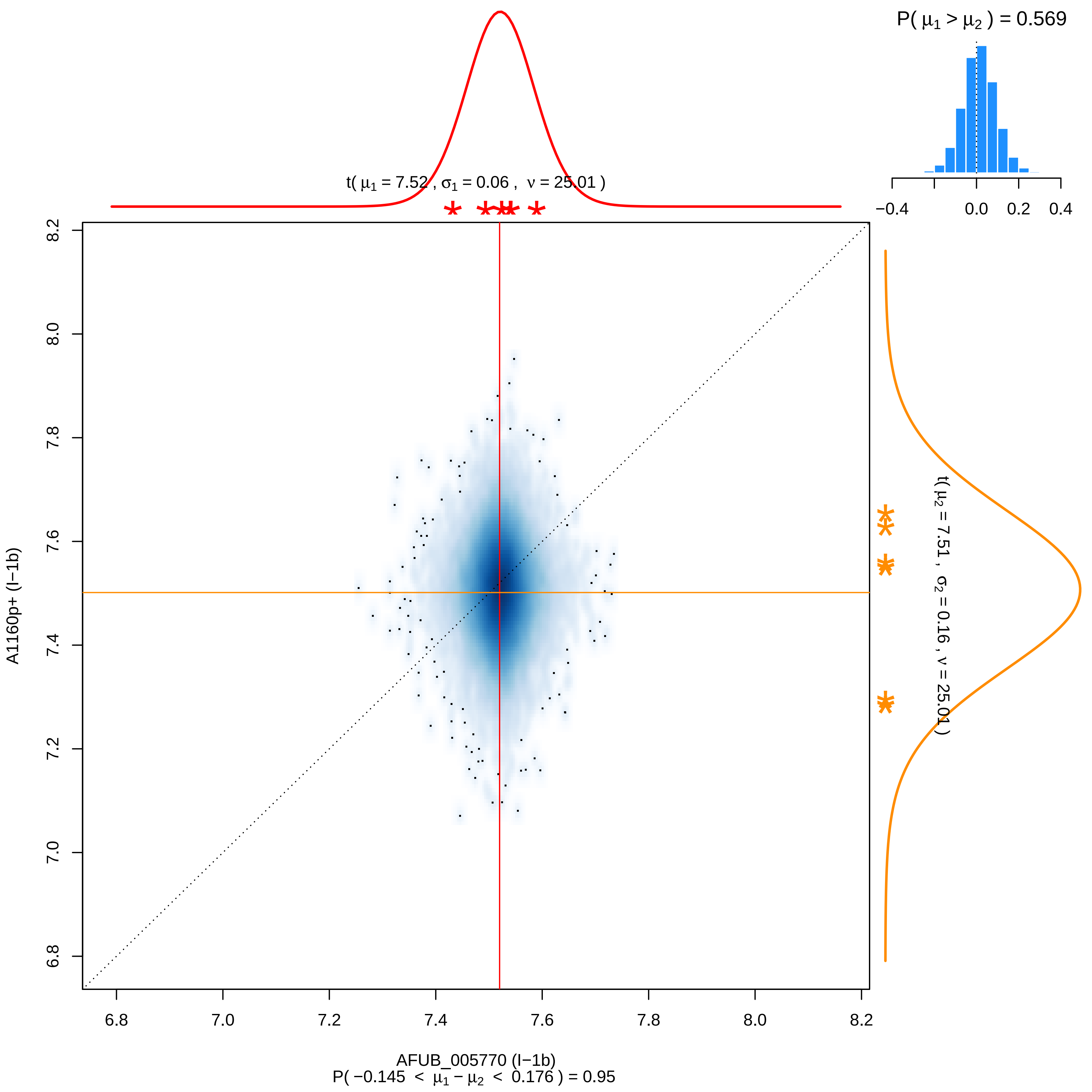

P( μ1 > μ2 ) = 0.569
−0.4
0.0
0.2
0.4
t( μ1 = 7.52 , σ1 = 0.06 , ν = 25.01 )
t( μ2 = 7.51 , σ2 = 0.16 , ν = 25.01 )
A1160p+ (I−1b)
8.2
8.0
7.8
7.6
7.4
7.2
7.0
6.8
AFUB_005770 (I−1b)
P( −0.145 < μ1 − μ2 < 0.176 ) = 0.95

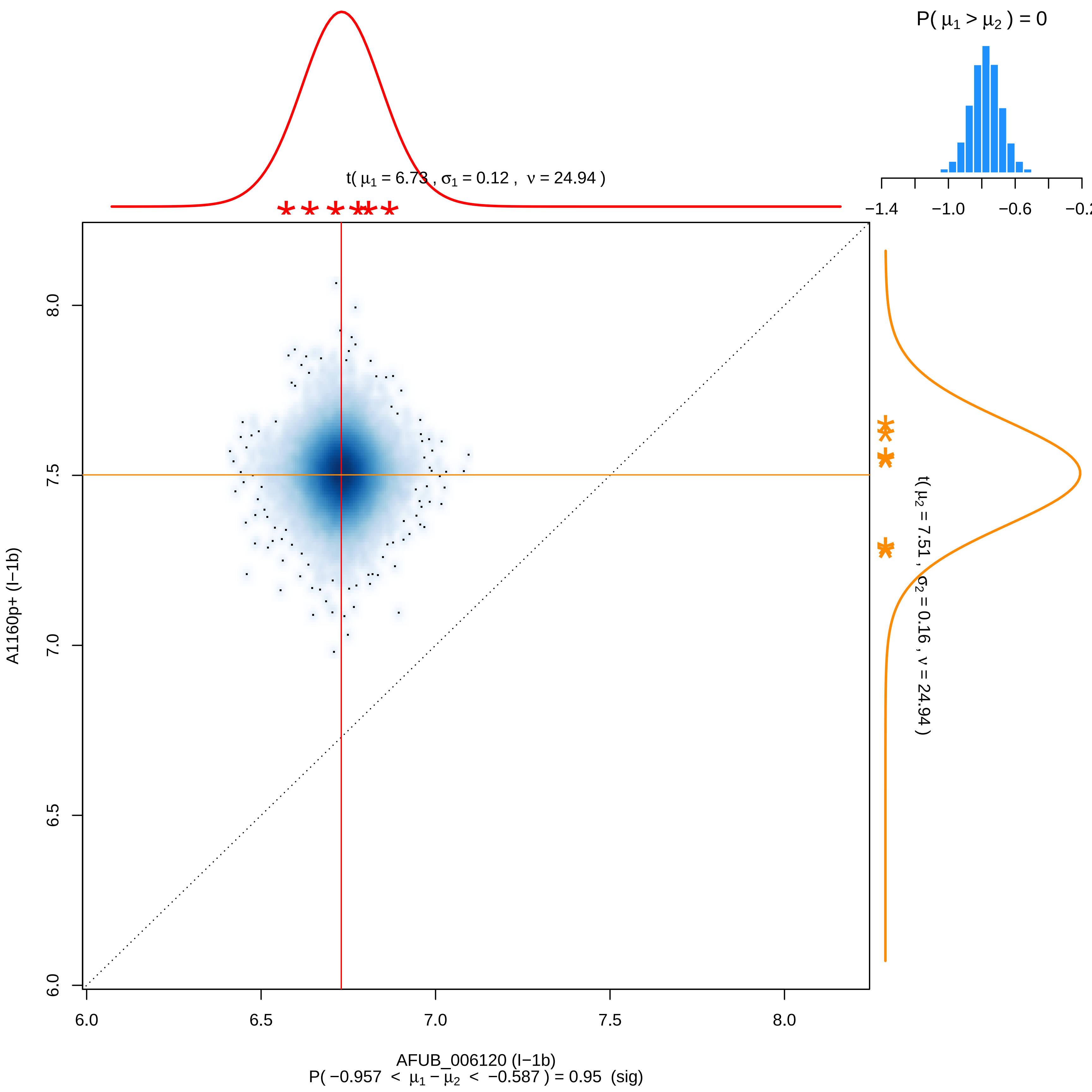

P( μ1 > μ2 ) = 0
−1.4
−1.0
−0.6
−0.2
t( μ1 = 6.73 , σ1 = 0.12 , ν = 24.94 )
t( μ2 = 7.51 , σ2 = 0.16 , ν = 24.94 )
A1160p+ (I−1b)
6.0
6.5
7.0
7.5
8.0
AFUB_006120 (I−1b)
P( −0.957 < μ1 − μ2 < −0.587 ) = 0.95 (sig)

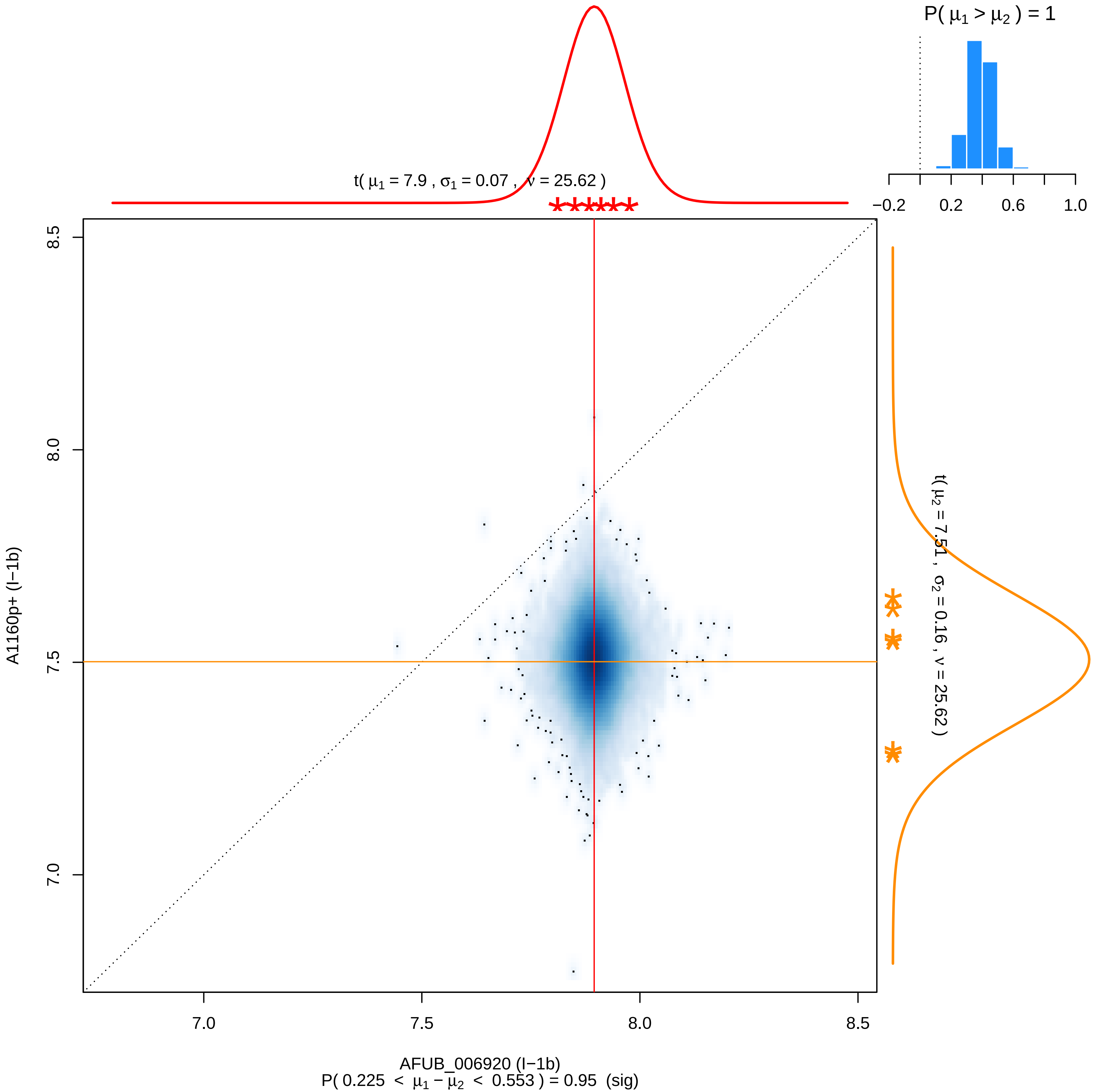

P( $\mu_1 > \mu_2$ ) = 1
−0.2
0.2
0.6
1.0
t( $\mu_1$ = 7.9 , $\sigma_1$ = 0.07 , $\nu$ = 25.62 )
t( $\mu_2$ = 7.51 , $\sigma_2$ = 0.16 , $\nu$ = 25.62 )
8.5
8.0
7.5
7.0
A1160p+ (I−1b)
7.0
7.5
8.0
8.5
AFUB_006920 (I−1b)
P( 0.225 < $\mu_1 - \mu_2$ < 0.553 ) = 0.95 (sig)

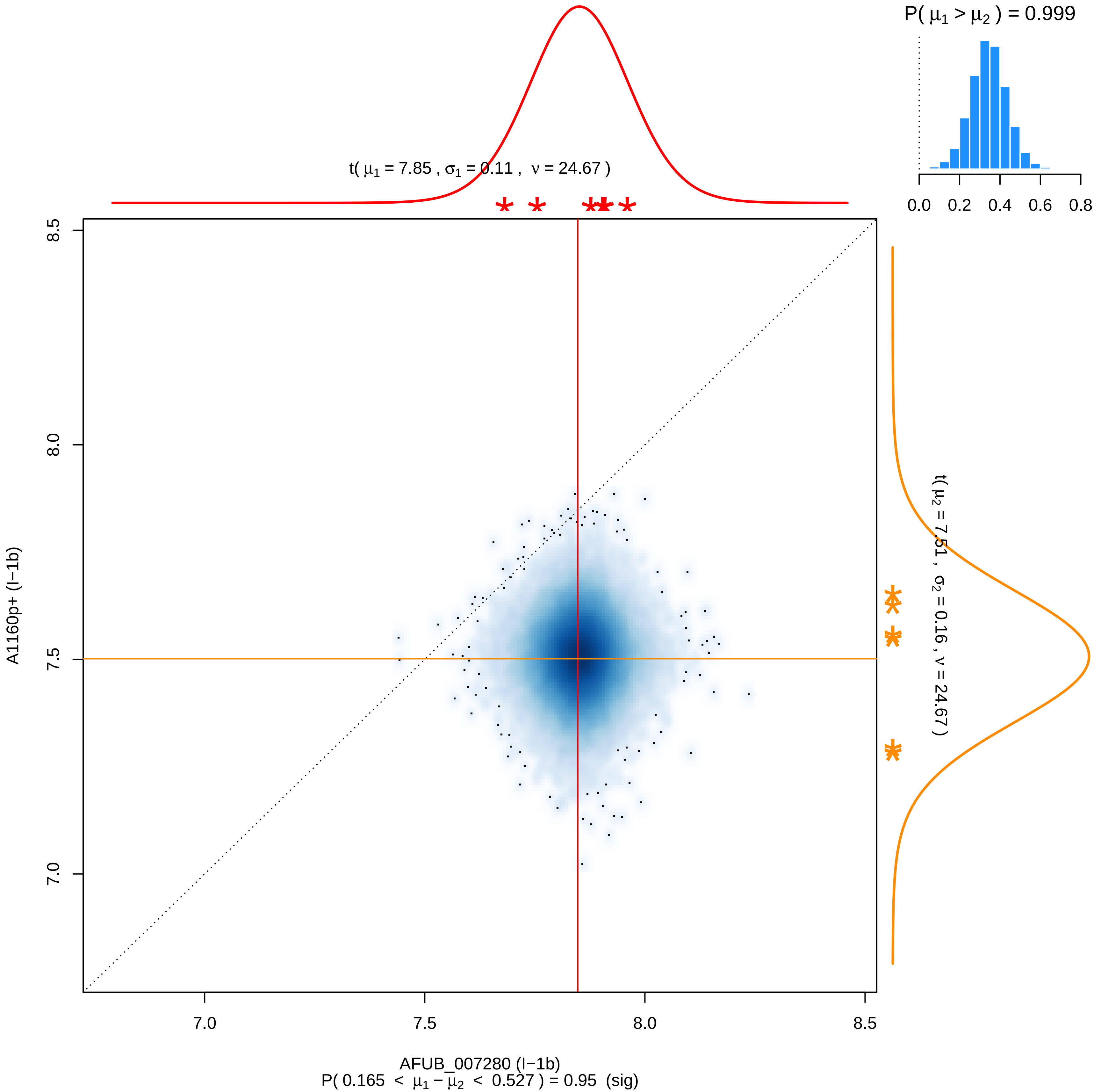
P( μ1 > μ2 ) = 0.999
0.0 0.2 0.4 0.6 0.8
t( μ1 = 7.85 , σ1 = 0.11 , ν = 24.67 )
t( μ2 = 7.51 , σ2 = 0.16 , ν = 24.67 )
8.5
8.0
7.5
7.0
A1160p+ (I−1b)
7.0 7.5 8.0 8.5
AFUB_007280 (I−1b)
P( 0.165 < μ1 − μ2 < 0.527 ) = 0.95 (sig)

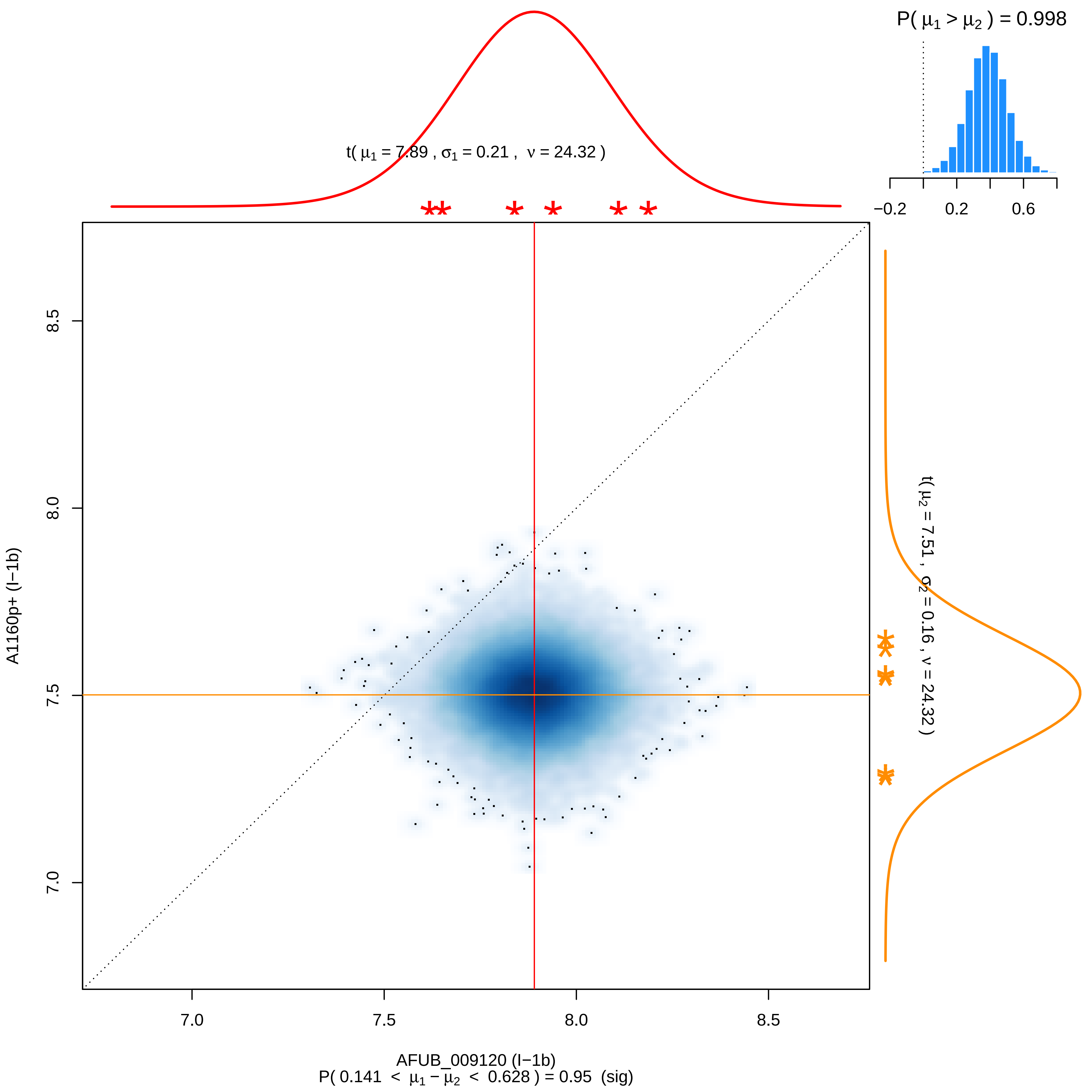

P( μ1 > μ2 ) = 0.998
t( μ1 = 7.89 , σ1 = 0.21 , ν = 24.32 )
−0.2
0.2
0.6
8.5
8.0
7.5
7.0
A1160p+ (I−1b)
t( μ2 = 7.51 , σ2 = 0.16 , ν = 24.32 )
7.0
7.5
8.0
8.5
AFUB_009120 (I−1b)
P( 0.141 < μ1 − μ2 < 0.628 ) = 0.95 (sig)

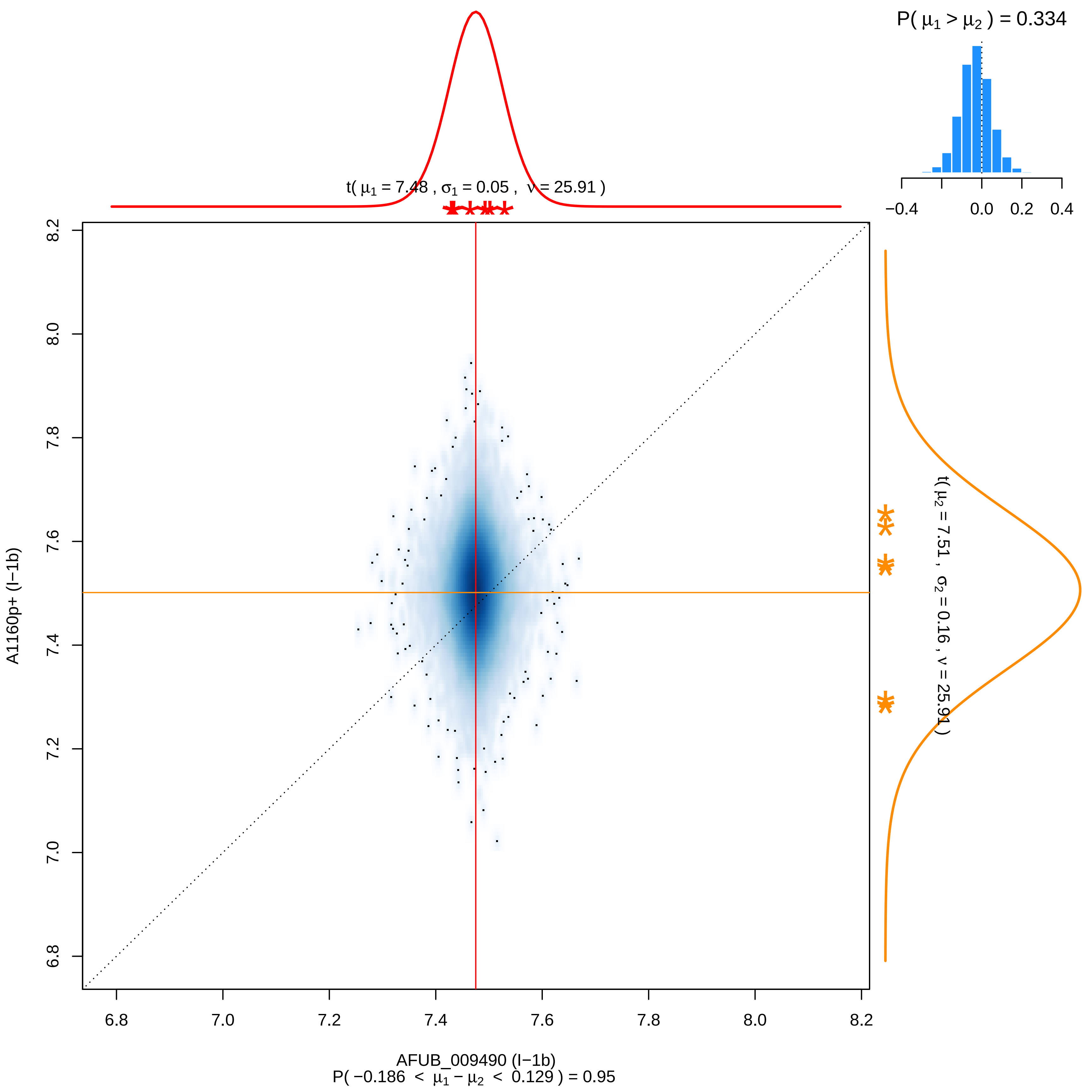

P( μ1 > μ2 ) = 0.334
−0.4
0.0
0.2
0.4
t( μ1 = 7.48 , σ1 = 0.05 , ν = 25.91 )
t( μ2 = 7.51 , σ2 = 0.16 , ν = 25.91 )
A1160p+ (I−1b)
AFUB_009490 (I−1b)
P( −0.186 < μ1 − μ2 < 0.129 ) = 0.95

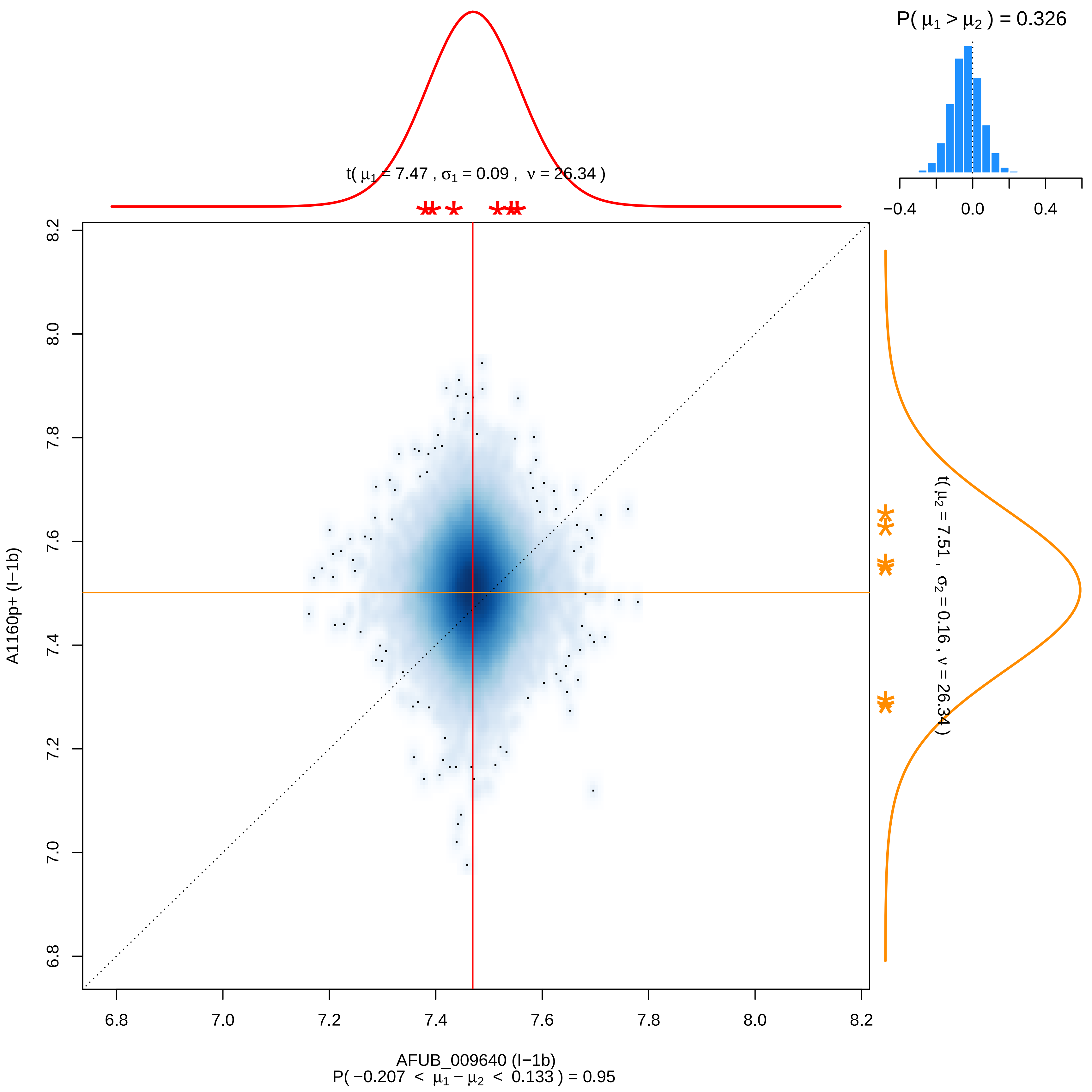

P( μ1 > μ2 ) = 0.326
−0.4
0.0
0.4
t( μ1 = 7.47 , σ1 = 0.09 , ν = 26.34 )
t( μ2 = 7.51 , σ2 = 0.16 , ν = 26.34 )
A1160p+ (I−1b)
8.2
8.0
7.8
7.6
7.4
7.2
7.0
6.8
6.8
7.0
7.2
7.4
7.6
7.8
8.0
8.2
AFUB_009640 (I−1b)
P( −0.207 < μ1 − μ2 < 0.133 ) = 0.95

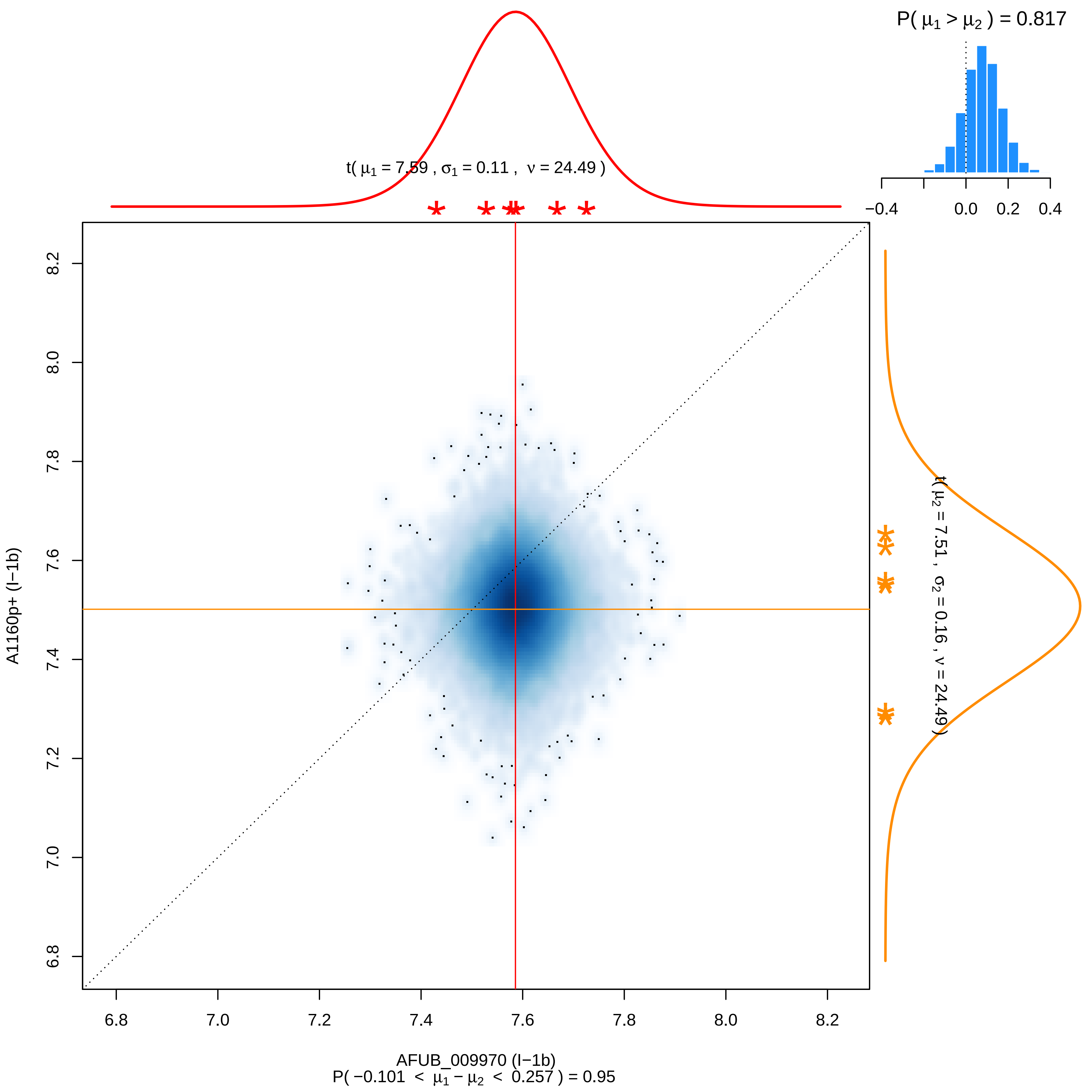

P( μ1 > μ2 ) = 0.817
−0.4
0.0
0.2
0.4
t( μ1 = 7.59 , σ1 = 0.11 , ν = 24.49 )
t( μ2 = 7.51 , σ2 = 0.16 , ν = 24.49 )
A1160p+ (I−1b)
AFUB_009970 (I−1b)
P( −0.101 < μ1 − μ2 < 0.257 ) = 0.95

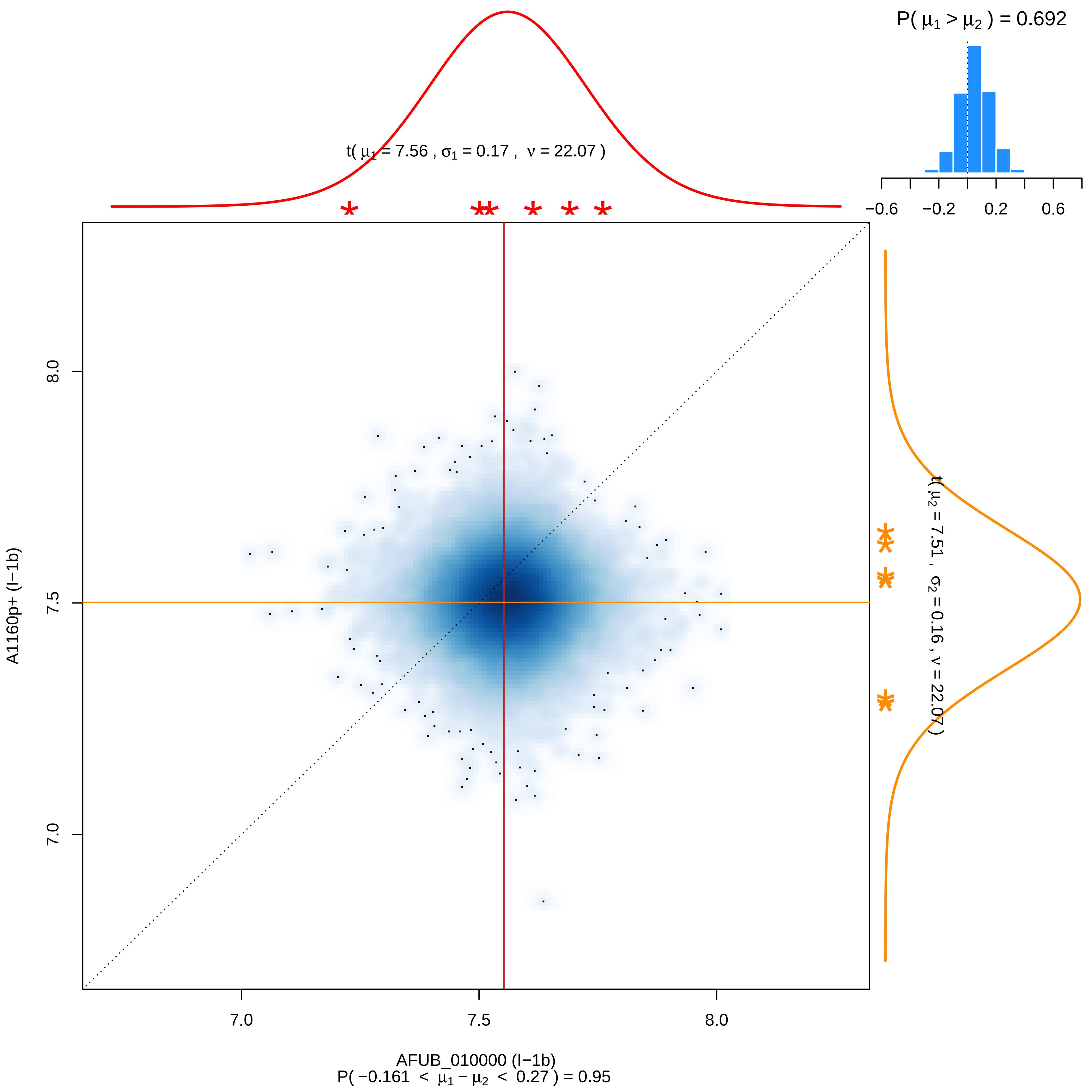
P( μ1 > μ2 ) = 0.692
−0.6
−0.2
0.2
0.6
t( μ1 = 7.56 , σ1 = 0.17 , ν = 22.07 )
t( μ2 = 7.51 , σ2 = 0.16 , ν = 22.07 )
8.0
7.5
7.0
A1160p+ (I−1b)
7.0
7.5
8.0
AFUB_010000 (I−1b)
P( −0.161 < μ1 − μ2 < 0.27 ) = 0.95

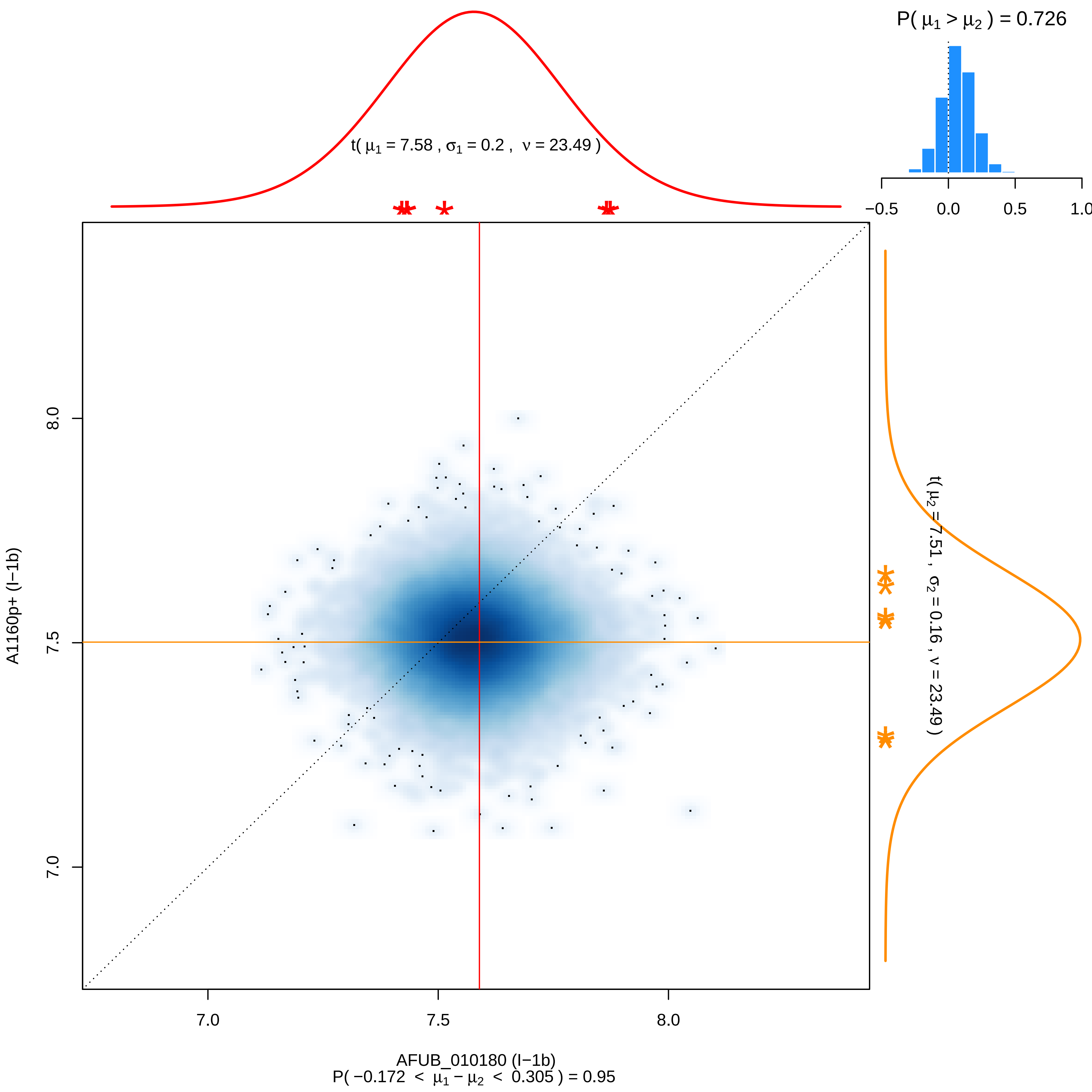
P( μ1 > μ2 ) = 0.726
−0.5
0.0
0.5
1.0
t( μ1 = 7.58 , σ1 = 0.2 , ν = 23.49 )
t( μ2 = 7.51 , σ2 = 0.16 , ν = 23.49 )
A1160p+ (I−1b)
8.0
7.5
7.0
7.0
7.5
8.0
AFUB_010180 (I−1b)
P( −0.172 < μ1 − μ2 < 0.305 ) = 0.95

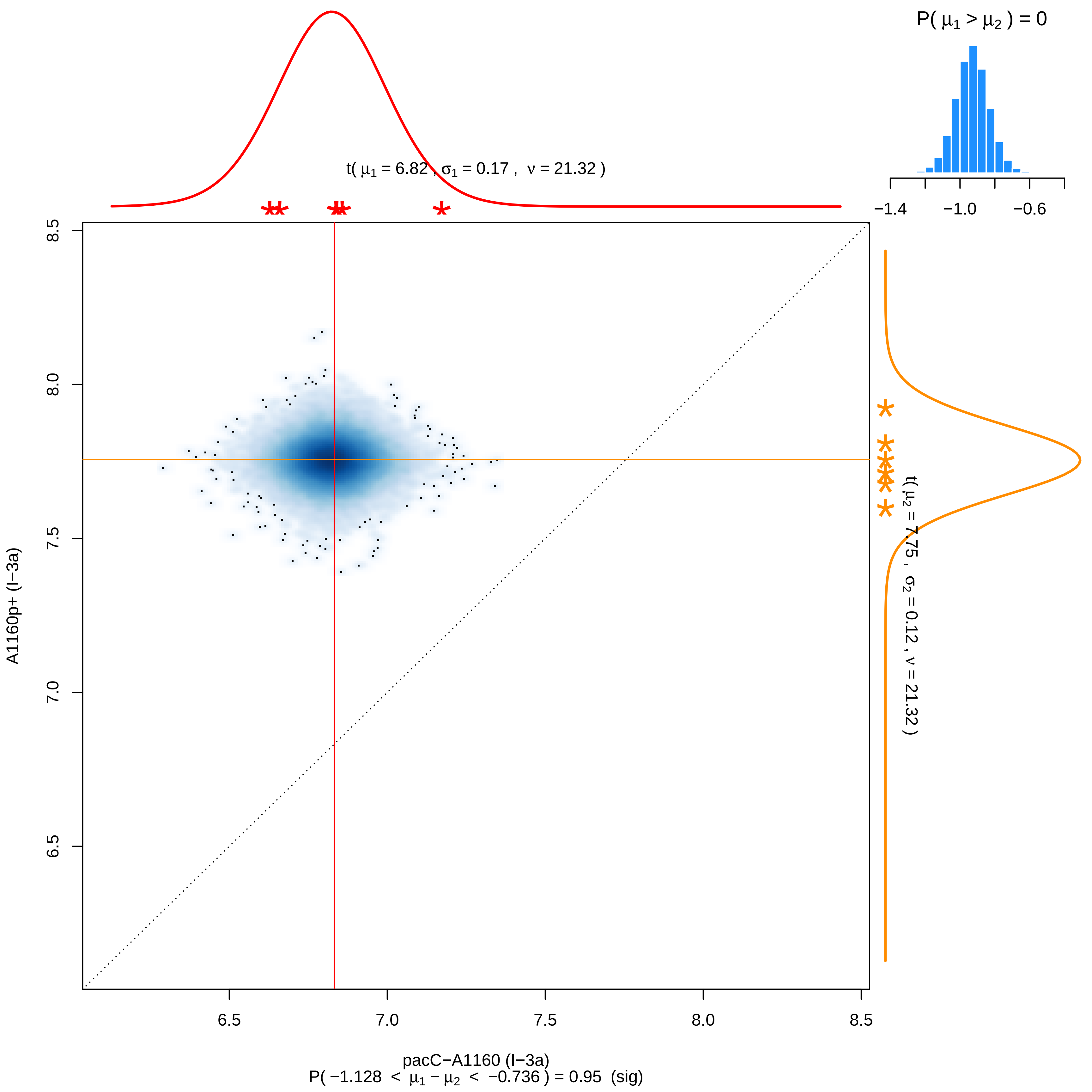

P( μ1 > μ2 ) = 0
−1.4
−1.0
−0.6
t( μ1 = 6.82 , σ1 = 0.17 , ν = 21.32 )
t( μ2 = 7.75 , σ2 = 0.12 , ν = 21.32 )
A1160p+ (I−3a)
8.5
8.0
7.5
7.0
6.5
6.5
7.0
7.5
8.0
8.5
pacC−A1160 (I−3a)
P( −1.128 < μ1 − μ2 < −0.736 ) = 0.95 (sig)

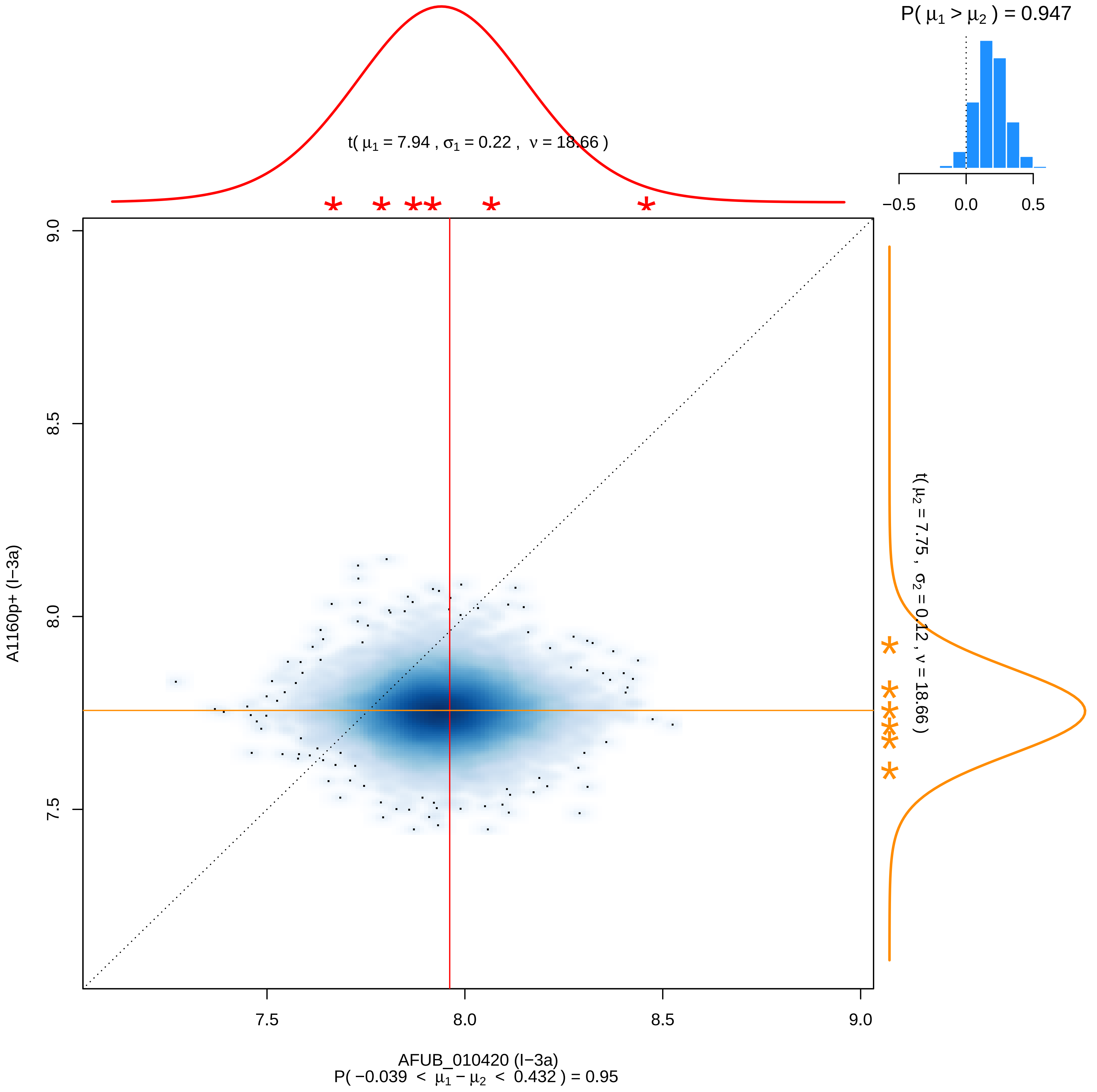

P( μ1 > μ2 ) = 0.947
t( μ1 = 7.94 , σ1 = 0.22 , ν = 18.66 )
t( μ2 = 7.75 , σ2 = 0.12 , ν = 18.66 )
−0.5
0.0
0.5
9.0
8.5
8.0
7.5
A1160p+ (I−3a)
AFUB_010420 (I−3a)
P( −0.039 < μ1 − μ2 < 0.432 ) = 0.95

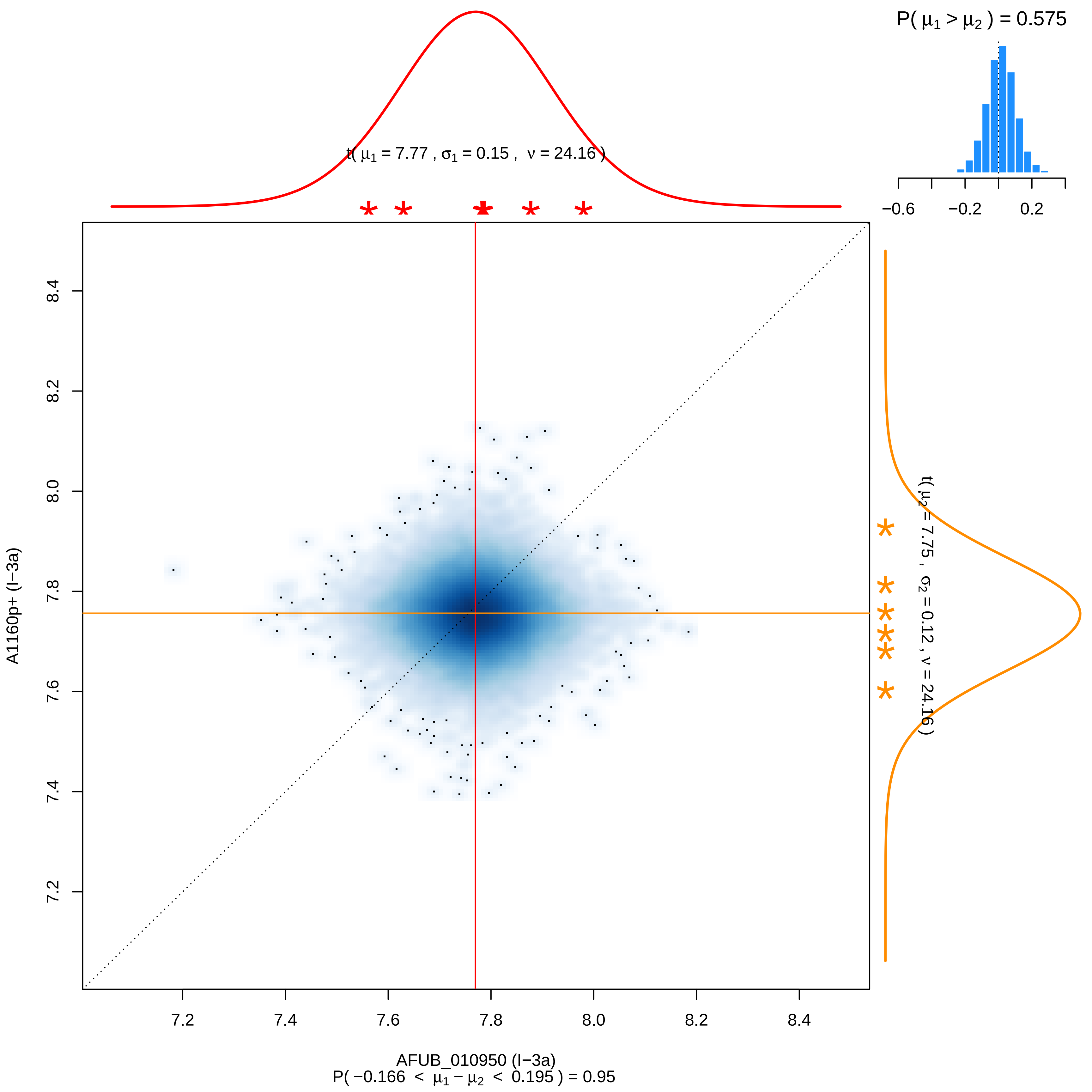

P( μ1 > μ2 ) = 0.575
−0.6
−0.2
0.2
t( μ1 = 7.77 , σ1 = 0.15 , ν = 24.16 )
t( μ2 = 7.75 , σ2 = 0.12 , ν = 24.16 )
A1160p+ (I−3a)
8.4
8.2
8.0
7.8
7.6
7.4
7.2
AFUB_010950 (I−3a)
P( −0.166 < μ1 − μ2 < 0.195 ) = 0.95

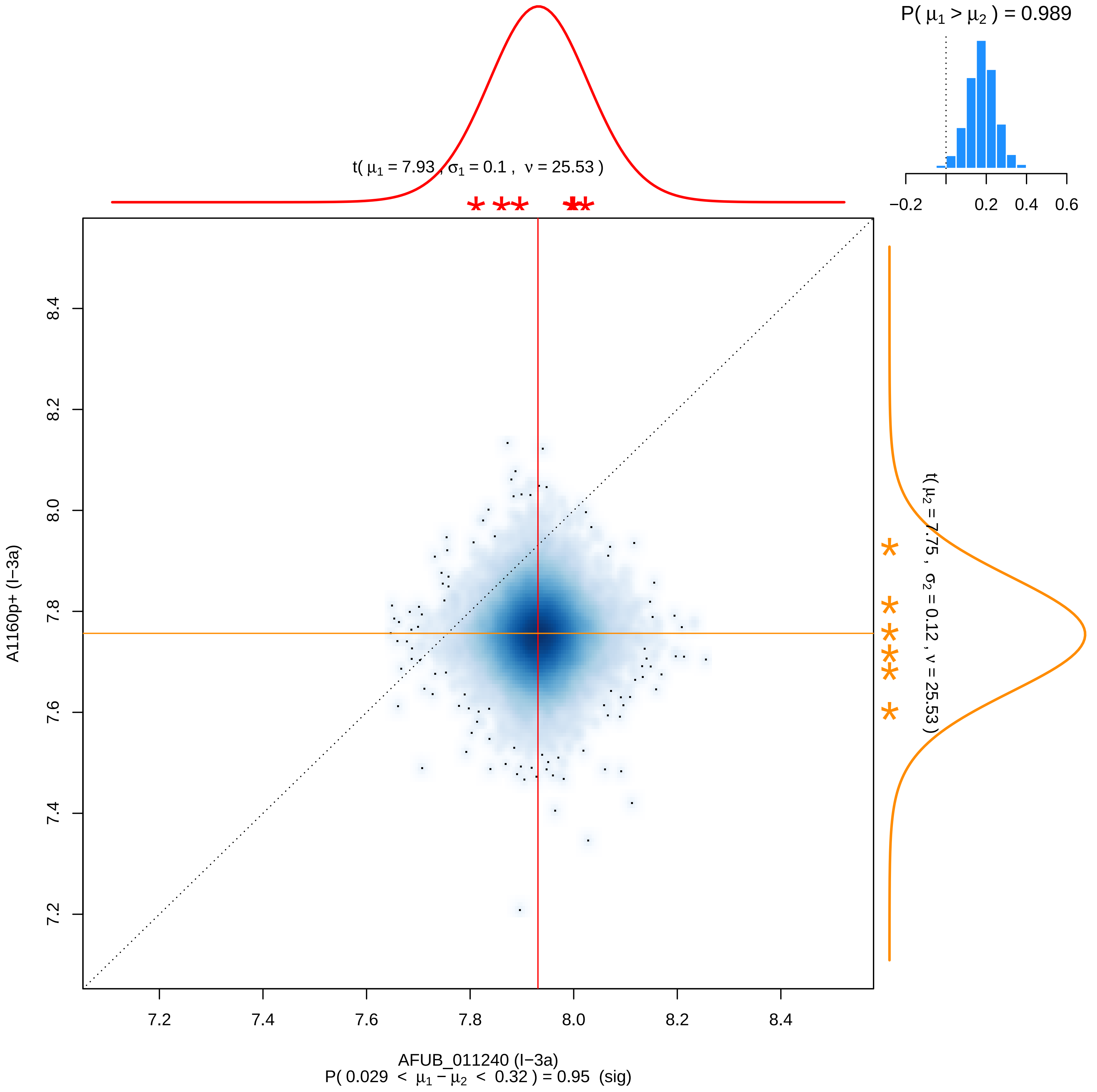
P( μ1 > μ2 ) = 0.989
−0.2
0.2
0.4
0.6
t( μ1 = 7.93 , σ1 = 0.1 , ν = 25.53 )
t( μ2 = 7.75 , σ2 = 0.12 , ν = 25.53 )
A1160p+ (I−3a)
7.2
7.4
7.6
7.8
8.0
8.2
8.4
AFUB_011240 (I−3a)
P( 0.029 < μ1 − μ2 < 0.32 ) = 0.95 (sig)

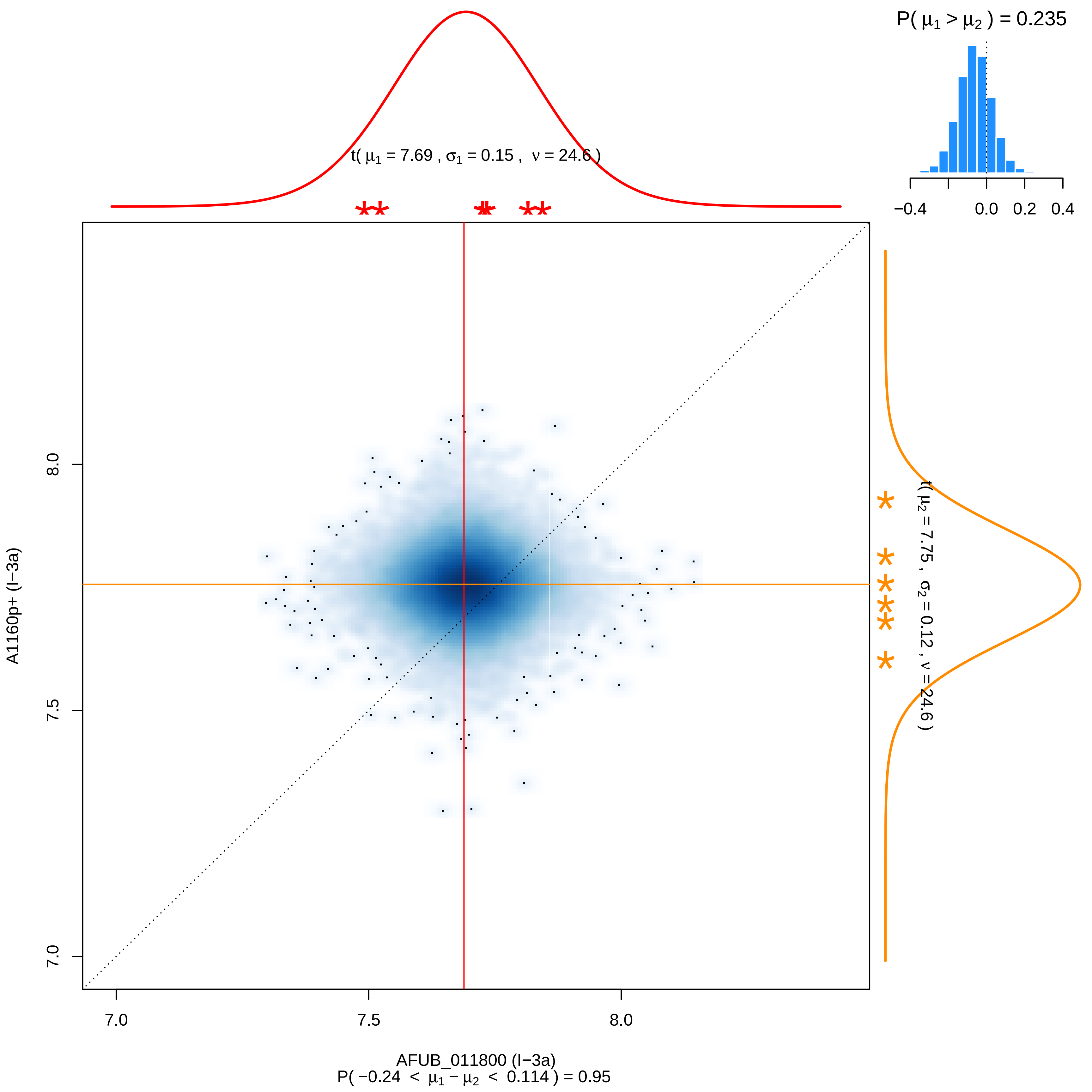

P( $\mu_1 > \mu_2$ ) = 0.235
−0.4
0.0
0.2
0.4
t( $\mu_1$ = 7.69 , $\sigma_1$ = 0.15 , $\nu$ = 24.6 )
t( $\mu_2$ = 7.75 , $\sigma_2$ = 0.12 , $\nu$ = 24.6 )
A1160p+ (I−3a)
7.0
7.5
8.0
AFUB_011800 (I−3a)
P( −0.24 < $\mu_1 - \mu_2$ < 0.114 ) = 0.95

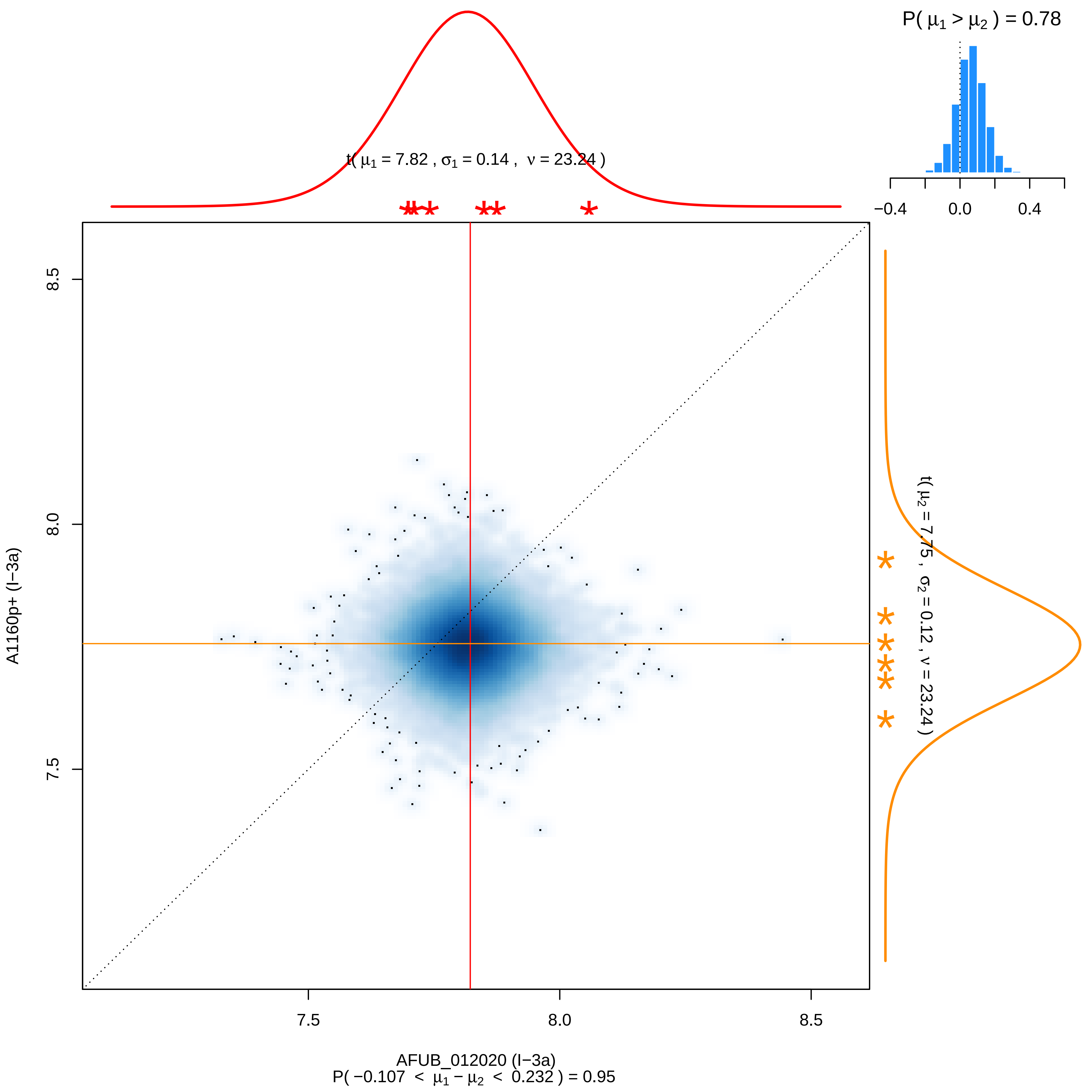
P( μ1 > μ2 ) = 0.78
−0.4
0.0
0.4
t( μ1 = 7.82 , σ1 = 0.14 , ν = 23.24 )
t( μ2 = 7.75 , σ2 = 0.12 , ν = 23.24 )
8.5
8.0
7.5
A1160p+ (I−3a)
7.5
8.0
8.5
AFUB_012020 (I−3a)
P( −0.107 < μ1 − μ2 < 0.232 ) = 0.95

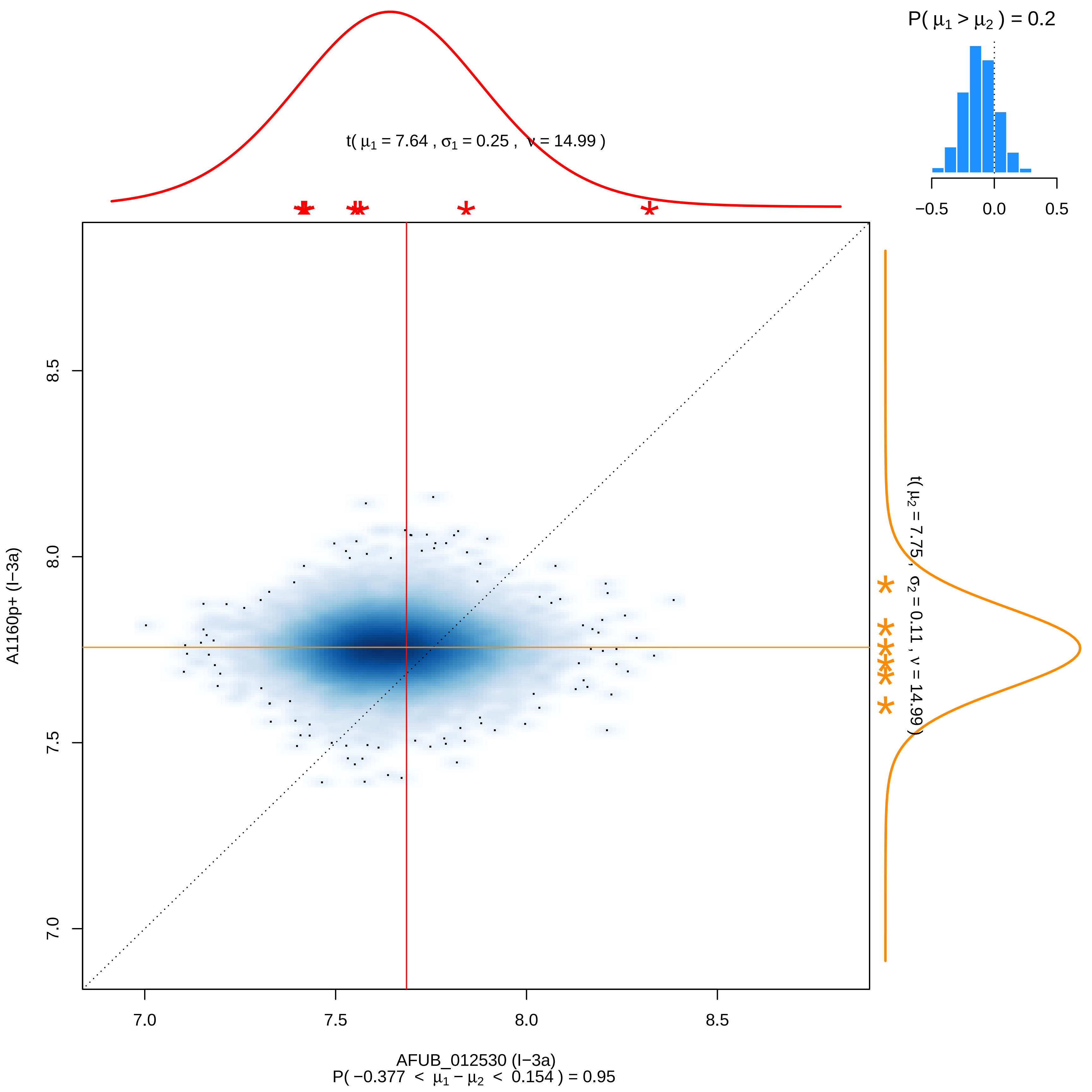

P( μ1 > μ2 ) = 0.2
−0.5
0.0
0.5
t( μ1 = 7.64 , σ1 = 0.25 , ν = 14.99 )
t( μ2 = 7.75 , σ2 = 0.11 , ν = 14.99 )
A1160p+ (I−3a)
8.5
8.0
7.5
7.0
7.0
7.5
8.0
8.5
AFUB_012530 (I−3a)
P( −0.377 < μ1 − μ2 < 0.154 ) = 0.95

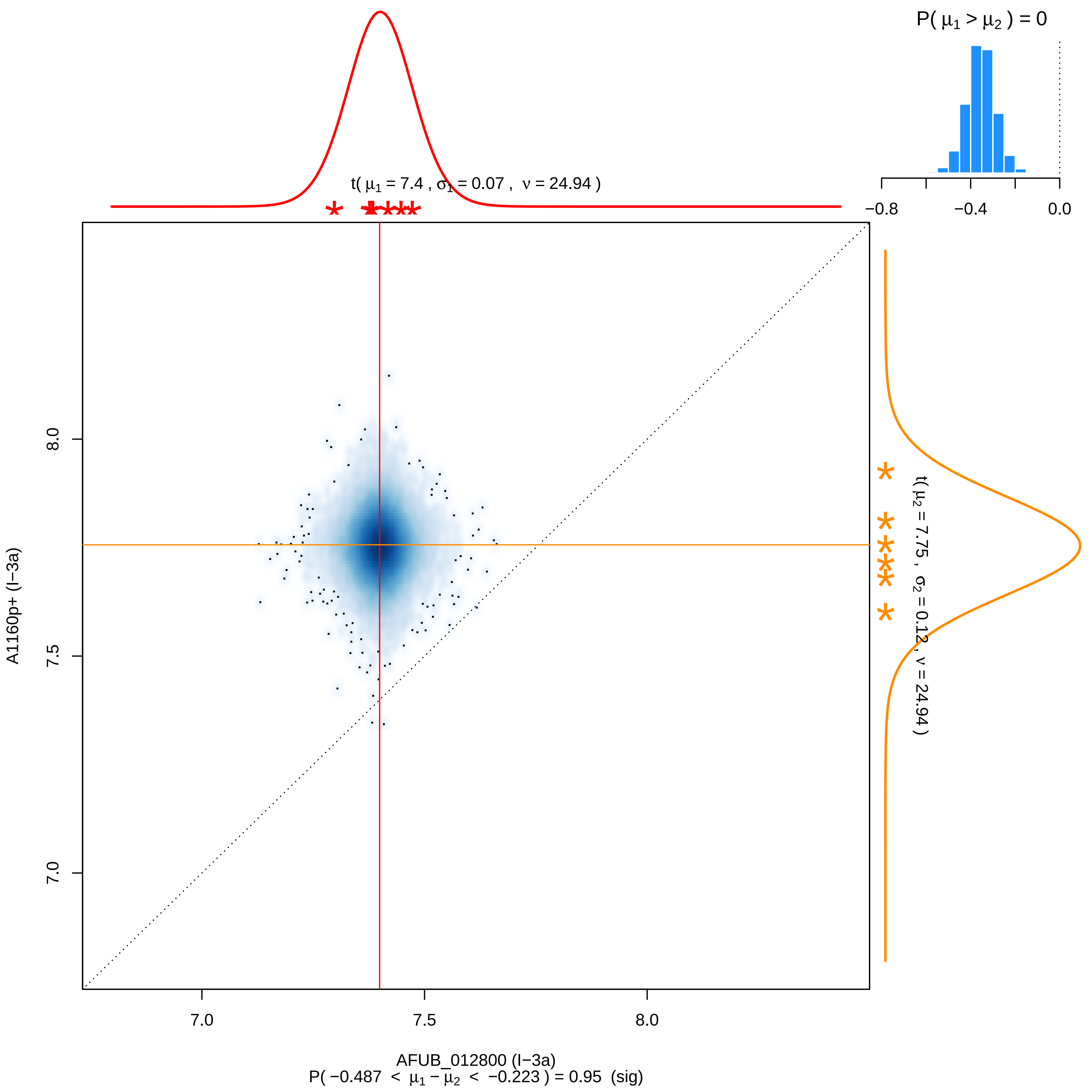

P( μ1 > μ2 ) = 0
−0.8
−0.4
0.0
t( μ1 = 7.4 , σ1 = 0.07 , ν = 24.94 )
t( μ2 = 7.75 , σ2 = 0.12 , ν = 24.94 )
A1160p+ (I−3a)
8.0
7.5
7.0
7.0
7.5
8.0
AFUB_012800 (I−3a)
P( −0.487 < μ1 − μ2 < −0.223 ) = 0.95 (sig)

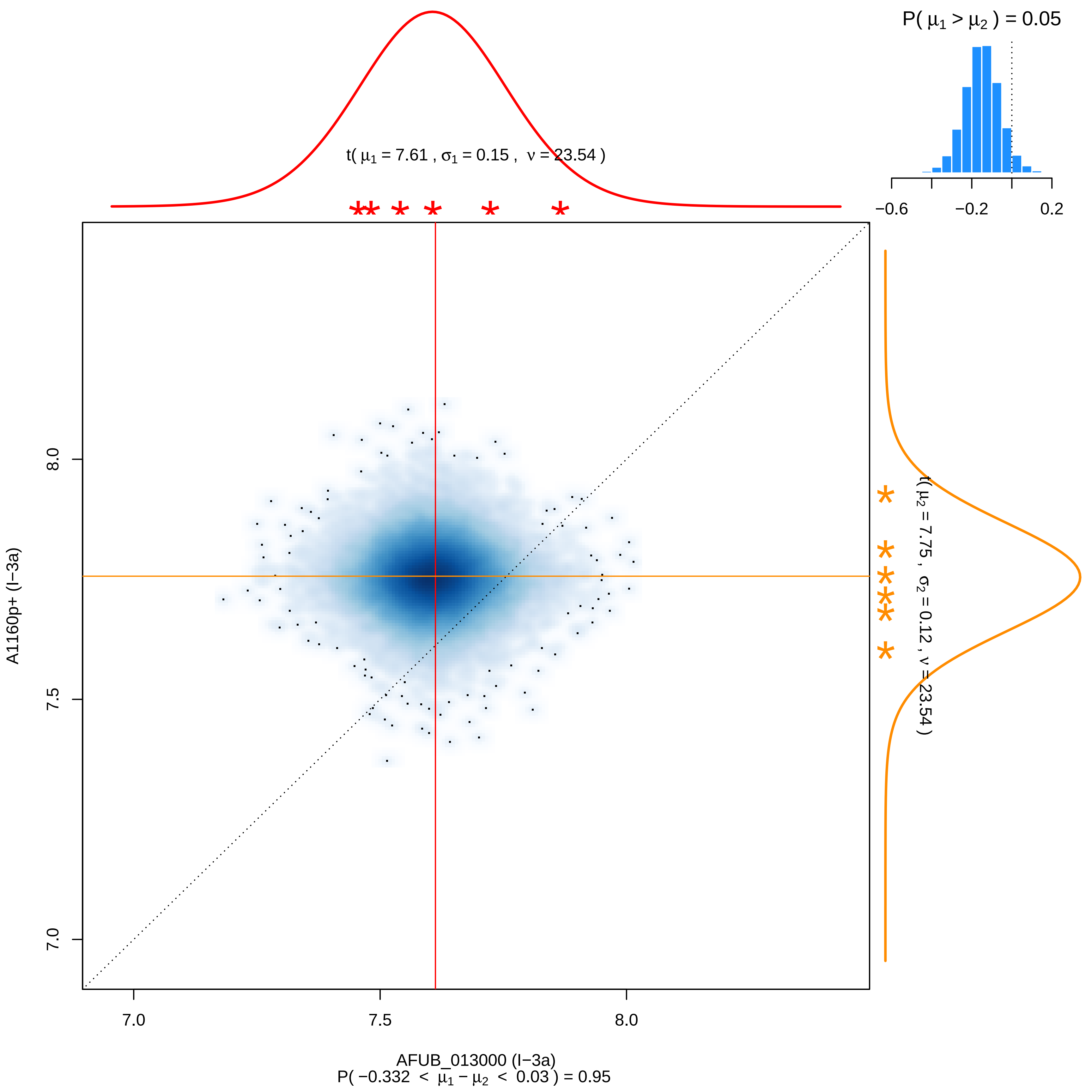

P( μ1 > μ2 ) = 0.05
−0.6
−0.2
0.2
t( μ1 = 7.61 , σ1 = 0.15 , ν = 23.54 )
t( μ2 = 7.75 , σ2 = 0.12 , ν = 23.54 )
A1160p+ (I−3a)
8.0
7.5
7.0
7.0
7.5
8.0
AFUB_013000 (I−3a)
P( −0.332 < μ1 − μ2 < 0.03 ) = 0.95

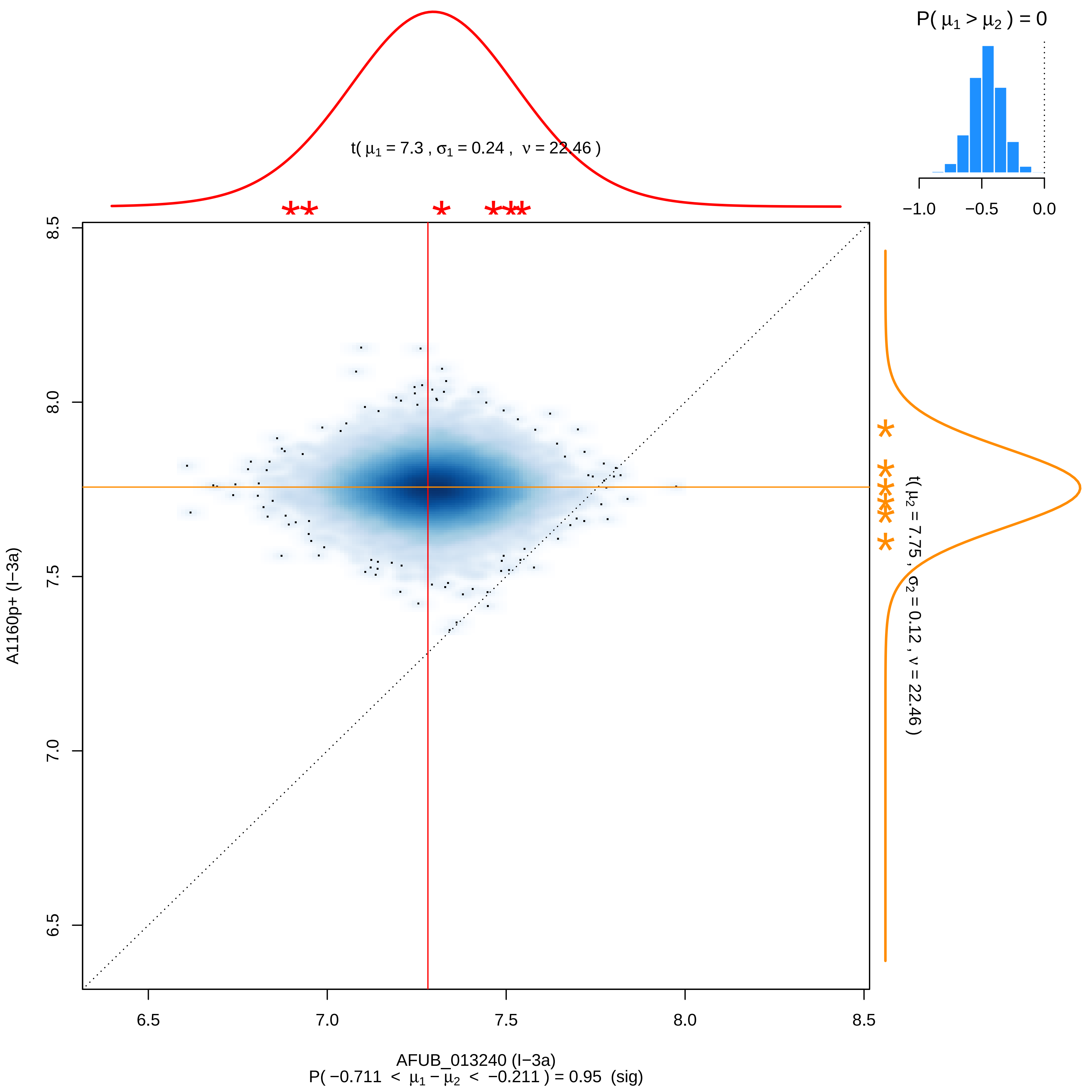

P( μ1 > μ2 ) = 0
−1.0
−0.5
0.0
t( μ1 = 7.3 , σ1 = 0.24 , ν = 22.46 )
t( μ2 = 7.75 , σ2 = 0.12 , ν = 22.46 )
A1160p+ (I−3a)
6.5
7.0
7.5
8.0
8.5
AFUB_013240 (I−3a)
P( −0.711 < μ1 − μ2 < −0.211 ) = 0.95 (sig)

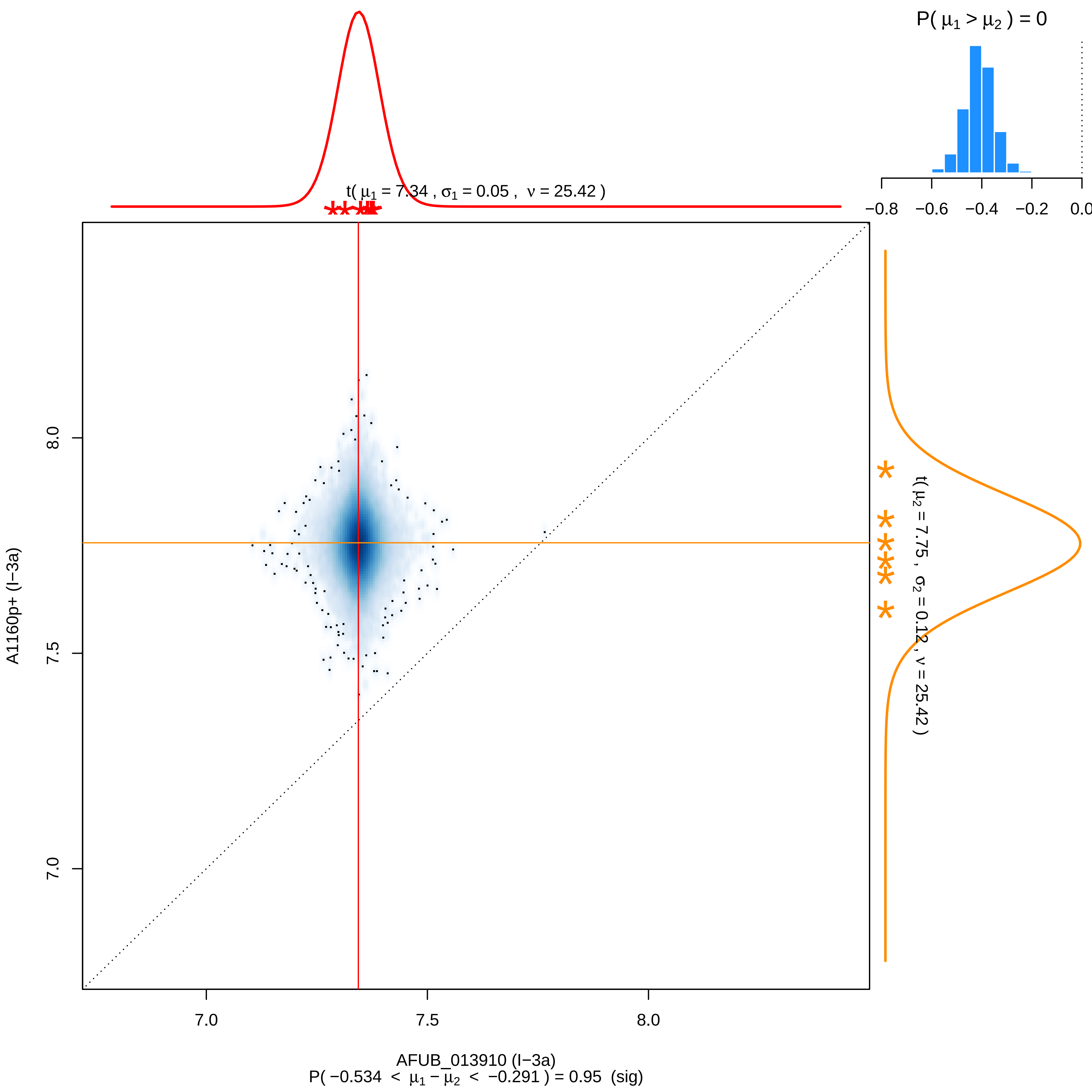

P( μ1 > μ2 ) = 0
−0.8
−0.6
−0.4
−0.2
0.0
t( μ1 = 7.34 , σ1 = 0.05 , ν = 25.42 )
t( μ2 = 7.75 , σ2 = 0.12 , ν = 25.42 )
8.0
7.5
7.0
A1160p+ (I−3a)
7.0
7.5
8.0
AFUB_013910 (I−3a)
P( −0.534 < μ1 − μ2 < −0.291 ) = 0.95 (sig)

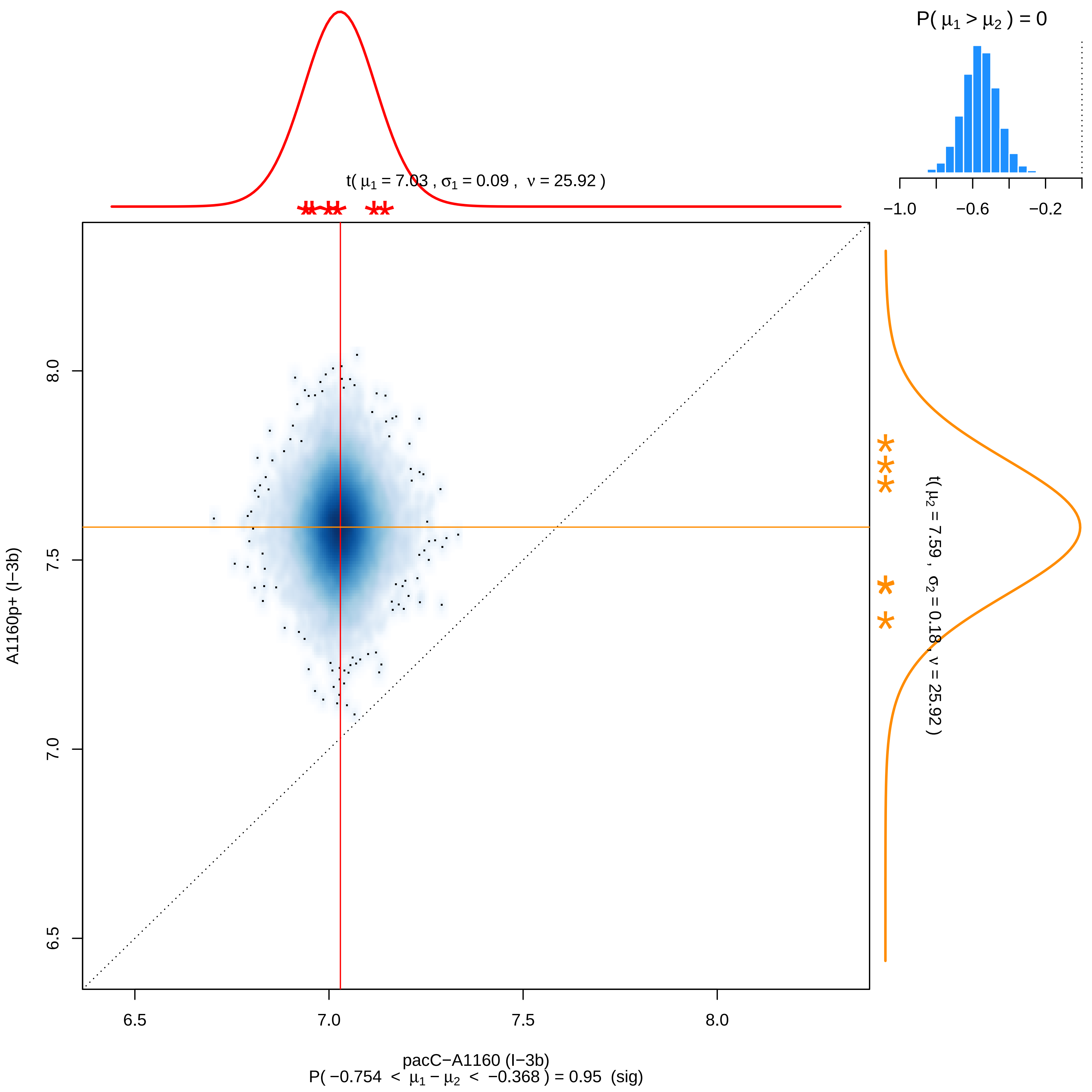
P( $\mu_1 > \mu_2$ ) = 0
−1.0
−0.6
−0.2
t( $\mu_1$ = 7.03 , $\sigma_1$ = 0.09 , $\nu$ = 25.92 )
t( $\mu_2$ = 7.59 , $\sigma_2$ = 0.18 , $\nu$ = 25.92 )
A1160p+ (I−3b)
8.0
7.5
7.0
6.5
6.5
7.0
7.5
8.0
pacC−A1160 (I−3b)
P( −0.754 < $\mu_1 - \mu_2$ < −0.368 ) = 0.95 (sig)

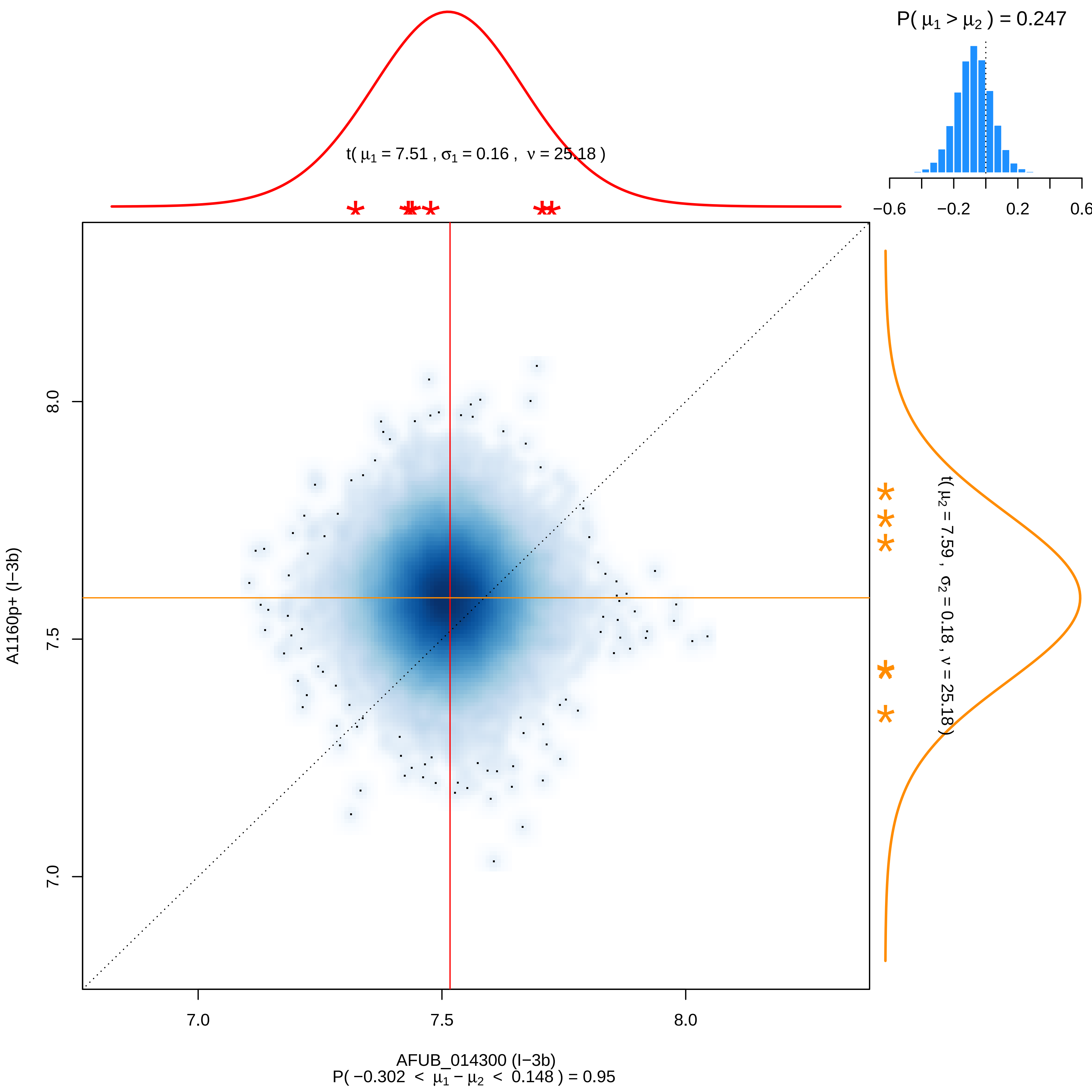

P( μ1 > μ2 ) = 0.247
−0.6
−0.2
0.2
0.6
t( μ1 = 7.51 , σ1 = 0.16 , ν = 25.18 )
t( μ2 = 7.59 , σ2 = 0.18 , ν = 25.18 )
8.0
7.5
7.0
A1160p+ (I−3b)
7.0
7.5
8.0
AFUB_014300 (I−3b)
P( −0.302 < μ1 − μ2 < 0.148 ) = 0.95

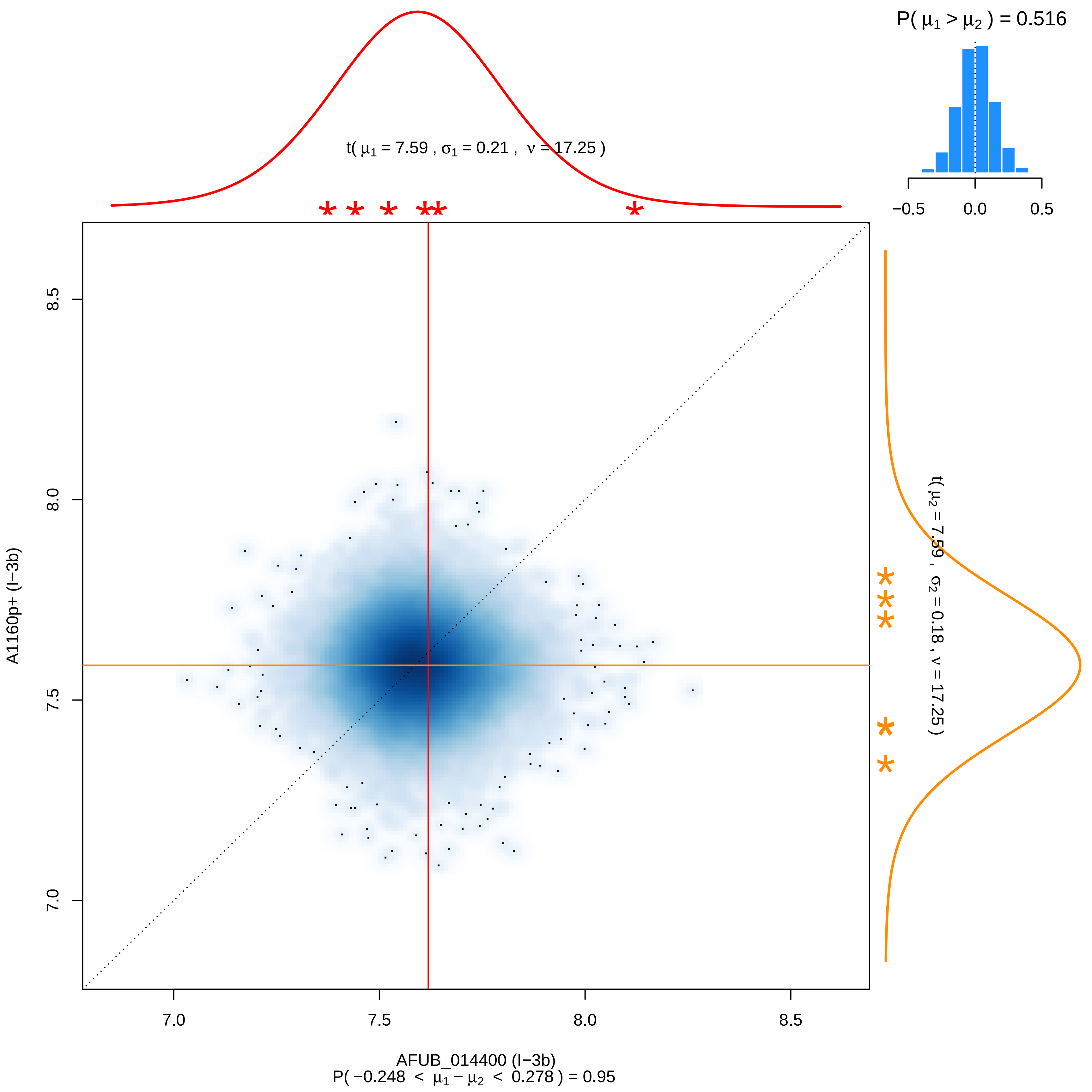

P( μ1 > μ2 ) = 0.516
−0.5
0.0
0.5
t( μ1 = 7.59 , σ1 = 0.21 , ν = 17.25 )
8.5
8.0
7.5
7.0
A1160p+ (I−3b)
t( μ2 = 7.59 , σ2 = 0.18 , ν = 17.25 )
7.0
7.5
8.0
8.5
AFUB_014400 (I−3b)
P( −0.248 < μ1 − μ2 < 0.278 ) = 0.95

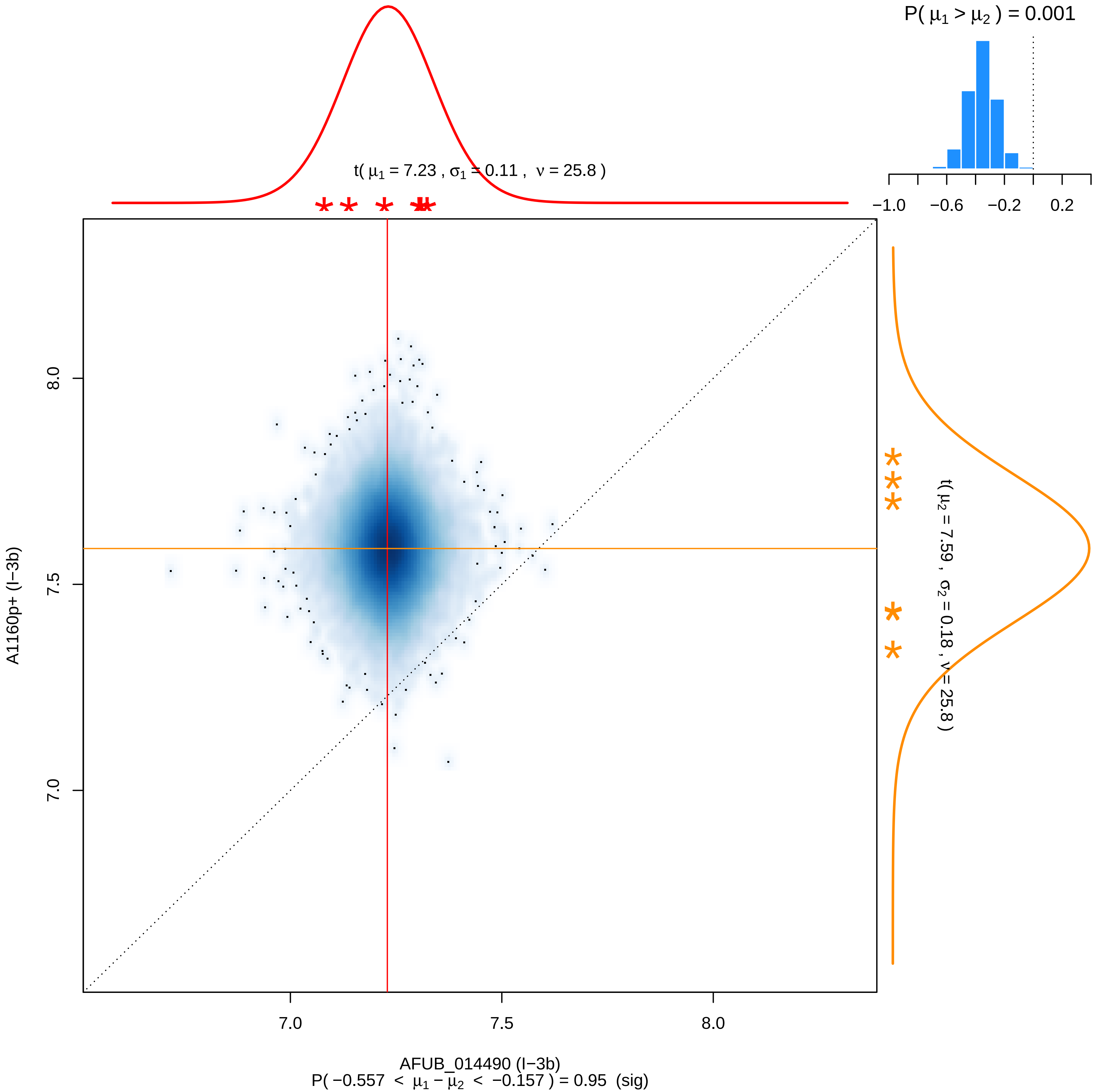

P( μ1 > μ2 ) = 0.001
−1.0
−0.6
−0.2
0.2
t( μ1 = 7.23 , σ1 = 0.11 , ν = 25.8 )
t( μ2 = 7.59 , σ2 = 0.18 , ν = 25.8 )
A1160p+ (I−3b)
8.0
7.5
7.0
7.0
7.5
8.0
AFUB_014490 (I−3b)
P( −0.557 < μ1 − μ2 < −0.157 ) = 0.95 (sig)

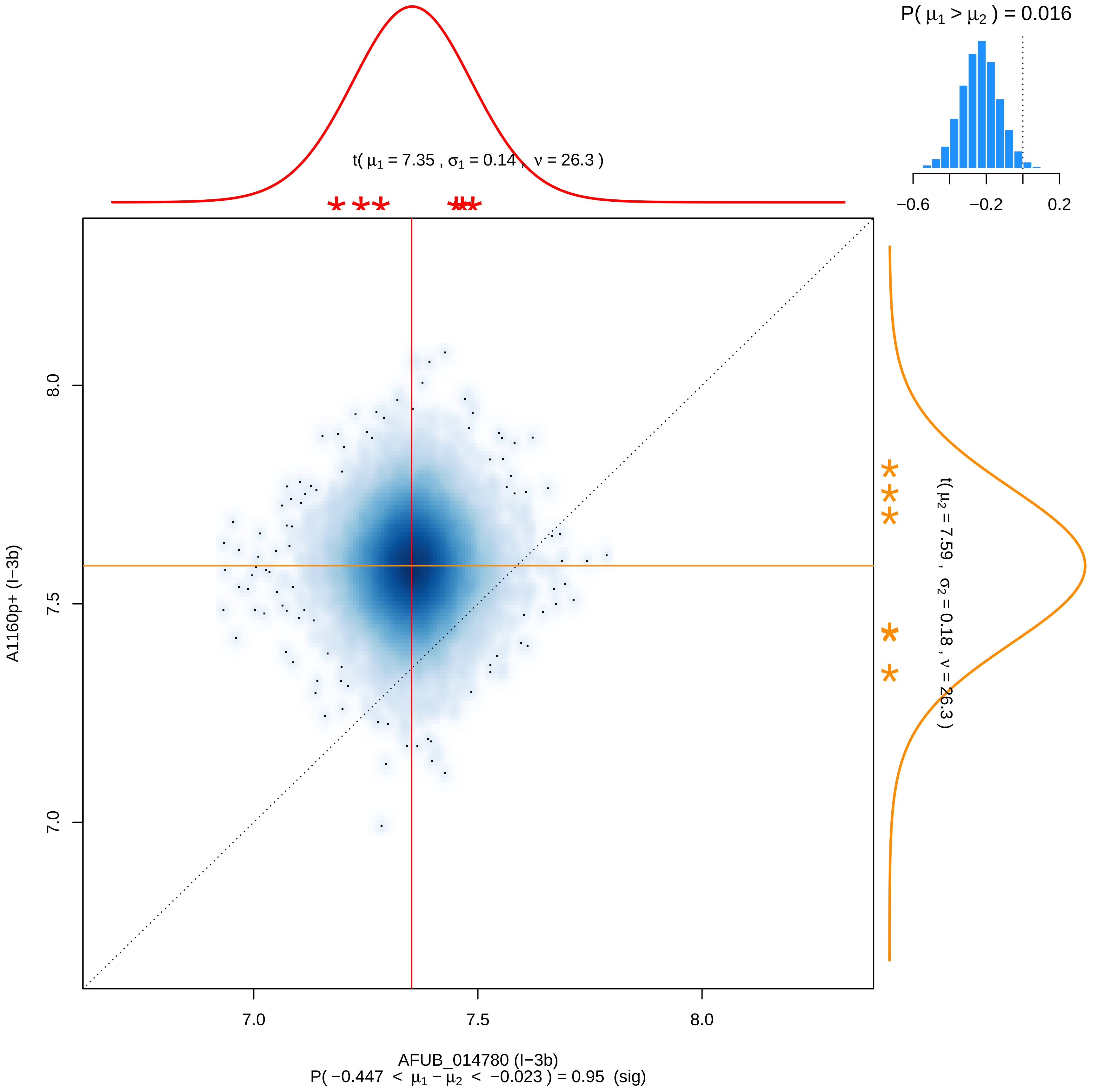

P( μ1 > μ2 ) = 0.016
−0.6
−0.2
0.2
t( μ1 = 7.35 , σ1 = 0.14 , ν = 26.3 )
t( μ2 = 7.59 , σ2 = 0.18 , ν = 26.3 )
A1160p+ (I−3b)
8.0
7.5
7.0
AFUB_014780 (I−3b)
P( −0.447 < μ1 − μ2 < −0.023 ) = 0.95 (sig)

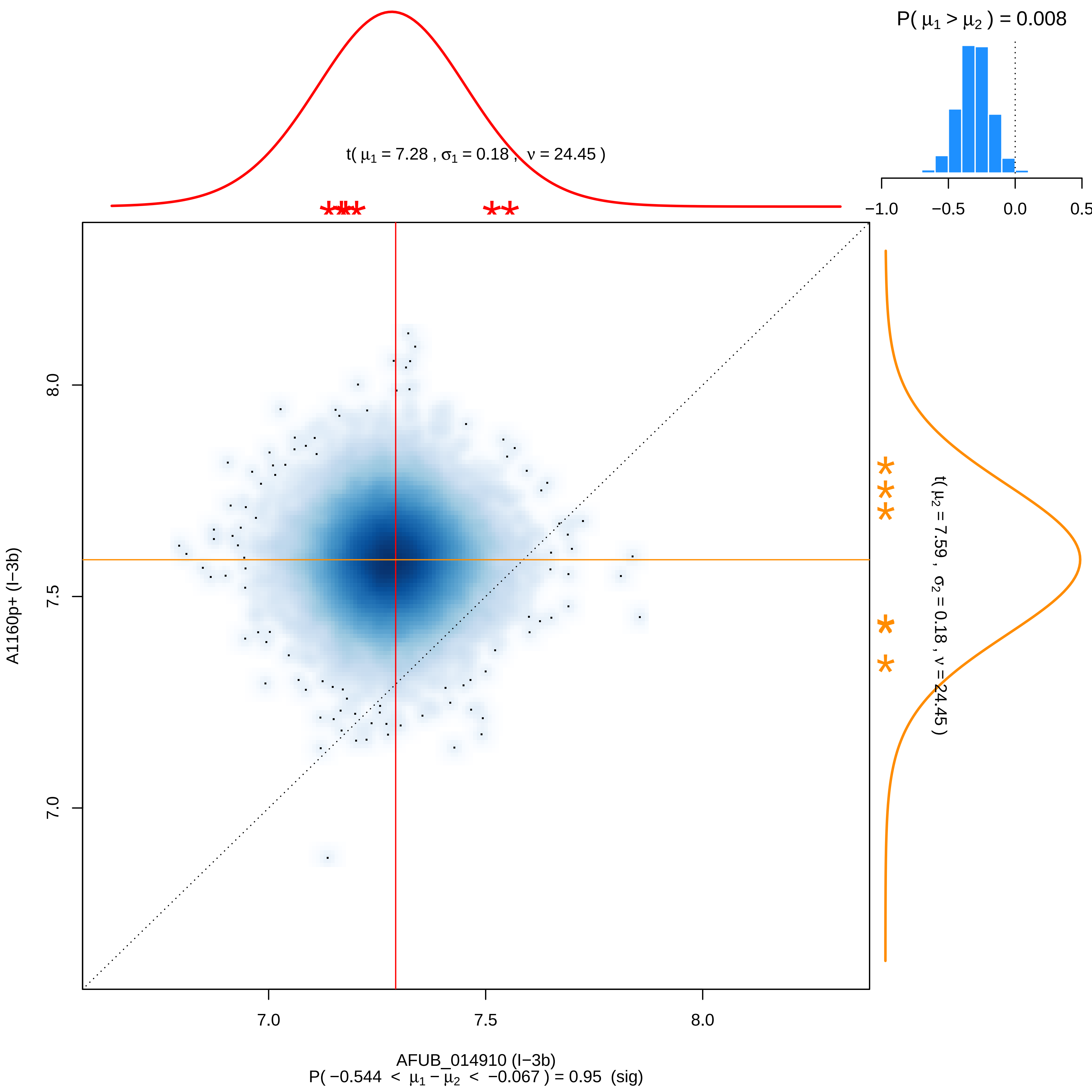

P( μ1 > μ2 ) = 0.008
−1.0
−0.5
0.0
0.5
t( μ1 = 7.28 , σ1 = 0.18 , ν = 24.45 )
t( μ2 = 7.59 , σ2 = 0.18 , ν = 24.45 )
8.0
7.5
7.0
A1160p+ (I−3b)
7.0
7.5
8.0
AFUB_014910 (I−3b)
P( −0.544 < μ1 − μ2 < −0.067 ) = 0.95 (sig)

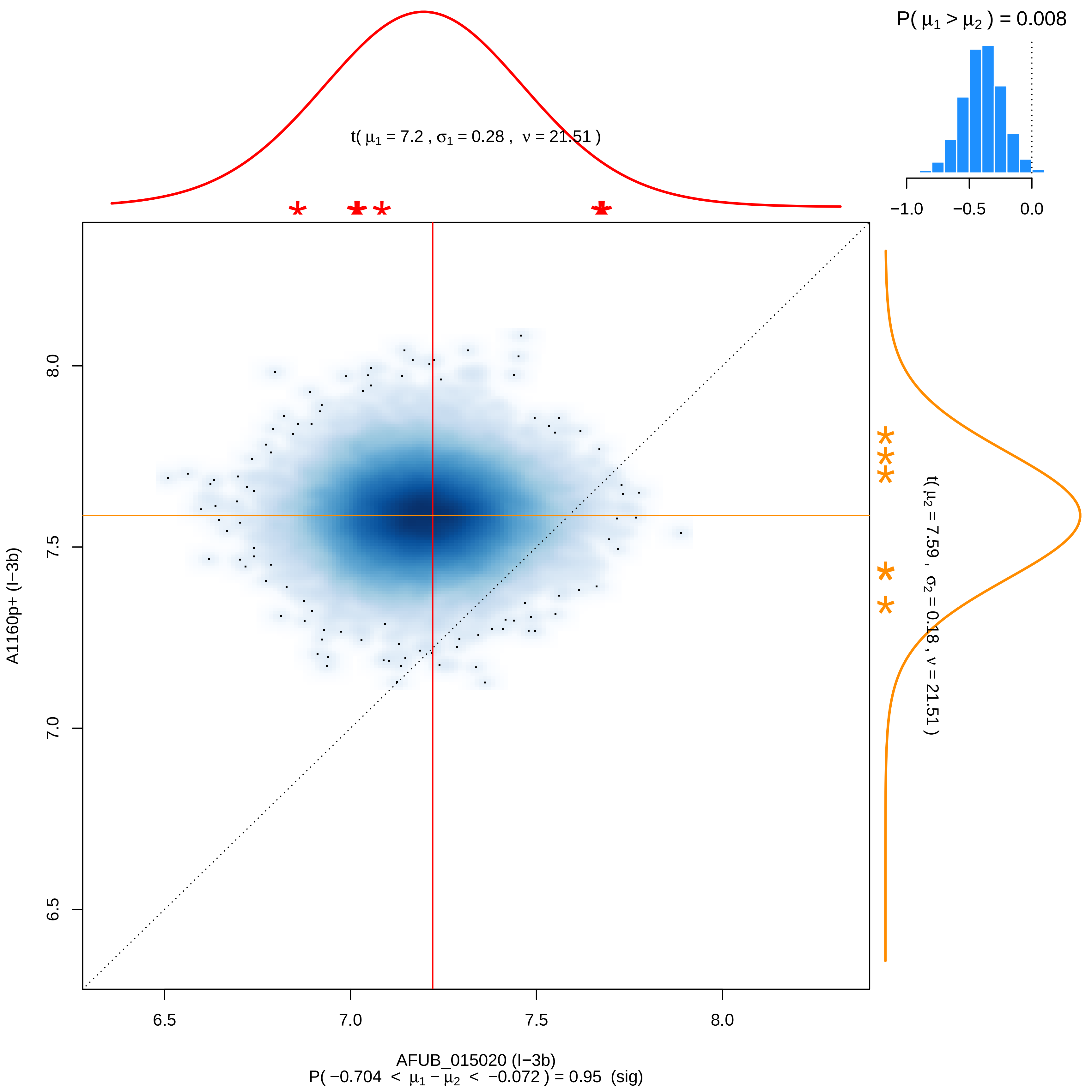

P( μ1 > μ2 ) = 0.008
−1.0
−0.5
0.0
t( μ1 = 7.2 , σ1 = 0.28 , ν = 21.51 )
t( μ2 = 7.59 , σ2 = 0.18 , ν = 21.51 )
A1160p+ (I−3b)
8.0
7.5
7.0
6.5
6.5
7.0
7.5
8.0
AFUB_015020 (I−3b)
P( −0.704 < μ1 − μ2 < −0.072 ) = 0.95 (sig)

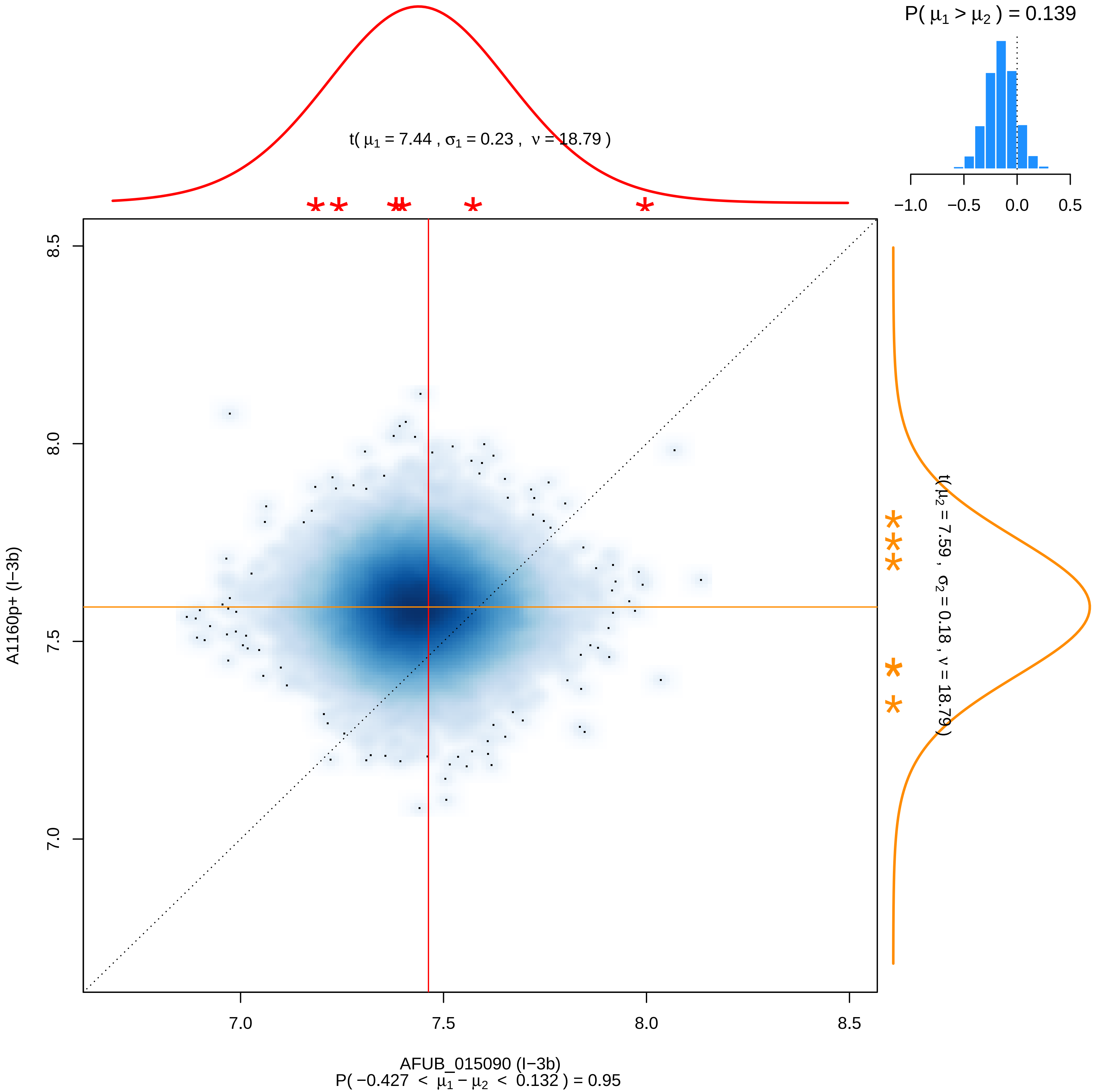

P( μ1 > μ2 ) = 0.139
−1.0 −0.5 0.0 0.5
t( μ1 = 7.44 , σ1 = 0.23 , ν = 18.79 )
t( μ2 = 7.59 , σ2 = 0.18 , ν = 18.79 )
8.5
8.0
7.5
7.0
A1160p+ (I−3b)
7.0 7.5 8.0 8.5
AFUB_015090 (I−3b)
P( −0.427 < μ1 − μ2 < 0.132 ) = 0.95

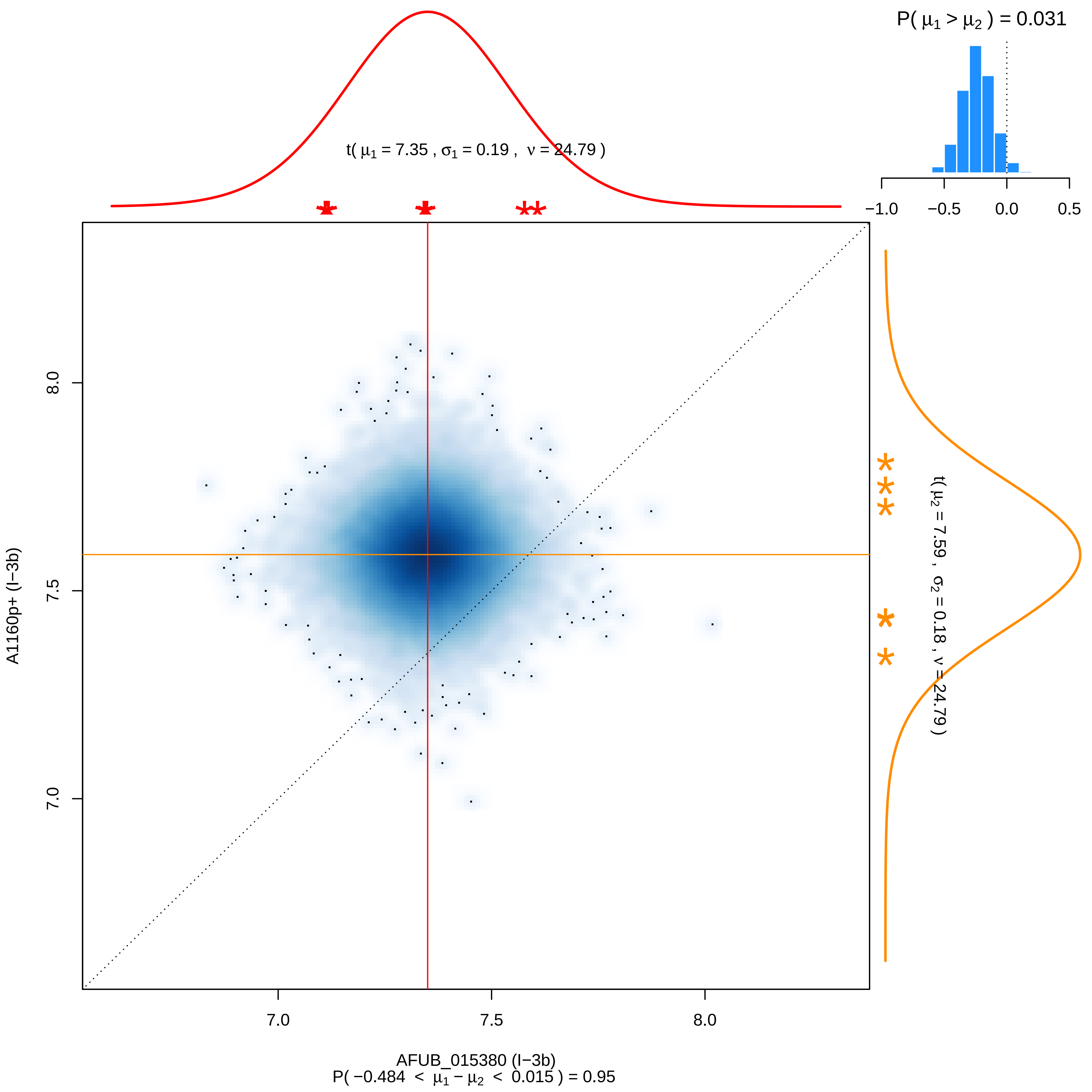

P( μ1 > μ2 ) = 0.031
−1.0
−0.5
0.0
0.5
t( μ1 = 7.35 , σ1 = 0.19 , ν = 24.79 )
t( μ2 = 7.59 , σ2 = 0.18 , ν = 24.79 )
8.0
7.5
7.0
A1160p+ (I−3b)
7.0
7.5
8.0
AFUB_015380 (I−3b)
P( −0.484 < μ1 − μ2 < 0.015 ) = 0.95

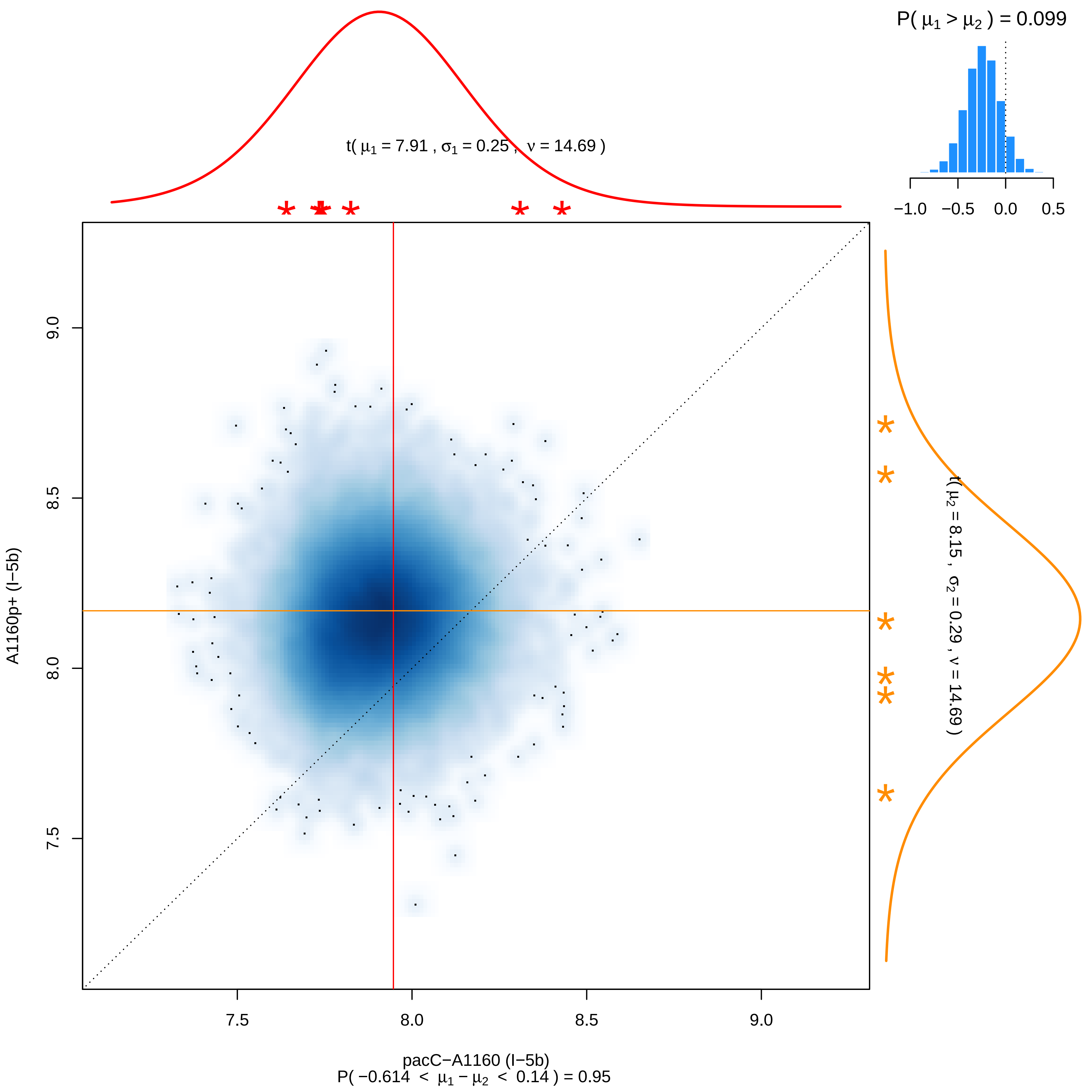

P( μ1 > μ2 ) = 0.099
−1.0 −0.5 0.0 0.5
t( μ1 = 7.91 , σ1 = 0.25 , ν = 14.69 )
t( μ2 = 8.15 , σ2 = 0.29 , ν = 14.69 )
A1160p+ (I−5b)
9.0
8.5
8.0
7.5
7.5 8.0 8.5 9.0
pacC−A1160 (I−5b)
P( −0.614 < μ1 − μ2 < 0.14 ) = 0.95

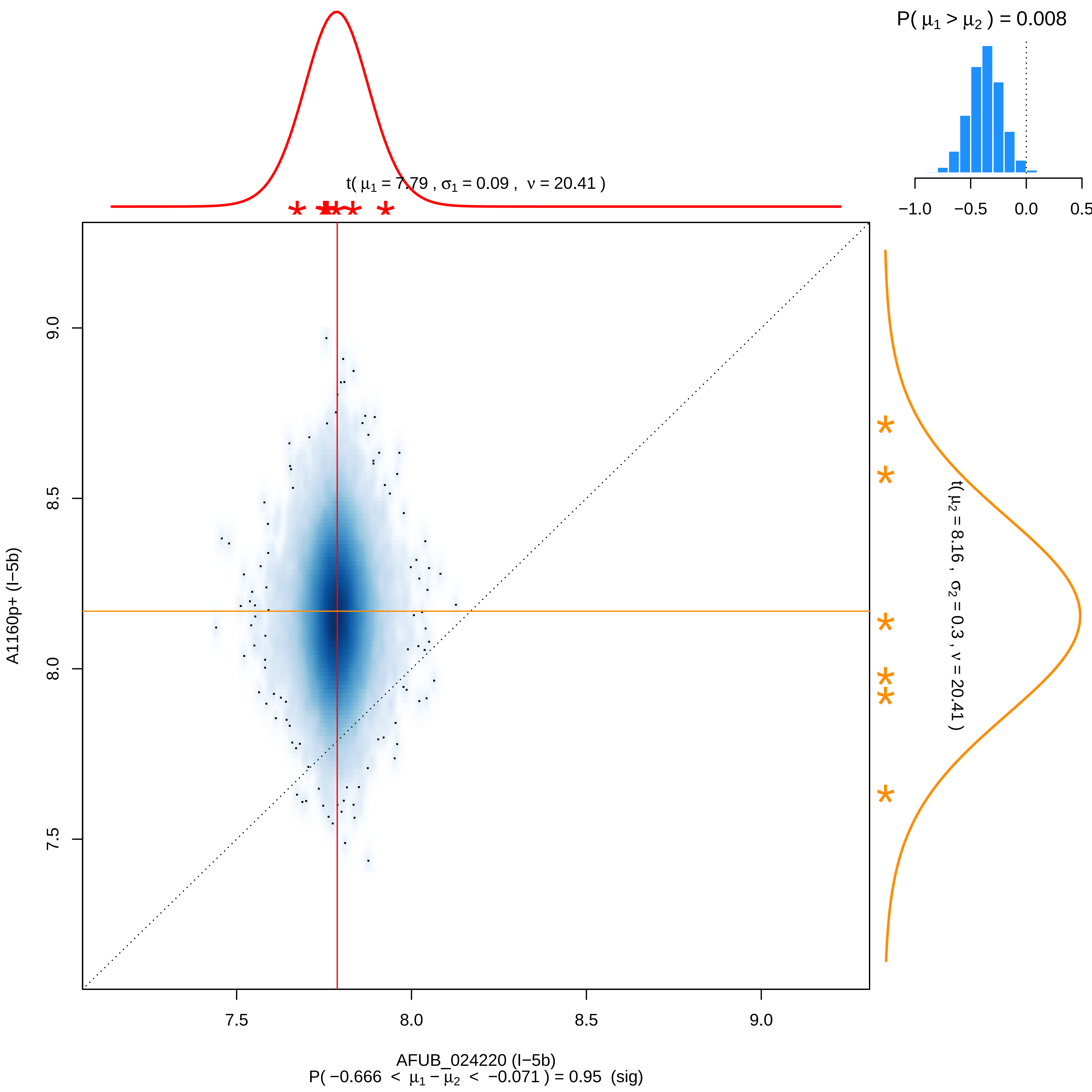

P( $\mu_1 > \mu_2$ ) = 0.008
−1.0
−0.5
0.0
0.5
t( $\mu_1$ = 7.79 , $\sigma_1$ = 0.09 , $\nu$ = 20.41 )
t( $\mu_2$ = 8.16 , $\sigma_2$ = 0.3 , $\nu$ = 20.41 )
9.0
8.5
8.0
7.5
A1160p+ (I−5b)
7.5
8.0
8.5
9.0
AFUB_024220 (I−5b)
P( −0.666 < $\mu_1 - \mu_2$ < −0.071 ) = 0.95 (sig)

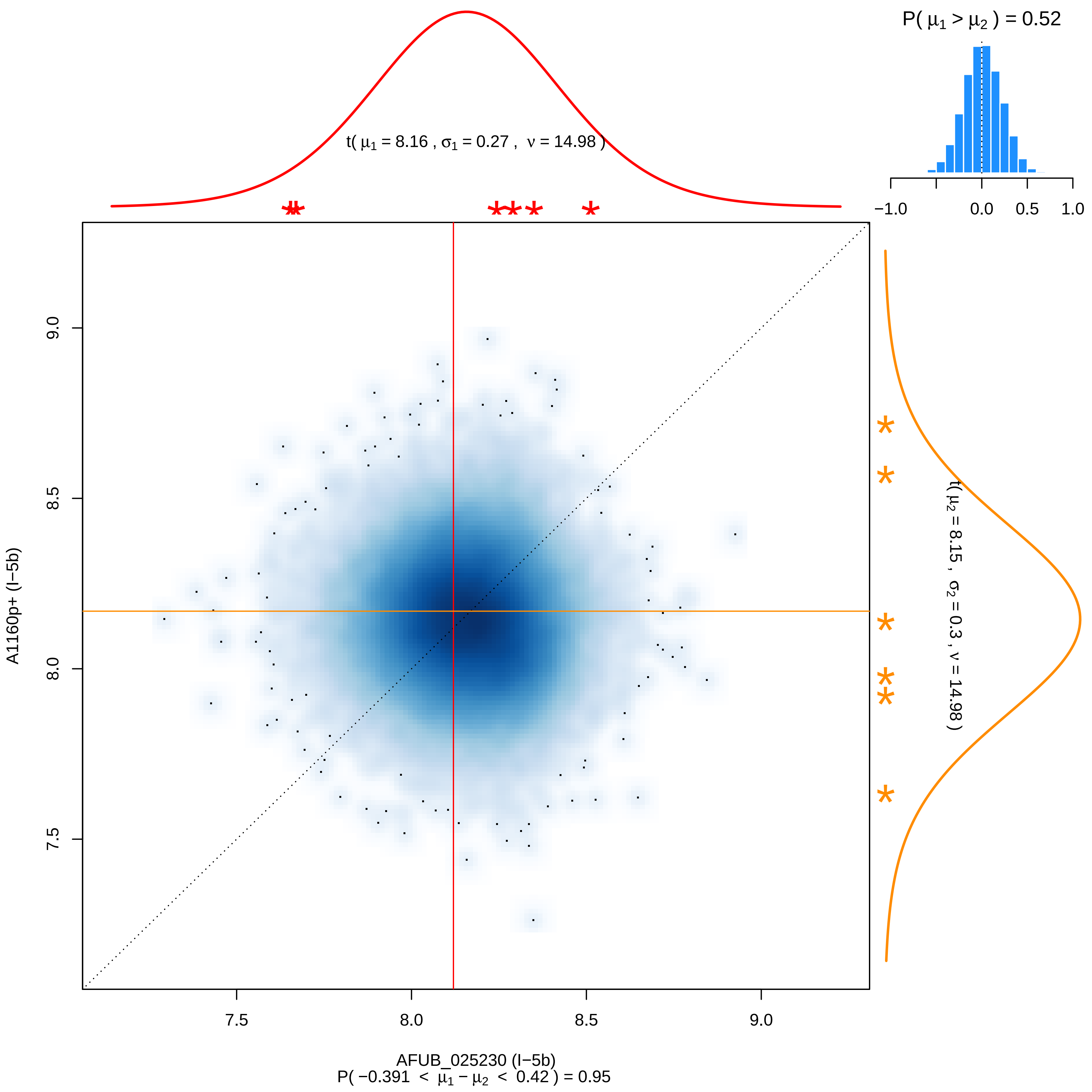

P( μ1 > μ2 ) = 0.52
−1.0
0.0
0.5
1.0
t( μ1 = 8.16 , σ1 = 0.27 , ν = 14.98 )
t( μ2 = 8.15 , σ2 = 0.3 , ν = 14.98 )
A1160p+ (I−5b)
9.0
8.5
8.0
7.5
7.5
8.0
8.5
9.0
AFUB_025230 (I−5b)
P( −0.391 < μ1 − μ2 < 0.42 ) = 0.95

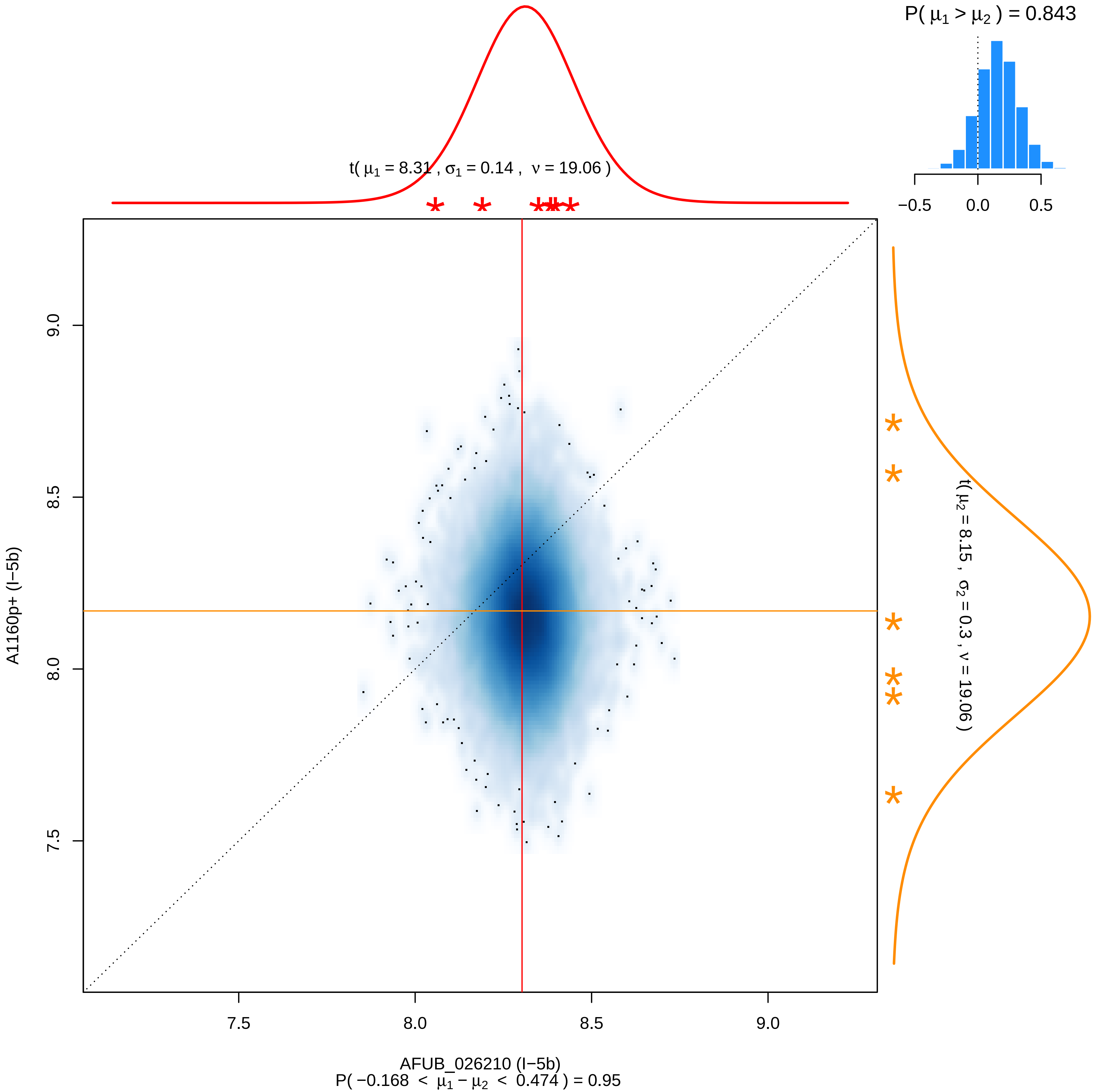

P( μ1 > μ2 ) = 0.843
−0.5
0.0
0.5
t( μ1 = 8.31 , σ1 = 0.14 , ν = 19.06 )
9.0
8.5
8.0
7.5
A1160p+ (I−5b)
t( μ2 = 8.15 , σ2 = 0.3 , ν = 19.06 )
7.5
8.0
8.5
9.0
AFUB_026210 (I−5b)
P( −0.168 < μ1 − μ2 < 0.474 ) = 0.95

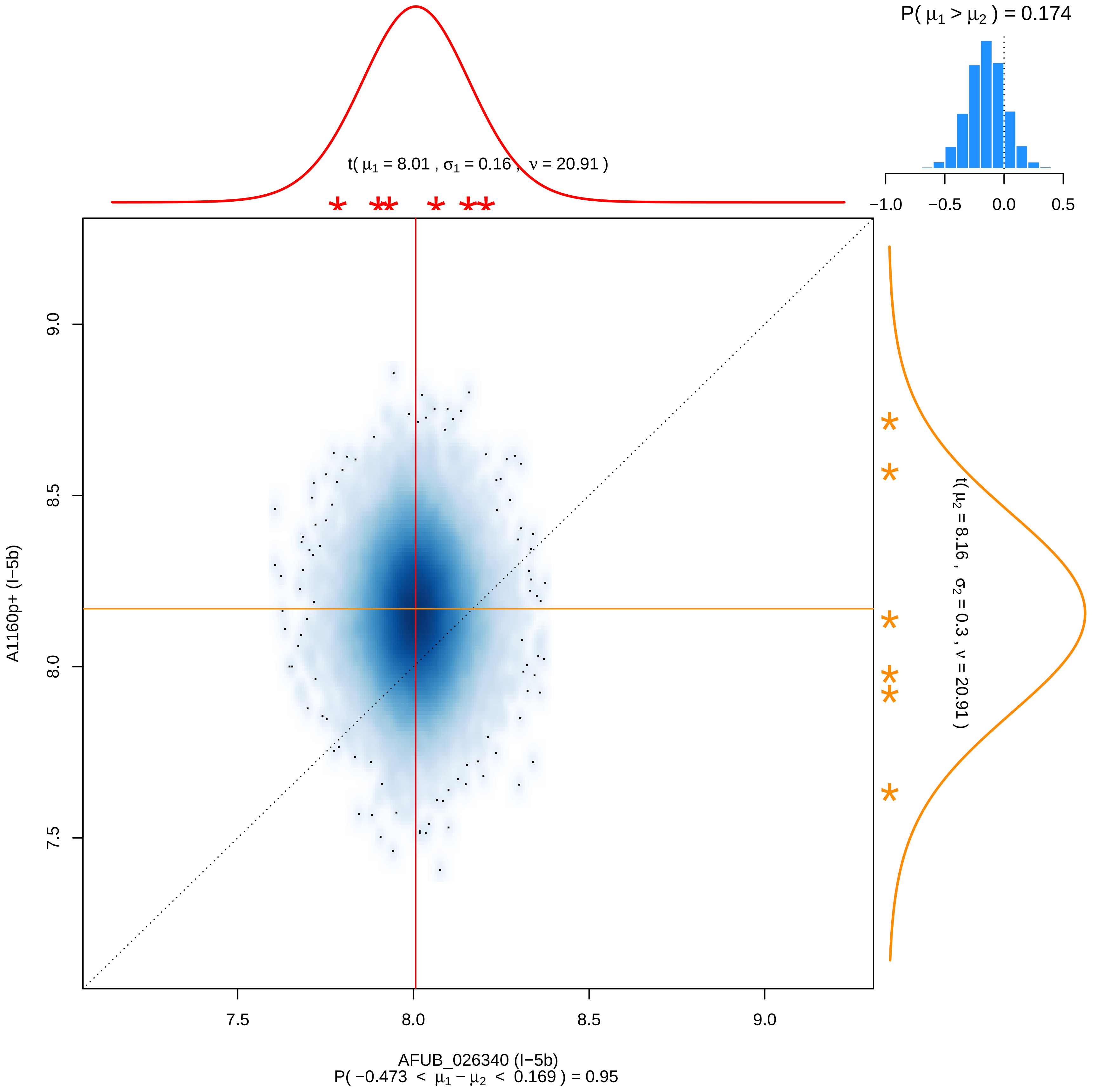

P( μ1 > μ2 ) = 0.174
t( μ1 = 8.01 , σ1 = 0.16 , ν = 20.91 )
−1.0
−0.5
0.0
0.5
9.0
8.5
8.0
7.5
A1160p+ (I−5b)
t( μ2 = 8.16 , σ2 = 0.3 , ν = 20.91 )
7.5
8.0
8.5
9.0
AFUB_026340 (I−5b)
P( −0.473 < μ1 − μ2 < 0.169 ) = 0.95

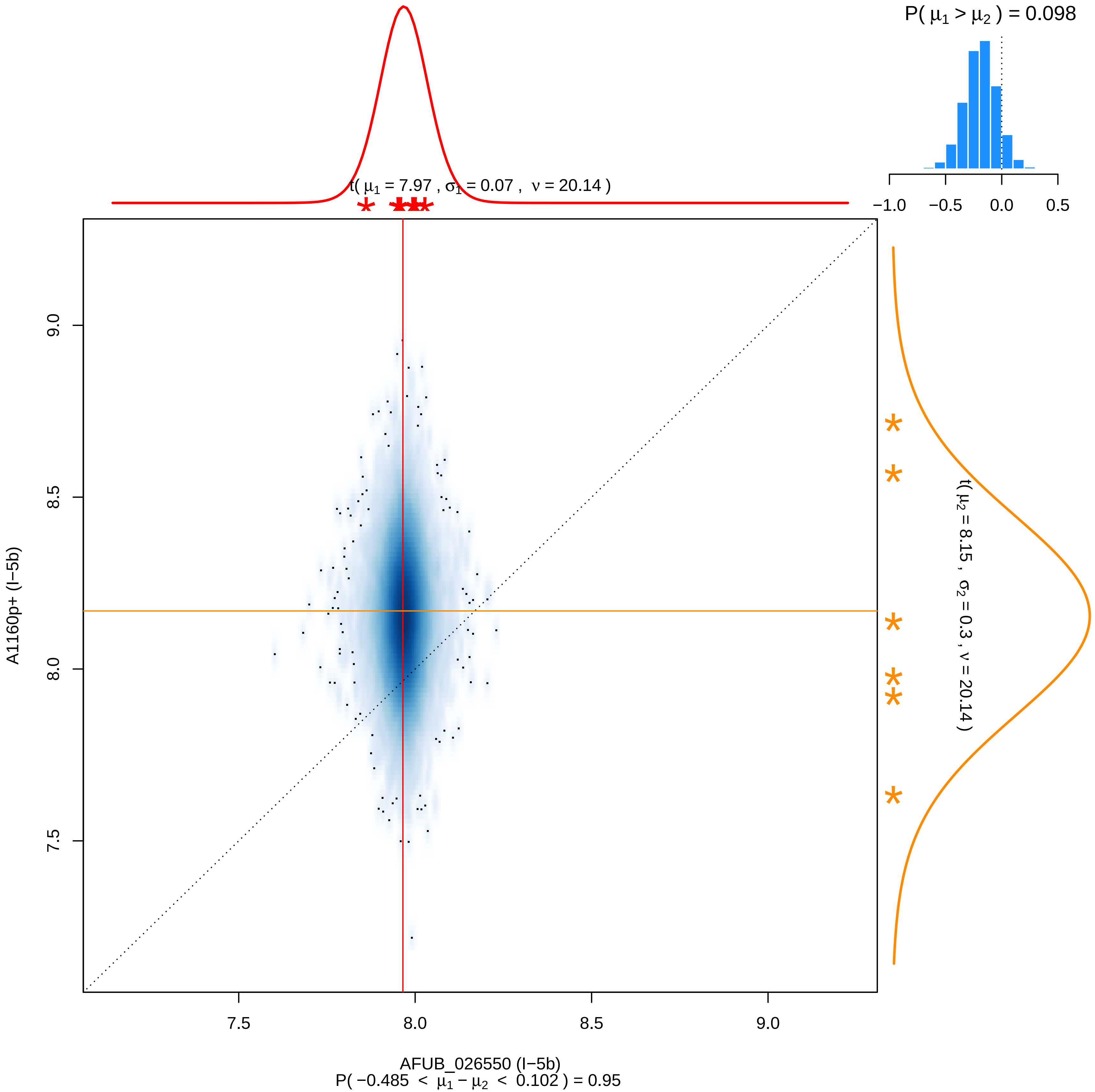

P( μ1 > μ2 ) = 0.098
−1.0
−0.5
0.0
0.5
t( μ1 = 7.97 , σ1 = 0.07 , ν = 20.14 )
9.0
8.5
8.0
7.5
A1160p+ (I−5b)
t( μ2 = 8.15 , σ2 = 0.3 , ν = 20.14 )
7.5
8.0
8.5
9.0
AFUB_026550 (I−5b)
P( −0.485 < μ1 − μ2 < 0.102 ) = 0.95

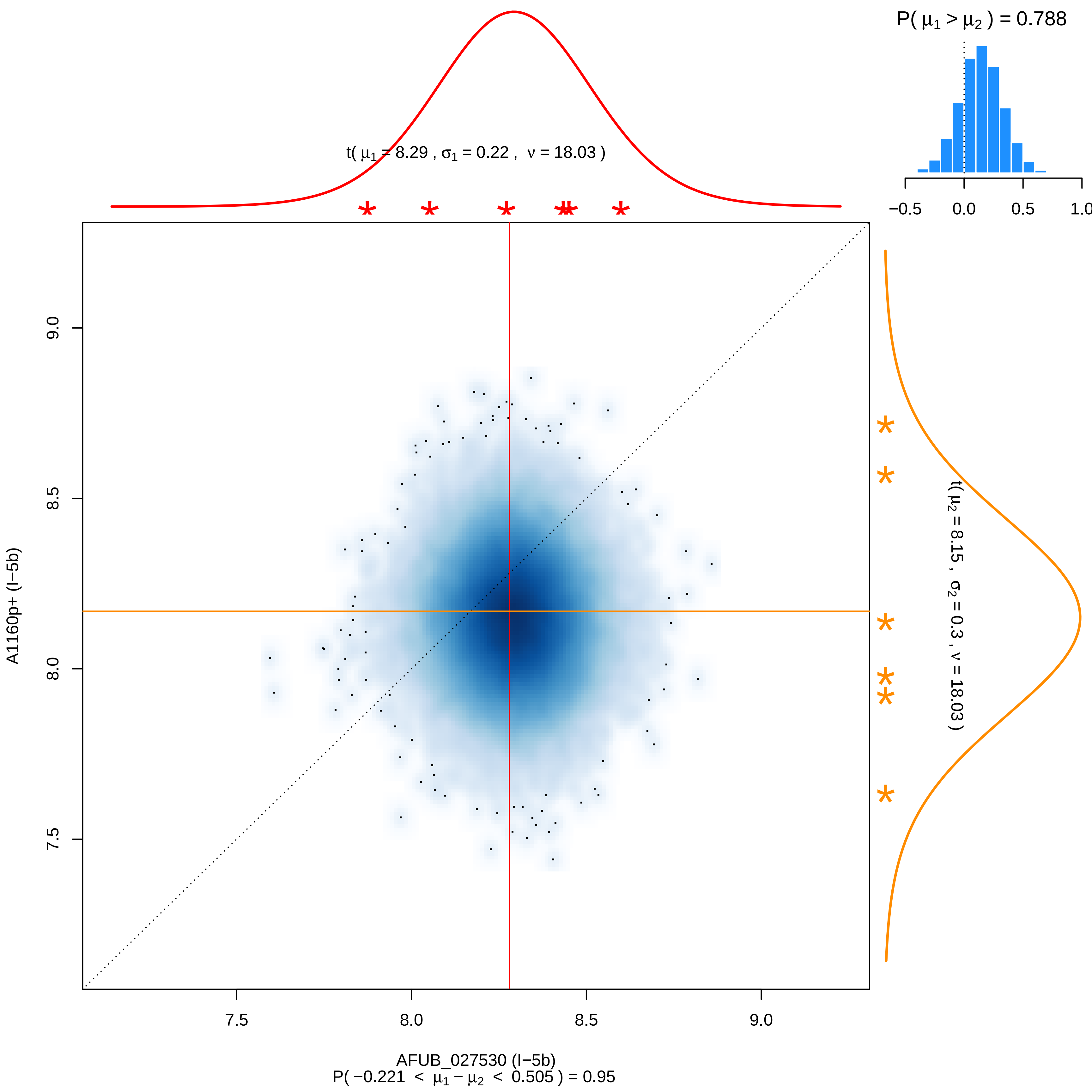

P( μ1 > μ2 ) = 0.788
−0.5
0.0
0.5
1.0
t( μ1 = 8.29 , σ1 = 0.22 , ν = 18.03 )
t( μ2 = 8.15 , σ2 = 0.3 , ν = 18.03 )
A1160p+ (I−5b)
9.0
8.5
8.0
7.5
7.5
8.0
8.5
9.0
AFUB_027530 (I−5b)
P( −0.221 < μ1 − μ2 < 0.505 ) = 0.95

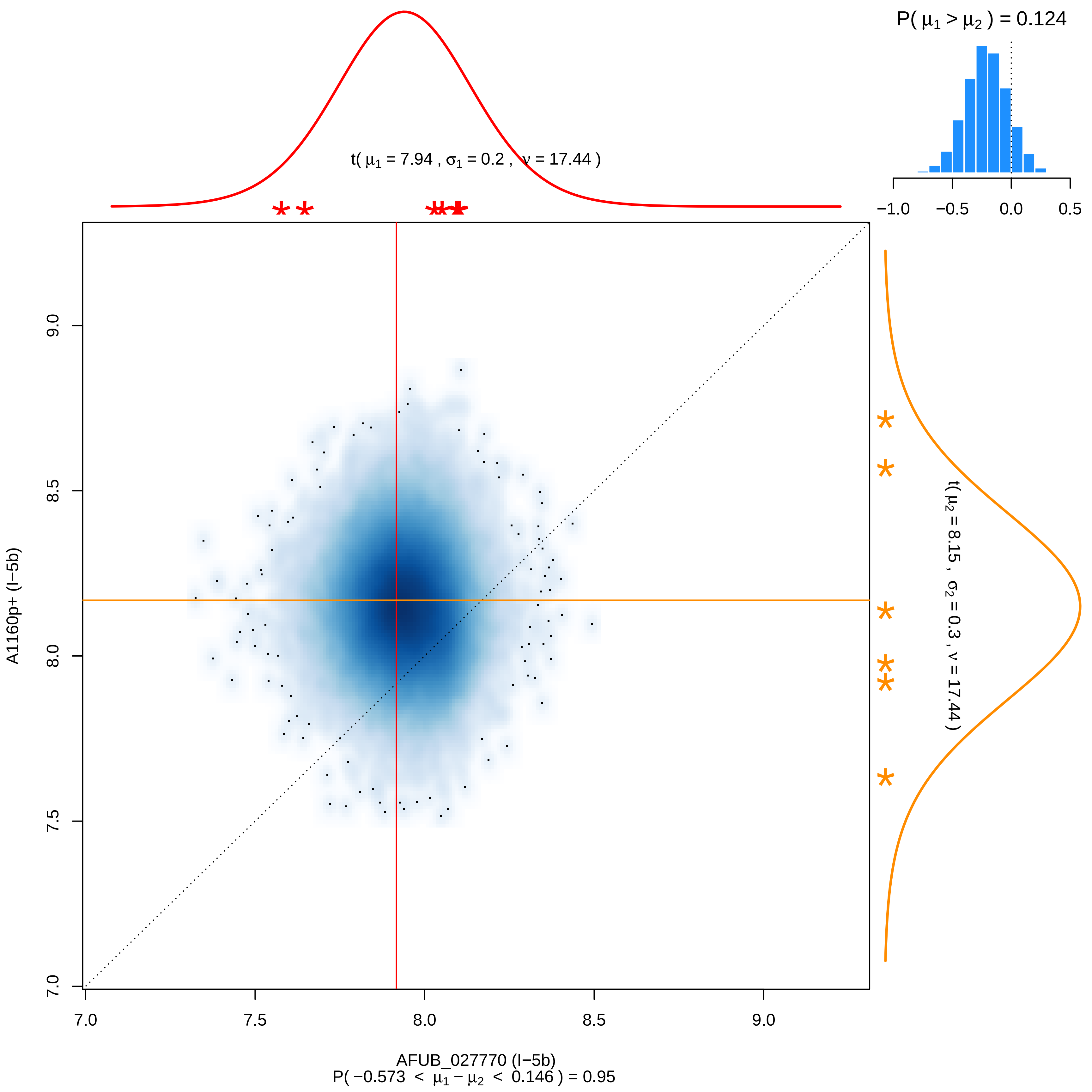

P( μ1 > μ2 ) = 0.124
−1.0
−0.5
0.0
0.5
t( μ1 = 7.94 , σ1 = 0.2 , ν = 17.44 )
t( μ2 = 8.15 , σ2 = 0.3 , ν = 17.44 )
A1160p+ (I−5b)
9.0
8.5
8.0
7.5
7.0
7.0
7.5
8.0
8.5
9.0
AFUB_027770 (I−5b)
P( −0.573 < μ1 − μ2 < 0.146 ) = 0.95

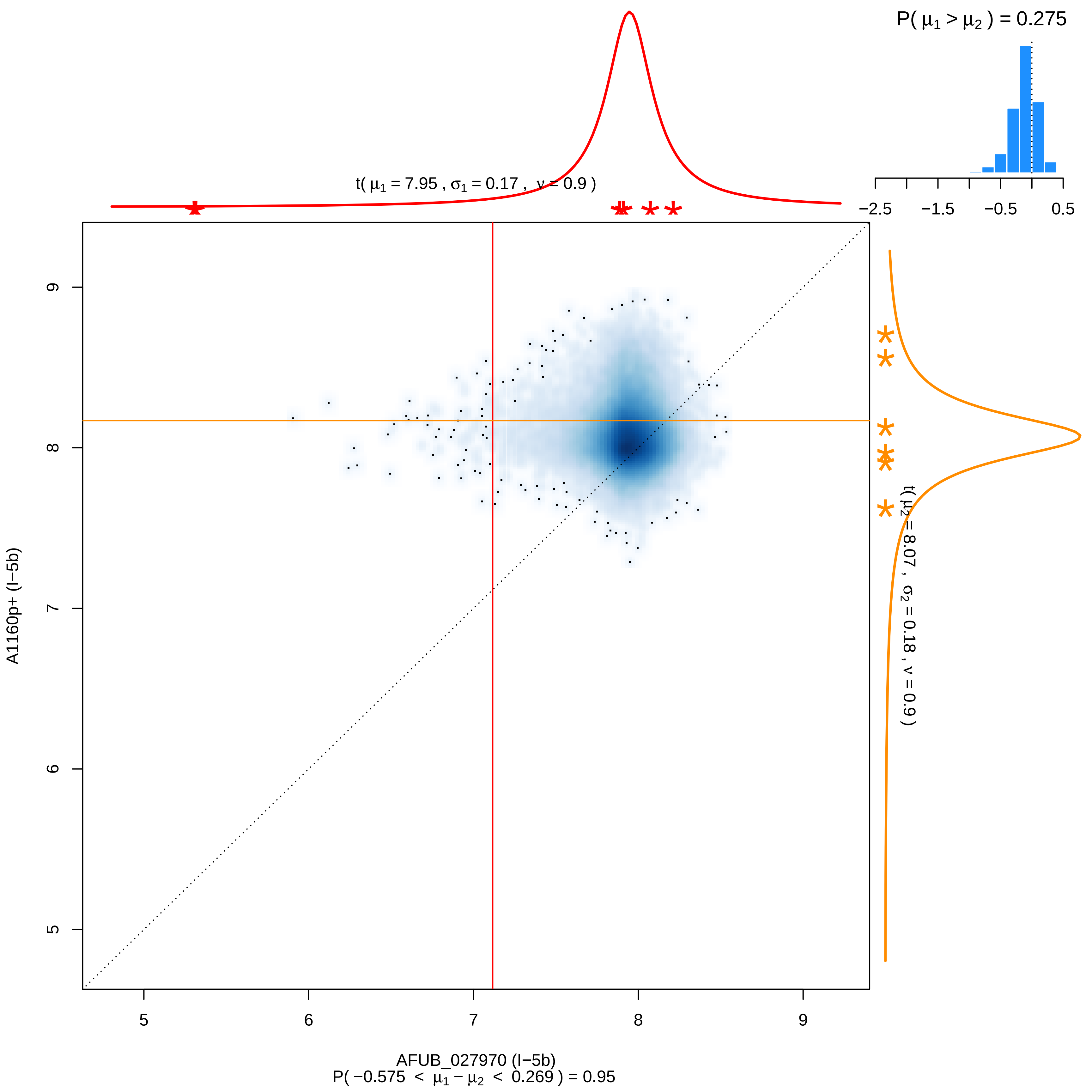

P( $\mu_1 > \mu_2$ ) = 0.275
t( $\mu_1$ = 7.95 , $\sigma_1$ = 0.17 , $\nu$ = 0.9 )
−2.5
−1.5
−0.5
0.5
t( $\mu_2$ = 8.07 , $\sigma_2$ = 0.18 , $\nu$ = 0.9 )
A1160p+ (I−5b)
AFUB_027970 (I−5b)
P( −0.575 < $\mu_1 - \mu_2$ < 0.269 ) = 0.95
5
6
7
8
9

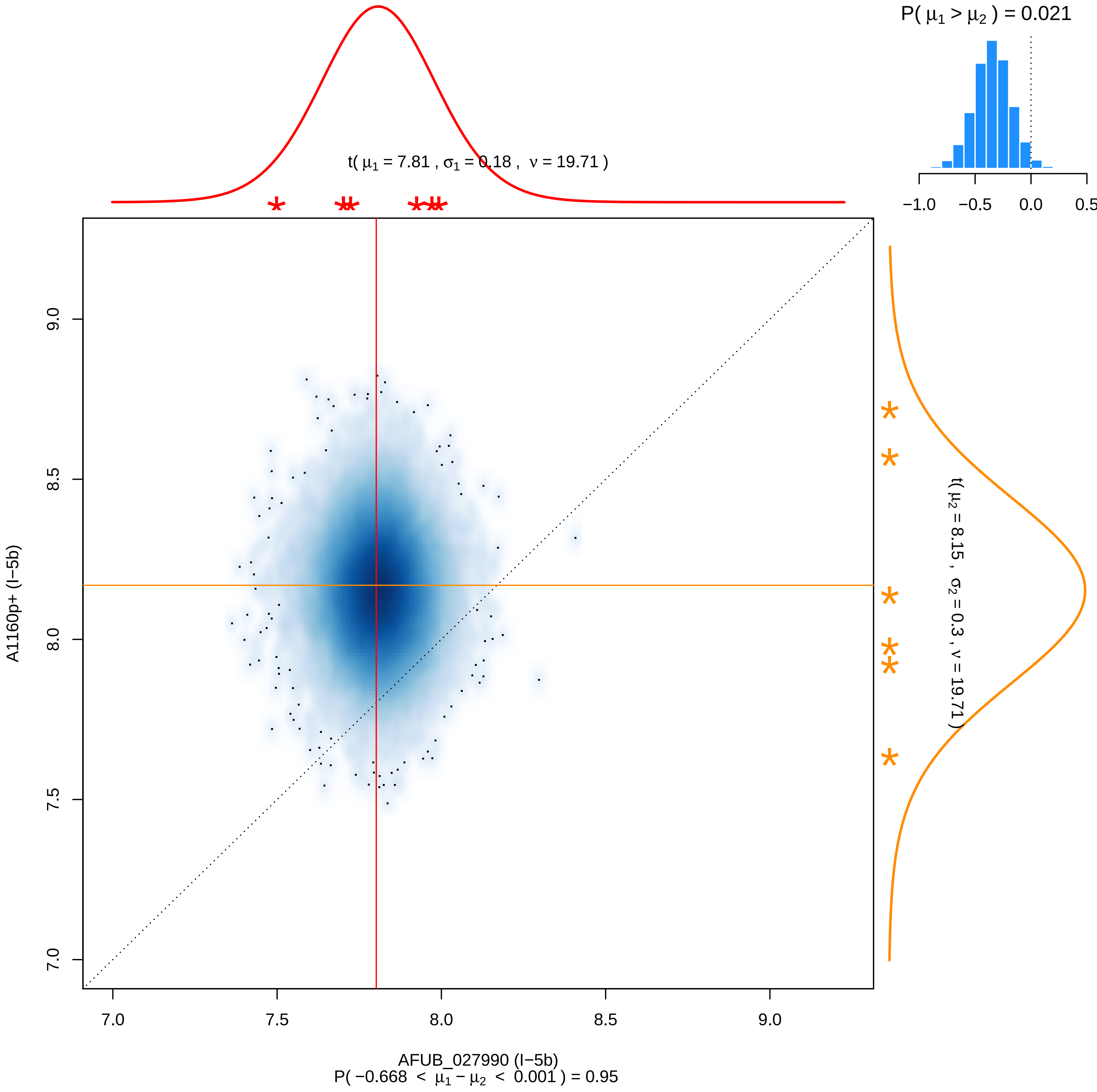

P( μ1 > μ2 ) = 0.021
−1.0 −0.5 0.0 0.5
t( μ1 = 7.81 , σ1 = 0.18 , ν = 19.71 )
t( μ2 = 8.15 , σ2 = 0.3 , ν = 19.71 )
A1160p+ (I−5b)
9.0
8.5
8.0
7.5
7.0
7.0 7.5 8.0 8.5 9.0
AFUB_027990 (I−5b)
P( −0.668 < μ1 − μ2 < 0.001 ) = 0.95

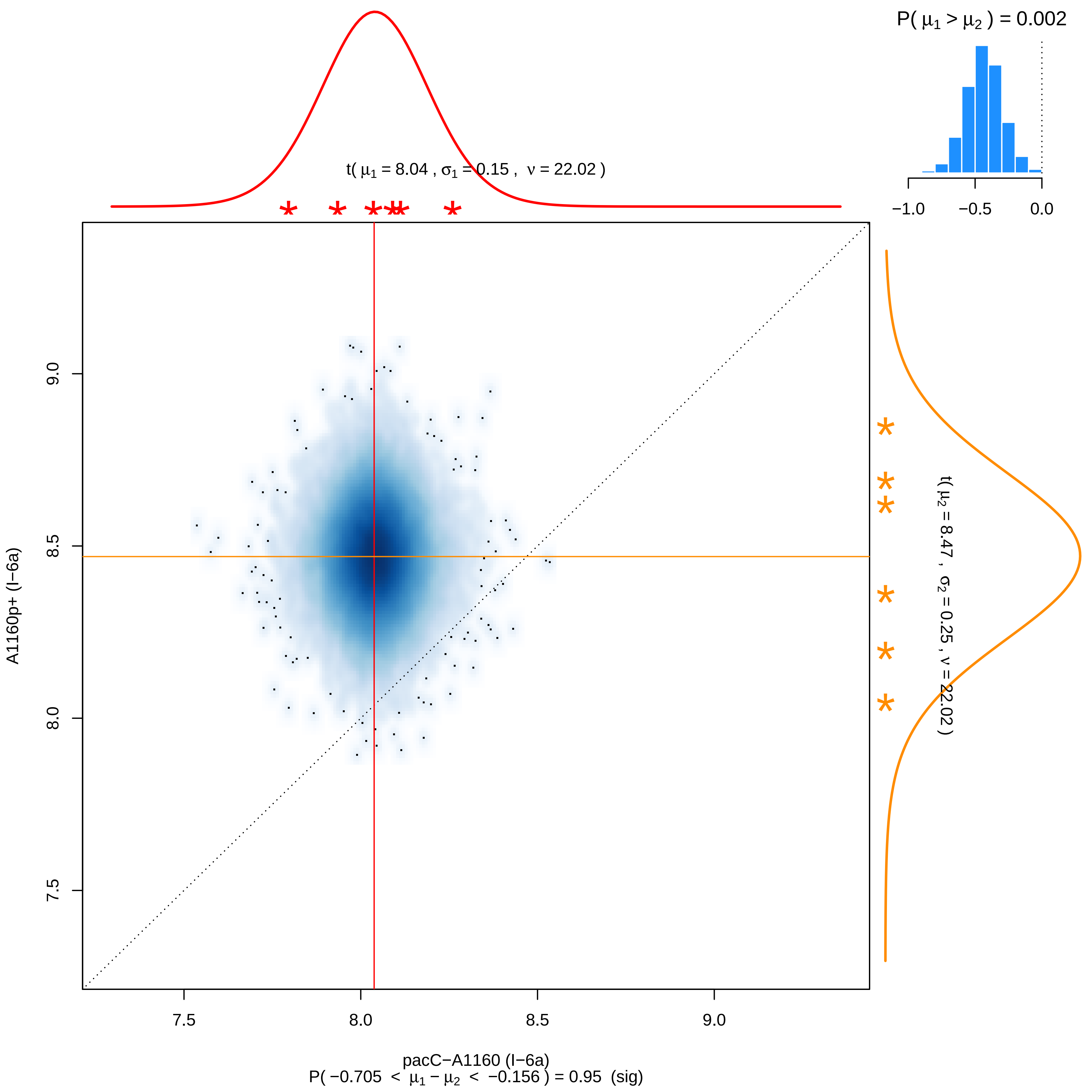

P( μ1 > μ2 ) = 0.002
−1.0
−0.5
0.0
t( μ1 = 8.04 , σ1 = 0.15 , ν = 22.02 )
t( μ2 = 8.47 , σ2 = 0.25 , ν = 22.02 )
9.0
8.5
8.0
7.5
A1160p+ (I−6a)
7.5
8.0
8.5
9.0
pacC−A1160 (I−6a)
P( −0.705 < μ1 − μ2 < −0.156 ) = 0.95 (sig)

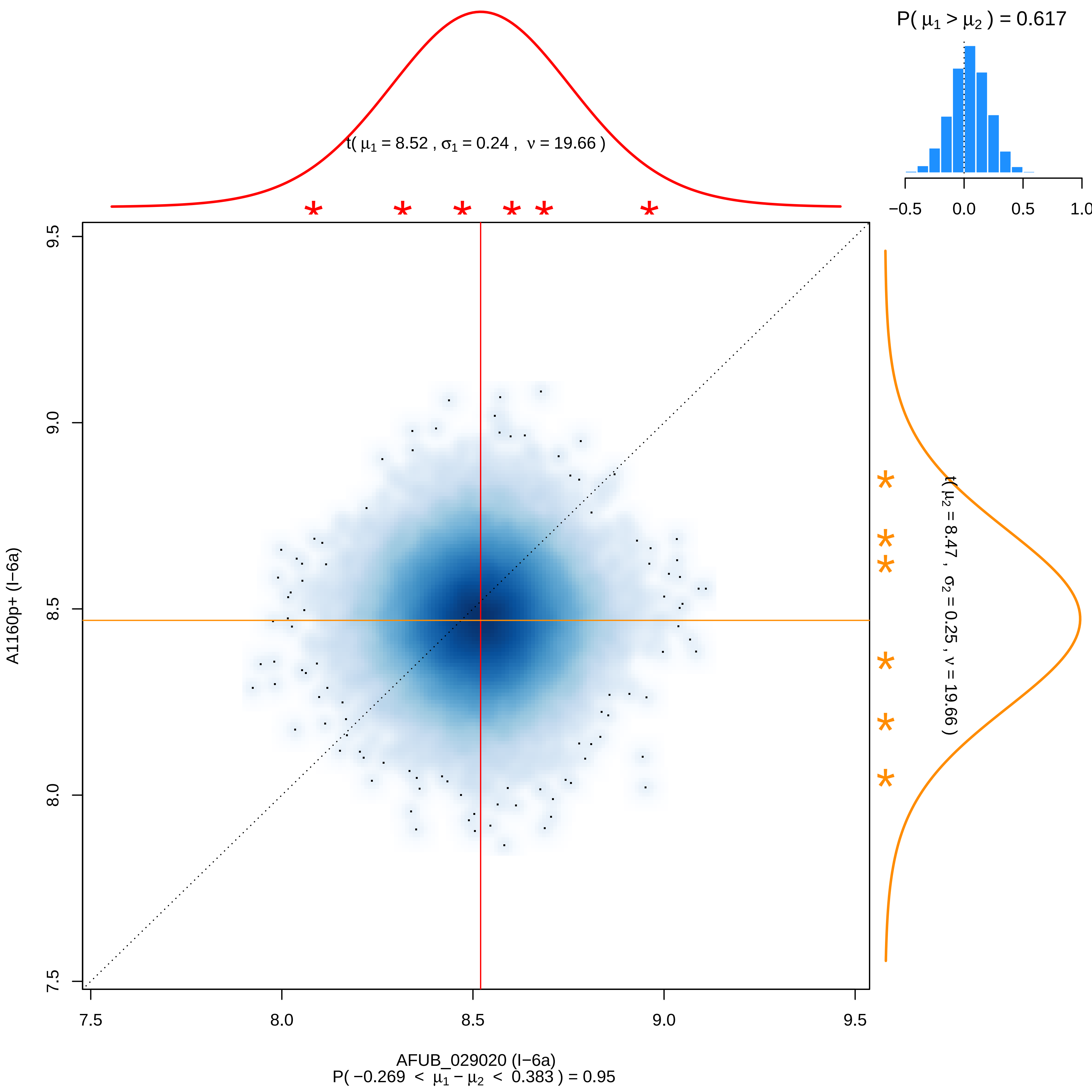

P( μ1 > μ2 ) = 0.617
−0.5
0.0
0.5
1.0
t( μ1 = 8.52 , σ1 = 0.24 , ν = 19.66 )
t( μ2 = 8.47 , σ2 = 0.25 , ν = 19.66 )
7.5
8.0
8.5
9.0
9.5
A1160p+ (I−6a)
AFUB_029020 (I−6a)
P( −0.269 < μ1 − μ2 < 0.383 ) = 0.95

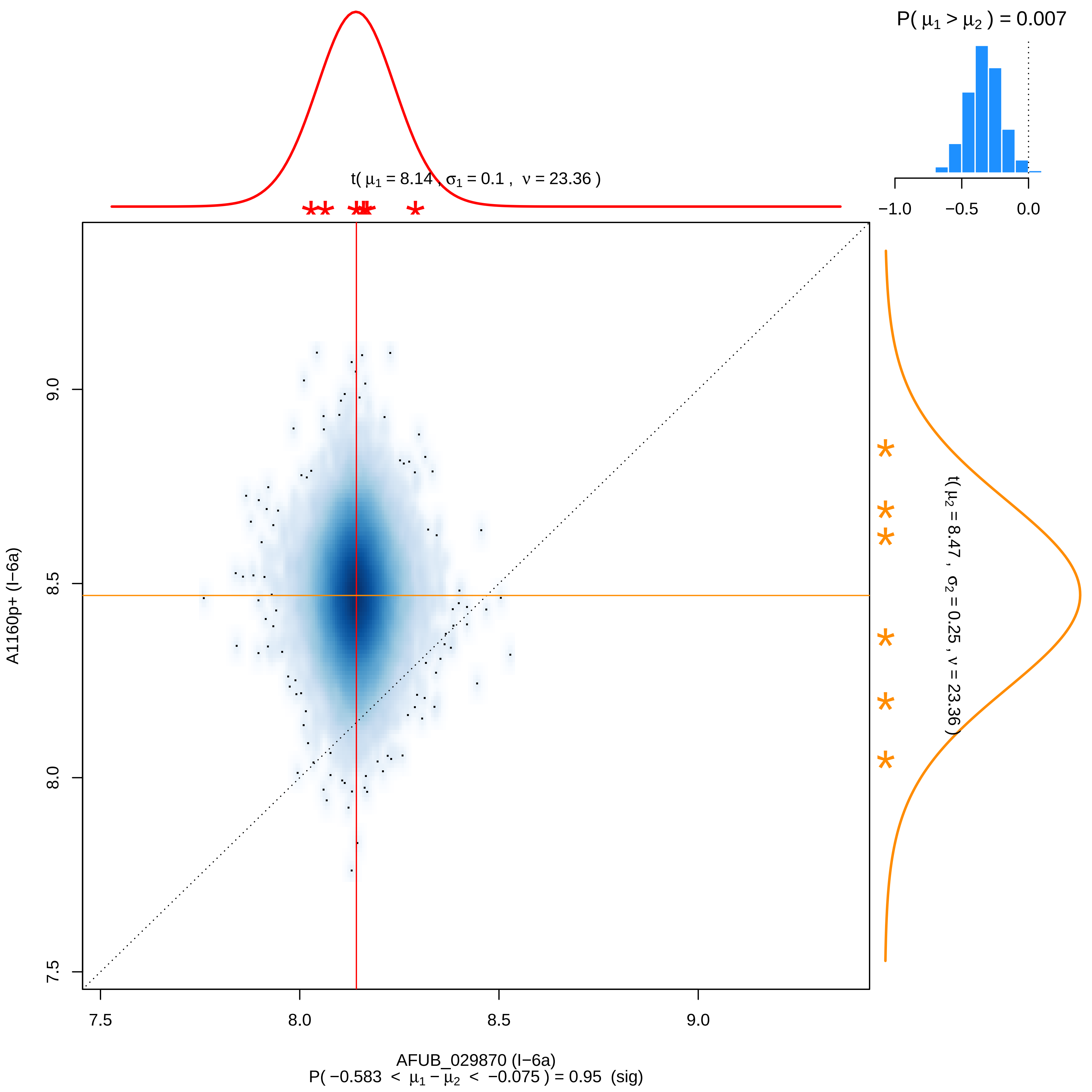

P( μ1 > μ2 ) = 0.007
−1.0
−0.5
0.0
t( μ1 = 8.14 , σ1 = 0.1 , ν = 23.36 )
t( μ2 = 8.47 , σ2 = 0.25 , ν = 23.36 )
A1160p+ (I−6a)
9.0
8.5
8.0
7.5
7.5
8.0
8.5
9.0
AFUB_029870 (I−6a)
P( −0.583 < μ1 − μ2 < −0.075 ) = 0.95 (sig)

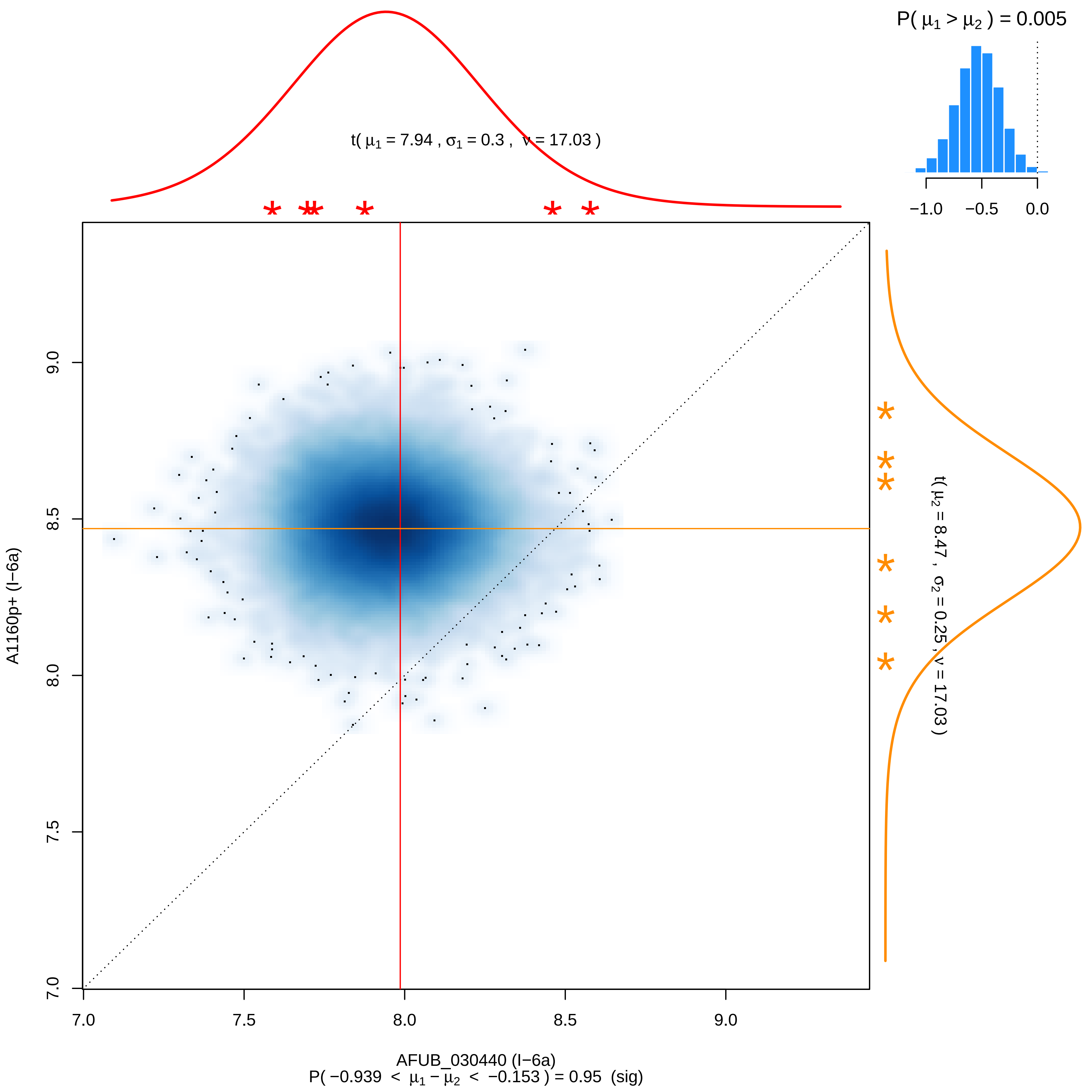

P( μ1 > μ2 ) = 0.005
−1.0
−0.5
0.0
t( μ1 = 7.94 , σ1 = 0.3 , ν = 17.03 )
t( μ2 = 8.47 , σ2 = 0.25 , ν = 17.03 )
A1160p+ (I−6a)
9.0
8.5
8.0
7.5
7.0
7.0
7.5
8.0
8.5
9.0
AFUB_030440 (I−6a)
P( −0.939 < μ1 − μ2 < −0.153 ) = 0.95 (sig)

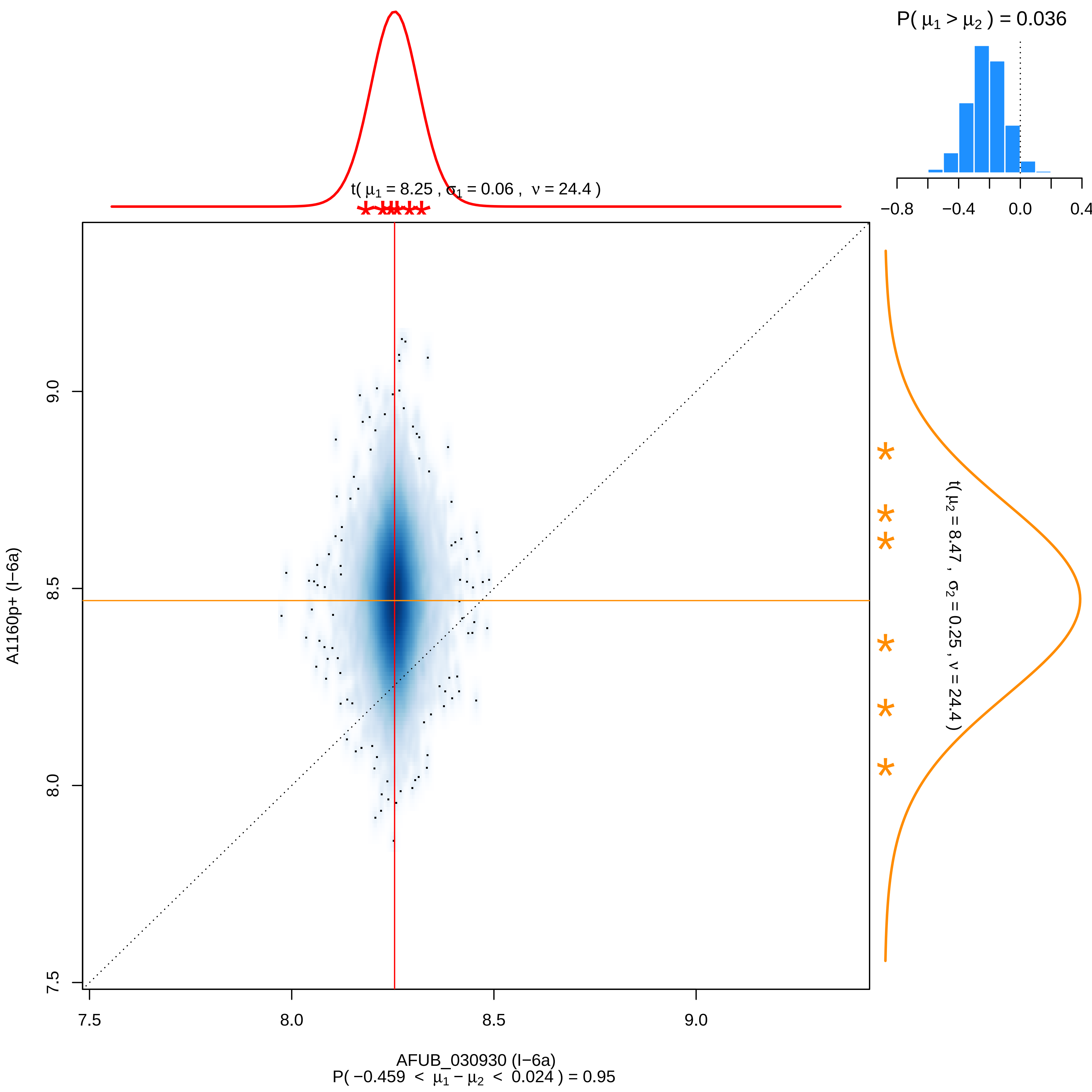

P( μ1 > μ2 ) = 0.036
−0.8
−0.4
0.0
0.4
t( μ1 = 8.25 , σ1 = 0.06 , ν = 24.4 )
t( μ2 = 8.47 , σ2 = 0.25 , ν = 24.4 )
9.0
8.5
8.0
7.5
A1160p+ (I−6a)
7.5
8.0
8.5
9.0
AFUB_030930 (I−6a)
P( −0.459 < μ1 − μ2 < 0.024 ) = 0.95

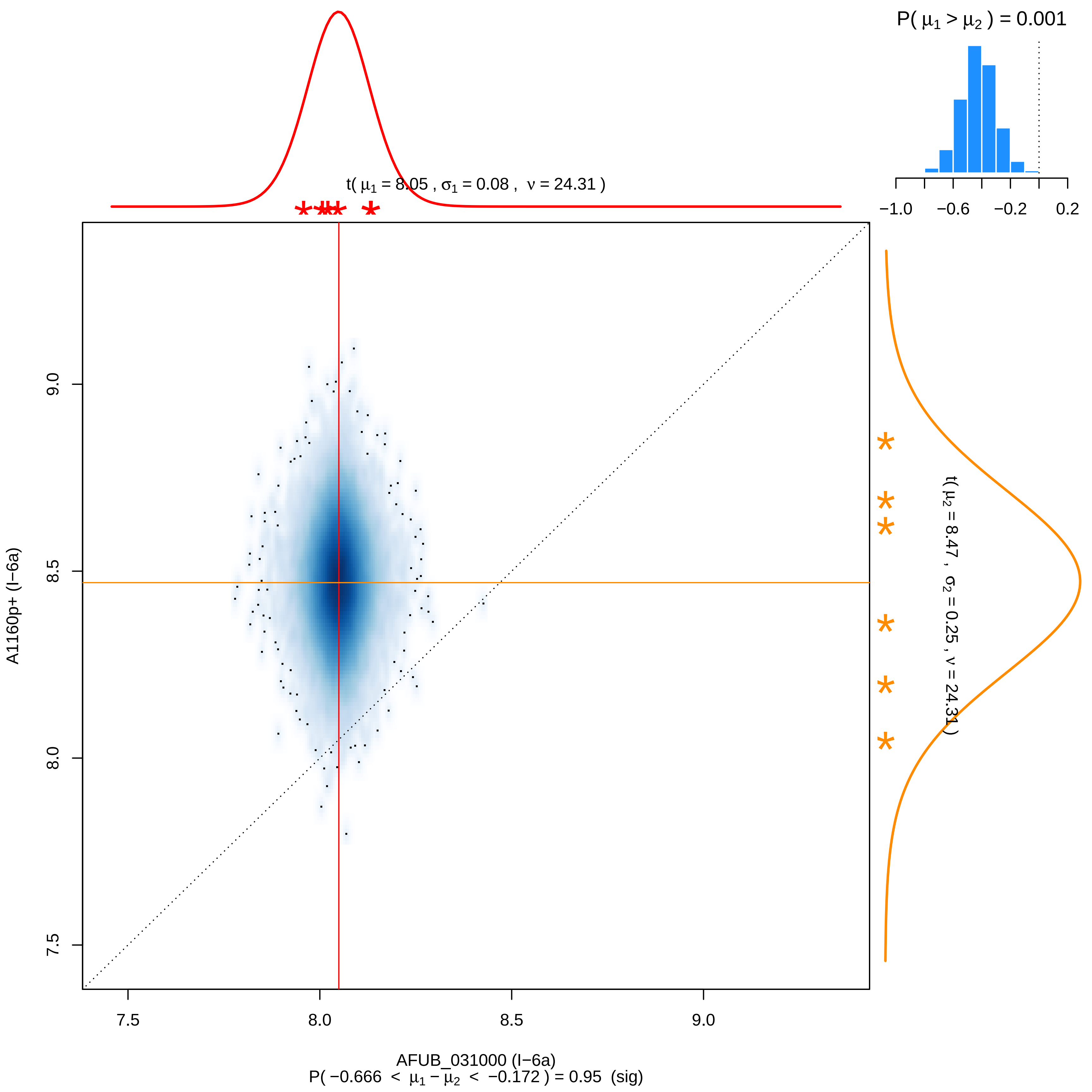

P( μ1 > μ2 ) = 0.001
−1.0
−0.6
−0.2
0.2
t( μ1 = 8.05 , σ1 = 0.08 , ν = 24.31 )
t( μ2 = 8.47 , σ2 = 0.25 , ν = 24.31 )
A1160p+ (I−6a)
9.0
8.5
8.0
7.5
7.5
8.0
8.5
9.0
AFUB_031000 (I−6a)
P( −0.666 < μ1 − μ2 < −0.172 ) = 0.95 (sig)

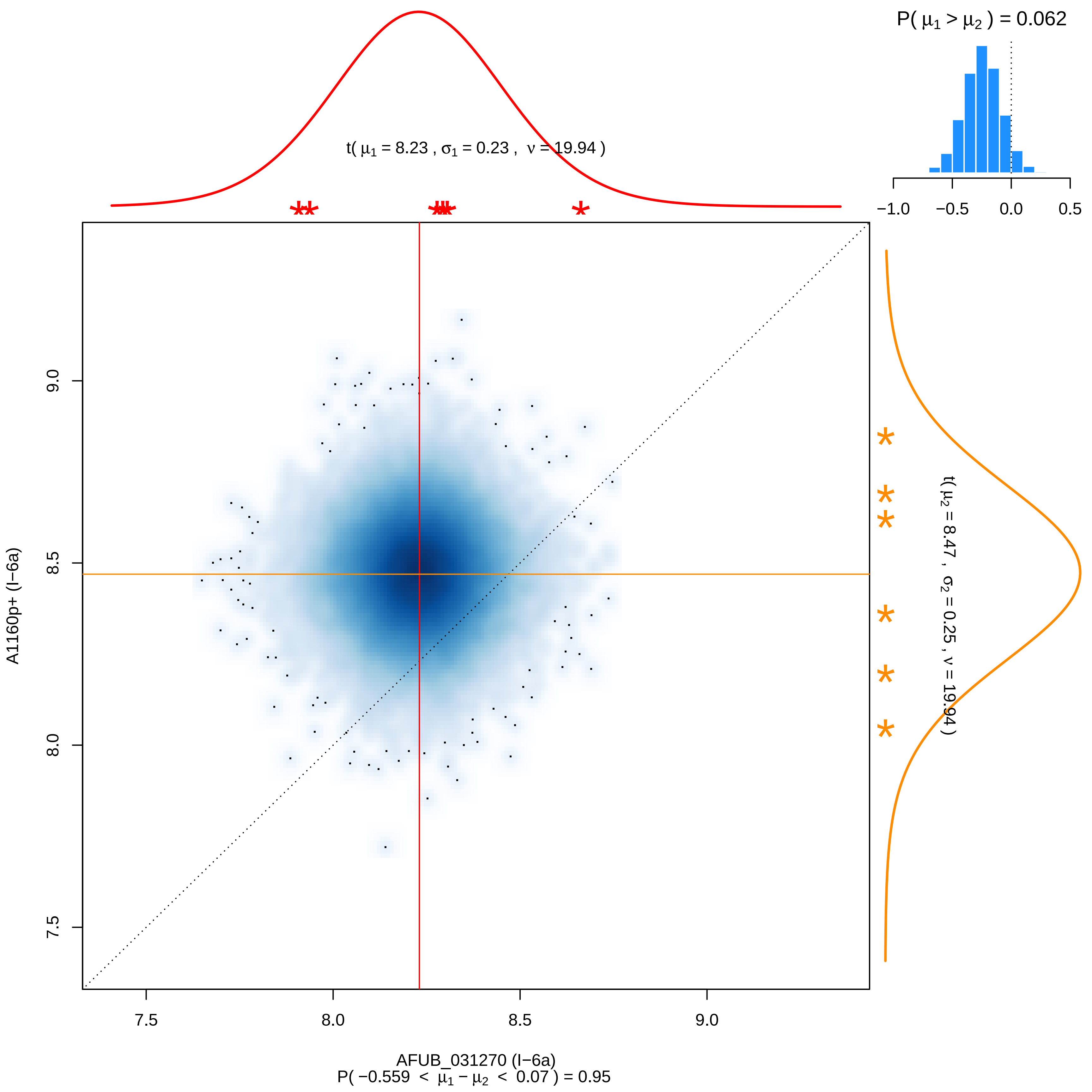

P( μ1 > μ2 ) = 0.062
−1.0 −0.5 0.0 0.5
t( μ1 = 8.23 , σ1 = 0.23 , ν = 19.94 )
t( μ2 = 8.47 , σ2 = 0.25 , ν = 19.94 )
A1160p+ (I−6a)
7.5 8.0 8.5 9.0
AFUB_031270 (I−6a)
P( −0.559 < μ1 − μ2 < 0.07 ) = 0.95

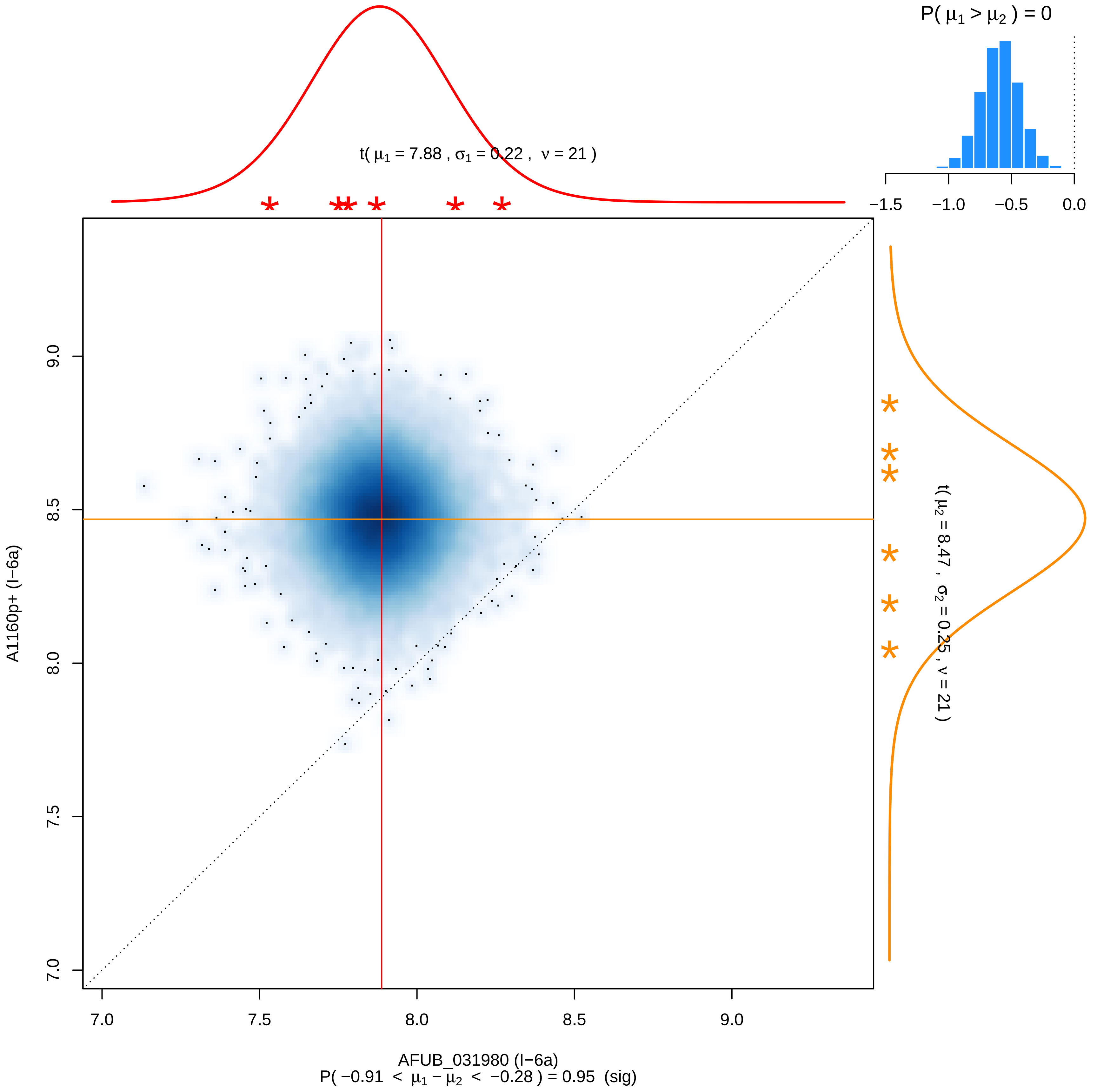

P( μ1 > μ2 ) = 0
−1.5
−1.0
−0.5
0.0
t( μ1 = 7.88 , σ1 = 0.22 , ν = 21 )
t( μ2 = 8.47 , σ2 = 0.25 , ν = 21 )
A1160p+ (I−6a)
7.0
7.5
8.0
8.5
9.0
AFUB_031980 (I−6a)
P( −0.91 < μ1 − μ2 < −0.28 ) = 0.95 (sig)

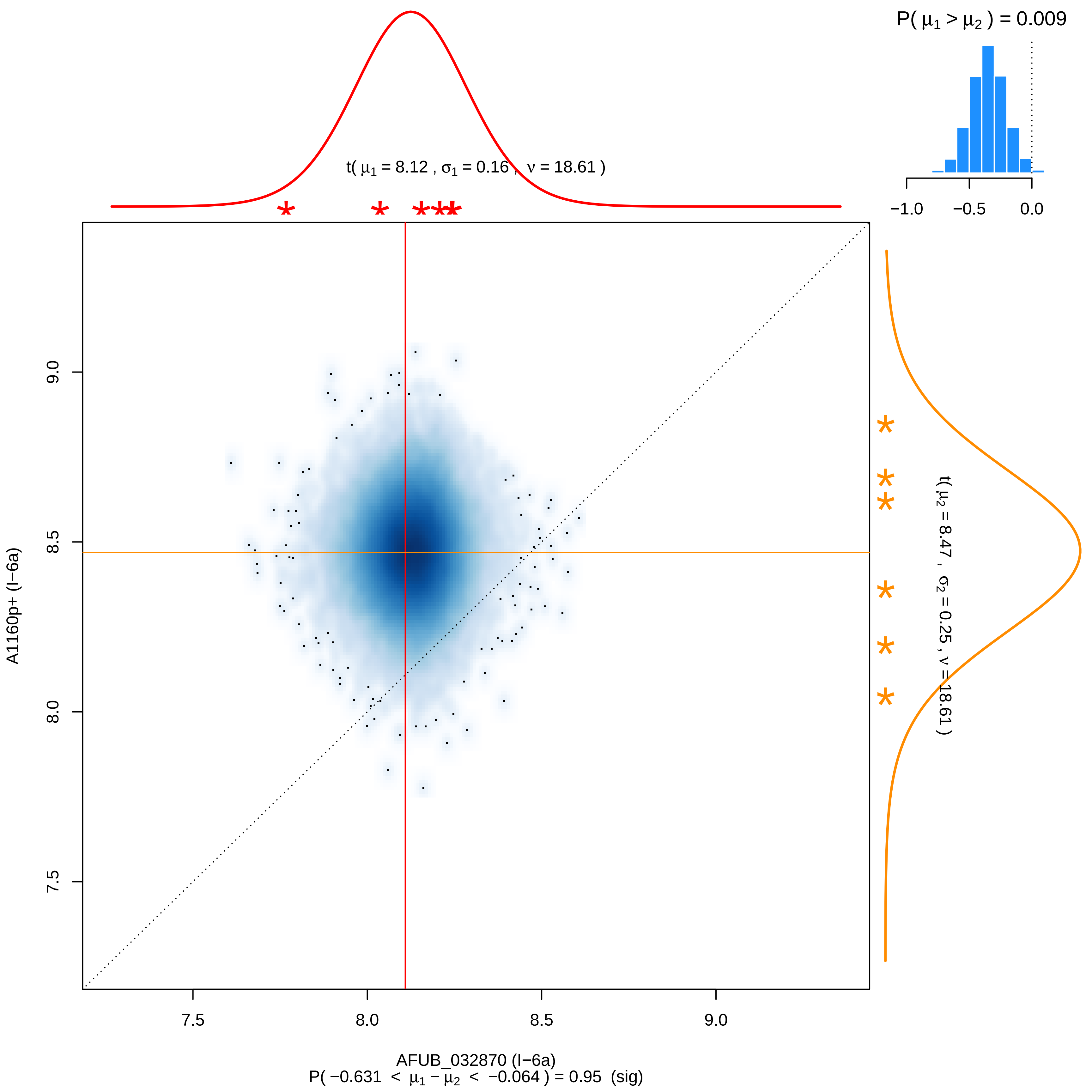

P( μ1 > μ2 ) = 0.009
−1.0
−0.5
0.0
t( μ1 = 8.12 , σ1 = 0.16 , ν = 18.61 )
9.0
8.5
8.0
7.5
A1160p+ (I−6a)
t( μ2 = 8.47 , σ2 = 0.25 , ν = 18.61 )
AFUB_032870 (I−6a)
P( −0.631 < μ1 − μ2 < −0.064 ) = 0.95 (sig)

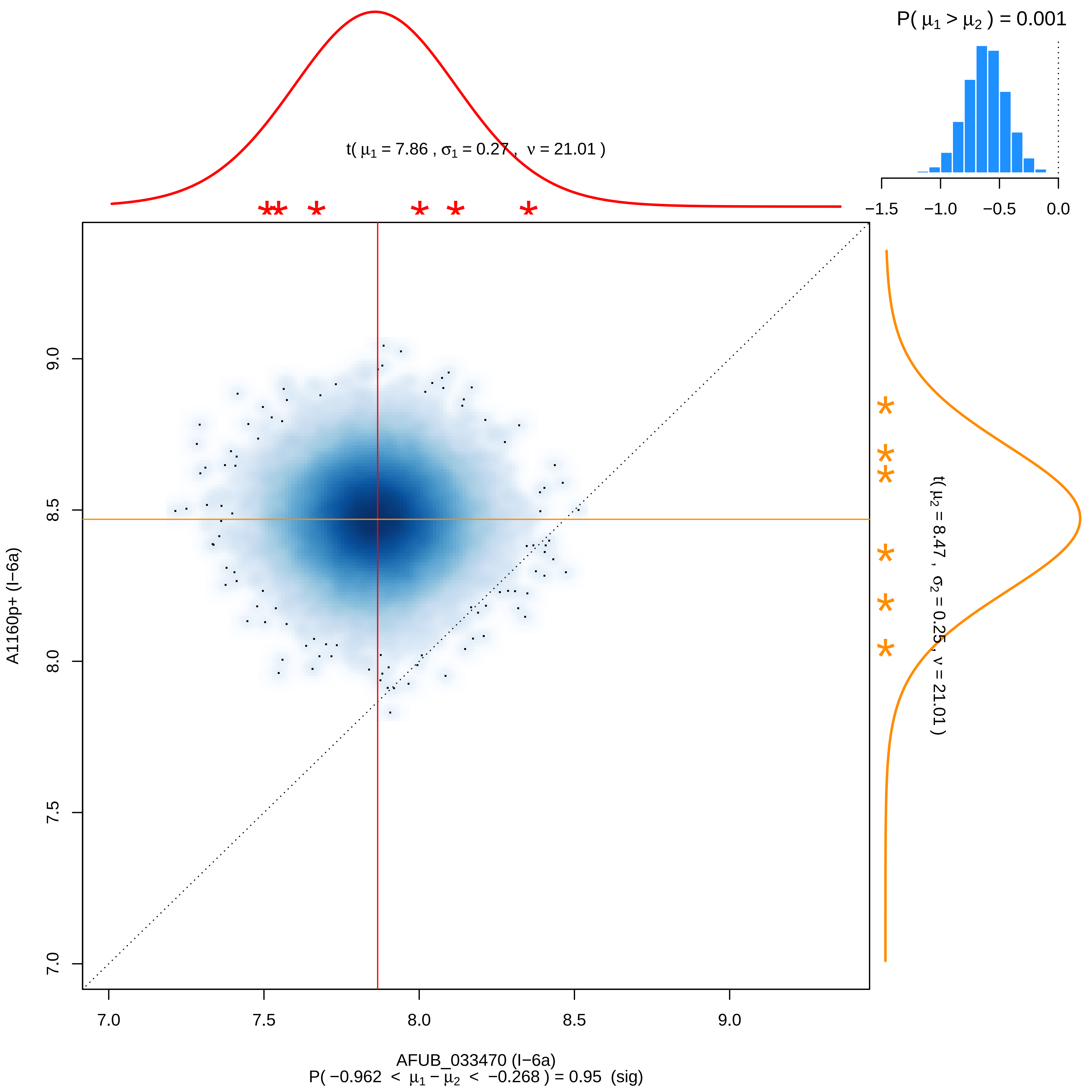

P( $\mu_1 > \mu_2$ ) = 0.001
−1.5
−1.0
−0.5
0.0
t( $\mu_1$ = 7.86 , $\sigma_1$ = 0.27 , $\nu$ = 21.01 )
t( $\mu_2$ = 8.47 , $\sigma_2$ = 0.25 , $\nu$ = 21.01 )
9.0
8.5
8.0
7.5
7.0
A1160p+ (I−6a)
7.0
7.5
8.0
8.5
9.0
AFUB_033470 (I−6a)
P( −0.962 < $\mu_1 - \mu_2$ < −0.268 ) = 0.95 (sig)

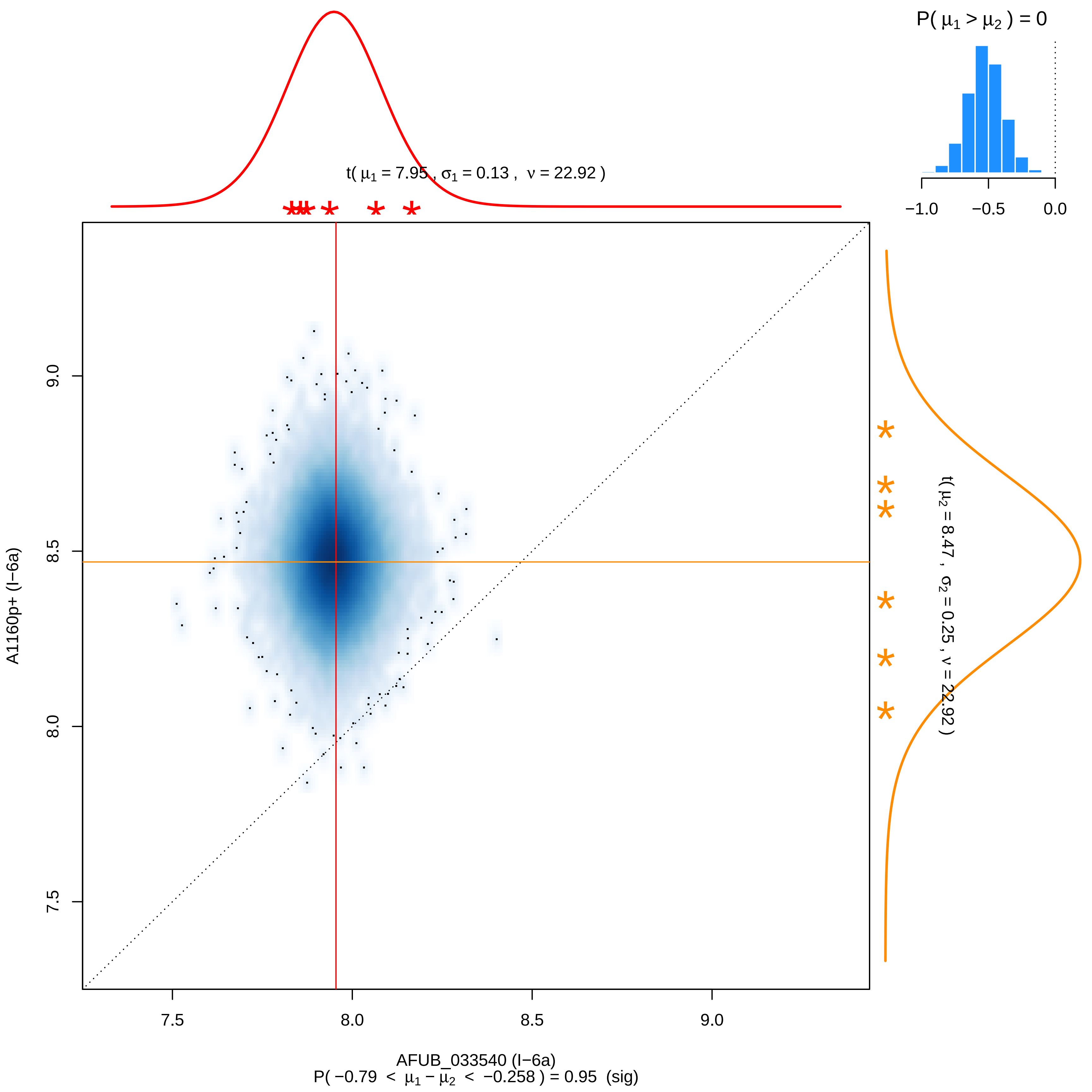
P( μ1 > μ2 ) = 0
−1.0
−0.5
0.0
t( μ1 = 7.95 , σ1 = 0.13 , ν = 22.92 )
A1160p+ (I−6a)
9.0
8.5
8.0
7.5
t( μ2 = 8.47 , σ2 = 0.25 , ν = 22.92 )
7.5
8.0
8.5
9.0
AFUB_033540 (I−6a)
P( −0.79 < μ1 − μ2 < −0.258 ) = 0.95 (sig)

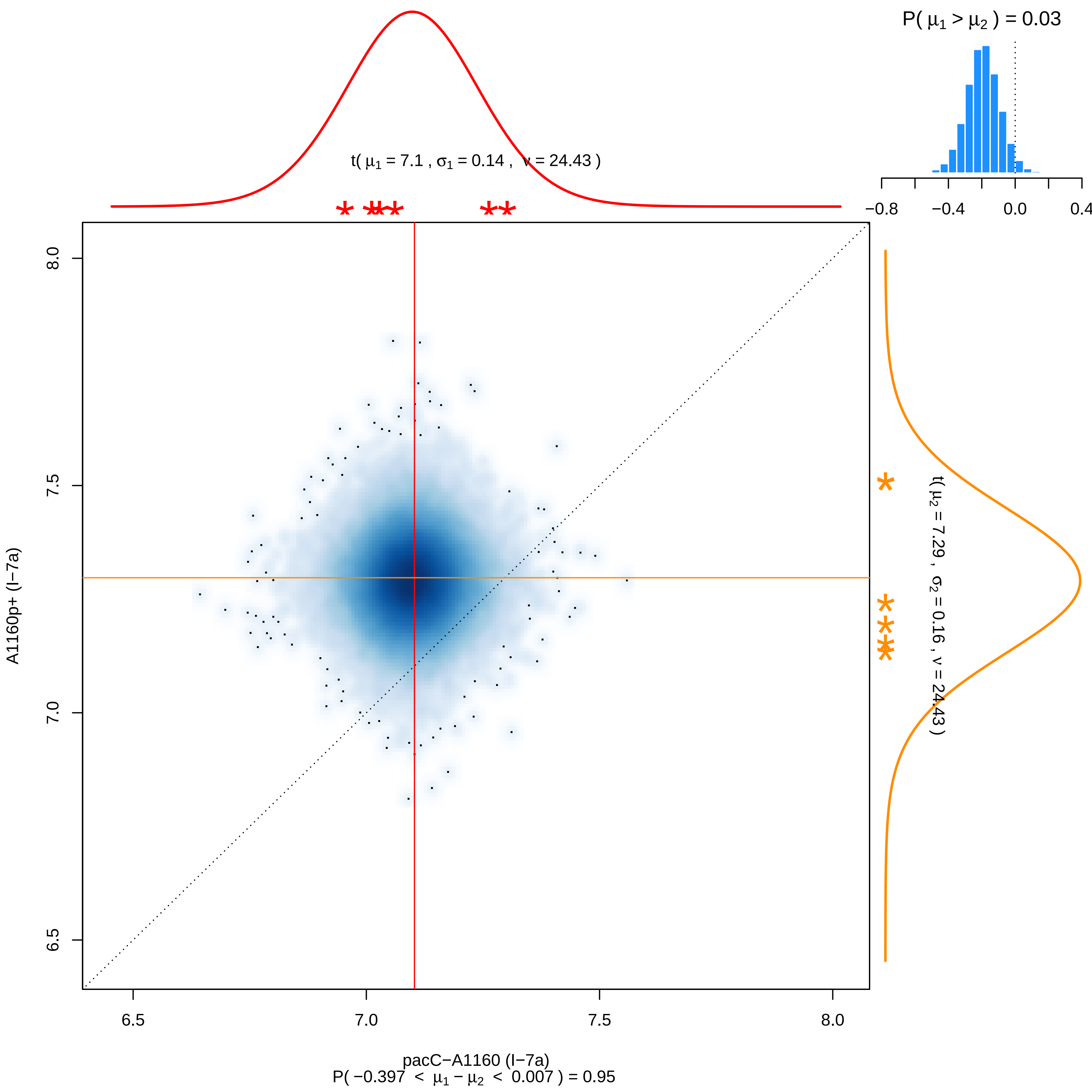

P( μ1 > μ2 ) = 0.03
−0.8
−0.4
0.0
0.4
t( μ1 = 7.1 , σ1 = 0.14 , ν = 24.43 )
t( μ2 = 7.29 , σ2 = 0.16 , ν = 24.43 )
8.0
7.5
7.0
6.5
A1160p+ (I−7a)
6.5
7.0
7.5
8.0
pacC−A1160 (I−7a)
P( −0.397 < μ1 − μ2 < 0.007 ) = 0.95

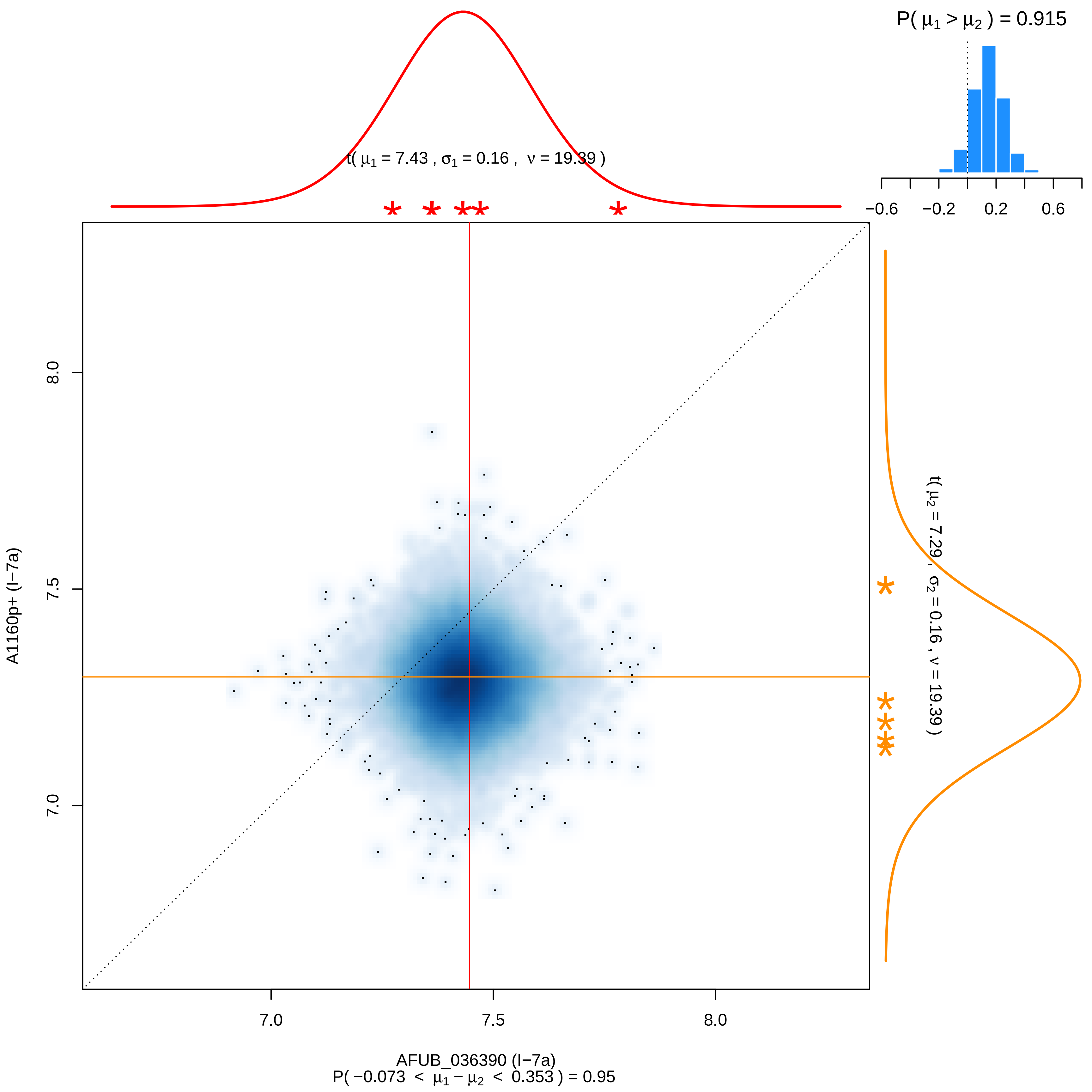

P( μ1 > μ2 ) = 0.915
−0.6
−0.2
0.2
0.6
t( μ1 = 7.43 , σ1 = 0.16 , ν = 19.39 )
t( μ2 = 7.29 , σ2 = 0.16 , ν = 19.39 )
A1160p+ (I−7a)
8.0
7.5
7.0
7.0
7.5
8.0
AFUB_036390 (I−7a)
P( −0.073 < μ1 − μ2 < 0.353 ) = 0.95

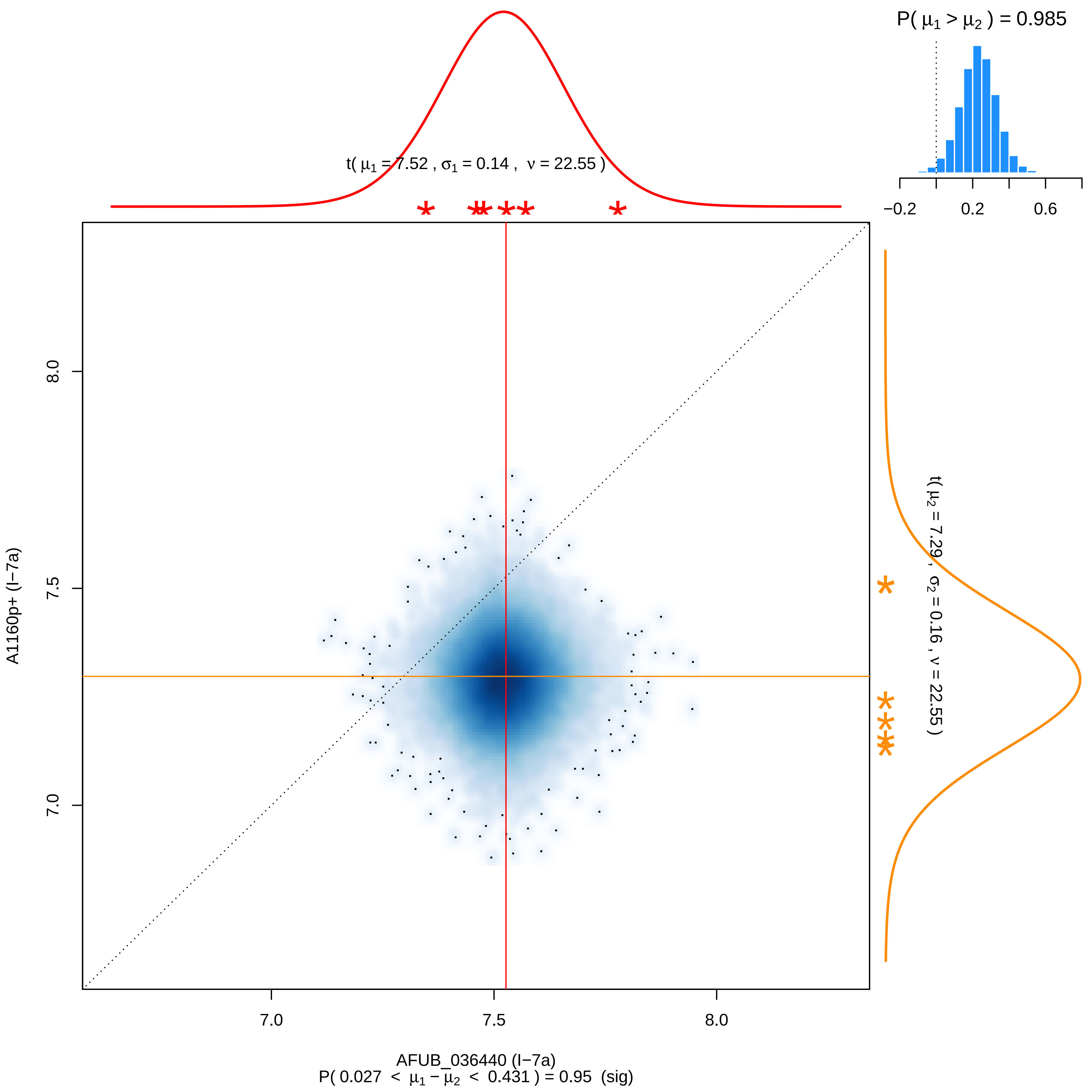

P( μ1 > μ2 ) = 0.985
−0.2
0.2
0.6
t( μ1 = 7.52 , σ1 = 0.14 , ν = 22.55 )
t( μ2 = 7.29 , σ2 = 0.16 , ν = 22.55 )
8.0
7.5
7.0
A1160p+ (I−7a)
7.0
7.5
8.0
AFUB_036440 (I−7a)
P( 0.027 < μ1 − μ2 < 0.431 ) = 0.95 (sig)

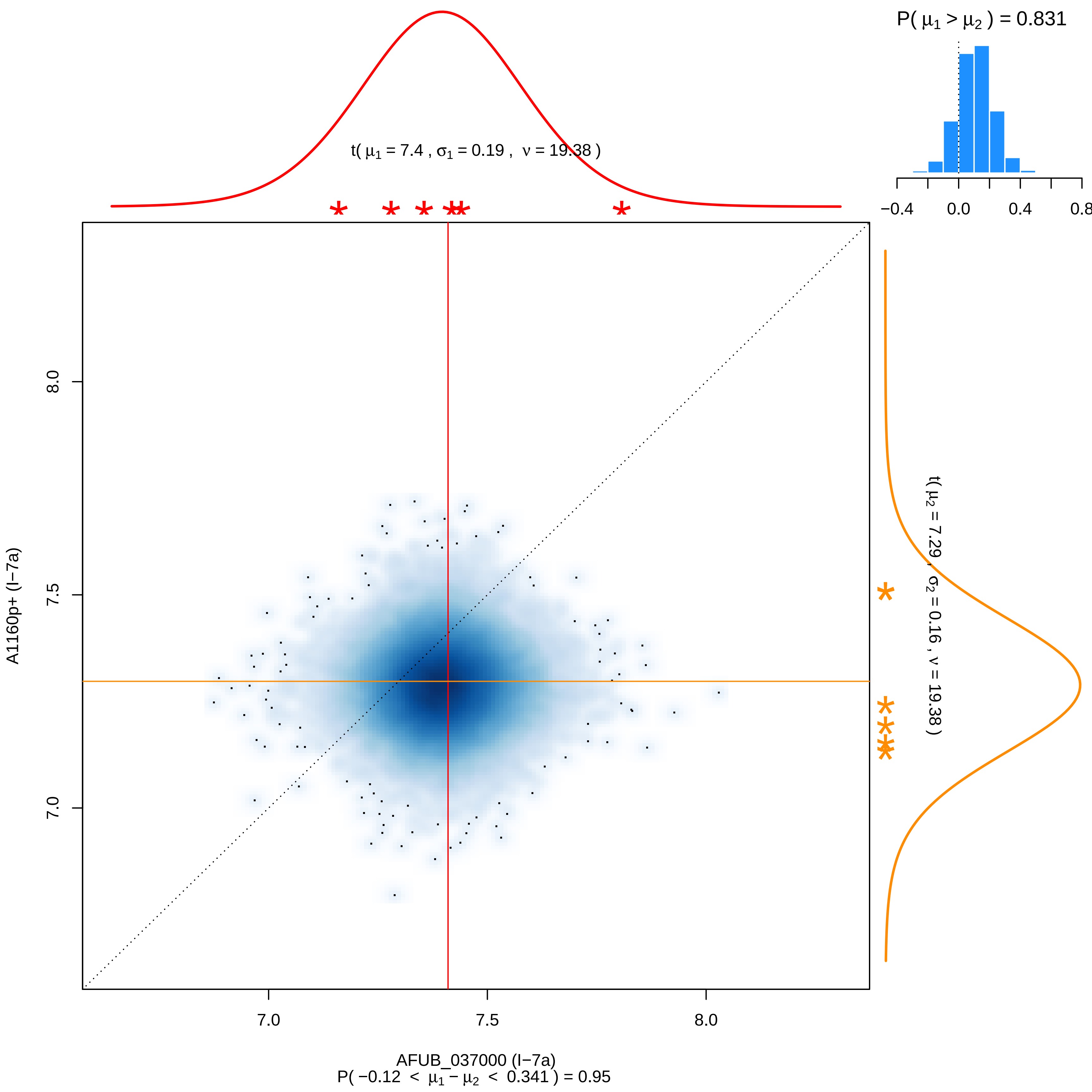

P( μ1 > μ2 ) = 0.831
−0.4
0.0
0.4
0.8
t( μ1 = 7.4 , σ1 = 0.19 , ν = 19.38 )
t( μ2 = 7.29 , σ2 = 0.16 , ν = 19.38 )
A1160p+ (I−7a)
8.0
7.5
7.0
AFUB_037000 (I−7a)
P( −0.12 < μ1 − μ2 < 0.341 ) = 0.95

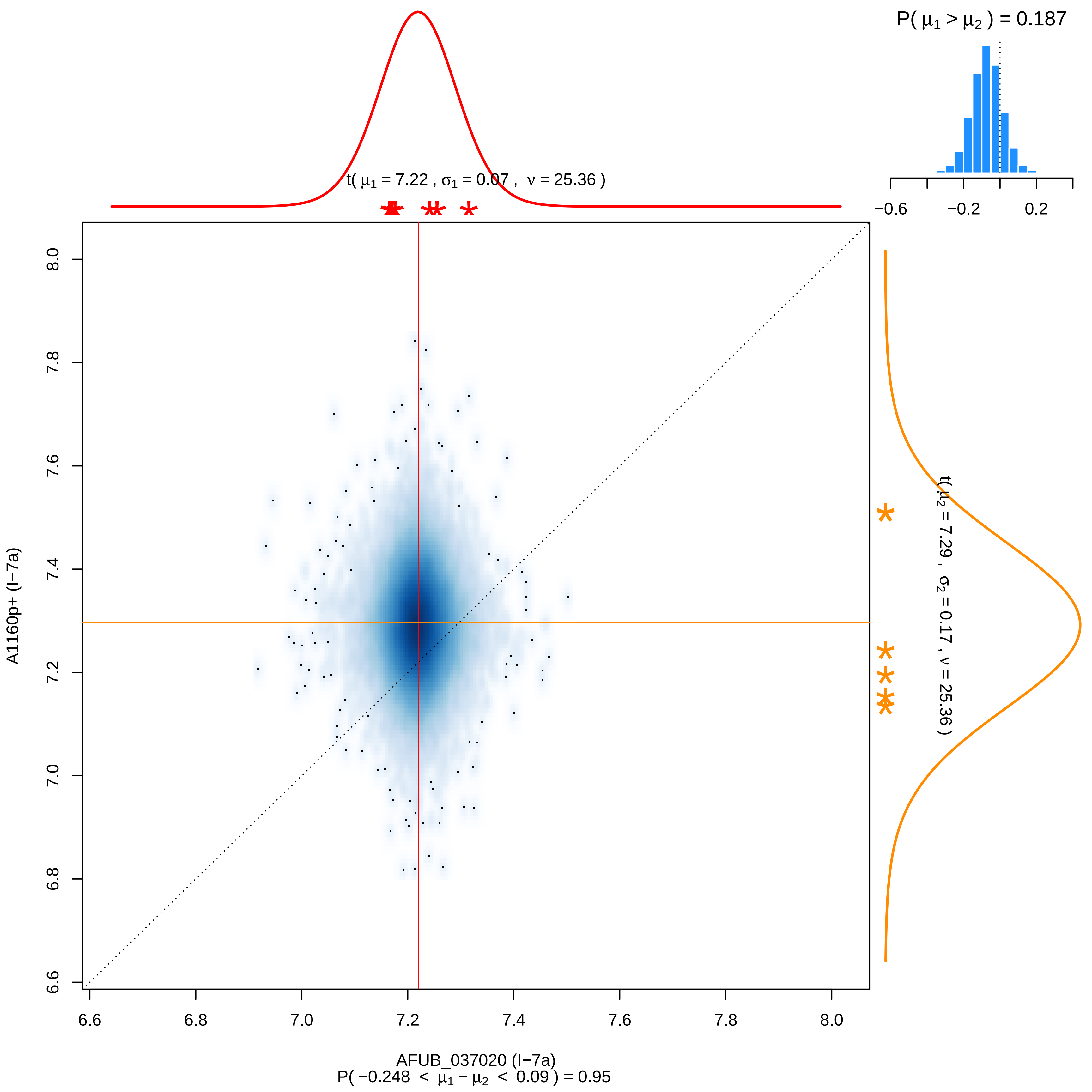

P( μ1 > μ2 ) = 0.187
−0.6
−0.2
0.2
t( μ1 = 7.22 , σ1 = 0.07 , ν = 25.36 )
t( μ2 = 7.29 , σ2 = 0.17 , ν = 25.36 )
A1160p+ (I−7a)
6.6
6.8
7.0
7.2
7.4
7.6
7.8
8.0
AFUB_037020 (I−7a)
P( −0.248 < μ1 − μ2 < 0.09 ) = 0.95

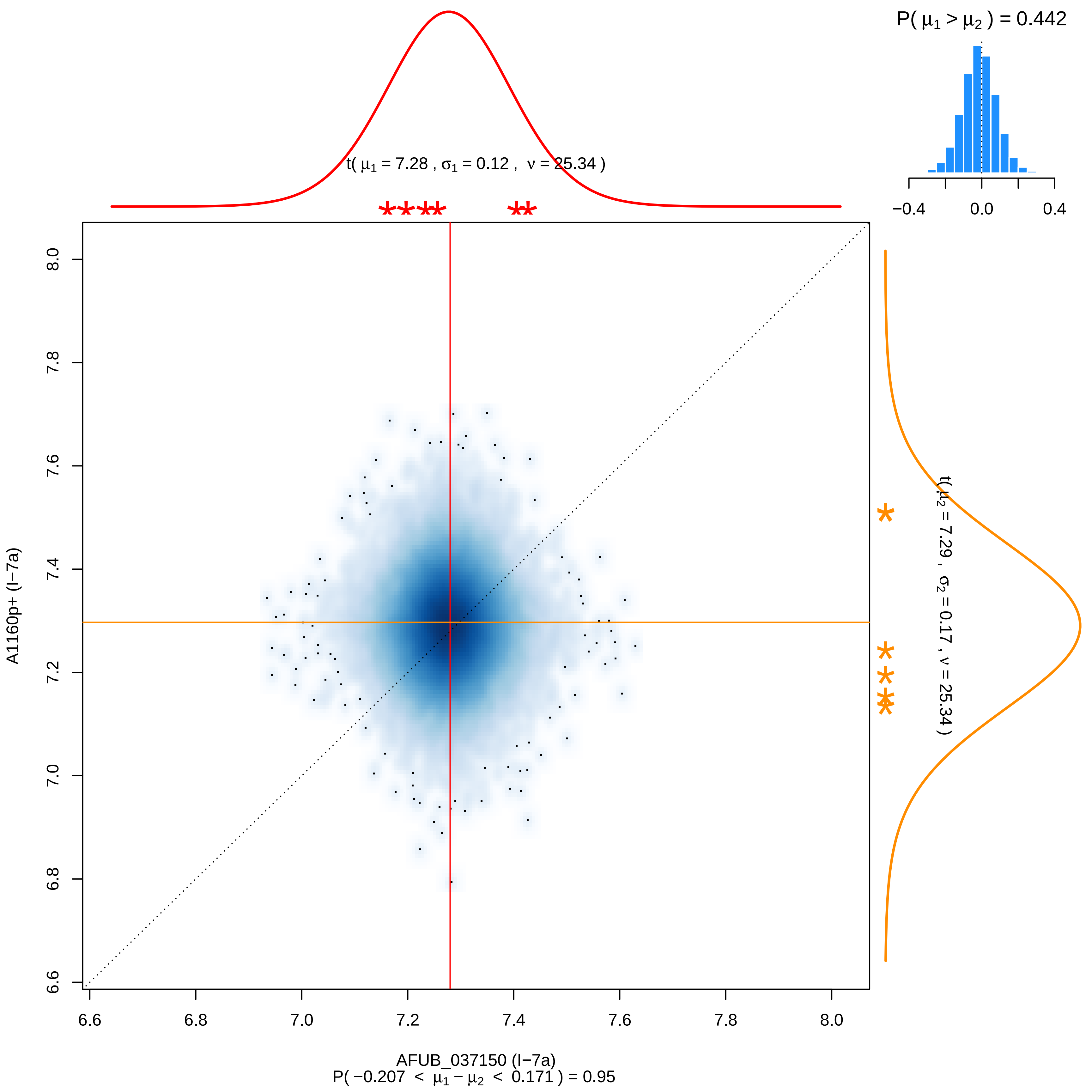

P( μ1 > μ2 ) = 0.442
−0.4
0.0
0.4
t( μ1 = 7.28 , σ1 = 0.12 , ν = 25.34 )
t( μ2 = 7.29 , σ2 = 0.17 , ν = 25.34 )
A1160p+ (I−7a)
6.6
6.8
7.0
7.2
7.4
7.6
7.8
8.0
AFUB_037150 (I−7a)
P( −0.207 < μ1 − μ2 < 0.171 ) = 0.95

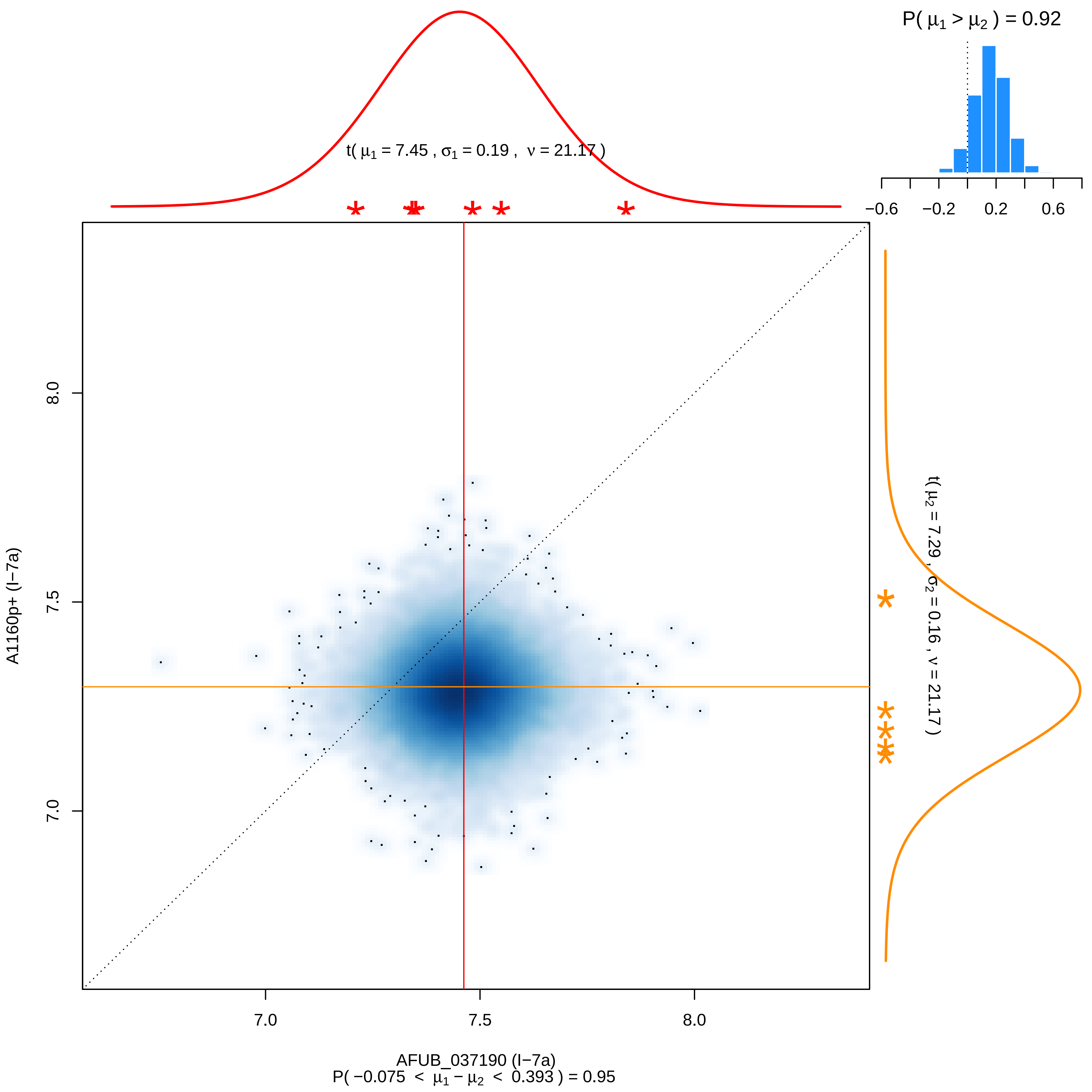

P( μ1 > μ2 ) = 0.92
−0.6
−0.2
0.2
0.6
t( μ1 = 7.45 , σ1 = 0.19 , ν = 21.17 )
t( μ2 = 7.29 , σ2 = 0.16 , ν = 21.17 )
8.0
7.5
7.0
A1160p+ (I−7a)
7.0
7.5
8.0
AFUB_037190 (I−7a)
P( −0.075 < μ1 − μ2 < 0.393 ) = 0.95

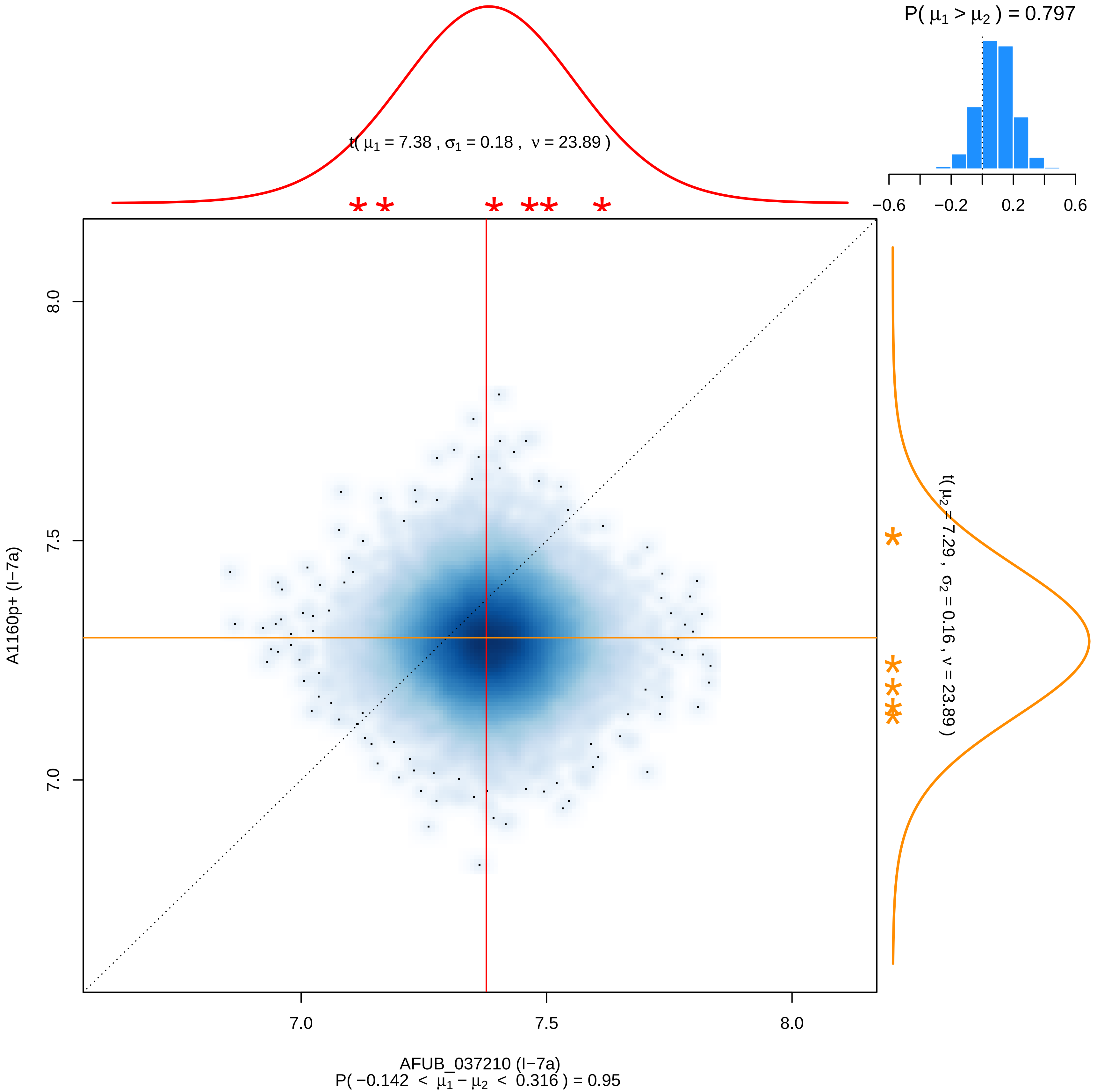

P( μ1 > μ2 ) = 0.797
−0.6
−0.2
0.2
0.6
t( μ1 = 7.38 , σ1 = 0.18 , ν = 23.89 )
t( μ2 = 7.29 , σ2 = 0.16 , ν = 23.89 )
8.0
7.5
7.0
A1160p+ (I−7a)
7.0
7.5
8.0
AFUB_037210 (I−7a)
P( −0.142 < μ1 − μ2 < 0.316 ) = 0.95

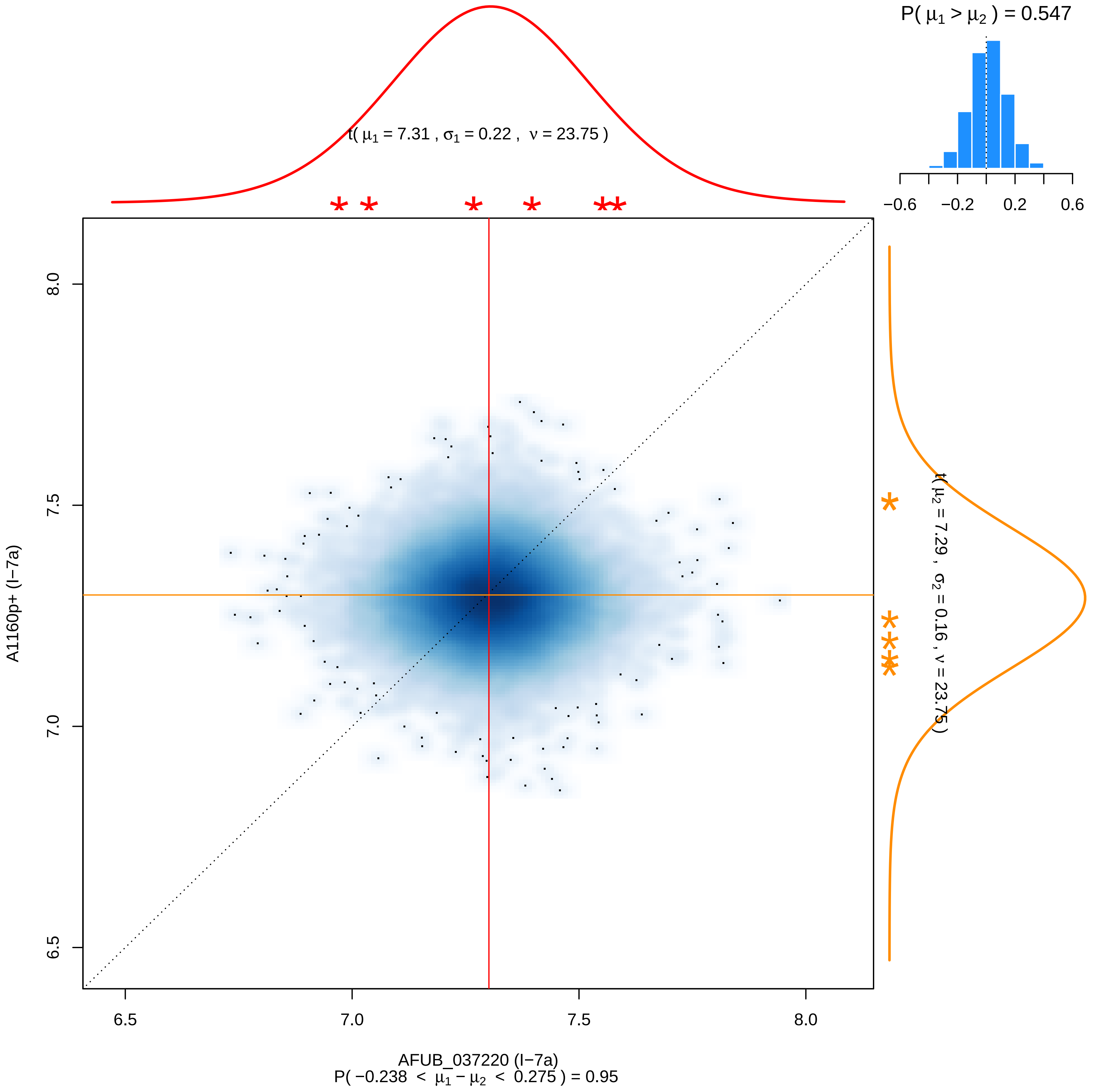

P( μ1 > μ2 ) = 0.547
t( μ1 = 7.31 , σ1 = 0.22 , ν = 23.75 )
−0.6
−0.2
0.2
0.6
t( μ2 = 7.29 , σ2 = 0.16 , ν = 23.75 )
8.0
7.5
7.0
6.5
A1160p+ (I−7a)
6.5
7.0
7.5
8.0
AFUB_037220 (I−7a)
P( −0.238 < μ1 − μ2 < 0.275 ) = 0.95

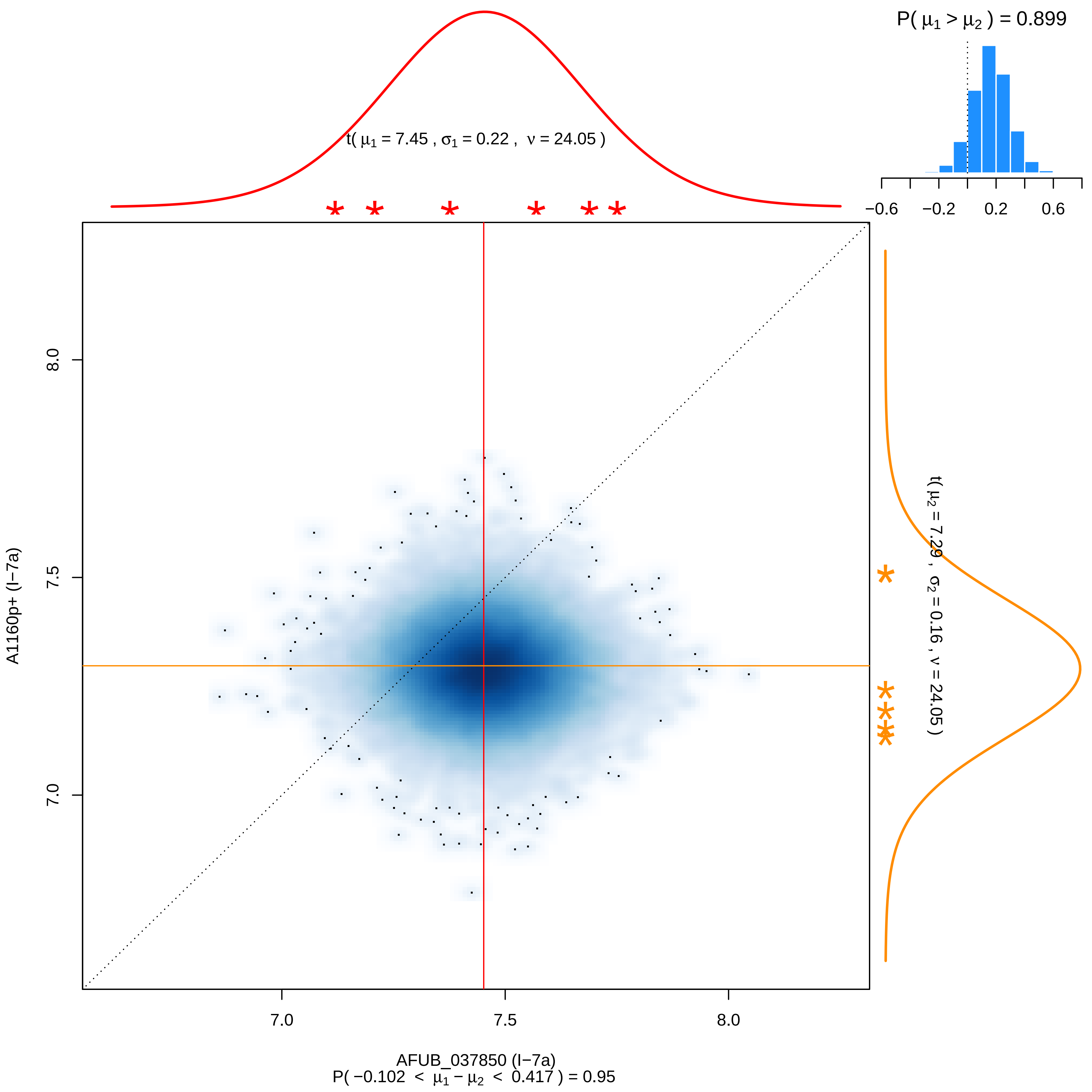

P( μ1 > μ2 ) = 0.899
t( μ1 = 7.45 , σ1 = 0.22 , ν = 24.05 )
−0.6
−0.2
0.2
0.6
t( μ2 = 7.29 , σ2 = 0.16 , ν = 24.05 )
8.0
7.5
7.0
A1160p+ (I−7a)
7.0
7.5
8.0
AFUB_037850 (I−7a)
P( −0.102 < μ1 − μ2 < 0.417 ) = 0.95

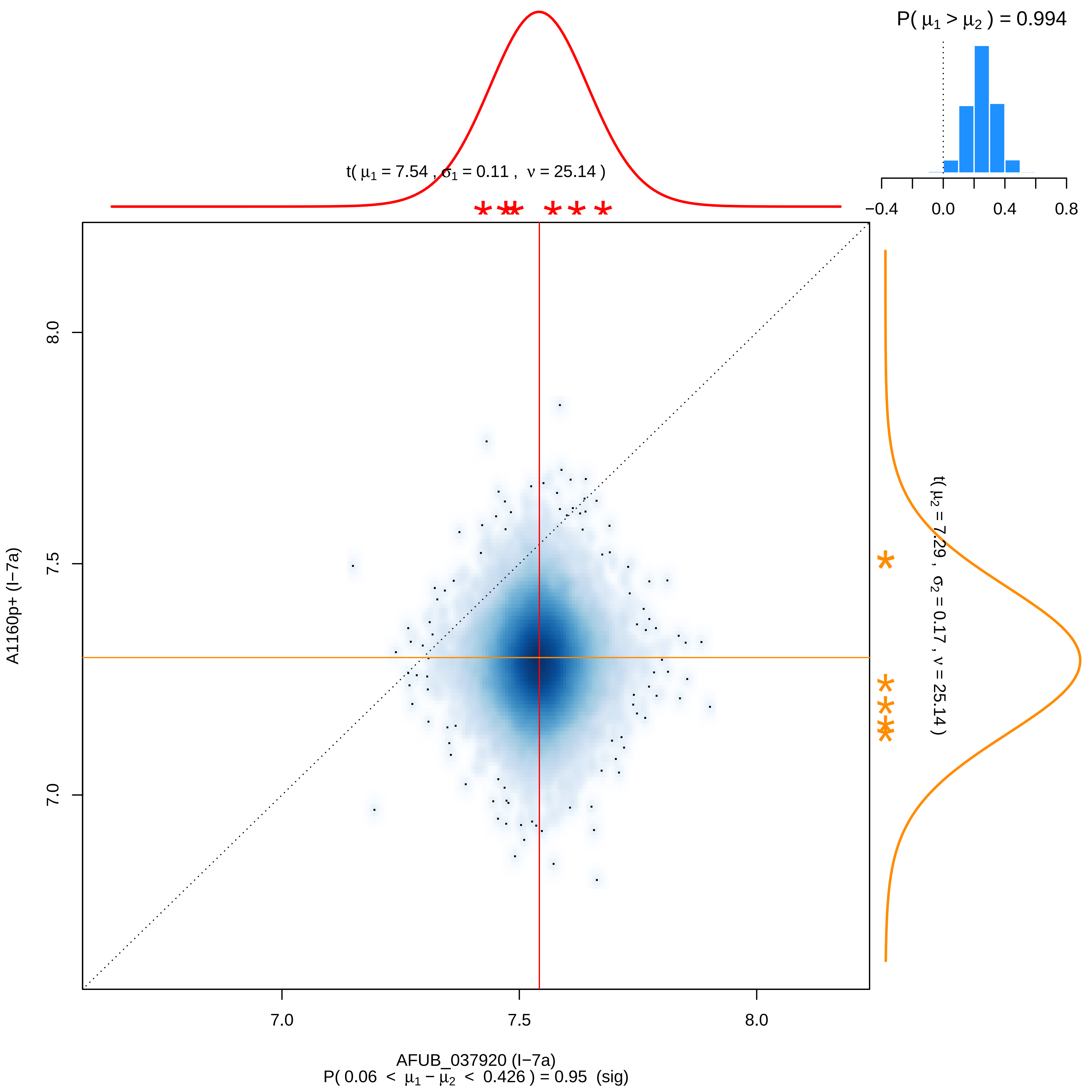

P( μ1 > μ2 ) = 0.994
−0.4 0.0 0.4 0.8
t( μ1 = 7.54 , σ1 = 0.11 , ν = 25.14 )
t( μ2 = 7.29 , σ2 = 0.17 , ν = 25.14 )
8.0
7.5
7.0
A1160p+ (I−7a)
7.0 7.5 8.0
AFUB_037920 (I−7a)
P( 0.06 < μ1 − μ2 < 0.426 ) = 0.95 (sig)

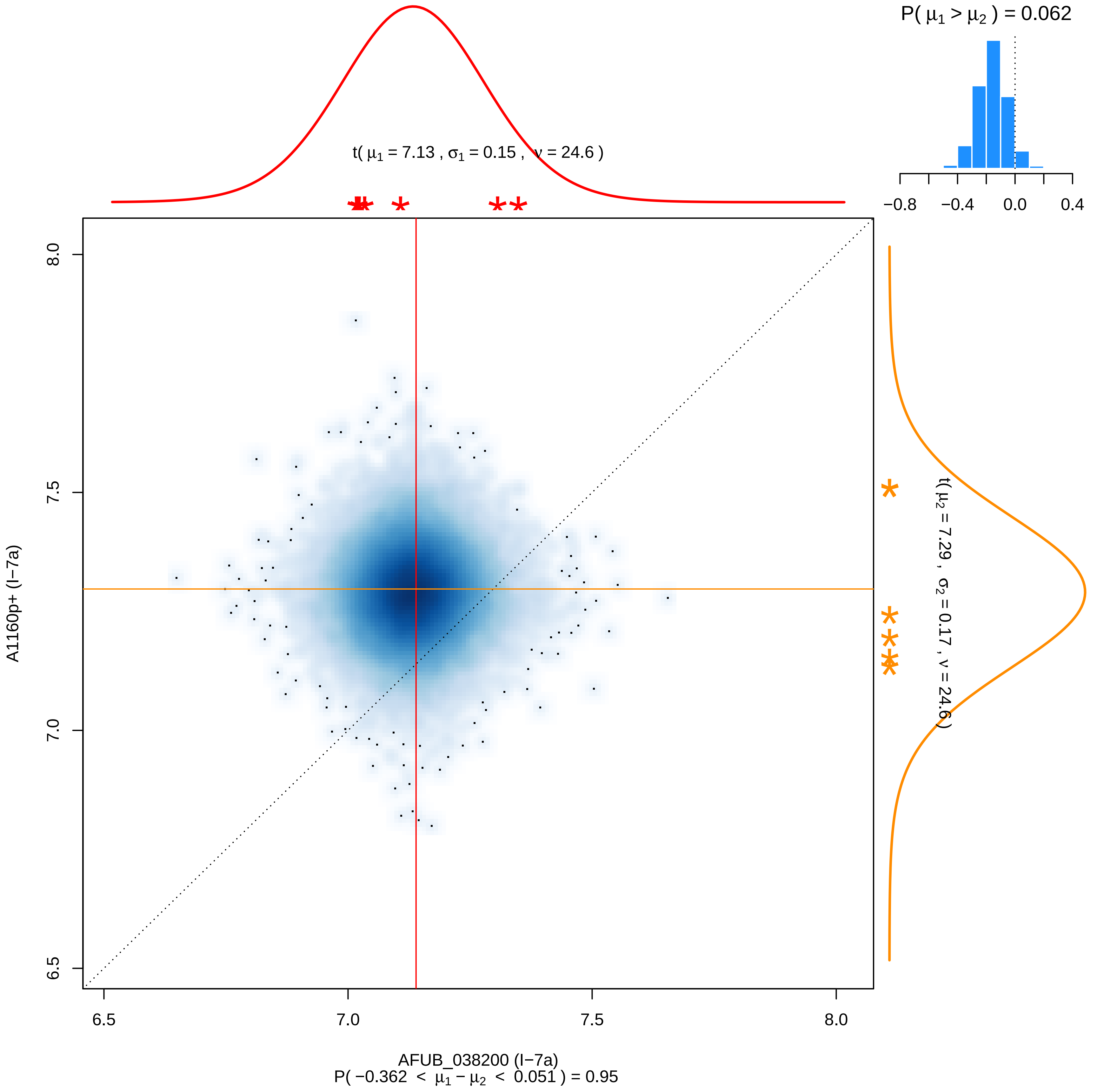

P( μ1 > μ2 ) = 0.062
−0.8
−0.4
0.0
0.4
t( μ1 = 7.13 , σ1 = 0.15 , ν = 24.6 )
t( μ2 = 7.29 , σ2 = 0.17 , ν = 24.6 )
A1160p+ (I−7a)
8.0
7.5
7.0
6.5
6.5
7.0
7.5
8.0
AFUB_038200 (I−7a)
P( −0.362 < μ1 − μ2 < 0.051 ) = 0.95

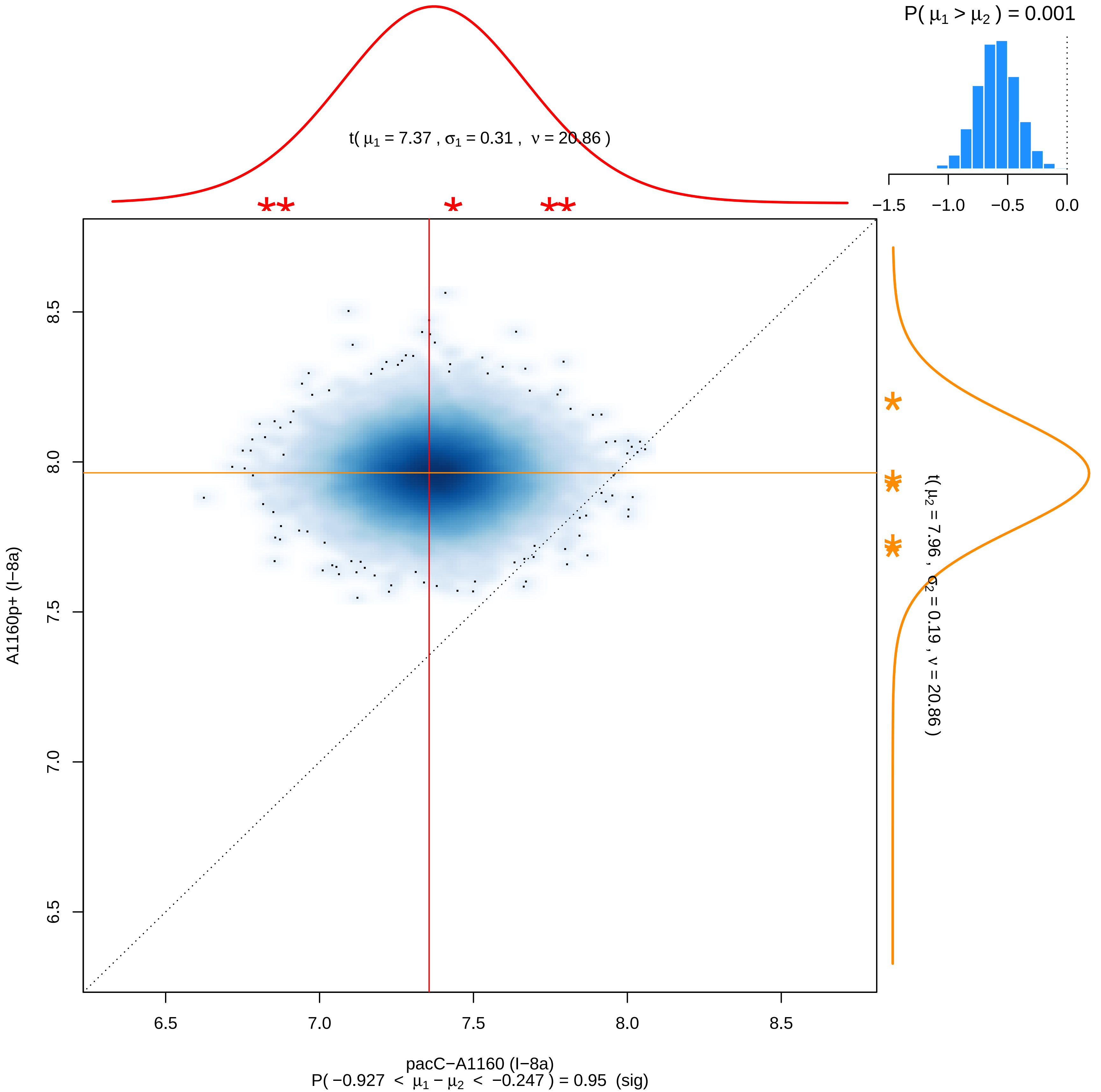

P( μ1 > μ2 ) = 0.001
−1.5
−1.0
−0.5
0.0
t( μ1 = 7.37 , σ1 = 0.31 , ν = 20.86 )
t( μ2 = 7.96 , σ2 = 0.19 , ν = 20.86 )
A1160p+ (I−8a)
8.5
8.0
7.5
7.0
6.5
6.5
7.0
7.5
8.0
8.5
pacC−A1160 (I−8a)
P( −0.927 < μ1 − μ2 < −0.247 ) = 0.95 (sig)

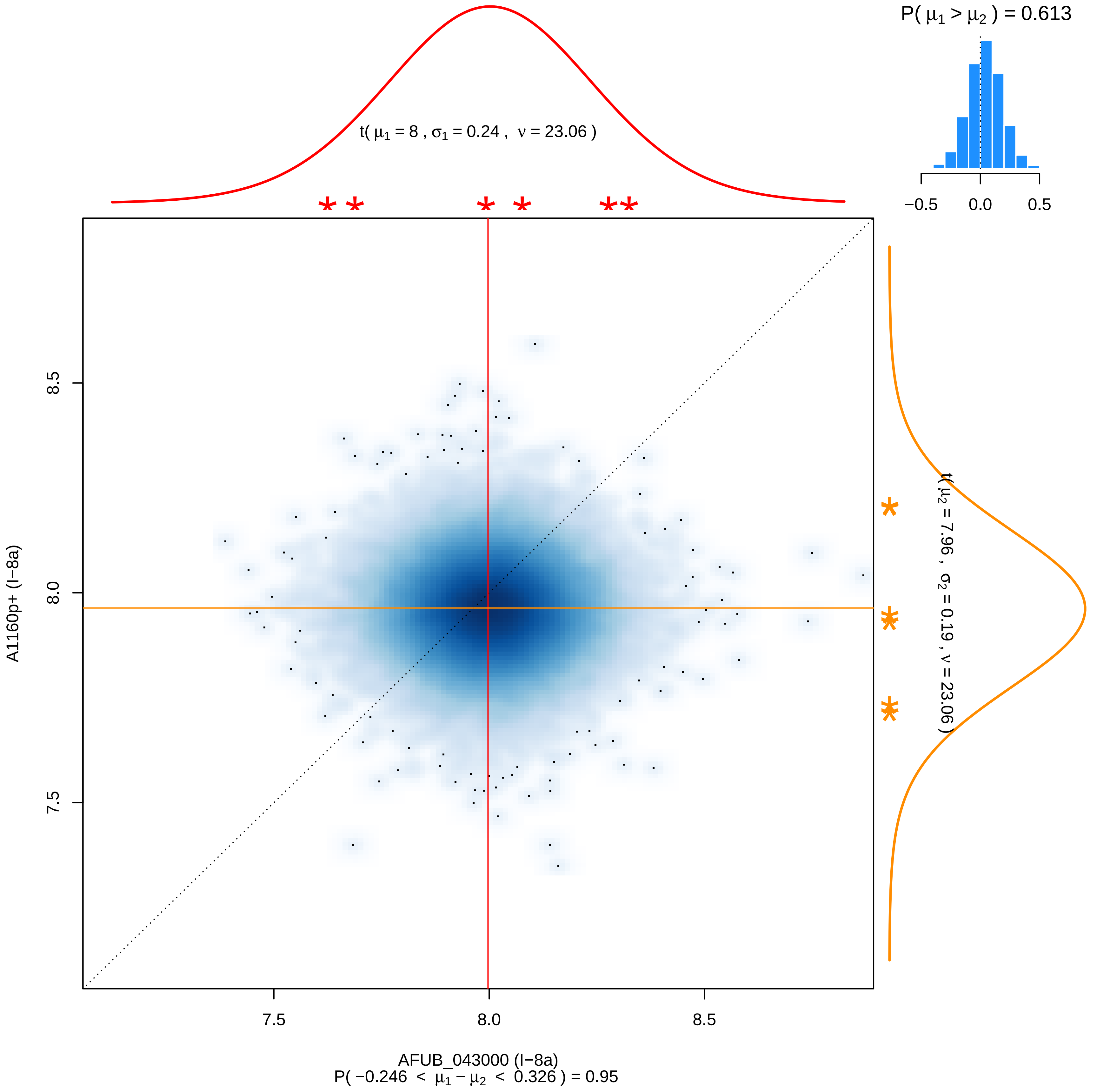

P( μ1 > μ2 ) = 0.613
−0.5
0.0
0.5
t( μ1 = 8 , σ1 = 0.24 , ν = 23.06 )
t( μ2 = 7.96 , σ2 = 0.19 , ν = 23.06 )
8.5
8.0
7.5
A1160p+ (I−8a)
7.5
8.0
8.5
AFUB_043000 (I−8a)
P( −0.246 < μ1 − μ2 < 0.326 ) = 0.95

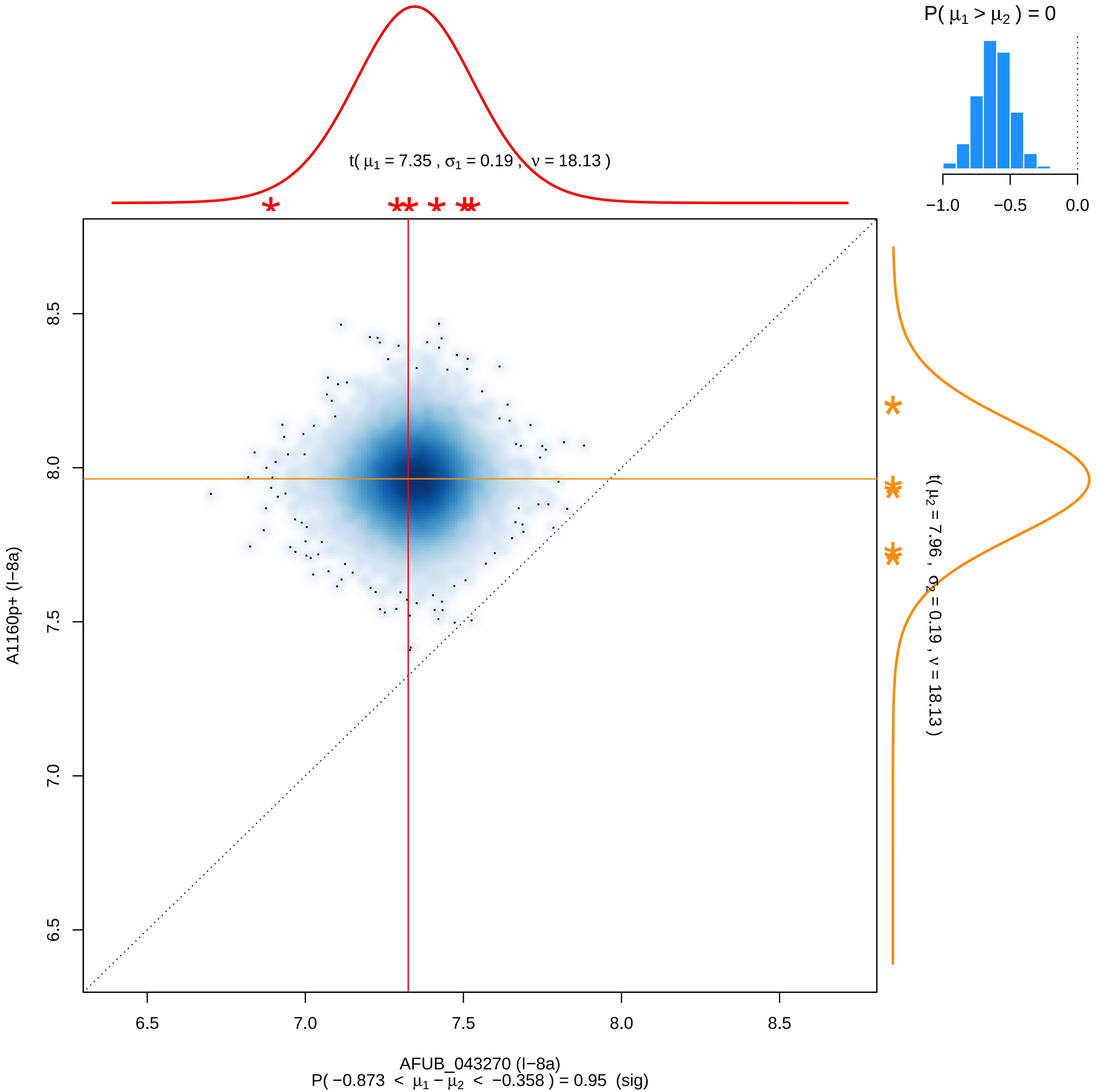

P( μ1 > μ2 ) = 0
−1.0
−0.5
0.0
t( μ1 = 7.35 , σ1 = 0.19 , ν = 18.13 )
t( μ2 = 7.96 , σ2 = 0.19 , ν = 18.13 )
A1160p+ (I−8a)
8.5
8.0
7.5
7.0
6.5
AFUB_043270 (I−8a)
P( −0.873 < μ1 − μ2 < −0.358 ) = 0.95 (sig)

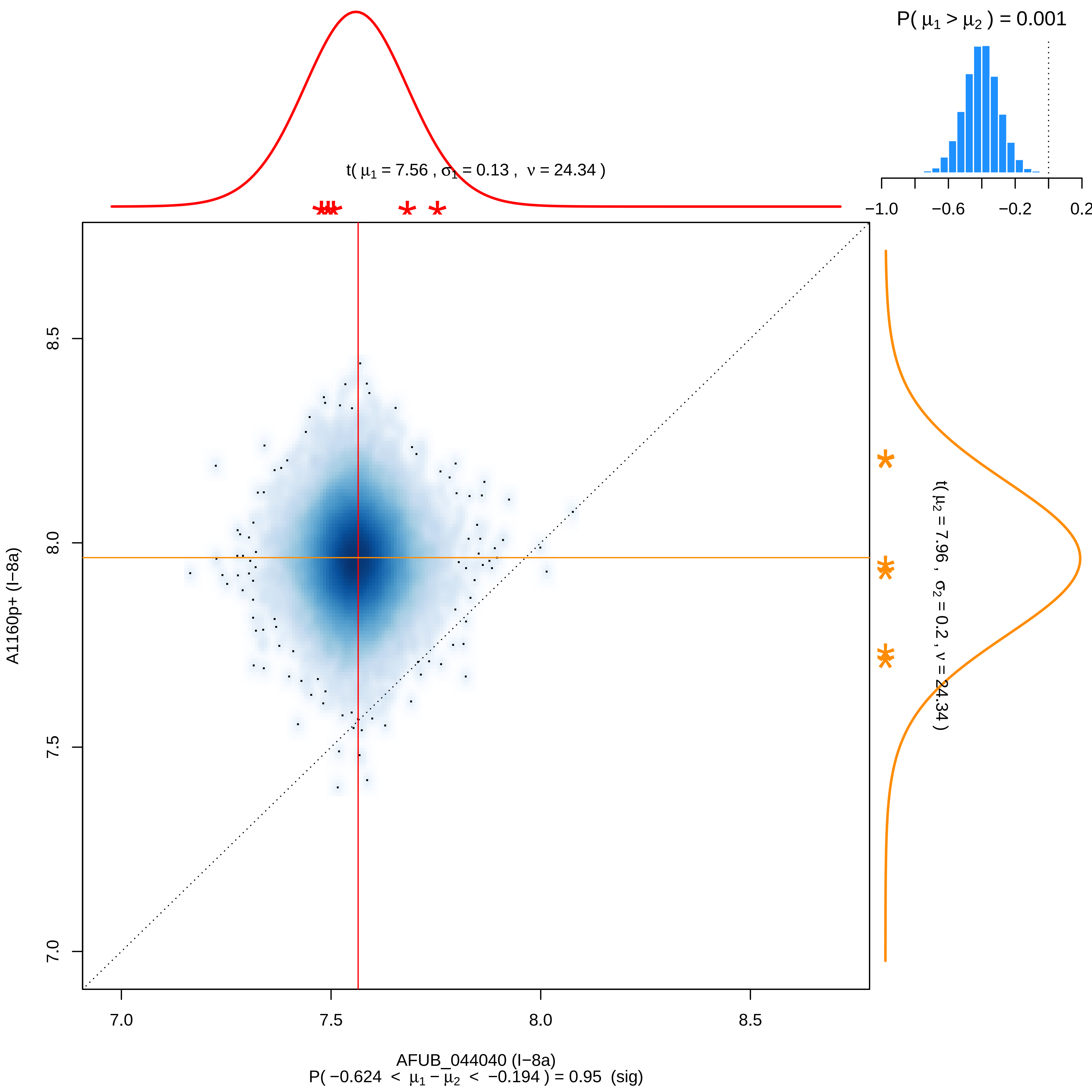

P( μ1 > μ2 ) = 0.001
−1.0
−0.6
−0.2
0.2
t( μ1 = 7.56 , σ1 = 0.13 , ν = 24.34 )
8.5
8.0
7.5
7.0
A1160p+ (I−8a)
t( μ2 = 7.96 , σ2 = 0.2 , ν = 24.34 )
7.0
7.5
8.0
8.5
AFUB_044040 (I−8a)
P( −0.624 < μ1 − μ2 < −0.194 ) = 0.95 (sig)

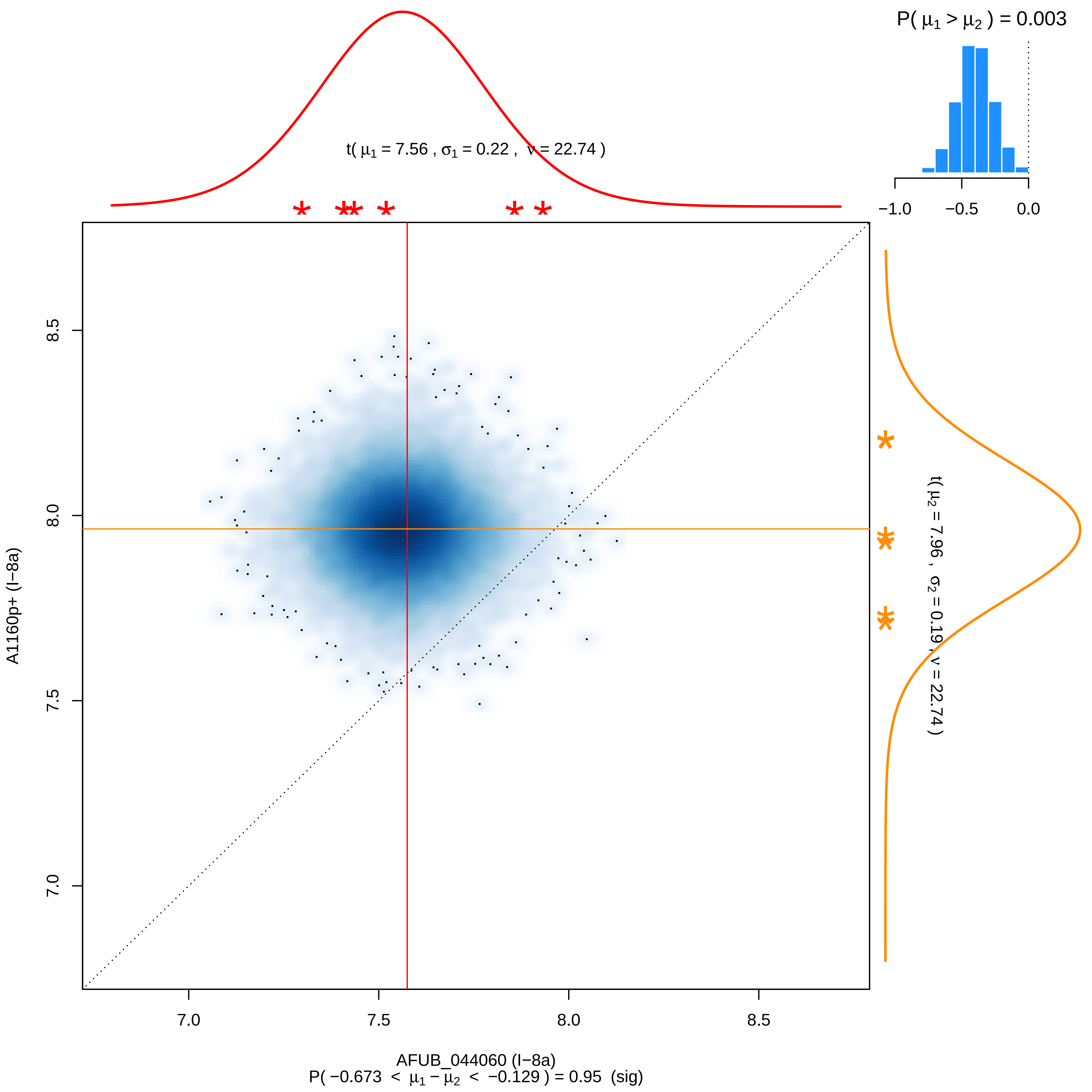

P( $\mu_1 > \mu_2$ ) = 0.003
t( $\mu_1$ = 7.56 , $\sigma_1$ = 0.22 , $\nu$ = 22.74 )
−1.0
−0.5
0.0
8.5
8.0
7.5
7.0
A1160p+ (I−8a)
t( $\mu_2$ = 7.96 , $\sigma_2$ = 0.19 , $\nu$ = 22.74 )
7.0
7.5
8.0
8.5
AFUB_044060 (I−8a)
P( −0.673 < $\mu_1 - \mu_2$ < −0.129 ) = 0.95 (sig)

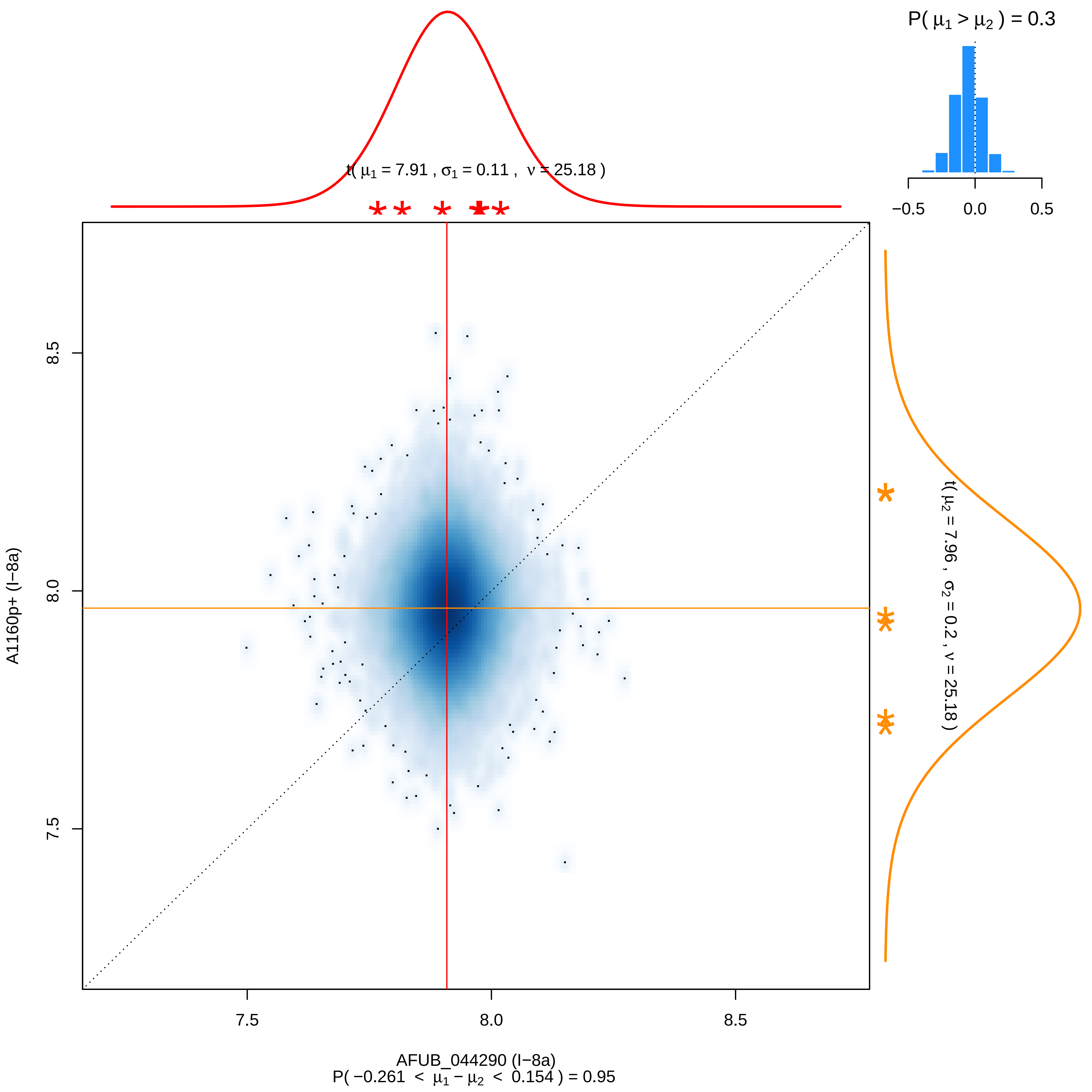

P( μ1 > μ2 ) = 0.3
−0.5
0.0
0.5
t( μ1 = 7.91 , σ1 = 0.11 , ν = 25.18 )
8.5
8.0
7.5
A1160p+ (I−8a)
t( μ2 = 7.96 , σ2 = 0.2 , ν = 25.18 )
7.5
8.0
8.5
AFUB_044290 (I−8a)
P( −0.261 < μ1 − μ2 < 0.154 ) = 0.95

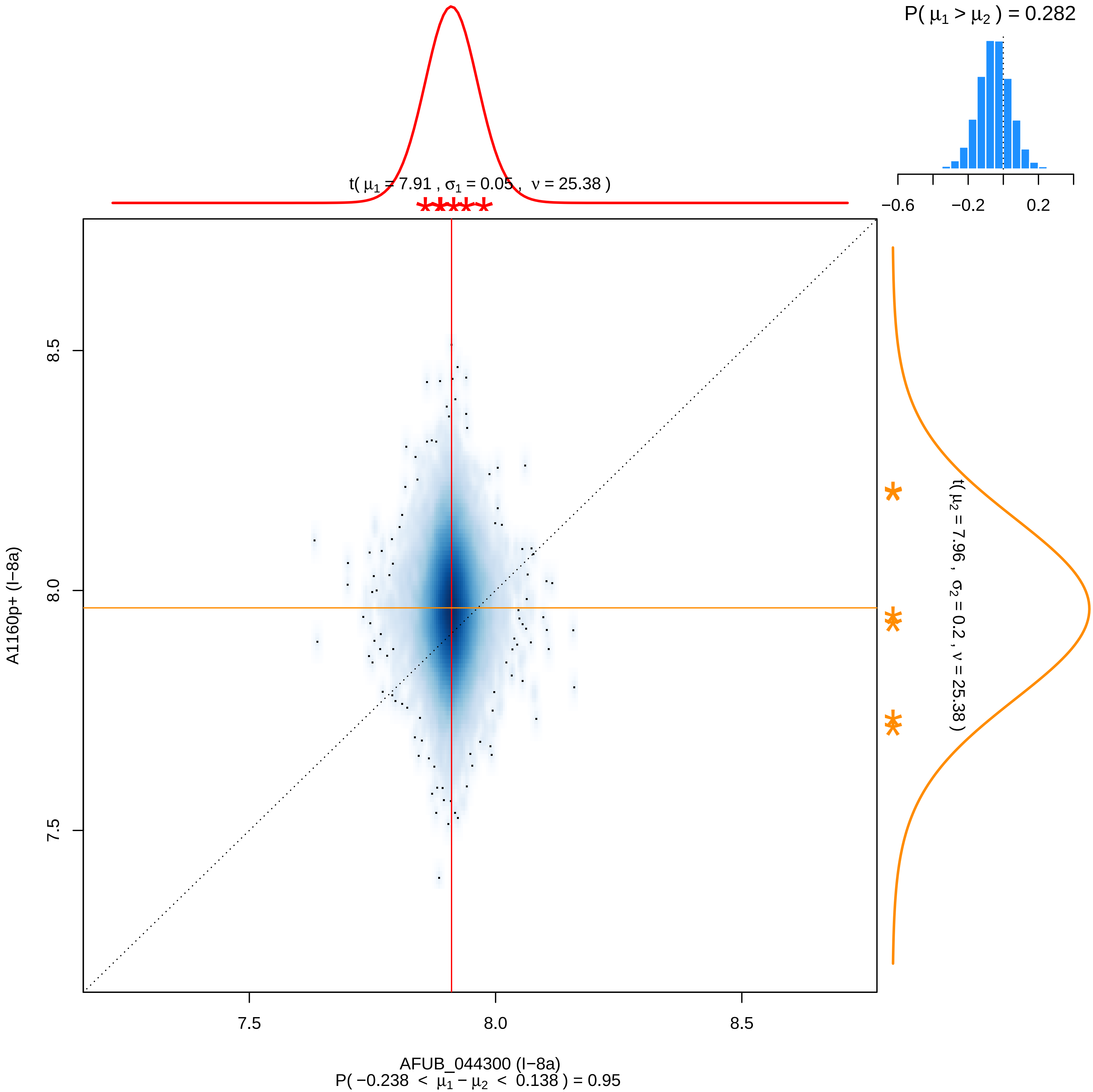

P( μ1 > μ2 ) = 0.282
−0.6
−0.2
0.2
t( μ1 = 7.91 , σ1 = 0.05 , ν = 25.38 )
t( μ2 = 7.96 , σ2 = 0.2 , ν = 25.38 )
A1160p+ (I−8a)
8.5
8.0
7.5
7.5
8.0
8.5
AFUB_044300 (I−8a)
P( −0.238 < μ1 − μ2 < 0.138 ) = 0.95

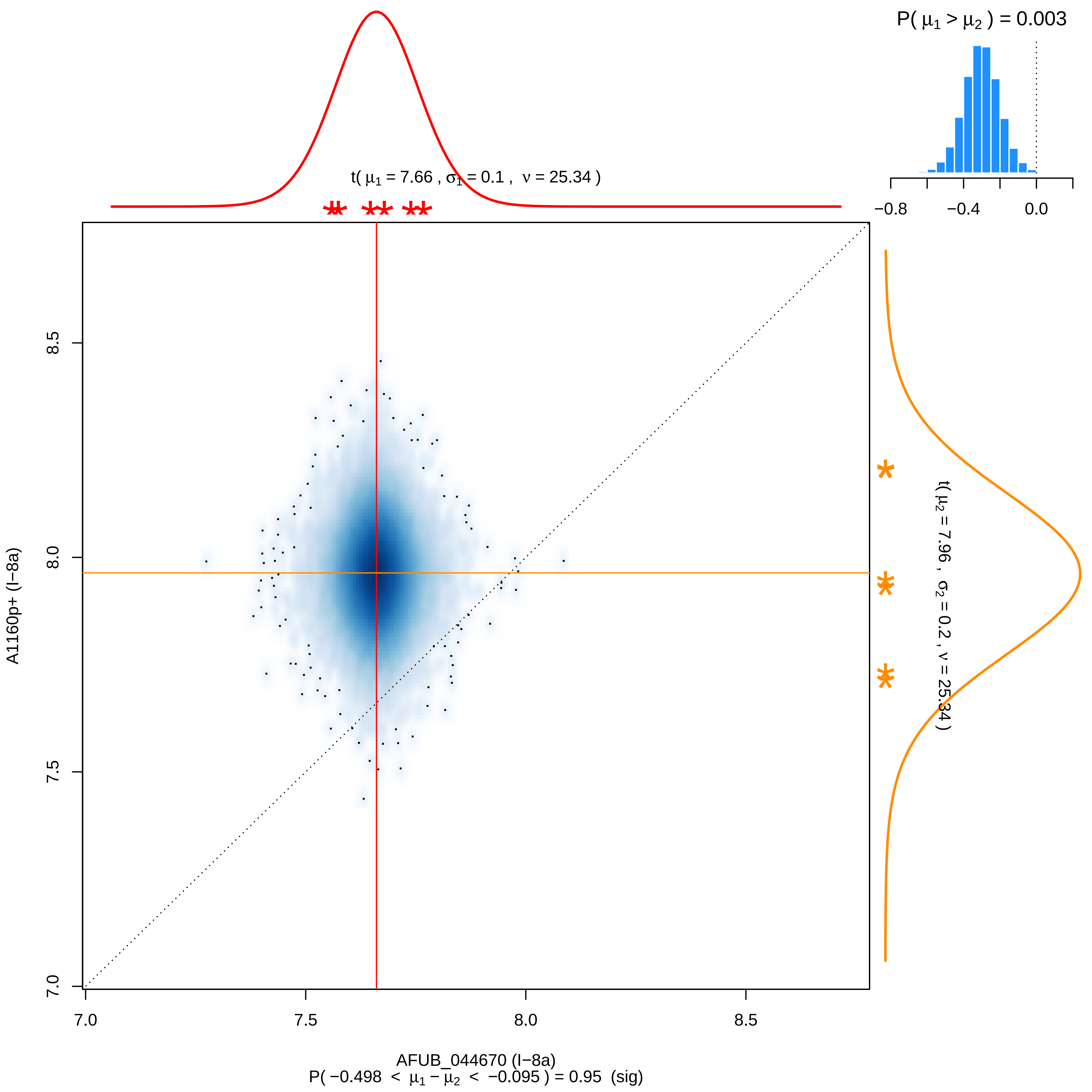

P( μ1 > μ2 ) = 0.003
−0.8
−0.4
0.0
t( μ1 = 7.66 , σ1 = 0.1 , ν = 25.34 )
t( μ2 = 7.96 , σ2 = 0.2 , ν = 25.34 )
A1160p+ (I−8a)
8.5
8.0
7.5
7.0
7.0
7.5
8.0
8.5
AFUB_044670 (I−8a)
P( −0.498 < μ1 − μ2 < −0.095 ) = 0.95 (sig)

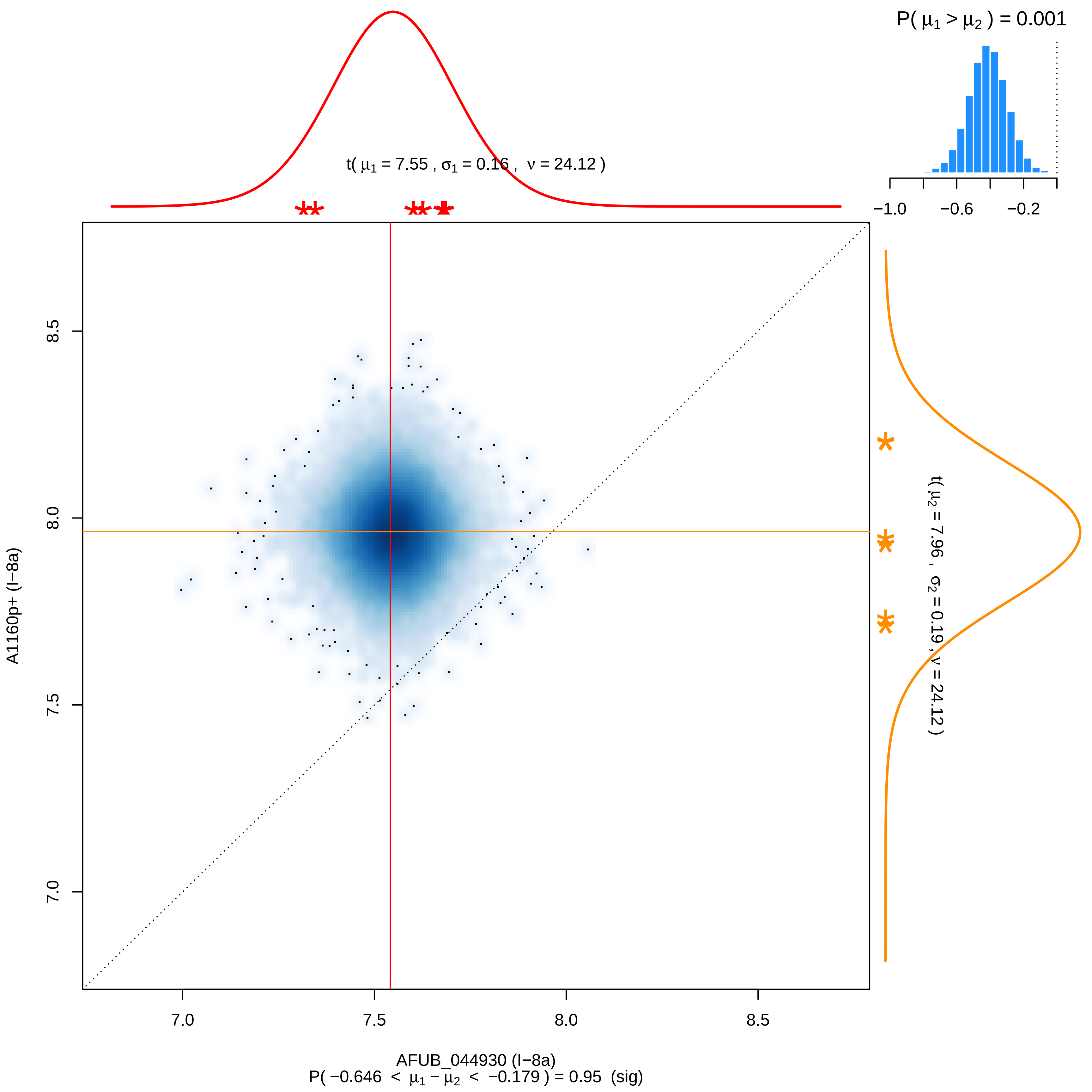

P( μ1 > μ2 ) = 0.001
−1.0
−0.6
−0.2
t( μ1 = 7.55 , σ1 = 0.16 , ν = 24.12 )
t( μ2 = 7.96 , σ2 = 0.19 , ν = 24.12 )
8.5
8.0
7.5
7.0
A1160p+ (I−8a)
7.0
7.5
8.0
8.5
AFUB_044930 (I−8a)
P( −0.646 < μ1 − μ2 < −0.179 ) = 0.95 (sig)

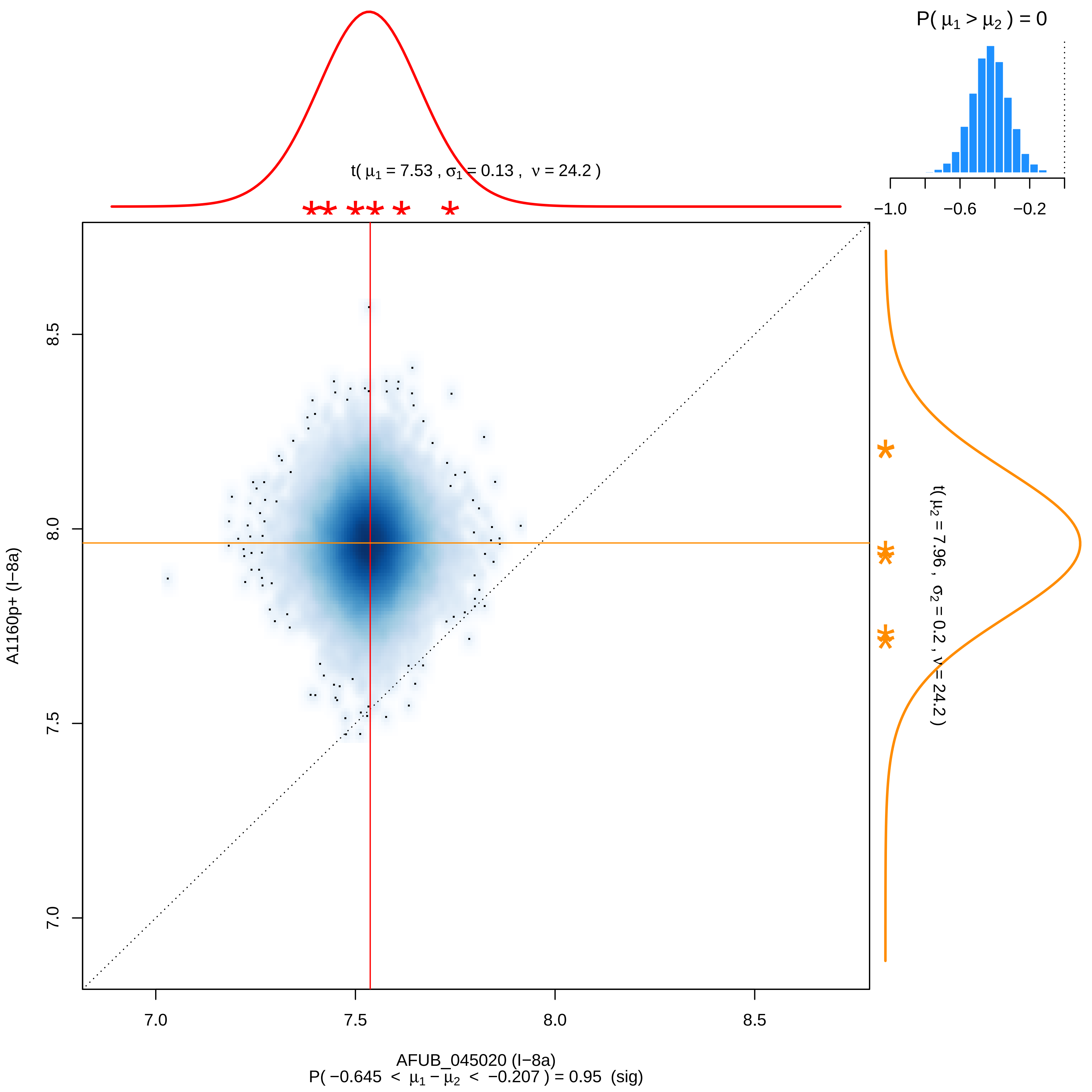

P( μ1 > μ2 ) = 0
−1.0
−0.6
−0.2
t( μ1 = 7.53 , σ1 = 0.13 , ν = 24.2 )
8.5
8.0
7.5
7.0
A1160p+ (I−8a)
t( μ2 = 7.96 , σ2 = 0.2 , ν = 24.2 )
7.0
7.5
8.0
8.5
AFUB_045020 (I−8a)
P( −0.645 < μ1 − μ2 < −0.207 ) = 0.95 (sig)

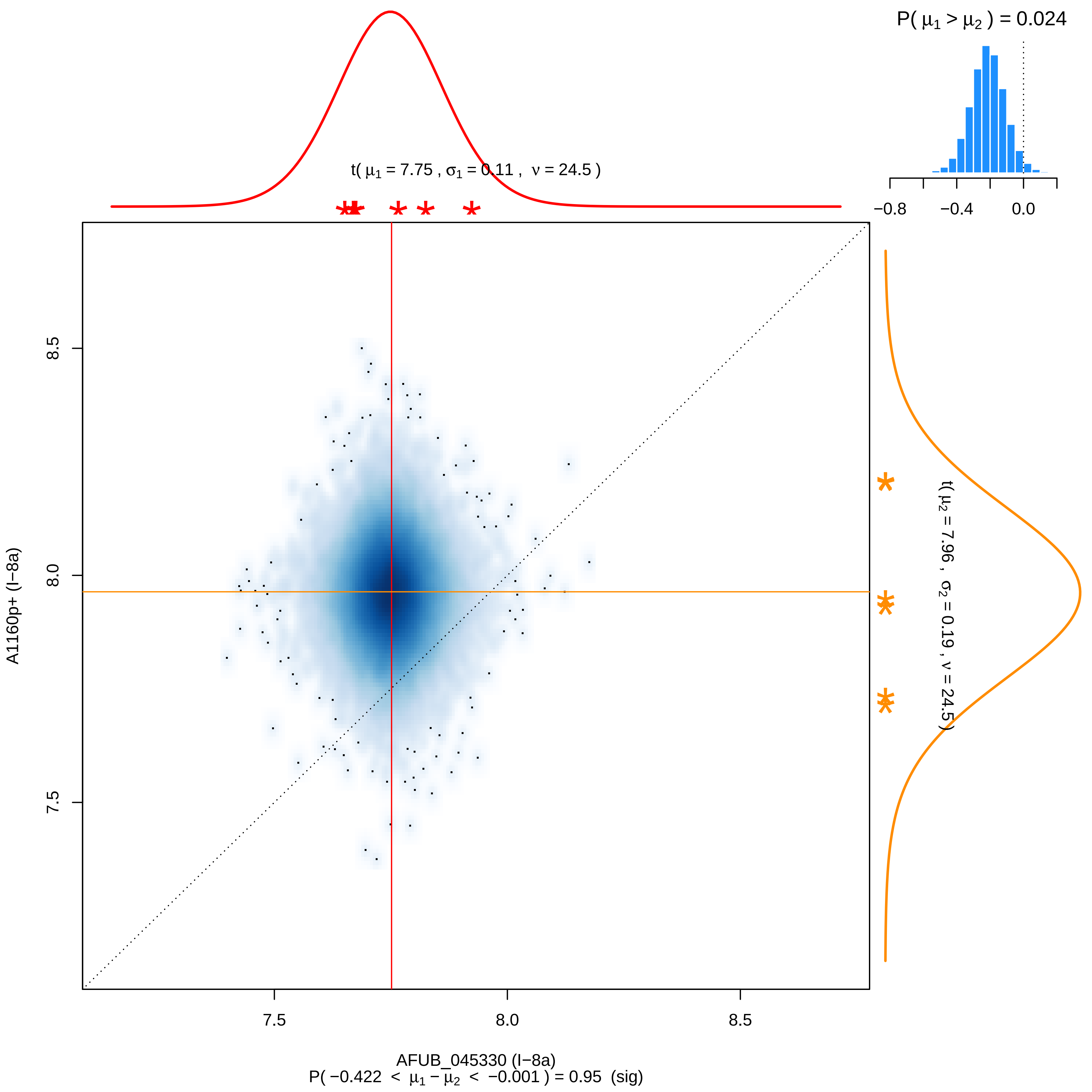

P( μ1 > μ2 ) = 0.024
−0.8
−0.4
0.0
t( μ1 = 7.75 , σ1 = 0.11 , ν = 24.5 )
t( μ2 = 7.96 , σ2 = 0.19 , ν = 24.5 )
8.5
8.0
7.5
A1160p+ (I−8a)
7.5
8.0
8.5
AFUB_045330 (I−8a)
P( −0.422 < μ1 − μ2 < −0.001 ) = 0.95 (sig)

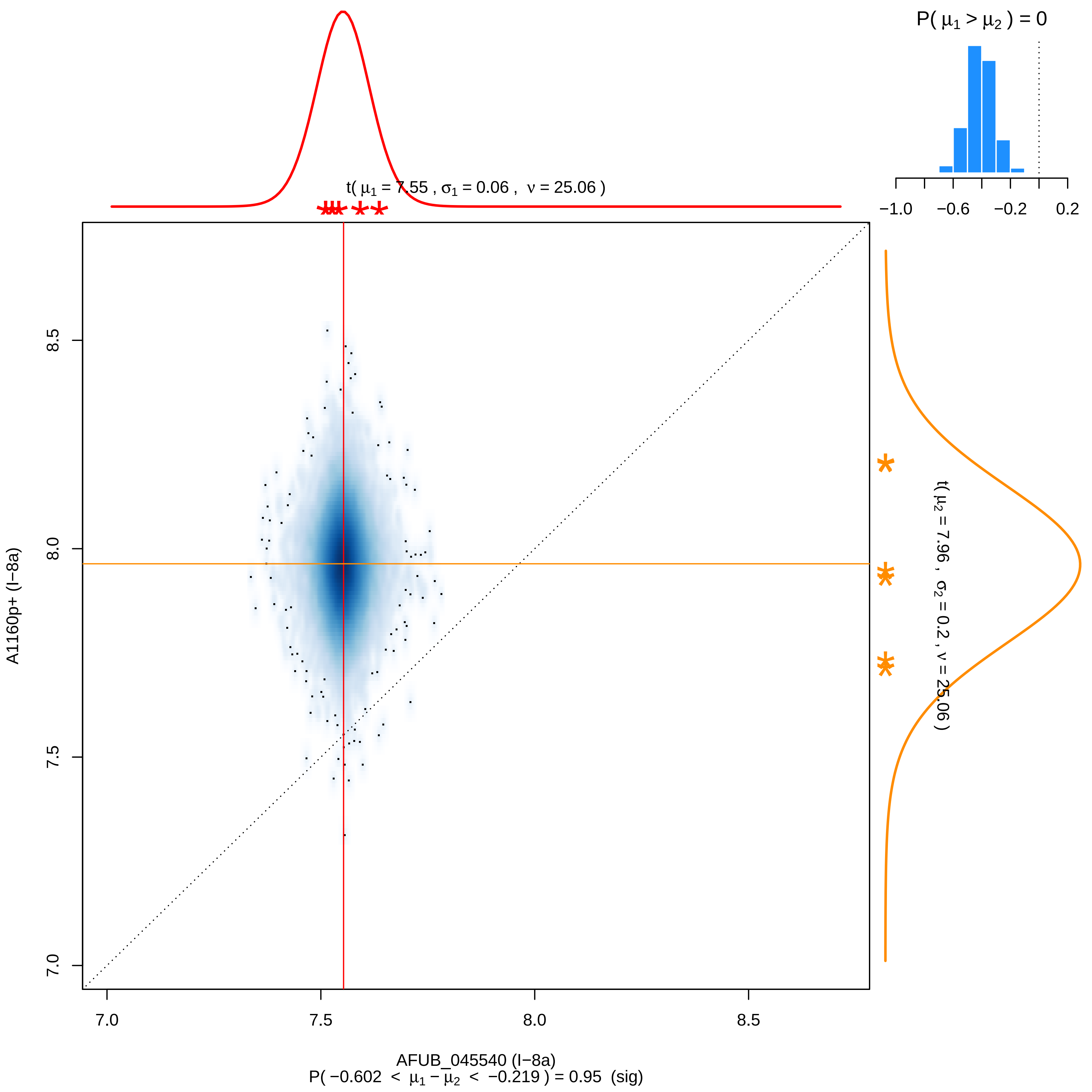

P( μ1 > μ2 ) = 0
−1.0
−0.6
−0.2
0.2
t( μ1 = 7.55 , σ1 = 0.06 , ν = 25.06 )
t( μ2 = 7.96 , σ2 = 0.2 , ν = 25.06 )
A1160p+ (I−8a)
7.0
7.5
8.0
8.5
AFUB_045540 (I−8a)
P( −0.602 < μ1 − μ2 < −0.219 ) = 0.95 (sig)

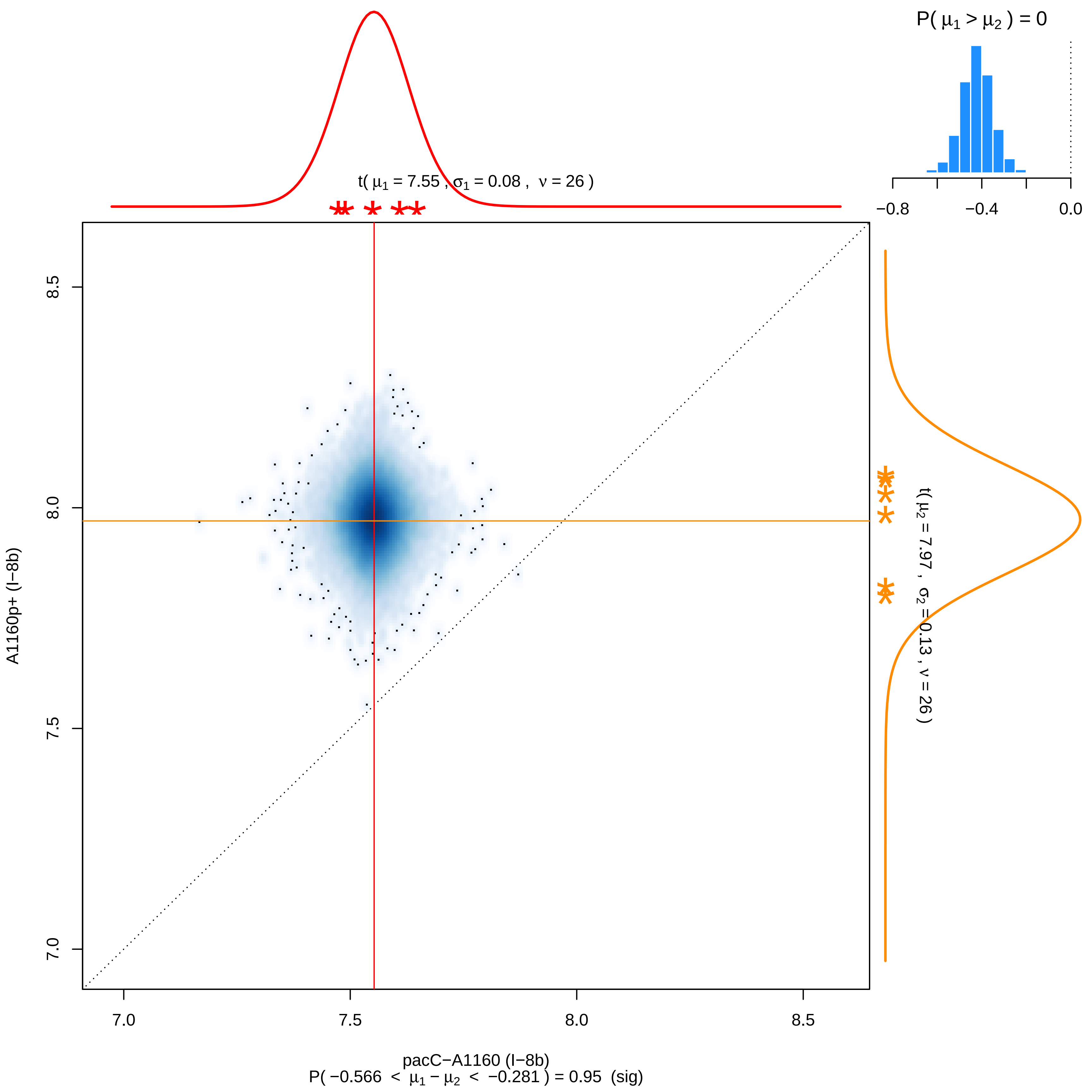

P( μ1 > μ2 ) = 0
−0.8
−0.4
0.0
t( μ1 = 7.55 , σ1 = 0.08 , ν = 26 )
t( μ2 = 7.97 , σ2 = 0.13 , ν = 26 )
8.5
8.0
7.5
7.0
A1160p+ (I−8b)
7.0
7.5
8.0
8.5
pacC−A1160 (I−8b)
P( −0.566 < μ1 − μ2 < −0.281 ) = 0.95 (sig)

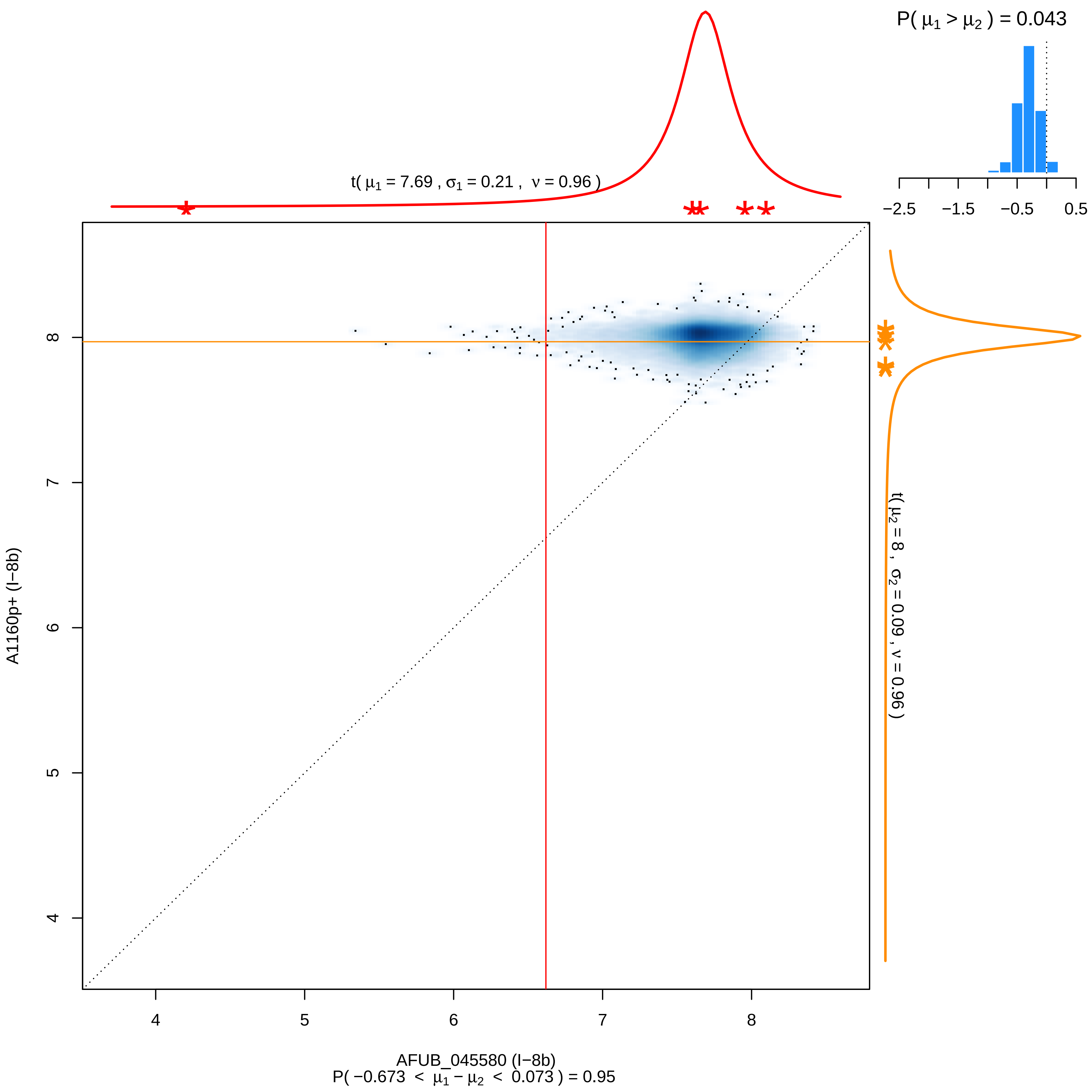

P( μ1 > μ2 ) = 0.043
−2.5
−1.5
−0.5
0.5
t( μ1 = 7.69 , σ1 = 0.21 , ν = 0.96 )
t( μ2 = 8 , σ2 = 0.09 , ν = 0.96 )
A1160p+ (I−8b)
4
5
6
7
8
AFUB_045580 (I−8b)
P( −0.673 < μ1 − μ2 < 0.073 ) = 0.95

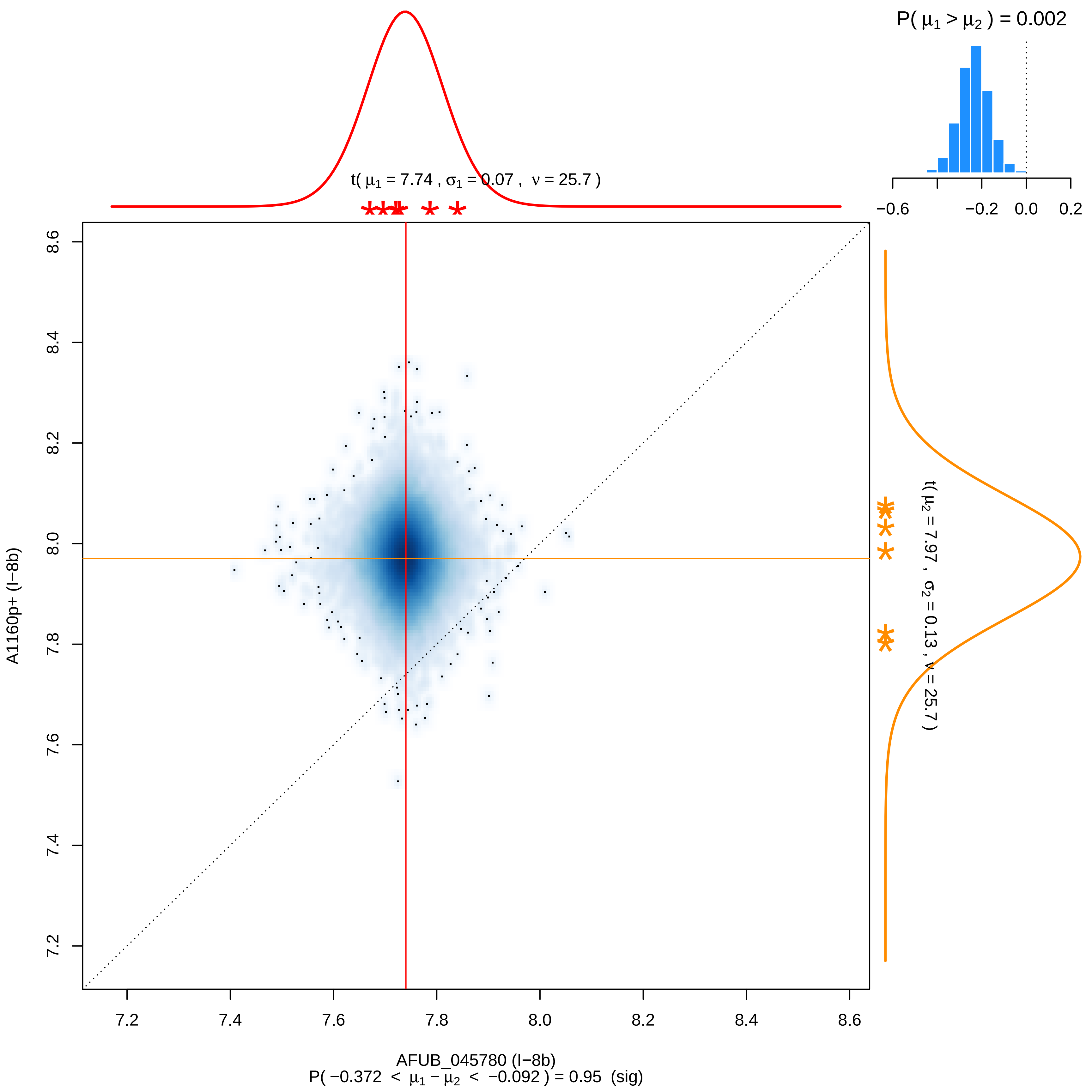

P( $\mu_1 > \mu_2$ ) = 0.002
−0.6
−0.2
0.0
0.2
t( $\mu_1$ = 7.74 , $\sigma_1$ = 0.07 , $\nu$ = 25.7 )
t( $\mu_2$ = 7.97 , $\sigma_2$ = 0.13 , $\nu$ = 25.7 )
A1160p+ (I−8b)
8.6
8.4
8.2
8.0
7.8
7.6
7.4
7.2
7.2
7.4
7.6
7.8
8.0
8.2
8.4
8.6
AFUB_045780 (I−8b)
P( −0.372 < $\mu_1 - \mu_2$ < −0.092 ) = 0.95 (sig)

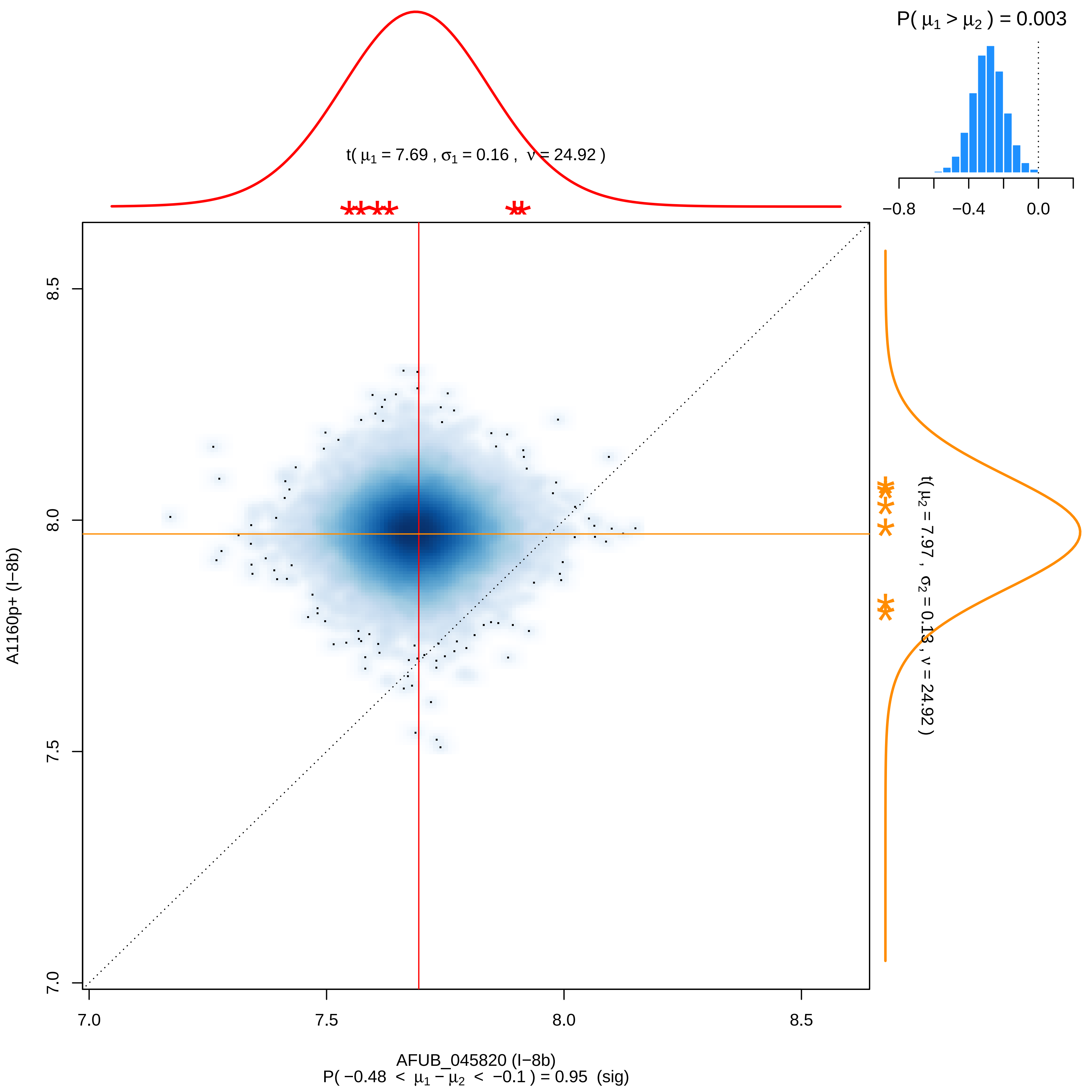
P( μ1 > μ2 ) = 0.003
−0.8
−0.4
0.0
t( μ1 = 7.69 , σ1 = 0.16 , ν = 24.92 )
t( μ2 = 7.97 , σ2 = 0.13 , ν = 24.92 )
8.5
8.0
7.5
7.0
A1160p+ (I−8b)
7.0
7.5
8.0
8.5
AFUB_045820 (I−8b)
P( −0.48 < μ1 − μ2 < −0.1 ) = 0.95 (sig)

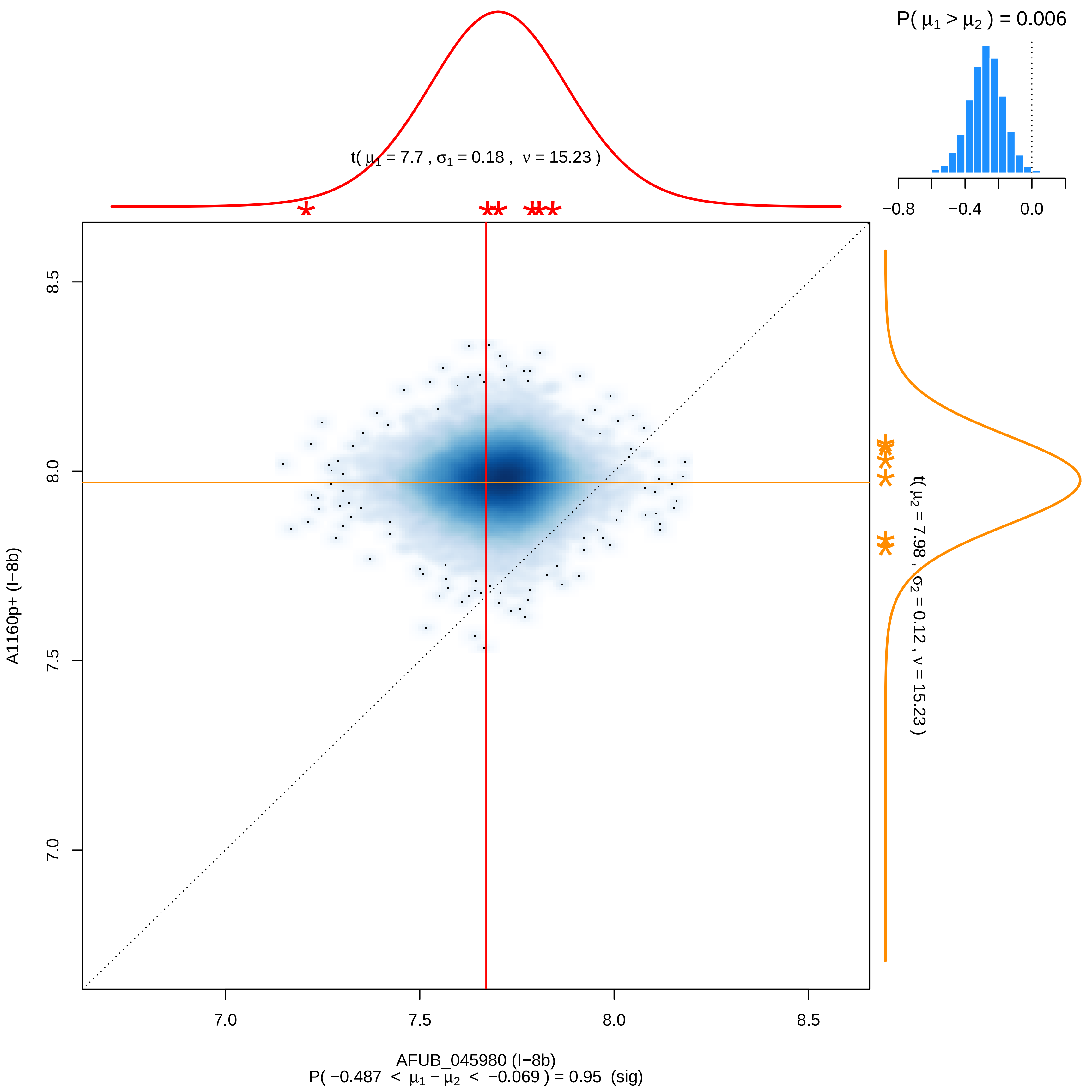

P( μ1 > μ2 ) = 0.006
−0.8
−0.4
0.0
t( μ1 = 7.7 , σ1 = 0.18 , ν = 15.23 )
8.5
8.0
7.5
7.0
A1160p+ (I−8b)
t( μ2 = 7.98 , σ2 = 0.12 , ν = 15.23 )
7.0
7.5
8.0
8.5
AFUB_045980 (I−8b)
P( −0.487 < μ1 − μ2 < −0.069 ) = 0.95 (sig)

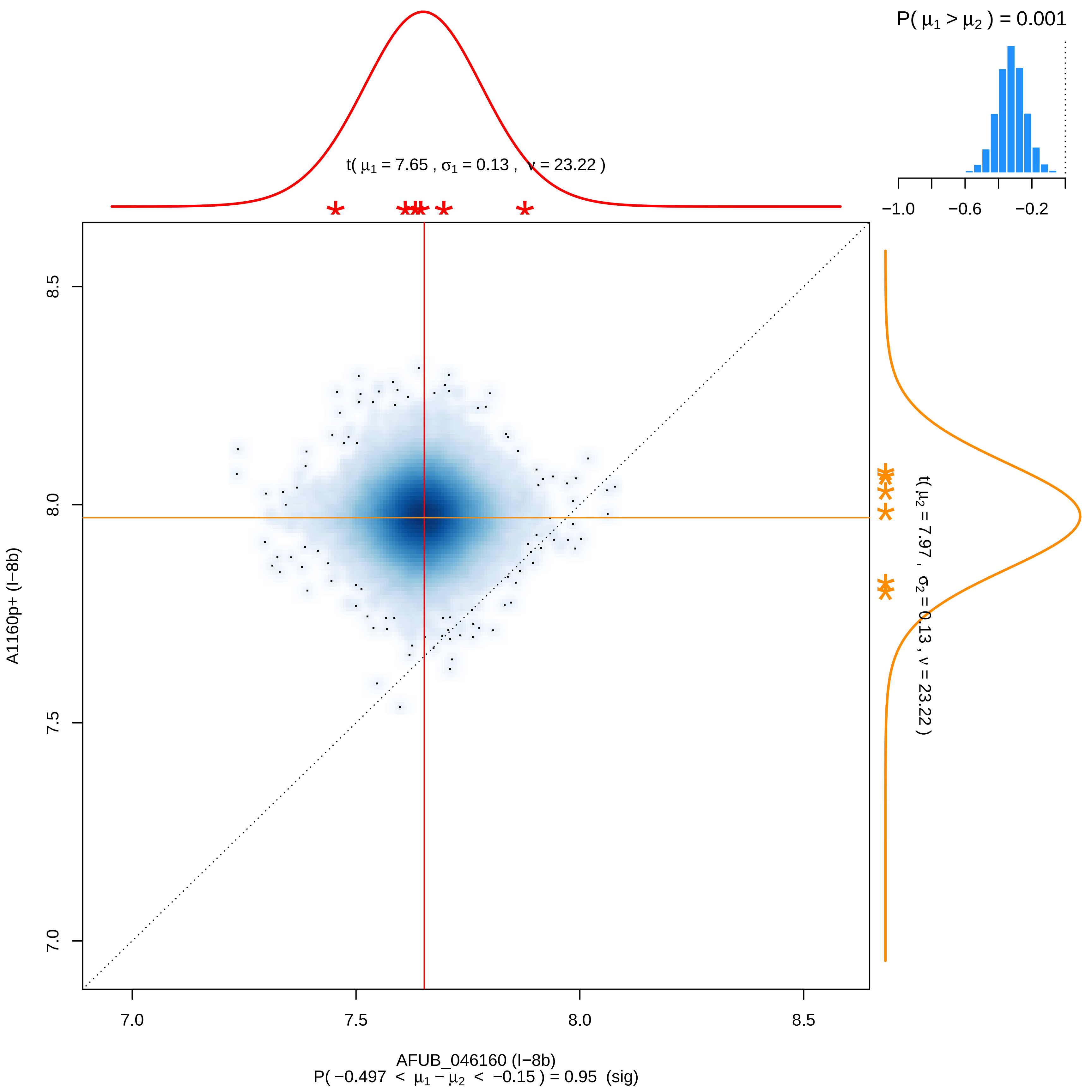

P( μ1 > μ2 ) = 0.001
−1.0
−0.6
−0.2
t( μ1 = 7.65 , σ1 = 0.13 , ν = 23.22 )
t( μ2 = 7.97 , σ2 = 0.13 , ν = 23.22 )
8.5
8.0
7.5
7.0
A1160p+ (I−8b)
7.0
7.5
8.0
8.5
AFUB_046160 (I−8b)
P( −0.497 < μ1 − μ2 < −0.15 ) = 0.95 (sig)

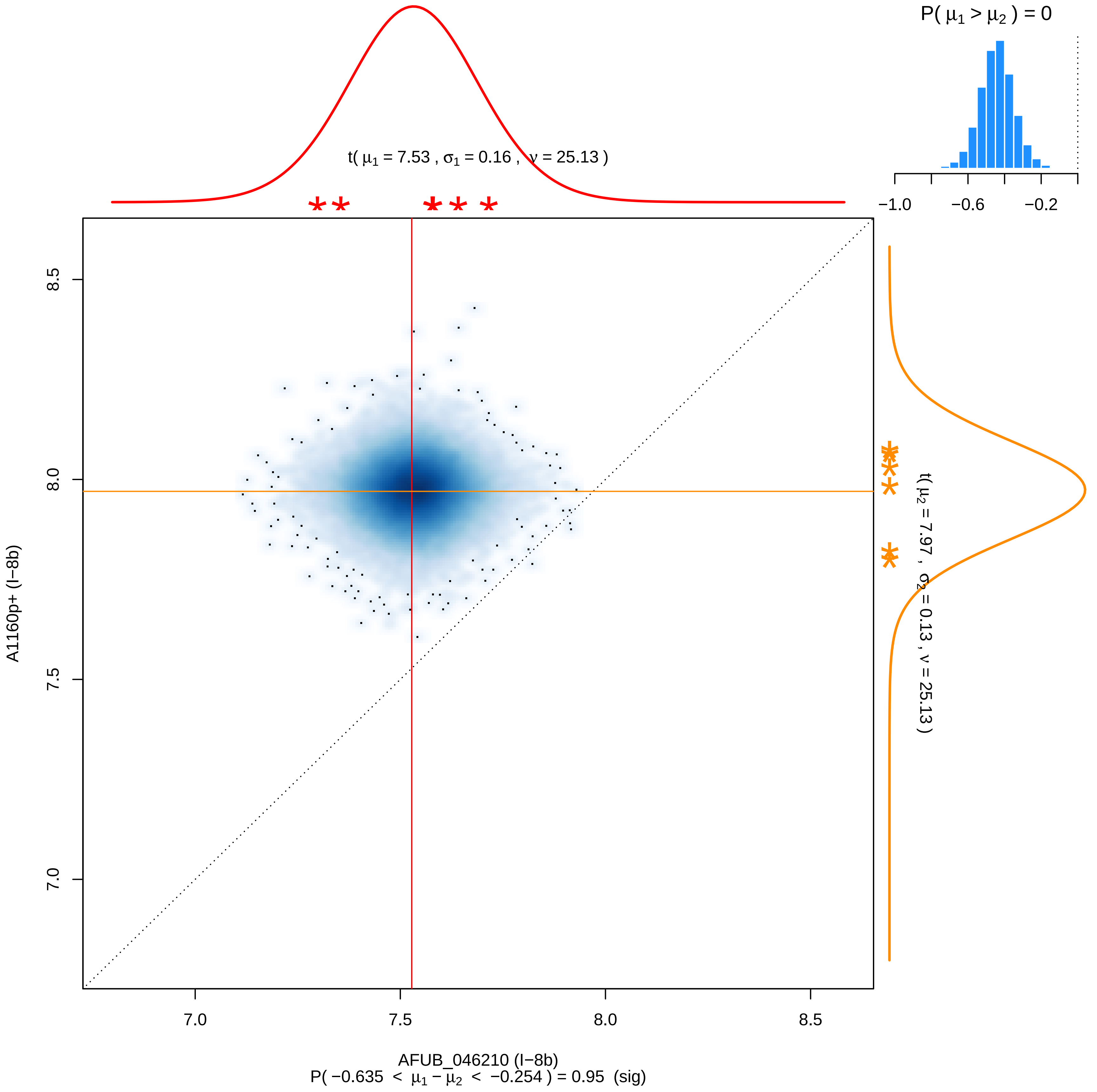

P( μ1 > μ2 ) = 0
−1.0
−0.6
−0.2
t( μ1 = 7.53 , σ1 = 0.16 , ν = 25.13 )
8.5
8.0
7.5
7.0
A1160p+ (I−8b)
t( μ2 = 7.97 , σ2 = 0.13 , ν = 25.13 )
7.0
7.5
8.0
8.5
AFUB_046210 (I−8b)
P( −0.635 < μ1 − μ2 < −0.254 ) = 0.95 (sig)

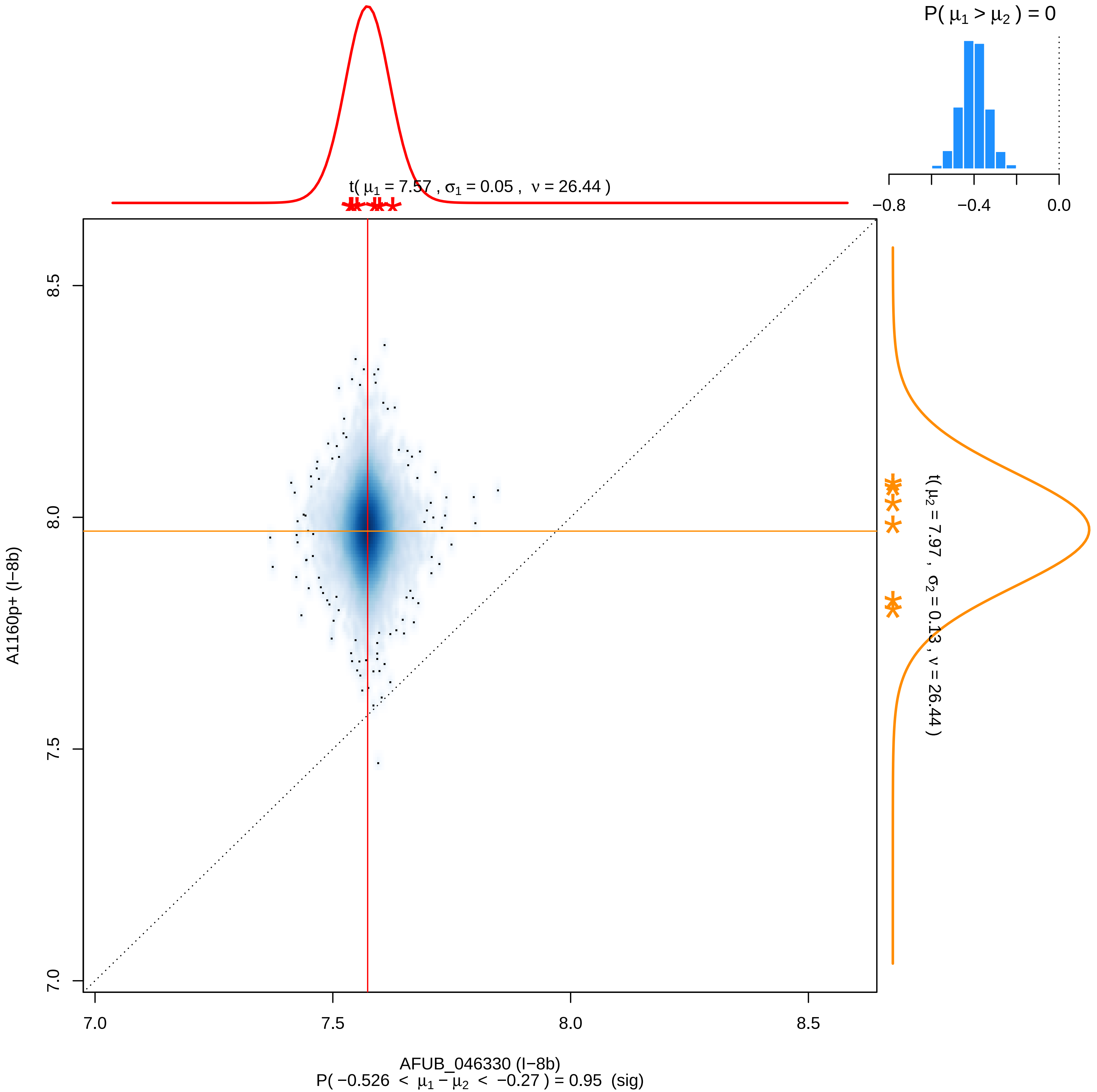

P( μ1 > μ2 ) = 0
−0.8
−0.4
0.0
t( μ1 = 7.57 , σ1 = 0.05 , ν = 26.44 )
t( μ2 = 7.97 , σ2 = 0.13 , ν = 26.44 )
A1160p+ (I−8b)
8.5
8.0
7.5
7.0
AFUB_046330 (I−8b)
P( −0.526 < μ1 − μ2 < −0.27 ) = 0.95 (sig)

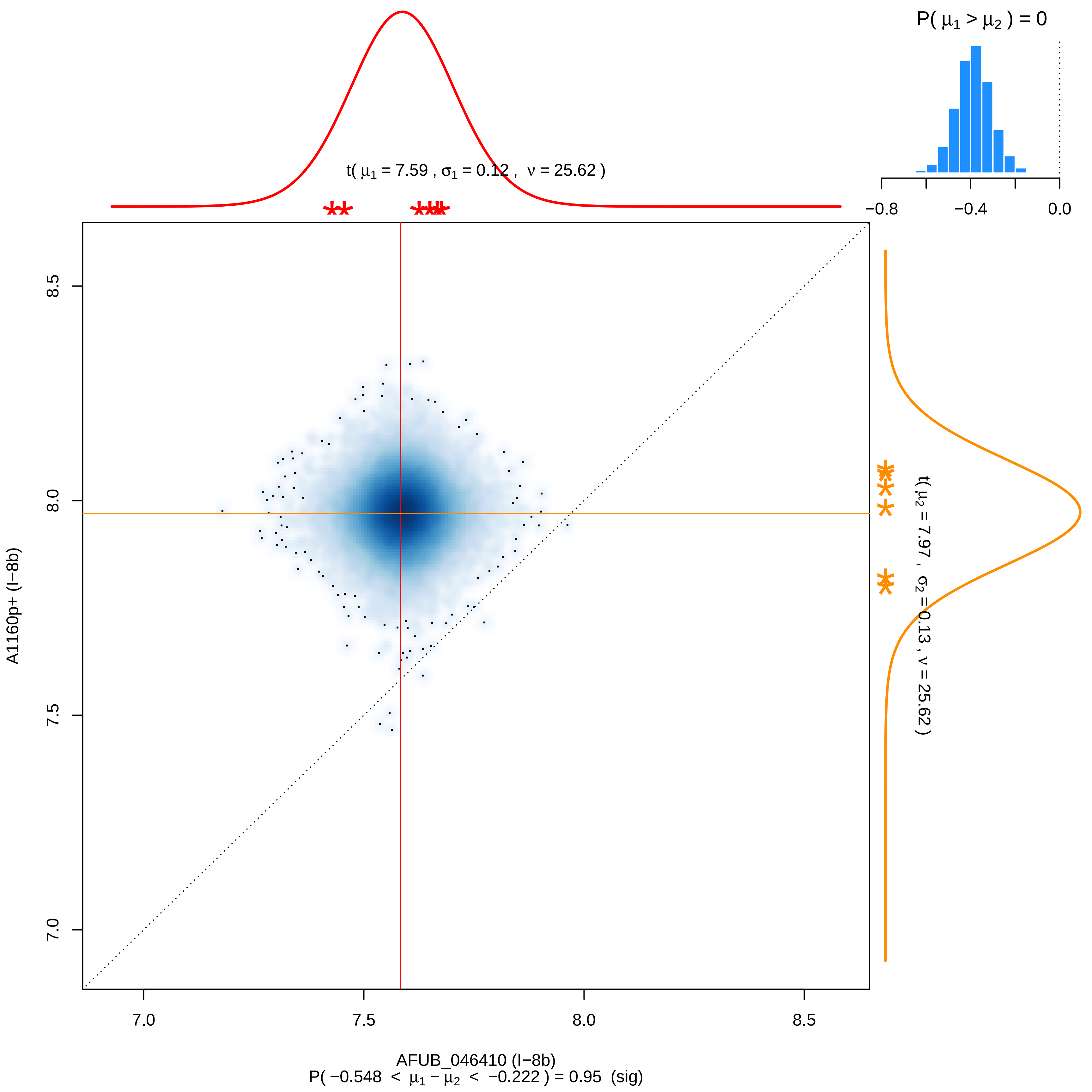

P( μ1 > μ2 ) = 0
−0.8
−0.4
0.0
t( μ1 = 7.59 , σ1 = 0.12 , ν = 25.62 )
t( μ2 = 7.97 , σ2 = 0.13 , ν = 25.62 )
8.5
8.0
7.5
7.0
A1160p+ (I−8b)
7.0
7.5
8.0
8.5
AFUB_046410 (I−8b)
P( −0.548 < μ1 − μ2 < −0.222 ) = 0.95 (sig)

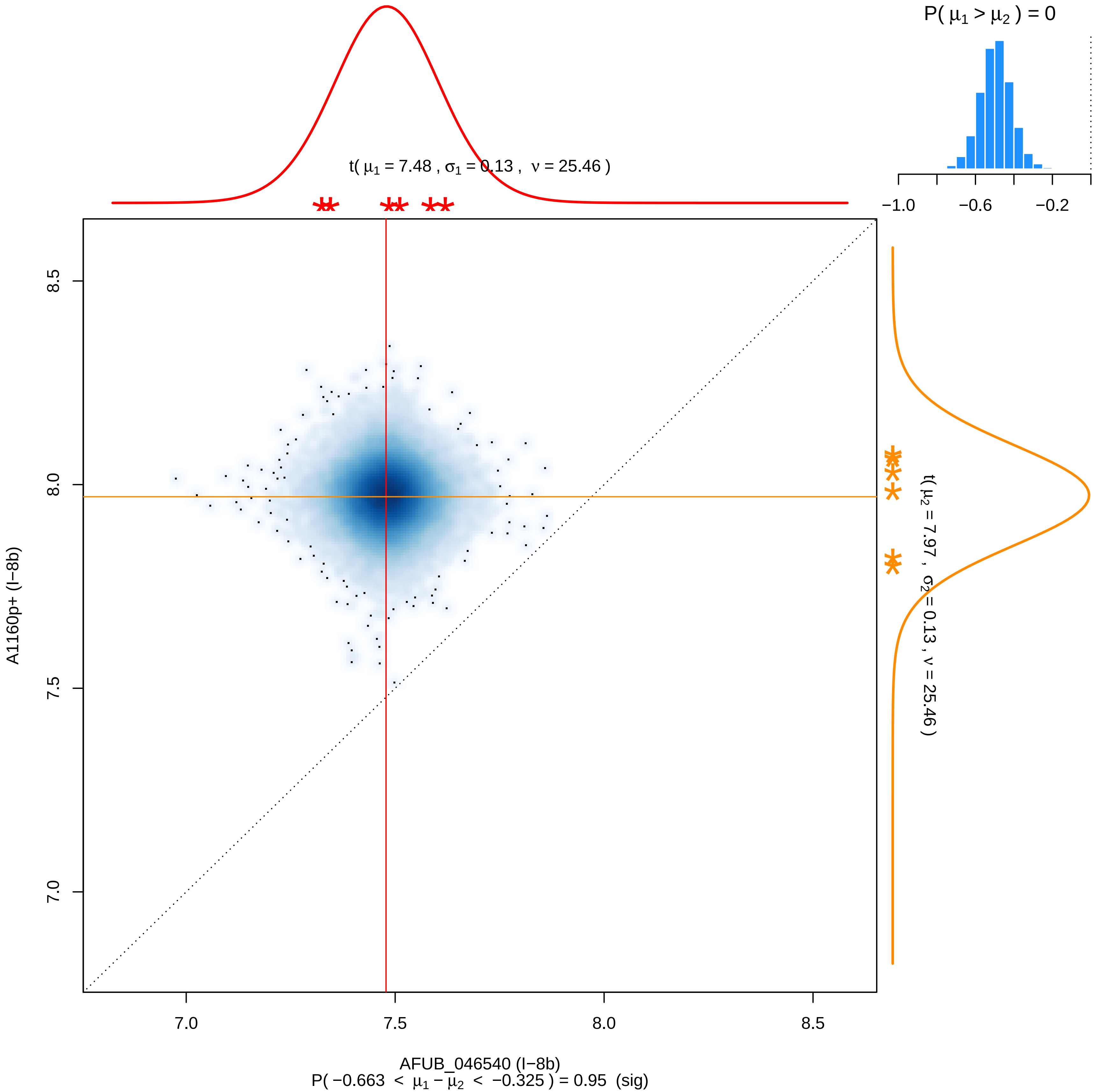

P( $\mu_1 > \mu_2$ ) = 0
−1.0
−0.6
−0.2
t( $\mu_1$ = 7.48 , $\sigma_1$ = 0.13 , $\nu$ = 25.46 )
t( $\mu_2$ = 7.97 , $\sigma_2$ = 0.13 , $\nu$ = 25.46 )
A1160p+ (I−8b)
8.5
8.0
7.5
7.0
AFUB_046540 (I−8b)
P( −0.663 < $\mu_1 - \mu_2$ < −0.325 ) = 0.95 (sig)

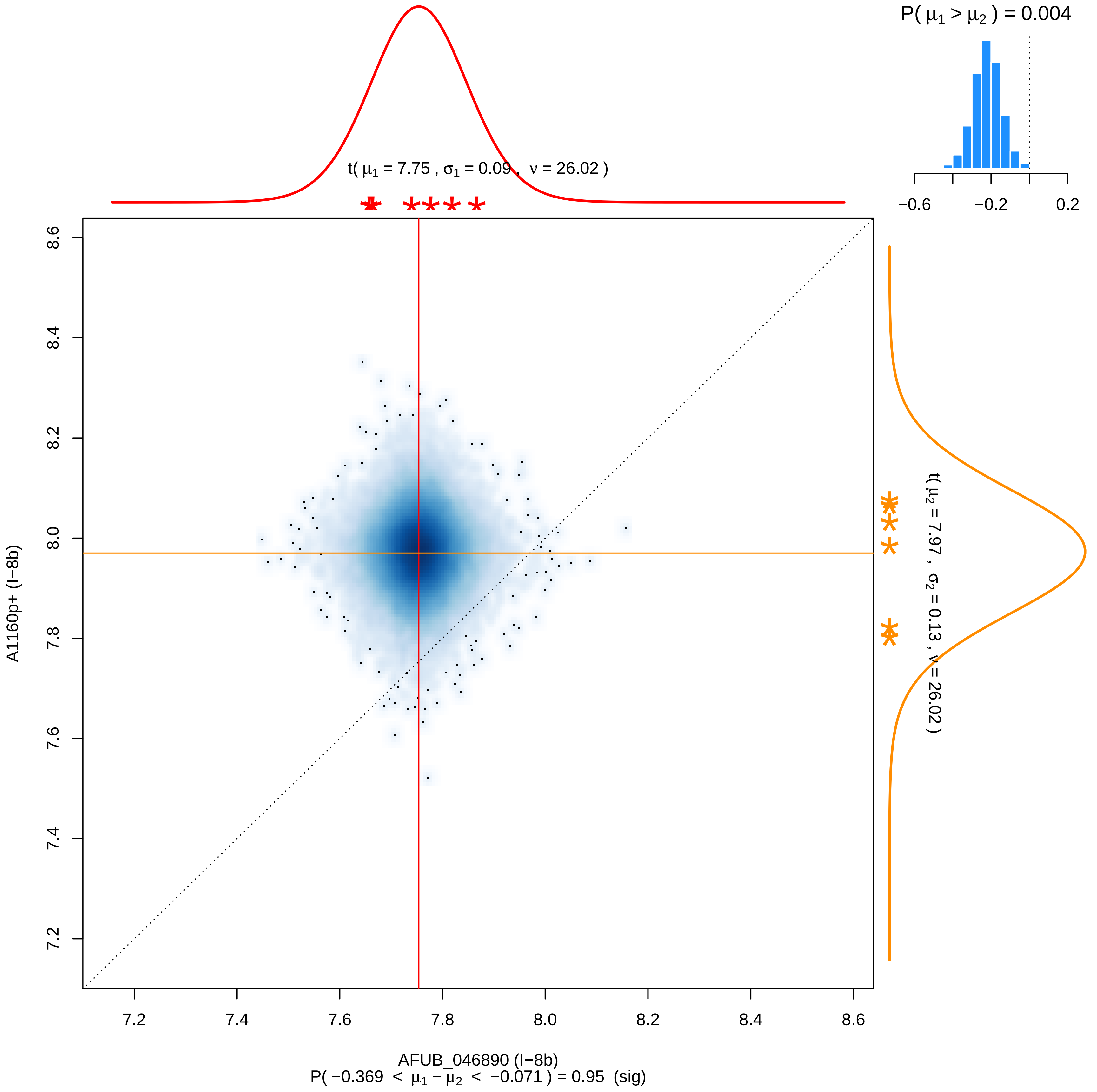

P( μ1 > μ2 ) = 0.004
−0.6
−0.2
0.2
t( μ1 = 7.75 , σ1 = 0.09 , ν = 26.02 )
t( μ2 = 7.97 , σ2 = 0.13 , ν = 26.02 )
A1160p+ (I−8b)
8.6
8.4
8.2
8.0
7.8
7.6
7.4
7.2
AFUB_046890 (I−8b)
P( −0.369 < μ1 − μ2 < −0.071 ) = 0.95 (sig)

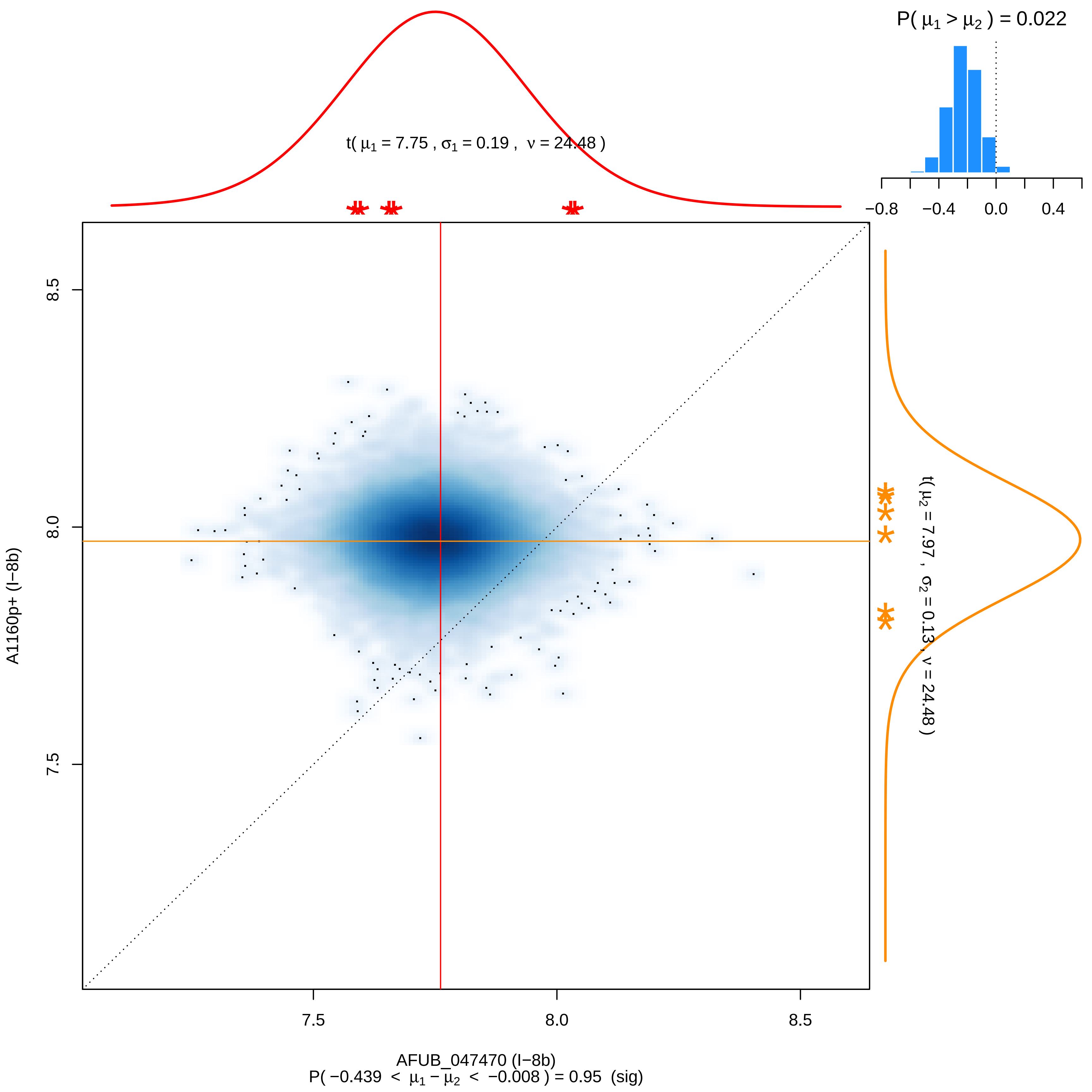

P( μ1 > μ2 ) = 0.022
−0.8
−0.4
0.0
0.4
t( μ1 = 7.75 , σ1 = 0.19 , ν = 24.48 )
t( μ2 = 7.97 , σ2 = 0.13 , ν = 24.48 )
8.5
8.0
7.5
A1160p+ (I−8b)
7.5
8.0
8.5
AFUB_047470 (I−8b)
P( −0.439 < μ1 − μ2 < −0.008 ) = 0.95 (sig)

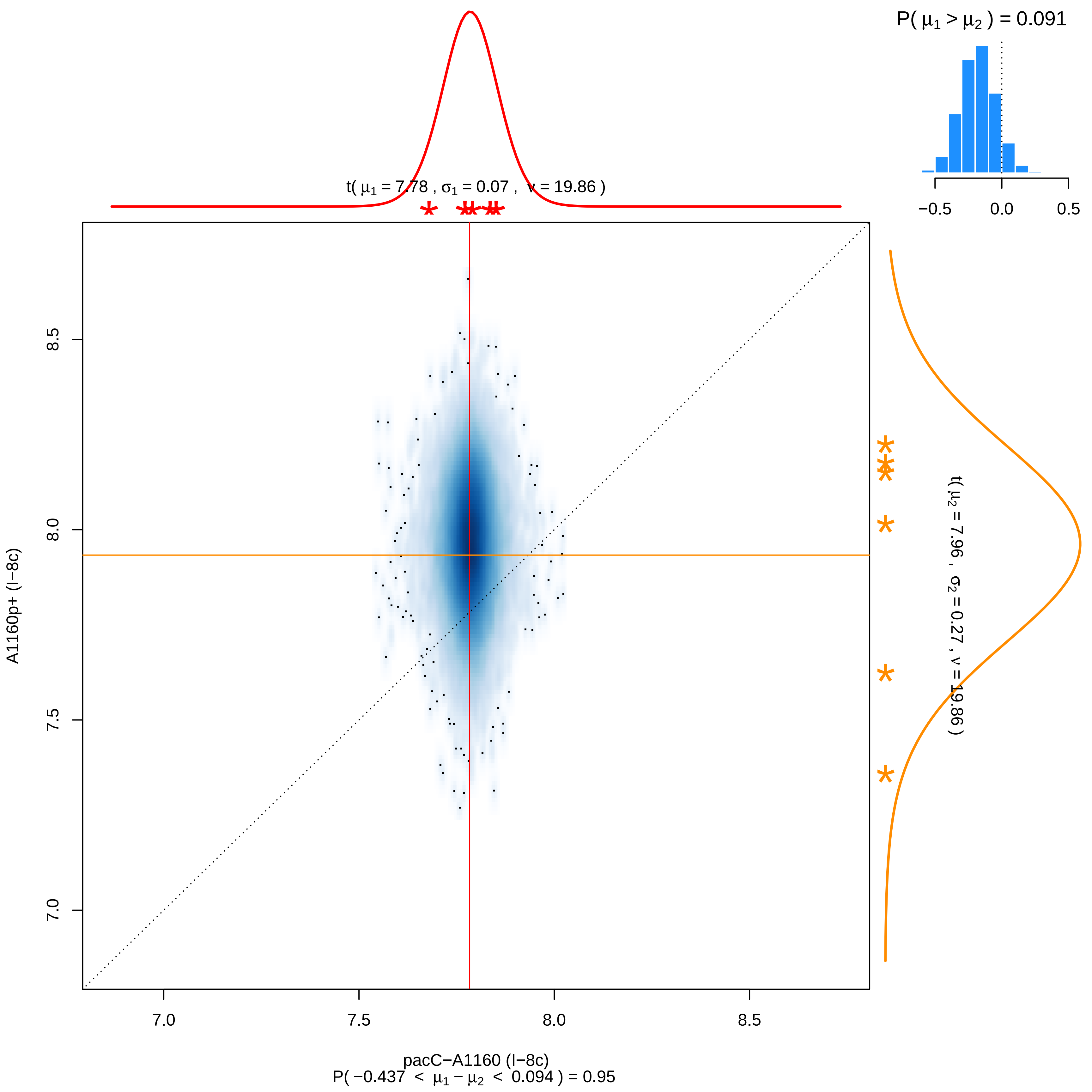

P( μ1 > μ2 ) = 0.091
−0.5
0.0
0.5
t( μ1 = 7.78 , σ1 = 0.07 , ν = 19.86 )
t( μ2 = 7.96 , σ2 = 0.27 , ν = 19.86 )
A1160p+ (I−8c)
7.0
7.5
8.0
8.5
pacC−A1160 (I−8c)
P( −0.437 < μ1 − μ2 < 0.094 ) = 0.95

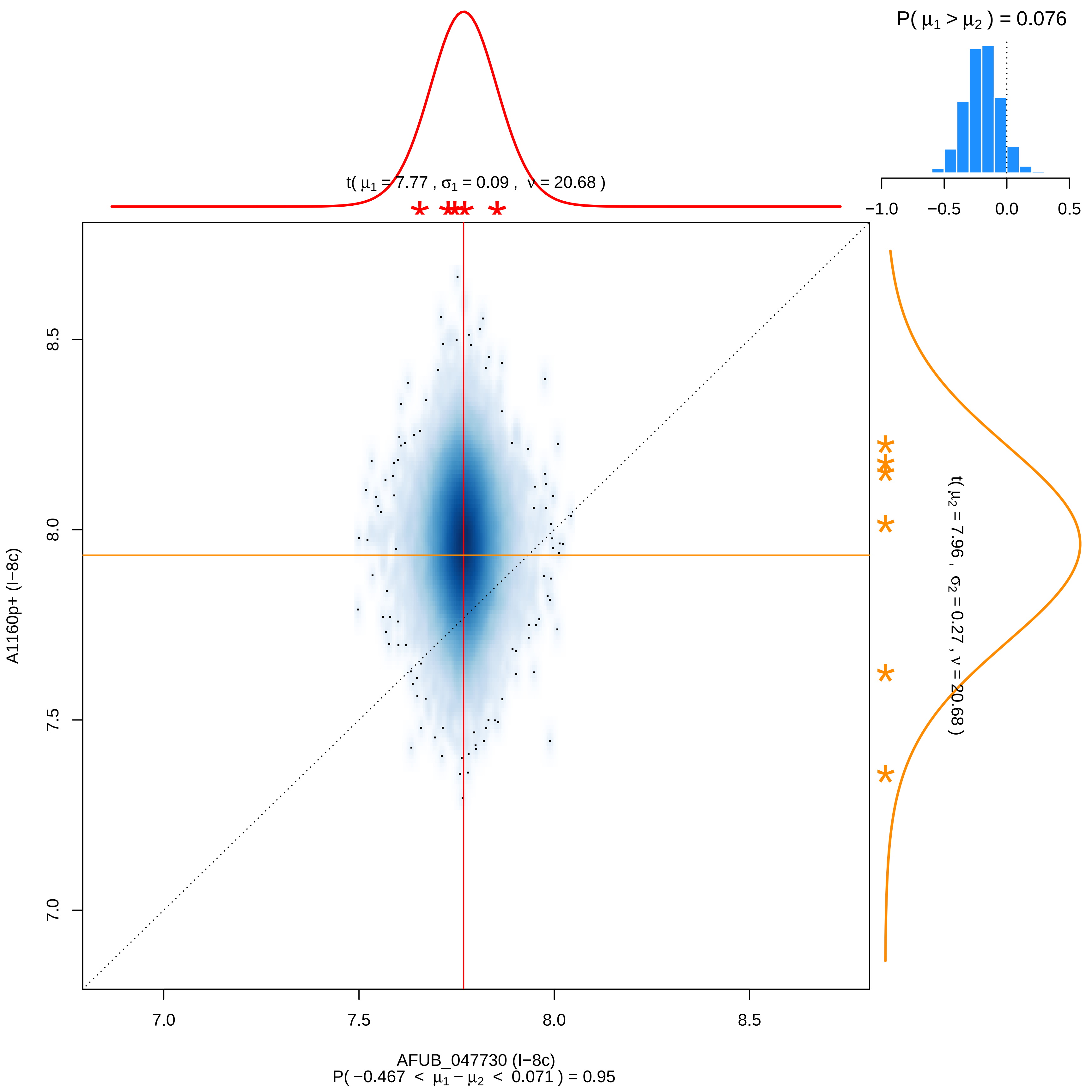

P( μ1 > μ2 ) = 0.076
−1.0
−0.5
0.0
0.5
t( μ1 = 7.77 , σ1 = 0.09 , ν = 20.68 )
8.5
8.0
7.5
7.0
A1160p+ (I−8c)
t( μ2 = 7.96 , σ2 = 0.27 , ν = 20.68 )
AFUB_047730 (I−8c)
P( −0.467 < μ1 − μ2 < 0.071 ) = 0.95

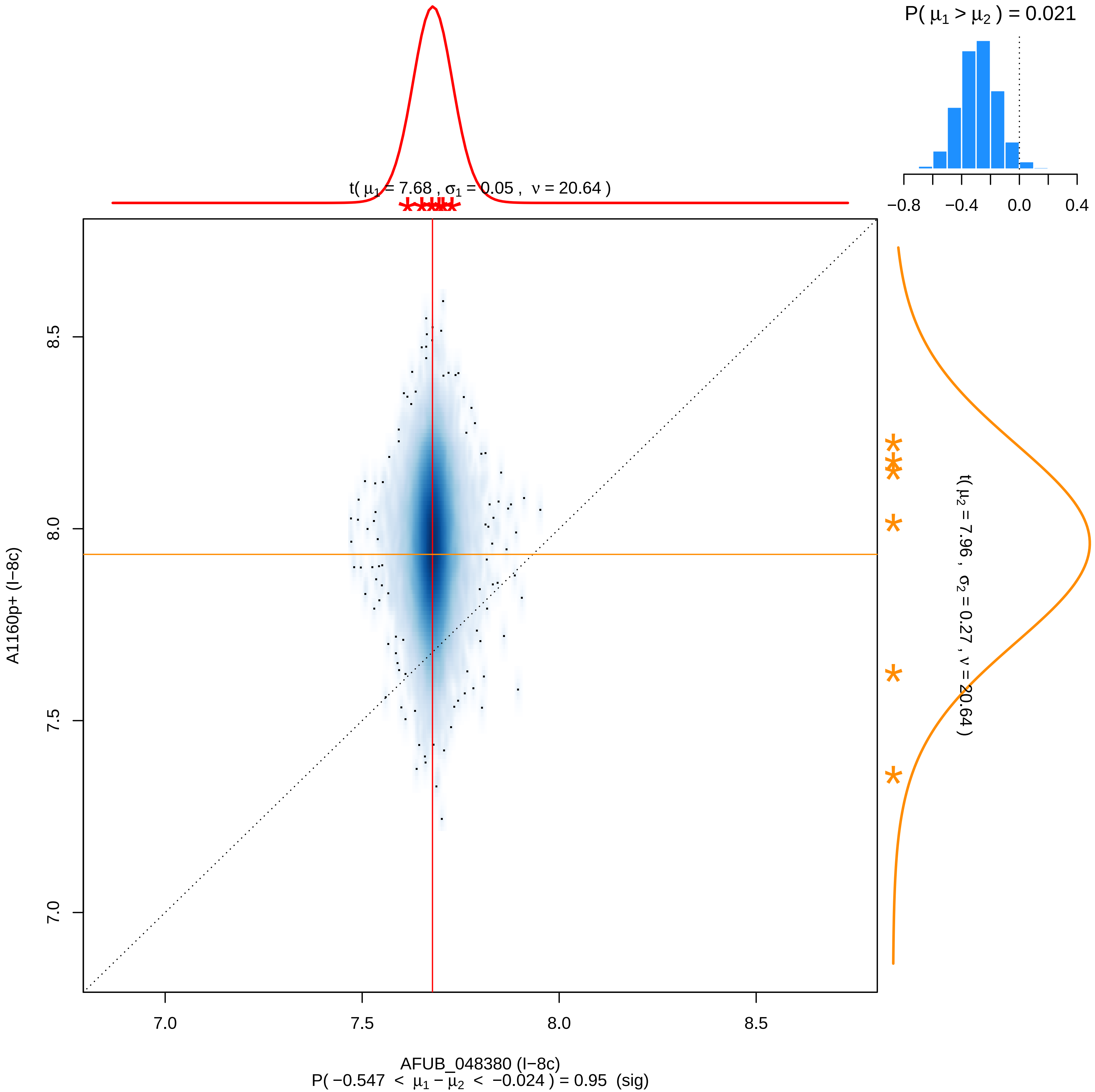

P( μ1 > μ2 ) = 0.021
−0.8
−0.4
0.0
0.4
t( μ1 = 7.68 , σ1 = 0.05 , ν = 20.64 )
t( μ2 = 7.96 , σ2 = 0.27 , ν = 20.64 )
A1160p+ (I−8c)
7.0
7.5
8.0
8.5
AFUB_048380 (I−8c)
P( −0.547 < μ1 − μ2 < −0.024 ) = 0.95 (sig)

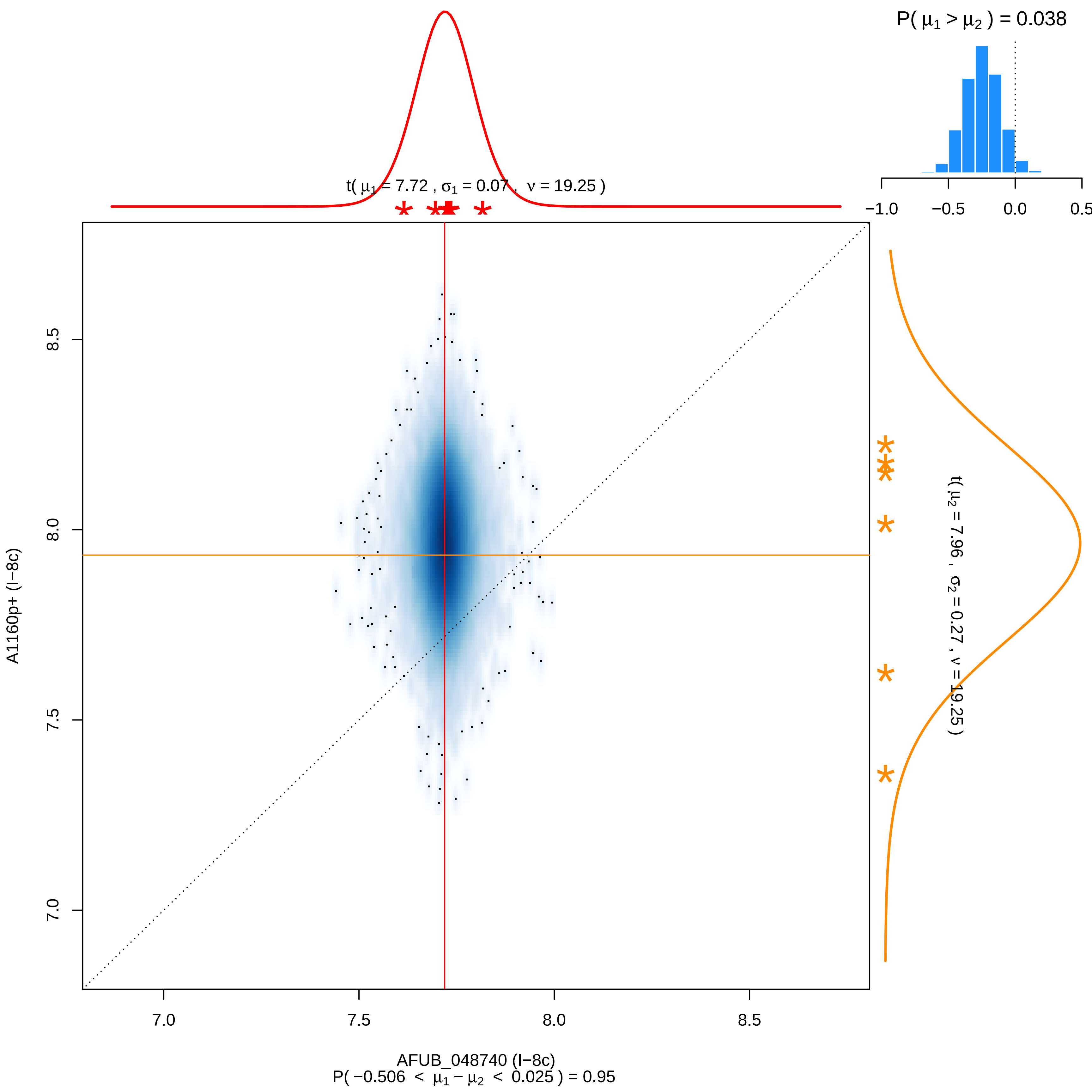

P( μ1 > μ2 ) = 0.038
−1.0
−0.5
0.0
0.5
t( μ1 = 7.72 , σ1 = 0.07 , ν = 19.25 )
t( μ2 = 7.96 , σ2 = 0.27 , ν = 19.25 )
8.5
8.0
7.5
7.0
A1160p+ (I−8c)
7.0
7.5
8.0
8.5
AFUB_048740 (I−8c)
P( −0.506 < μ1 − μ2 < 0.025 ) = 0.95

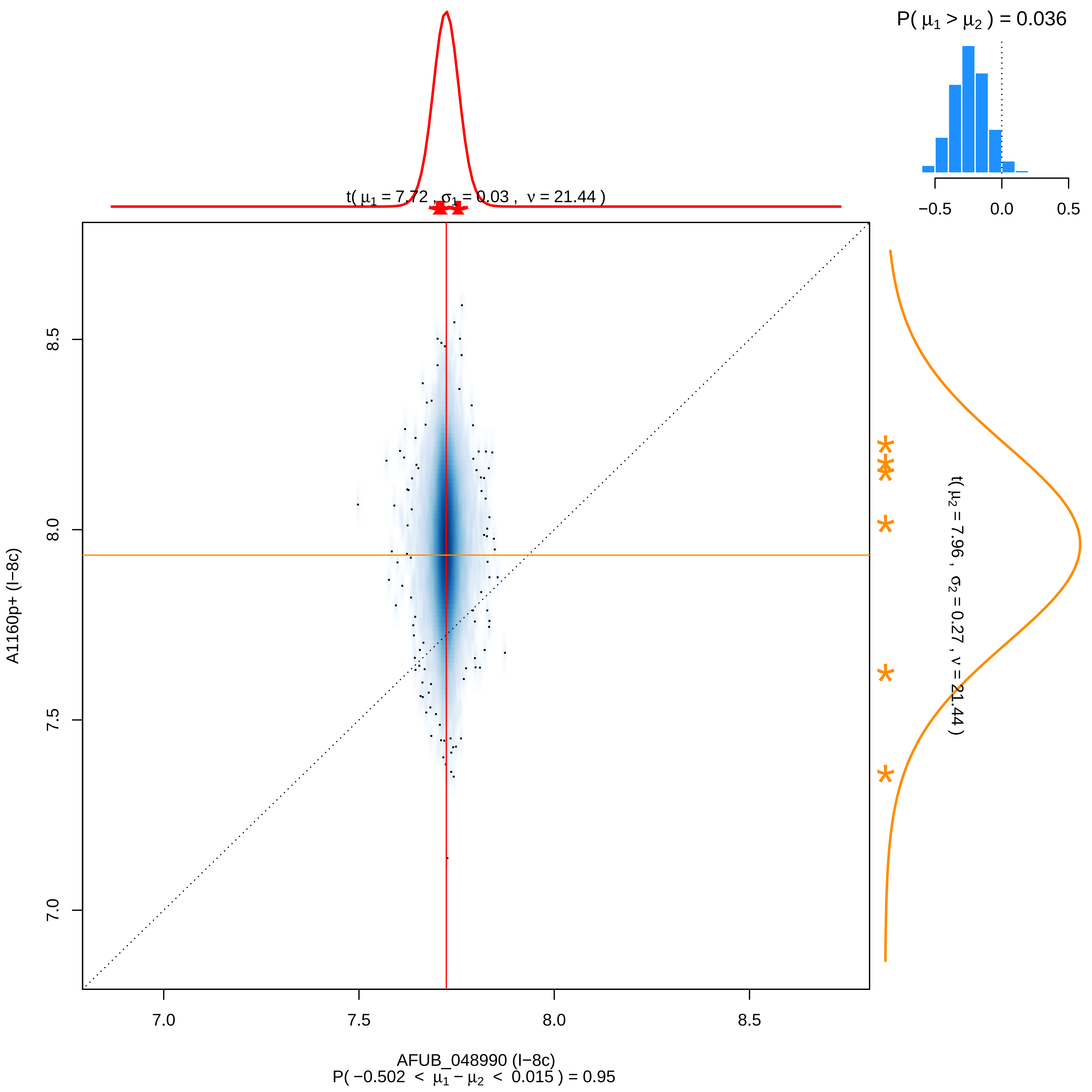

P( μ1 > μ2 ) = 0.036
−0.5
0.0
0.5
t( μ1 = 7.72 , σ1 = 0.03 , ν = 21.44 )
t( μ2 = 7.96 , σ2 = 0.27 , ν = 21.44 )
8.5
8.0
7.5
7.0
A1160p+ (I−8c)
AFUB_048990 (I−8c)
P( −0.502 < μ1 − μ2 < 0.015 ) = 0.95

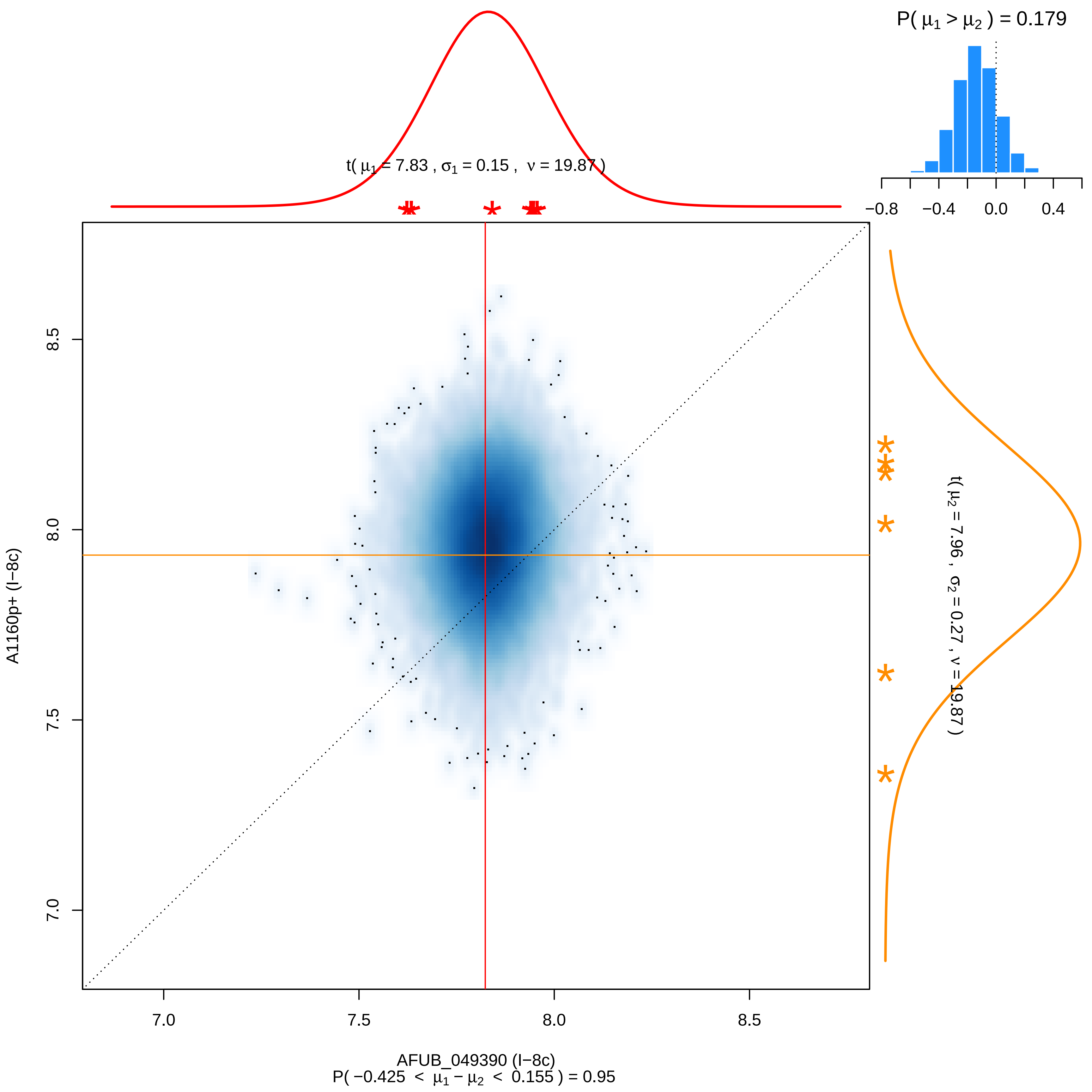

P( μ1 > μ2 ) = 0.179
−0.8
−0.4
0.0
0.4
t( μ1 = 7.83 , σ1 = 0.15 , ν = 19.87 )
A1160p+ (I−8c)
8.5
8.0
7.5
7.0
t( μ2 = 7.96 , σ2 = 0.27 , ν = 19.87 )
AFUB_049390 (I−8c)
P( −0.425 < μ1 − μ2 < 0.155 ) = 0.95

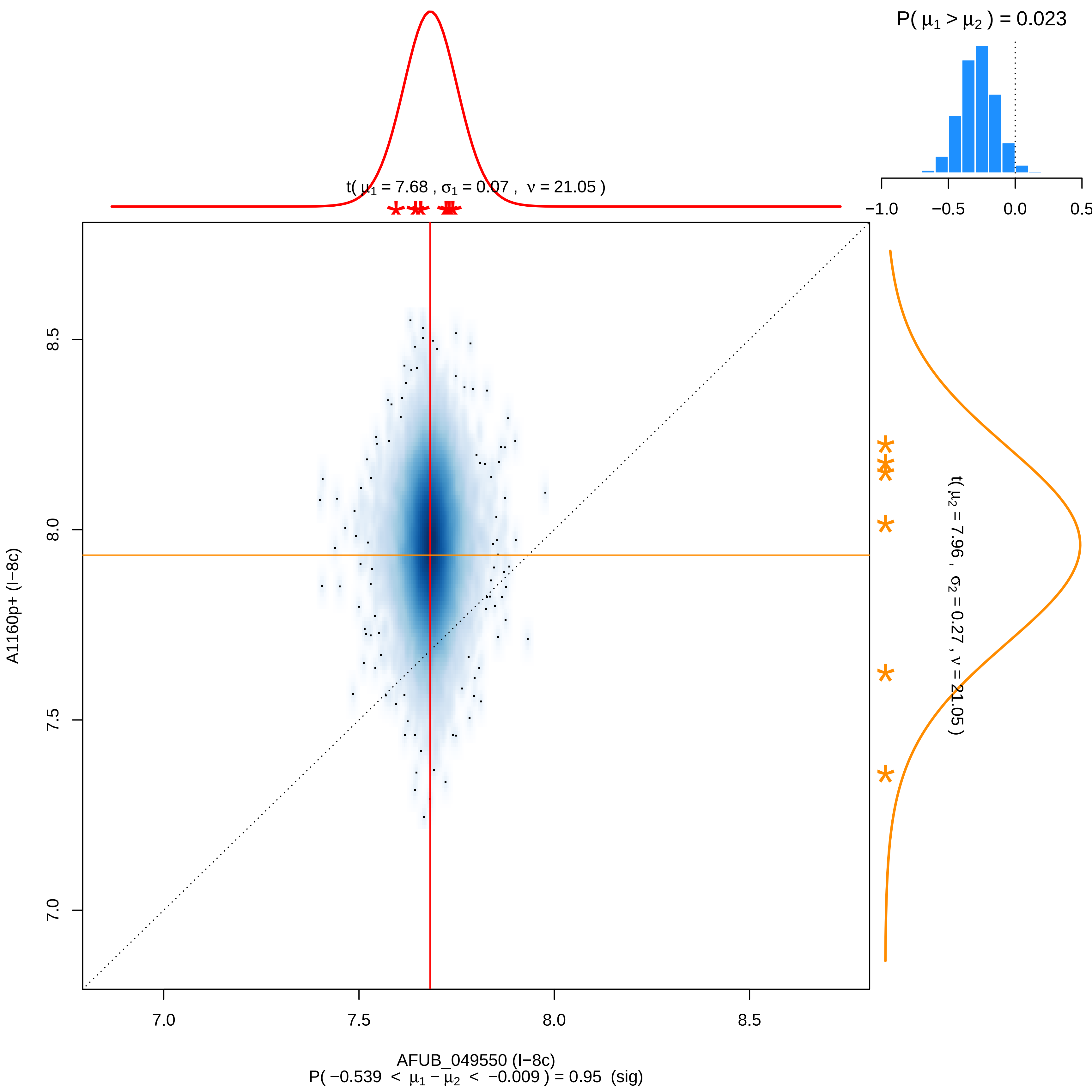

P( μ1 > μ2 ) = 0.023
−1.0
−0.5
0.0
0.5
t( μ1 = 7.68 , σ1 = 0.07 , ν = 21.05 )
t( μ2 = 7.96 , σ2 = 0.27 , ν = 21.05 )
8.5
8.0
7.5
7.0
A1160p+ (I−8c)
AFUB_049550 (I−8c)
P( −0.539 < μ1 − μ2 < −0.009 ) = 0.95 (sig)

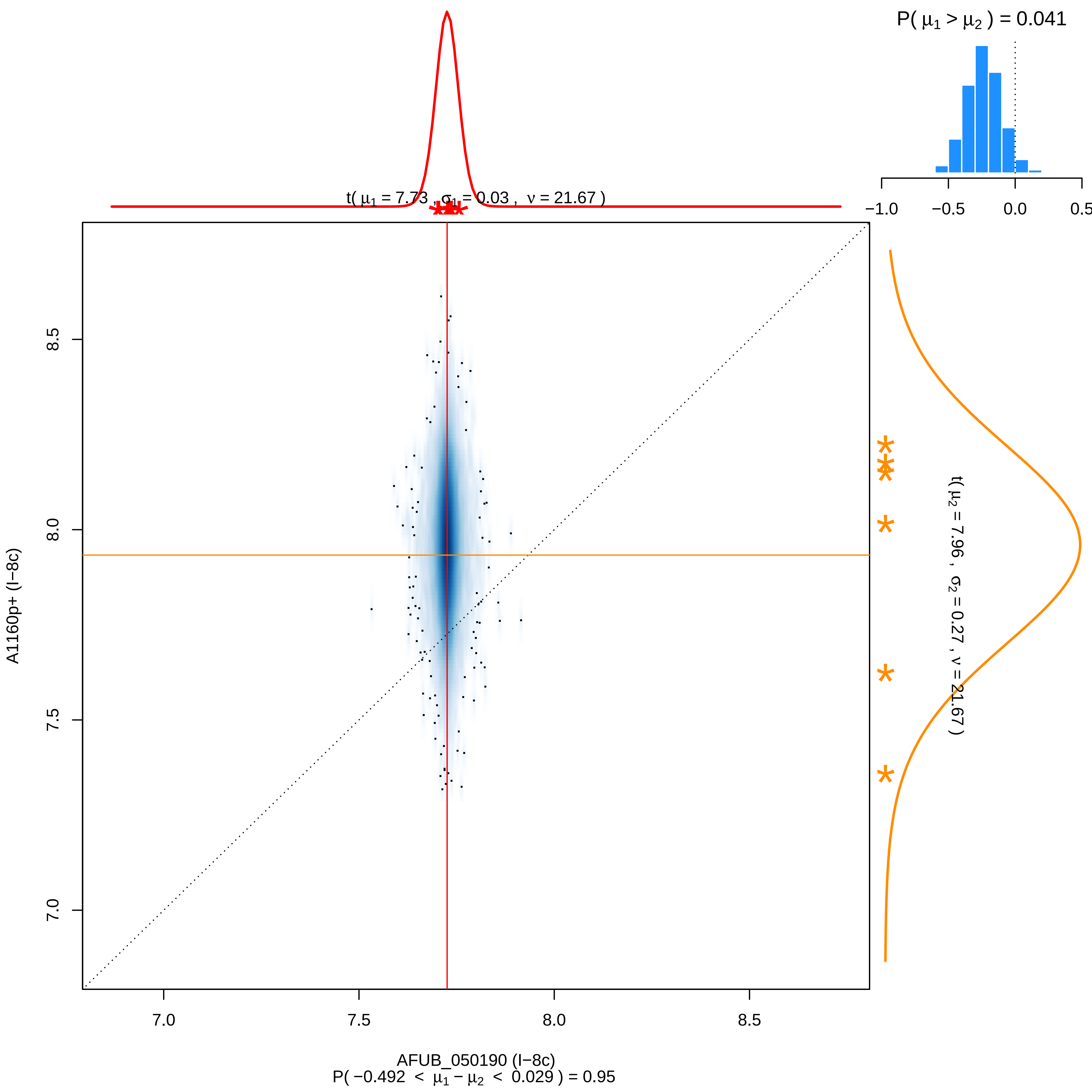
P( μ1 > μ2 ) = 0.041
−1.0
−0.5
0.0
0.5
t( μ1 = 7.73 , σ1 = 0.03 , ν = 21.67 )
t( μ2 = 7.96 , σ2 = 0.27 , ν = 21.67 )
8.5
8.0
7.5
7.0
A1160p+ (I−8c)
AFUB_050190 (I−8c)
P( −0.492 < μ1 − μ2 < 0.029 ) = 0.95

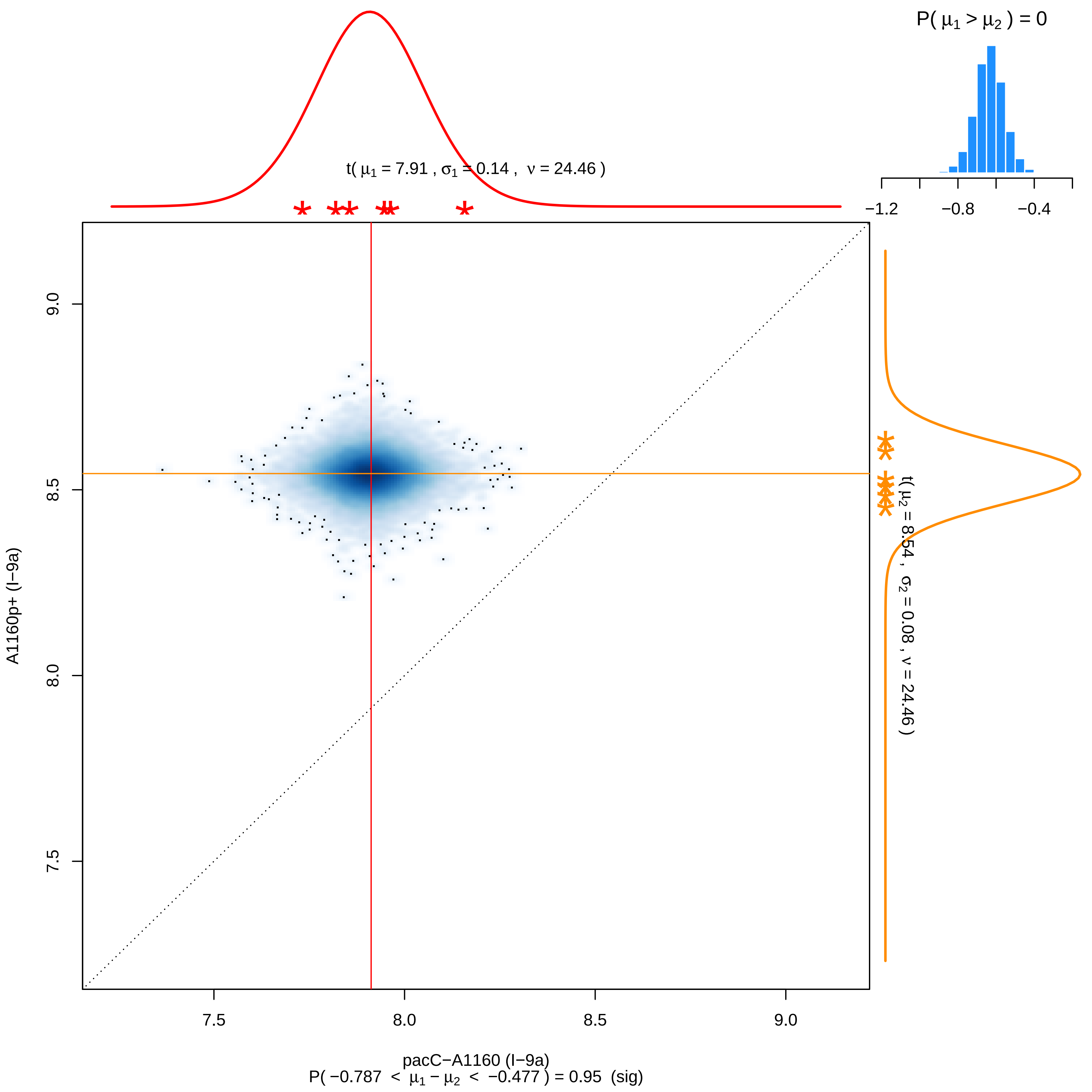

P( μ1 > μ2 ) = 0
−1.2
−0.8
−0.4
t( μ1 = 7.91 , σ1 = 0.14 , ν = 24.46 )
t( μ2 = 8.54 , σ2 = 0.08 , ν = 24.46 )
A1160p+ (I−9a)
9.0
8.5
8.0
7.5
7.5
8.0
8.5
9.0
pacC−A1160 (I−9a)
P( −0.787 < μ1 − μ2 < −0.477 ) = 0.95 (sig)

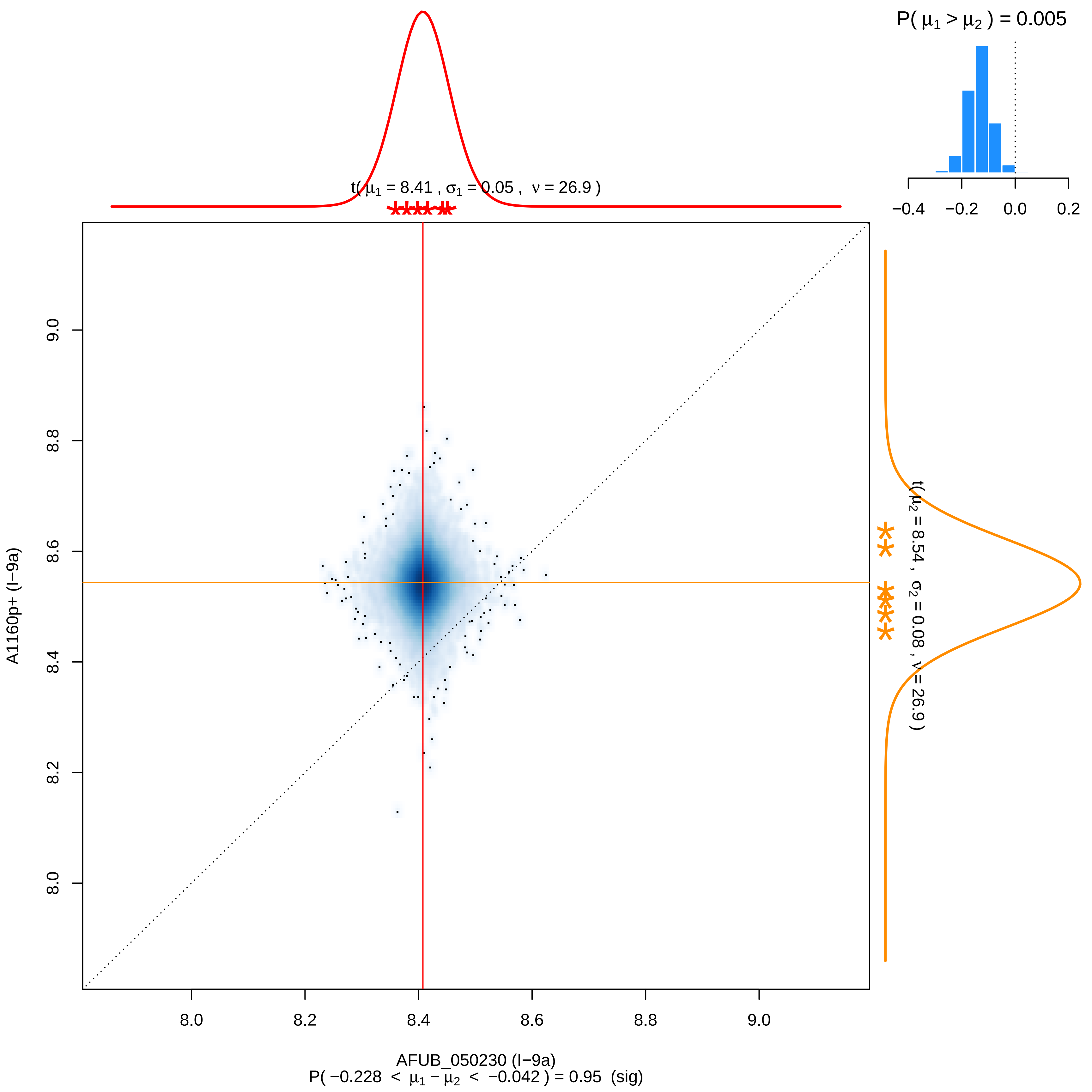

P( μ1 > μ2 ) = 0.005
−0.4
−0.2
0.0
0.2
t( μ1 = 8.41 , σ1 = 0.05 , ν = 26.9 )
t( μ2 = 8.54 , σ2 = 0.08 , ν = 26.9 )
A1160p+ (I−9a)
8.0
8.2
8.4
8.6
8.8
9.0
AFUB_050230 (I−9a)
P( −0.228 < μ1 − μ2 < −0.042 ) = 0.95 (sig)

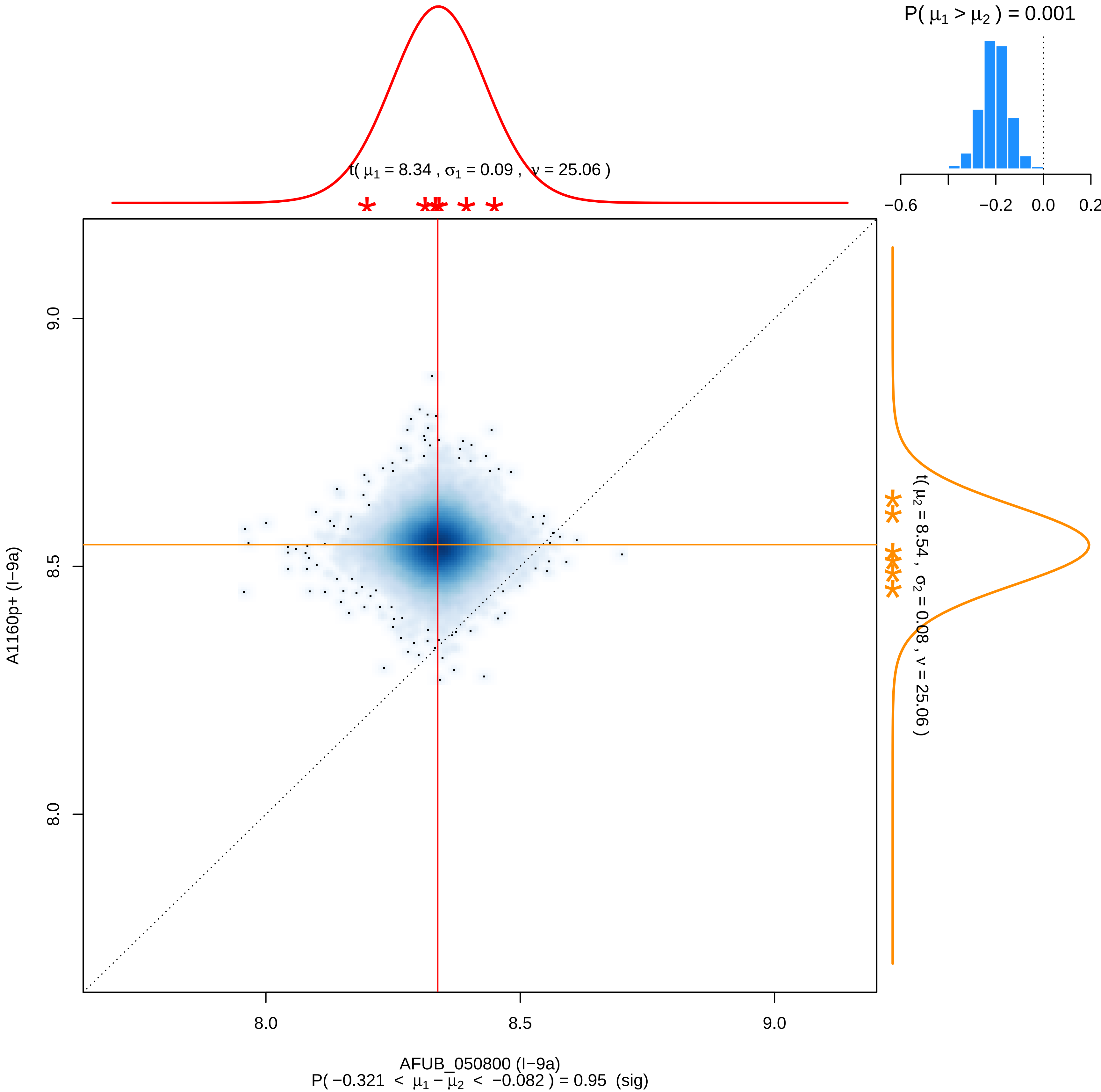

P( μ1 > μ2 ) = 0.001
−0.6
−0.2
0.0
0.2
t( μ1 = 8.34 , σ1 = 0.09 , ν = 25.06 )
t( μ2 = 8.54 , σ2 = 0.08 , ν = 25.06 )
9.0
8.5
8.0
A1160p+ (I−9a)
8.0
8.5
9.0
AFUB_050800 (I−9a)
P( −0.321 < μ1 − μ2 < −0.082 ) = 0.95 (sig)

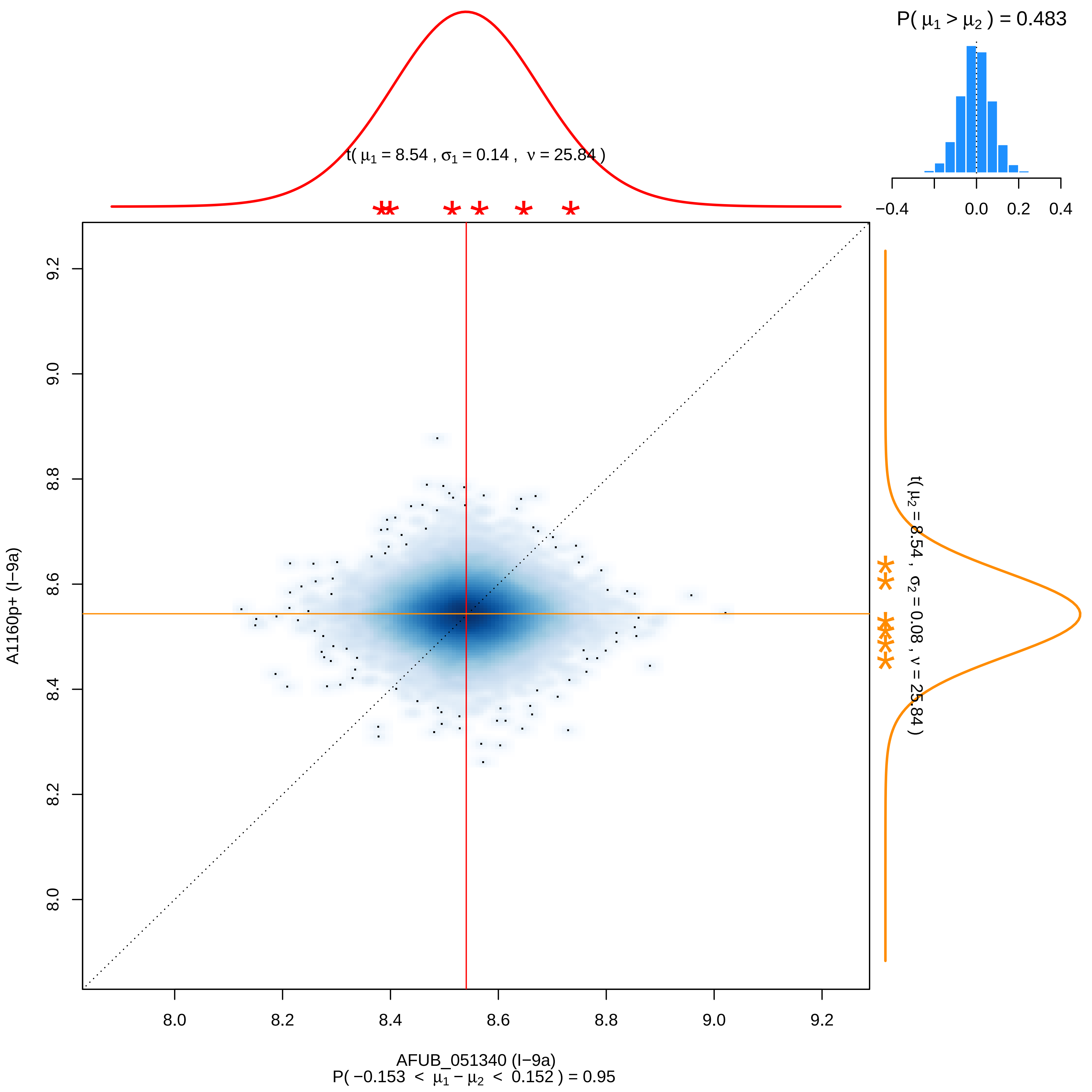

P( μ1 > μ2 ) = 0.483
−0.4
0.0
0.2
0.4
t( μ1 = 8.54 , σ1 = 0.14 , ν = 25.84 )
t( μ2 = 8.54 , σ2 = 0.08 , ν = 25.84 )
A1160p+ (I−9a)
9.2
9.0
8.8
8.6
8.4
8.2
8.0
8.0
8.2
8.4
8.6
8.8
9.0
9.2
AFUB_051340 (I−9a)
P( −0.153 < μ1 − μ2 < 0.152 ) = 0.95

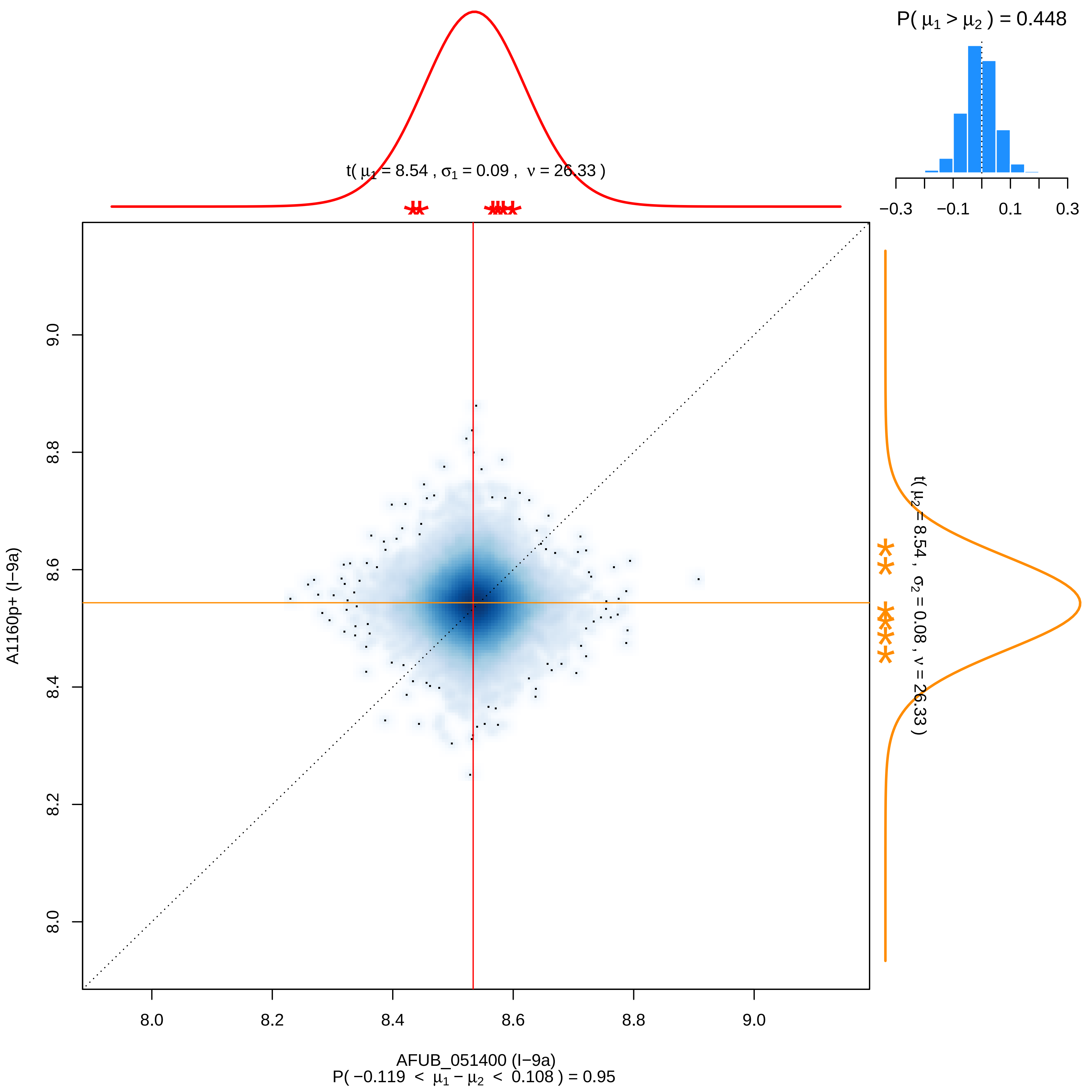

P( μ1 > μ2 ) = 0.448
−0.3 −0.1 0.1 0.3
t( μ1 = 8.54 , σ1 = 0.09 , ν = 26.33 )
t( μ2 = 8.54 , σ2 = 0.08 , ν = 26.33 )
A1160p+ (I−9a)
8.0 8.2 8.4 8.6 8.8 9.0
AFUB_051400 (I−9a)
P( −0.119 < μ1 − μ2 < 0.108 ) = 0.95

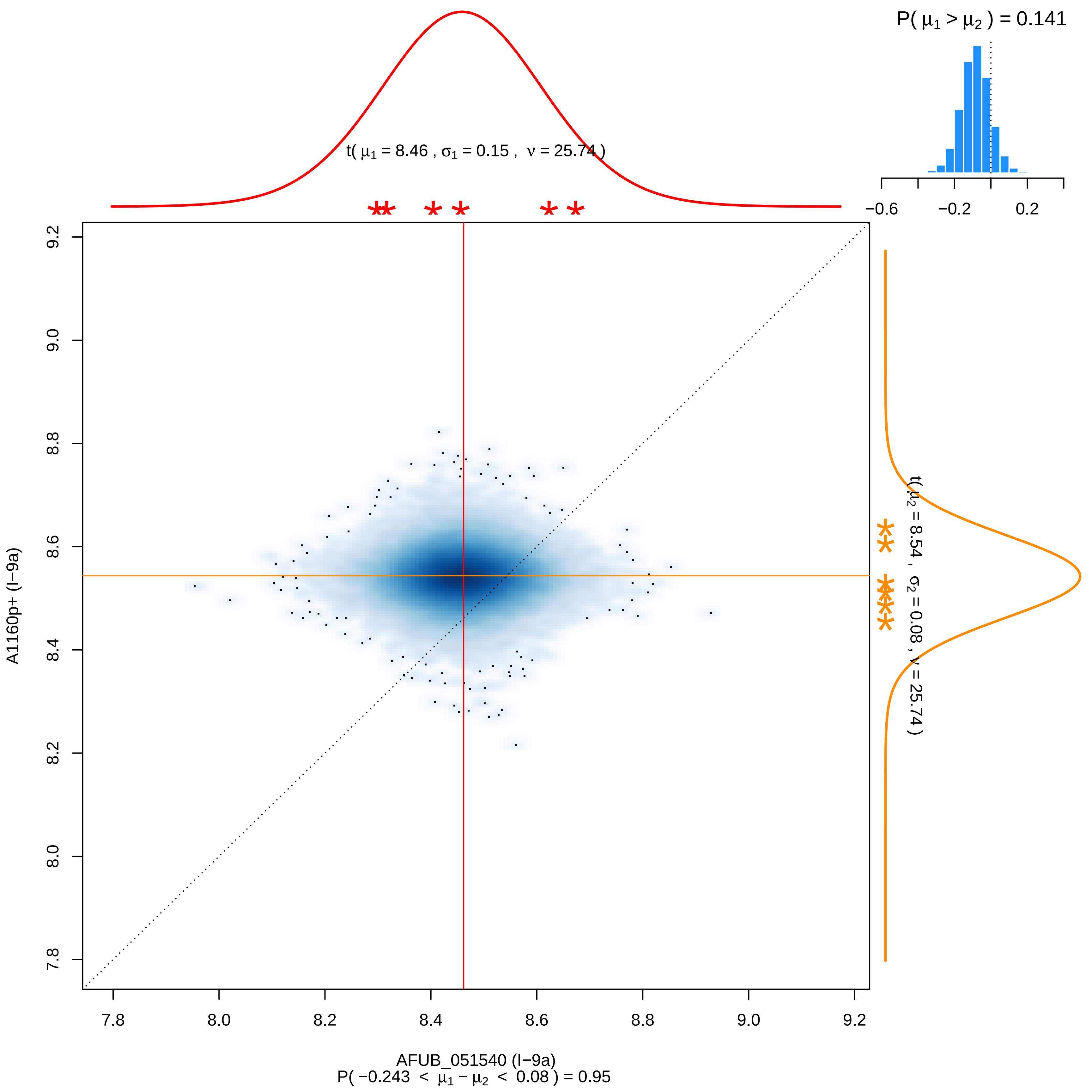

P( μ1 > μ2 ) = 0.141
−0.6
−0.2
0.2
t( μ1 = 8.46 , σ1 = 0.15 , ν = 25.74 )
t( μ2 = 8.54 , σ2 = 0.08 , ν = 25.74 )
A1160p+ (I−9a)
7.8
8.0
8.2
8.4
8.6
8.8
9.0
9.2
AFUB_051540 (I−9a)
P( −0.243 < μ1 − μ2 < 0.08 ) = 0.95

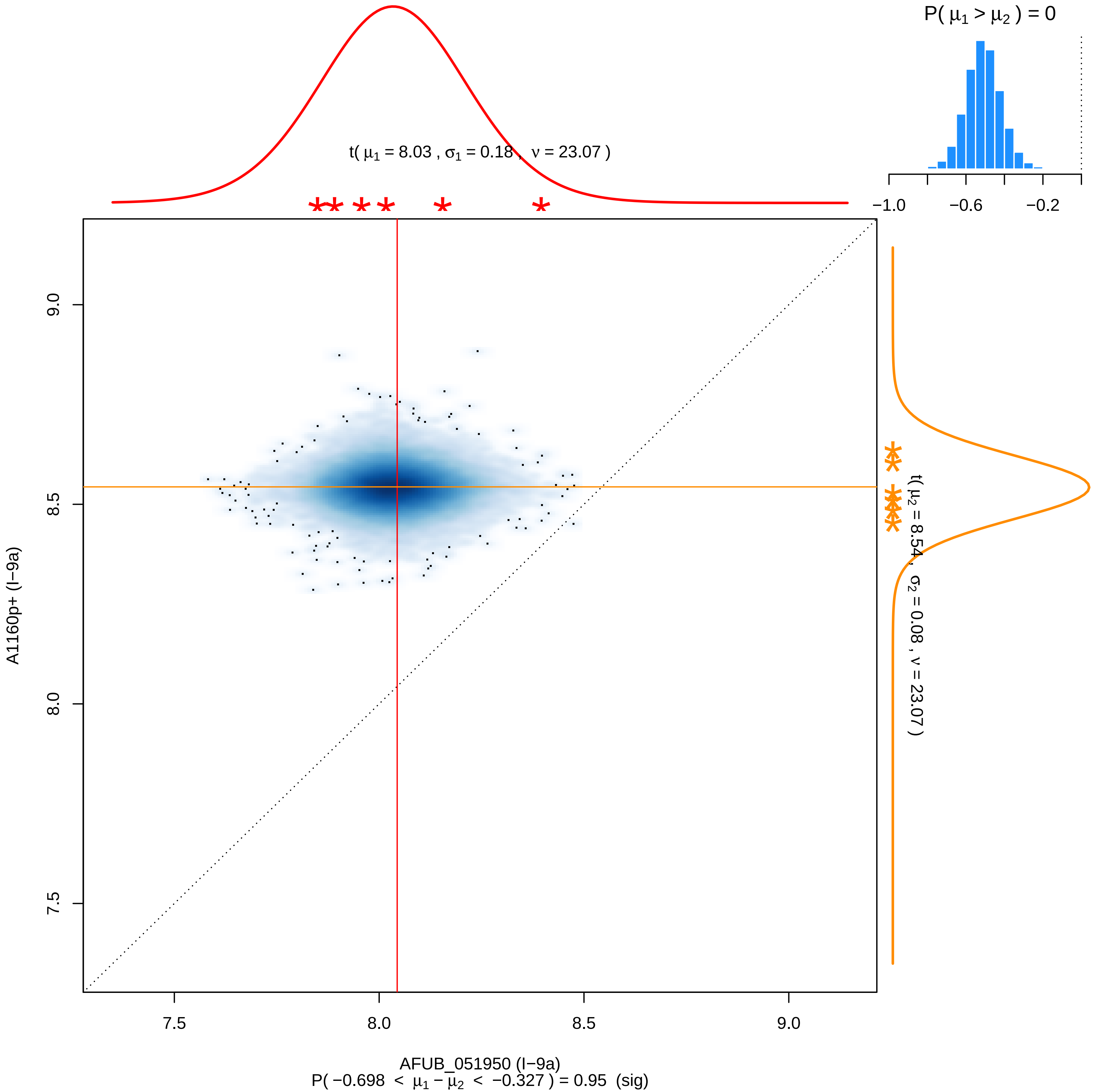
P( μ1 > μ2 ) = 0
−1.0
−0.6
−0.2
t( μ1 = 8.03 , σ1 = 0.18 , ν = 23.07 )
t( μ2 = 8.54 , σ2 = 0.08 , ν = 23.07 )
A1160p+ (I−9a)
9.0
8.5
8.0
7.5
7.5
8.0
8.5
9.0
AFUB_051950 (I−9a)
P( −0.698 < μ1 − μ2 < −0.327 ) = 0.95 (sig)

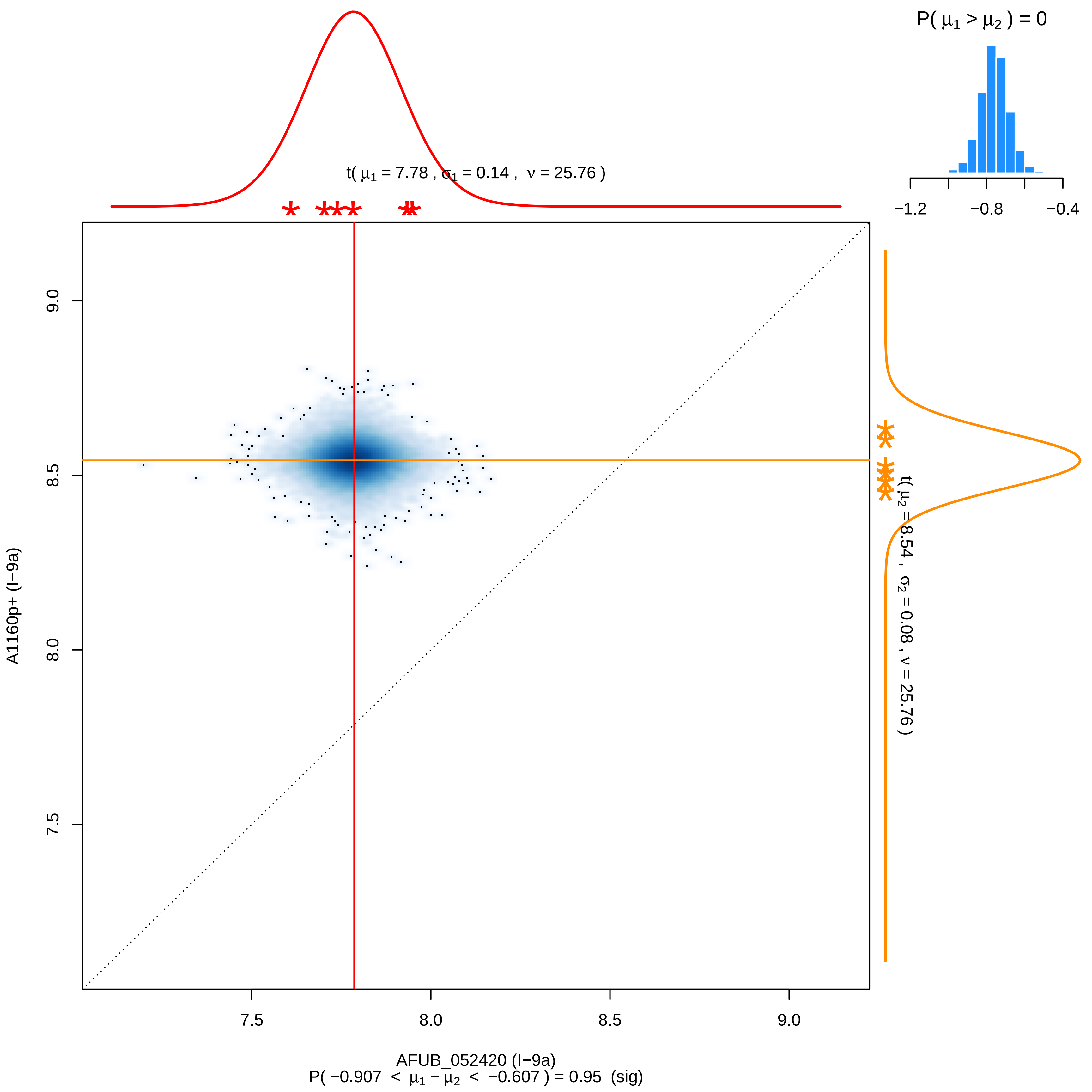

P( μ1 > μ2 ) = 0
−1.2
−0.8
−0.4
t( μ1 = 7.78 , σ1 = 0.14 , ν = 25.76 )
t( μ2 = 8.54 , σ2 = 0.08 , ν = 25.76 )
9.0
8.5
8.0
7.5
A1160p+ (I−9a)
7.5
8.0
8.5
9.0
AFUB_052420 (I−9a)
P( −0.907 < μ1 − μ2 < −0.607 ) = 0.95 (sig)

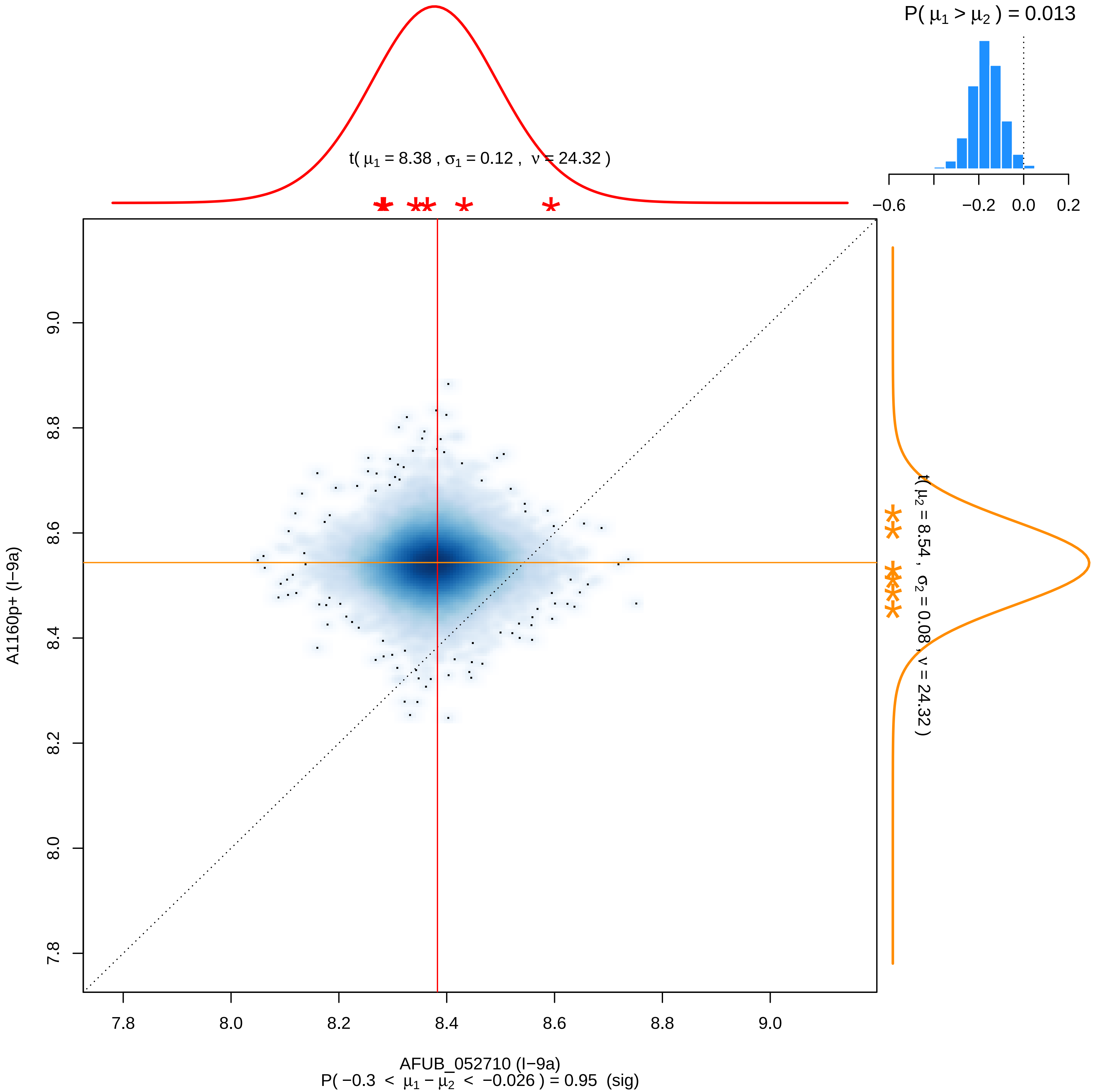

P( μ1 > μ2 ) = 0.013
−0.6
−0.2
0.0
0.2
t( μ1 = 8.38 , σ1 = 0.12 , ν = 24.32 )
t( μ2 = 8.54 , σ2 = 0.08 , ν = 24.32 )
9.0
8.8
8.6
8.4
8.2
8.0
7.8
A1160p+ (I−9a)
7.8
8.0
8.2
8.4
8.6
8.8
9.0
AFUB_052710 (I−9a)
P( −0.3 < μ1 − μ2 < −0.026 ) = 0.95 (sig)

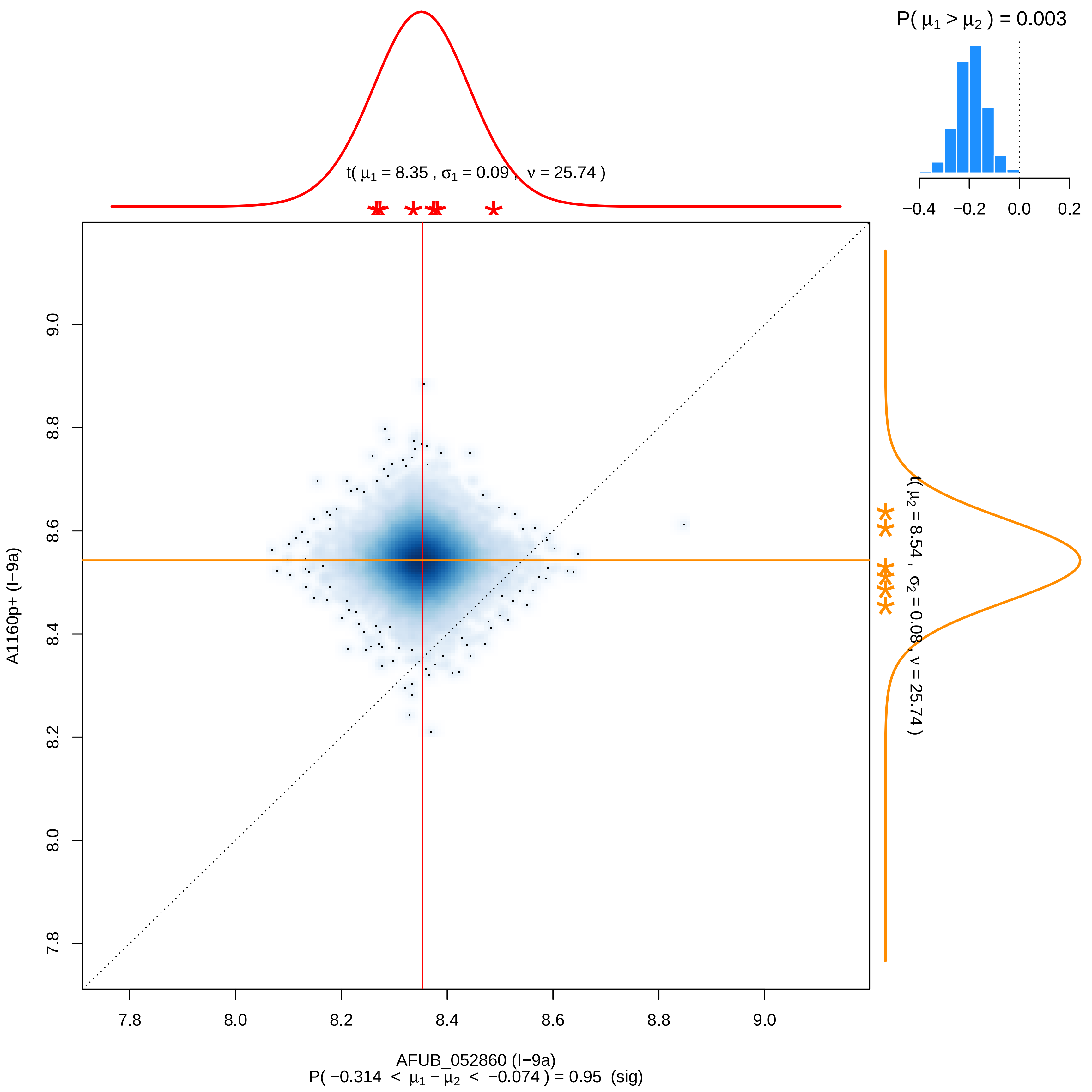

P( μ1 > μ2 ) = 0.003
−0.4
−0.2
0.0
0.2
t( μ1 = 8.35 , σ1 = 0.09 , ν = 25.74 )
t( μ2 = 8.54 , σ2 = 0.08 , ν = 25.74 )
9.0
8.8
8.6
8.4
8.2
8.0
7.8
A1160p+ (I−9a)
AFUB_052860 (I−9a)
P( −0.314 < μ1 − μ2 < −0.074 ) = 0.95 (sig)

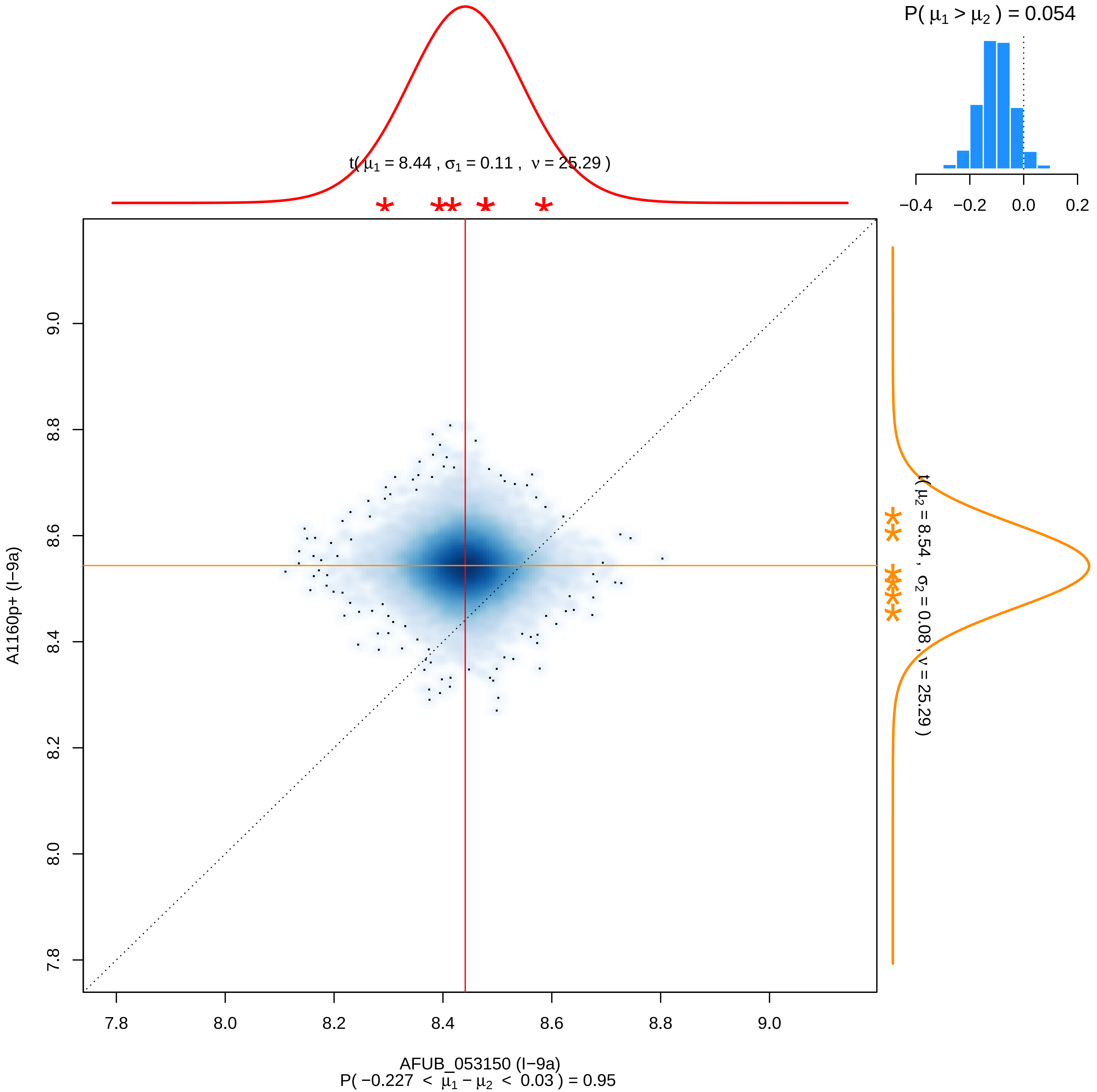

P( μ1 > μ2 ) = 0.054
−0.4
−0.2
0.0
0.2
t( μ1 = 8.44 , σ1 = 0.11 , ν = 25.29 )
t( μ2 = 8.54 , σ2 = 0.08 , ν = 25.29 )
A1160p+ (I−9a)
7.8
8.0
8.2
8.4
8.6
8.8
9.0
AFUB_053150 (I−9a)
P( −0.227 < μ1 − μ2 < 0.03 ) = 0.95
