## Supplementary material for "Distinct cohorts of *Aspergillus fumigatus* transcription factors are required for epithelial damage occurring via contact- or soluble effector-mediated mechanisms": TABLE S2

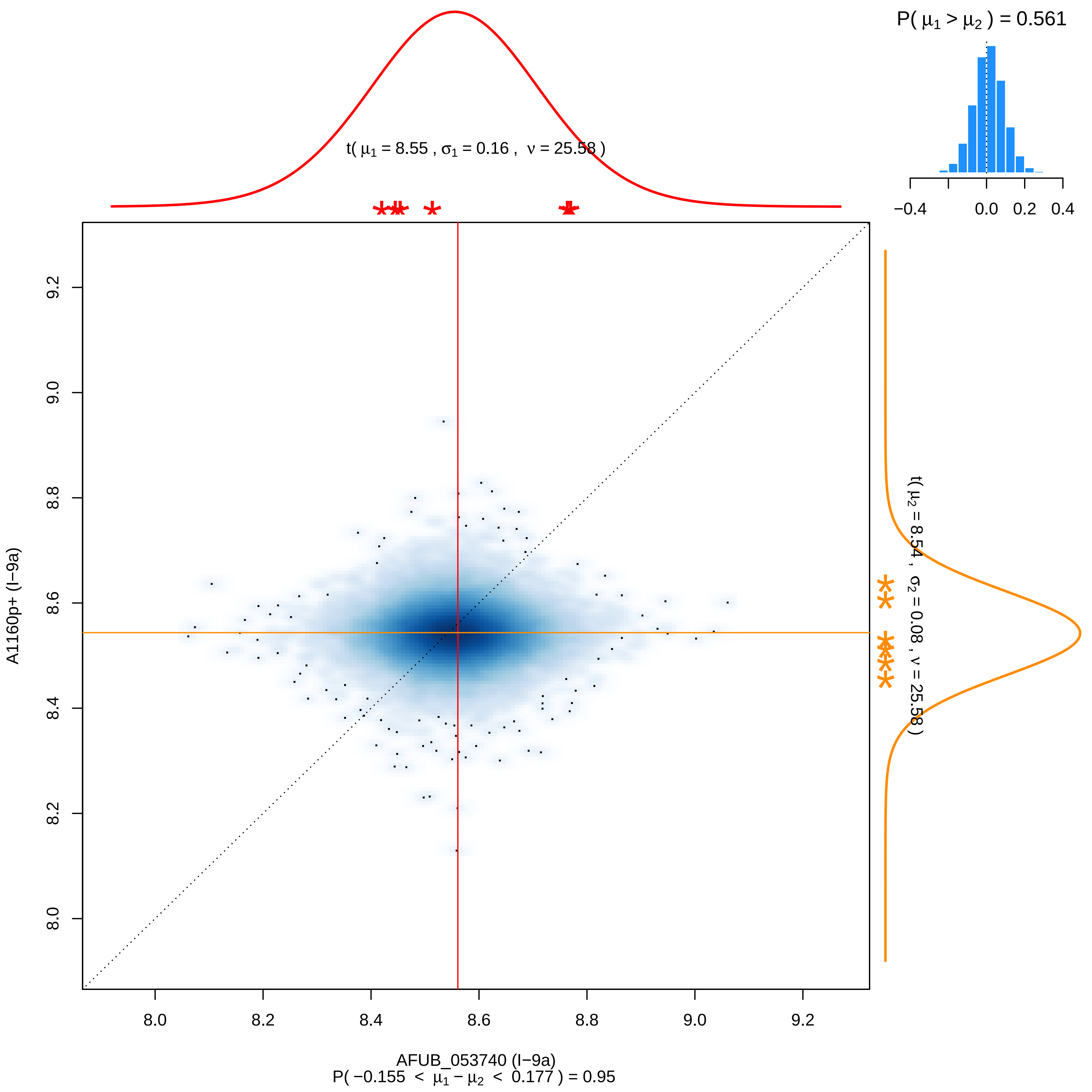
P( μ1 > μ2 ) = 0.561
−0.4
0.0
0.2
0.4
t( μ1 = 8.55 , σ1 = 0.16 , ν = 25.58 )
t( μ2 = 8.54 , σ2 = 0.08 , ν = 25.58 )
A1160p+ (I−9a)
8.0
8.2
8.4
8.6
8.8
9.0
9.2
AFUB_053740 (I−9a)
P( −0.155 < μ1 − μ2 < 0.177 ) = 0.95

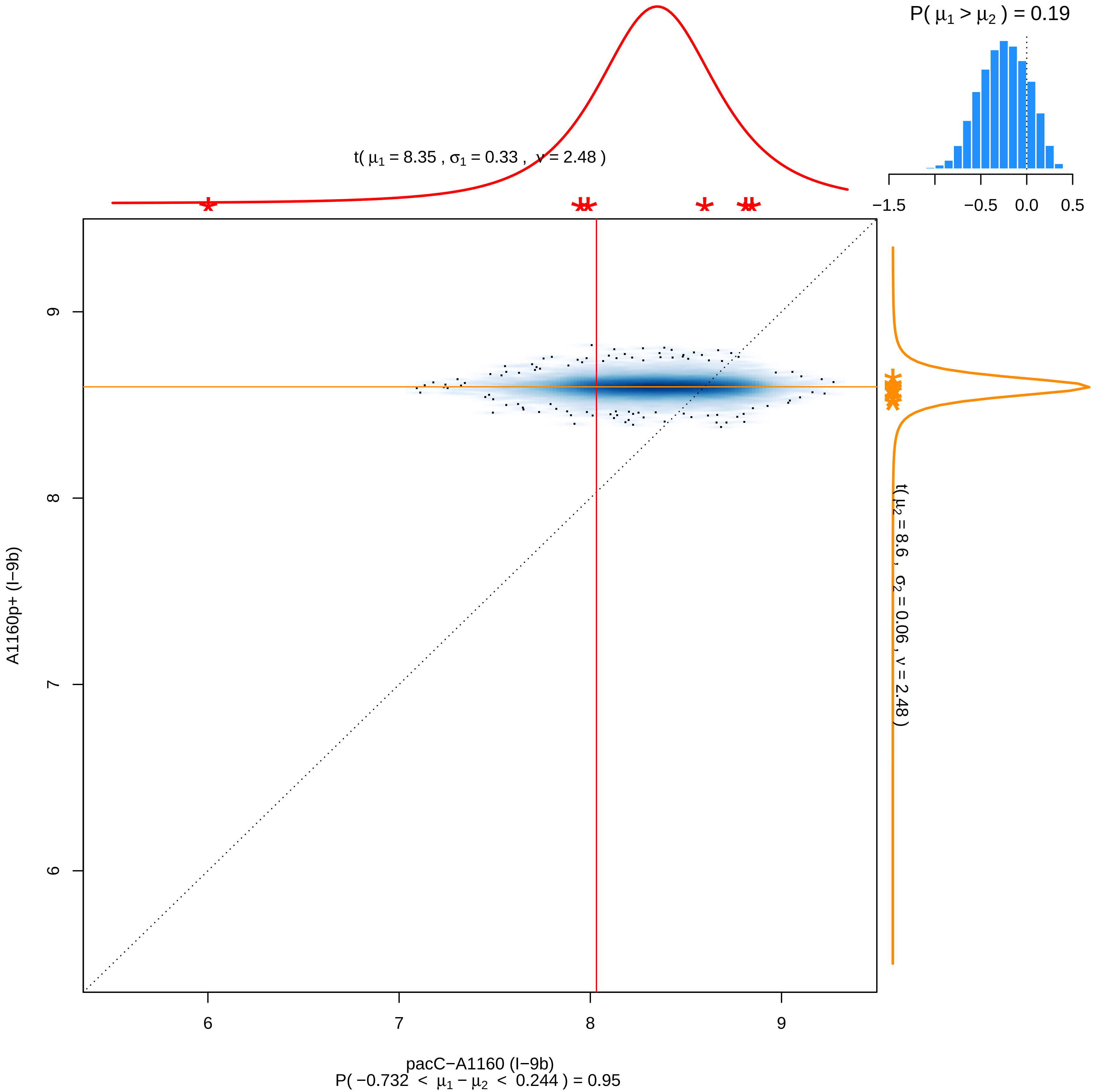

P( μ1 > μ2 ) = 0.19
−1.5
−0.5
0.0
0.5
t( μ1 = 8.35 , σ1 = 0.33 , ν = 2.48 )
t( μ2 = 8.6 , σ2 = 0.06 , ν = 2.48 )
A1160p+ (I−9b)
9
8
7
6
6
7
8
9
pacC−A1160 (I−9b)
P( −0.732 < μ1 − μ2 < 0.244 ) = 0.95

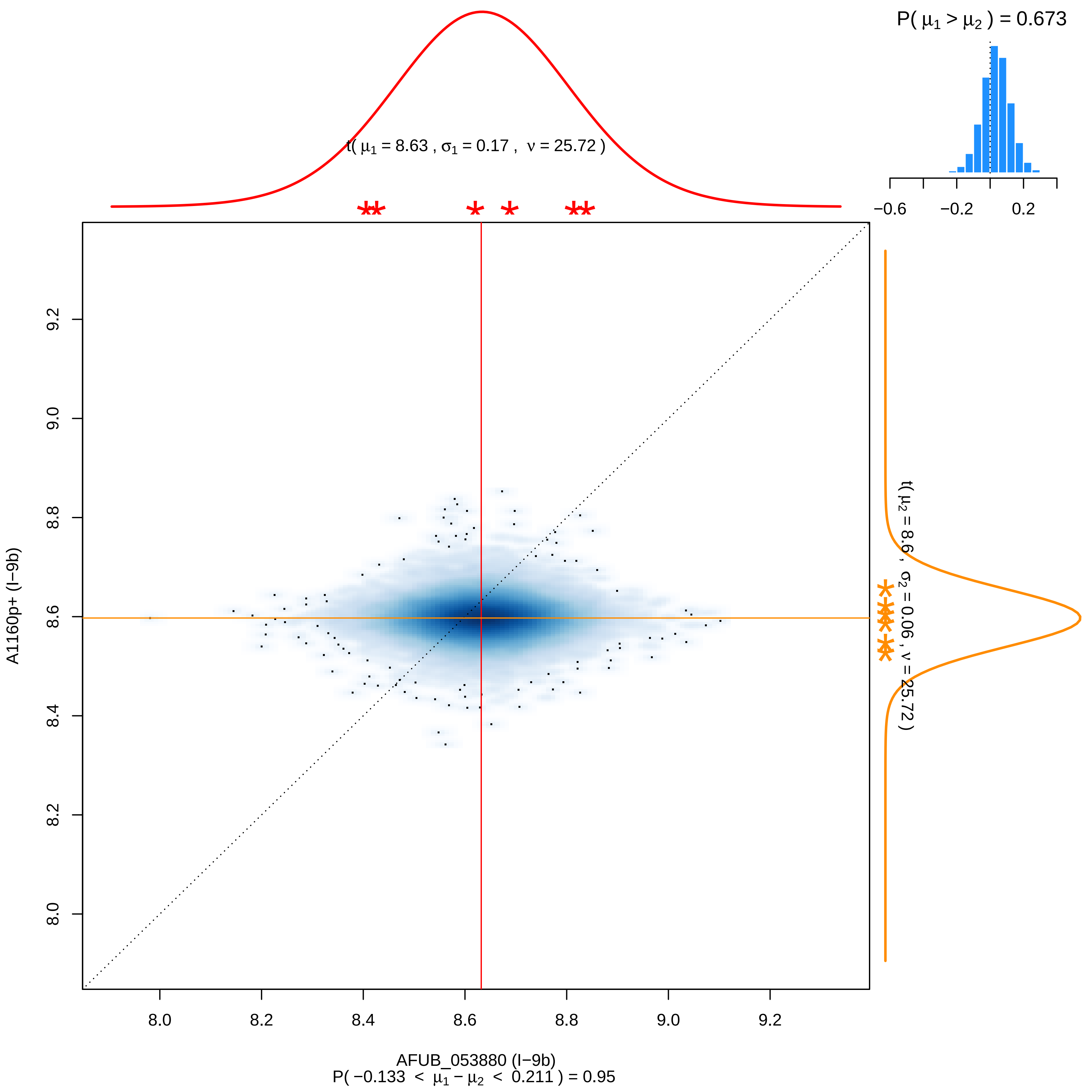

P( μ1 > μ2 ) = 0.673
t( μ1 = 8.63 , σ1 = 0.17 , ν = 25.72 )
−0.6
−0.2
0.2
t( μ2 = 8.6 , σ2 = 0.06 , ν = 25.72 )
A1160p+ (I−9b)
8.0
8.2
8.4
8.6
8.8
9.0
9.2
AFUB_053880 (I−9b)
P( −0.133 < μ1 − μ2 < 0.211 ) = 0.95

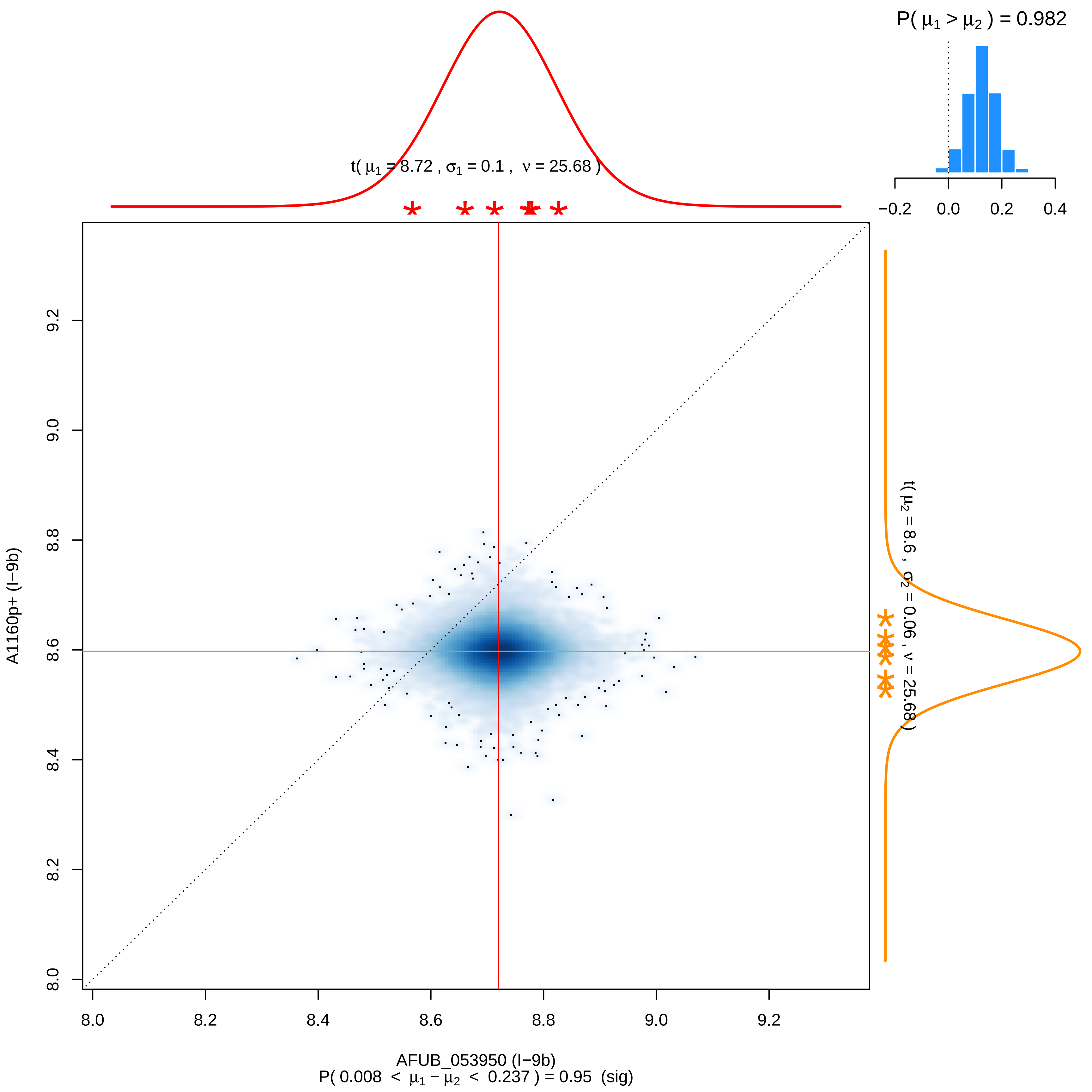

P( μ1 > μ2 ) = 0.982
−0.2 0.0 0.2 0.4
t( μ1 = 8.72 , σ1 = 0.1 , ν = 25.68 )
t( μ2 = 8.6 , σ2 = 0.06 , ν = 25.68 )
8.0 8.2 8.4 8.6 8.8 9.0 9.2
A1160p+ (I−9b)
AFUB_053950 (I−9b)
P( 0.008 < μ1 − μ2 < 0.237 ) = 0.95 (sig)

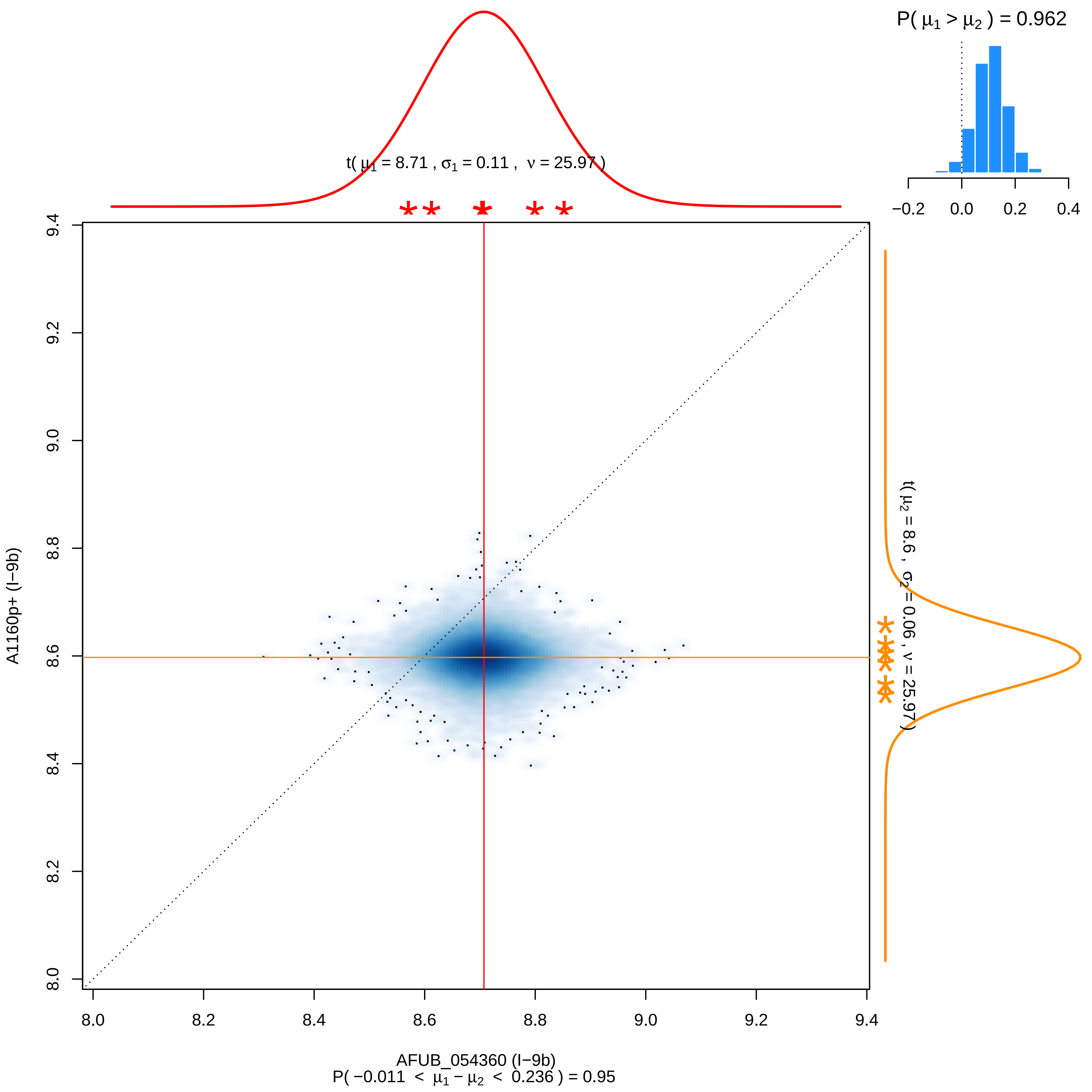

P( μ1 > μ2 ) = 0.962
−0.2 0.0 0.2 0.4
t( μ1 = 8.71 , σ1 = 0.11 , ν = 25.97 )
t( μ2 = 8.6 , σ2 = 0.06 , ν = 25.97 )
A1160p+ (I−9b)
8.0 8.2 8.4 8.6 8.8 9.0 9.2 9.4
AFUB_054360 (I−9b)
P( −0.011 < μ1 − μ2 < 0.236 ) = 0.95

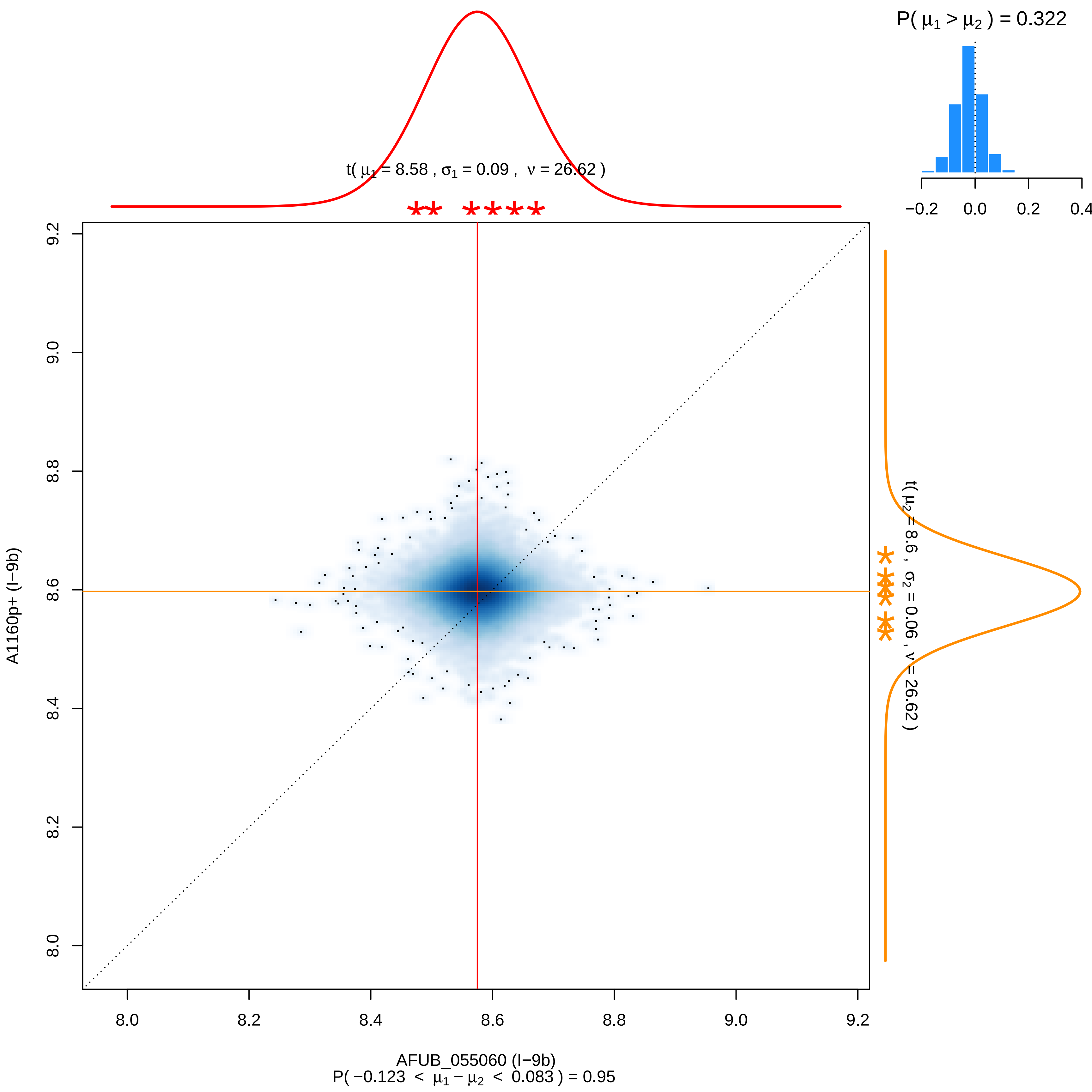

P( $\mu_1 > \mu_2$ ) = 0.322
−0.2
0.0
0.2
0.4
t( $\mu_1$ = 8.58 , $\sigma_1$ = 0.09 , $\nu$ = 26.62 )
t( $\mu_2$ = 8.6 , $\sigma_2$ = 0.06 , $\nu$ = 26.62 )
A1160p+ (I−9b)
9.2
9.0
8.8
8.6
8.4
8.2
8.0
8.0
8.2
8.4
8.6
8.8
9.0
9.2
AFUB_055060 (I−9b)
P( −0.123 < $\mu_1 - \mu_2$ < 0.083 ) = 0.95

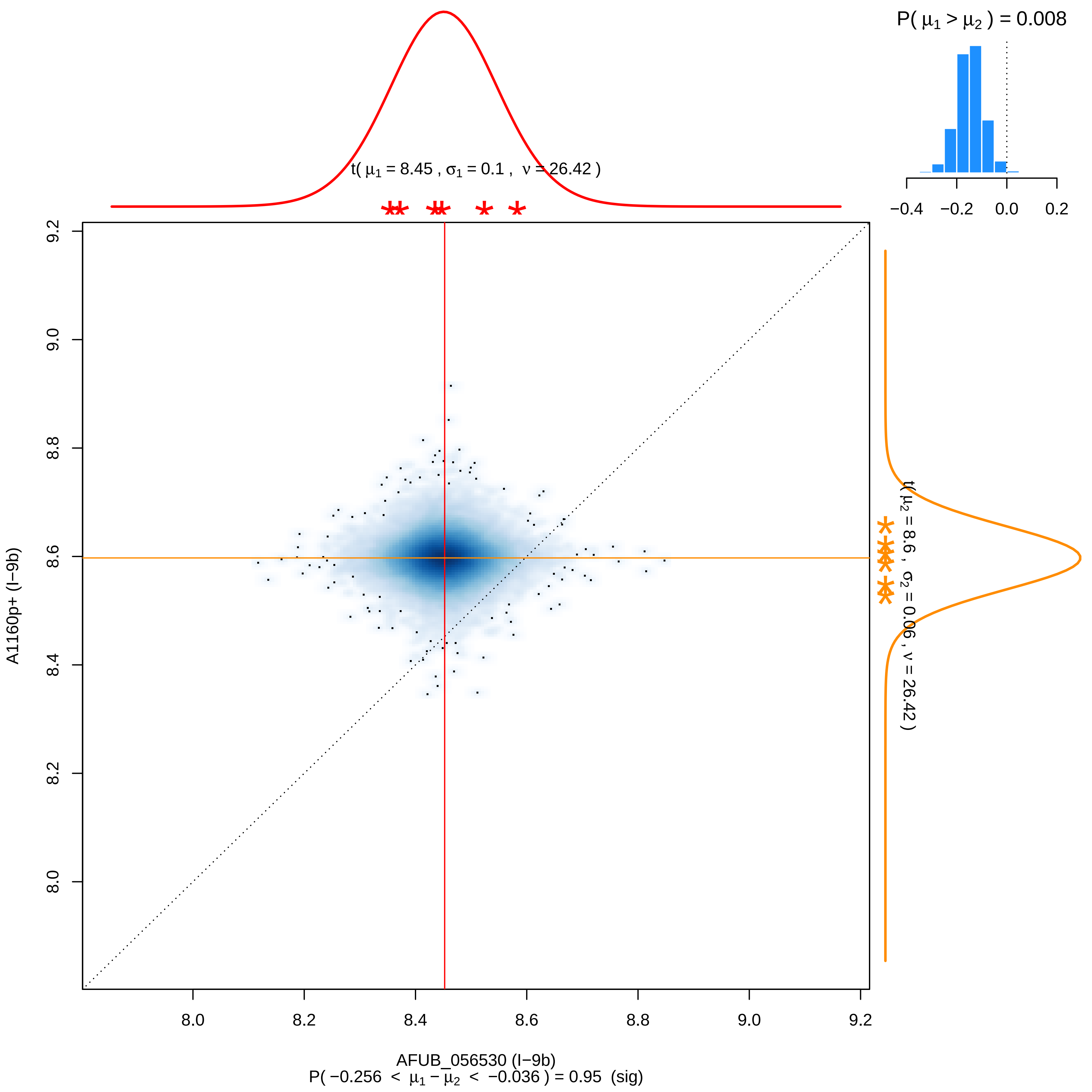

P( μ1 > μ2 ) = 0.008
−0.4 −0.2 0.0 0.2
t( μ1 = 8.45 , σ1 = 0.1 , ν = 26.42 )
t( μ2 = 8.6 , σ2 = 0.06 , ν = 26.42 )
A1160p+ (I−9b)
8.0 8.2 8.4 8.6 8.8 9.0 9.2
AFUB_056530 (I−9b)
P( −0.256 < μ1 − μ2 < −0.036 ) = 0.95 (sig)

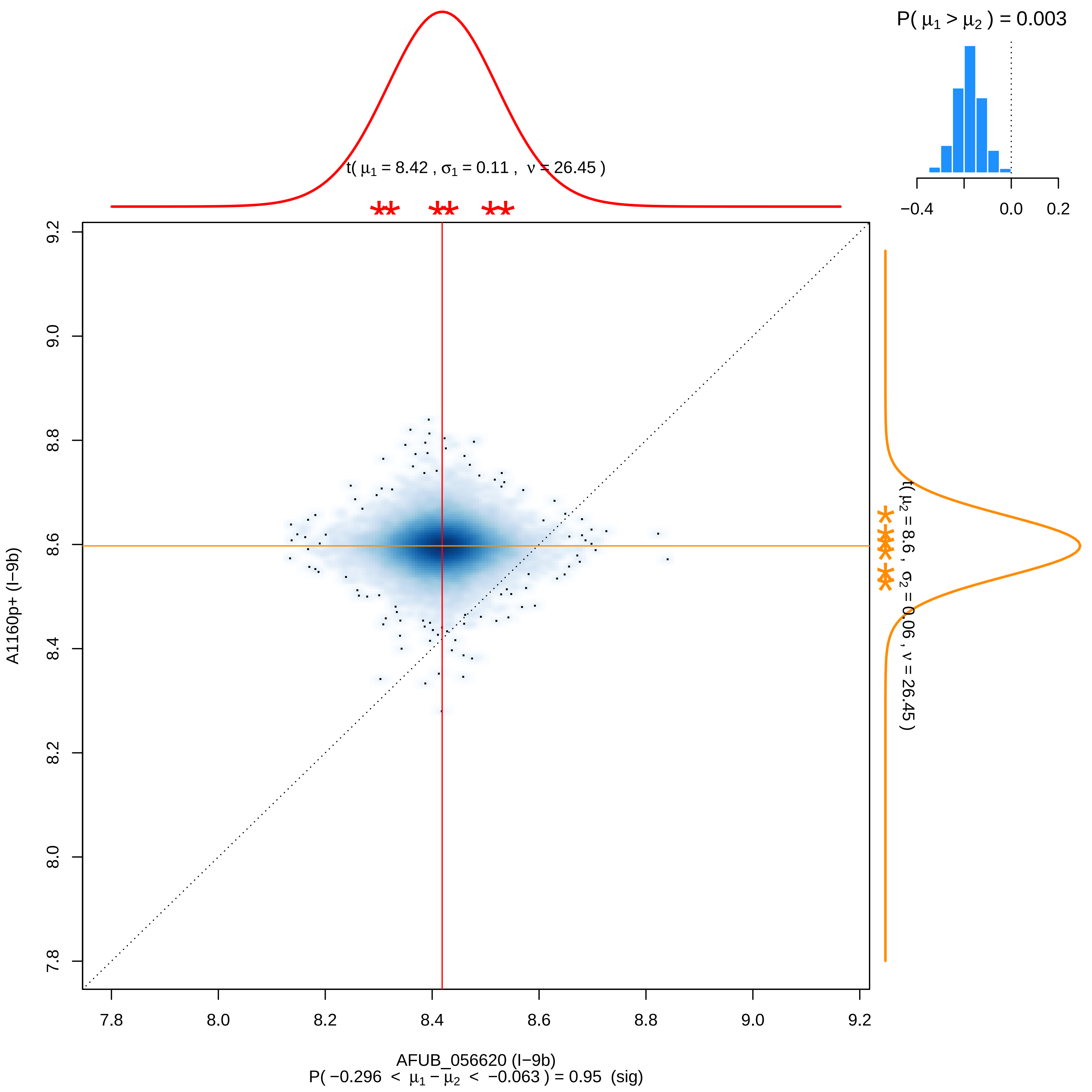

P( μ1 > μ2 ) = 0.003
−0.4
0.0
0.2
t( μ1 = 8.42 , σ1 = 0.11 , ν = 26.45 )
t( μ2 = 8.6 , σ2 = 0.06 , ν = 26.45 )
A1160p+ (I−9b)
9.2
9.0
8.8
8.6
8.4
8.2
8.0
7.8
7.8
8.0
8.2
8.4
8.6
8.8
9.0
9.2
AFUB_056620 (I−9b)
P( −0.296 < μ1 − μ2 < −0.063 ) = 0.95 (sig)

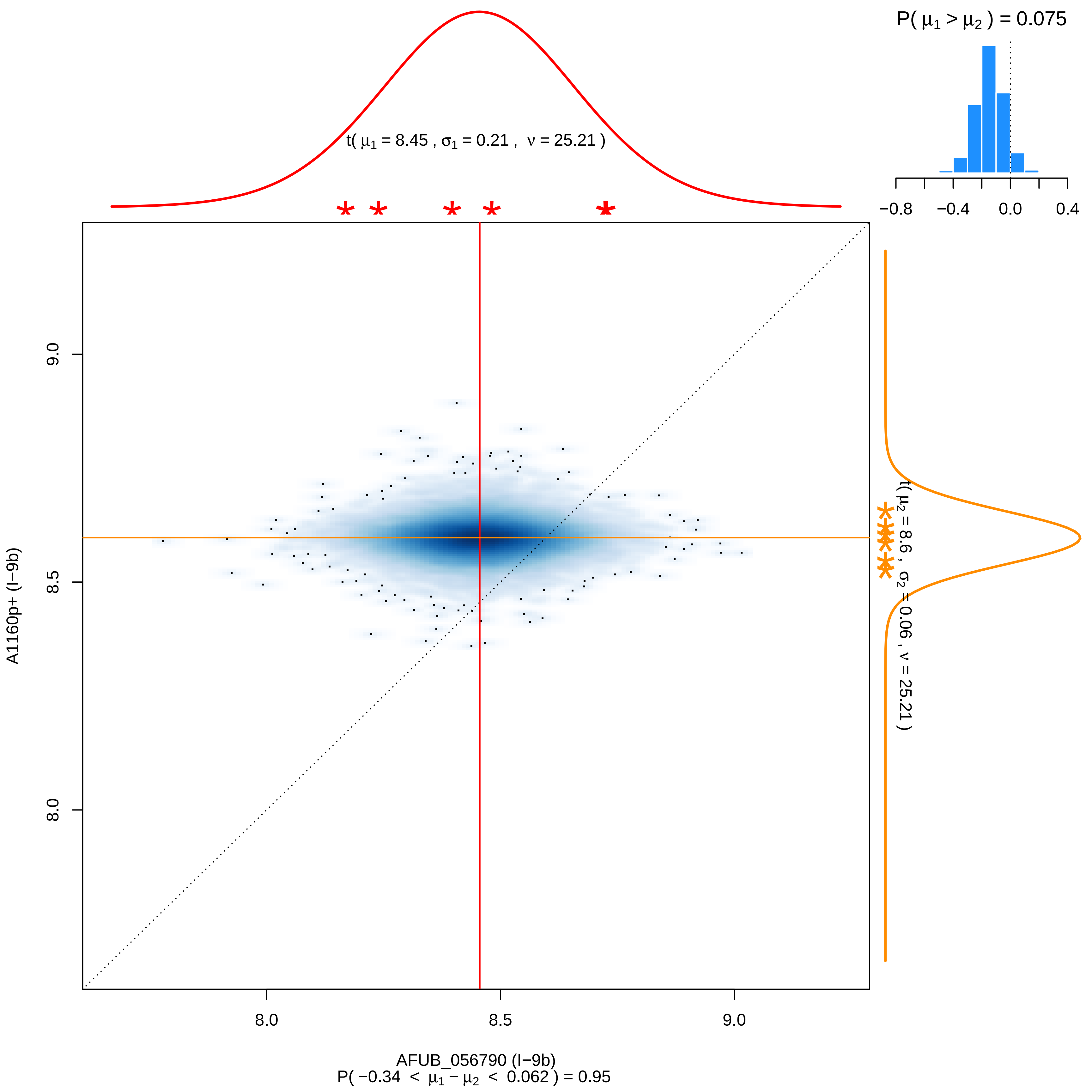

P( μ1 > μ2 ) = 0.075
−0.8
−0.4
0.0
0.4
t( μ1 = 8.45 , σ1 = 0.21 , ν = 25.21 )
t( μ2 = 8.6 , σ2 = 0.06 , ν = 25.21 )
A1160p+ (I−9b)
9.0
8.5
8.0
8.0
8.5
9.0
AFUB_056790 (I−9b)
P( −0.34 < μ1 − μ2 < 0.062 ) = 0.95

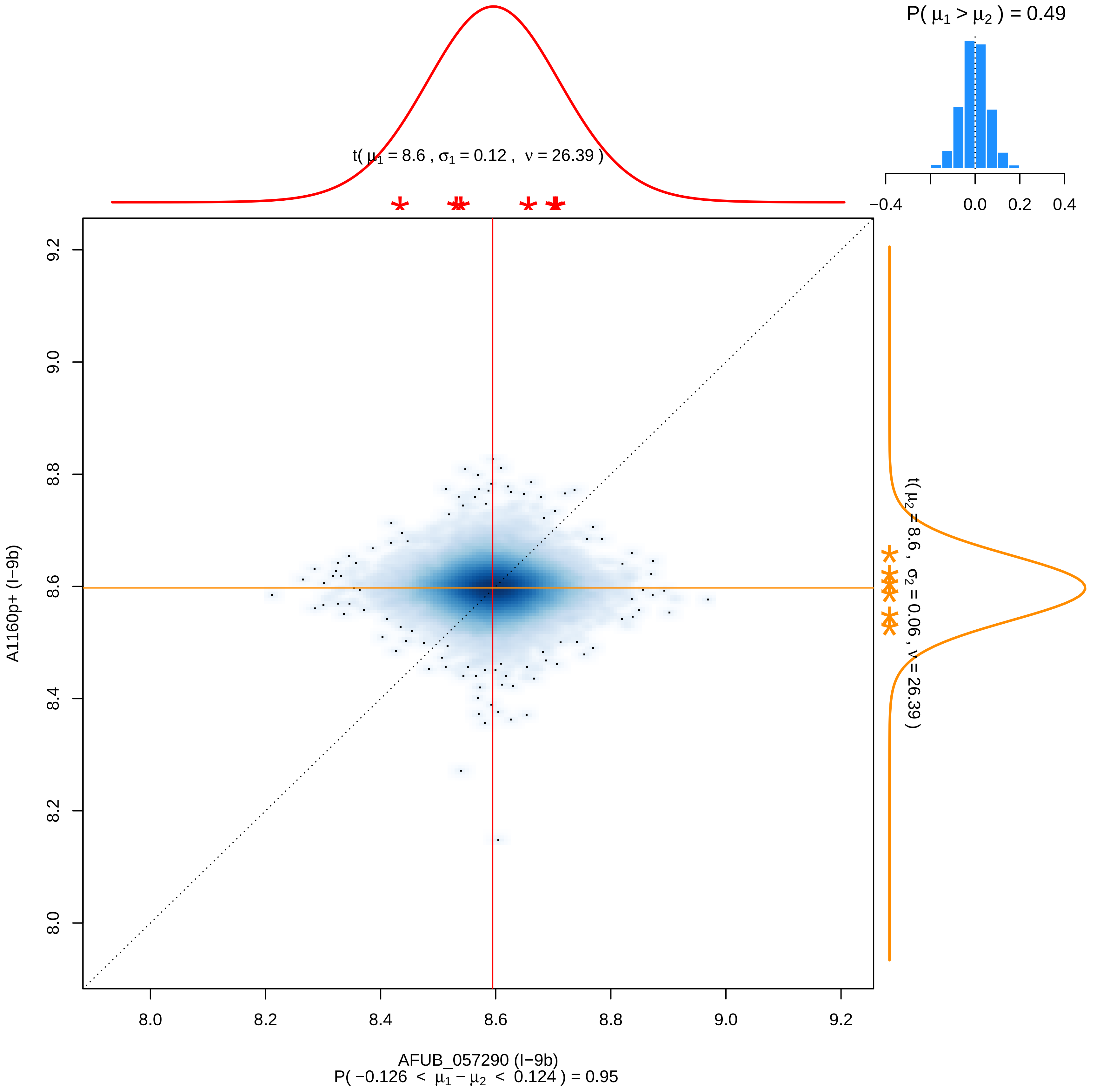

P( μ1 > μ2 ) = 0.49
−0.4
0.0
0.2
0.4
t( μ1 = 8.6 , σ1 = 0.12 , ν = 26.39 )
t( μ2 = 8.6 , σ2 = 0.06 , ν = 26.39 )
9.2
9.0
8.8
8.6
8.4
8.2
8.0
A1160p+ (I−9b)
8.0
8.2
8.4
8.6
8.8
9.0
9.2
AFUB_057290 (I−9b)
P( −0.126 < μ1 − μ2 < 0.124 ) = 0.95

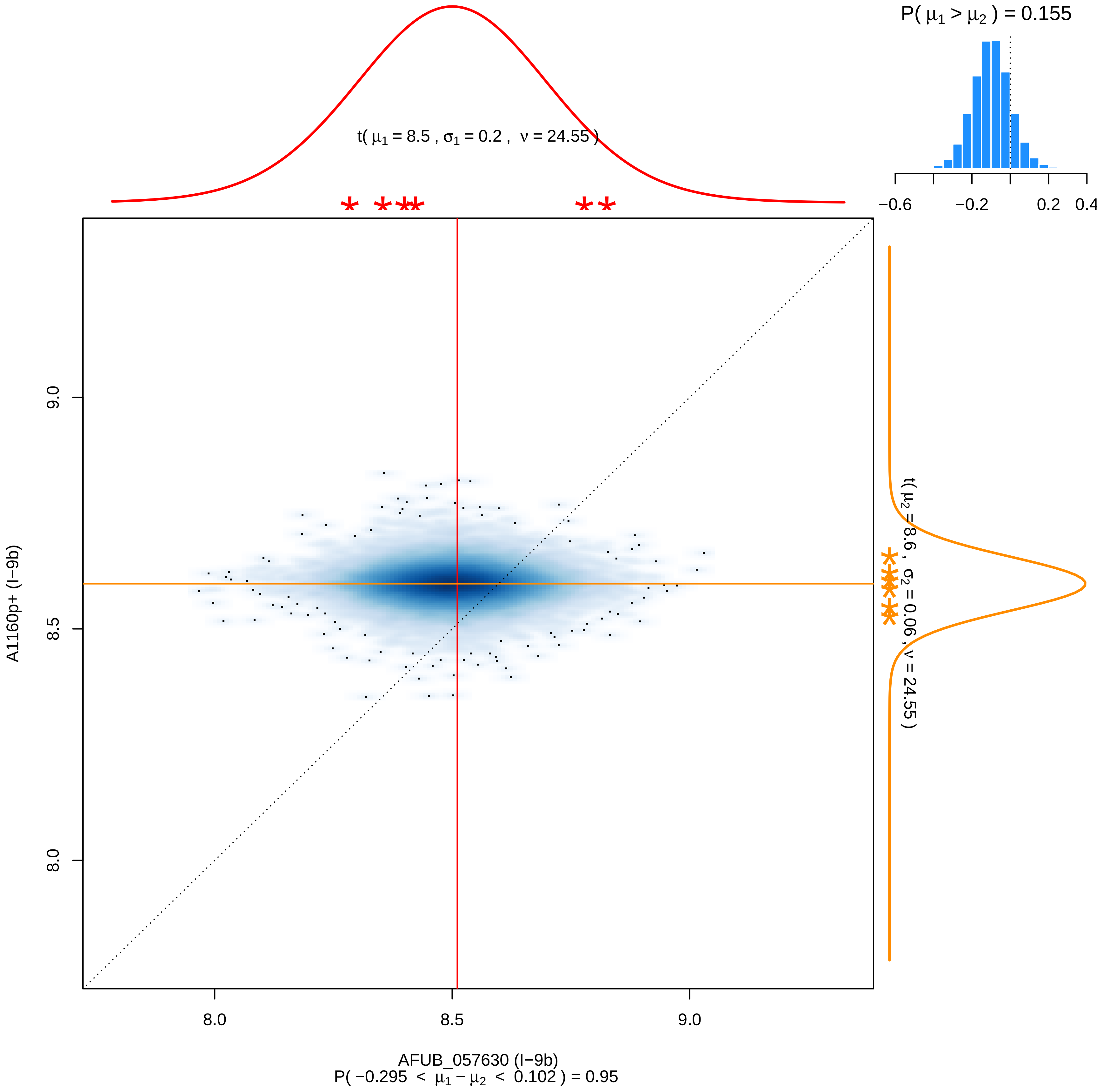

P( μ1 > μ2 ) = 0.155
−0.6
−0.2
0.2
0.4
t( μ1 = 8.5 , σ1 = 0.2 , ν = 24.55 )
t( μ2 = 8.6 , σ2 = 0.06 , ν = 24.55 )
9.0
8.5
8.0
A1160p+ (I−9b)
8.0
8.5
9.0
AFUB_057630 (I−9b)
P( −0.295 < μ1 − μ2 < 0.102 ) = 0.95

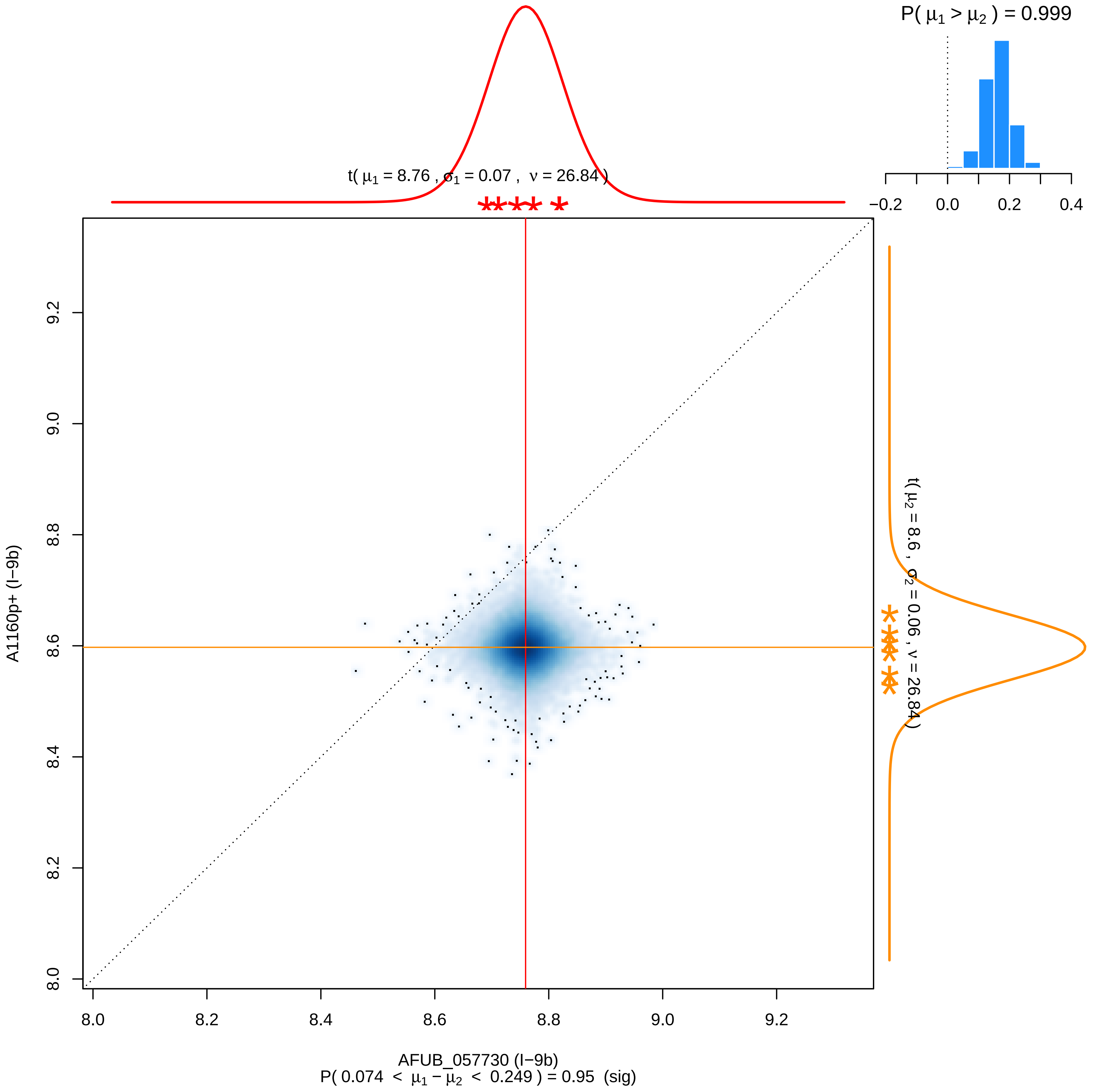

P( $\mu_1 > \mu_2$ ) = 0.999
−0.2
0.0
0.2
0.4
t( $\mu_1$ = 8.76 , $\sigma_1$ = 0.07 , $\nu$ = 26.84 )
t( $\mu_2$ = 8.6 , $\sigma_2$ = 0.06 , $\nu$ = 26.84 )
A1160p+ (I−9b)
8.0
8.2
8.4
8.6
8.8
9.0
9.2
AFUB_057730 (I−9b)
P( 0.074 < $\mu_1 - \mu_2$ < 0.249 ) = 0.95 (sig)

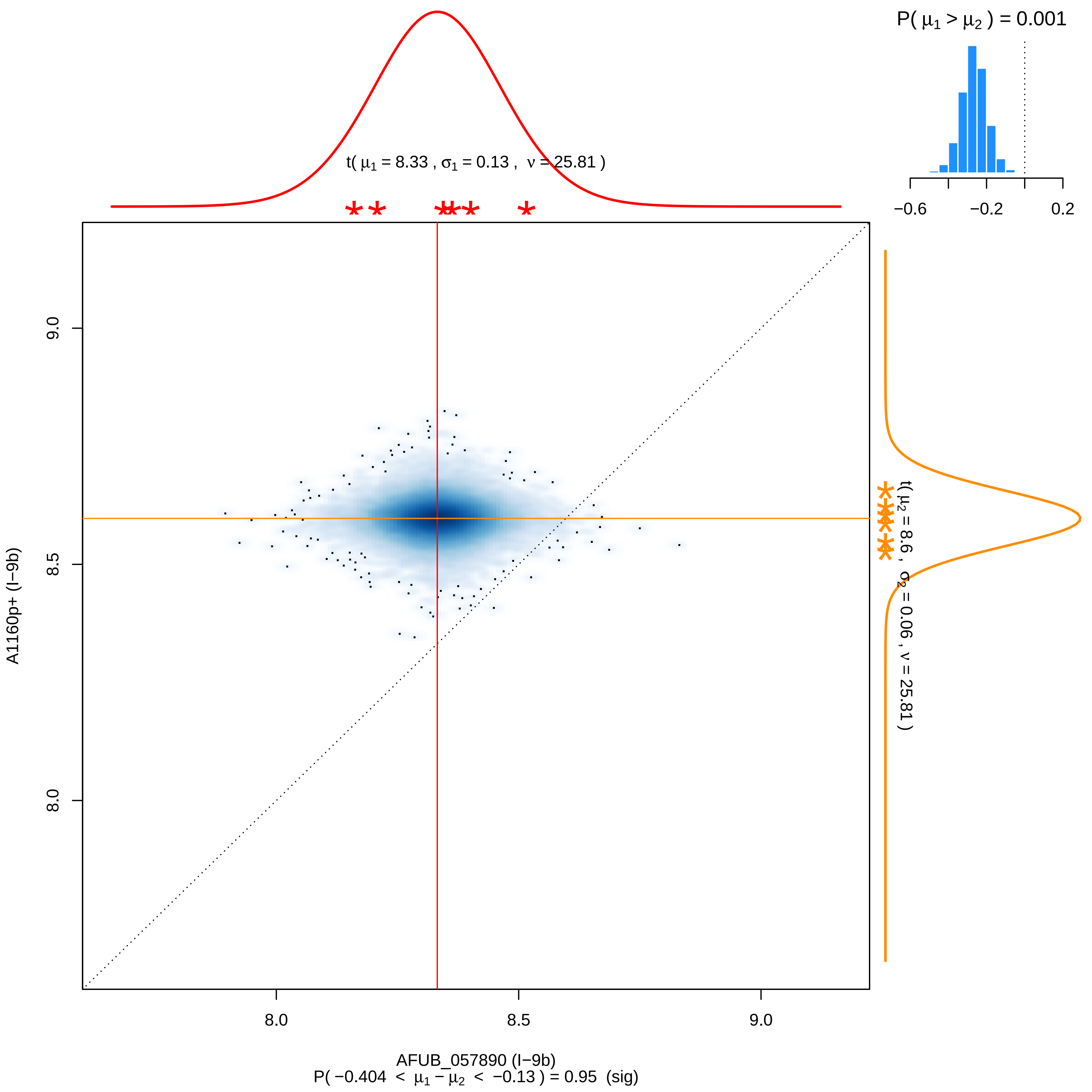

P( μ1 > μ2 ) = 0.001
−0.6
−0.2
0.2
t( μ1 = 8.33 , σ1 = 0.13 , ν = 25.81 )
t( μ2 = 8.6 , σ2 = 0.06 , ν = 25.81 )
9.0
8.5
8.0
A1160p+ (I−9b)
8.0
8.5
9.0
AFUB_057890 (I−9b)
P( −0.404 < μ1 − μ2 < −0.13 ) = 0.95 (sig)

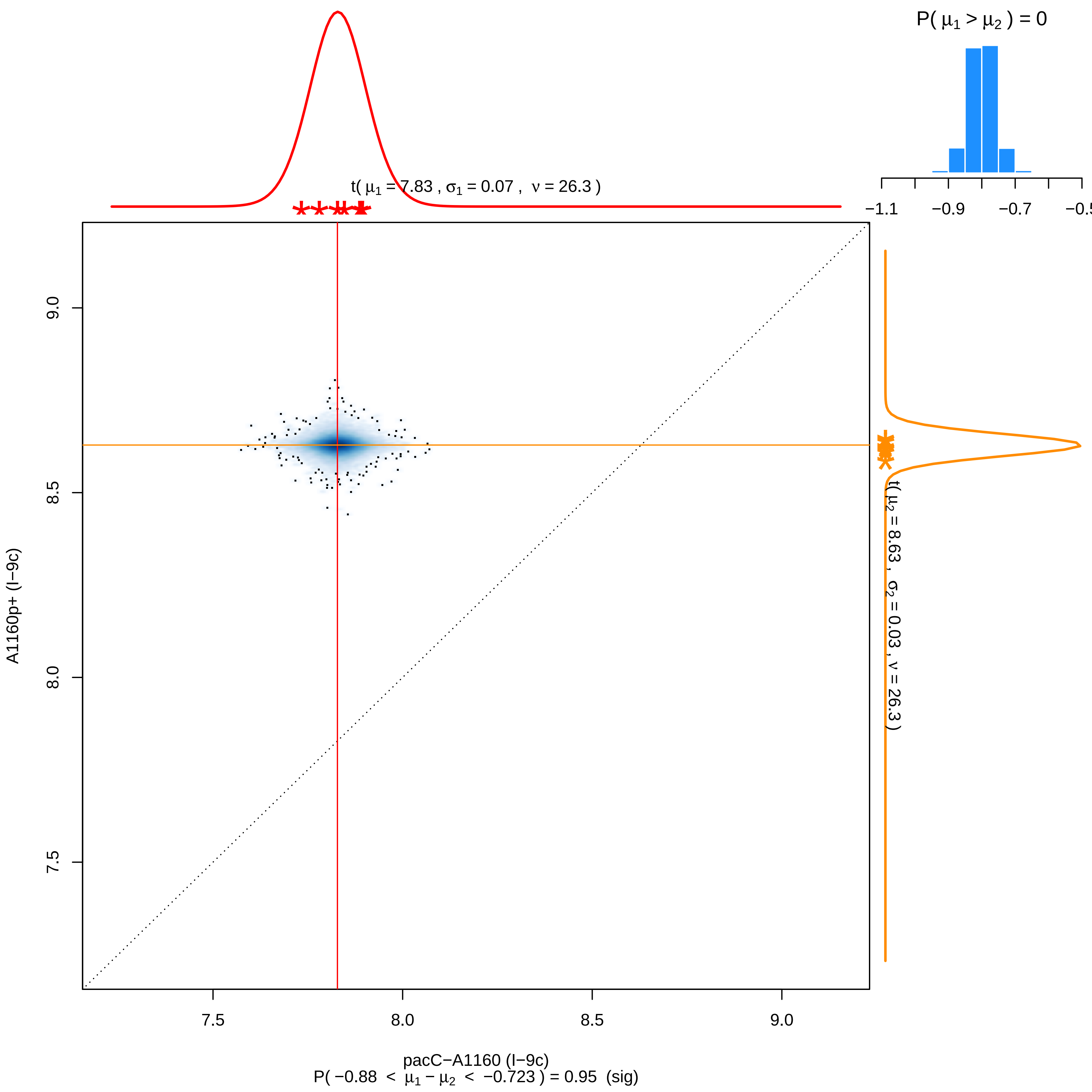
P( $\mu_1 > \mu_2$ ) = 0
−1.1
−0.9
−0.7
−0.5
t( $\mu_1$ = 7.83 , $\sigma_1$ = 0.07 , $\nu$ = 26.3 )
t( $\mu_2$ = 8.63 , $\sigma_2$ = 0.03 , $\nu$ = 26.3 )
A1160p+ (I−9c)
9.0
8.5
8.0
7.5
7.5
8.0
8.5
9.0
pacC−A1160 (I−9c)
P( −0.88 < $\mu_1 - \mu_2$ < −0.723 ) = 0.95 (sig)

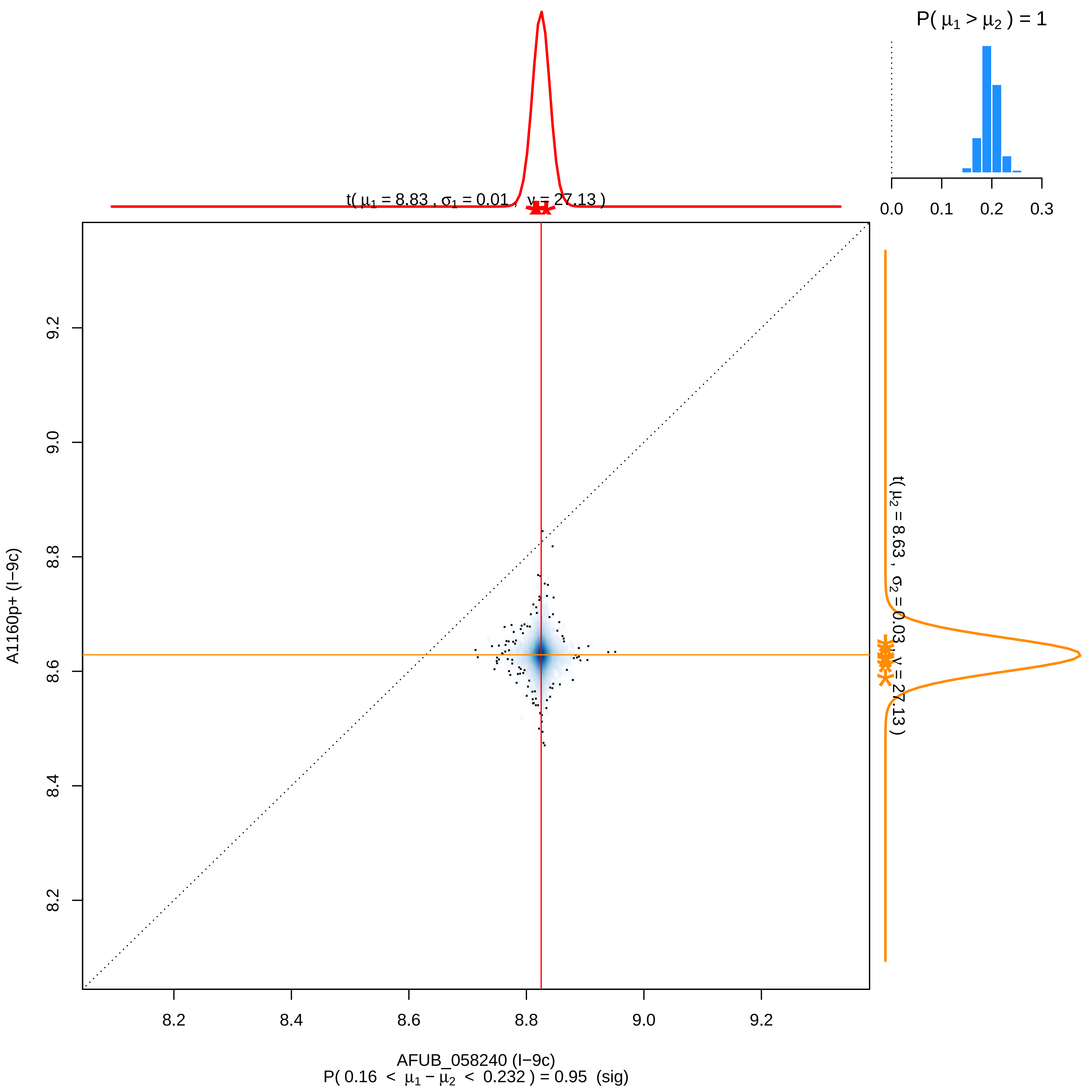

P( μ1 > μ2 ) = 1
0.0
0.1
0.2
0.3
t( μ1 = 8.83 , σ1 = 0.01 , ν = 27.13 )
t( μ2 = 8.63 , σ2 = 0.03 , ν = 27.13 )
9.2
9.0
8.8
8.6
8.4
8.2
A1160p+ (I−9c)
8.2
8.4
8.6
8.8
9.0
9.2
AFUB_058240 (I−9c)
P( 0.16 < μ1 − μ2 < 0.232 ) = 0.95 (sig)

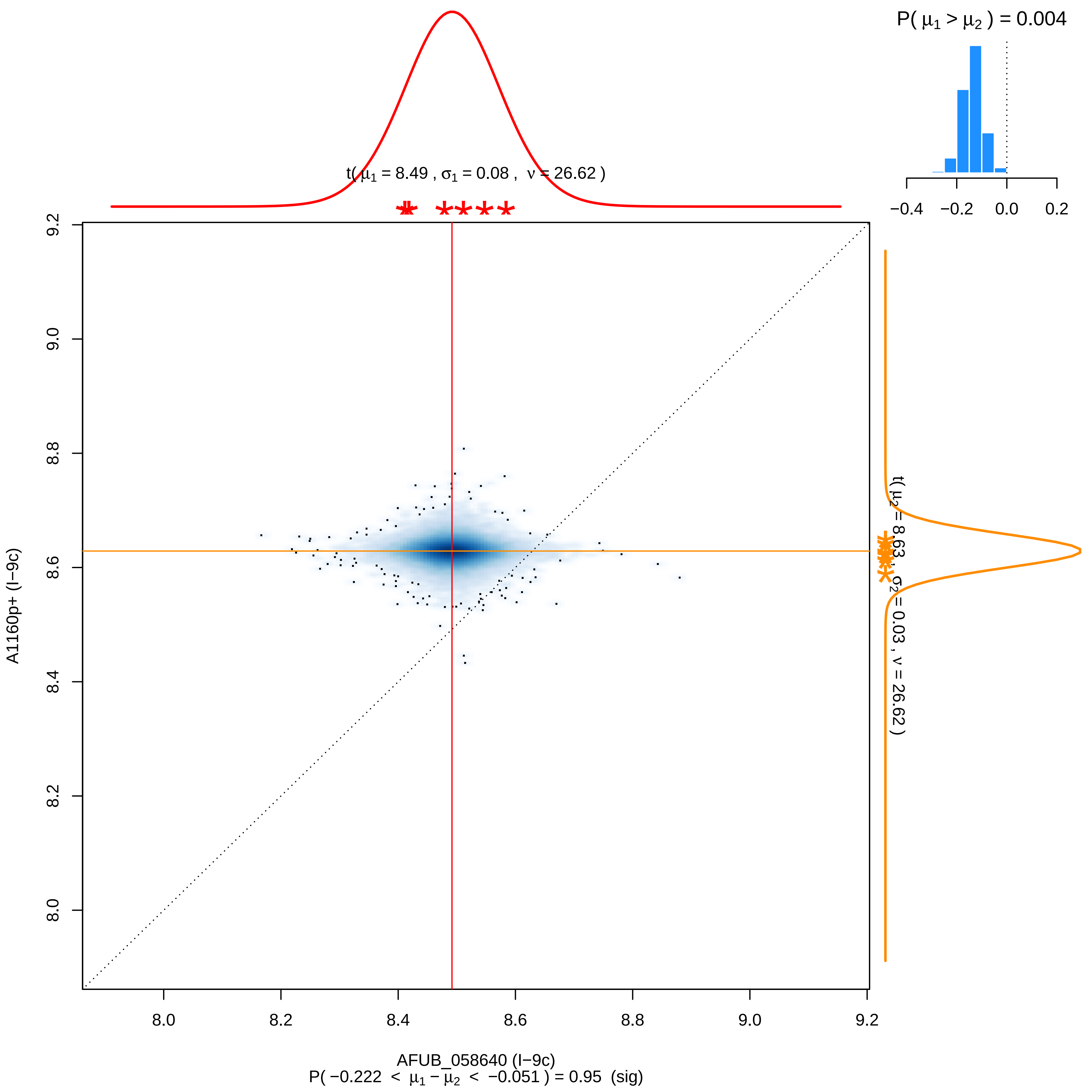

P( μ1 > μ2 ) = 0.004
−0.4
−0.2
0.0
0.2
t( μ1 = 8.49 , σ1 = 0.08 , ν = 26.62 )
t( μ2 = 8.63 , σ2 = 0.03 , ν = 26.62 )
9.2
9.0
8.8
8.6
8.4
8.2
8.0
A1160p+ (I−9c)
8.0
8.2
8.4
8.6
8.8
9.0
9.2
AFUB_058640 (I−9c)
P( −0.222 < μ1 − μ2 < −0.051 ) = 0.95 (sig)

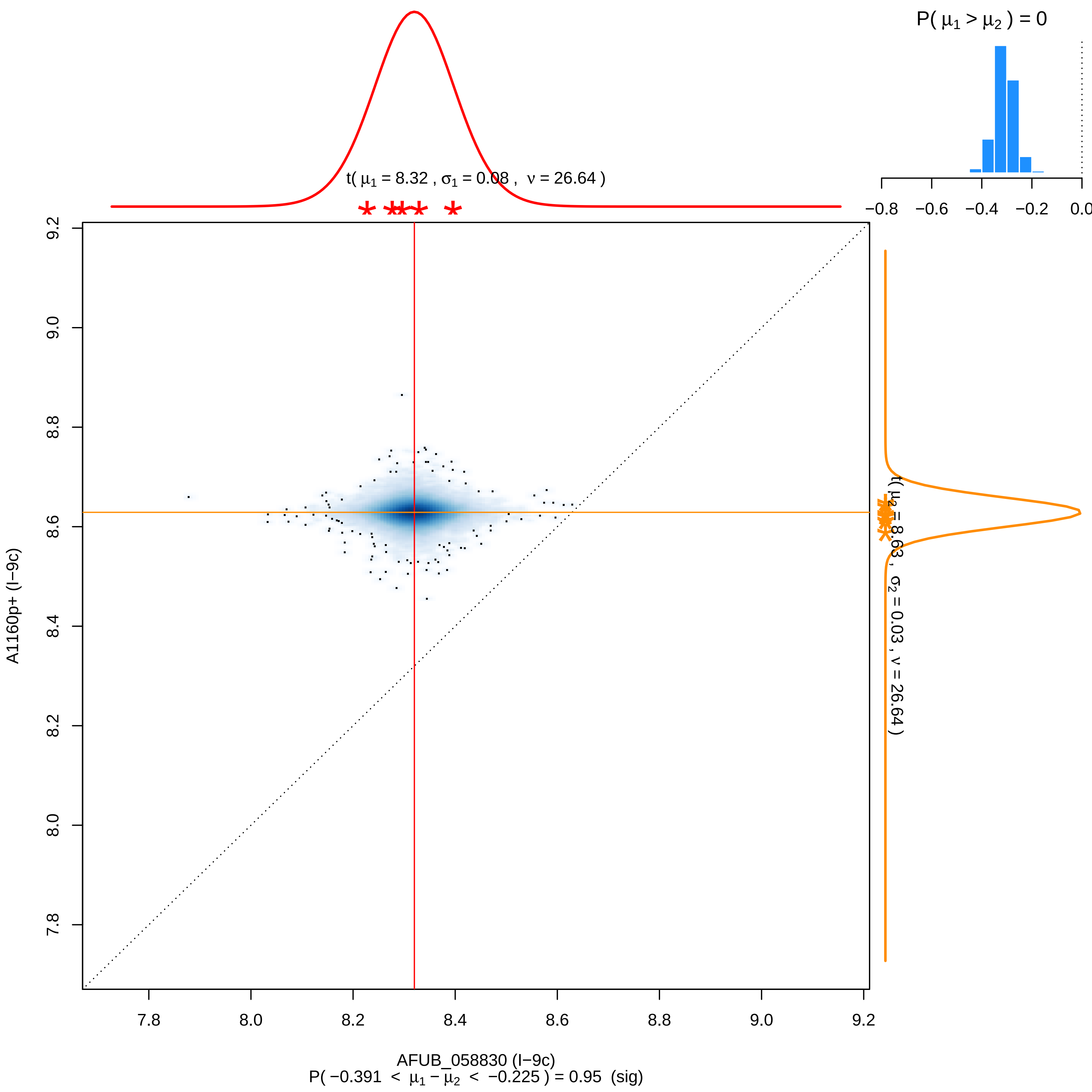

P( $\mu_1 > \mu_2$ ) = 0
−0.8
−0.6
−0.4
−0.2
0.0
t( $\mu_1$ = 8.32 , $\sigma_1$ = 0.08 , $\nu$ = 26.64 )
t( $\mu_2$ = 8.63 , $\sigma_2$ = 0.03 , $\nu$ = 26.64 )
A1160p+ (I−9c)
AFUB_058830 (I−9c)
P( −0.391 < $\mu_1 - \mu_2$ < −0.225 ) = 0.95 (sig)
7.8
8.0
8.2
8.4
8.6
8.8
9.0
9.2

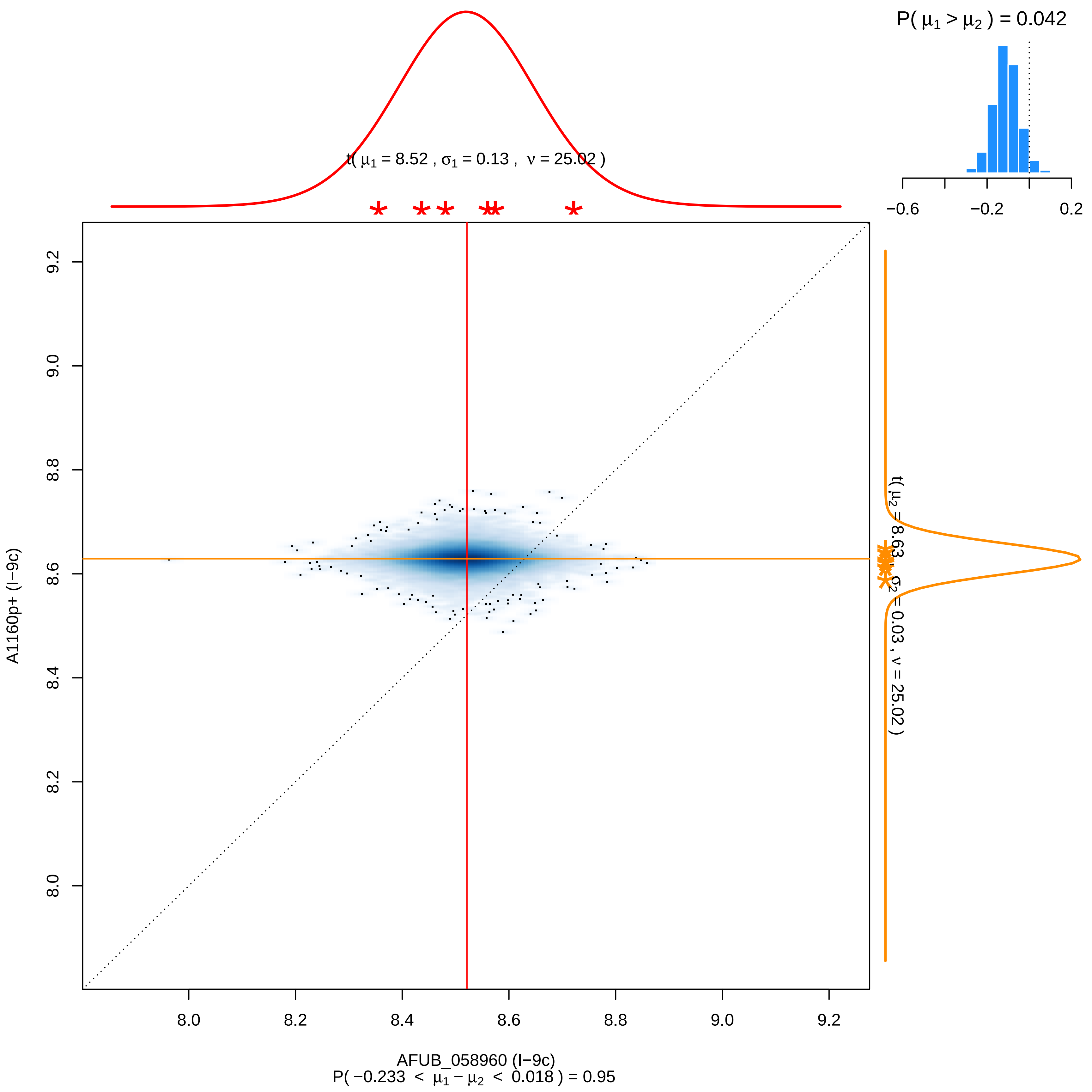

P( μ1 > μ2 ) = 0.042
−0.6
−0.2
0.2
t( μ1 = 8.52 , σ1 = 0.13 , ν = 25.02 )
t( μ2 = 8.63 , σ2 = 0.03 , ν = 25.02 )
A1160p+ (I−9c)
AFUB_058960 (I−9c)
P( −0.233 < μ1 − μ2 < 0.018 ) = 0.95
8.0
8.2
8.4
8.6
8.8
9.0
9.2

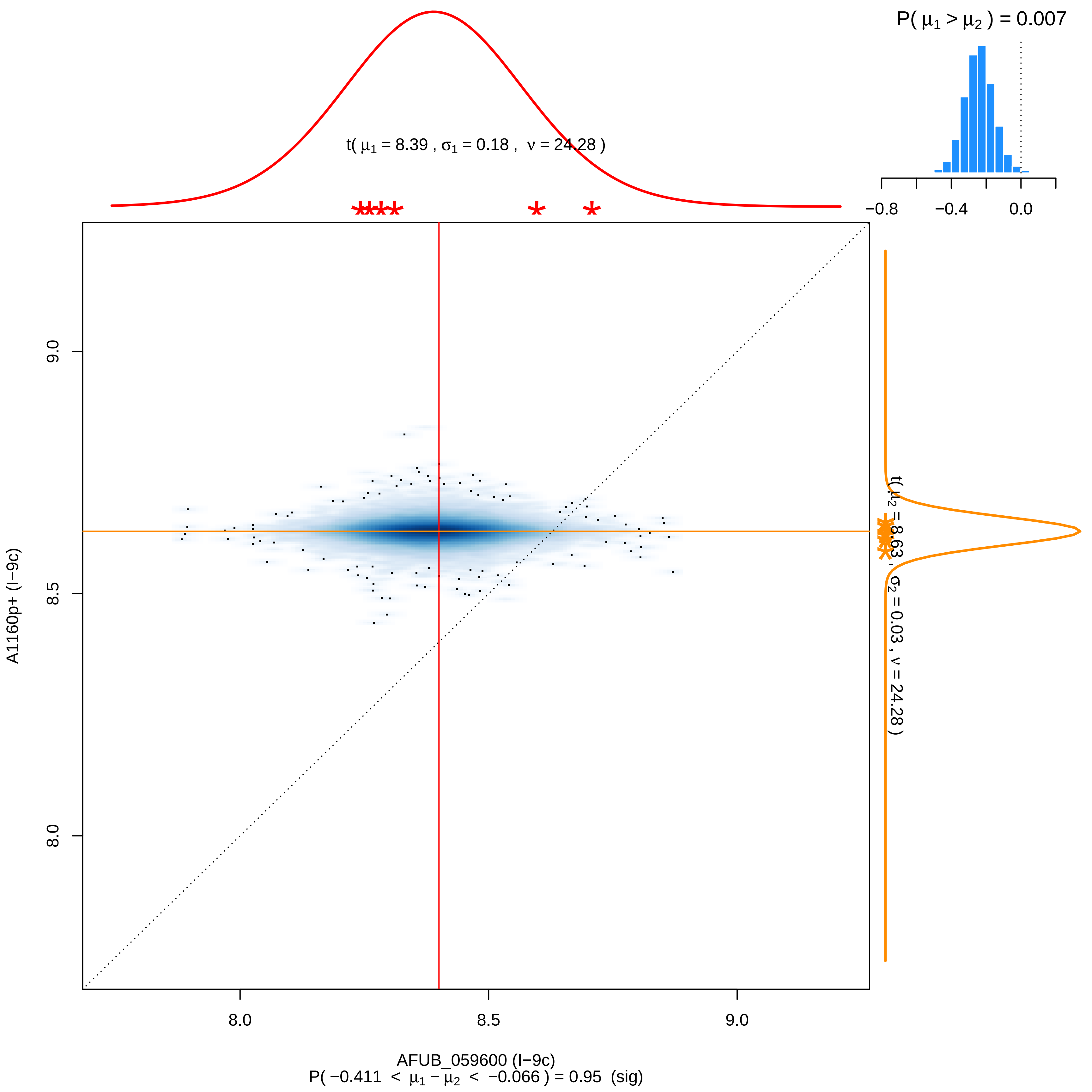

P( μ1 > μ2 ) = 0.007
−0.8
−0.4
0.0
t( μ1 = 8.39 , σ1 = 0.18 , ν = 24.28 )
t( μ2 = 8.63 , σ2 = 0.03 , ν = 24.28 )
A1160p+ (I−9c)
9.0
8.5
8.0
AFUB_059600 (I−9c)
P( −0.411 < μ1 − μ2 < −0.066 ) = 0.95 (sig)

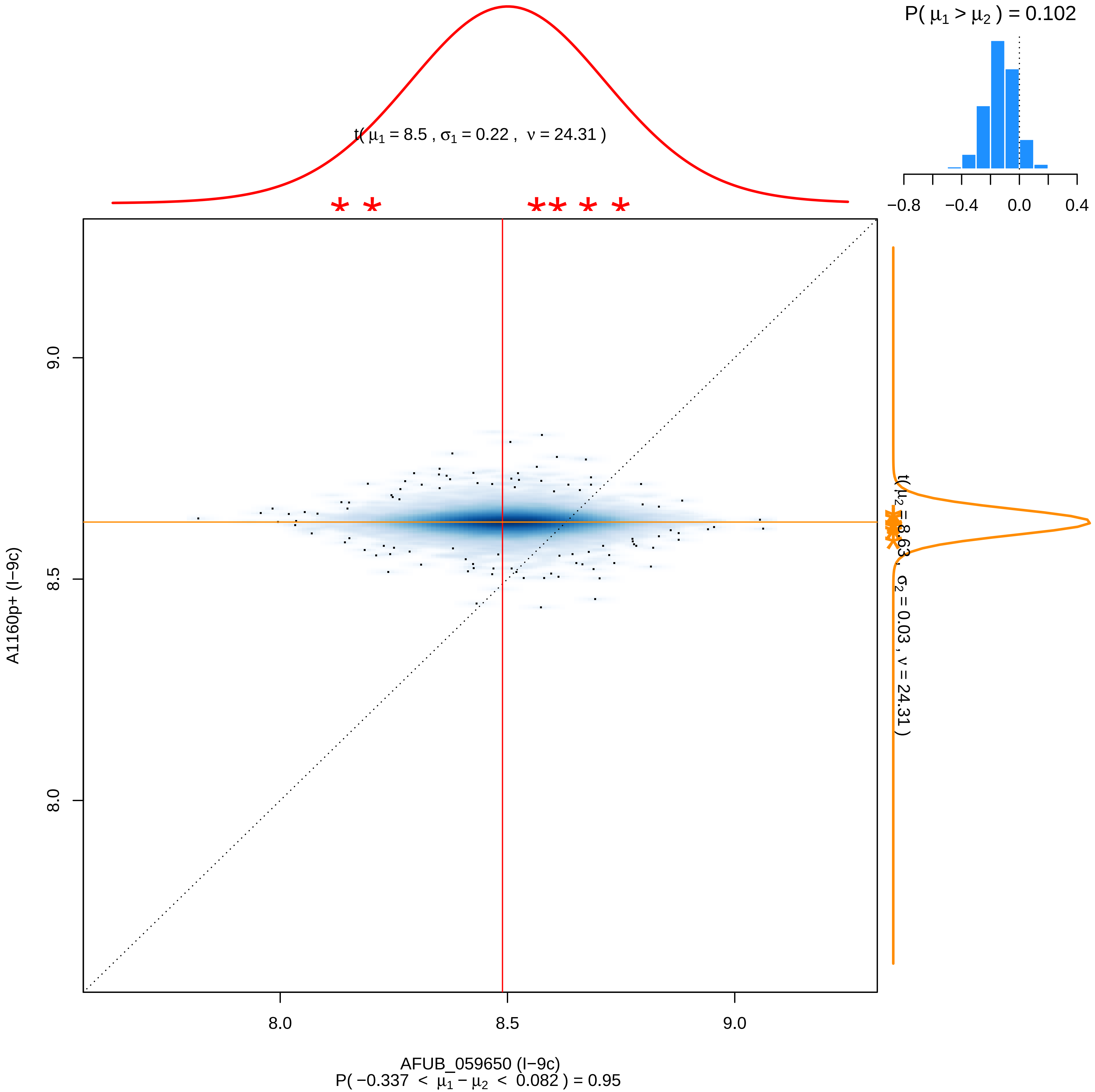

P( μ1 > μ2 ) = 0.102
−0.8
−0.4
0.0
0.4
t( μ1 = 8.5 , σ1 = 0.22 , ν = 24.31 )
t( μ2 = 8.63 , σ2 = 0.03 , ν = 24.31 )
9.0
8.5
8.0
A1160p+ (I−9c)
8.0
8.5
9.0
AFUB_059650 (I−9c)
P( −0.337 < μ1 − μ2 < 0.082 ) = 0.95

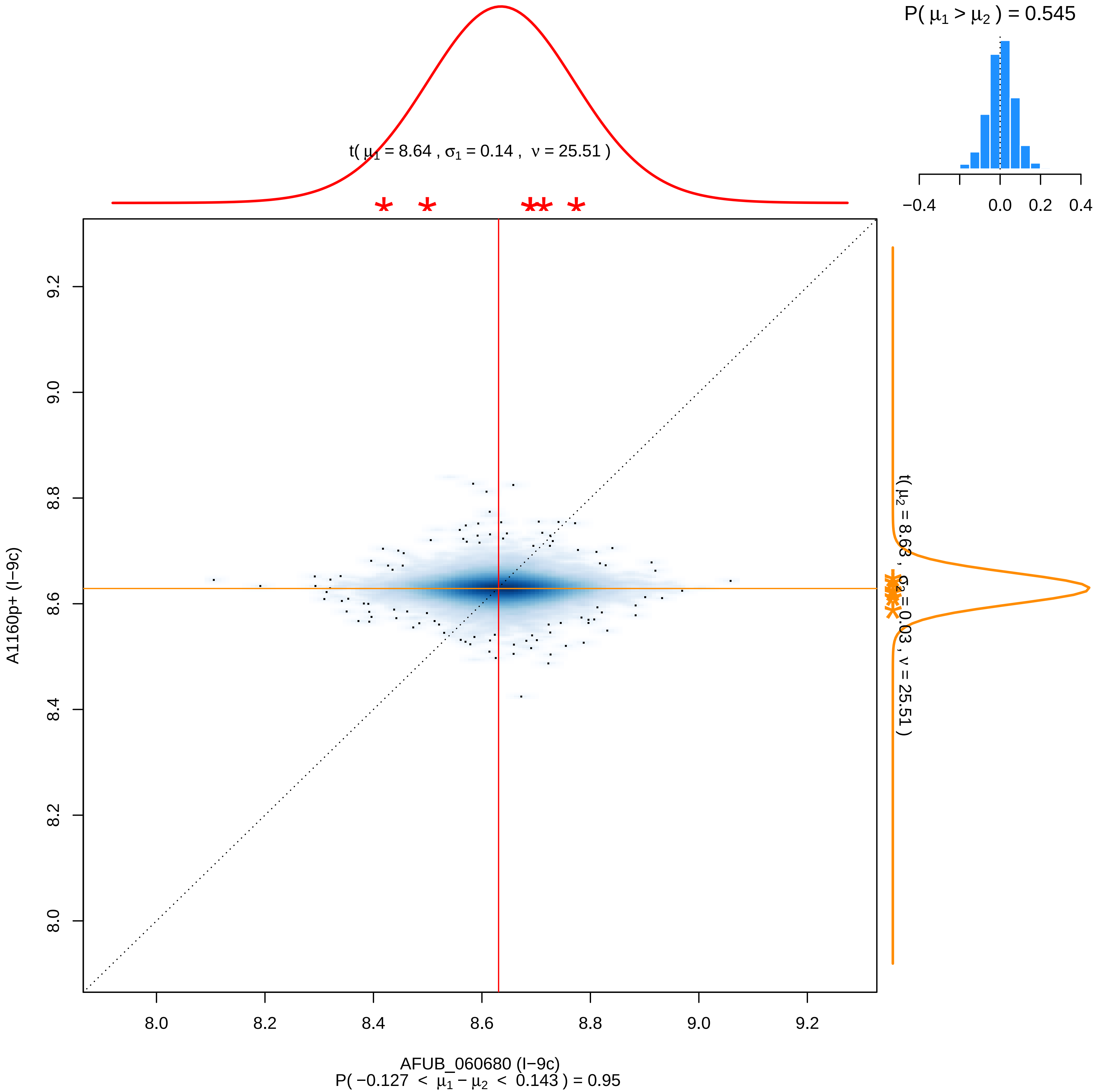

P( μ1 > μ2 ) = 0.545
−0.4
0.0
0.2
0.4
t( μ1 = 8.64 , σ1 = 0.14 , ν = 25.51 )
t( μ2 = 8.63 , σ2 = 0.03 , ν = 25.51 )
A1160p+ (I−9c)
9.2
9.0
8.8
8.6
8.4
8.2
8.0
AFUB_060680 (I−9c)
P( −0.127 < μ1 − μ2 < 0.143 ) = 0.95

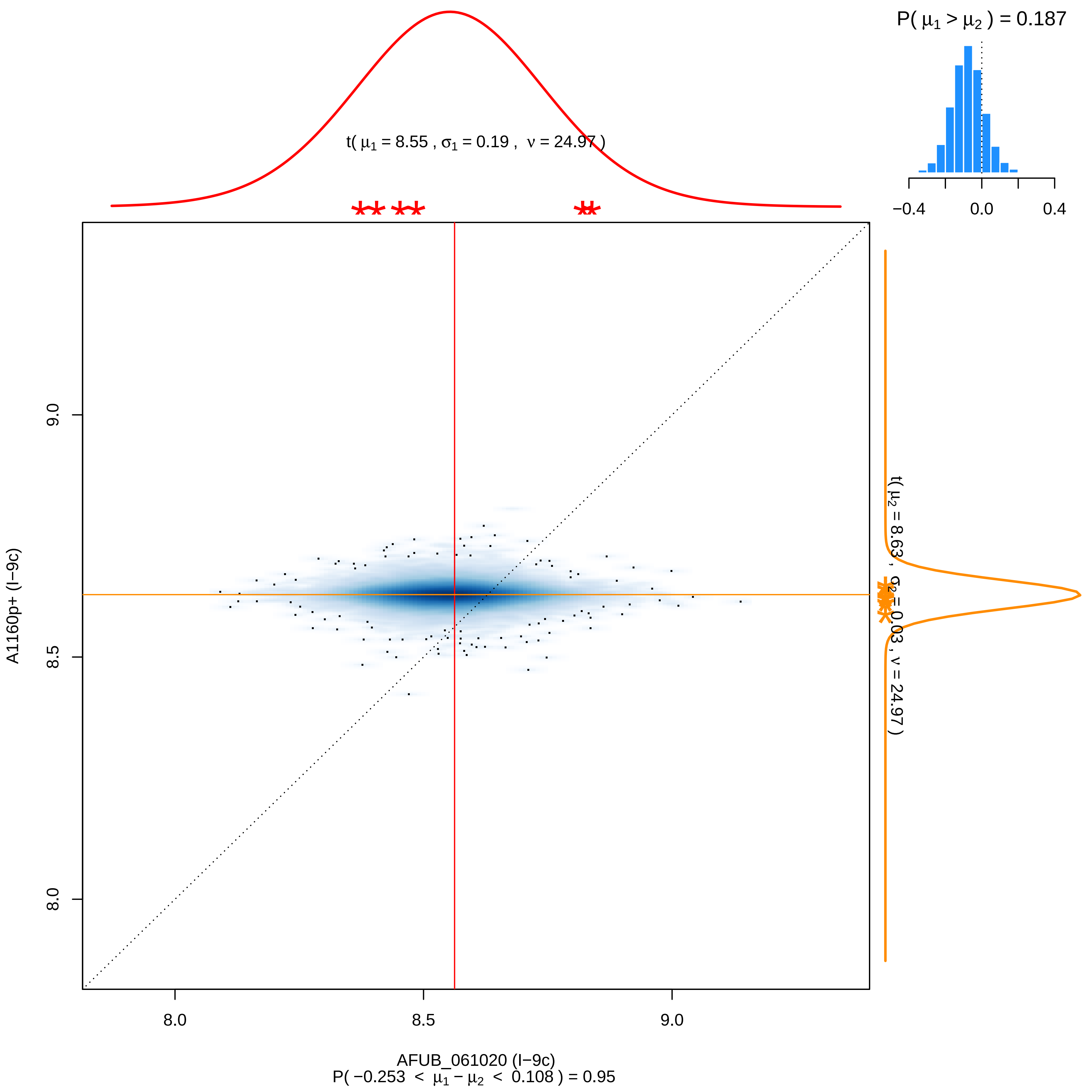

P( μ1 > μ2 ) = 0.187
−0.4
0.0
0.4
t( μ1 = 8.55 , σ1 = 0.19 , ν = 24.97 )
t( μ2 = 8.63 , σ2 = 0.03 , ν = 24.97 )
9.0
8.5
8.0
A1160p+ (I−9c)
8.0
8.5
9.0
AFUB_061020 (I−9c)
P( −0.253 < μ1 − μ2 < 0.108 ) = 0.95

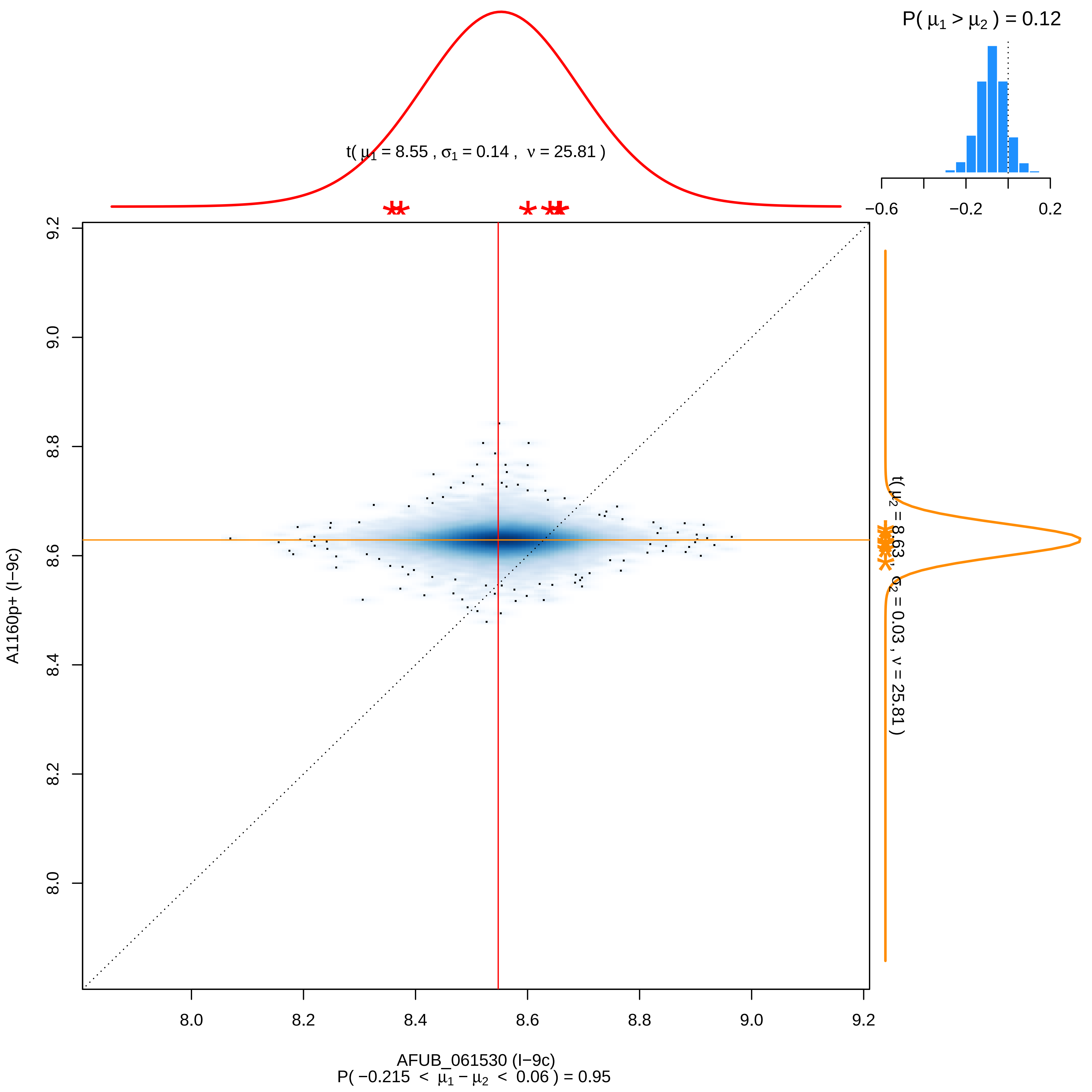

P( μ1 > μ2 ) = 0.12
−0.6
−0.2
0.2
t( μ1 = 8.55 , σ1 = 0.14 , ν = 25.81 )
t( μ2 = 8.63 , σ2 = 0.03 , ν = 25.81 )
A1160p+ (I−9c)
9.2
9.0
8.8
8.6
8.4
8.2
8.0
AFUB_061530 (I−9c)
P( −0.215 < μ1 − μ2 < 0.06 ) = 0.95

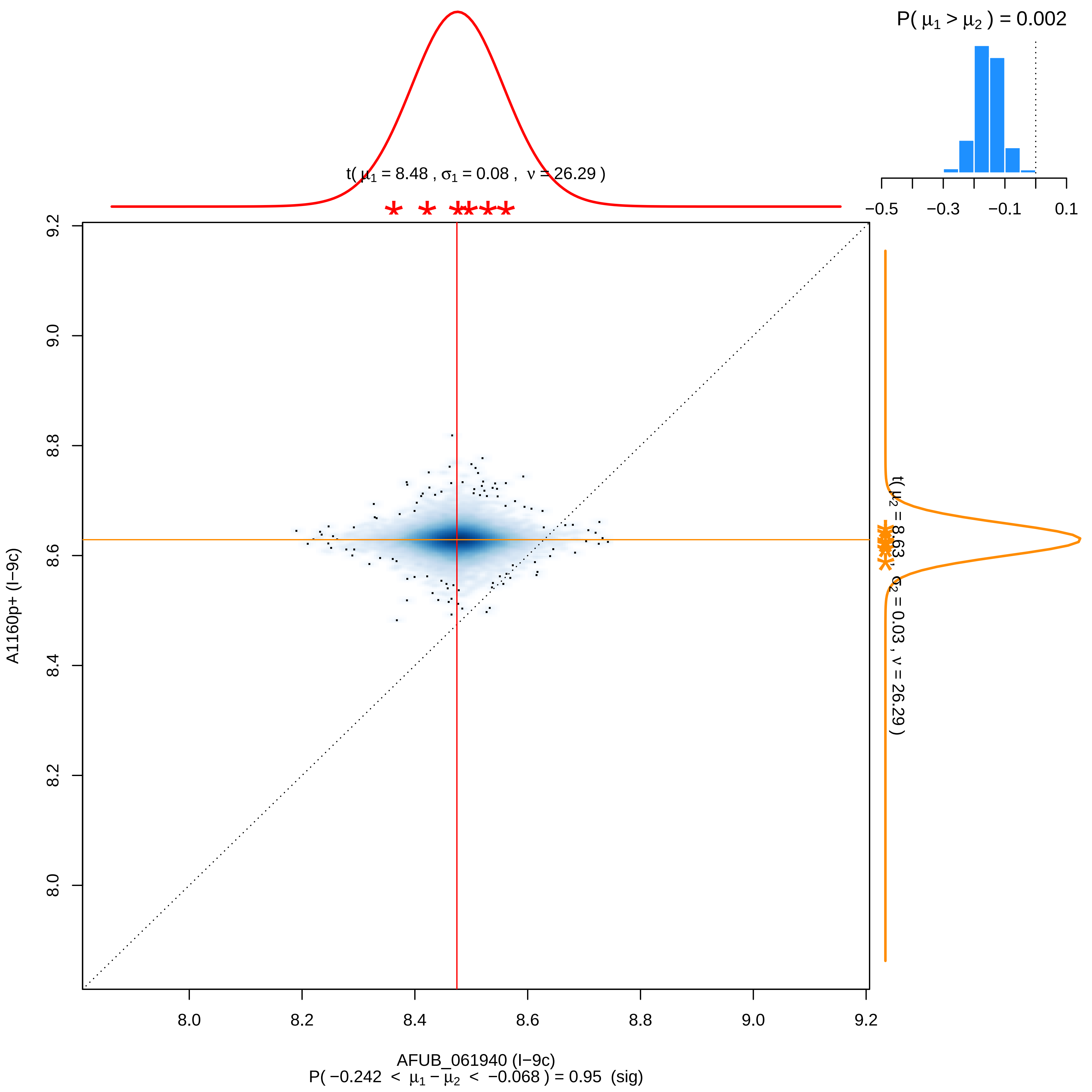

P( μ1 > μ2 ) = 0.002
−0.5
−0.3
−0.1
0.1
t( μ1 = 8.48 , σ1 = 0.08 , ν = 26.29 )
t( μ2 = 8.63 , σ2 = 0.03 , ν = 26.29 )
A1160p+ (I−9c)
8.0
8.2
8.4
8.6
8.8
9.0
9.2
AFUB_061940 (I−9c)
P( −0.242 < μ1 − μ2 < −0.068 ) = 0.95 (sig)

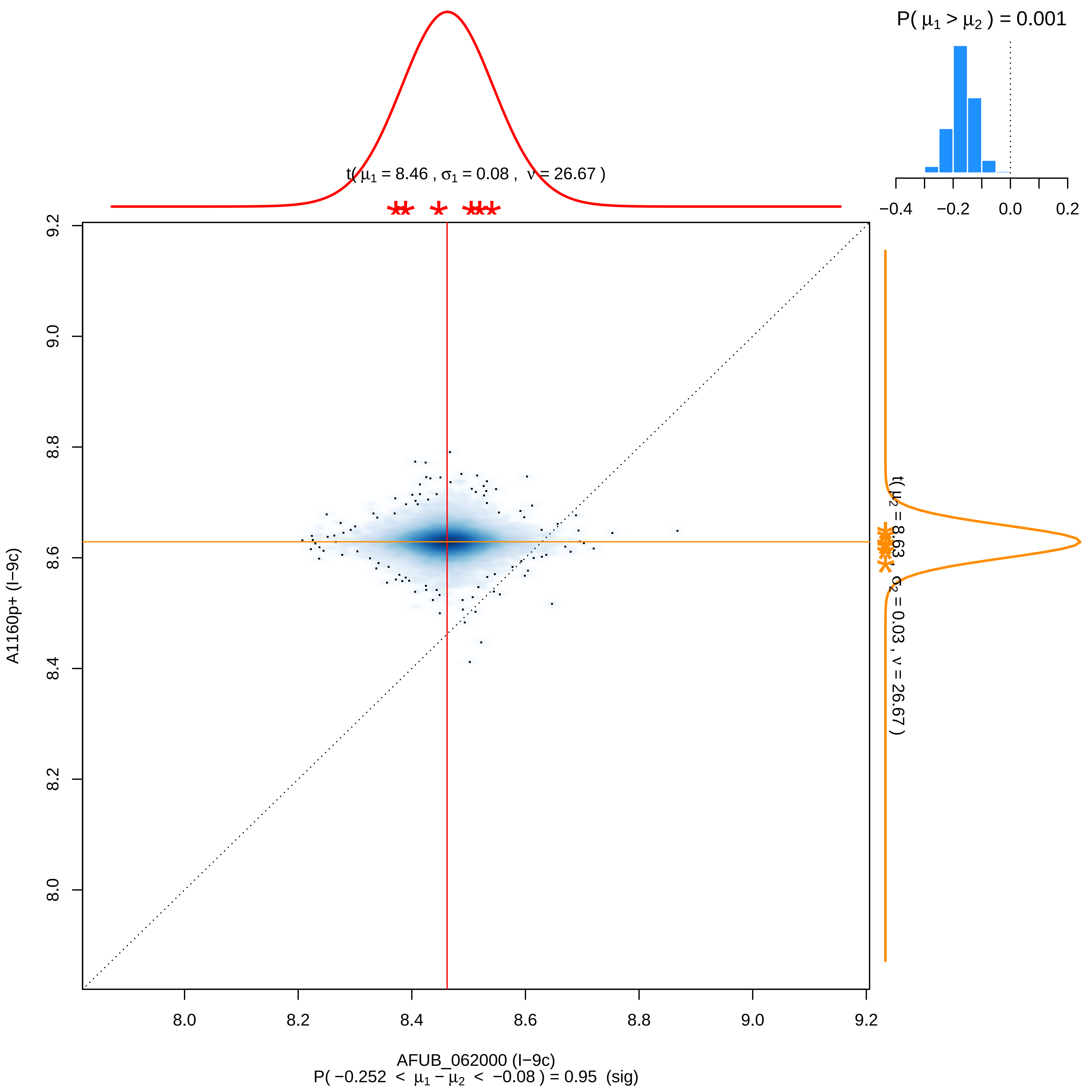

P( μ1 > μ2 ) = 0.001
−0.4
−0.2
0.0
0.2
t( μ1 = 8.46 , σ1 = 0.08 , ν = 26.67 )
t( μ2 = 8.63 , σ2 = 0.03 , ν = 26.67 )
9.2
9.0
8.8
8.6
8.4
8.2
8.0
A1160p+ (I−9c)
8.0
8.2
8.4
8.6
8.8
9.0
9.2
AFUB_062000 (I−9c)
P( −0.252 < μ1 − μ2 < −0.08 ) = 0.95 (sig)

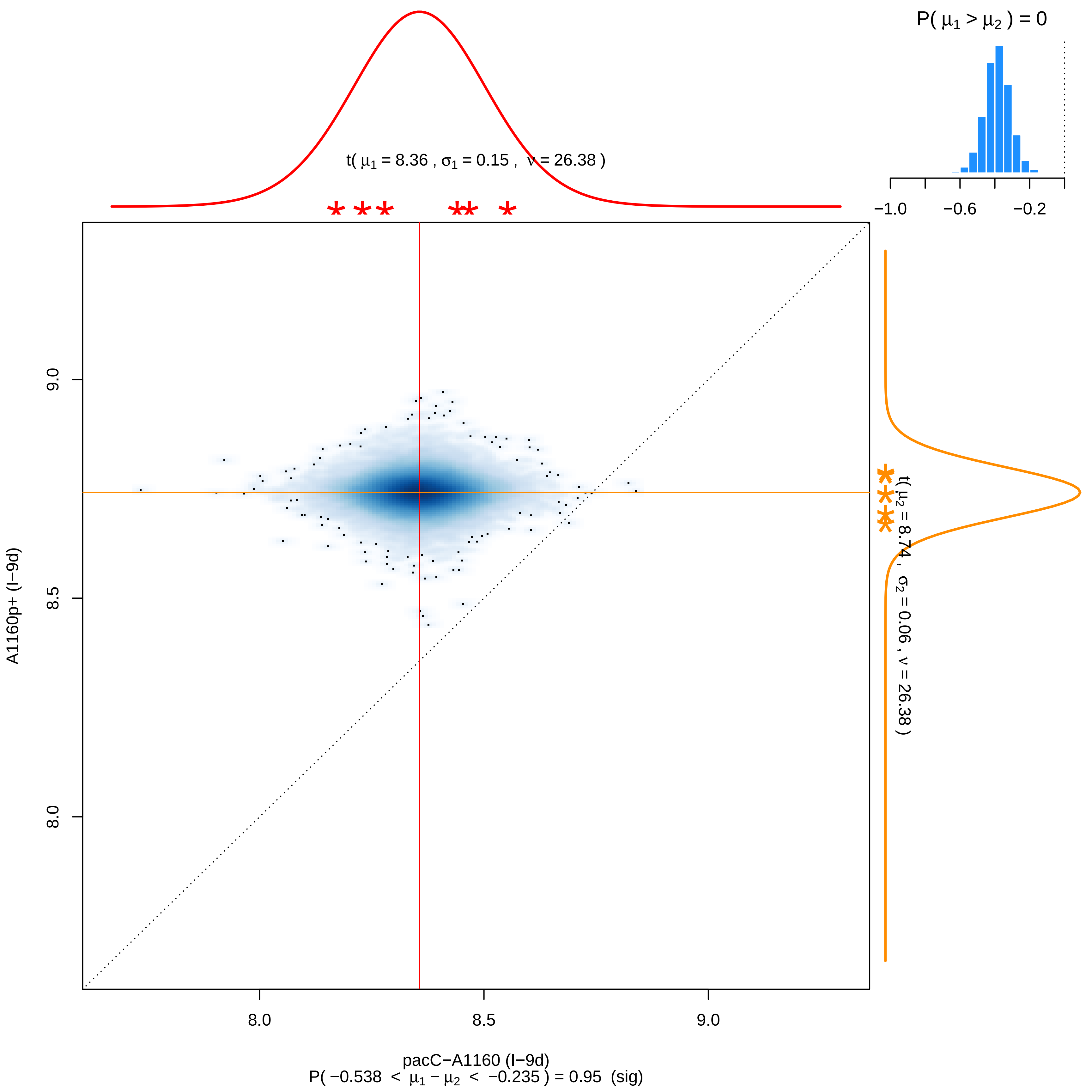

P( μ1 > μ2 ) = 0
−1.0
−0.6
−0.2
t( μ1 = 8.36 , σ1 = 0.15 , ν = 26.38 )
t( μ2 = 8.74 , σ2 = 0.06 , ν = 26.38 )
9.0
8.5
8.0
A1160p+ (I−9d)
8.0
8.5
9.0
pacC−A1160 (I−9d)
P( −0.538 < μ1 − μ2 < −0.235 ) = 0.95 (sig)

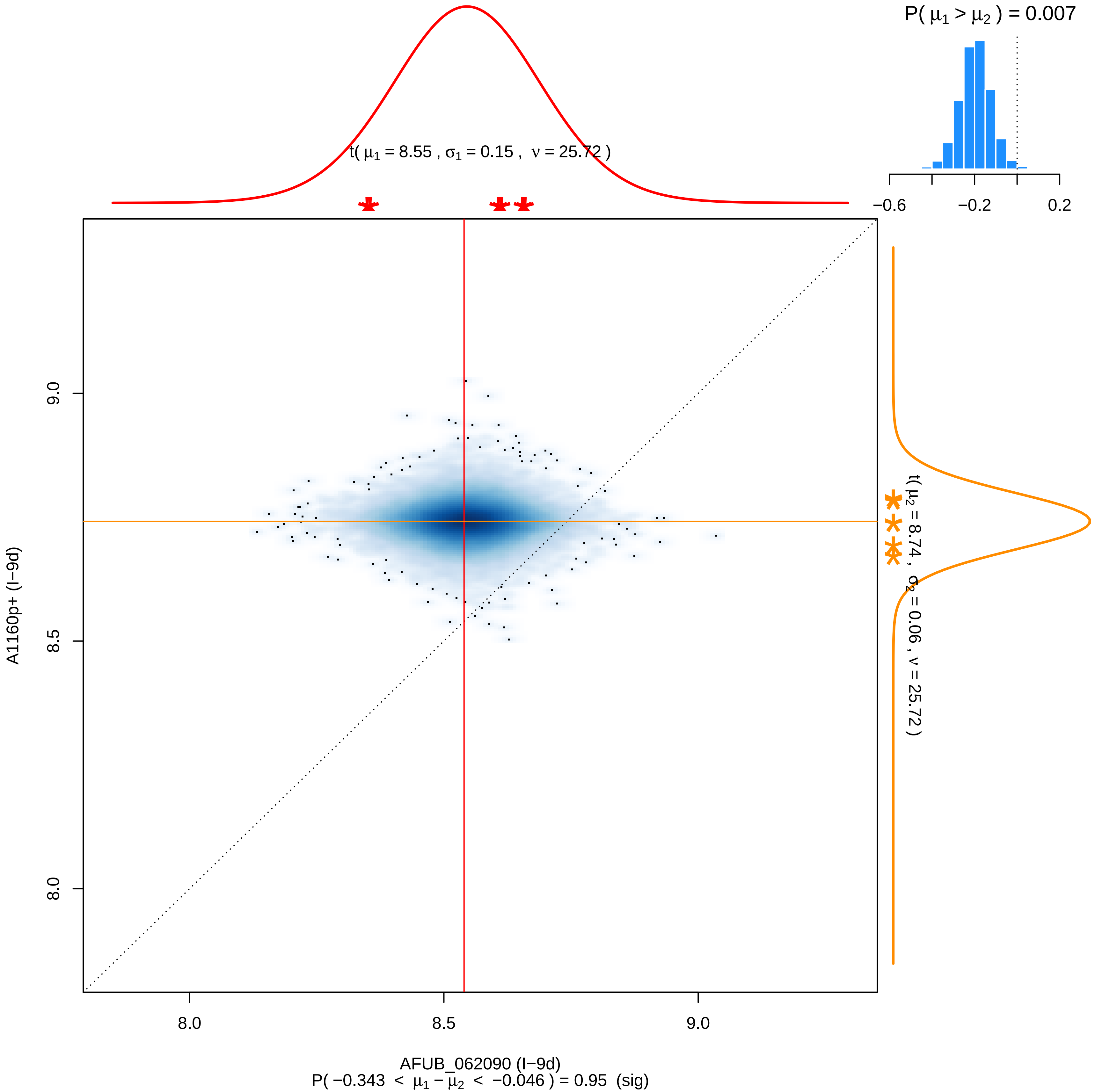

P( μ1 > μ2 ) = 0.007
−0.6
−0.2
0.2
t( μ1 = 8.55 , σ1 = 0.15 , ν = 25.72 )
t( μ2 = 8.74 , σ2 = 0.06 , ν = 25.72 )
A1160p+ (I−9d)
9.0
8.5
8.0
8.0
8.5
9.0
AFUB_062090 (I−9d)
P( −0.343 < μ1 − μ2 < −0.046 ) = 0.95 (sig)

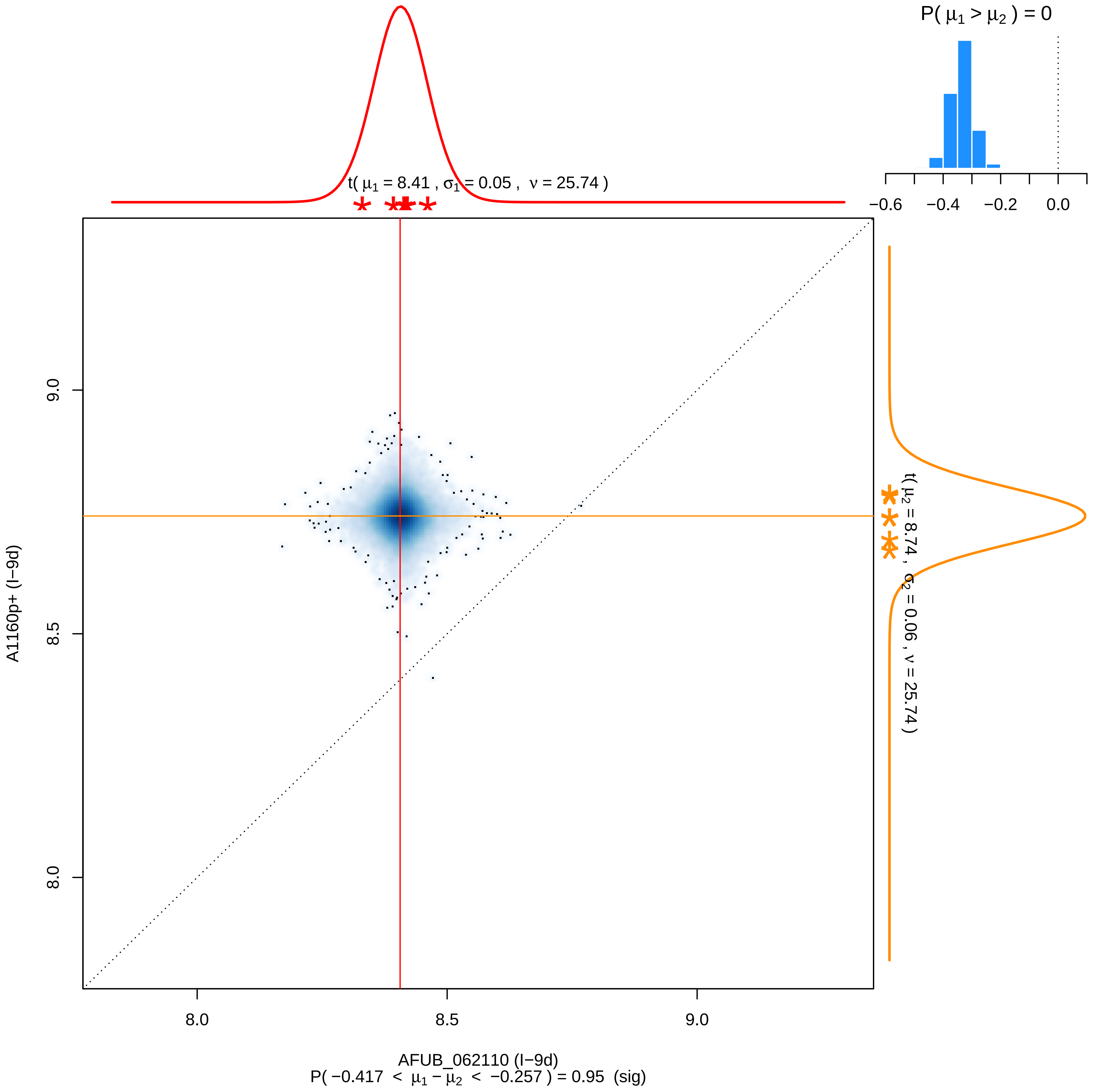

P( μ1 > μ2 ) = 0
−0.6
−0.4
−0.2
0.0
t( μ1 = 8.41 , σ1 = 0.05 , ν = 25.74 )
t( μ2 = 8.74 , σ2 = 0.06 , ν = 25.74 )
9.0
8.5
8.0
A1160p+ (I−9d)
8.0
8.5
9.0
AFUB_062110 (I−9d)
P( −0.417 < μ1 − μ2 < −0.257 ) = 0.95 (sig)

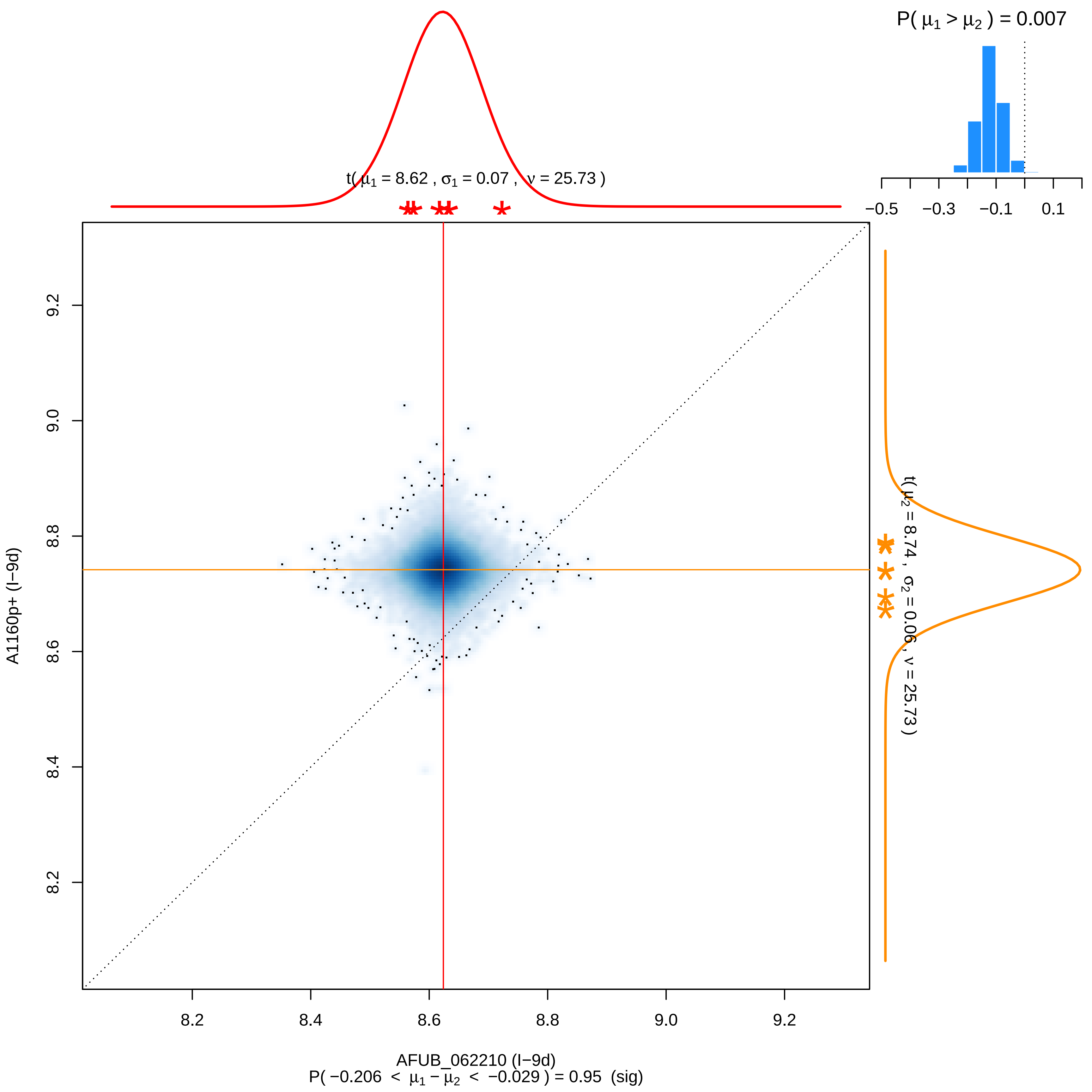

P( μ1 > μ2 ) = 0.007
−0.5
−0.3
−0.1
0.1
t( μ1 = 8.62 , σ1 = 0.07 , ν = 25.73 )
t( μ2 = 8.74 , σ2 = 0.06 , ν = 25.73 )
A1160p+ (I−9d)
9.2
9.0
8.8
8.6
8.4
8.2
AFUB_062210 (I−9d)
P( −0.206 < μ1 − μ2 < −0.029 ) = 0.95 (sig)

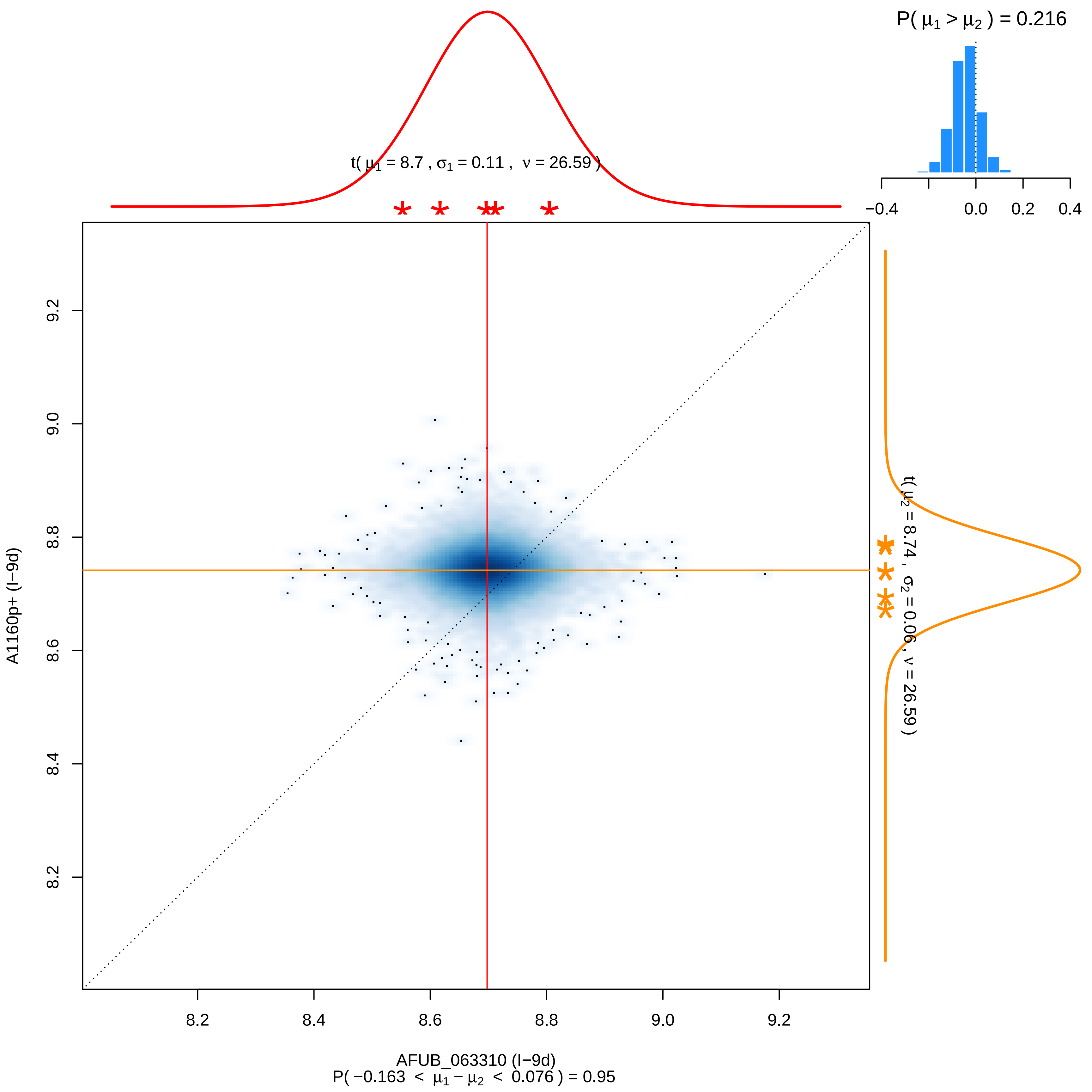
P( μ1 > μ2 ) = 0.216
−0.4
0.0
0.2
0.4
t( μ1 = 8.7 , σ1 = 0.11 , ν = 26.59 )
t( μ2 = 8.74 , σ2 = 0.06 , ν = 26.59 )
9.2
9.0
8.8
8.6
8.4
8.2
A1160p+ (I−9d)
8.2
8.4
8.6
8.8
9.0
9.2
AFUB_063310 (I−9d)
P( −0.163 < μ1 − μ2 < 0.076 ) = 0.95

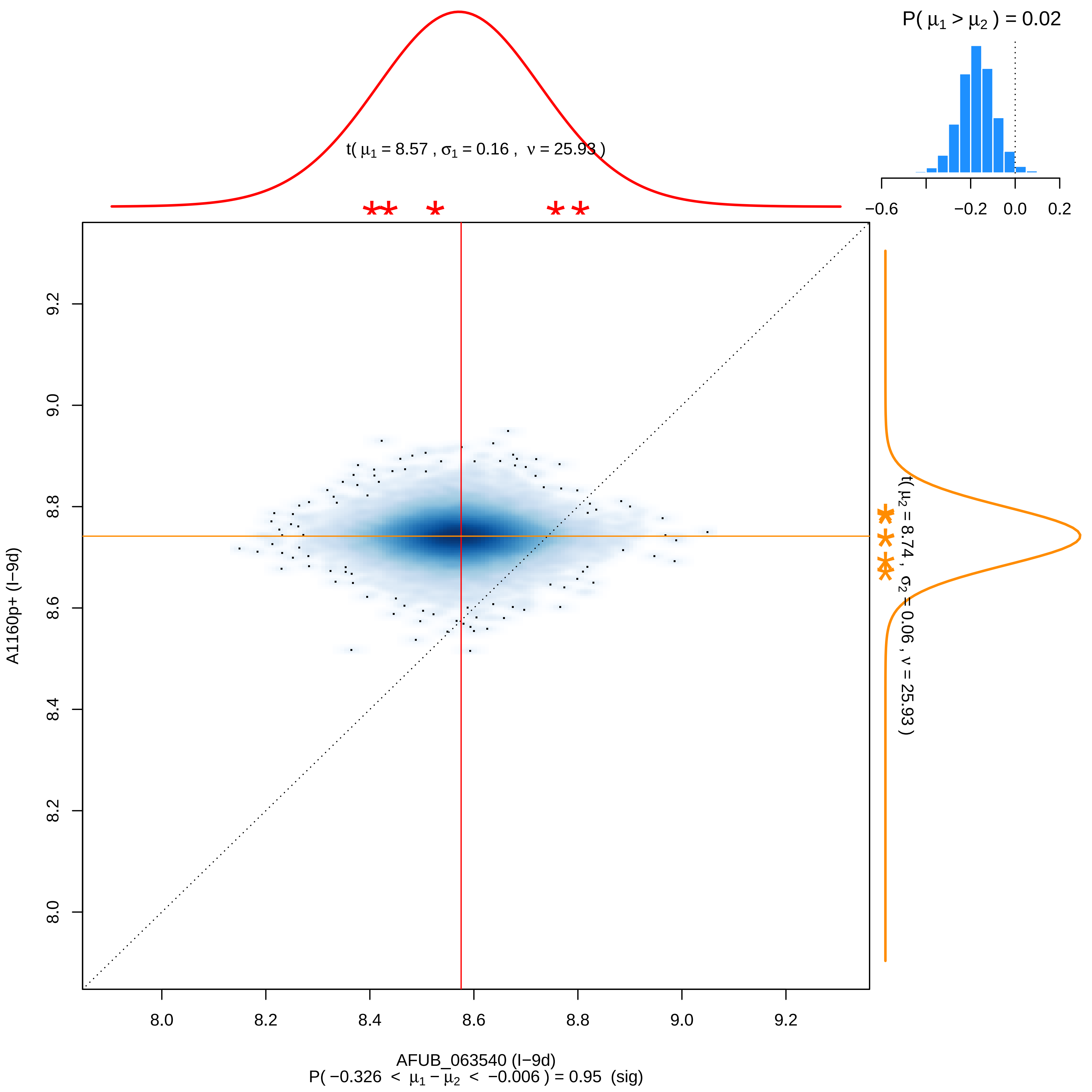

P( μ1 > μ2 ) = 0.02
−0.6
−0.2
0.0
0.2
t( μ1 = 8.57 , σ1 = 0.16 , ν = 25.93 )
t( μ2 = 8.74 , σ2 = 0.06 , ν = 25.93 )
A1160p+ (I−9d)
9.2
9.0
8.8
8.6
8.4
8.2
8.0
8.0
8.2
8.4
8.6
8.8
9.0
9.2
AFUB_063540 (I−9d)
P( −0.326 < μ1 − μ2 < −0.006 ) = 0.95 (sig)

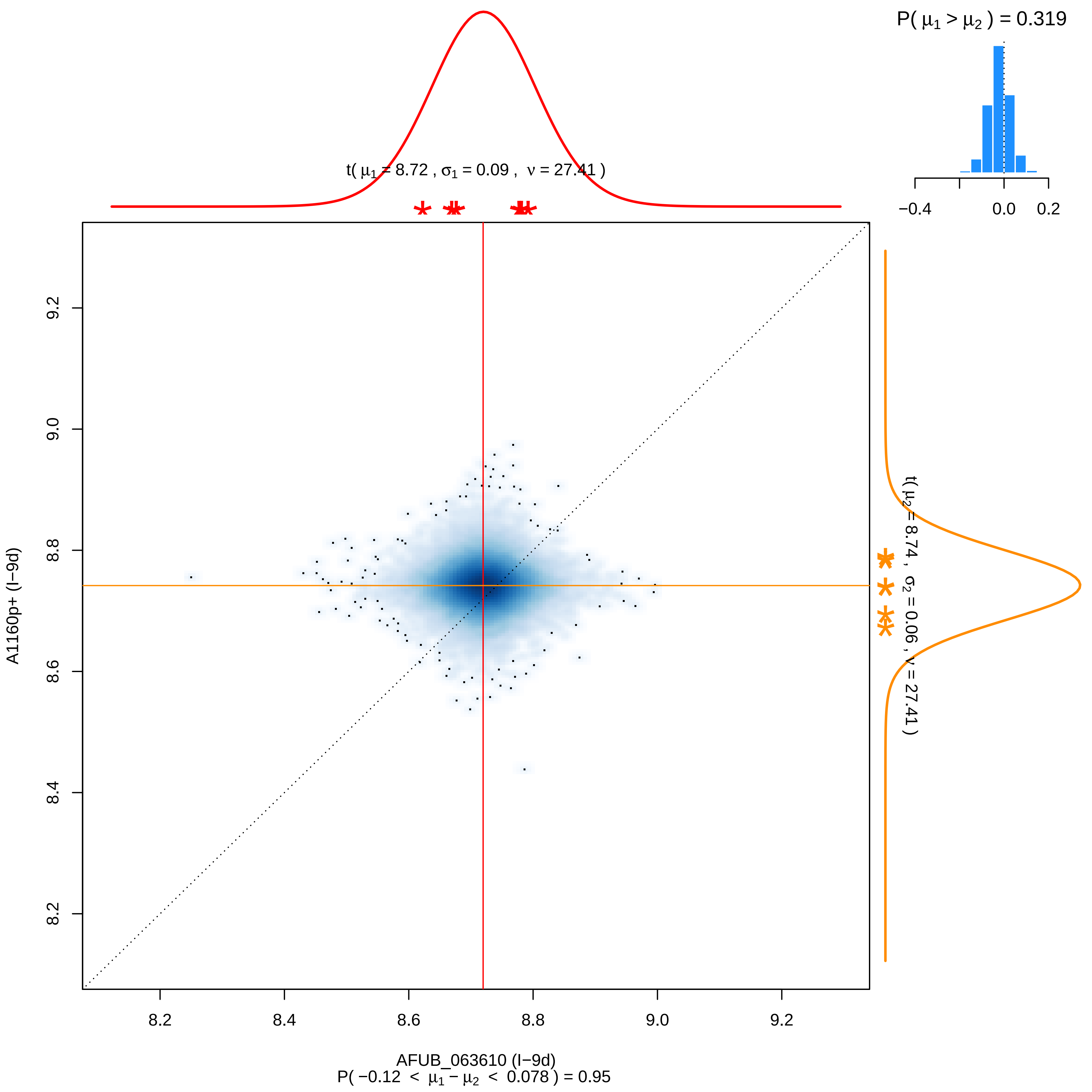

P( μ1 > μ2 ) = 0.319
−0.4
0.0
0.2
t( μ1 = 8.72 , σ1 = 0.09 , ν = 27.41 )
t( μ2 = 8.74 , σ2 = 0.06 , ν = 27.41 )
A1160p+ (I−9d)
9.2
9.0
8.8
8.6
8.4
8.2
8.2
8.4
8.6
8.8
9.0
9.2
AFUB_063610 (I−9d)
P( −0.12 < μ1 − μ2 < 0.078 ) = 0.95

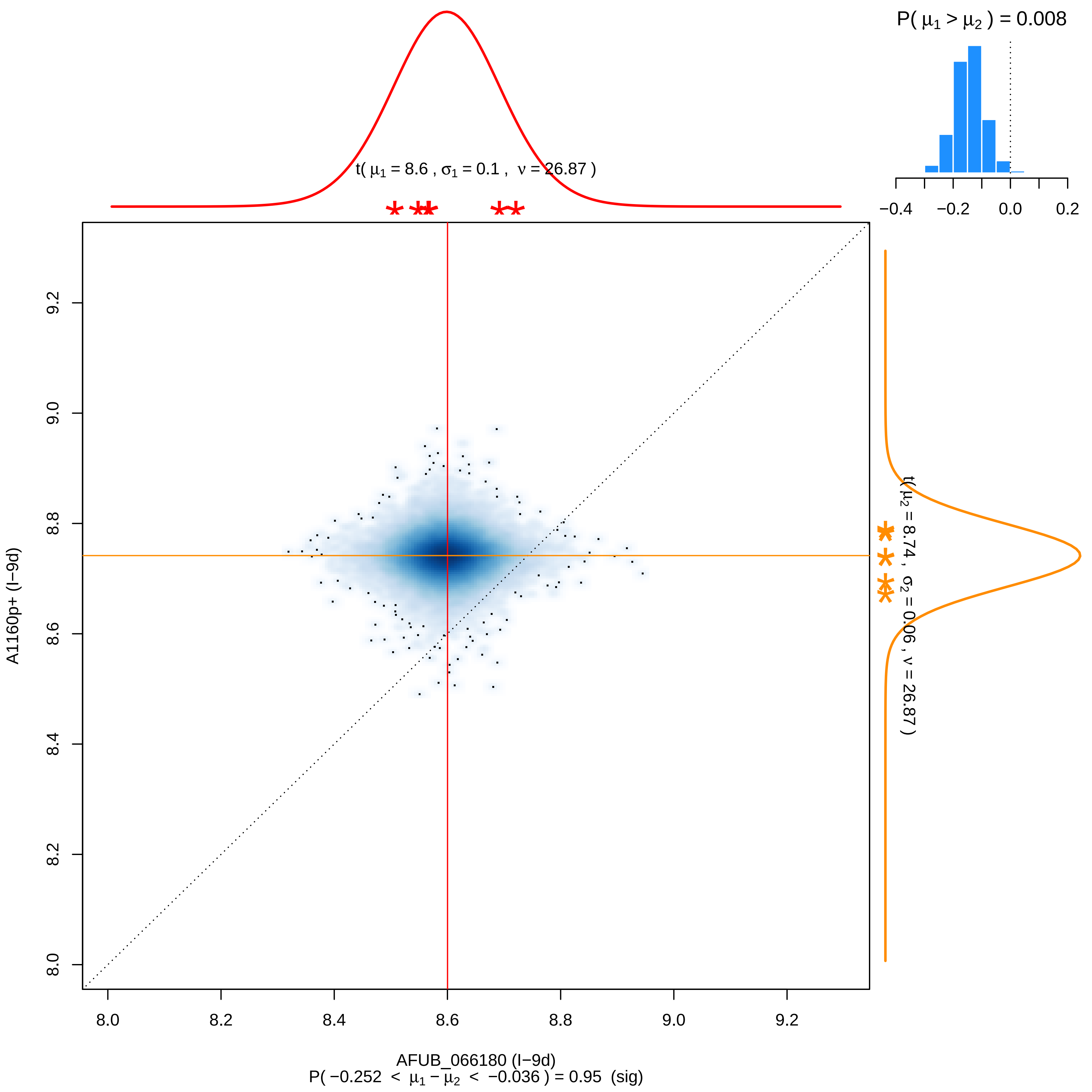

P( $\mu_1 > \mu_2$ ) = 0.008
−0.4 −0.2 0.0 0.2
t( $\mu_1$ = 8.6 , $\sigma_1$ = 0.1 , $\nu$ = 26.87 )
t( $\mu_2$ = 8.74 , $\sigma_2$ = 0.06 , $\nu$ = 26.87 )
8.0 8.2 8.4 8.6 8.8 9.0 9.2
A1160p+ (I−9d)
AFUB_066180 (I−9d)
P( −0.252 < $\mu_1 - \mu_2$ < −0.036 ) = 0.95 (sig)

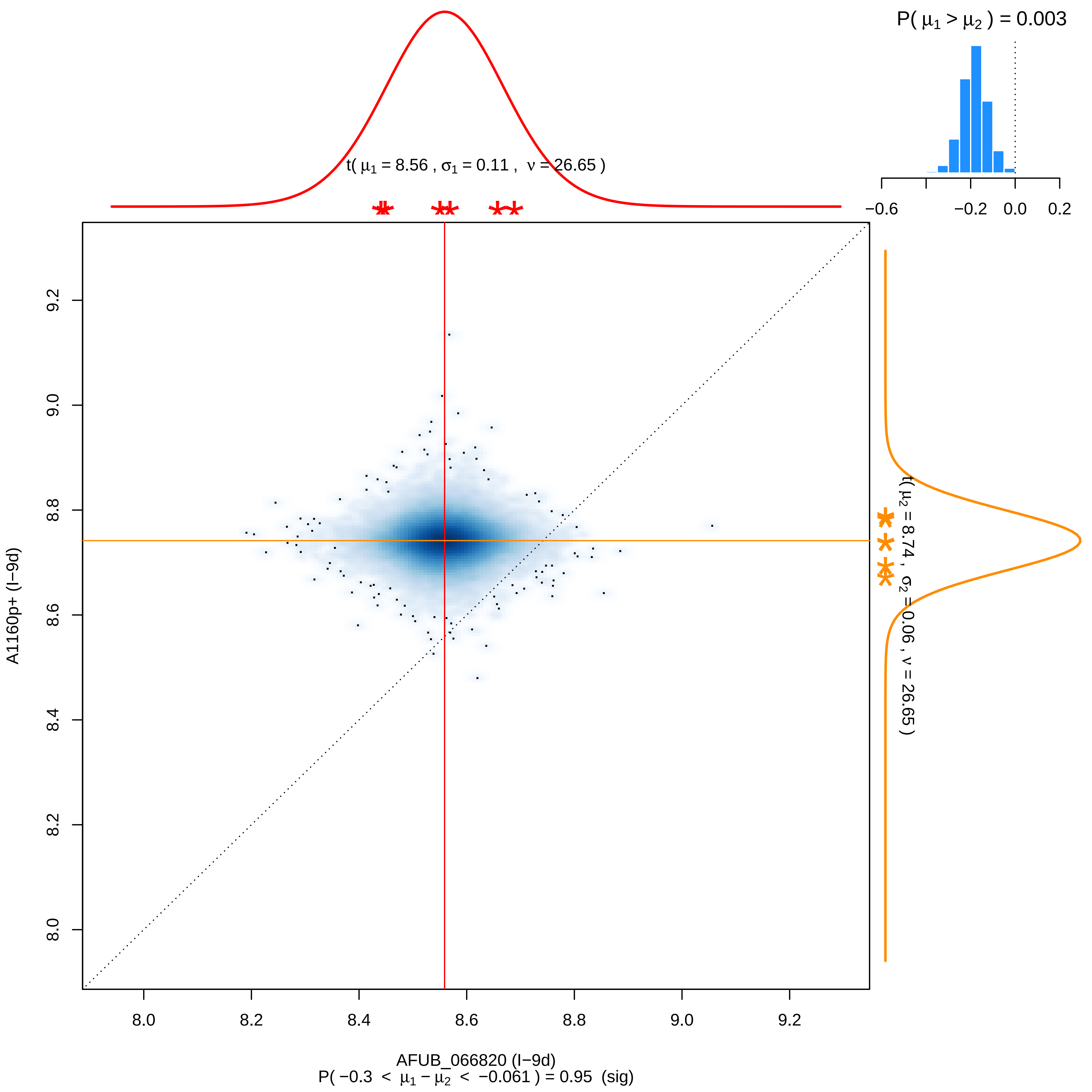

P( μ1 > μ2 ) = 0.003
−0.6
−0.2
0.0
0.2
t( μ1 = 8.56 , σ1 = 0.11 , ν = 26.65 )
t( μ2 = 8.74 , σ2 = 0.06 , ν = 26.65 )
A1160p+ (I−9d)
9.2
9.0
8.8
8.6
8.4
8.2
8.0
8.0
8.2
8.4
8.6
8.8
9.0
9.2
AFUB_066820 (I−9d)
P( −0.3 < μ1 − μ2 < −0.061 ) = 0.95 (sig)

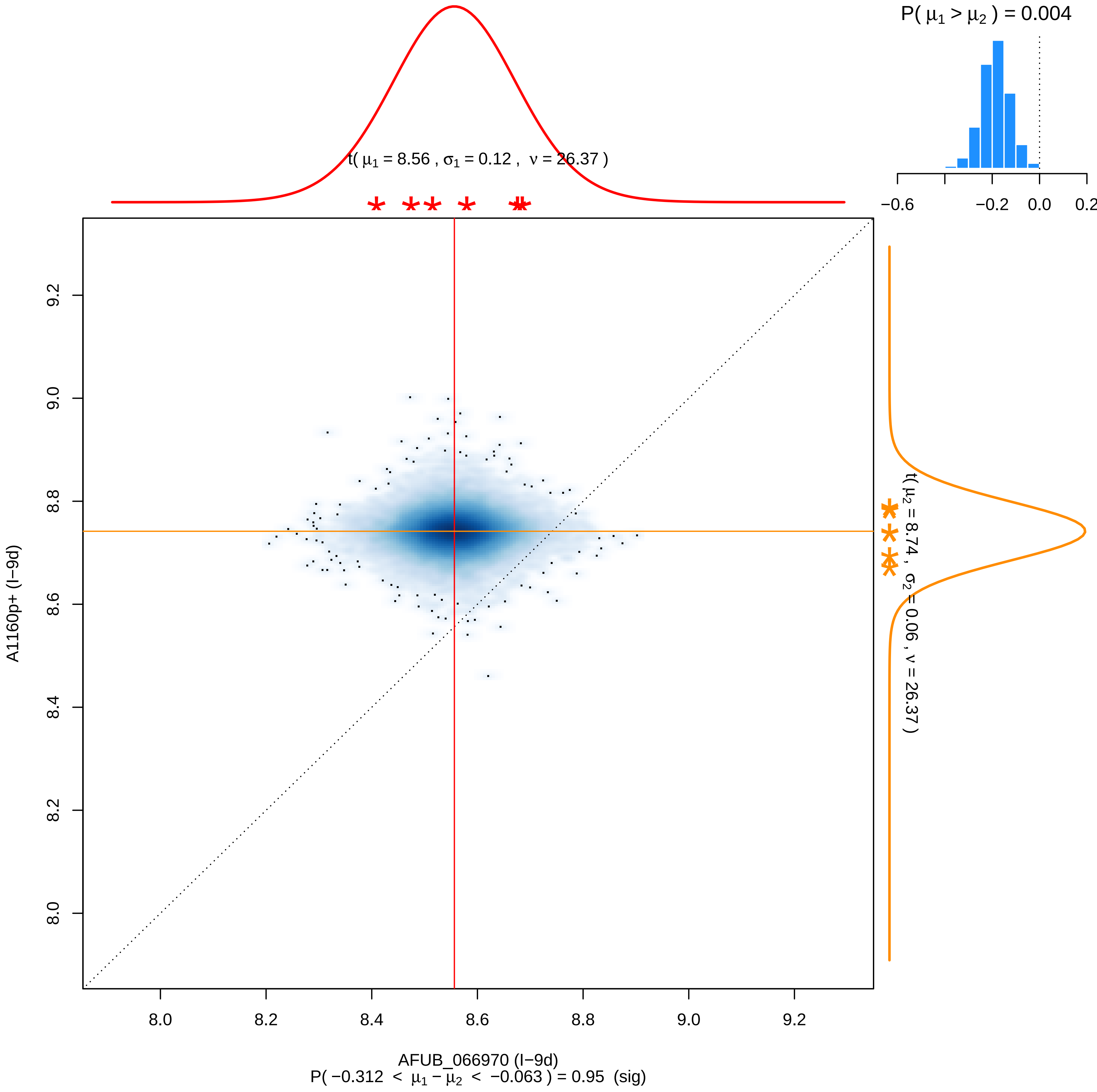

P( μ1 > μ2 ) = 0.004
−0.6
−0.2
0.0
0.2
t( μ1 = 8.56 , σ1 = 0.12 , ν = 26.37 )
t( μ2 = 8.74 , σ2 = 0.06 , ν = 26.37 )
9.2
9.0
8.8
8.6
8.4
8.2
8.0
A1160p+ (I−9d)
8.0
8.2
8.4
8.6
8.8
9.0
9.2
AFUB_066970 (I−9d)
P( −0.312 < μ1 − μ2 < −0.063 ) = 0.95 (sig)

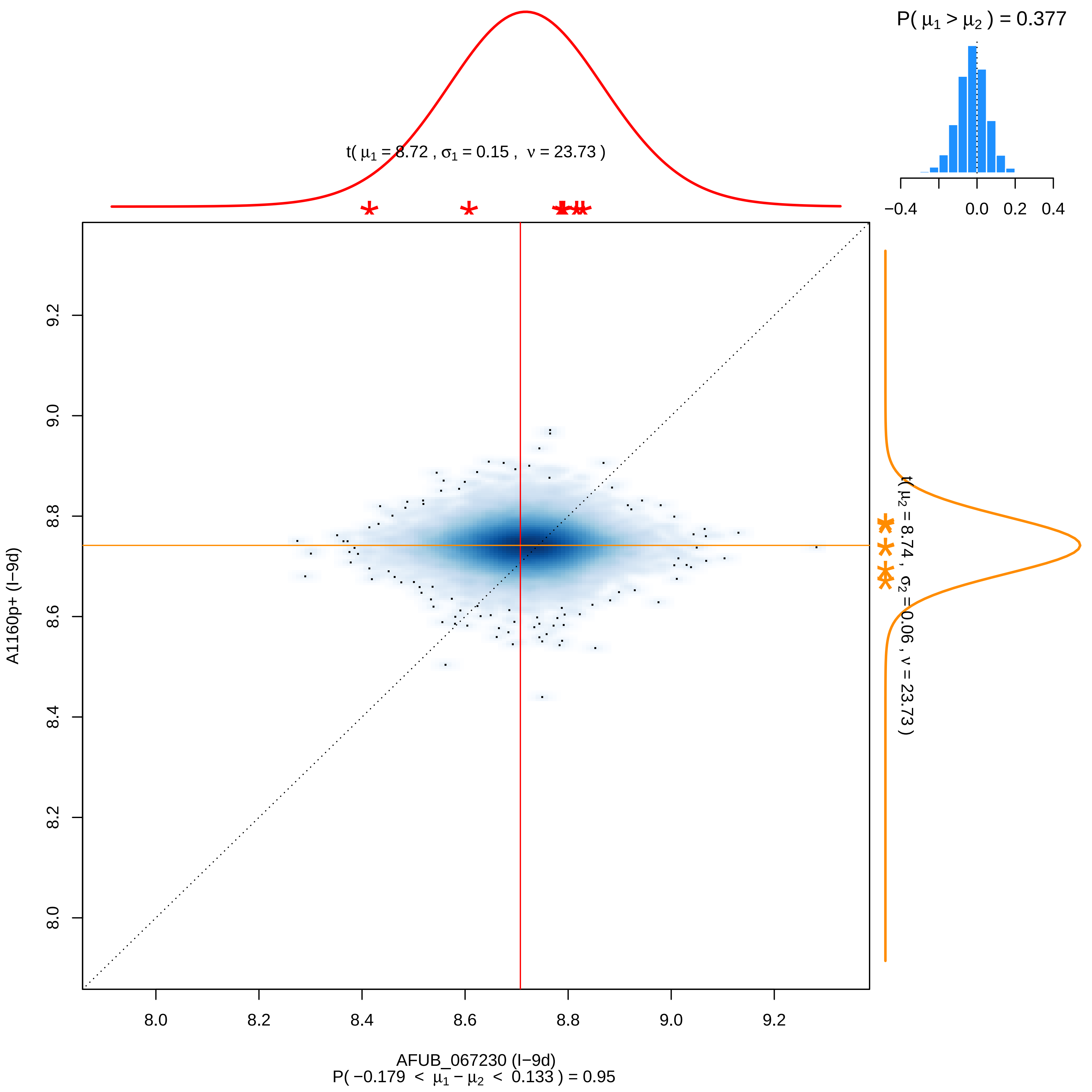

P( μ1 > μ2 ) = 0.377
−0.4 0.0 0.2 0.4
t( μ1 = 8.72 , σ1 = 0.15 , ν = 23.73 )
t( μ2 = 8.74 , σ2 = 0.06 , ν = 23.73 )
A1160p+ (I−9d)
8.0 8.2 8.4 8.6 8.8 9.0 9.2
AFUB_067230 (I−9d)
P( −0.179 < μ1 − μ2 < 0.133 ) = 0.95

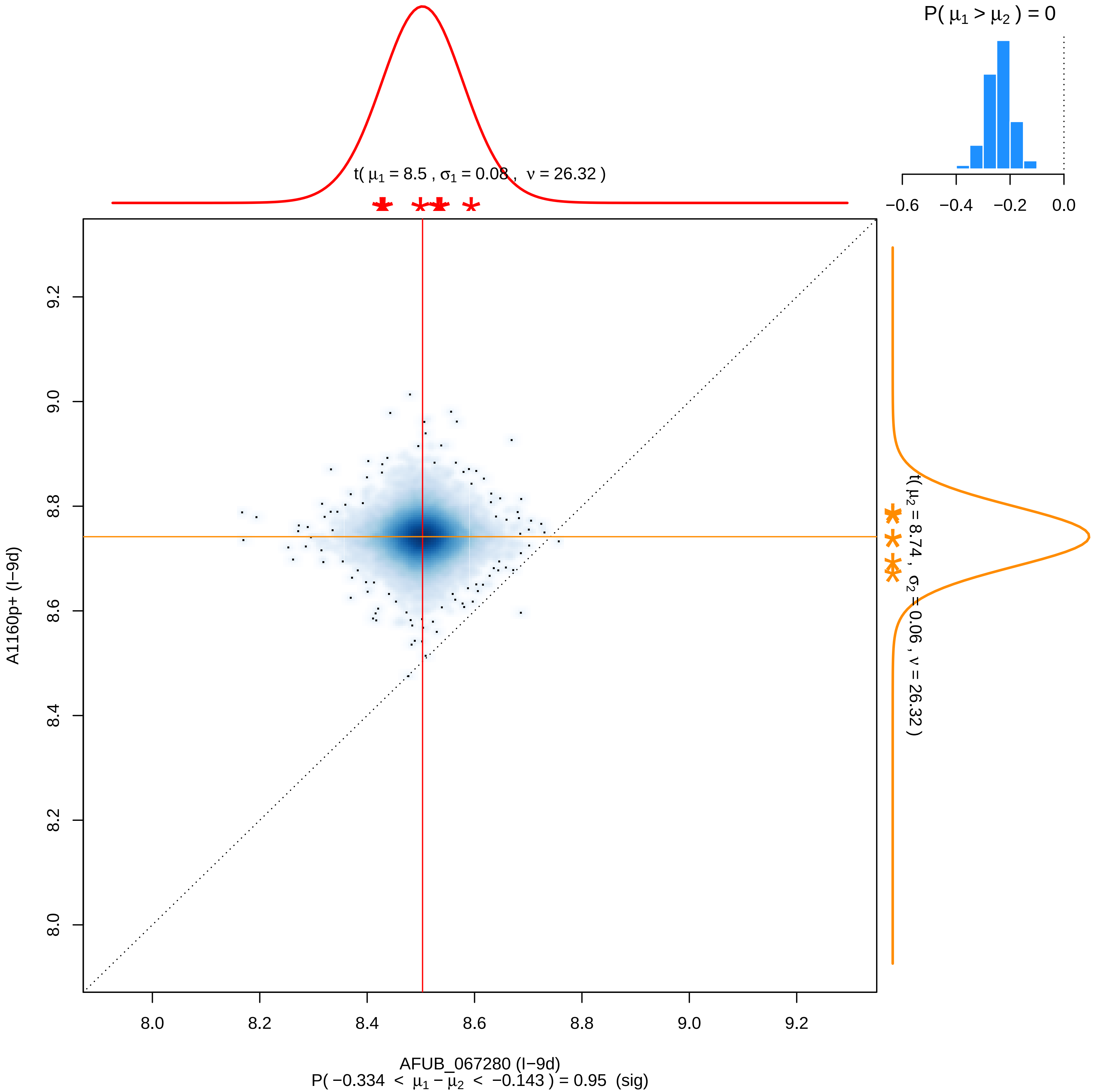

P( μ1 > μ2 ) = 0
−0.6 −0.4 −0.2 0.0
t( μ1 = 8.5 , σ1 = 0.08 , ν = 26.32 )
t( μ2 = 8.74 , σ2 = 0.06 , ν = 26.32 )
A1160p+ (I−9d)
9.2 9.0 8.8 8.6 8.4 8.2 8.0
8.0 8.2 8.4 8.6 8.8 9.0 9.2
AFUB_067280 (I−9d)
P( −0.334 < μ1 − μ2 < −0.143 ) = 0.95 (sig)

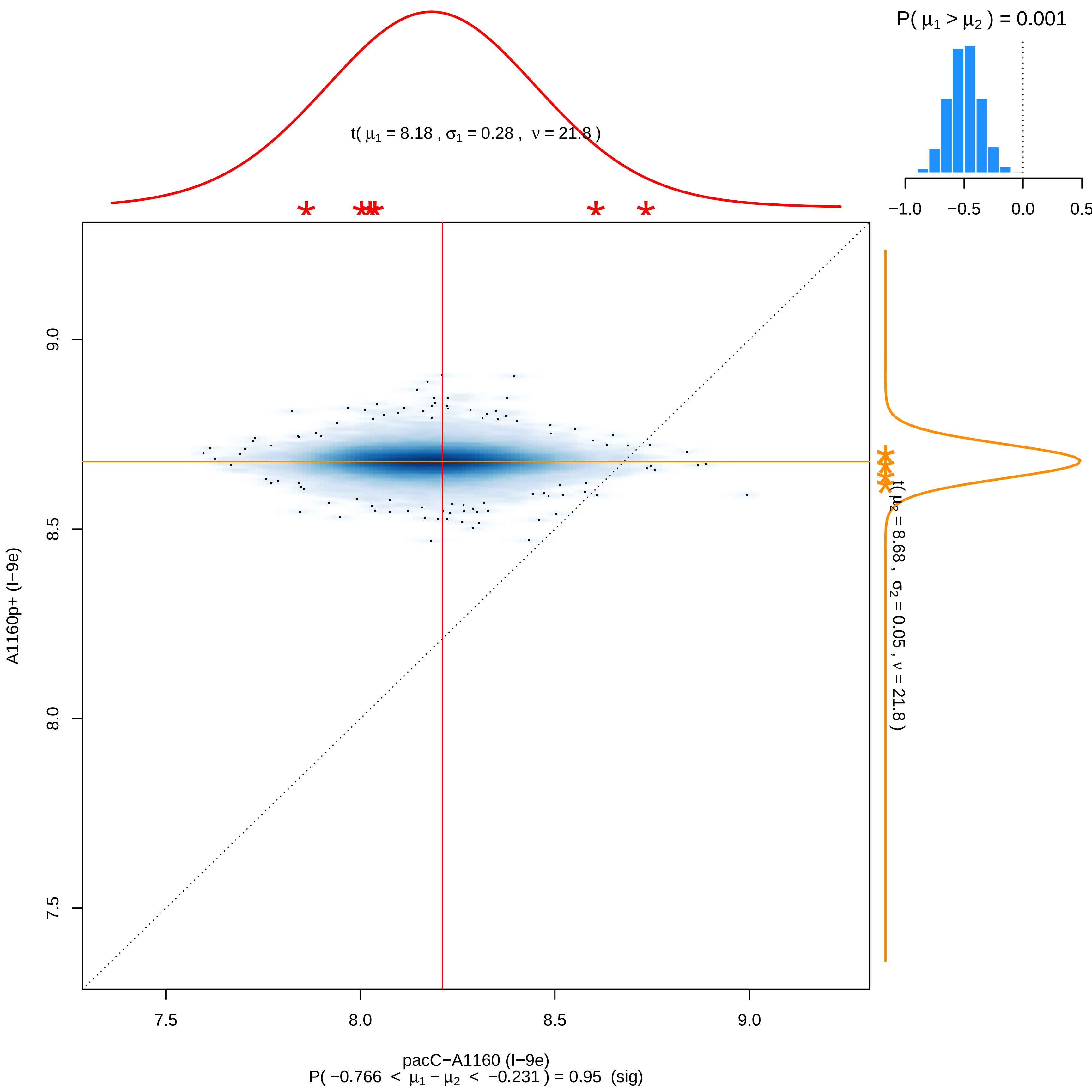

P( μ1 > μ2 ) = 0.001
−1.0
−0.5
0.0
0.5
t( μ1 = 8.18 , σ1 = 0.28 , ν = 21.8 )
9.0
8.5
8.0
7.5
A1160p+ (I−9e)
t( μ2 = 8.68 , σ2 = 0.05 , ν = 21.8 )
pacC−A1160 (I−9e)
P( −0.766 < μ1 − μ2 < −0.231 ) = 0.95 (sig)

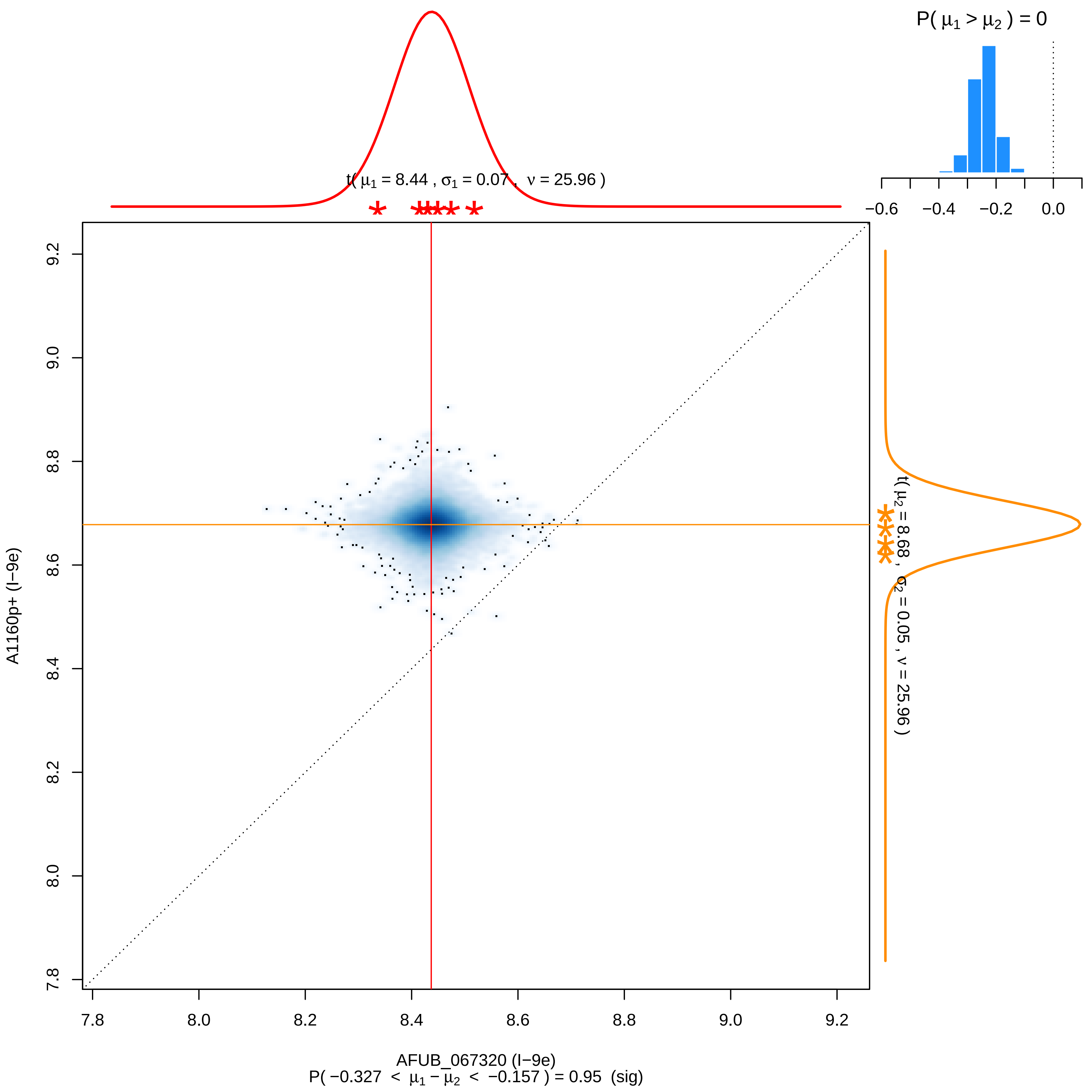

P( μ1 > μ2 ) = 0
−0.6
−0.4
−0.2
0.0
t( μ1 = 8.44 , σ1 = 0.07 , ν = 25.96 )
t( μ2 = 8.68 , σ2 = 0.05 , ν = 25.96 )
A1160p+ (I−9e)
AFUB_067320 (I−9e)
7.8
8.0
8.2
8.4
8.6
8.8
9.0
9.2
P( −0.327 < μ1 − μ2 < −0.157 ) = 0.95 (sig)

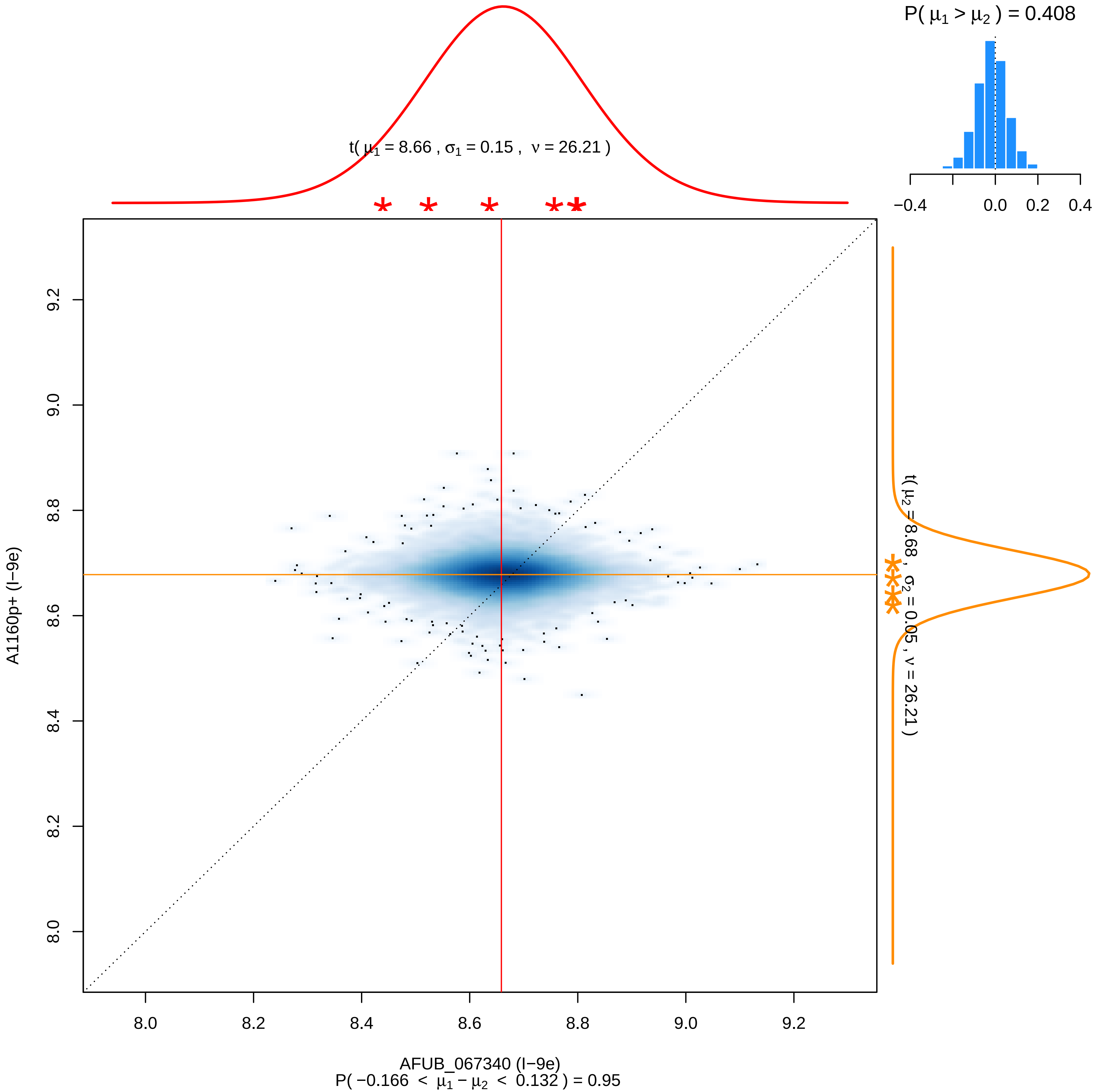

P( μ1 > μ2 ) = 0.408
−0.4
0.0
0.2
0.4
t( μ1 = 8.66 , σ1 = 0.15 , ν = 26.21 )
t( μ2 = 8.68 , σ2 = 0.05 , ν = 26.21 )
8.0
8.2
8.4
8.6
8.8
9.0
9.2
A1160p+ (I−9e)
AFUB_067340 (I−9e)
P( −0.166 < μ1 − μ2 < 0.132 ) = 0.95

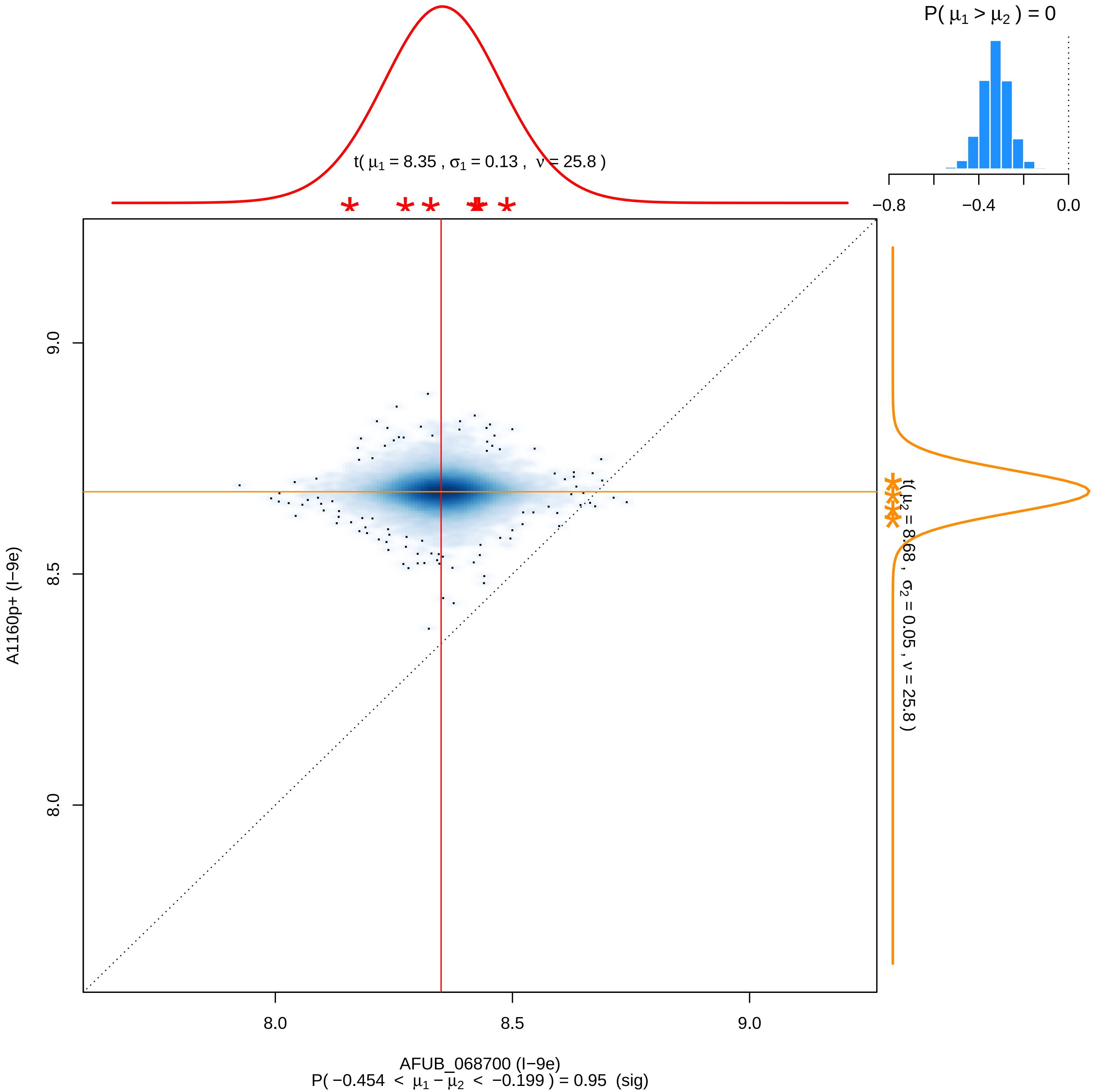

P( μ1 > μ2 ) = 0
−0.8
−0.4
0.0
t( μ1 = 8.35 , σ1 = 0.13 , ν = 25.8 )
t( μ2 = 8.68 , σ2 = 0.05 , ν = 25.8 )
9.0
8.5
8.0
A1160p+ (I−9e)
8.0
8.5
9.0
AFUB_068700 (I−9e)
P( −0.454 < μ1 − μ2 < −0.199 ) = 0.95 (sig)

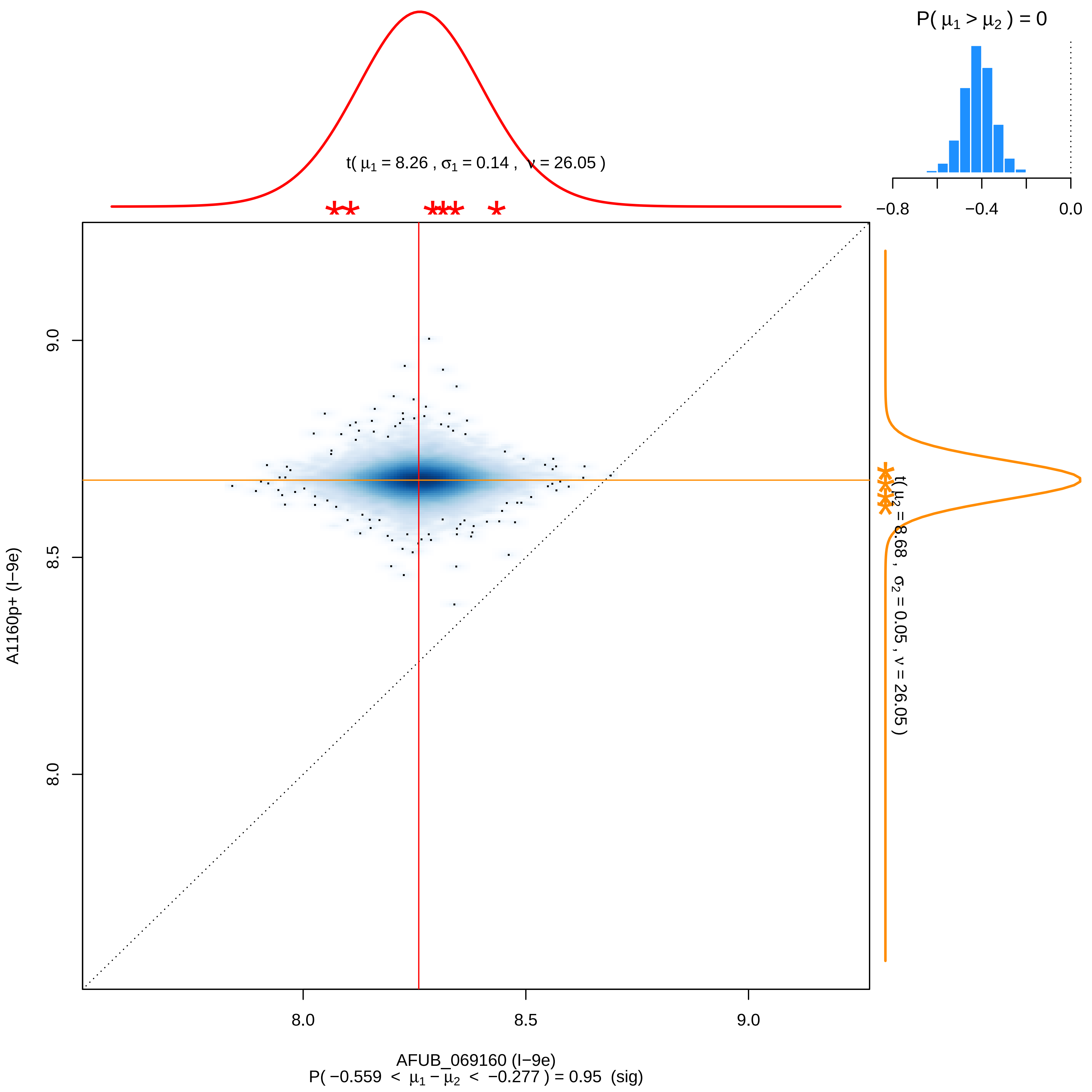

P( μ1 > μ2 ) = 0
−0.8
−0.4
0.0
t( μ1 = 8.26 , σ1 = 0.14 , ν = 26.05 )
t( μ2 = 8.68 , σ2 = 0.05 , ν = 26.05 )
A1160p+ (I−9e)
9.0
8.5
8.0
8.0
8.5
9.0
AFUB_069160 (I−9e)
P( −0.559 < μ1 − μ2 < −0.277 ) = 0.95 (sig)

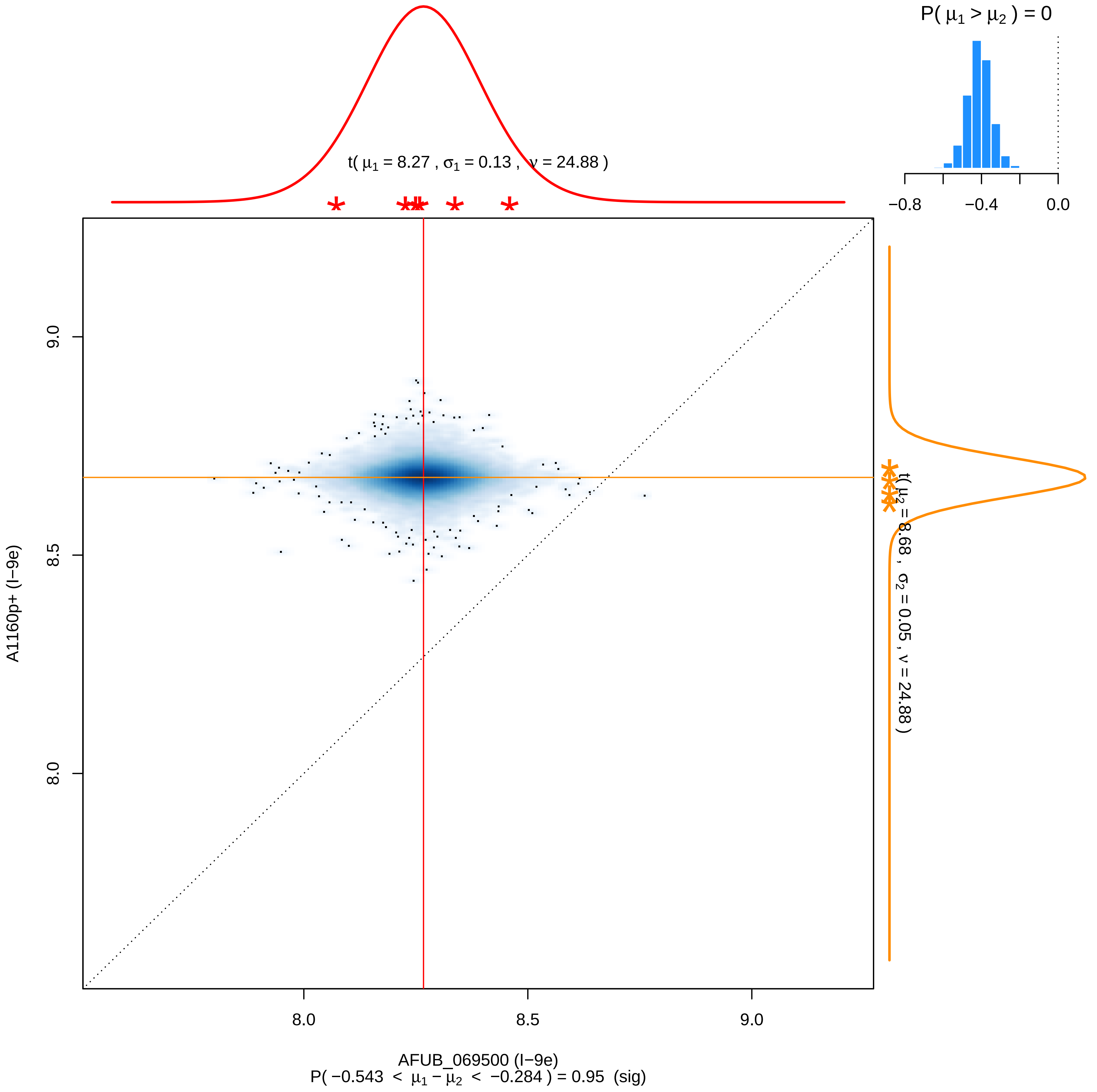

P( μ1 > μ2 ) = 0
−0.8
−0.4
0.0
t( μ1 = 8.27 , σ1 = 0.13 , ν = 24.88 )
t( μ2 = 8.68 , σ2 = 0.05 , ν = 24.88 )
9.0
8.5
8.0
A1160p+ (I−9e)
8.0
8.5
9.0
AFUB_069500 (I−9e)
P( −0.543 < μ1 − μ2 < −0.284 ) = 0.95 (sig)

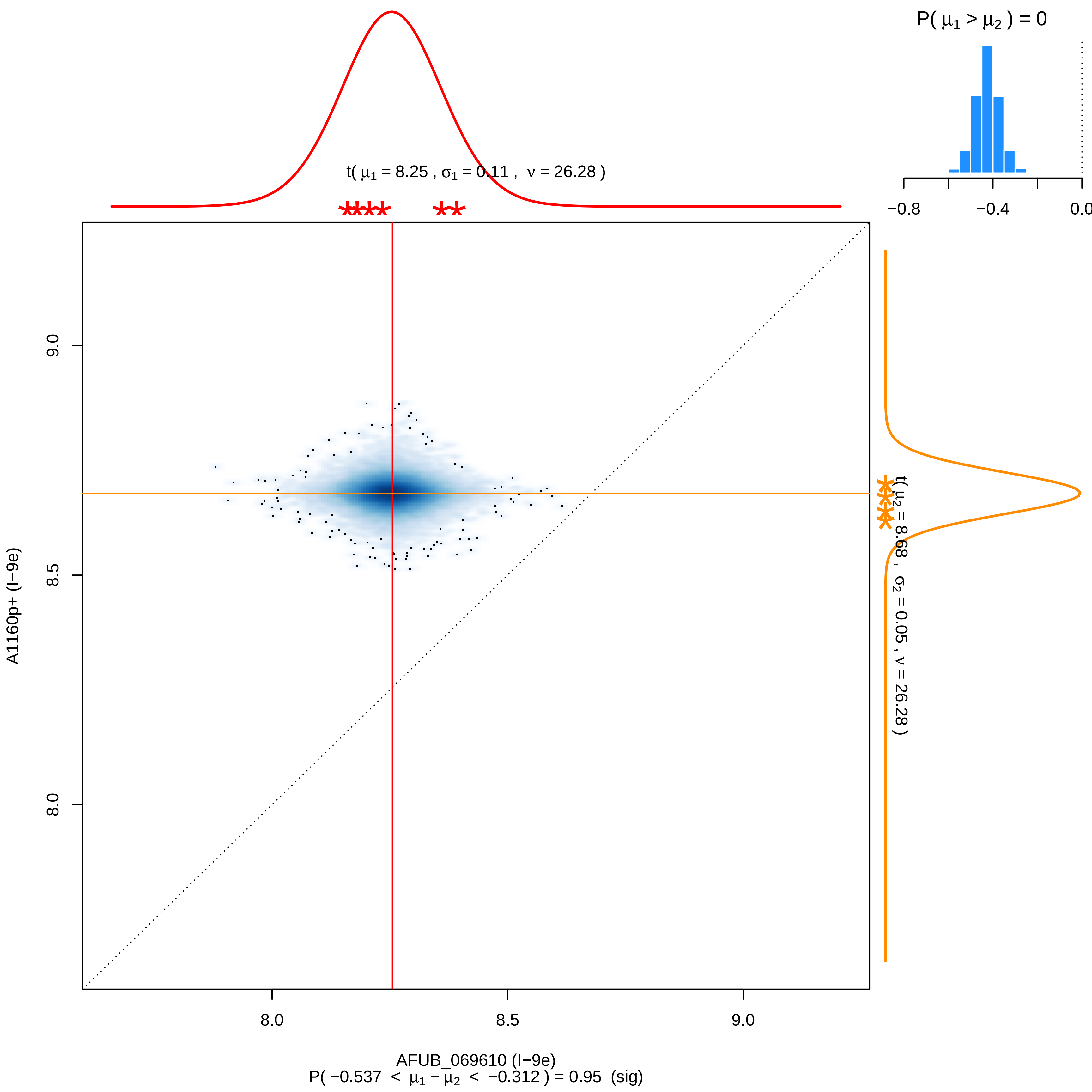

P( μ1 > μ2 ) = 0
−0.8
−0.4
0.0
t( μ1 = 8.25 , σ1 = 0.11 , ν = 26.28 )
t( μ2 = 8.68 , σ2 = 0.05 , ν = 26.28 )
9.0
8.5
8.0
A1160p+ (I−9e)
AFUB_069610 (I−9e)
P( −0.537 < μ1 − μ2 < −0.312 ) = 0.95 (sig)

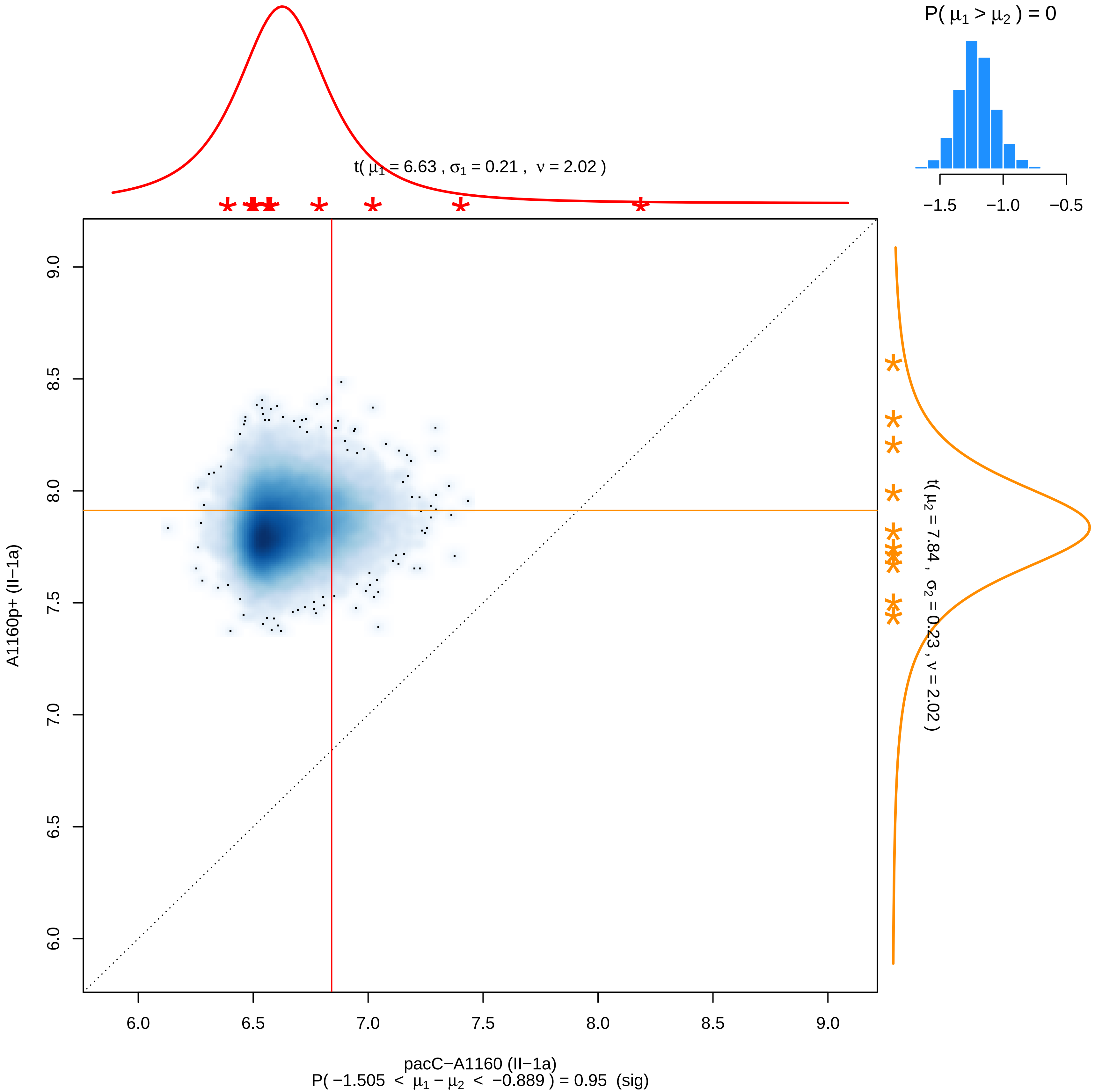

P( μ1 > μ2 ) = 0
−1.5
−1.0
−0.5
t( μ1 = 6.63 , σ1 = 0.21 , ν = 2.02 )
t( μ2 = 7.84 , σ2 = 0.23 , ν = 2.02 )
A1160p+ (II−1a)
6.0
6.5
7.0
7.5
8.0
8.5
9.0
pacC−A1160 (II−1a)
P( −1.505 < μ1 − μ2 < −0.889 ) = 0.95 (sig)

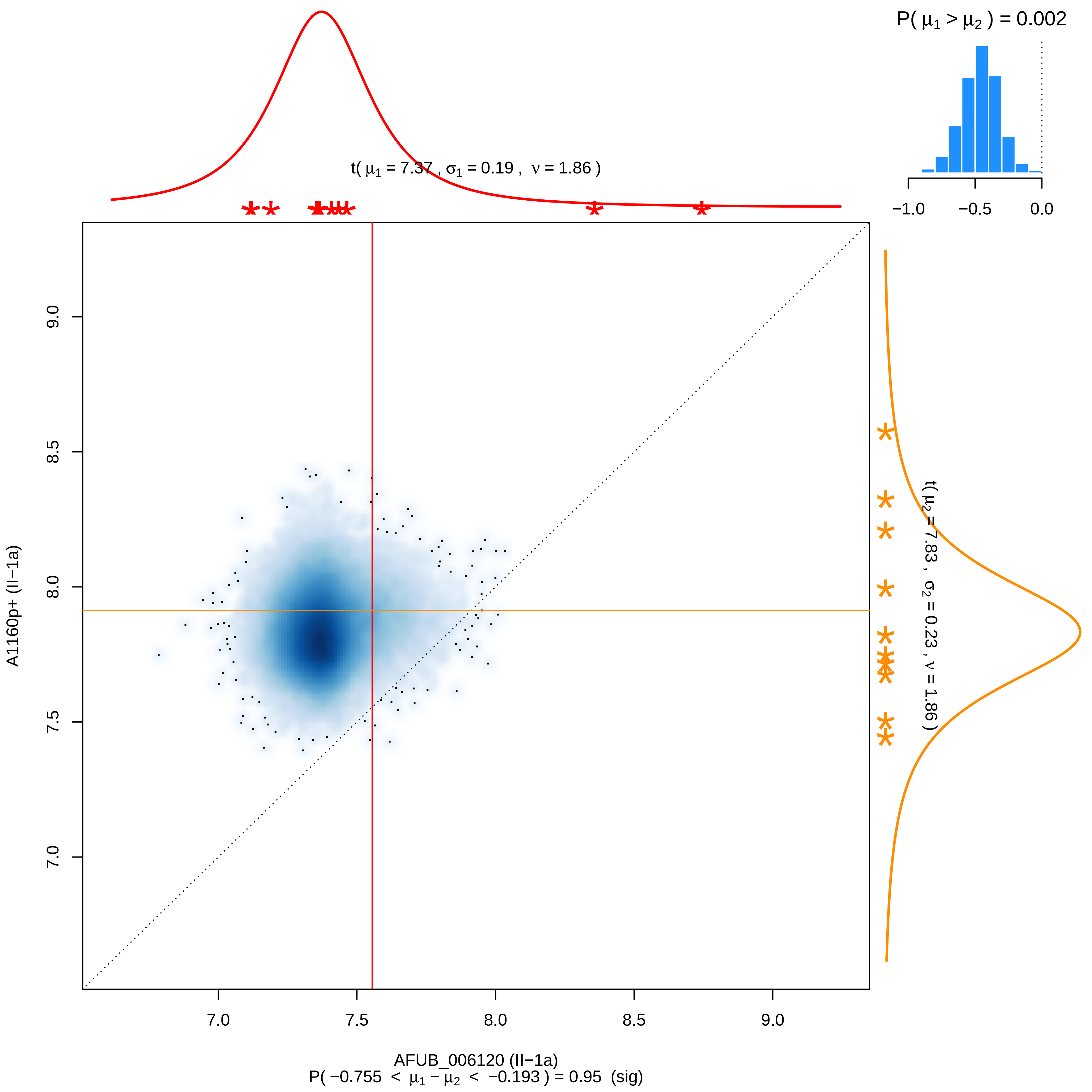

P( μ1 > μ2 ) = 0.002
−1.0
−0.5
0.0
t( μ1 = 7.37 , σ1 = 0.19 , ν = 1.86 )
t( μ2 = 7.83 , σ2 = 0.23 , ν = 1.86 )
9.0
8.5
8.0
7.5
7.0
A1160p+ (II−1a)
7.0
7.5
8.0
8.5
9.0
AFUB_006120 (II−1a)
P( −0.755 < μ1 − μ2 < −0.193 ) = 0.95 (sig)

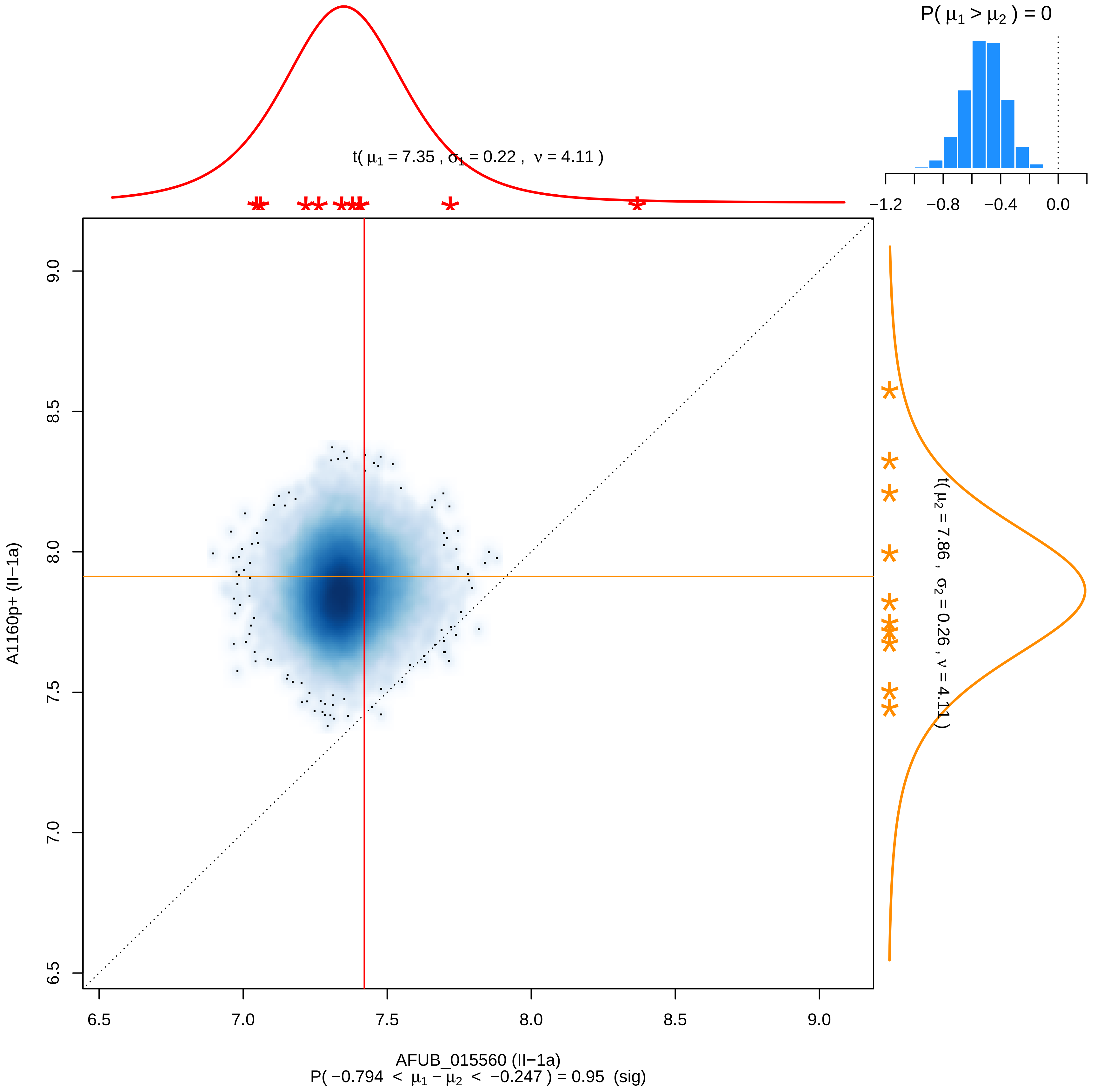
P( μ1 > μ2 ) = 0
−1.2
−0.8
−0.4
0.0
t( μ1 = 7.35 , σ1 = 0.22 , ν = 4.11 )
t( μ2 = 7.86 , σ2 = 0.26 , ν = 4.11 )
9.0
8.5
8.0
7.5
7.0
6.5
A1160p+ (II−1a)
6.5
7.0
7.5
8.0
8.5
9.0
AFUB_015560 (II−1a)
P( −0.794 < μ1 − μ2 < −0.247 ) = 0.95 (sig)

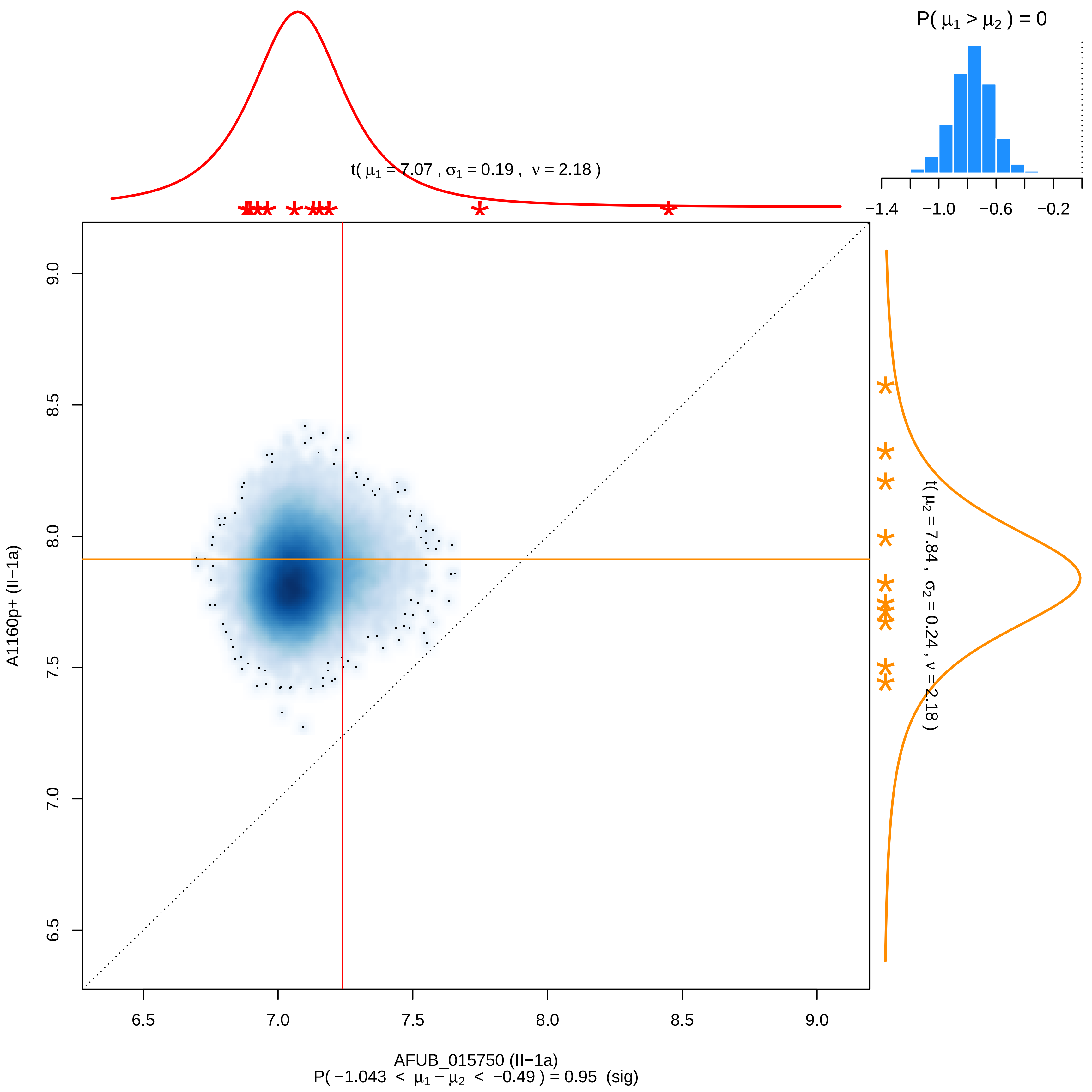

P( μ1 > μ2 ) = 0
−1.4
−1.0
−0.6
−0.2
t( μ1 = 7.07 , σ1 = 0.19 , ν = 2.18 )
t( μ2 = 7.84 , σ2 = 0.24 , ν = 2.18 )
A1160p+ (II−1a)
9.0
8.5
8.0
7.5
7.0
6.5
6.5
7.0
7.5
8.0
8.5
9.0
AFUB_015750 (II−1a)
P( −1.043 < μ1 − μ2 < −0.49 ) = 0.95 (sig)

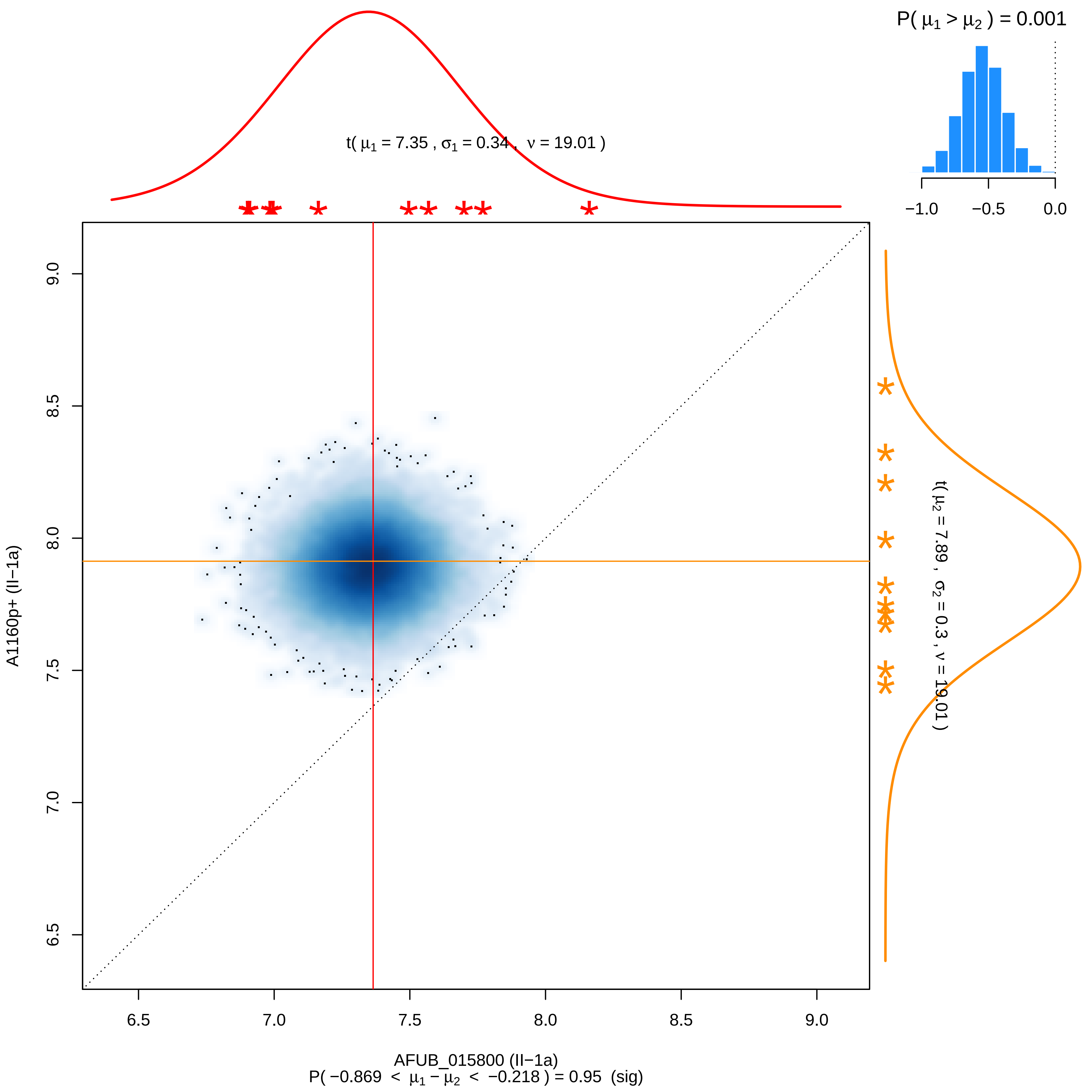

P( μ1 > μ2 ) = 0.001
−1.0
−0.5
0.0
t( μ1 = 7.35 , σ1 = 0.34 , ν = 19.01 )
t( μ2 = 7.89 , σ2 = 0.3 , ν = 19.01 )
A1160p+ (II−1a)
AFUB_015800 (II−1a)
P( −0.869 < μ1 − μ2 < −0.218 ) = 0.95 (sig)
6.5
7.0
7.5
8.0
8.5
9.0

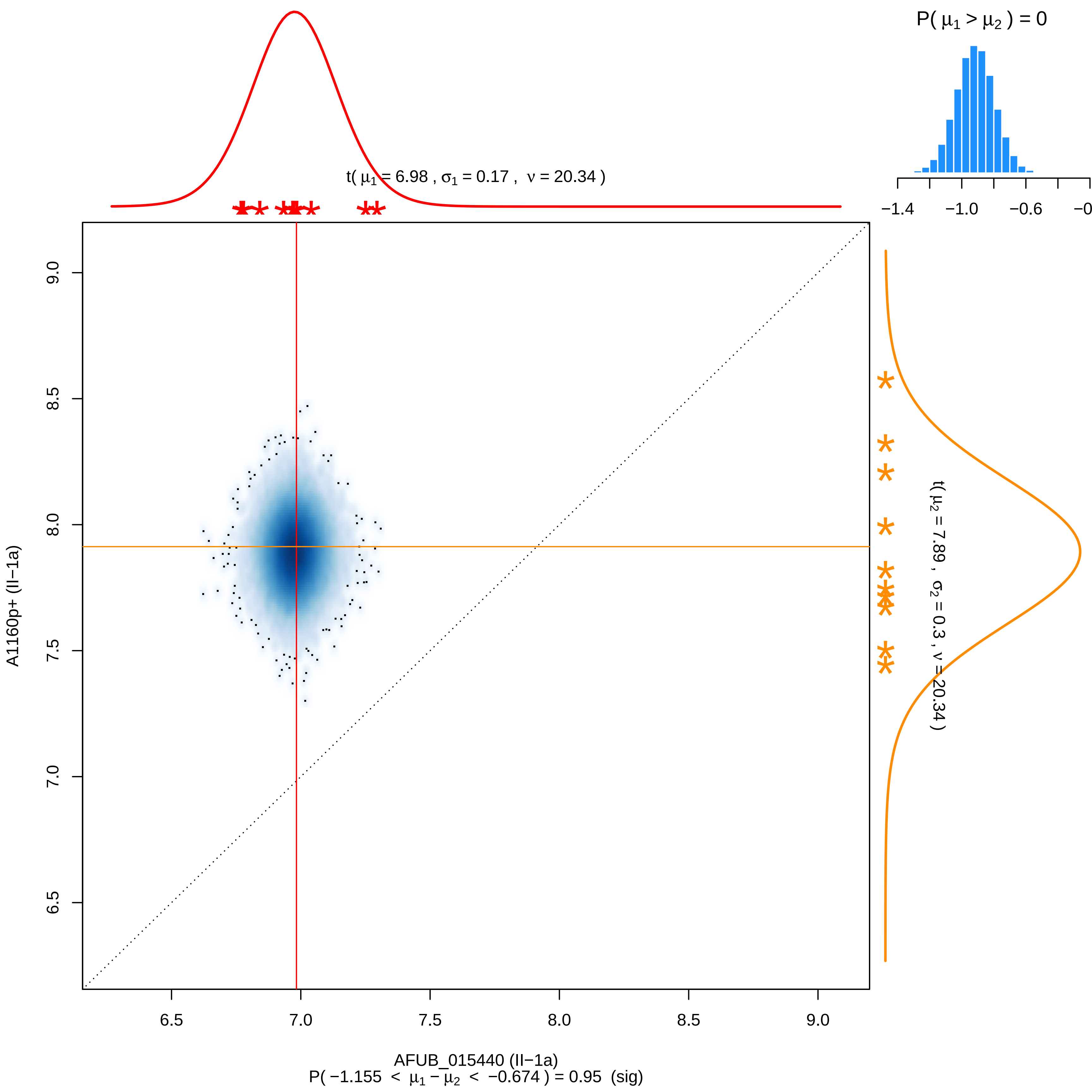

P( μ1 > μ2 ) = 0
−1.4
−1.0
−0.6
t( μ1 = 6.98 , σ1 = 0.17 , ν = 20.34 )
t( μ2 = 7.89 , σ2 = 0.3 , ν = 20.34 )
A1160p+ (II−1a)
AFUB_015440 (II−1a)
6.5
7.0
7.5
8.0
8.5
9.0
P( −1.155 < μ1 − μ2 < −0.674 ) = 0.95 (sig)

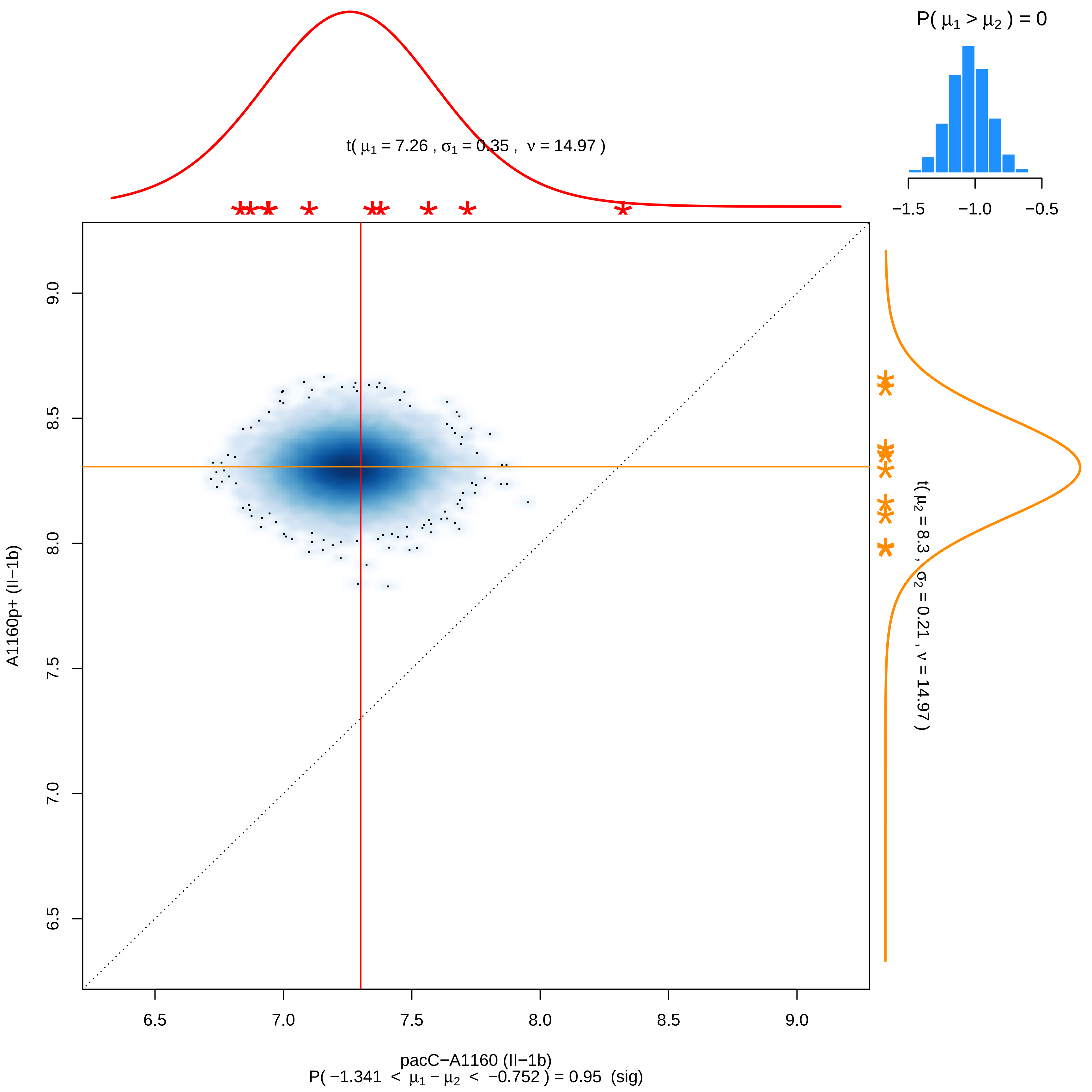

P( μ1 > μ2 ) = 0
−1.5
−1.0
−0.5
t( μ1 = 7.26 , σ1 = 0.35 , ν = 14.97 )
t( μ2 = 8.3 , σ2 = 0.21 , ν = 14.97 )
A1160p+ (II−1b)
9.0
8.5
8.0
7.5
7.0
6.5
6.5
7.0
7.5
8.0
8.5
9.0
pacC−A1160 (II−1b)
P( −1.341 < μ1 − μ2 < −0.752 ) = 0.95 (sig)

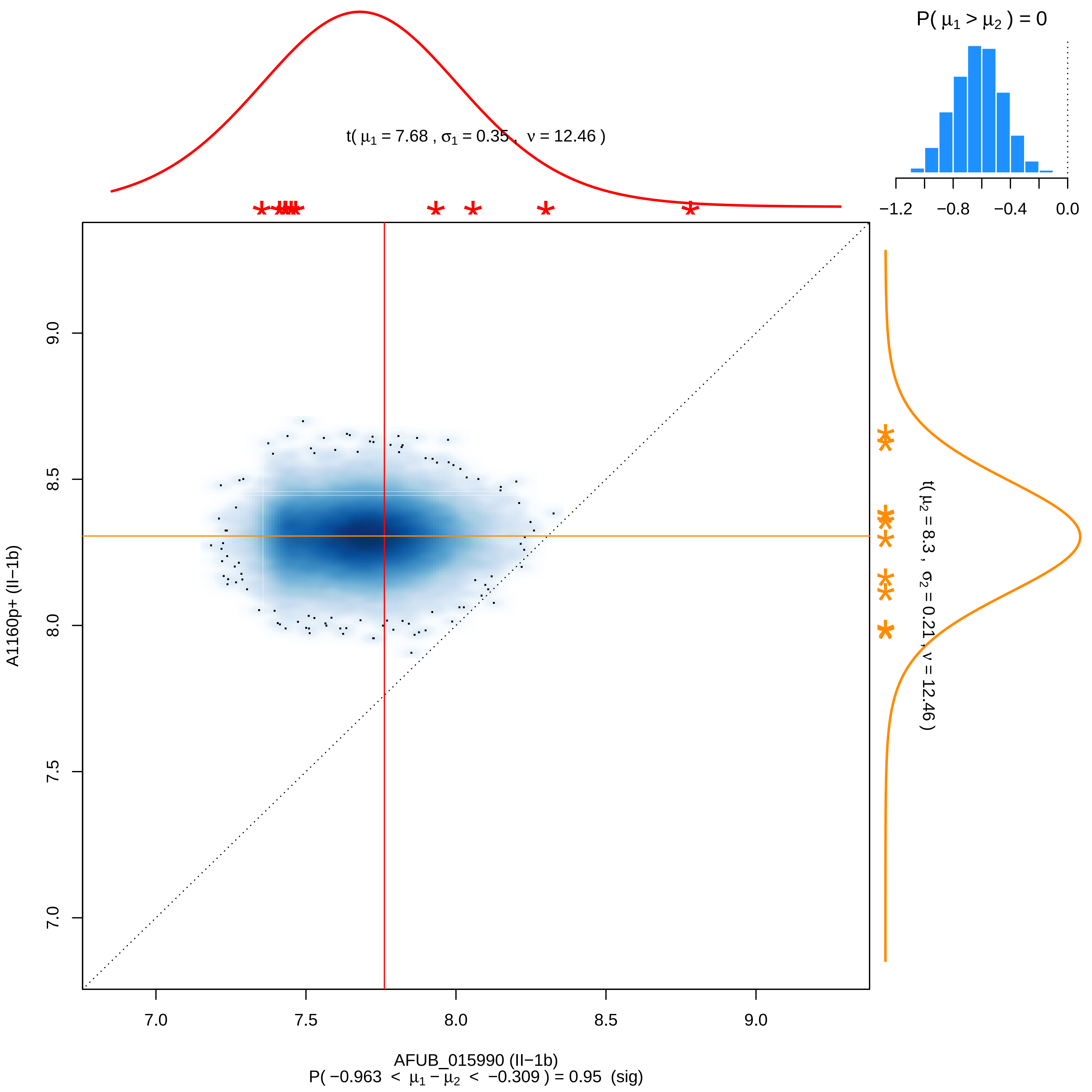

P( μ1 > μ2 ) = 0
−1.2
−0.8
−0.4
0.0
t( μ1 = 7.68 , σ1 = 0.35 , ν = 12.46 )
t( μ2 = 8.3 , σ2 = 0.21 , ν = 12.46 )
9.0
8.5
8.0
7.5
7.0
A1160p+ (II−1b)
7.0
7.5
8.0
8.5
9.0
AFUB_015990 (II−1b)
P( −0.963 < μ1 − μ2 < −0.309 ) = 0.95 (sig)

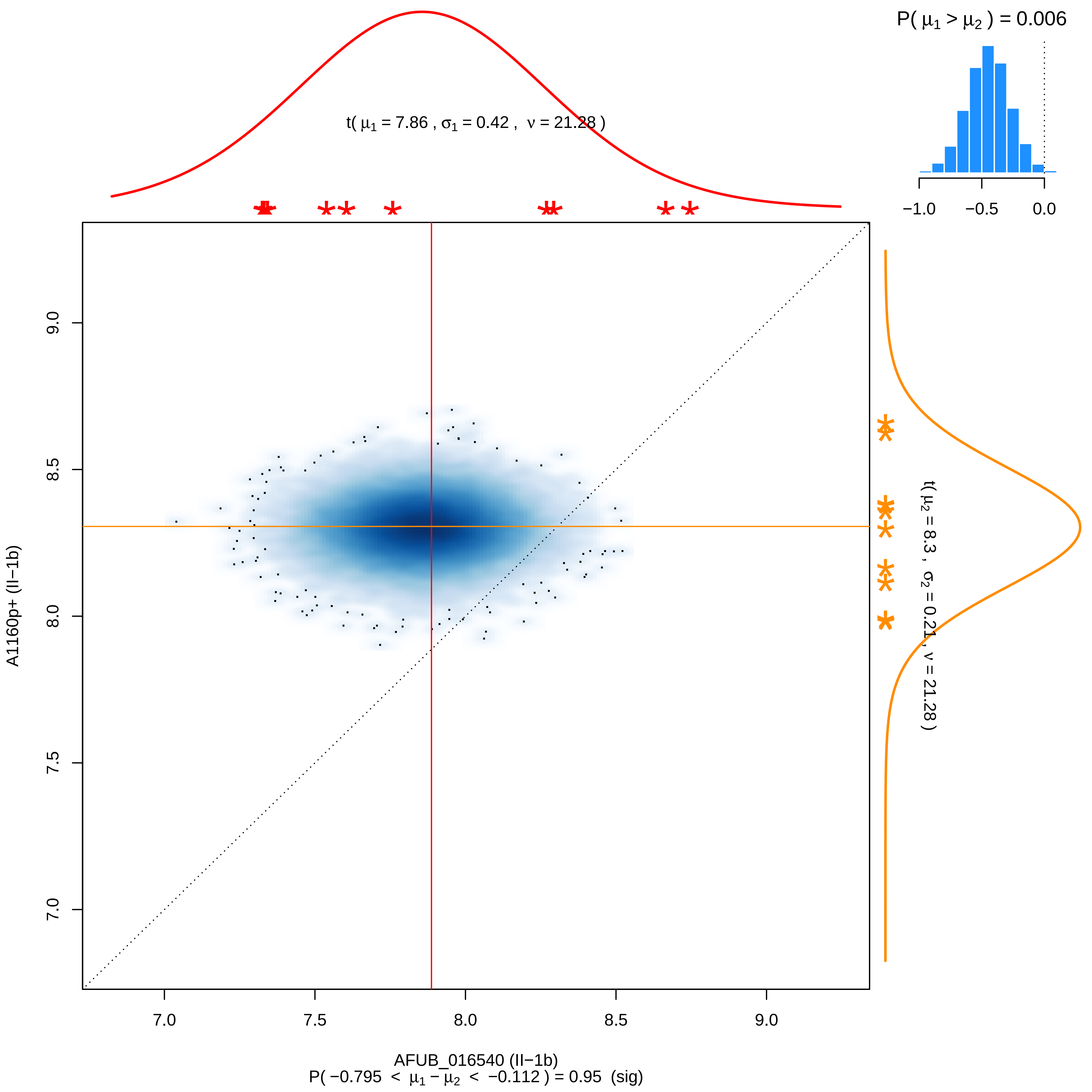

P( μ1 > μ2 ) = 0.006
−1.0
−0.5
0.0
t( μ1 = 7.86 , σ1 = 0.42 , ν = 21.28 )
t( μ2 = 8.3 , σ2 = 0.21 , ν = 21.28 )
9.0
8.5
8.0
7.5
7.0
A1160p+ (II−1b)
7.0
7.5
8.0
8.5
9.0
AFUB_016540 (II−1b)
P( −0.795 < μ1 − μ2 < −0.112 ) = 0.95 (sig)

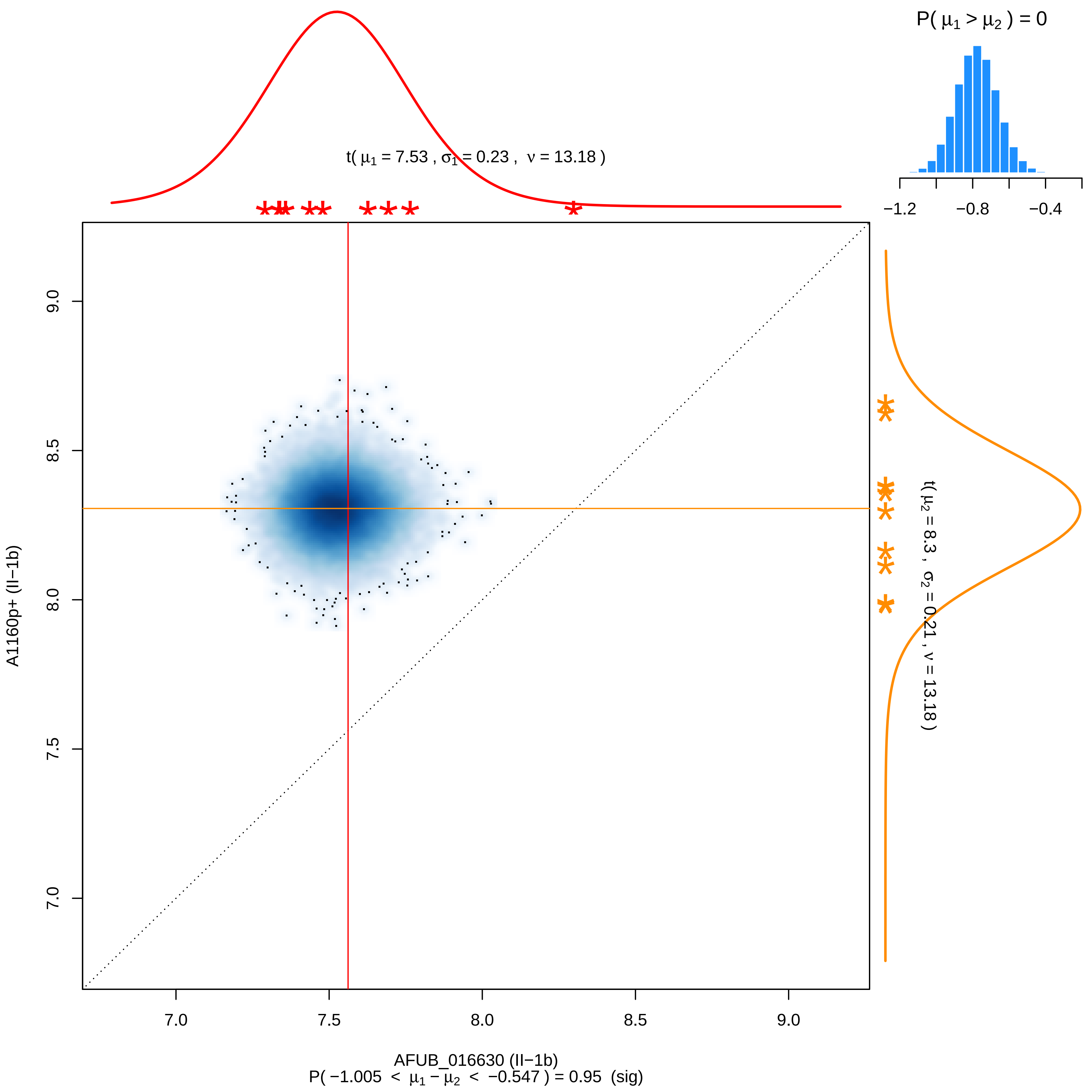

P( μ1 > μ2 ) = 0
−1.2
−0.8
−0.4
t( μ1 = 7.53 , σ1 = 0.23 , ν = 13.18 )
t( μ2 = 8.3 , σ2 = 0.21 , ν = 13.18 )
A1160p+ (II−1b)
9.0
8.5
8.0
7.5
7.0
7.0
7.5
8.0
8.5
9.0
AFUB_016630 (II−1b)
P( −1.005 < μ1 − μ2 < −0.547 ) = 0.95 (sig)

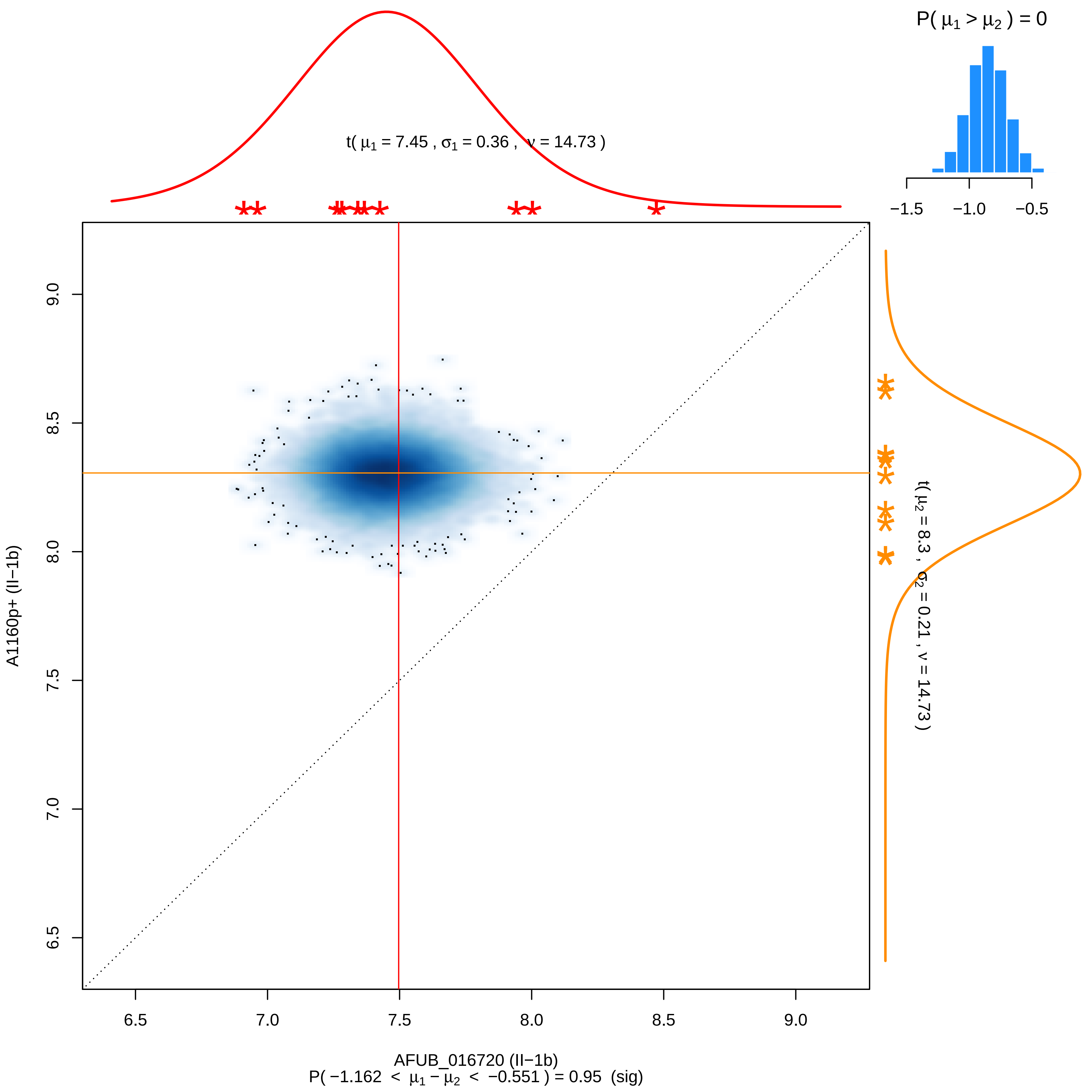

P( μ1 > μ2 ) = 0
−1.5
−1.0
−0.5
t( μ1 = 7.45 , σ1 = 0.36 , ν = 14.73 )
t( μ2 = 8.3 , σ2 = 0.21 , ν = 14.73 )
A1160p+ (II−1b)
6.5
7.0
7.5
8.0
8.5
9.0
AFUB_016720 (II−1b)
P( −1.162 < μ1 − μ2 < −0.551 ) = 0.95 (sig)

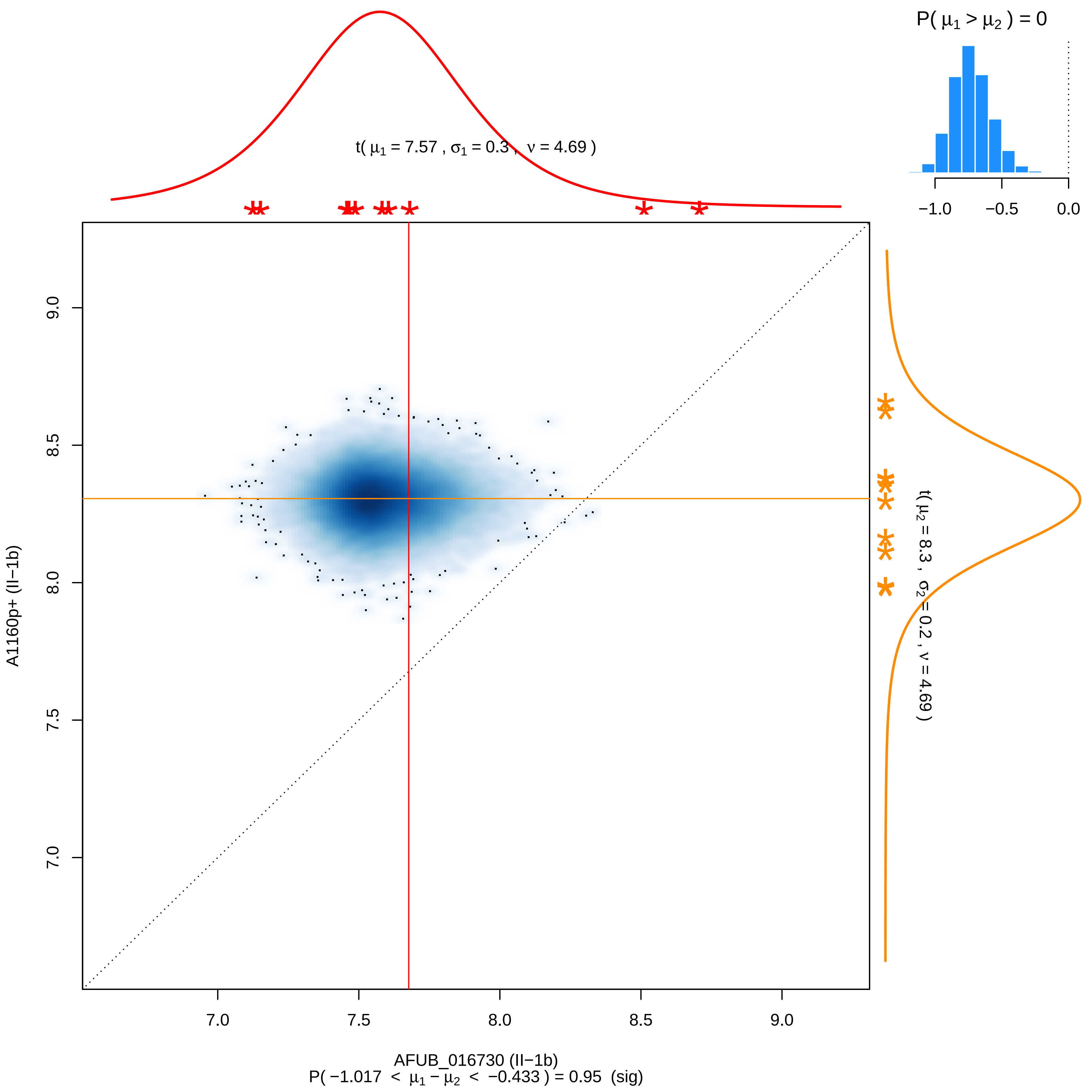

P( μ1 > μ2 ) = 0
−1.0
−0.5
0.0
t( μ1 = 7.57 , σ1 = 0.3 , ν = 4.69 )
t( μ2 = 8.3 , σ2 = 0.2 , ν = 4.69 )
A1160p+ (II−1b)
9.0
8.5
8.0
7.5
7.0
7.0
7.5
8.0
8.5
9.0
AFUB_016730 (II−1b)
P( −1.017 < μ1 − μ2 < −0.433 ) = 0.95 (sig)

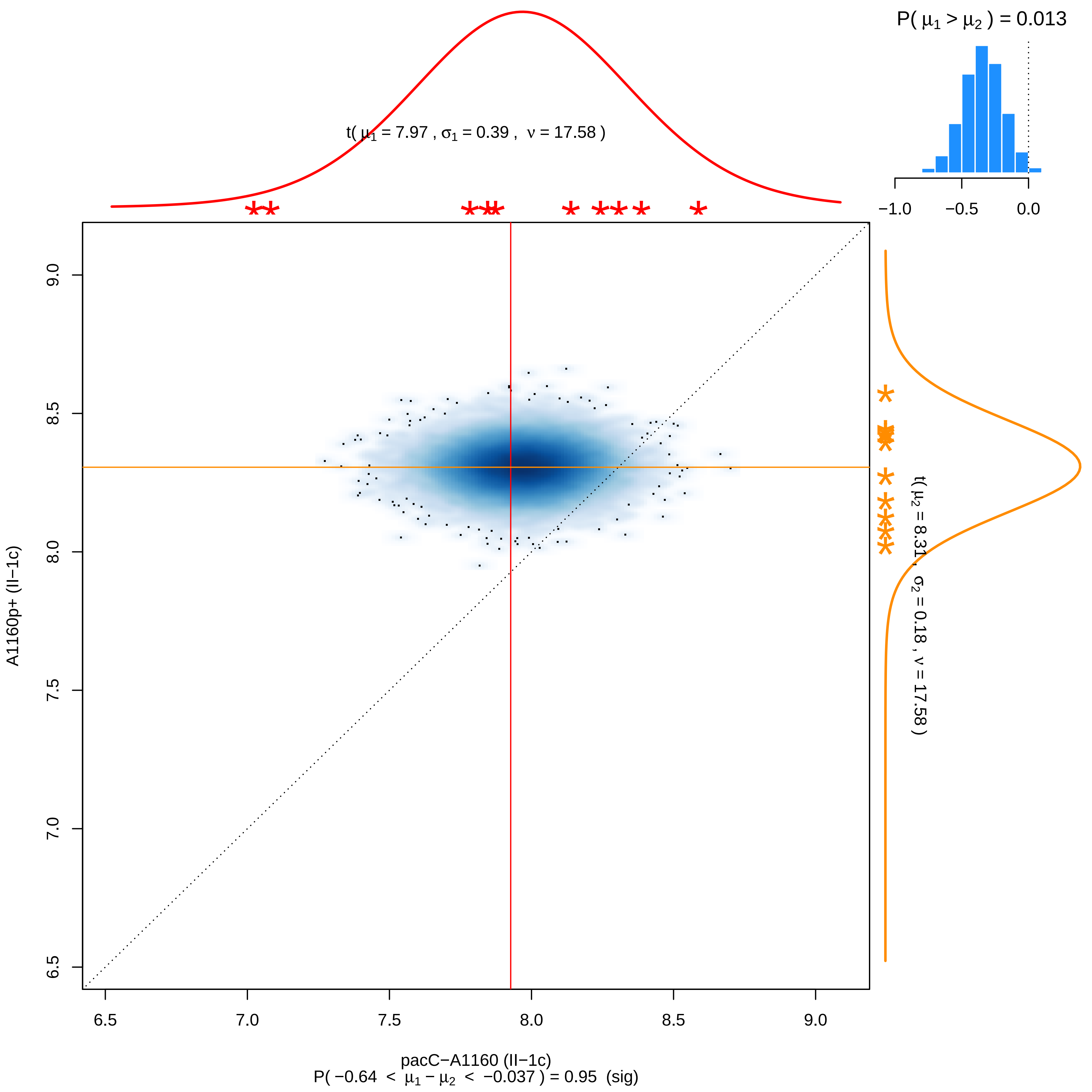

P( μ1 > μ2 ) = 0.013
−1.0
−0.5
0.0
t( μ1 = 7.97 , σ1 = 0.39 , ν = 17.58 )
t( μ2 = 8.31 , σ2 = 0.18 , ν = 17.58 )
A1160p+ (II−1c)
9.0
8.5
8.0
7.5
7.0
6.5
6.5
7.0
7.5
8.0
8.5
9.0
pacC−A1160 (II−1c)
P( −0.64 < μ1 − μ2 < −0.037 ) = 0.95 (sig)

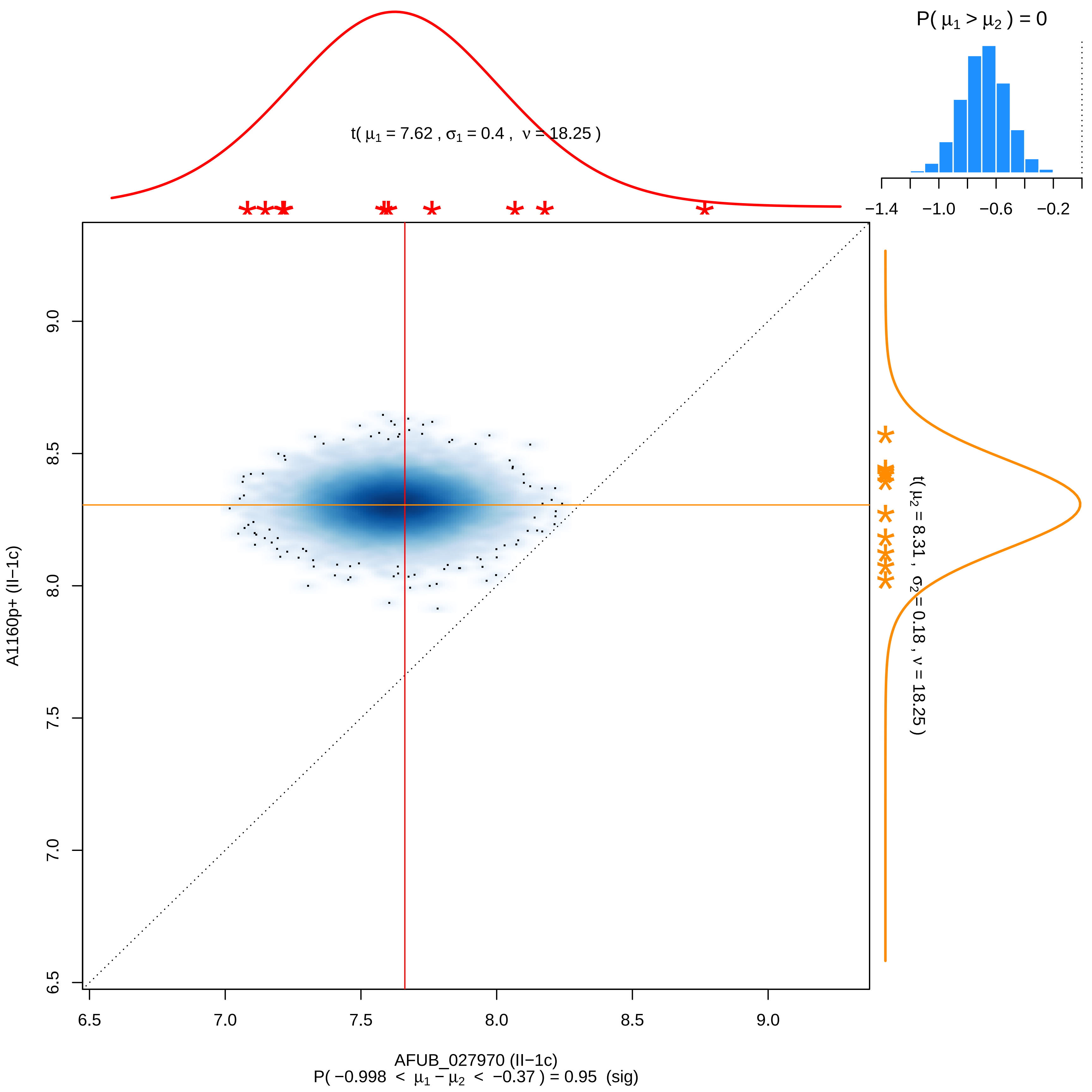

P( μ1 > μ2 ) = 0
−1.4
−1.0
−0.6
−0.2
t( μ1 = 7.62 , σ1 = 0.4 , ν = 18.25 )
t( μ2 = 8.31 , σ2 = 0.18 , ν = 18.25 )
A1160p+ (II−1c)
6.5
7.0
7.5
8.0
8.5
9.0
AFUB_027970 (II−1c)
P( −0.998 < μ1 − μ2 < −0.37 ) = 0.95 (sig)

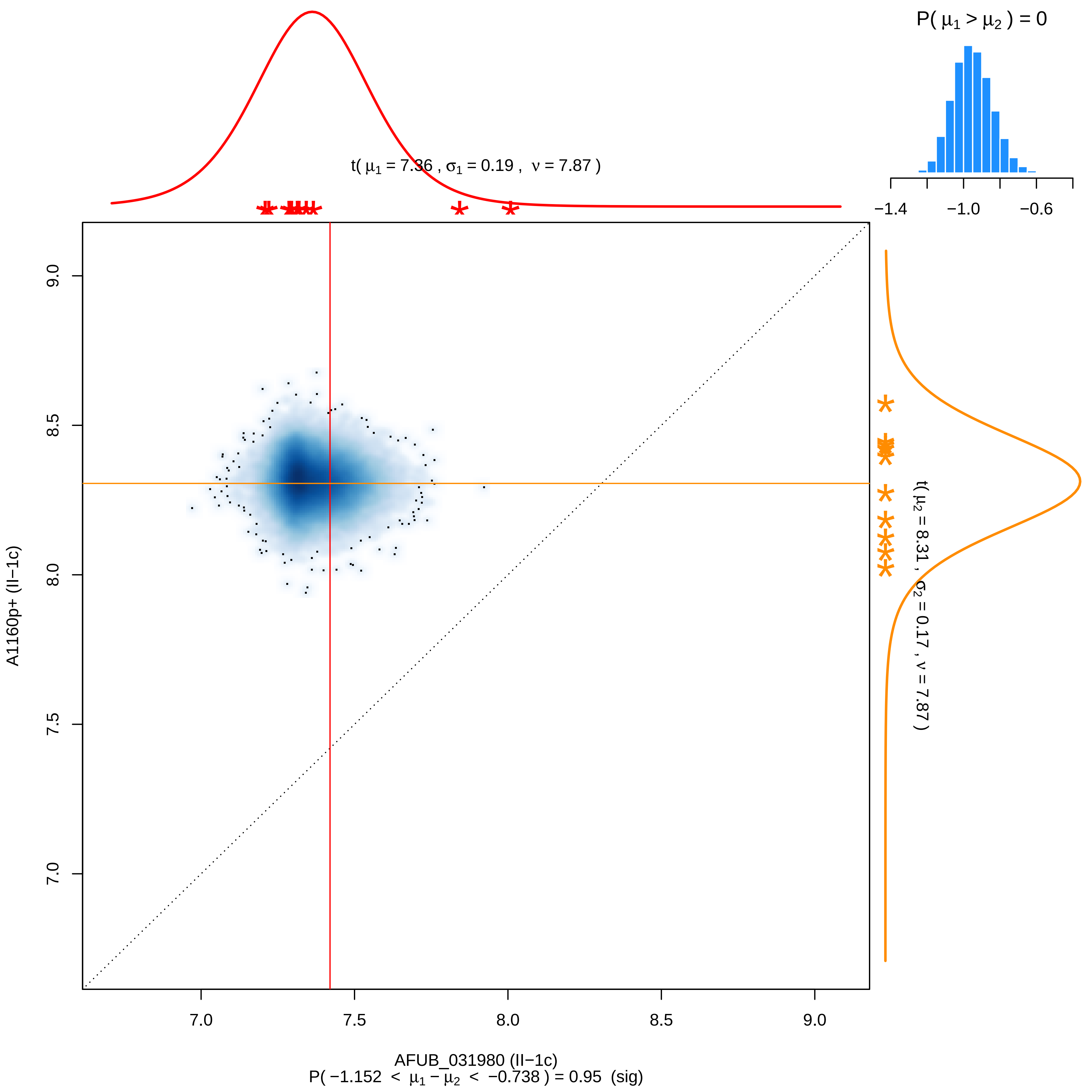

P( μ1 > μ2 ) = 0
−1.4
−1.0
−0.6
t( μ1 = 7.36 , σ1 = 0.19 , ν = 7.87 )
t( μ2 = 8.31 , σ2 = 0.17 , ν = 7.87 )
9.0
8.5
8.0
7.5
7.0
A1160p+ (II−1c)
7.0
7.5
8.0
8.5
9.0
AFUB_031980 (II−1c)
P( −1.152 < μ1 − μ2 < −0.738 ) = 0.95 (sig)

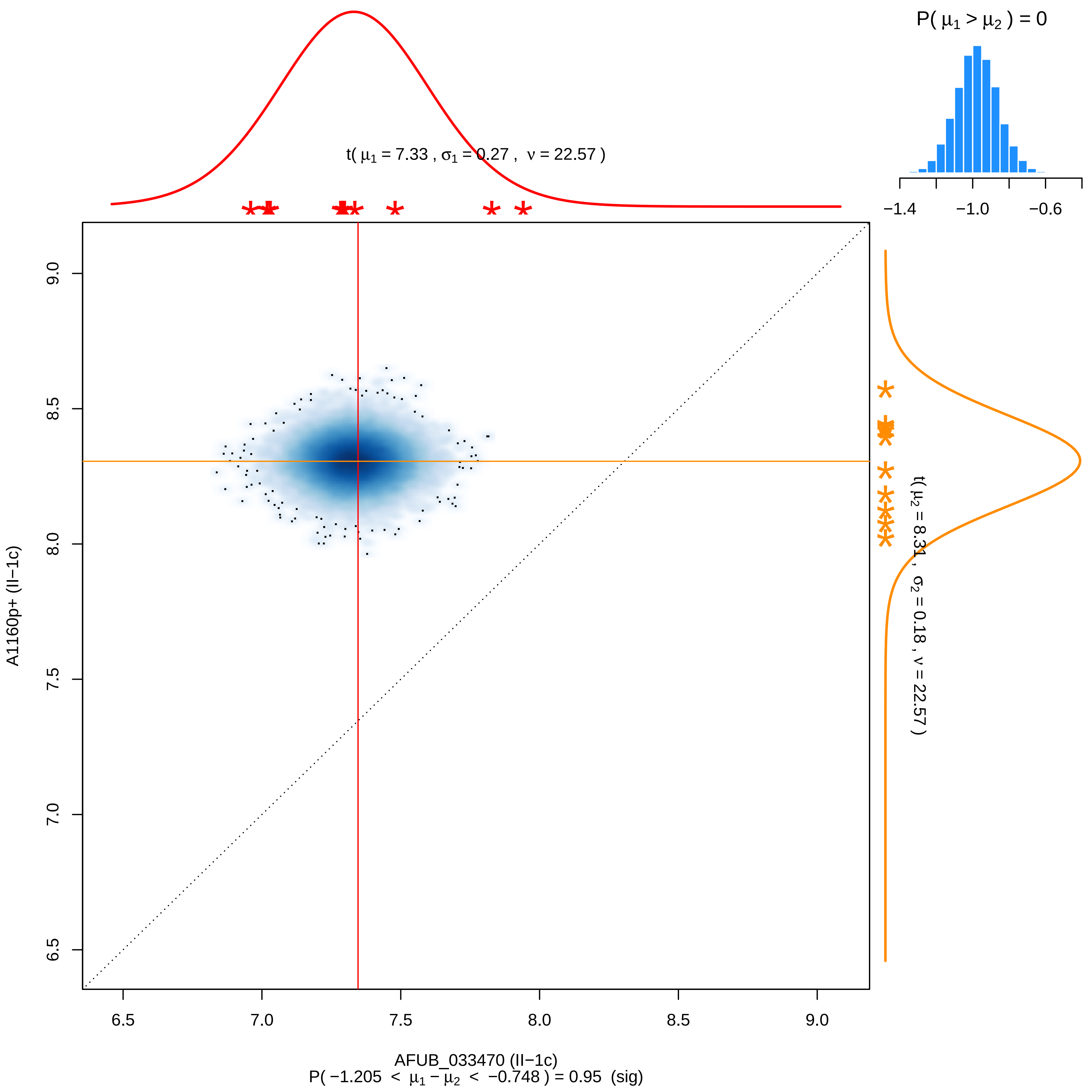

P( μ1 > μ2 ) = 0
−1.4
−1.0
−0.6
t( μ1 = 7.33 , σ1 = 0.27 , ν = 22.57 )
t( μ2 = 8.31 , σ2 = 0.18 , ν = 22.57 )
A1160p+ (II−1c)
9.0
8.5
8.0
7.5
7.0
6.5
6.5
7.0
7.5
8.0
8.5
9.0
AFUB_033470 (II−1c)
P( −1.205 < μ1 − μ2 < −0.748 ) = 0.95 (sig)

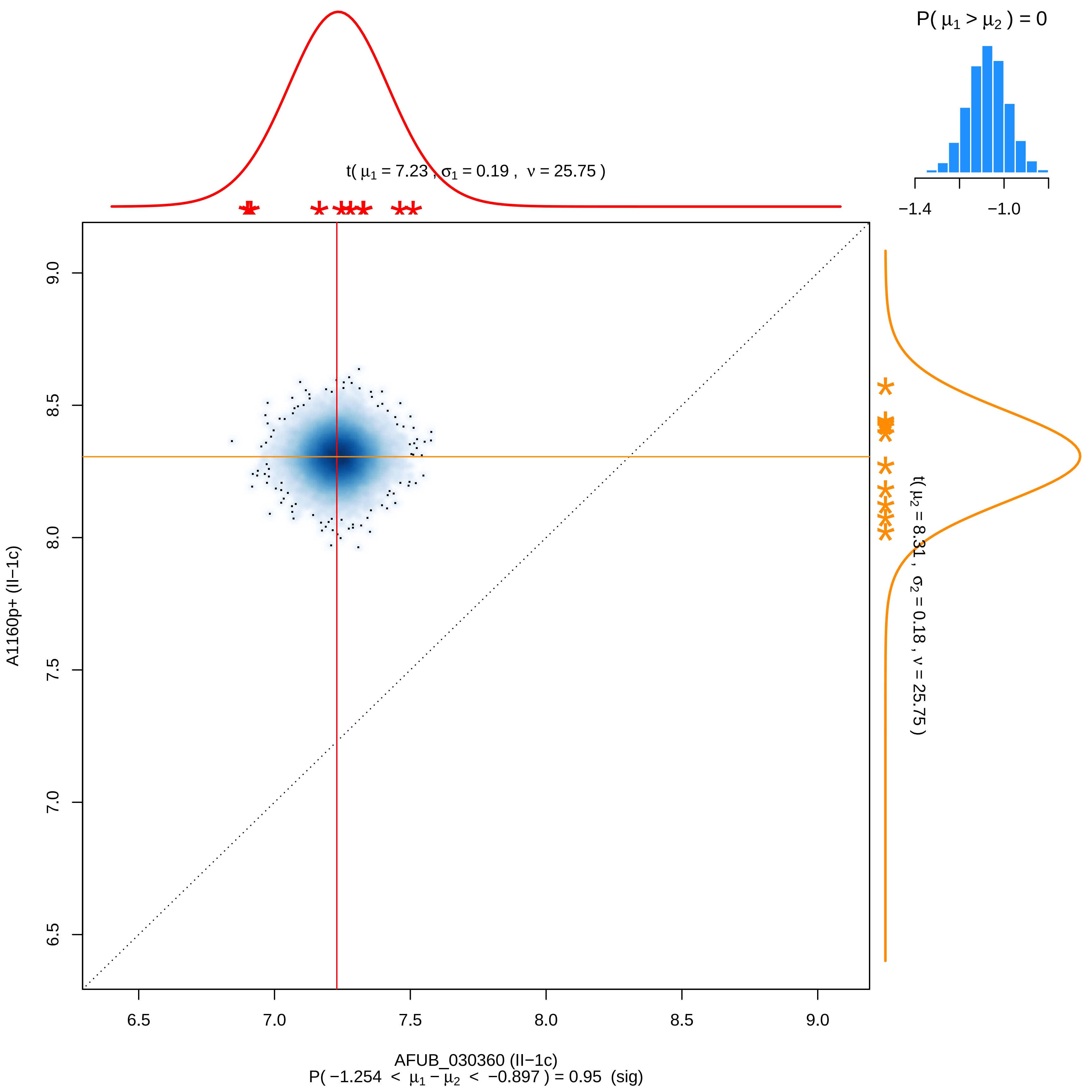
P( $\mu_1 > \mu_2$ ) = 0
−1.4
−1.0
t( $\mu_1$ = 7.23 , $\sigma_1$ = 0.19 , $\nu$ = 25.75 )
t( $\mu_2$ = 8.31 , $\sigma_2$ = 0.18 , $\nu$ = 25.75 )
A1160p+ (II−1c)
9.0
8.5
8.0
7.5
7.0
6.5
6.5
7.0
7.5
8.0
8.5
9.0
AFUB_030360 (II−1c)
P( −1.254 < $\mu_1 - \mu_2$ < −0.897 ) = 0.95 (sig)

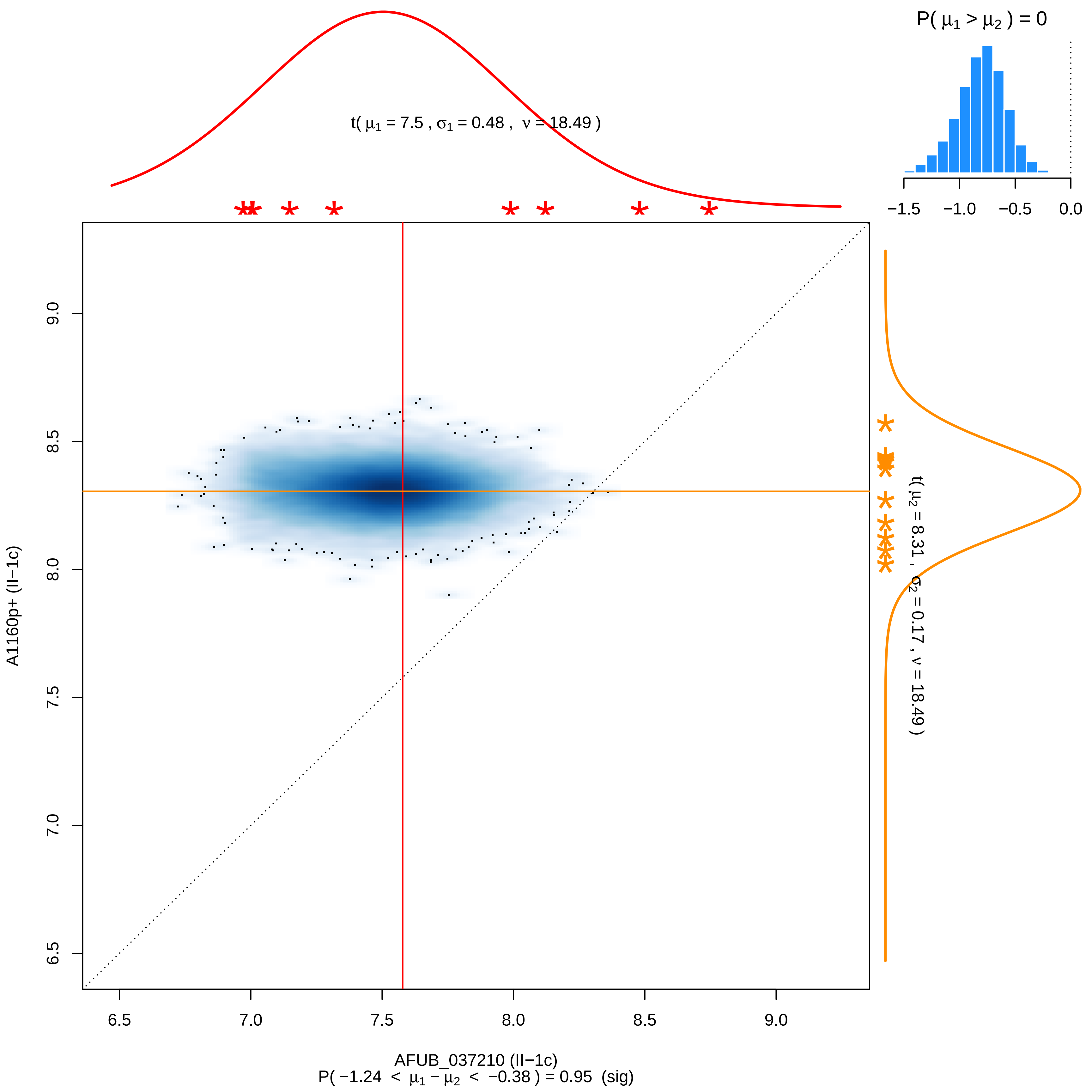

P( μ1 > μ2 ) = 0
−1.5 −1.0 −0.5 0.0
t( μ1 = 7.5 , σ1 = 0.48 , ν = 18.49 )
t( μ2 = 8.31 , σ2 = 0.17 , ν = 18.49 )
A1160p+ (II−1c)
AFUB_037210 (II−1c)
P( −1.24 < μ1 − μ2 < −0.38 ) = 0.95 (sig)
6.5 7.0 7.5 8.0 8.5 9.0

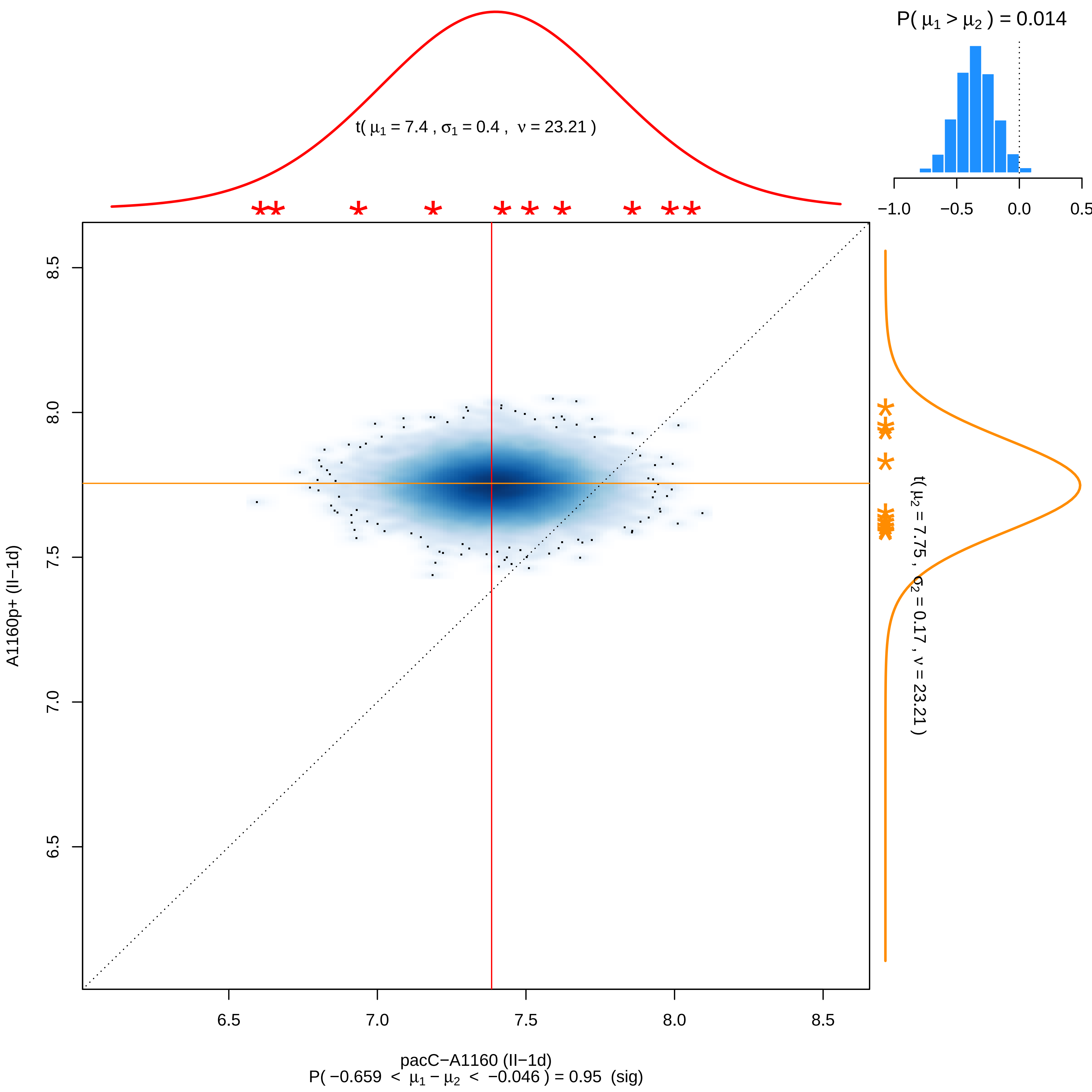

P( μ1 > μ2 ) = 0.014
−1.0
−0.5
0.0
0.5
t( μ1 = 7.4 , σ1 = 0.4 , ν = 23.21 )
t( μ2 = 7.75 , σ2 = 0.17 , ν = 23.21 )
A1160p+ (II−1d)
8.5
8.0
7.5
7.0
6.5
6.5
7.0
7.5
8.0
8.5
pacC−A1160 (II−1d)
P( −0.659 < μ1 − μ2 < −0.046 ) = 0.95 (sig)

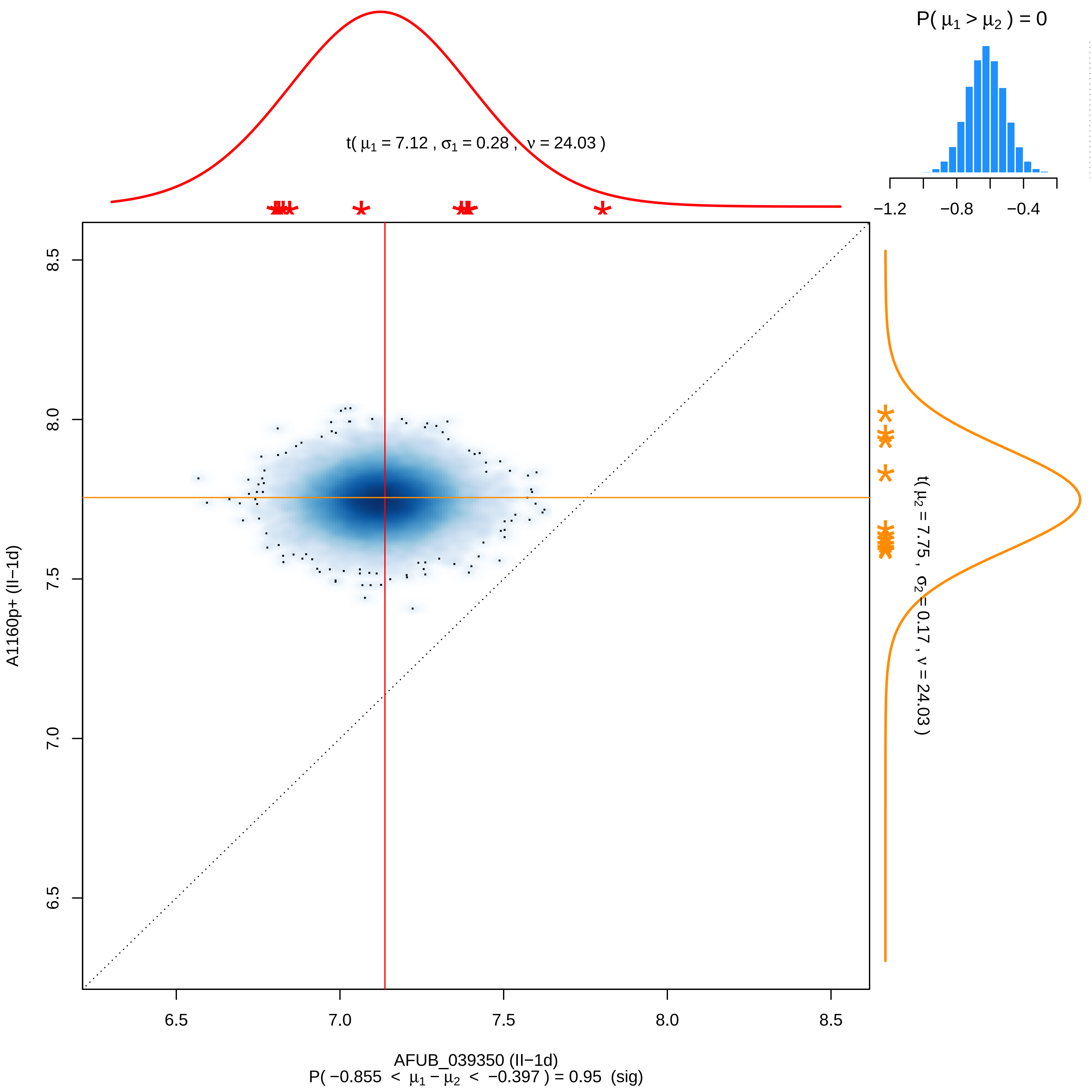
P( μ1 > μ2 ) = 0
−1.2
−0.8
−0.4
t( μ1 = 7.12 , σ1 = 0.28 , ν = 24.03 )
t( μ2 = 7.75 , σ2 = 0.17 , ν = 24.03 )
A1160p+ (II−1d)
8.5
8.0
7.5
7.0
6.5
6.5
7.0
7.5
8.0
8.5
AFUB_039350 (II−1d)
P( −0.855 < μ1 − μ2 < −0.397 ) = 0.95 (sig)

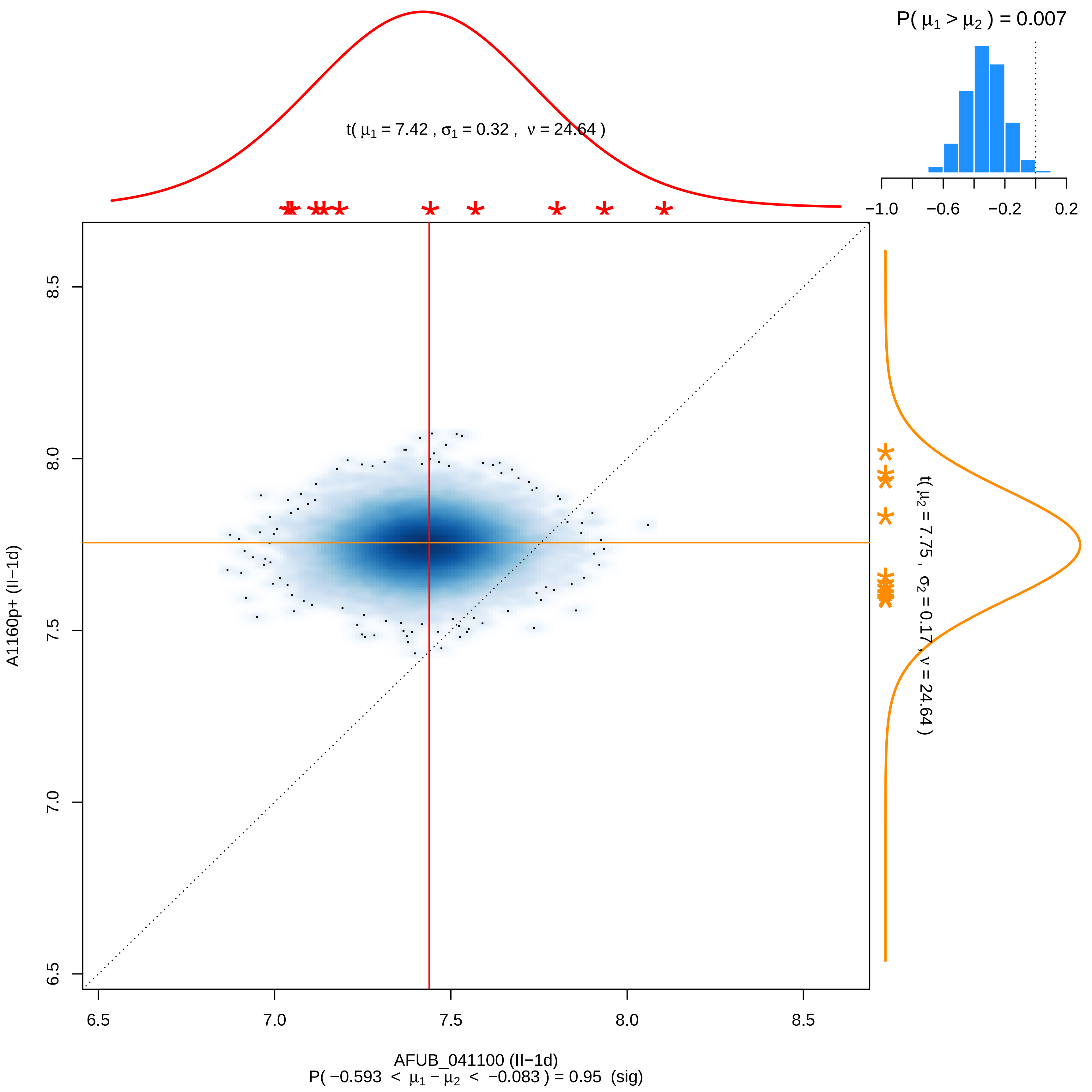

P( μ1 > μ2 ) = 0.007
−1.0 −0.6 −0.2 0.2
t( μ1 = 7.42 , σ1 = 0.32 , ν = 24.64 )
t( μ2 = 7.75 , σ2 = 0.17 , ν = 24.64 )
8.5
8.0
7.5
7.0
6.5
A1160p+ (II−1d)
6.5 7.0 7.5 8.0 8.5
AFUB_041100 (II−1d)
P( −0.593 < μ1 − μ2 < −0.083 ) = 0.95 (sig)

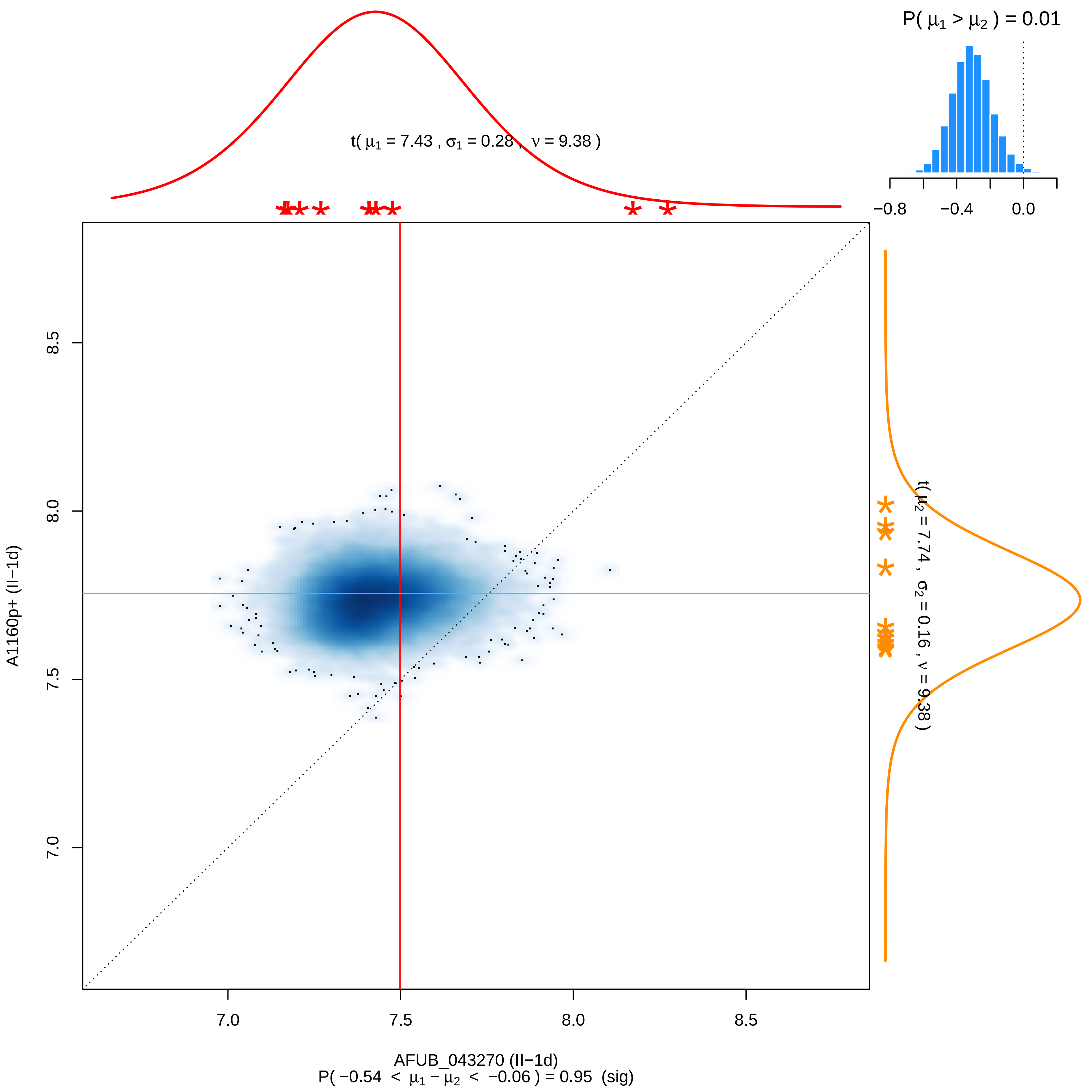

P( μ1 > μ2 ) = 0.01
−0.8
−0.4
0.0
t( μ1 = 7.43 , σ1 = 0.28 , ν = 9.38 )
t( μ2 = 7.74 , σ2 = 0.16 , ν = 9.38 )
8.5
8.0
7.5
7.0
A1160p+ (II−1d)
7.0
7.5
8.0
8.5
AFUB_043270 (II−1d)
P( −0.54 < μ1 − μ2 < −0.06 ) = 0.95 (sig)

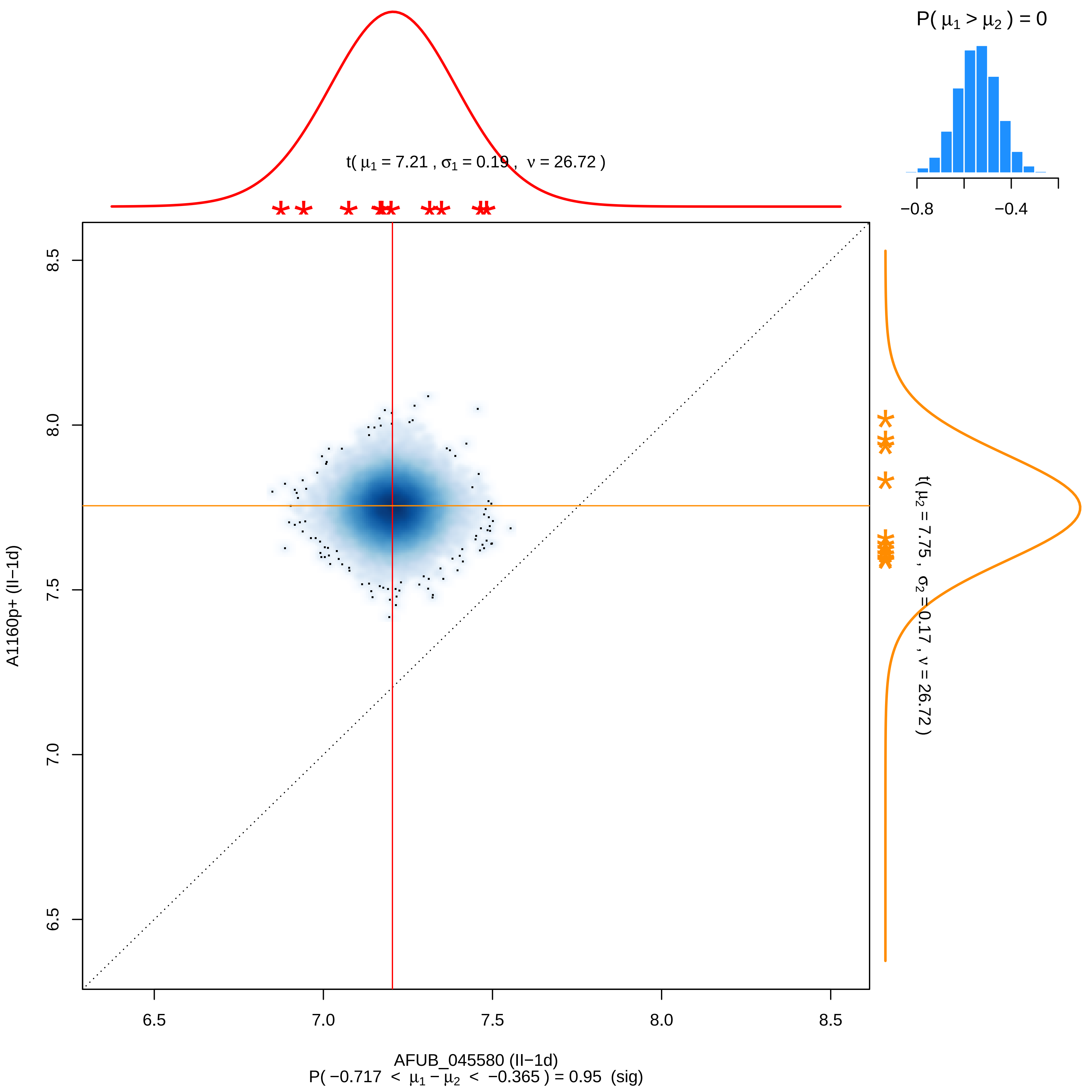

P( μ1 > μ2 ) = 0
−0.8
−0.4
t( μ1 = 7.21 , σ1 = 0.19 , ν = 26.72 )
8.5
8.0
7.5
7.0
6.5
A1160p+ (II−1d)
t( μ2 = 7.75 , σ2 = 0.17 , ν = 26.72 )
6.5
7.0
7.5
8.0
8.5
AFUB_045580 (II−1d)
P( −0.717 < μ1 − μ2 < −0.365 ) = 0.95 (sig)

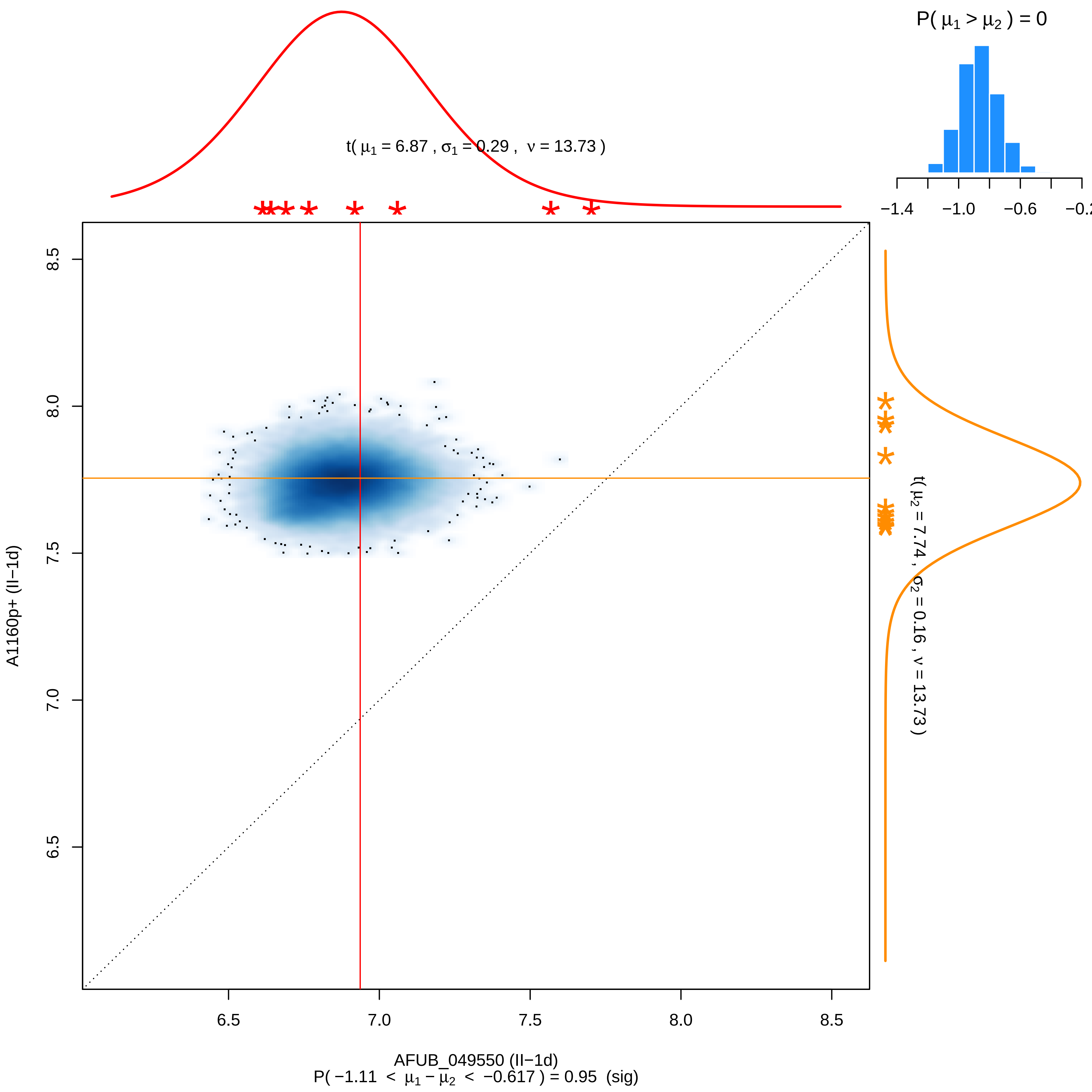

P( μ1 > μ2 ) = 0
−1.4
−1.0
−0.6
−0.2
t( μ1 = 6.87 , σ1 = 0.29 , ν = 13.73 )
t( μ2 = 7.74 , σ2 = 0.16 , ν = 13.73 )
8.5
8.0
7.5
7.0
6.5
A1160p+ (II−1d)
6.5
7.0
7.5
8.0
8.5
AFUB_049550 (II−1d)
P( −1.11 < μ1 − μ2 < −0.617 ) = 0.95 (sig)

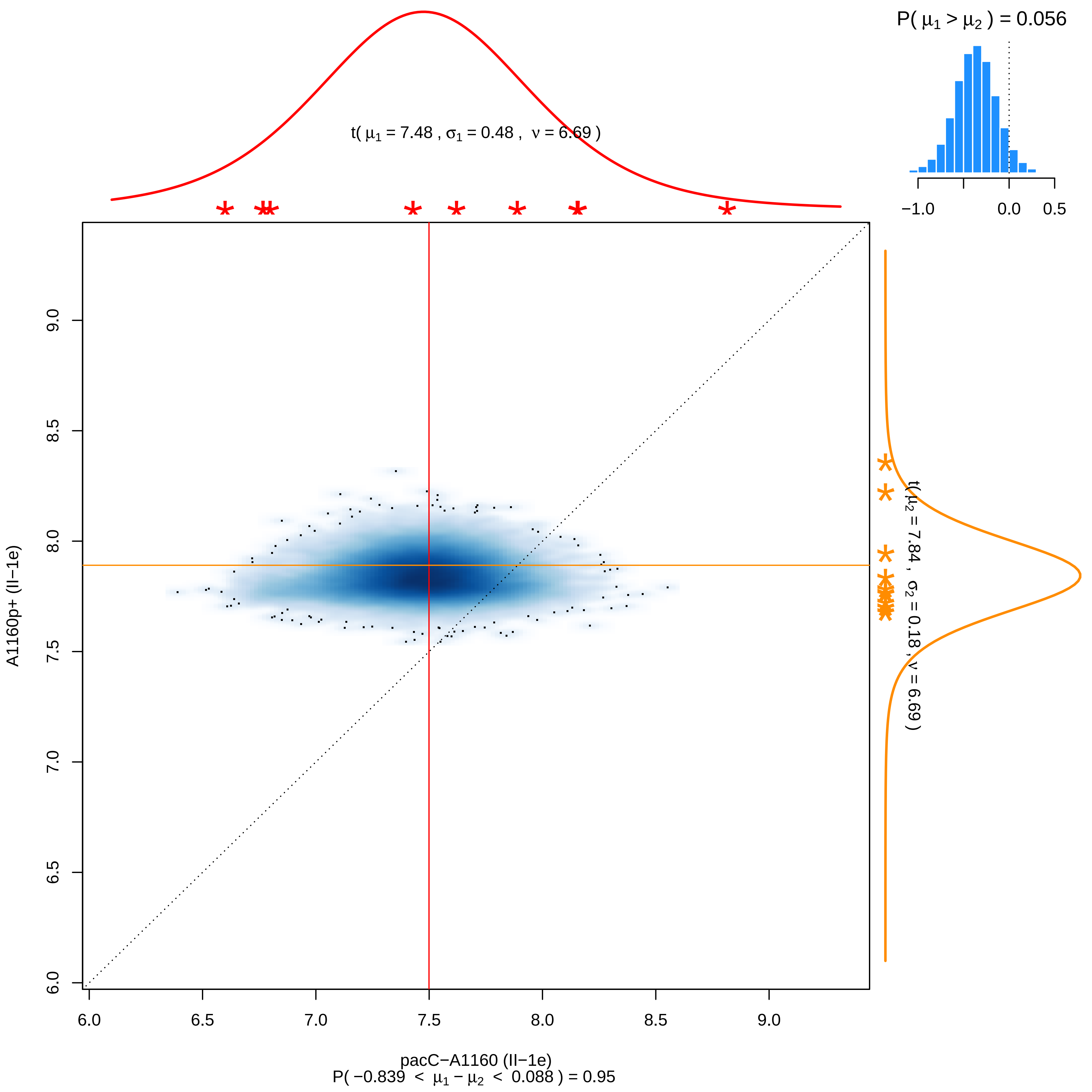

P( μ1 > μ2 ) = 0.056
−1.0
0.0
0.5
t( μ1 = 7.48 , σ1 = 0.48 , ν = 6.69 )
t( μ2 = 7.84 , σ2 = 0.18 , ν = 6.69 )
A1160p+ (II−1e)
6.0
6.5
7.0
7.5
8.0
8.5
9.0
pacC−A1160 (II−1e)
P( −0.839 < μ1 − μ2 < 0.088 ) = 0.95

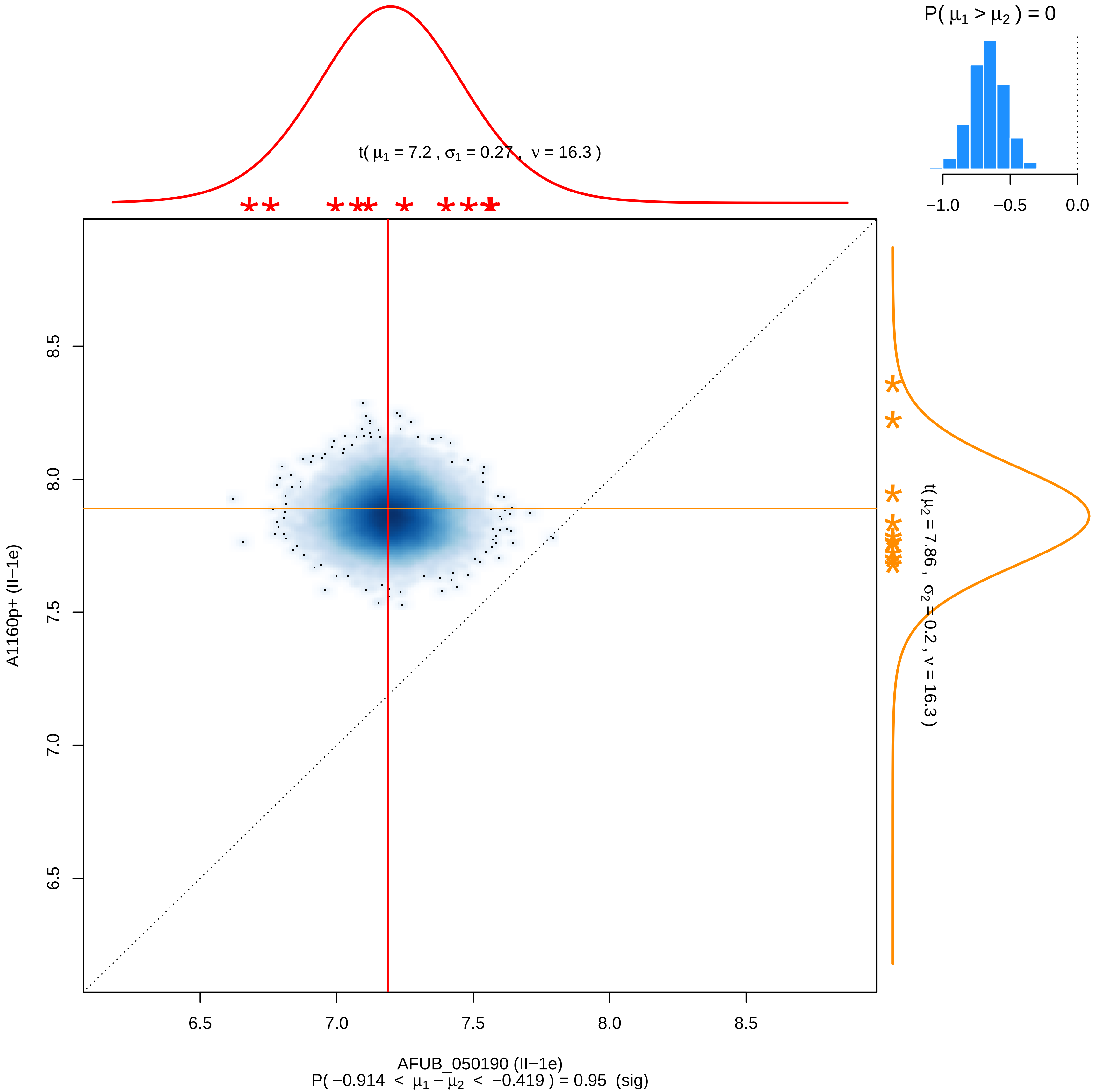

P( μ1 > μ2 ) = 0
−1.0
−0.5
0.0
t( μ1 = 7.2 , σ1 = 0.27 , ν = 16.3 )
t( μ2 = 7.86 , σ2 = 0.2 , ν = 16.3 )
A1160p+ (II−1e)
6.5
7.0
7.5
8.0
8.5
AFUB_050190 (II−1e)
P( −0.914 < μ1 − μ2 < −0.419 ) = 0.95 (sig)

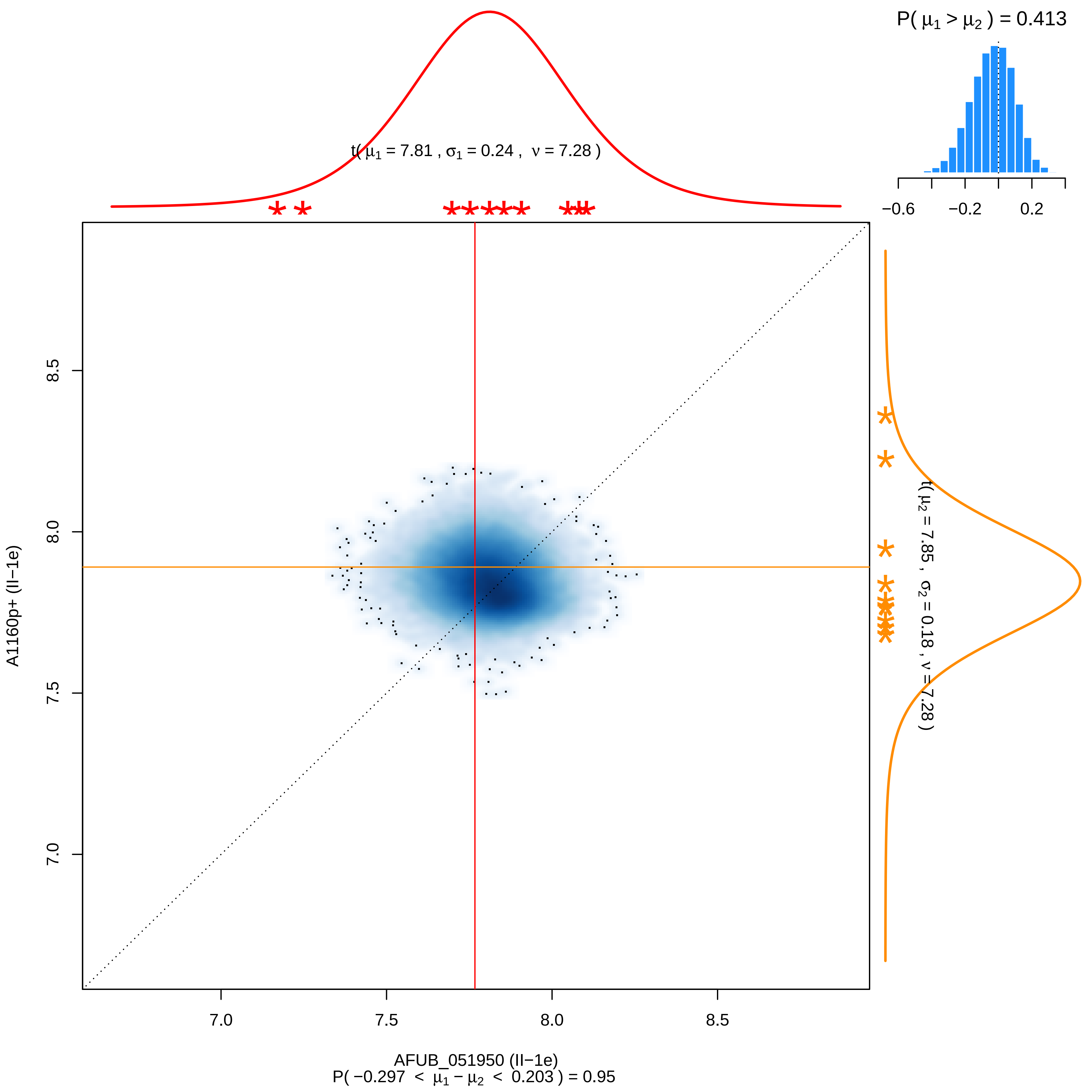

P( μ1 > μ2 ) = 0.413
−0.6
−0.2
0.2
t( μ1 = 7.81 , σ1 = 0.24 , ν = 7.28 )
t( μ2 = 7.85 , σ2 = 0.18 , ν = 7.28 )
8.5
8.0
7.5
7.0
A1160p+ (II−1e)
7.0
7.5
8.0
8.5
AFUB_051950 (II−1e)
P( −0.297 < μ1 − μ2 < 0.203 ) = 0.95

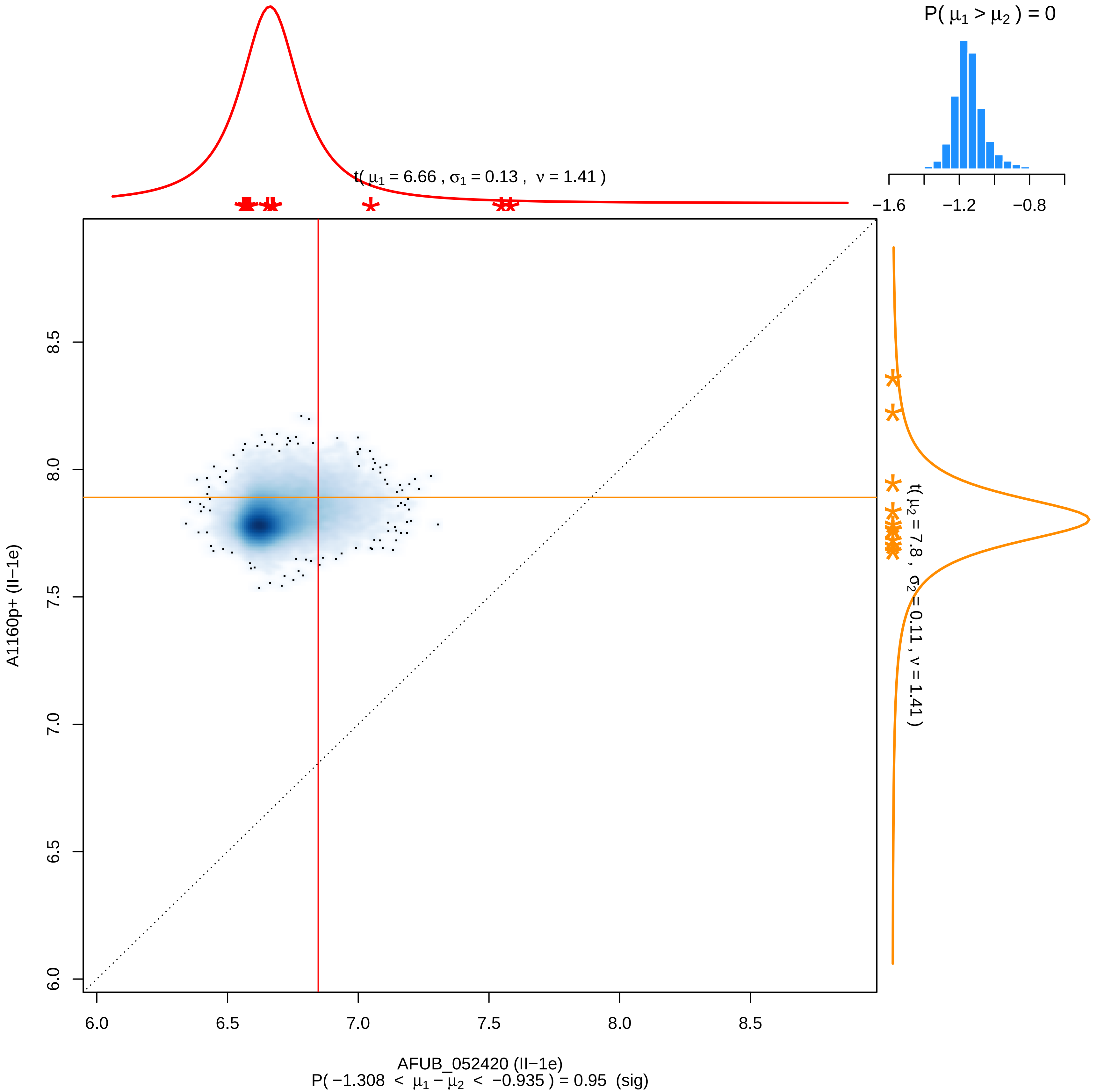

P( μ1 > μ2 ) = 0
−1.6
−1.2
−0.8
t( μ1 = 6.66 , σ1 = 0.13 , ν = 1.41 )
t( μ2 = 7.8 , σ2 = 0.11 , ν = 1.41 )
8.5
8.0
7.5
7.0
6.5
6.0
A1160p+ (II−1e)
6.0
6.5
7.0
7.5
8.0
8.5
AFUB_052420 (II−1e)
P( −1.308 < μ1 − μ2 < −0.935 ) = 0.95 (sig)

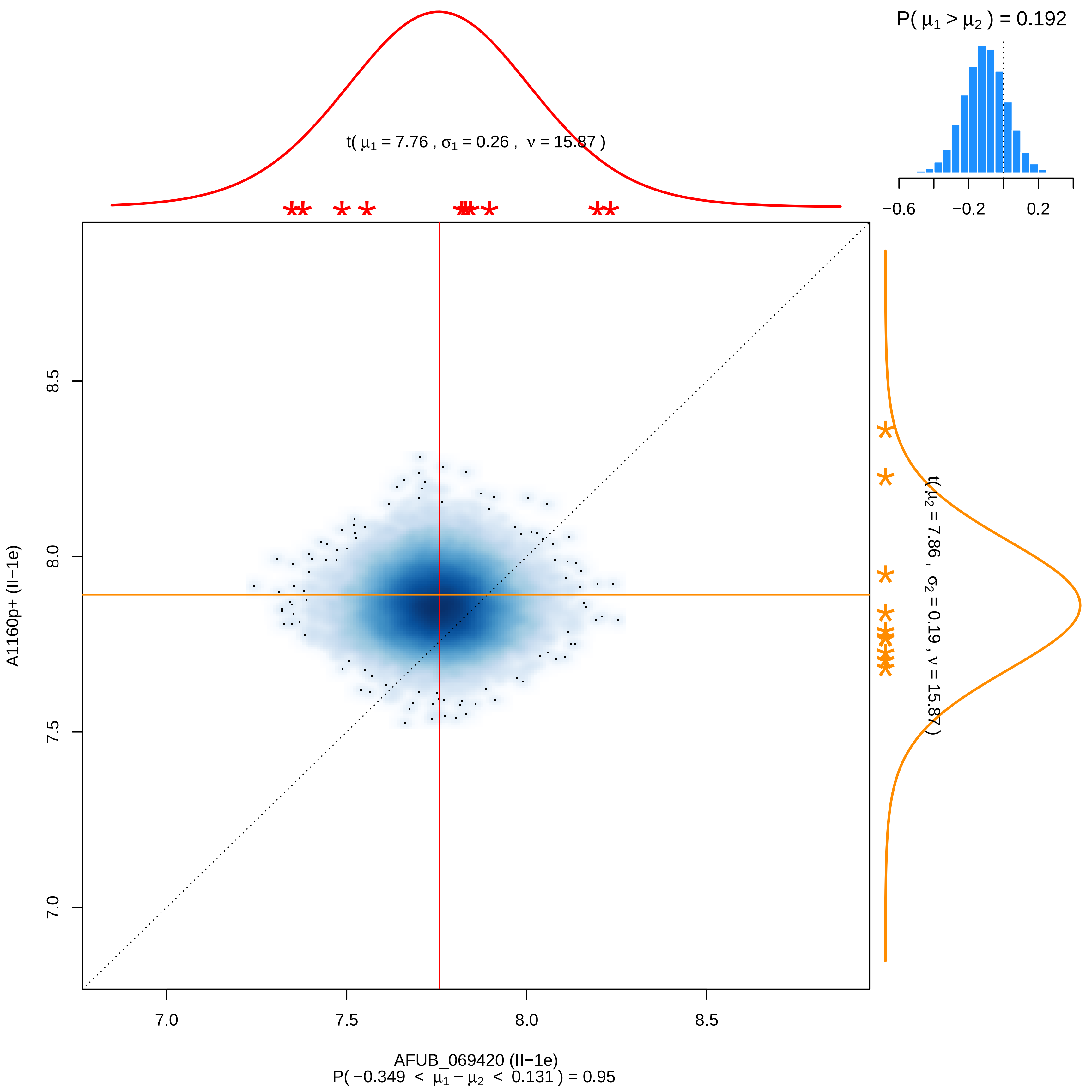

P( μ1 > μ2 ) = 0.192
−0.6
−0.2
0.2
t( μ1 = 7.76 , σ1 = 0.26 , ν = 15.87 )
t( μ2 = 7.86 , σ2 = 0.19 , ν = 15.87 )
A1160p+ (II−1e)
8.5
8.0
7.5
7.0
AFUB_069420 (II−1e)
P( −0.349 < μ1 − μ2 < 0.131 ) = 0.95

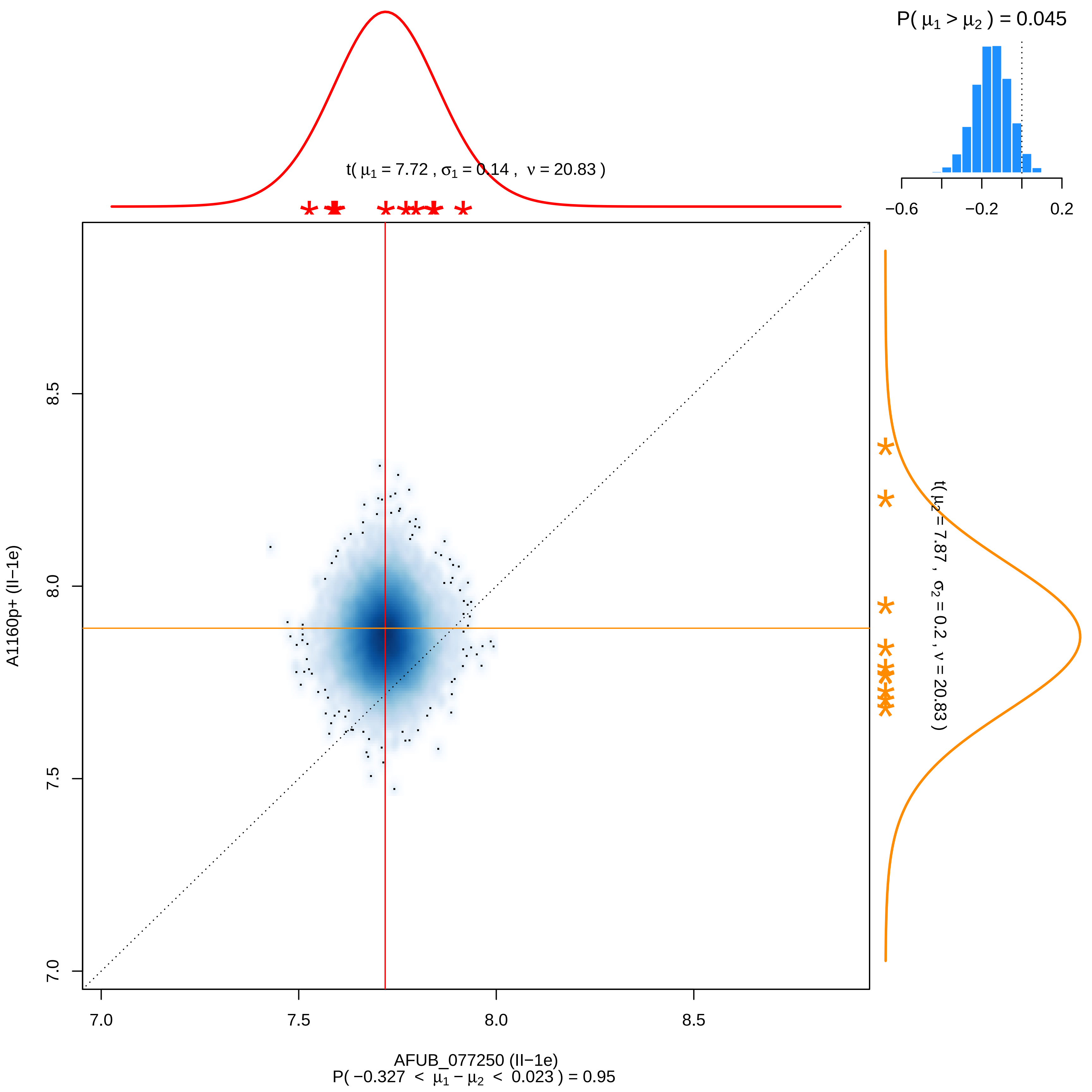

P( μ1 > μ2 ) = 0.045
−0.6
−0.2
0.2
t( μ1 = 7.72 , σ1 = 0.14 , ν = 20.83 )
t( μ2 = 7.87 , σ2 = 0.2 , ν = 20.83 )
8.5
8.0
7.5
7.0
A1160p+ (II−1e)
7.0
7.5
8.0
8.5
AFUB_077250 (II−1e)
P( −0.327 < μ1 − μ2 < 0.023 ) = 0.95

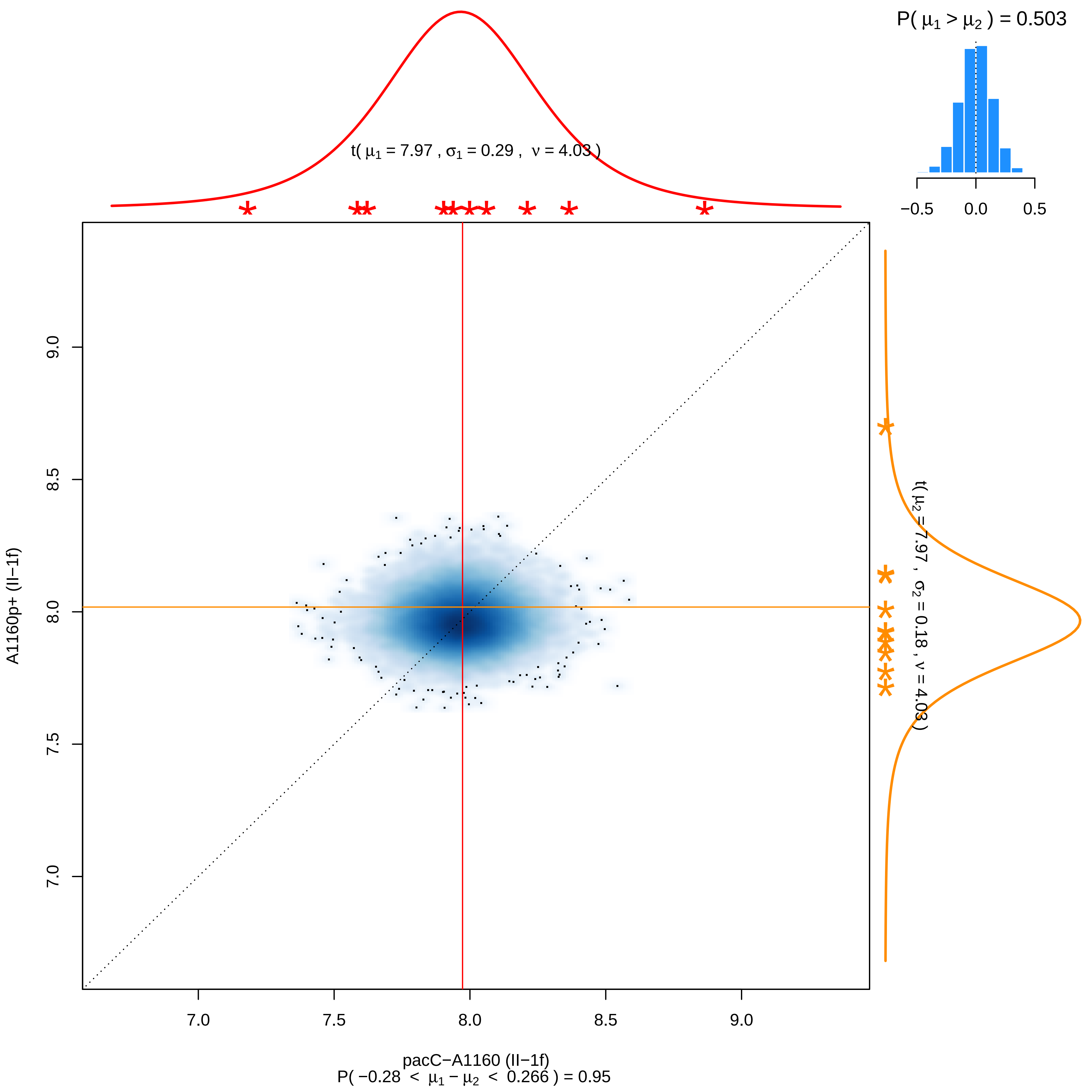

P( μ1 > μ2 ) = 0.503
−0.5
0.0
0.5
t( μ1 = 7.97 , σ1 = 0.29 , ν = 4.03 )
t( μ2 = 7.97 , σ2 = 0.18 , ν = 4.03 )
A1160p+ (II−1f)
9.0
8.5
8.0
7.5
7.0
7.0
7.5
8.0
8.5
9.0
pacC−A1160 (II−1f)
P( −0.28 < μ1 − μ2 < 0.266 ) = 0.95

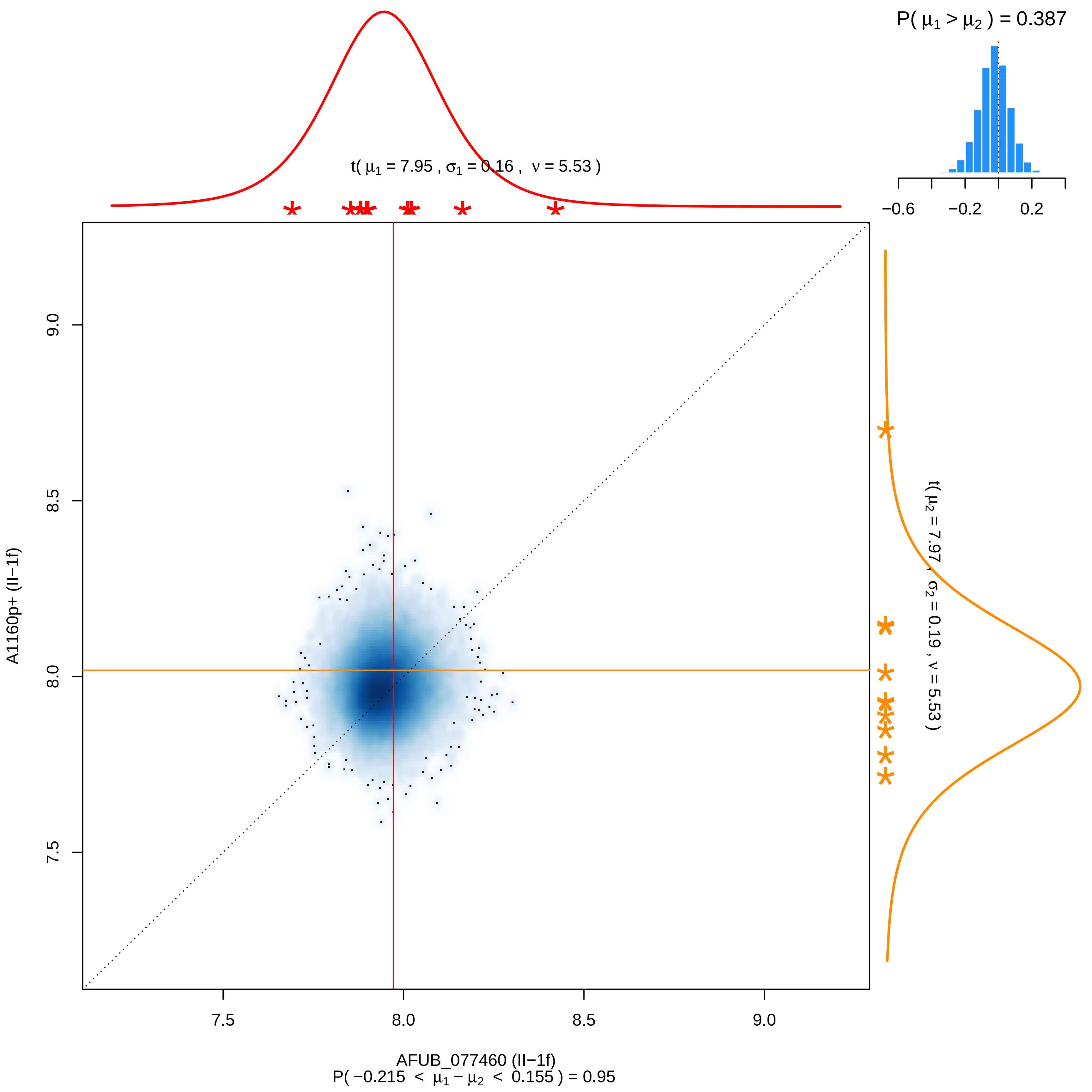

P( μ1 > μ2 ) = 0.387
−0.6
−0.2
0.2
t( μ1 = 7.95 , σ1 = 0.16 , ν = 5.53 )
t( μ2 = 7.97 , σ2 = 0.19 , ν = 5.53 )
A1160p+ (II−1f)
9.0
8.5
8.0
7.5
7.5
8.0
8.5
9.0
AFUB_077460 (II−1f)
P( −0.215 < μ1 − μ2 < 0.155 ) = 0.95

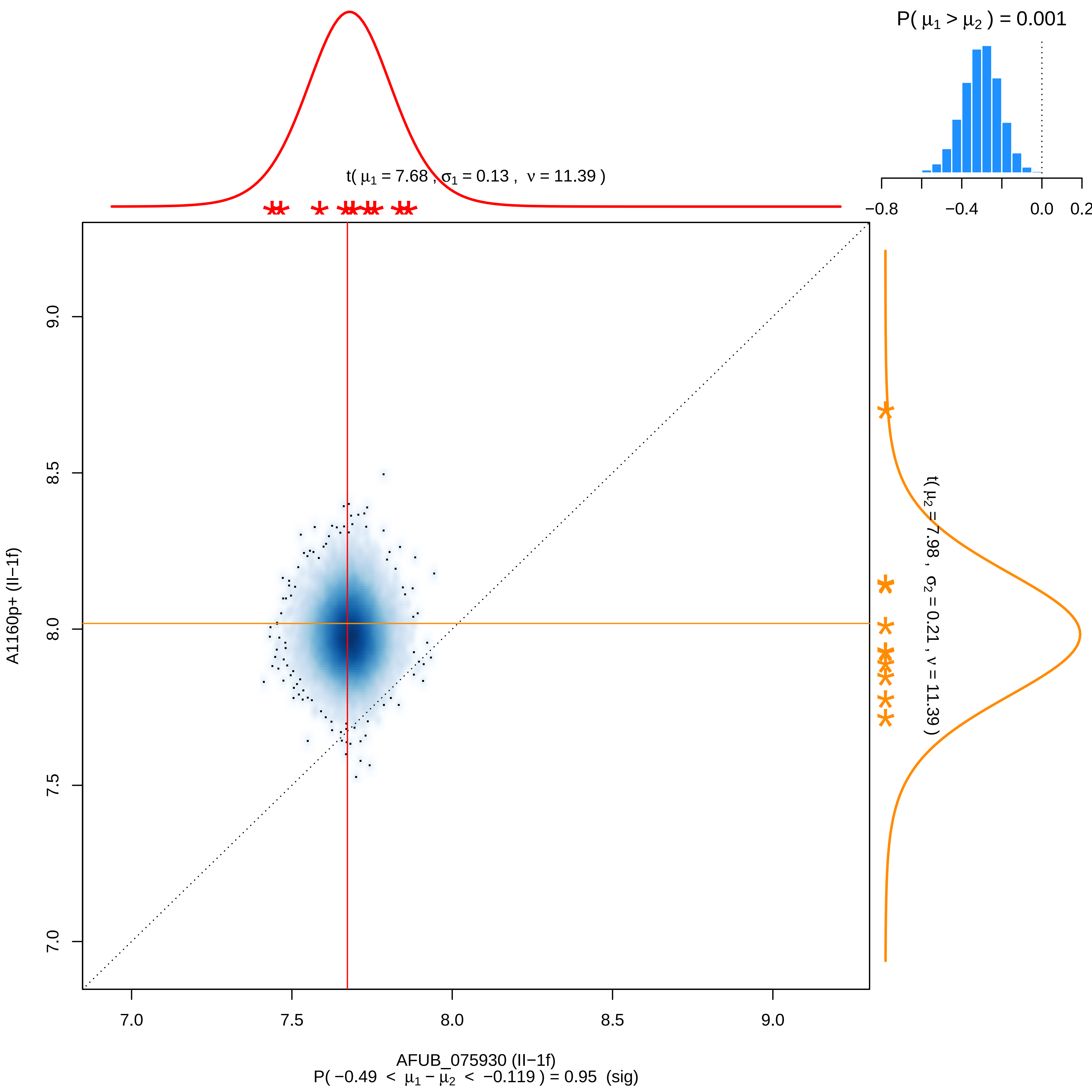

P( μ1 > μ2 ) = 0.001
−0.8
−0.4
0.0
0.2
t( μ1 = 7.68 , σ1 = 0.13 , ν = 11.39 )
t( μ2 = 7.98 , σ2 = 0.21 , ν = 11.39 )
A1160p+ (II−1f)
9.0
8.5
8.0
7.5
7.0
AFUB_075930 (II−1f)
P( −0.49 < μ1 − μ2 < −0.119 ) = 0.95 (sig)

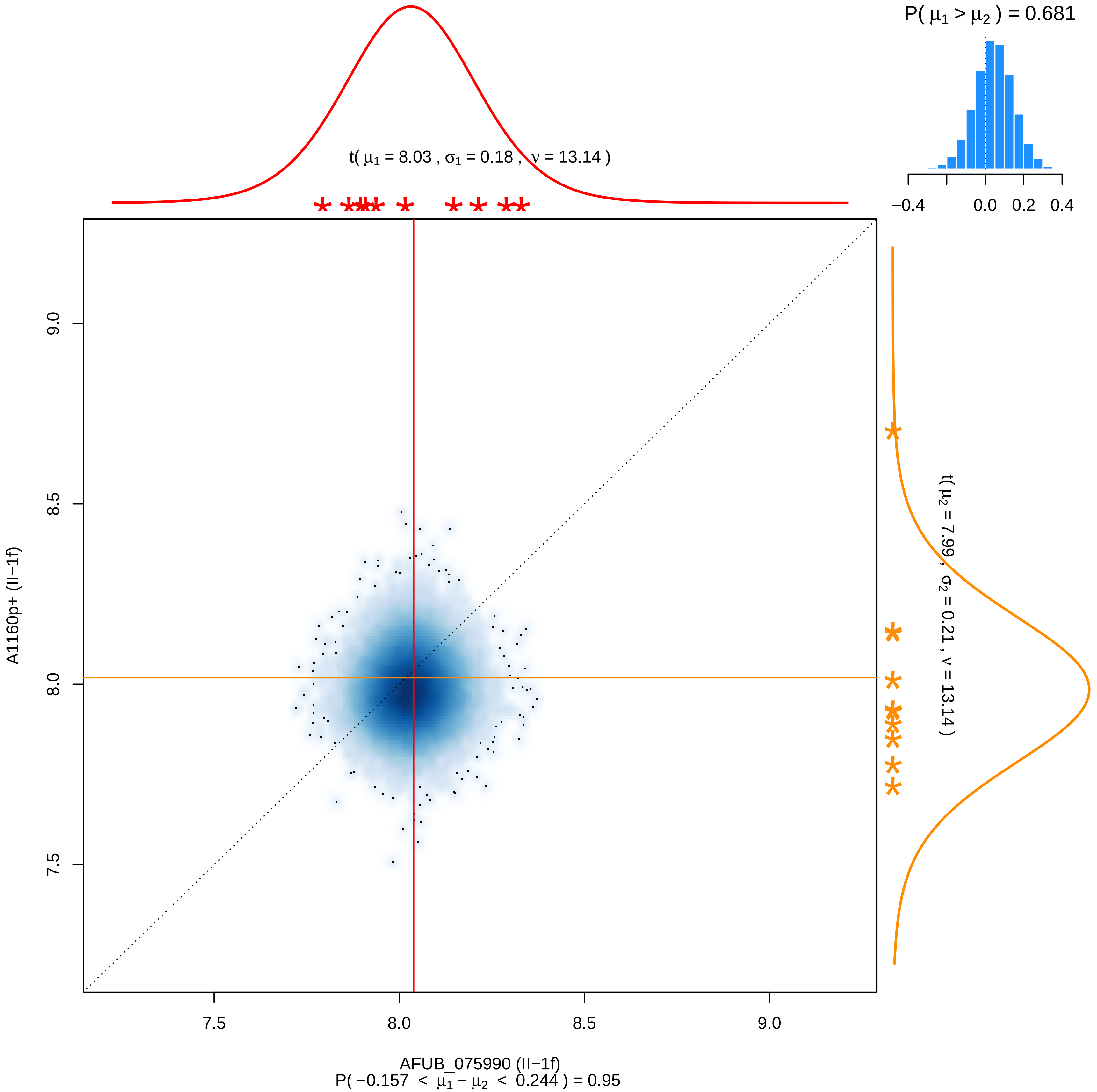

P( μ1 > μ2 ) = 0.681
−0.4
0.0
0.2
0.4
t( μ1 = 8.03 , σ1 = 0.18 , ν = 13.14 )
t( μ2 = 7.99 , σ2 = 0.21 , ν = 13.14 )
9.0
8.5
8.0
7.5
A1160p+ (II−1f)
7.5
8.0
8.5
9.0
AFUB_075990 (II−1f)
P( −0.157 < μ1 − μ2 < 0.244 ) = 0.95

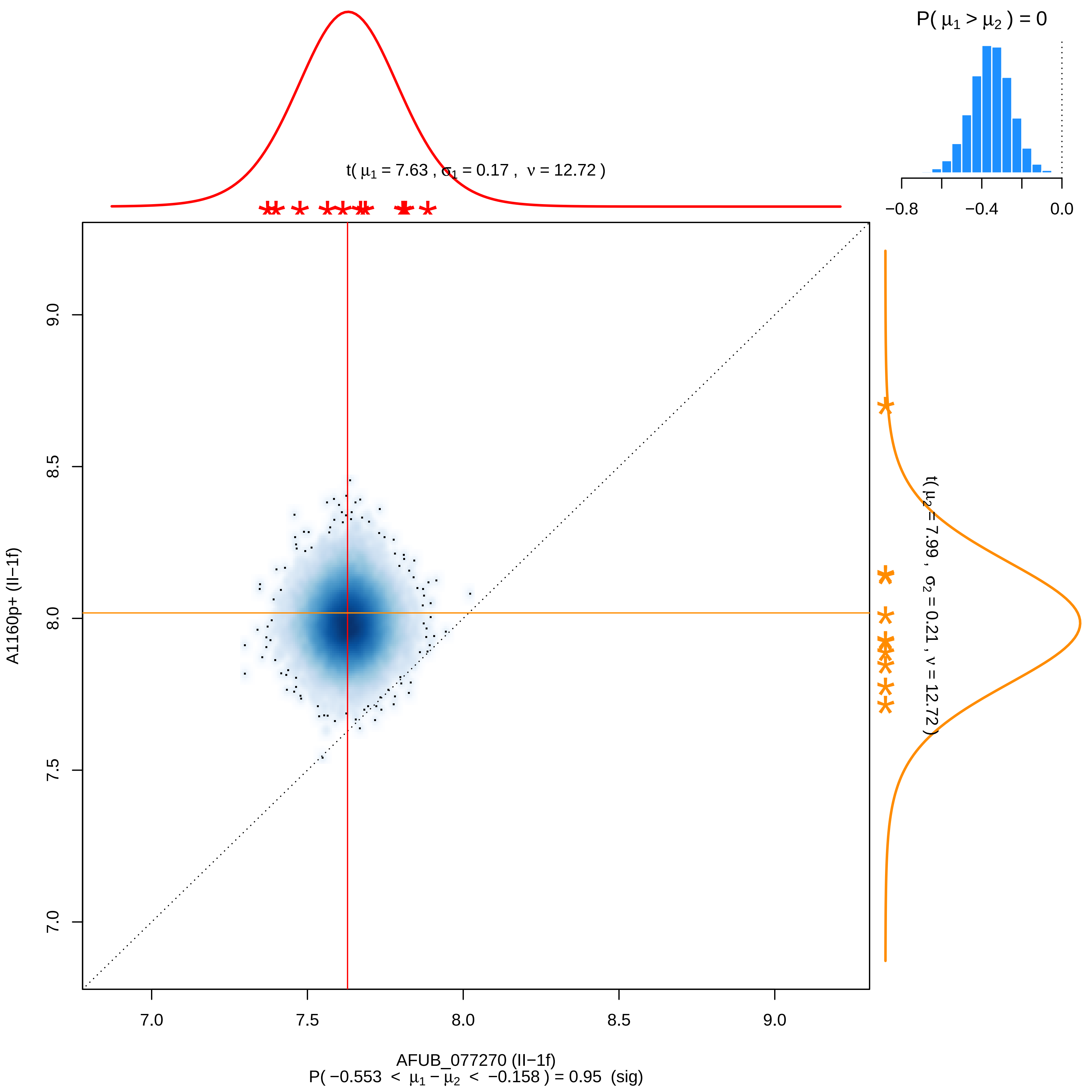

P( μ1 > μ2 ) = 0
−0.8
−0.4
0.0
t( μ1 = 7.63 , σ1 = 0.17 , ν = 12.72 )
t( μ2 = 7.99 , σ2 = 0.21 , ν = 12.72 )
A1160p+ (II−1f)
9.0
8.5
8.0
7.5
7.0
7.0
7.5
8.0
8.5
9.0
AFUB_077270 (II−1f)
P( −0.553 < μ1 − μ2 < −0.158 ) = 0.95 (sig)

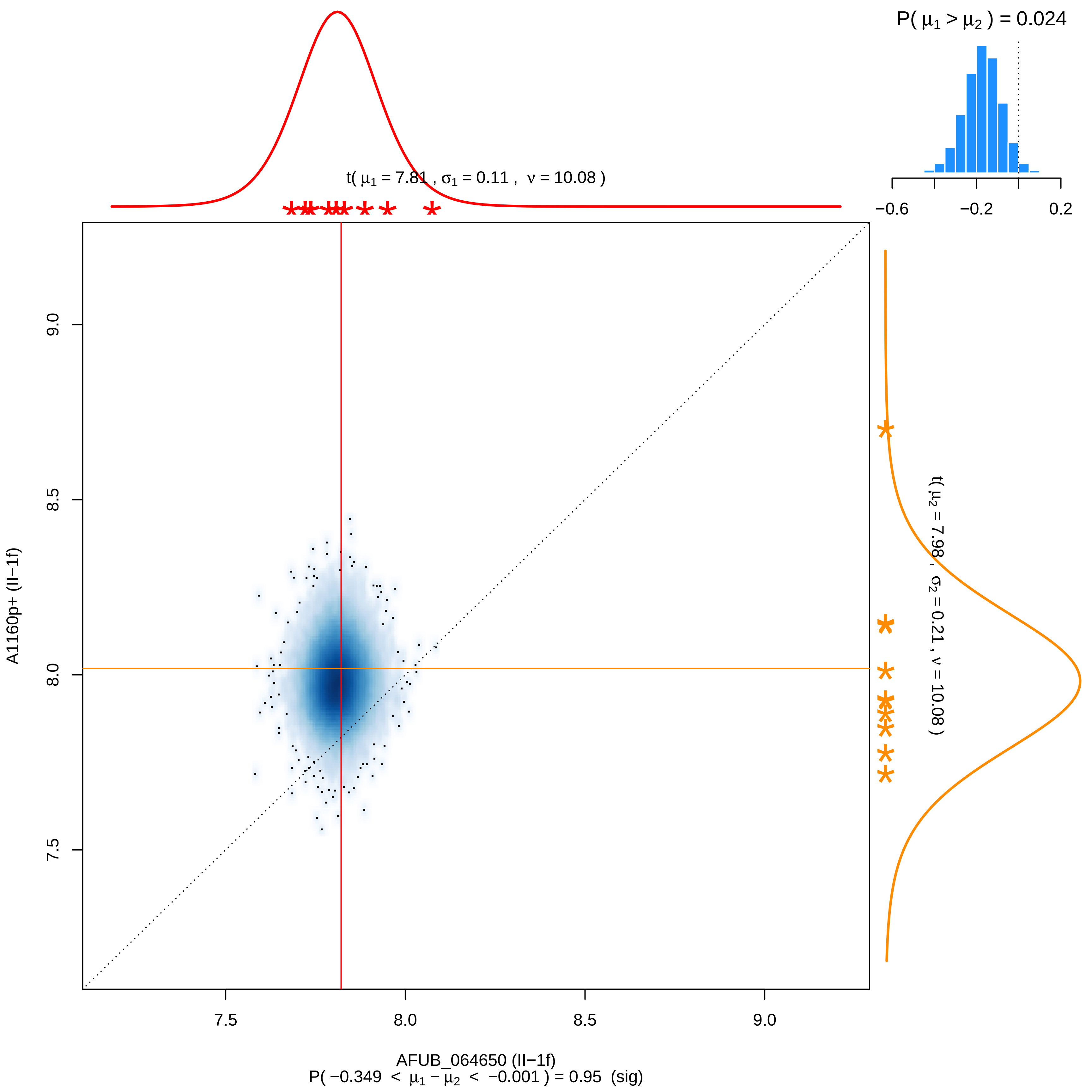

P( μ1 > μ2 ) = 0.024
−0.6
−0.2
0.2
t( μ1 = 7.81 , σ1 = 0.11 , ν = 10.08 )
t( μ2 = 7.98 , σ2 = 0.21 , ν = 10.08 )
9.0
8.5
8.0
7.5
A1160p+ (II−1f)
7.5
8.0
8.5
9.0
AFUB_064650 (II−1f)
P( −0.349 < μ1 − μ2 < −0.001 ) = 0.95 (sig)

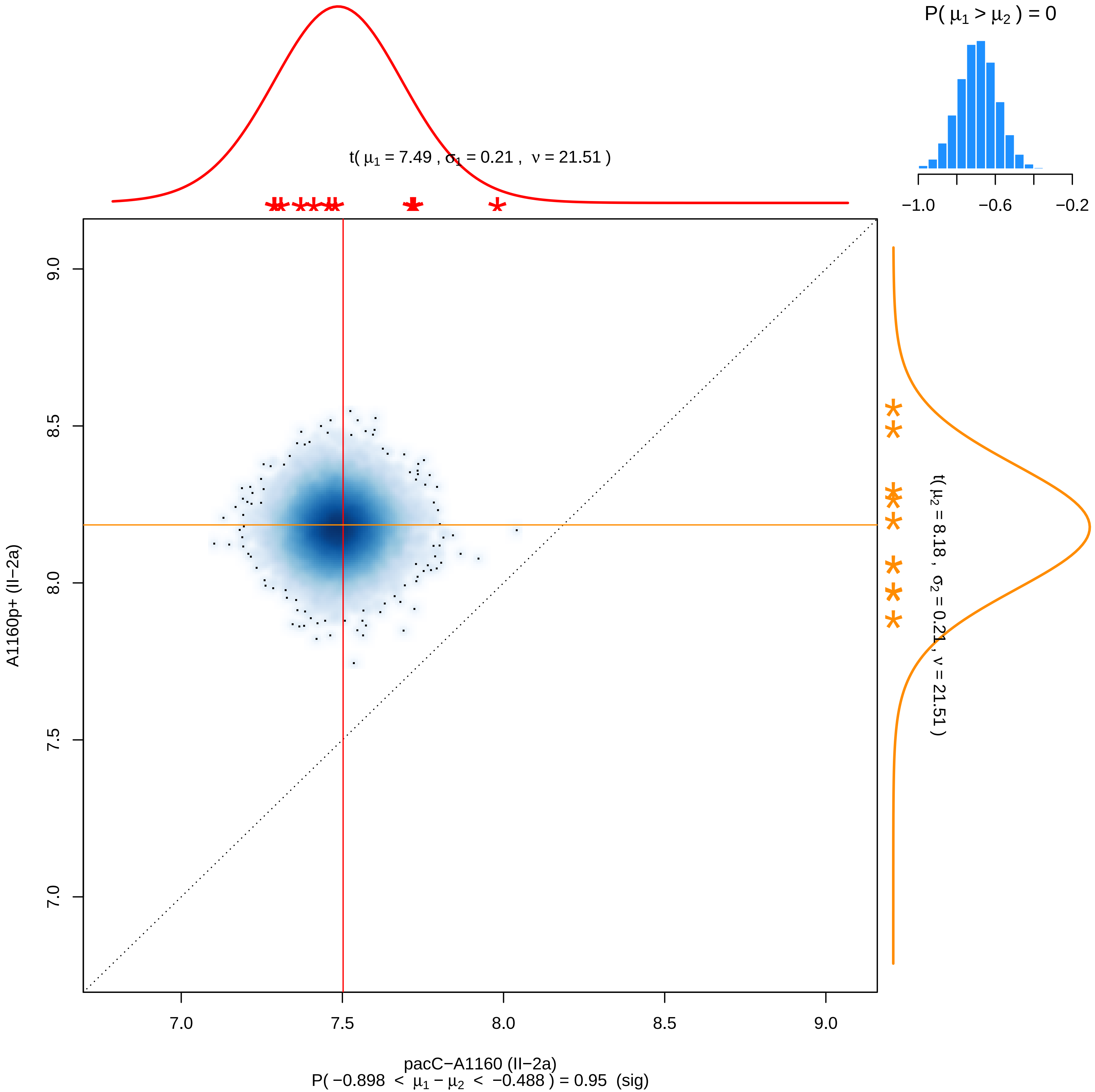

P( μ1 > μ2 ) = 0
−1.0
−0.6
−0.2
t( μ1 = 7.49 , σ1 = 0.21 , ν = 21.51 )
t( μ2 = 8.18 , σ2 = 0.21 , ν = 21.51 )
9.0
8.5
8.0
7.5
7.0
A1160p+ (II−2a)
7.0
7.5
8.0
8.5
9.0
pacC−A1160 (II−2a)
P( −0.898 < μ1 − μ2 < −0.488 ) = 0.95 (sig)

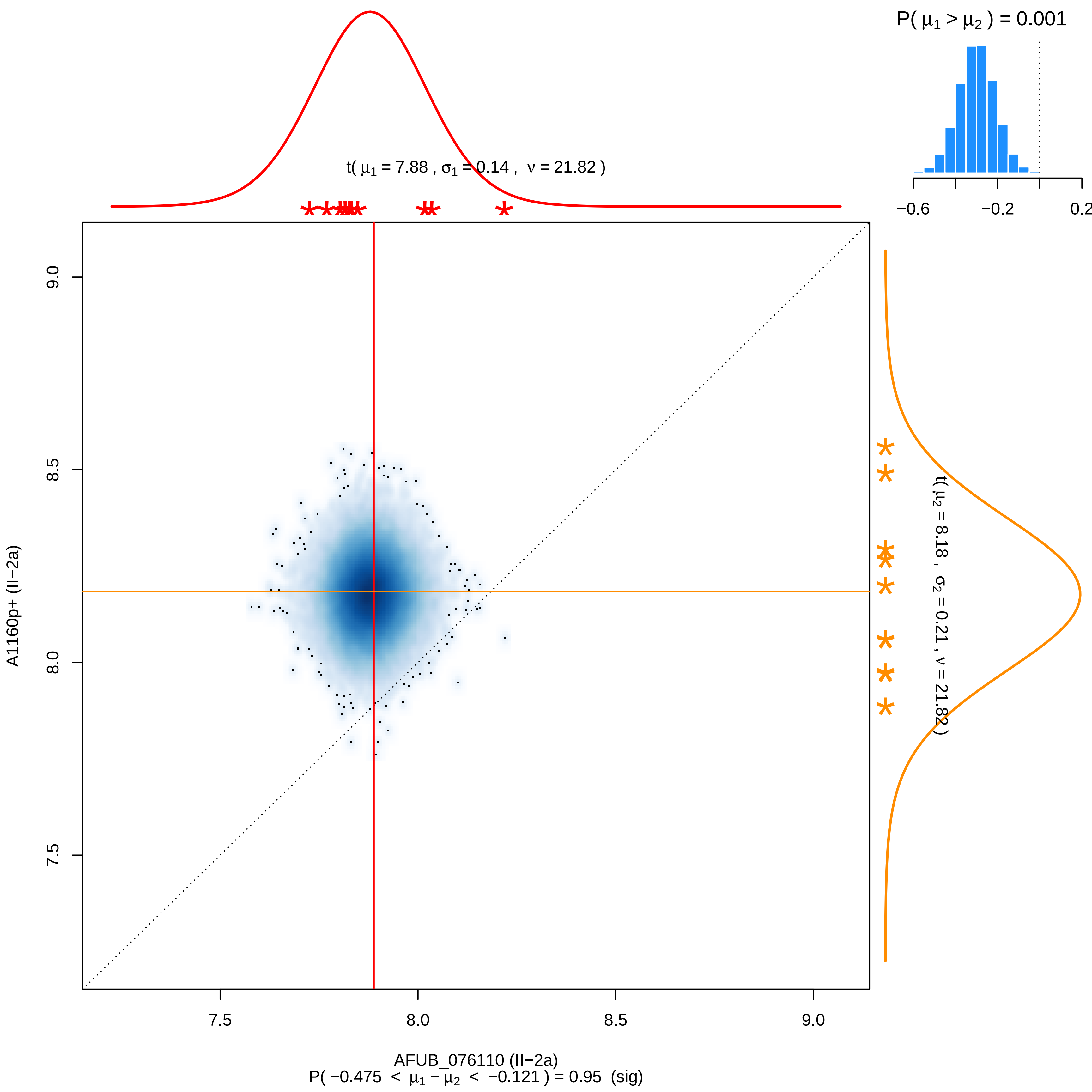

P( μ1 > μ2 ) = 0.001
−0.6
−0.2
0.2
t( μ1 = 7.88 , σ1 = 0.14 , ν = 21.82 )
t( μ2 = 8.18 , σ2 = 0.21 , ν = 21.82 )
9.0
8.5
8.0
7.5
A1160p+ (II−2a)
7.5
8.0
8.5
9.0
AFUB_076110 (II−2a)
P( −0.475 < μ1 − μ2 < −0.121 ) = 0.95 (sig)

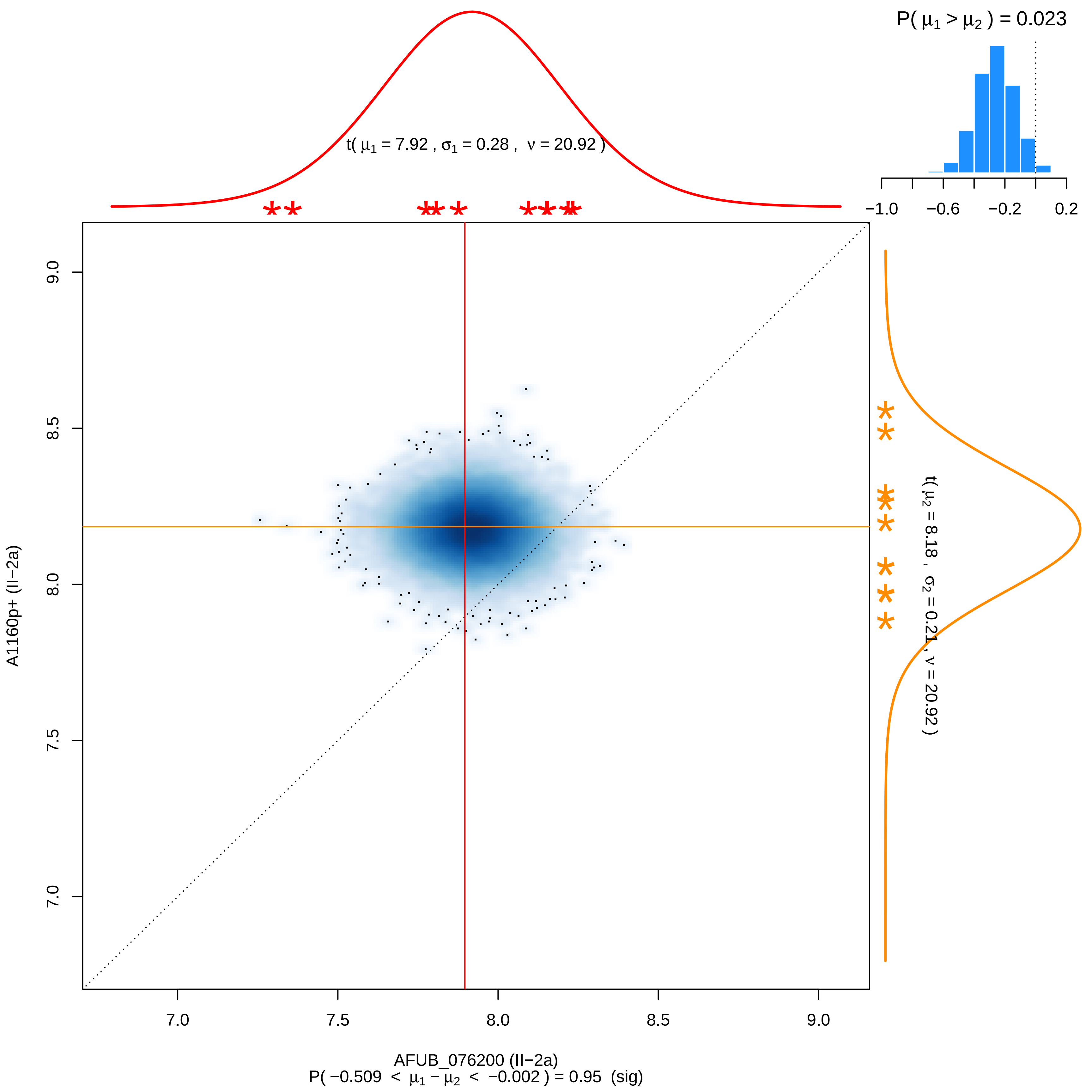

P( μ1 > μ2 ) = 0.023
−1.0
−0.6
−0.2
0.2
t( μ1 = 7.92 , σ1 = 0.28 , ν = 20.92 )
9.0
8.5
8.0
7.5
7.0
A1160p+ (II−2a)
t( μ2 = 8.18 , σ2 = 0.21 , ν = 20.92 )
7.0
7.5
8.0
8.5
9.0
AFUB_076200 (II−2a)
P( −0.509 < μ1 − μ2 < −0.002 ) = 0.95 (sig)

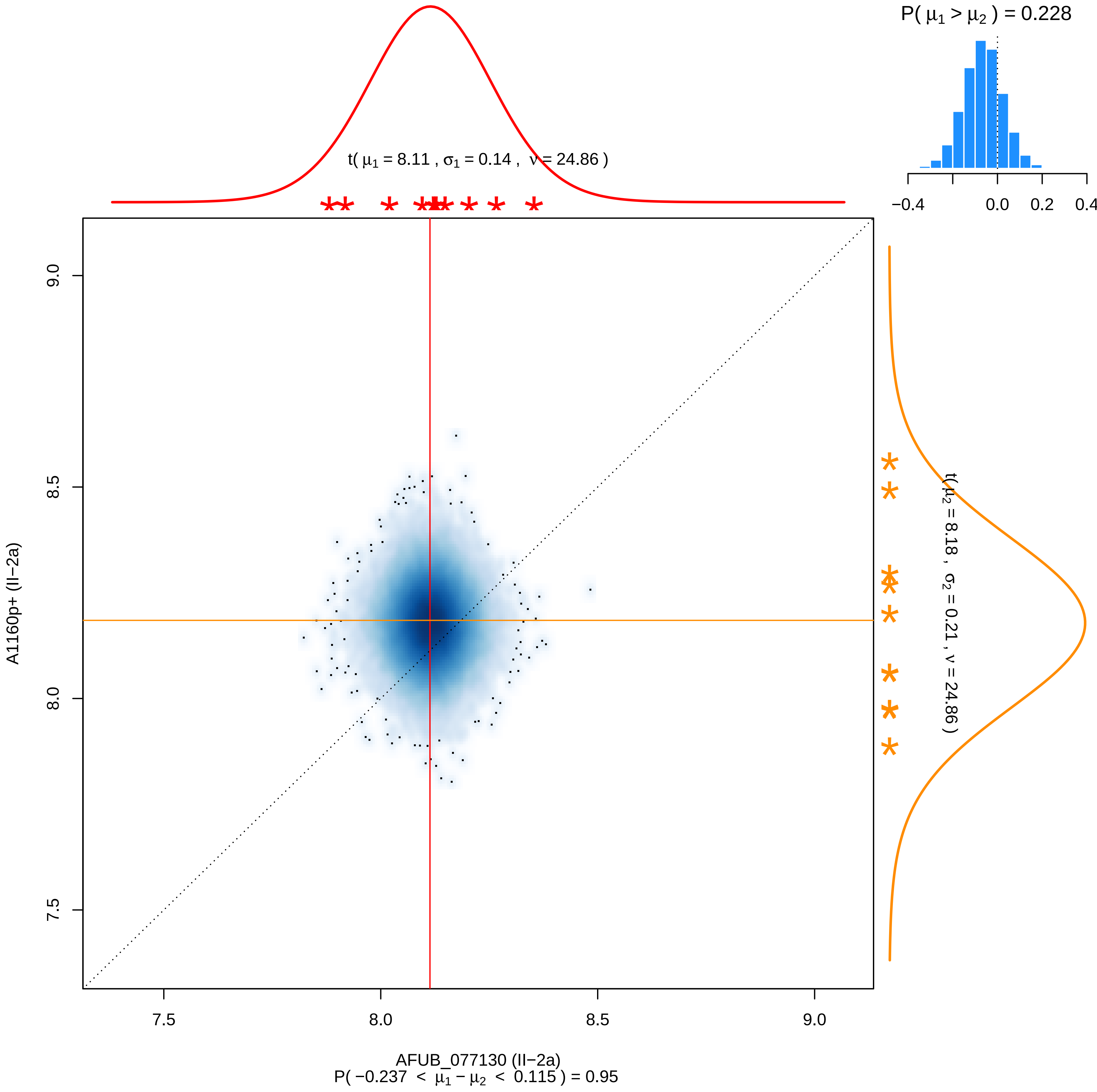

P( μ1 > μ2 ) = 0.228
−0.4
0.0
0.2
0.4
t( μ1 = 8.11 , σ1 = 0.14 , ν = 24.86 )
t( μ2 = 8.18 , σ2 = 0.21 , ν = 24.86 )
9.0
8.5
8.0
7.5
A1160p+ (II−2a)
7.5
8.0
8.5
9.0
AFUB_077130 (II−2a)
P( −0.237 < μ1 − μ2 < 0.115 ) = 0.95

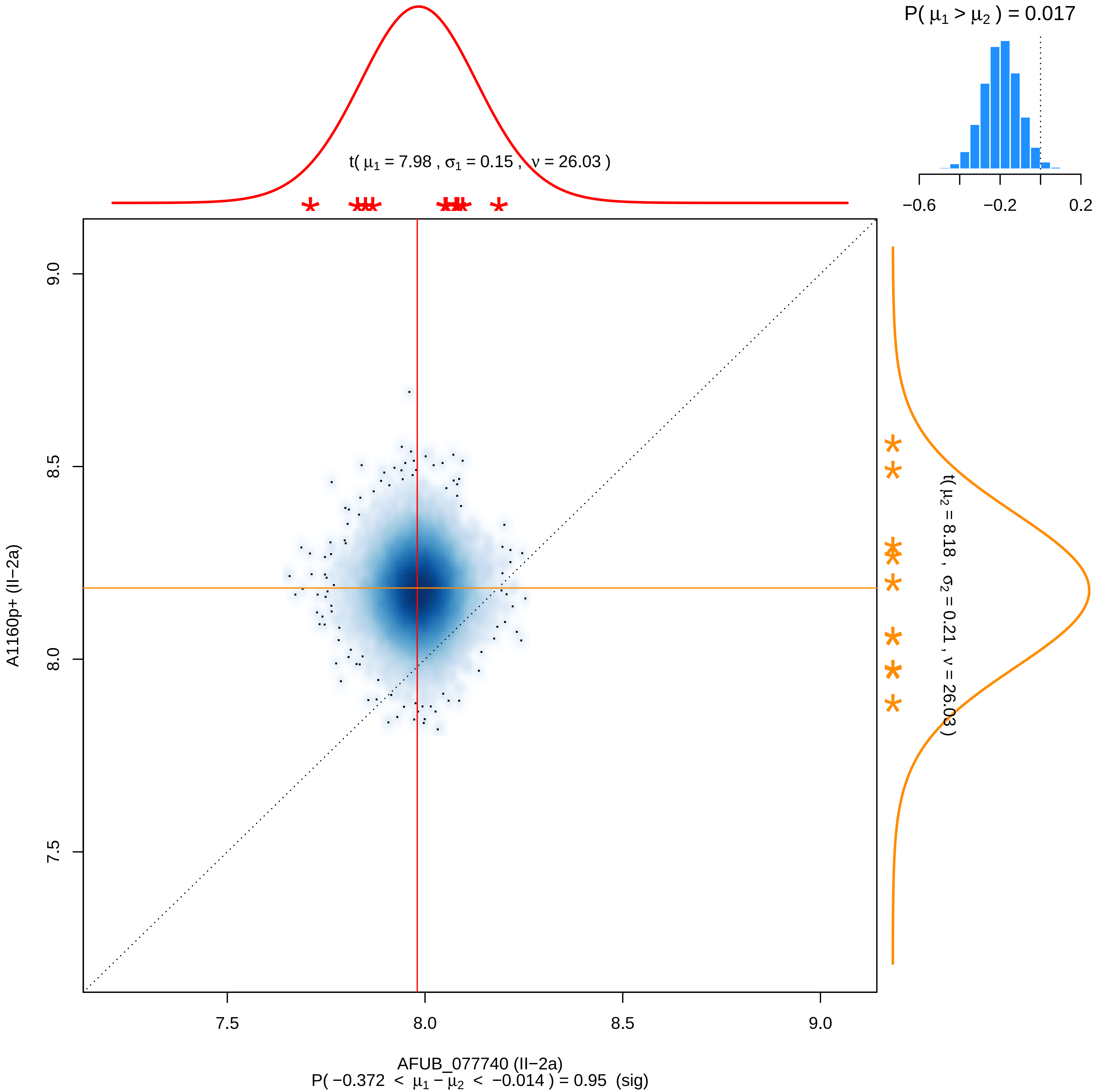

P( μ1 > μ2 ) = 0.017
−0.6
−0.2
0.2
t( μ1 = 7.98 , σ1 = 0.15 , ν = 26.03 )
t( μ2 = 8.18 , σ2 = 0.21 , ν = 26.03 )
9.0
8.5
8.0
7.5
A1160p+ (II−2a)
7.5
8.0
8.5
9.0
AFUB_077740 (II−2a)
P( −0.372 < μ1 − μ2 < −0.014 ) = 0.95 (sig)

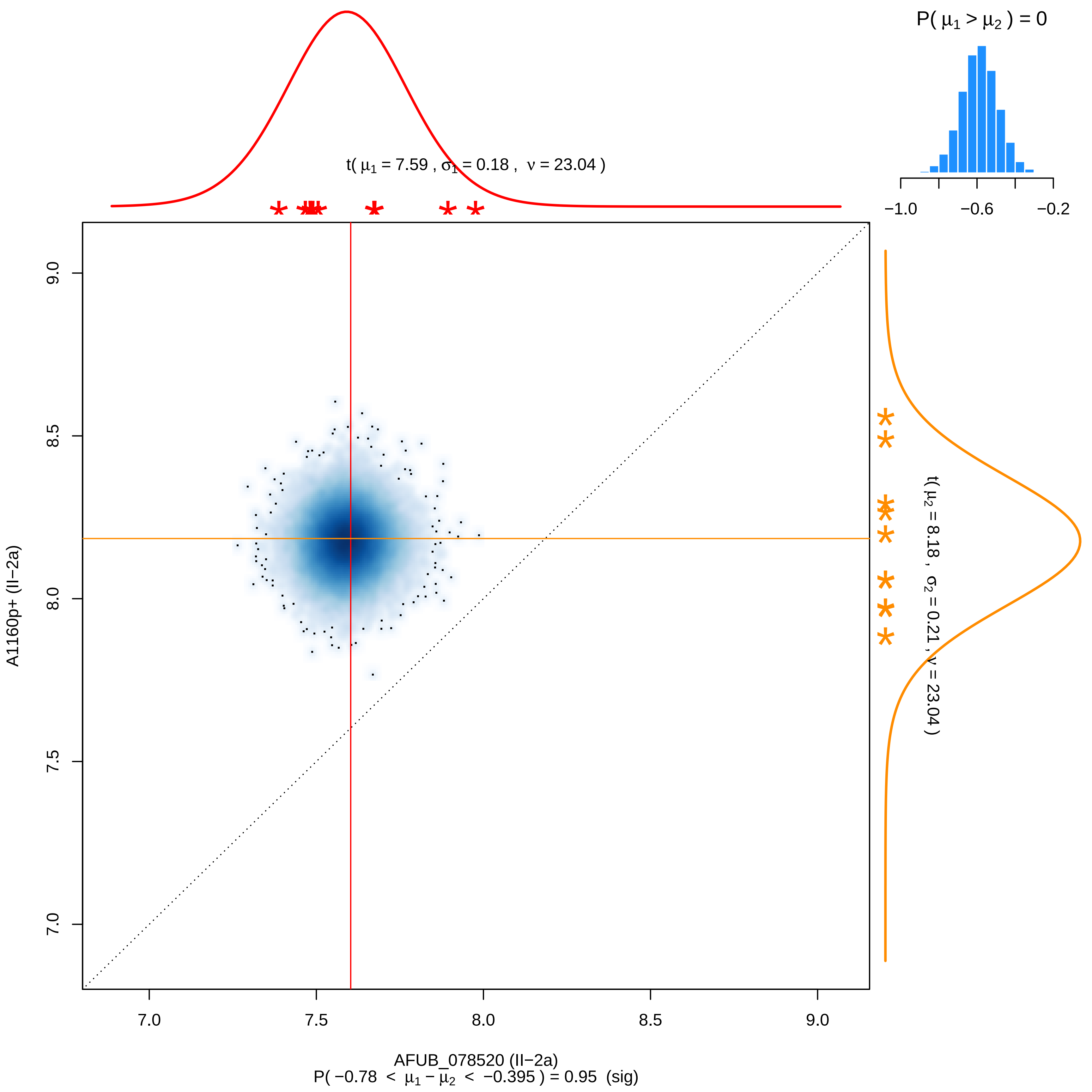

P( μ1 > μ2 ) = 0
−1.0
−0.6
−0.2
t( μ1 = 7.59 , σ1 = 0.18 , ν = 23.04 )
t( μ2 = 8.18 , σ2 = 0.21 , ν = 23.04 )
A1160p+ (II−2a)
7.0
7.5
8.0
8.5
9.0
AFUB_078520 (II−2a)
P( −0.78 < μ1 − μ2 < −0.395 ) = 0.95 (sig)

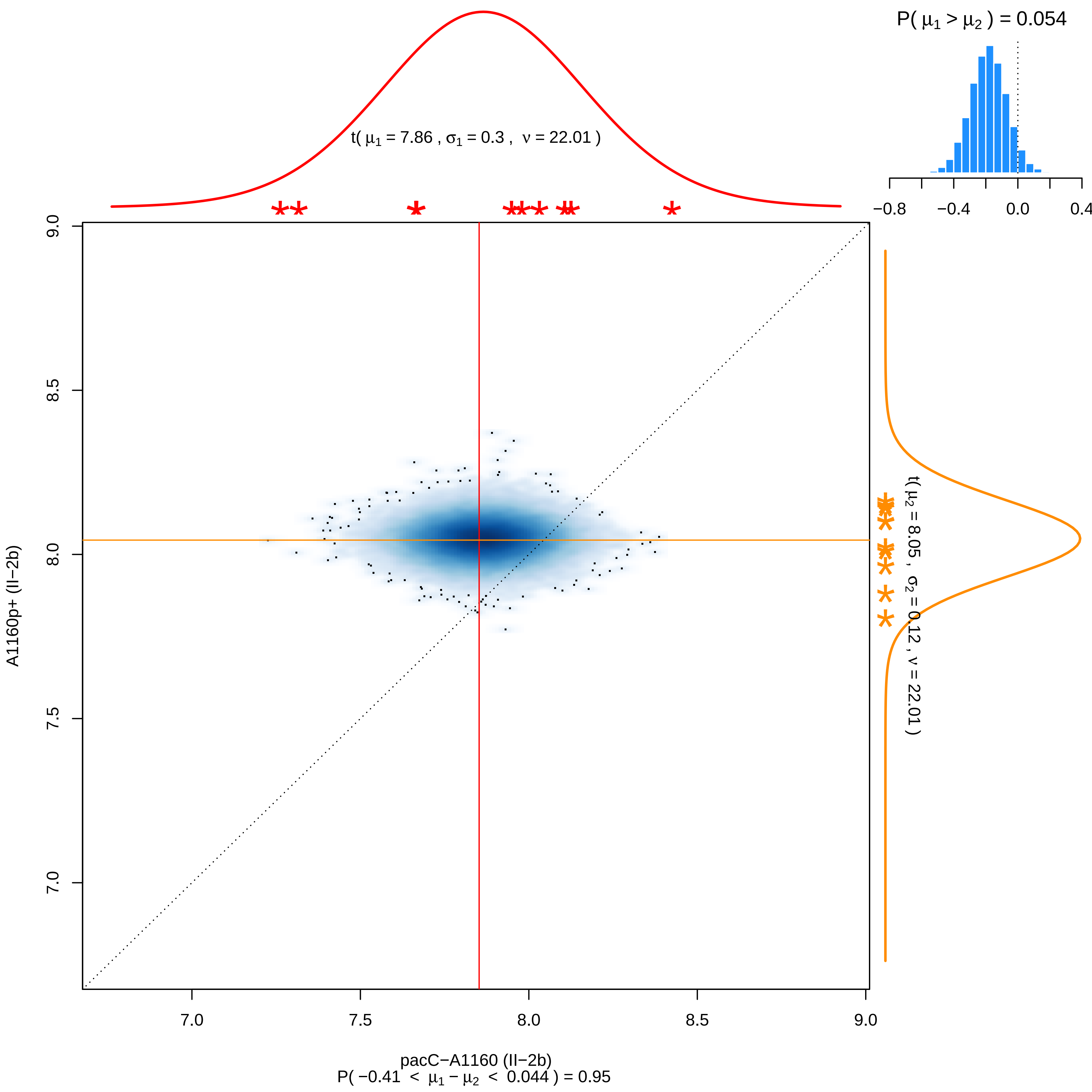

P( μ1 > μ2 ) = 0.054
−0.8
−0.4
0.0
0.4
t( μ1 = 7.86 , σ1 = 0.3 , ν = 22.01 )
t( μ2 = 8.05 , σ2 = 0.12 , ν = 22.01 )
A1160p+ (II−2b)
9.0
8.5
8.0
7.5
7.0
7.0
7.5
8.0
8.5
9.0
pacC−A1160 (II−2b)
P( −0.41 < μ1 − μ2 < 0.044 ) = 0.95

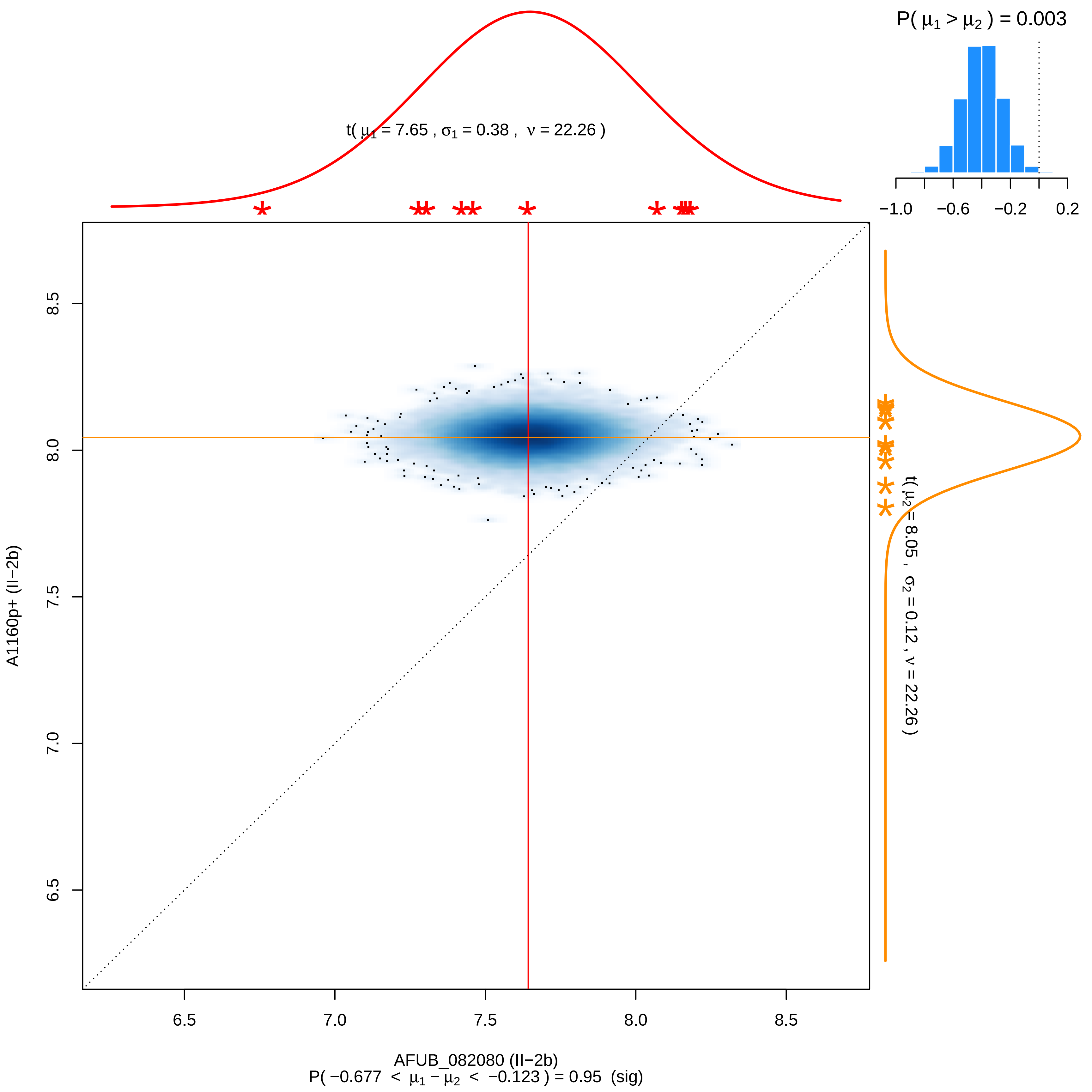

P( μ1 > μ2 ) = 0.003
−1.0 −0.6 −0.2 0.2
t( μ1 = 7.65 , σ1 = 0.38 , ν = 22.26 )
t( μ2 = 8.05 , σ2 = 0.12 , ν = 22.26 )
8.5
8.0
7.5
7.0
6.5
A1160p+ (II−2b)
6.5 7.0 7.5 8.0 8.5
AFUB_082080 (II−2b)
P( −0.677 < μ1 − μ2 < −0.123 ) = 0.95 (sig)

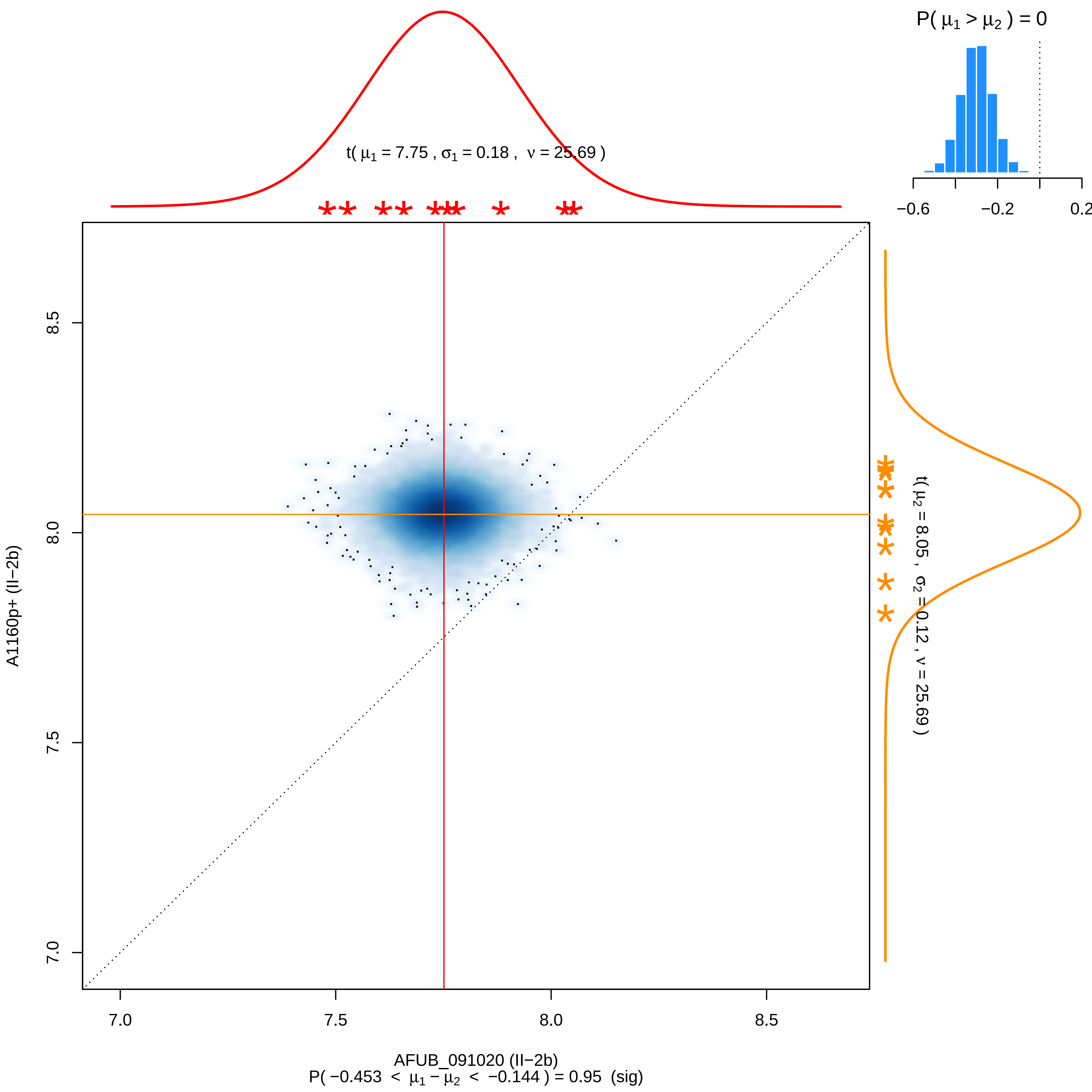

P( μ1 > μ2 ) = 0
−0.6
−0.2
0.2
t( μ1 = 7.75 , σ1 = 0.18 , ν = 25.69 )
t( μ2 = 8.05 , σ2 = 0.12 , ν = 25.69 )
8.5
8.0
7.5
7.0
A1160p+ (II−2b)
7.0
7.5
8.0
8.5
AFUB_091020 (II−2b)
P( −0.453 < μ1 − μ2 < −0.144 ) = 0.95 (sig)

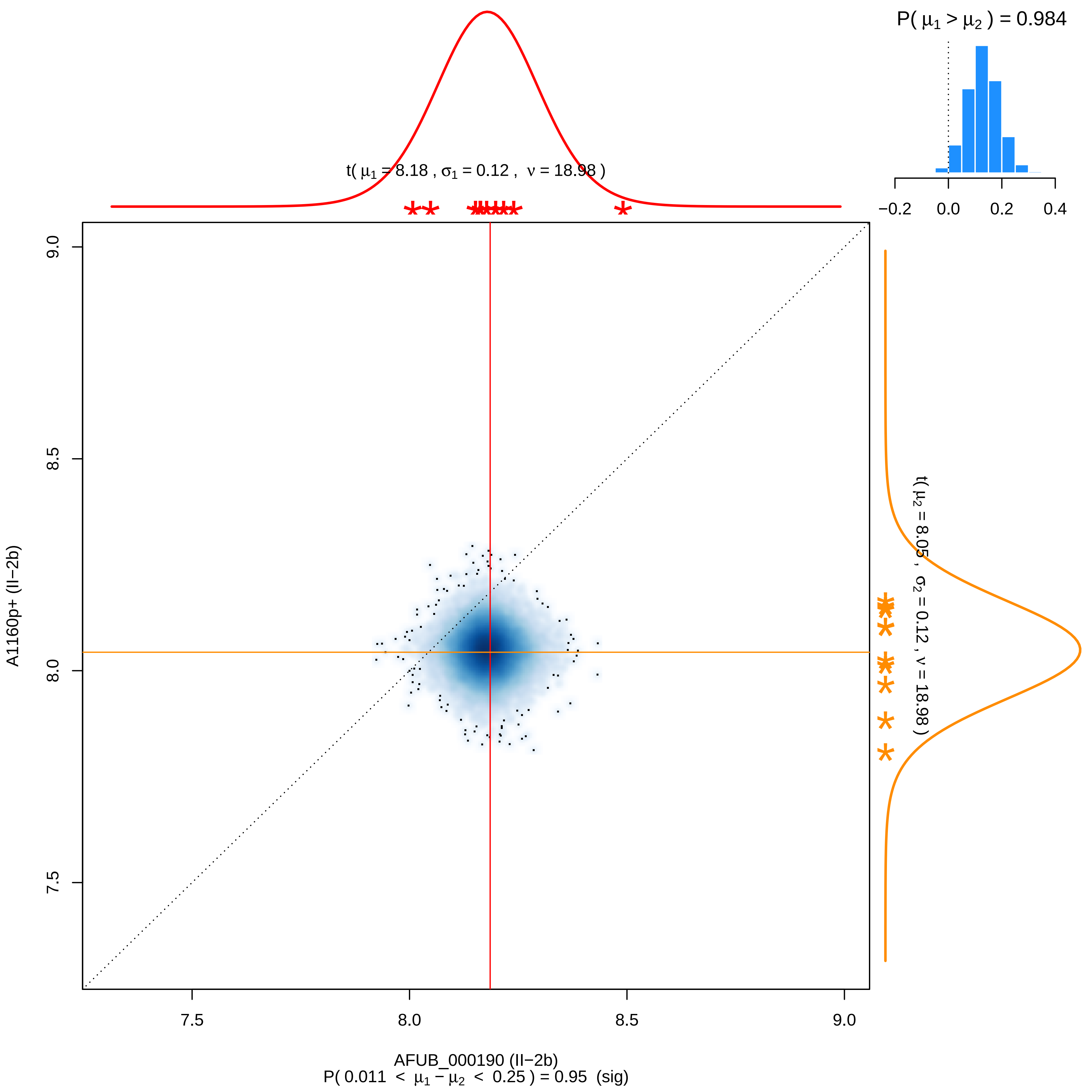

P( μ1 > μ2 ) = 0.984
−0.2
0.0
0.2
0.4
t( μ1 = 8.18 , σ1 = 0.12 , ν = 18.98 )
t( μ2 = 8.05 , σ2 = 0.12 , ν = 18.98 )
A1160p+ (II−2b)
9.0
8.5
8.0
7.5
7.5
8.0
8.5
9.0
AFUB_000190 (II−2b)
P( 0.011 < μ1 − μ2 < 0.25 ) = 0.95 (sig)

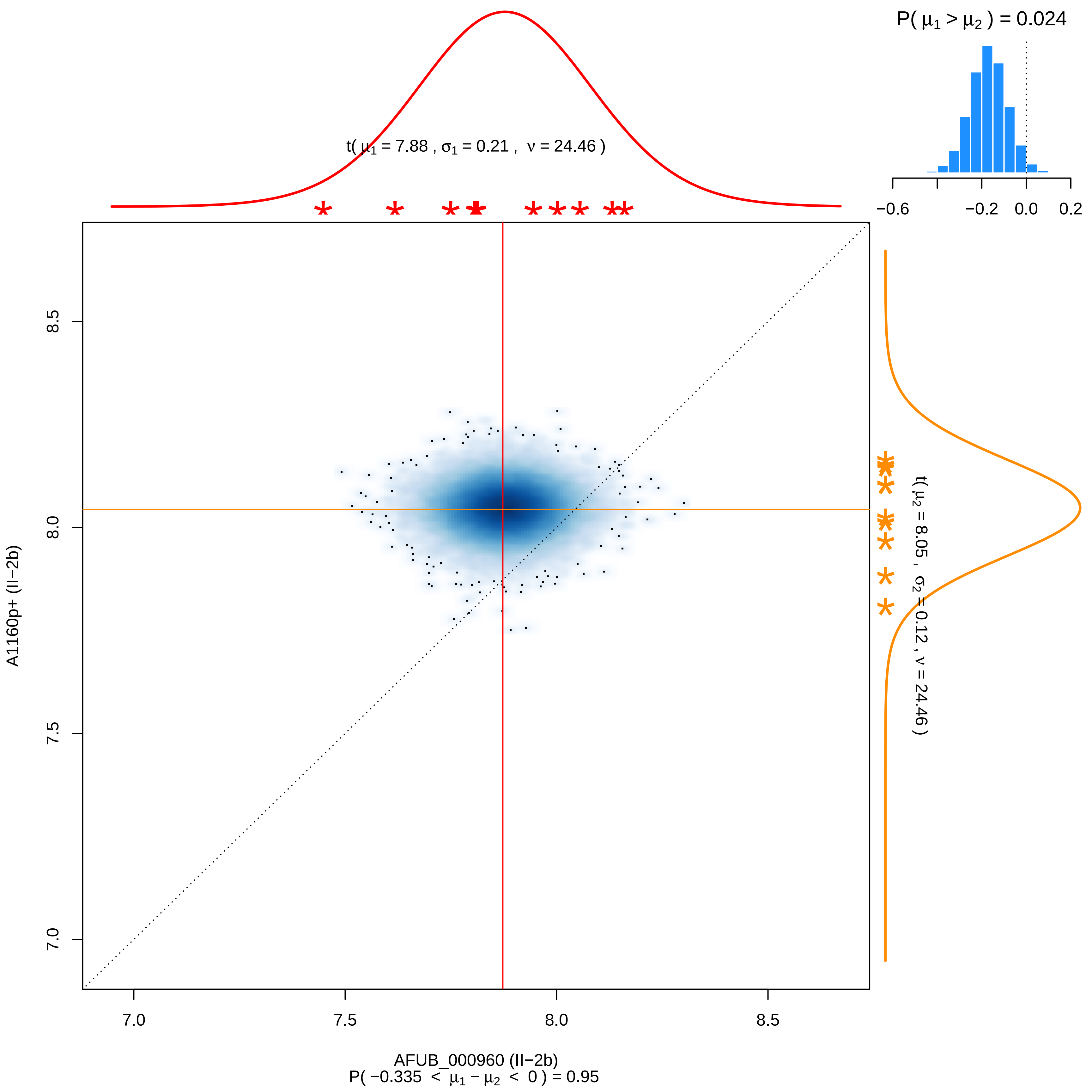

P( μ1 > μ2 ) = 0.024
−0.6
−0.2
0.0
0.2
t( μ1 = 7.88 , σ1 = 0.21 , ν = 24.46 )
t( μ2 = 8.05 , σ2 = 0.12 , ν = 24.46 )
A1160p+ (II−2b)
8.5
8.0
7.5
7.0
7.0
7.5
8.0
8.5
AFUB_000960 (II−2b)
P( −0.335 < μ1 − μ2 < 0 ) = 0.95

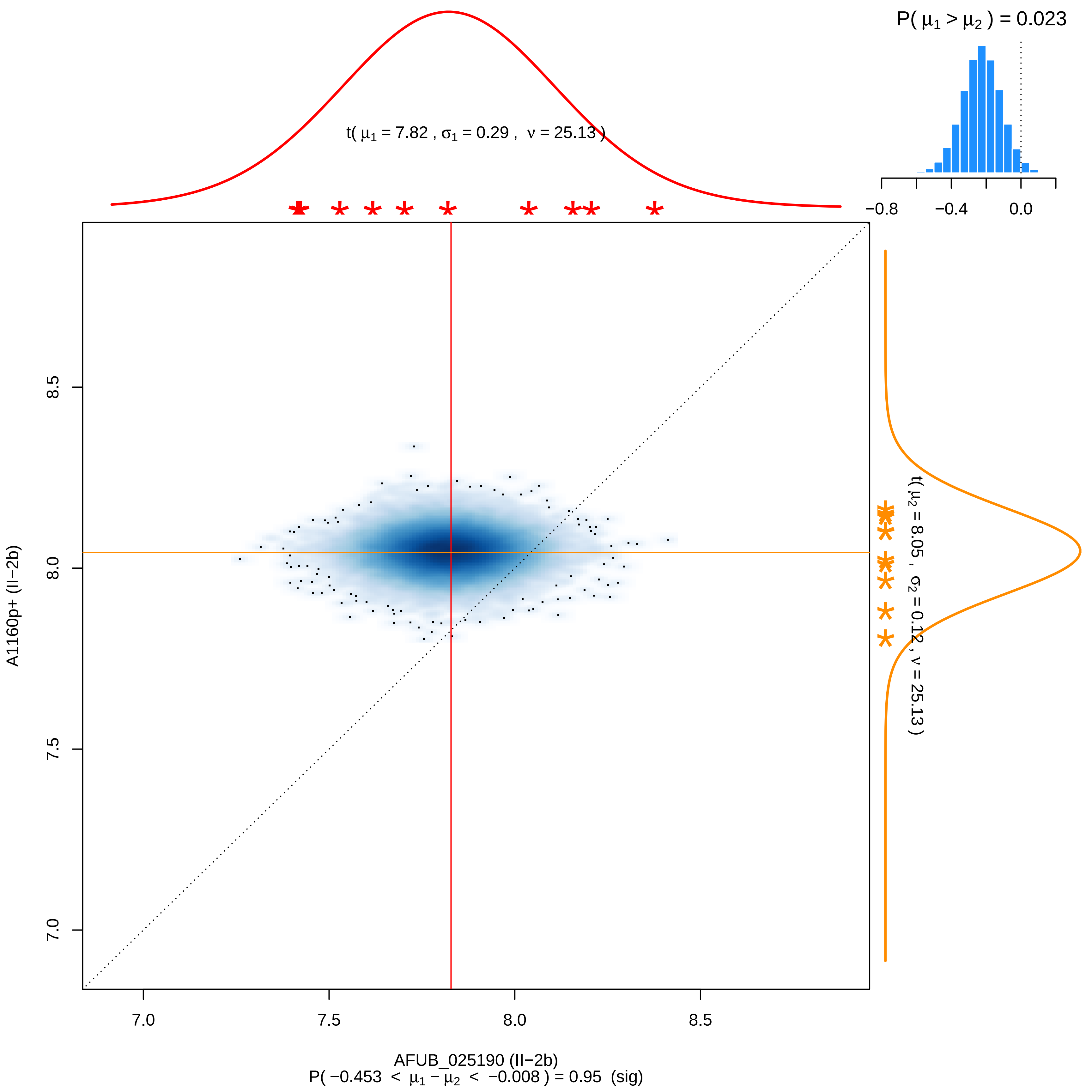

P( μ1 > μ2 ) = 0.023
t( μ1 = 7.82 , σ1 = 0.29 , ν = 25.13 )
−0.8
−0.4
0.0
t( μ2 = 8.05 , σ2 = 0.12 , ν = 25.13 )
A1160p+ (II−2b)
7.0
7.5
8.0
8.5
AFUB_025190 (II−2b)
P( −0.453 < μ1 − μ2 < −0.008 ) = 0.95 (sig)

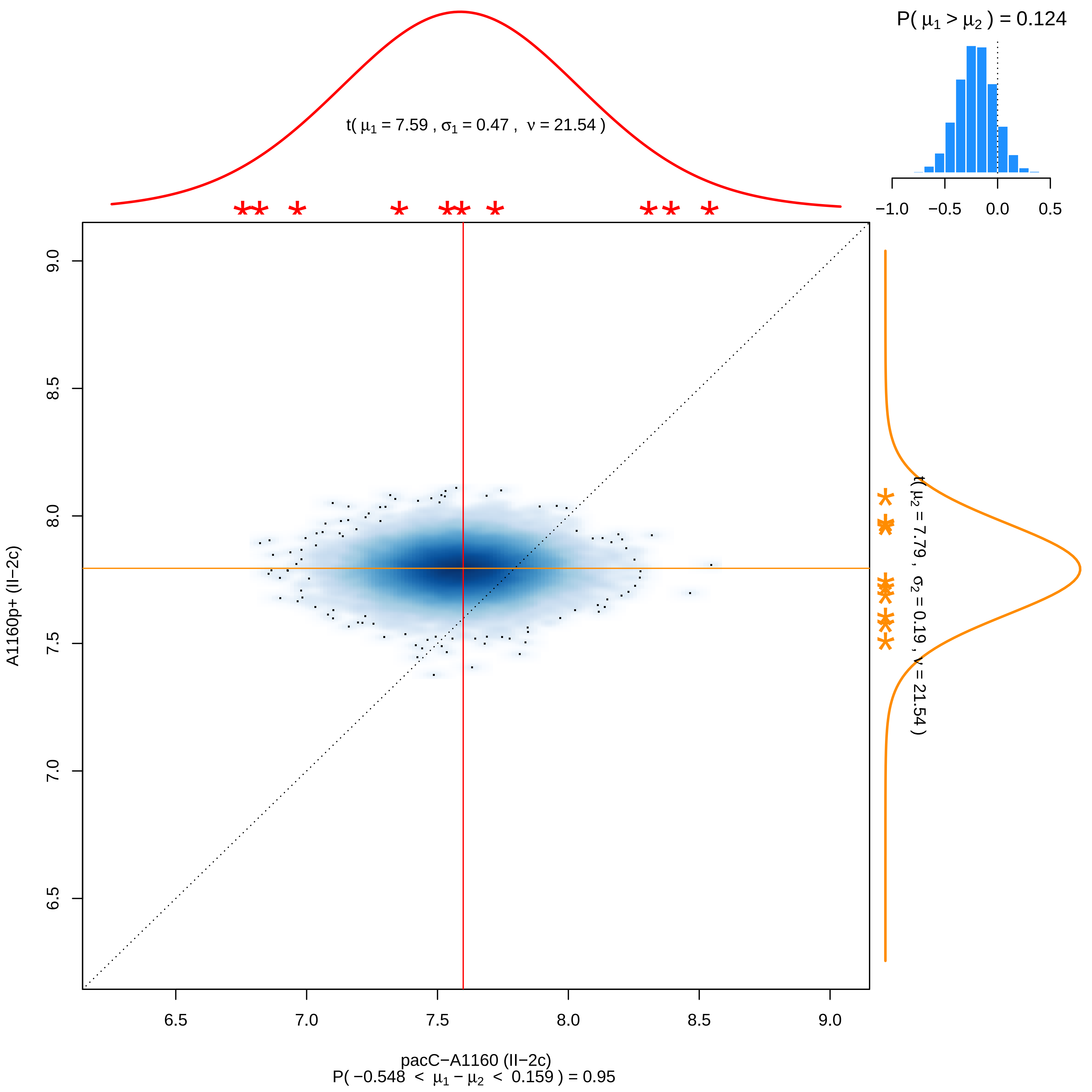

P( μ1 > μ2 ) = 0.124
−1.0 −0.5 0.0 0.5
t( μ1 = 7.59 , σ1 = 0.47 , ν = 21.54 )
t( μ2 = 7.79 , σ2 = 0.19 , ν = 21.54 )
9.0 8.5 8.0 7.5 7.0 6.5
A1160p+ (II−2c)
6.5 7.0 7.5 8.0 8.5 9.0
pacC−A1160 (II−2c)
P( −0.548 < μ1 − μ2 < 0.159 ) = 0.95

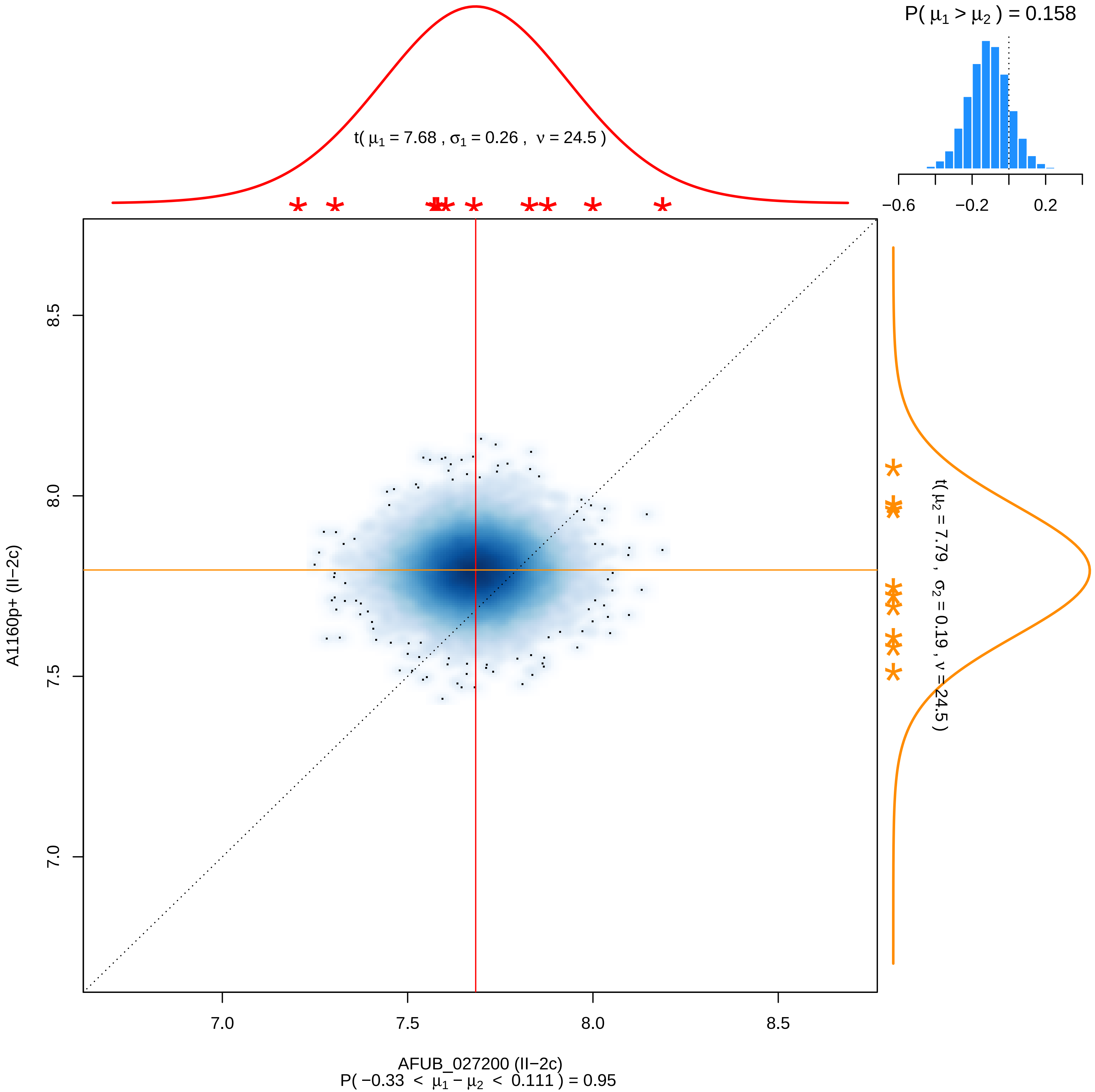

P( μ1 > μ2 ) = 0.158
−0.6
−0.2
0.2
t( μ1 = 7.68 , σ1 = 0.26 , ν = 24.5 )
t( μ2 = 7.79 , σ2 = 0.19 , ν = 24.5 )
8.5
8.0
7.5
7.0
A1160p+ (II−2c)
7.0
7.5
8.0
8.5
AFUB_027200 (II−2c)
P( −0.33 < μ1 − μ2 < 0.111 ) = 0.95

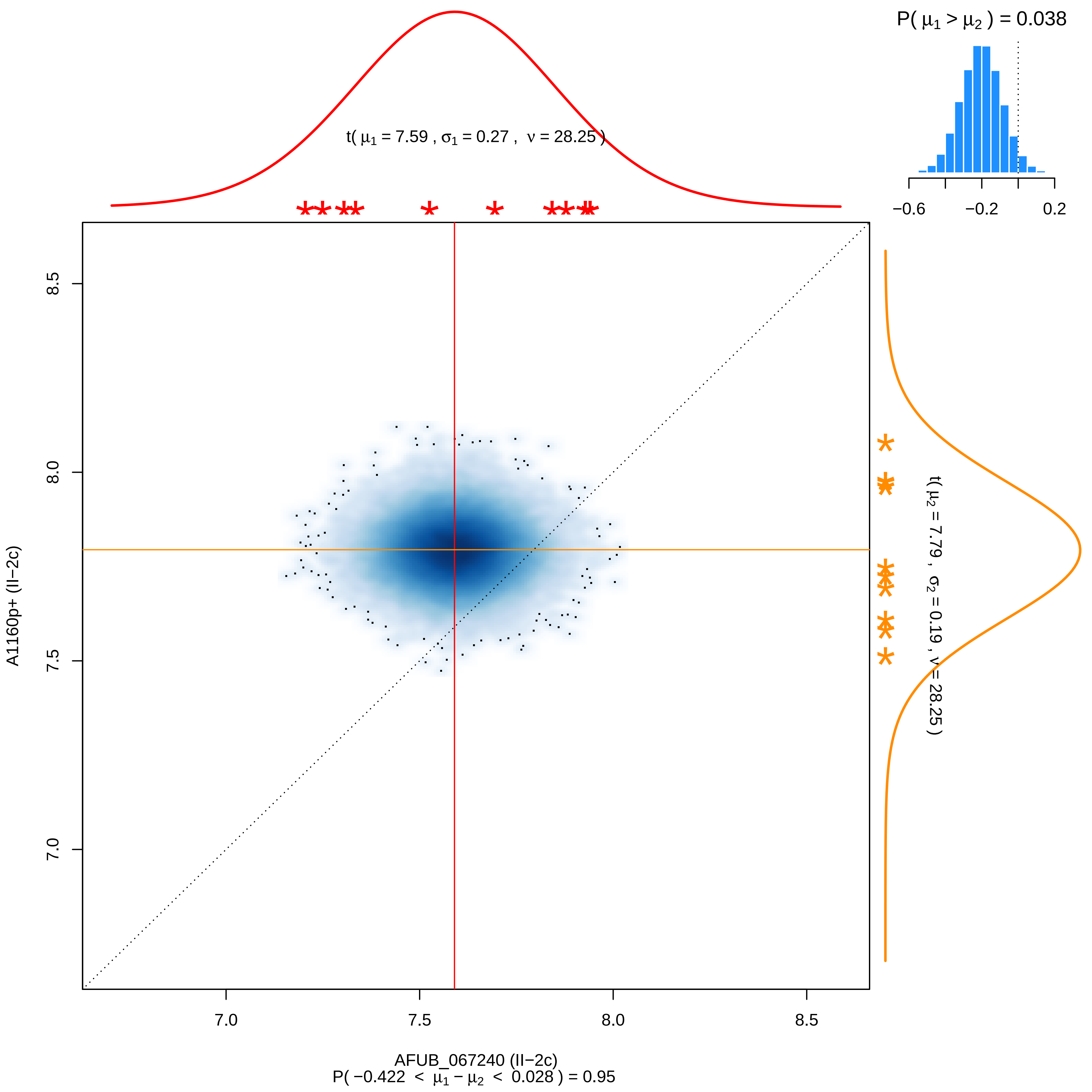

P( μ1 > μ2 ) = 0.038
−0.6
−0.2
0.2
t( μ1 = 7.59 , σ1 = 0.27 , ν = 28.25 )
t( μ2 = 7.79 , σ2 = 0.19 , ν = 28.25 )
8.5
8.0
7.5
7.0
A1160p+ (II−2c)
7.0
7.5
8.0
8.5
AFUB_067240 (II−2c)
P( −0.422 < μ1 − μ2 < 0.028 ) = 0.95

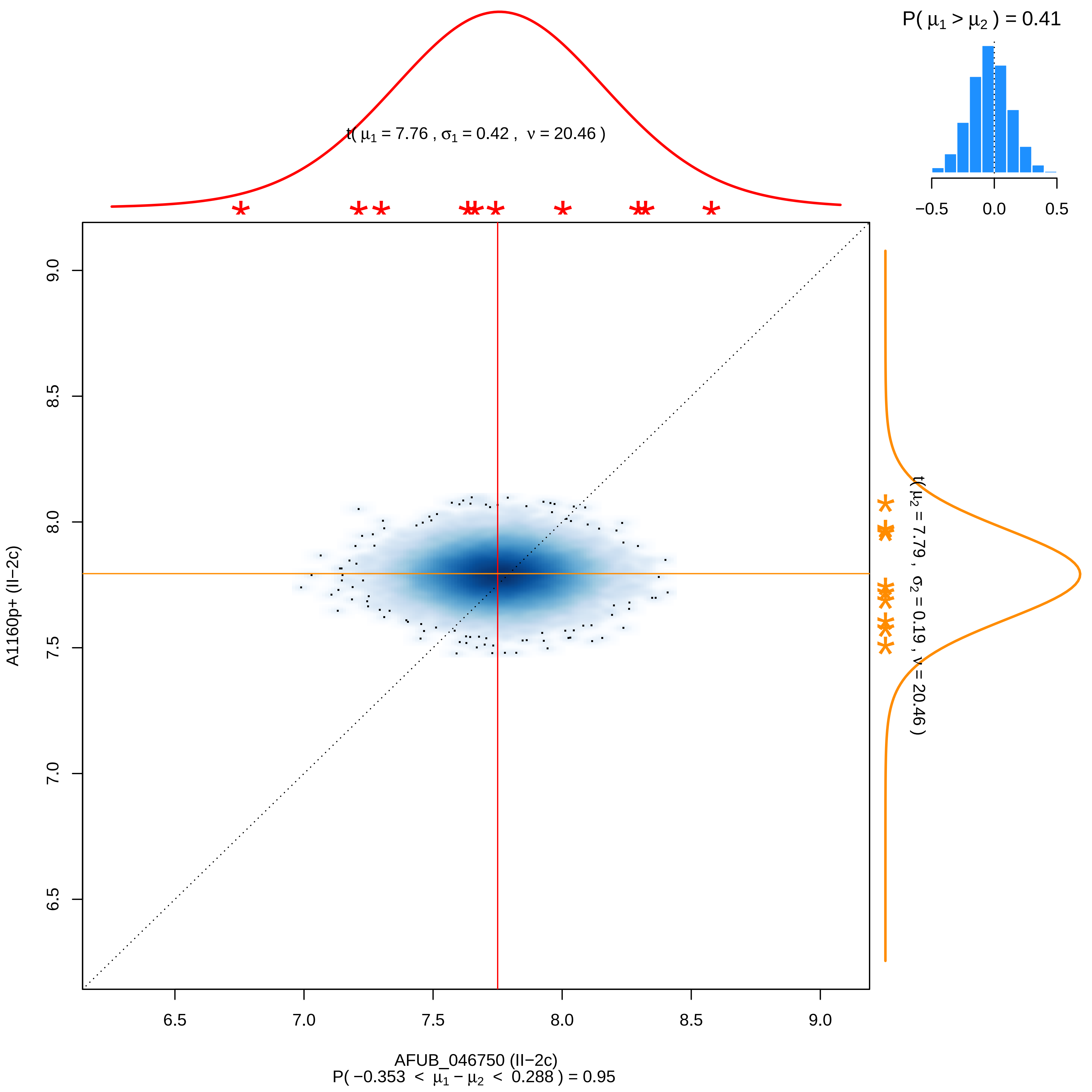

P( $\mu_1 > \mu_2$ ) = 0.41
−0.5
0.0
0.5
t( $\mu_1$ = 7.76 , $\sigma_1$ = 0.42 , $\nu$ = 20.46 )
t( $\mu_2$ = 7.79 , $\sigma_2$ = 0.19 , $\nu$ = 20.46 )
A1160p+ (II−2c)
6.5
7.0
7.5
8.0
8.5
9.0
AFUB_046750 (II−2c)
P( −0.353 < $\mu_1 - \mu_2$ < 0.288 ) = 0.95

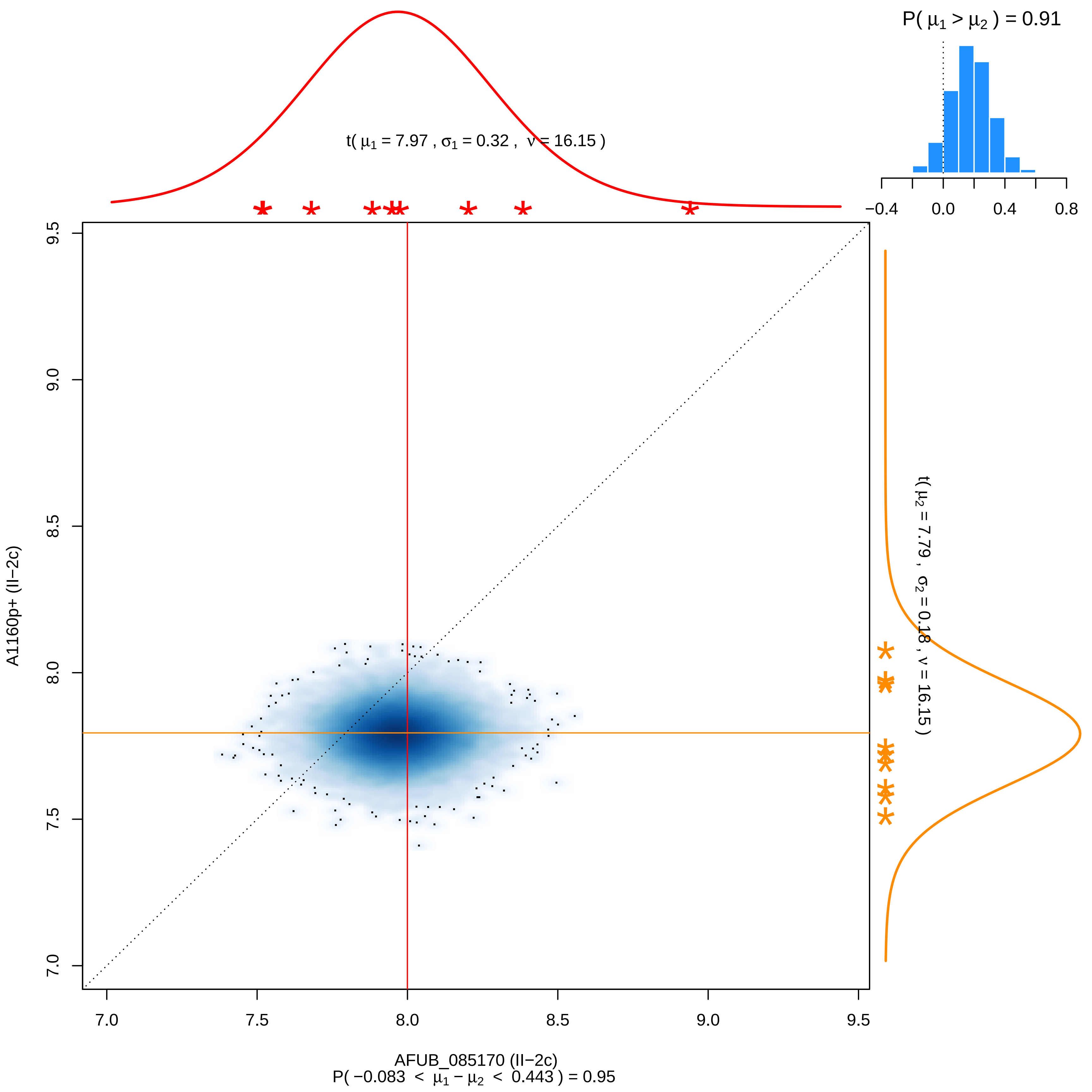

P( $\mu_1 > \mu_2$ ) = 0.91
t( $\mu_1$ = 7.97 , $\sigma_1$ = 0.32 , $\nu$ = 16.15 )
t( $\mu_2$ = 7.79 , $\sigma_2$ = 0.18 , $\nu$ = 16.15 )
A1160p+ (II−2c)
AFUB_085170 (II−2c)
P( −0.083 < $\mu_1 - \mu_2$ < 0.443 ) = 0.95

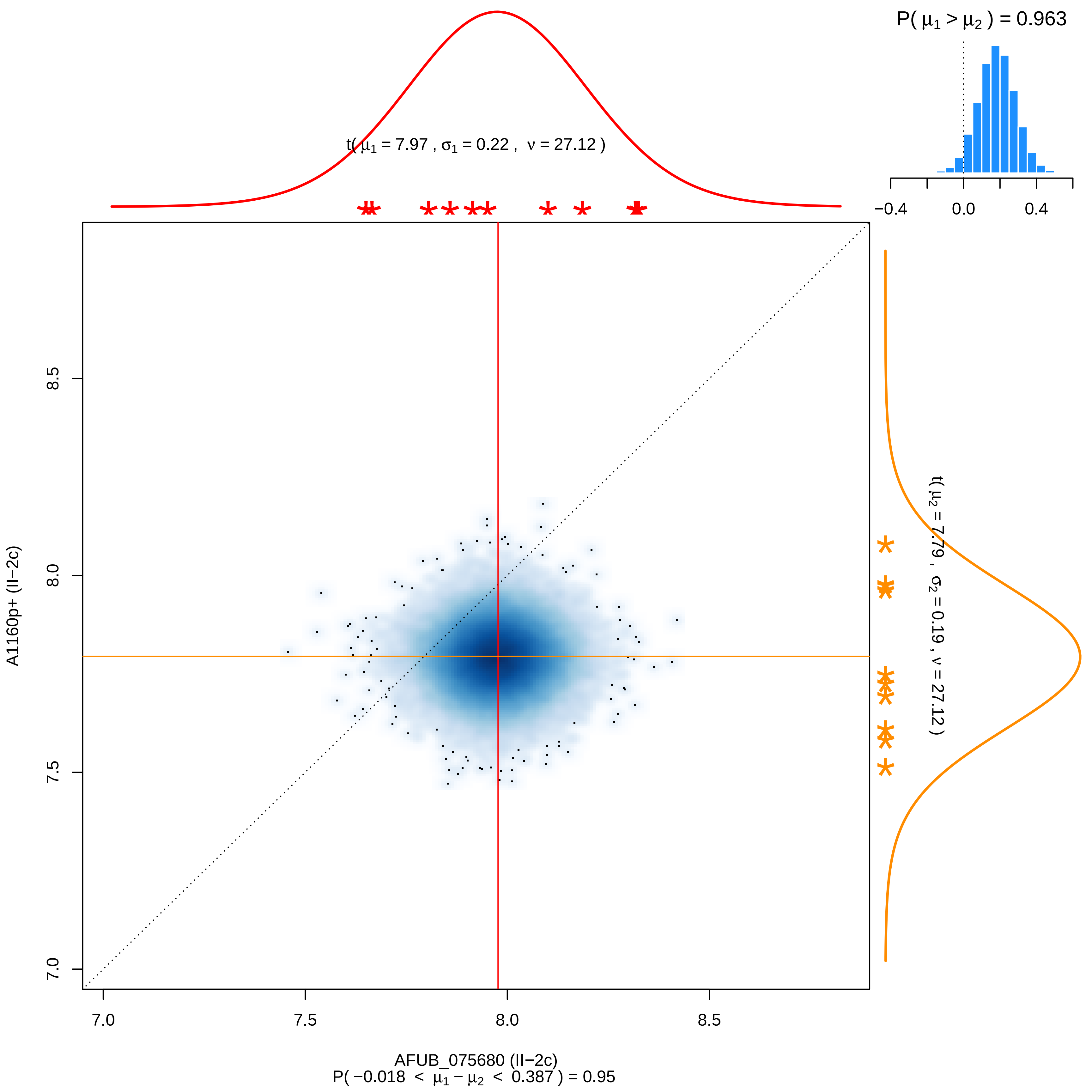

P( μ1 > μ2 ) = 0.963
−0.4
0.0
0.4
t( μ1 = 7.97 , σ1 = 0.22 , ν = 27.12 )
t( μ2 = 7.79 , σ2 = 0.19 , ν = 27.12 )
A1160p+ (II−2c)
8.5
8.0
7.5
7.0
7.0
7.5
8.0
8.5
AFUB_075680 (II−2c)
P( −0.018 < μ1 − μ2 < 0.387 ) = 0.95

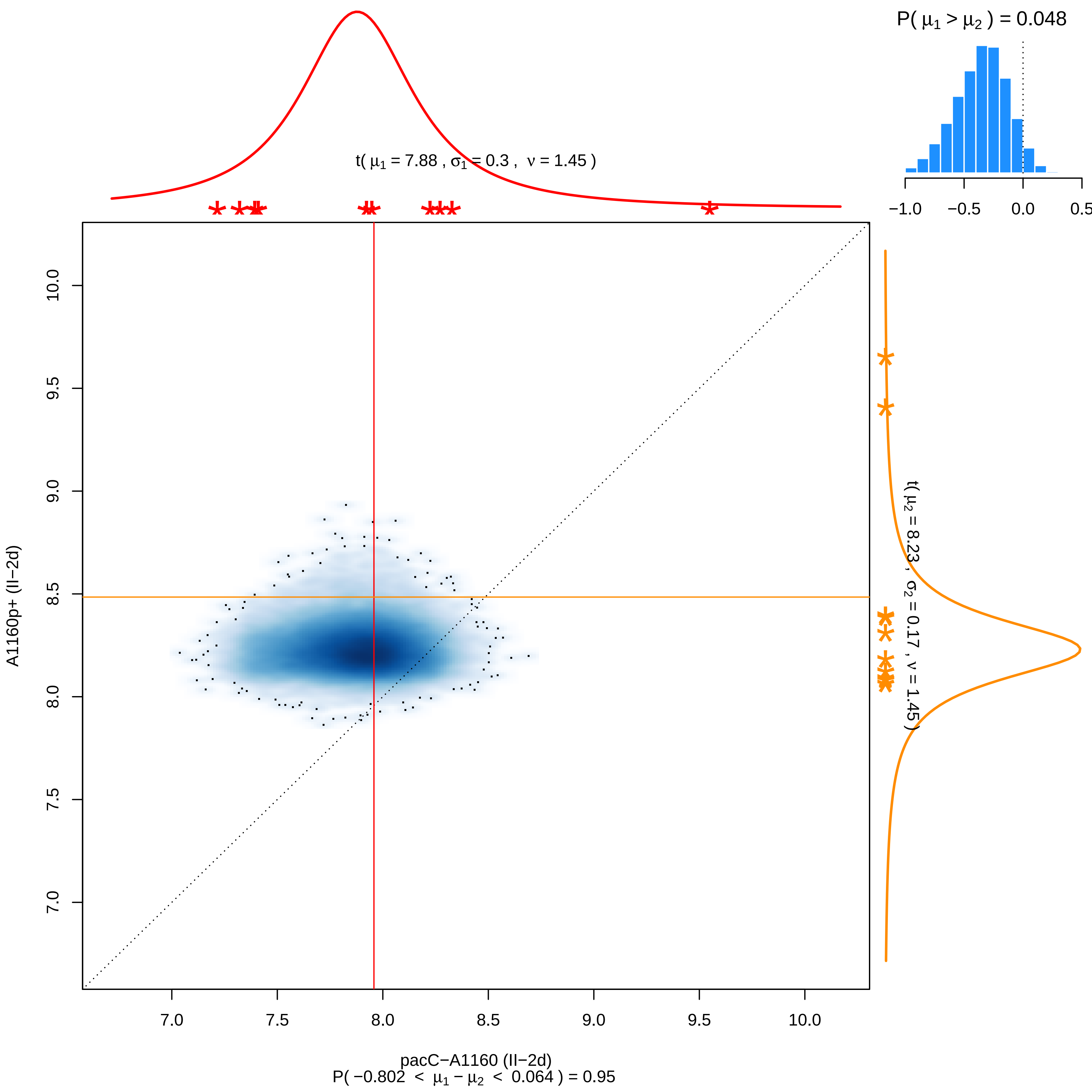

P( μ1 > μ2 ) = 0.048
−1.0
−0.5
0.0
0.5
t( μ1 = 7.88 , σ1 = 0.3 , ν = 1.45 )
t( μ2 = 8.23 , σ2 = 0.17 , ν = 1.45 )
A1160p+ (II−2d)
10.0
9.5
9.0
8.5
8.0
7.5
7.0
7.0
7.5
8.0
8.5
9.0
9.5
10.0
pacC−A1160 (II−2d)
P( −0.802 < μ1 − μ2 < 0.064 ) = 0.95

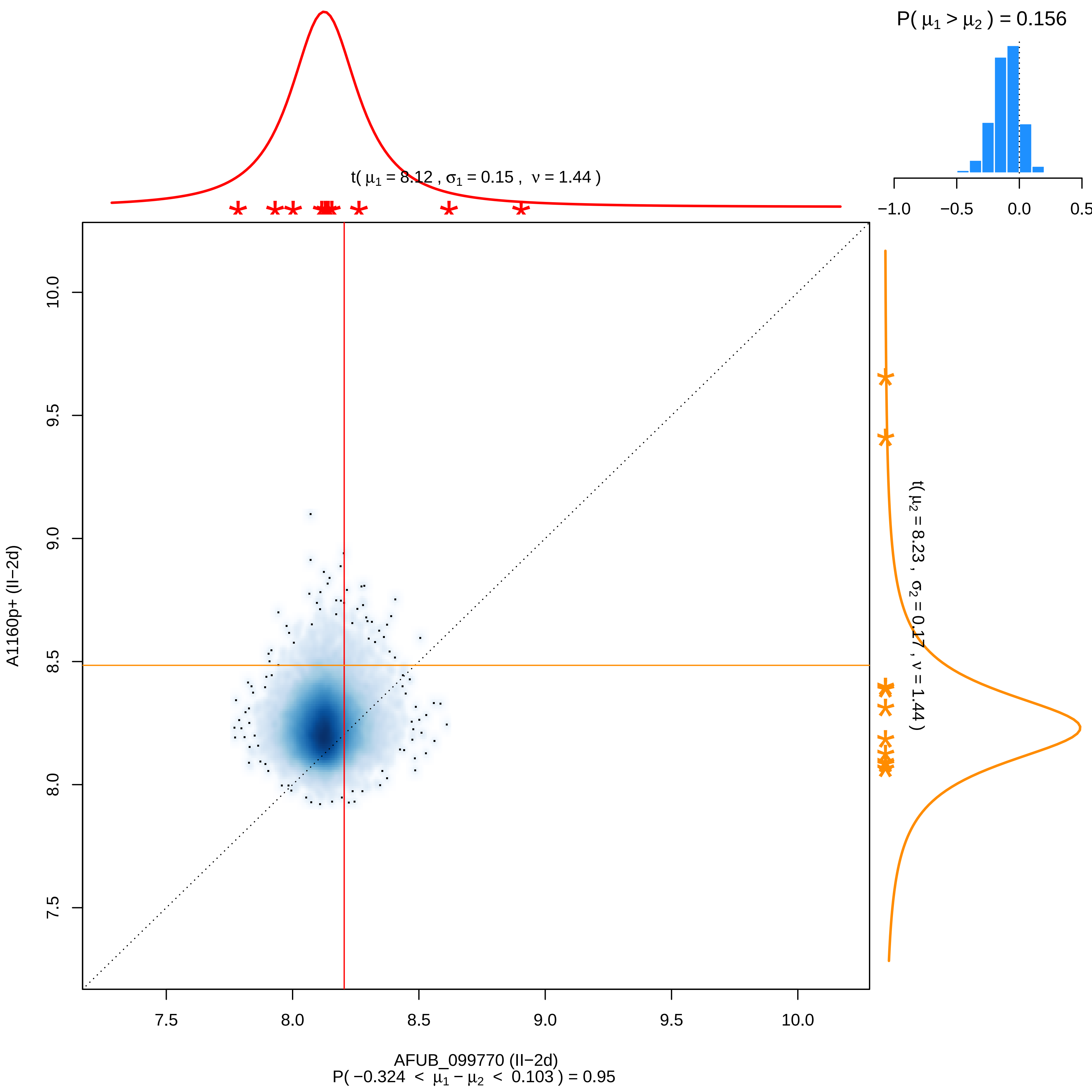

P( μ1 > μ2 ) = 0.156
−1.0
−0.5
0.0
0.5
t( μ1 = 8.12 , σ1 = 0.15 , ν = 1.44 )
t( μ2 = 8.23 , σ2 = 0.17 , ν = 1.44 )
A1160p+ (II−2d)
10.0
9.5
9.0
8.5
8.0
7.5
7.5
8.0
8.5
9.0
9.5
10.0
AFUB_099770 (II−2d)
P( −0.324 < μ1 − μ2 < 0.103 ) = 0.95

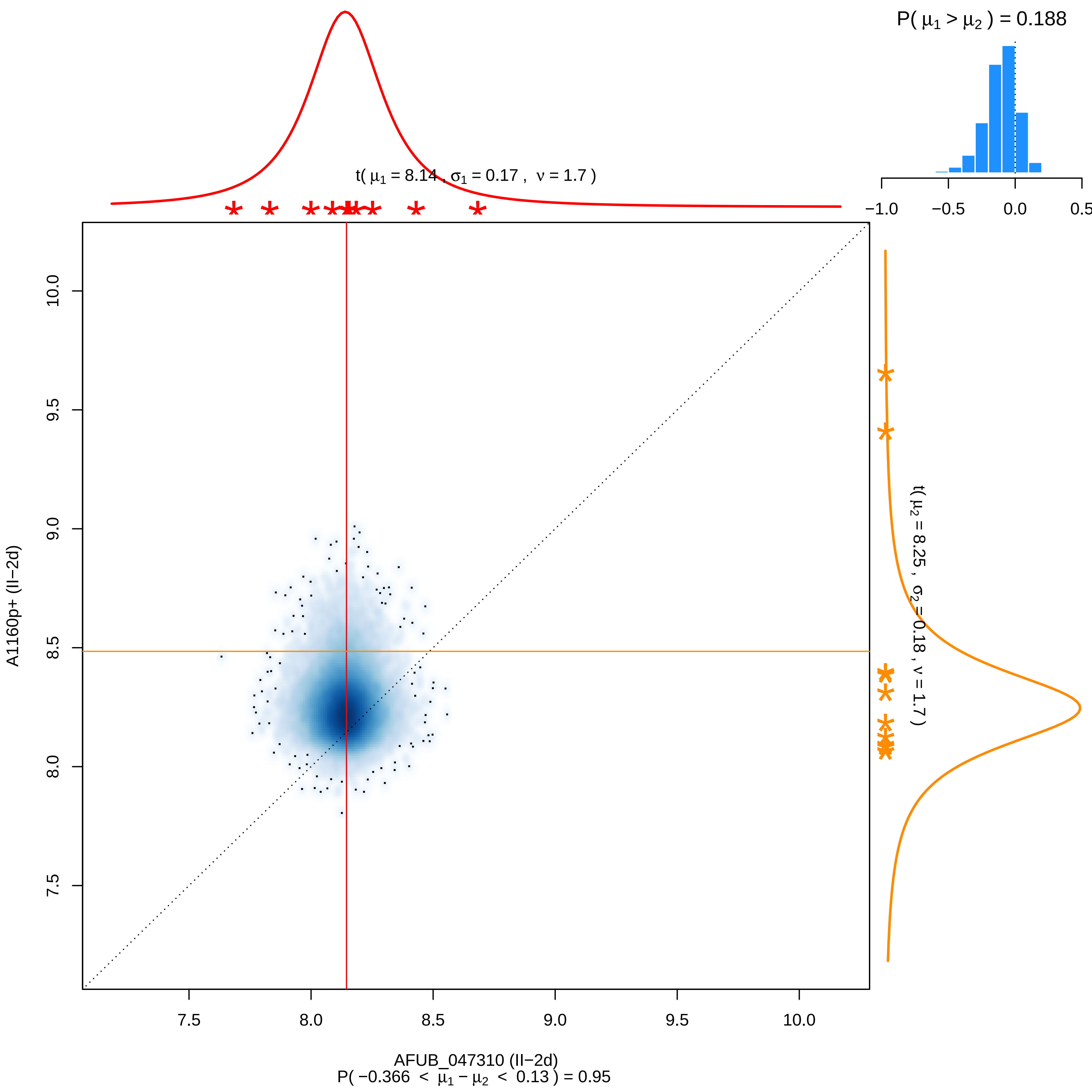

P( μ1 > μ2 ) = 0.188
−1.0 −0.5 0.0 0.5
t( μ1 = 8.14 , σ1 = 0.17 , ν = 1.7 )
t( μ2 = 8.25 , σ2 = 0.18 , ν = 1.7 )
A1160p+ (II−2d)
10.0 9.5 9.0 8.5 8.0 7.5
7.5 8.0 8.5 9.0 9.5 10.0
AFUB_047310 (II−2d)
P( −0.366 < μ1 − μ2 < 0.13 ) = 0.95

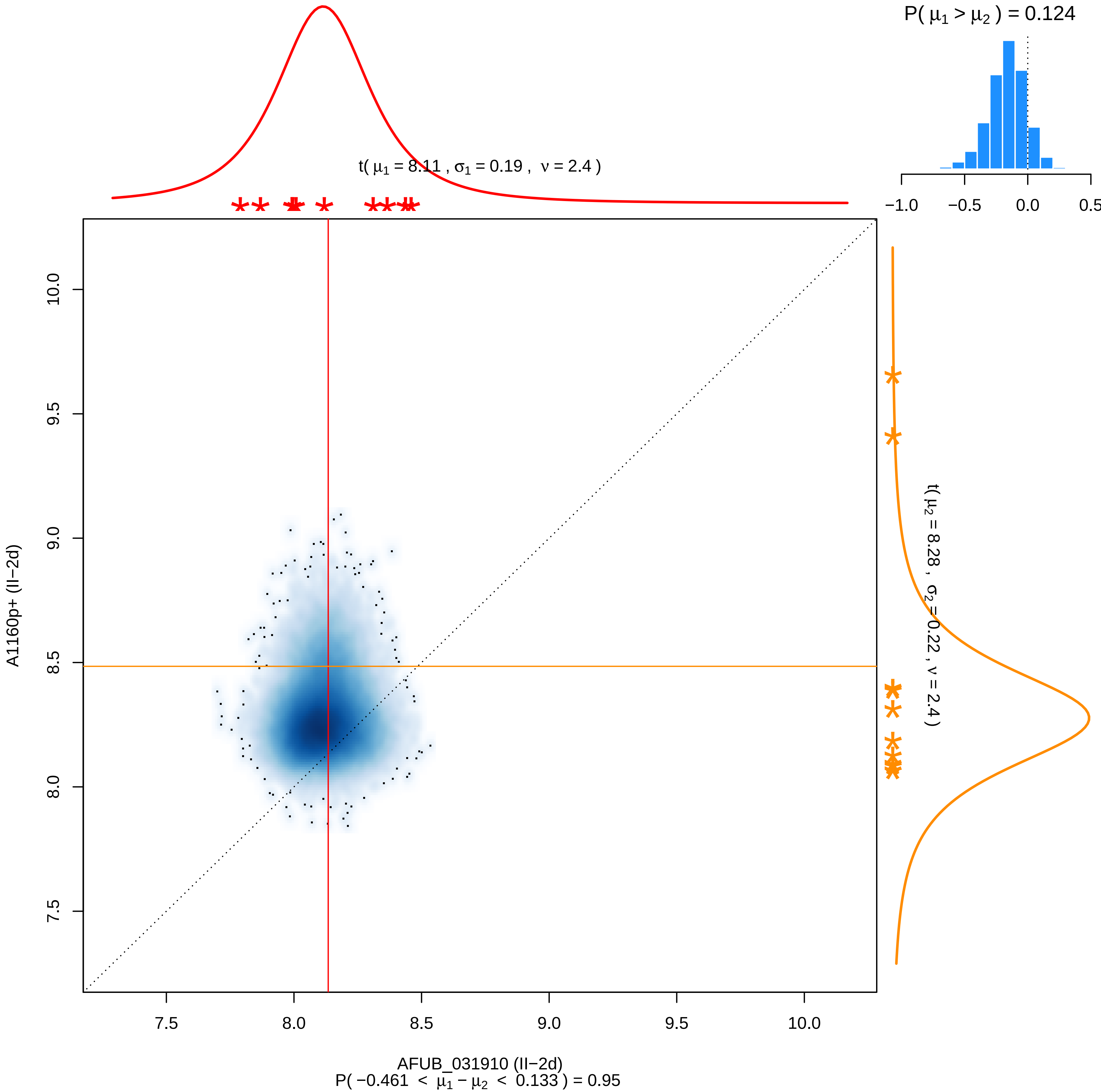

P( μ1 > μ2 ) = 0.124
t( μ1 = 8.11 , σ1 = 0.19 , ν = 2.4 )
−1.0
−0.5
0.0
0.5
10.0
9.5
9.0
8.5
8.0
7.5
A1160p+ (II−2d)
t( μ2 = 8.28 , σ2 = 0.22 , ν = 2.4 )
7.5
8.0
8.5
9.0
9.5
10.0
AFUB_031910 (II−2d)
P( −0.461 < μ1 − μ2 < 0.133 ) = 0.95

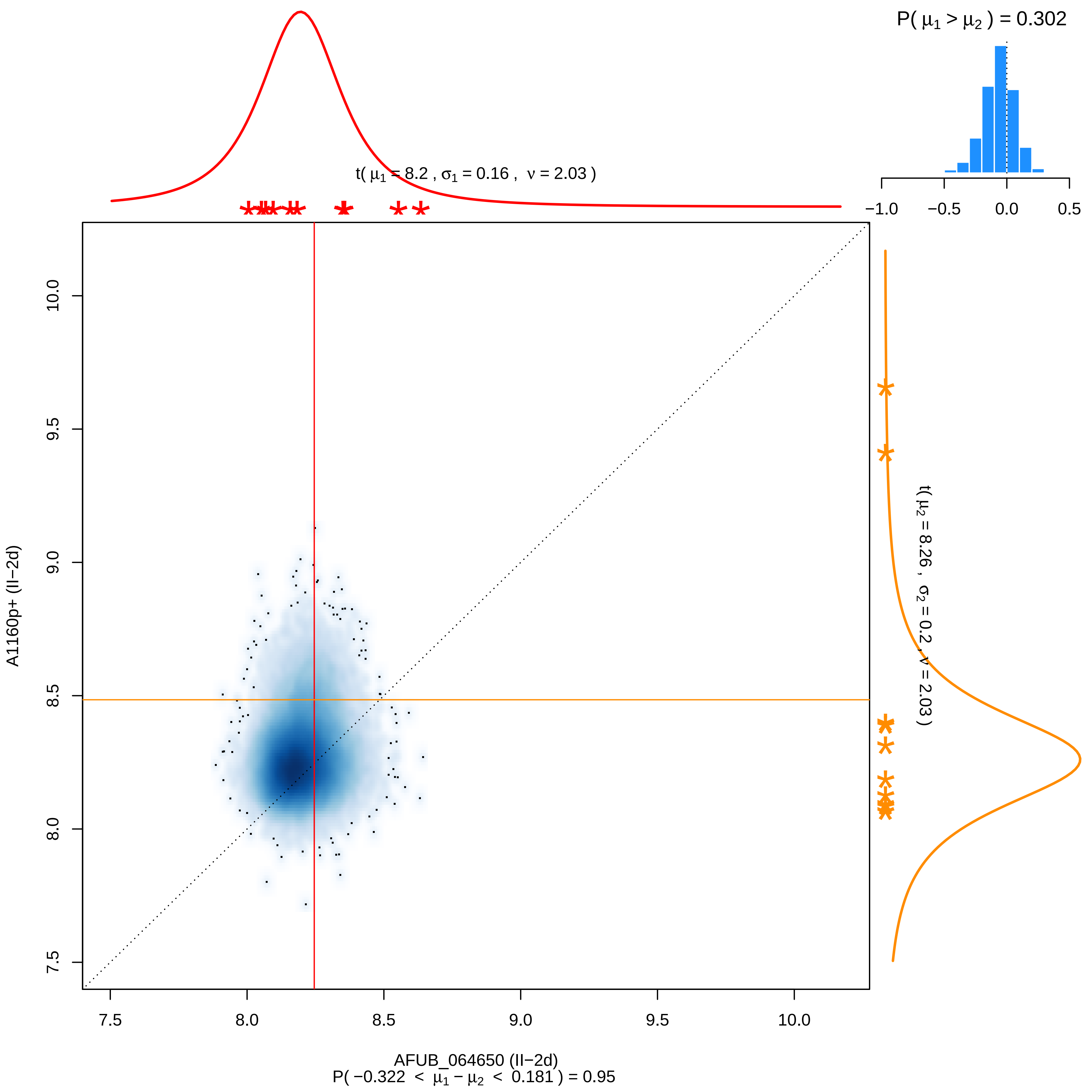

P( μ1 > μ2 ) = 0.302
t( μ1 = 8.2 , σ1 = 0.16 , ν = 2.03 )
t( μ2 = 8.26 , σ2 = 0.2 , ν = 2.03 )
A1160p+ (II−2d)
AFUB_064650 (II−2d)
P( −0.322 < μ1 − μ2 < 0.181 ) = 0.95

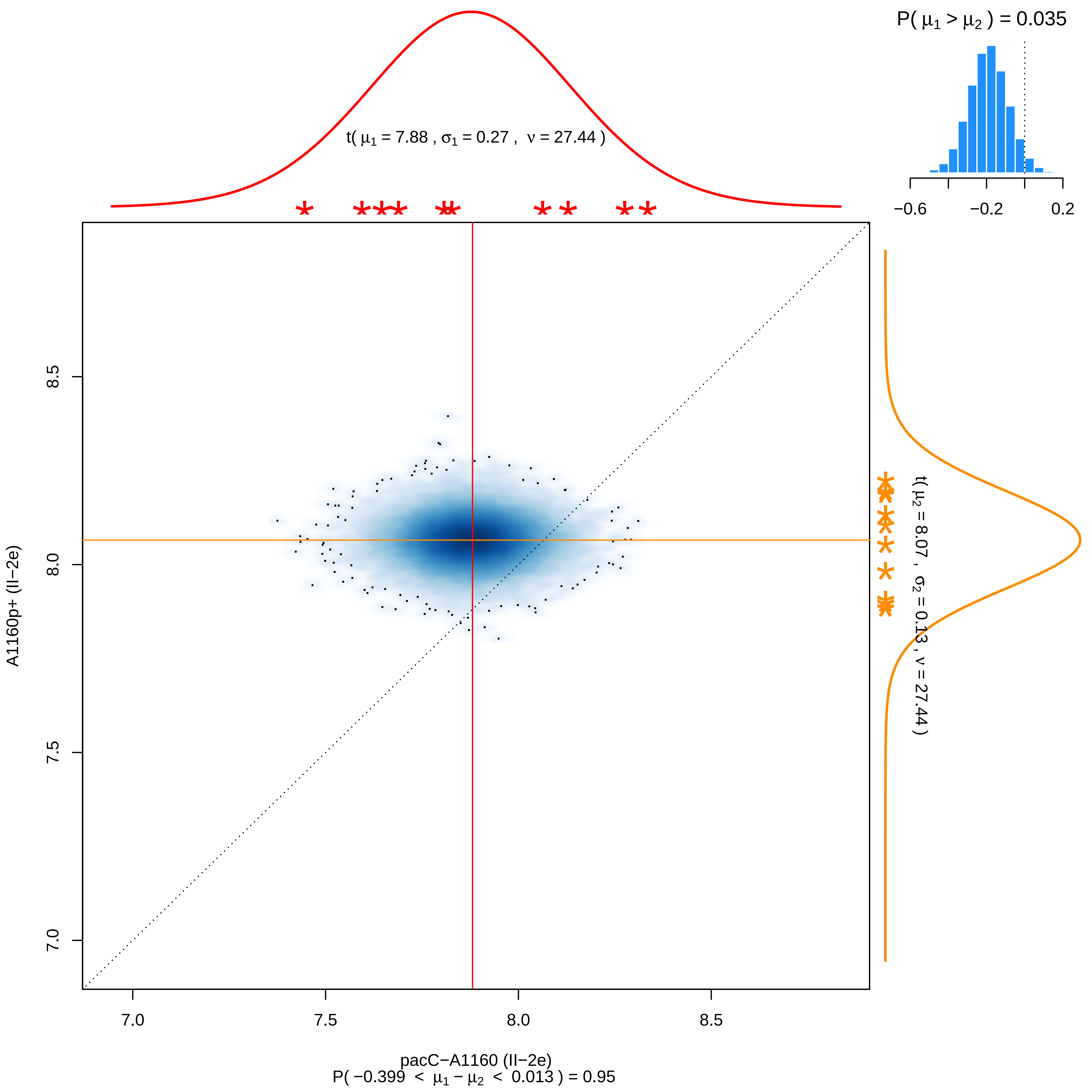

P( μ1 > μ2 ) = 0.035
−0.6
−0.2
0.2
t( μ1 = 7.88 , σ1 = 0.27 , ν = 27.44 )
t( μ2 = 8.07 , σ2 = 0.13 , ν = 27.44 )
A1160p+ (II−2e)
7.0
7.5
8.0
8.5
pacC−A1160 (II−2e)
P( −0.399 < μ1 − μ2 < 0.013 ) = 0.95

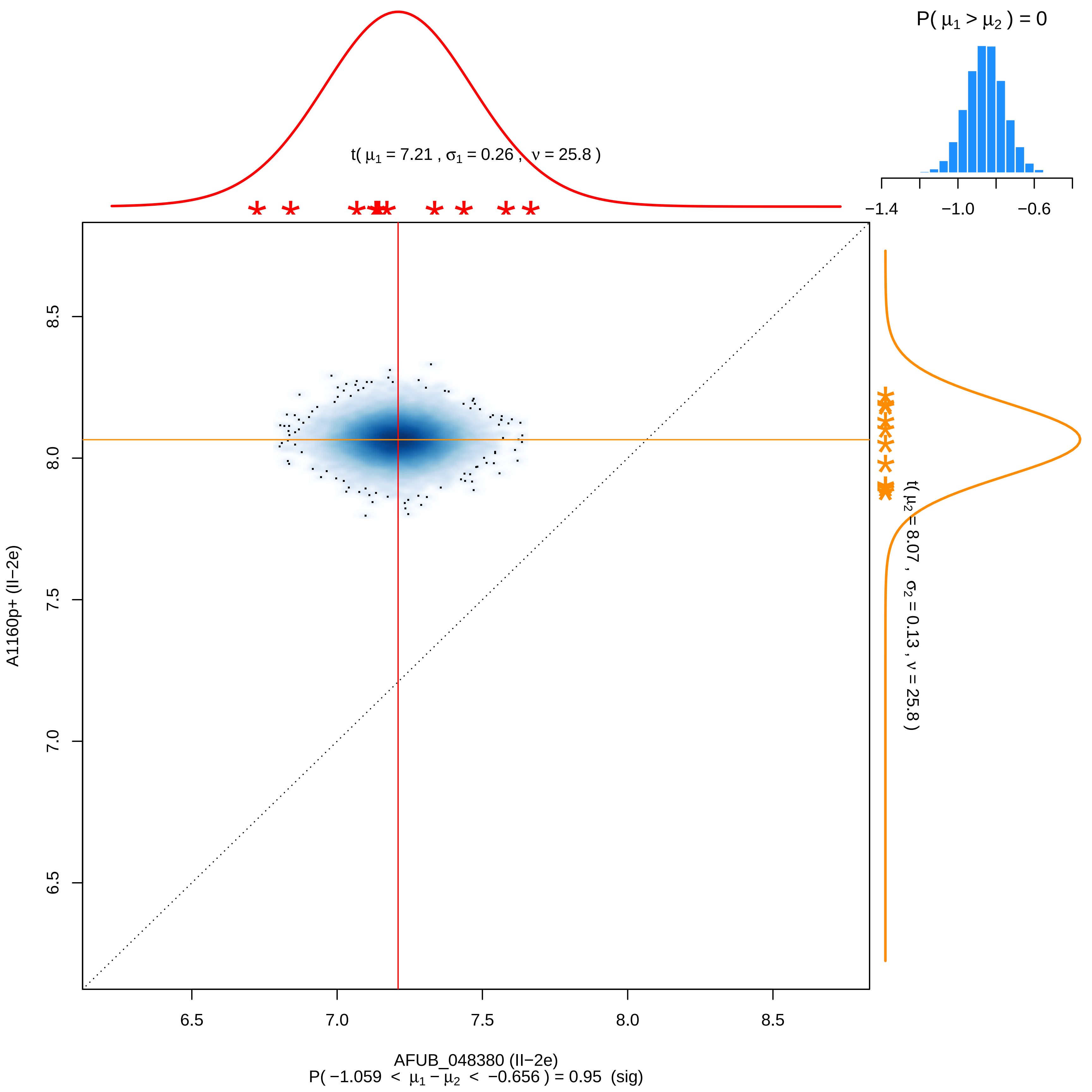

P( μ1 > μ2 ) = 0
−1.4
−1.0
−0.6
t( μ1 = 7.21 , σ1 = 0.26 , ν = 25.8 )
t( μ2 = 8.07 , σ2 = 0.13 , ν = 25.8 )
A1160p+ (II−2e)
8.5
8.0
7.5
7.0
6.5
AFUB_048380 (II−2e)
P( −1.059 < μ1 − μ2 < −0.656 ) = 0.95 (sig)

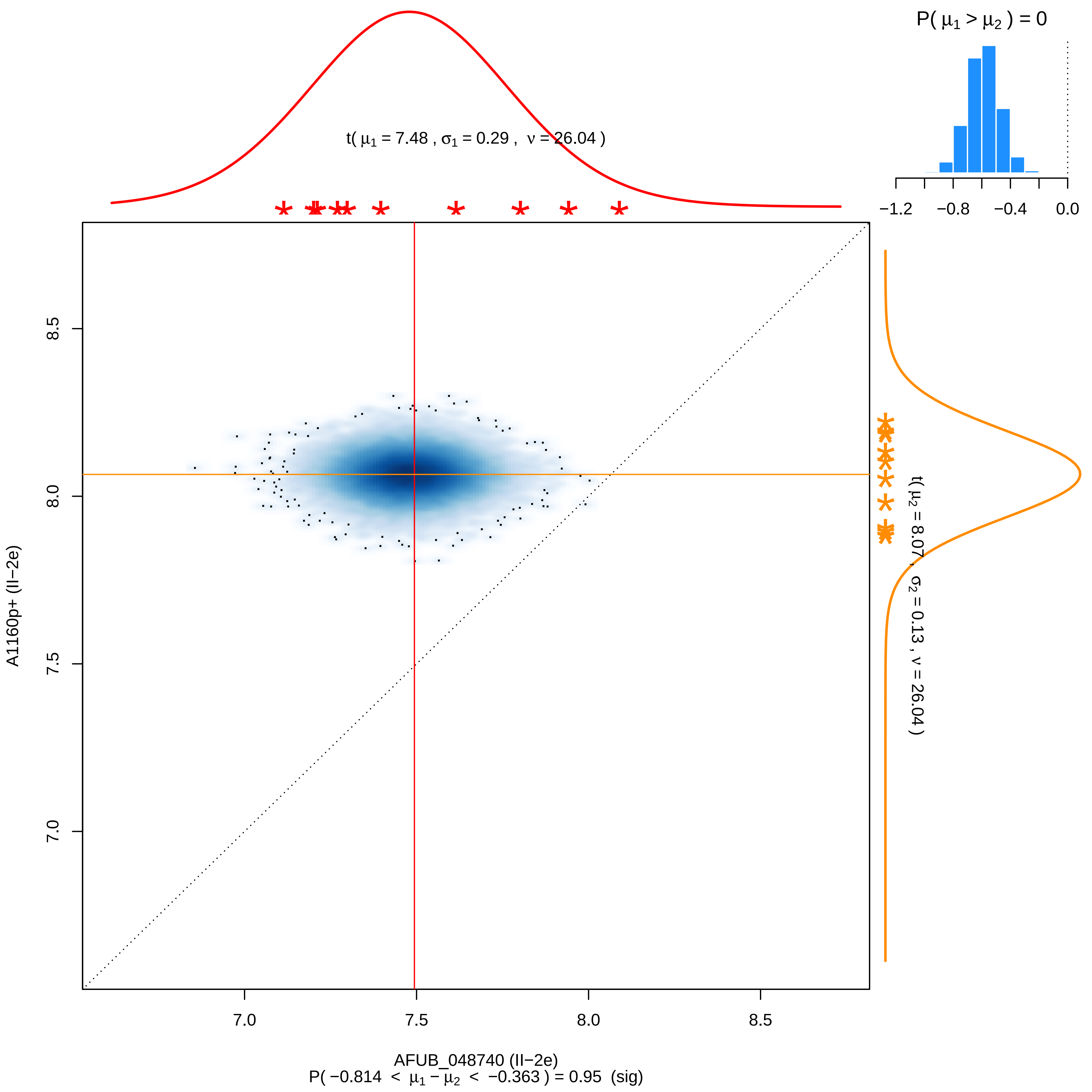

P( μ1 > μ2 ) = 0
−1.2
−0.8
−0.4
0.0
t( μ1 = 7.48 , σ1 = 0.29 , ν = 26.04 )
t( μ2 = 8.07 , σ2 = 0.13 , ν = 26.04 )
A1160p+ (II−2e)
7.0
7.5
8.0
8.5
AFUB_048740 (II−2e)
P( −0.814 < μ1 − μ2 < −0.363 ) = 0.95 (sig)

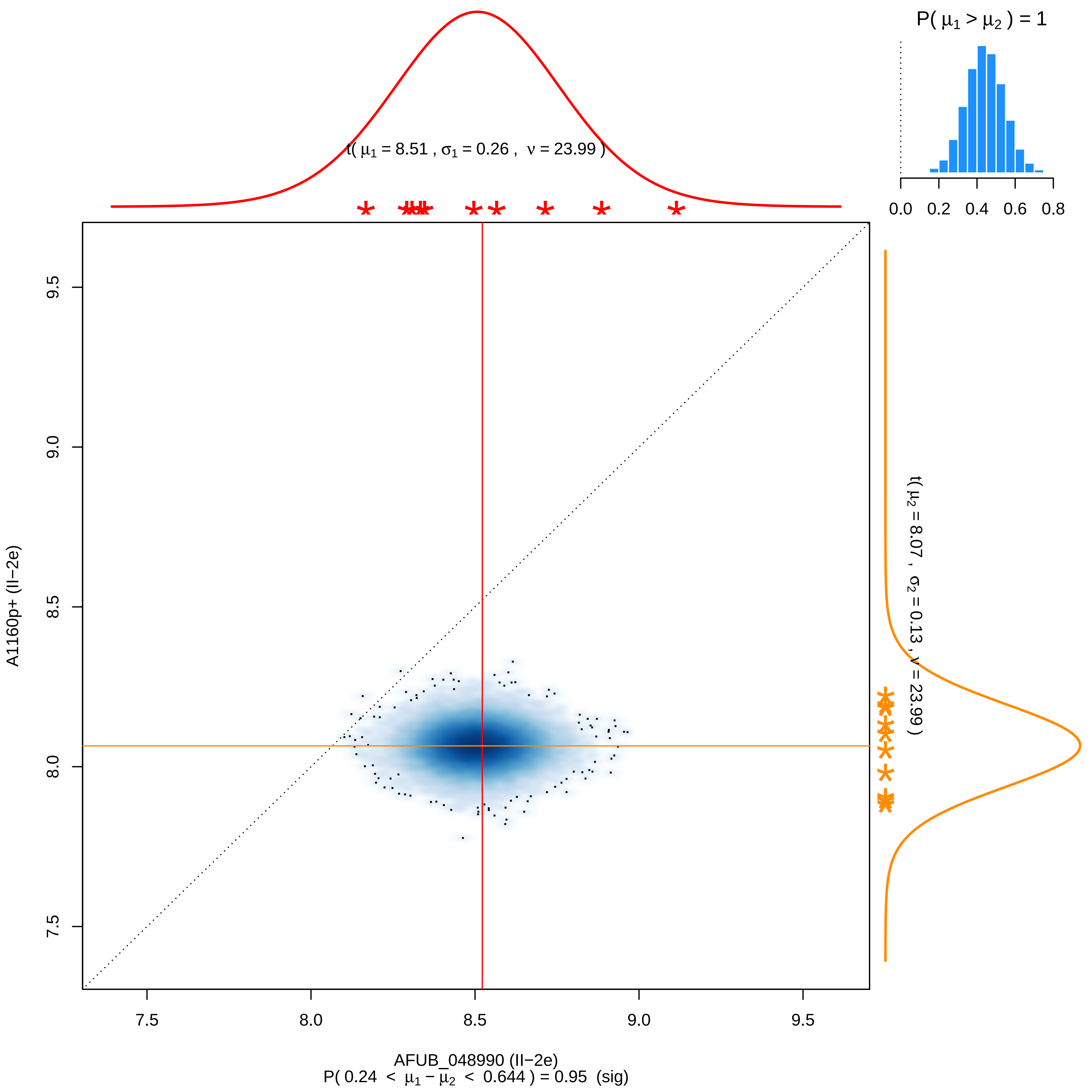
P( μ1 > μ2 ) = 1
0.0 0.2 0.4 0.6 0.8
t( μ1 = 8.51 , σ1 = 0.26 , ν = 23.99 )
t( μ2 = 8.07 , σ2 = 0.13 , ν = 23.99 )
A1160p+ (II−2e)
9.5
9.0
8.5
8.0
7.5
7.5 8.0 8.5 9.0 9.5
AFUB_048990 (II−2e)
P( 0.24 < μ1 − μ2 < 0.644 ) = 0.95 (sig)

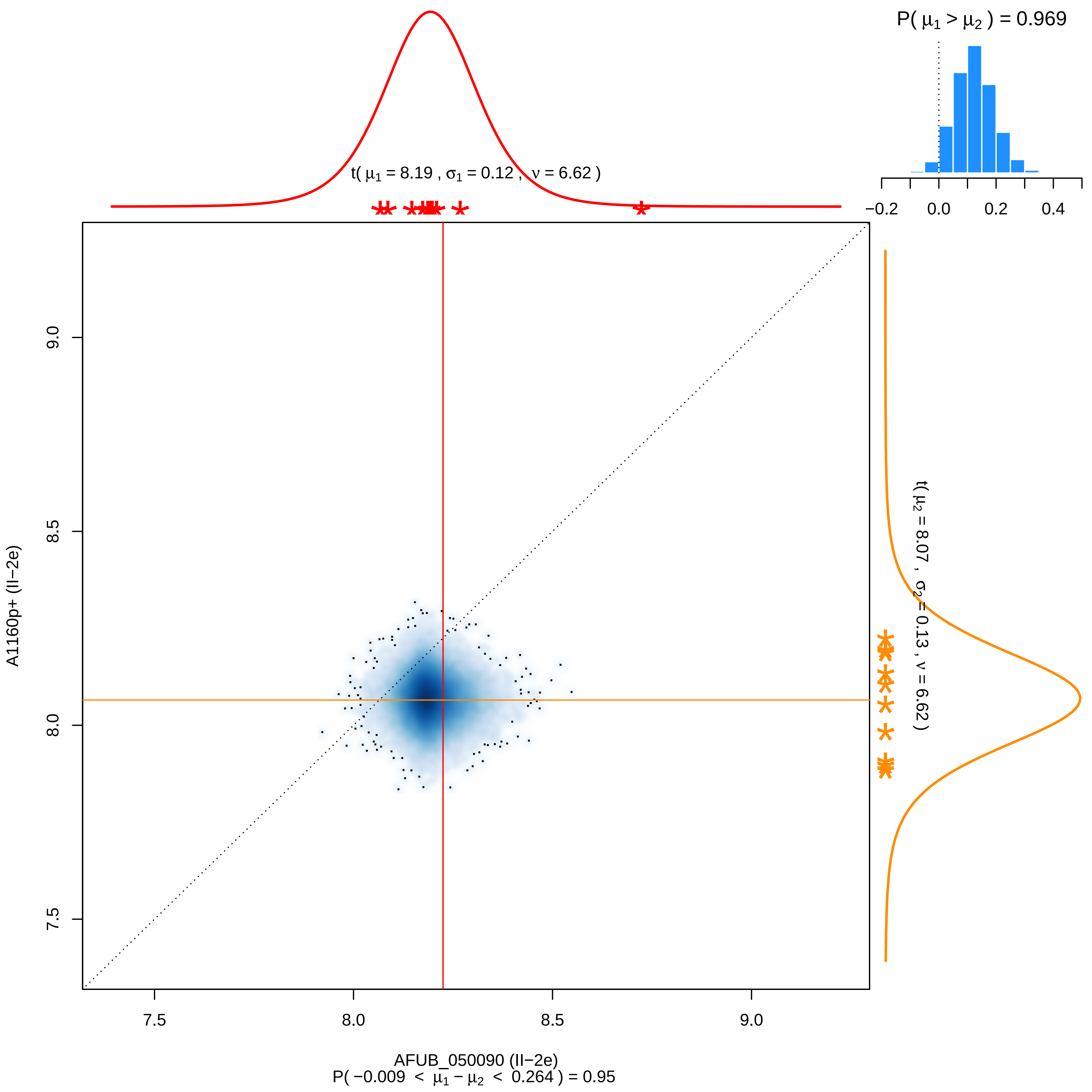

P( μ1 > μ2 ) = 0.969
t( μ1 = 8.19 , σ1 = 0.12 , ν = 6.62 )
t( μ2 = 8.07 , σ2 = 0.13 , ν = 6.62 )
A1160p+ (II−2e)
AFUB_050090 (II−2e)
P( −0.009 < μ1 − μ2 < 0.264 ) = 0.95
−0.2
0.0
0.2
0.4
7.5
8.0
8.5
9.0

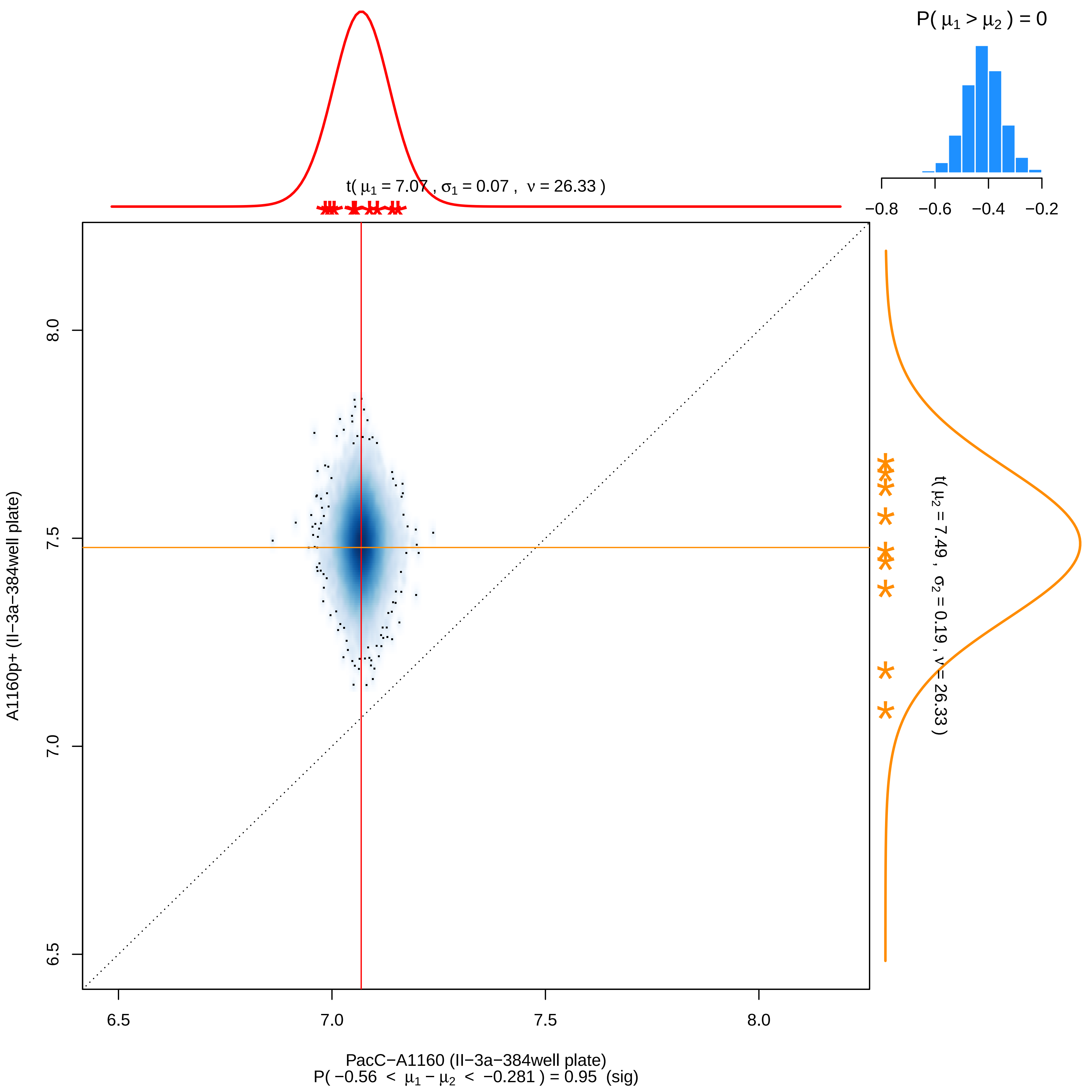

P( μ1 > μ2 ) = 0
−0.8 −0.6 −0.4 −0.2
t( μ1 = 7.07 , σ1 = 0.07 , ν = 26.33 )
t( μ2 = 7.49 , σ2 = 0.19 , ν = 26.33 )
A1160p+ (II−3a−384well plate)
8.0
7.5
7.0
6.5
6.5
7.0
7.5
8.0
PacC−A1160 (II−3a−384well plate)
P( −0.56 < μ1 − μ2 < −0.281 ) = 0.95 (sig)

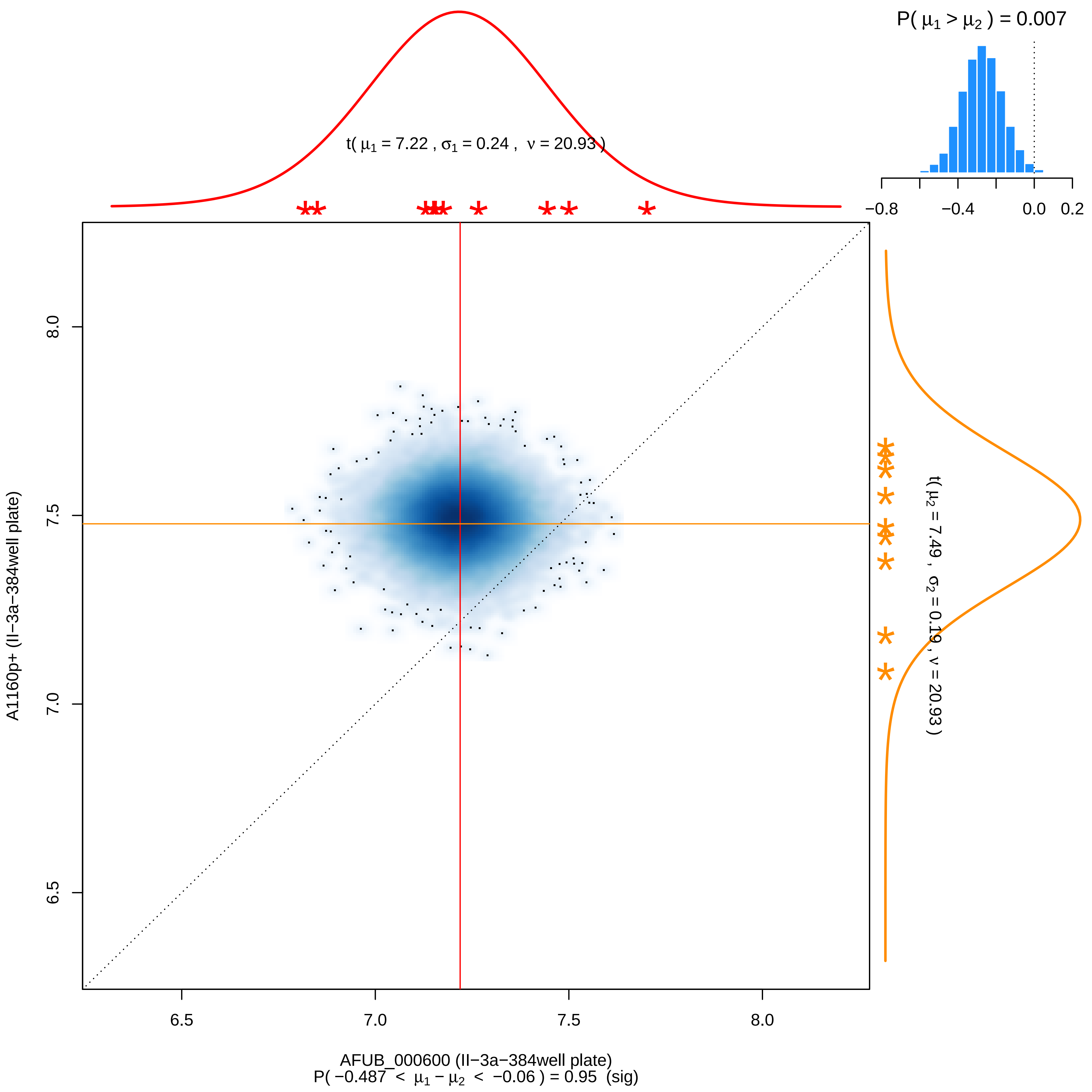

P( μ1 > μ2 ) = 0.007
−0.8
−0.4
0.0
0.2
t( μ1 = 7.22 , σ1 = 0.24 , ν = 20.93 )
t( μ2 = 7.49 , σ2 = 0.19 , ν = 20.93 )
A1160p+ (II−3a−384well plate)
6.5
7.0
7.5
8.0
AFUB_000600 (II−3a−384well plate)
P( −0.487 < μ1 − μ2 < −0.06 ) = 0.95 (sig)

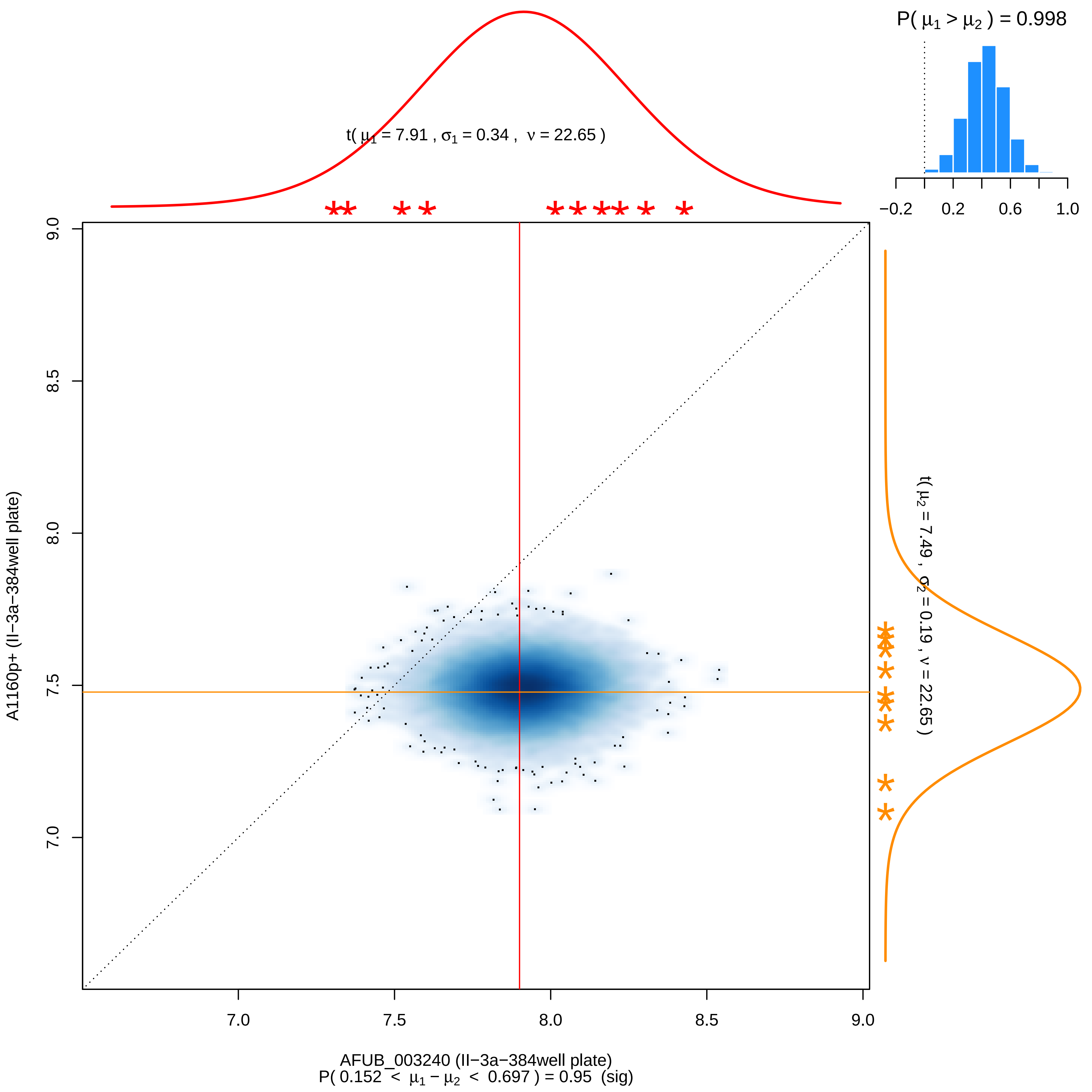
P( μ1 > μ2 ) = 0.998
−0.2
0.2
0.6
1.0
t( μ1 = 7.91 , σ1 = 0.34 , ν = 22.65 )
t( μ2 = 7.49 , σ2 = 0.19 , ν = 22.65 )
9.0
8.5
8.0
7.5
7.0
A1160p+ (II−3a−384well plate)
7.0
7.5
8.0
8.5
9.0
AFUB_003240 (II−3a−384well plate)
P( 0.152 < μ1 − μ2 < 0.697 ) = 0.95 (sig)

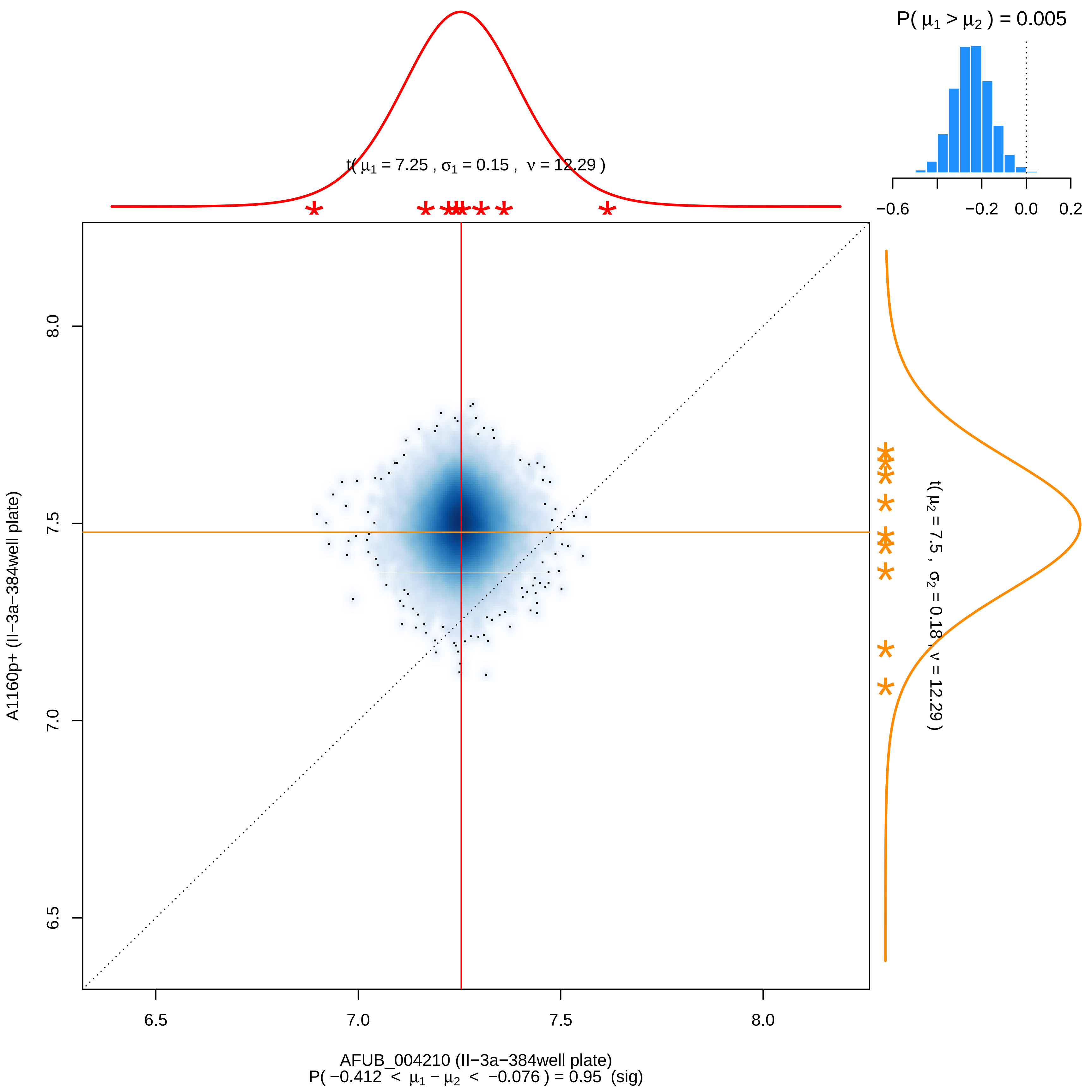

P( μ1 > μ2 ) = 0.005
−0.6
−0.2
0.0
0.2
t( μ1 = 7.25 , σ1 = 0.15 , ν = 12.29 )
t( μ2 = 7.5 , σ2 = 0.18 , ν = 12.29 )
A1160p+ (II−3a−384well plate)
8.0
7.5
7.0
6.5
6.5
7.0
7.5
8.0
AFUB_004210 (II−3a−384well plate)
P( −0.412 < μ1 − μ2 < −0.076 ) = 0.95 (sig)

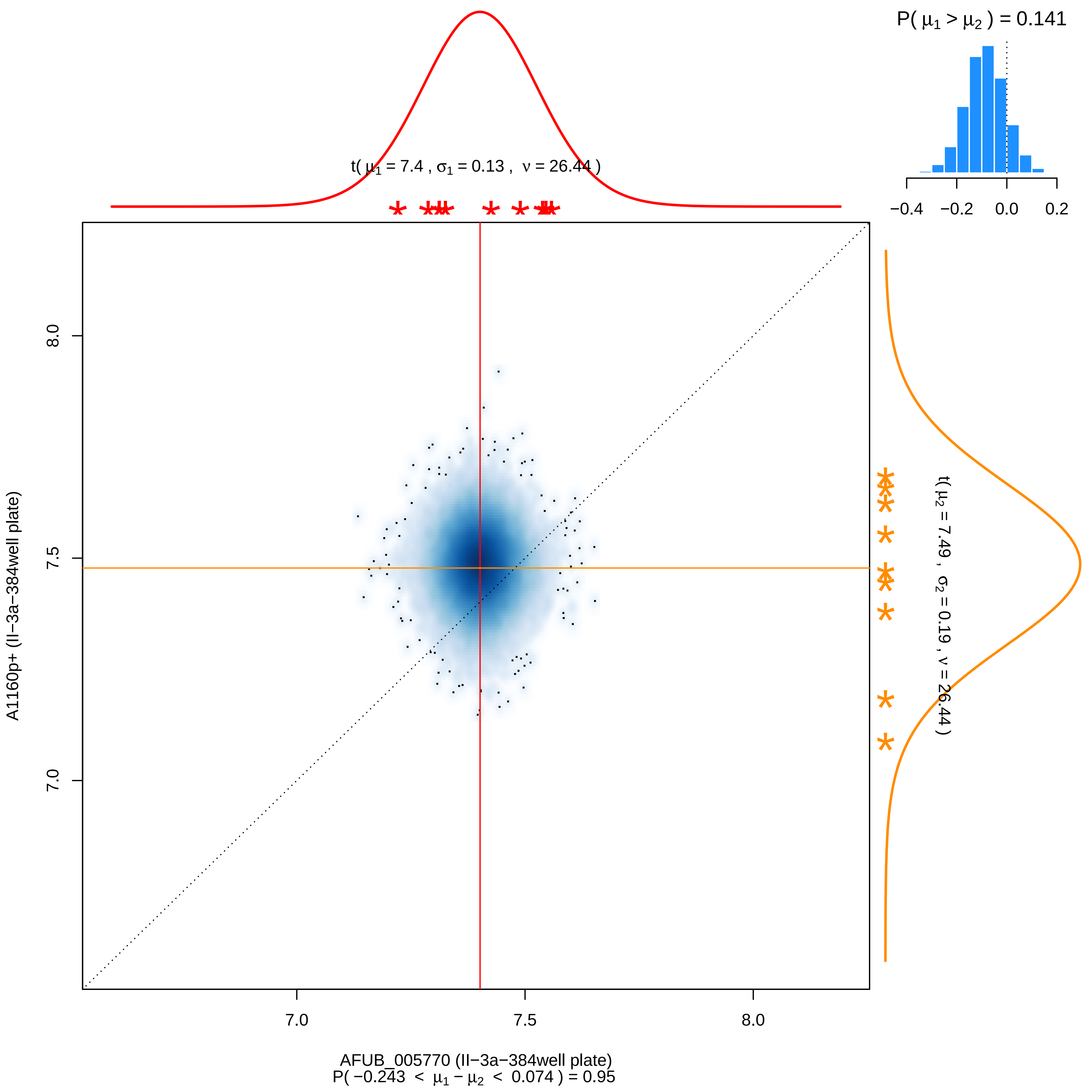

P( μ1 > μ2 ) = 0.141
−0.4
−0.2
0.0
0.2
t( μ1 = 7.4 , σ1 = 0.13 , ν = 26.44 )
t( μ2 = 7.49 , σ2 = 0.19 , ν = 26.44 )
8.0
7.5
7.0
A1160p+ (II−3a−384well plate)
7.0
7.5
8.0
AFUB_005770 (II−3a−384well plate)
P( −0.243 < μ1 − μ2 < 0.074 ) = 0.95

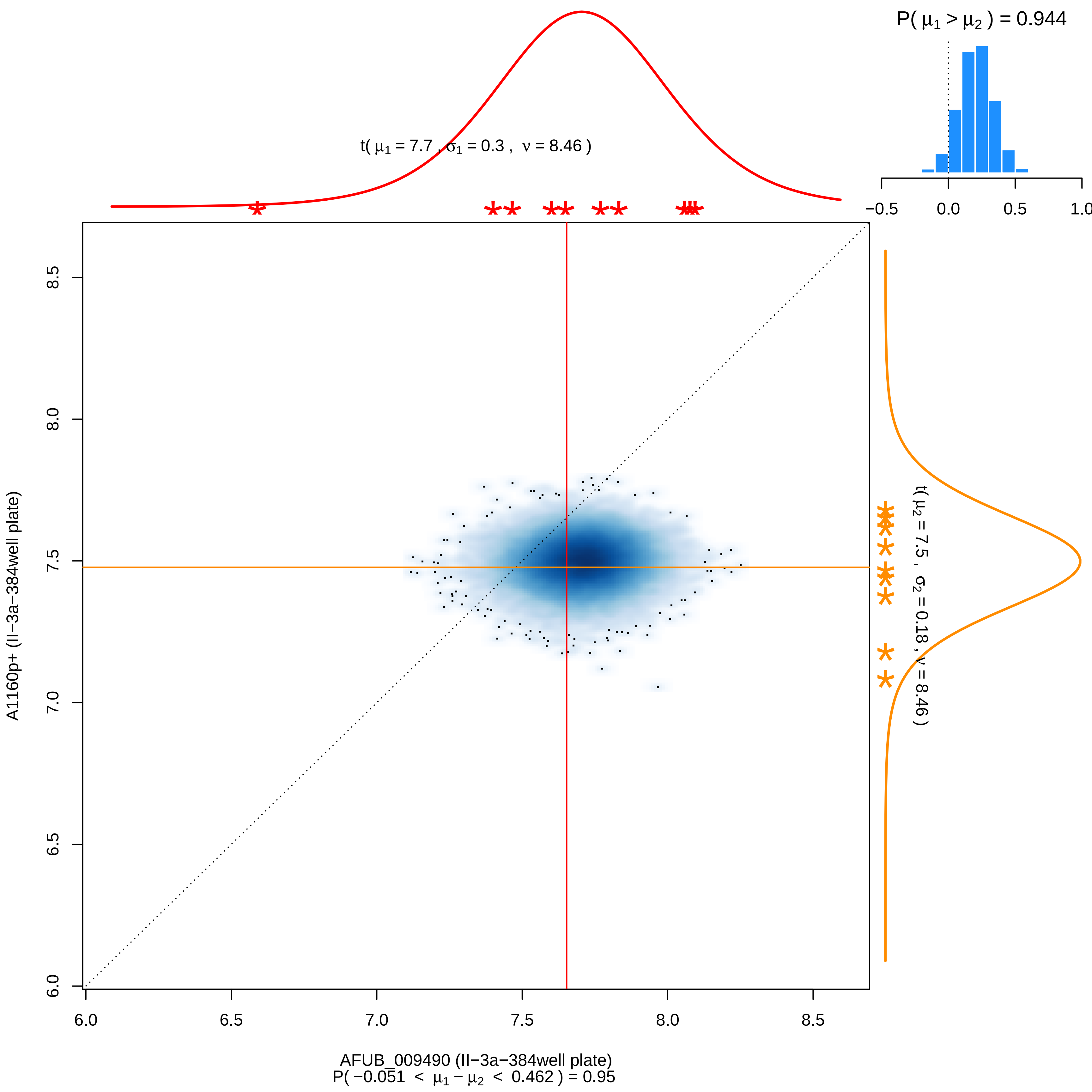

P( μ1 > μ2 ) = 0.944
t( μ1 = 7.7 , σ1 = 0.3 , ν = 8.46 )
t( μ2 = 7.5 , σ2 = 0.18 , ν = 8.46 )
−0.5 0.0 0.5 1.0
A1160p+ (II−3a−384well plate)
AFUB_009490 (II−3a−384well plate)
P( −0.051 < μ1 − μ2 < 0.462 ) = 0.95
6.0 6.5 7.0 7.5 8.0 8.5

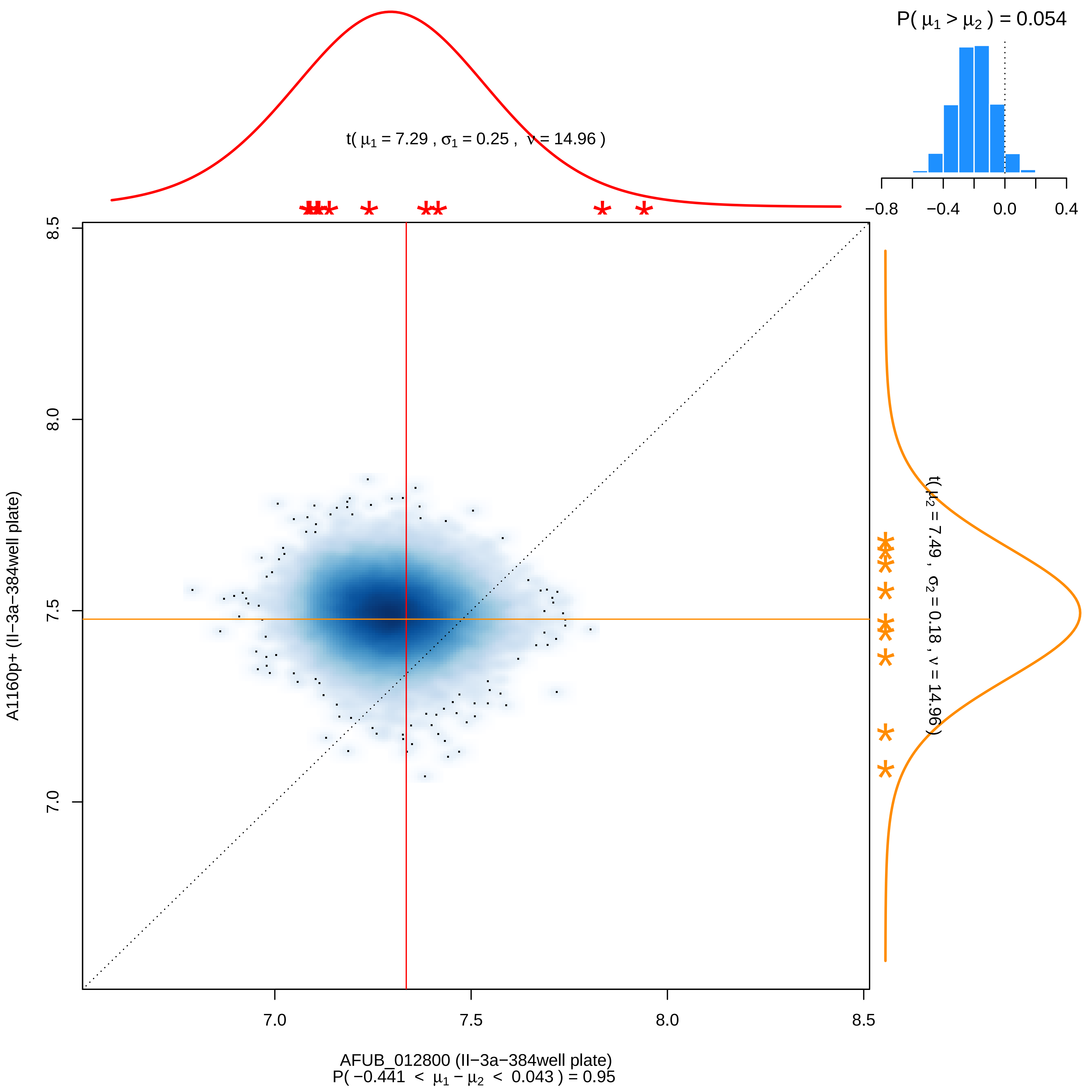

P( μ1 > μ2 ) = 0.054
−0.8 −0.4 0.0 0.4
t( μ1 = 7.29 , σ1 = 0.25 , ν = 14.96 )
t( μ2 = 7.49 , σ2 = 0.18 , ν = 14.96 )
8.5
8.0
7.5
7.0
A1160p+ (II−3a−384well plate)
7.0 7.5 8.0 8.5
AFUB_012800 (II−3a−384well plate)
P( −0.441 < μ1 − μ2 < 0.043 ) = 0.95

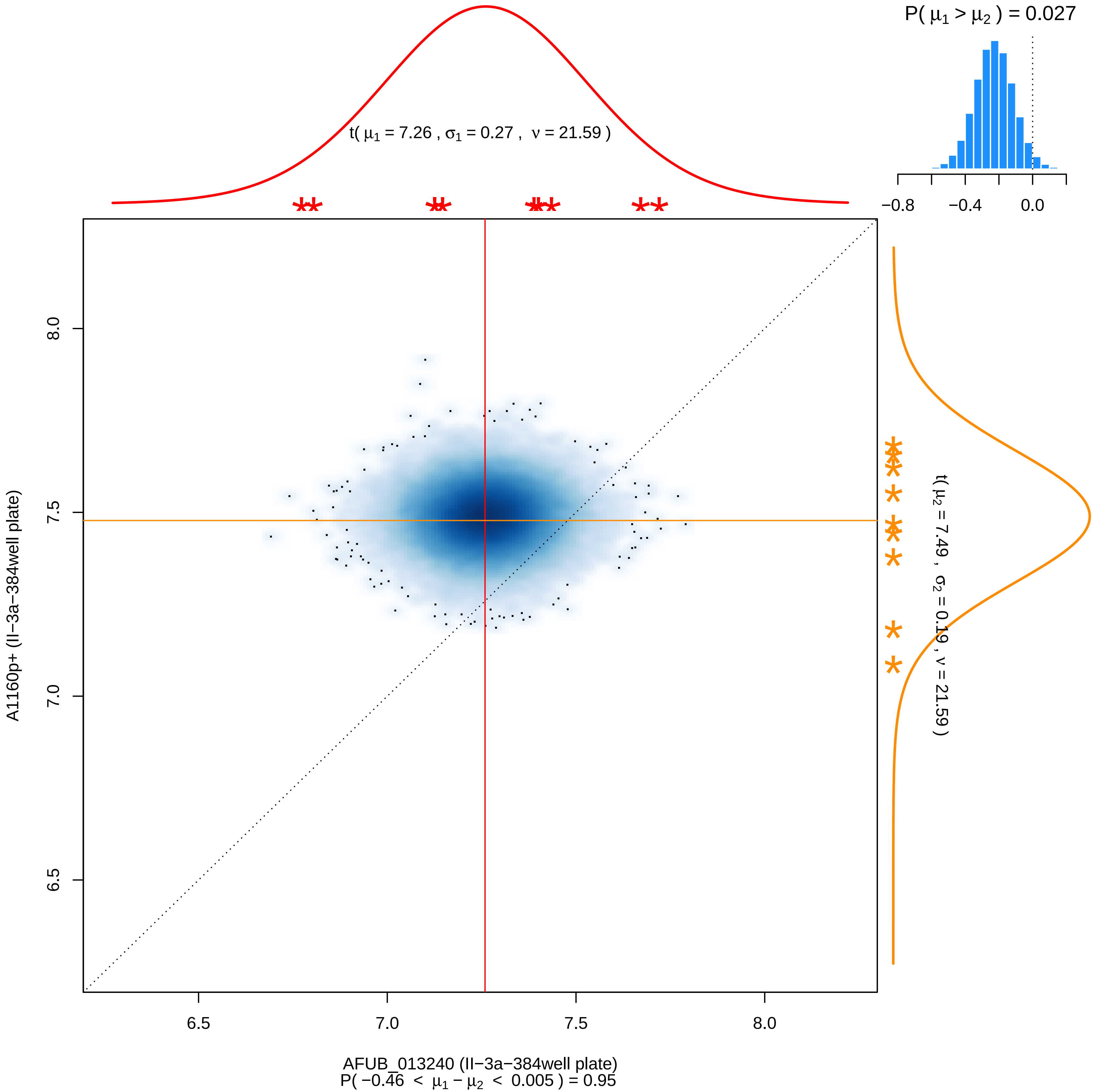

P( μ1 > μ2 ) = 0.027
−0.8
−0.4
0.0
t( μ1 = 7.26 , σ1 = 0.27 , ν = 21.59 )
8.0
7.5
7.0
6.5
A1160p+ (II−3a−384well plate)
t( μ2 = 7.49 , σ2 = 0.19 , ν = 21.59 )
6.5
7.0
7.5
8.0
AFUB_013240 (II−3a−384well plate)
P( −0.46 < μ1 − μ2 < 0.005 ) = 0.95

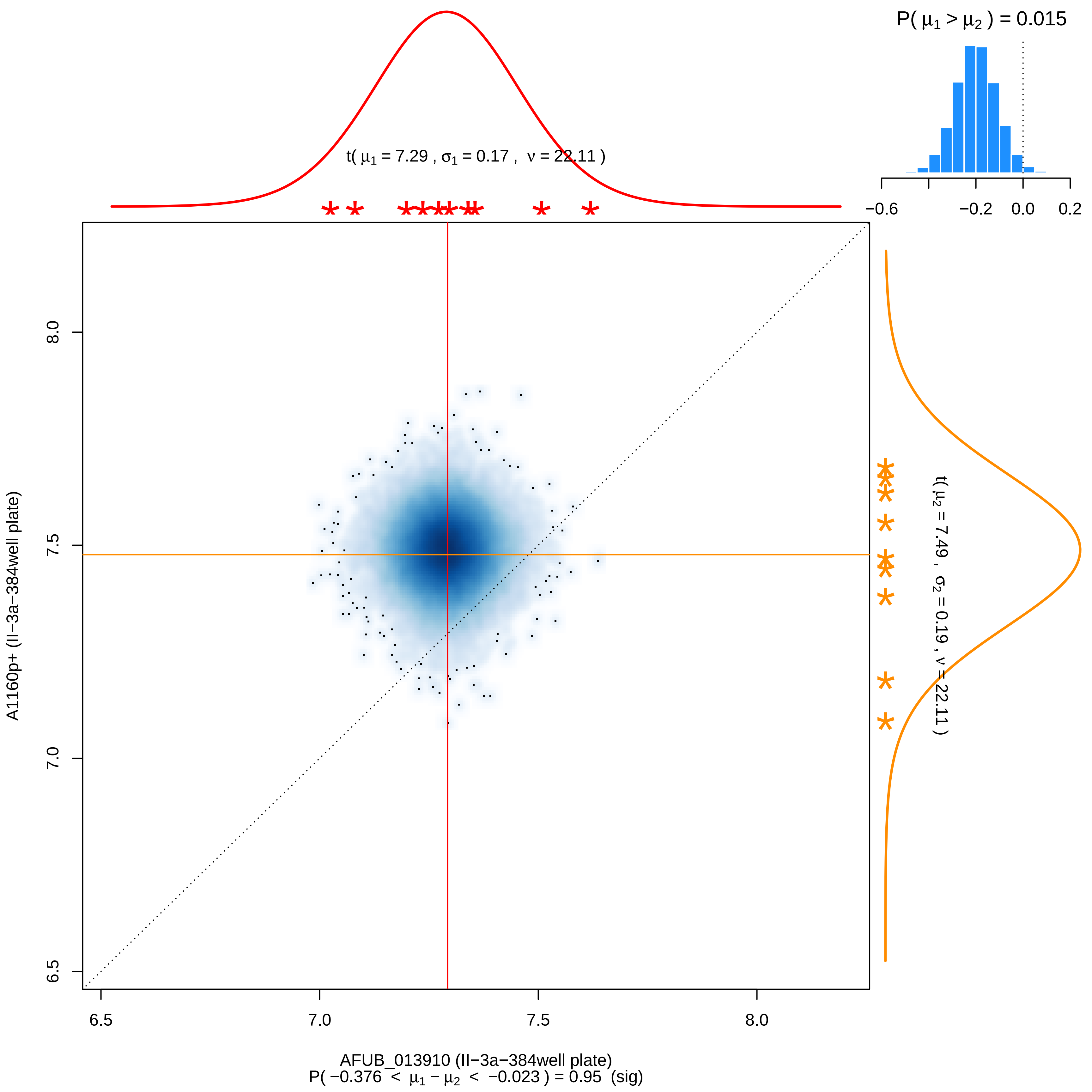

P( μ1 > μ2 ) = 0.015
−0.6
−0.2
0.0
0.2
t( μ1 = 7.29 , σ1 = 0.17 , ν = 22.11 )
t( μ2 = 7.49 , σ2 = 0.19 , ν = 22.11 )
A1160p+ (II−3a−384well plate)
8.0
7.5
7.0
6.5
6.5
7.0
7.5
8.0
AFUB_013910 (II−3a−384well plate)
P( −0.376 < μ1 − μ2 < −0.023 ) = 0.95 (sig)

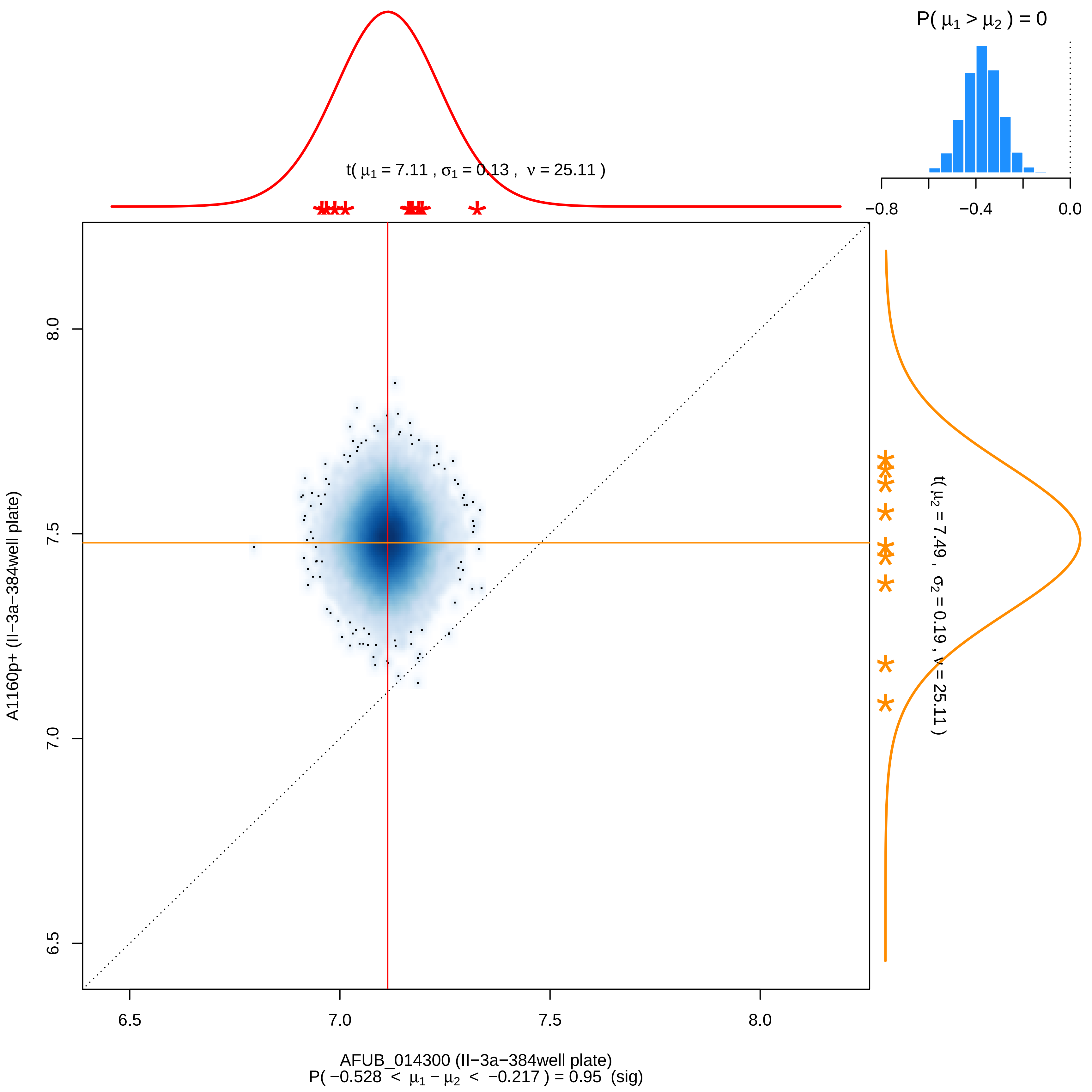

P( μ1 > μ2 ) = 0
−0.8
−0.4
0.0
t( μ1 = 7.11 , σ1 = 0.13 , ν = 25.11 )
t( μ2 = 7.49 , σ2 = 0.19 , ν = 25.11 )
A1160p+ (II−3a−384well plate)
8.0
7.5
7.0
6.5
6.5
7.0
7.5
8.0
AFUB_014300 (II−3a−384well plate)
P( −0.528 < μ1 − μ2 < −0.217 ) = 0.95 (sig)

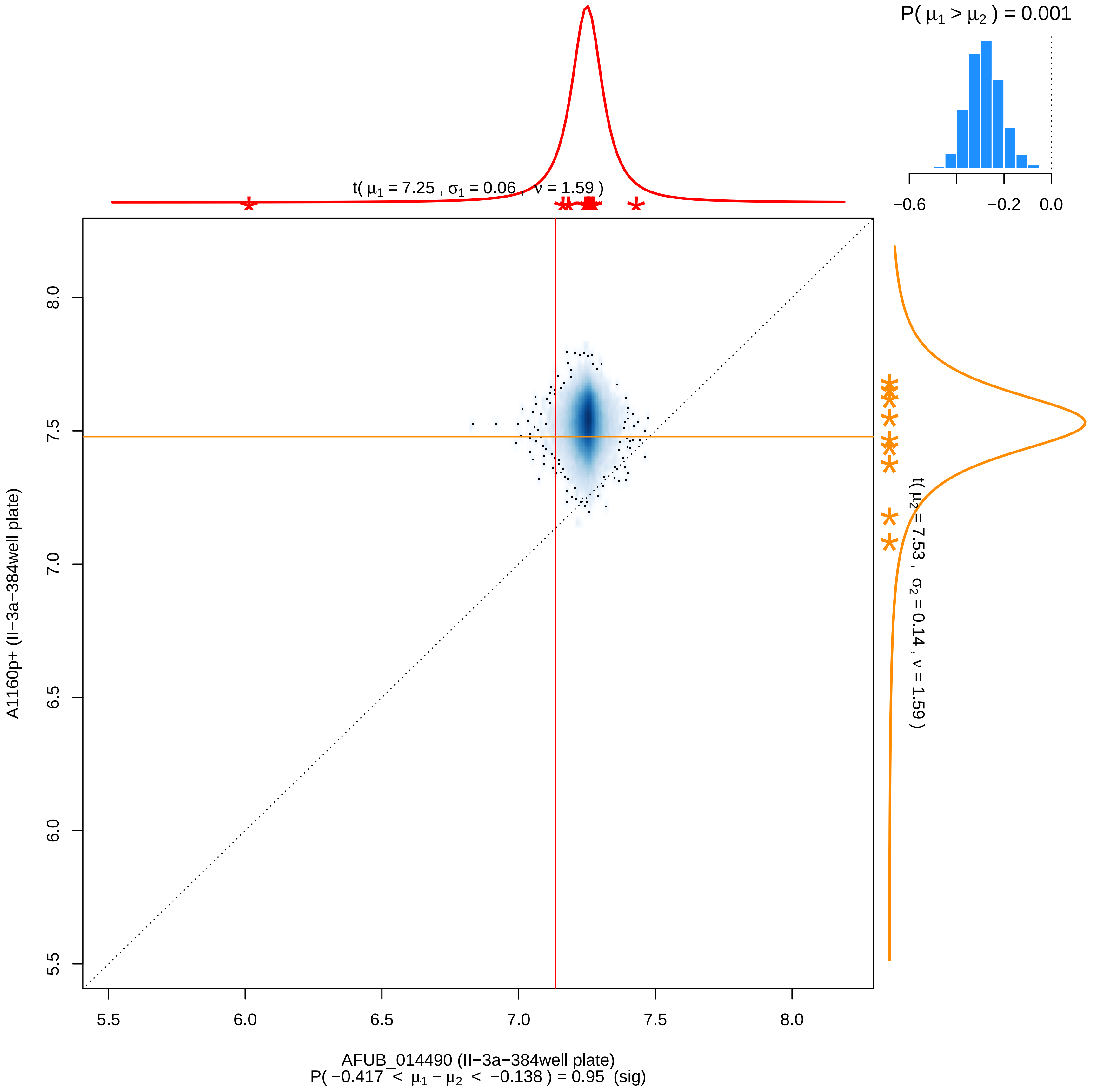

P( μ1 > μ2 ) = 0.001
−0.6
−0.2
0.0
t( μ1 = 7.25 , σ1 = 0.06 , ν = 1.59 )
t( μ2 = 7.53 , σ2 = 0.14 , ν = 1.59 )
5.5
6.0
6.5
7.0
7.5
8.0
A1160p+ (II−3a−384well plate)
AFUB_014490 (II−3a−384well plate)
P( −0.417 < μ1 − μ2 < −0.138 ) = 0.95 (sig)

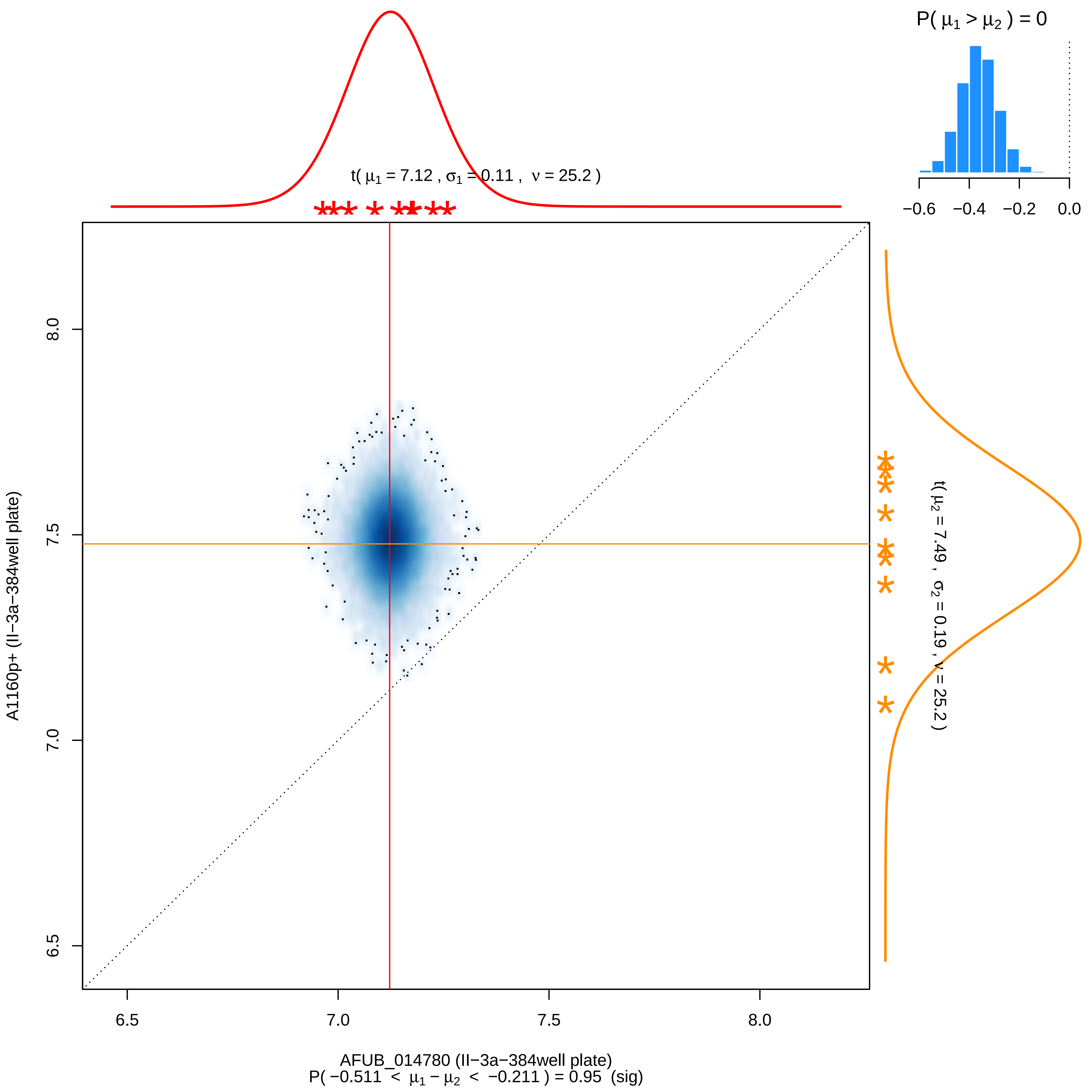

P( μ1 > μ2 ) = 0
−0.6
−0.4
−0.2
0.0
t( μ1 = 7.12 , σ1 = 0.11 , ν = 25.2 )
t( μ2 = 7.49 , σ2 = 0.19 , ν = 25.2 )
A1160p+ (II−3a−384well plate)
8.0
7.5
7.0
6.5
6.5
7.0
7.5
8.0
AFUB_014780 (II−3a−384well plate)
P( −0.511 < μ1 − μ2 < −0.211 ) = 0.95 (sig)

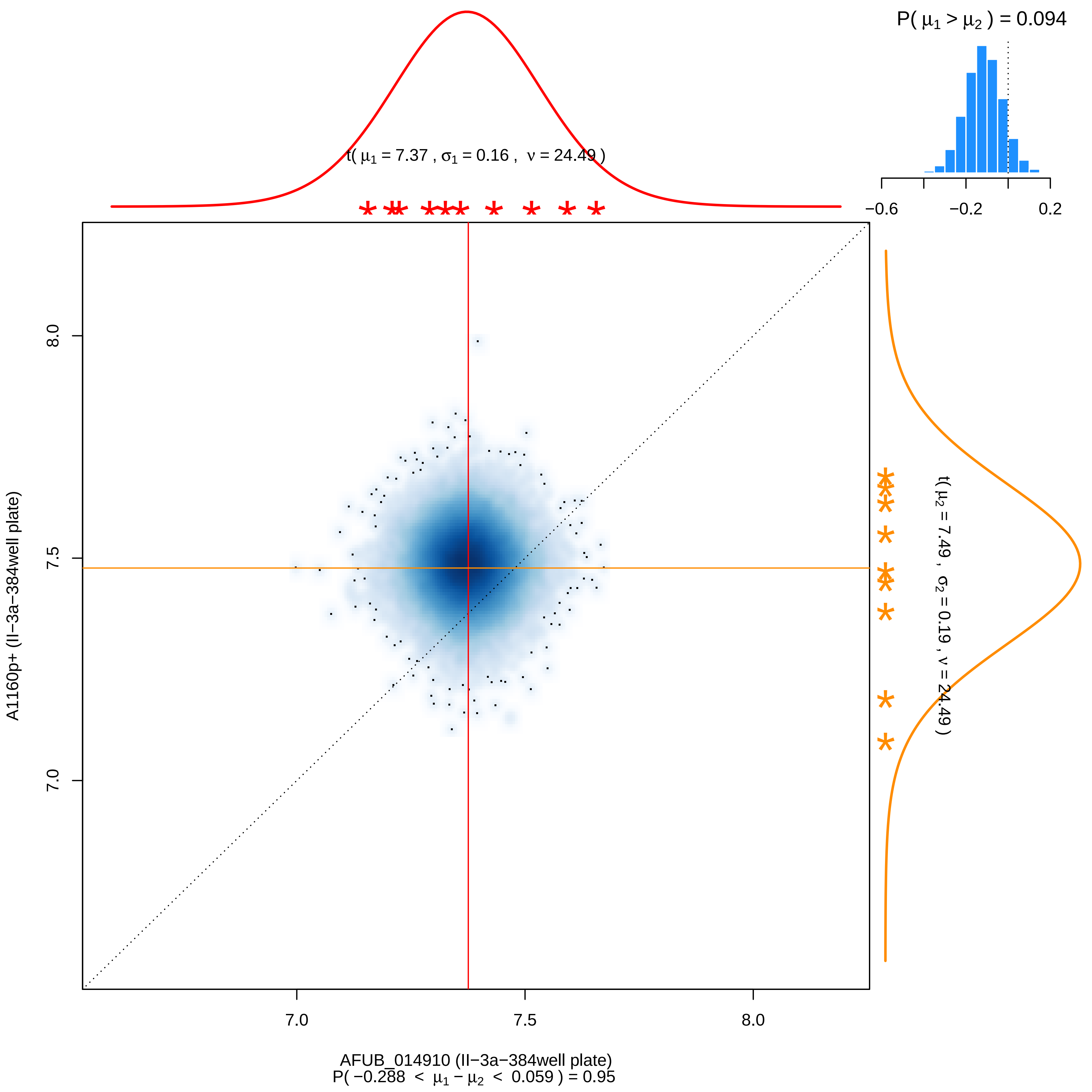

P( μ1 > μ2 ) = 0.094
−0.6
−0.2
0.2
t( μ1 = 7.37 , σ1 = 0.16 , ν = 24.49 )
t( μ2 = 7.49 , σ2 = 0.19 , ν = 24.49 )
8.0
7.5
7.0
A1160p+ (II−3a−384well plate)
7.0
7.5
8.0
AFUB_014910 (II−3a−384well plate)
P( −0.288 < μ1 − μ2 < 0.059 ) = 0.95

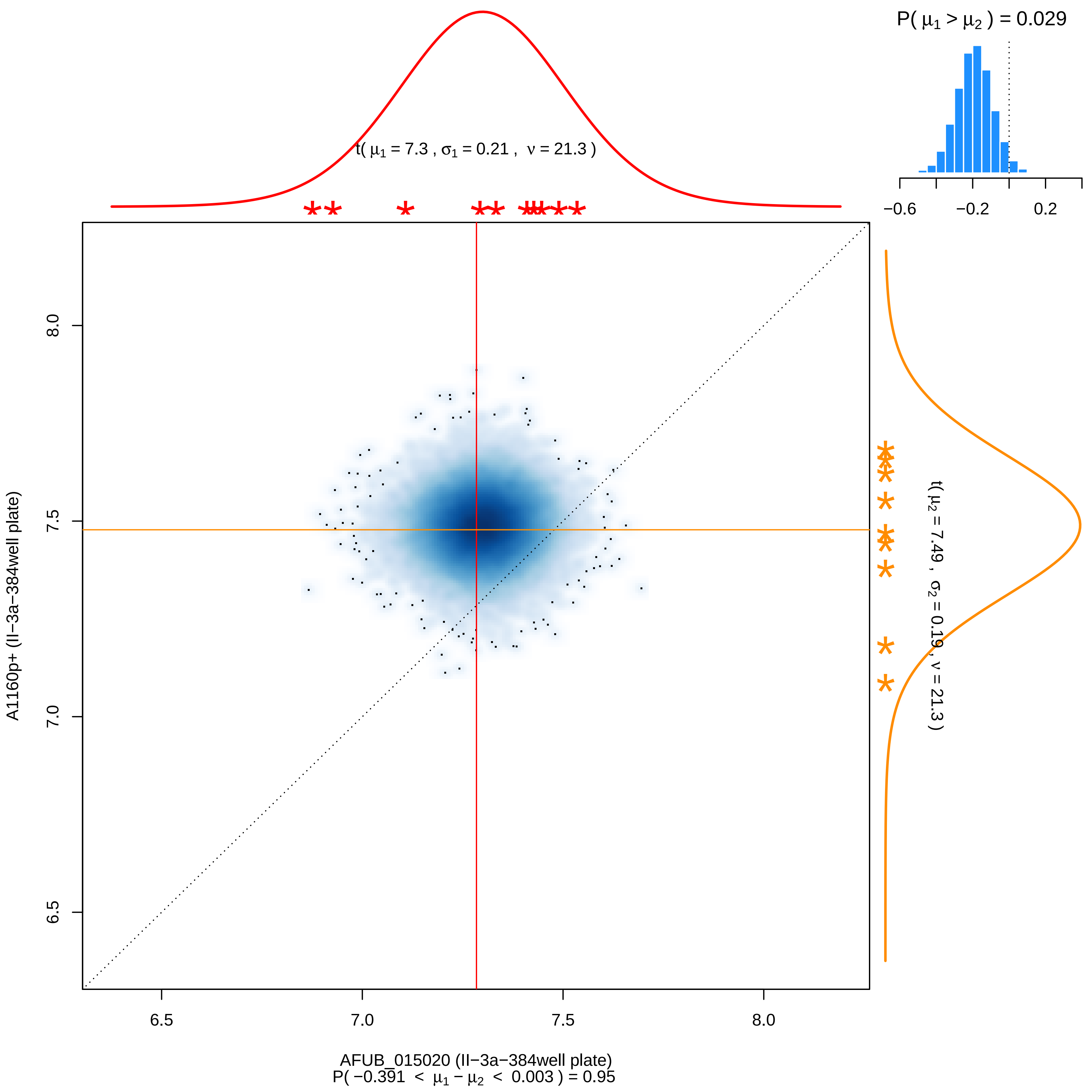

P( μ1 > μ2 ) = 0.029
−0.6
−0.2
0.2
t( μ1 = 7.3 , σ1 = 0.21 , ν = 21.3 )
t( μ2 = 7.49 , σ2 = 0.19 , ν = 21.3 )
A1160p+ (II−3a−384well plate)
8.0
7.5
7.0
6.5
AFUB_015020 (II−3a−384well plate)
P( −0.391 < μ1 − μ2 < 0.003 ) = 0.95

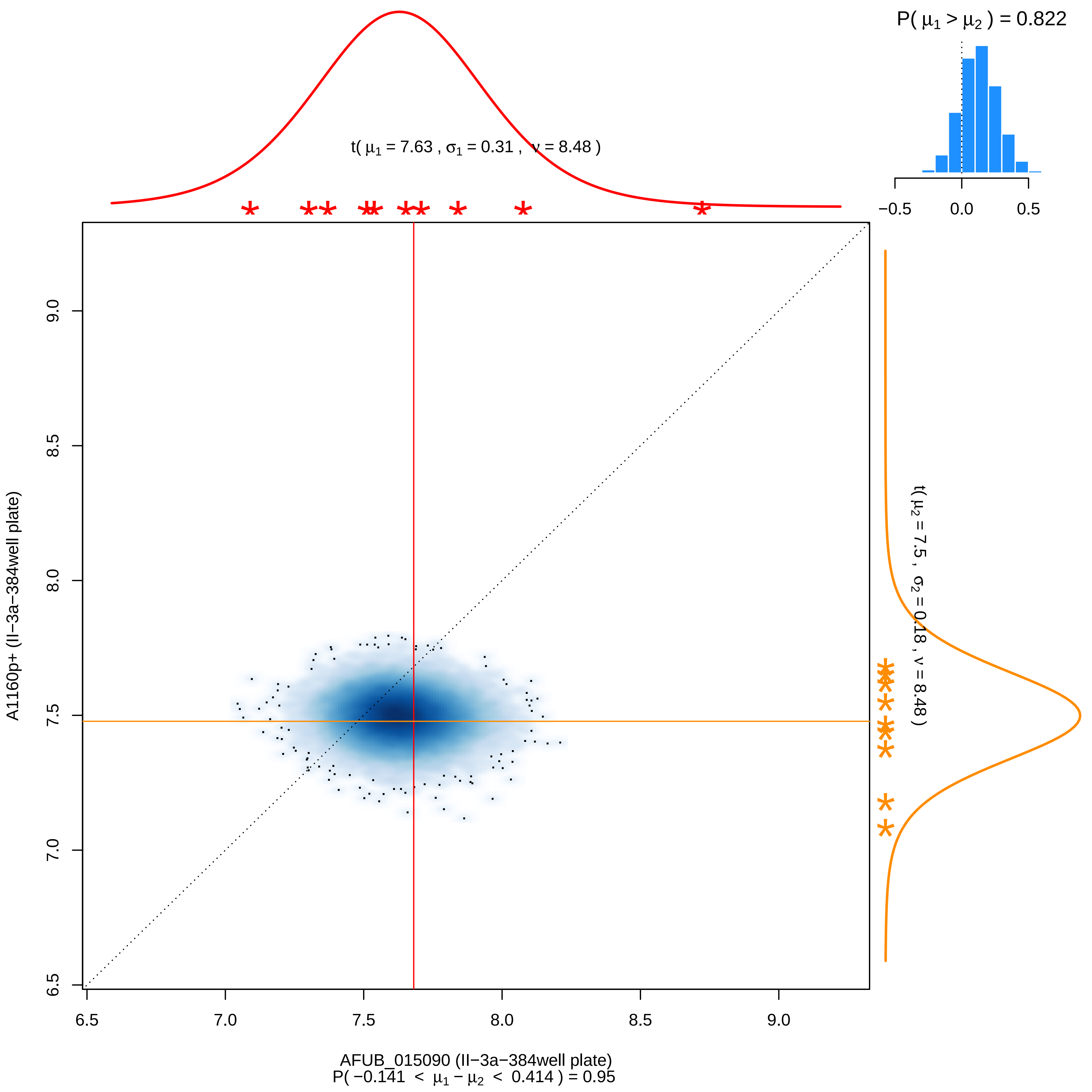

P( μ1 > μ2 ) = 0.822
−0.5
0.0
0.5
t( μ1 = 7.63 , σ1 = 0.31 , ν = 8.48 )
t( μ2 = 7.5 , σ2 = 0.18 , ν = 8.48 )
A1160p+ (II−3a−384well plate)
9.0
8.5
8.0
7.5
7.0
6.5
6.5
7.0
7.5
8.0
8.5
9.0
AFUB_015090 (II−3a−384well plate)
P( −0.141 < μ1 − μ2 < 0.414 ) = 0.95

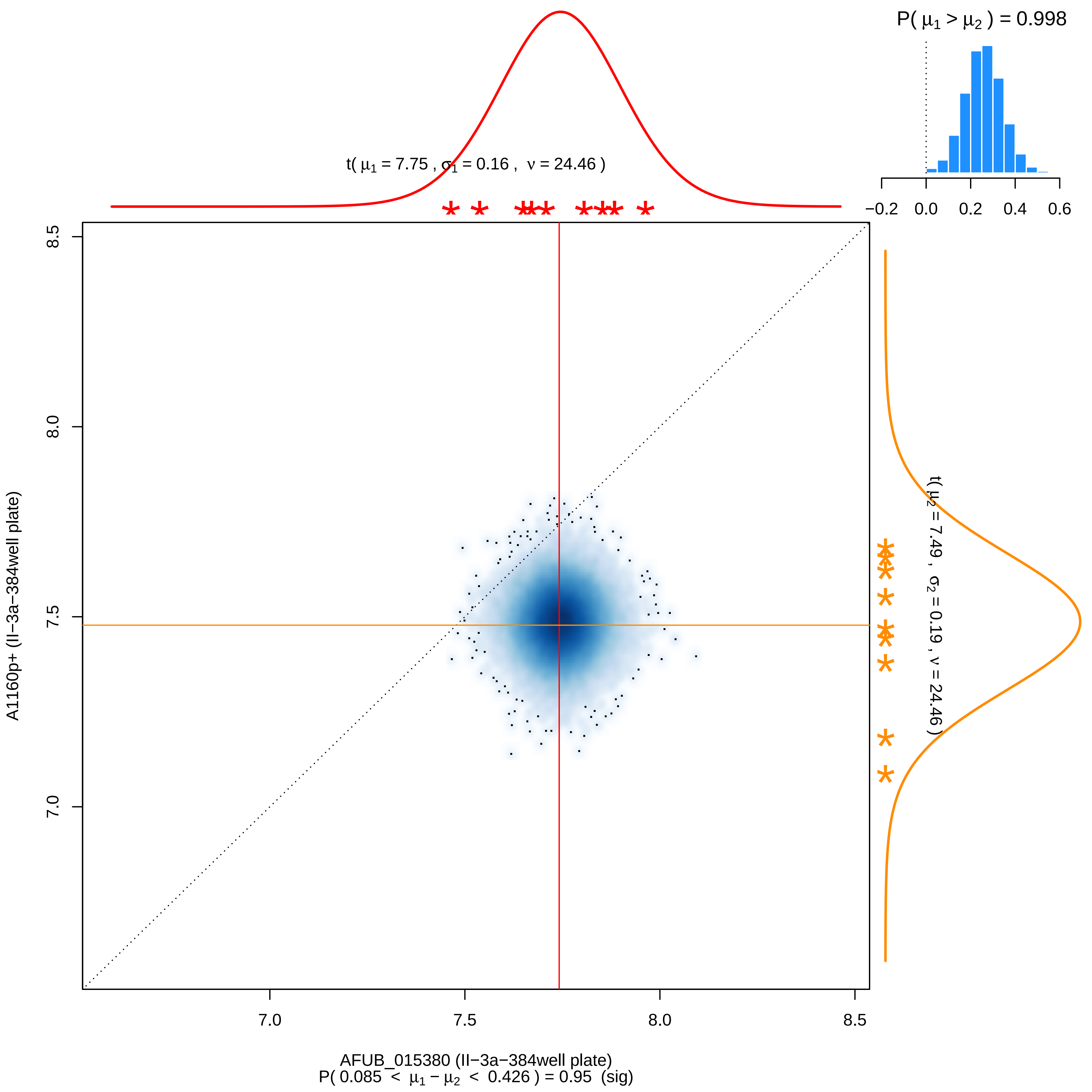

P( μ1 > μ2 ) = 0.998
−0.2 0.0 0.2 0.4 0.6
t( μ1 = 7.75 , σ1 = 0.16 , ν = 24.46 )
t( μ2 = 7.49 , σ2 = 0.19 , ν = 24.46 )
8.5
8.0
7.5
7.0
A1160p+ (II−3a−384well plate)
7.0 7.5 8.0 8.5
AFUB_015380 (II−3a−384well plate)
P( 0.085 < μ1 − μ2 < 0.426 ) = 0.95 (sig)

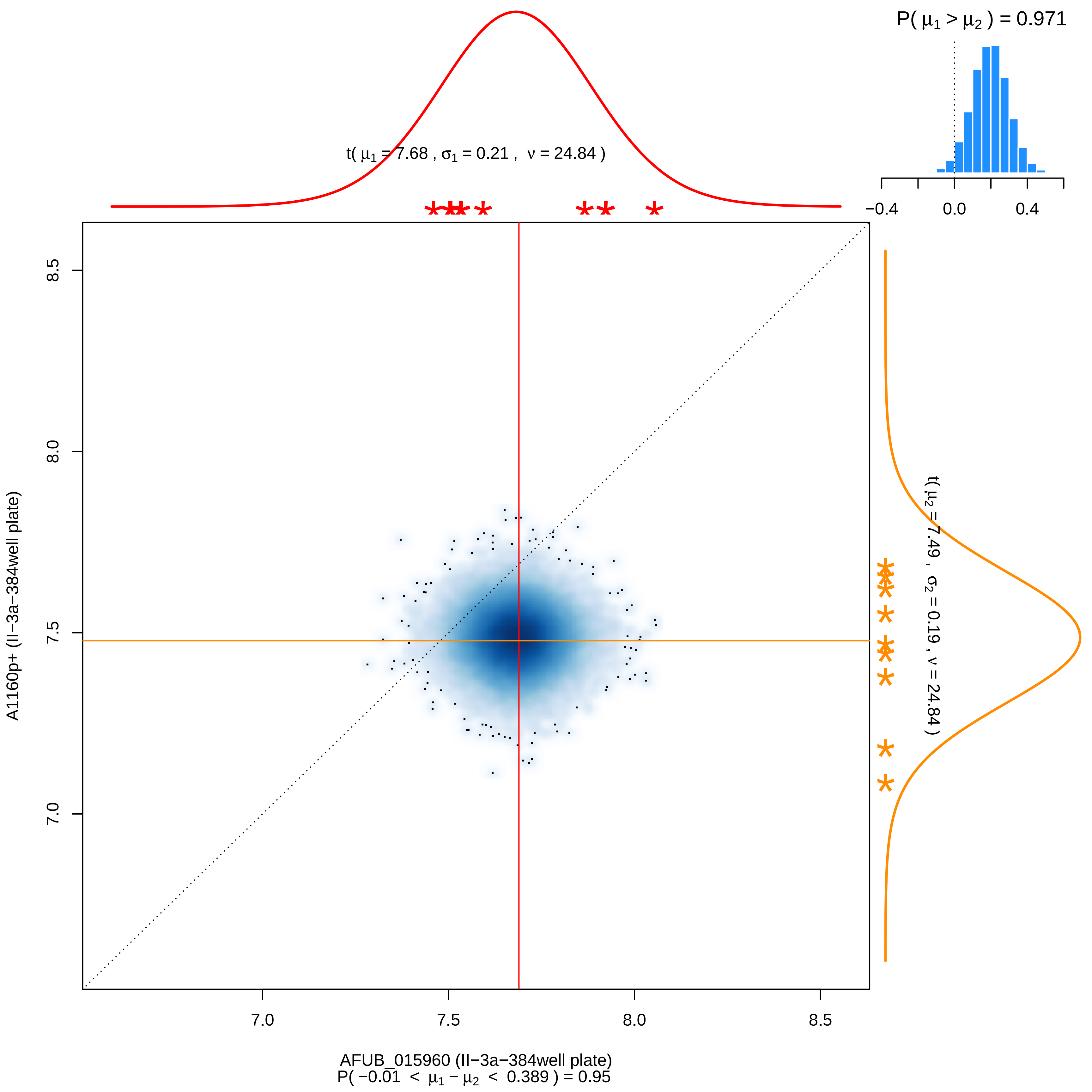

P( $\mu_1 > \mu_2$ ) = 0.971
−0.4
0.0
0.4
t( $\mu_1$ = 7.68 , $\sigma_1$ = 0.21 , $\nu$ = 24.84 )
t( $\mu_2$ = 7.49 , $\sigma_2$ = 0.19 , $\nu$ = 24.84 )
8.5
8.0
7.5
7.0
A1160p+ (II−3a−384well plate)
7.0
7.5
8.0
8.5
AFUB_015960 (II−3a−384well plate)
P( −0.01 < $\mu_1 - \mu_2$ < 0.389 ) = 0.95

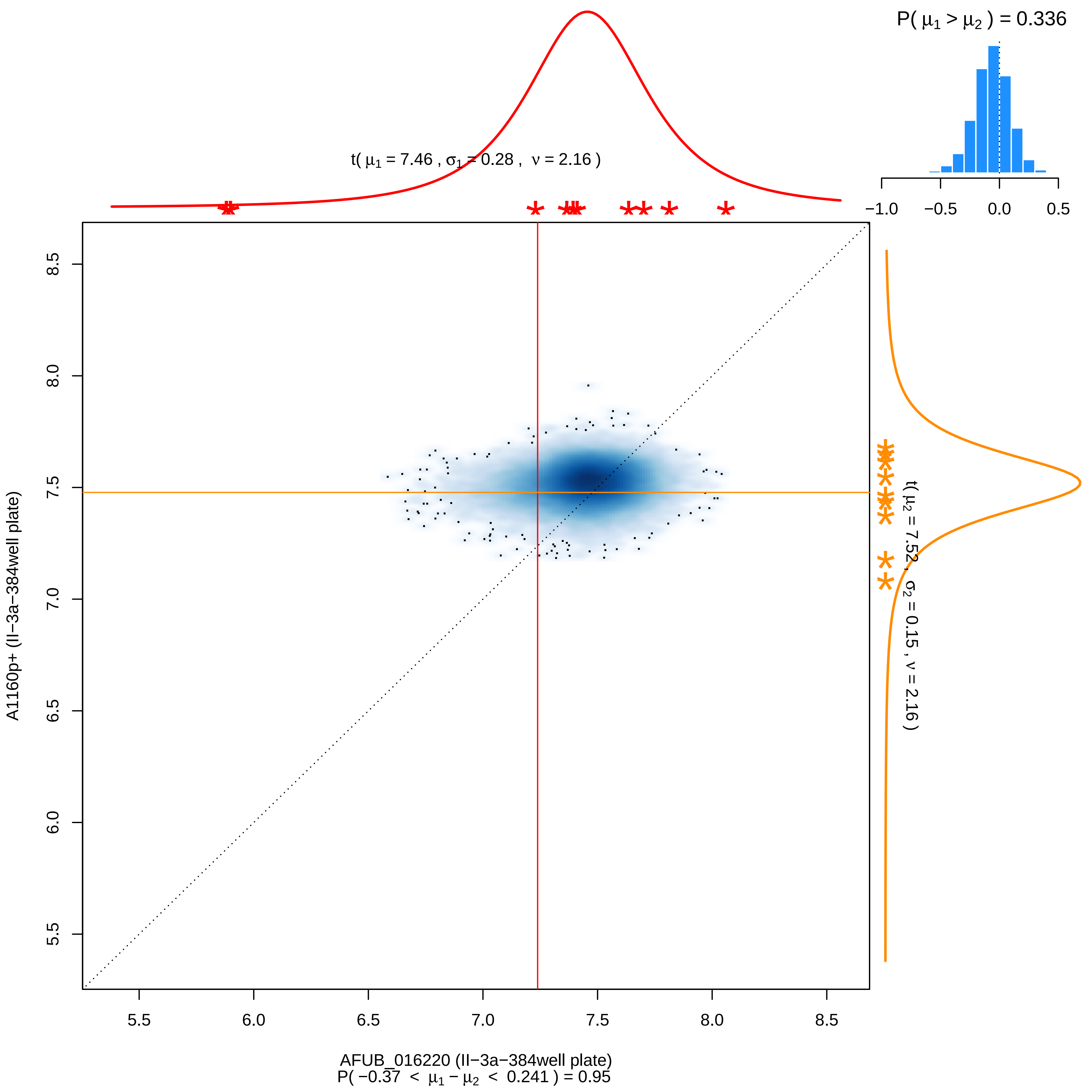
P( $\mu_1 > \mu_2$ ) = 0.336
−1.0 −0.5 0.0 0.5
t( $\mu_1$ = 7.46 , $\sigma_1$ = 0.28 , $\nu$ = 2.16 )
t( $\mu_2$ = 7.52 , $\sigma_2$ = 0.15 , $\nu$ = 2.16 )
A1160p+ (II−3a−384well plate)
5.5 6.0 6.5 7.0 7.5 8.0 8.5
AFUB_016220 (II−3a−384well plate)
P( −0.37 < $\mu_1 - \mu_2$ < 0.241 ) = 0.95

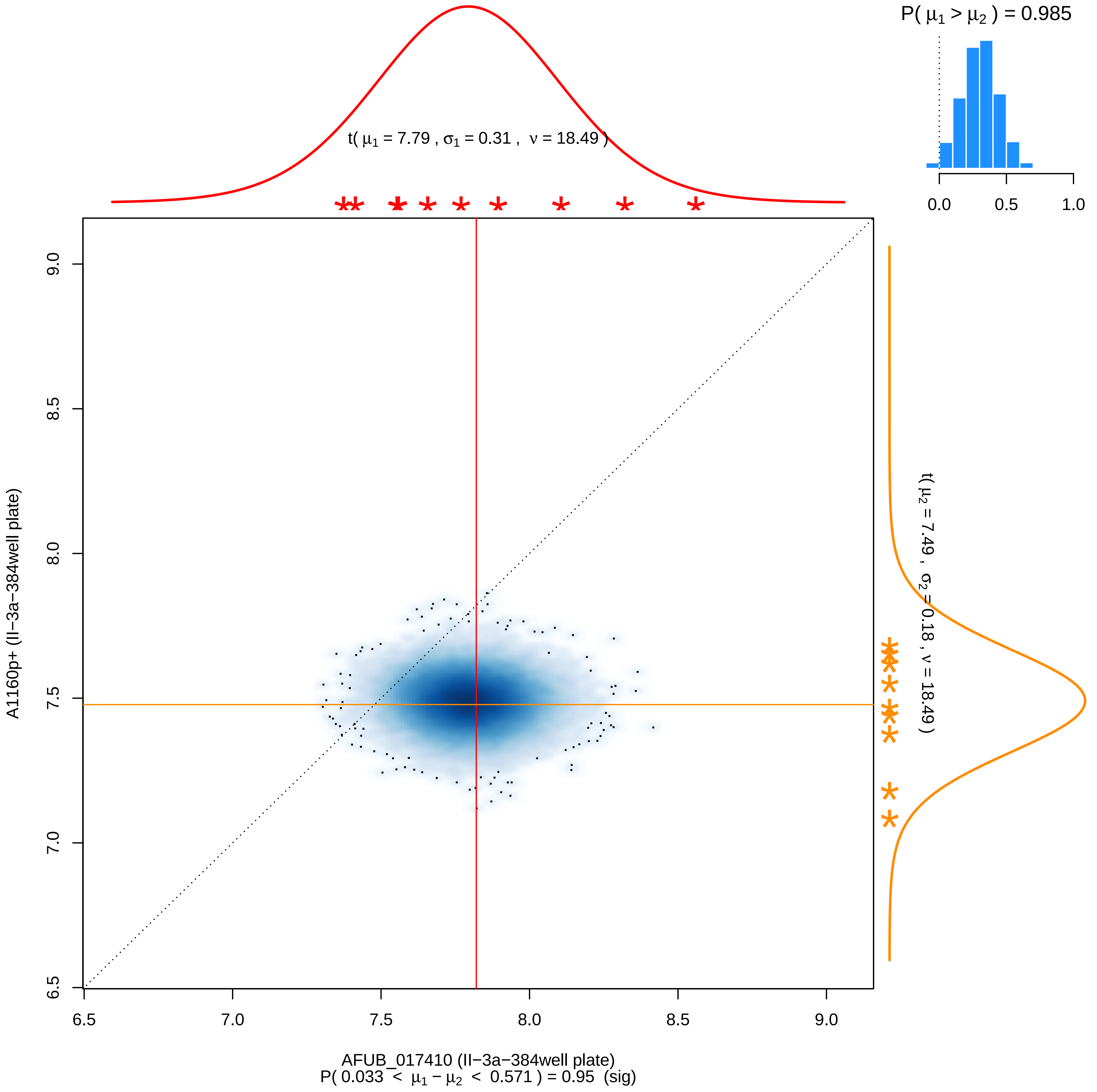

P( μ1 > μ2 ) = 0.985
0.0
0.5
1.0
t( μ1 = 7.79 , σ1 = 0.31 , ν = 18.49 )
t( μ2 = 7.49 , σ2 = 0.18 , ν = 18.49 )
A1160p+ (II−3a−384well plate)
9.0
8.5
8.0
7.5
7.0
6.5
6.5
7.0
7.5
8.0
8.5
9.0
AFUB_017410 (II−3a−384well plate)
P( 0.033 < μ1 − μ2 < 0.571 ) = 0.95 (sig)

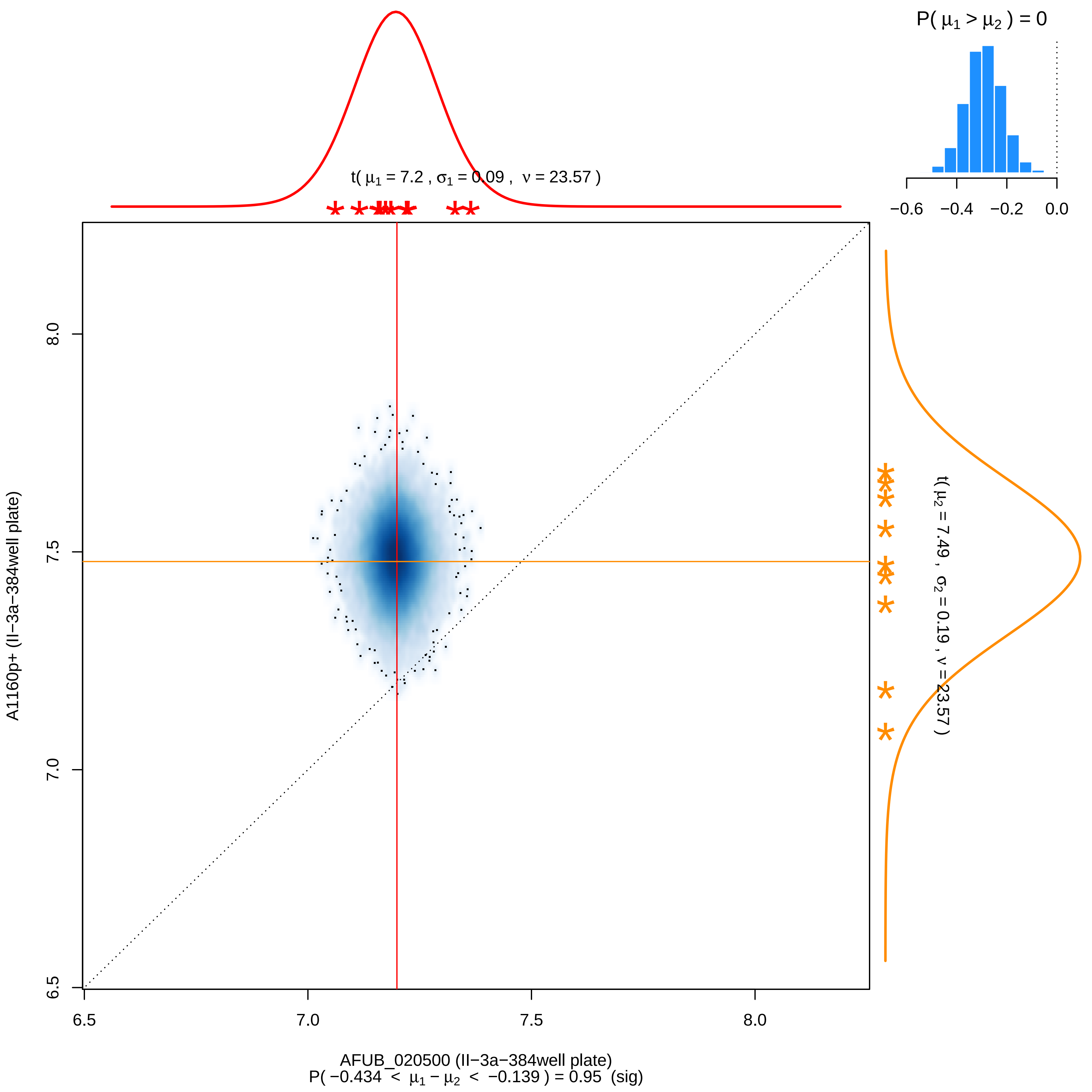

P( μ1 > μ2 ) = 0
−0.6 −0.4 −0.2 0.0
t( μ1 = 7.2 , σ1 = 0.09 , ν = 23.57 )
t( μ2 = 7.49 , σ2 = 0.19 , ν = 23.57 )
A1160p+ (II−3a−384well plate)
6.5 7.0 7.5 8.0
AFUB_020500 (II−3a−384well plate)
P( −0.434 < μ1 − μ2 < −0.139 ) = 0.95 (sig)

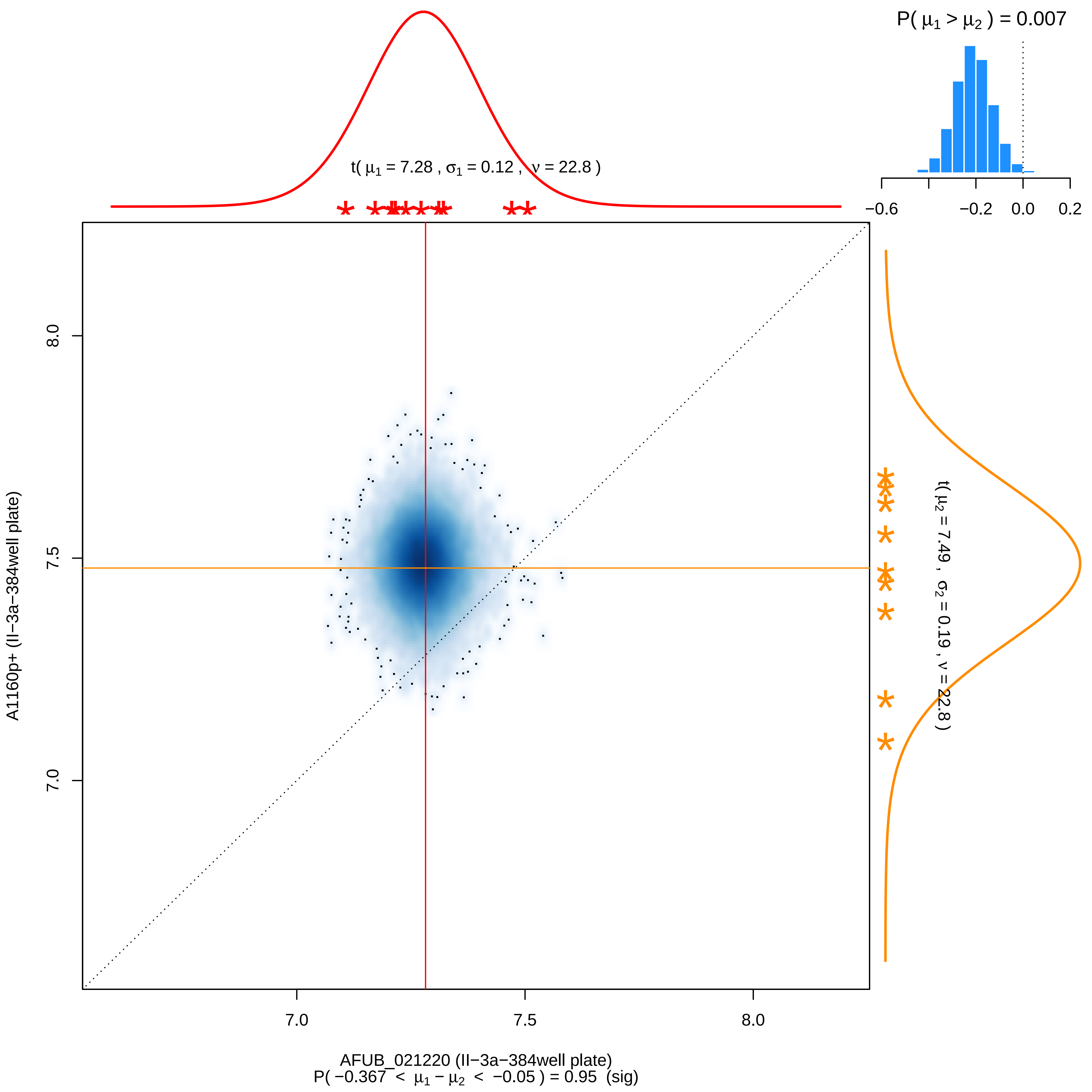
P( μ1 > μ2 ) = 0.007
−0.6
−0.2
0.0
0.2
t( μ1 = 7.28 , σ1 = 0.12 , ν = 22.8 )
t( μ2 = 7.49 , σ2 = 0.19 , ν = 22.8 )
A1160p+ (II−3a−384well plate)
8.0
7.5
7.0
AFUB_021220 (II−3a−384well plate)
P( −0.367 < μ1 − μ2 < −0.05 ) = 0.95 (sig)

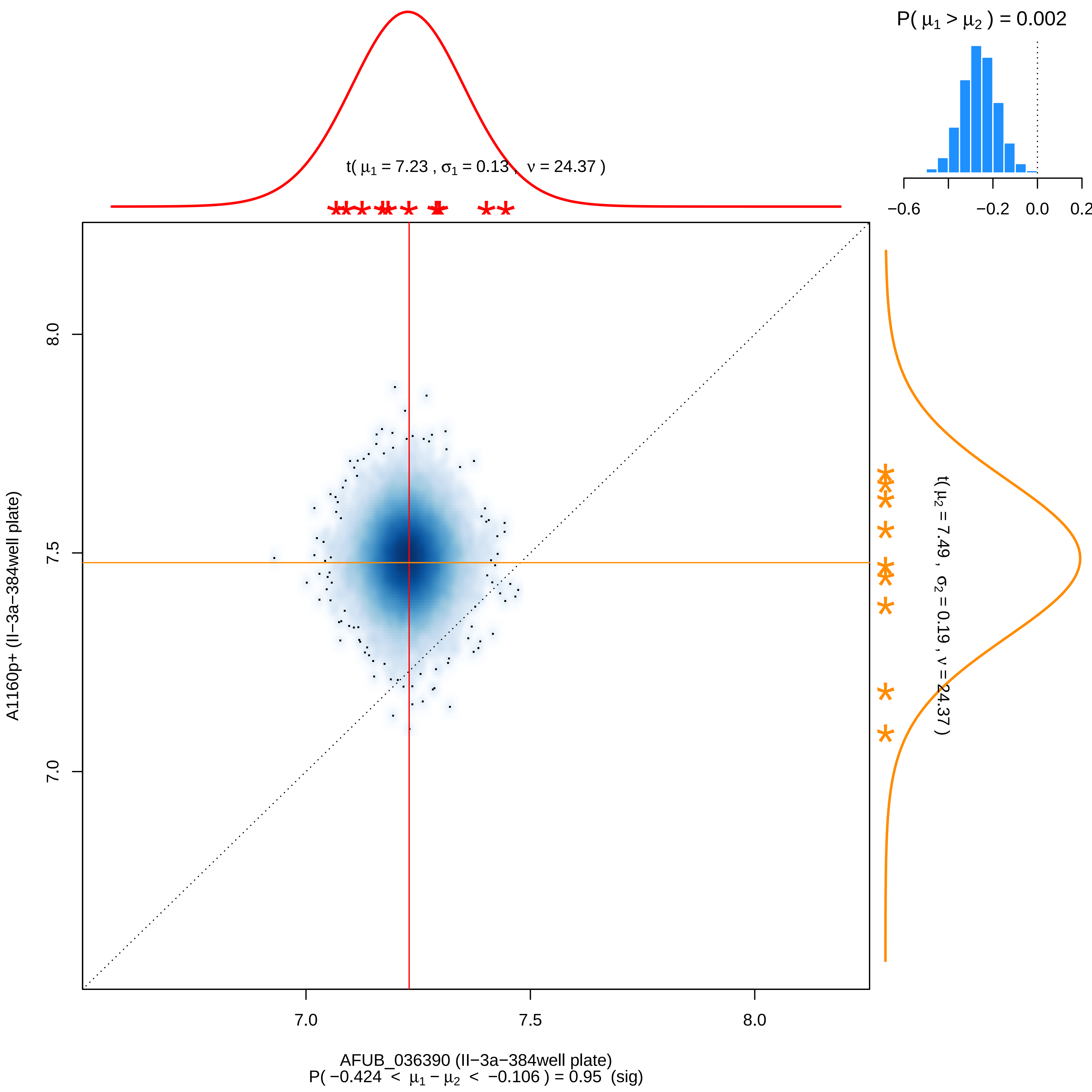
P( μ1 > μ2 ) = 0.002
−0.6
−0.2
0.0
0.2
t( μ1 = 7.23 , σ1 = 0.13 , ν = 24.37 )
t( μ2 = 7.49 , σ2 = 0.19 , ν = 24.37 )
A1160p+ (II−3a−384well plate)
8.0
7.5
7.0
7.0
7.5
8.0
AFUB_036390 (II−3a−384well plate)
P( −0.424 < μ1 − μ2 < −0.106 ) = 0.95 (sig)

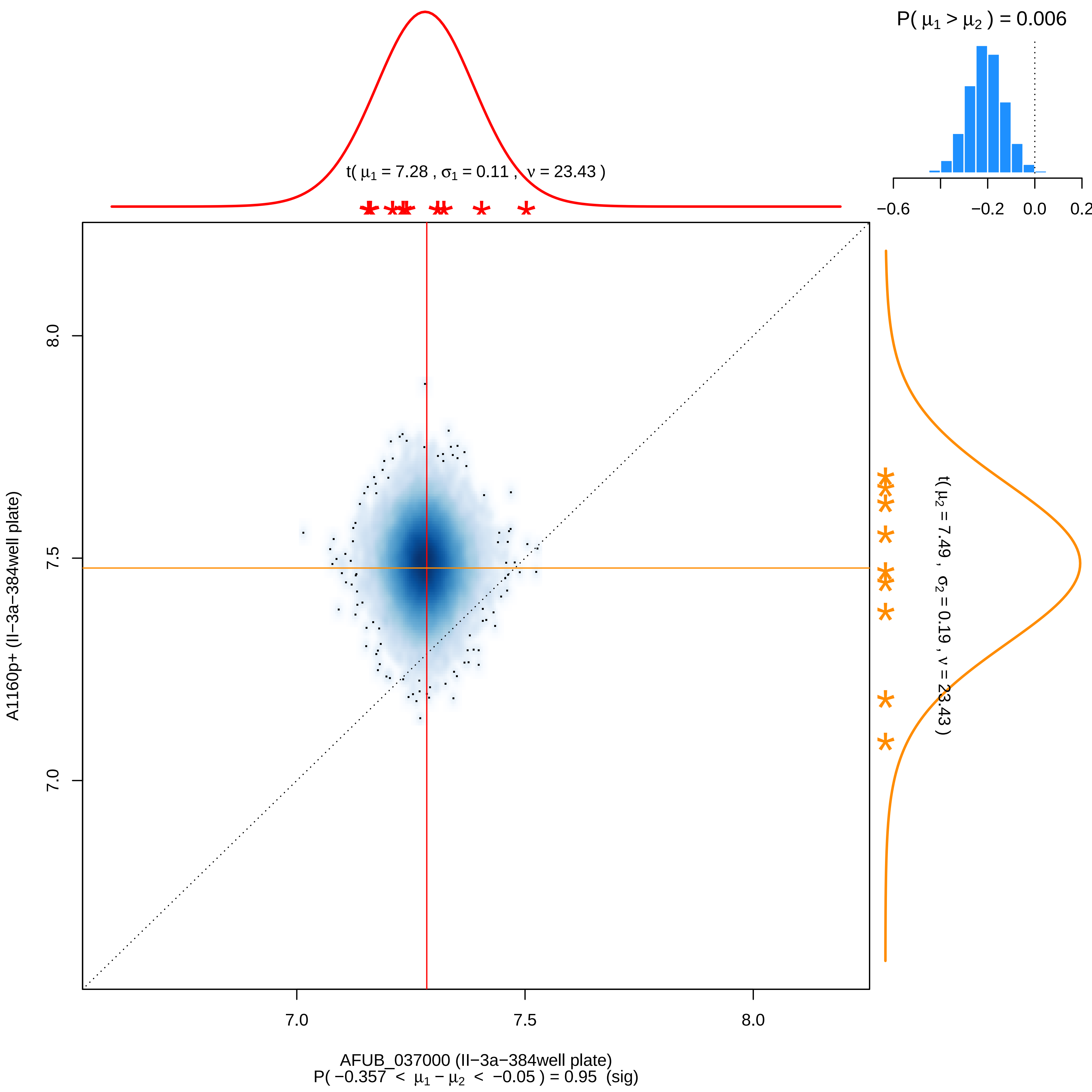

P( μ1 > μ2 ) = 0.006
−0.6
−0.2
0.0
0.2
t( μ1 = 7.28 , σ1 = 0.11 , ν = 23.43 )
t( μ2 = 7.49 , σ2 = 0.19 , ν = 23.43 )
8.0
7.5
7.0
A1160p+ (II−3a−384well plate)
7.0
7.5
8.0
AFUB_037000 (II−3a−384well plate)
P( −0.357 < μ1 − μ2 < −0.05 ) = 0.95 (sig)

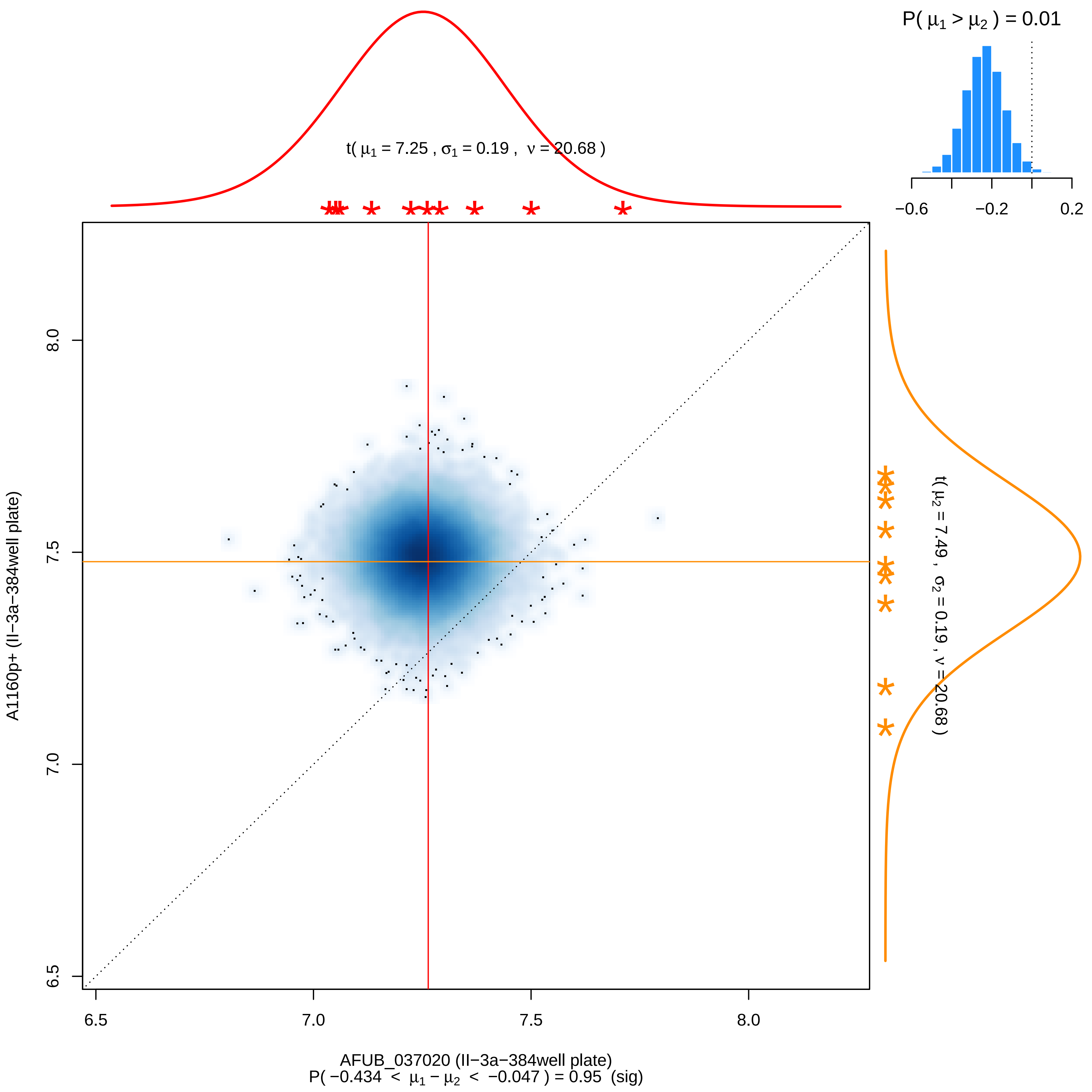

P( μ1 > μ2 ) = 0.01
−0.6
−0.2
0.2
t( μ1 = 7.25 , σ1 = 0.19 , ν = 20.68 )
t( μ2 = 7.49 , σ2 = 0.19 , ν = 20.68 )
A1160p+ (II−3a−384well plate)
8.0
7.5
7.0
6.5
6.5
7.0
7.5
8.0
AFUB_037020 (II−3a−384well plate)
P( −0.434 < μ1 − μ2 < −0.047 ) = 0.95 (sig)

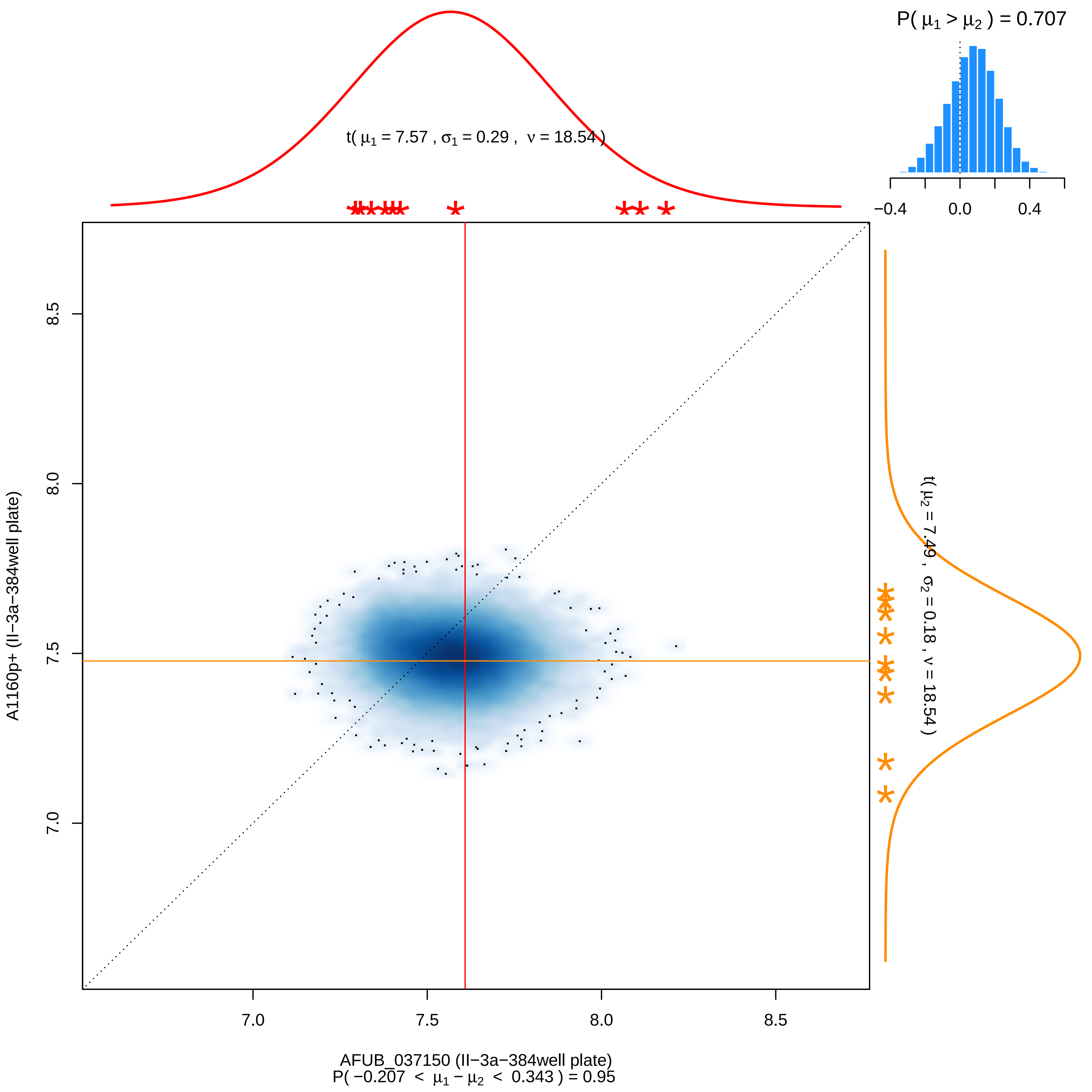
P( μ1 > μ2 ) = 0.707
−0.4
0.0
0.4
t( μ1 = 7.57 , σ1 = 0.29 , ν = 18.54 )
t( μ2 = 7.49 , σ2 = 0.18 , ν = 18.54 )
8.5
8.0
7.5
7.0
A1160p+ (II−3a−384well plate)
7.0
7.5
8.0
8.5
AFUB_037150 (II−3a−384well plate)
P( −0.207 < μ1 − μ2 < 0.343 ) = 0.95

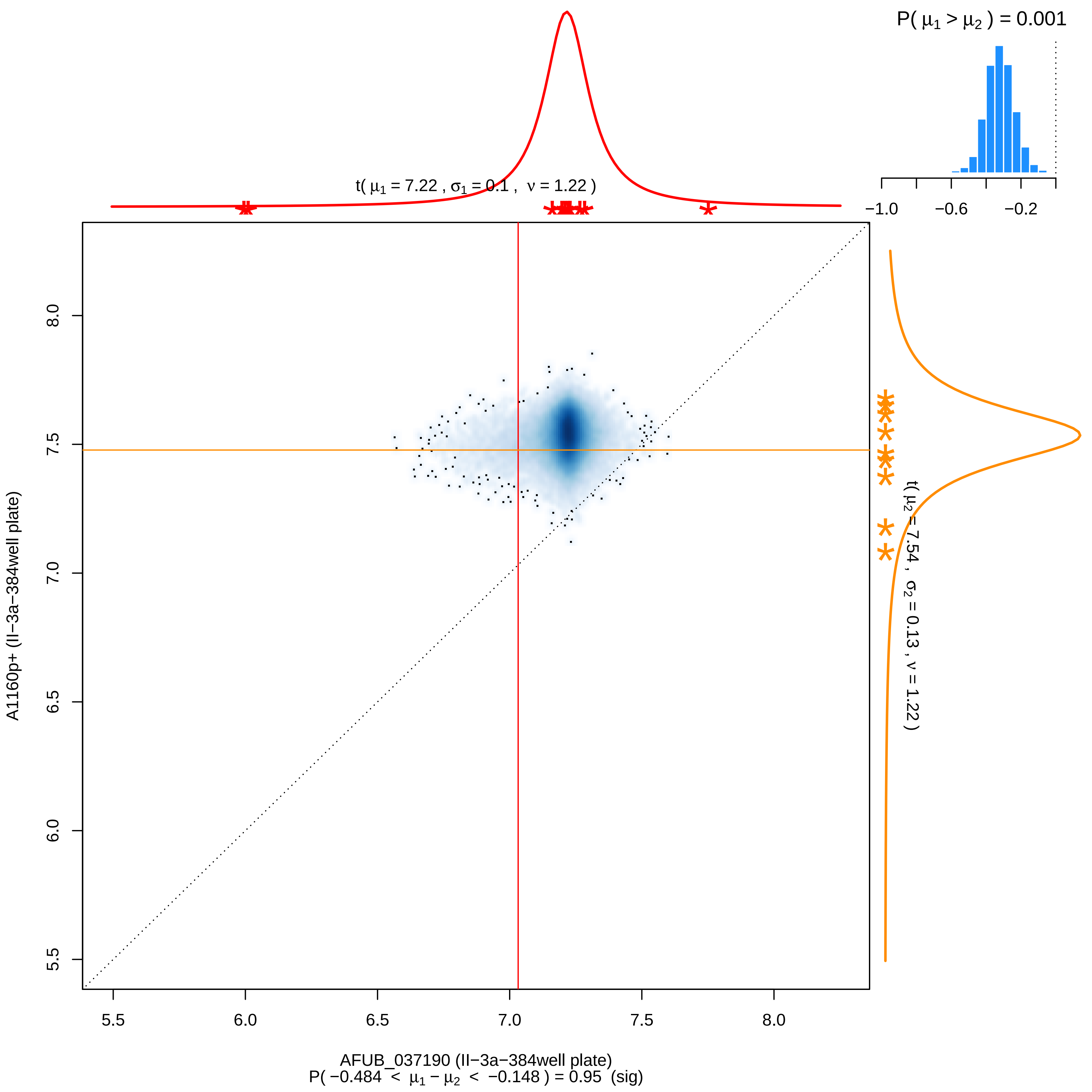

P( μ1 > μ2 ) = 0.001
−1.0
−0.6
−0.2
t( μ1 = 7.22 , σ1 = 0.1 , ν = 1.22 )
t( μ2 = 7.54 , σ2 = 0.13 , ν = 1.22 )
A1160p+ (II−3a−384well plate)
8.0
7.5
7.0
6.5
6.0
5.5
AFUB_037190 (II−3a−384well plate)
P( −0.484 < μ1 − μ2 < −0.148 ) = 0.95 (sig)

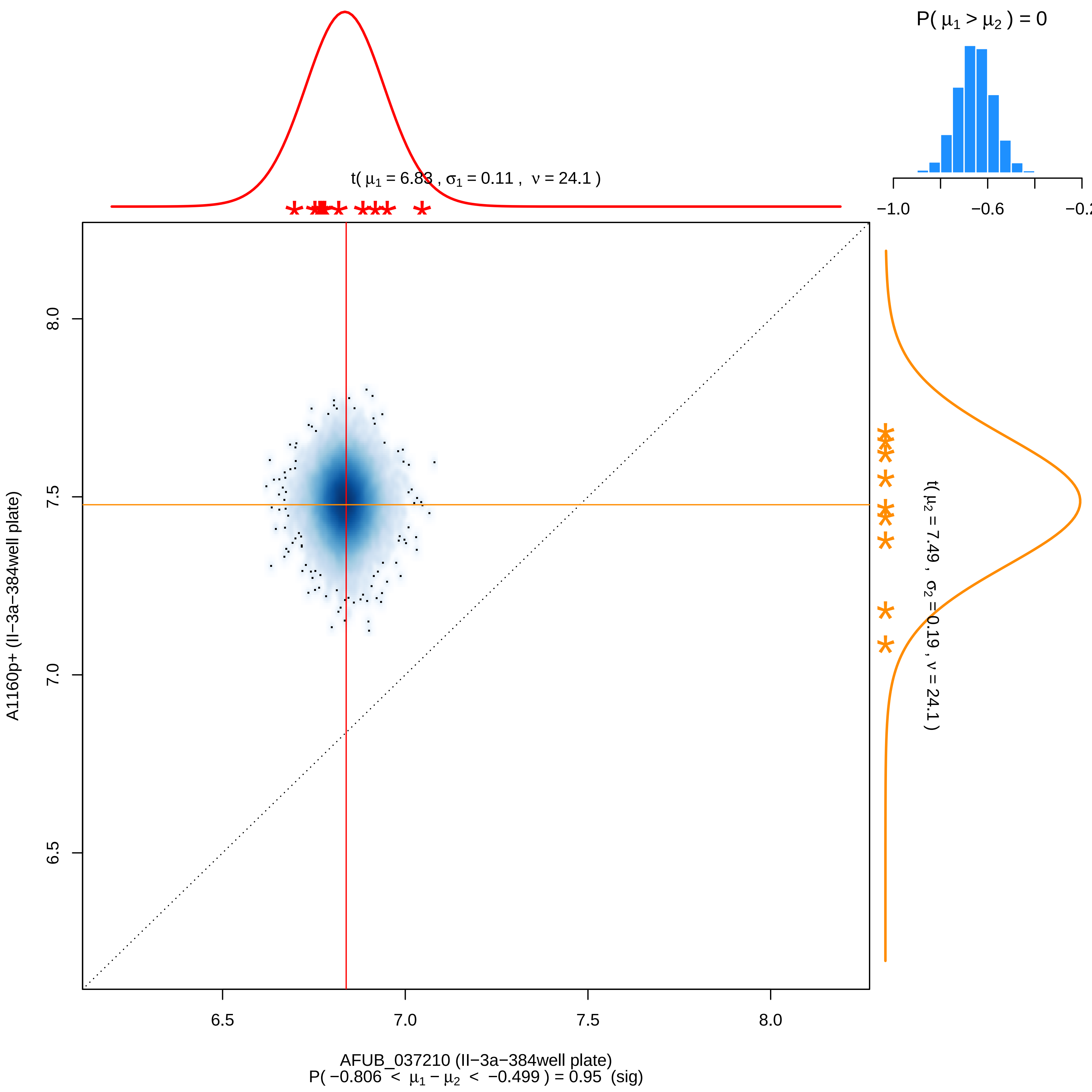

P( μ1 > μ2 ) = 0
−1.0
−0.6
−0.2
t( μ1 = 6.83 , σ1 = 0.11 , ν = 24.1 )
t( μ2 = 7.49 , σ2 = 0.19 , ν = 24.1 )
A1160p+ (II−3a−384well plate)
8.0
7.5
7.0
6.5
AFUB_037210 (II−3a−384well plate)
P( −0.806 < μ1 − μ2 < −0.499 ) = 0.95 (sig)

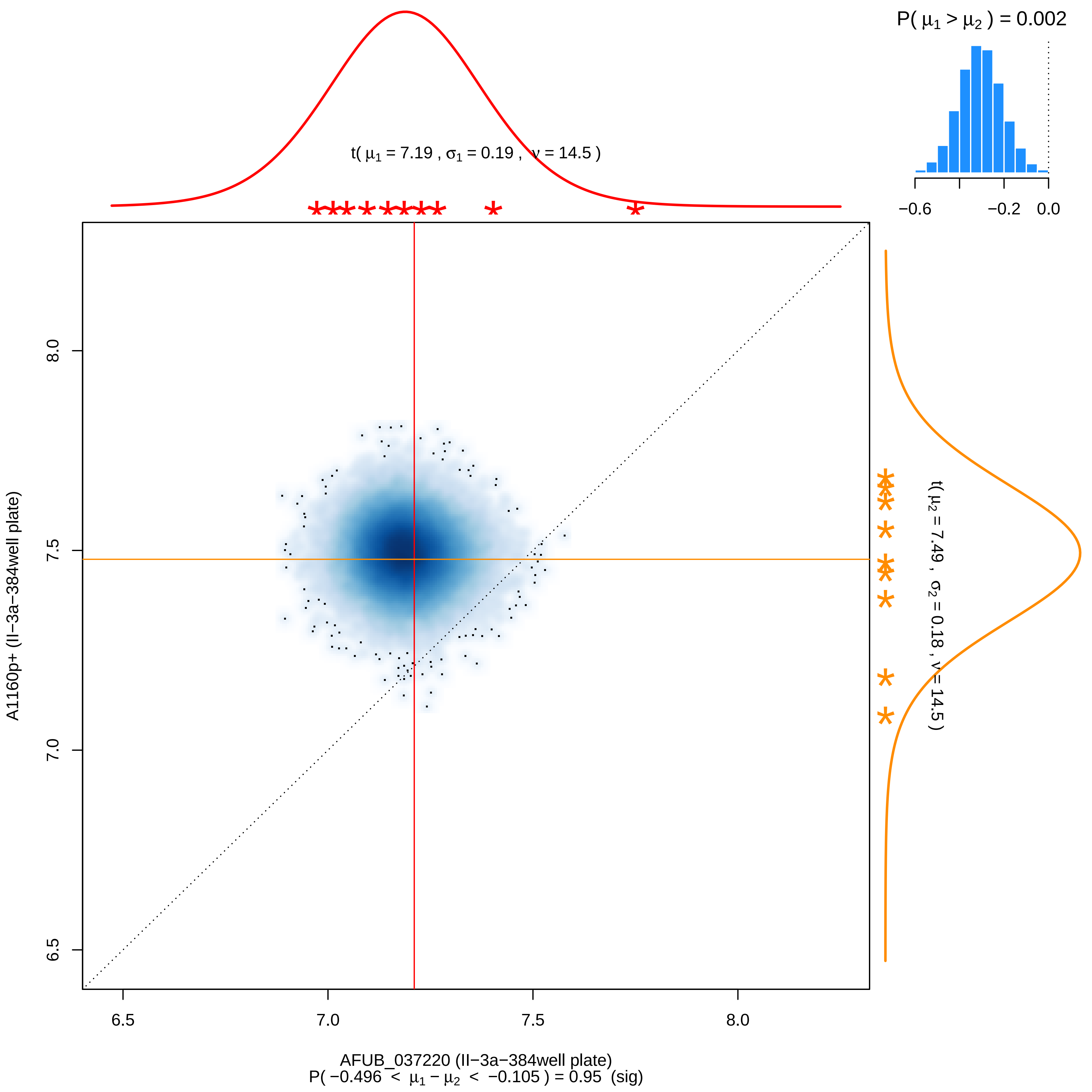

P( μ1 > μ2 ) = 0.002
−0.6
−0.2
0.0
t( μ1 = 7.19 , σ1 = 0.19 , ν = 14.5 )
t( μ2 = 7.49 , σ2 = 0.18 , ν = 14.5 )
A1160p+ (II−3a−384well plate)
6.5
7.0
7.5
8.0
AFUB_037220 (II−3a−384well plate)
P( −0.496 < μ1 − μ2 < −0.105 ) = 0.95 (sig)

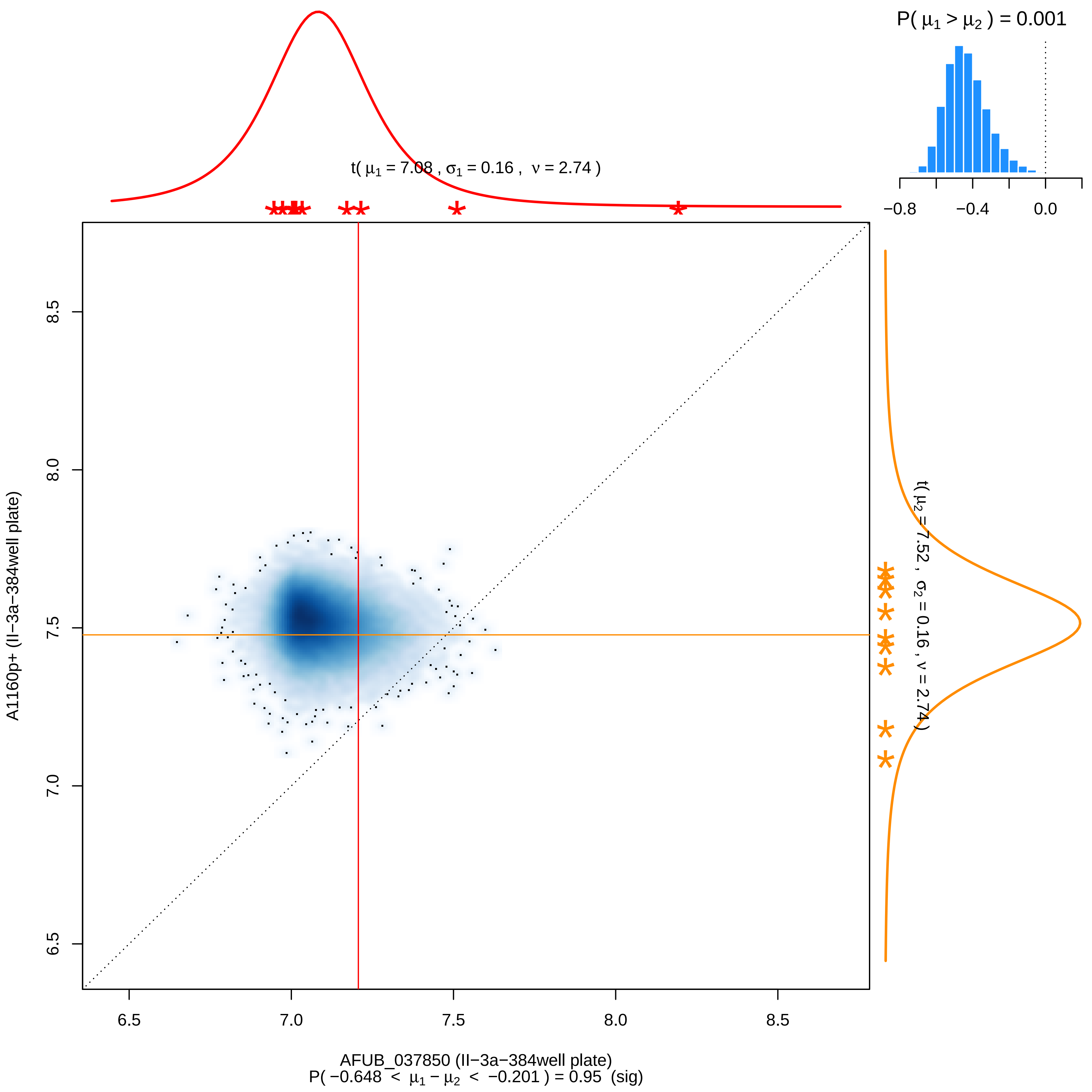

P( μ1 > μ2 ) = 0.001
−0.8
−0.4
0.0
t( μ1 = 7.08 , σ1 = 0.16 , ν = 2.74 )
t( μ2 = 7.52 , σ2 = 0.16 , ν = 2.74 )
6.5
7.0
7.5
8.0
8.5
A1160p+ (II−3a−384well plate)
AFUB_037850 (II−3a−384well plate)
P( −0.648 < μ1 − μ2 < −0.201 ) = 0.95 (sig)

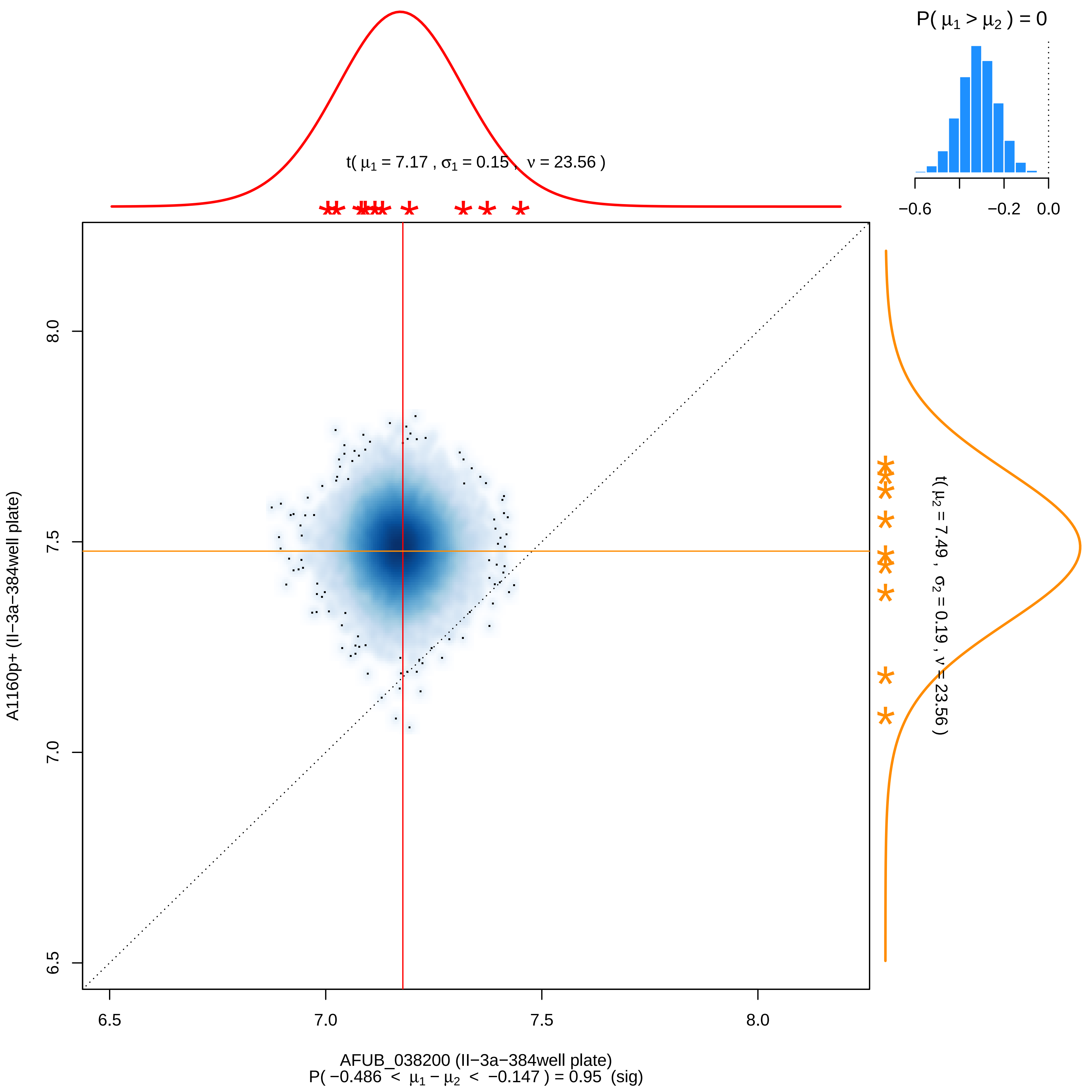

P( μ1 > μ2 ) = 0
−0.6
−0.2
0.0
t( μ1 = 7.17 , σ1 = 0.15 , ν = 23.56 )
t( μ2 = 7.49 , σ2 = 0.19 , ν = 23.56 )
A1160p+ (II−3a−384well plate)
8.0
7.5
7.0
6.5
AFUB_038200 (II−3a−384well plate)
P( −0.486 < μ1 − μ2 < −0.147 ) = 0.95 (sig)

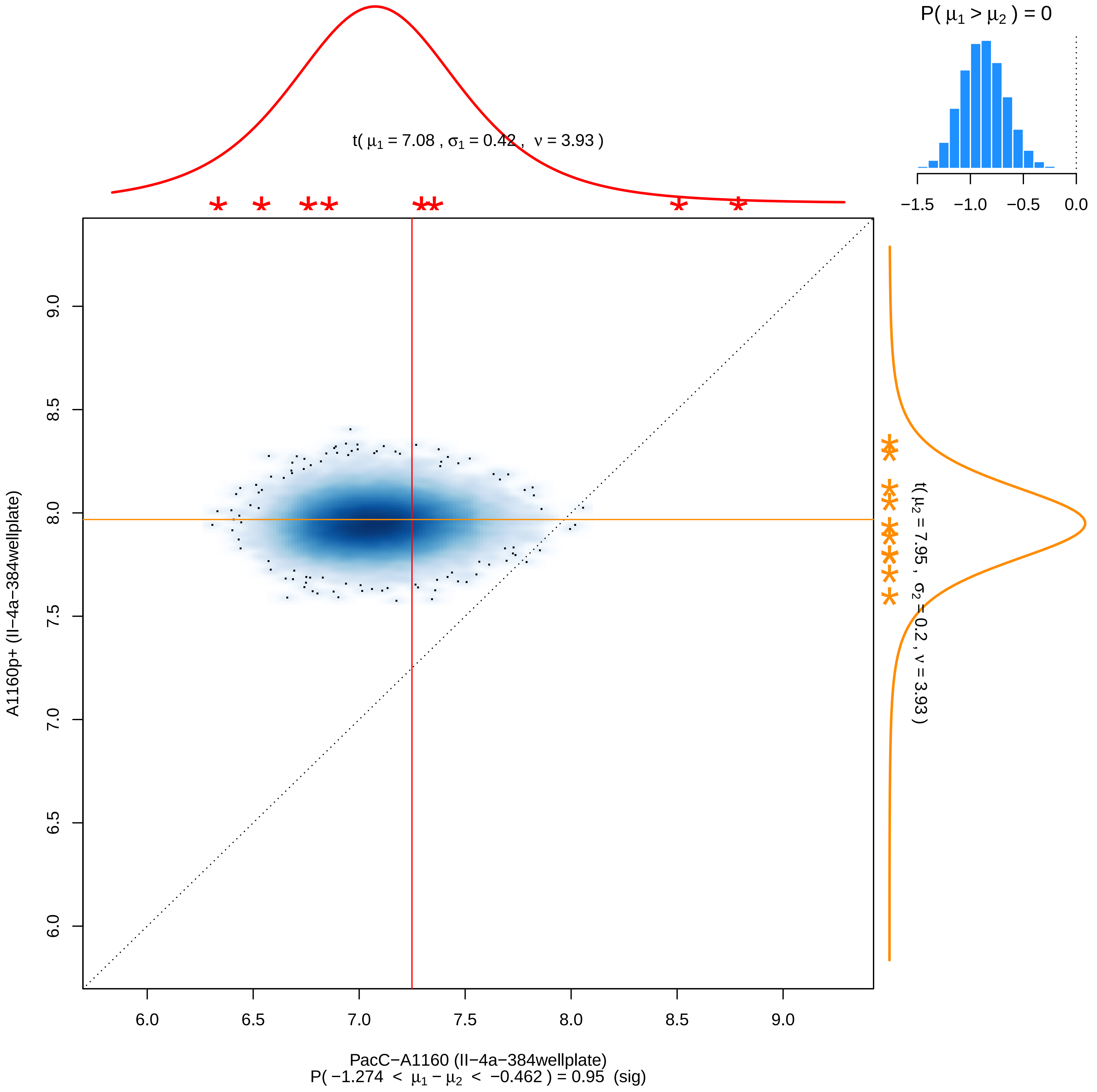
P( μ1 > μ2 ) = 0
−1.5 −1.0 −0.5 0.0
t( μ1 = 7.08 , σ1 = 0.42 , ν = 3.93 )
t( μ2 = 7.95 , σ2 = 0.2 , ν = 3.93 )
A1160p+ (II−4a−384wellplate)
9.0 8.5 8.0 7.5 7.0 6.5 6.0
6.0 6.5 7.0 7.5 8.0 8.5 9.0
PacC−A1160 (II−4a−384wellplate)
P( −1.274 < μ1 − μ2 < −0.462 ) = 0.95 (sig)

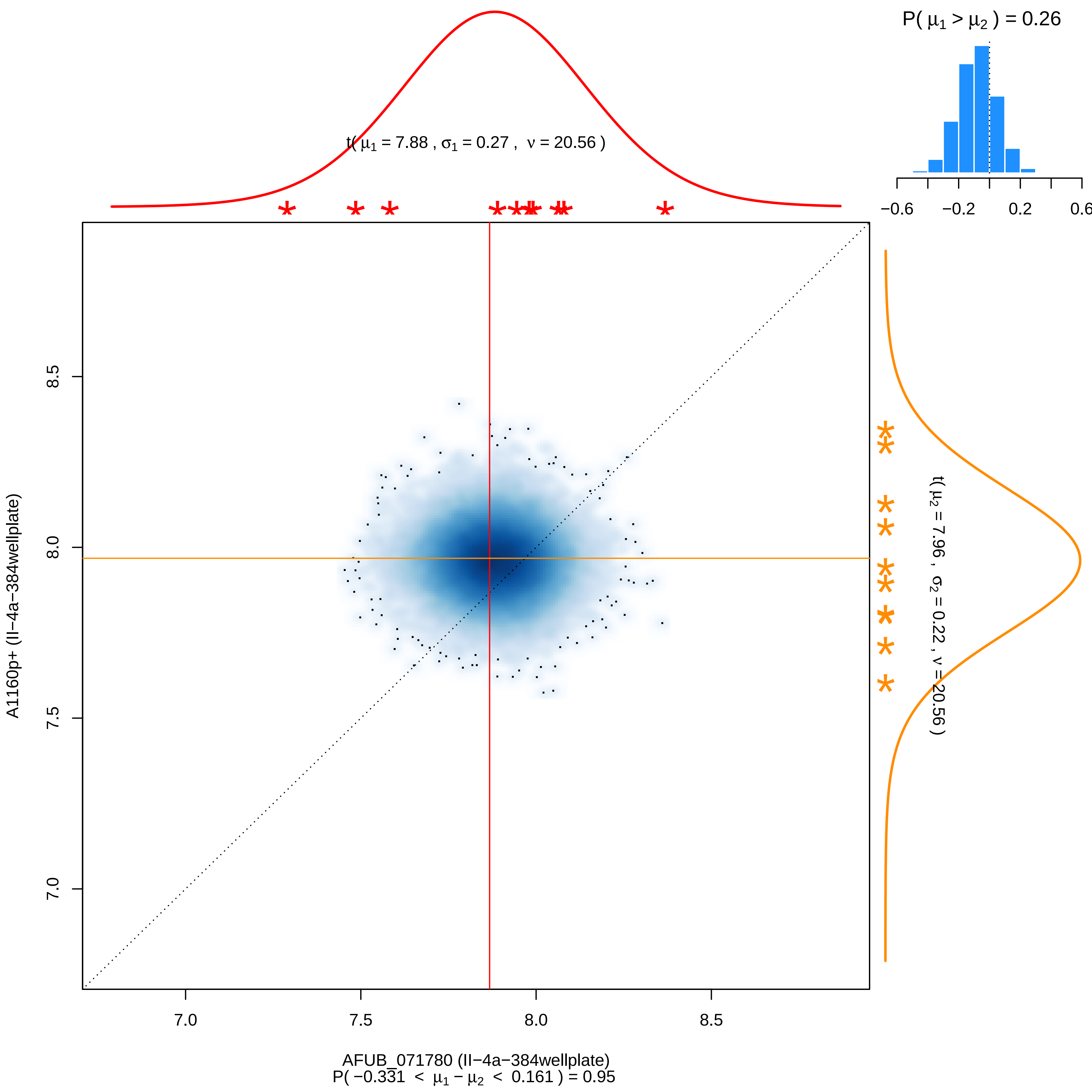

P( μ1 > μ2 ) = 0.26
−0.6
−0.2
0.2
0.6
t( μ1 = 7.88 , σ1 = 0.27 , ν = 20.56 )
t( μ2 = 7.96 , σ2 = 0.22 , ν = 20.56 )
A1160p+ (II−4a−384wellplate)
8.5
8.0
7.5
7.0
AFUB_071780 (II−4a−384wellplate)
P( −0.331 < μ1 − μ2 < 0.161 ) = 0.95

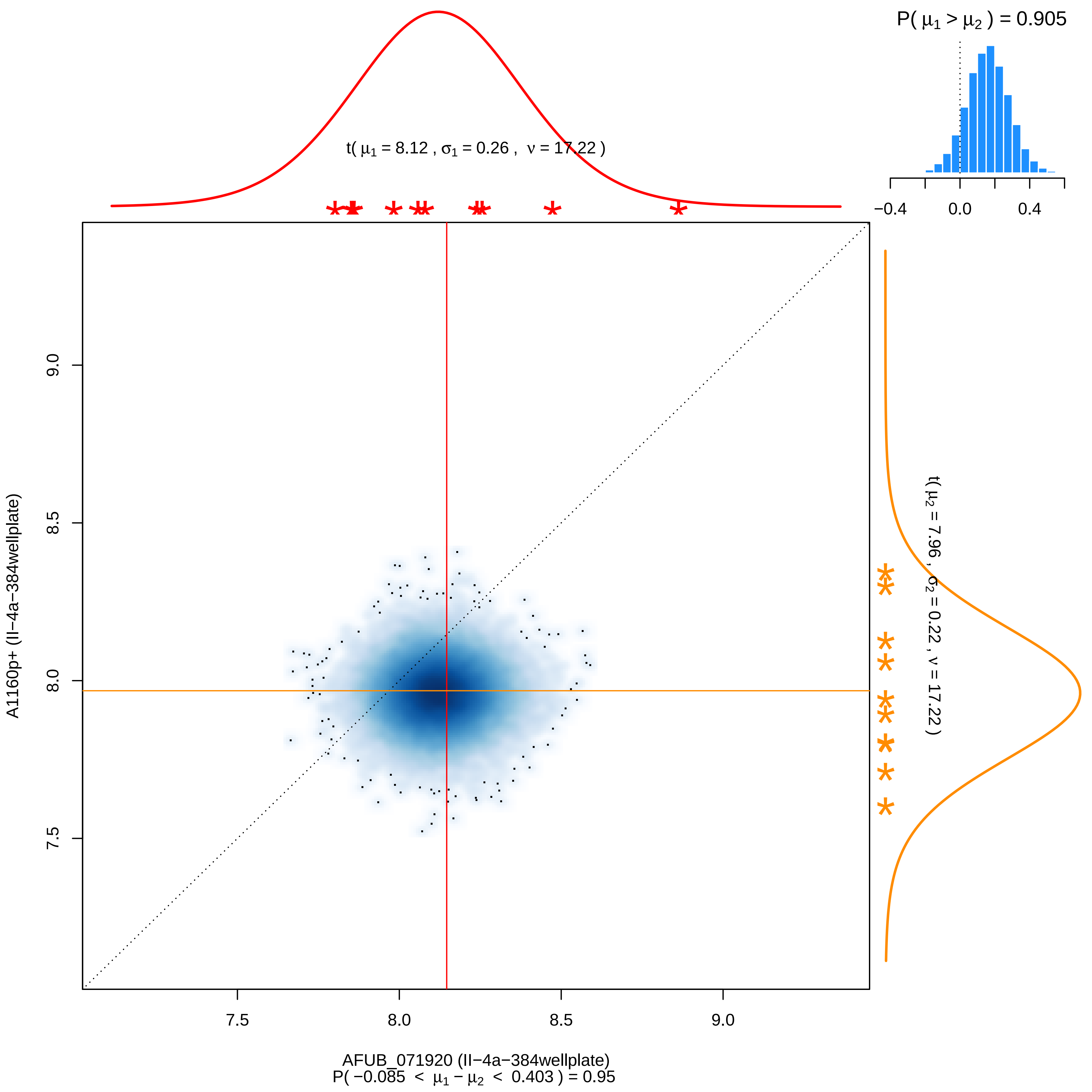

P( μ1 > μ2 ) = 0.905
−0.4
0.0
0.4
t( μ1 = 8.12 , σ1 = 0.26 , ν = 17.22 )
t( μ2 = 7.96 , σ2 = 0.22 , ν = 17.22 )
9.0
8.5
8.0
7.5
A1160p+ (II−4a−384wellplate)
7.5
8.0
8.5
9.0
AFUB_071920 (II−4a−384wellplate)
P( −0.085 < μ1 − μ2 < 0.403 ) = 0.95

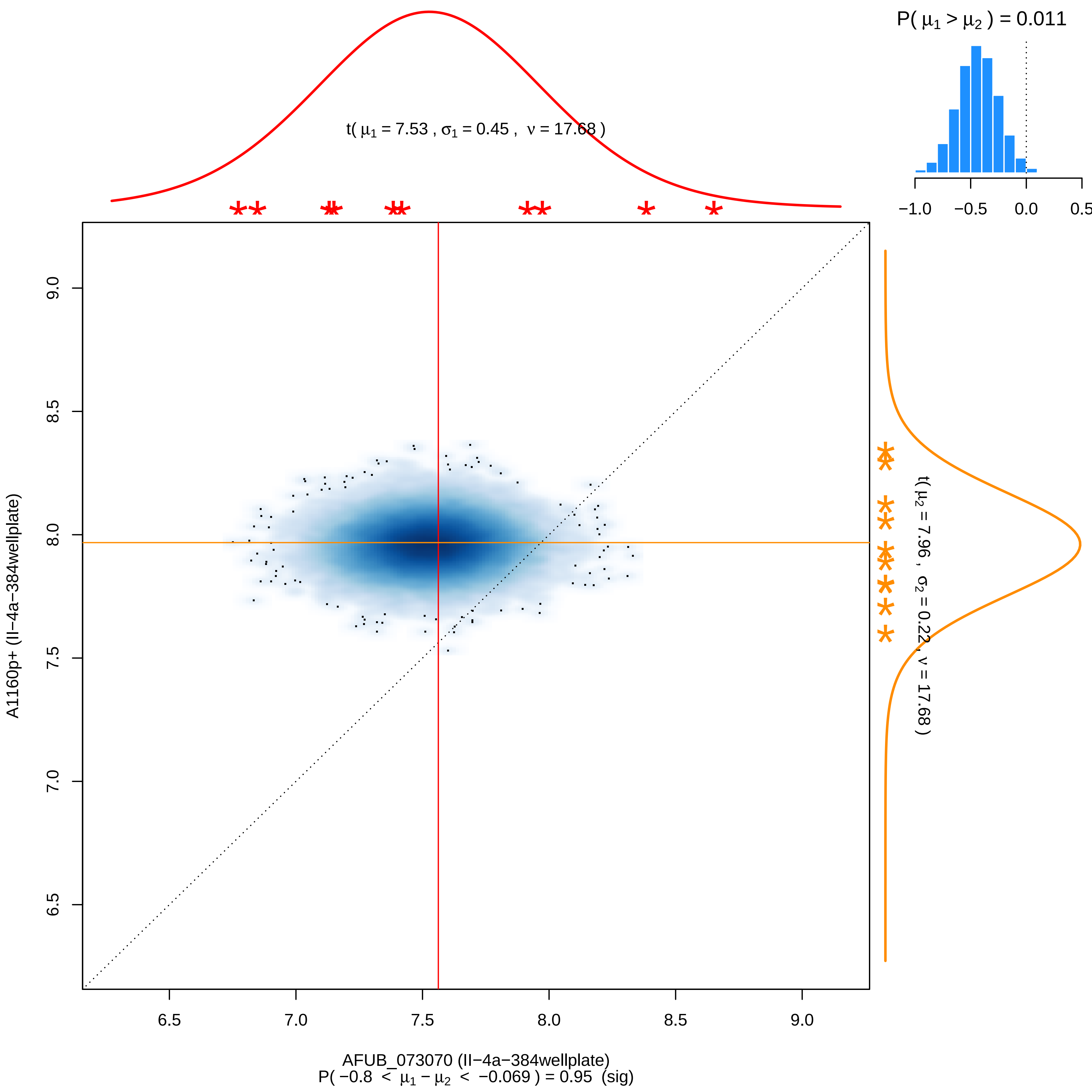

P( μ1 > μ2 ) = 0.011
−1.0
−0.5
0.0
0.5
t( μ1 = 7.53 , σ1 = 0.45 , ν = 17.68 )
t( μ2 = 7.96 , σ2 = 0.22 , ν = 17.68 )
A1160p+ (II−4a−384wellplate)
9.0
8.5
8.0
7.5
7.0
6.5
6.5
7.0
7.5
8.0
8.5
9.0
AFUB_073070 (II−4a−384wellplate)
P( −0.8 < μ1 − μ2 < −0.069 ) = 0.95 (sig)

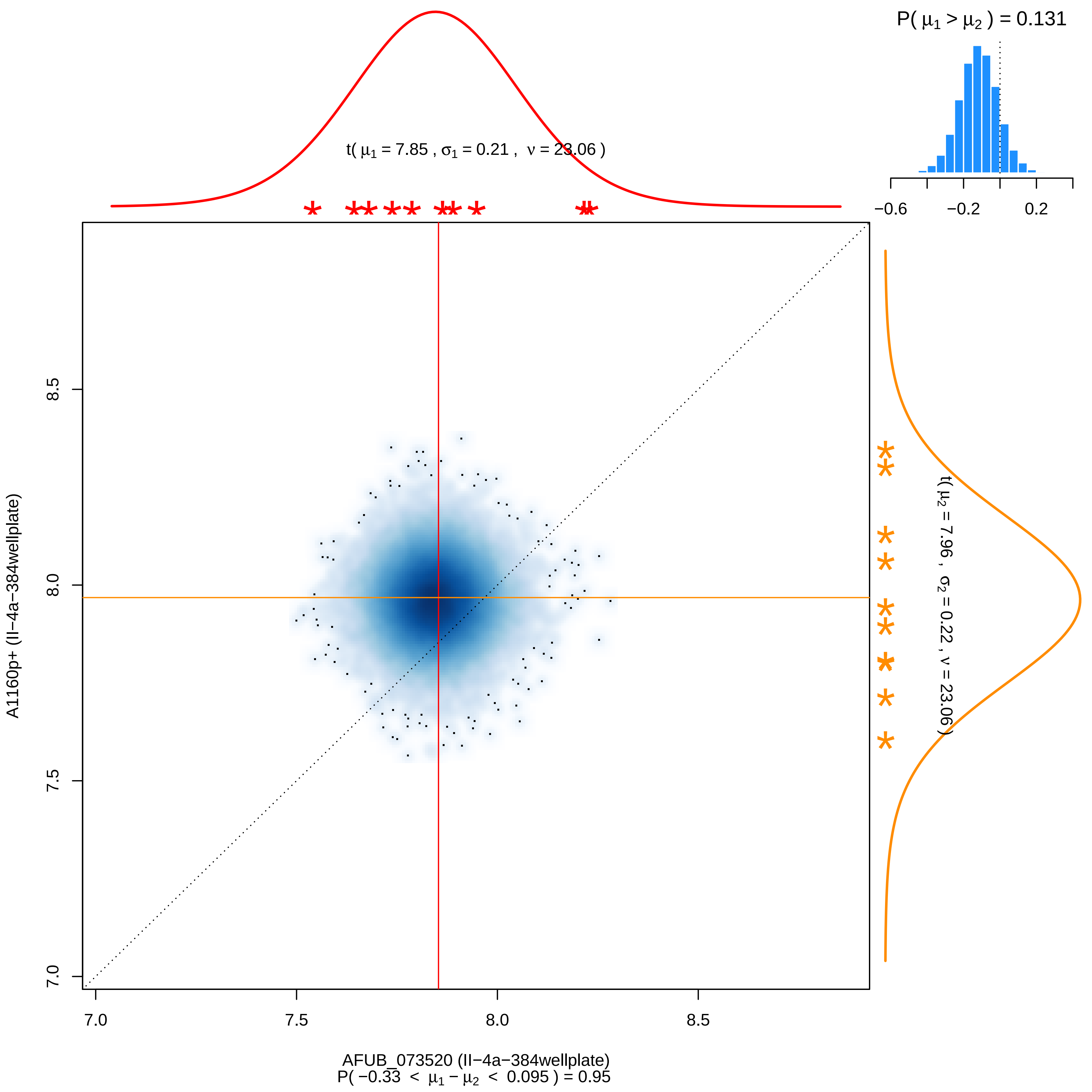

P( μ1 > μ2 ) = 0.131
−0.6
−0.2
0.2
t( μ1 = 7.85 , σ1 = 0.21 , ν = 23.06 )
t( μ2 = 7.96 , σ2 = 0.22 , ν = 23.06 )
7.0
7.5
8.0
8.5
A1160p+ (II−4a−384wellplate)
AFUB_073520 (II−4a−384wellplate)
P( −0.33 < μ1 − μ2 < 0.095 ) = 0.95

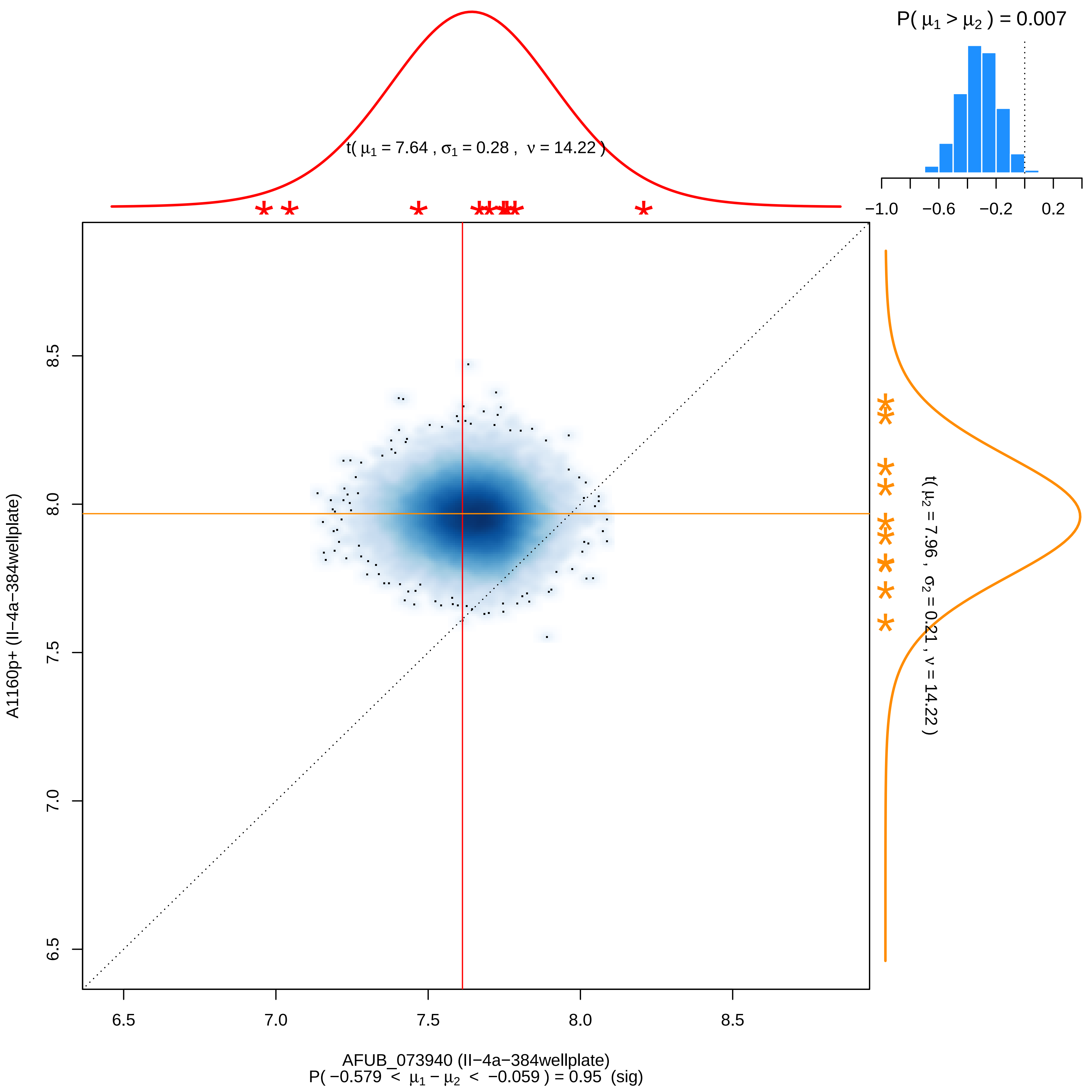

P( μ1 > μ2 ) = 0.007
−1.0
−0.6
−0.2
0.2
t( μ1 = 7.64 , σ1 = 0.28 , ν = 14.22 )
t( μ2 = 7.96 , σ2 = 0.21 , ν = 14.22 )
A1160p+ (II−4a−384wellplate)
8.5
8.0
7.5
7.0
6.5
6.5
7.0
7.5
8.0
8.5
AFUB_073940 (II−4a−384wellplate)
P( −0.579 < μ1 − μ2 < −0.059 ) = 0.95 (sig)

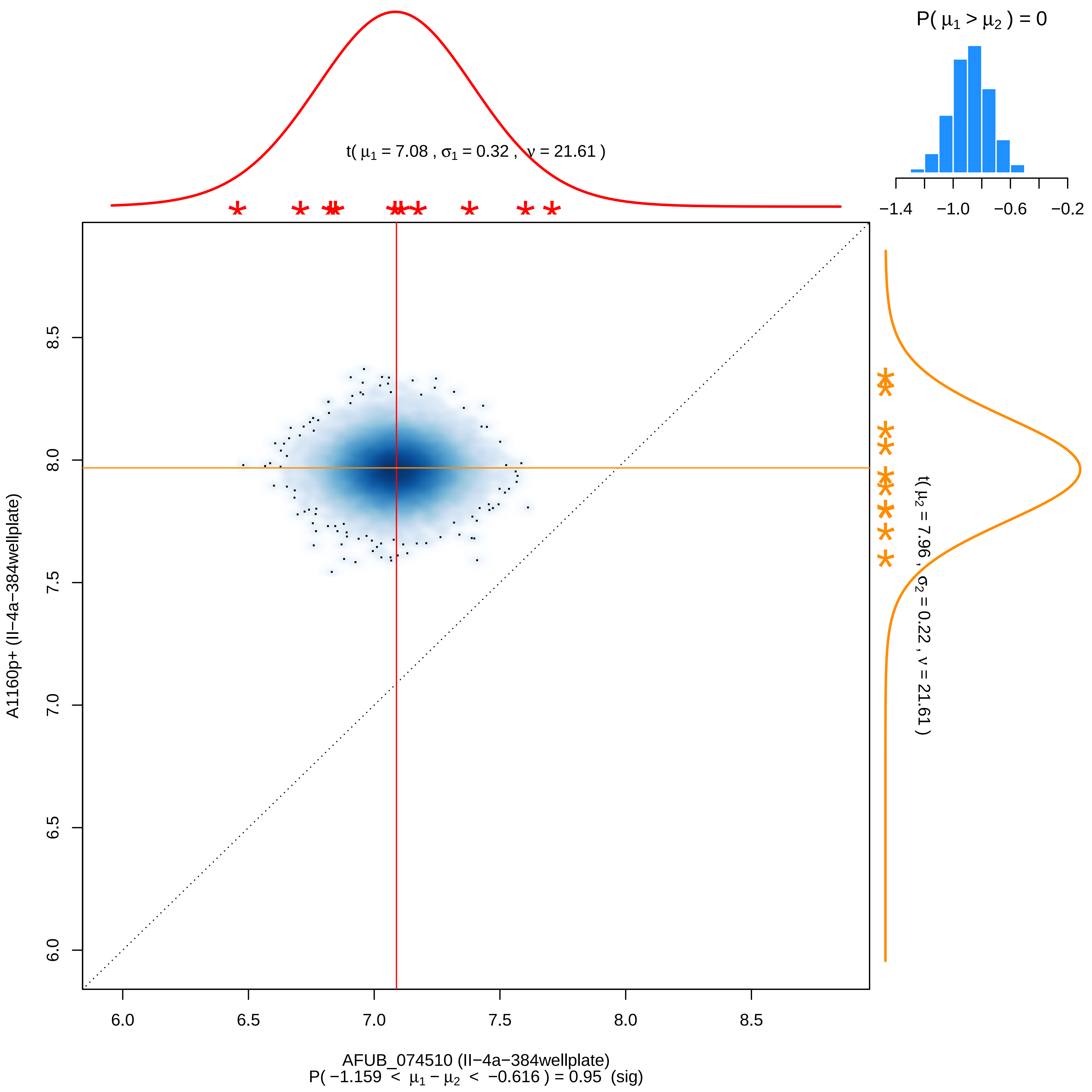

P( μ1 > μ2 ) = 0
−1.4 −1.0 −0.6 −0.2
t( μ1 = 7.08 , σ1 = 0.32 , ν = 21.61 )
t( μ2 = 7.96 , σ2 = 0.22 , ν = 21.61 )
A1160p+ (II−4a−384wellplate)
6.0 6.5 7.0 7.5 8.0 8.5
AFUB_074510 (II−4a−384wellplate)
P( −1.159 < μ1 − μ2 < −0.616 ) = 0.95 (sig)

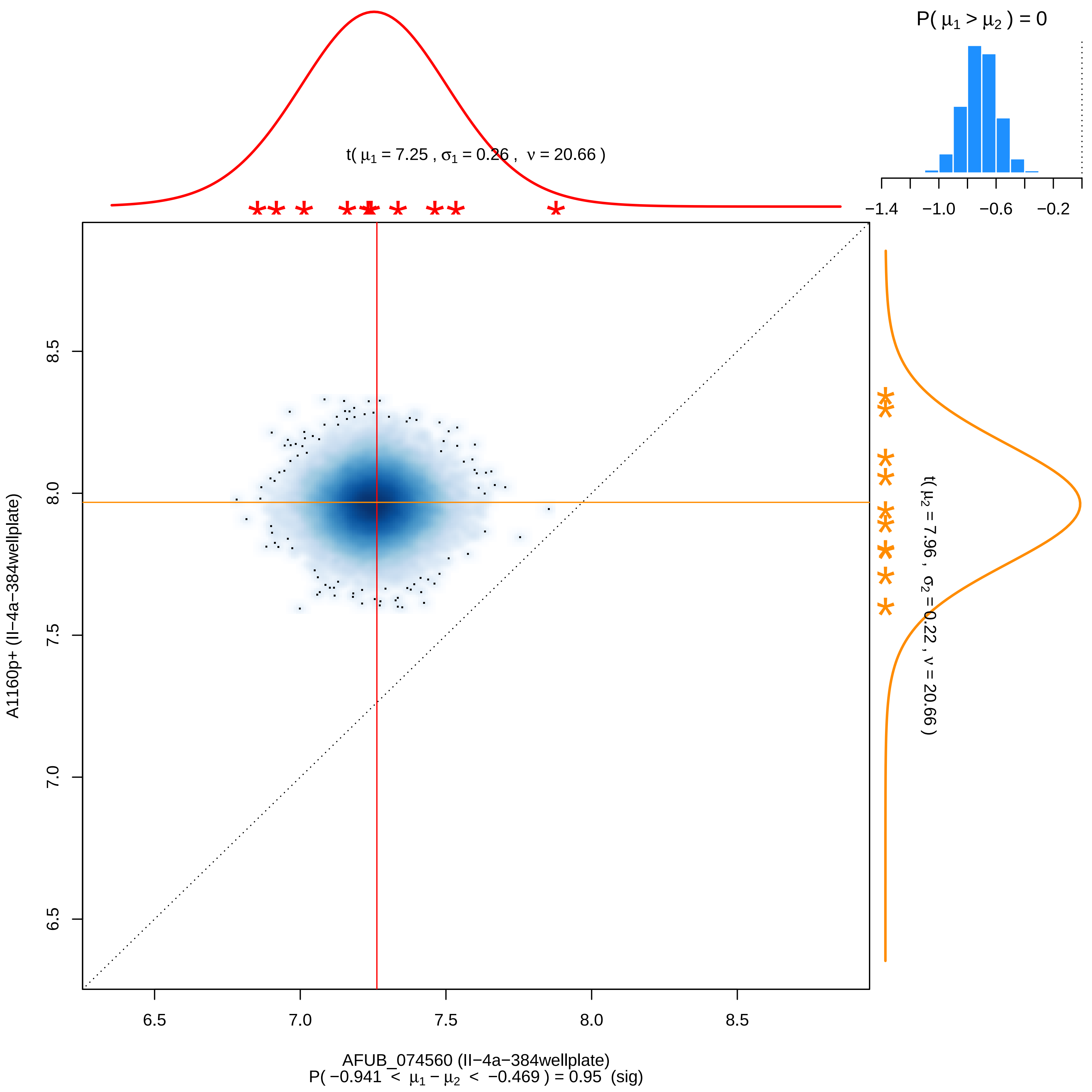

P( μ1 > μ2 ) = 0
−1.4
−1.0
−0.6
−0.2
t( μ1 = 7.25 , σ1 = 0.26 , ν = 20.66 )
t( μ2 = 7.96 , σ2 = 0.22 , ν = 20.66 )
8.5
8.0
7.5
7.0
6.5
A1160p+ (II−4a−384wellplate)
6.5
7.0
7.5
8.0
8.5
AFUB_074560 (II−4a−384wellplate)
P( −0.941 < μ1 − μ2 < −0.469 ) = 0.95 (sig)

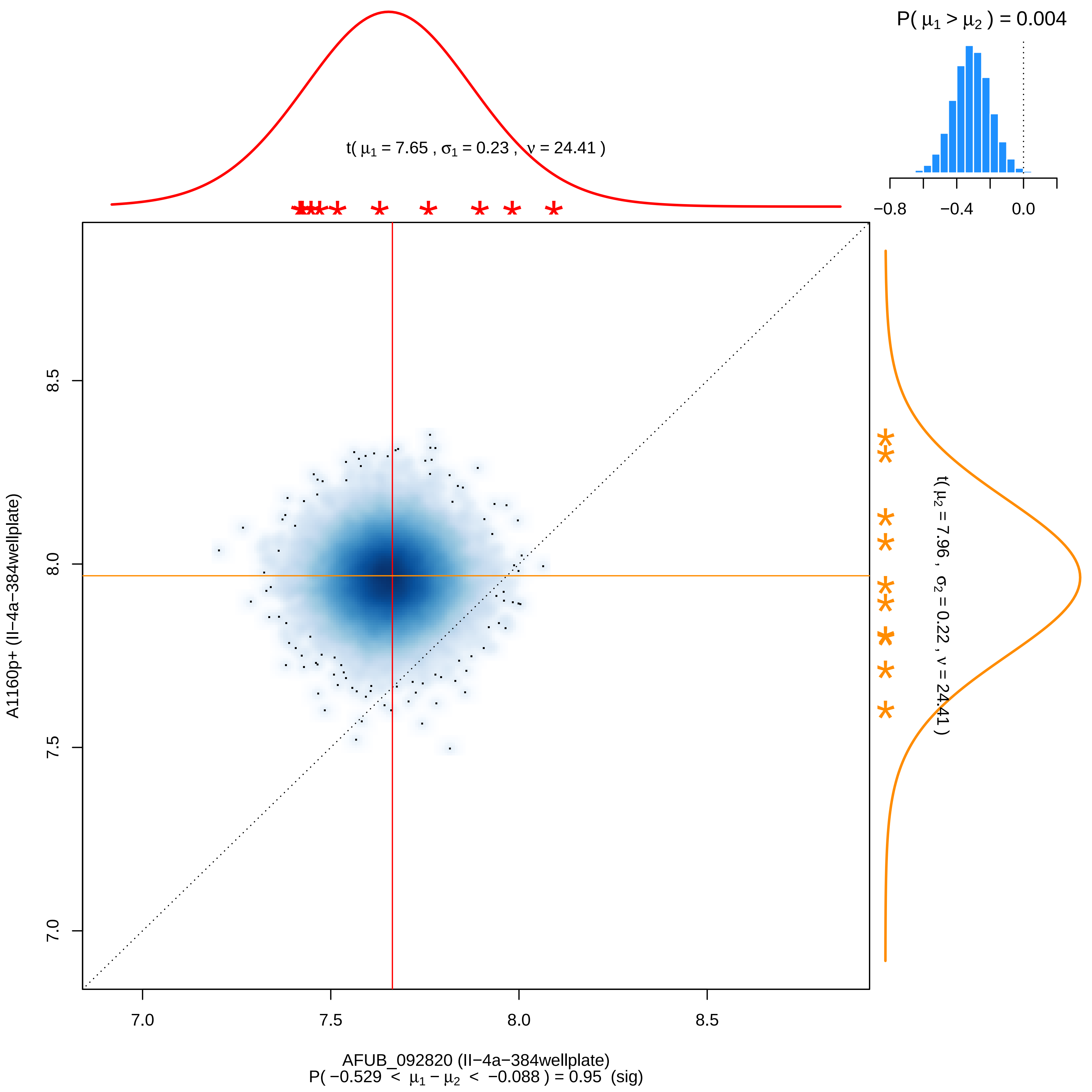

P( μ1 > μ2 ) = 0.004
−0.8
−0.4
0.0
t( μ1 = 7.65 , σ1 = 0.23 , ν = 24.41 )
t( μ2 = 7.96 , σ2 = 0.22 , ν = 24.41 )
A1160p+ (II−4a−384wellplate)
8.5
8.0
7.5
7.0
AFUB_092820 (II−4a−384wellplate)
P( −0.529 < μ1 − μ2 < −0.088 ) = 0.95 (sig)

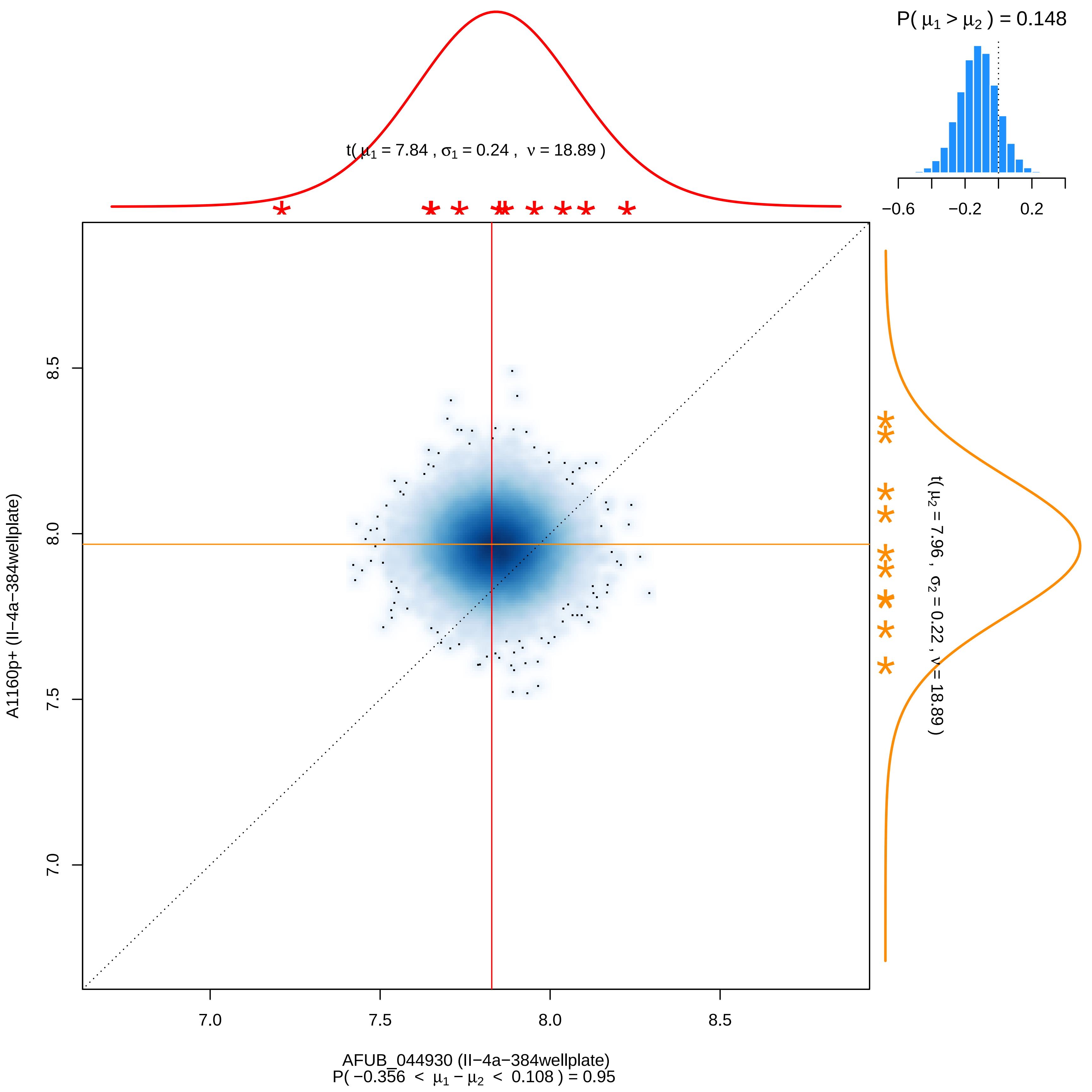

P( μ1 > μ2 ) = 0.148
−0.6
−0.2
0.2
t( μ1 = 7.84 , σ1 = 0.24 , ν = 18.89 )
t( μ2 = 7.96 , σ2 = 0.22 , ν = 18.89 )
A1160p+ (II−4a−384wellplate)
7.0
7.5
8.0
8.5
AFUB_044930 (II−4a−384wellplate)
P( −0.356 < μ1 − μ2 < 0.108 ) = 0.95

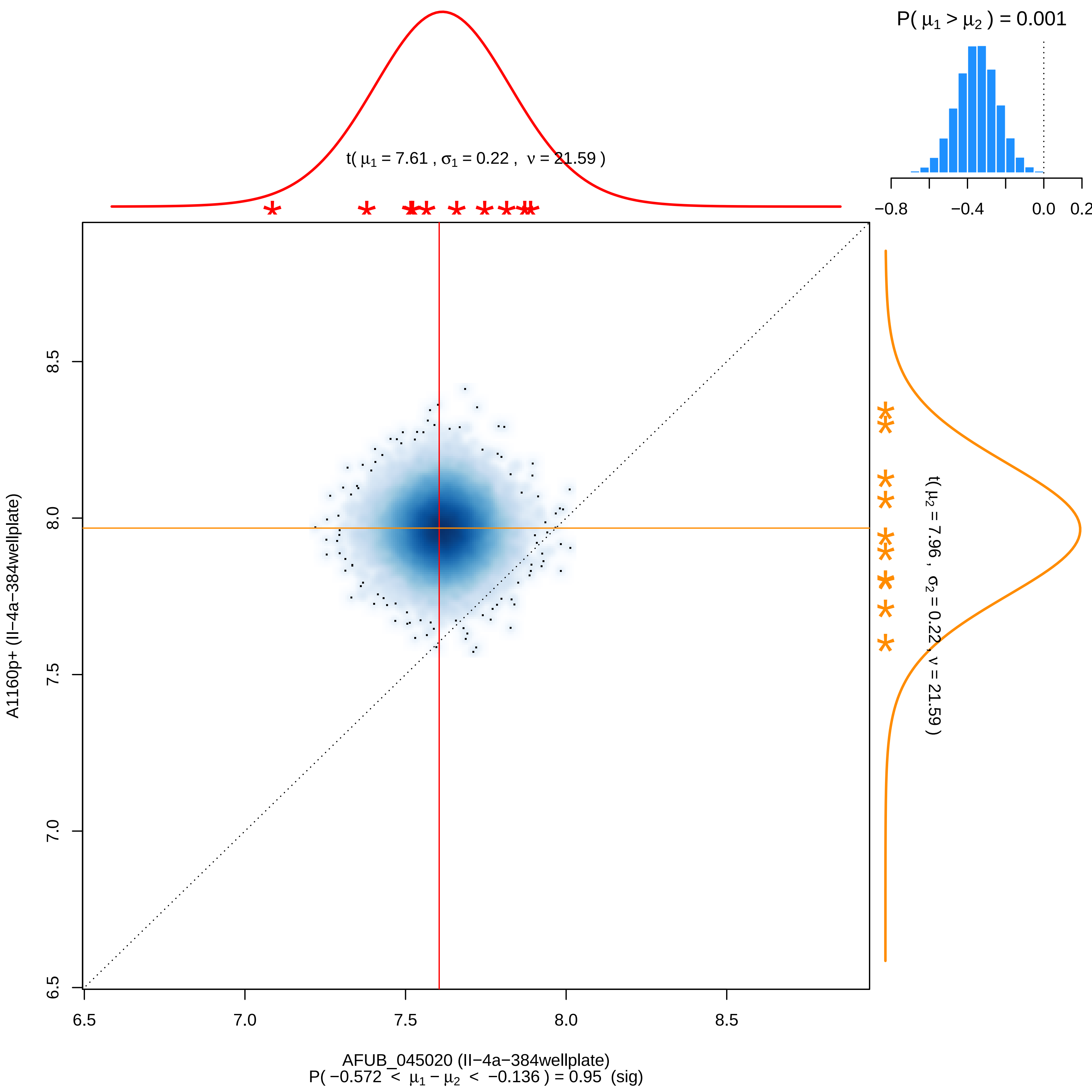

P( μ1 > μ2 ) = 0.001
−0.8
−0.4
0.0
0.2
t( μ1 = 7.61 , σ1 = 0.22 , ν = 21.59 )
t( μ2 = 7.96 , σ2 = 0.22 , ν = 21.59 )
A1160p+ (II−4a−384wellplate)
6.5
7.0
7.5
8.0
8.5
AFUB_045020 (II−4a−384wellplate)
P( −0.572 < μ1 − μ2 < −0.136 ) = 0.95 (sig)

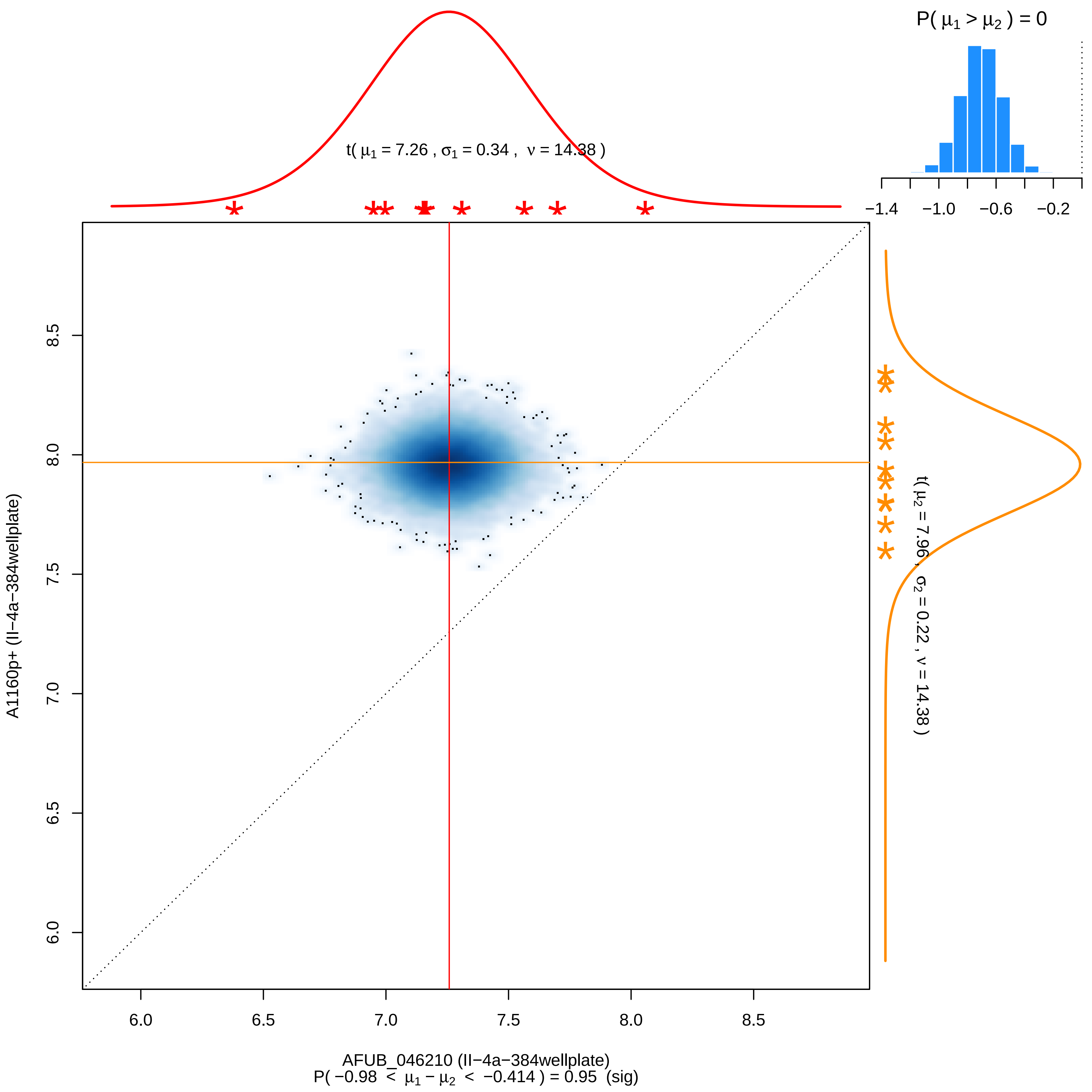

P( μ1 > μ2 ) = 0
−1.4
−1.0
−0.6
−0.2
t( μ1 = 7.26 , σ1 = 0.34 , ν = 14.38 )
t( μ2 = 7.96 , σ2 = 0.22 , ν = 14.38 )
A1160p+ (II−4a−384wellplate)
6.0
6.5
7.0
7.5
8.0
8.5
AFUB_046210 (II−4a−384wellplate)
P( −0.98 < μ1 − μ2 < −0.414 ) = 0.95 (sig)

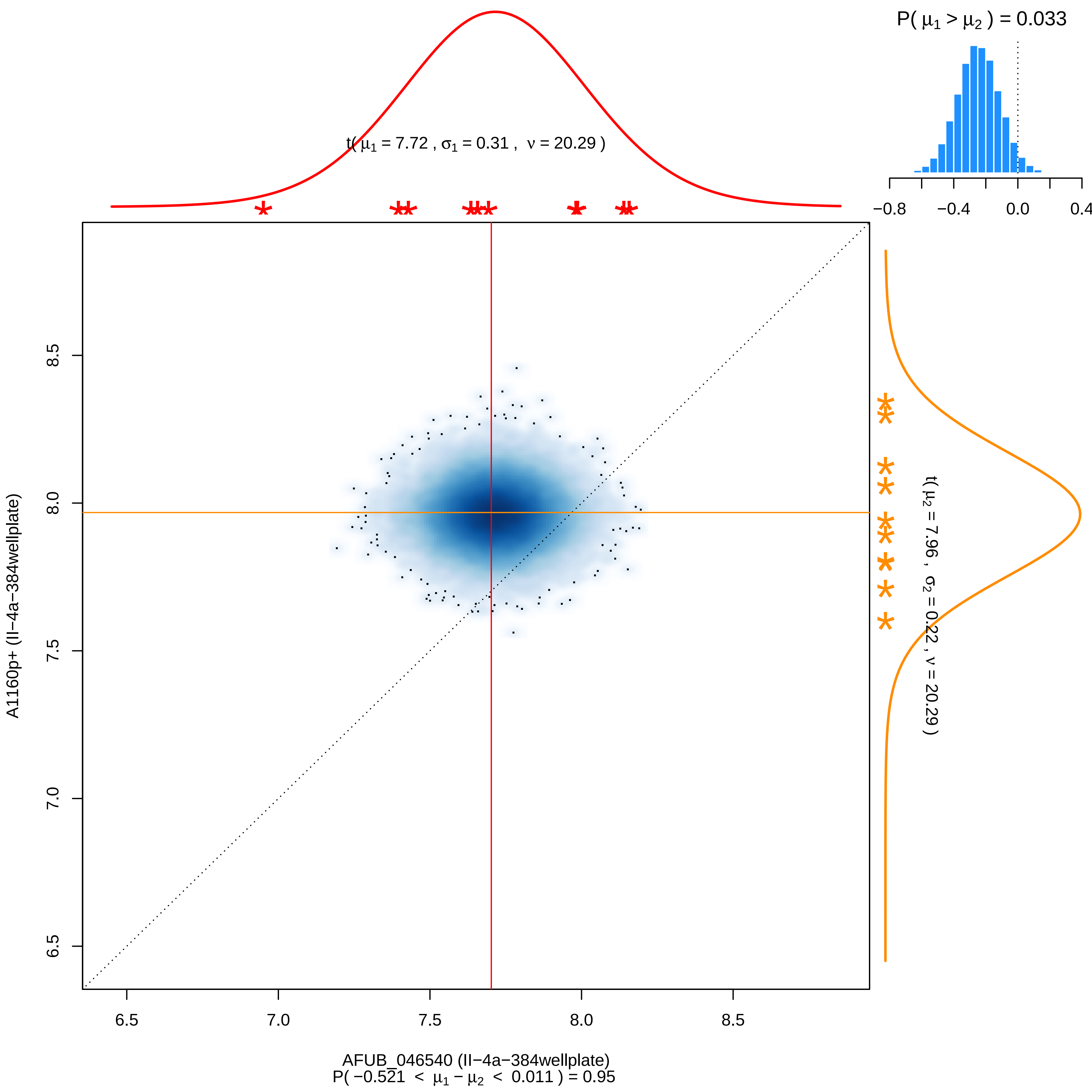

P( $\mu_1 > \mu_2$ ) = 0.033
−0.8
−0.4
0.0
0.4
t( $\mu_1$ = 7.72 , $\sigma_1$ = 0.31 , $\nu$ = 20.29 )
t( $\mu_2$ = 7.96 , $\sigma_2$ = 0.22 , $\nu$ = 20.29 )
8.5
8.0
7.5
7.0
6.5
A1160p+ (II−4a−384wellplate)
6.5
7.0
7.5
8.0
8.5
AFUB_046540 (II−4a−384wellplate)
P( −0.521 < $\mu_1 - \mu_2$ < 0.011 ) = 0.95

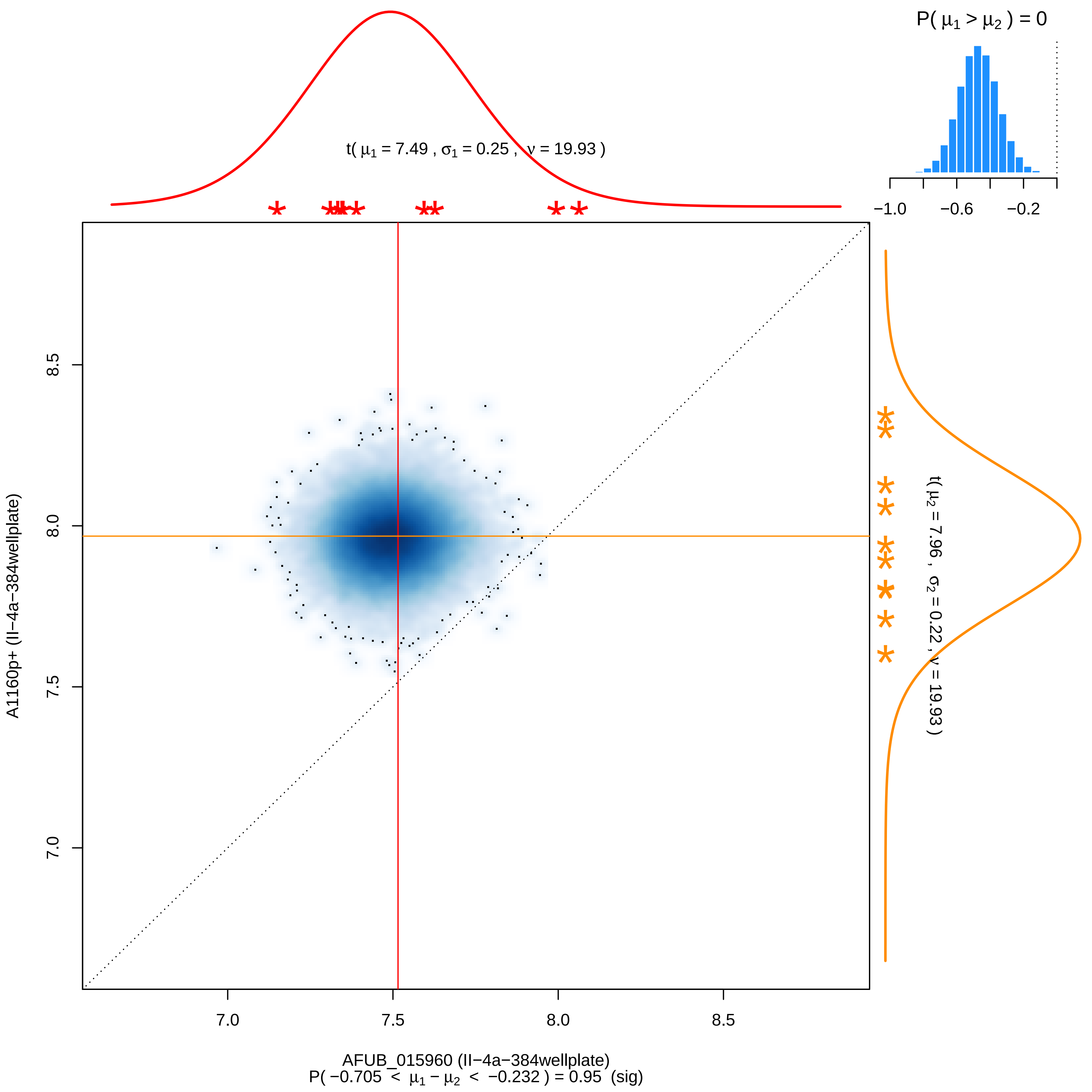

P( μ1 > μ2 ) = 0
−1.0
−0.6
−0.2
t( μ1 = 7.49 , σ1 = 0.25 , ν = 19.93 )
t( μ2 = 7.96 , σ2 = 0.22 , ν = 19.93 )
8.5
8.0
7.5
7.0
A1160p+ (II−4a−384wellplate)
7.0
7.5
8.0
8.5
AFUB_015960 (II−4a−384wellplate)
P( −0.705 < μ1 − μ2 < −0.232 ) = 0.95 (sig)

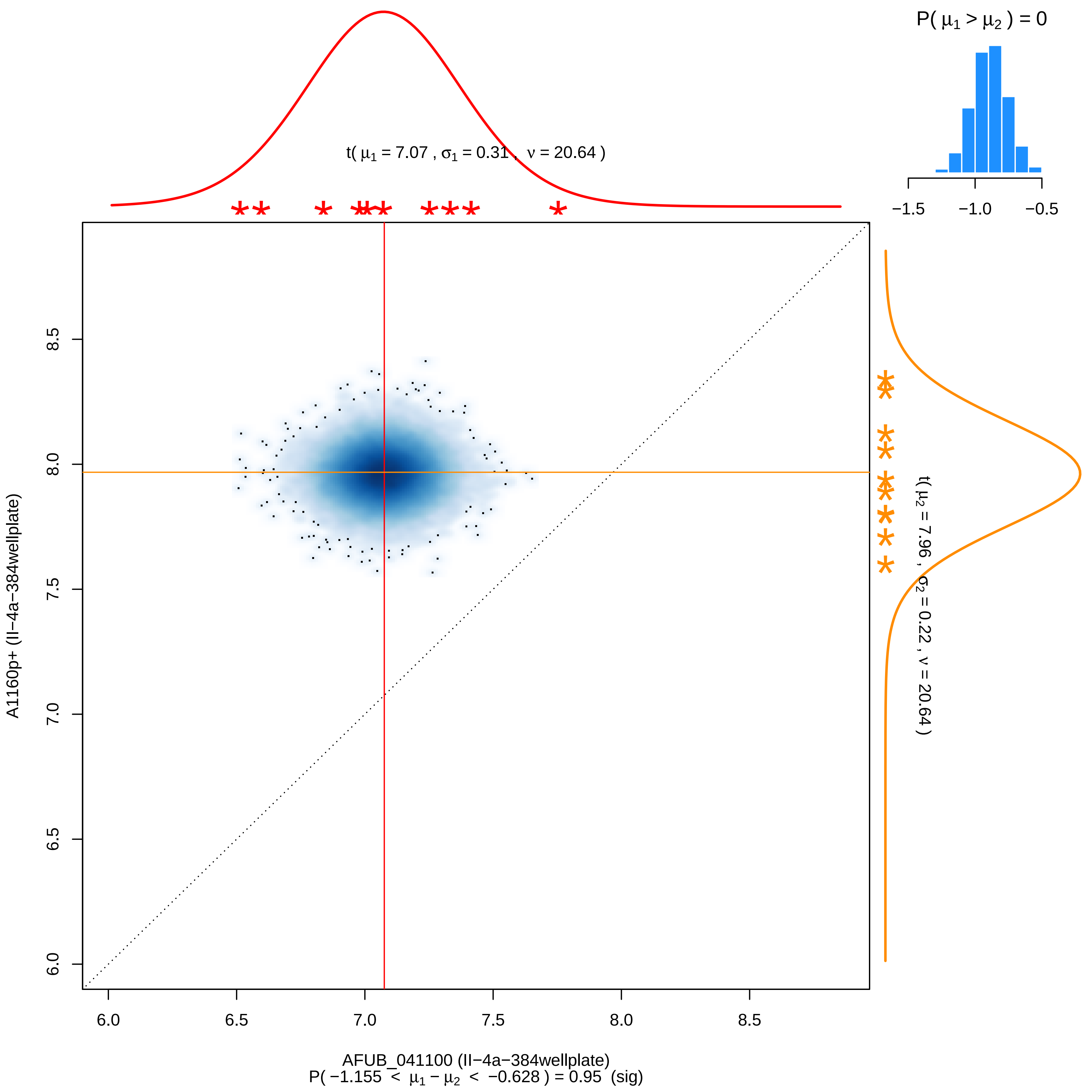

P( μ1 > μ2 ) = 0
−1.5
−1.0
−0.5
t( μ1 = 7.07 , σ1 = 0.31 , ν = 20.64 )
t( μ2 = 7.96 , σ2 = 0.22 , ν = 20.64 )
A1160p+ (II−4a−384wellplate)
6.0
6.5
7.0
7.5
8.0
8.5
AFUB_041100 (II−4a−384wellplate)
P( −1.155 < μ1 − μ2 < −0.628 ) = 0.95 (sig)

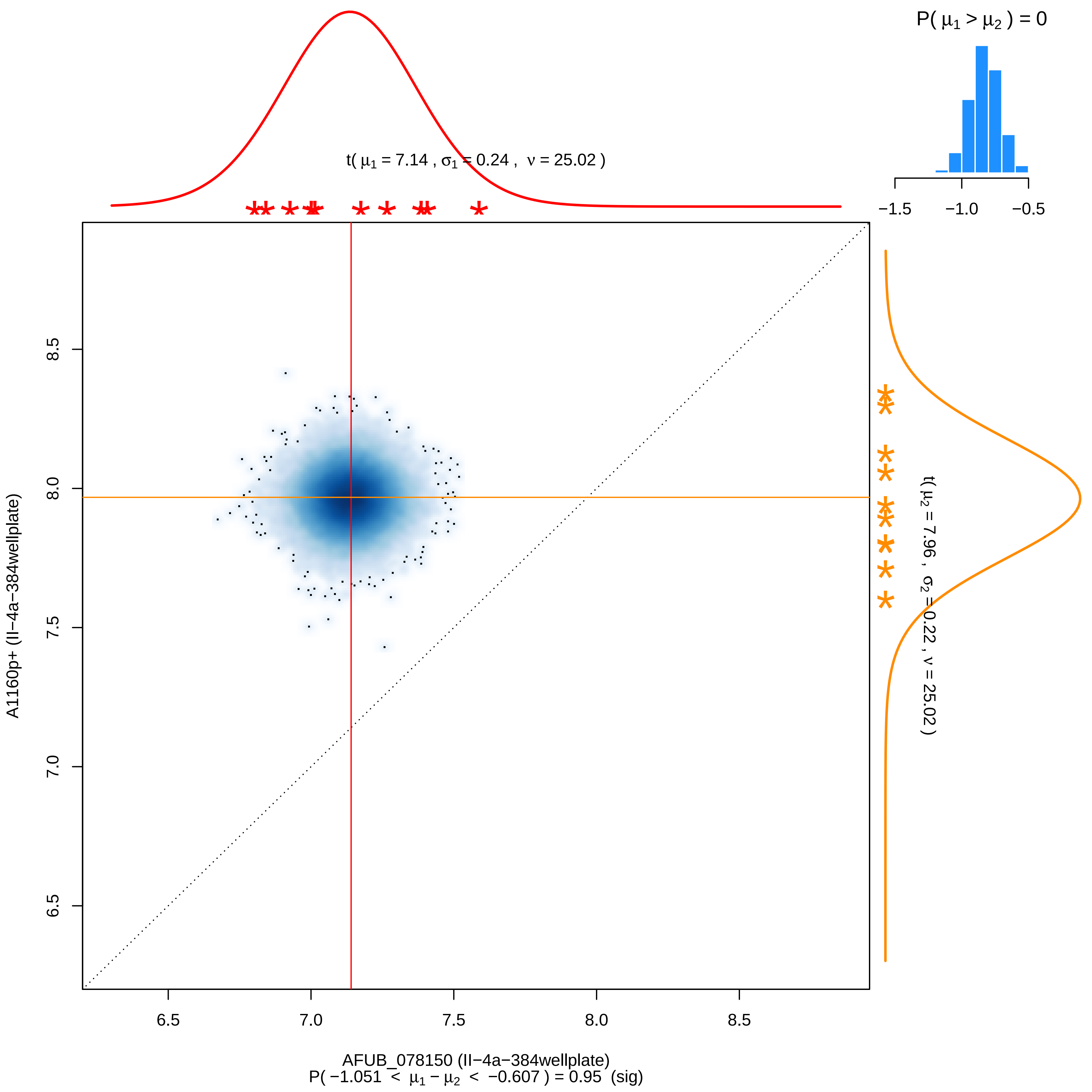

P( μ1 > μ2 ) = 0
−1.5
−1.0
−0.5
t( μ1 = 7.14 , σ1 = 0.24 , ν = 25.02 )
t( μ2 = 7.96 , σ2 = 0.22 , ν = 25.02 )
A1160p+ (II−4a−384wellplate)
8.5
8.0
7.5
7.0
6.5
6.5
7.0
7.5
8.0
8.5
AFUB_078150 (II−4a−384wellplate)
P( −1.051 < μ1 − μ2 < −0.607 ) = 0.95 (sig)

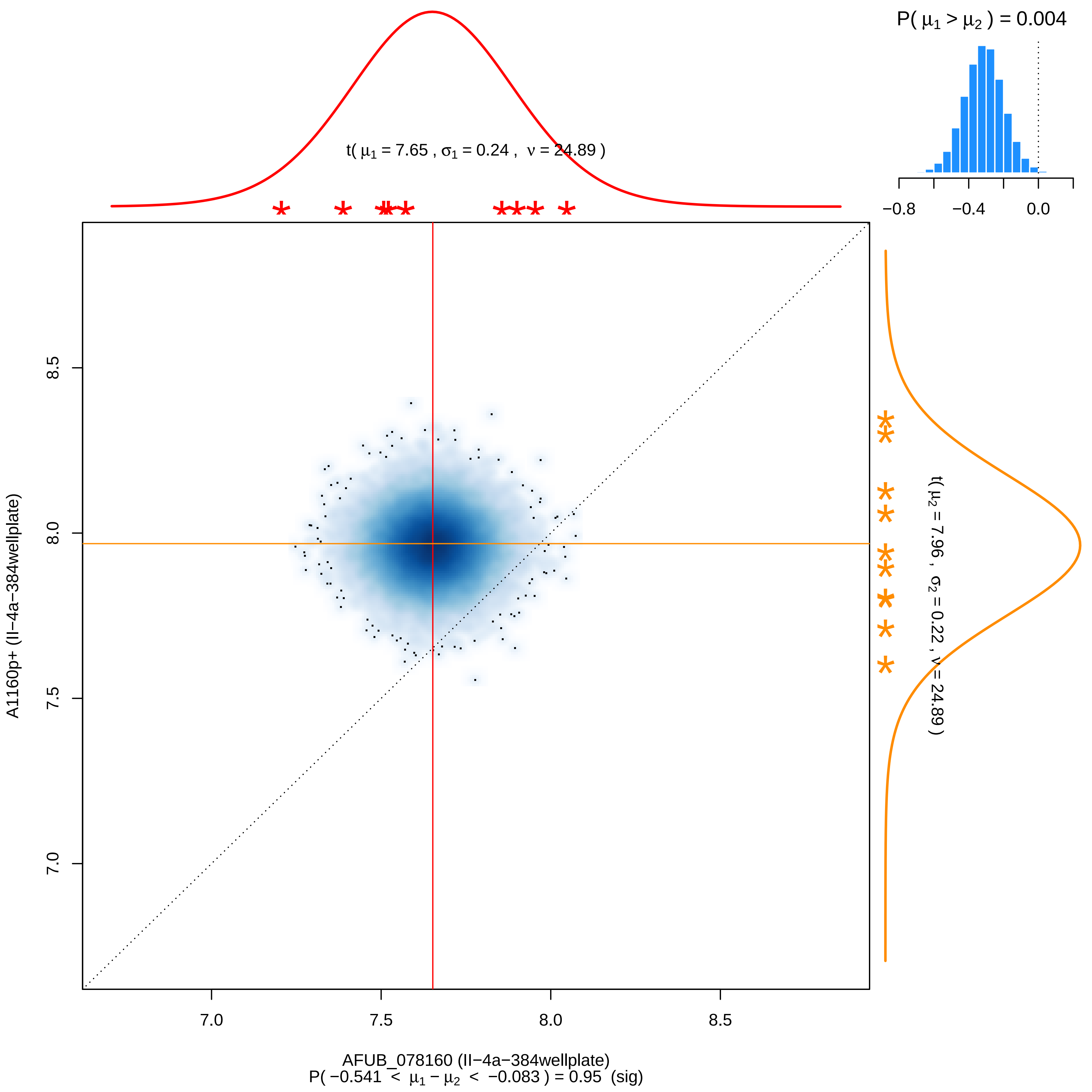

P( μ1 > μ2 ) = 0.004
−0.8
−0.4
0.0
t( μ1 = 7.65 , σ1 = 0.24 , ν = 24.89 )
t( μ2 = 7.96 , σ2 = 0.22 , ν = 24.89 )
8.5
8.0
7.5
7.0
A1160p+ (II−4a−384wellplate)
7.0
7.5
8.0
8.5
AFUB_078160 (II−4a−384wellplate)
P( −0.541 < μ1 − μ2 < −0.083 ) = 0.95 (sig)

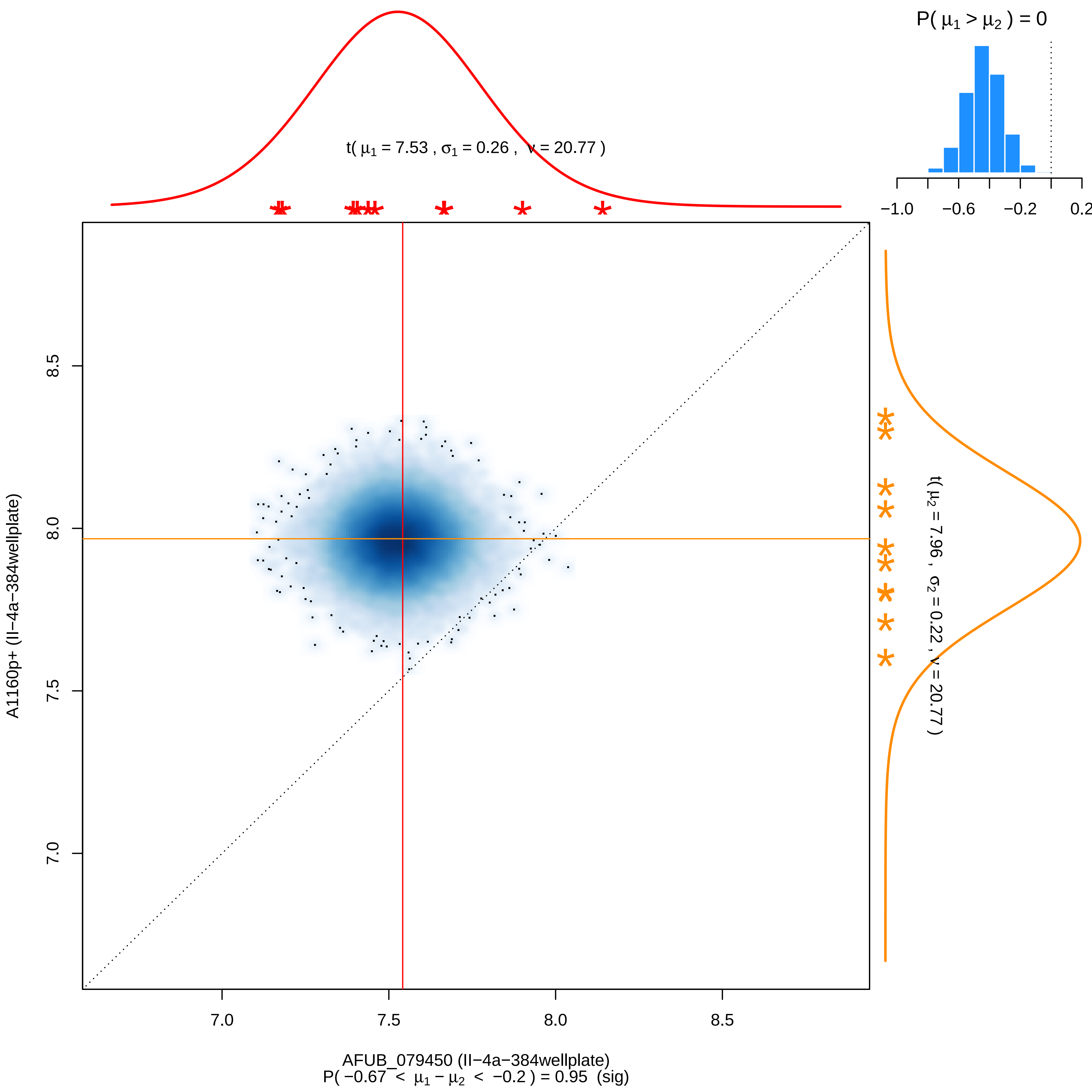

P( μ1 > μ2 ) = 0
−1.0
−0.6
−0.2
0.2
t( μ1 = 7.53 , σ1 = 0.26 , ν = 20.77 )
t( μ2 = 7.96 , σ2 = 0.22 , ν = 20.77 )
A1160p+ (II−4a−384wellplate)
7.0
7.5
8.0
8.5
AFUB_079450 (II−4a−384wellplate)
P( −0.67 < μ1 − μ2 < −0.2 ) = 0.95 (sig)

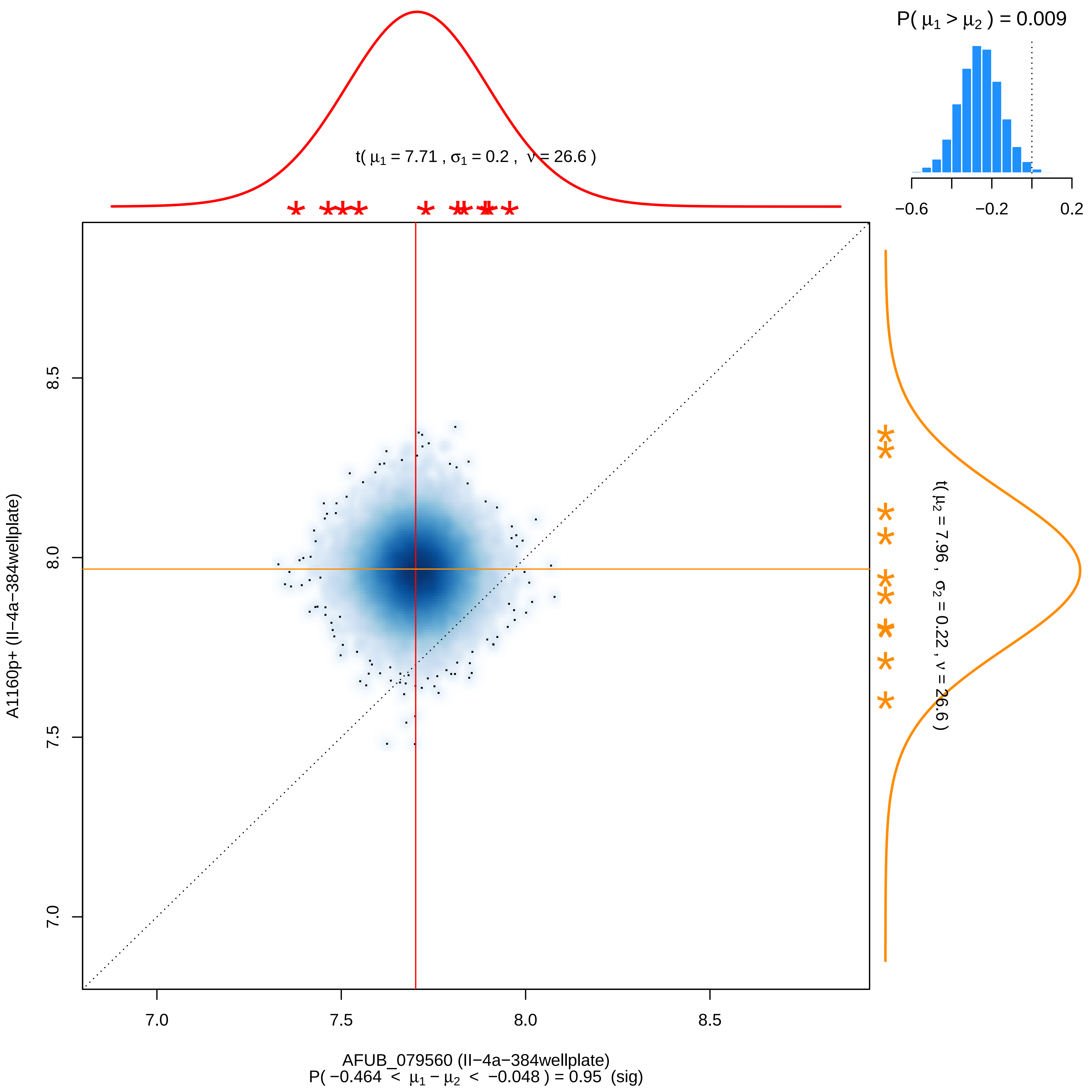

P( μ1 > μ2 ) = 0.009
−0.6
−0.2
0.2
t( μ1 = 7.71 , σ1 = 0.2 , ν = 26.6 )
t( μ2 = 7.96 , σ2 = 0.22 , ν = 26.6 )
A1160p+ (II−4a−384wellplate)
8.5
8.0
7.5
7.0
7.0
7.5
8.0
8.5
AFUB_079560 (II−4a−384wellplate)
P( −0.464 < μ1 − μ2 < −0.048 ) = 0.95 (sig)

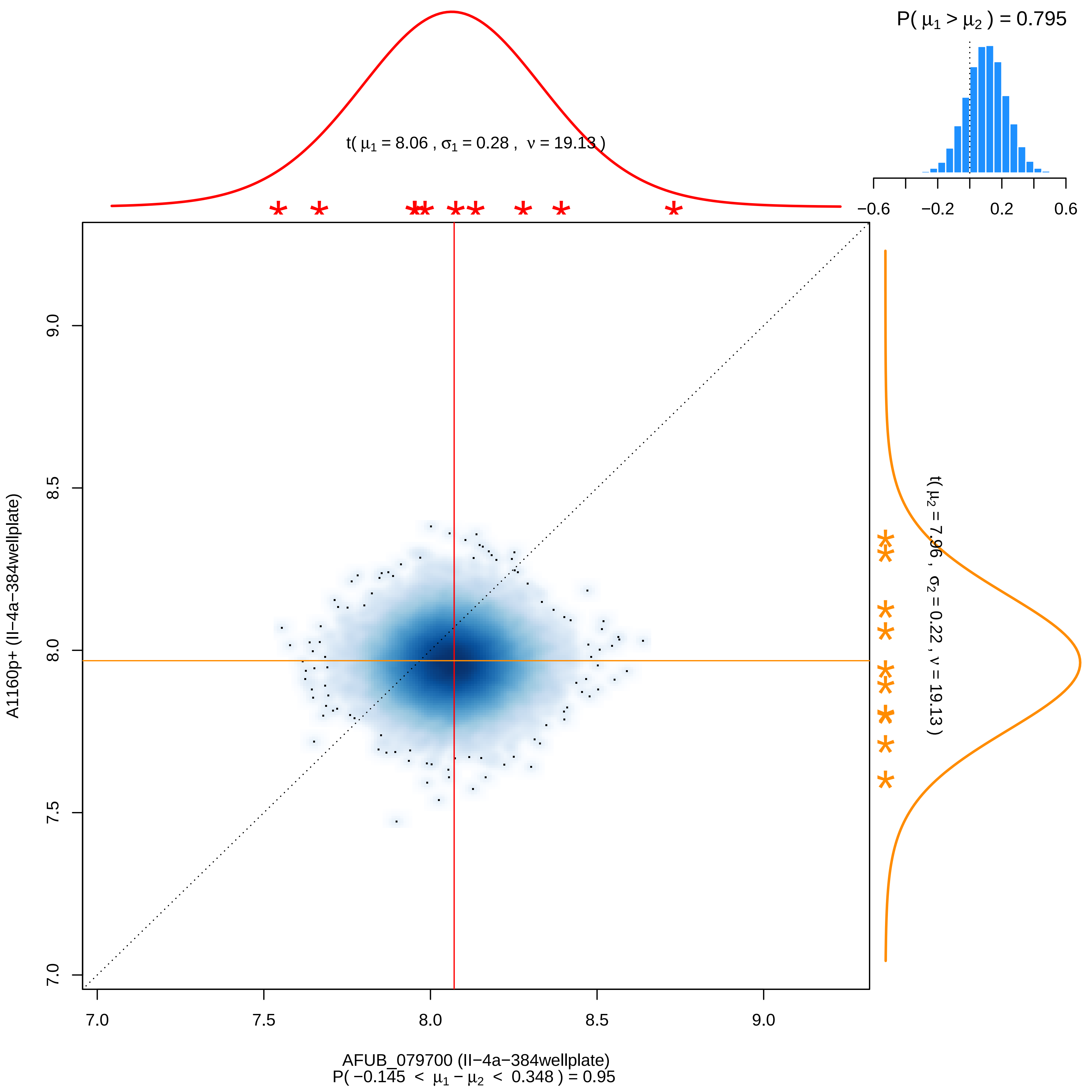

P( μ1 > μ2 ) = 0.795
t( μ1 = 8.06 , σ1 = 0.28 , ν = 19.13 )
−0.6
−0.2
0.2
0.6
t( μ2 = 7.96 , σ2 = 0.22 , ν = 19.13 )
A1160p+ (II−4a−384wellplate)
7.0
7.5
8.0
8.5
9.0
AFUB_079700 (II−4a−384wellplate)
P( −0.145 < μ1 − μ2 < 0.348 ) = 0.95

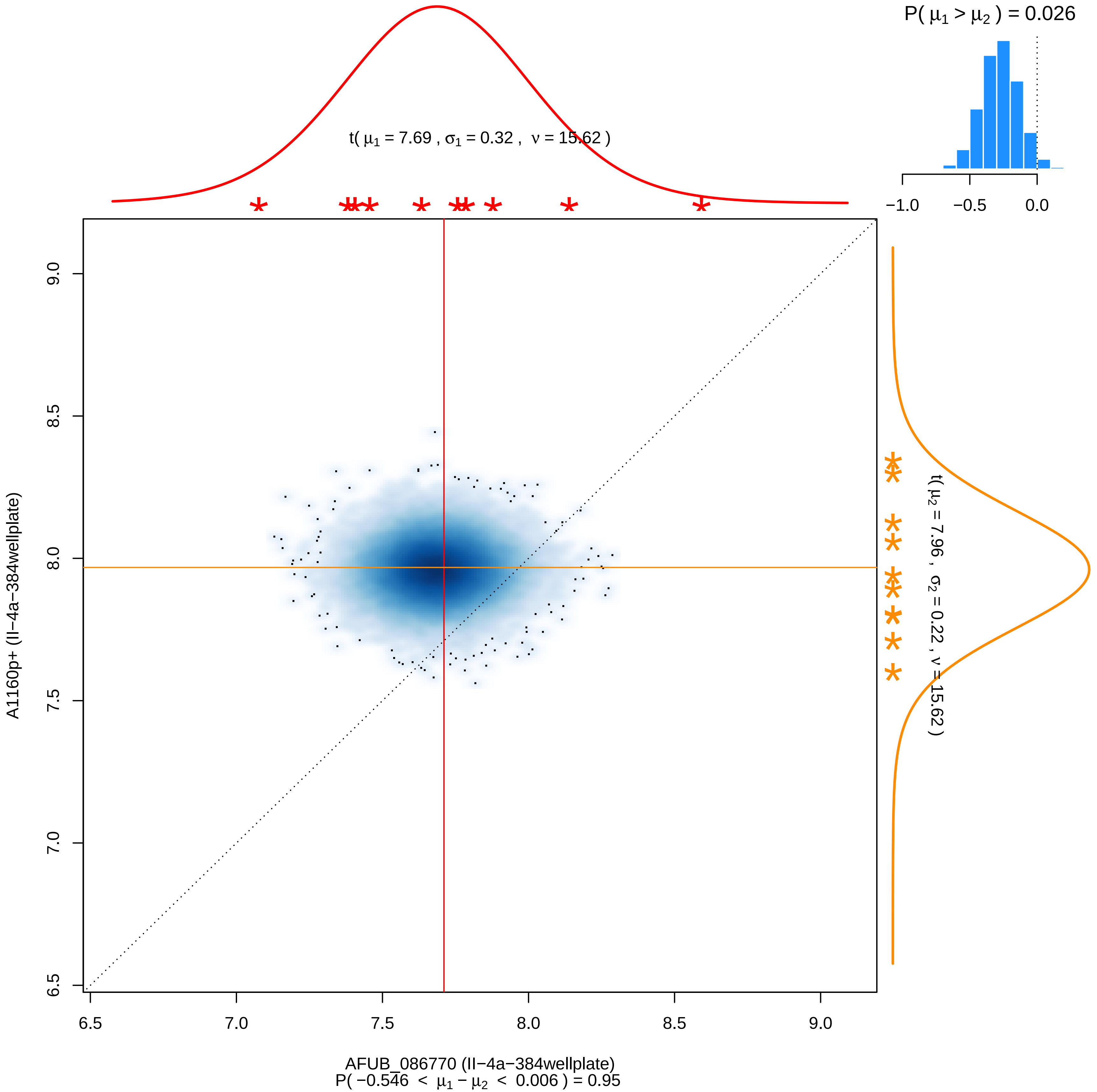

P( μ1 > μ2 ) = 0.026
−1.0
−0.5
0.0
t( μ1 = 7.69 , σ1 = 0.32 , ν = 15.62 )
t( μ2 = 7.96 , σ2 = 0.22 , ν = 15.62 )
A1160p+ (II−4a−384wellplate)
9.0
8.5
8.0
7.5
7.0
6.5
6.5
7.0
7.5
8.0
8.5
9.0
AFUB_086770 (II−4a−384wellplate)
P( −0.546 < μ1 − μ2 < 0.006 ) = 0.95

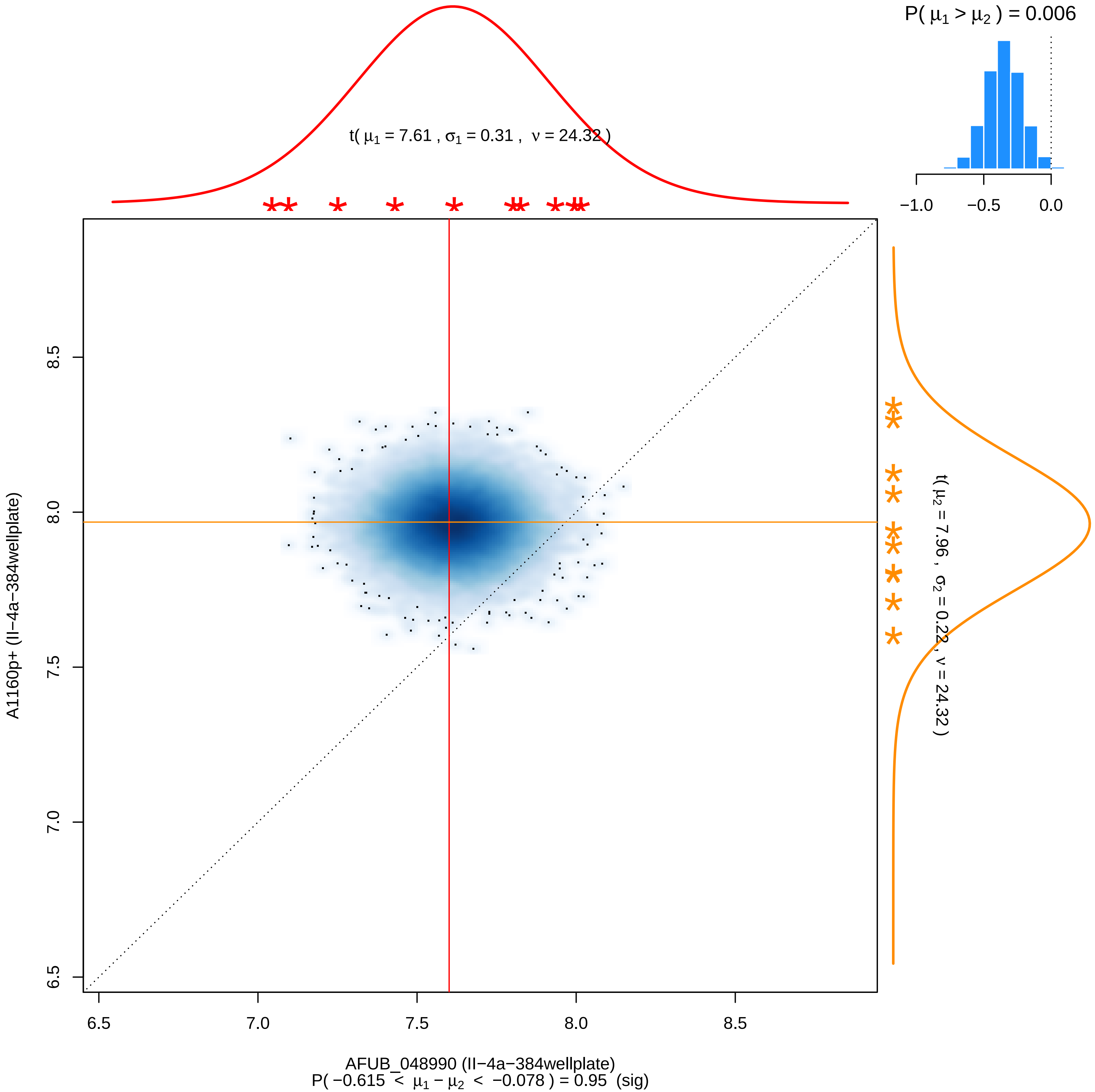

P( μ1 > μ2 ) = 0.006
−1.0
−0.5
0.0
t( μ1 = 7.61 , σ1 = 0.31 , ν = 24.32 )
8.5
8.0
7.5
7.0
6.5
A1160p+ (II−4a−384wellplate)
t( μ2 = 7.96 , σ2 = 0.22 , ν = 24.32 )
6.5
7.0
7.5
8.0
8.5
AFUB_048990 (II−4a−384wellplate)
P( −0.615 < μ1 − μ2 < −0.078 ) = 0.95 (sig)

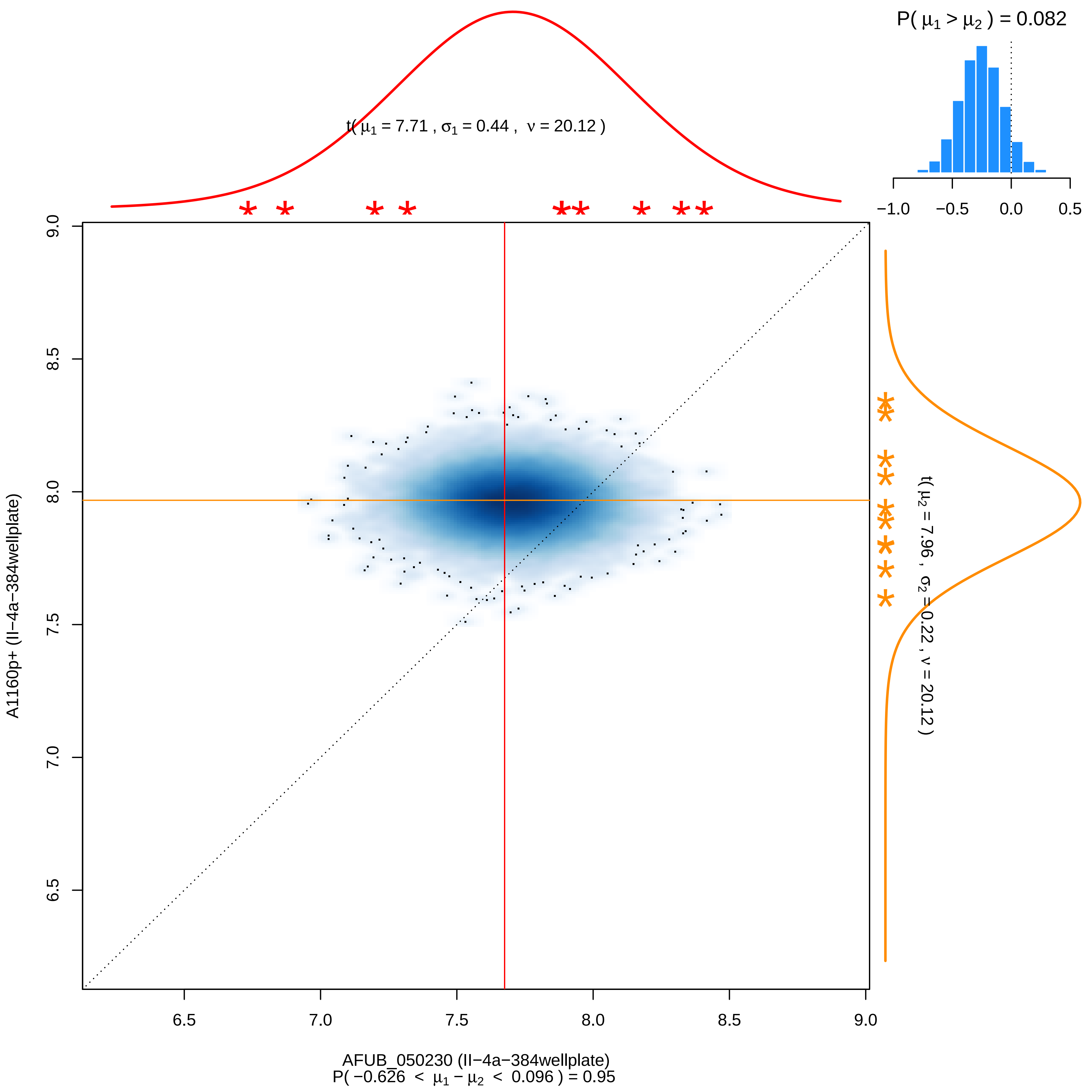
P( μ1 > μ2 ) = 0.082
−1.0 −0.5 0.0 0.5
t( μ1 = 7.71 , σ1 = 0.44 , ν = 20.12 )
t( μ2 = 7.96 , σ2 = 0.22 , ν = 20.12 )
A1160p+ (II−4a−384wellplate)
6.5 7.0 7.5 8.0 8.5 9.0
AFUB_050230 (II−4a−384wellplate)
P( −0.626 < μ1 − μ2 < 0.096 ) = 0.95

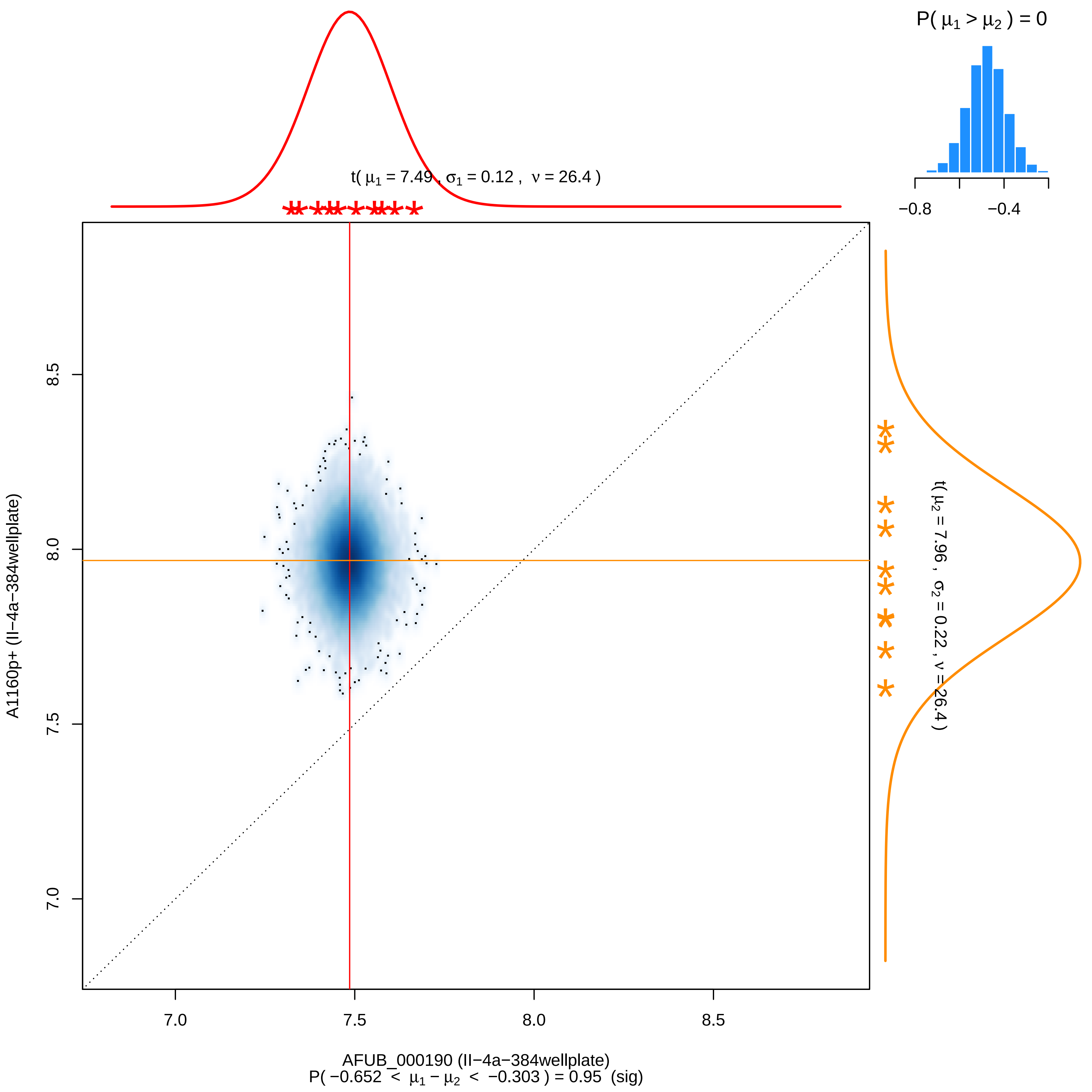

P( μ1 > μ2 ) = 0
−0.8
−0.4
t( μ1 = 7.49 , σ1 = 0.12 , ν = 26.4 )
t( μ2 = 7.96 , σ2 = 0.22 , ν = 26.4 )
8.5
8.0
7.5
7.0
A1160p+ (II−4a−384wellplate)
7.0
7.5
8.0
8.5
AFUB_000190 (II−4a−384wellplate)
P( −0.652 < μ1 − μ2 < −0.303 ) = 0.95 (sig)

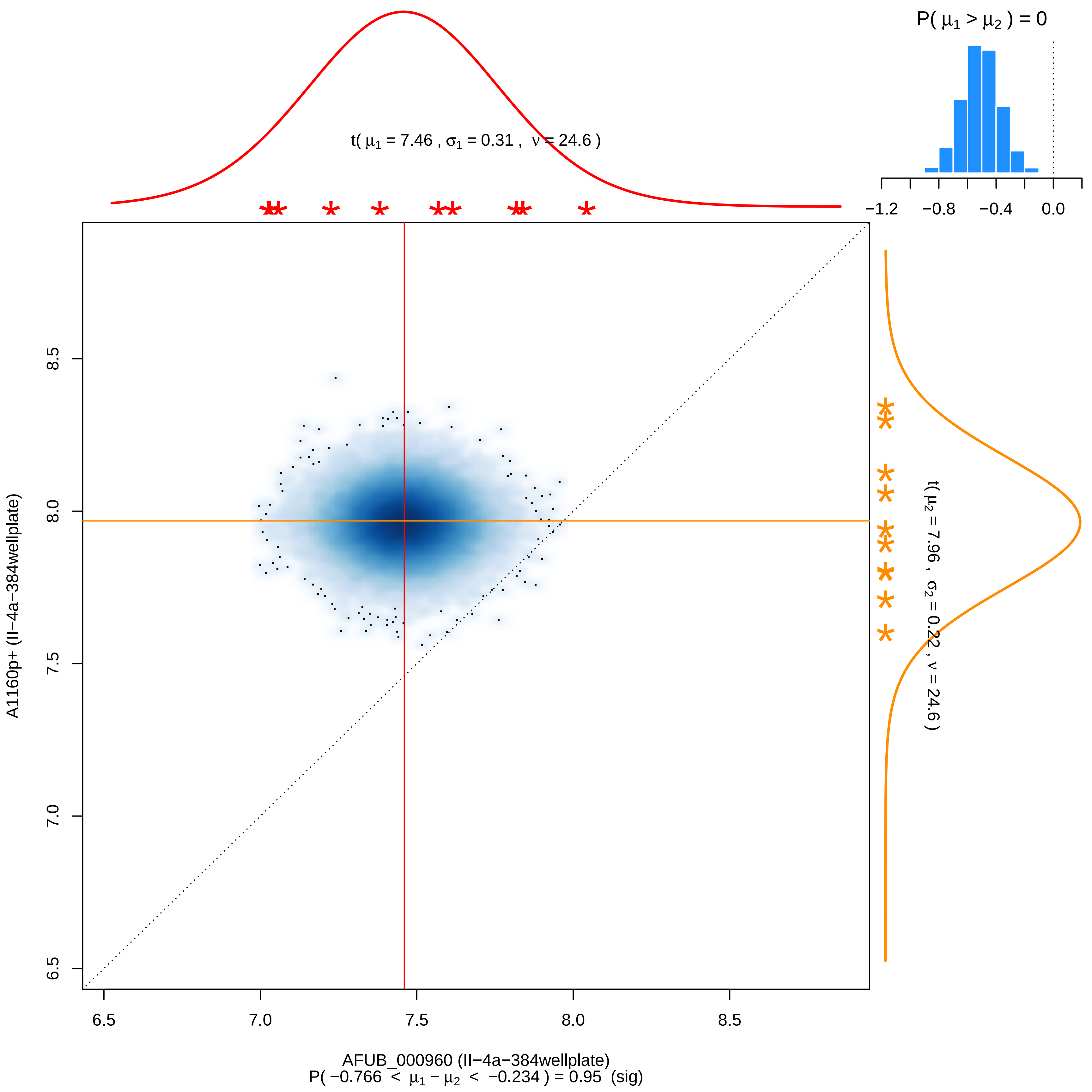

P( μ1 > μ2 ) = 0
−1.2
−0.8
−0.4
0.0
t( μ1 = 7.46 , σ1 = 0.31 , ν = 24.6 )
t( μ2 = 7.96 , σ2 = 0.22 , ν = 24.6 )
A1160p+ (II−4a−384wellplate)
8.5
8.0
7.5
7.0
6.5
6.5
7.0
7.5
8.0
8.5
AFUB_000960 (II−4a−384wellplate)
P( −0.766 < μ1 − μ2 < −0.234 ) = 0.95 (sig)

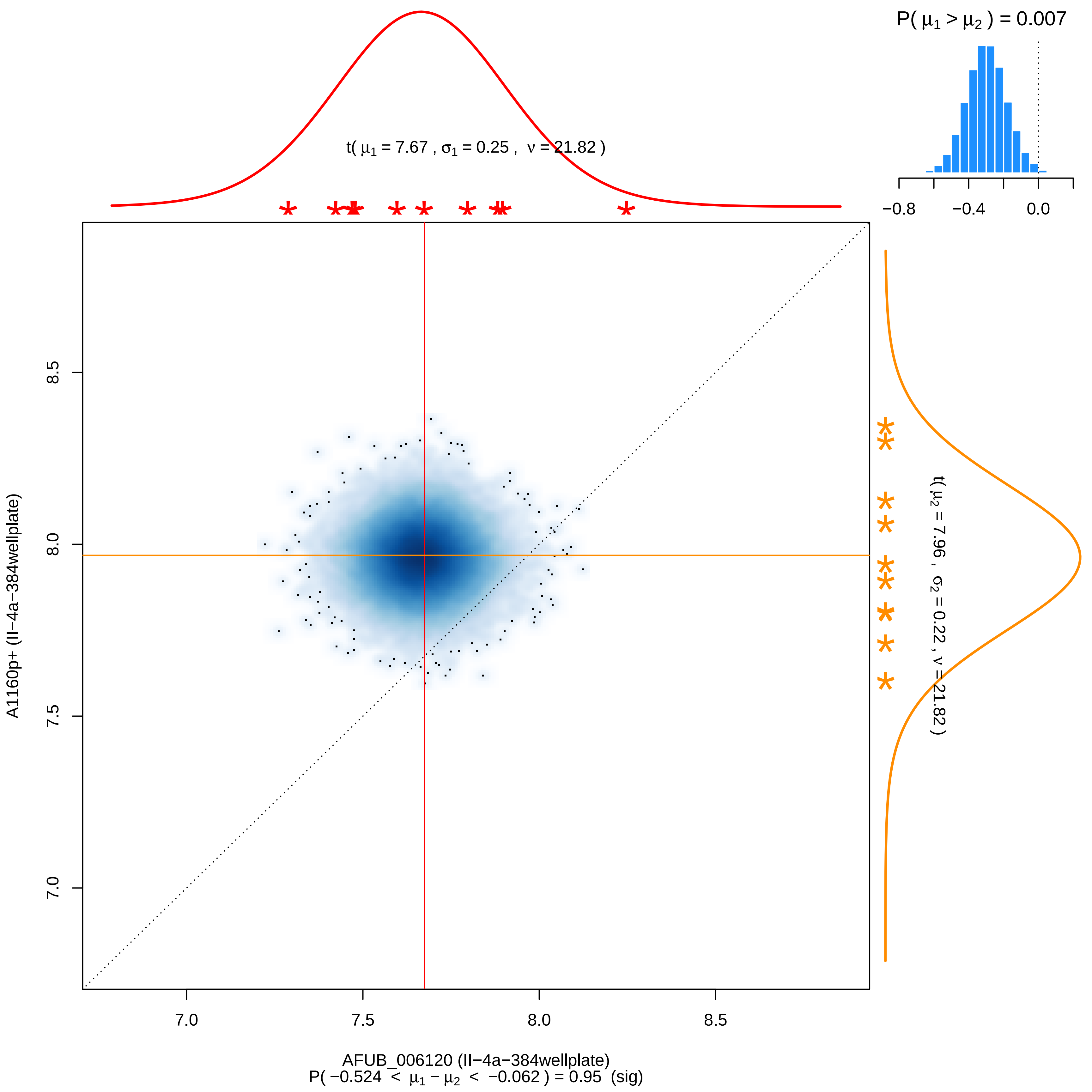

P( μ1 > μ2 ) = 0.007
−0.8
−0.4
0.0
t( μ1 = 7.67 , σ1 = 0.25 , ν = 21.82 )
t( μ2 = 7.96 , σ2 = 0.22 , ν = 21.82 )
8.5
8.0
7.5
7.0
A1160p+ (II−4a−384wellplate)
7.0
7.5
8.0
8.5
AFUB_006120 (II−4a−384wellplate)
P( −0.524 < μ1 − μ2 < −0.062 ) = 0.95 (sig)

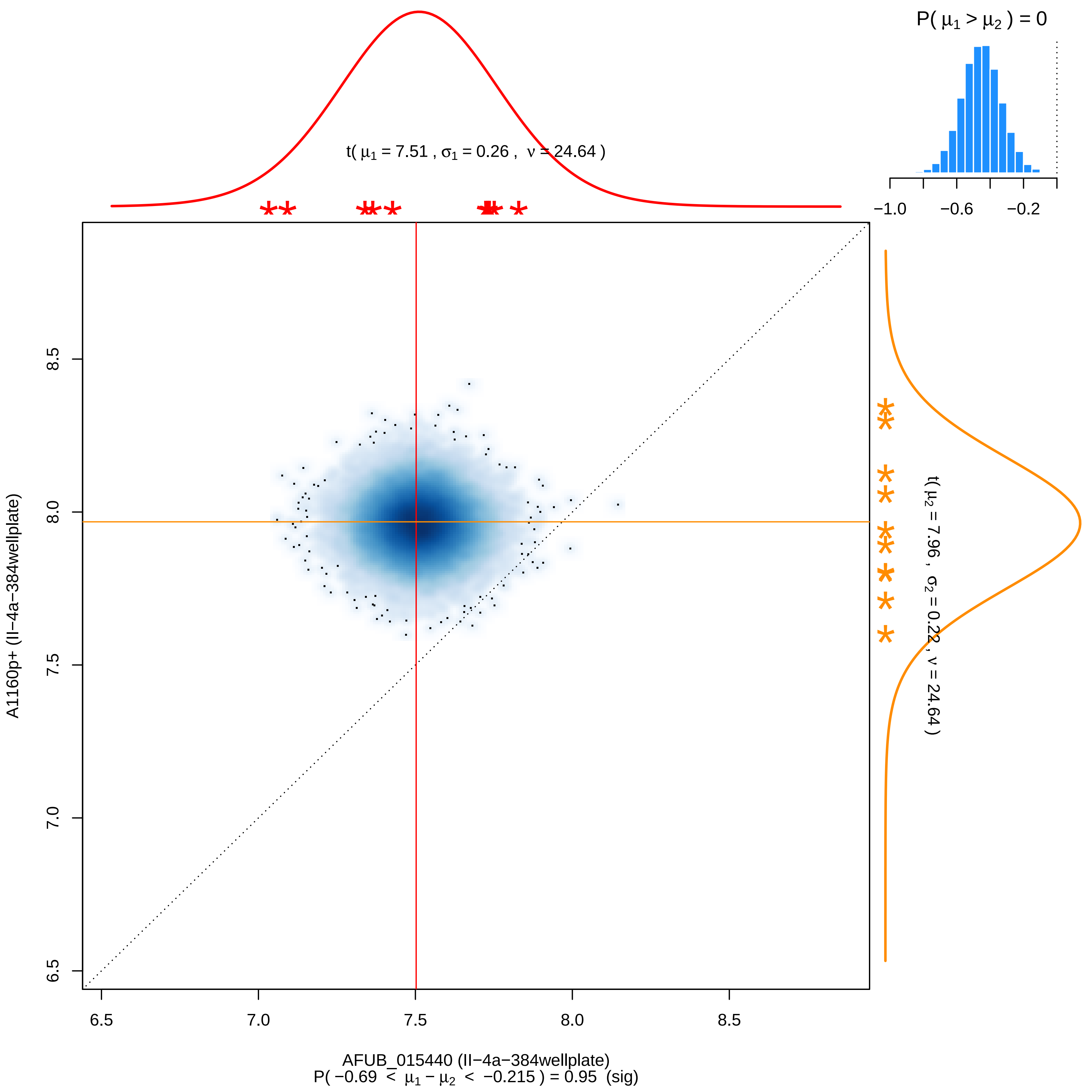

P( μ1 > μ2 ) = 0
−1.0
−0.6
−0.2
t( μ1 = 7.51 , σ1 = 0.26 , ν = 24.64 )
t( μ2 = 7.96 , σ2 = 0.22 , ν = 24.64 )
8.5
8.0
7.5
7.0
6.5
A1160p+ (II−4a−384wellplate)
6.5
7.0
7.5
8.0
8.5
AFUB_015440 (II−4a−384wellplate)
P( −0.69 < μ1 − μ2 < −0.215 ) = 0.95 (sig)

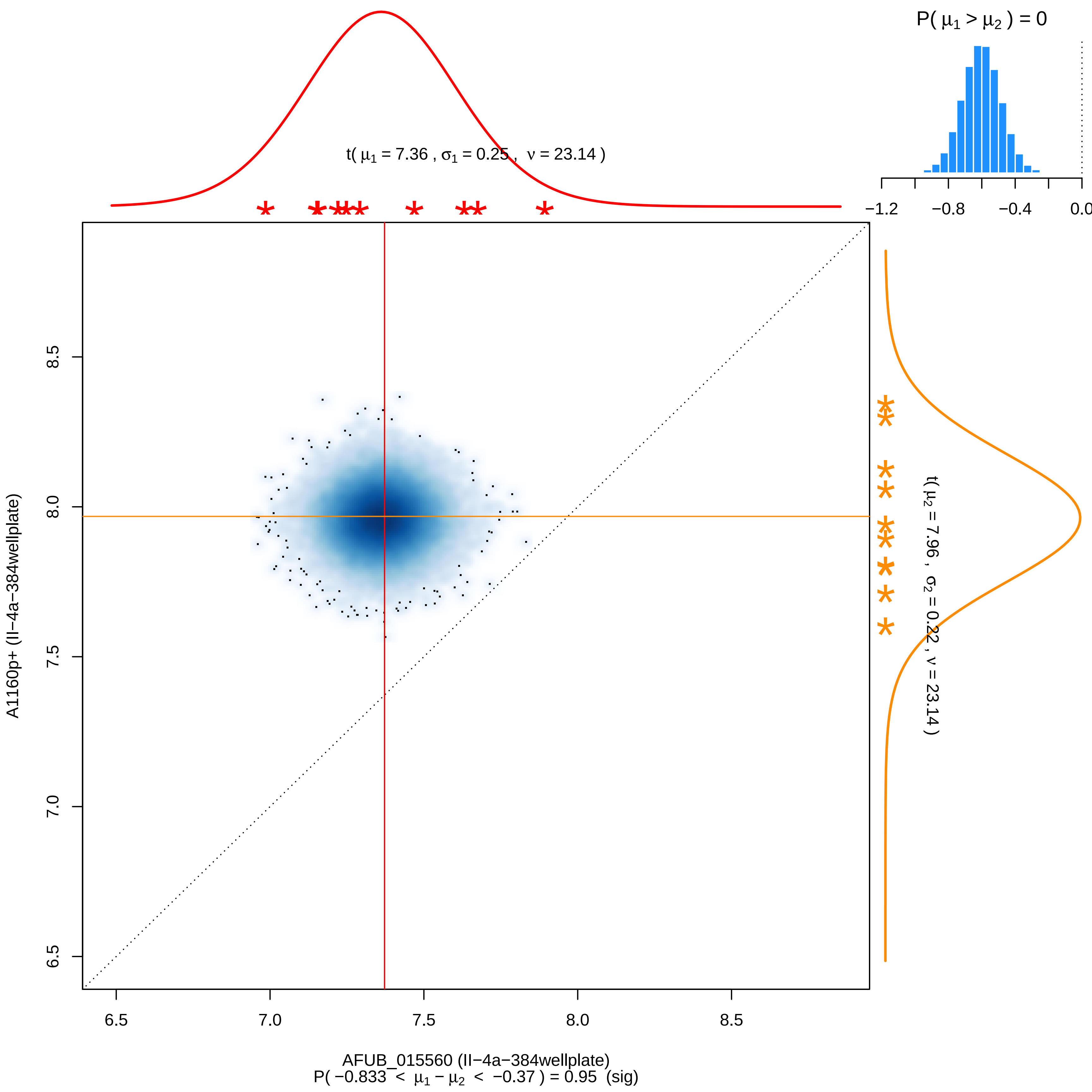

P( μ1 > μ2 ) = 0
−1.2
−0.8
−0.4
0.0
t( μ1 = 7.36 , σ1 = 0.25 , ν = 23.14 )
t( μ2 = 7.96 , σ2 = 0.22 , ν = 23.14 )
A1160p+ (II−4a−384wellplate)
8.5
8.0
7.5
7.0
6.5
6.5
7.0
7.5
8.0
8.5
AFUB_015560 (II−4a−384wellplate)
P( −0.833 < μ1 − μ2 < −0.37 ) = 0.95 (sig)

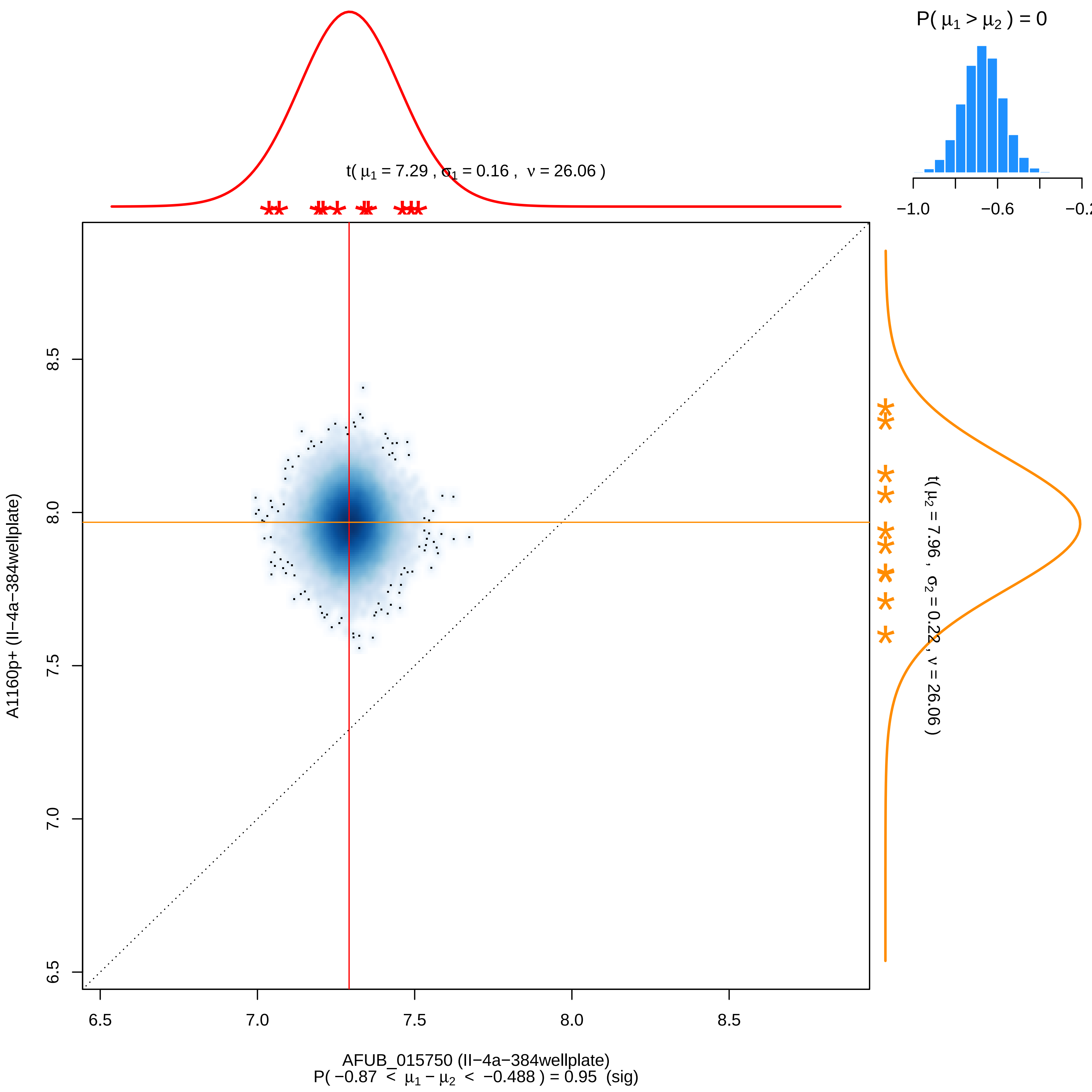

P( μ1 > μ2 ) = 0
−1.0
−0.6
−0.2
t( μ1 = 7.29 , σ1 = 0.16 , ν = 26.06 )
t( μ2 = 7.96 , σ2 = 0.22 , ν = 26.06 )
A1160p+ (II−4a−384wellplate)
6.5
7.0
7.5
8.0
8.5
AFUB_015750 (II−4a−384wellplate)
P( −0.87 < μ1 − μ2 < −0.488 ) = 0.95 (sig)

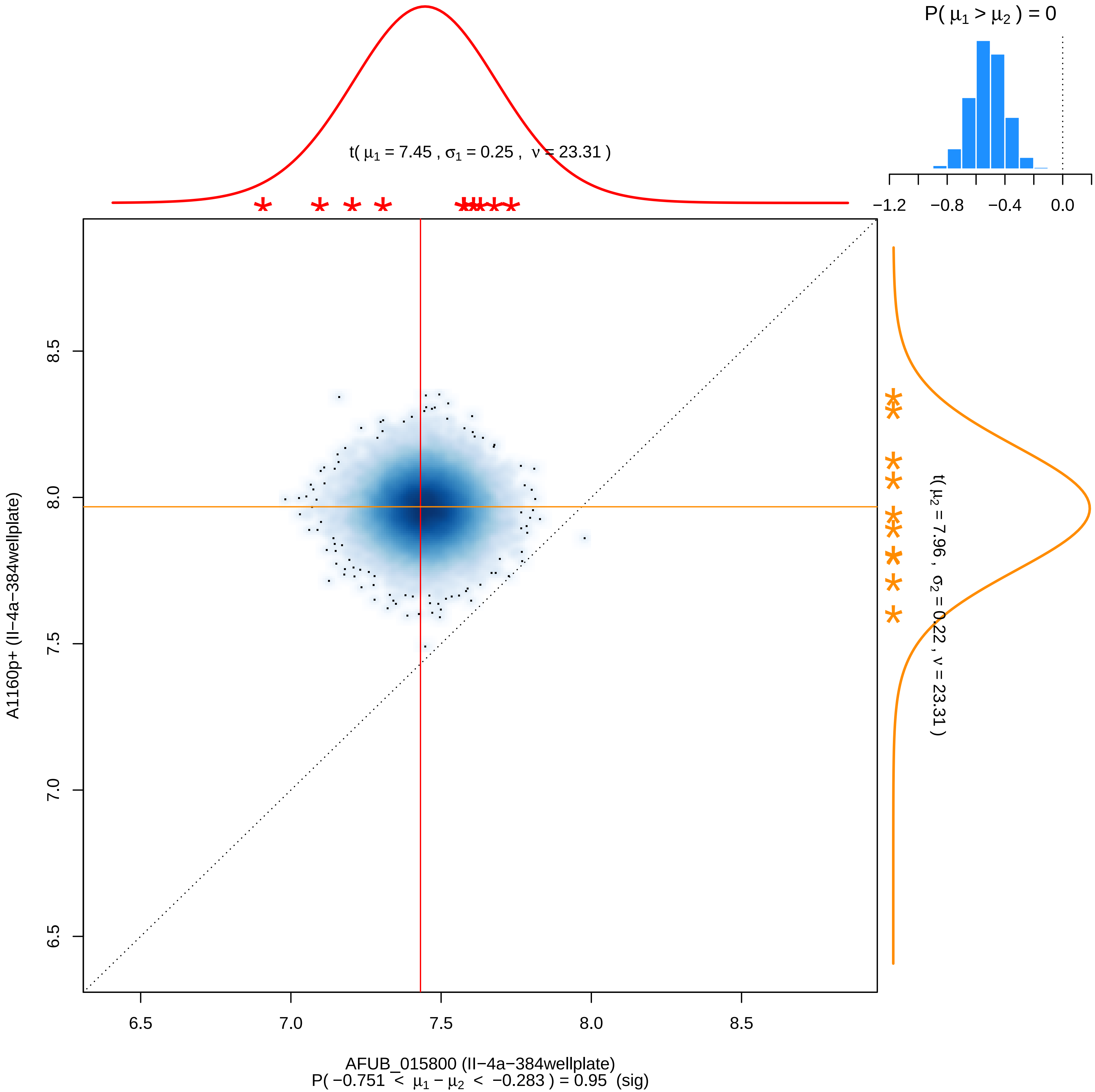

P( μ1 > μ2 ) = 0
−1.2
−0.8
−0.4
0.0
t( μ1 = 7.45 , σ1 = 0.25 , ν = 23.31 )
t( μ2 = 7.96 , σ2 = 0.22 , ν = 23.31 )
8.5
8.0
7.5
7.0
6.5
A1160p+ (II−4a−384wellplate)
6.5
7.0
7.5
8.0
8.5
AFUB_015800 (II−4a−384wellplate)
P( −0.751 < μ1 − μ2 < −0.283 ) = 0.95 (sig)

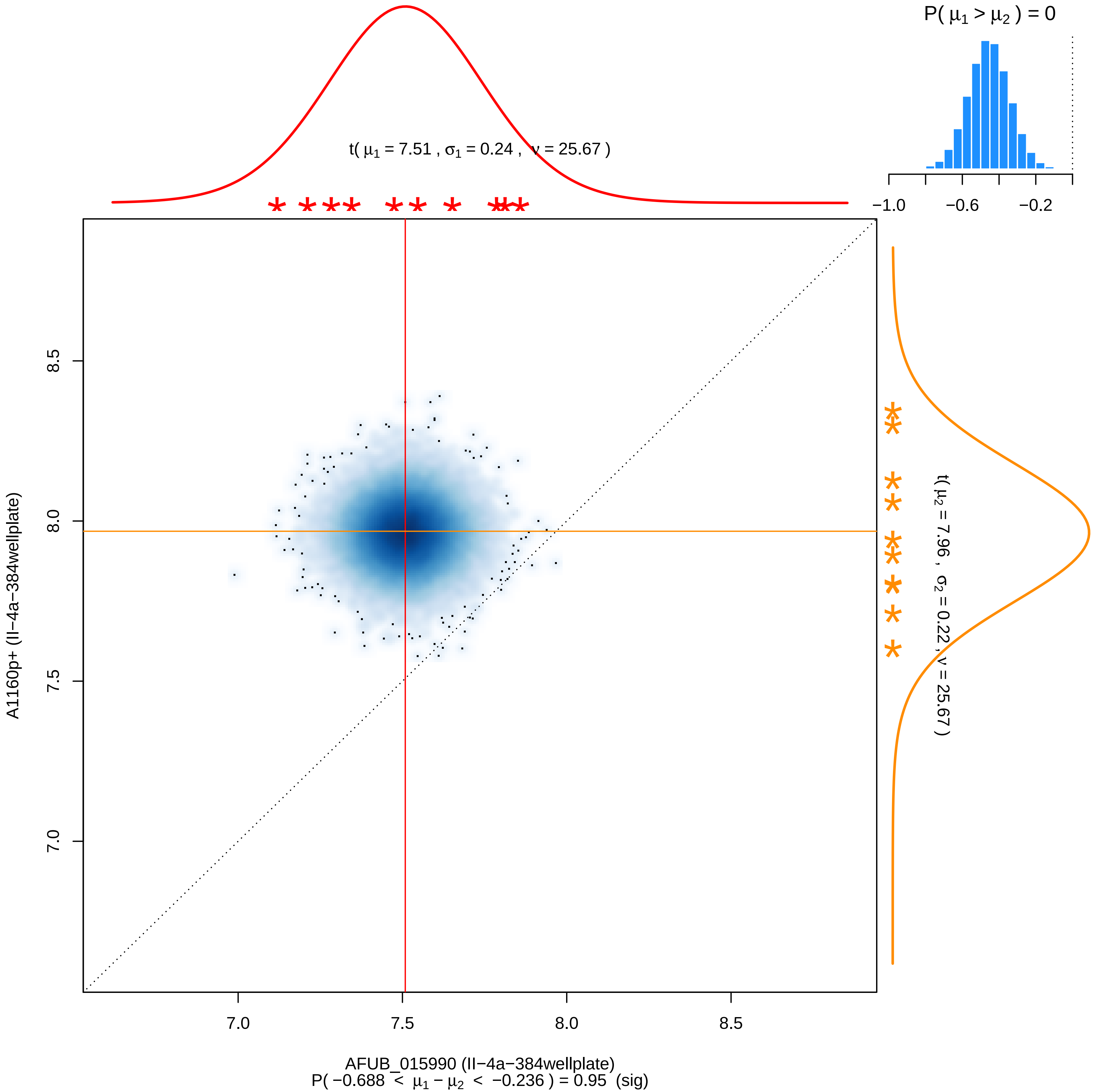

P( μ1 > μ2 ) = 0
−1.0
−0.6
−0.2
t( μ1 = 7.51 , σ1 = 0.24 , ν = 25.67 )
t( μ2 = 7.96 , σ2 = 0.22 , ν = 25.67 )
8.5
8.0
7.5
7.0
A1160p+ (II−4a−384wellplate)
7.0
7.5
8.0
8.5
AFUB_015990 (II−4a−384wellplate)
P( −0.688 < μ1 − μ2 < −0.236 ) = 0.95 (sig)

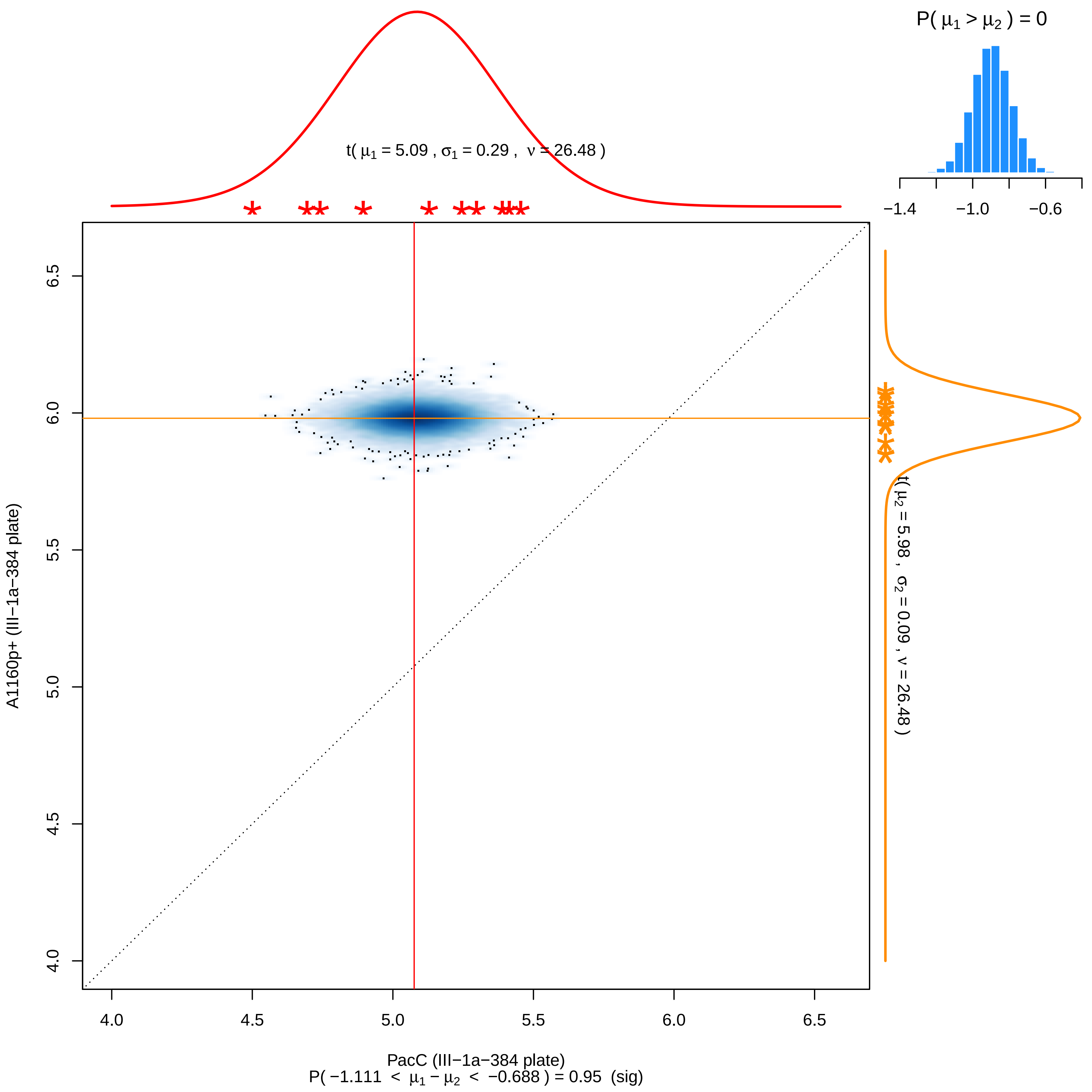

P( μ1 > μ2 ) = 0
−1.4
−1.0
−0.6
t( μ1 = 5.09 , σ1 = 0.29 , ν = 26.48 )
t( μ2 = 5.98 , σ2 = 0.09 , ν = 26.48 )
A1160p+ (III−1a−384 plate)
6.5
6.0
5.5
5.0
4.5
4.0
4.0
4.5
5.0
5.5
6.0
6.5
PacC (III−1a−384 plate)
P( −1.111 < μ1 − μ2 < −0.688 ) = 0.95 (sig)

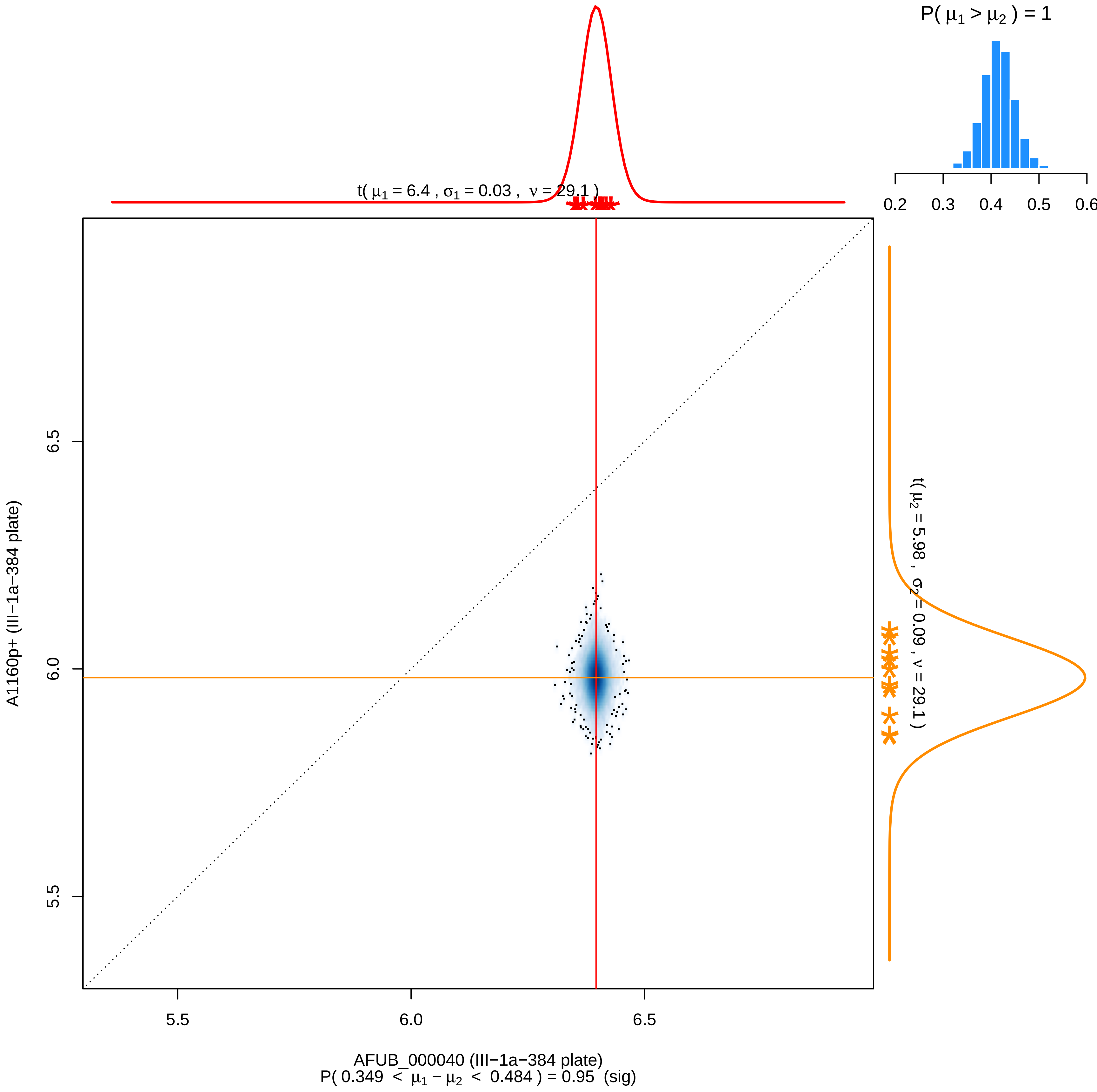

P( μ1 > μ2 ) = 1
0.2
0.3
0.4
0.5
0.6
t( μ1 = 6.4 , σ1 = 0.03 , ν = 29.1 )
t( μ2 = 5.98 , σ2 = 0.09 , ν = 29.1 )
A1160p+ (III−1a−384 plate)
6.5
6.0
5.5
5.5
6.0
6.5
AFUB_000040 (III−1a−384 plate)
P( 0.349 < μ1 − μ2 < 0.484 ) = 0.95 (sig)

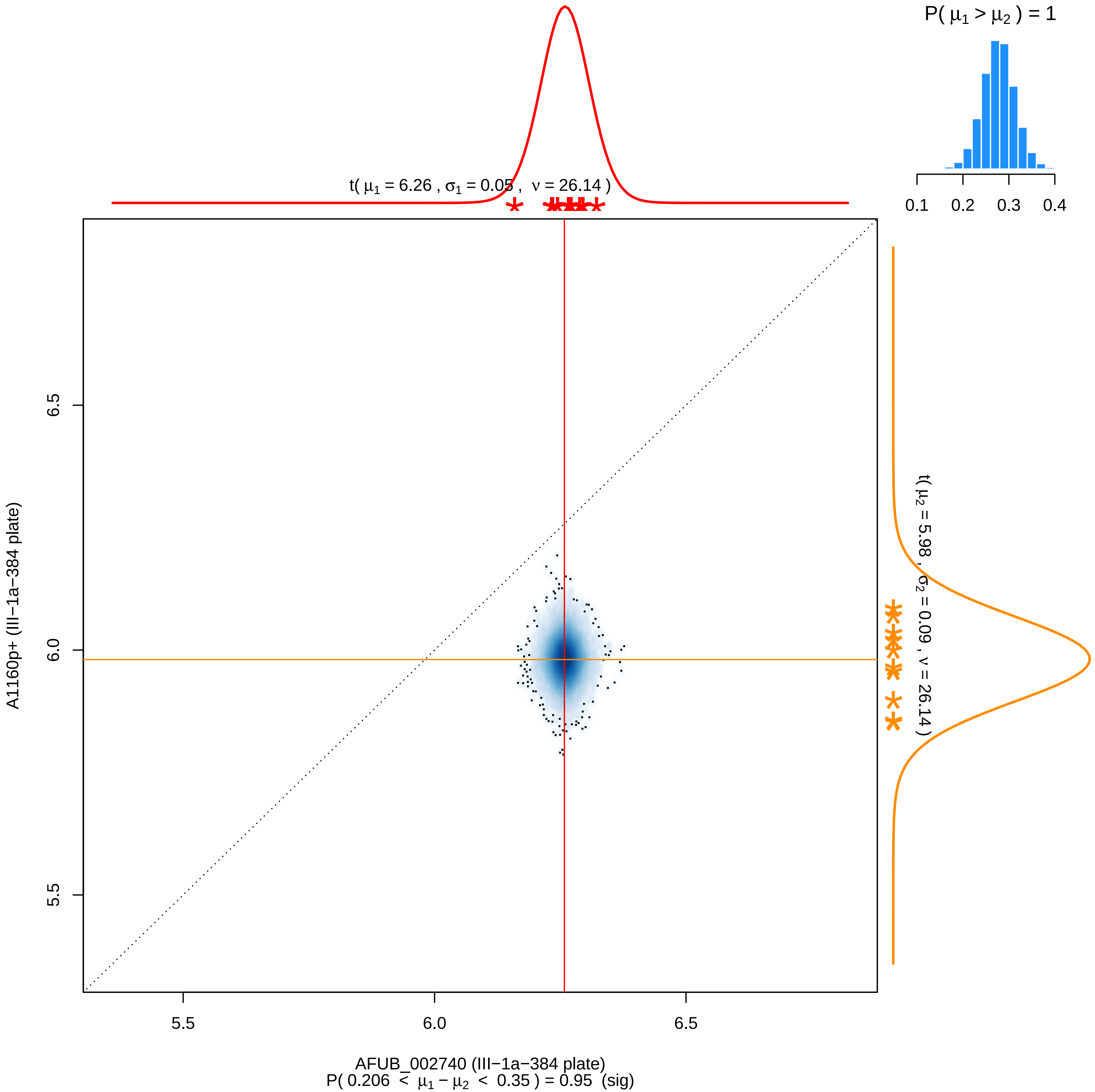
P( μ1 > μ2 ) = 1
0.1 0.2 0.3 0.4
t( μ1 = 6.26 , σ1 = 0.05 , ν = 26.14 )
t( μ2 = 5.98 , σ2 = 0.09 , ν = 26.14 )
6.5
6.0
5.5
A1160p+ (III−1a−384 plate)
5.5
6.0
6.5
AFUB_002740 (III−1a−384 plate)
P( 0.206 < μ1 − μ2 < 0.35 ) = 0.95 (sig)

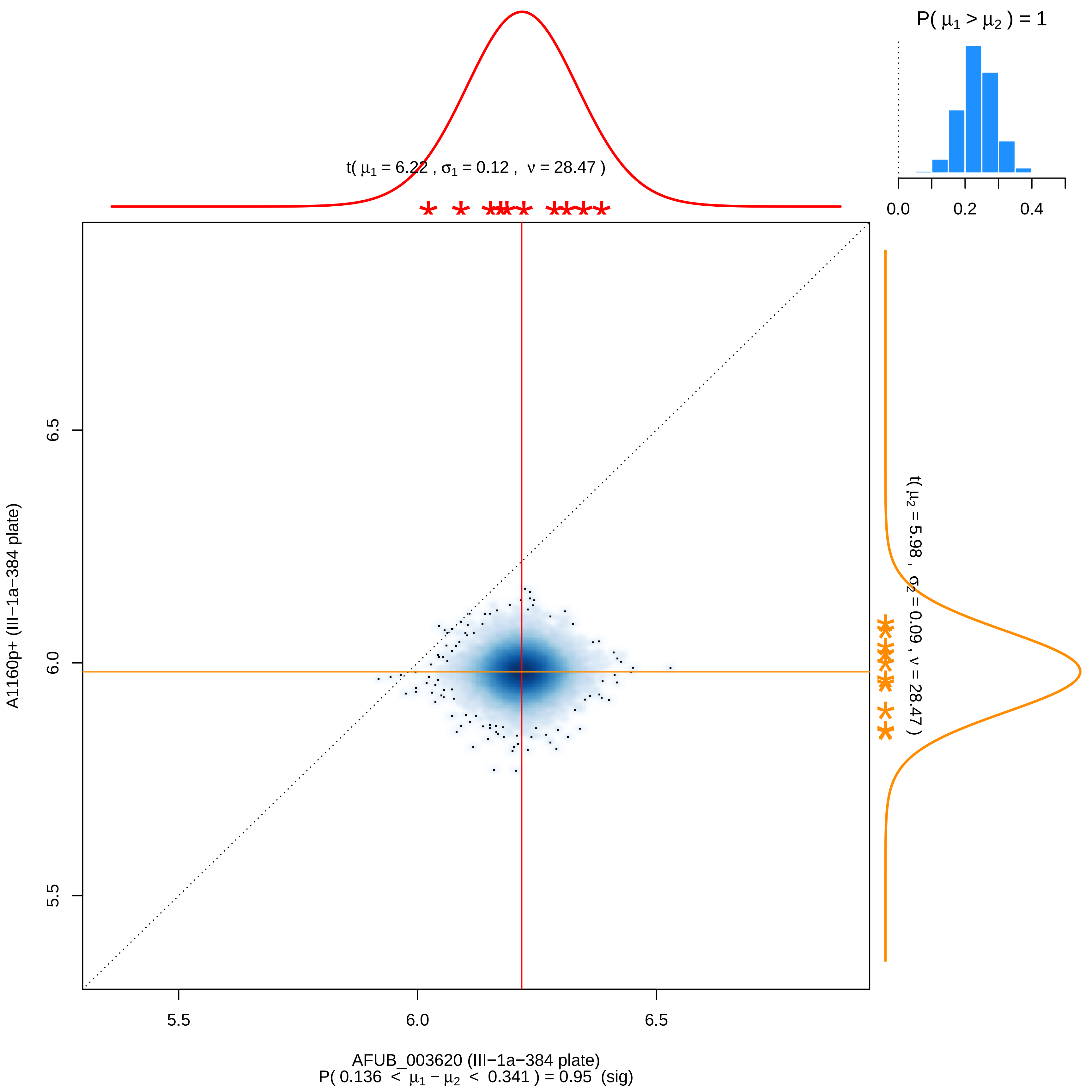

P( μ1 > μ2 ) = 1
0.0
0.2
0.4
t( μ1 = 6.22 , σ1 = 0.12 , ν = 28.47 )
t( μ2 = 5.98 , σ2 = 0.09 , ν = 28.47 )
6.5
6.0
5.5
A1160p+ (III−1a−384 plate)
5.5
6.0
6.5
AFUB_003620 (III−1a−384 plate)
P( 0.136 < μ1 − μ2 < 0.341 ) = 0.95 (sig)

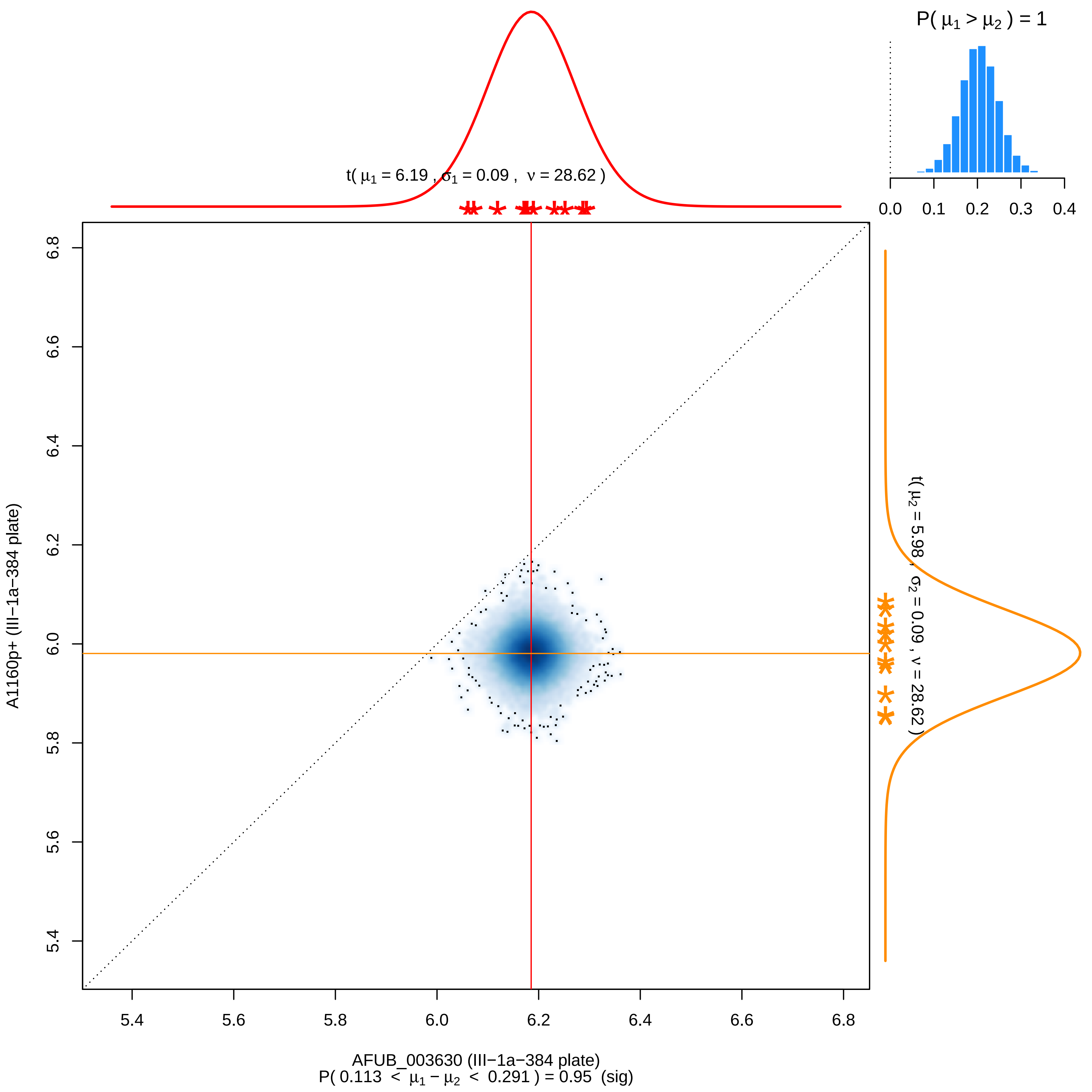

P( μ1 > μ2 ) = 1
0.0 0.1 0.2 0.3 0.4
t( μ1 = 6.19 , σ1 = 0.09 , ν = 28.62 )
t( μ2 = 5.98 , σ2 = 0.09 , ν = 28.62 )
6.8 6.6 6.4 6.2 6.0 5.8 5.6 5.4
A1160p+ (III−1a−384 plate)
5.4 5.6 5.8 6.0 6.2 6.4 6.6 6.8
AFUB_003630 (III−1a−384 plate)
P( 0.113 < μ1 − μ2 < 0.291 ) = 0.95 (sig)

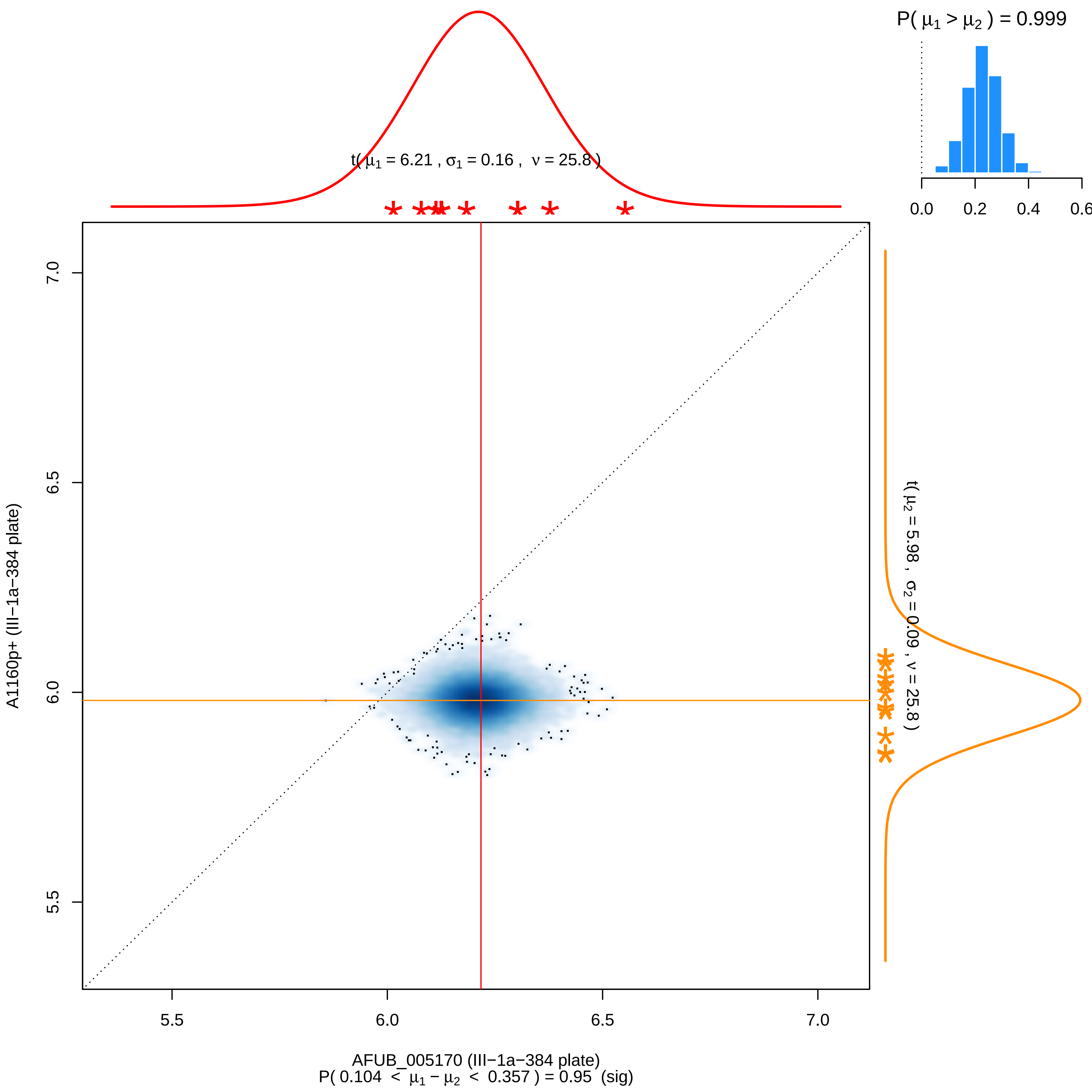

P( $\mu_1 > \mu_2$ ) = 0.999
0.0
0.2
0.4
0.6
t( $\mu_1$ = 6.21 , $\sigma_1$ = 0.16 , $\nu$ = 25.8 )
t( $\mu_2$ = 5.98 , $\sigma_2$ = 0.09 , $\nu$ = 25.8 )
A1160p+ (III−1a−384 plate)
7.0
6.5
6.0
5.5
AFUB_005170 (III−1a−384 plate)
P( 0.104 < $\mu_1 - \mu_2$ < 0.357 ) = 0.95 (sig)

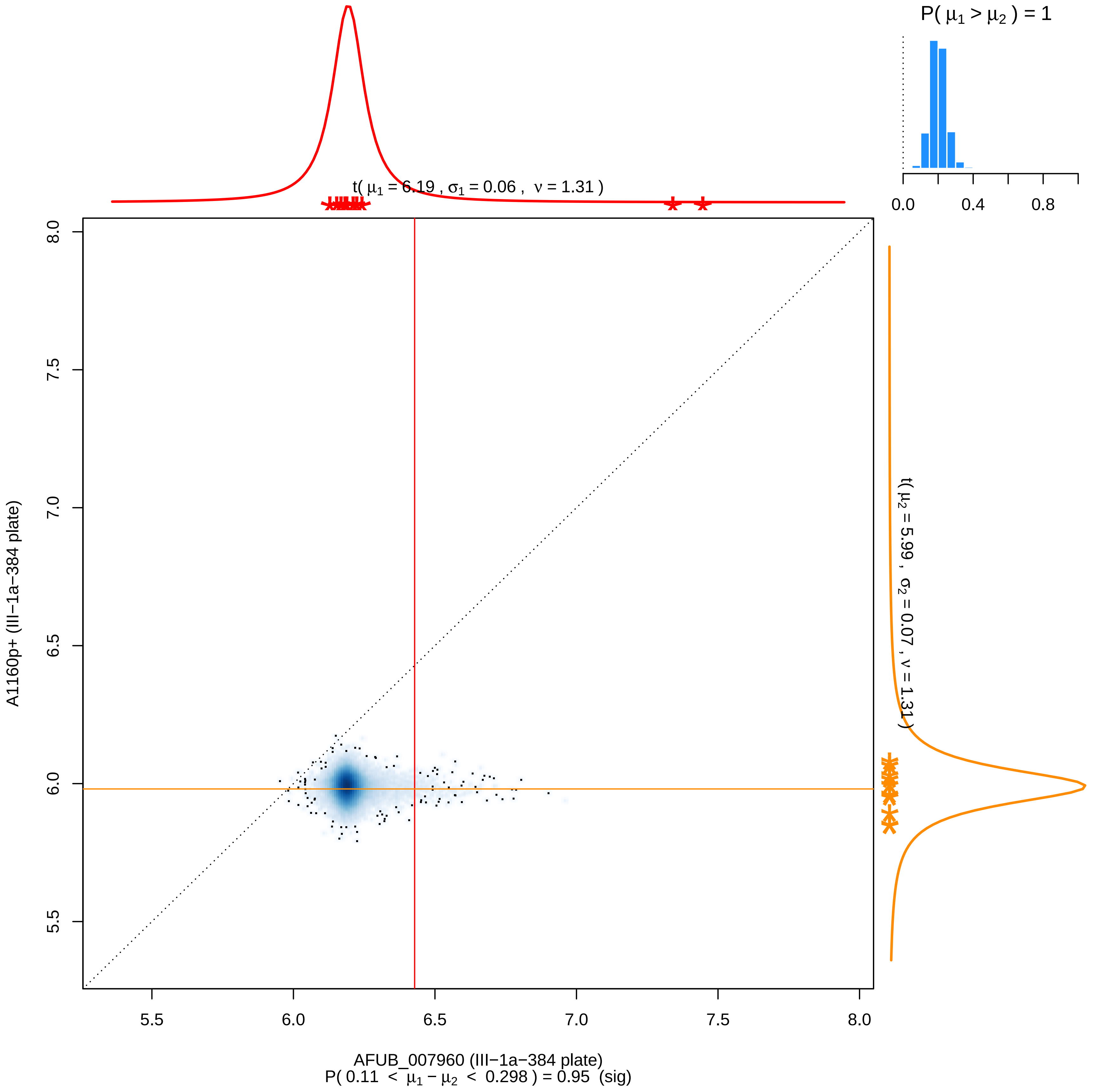

P( μ1 > μ2 ) = 1
0.0
0.4
0.8
t( μ1 = 6.19 , σ1 = 0.06 , ν = 1.31 )
t( μ2 = 5.99 , σ2 = 0.07 , ν = 1.31 )
A1160p+ (III−1a−384 plate)
5.5
6.0
6.5
7.0
7.5
8.0
AFUB_007960 (III−1a−384 plate)
P( 0.11 < μ1 − μ2 < 0.298 ) = 0.95 (sig)

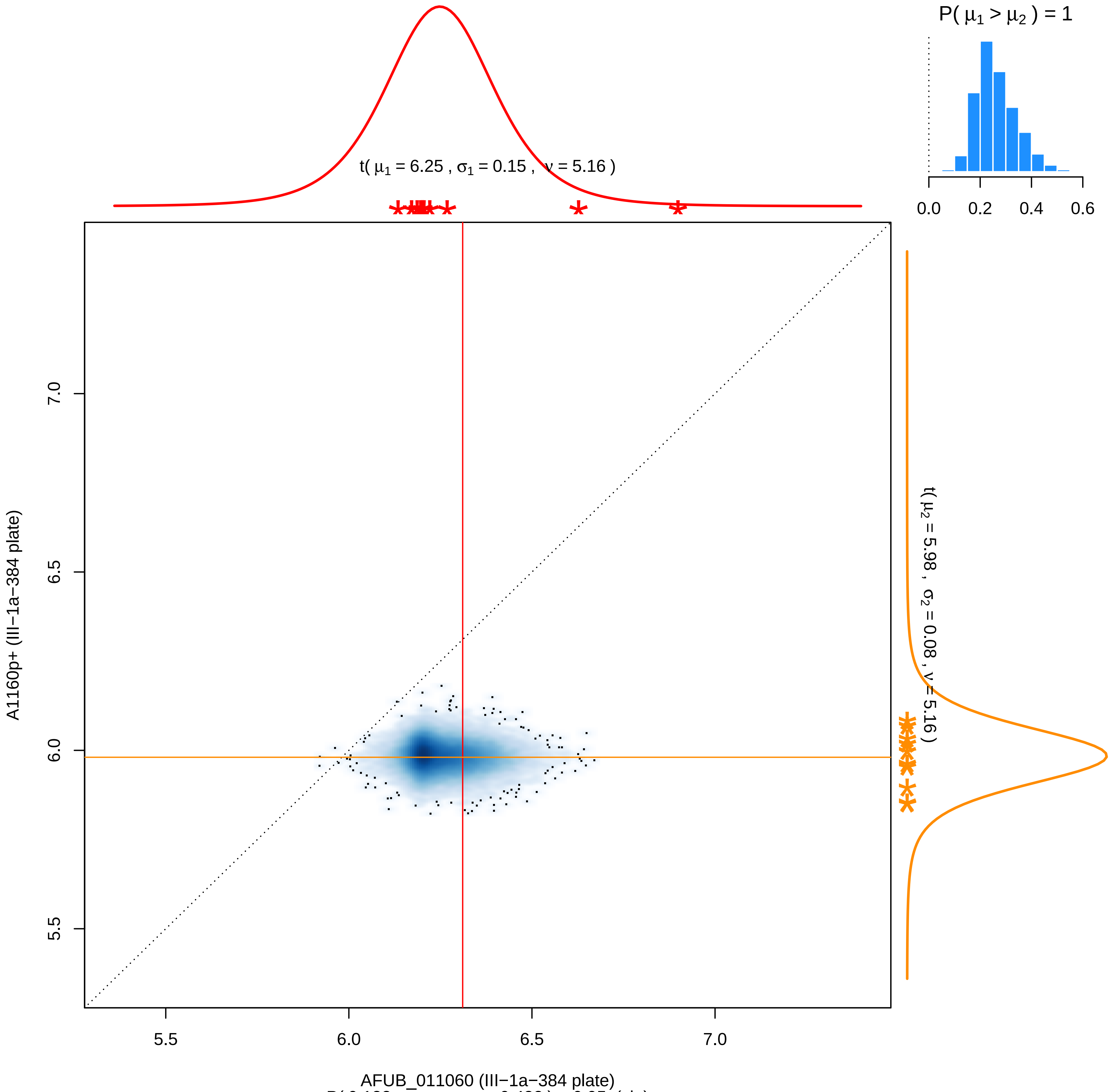

P( 0.132 < $\mu_1 - \mu_2$ < 0.432 ) = 0.95 (sig)

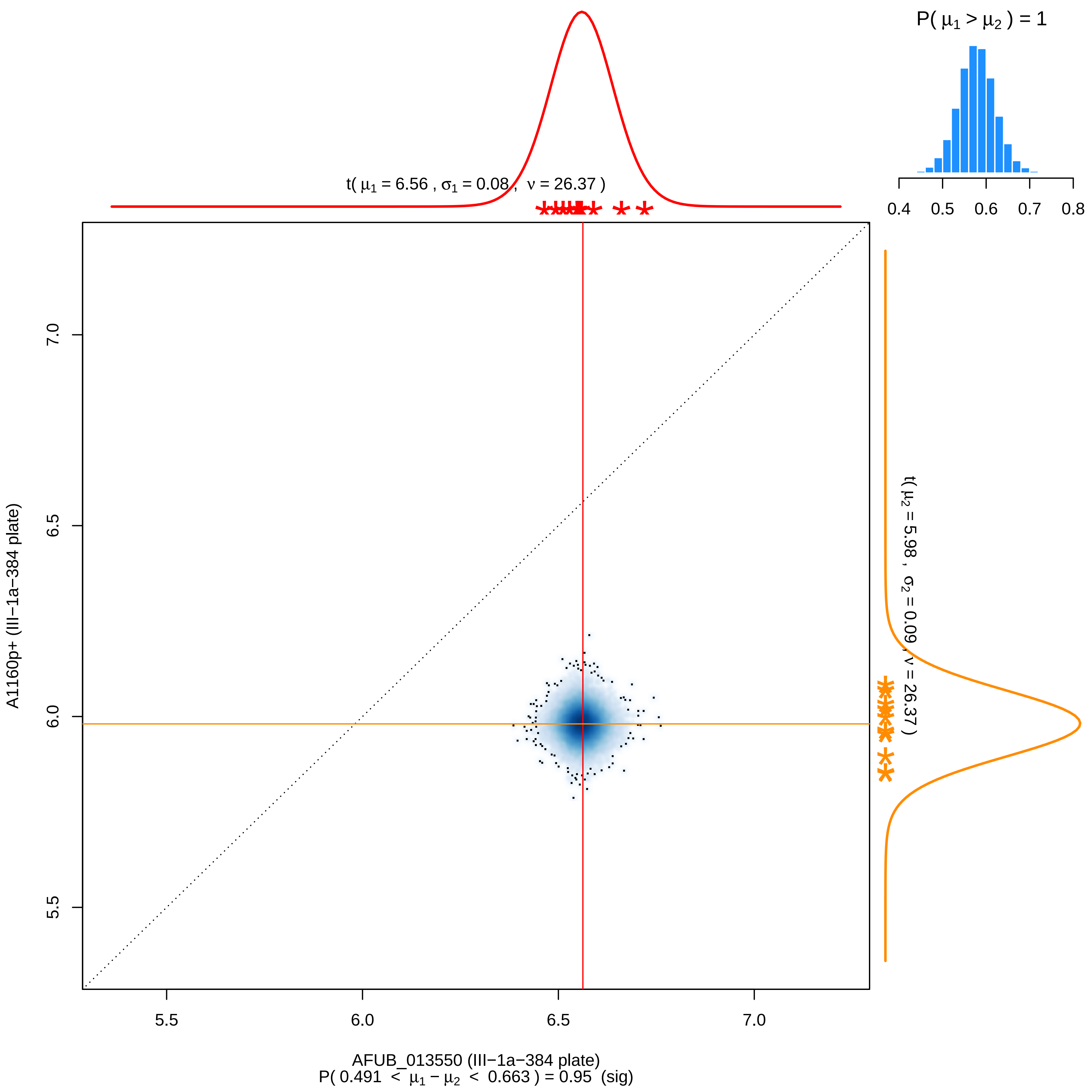

P( μ1 > μ2 ) = 1
0.4 0.5 0.6 0.7 0.8
t( μ1 = 6.56 , σ1 = 0.08 , ν = 26.37 )
t( μ2 = 5.98 , σ2 = 0.09 , ν = 26.37 )
A1160p+ (III−1a−384 plate)
7.0
6.5
6.0
5.5
5.5 6.0 6.5 7.0
AFUB_013550 (III−1a−384 plate)
P( 0.491 < μ1 − μ2 < 0.663 ) = 0.95 (sig)

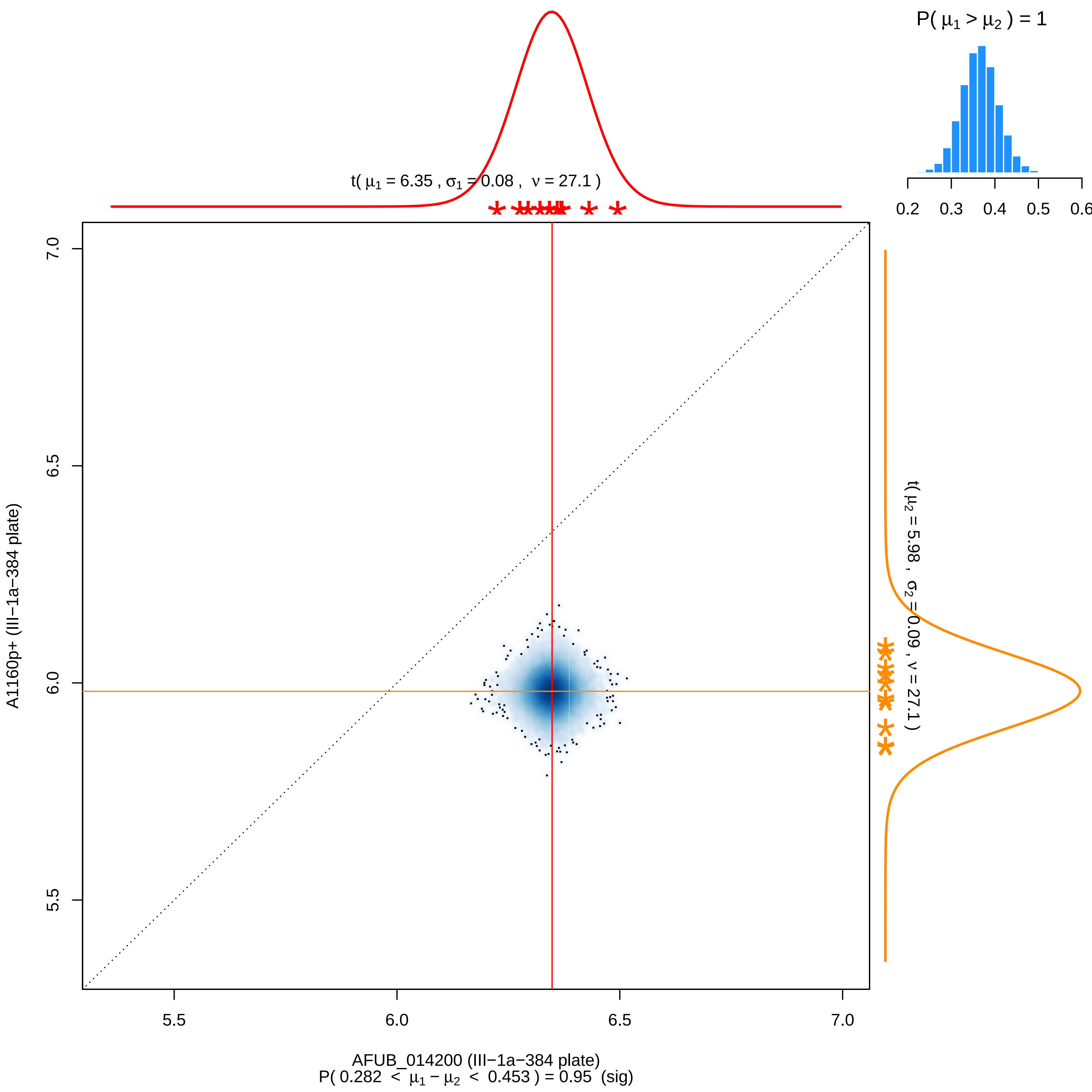
P( μ1 > μ2 ) = 1
0.2 0.3 0.4 0.5 0.6
t( μ1 = 6.35 , σ1 = 0.08 , ν = 27.1 )
t( μ2 = 5.98 , σ2 = 0.09 , ν = 27.1 )
7.0
6.5
6.0
5.5
A1160p+ (III−1a−384 plate)
5.5 6.0 6.5 7.0
AFUB_014200 (III−1a−384 plate)
P( 0.282 < μ1 − μ2 < 0.453 ) = 0.95 (sig)

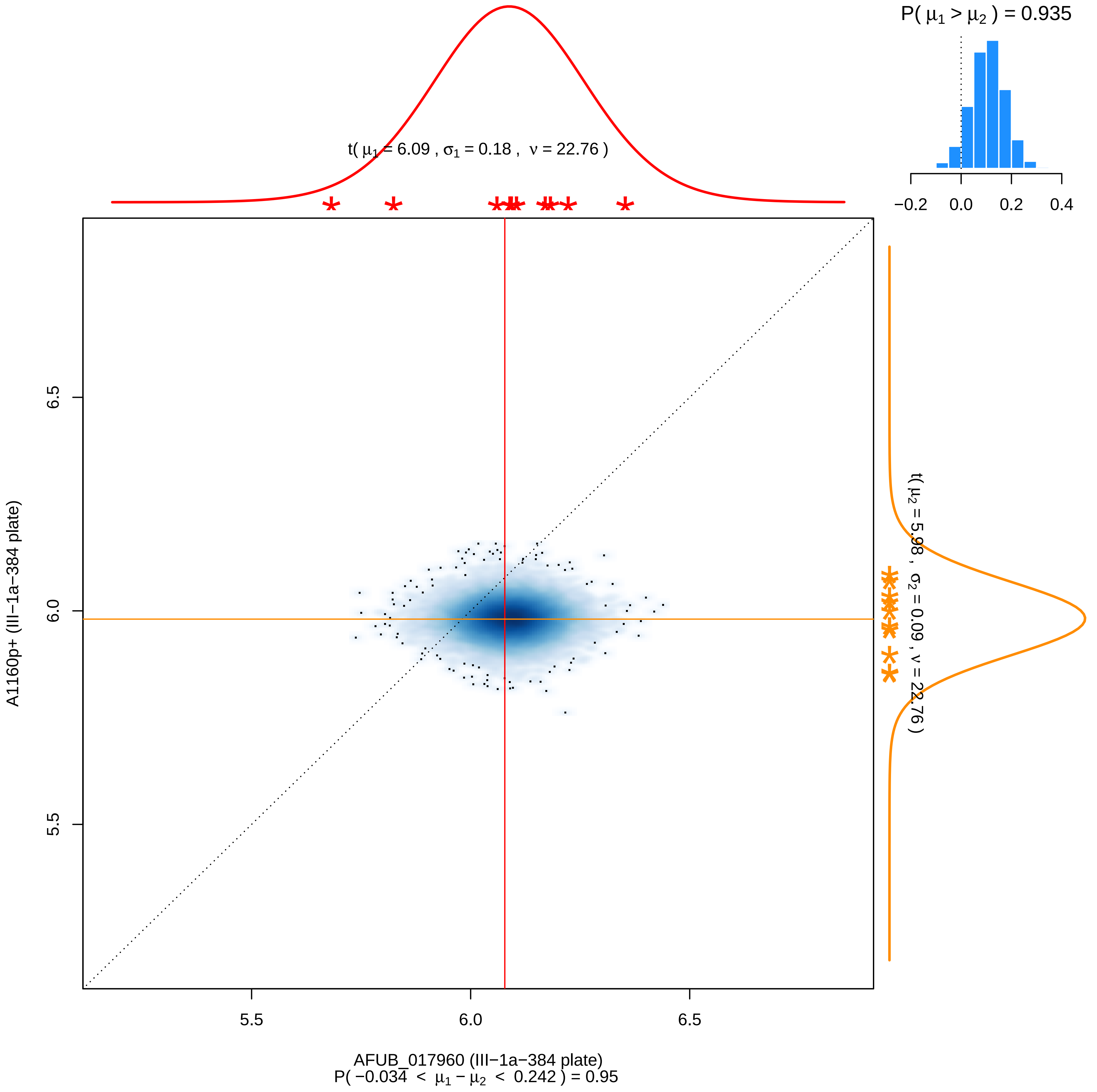

P( μ1 > μ2 ) = 0.935
−0.2 0.0 0.2 0.4
t( μ1 = 6.09 , σ1 = 0.18 , ν = 22.76 )
t( μ2 = 5.98 , σ2 = 0.09 , ν = 22.76 )
6.5
6.0
5.5
A1160p+ (III−1a−384 plate)
5.5 6.0 6.5
AFUB_017960 (III−1a−384 plate)
P( −0.034 < μ1 − μ2 < 0.242 ) = 0.95

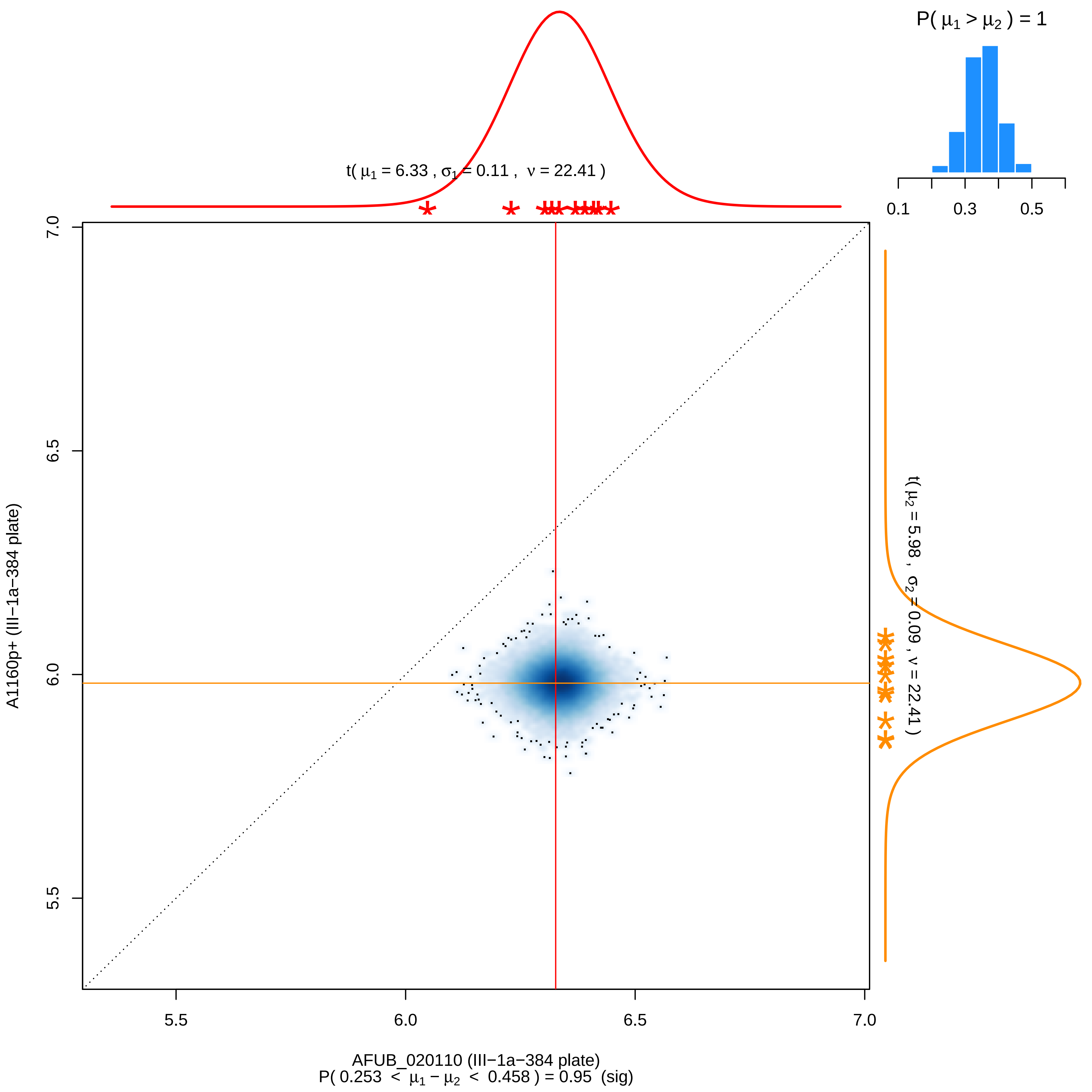
P( μ1 > μ2 ) = 1
0.1
0.3
0.5
t( μ1 = 6.33 , σ1 = 0.11 , ν = 22.41 )
t( μ2 = 5.98 , σ2 = 0.09 , ν = 22.41 )
7.0
6.5
6.0
5.5
A1160p+ (III−1a−384 plate)
5.5
6.0
6.5
7.0
AFUB_020110 (III−1a−384 plate)
P( 0.253 < μ1 − μ2 < 0.458 ) = 0.95 (sig)

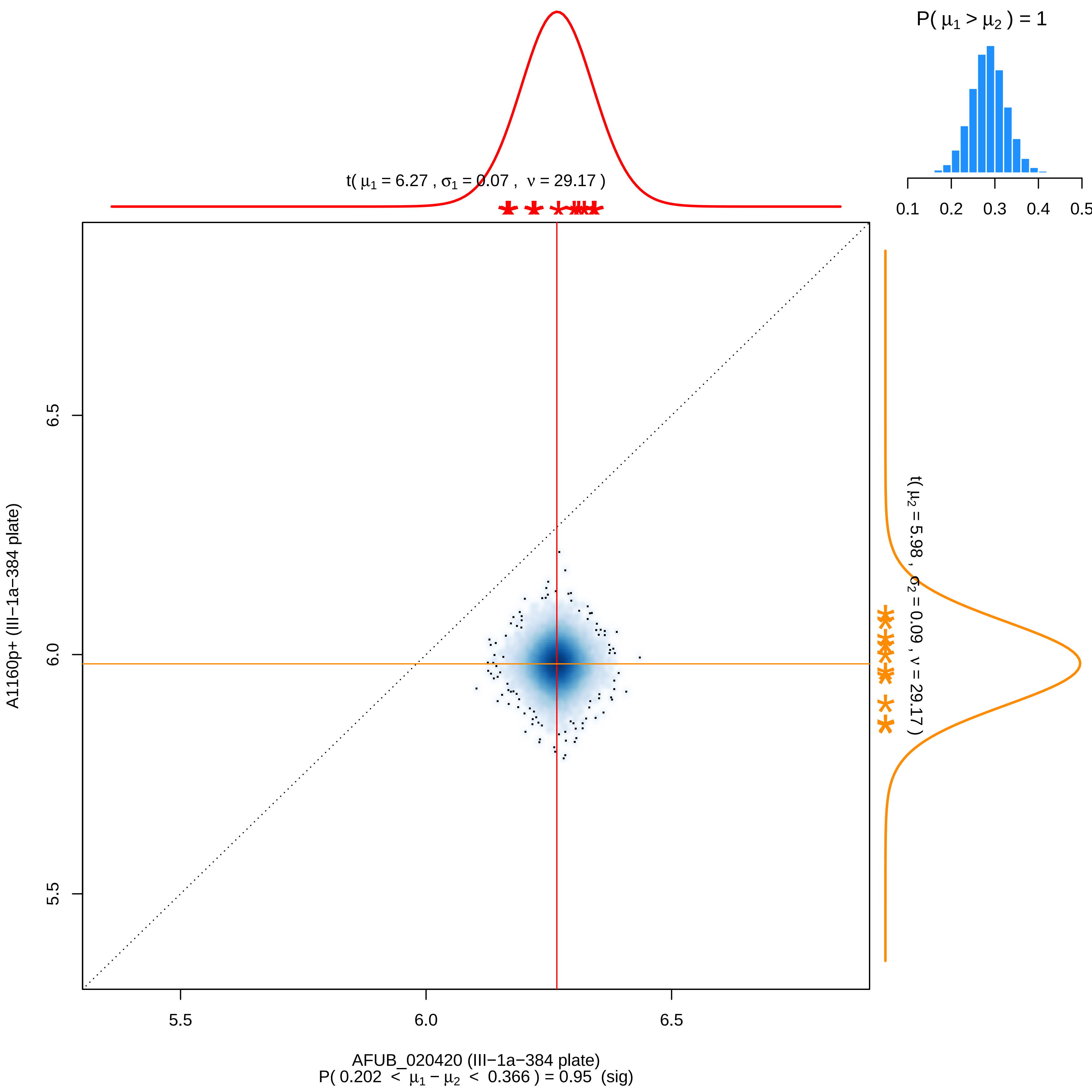

P( μ1 > μ2 ) = 1
0.1 0.2 0.3 0.4 0.5
t( μ1 = 6.27 , σ1 = 0.07 , ν = 29.17 )
t( μ2 = 5.98 , σ2 = 0.09 , ν = 29.17 )
A1160p+ (III−1a−384 plate)
5.5
6.0
6.5
AFUB_020420 (III−1a−384 plate)
P( 0.202 < μ1 − μ2 < 0.366 ) = 0.95 (sig)

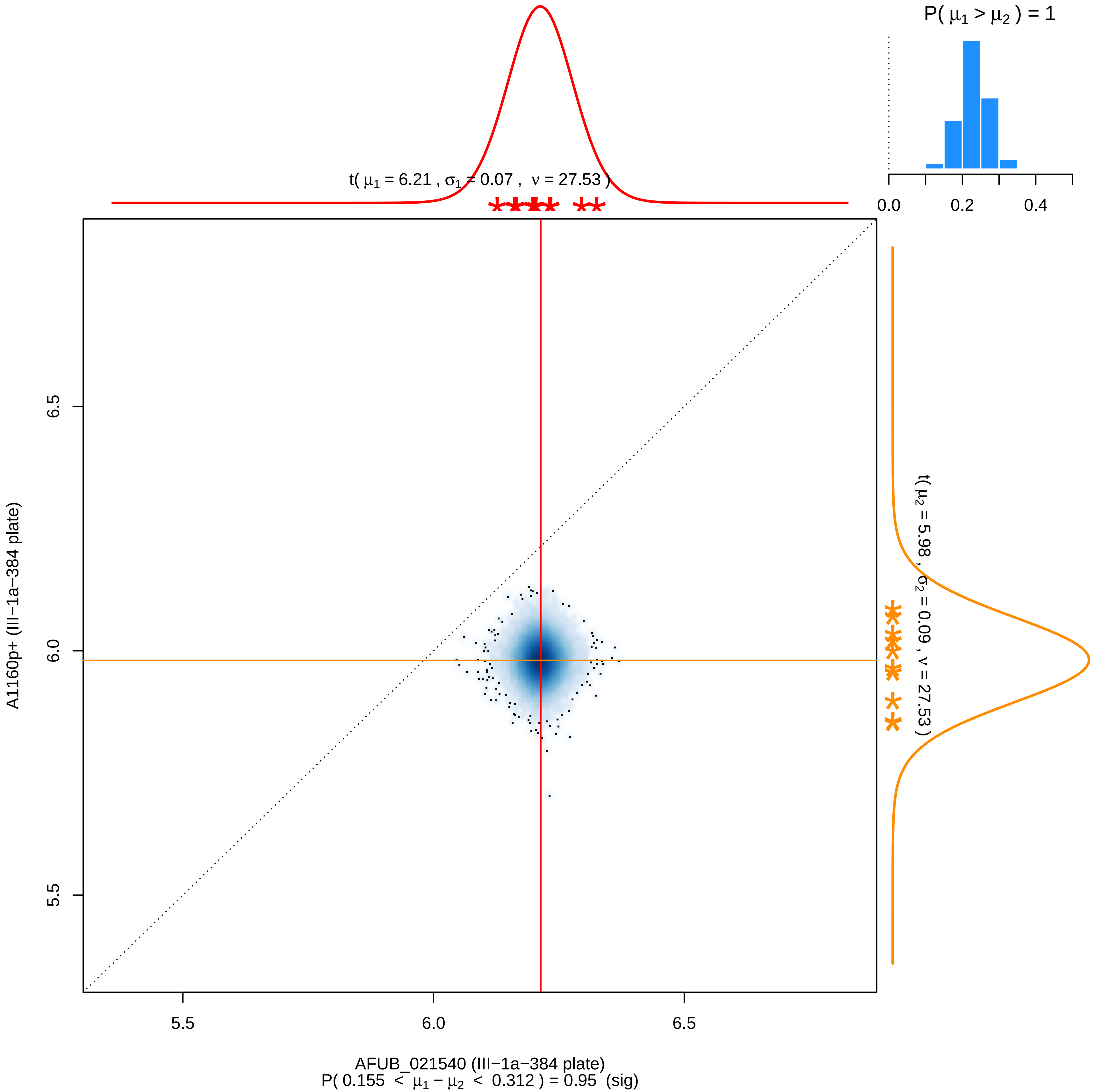

P( μ1 > μ2 ) = 1
0.0
0.2
0.4
t( μ1 = 6.21 , σ1 = 0.07 , ν = 27.53 )
t( μ2 = 5.98 , σ2 = 0.09 , ν = 27.53 )
6.5
6.0
5.5
A1160p+ (III−1a−384 plate)
AFUB_021540 (III−1a−384 plate)
P( 0.155 < μ1 − μ2 < 0.312 ) = 0.95 (sig)

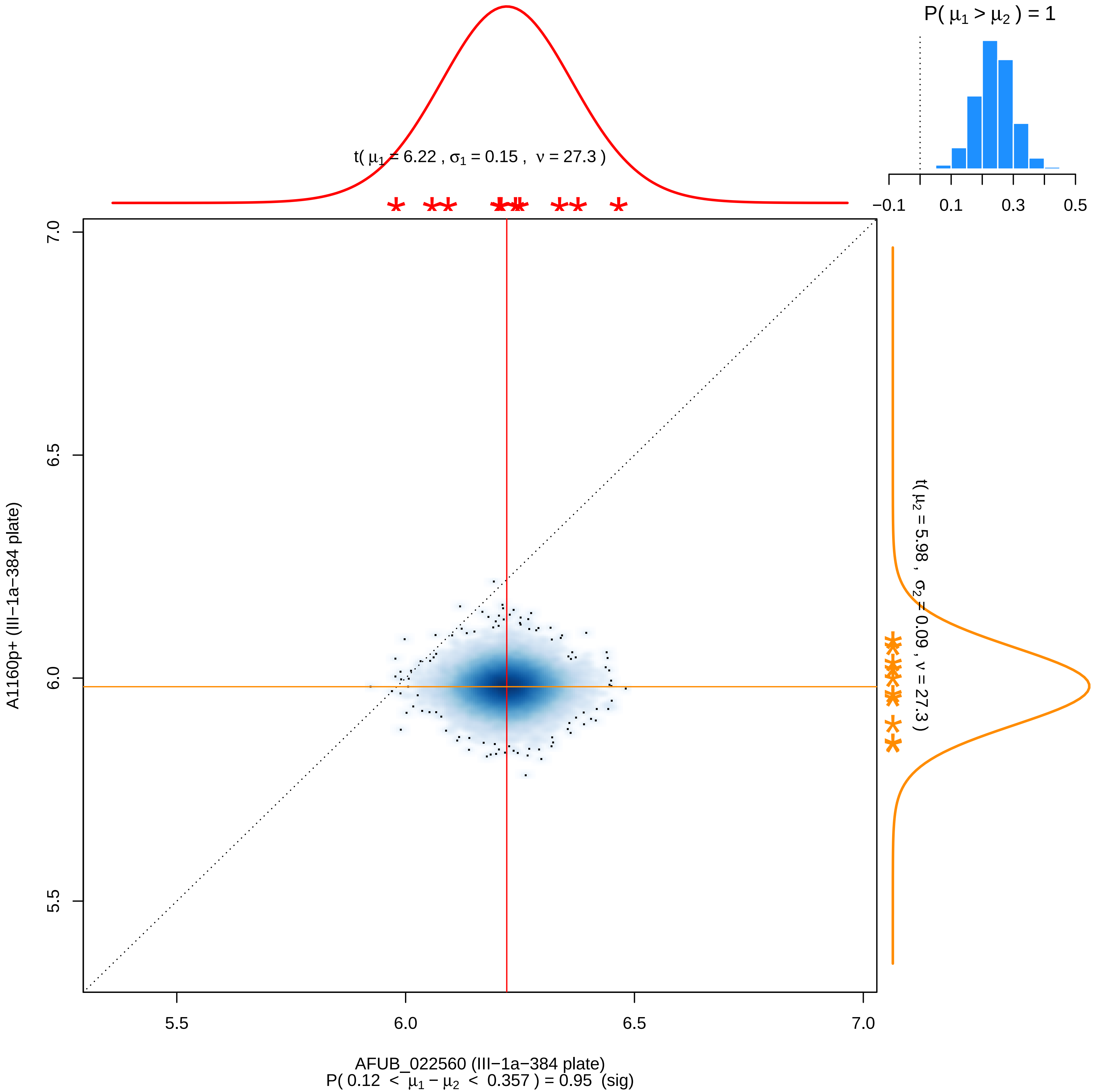

P( μ1 > μ2 ) = 1
−0.1
0.1
0.3
0.5
t( μ1 = 6.22 , σ1 = 0.15 , ν = 27.3 )
t( μ2 = 5.98 , σ2 = 0.09 , ν = 27.3 )
7.0
6.5
6.0
5.5
A1160p+ (III−1a−384 plate)
5.5
6.0
6.5
7.0
AFUB_022560 (III−1a−384 plate)
P( 0.12 < μ1 − μ2 < 0.357 ) = 0.95 (sig)

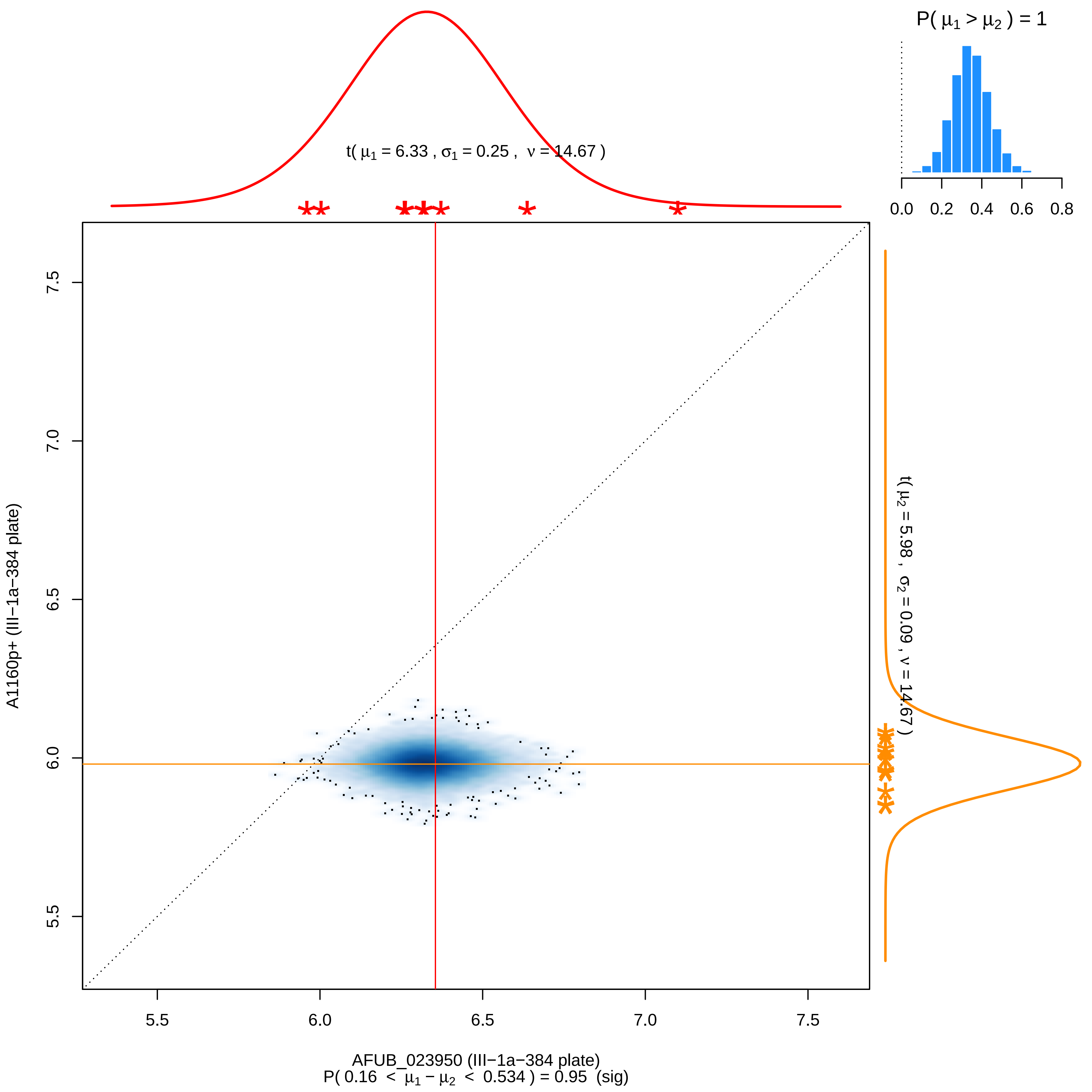
P( μ1 > μ2 ) = 1
0.0 0.2 0.4 0.6 0.8
t( μ1 = 6.33 , σ1 = 0.25 , ν = 14.67 )
t( μ2 = 5.98 , σ2 = 0.09 , ν = 14.67 )
A1160p+ (III−1a−384 plate)
5.5 6.0 6.5 7.0 7.5
AFUB_023950 (III−1a−384 plate)
P( 0.16 < μ1 − μ2 < 0.534 ) = 0.95 (sig)

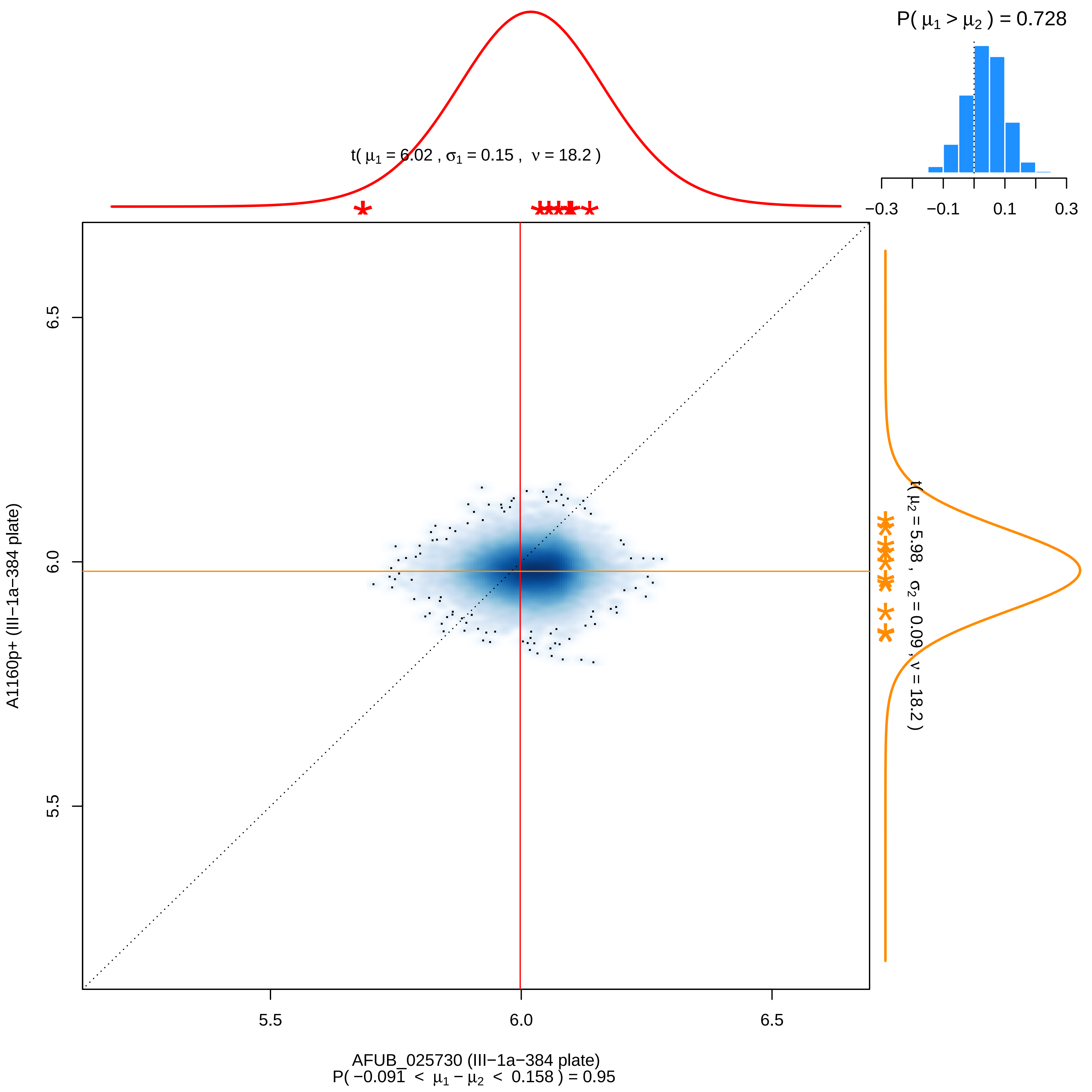

P( μ1 > μ2 ) = 0.728
−0.3
−0.1
0.1
0.3
t( μ1 = 6.02 , σ1 = 0.15 , ν = 18.2 )
t( μ2 = 5.98 , σ2 = 0.09 , ν = 18.2 )
6.5
6.0
5.5
A1160p+ (III−1a−384 plate)
5.5
6.0
6.5
AFUB_025730 (III−1a−384 plate)
P( −0.091 < μ1 − μ2 < 0.158 ) = 0.95

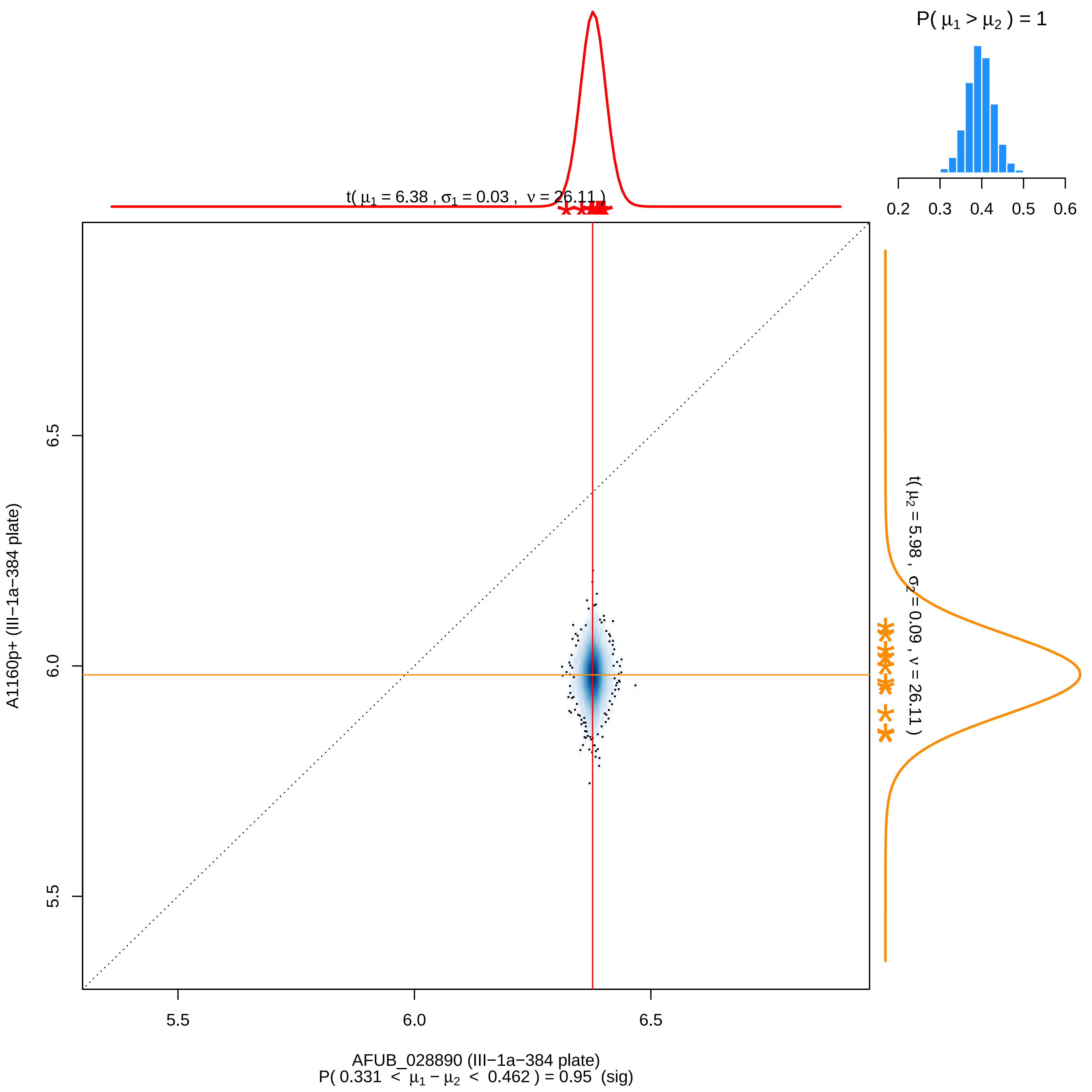

P( μ1 > μ2 ) = 1
0.2 0.3 0.4 0.5 0.6
t( μ1 = 6.38 , σ1 = 0.03 , ν = 26.11 )
t( μ2 = 5.98 , σ2 = 0.09 , ν = 26.11 )
A1160p+ (III−1a−384 plate)
5.5
6.0
6.5
AFUB_028890 (III−1a−384 plate)
P( 0.331 < μ1 − μ2 < 0.462 ) = 0.95 (sig)

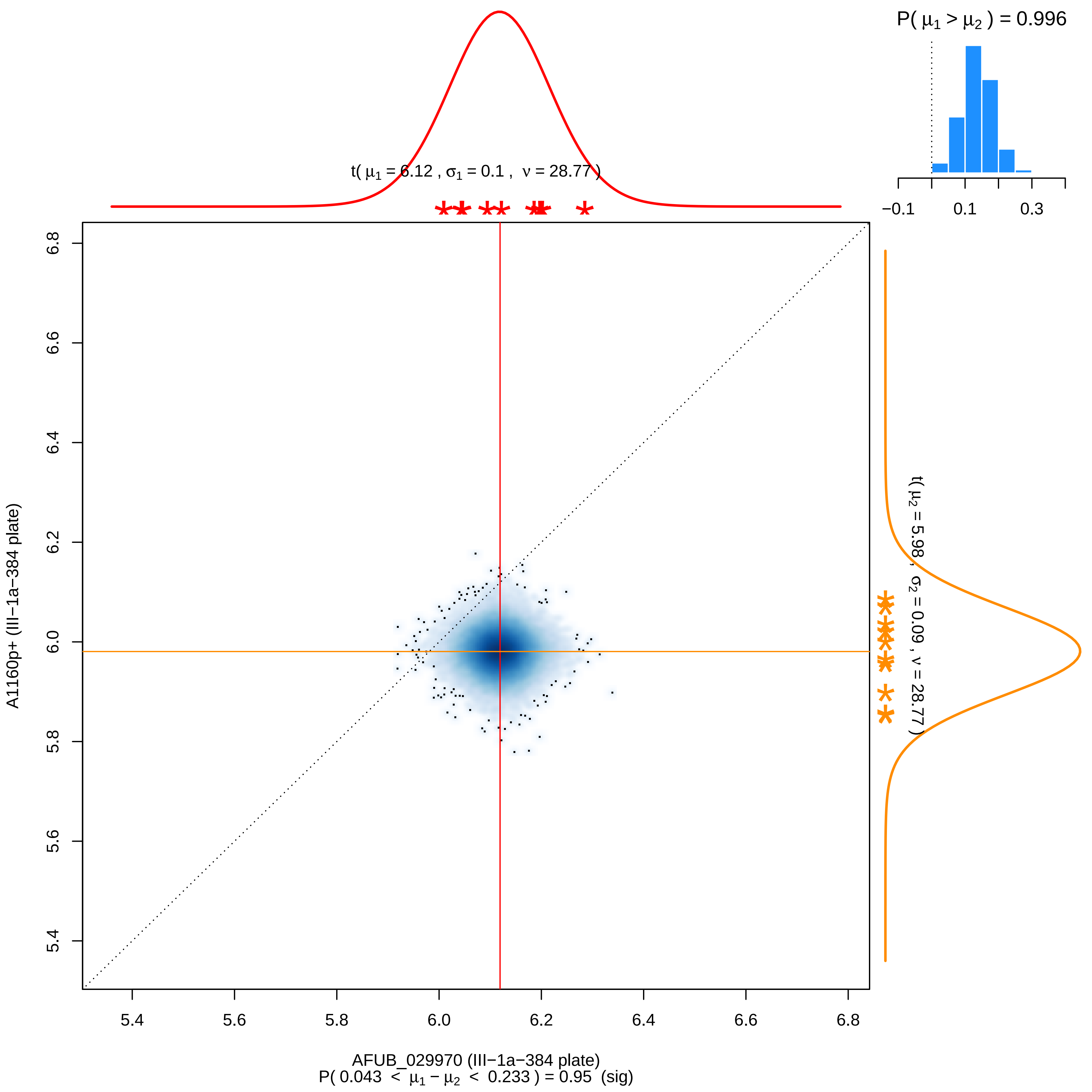

P( $\mu_1 > \mu_2$ ) = 0.996
−0.1
0.1
0.3
t( $\mu_1$ = 6.12 , $\sigma_1$ = 0.1 , $\nu$ = 28.77 )
t( $\mu_2$ = 5.98 , $\sigma_2$ = 0.09 , $\nu$ = 28.77 )
6.8
6.6
6.4
6.2
6.0
5.8
5.6
5.4
A1160p+ (III−1a−384 plate)
5.4
5.6
5.8
6.0
6.2
6.4
6.6
6.8
AFUB_029970 (III−1a−384 plate)
P( 0.043 < $\mu_1 - \mu_2$ < 0.233 ) = 0.95 (sig)

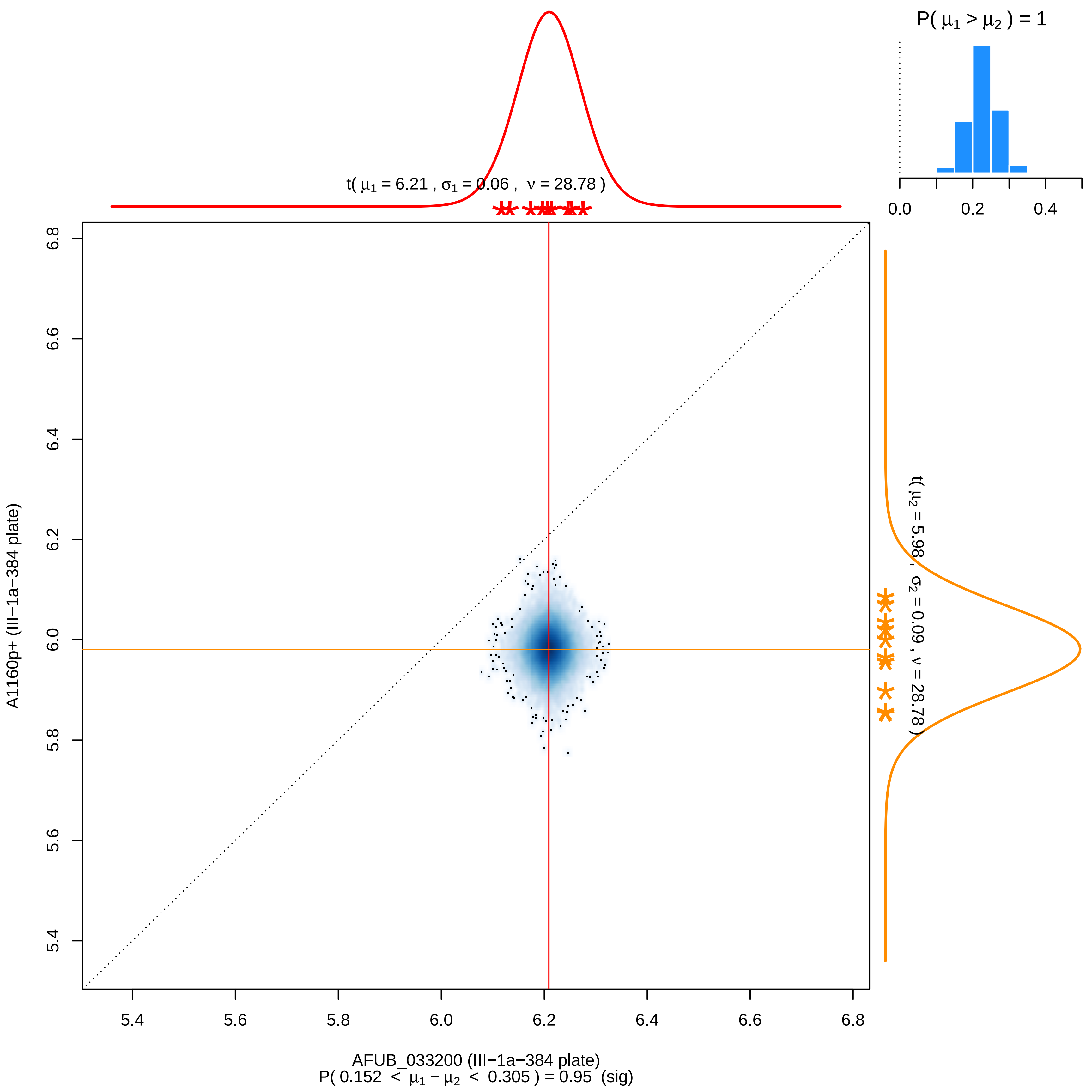

P( μ1 > μ2 ) = 1
0.0
0.2
0.4
t( μ1 = 6.21 , σ1 = 0.06 , ν = 28.78 )
t( μ2 = 5.98 , σ2 = 0.09 , ν = 28.78 )
6.8
6.6
6.4
6.2
6.0
5.8
5.6
5.4
A1160p+ (III−1a−384 plate)
5.4
5.6
5.8
6.0
6.2
6.4
6.6
6.8
AFUB_033200 (III−1a−384 plate)
P( 0.152 < μ1 − μ2 < 0.305 ) = 0.95 (sig)

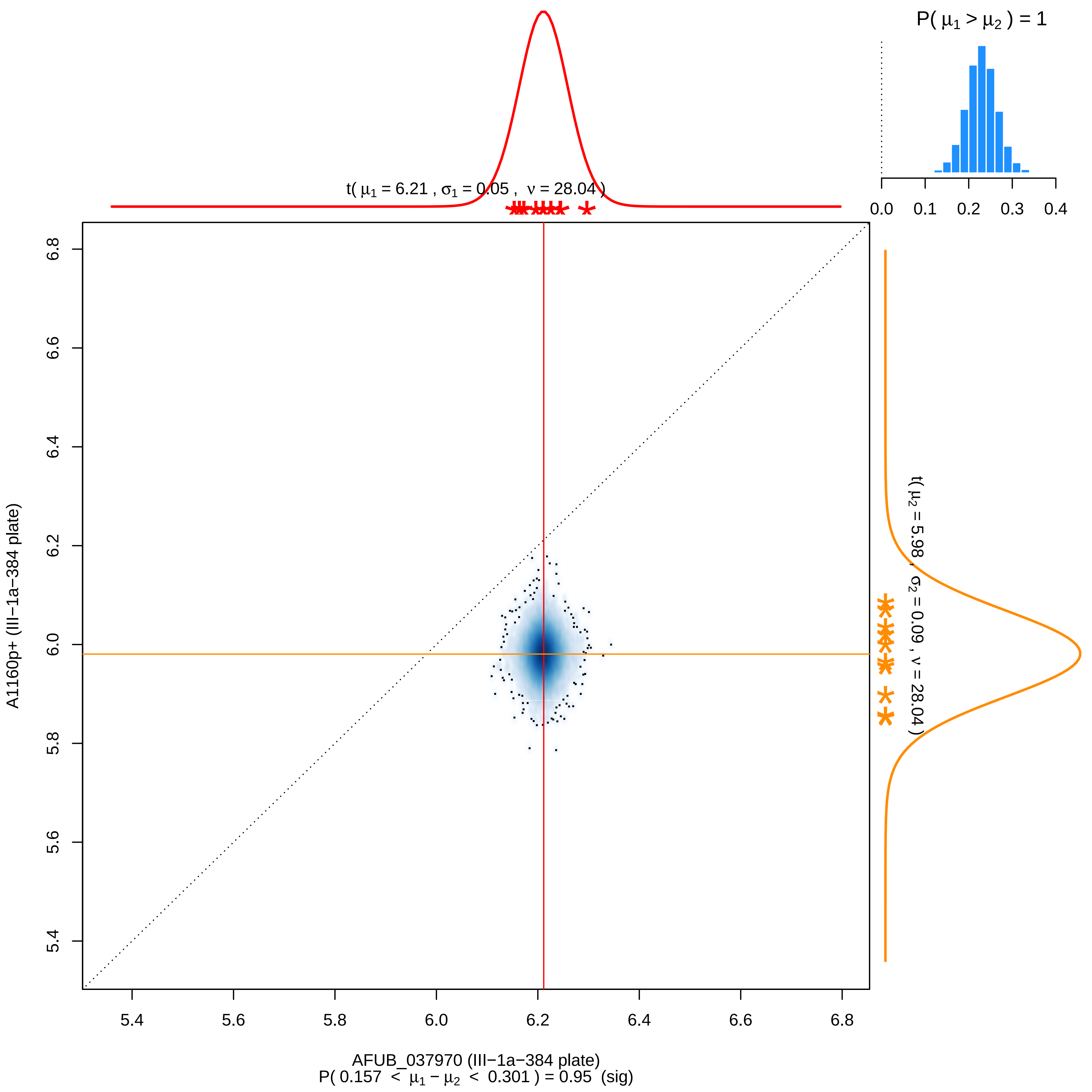

P( μ1 > μ2 ) = 1
0.0 0.1 0.2 0.3 0.4
t( μ1 = 6.21 , σ1 = 0.05 , ν = 28.04 )
t( μ2 = 5.98 , σ2 = 0.09 , ν = 28.04 )
A1160p+ (III−1a−384 plate)
6.8 6.6 6.4 6.2 6.0 5.8 5.6 5.4
5.4 5.6 5.8 6.0 6.2 6.4 6.6 6.8
AFUB_037970 (III−1a−384 plate)
P( 0.157 < μ1 − μ2 < 0.301 ) = 0.95 (sig)

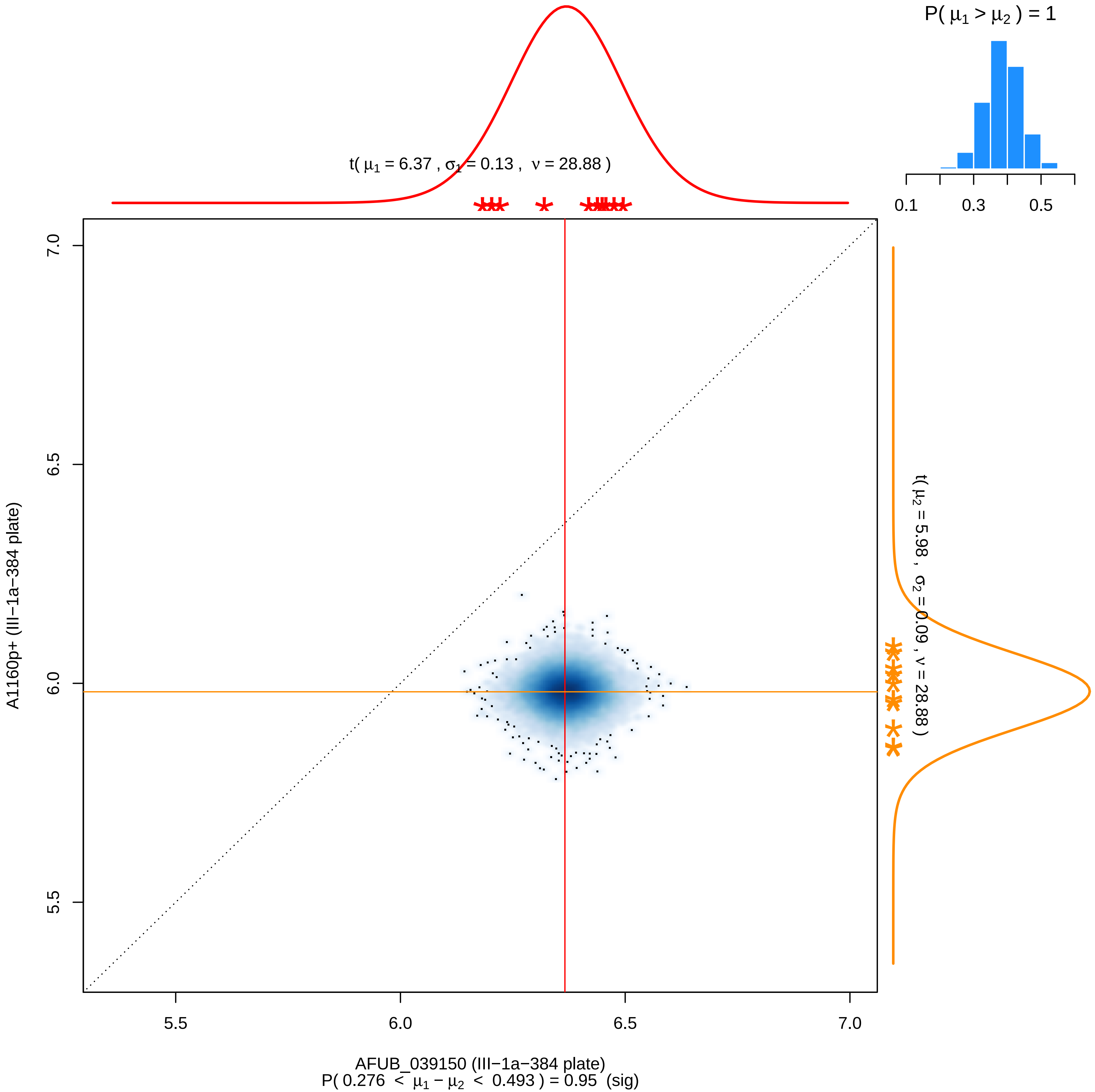

P( μ1 > μ2 ) = 1
0.1
0.3
0.5
t( μ1 = 6.37 , σ1 = 0.13 , ν = 28.88 )
t( μ2 = 5.98 , σ2 = 0.09 , ν = 28.88 )
A1160p+ (III−1a−384 plate)
7.0
6.5
6.0
5.5
AFUB_039150 (III−1a−384 plate)
P( 0.276 < μ1 − μ2 < 0.493 ) = 0.95 (sig)

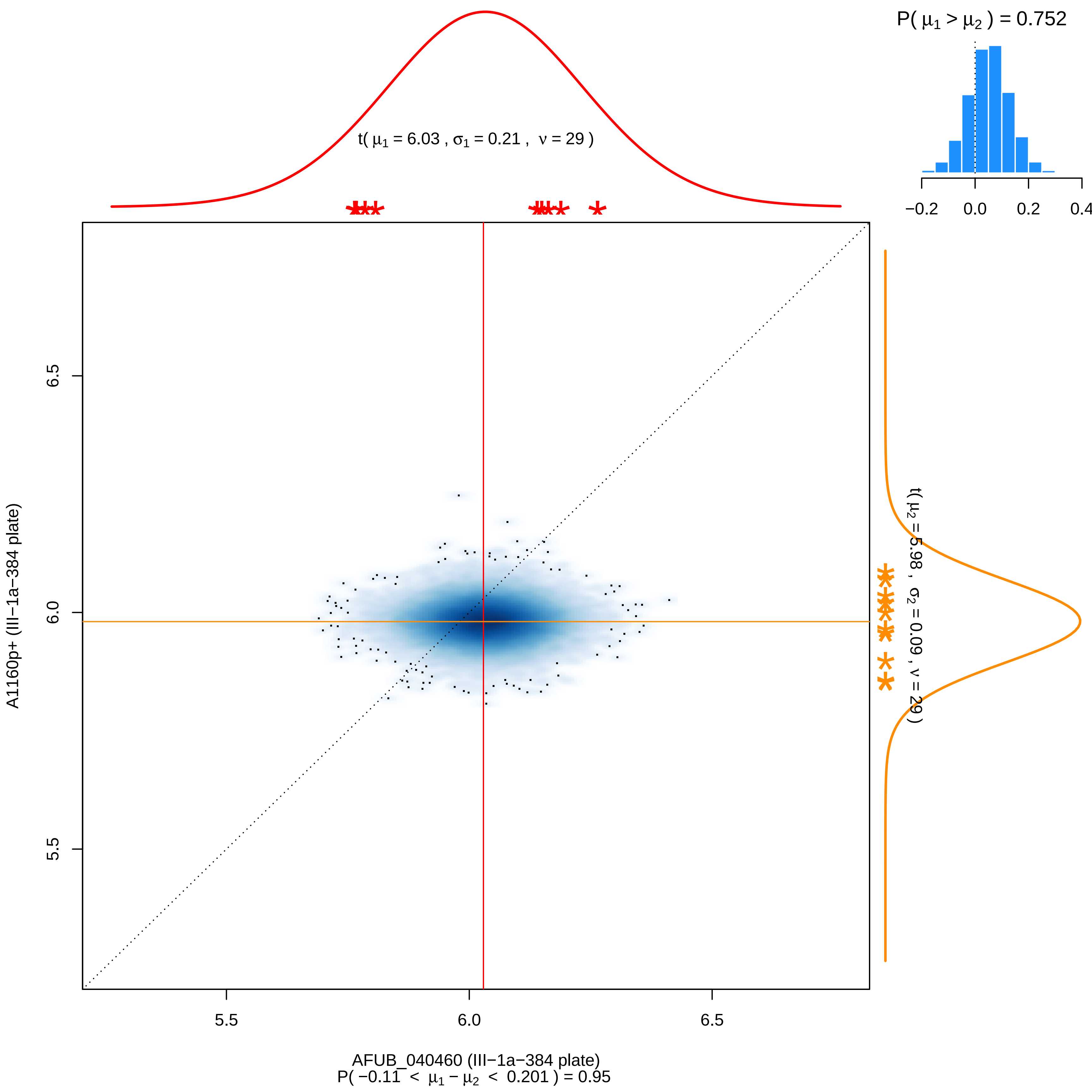
P( μ1 > μ2 ) = 0.752
−0.2
0.0
0.2
0.4
t( μ1 = 6.03 , σ1 = 0.21 , ν = 29 )
6.5
6.0
5.5
A1160p+ (III−1a−384 plate)
t( μ2 = 5.98 , σ2 = 0.09 , ν = 29 )
5.5
6.0
6.5
AFUB_040460 (III−1a−384 plate)
P( −0.11 < μ1 − μ2 < 0.201 ) = 0.95

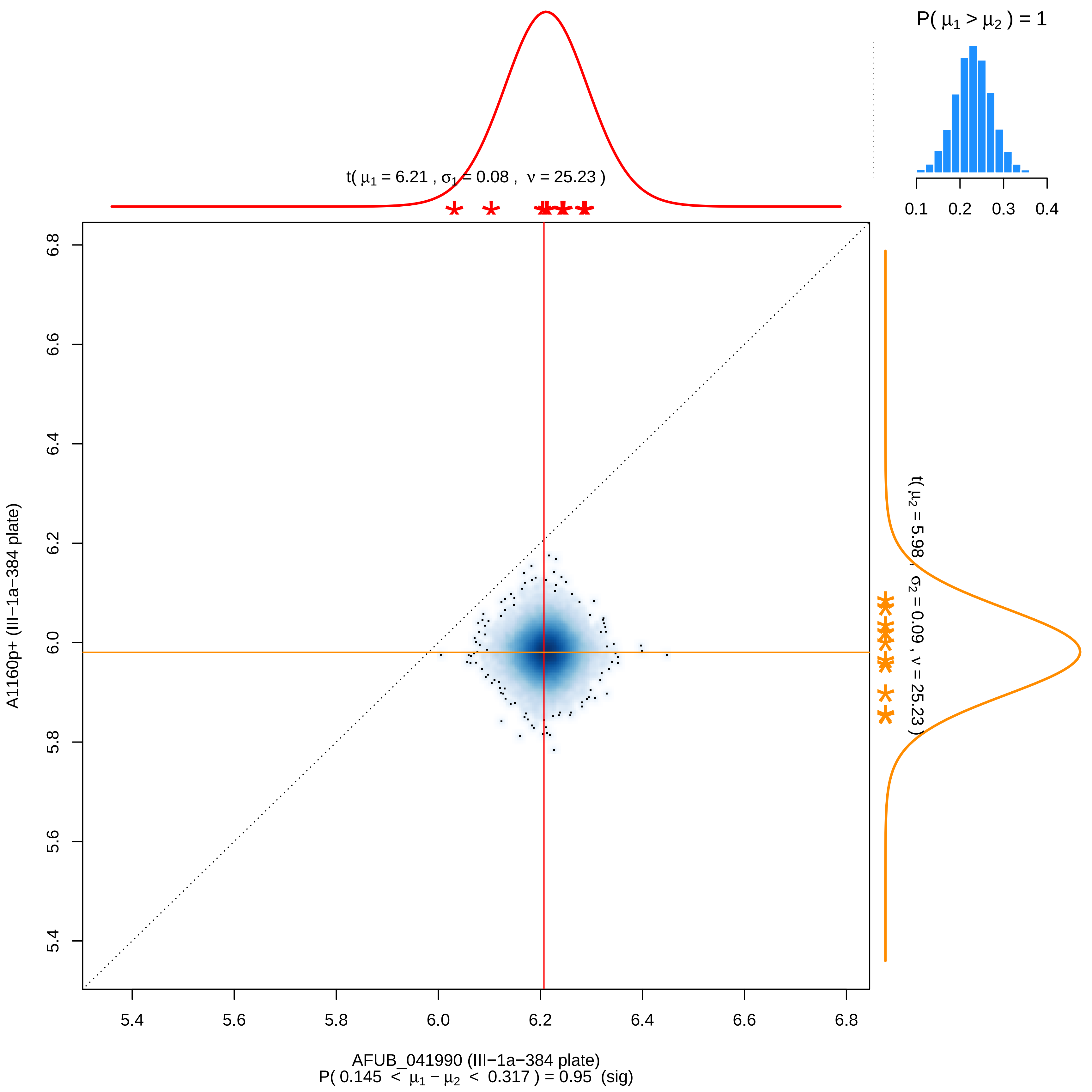
P( μ1 > μ2 ) = 1
0.1 0.2 0.3 0.4
t( μ1 = 6.21 , σ1 = 0.08 , ν = 25.23 )
t( μ2 = 5.98 , σ2 = 0.09 , ν = 25.23 )
A1160p+ (III−1a−384 plate)
6.8
6.6
6.4
6.2
6.0
5.8
5.6
5.4
5.4 5.6 5.8 6.0 6.2 6.4 6.6 6.8
AFUB_041990 (III−1a−384 plate)
P( 0.145 < μ1 − μ2 < 0.317 ) = 0.95 (sig)

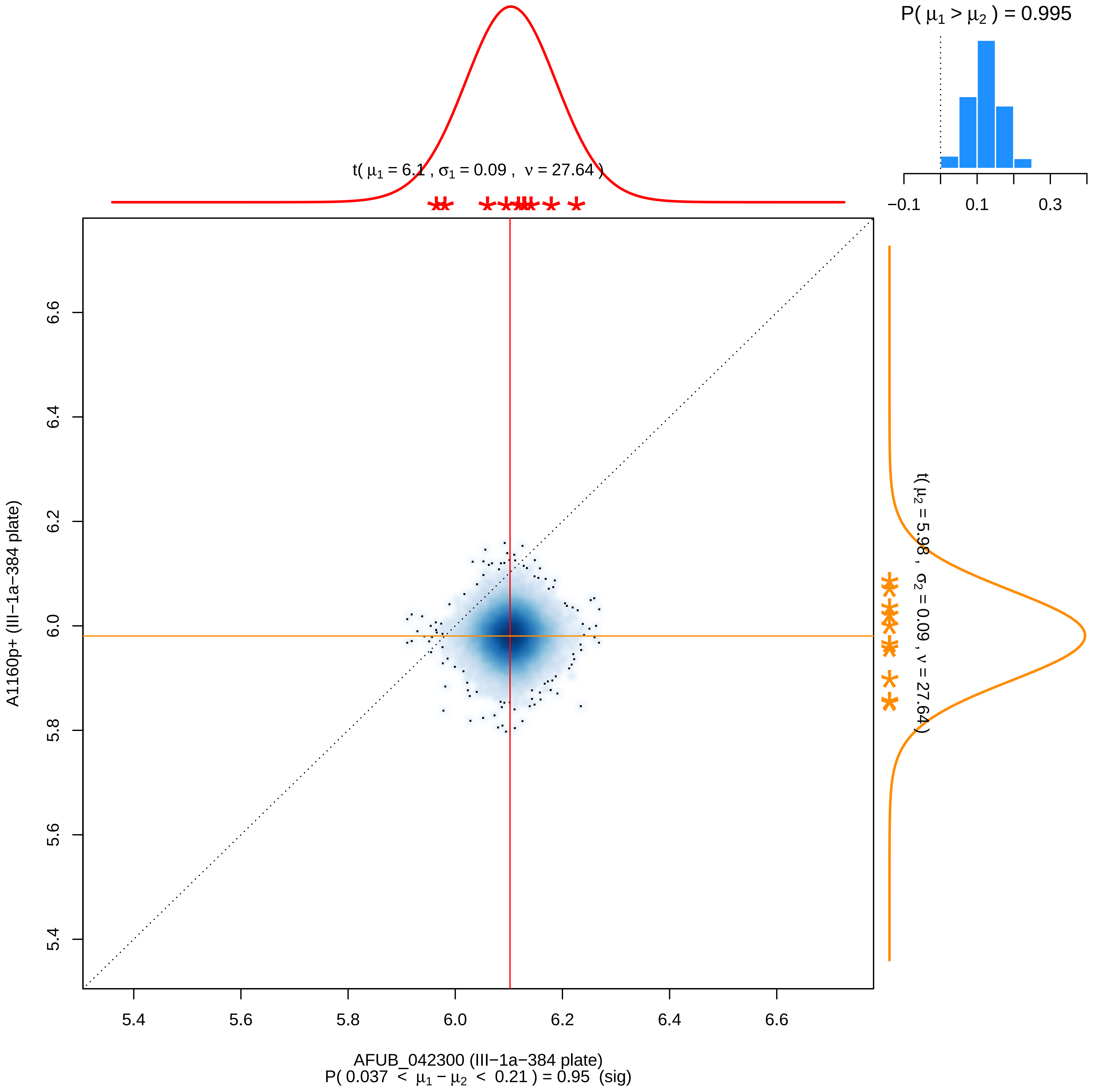

P( μ1 > μ2 ) = 0.995
−0.1
0.1
0.3
t( μ1 = 6.1 , σ1 = 0.09 , ν = 27.64 )
t( μ2 = 5.98 , σ2 = 0.09 , ν = 27.64 )
A1160p+ (III−1a−384 plate)
6.6
6.4
6.2
6.0
5.8
5.6
5.4
5.4
5.6
5.8
6.0
6.2
6.4
6.6
AFUB_042300 (III−1a−384 plate)
P( 0.037 < μ1 − μ2 < 0.21 ) = 0.95 (sig)

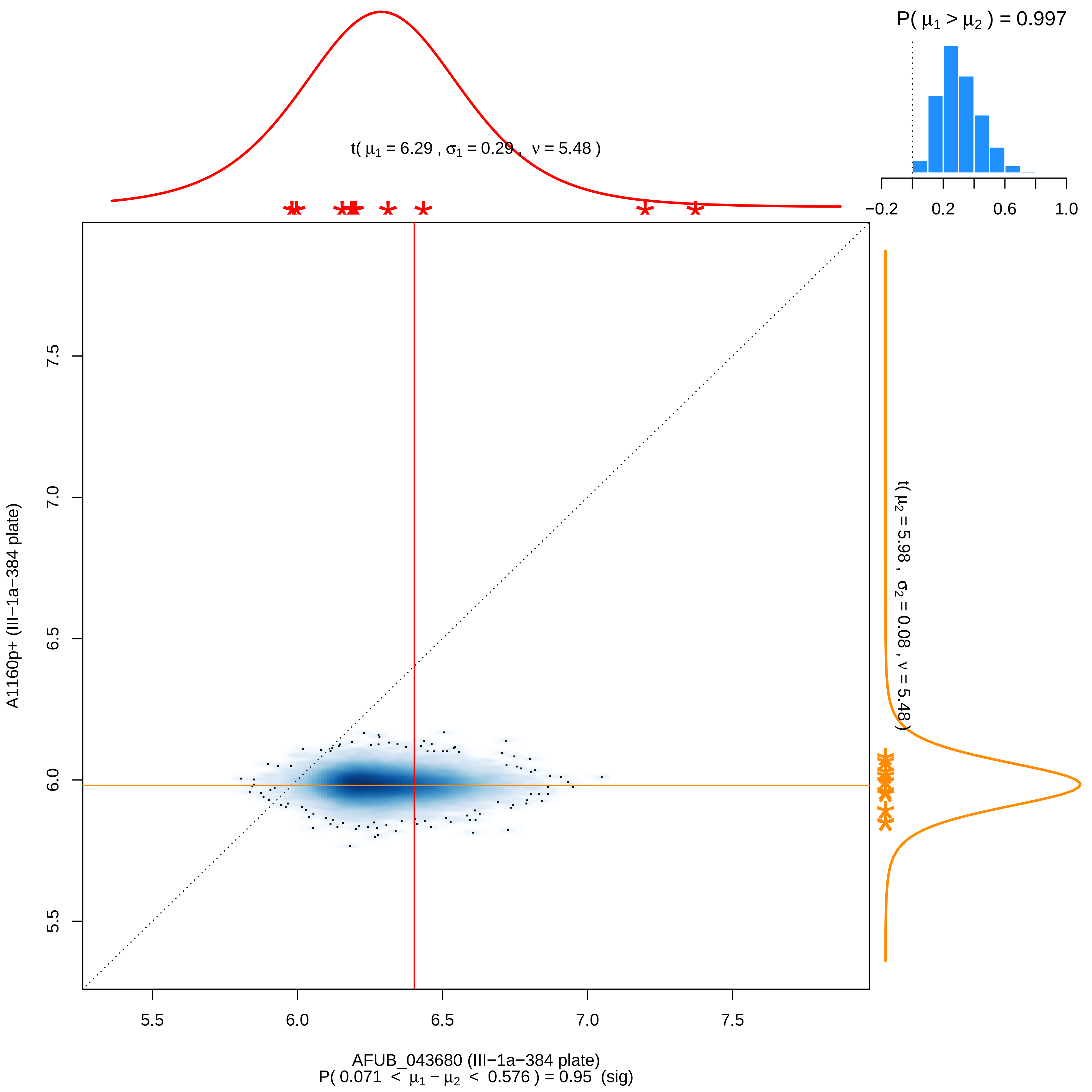
P( μ1 > μ2 ) = 0.997
−0.2 0.2 0.6 1.0
t( μ1 = 6.29 , σ1 = 0.29 , ν = 5.48 )
t( μ2 = 5.98 , σ2 = 0.08 , ν = 5.48 )
A1160p+ (III−1a−384 plate)
5.5 6.0 6.5 7.0 7.5
AFUB_043680 (III−1a−384 plate)
P( 0.071 < μ1 − μ2 < 0.576 ) = 0.95 (sig)

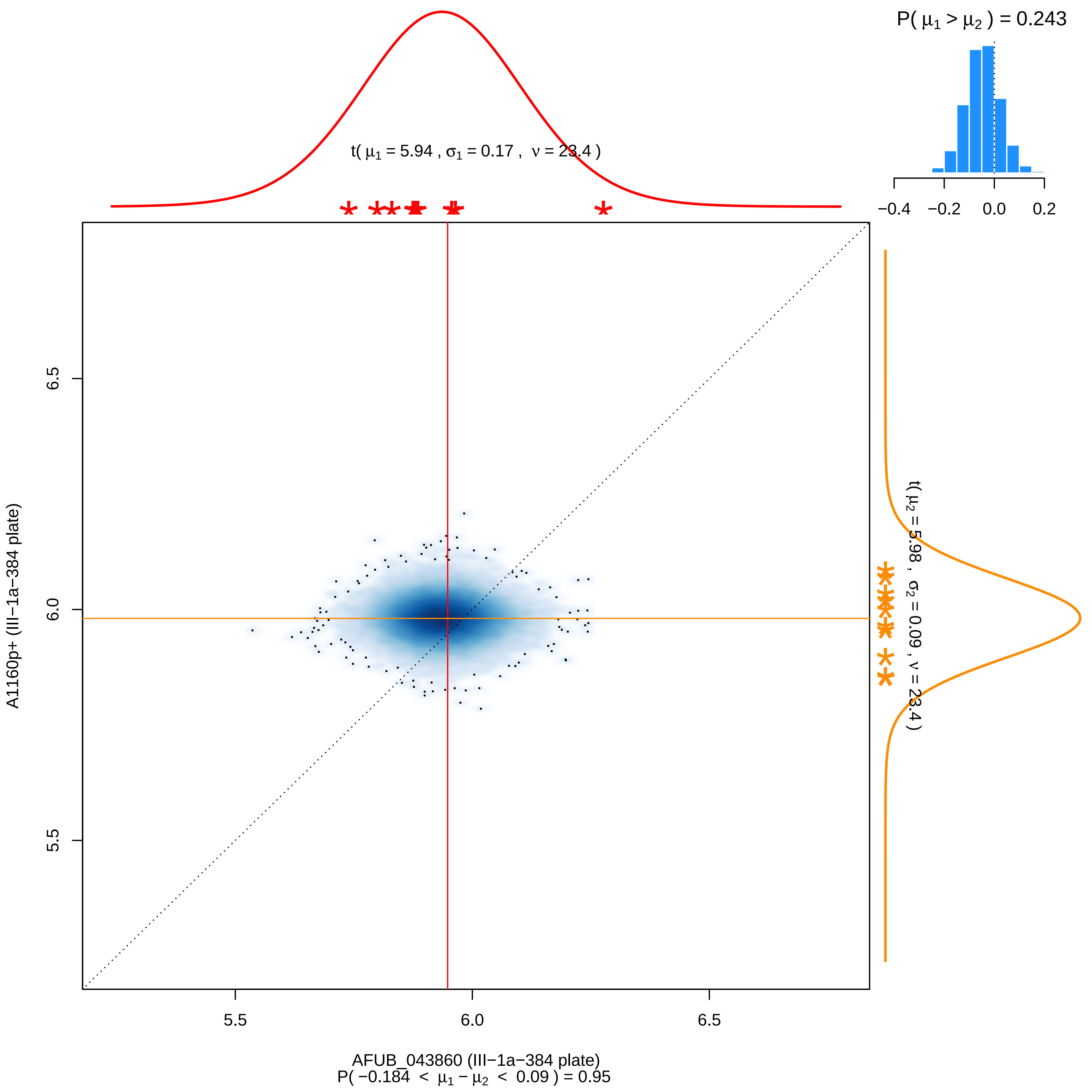

P( μ1 > μ2 ) = 0.243
−0.4 −0.2 0.0 0.2
t( μ1 = 5.94 , σ1 = 0.17 , ν = 23.4 )
t( μ2 = 5.98 , σ2 = 0.09 , ν = 23.4 )
6.5
6.0
5.5
A1160p+ (III−1a−384 plate)
5.5 6.0 6.5
AFUB_043860 (III−1a−384 plate)
P( −0.184 < μ1 − μ2 < 0.09 ) = 0.95

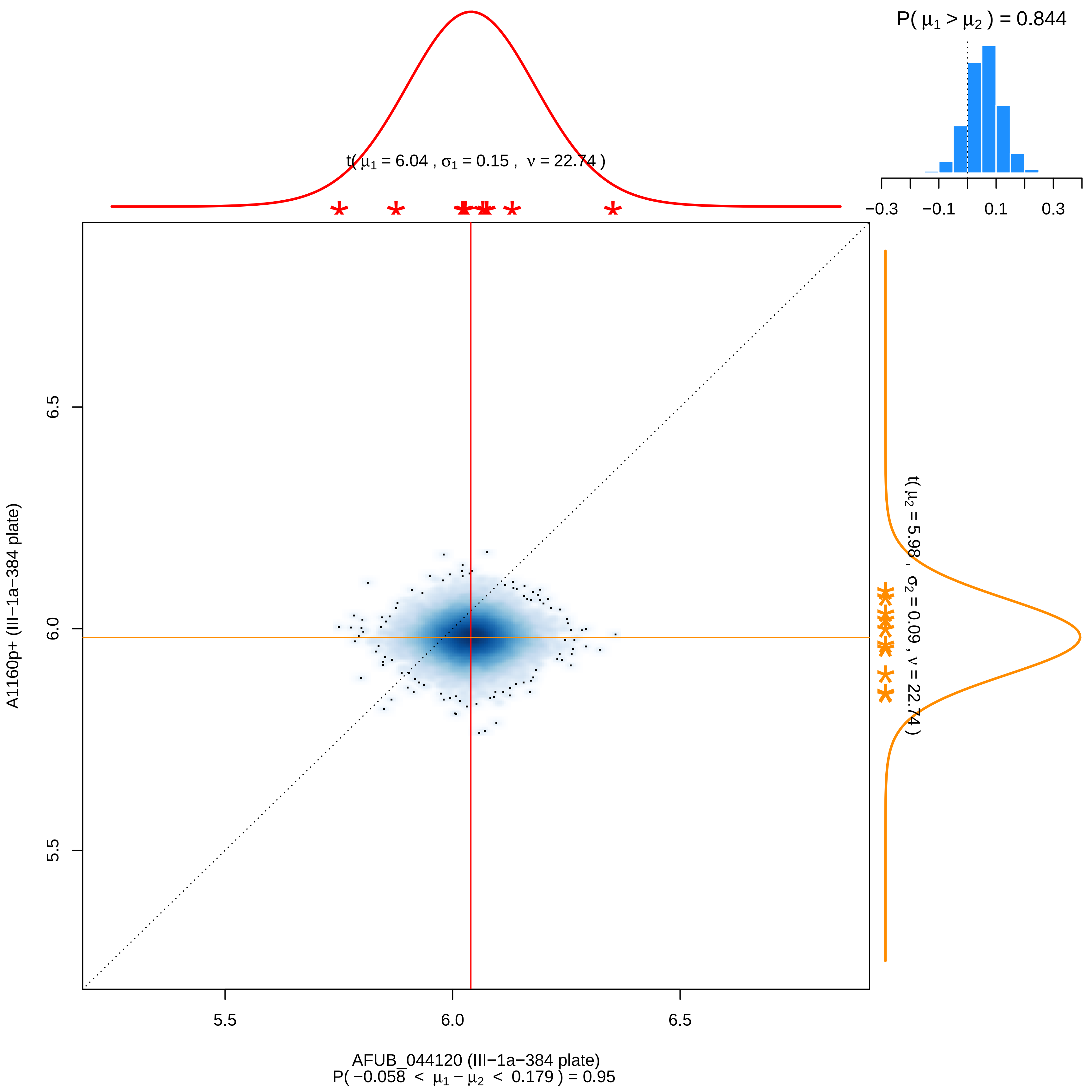

P( μ1 > μ2 ) = 0.844
−0.3
−0.1
0.1
0.3
t( μ1 = 6.04 , σ1 = 0.15 , ν = 22.74 )
t( μ2 = 5.98 , σ2 = 0.09 , ν = 22.74 )
6.5
6.0
5.5
A1160p+ (III−1a−384 plate)
5.5
6.0
6.5
AFUB_044120 (III−1a−384 plate)
P( −0.058 < μ1 − μ2 < 0.179 ) = 0.95

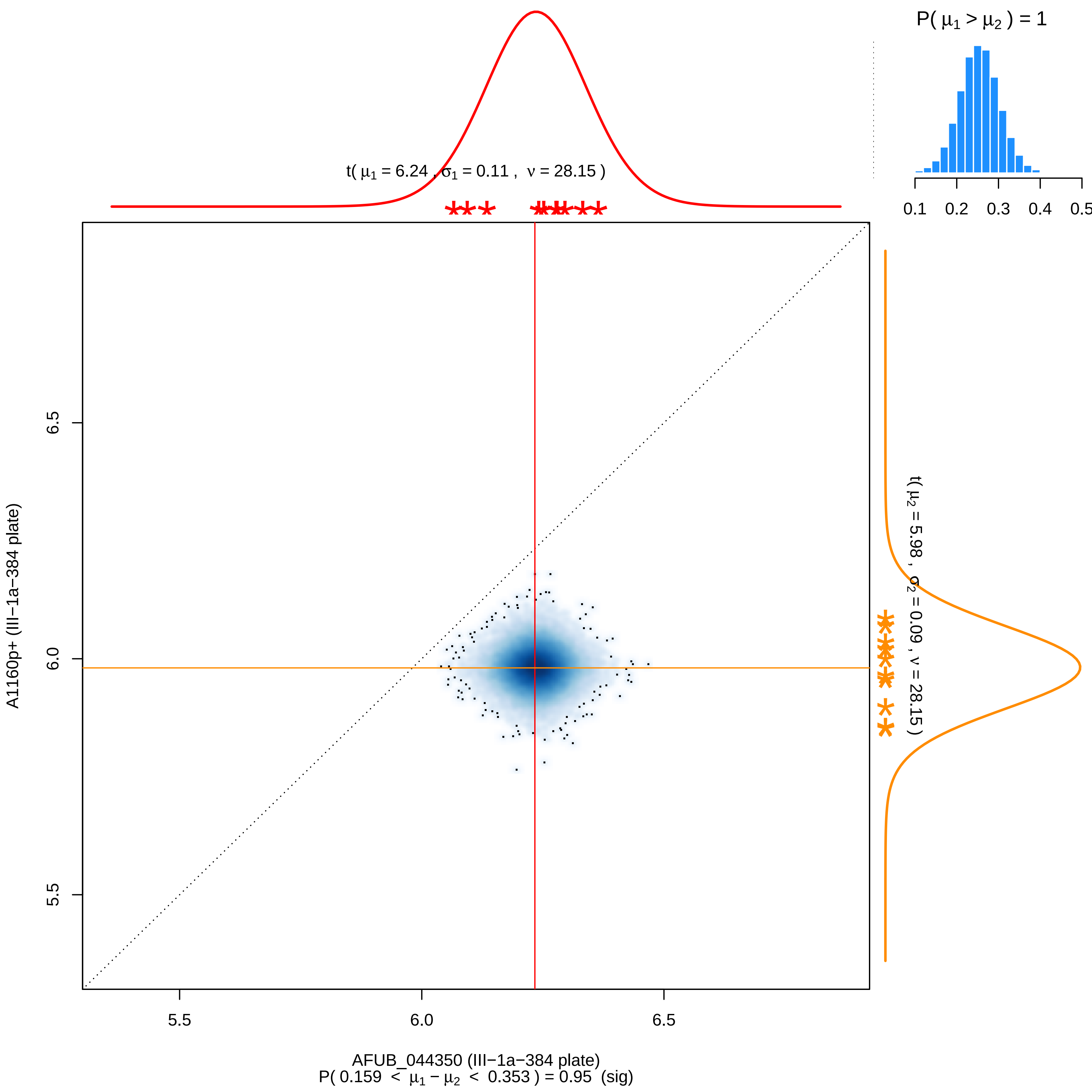

P( μ1 > μ2 ) = 1
0.1
0.2
0.3
0.4
0.5
t( μ1 = 6.24 , σ1 = 0.11 , ν = 28.15 )
t( μ2 = 5.98 , σ2 = 0.09 , ν = 28.15 )
6.5
6.0
5.5
A1160p+ (III−1a−384 plate)
5.5
6.0
6.5
AFUB_044350 (III−1a−384 plate)
P( 0.159 < μ1 − μ2 < 0.353 ) = 0.95 (sig)

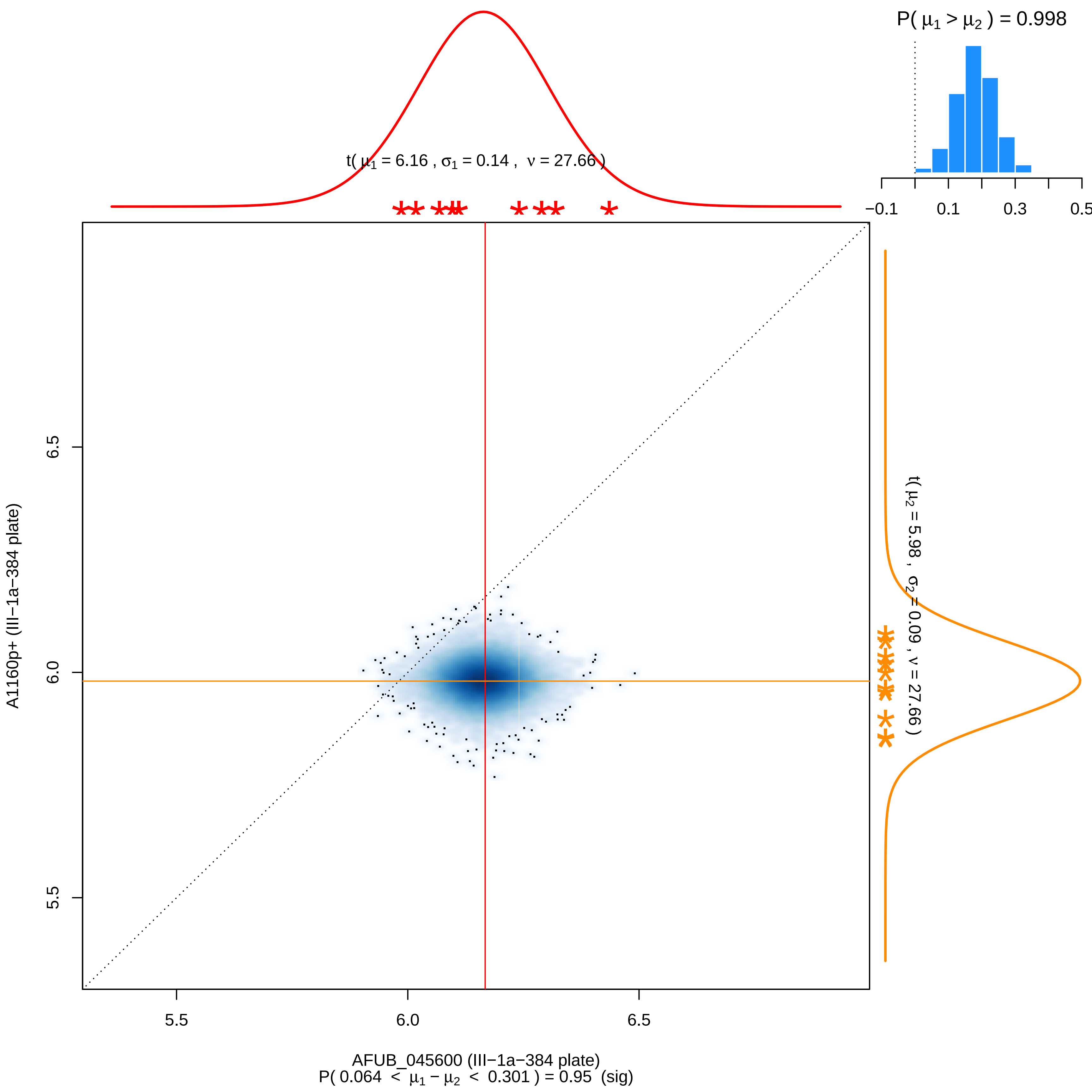

P( μ1 > μ2 ) = 0.998
t( μ1 = 6.16 , σ1 = 0.14 , ν = 27.66 )
−0.1
0.1
0.3
0.5
t( μ2 = 5.98 , σ2 = 0.09 , ν = 27.66 )
A1160p+ (III−1a−384 plate)
6.5
6.0
5.5
5.5
6.0
6.5
AFUB_045600 (III−1a−384 plate)
P( 0.064 < μ1 − μ2 < 0.301 ) = 0.95 (sig)

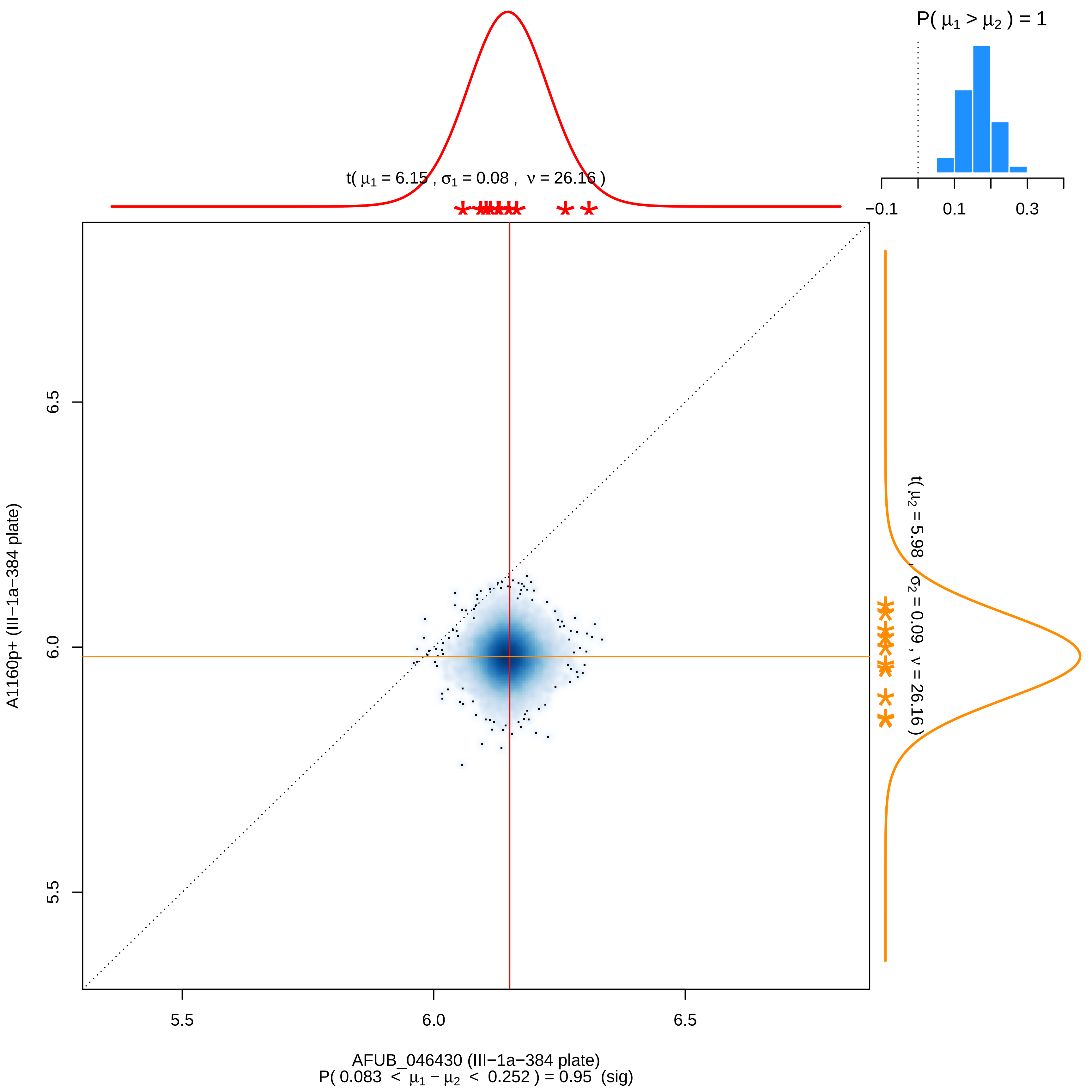

P( μ1 > μ2 ) = 1
−0.1
0.1
0.3
t( μ1 = 6.15 , σ1 = 0.08 , ν = 26.16 )
t( μ2 = 5.98 , σ2 = 0.09 , ν = 26.16 )
6.5
6.0
5.5
A1160p+ (III−1a−384 plate)
5.5
6.0
6.5
AFUB_046430 (III−1a−384 plate)
P( 0.083 < μ1 − μ2 < 0.252 ) = 0.95 (sig)

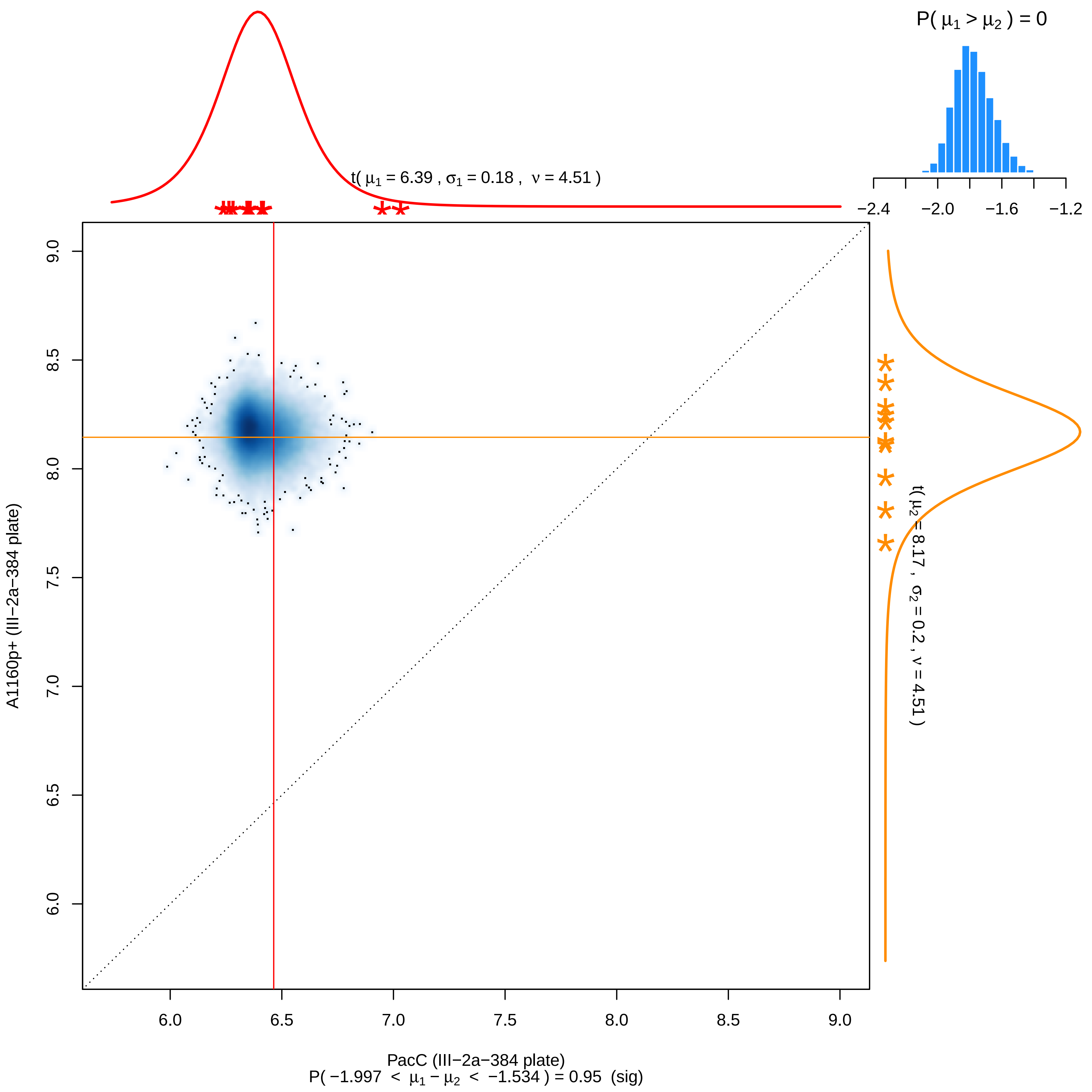

P( μ1 > μ2 ) = 0
−2.4 −2.0 −1.6 −1.2
t( μ1 = 6.39 , σ1 = 0.18 , ν = 4.51 )
t( μ2 = 8.17 , σ2 = 0.2 , ν = 4.51 )
9.0 8.5 8.0 7.5 7.0 6.5 6.0
A1160p+ (III−2a−384 plate)
6.0 6.5 7.0 7.5 8.0 8.5 9.0
PacC (III−2a−384 plate)
P( −1.997 < μ1 − μ2 < −1.534 ) = 0.95 (sig)

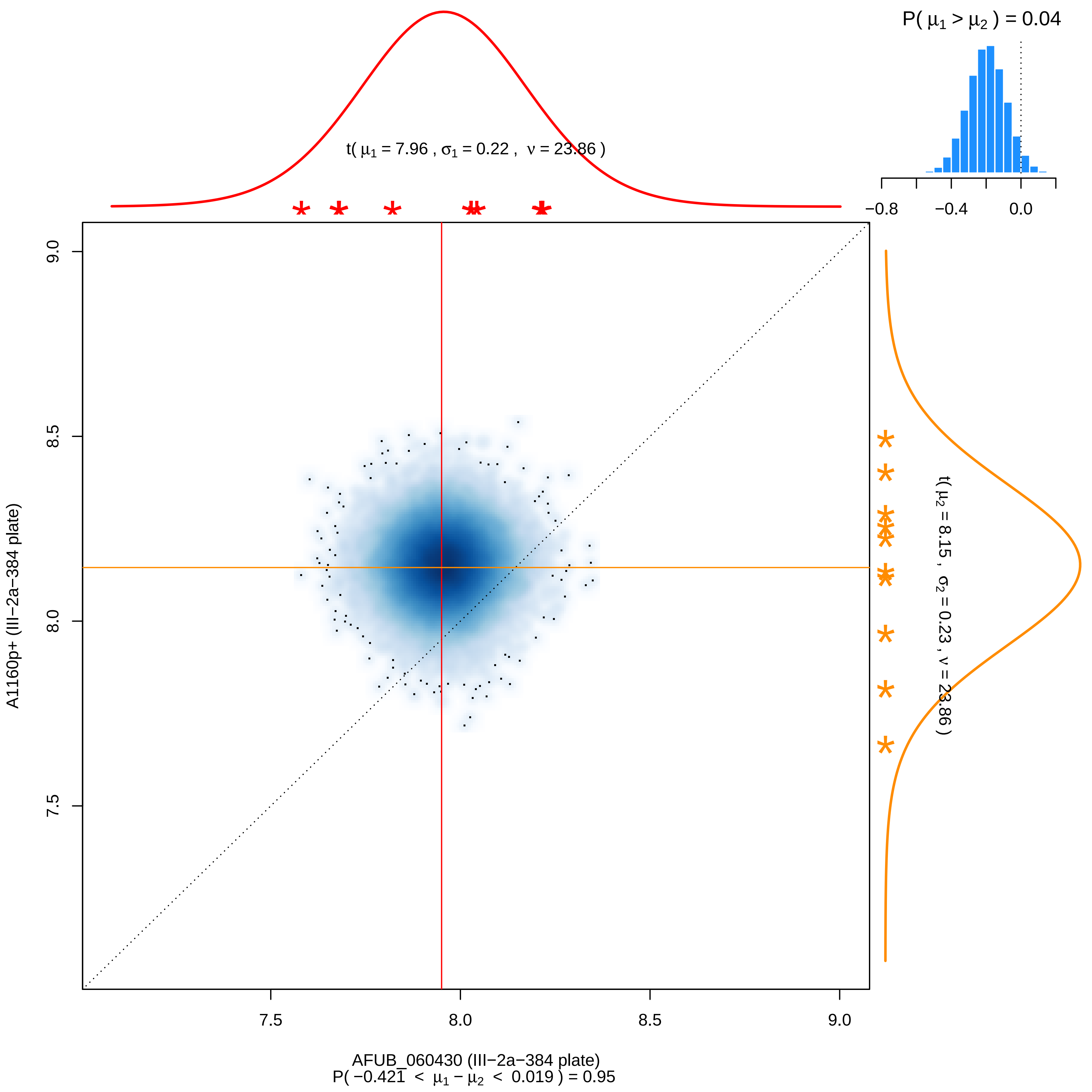

P( μ1 > μ2 ) = 0.04
−0.8
−0.4
0.0
t( μ1 = 7.96 , σ1 = 0.22 , ν = 23.86 )
t( μ2 = 8.15 , σ2 = 0.23 , ν = 23.86 )
9.0
8.5
8.0
7.5
A1160p+ (III−2a−384 plate)
7.5
8.0
8.5
9.0
AFUB_060430 (III−2a−384 plate)
P( −0.421 < μ1 − μ2 < 0.019 ) = 0.95

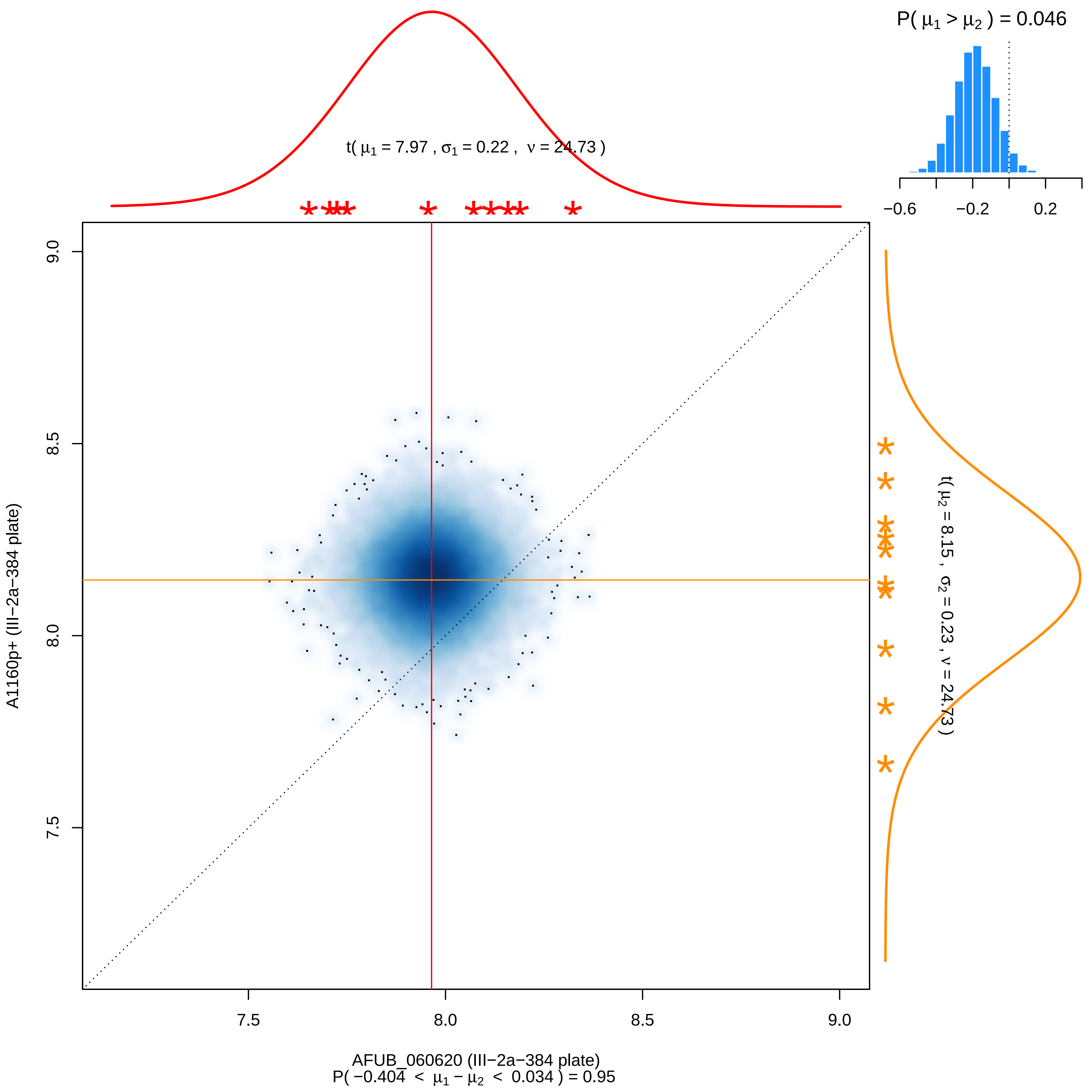

P( μ1 > μ2 ) = 0.046
−0.6
−0.2
0.2
t( μ1 = 7.97 , σ1 = 0.22 , ν = 24.73 )
t( μ2 = 8.15 , σ2 = 0.23 , ν = 24.73 )
A1160p+ (III−2a−384 plate)
9.0
8.5
8.0
7.5
AFUB_060620 (III−2a−384 plate)
P( −0.404 < μ1 − μ2 < 0.034 ) = 0.95

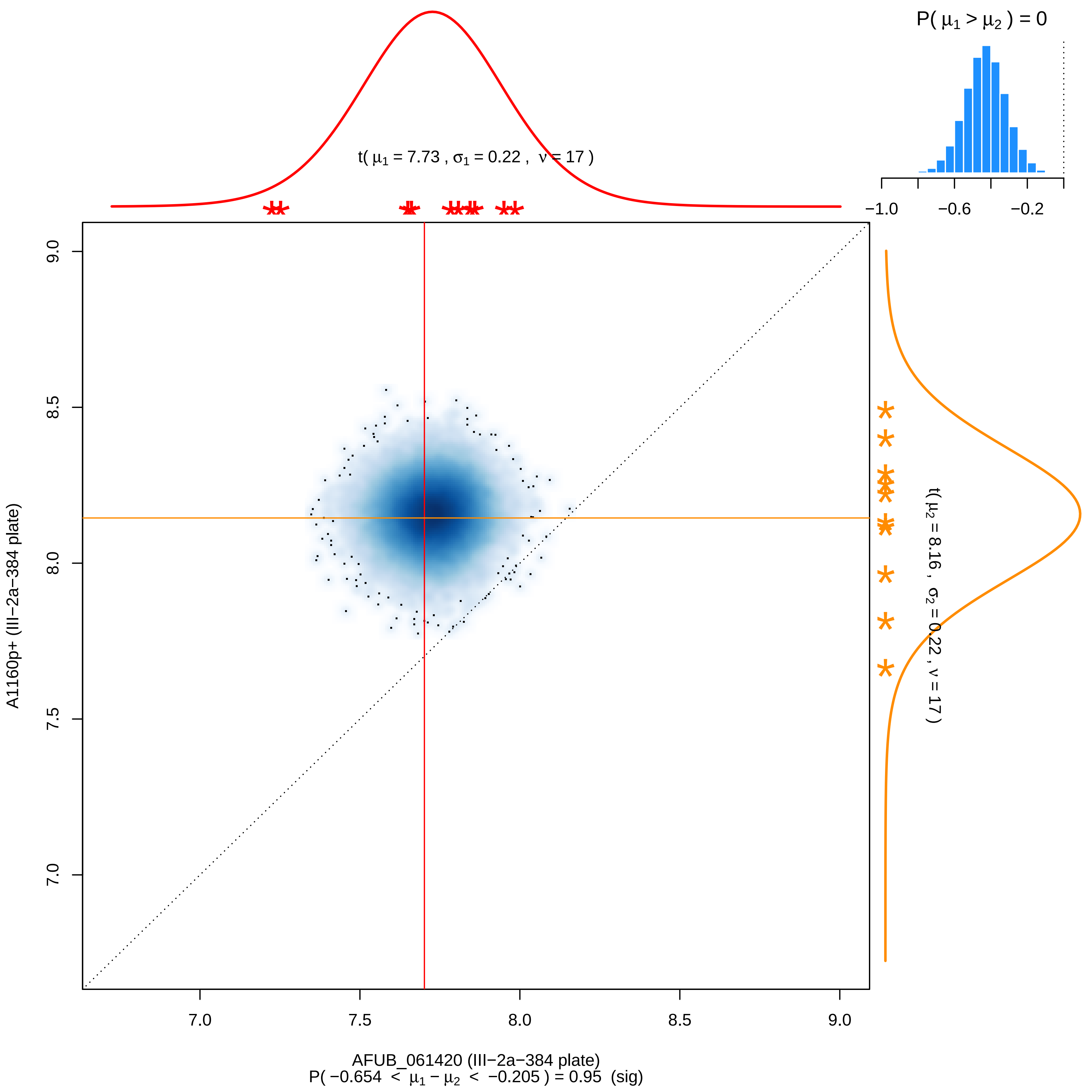

P( μ1 > μ2 ) = 0
−1.0
−0.6
−0.2
t( μ1 = 7.73 , σ1 = 0.22 , ν = 17 )
t( μ2 = 8.16 , σ2 = 0.22 , ν = 17 )
A1160p+ (III−2a−384 plate)
9.0
8.5
8.0
7.5
7.0
AFUB_061420 (III−2a−384 plate)
P( −0.654 < μ1 − μ2 < −0.205 ) = 0.95 (sig)

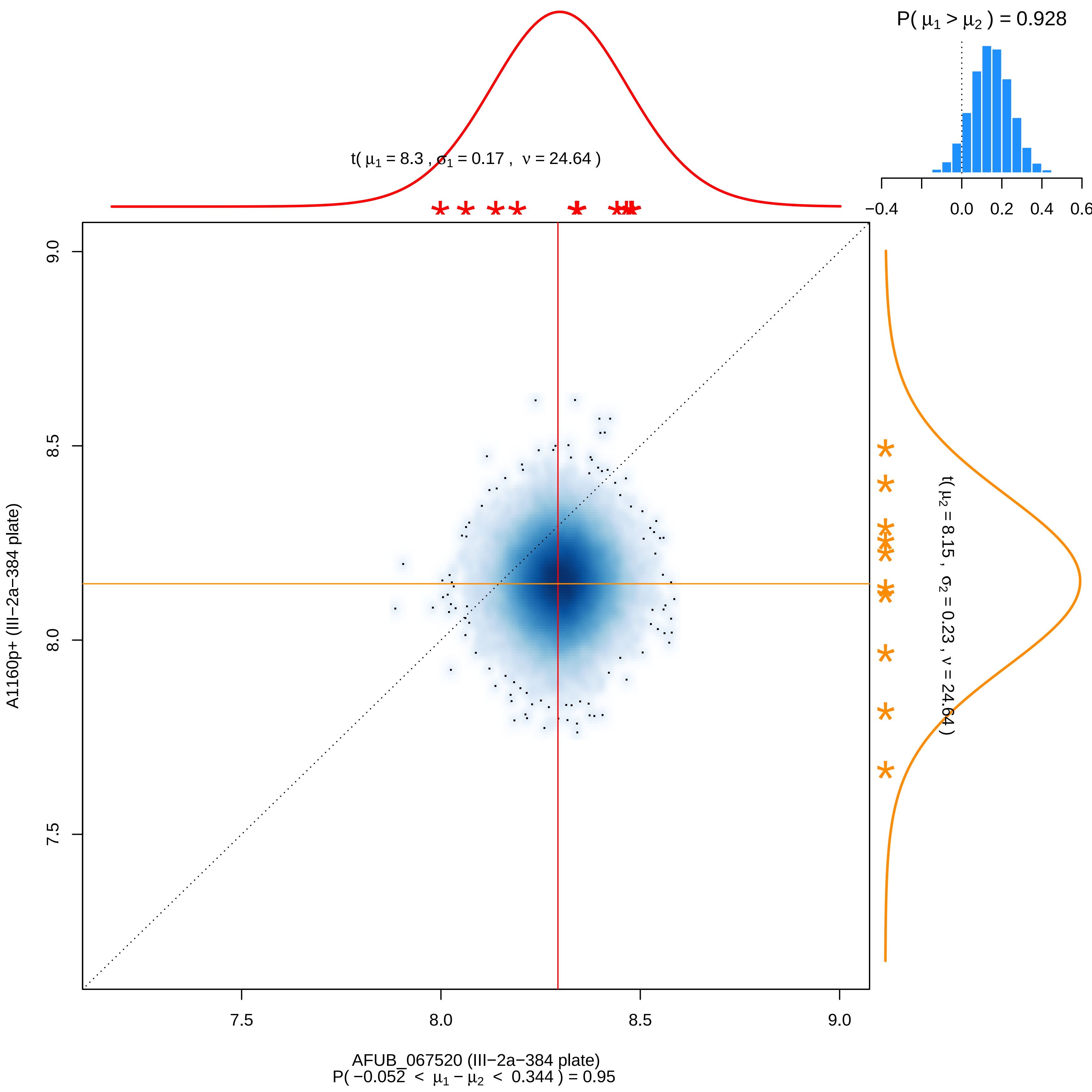

P( μ1 > μ2 ) = 0.928
−0.4
0.0
0.2
0.4
0.6
t( μ1 = 8.3 , σ1 = 0.17 , ν = 24.64 )
9.0
8.5
8.0
7.5
A1160p+ (III−2a−384 plate)
t( μ2 = 8.15 , σ2 = 0.23 , ν = 24.64 )
7.5
8.0
8.5
9.0
AFUB_067520 (III−2a−384 plate)
P( −0.052 < μ1 − μ2 < 0.344 ) = 0.95

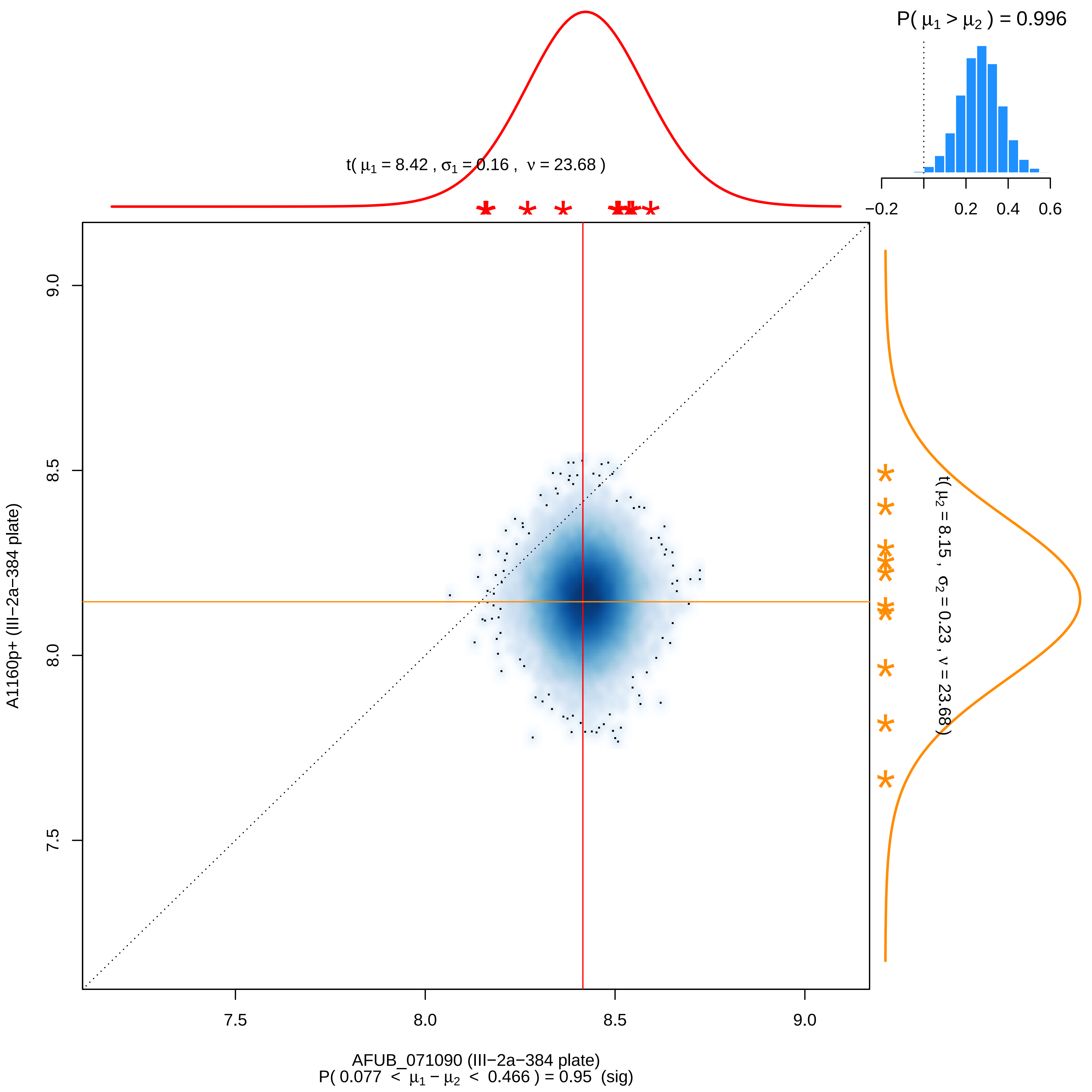

P( μ1 > μ2 ) = 0.996
−0.2
0.2
0.4
0.6
t( μ1 = 8.42 , σ1 = 0.16 , ν = 23.68 )
t( μ2 = 8.15 , σ2 = 0.23 , ν = 23.68 )
A1160p+ (III−2a−384 plate)
9.0
8.5
8.0
7.5
7.5
8.0
8.5
9.0
AFUB_071090 (III−2a−384 plate)
P( 0.077 < μ1 − μ2 < 0.466 ) = 0.95 (sig)

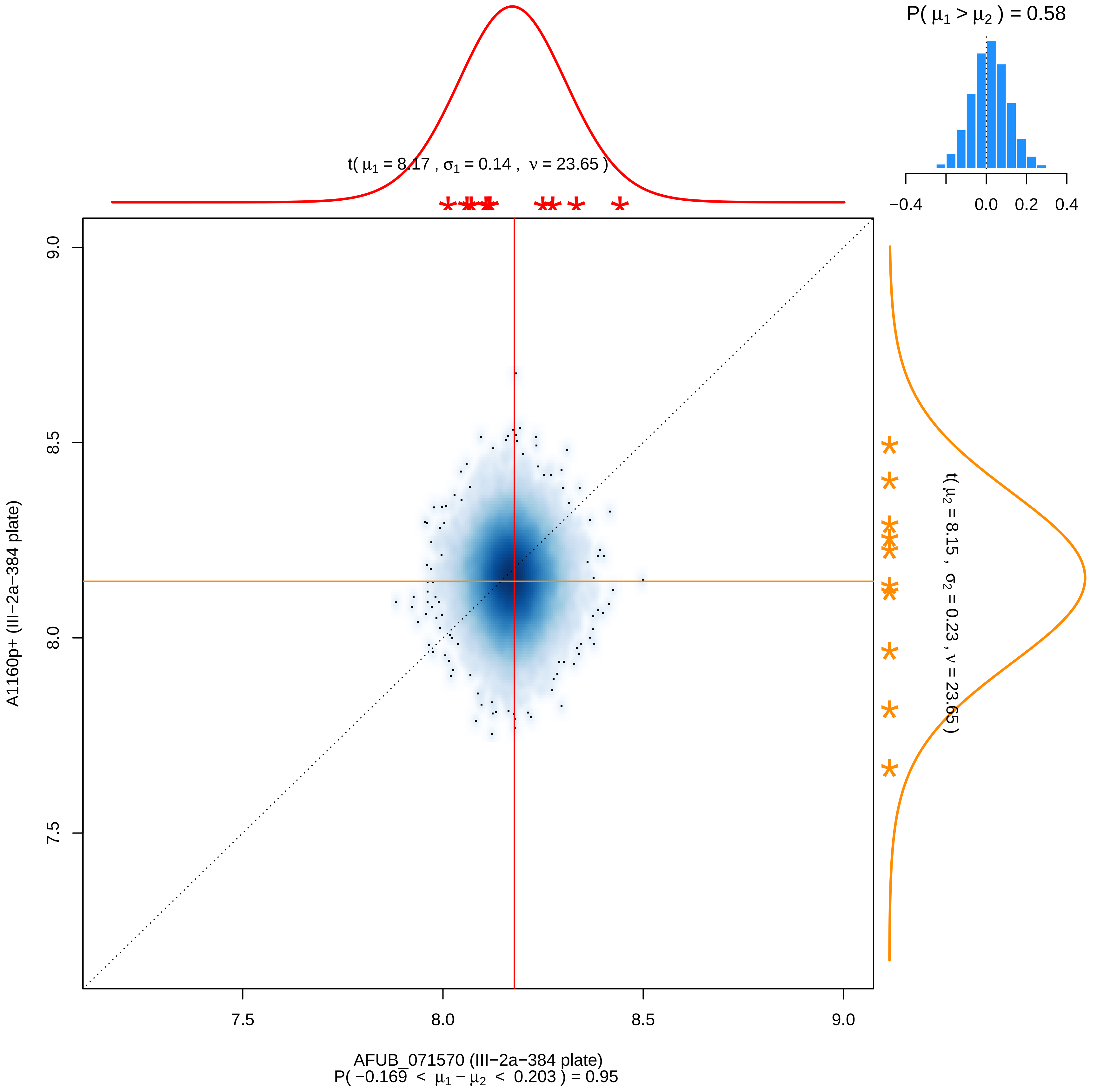

P( μ1 > μ2 ) = 0.58
−0.4
0.0
0.2
0.4
t( μ1 = 8.17 , σ1 = 0.14 , ν = 23.65 )
t( μ2 = 8.15 , σ2 = 0.23 , ν = 23.65 )
9.0
8.5
8.0
7.5
A1160p+ (III−2a−384 plate)
7.5
8.0
8.5
9.0
AFUB_071570 (III−2a−384 plate)
P( −0.169 < μ1 − μ2 < 0.203 ) = 0.95

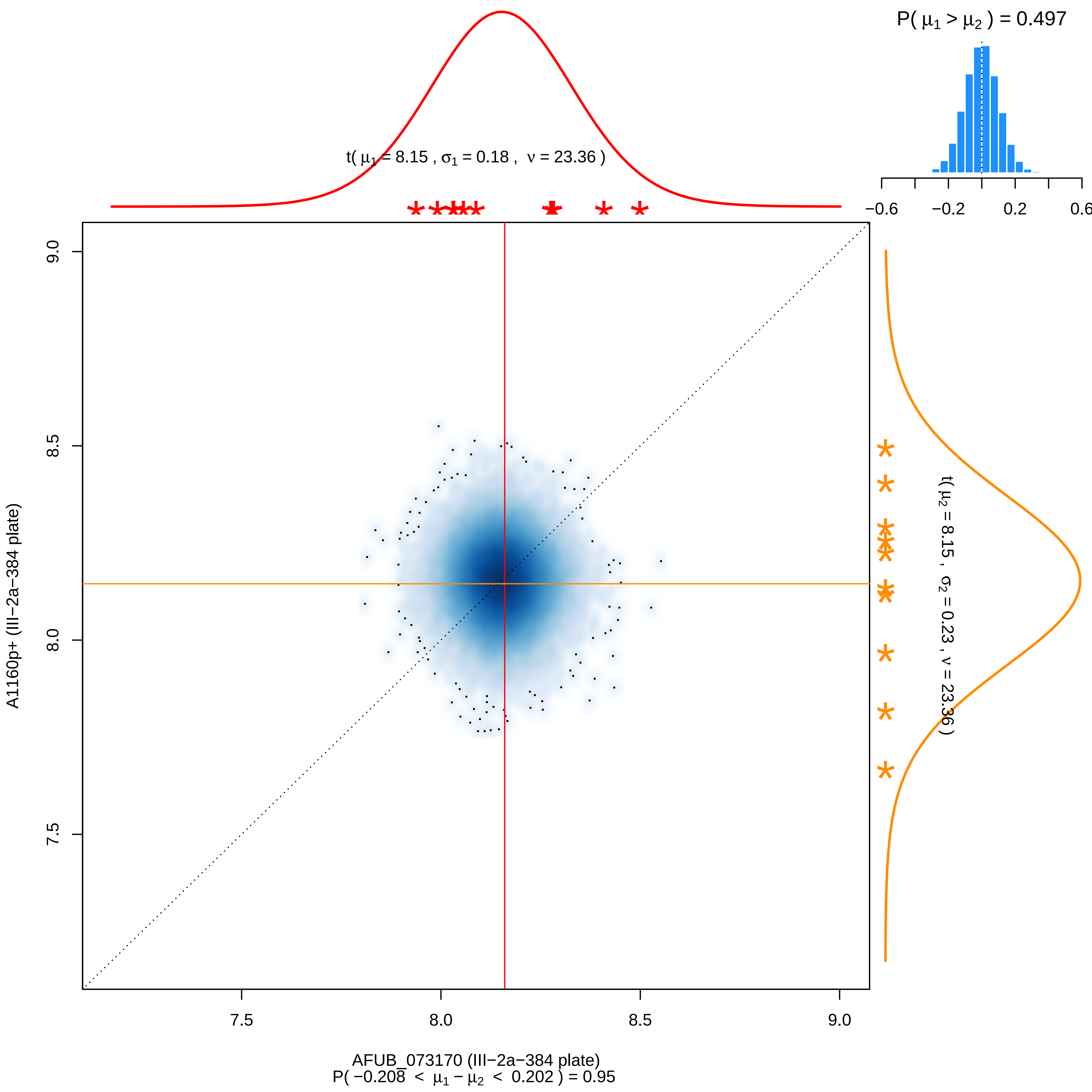

P( μ1 > μ2 ) = 0.497
−0.6
−0.2
0.2
0.6
t( μ1 = 8.15 , σ1 = 0.18 , ν = 23.36 )
t( μ2 = 8.15 , σ2 = 0.23 , ν = 23.36 )
9.0
8.5
8.0
7.5
A1160p+ (III−2a−384 plate)
AFUB_073170 (III−2a−384 plate)
P( −0.208 < μ1 − μ2 < 0.202 ) = 0.95

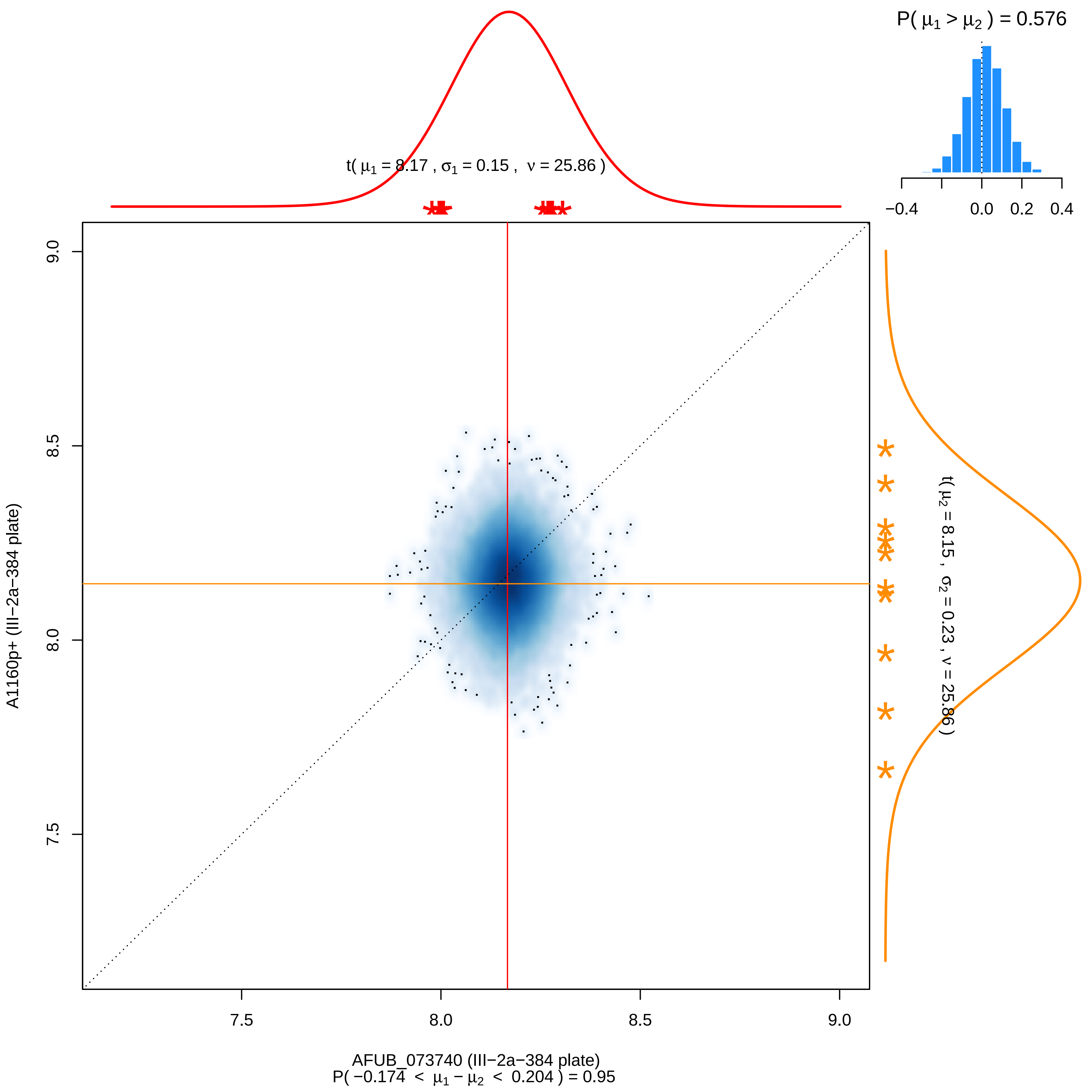

P( μ1 > μ2 ) = 0.576
−0.4
0.0
0.2
0.4
t( μ1 = 8.17 , σ1 = 0.15 , ν = 25.86 )
t( μ2 = 8.15 , σ2 = 0.23 , ν = 25.86 )
9.0
8.5
8.0
7.5
A1160p+ (III−2a−384 plate)
7.5
8.0
8.5
9.0
AFUB_073740 (III−2a−384 plate)
P( −0.174 < μ1 − μ2 < 0.204 ) = 0.95

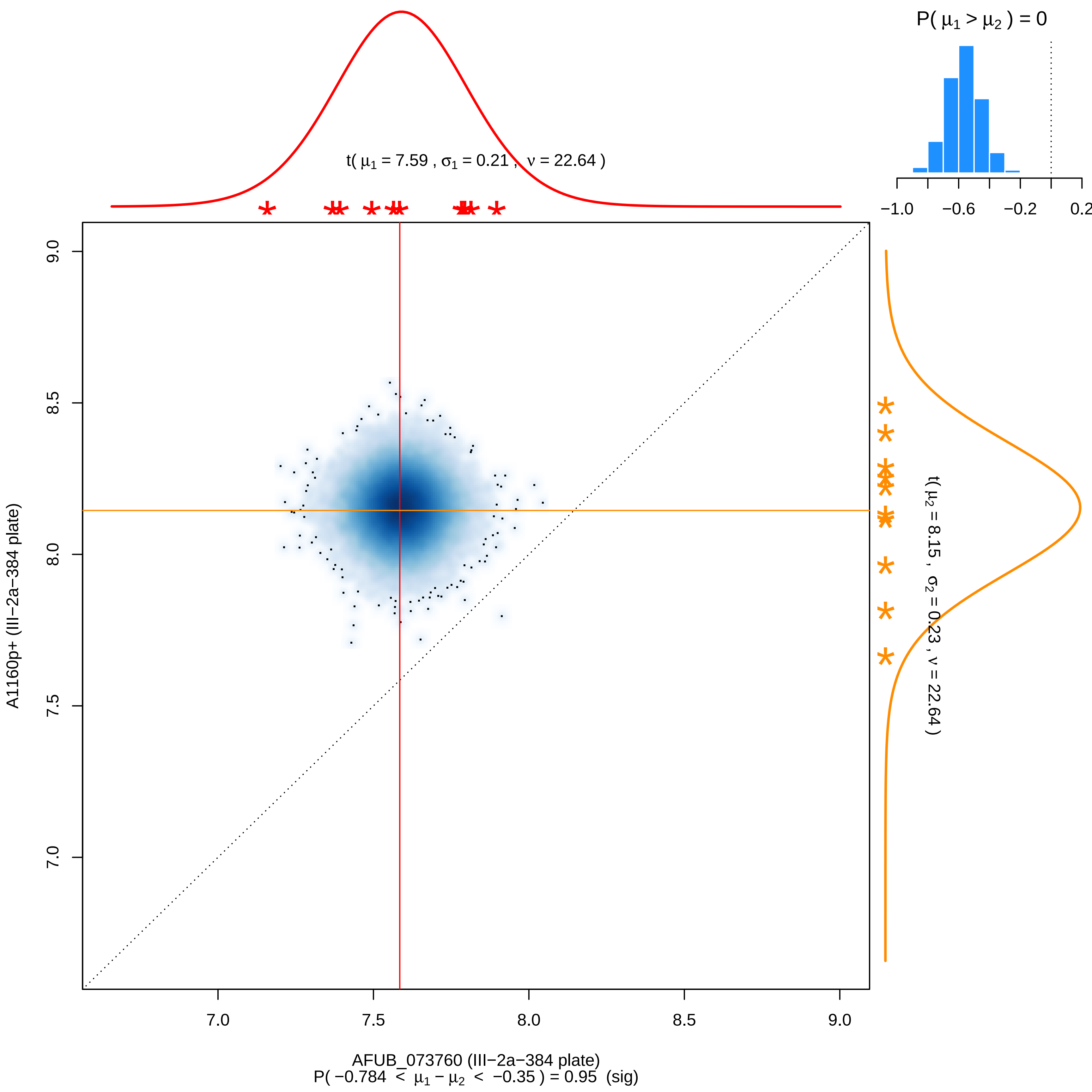

P( $\mu_1 > \mu_2$ ) = 0
−1.0
−0.6
−0.2
0.2
t( $\mu_1$ = 7.59 , $\sigma_1$ = 0.21 , $\nu$ = 22.64 )
9.0
8.5
8.0
7.5
7.0
A1160p+ (III−2a−384 plate)
t( $\mu_2$ = 8.15 , $\sigma_2$ = 0.23 , $\nu$ = 22.64 )
7.0
7.5
8.0
8.5
9.0
AFUB_073760 (III−2a−384 plate)
P( −0.784 < $\mu_1 - \mu_2$ < −0.35 ) = 0.95 (sig)

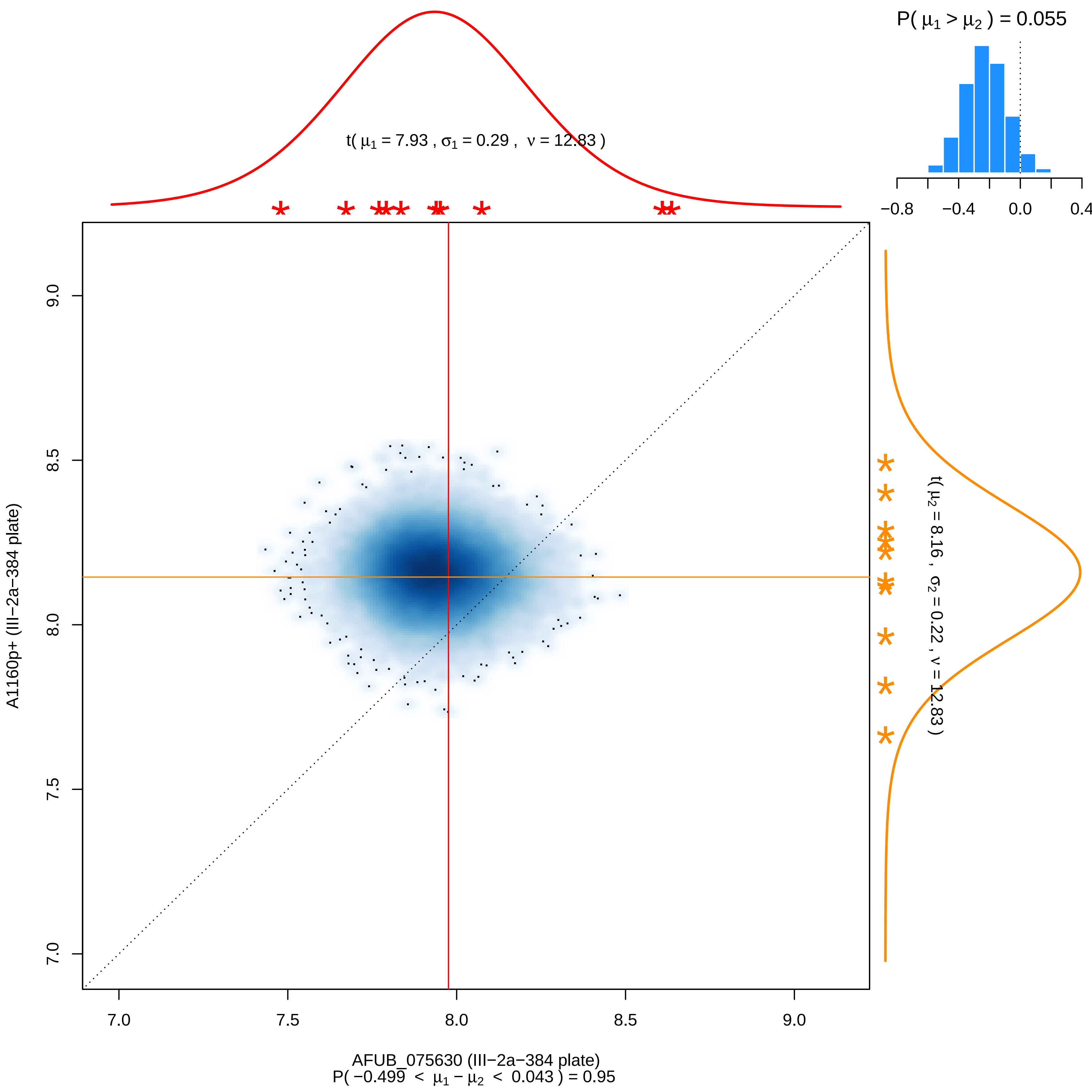

P( μ1 > μ2 ) = 0.055
−0.8
−0.4
0.0
0.4
t( μ1 = 7.93 , σ1 = 0.29 , ν = 12.83 )
t( μ2 = 8.16 , σ2 = 0.22 , ν = 12.83 )
9.0
8.5
8.0
7.5
7.0
A1160p+ (III−2a−384 plate)
7.0
7.5
8.0
8.5
9.0
AFUB_075630 (III−2a−384 plate)
P( −0.499 < μ1 − μ2 < 0.043 ) = 0.95

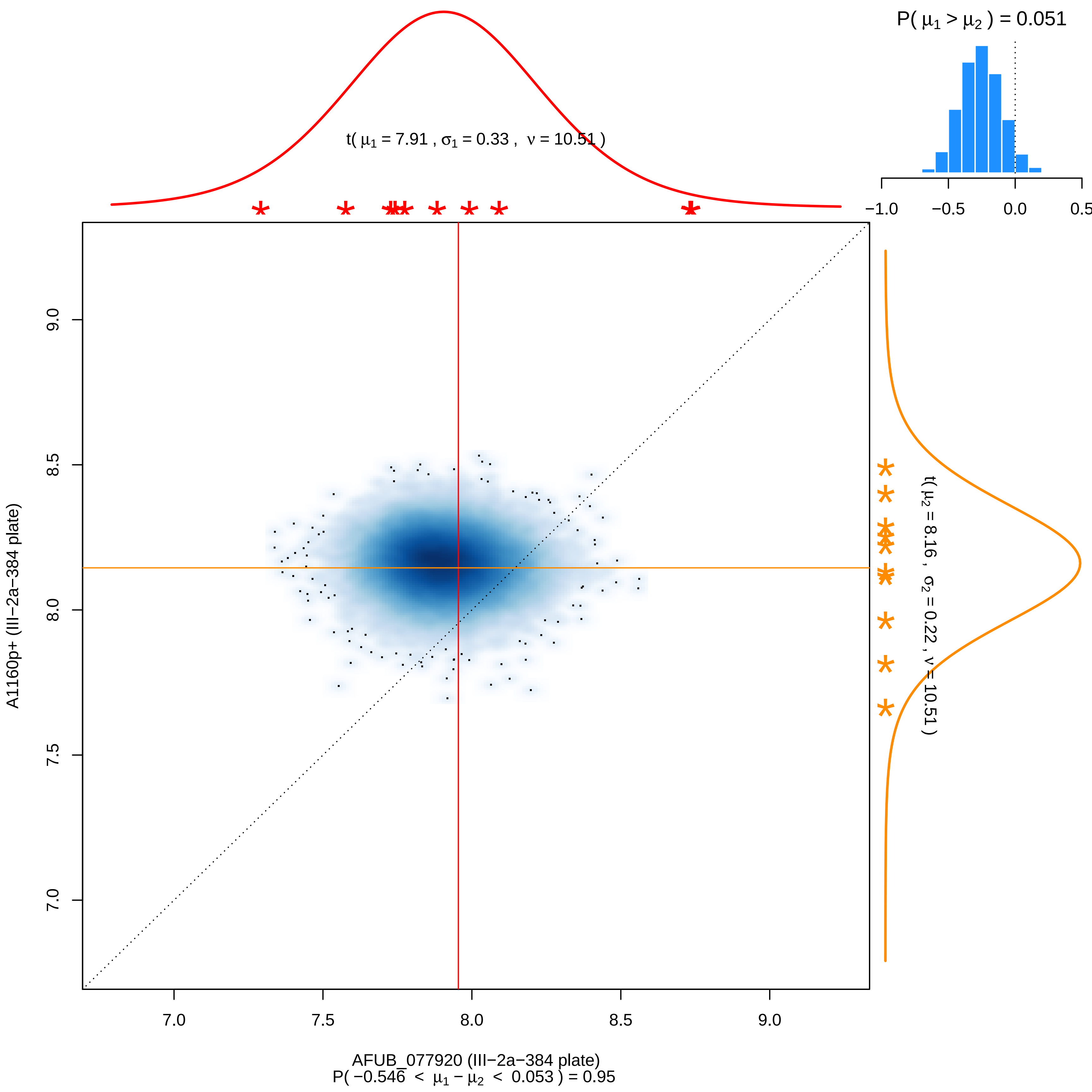

P( μ1 > μ2 ) = 0.051
t( μ1 = 7.91 , σ1 = 0.33 , ν = 10.51 )
−1.0
−0.5
0.0
0.5
9.0
8.5
8.0
7.5
7.0
A1160p+ (III−2a−384 plate)
t( μ2 = 8.16 , σ2 = 0.22 , ν = 10.51 )
7.0
7.5
8.0
8.5
9.0
AFUB_077920 (III−2a−384 plate)
P( −0.546 < μ1 − μ2 < 0.053 ) = 0.95

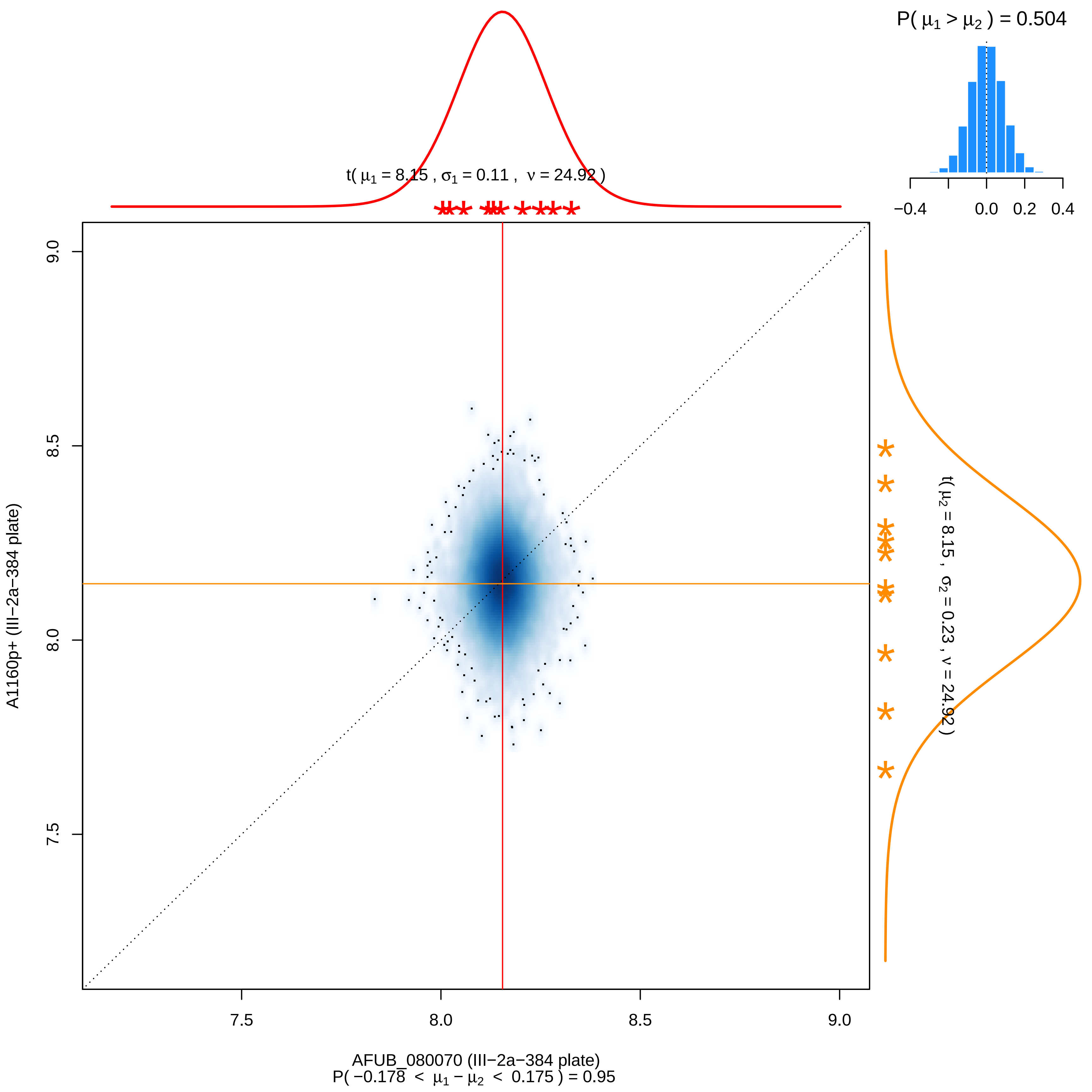

P( μ1 > μ2 ) = 0.504
−0.4
0.0
0.2
0.4
t( μ1 = 8.15 , σ1 = 0.11 , ν = 24.92 )
t( μ2 = 8.15 , σ2 = 0.23 , ν = 24.92 )
9.0
8.5
8.0
7.5
A1160p+ (III−2a−384 plate)
AFUB_080070 (III−2a−384 plate)
P( −0.178 < μ1 − μ2 < 0.175 ) = 0.95

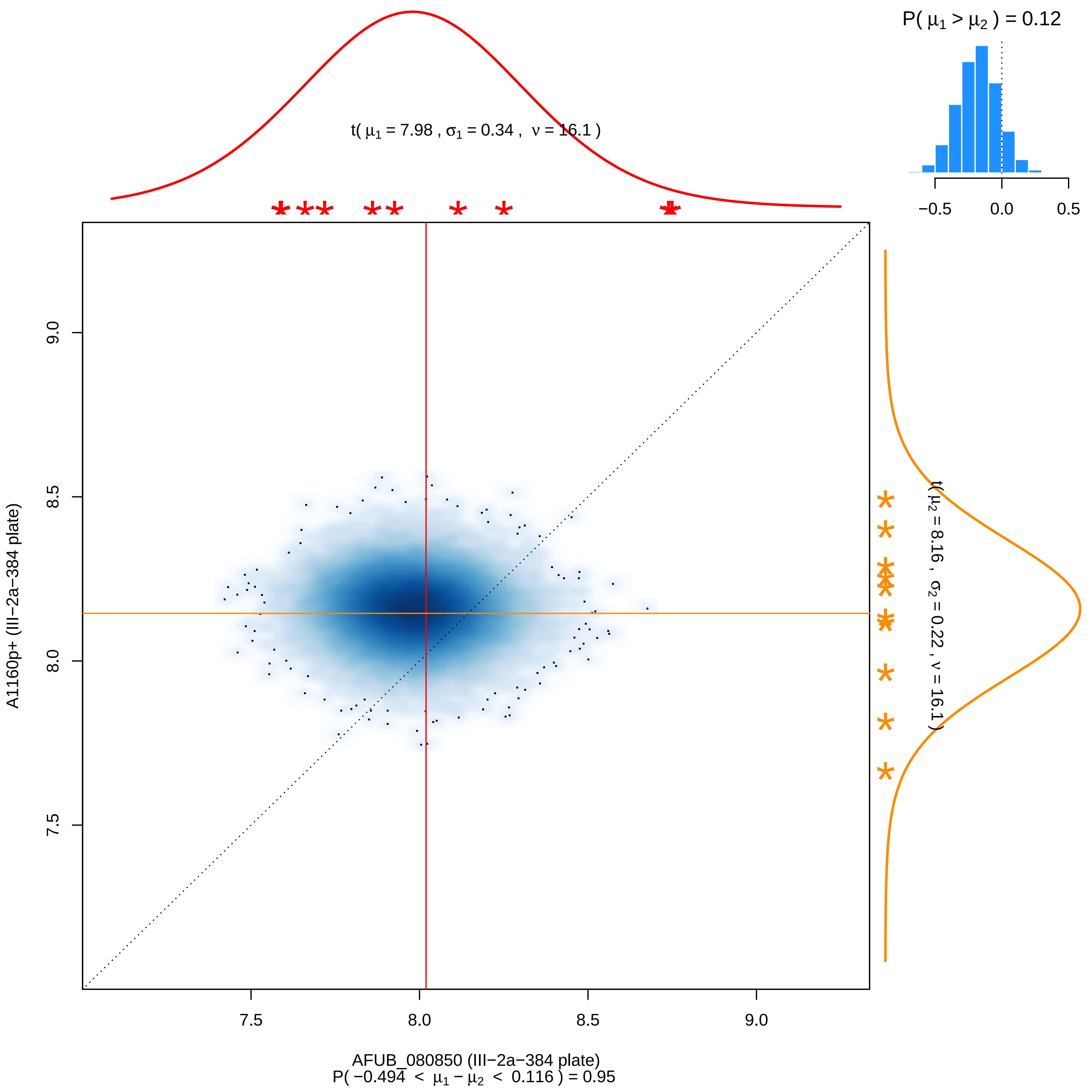

P( μ1 > μ2 ) = 0.12
−0.5
0.0
0.5
t( μ1 = 7.98 , σ1 = 0.34 , ν = 16.1 )
t( μ2 = 8.16 , σ2 = 0.22 , ν = 16.1 )
9.0
8.5
8.0
7.5
A1160p+ (III−2a−384 plate)
7.5
8.0
8.5
9.0
AFUB_080850 (III−2a−384 plate)
P( −0.494 < μ1 − μ2 < 0.116 ) = 0.95

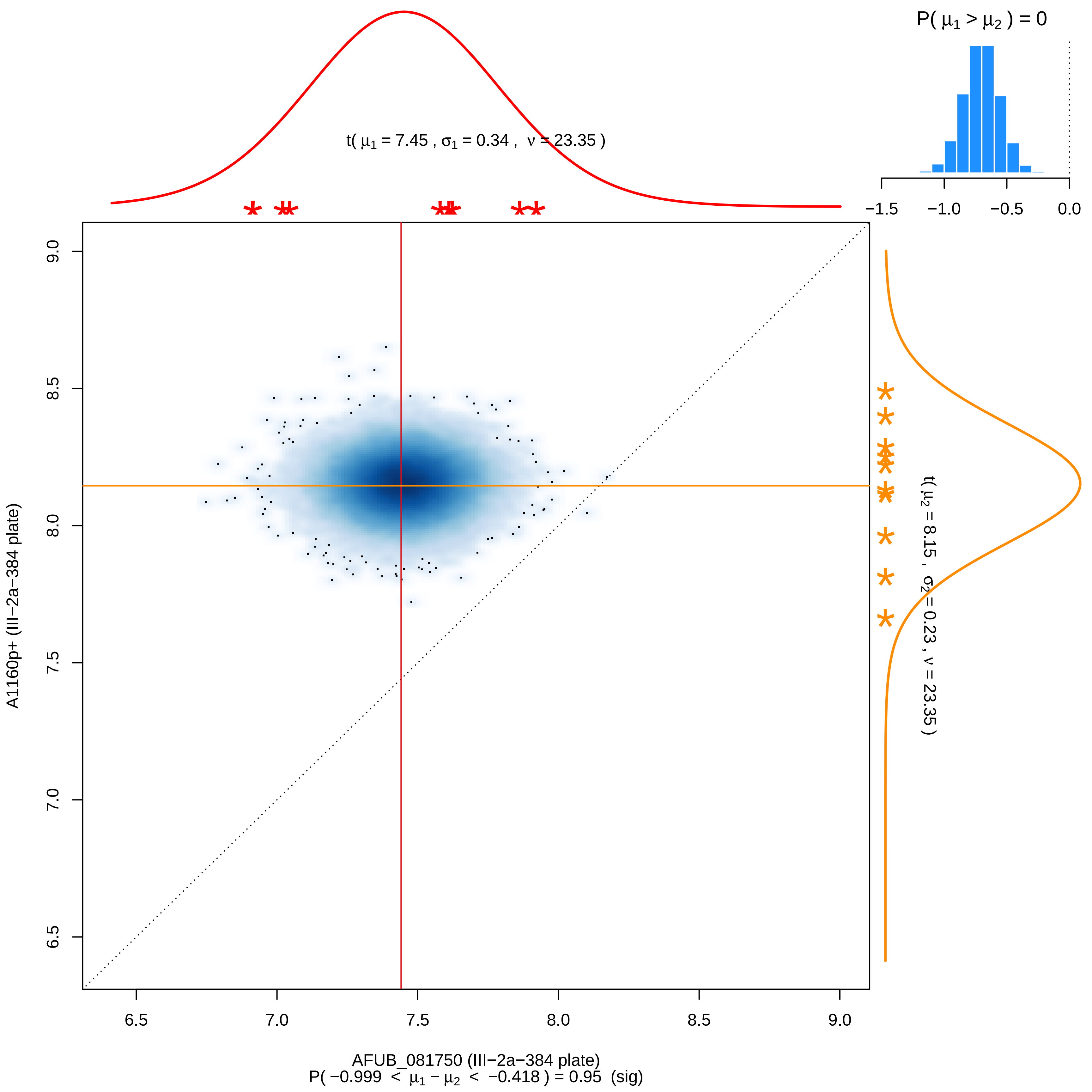

P( μ1 > μ2 ) = 0
−1.5
−1.0
−0.5
0.0
t( μ1 = 7.45 , σ1 = 0.34 , ν = 23.35 )
t( μ2 = 8.15 , σ2 = 0.23 , ν = 23.35 )
9.0
8.5
8.0
7.5
7.0
6.5
A1160p+ (III−2a−384 plate)
6.5
7.0
7.5
8.0
8.5
9.0
AFUB_081750 (III−2a−384 plate)
P( −0.999 < μ1 − μ2 < −0.418 ) = 0.95 (sig)

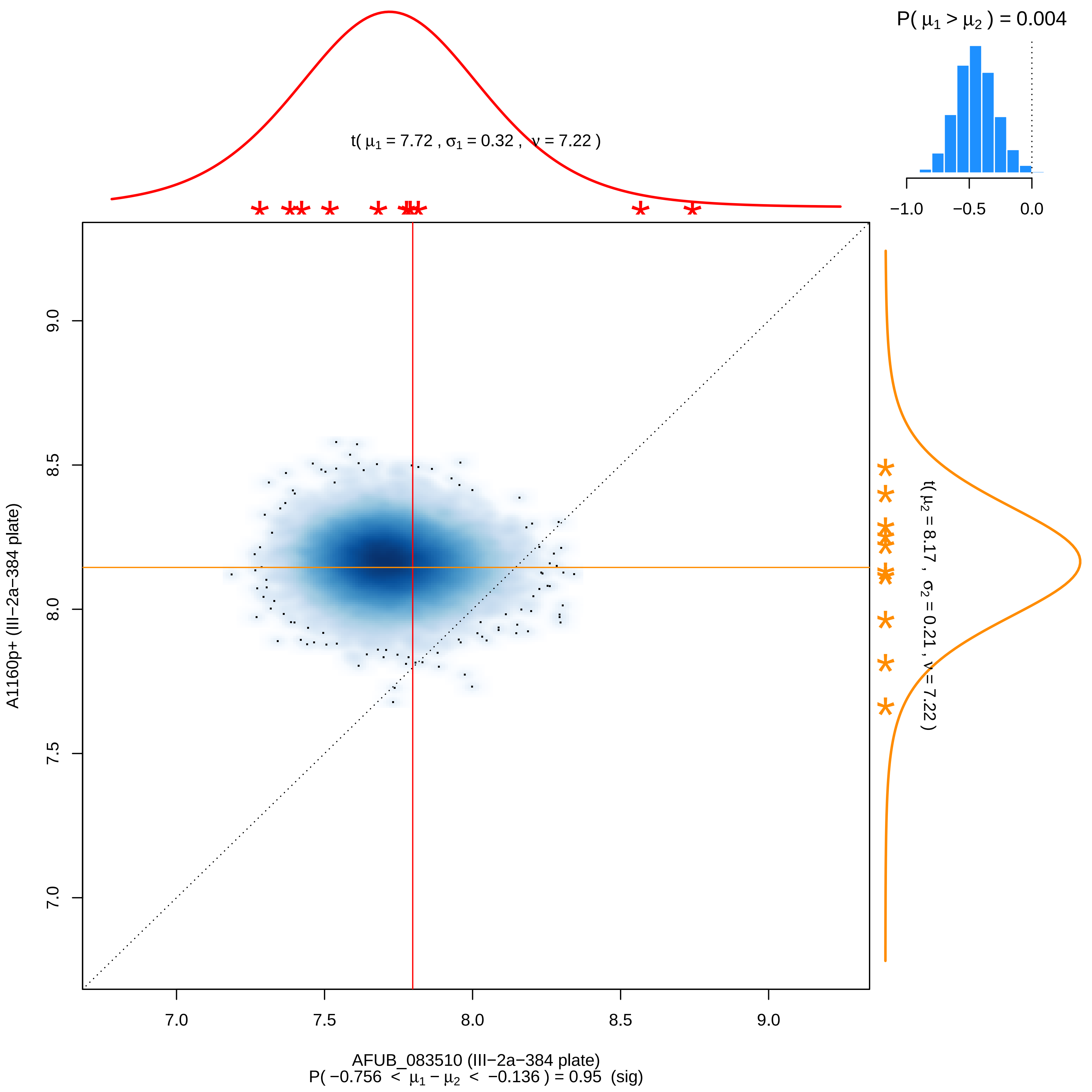

P( μ1 > μ2 ) = 0.004
−1.0
−0.5
0.0
t( μ1 = 7.72 , σ1 = 0.32 , ν = 7.22 )
t( μ2 = 8.17 , σ2 = 0.21 , ν = 7.22 )
9.0
8.5
8.0
7.5
7.0
A1160p+ (III−2a−384 plate)
7.0
7.5
8.0
8.5
9.0
AFUB_083510 (III−2a−384 plate)
P( −0.756 < μ1 − μ2 < −0.136 ) = 0.95 (sig)

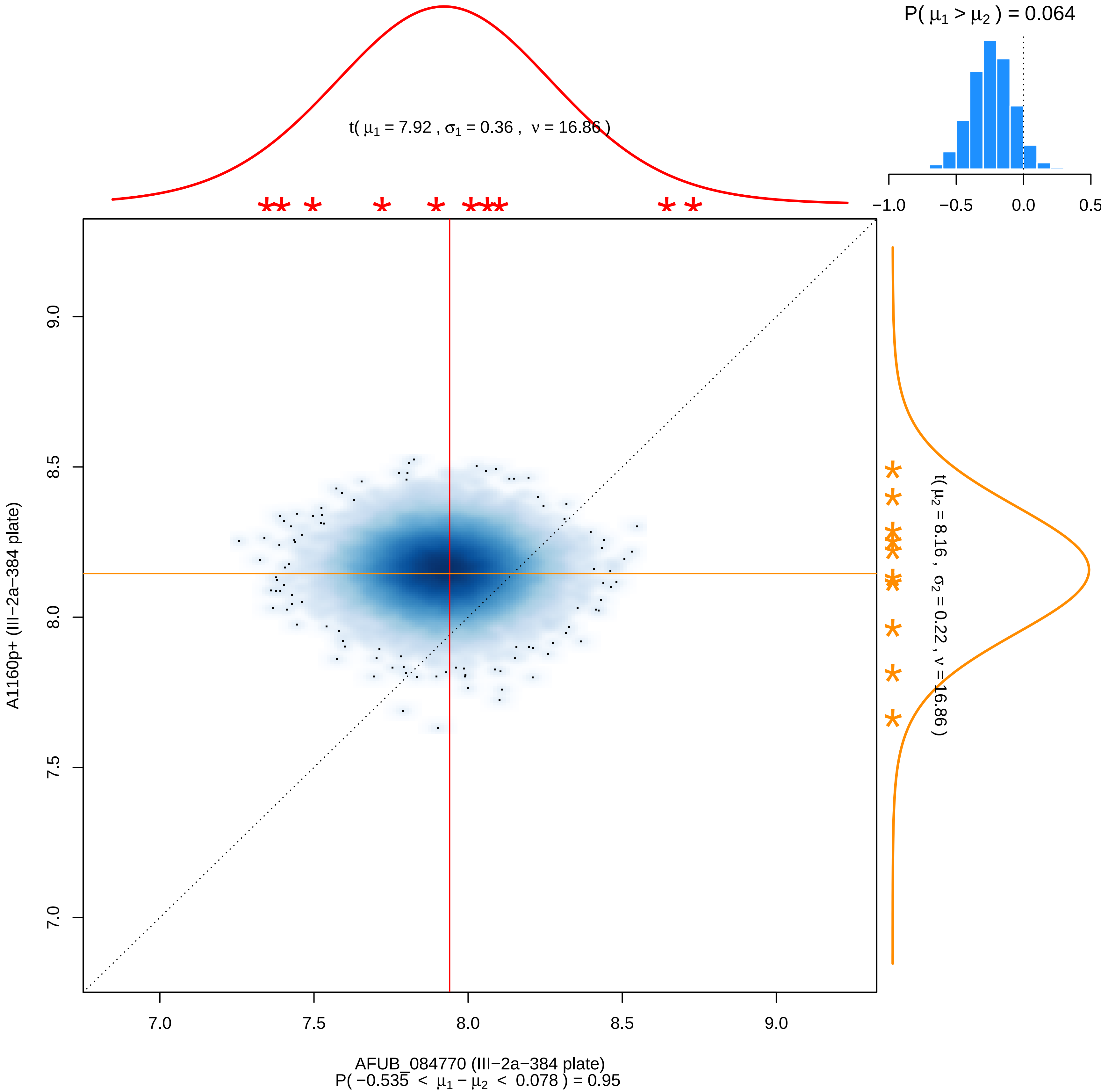

P( μ1 > μ2 ) = 0.064
t( μ1 = 7.92 , σ1 = 0.36 , ν = 16.86 )
−1.0
−0.5
0.0
0.5
t( μ2 = 8.16 , σ2 = 0.22 , ν = 16.86 )
7.0
7.5
8.0
8.5
9.0
A1160p+ (III−2a−384 plate)
AFUB_084770 (III−2a−384 plate)
P( −0.535 < μ1 − μ2 < 0.078 ) = 0.95

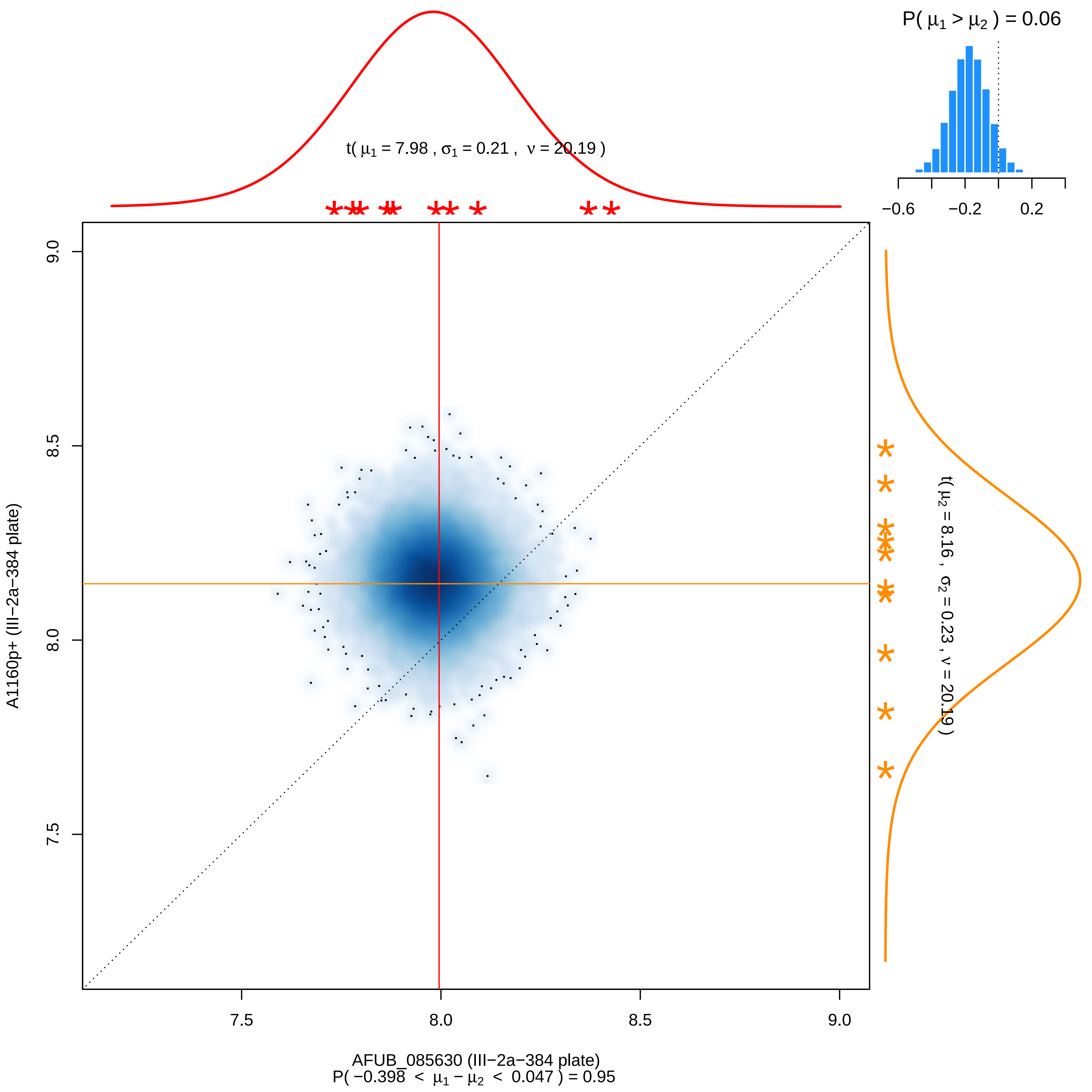
P( μ1 > μ2 ) = 0.06
−0.6
−0.2
0.2
t( μ1 = 7.98 , σ1 = 0.21 , ν = 20.19 )
t( μ2 = 8.16 , σ2 = 0.23 , ν = 20.19 )
9.0
8.5
8.0
7.5
A1160p+ (III−2a−384 plate)
7.5
8.0
8.5
9.0
AFUB_085630 (III−2a−384 plate)
P( −0.398 < μ1 − μ2 < 0.047 ) = 0.95

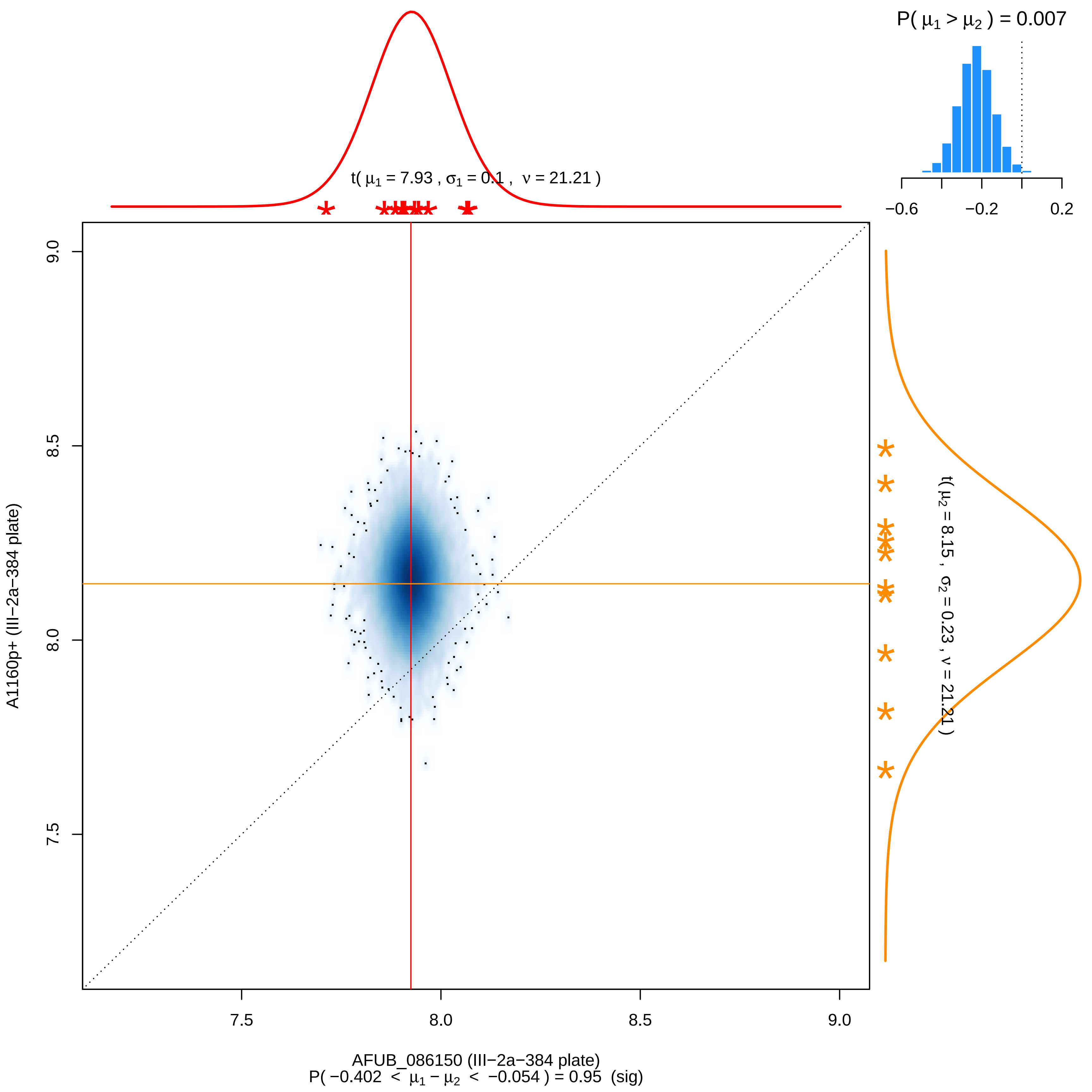

P( μ1 > μ2 ) = 0.007
−0.6
−0.2
0.2
t( μ1 = 7.93 , σ1 = 0.1 , ν = 21.21 )
t( μ2 = 8.15 , σ2 = 0.23 , ν = 21.21 )
9.0
8.5
8.0
7.5
A1160p+ (III−2a−384 plate)
7.5
8.0
8.5
9.0
AFUB_086150 (III−2a−384 plate)
P( −0.402 < μ1 − μ2 < −0.054 ) = 0.95 (sig)

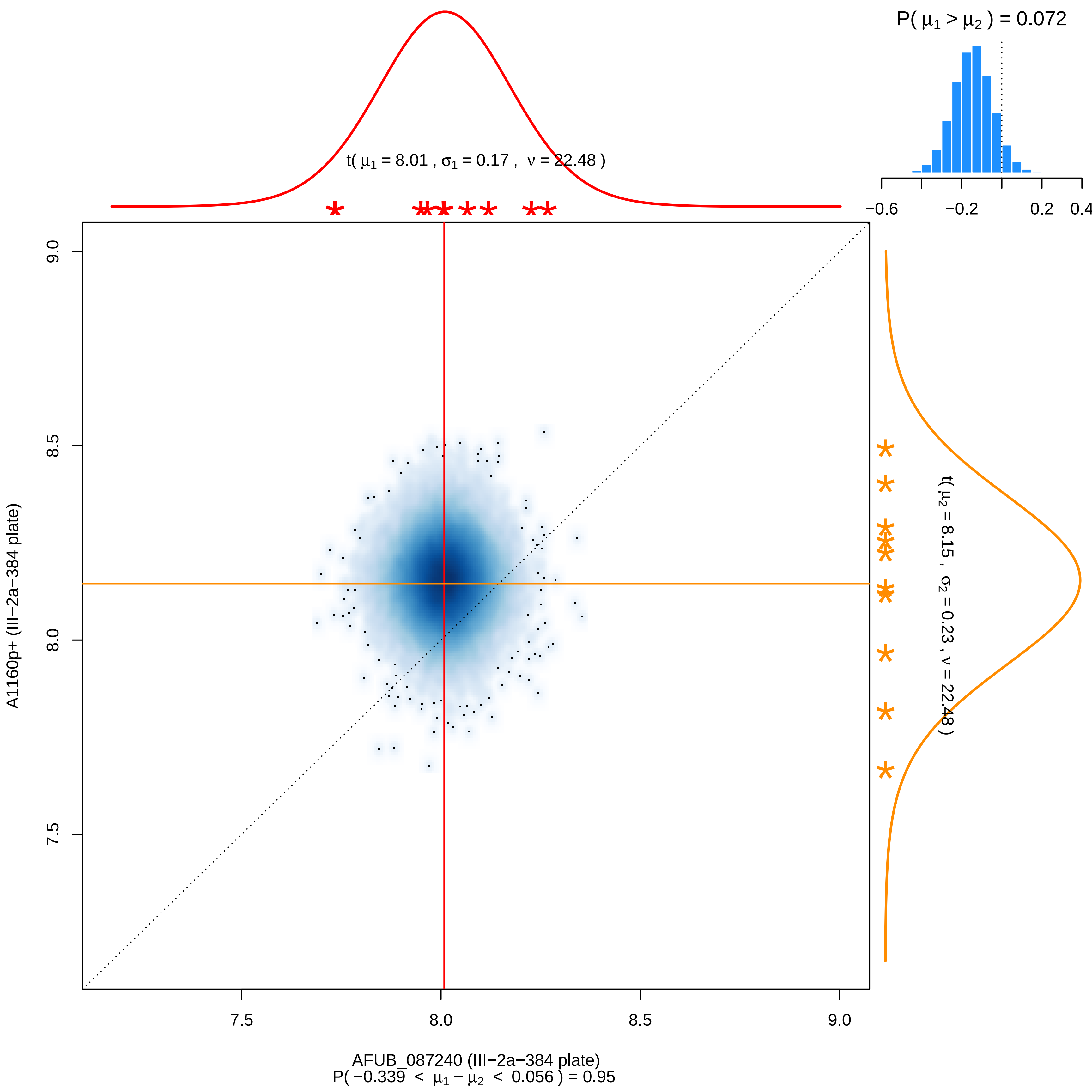

P( μ1 > μ2 ) = 0.072
−0.6
−0.2
0.2
0.4
t( μ1 = 8.01 , σ1 = 0.17 , ν = 22.48 )
t( μ2 = 8.15 , σ2 = 0.23 , ν = 22.48 )
9.0
8.5
8.0
7.5
A1160p+ (III−2a−384 plate)
7.5
8.0
8.5
9.0
AFUB_087240 (III−2a−384 plate)
P( −0.339 < μ1 − μ2 < 0.056 ) = 0.95

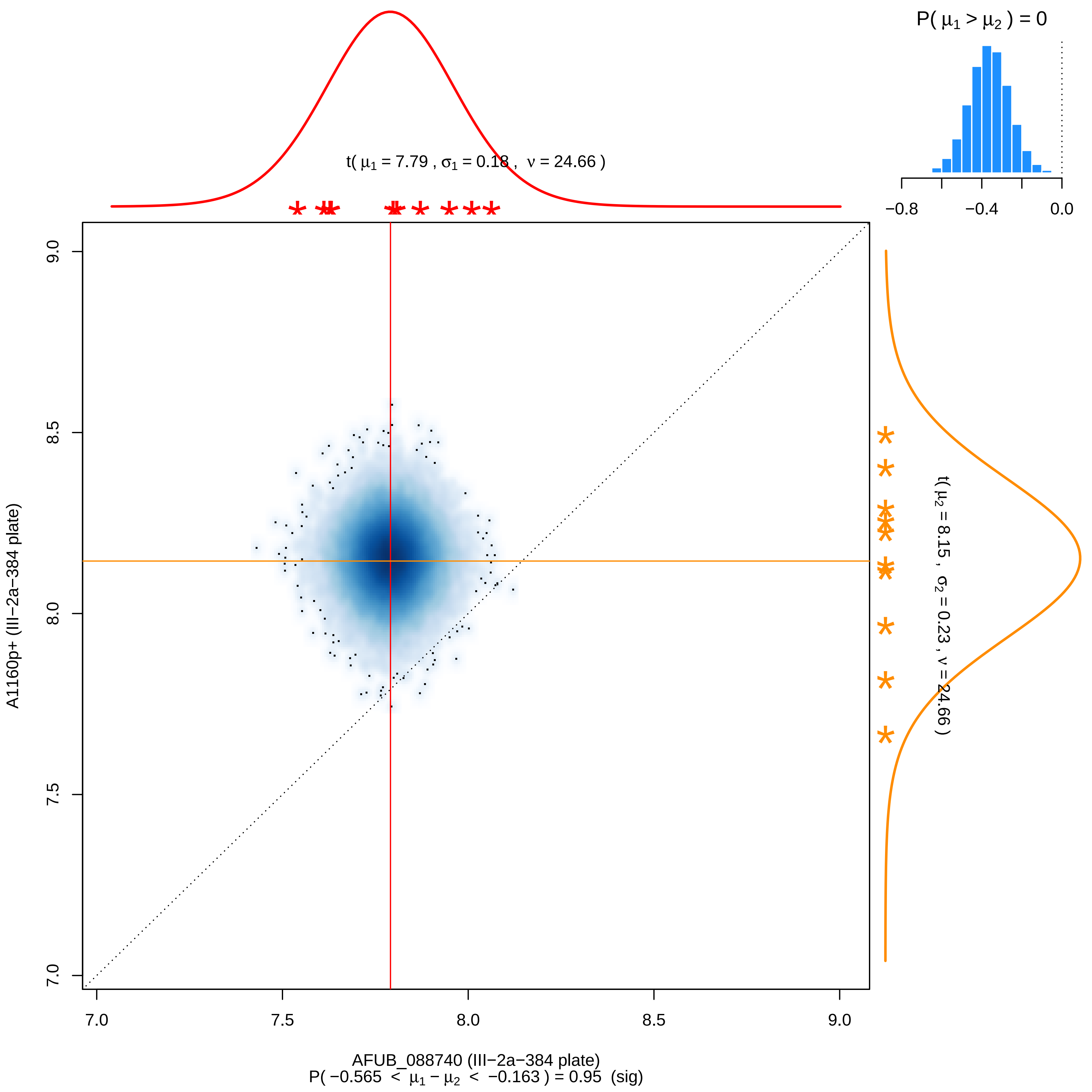

P( μ1 > μ2 ) = 0
−0.8
−0.4
0.0
t( μ1 = 7.79 , σ1 = 0.18 , ν = 24.66 )
t( μ2 = 8.15 , σ2 = 0.23 , ν = 24.66 )
9.0
8.5
8.0
7.5
7.0
A1160p+ (III−2a−384 plate)
7.0
7.5
8.0
8.5
9.0
AFUB_088740 (III−2a−384 plate)
P( −0.565 < μ1 − μ2 < −0.163 ) = 0.95 (sig)

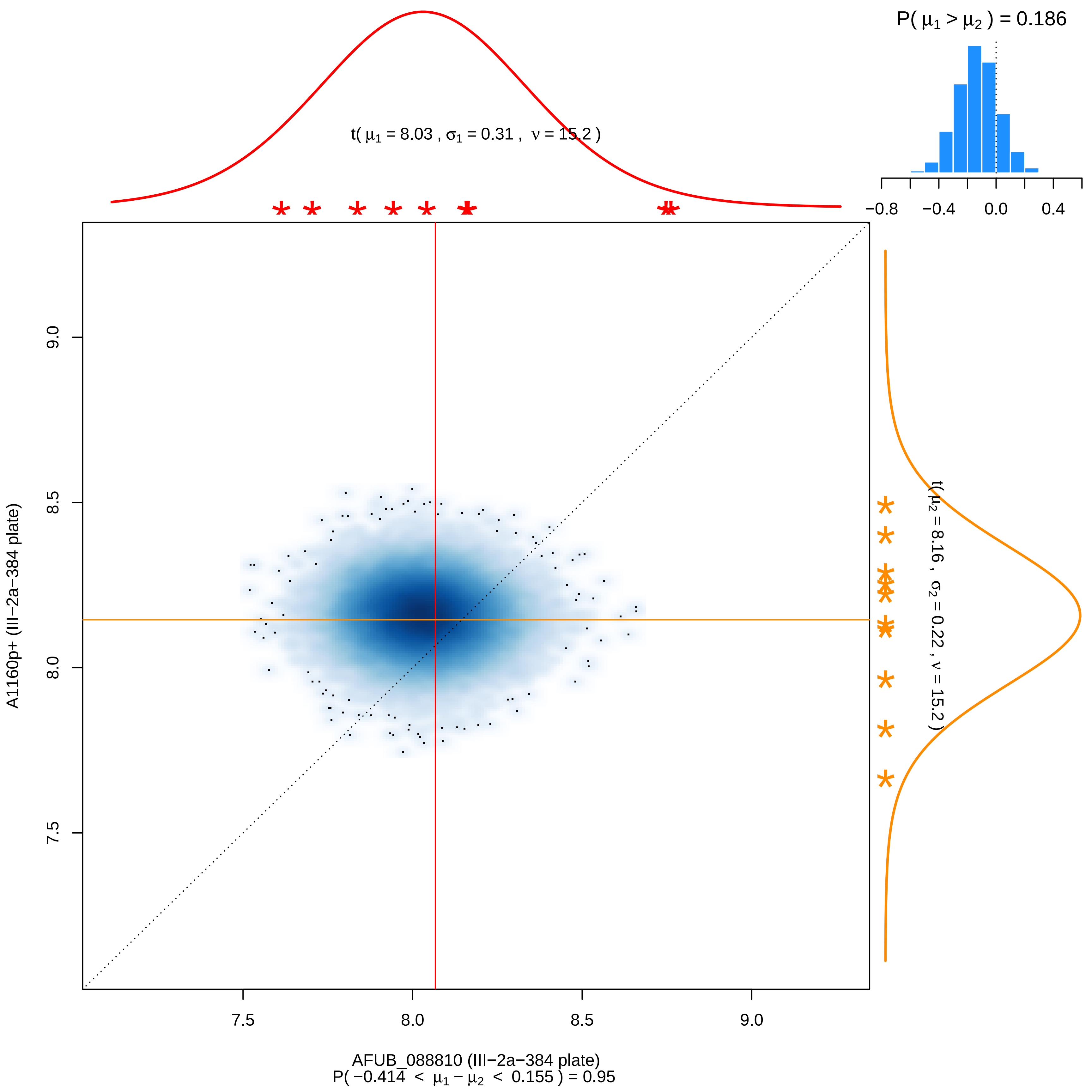

P( μ1 > μ2 ) = 0.186
−0.8
−0.4
0.0
0.4
t( μ1 = 8.03 , σ1 = 0.31 , ν = 15.2 )
t( μ2 = 8.16 , σ2 = 0.22 , ν = 15.2 )
A1160p+ (III−2a−384 plate)
9.0
8.5
8.0
7.5
7.5
8.0
8.5
9.0
AFUB_088810 (III−2a−384 plate)
P( −0.414 < μ1 − μ2 < 0.155 ) = 0.95

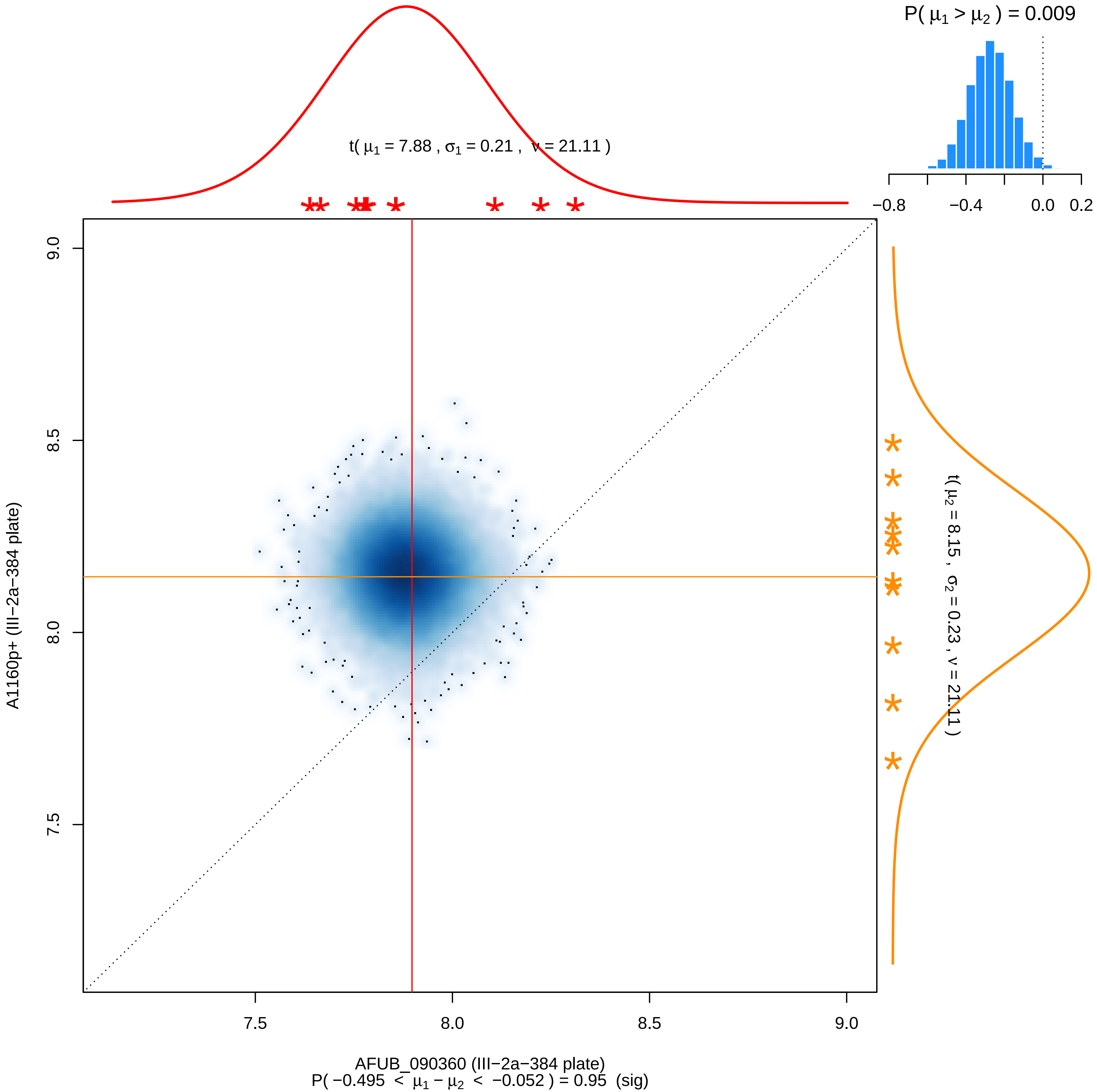

P( μ1 > μ2 ) = 0.009
−0.8
−0.4
0.0
0.2
t( μ1 = 7.88 , σ1 = 0.21 , ν = 21.11 )
t( μ2 = 8.15 , σ2 = 0.23 , ν = 21.11 )
9.0
8.5
8.0
7.5
A1160p+ (III−2a−384 plate)
7.5
8.0
8.5
9.0
AFUB_090360 (III−2a−384 plate)
P( −0.495 < μ1 − μ2 < −0.052 ) = 0.95 (sig)

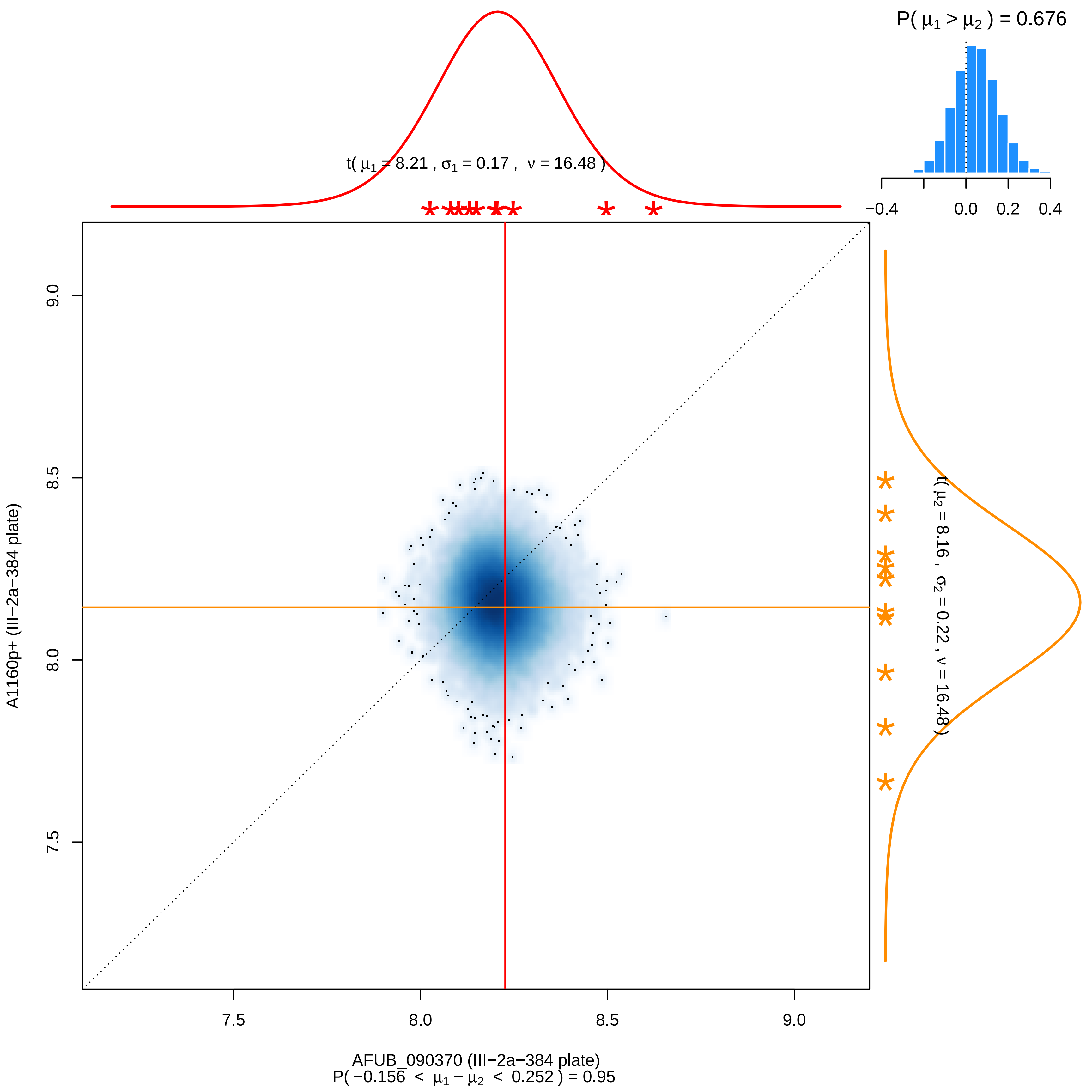

P( μ1 > μ2 ) = 0.676
−0.4
0.0
0.2
0.4
t( μ1 = 8.21 , σ1 = 0.17 , ν = 16.48 )
t( μ2 = 8.16 , σ2 = 0.22 , ν = 16.48 )
9.0
8.5
8.0
7.5
A1160p+ (III−2a−384 plate)
7.5
8.0
8.5
9.0
AFUB_090370 (III−2a−384 plate)
P( −0.156 < μ1 − μ2 < 0.252 ) = 0.95

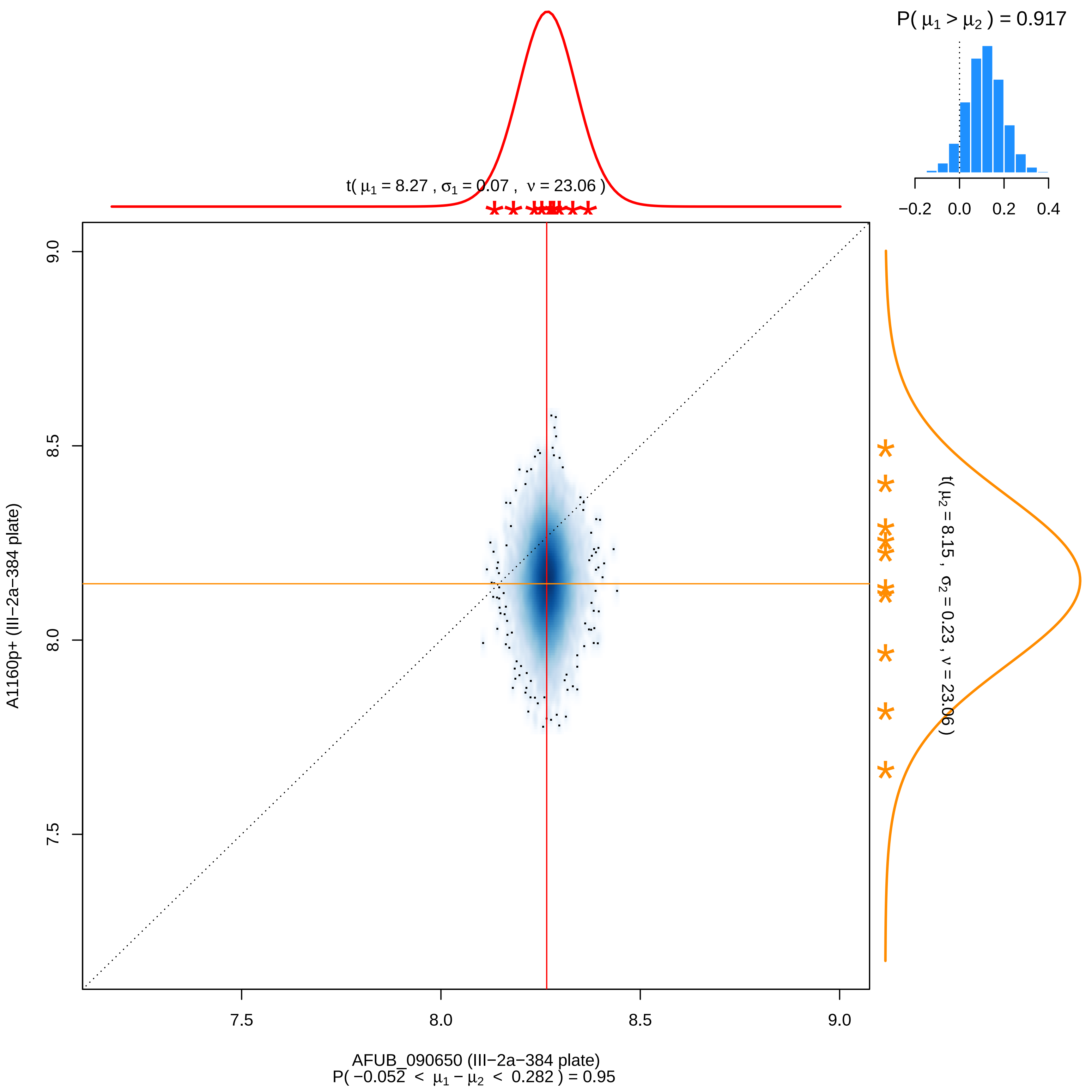

P( μ1 > μ2 ) = 0.917
−0.2 0.0 0.2 0.4
t( μ1 = 8.27 , σ1 = 0.07 , ν = 23.06 )
9.0
8.5
8.0
7.5
A1160p+ (III−2a−384 plate)
t( μ2 = 8.15 , σ2 = 0.23 , ν = 23.06 )
7.5 8.0 8.5 9.0
AFUB_090650 (III−2a−384 plate)
P( −0.052 < μ1 − μ2 < 0.282 ) = 0.95

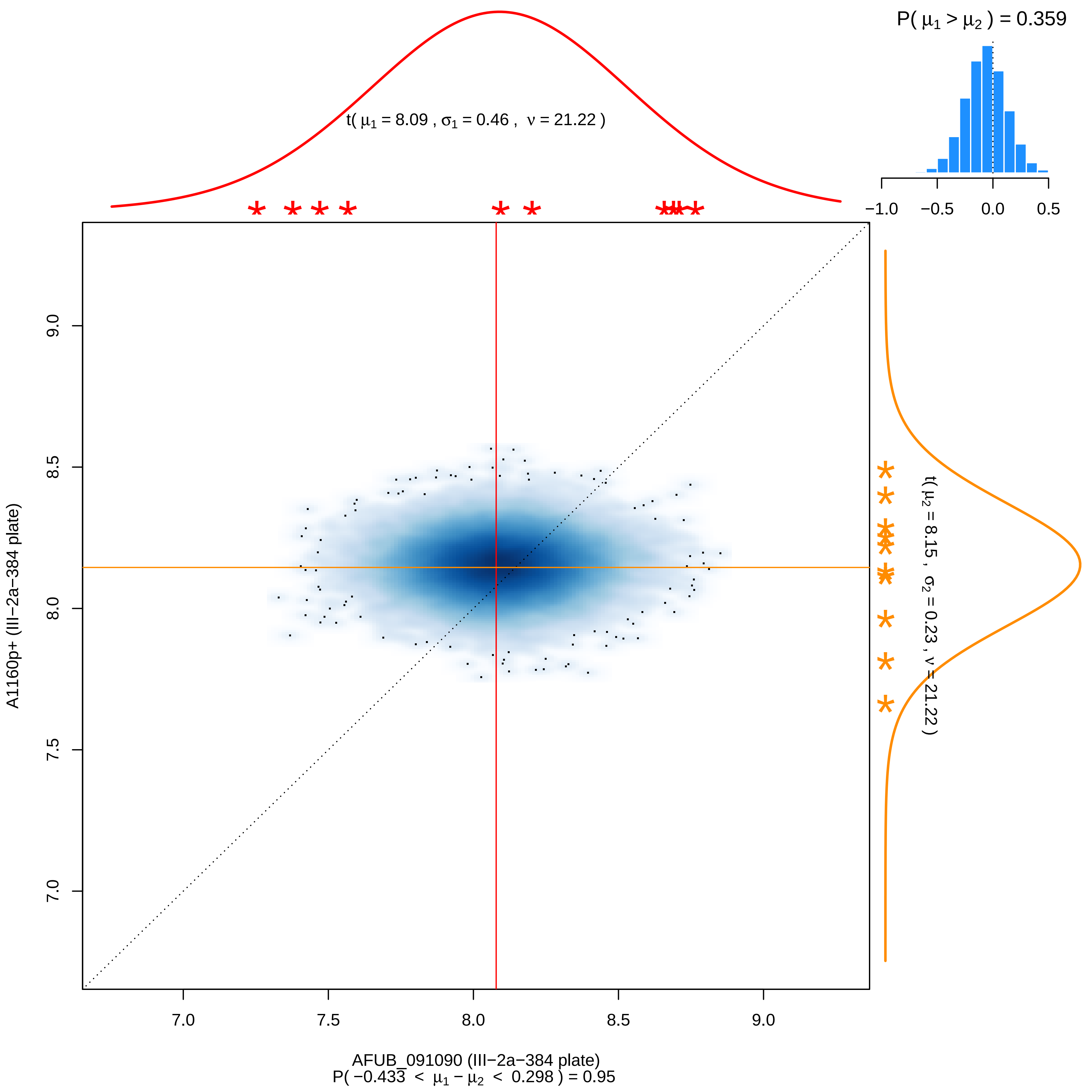

P( μ1 > μ2 ) = 0.359
t( μ1 = 8.09 , σ1 = 0.46 , ν = 21.22 )
−1.0
−0.5
0.0
0.5
t( μ2 = 8.15 , σ2 = 0.23 , ν = 21.22 )
A1160p+ (III−2a−384 plate)
7.0
7.5
8.0
8.5
9.0
AFUB_091090 (III−2a−384 plate)
P( −0.433 < μ1 − μ2 < 0.298 ) = 0.95

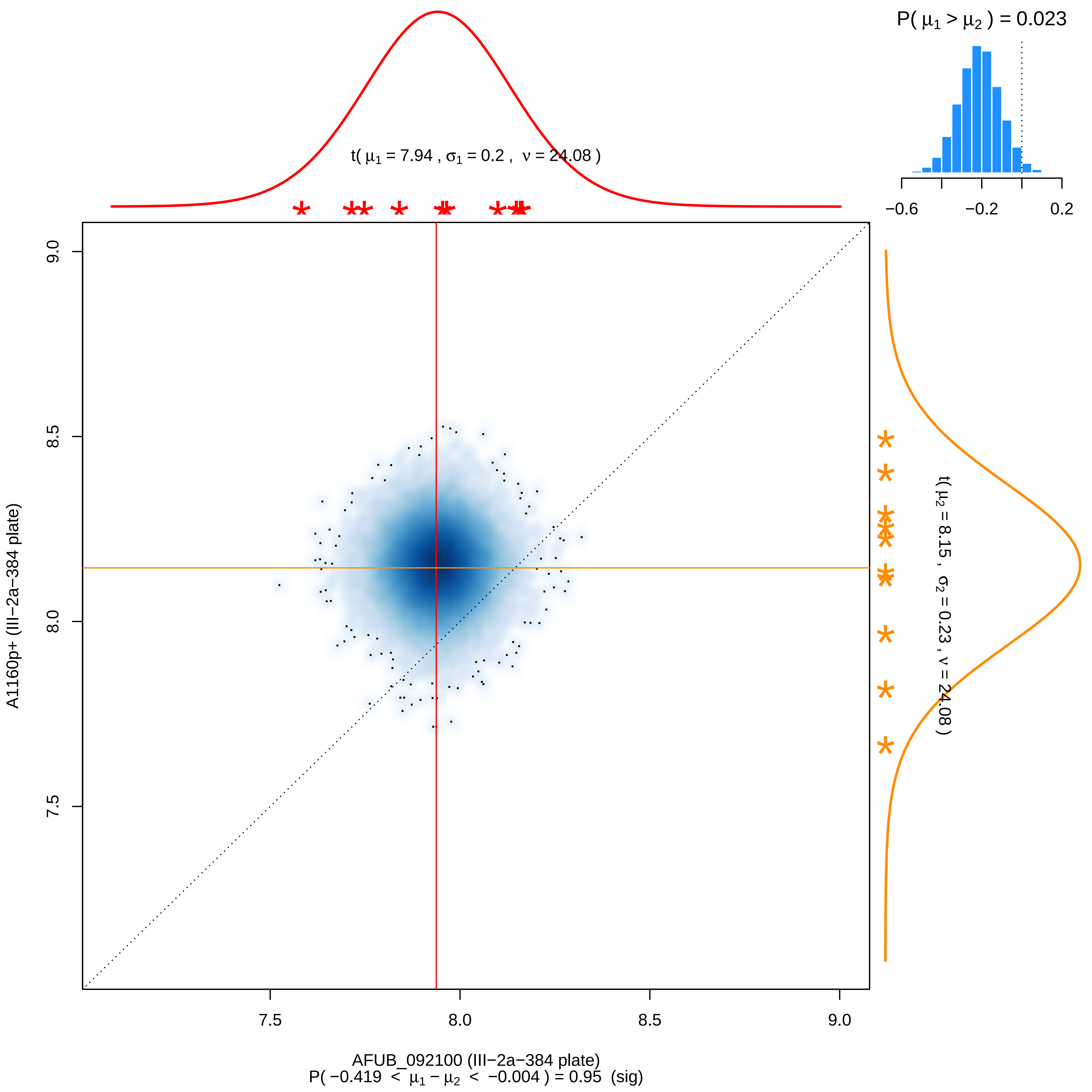

P( μ1 > μ2 ) = 0.023
−0.6
−0.2
0.2
t( μ1 = 7.94 , σ1 = 0.2 , ν = 24.08 )
9.0
8.5
8.0
7.5
A1160p+ (III−2a−384 plate)
t( μ2 = 8.15 , σ2 = 0.23 , ν = 24.08 )
7.5
8.0
8.5
9.0
AFUB_092100 (III−2a−384 plate)
P( −0.419 < μ1 − μ2 < −0.004 ) = 0.95 (sig)

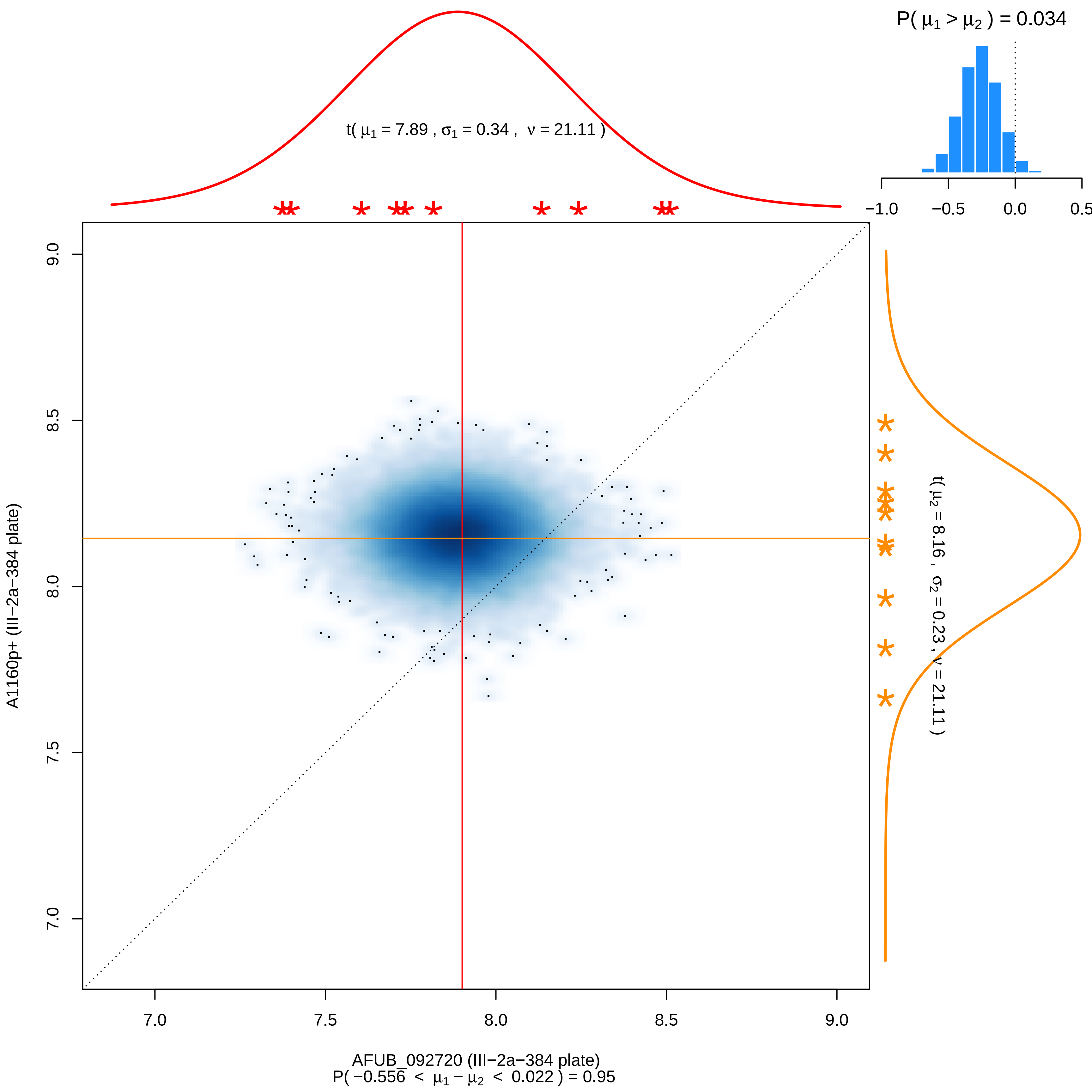
P( μ1 > μ2 ) = 0.034
−1.0
−0.5
0.0
0.5
t( μ1 = 7.89 , σ1 = 0.34 , ν = 21.11 )
t( μ2 = 8.16 , σ2 = 0.23 , ν = 21.11 )
A1160p+ (III−2a−384 plate)
9.0
8.5
8.0
7.5
7.0
7.0
7.5
8.0
8.5
9.0
AFUB_092720 (III−2a−384 plate)
P( −0.556 < μ1 − μ2 < 0.022 ) = 0.95

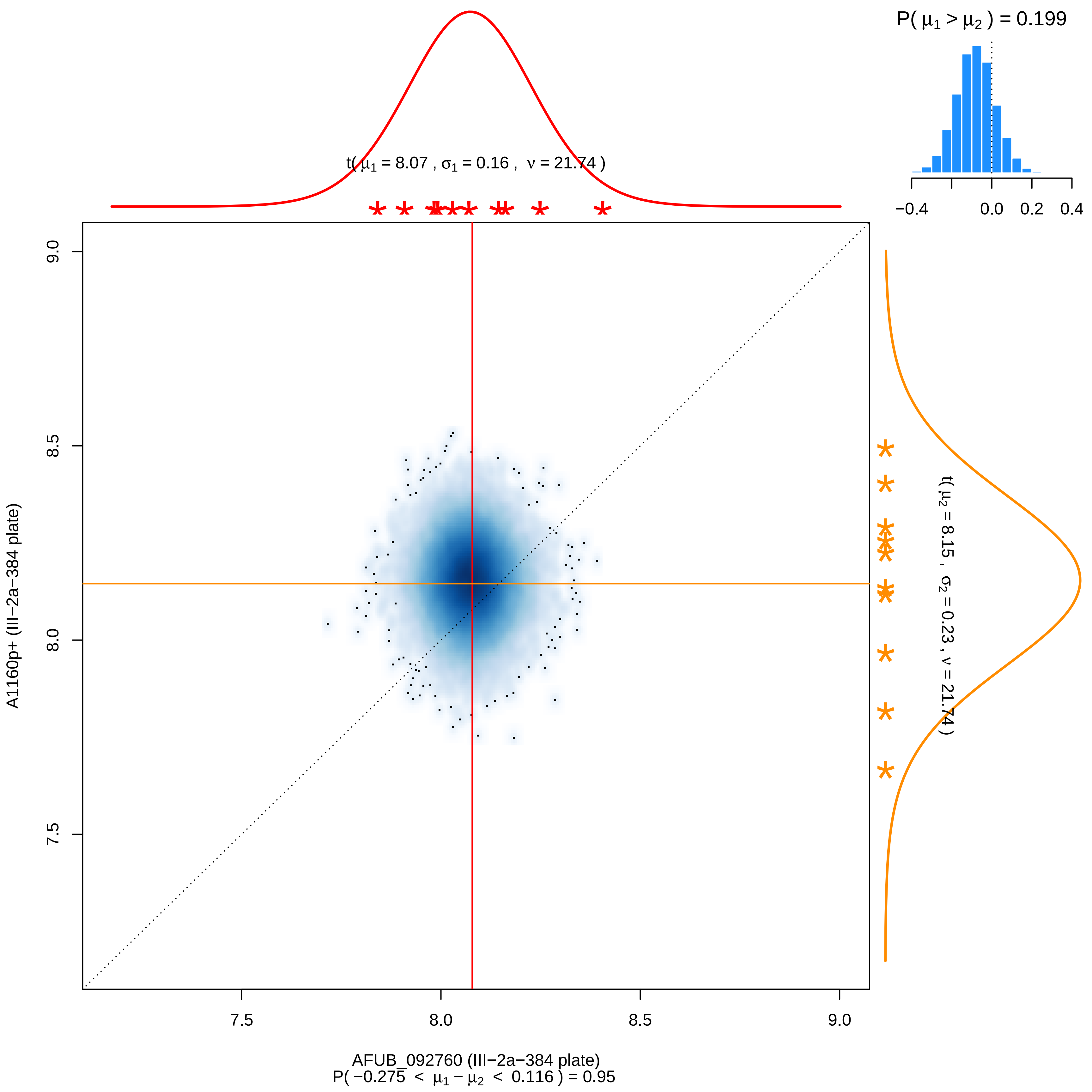

P( μ1 > μ2 ) = 0.199
−0.4
0.0
0.2
0.4
t( μ1 = 8.07 , σ1 = 0.16 , ν = 21.74 )
9.0
8.5
8.0
7.5
A1160p+ (III−2a−384 plate)
t( μ2 = 8.15 , σ2 = 0.23 , ν = 21.74 )
7.5
8.0
8.5
9.0
AFUB_092760 (III−2a−384 plate)
P( −0.275 < μ1 − μ2 < 0.116 ) = 0.95

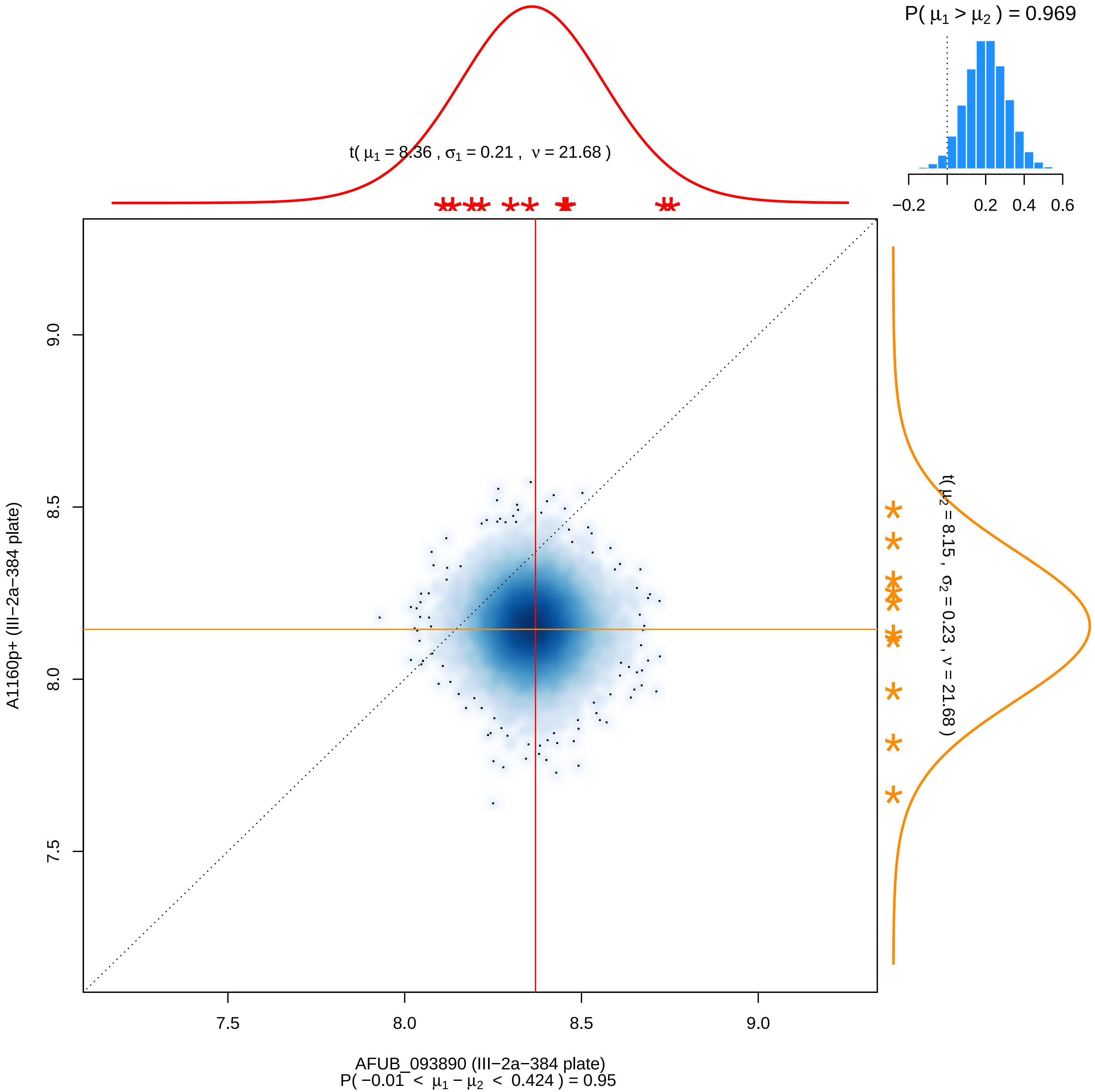

P( μ1 > μ2 ) = 0.969
−0.2
0.2
0.4
0.6
t( μ1 = 8.36 , σ1 = 0.21 , ν = 21.68 )
t( μ2 = 8.15 , σ2 = 0.23 , ν = 21.68 )
9.0
8.5
8.0
7.5
A1160p+ (III−2a−384 plate)
7.5
8.0
8.5
9.0
AFUB_093890 (III−2a−384 plate)
P( −0.01 < μ1 − μ2 < 0.424 ) = 0.95

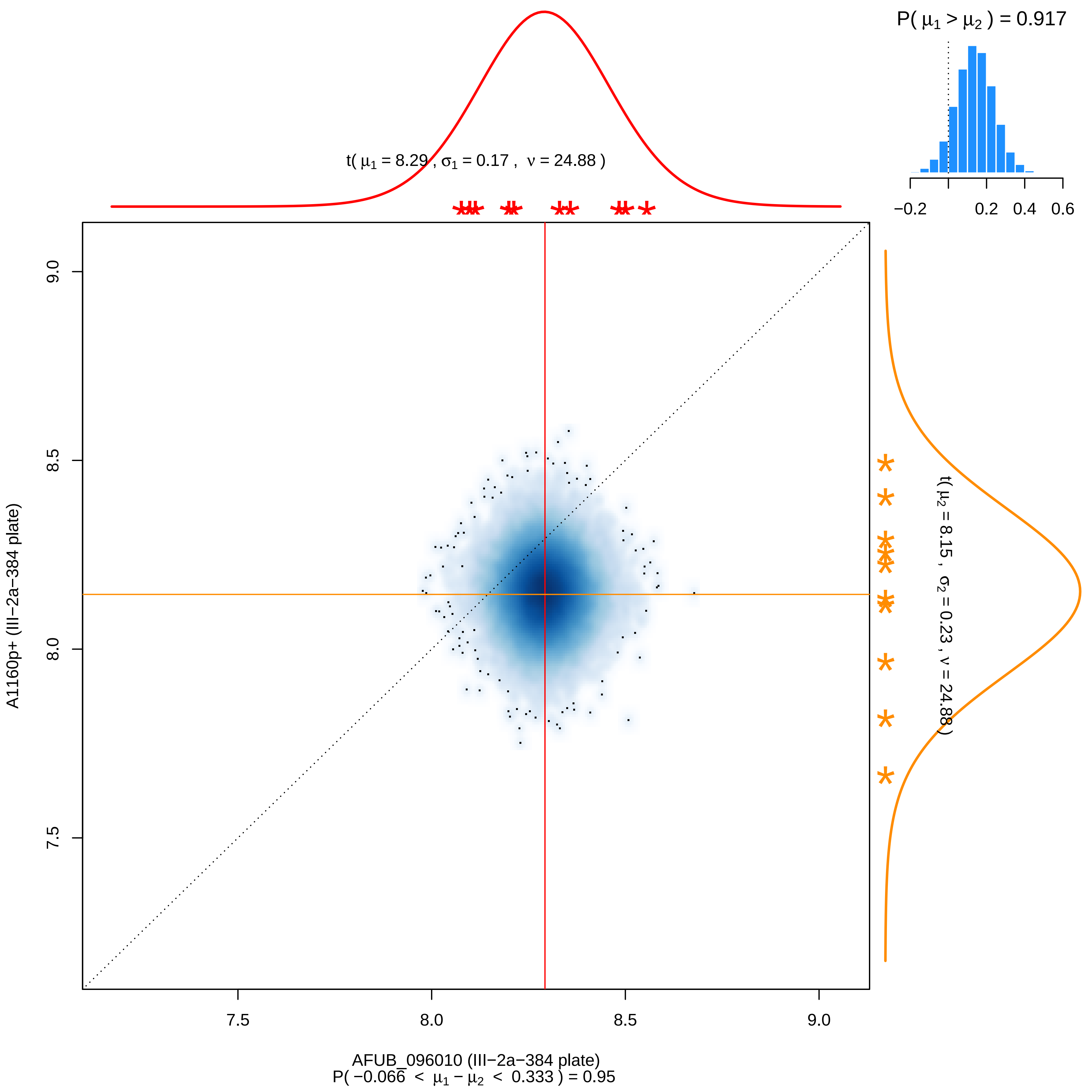

P( μ1 > μ2 ) = 0.917
−0.2
0.2
0.4
0.6
t( μ1 = 8.29 , σ1 = 0.17 , ν = 24.88 )
t( μ2 = 8.15 , σ2 = 0.23 , ν = 24.88 )
9.0
8.5
8.0
7.5
A1160p+ (III−2a−384 plate)
7.5
8.0
8.5
9.0
AFUB_096010 (III−2a−384 plate)
P( −0.066 < μ1 − μ2 < 0.333 ) = 0.95

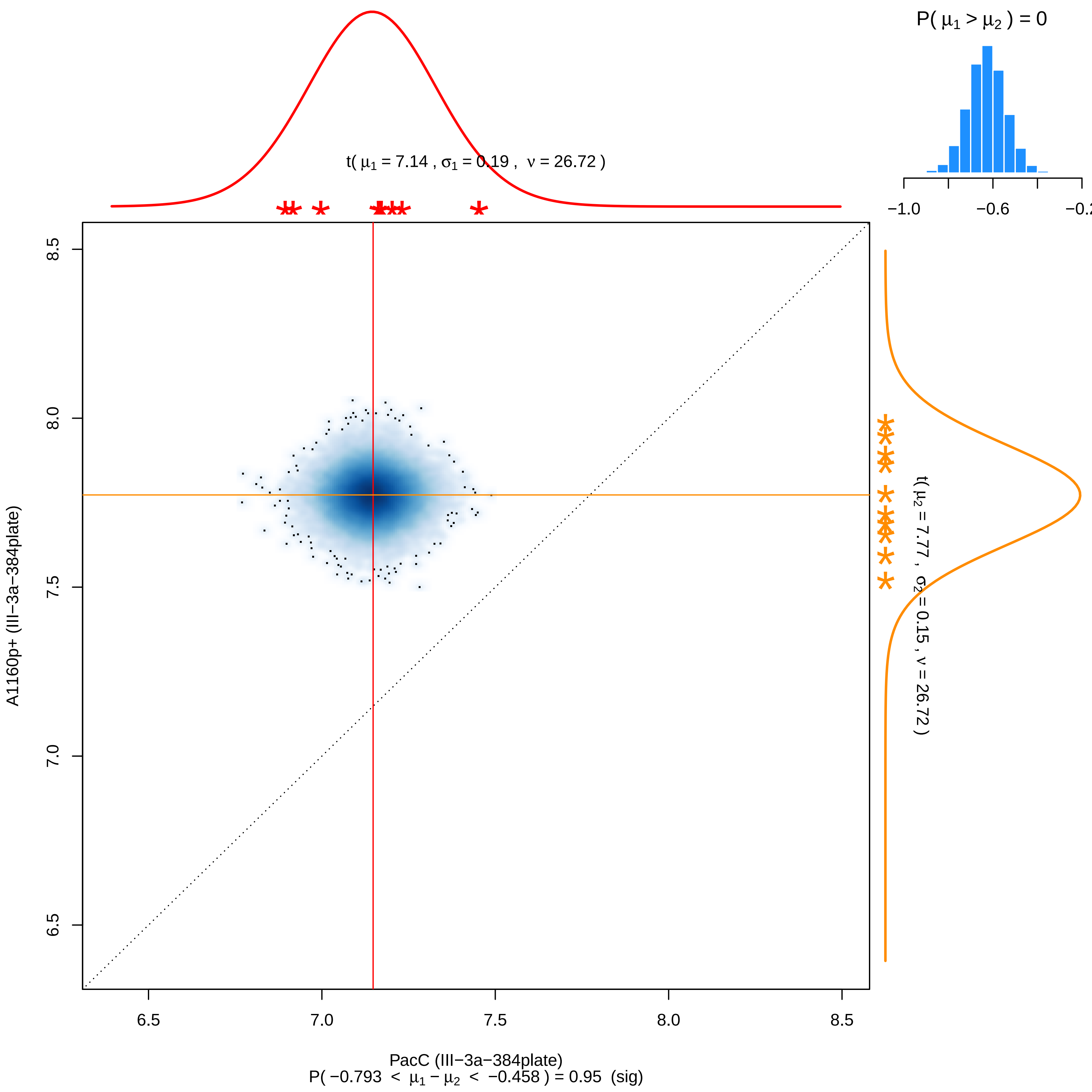

P( μ1 > μ2 ) = 0
−1.0
−0.6
−0.2
t( μ1 = 7.14 , σ1 = 0.19 , ν = 26.72 )
t( μ2 = 7.77 , σ2 = 0.15 , ν = 26.72 )
8.5
8.0
7.5
7.0
6.5
A1160p+ (III−3a−384plate)
6.5
7.0
7.5
8.0
8.5
PacC (III−3a−384plate)
P( −0.793 < μ1 − μ2 < −0.458 ) = 0.95 (sig)

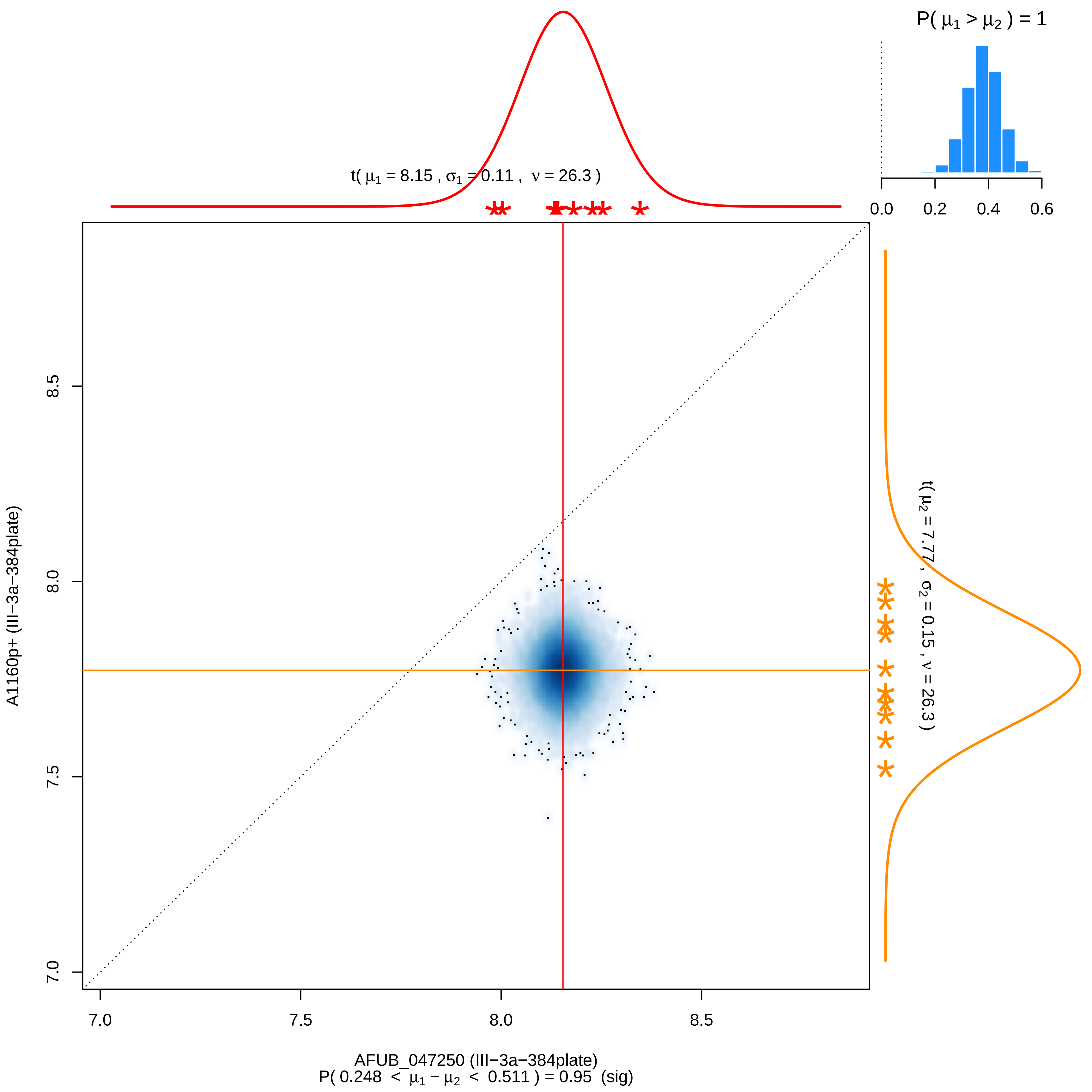
P( μ1 > μ2 ) = 1
0.0
0.2
0.4
0.6
t( μ1 = 8.15 , σ1 = 0.11 , ν = 26.3 )
t( μ2 = 7.77 , σ2 = 0.15 , ν = 26.3 )
A1160p+ (III−3a−384plate)
7.0
7.5
8.0
8.5
AFUB_047250 (III−3a−384plate)
P( 0.248 < μ1 − μ2 < 0.511 ) = 0.95 (sig)

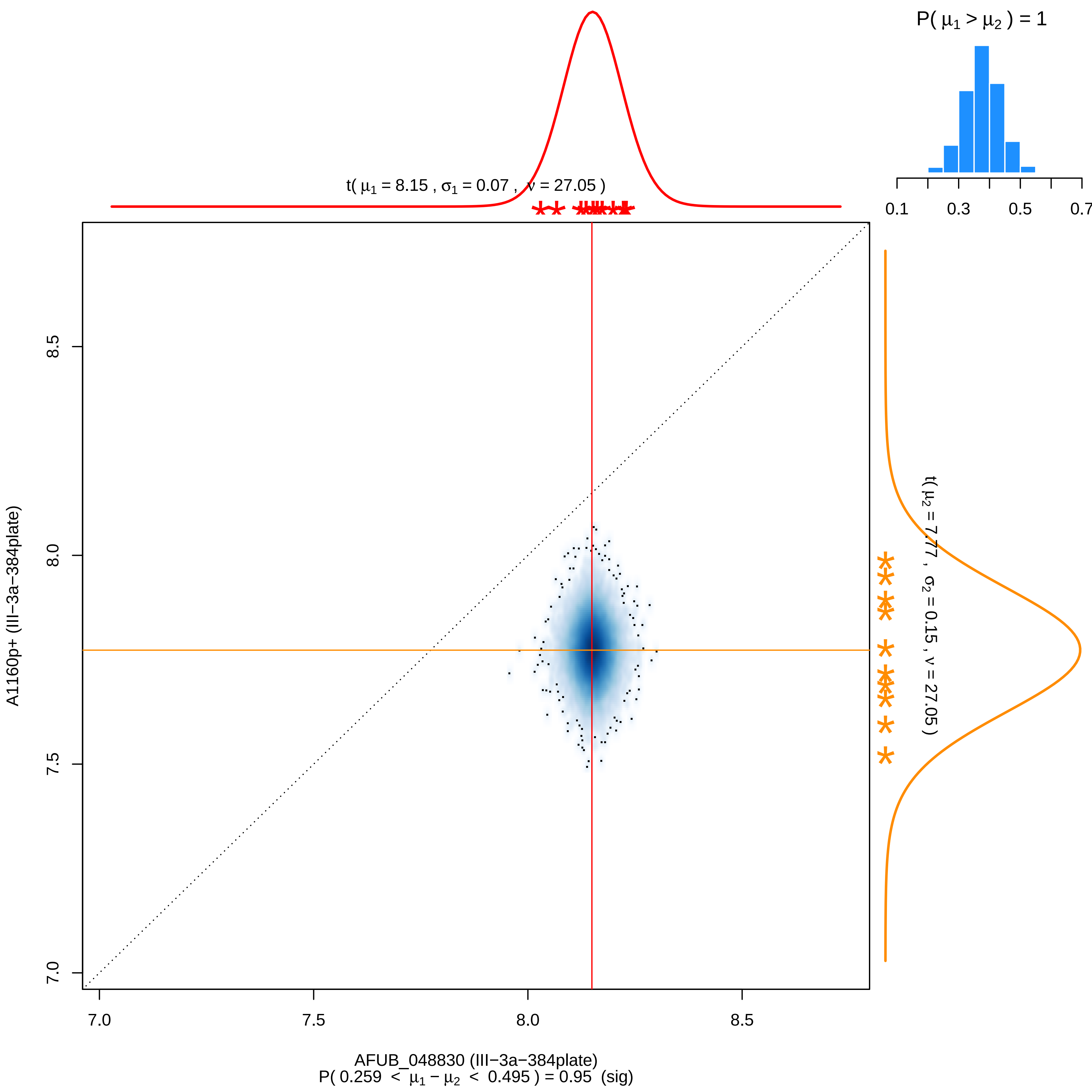
P( μ1 > μ2 ) = 1
0.1
0.3
0.5
0.7
t( μ1 = 8.15 , σ1 = 0.07 , ν = 27.05 )
t( μ2 = 7.77 , σ2 = 0.15 , ν = 27.05 )
8.5
8.0
7.5
7.0
A1160p+ (III−3a−384plate)
7.0
7.5
8.0
8.5
AFUB_048830 (III−3a−384plate)
P( 0.259 < μ1 − μ2 < 0.495 ) = 0.95 (sig)

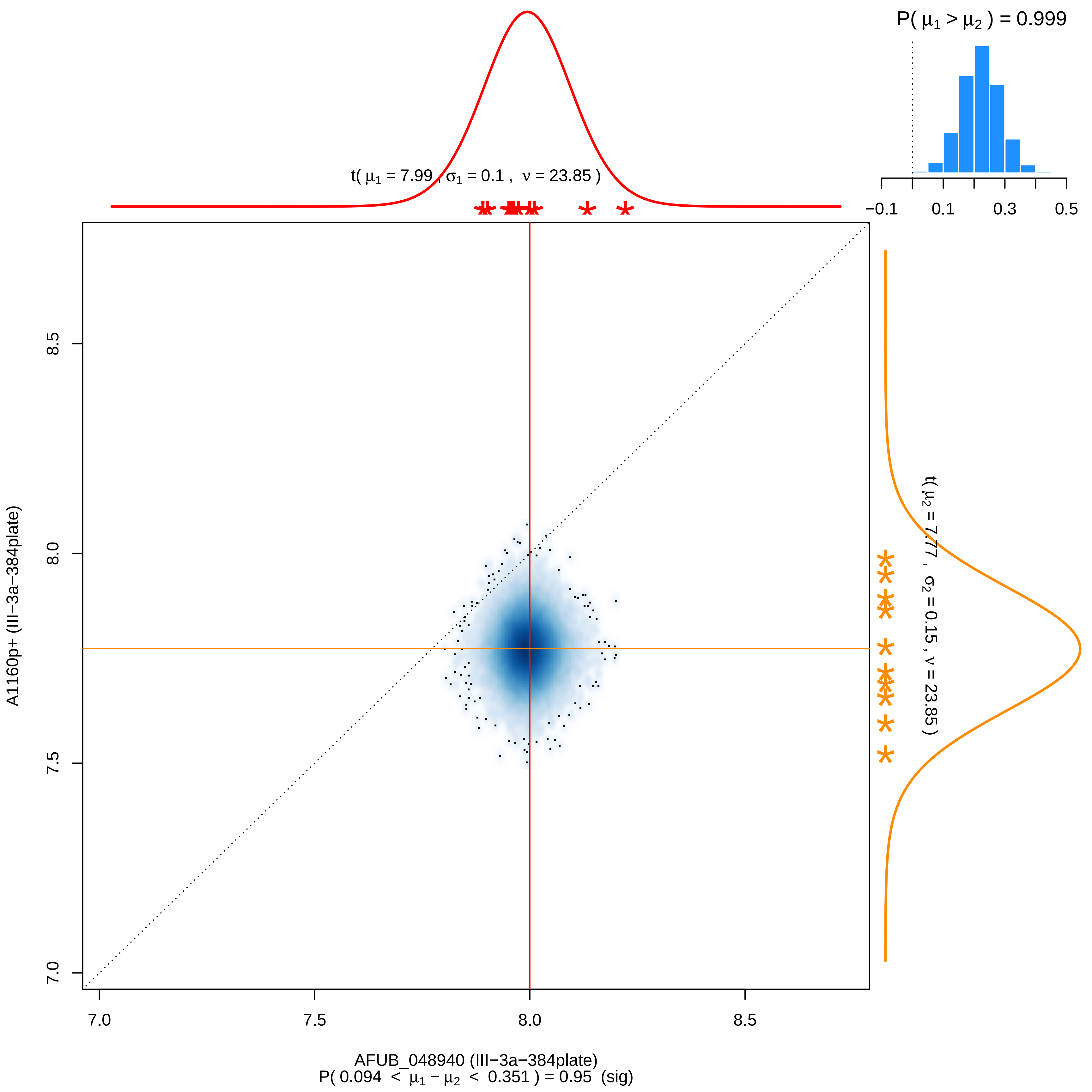

P( $\mu_1 > \mu_2$ ) = 0.999
−0.1
0.1
0.3
0.5
t( $\mu_1$ = 7.99 , $\sigma_1$ = 0.1 , $\nu$ = 23.85 )
t( $\mu_2$ = 7.77 , $\sigma_2$ = 0.15 , $\nu$ = 23.85 )
8.5
8.0
7.5
7.0
A1160p+ (III−3a−384plate)
7.0
7.5
8.0
8.5
AFUB_048940 (III−3a−384plate)
P( 0.094 < $\mu_1 - \mu_2$ < 0.351 ) = 0.95 (sig)

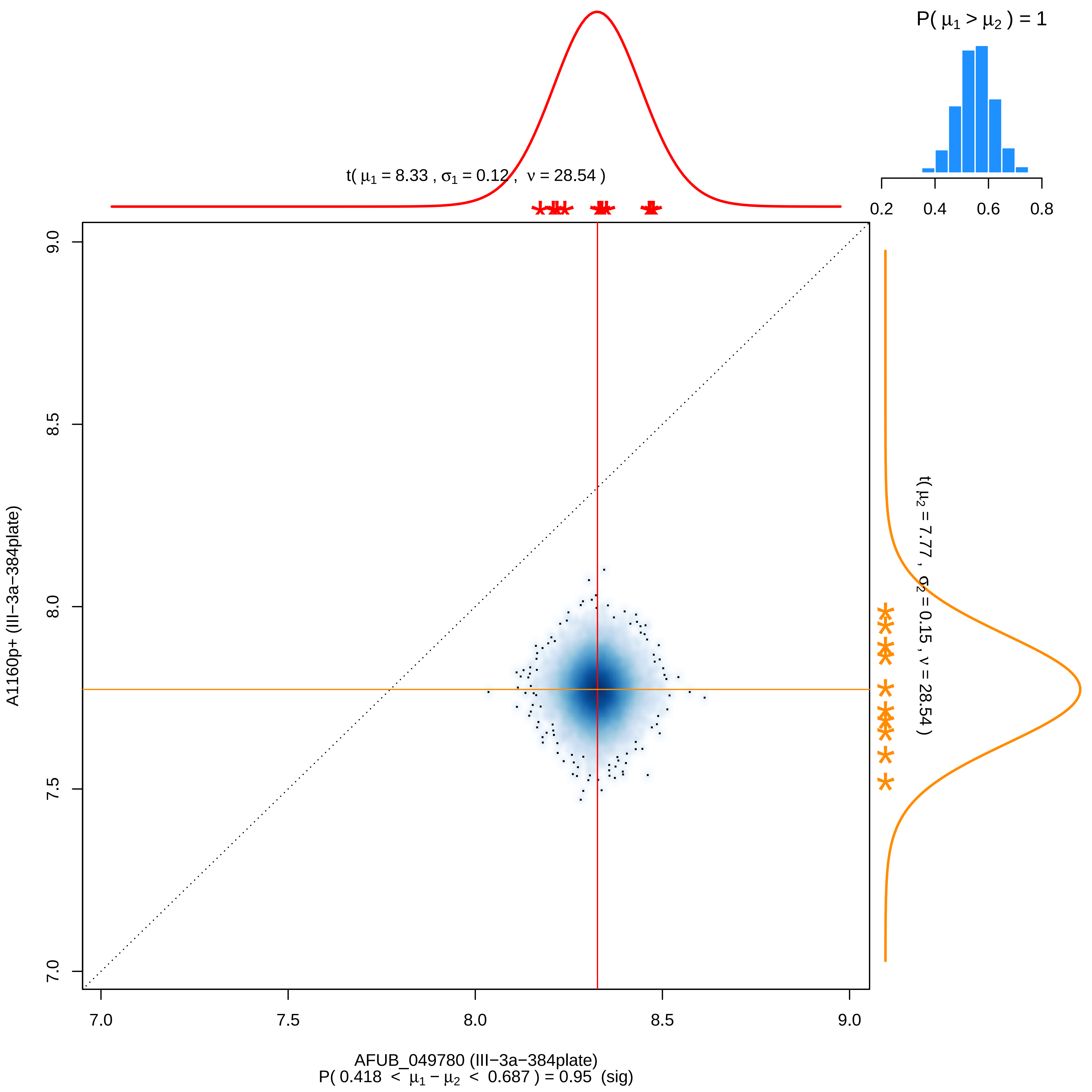

P( μ1 > μ2 ) = 1
0.2
0.4
0.6
0.8
t( μ1 = 8.33 , σ1 = 0.12 , ν = 28.54 )
t( μ2 = 7.77 , σ2 = 0.15 , ν = 28.54 )
A1160p+ (III−3a−384plate)
9.0
8.5
8.0
7.5
7.0
7.0
7.5
8.0
8.5
9.0
AFUB_049780 (III−3a−384plate)
P( 0.418 < μ1 − μ2 < 0.687 ) = 0.95 (sig)

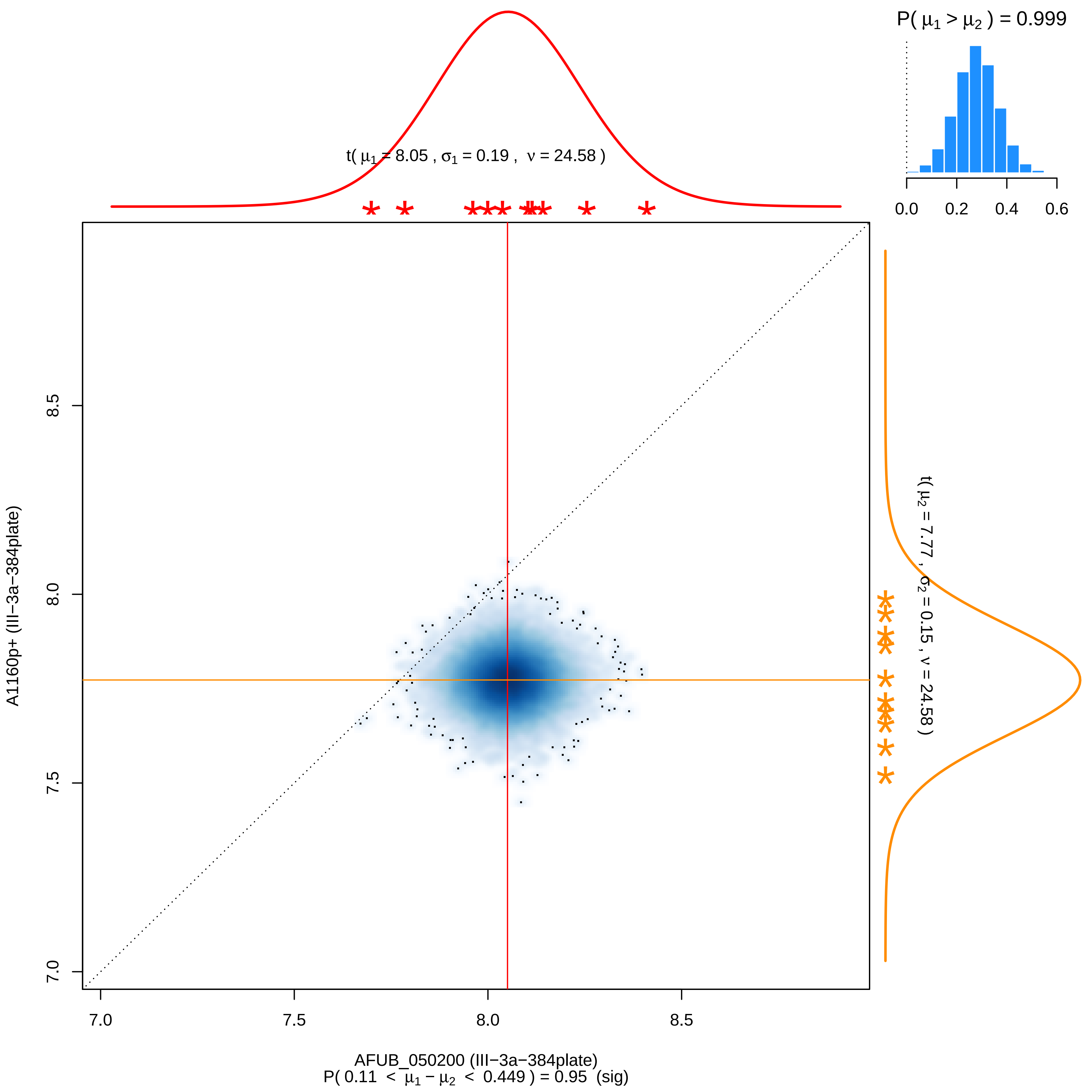

P( μ1 > μ2 ) = 0.999
0.0
0.2
0.4
0.6
t( μ1 = 8.05 , σ1 = 0.19 , ν = 24.58 )
t( μ2 = 7.77 , σ2 = 0.15 , ν = 24.58 )
8.5
8.0
7.5
7.0
A1160p+ (III−3a−384plate)
7.0
7.5
8.0
8.5
AFUB_050200 (III−3a−384plate)
P( 0.11 < μ1 − μ2 < 0.449 ) = 0.95 (sig)

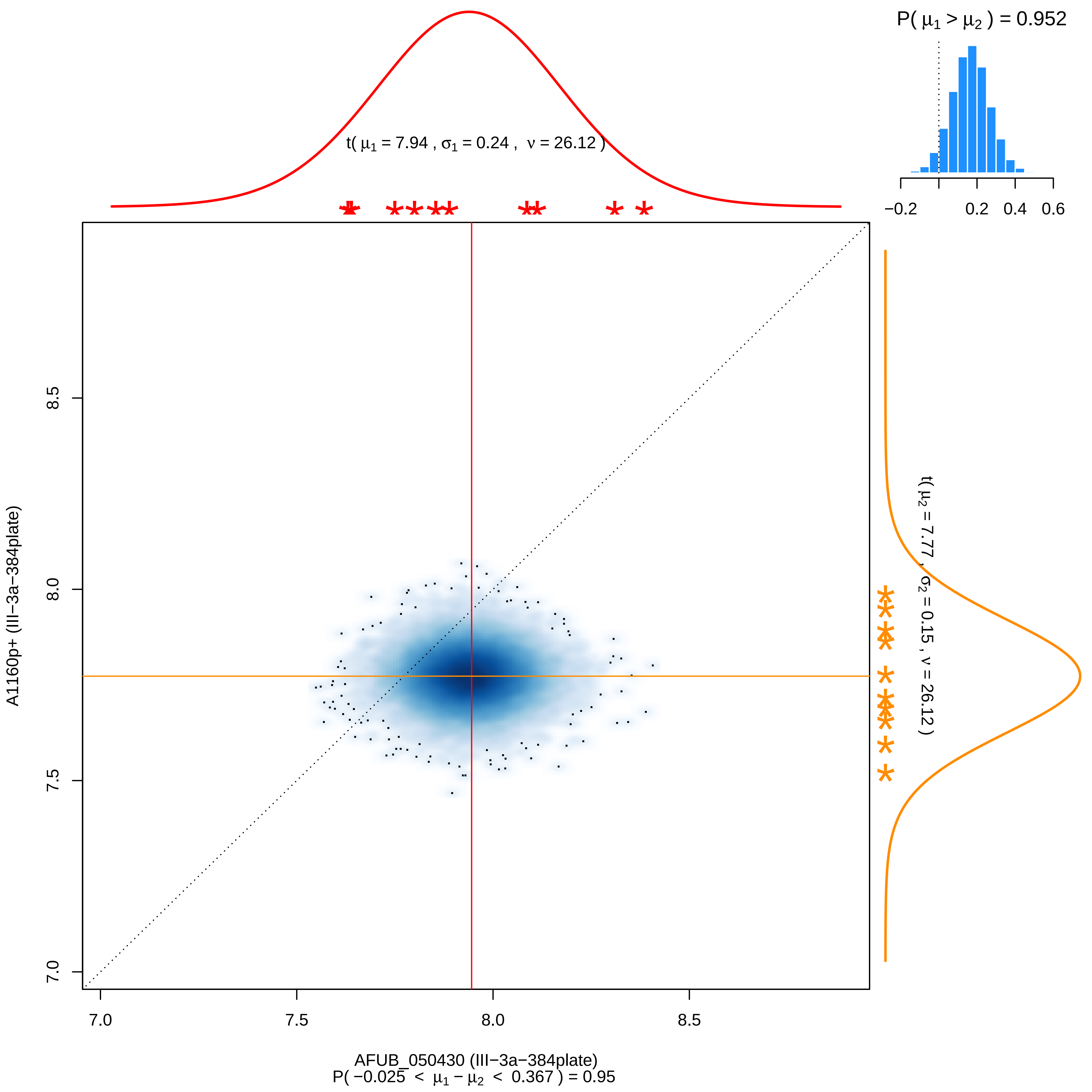

P( $\mu_1 > \mu_2$ ) = 0.952
−0.2
0.2
0.4
0.6
t( $\mu_1$ = 7.94 , $\sigma_1$ = 0.24 , $\nu$ = 26.12 )
t( $\mu_2$ = 7.77 , $\sigma_2$ = 0.15 , $\nu$ = 26.12 )
A1160p+ (III−3a−384plate)
7.0
7.5
8.0
8.5
AFUB_050430 (III−3a−384plate)
P( −0.025 < $\mu_1 - \mu_2$ < 0.367 ) = 0.95

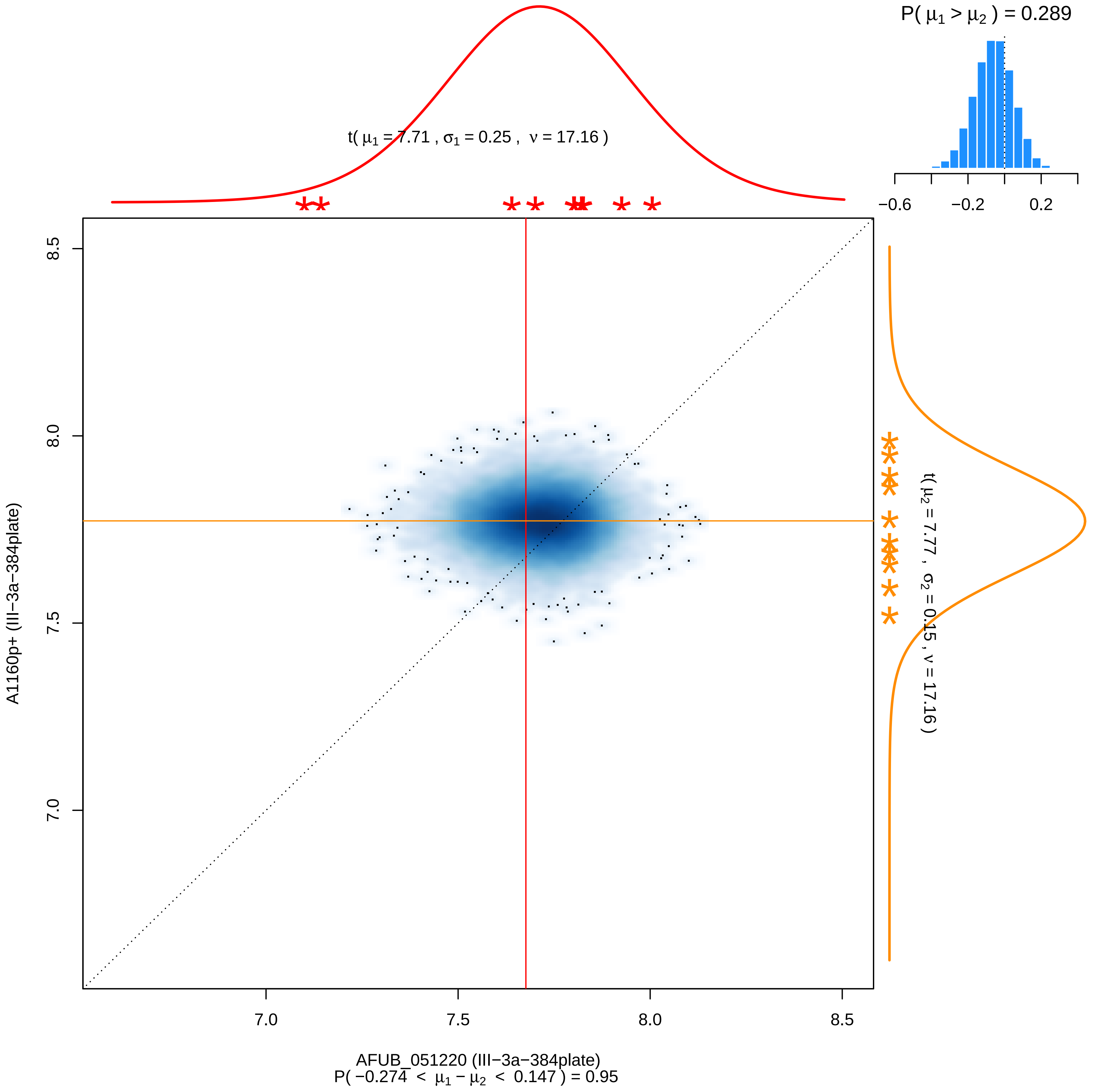

P( μ1 > μ2 ) = 0.289
−0.6
−0.2
0.2
t( μ1 = 7.71 , σ1 = 0.25 , ν = 17.16 )
t( μ2 = 7.77 , σ2 = 0.15 , ν = 17.16 )
8.5
8.0
7.5
7.0
A1160p+ (III−3a−384plate)
7.0
7.5
8.0
8.5
AFUB_051220 (III−3a−384plate)
P( −0.274 < μ1 − μ2 < 0.147 ) = 0.95

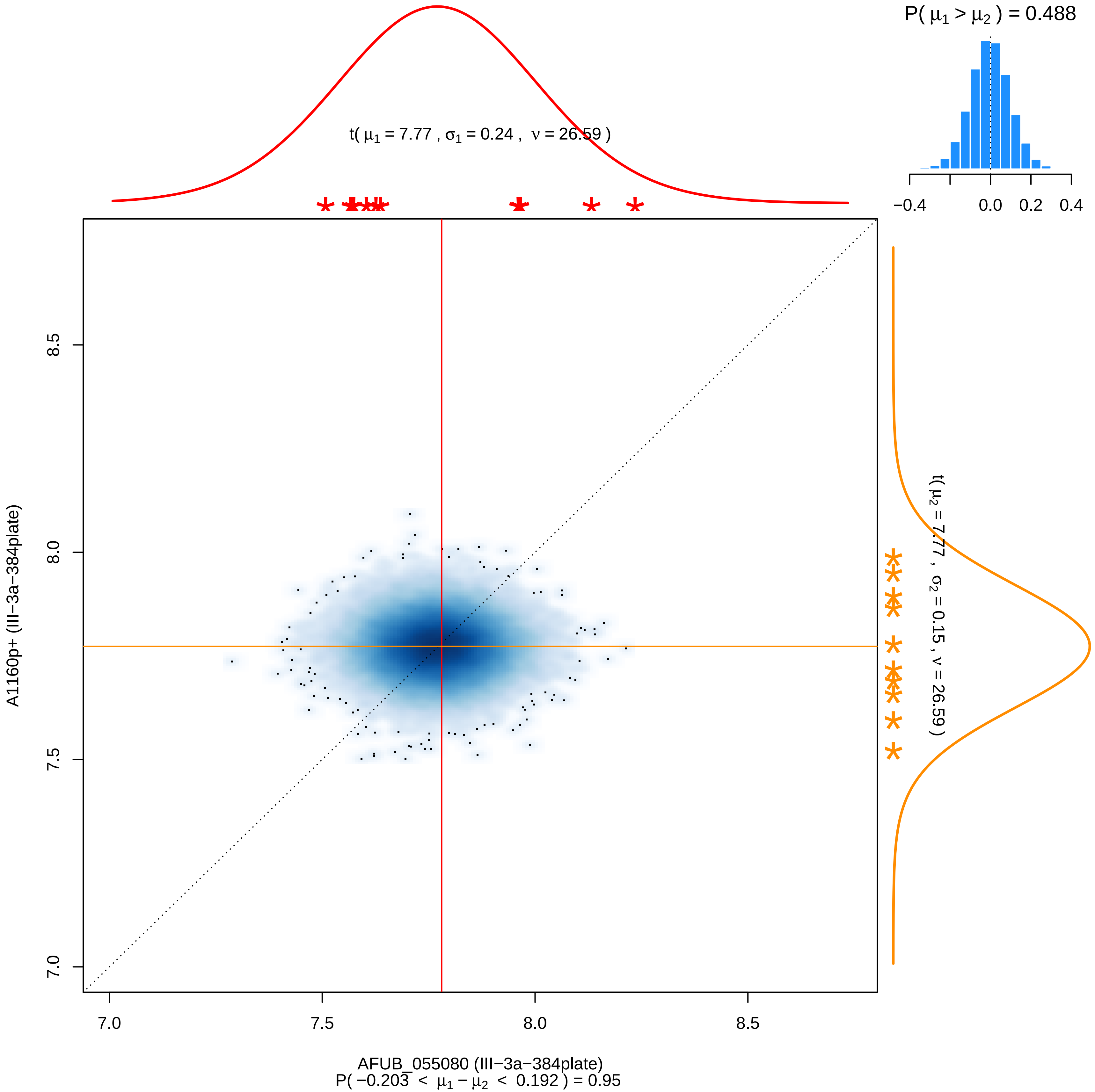

P( μ1 > μ2 ) = 0.488
−0.4
0.0
0.2
0.4
t( μ1 = 7.77 , σ1 = 0.24 , ν = 26.59 )
t( μ2 = 7.77 , σ2 = 0.15 , ν = 26.59 )
A1160p+ (III−3a−384plate)
8.5
8.0
7.5
7.0
7.0
7.5
8.0
8.5
AFUB_055080 (III−3a−384plate)
P( −0.203 < μ1 − μ2 < 0.192 ) = 0.95

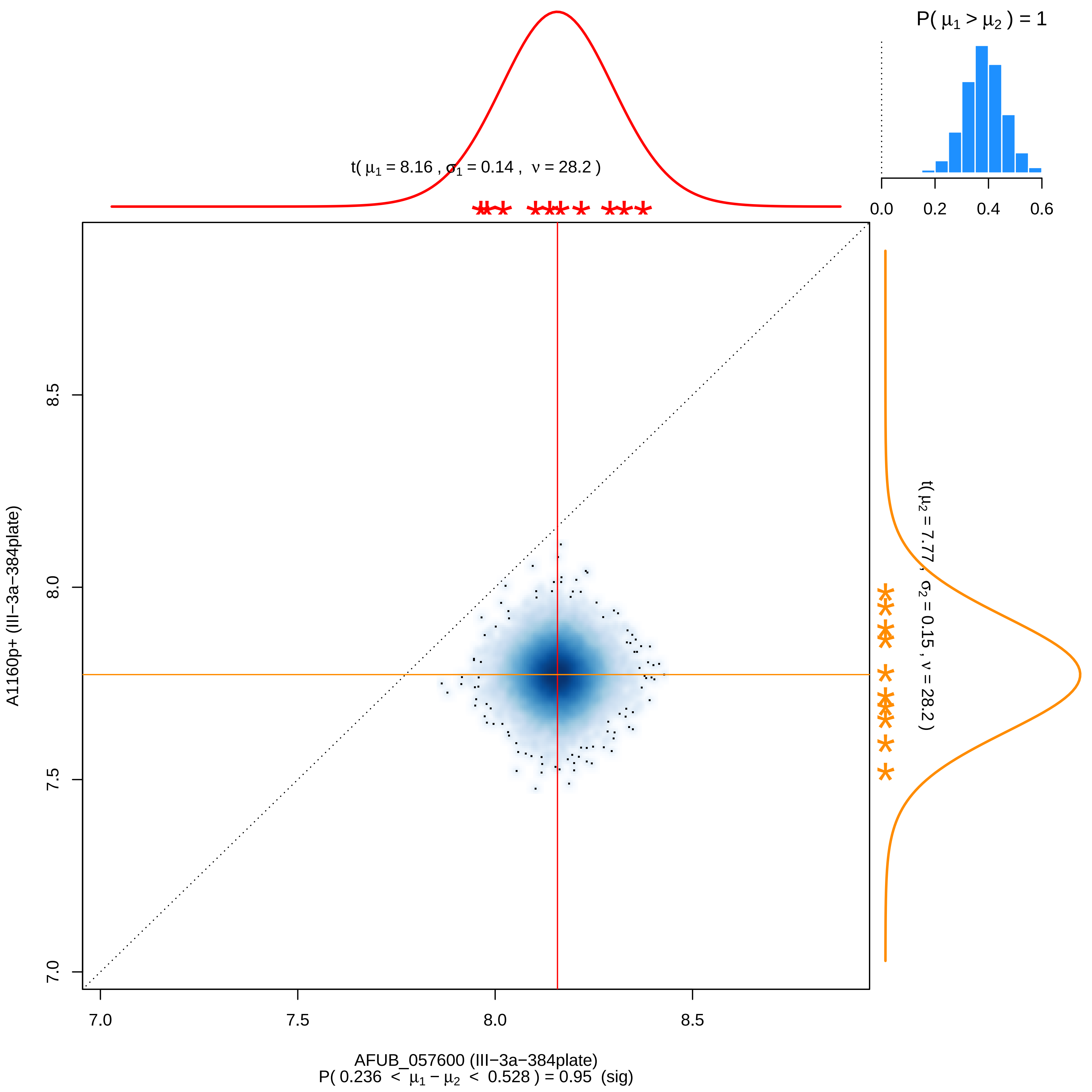

P( μ1 > μ2 ) = 1
0.0
0.2
0.4
0.6
t( μ1 = 8.16 , σ1 = 0.14 , ν = 28.2 )
t( μ2 = 7.77 , σ2 = 0.15 , ν = 28.2 )
A1160p+ (III−3a−384plate)
7.0
7.5
8.0
8.5
AFUB_057600 (III−3a−384plate)
P( 0.236 < μ1 − μ2 < 0.528 ) = 0.95 (sig)

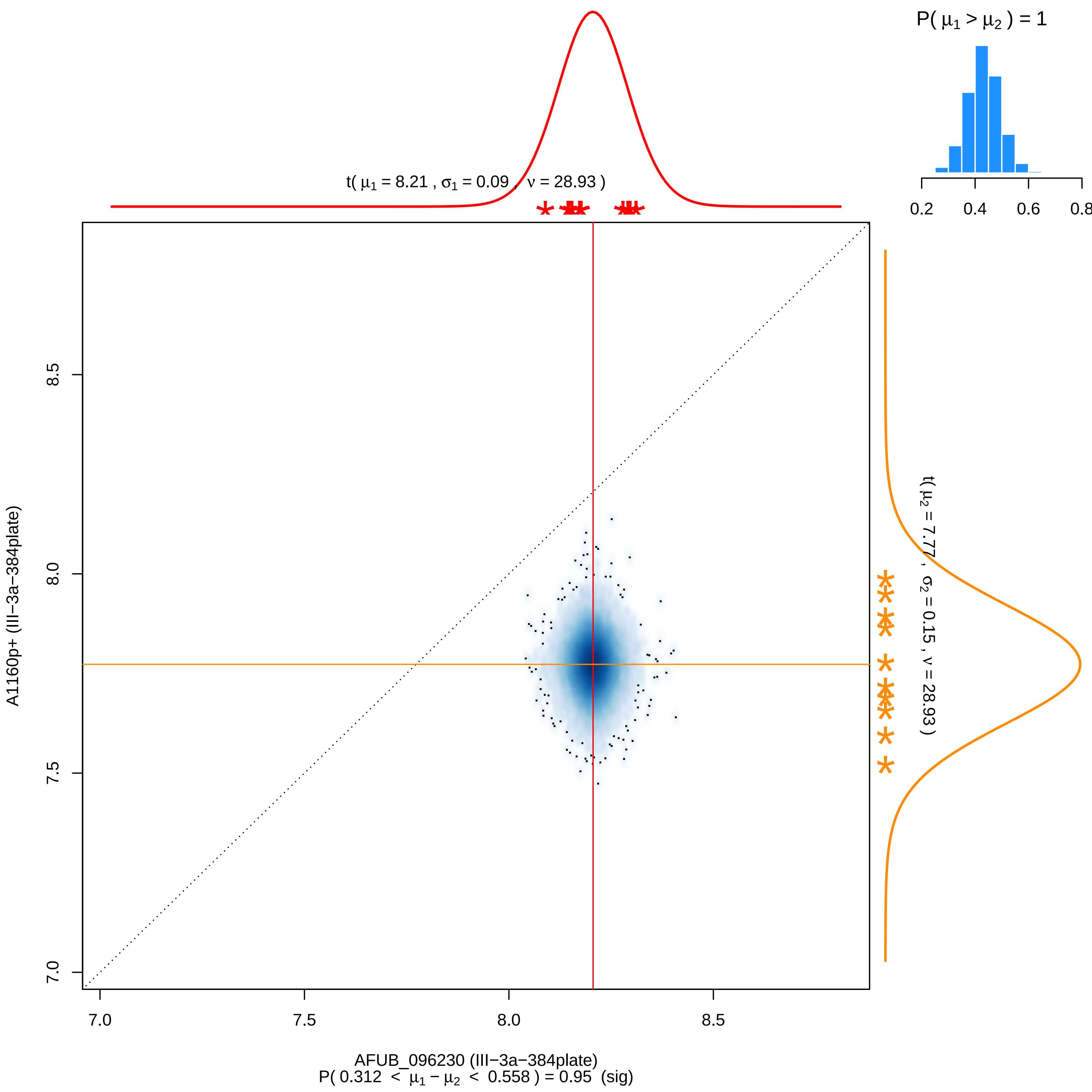

P( μ1 > μ2 ) = 1
0.2
0.4
0.6
0.8
t( μ1 = 8.21 , σ1 = 0.09 , ν = 28.93 )
t( μ2 = 7.77 , σ2 = 0.15 , ν = 28.93 )
A1160p+ (III−3a−384plate)
7.0
7.5
8.0
8.5
AFUB_096230 (III−3a−384plate)
P( 0.312 < μ1 − μ2 < 0.558 ) = 0.95 (sig)

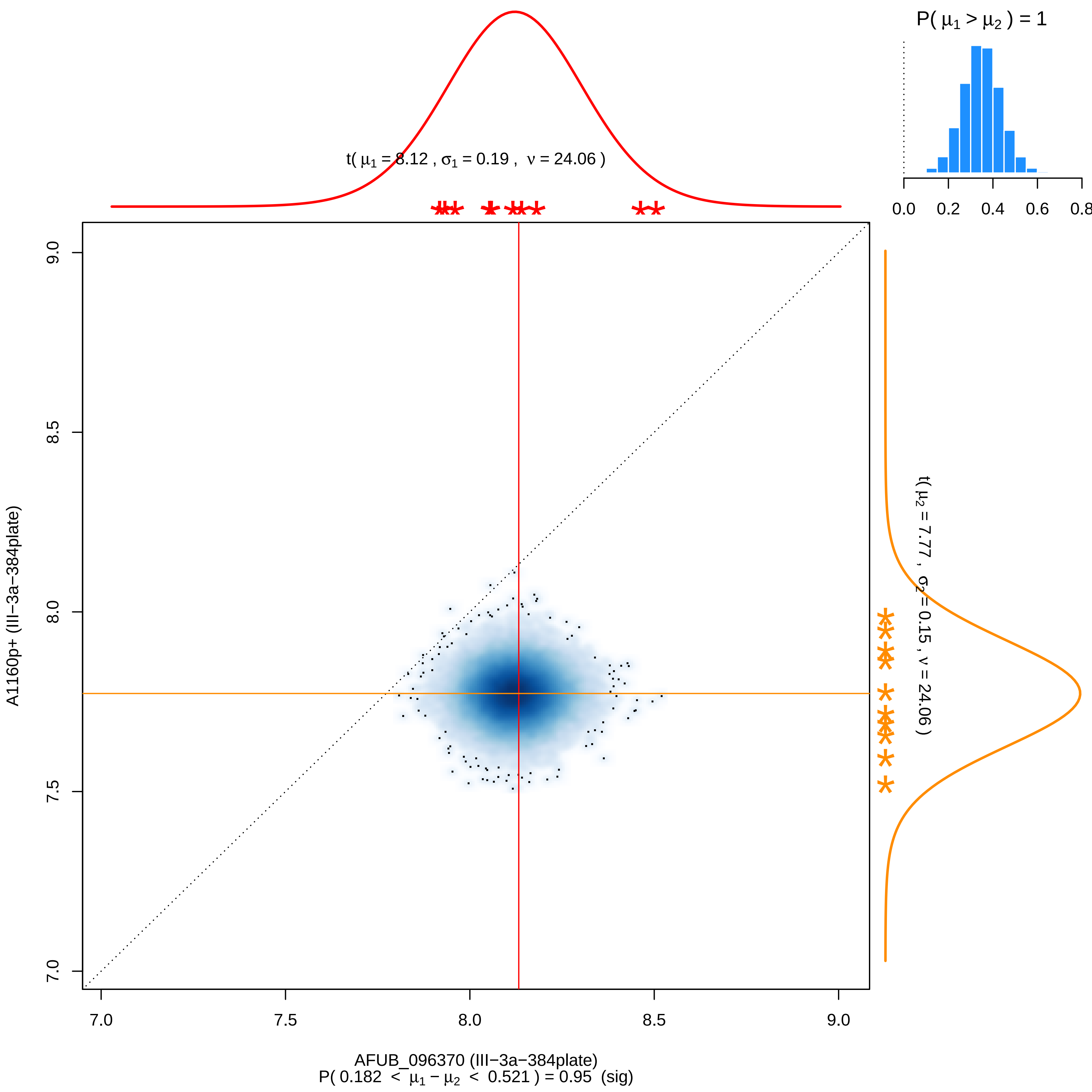

P( μ1 > μ2 ) = 1
0.0
0.2
0.4
0.6
0.8
t( μ1 = 8.12 , σ1 = 0.19 , ν = 24.06 )
t( μ2 = 7.77 , σ2 = 0.15 , ν = 24.06 )
9.0
8.5
8.0
7.5
7.0
A1160p+ (III−3a−384plate)
7.0
7.5
8.0
8.5
9.0
AFUB_096370 (III−3a−384plate)
P( 0.182 < μ1 − μ2 < 0.521 ) = 0.95 (sig)

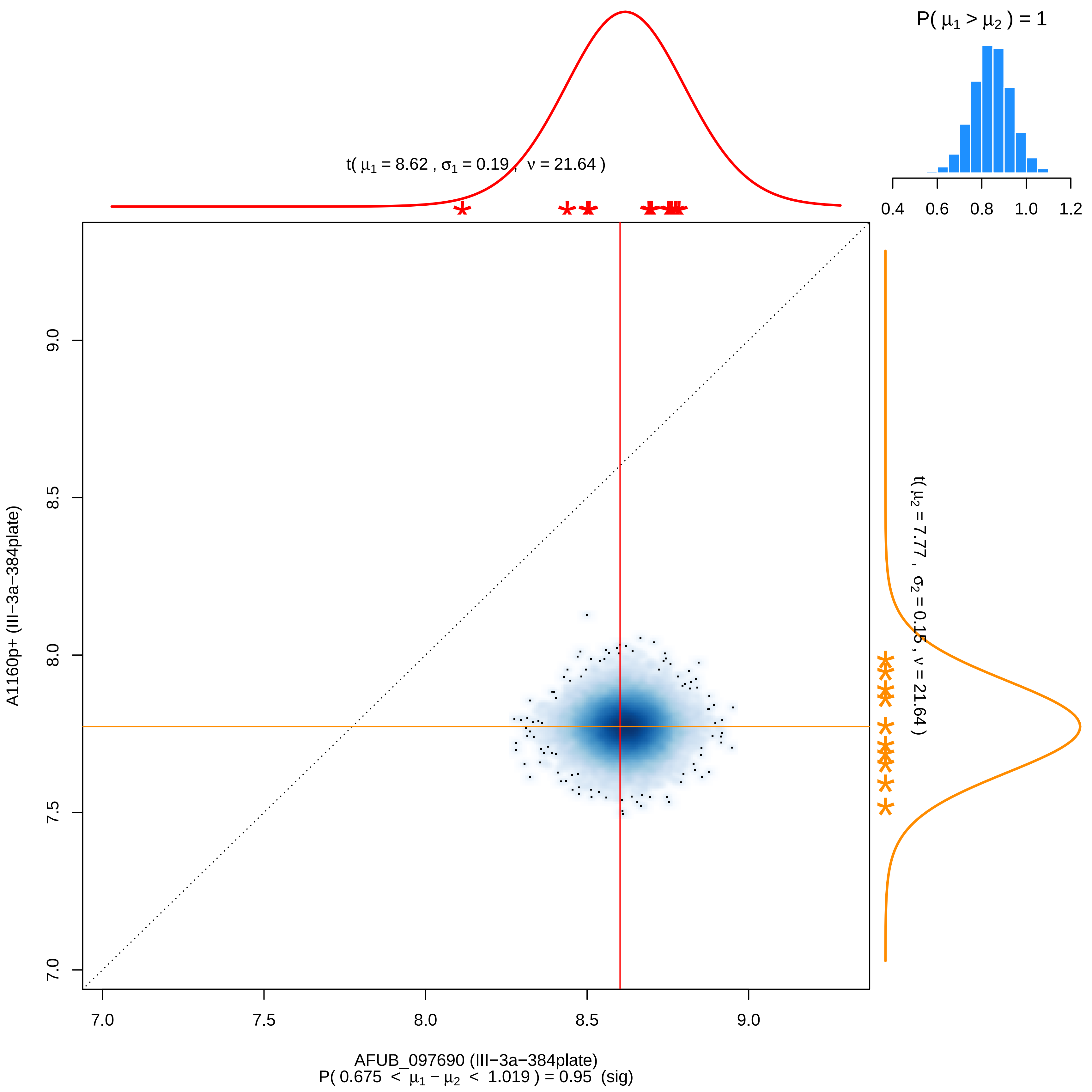

P( μ1 > μ2 ) = 1
0.4
0.6
0.8
1.0
1.2
t( μ1 = 8.62 , σ1 = 0.19 , ν = 21.64 )
t( μ2 = 7.77 , σ2 = 0.15 , ν = 21.64 )
A1160p+ (III−3a−384plate)
9.0
8.5
8.0
7.5
7.0
7.0
7.5
8.0
8.5
9.0
AFUB_097690 (III−3a−384plate)
P( 0.675 < μ1 − μ2 < 1.019 ) = 0.95 (sig)

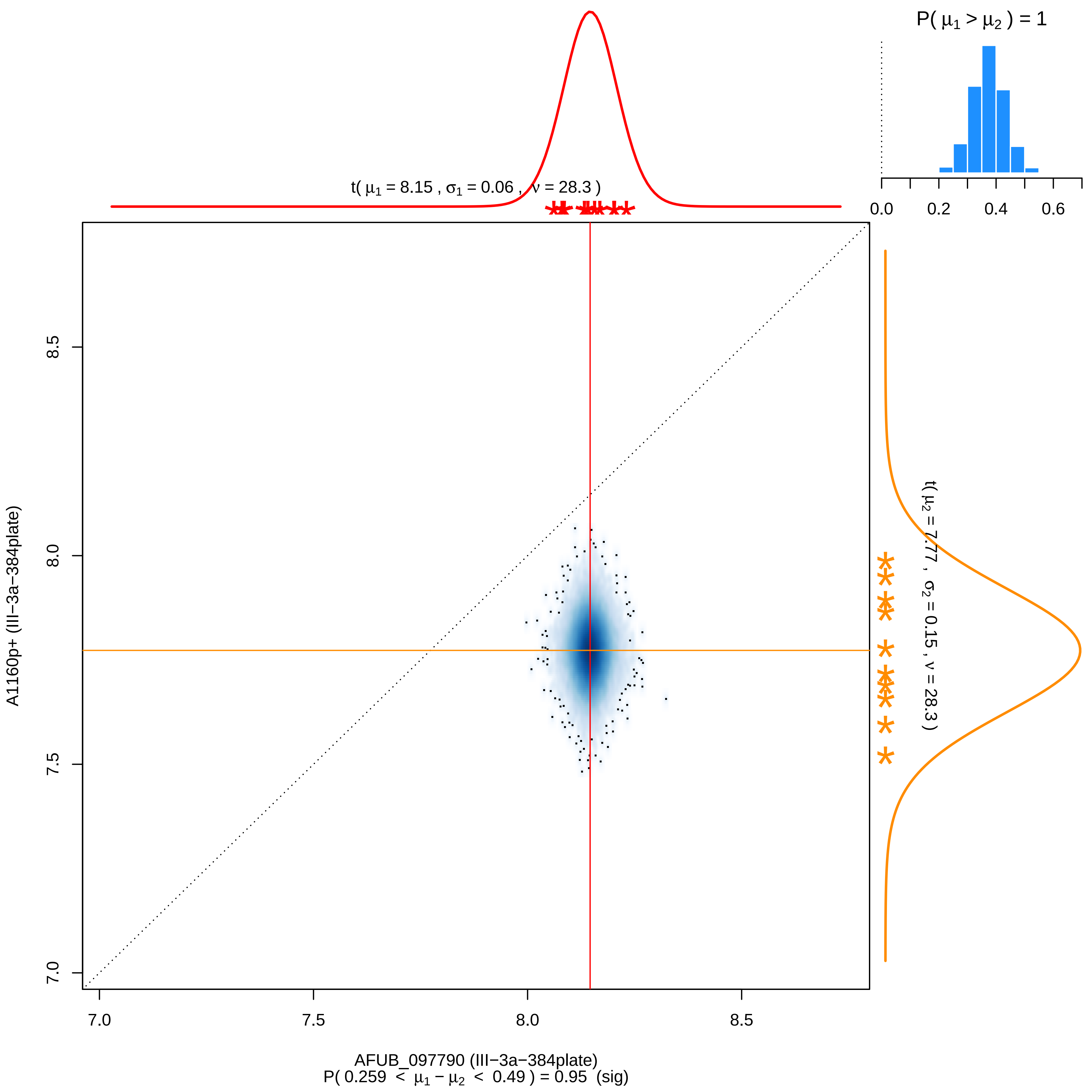

P( μ1 > μ2 ) = 1
0.0
0.2
0.4
0.6
t( μ1 = 8.15 , σ1 = 0.06 , ν = 28.3 )
t( μ2 = 7.77 , σ2 = 0.15 , ν = 28.3 )
8.5
8.0
7.5
7.0
A1160p+ (III−3a−384plate)
7.0
7.5
8.0
8.5
AFUB_097790 (III−3a−384plate)
P( 0.259 < μ1 − μ2 < 0.49 ) = 0.95 (sig)

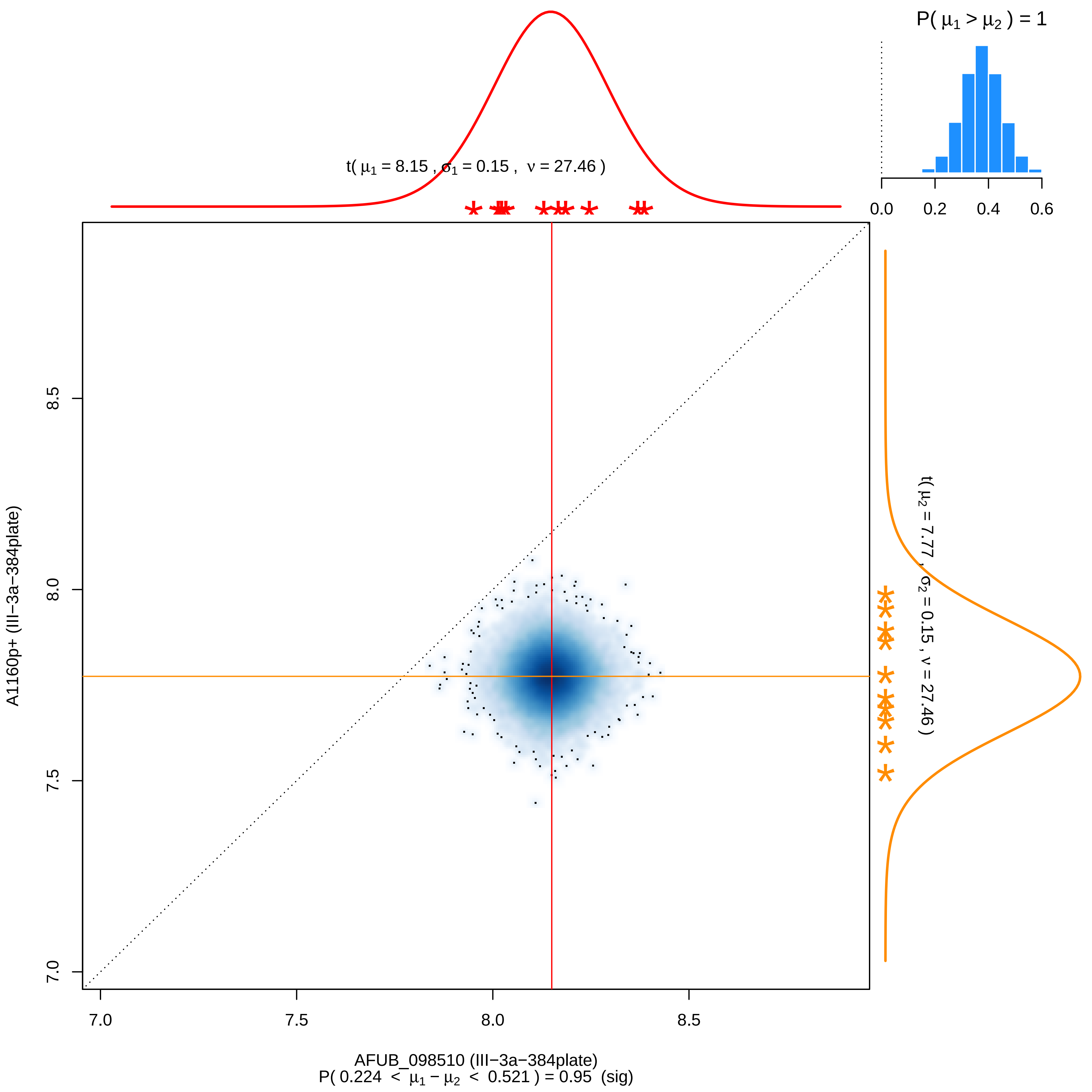

P( μ1 > μ2 ) = 1
0.0
0.2
0.4
0.6
t( μ1 = 8.15 , σ1 = 0.15 , ν = 27.46 )
8.5
8.0
7.5
7.0
A1160p+ (III−3a−384plate)
t( μ2 = 7.77 , σ2 = 0.15 , ν = 27.46 )
7.0
7.5
8.0
8.5
AFUB_098510 (III−3a−384plate)
P( 0.224 < μ1 − μ2 < 0.521 ) = 0.95 (sig)

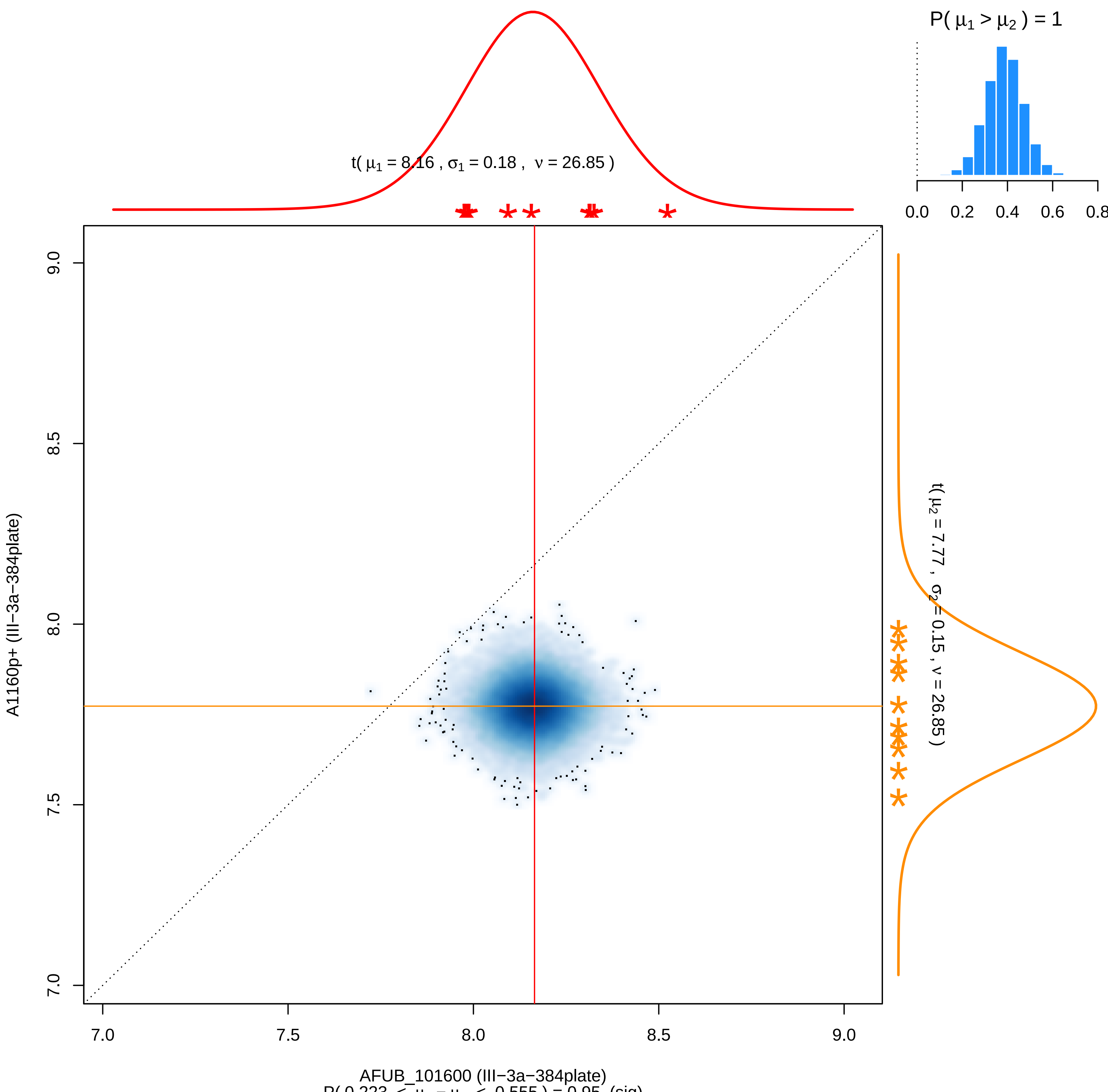


P( 0.223 < $\mu_1 - \mu_2$ < 0.555 ) = 0.95 (sig)

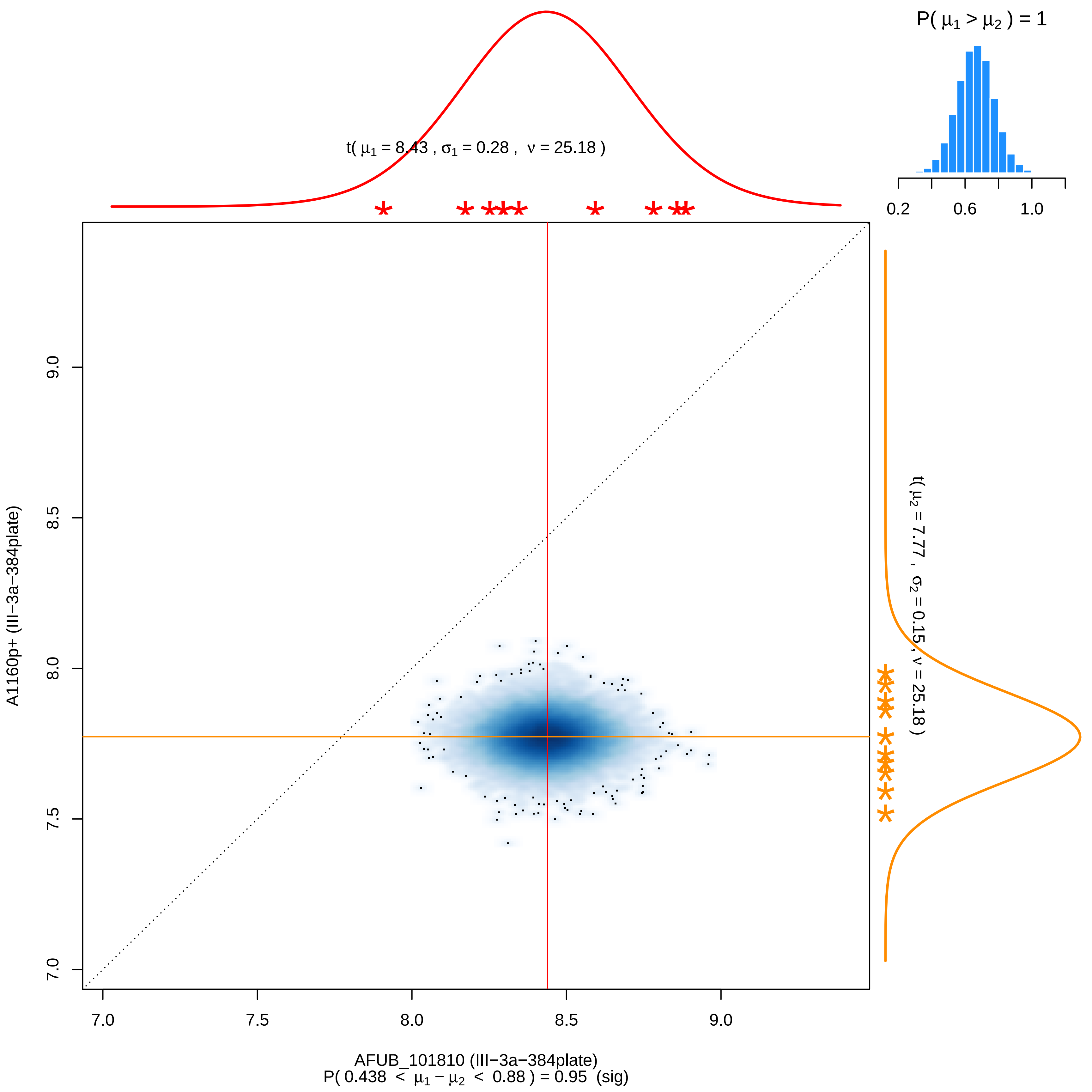

P( μ1 > μ2 ) = 1
0.2
0.6
1.0
t( μ1 = 8.43 , σ1 = 0.28 , ν = 25.18 )
t( μ2 = 7.77 , σ2 = 0.15 , ν = 25.18 )
9.0
8.5
8.0
7.5
7.0
A1160p+ (III−3a−384plate)
7.0
7.5
8.0
8.5
9.0
AFUB_101810 (III−3a−384plate)
P( 0.438 < μ1 − μ2 < 0.88 ) = 0.95 (sig)

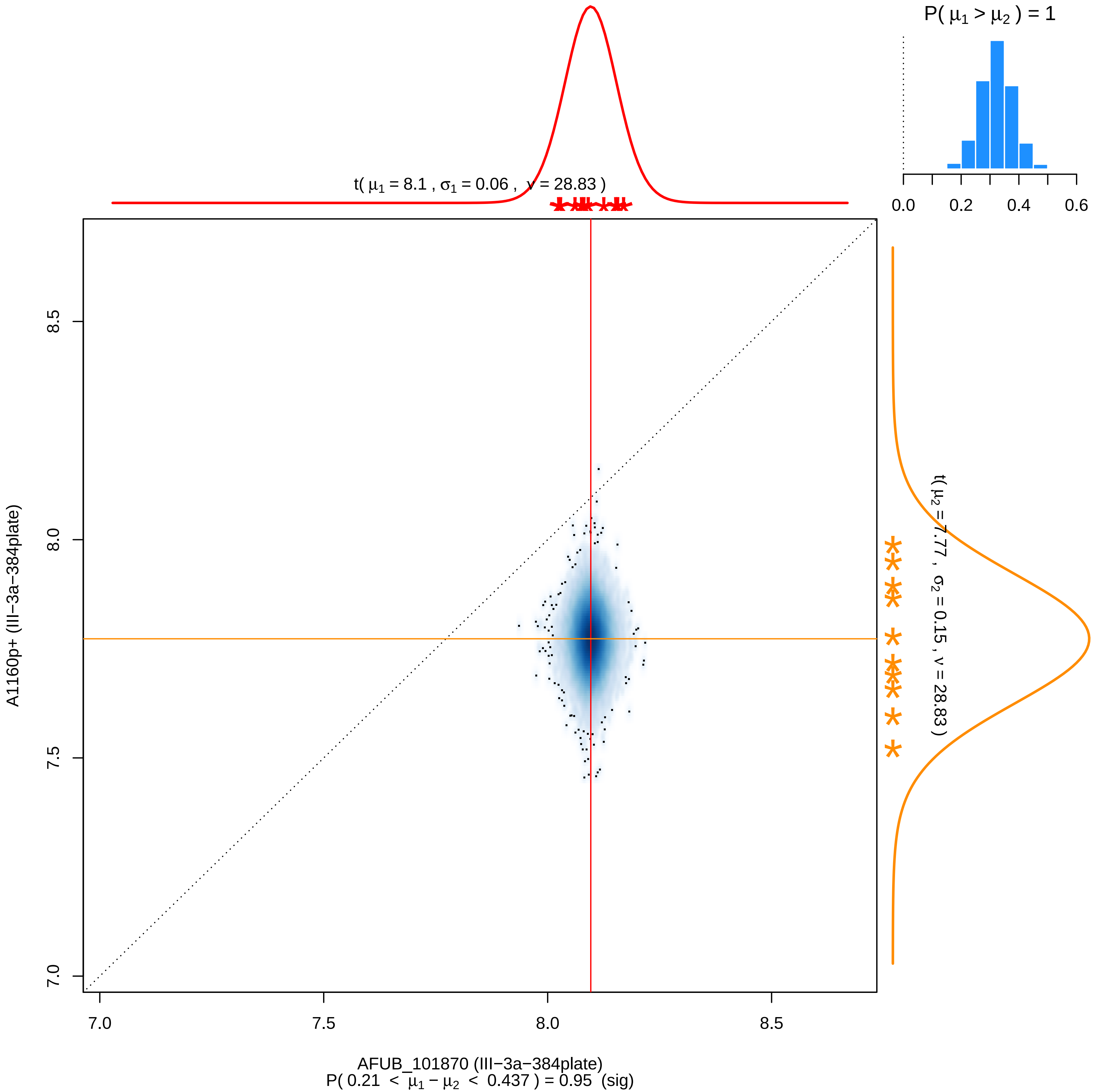
P( μ1 > μ2 ) = 1
0.0
0.2
0.4
0.6
t( μ1 = 8.1 , σ1 = 0.06 , ν = 28.83 )
t( μ2 = 7.77 , σ2 = 0.15 , ν = 28.83 )
8.5
8.0
7.5
7.0
A1160p+ (III−3a−384plate)
7.0
7.5
8.0
8.5
AFUB_101870 (III−3a−384plate)
P( 0.21 < μ1 − μ2 < 0.437 ) = 0.95 (sig)

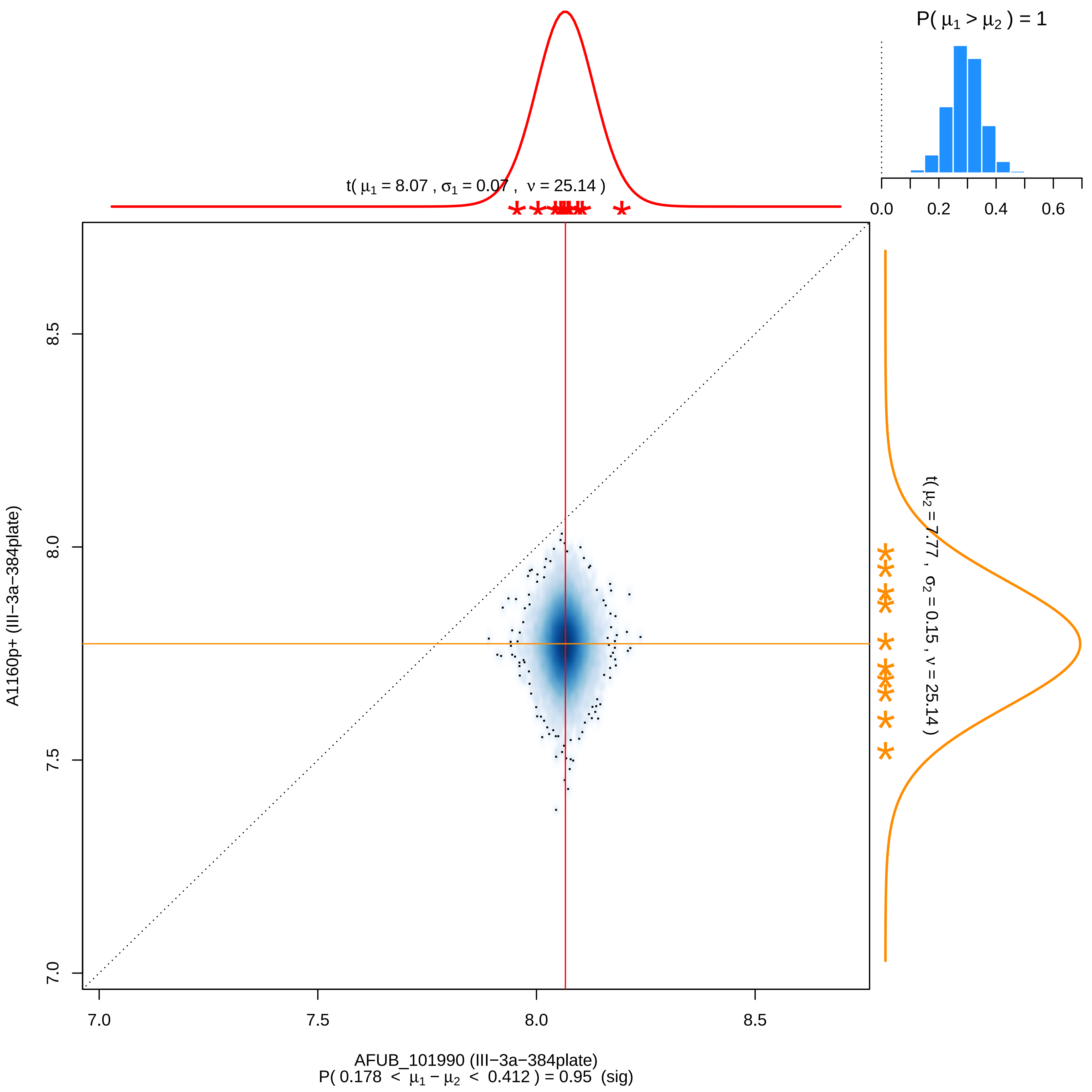

P( μ1 > μ2 ) = 1
0.0
0.2
0.4
0.6
t( μ1 = 8.07 , σ1 = 0.07 , ν = 25.14 )
t( μ2 = 7.77 , σ2 = 0.15 , ν = 25.14 )
7.0
7.5
8.0
8.5
A1160p+ (III−3a−384plate)
AFUB_101990 (III−3a−384plate)
P( 0.178 < μ1 − μ2 < 0.412 ) = 0.95 (sig)

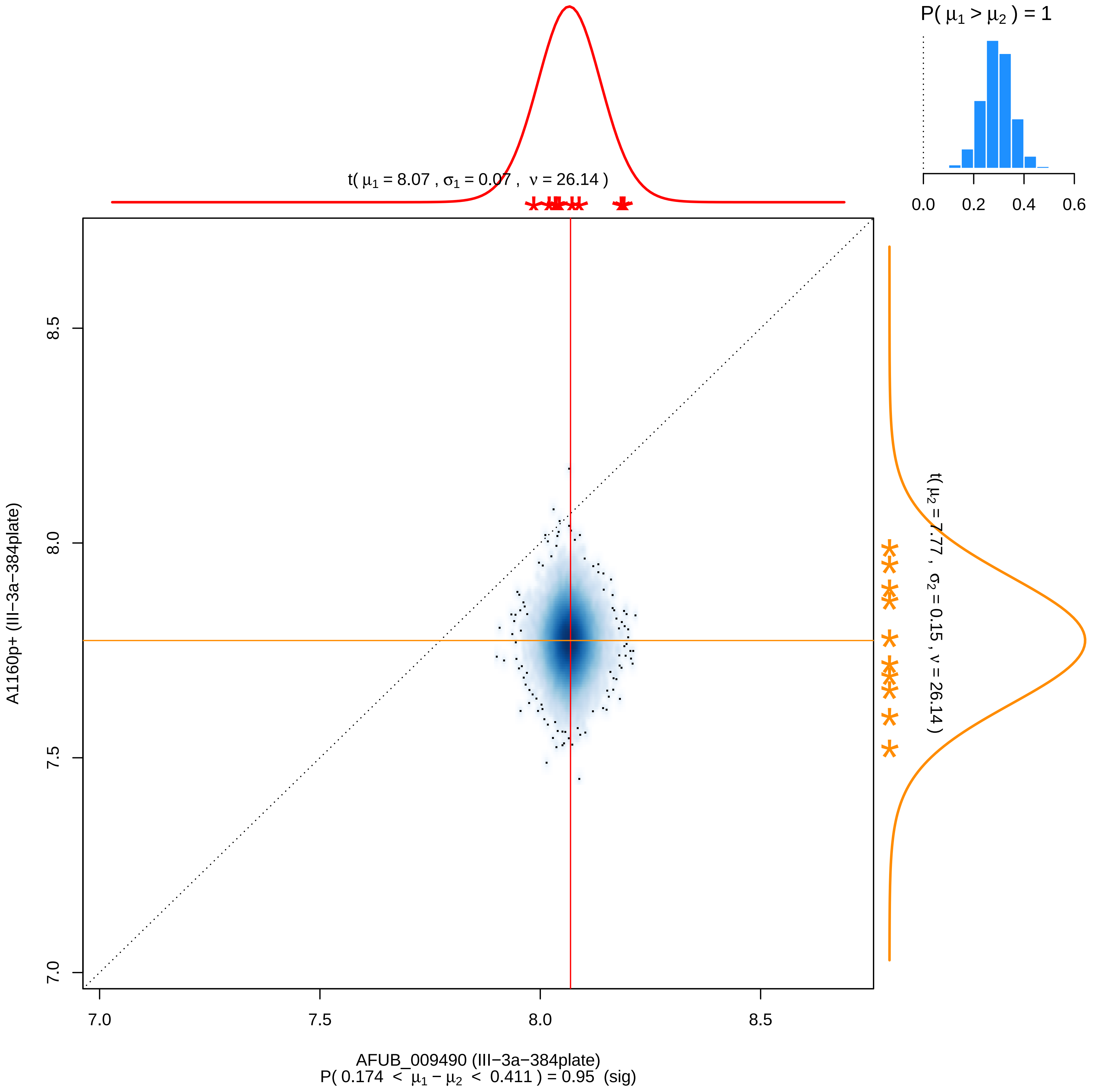
P( $\mu_1 > \mu_2$ ) = 1
0.0
0.2
0.4
0.6
t( $\mu_1$ = 8.07 , $\sigma_1$ = 0.07 , $\nu$ = 26.14 )
t( $\mu_2$ = 7.77 , $\sigma_2$ = 0.15 , $\nu$ = 26.14 )
A1160p+ (III−3a−384plate)
8.5
8.0
7.5
7.0
7.0
7.5
8.0
8.5
AFUB_009490 (III−3a−384plate)
P( 0.174 < $\mu_1 - \mu_2$ < 0.411 ) = 0.95 (sig)
